# Genome-Wide Mapping of Cisplatin Damaged Gene Loci

**DOI:** 10.1101/2020.07.27.222752

**Authors:** Luyu Qi, Qun Luo, Yan Xu, Wanchen Yu, Xingkai Liu, Yanyan Zhang, Feifei Jia, Tiantian Fang, Shijun Wang, Xiangjun Li, Yao Zhao, Fuyi Wang

## Abstract

Cisplatin is a DNA targeting anticancer drug, yet its damaged gene loci have remained unclear. In the present work, combining affinity isolation and high throughput sequencing, we genome-widely mapped 17729 gene loci containing platination lesions, which mainly function as enzymes, transcription regulators, transporters and kinases, and of which 445 genes account for 71% of potential gene targets for cancer therapy reported in the literature. The most related core signaling pathway, disease and tissue toxicity of 7578 genes with an enrichment fold (EF_G_) of >12, where EF_G_ refers to the ratio of total read counts of a gene detected in cells with and without cisplatin treatment, are sperm motility, cancer and hepatotoxicity with association *P* values of < 1×10^−22^. Among 616 kinase genes damaged by cisplatin, 427 are protein kinases which account for 82% of putative protein kinases, suggesting that cisplatin may act as broad-spectrum protein kinase inhibitor. Western Blot assays verified that expression of 8 important protein kinase genes was significantly reduced due to cisplatin damage. *SPAG9* is closely related to 147 of 361 cancer diseases which the cisplatin damaged genes are associated with and was severely damaged by cisplatin. Given *SPAG9* abundantly expresses JIP-4, a upstream mediator of protein kinase signaling, in testis, it may be responsible for the high sensitivity of testicular cancer to cisplatin, thus being a potential therapeutic target for precise treatment of testicular cancer. These findings provide novel insights into better understanding in molecular mechanism of anticancer activity and toxicity of cisplatin, more importantly inspire further studies in prioritizing gene targets for precise treatment of cancers.

## Introduction

Cisplatin, cis-diamminedichloroplatinum,^1^ has been widely used as a first line chemotherapeutic drug since its approval by the FAD in the late 1970’s.^2,3^ Although a number of molecularly targeting anticancer drugs including small molecular drugs and antibody drugs, for example protein tyrosine kinase inhibitor gefitinib^4^ and PD-1 antibody^5^, have been developed and applied in recent years, cisplatin is still involved extensively in clinic treatment, in particular systemic and combining chemotherapy of solid tumors.^6,7^ After hydrolysis inside cells, cisplatin can bind to a variety of biological molecules, e.g. proteins, phospholipids, RNA, carbohydrates, thiol-containing small molecules and nuclei acids.^3,8^ However, it is generally believed that attacking at nuclear DNA is the main pathway of cisplatin as an anticancer drug. The platinum center of cisplatin can bind to N7 atom on purine ring of nucleobases to form crosslinked DNA complexes, which include 1,2-intrastrand -GG- and/or -AG-adducts (ca. 90%) and 1,3-intrastrand -GNG-adducts (5-10%), in addition to a low percentage of monofunctional and interstranded cross-links.^9^ The coordination of cisplatin with DNA bases partially unwinds and bends DNA helices, as a consequence inhibiting DNA replication and transcription, thus inducing cell apoptosis and death.^10^ Previous studies have characterized the distribution of cisplatin binding sites in gene regions at the whole genome scale.^11–13^ However, the exact gene loci attacking by cisplatin has remain unclear yet, limiting deeper understanding in action mechanism of cisplatin.

On the other hand, despite its great success in clinic treatment of solid tumors, the clinic application of cisplatin has been hampered due to severe side effects or non-tissue specific toxicity, such as nephrotoxicity, ototoxicity, neurotoxicity, gastrointestinal toxicity, hematological toxicity, cardiotoxicity, hepatotoxicity, and etc.^14,15^ Given that DNA is the ultimate target of cisplatin, the non-specific attacking of cisplatin on gene DNA is most probably associated with its toxicity, though the direct interactions of cisplatin with proteins may be also implicated in the toxicity of the drug. It has been reported that patients with different genotypes showed different frequencies of various side effects.^15^ Therefore, it is also very important to identify which genes damaged by cisplatin cause specific toxicity.

In order to localize cisplatin damaged gene loci at the whole genome scale, in this work, we construct a magnetic affinity probe by functionalizing Fe_3_O_4_@Ni core-shell micro-beads with HMGB1a, referring to the domain A of high mobility group box protein 1 (HMGB1) which is a non-specific DNA-binding protein and showed specific-binding to 1,2-cisplatin crosslinked DNA,^3,9,16^ to capture and isolate cisplatin-damaged DNA fragments extracted from cisplatin-treated cells, followed by high throughput sequencing (Figure 1). Due to non-specific binding of cisplatin to gene DNA, it is not surprising that we identified more than 17000 genes which may be damaged by cisplatin. The bioinformatic analysis of 7578 cisplatin-damaged genes with an enrichment fold of >12 in comparison with the control samples showed that the most relevant signaling pathway of cisplatin-damaged genes is sperm mobility, and the most relevant disease is cancer. Combined with the finding that *SPAG9* gene, which abundantly expresses C-jun-amino-terminal kinase-interacting protein 4 (JIP-4) in testicular haploid germ cells, was heavily attacked by cisplatin, our study herein provides novel insights into better understanding in the high cure rate of cisplatin towards testicular cancer.

**Figure 1.**
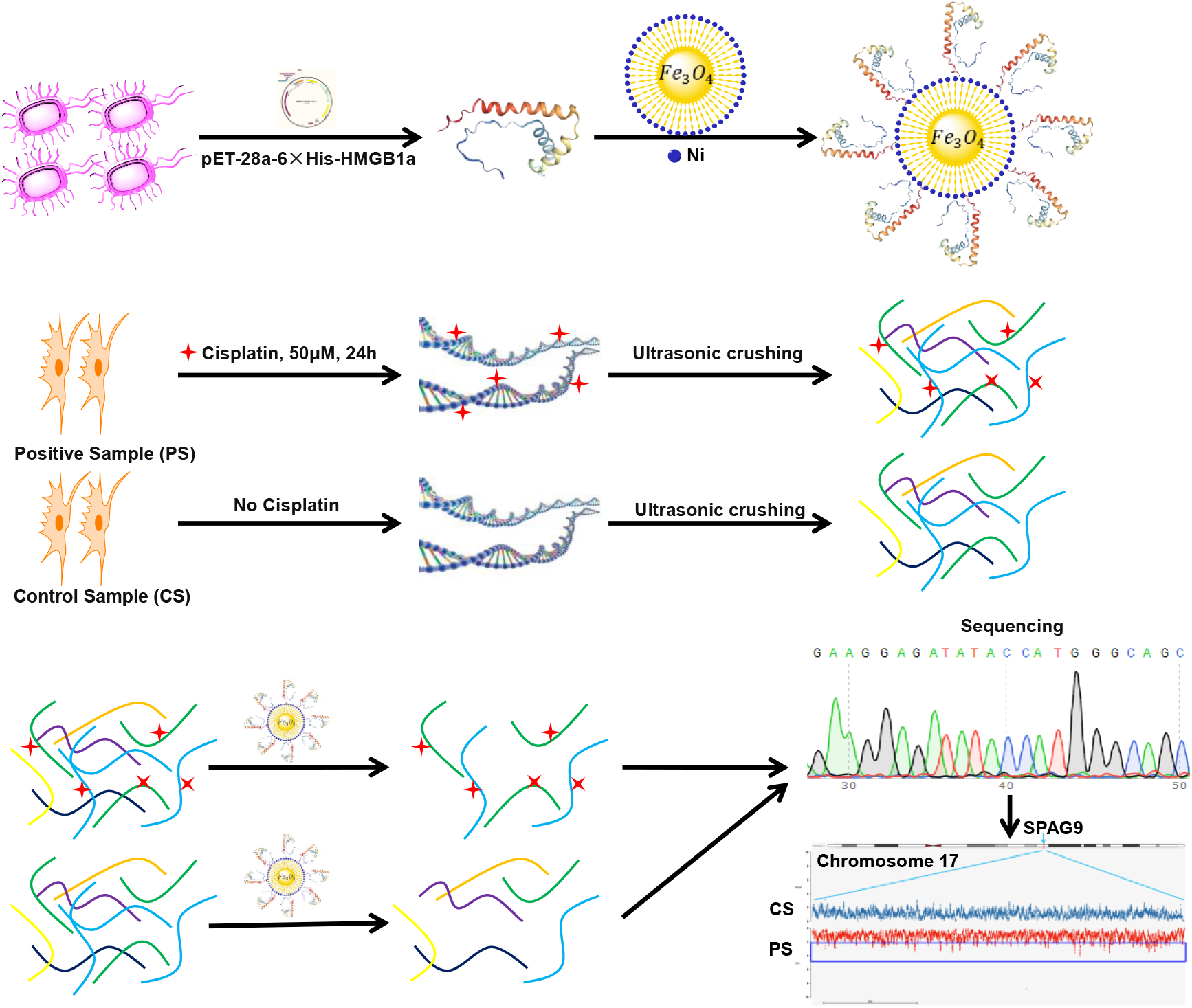
Diagrammatic illustration of the workflow for capture and identification of gene loci attacked by cisplatin. (A) HMGB1a expression and construction of HMGB1 affinity microprobe. (B) Cell culture, and the extraction and fragmentation of DNA. (C) Capture, sequencing and alignment of cisplatin damaged DNA fragments.

## Experimental Procedures

### Expression and purification of HMGB1a

The cDNA of human HMGB1 domain A (HMGB1a, Supplementary Chart 1) was synthesized by Proteintech (Rosemont, USA) and cloned into pET-28a plasmid. The resulting plasmid encoding a his_6_-tag was transfected into BL21 (DE3) *E. Coli* cells (Tian Gen Biotech, China), which were grown in a small volume of 2YT medium containing Bacto™ Peptone (16 mg/mL), Bacto™ Yeast Extract (10 mg/mL, Becton, Dickinson and company, USA), Sodium chloride (5 mg/mL, Beijing Chemical Works, China) and kanamycin (50 μg/mL, Solarbio, China) overnight at 37 °C and 180 rpm. Then the cells were transferred to a big volume of fresh 2YT medium at 37 °C and 180 rpm. When the OD_600_ value of the cells reached 0.5 – 1.0, IPTG (Sigma-Aldrich, USA) was added at a final concentration of 1 mM to induce gene expression. After incubated at 37 °C and 180 rpm for further 12 h, the cells were harvested and washed by PBS (Solarbio, China) for one time, and frozen at −80°C for at least one hour for better sonication. The deep frozen cells were resuspended in 10 mL cold lysis buffer (PBS and 5 mM imidazole, pH 7.4), and lysed by sonication for 20 min at 0 °C, followed by centrifugation for 30 min at 4 °C and 12000 rpm. The supernatant was purified by His Trap™ FF crude column (GE Healthcare) which was eluted first by 50 mM imidazole (Innochem), then by 300 mM imidazolein phosphate buffer. The eluted HMGB1a containing a his_6_-tag was characterized by SDS-PAGE and mass spectrometry (Supplementary Figure 1).

### Assembly and Characterization of HMGB1a-functionilizaed affinity microprobe

To construct affinity microprobe for cisplatin crosslinked DNA, 2.1 mg purified HMGB1a was incubated with 7.5 mg nickel-modified magnetic beads (MAg25K/IDA Ni, Enriching Biotechnology, Shanghai) suspended in 1 mL PBS, and rotated for 1 h at 4 °C. The beads were then separated by magnetic force and washed twice with PBS for 10 min. As shown in Supplementary Figure 2, the absorbance of the supernatant over the wavelength ranging from 250 – 300 nm significantly decreased subject to loading of HMGB1a into the magnetic bead. This indicated successful functionalization of the magnetic beads by HMGB1a.

### Cell culture and genomic DNA extraction

The A549 human non-small cell lung cancer cells obtained from the Centre for Cell Resource of Peking Union Medical College Hospital were cultured in DMEM medium (Gibco) supplemented with 10% FBS (Gibco) and 1% penicillin and streptomycin (GE Healthcare) at 310 K. When the cell density reached confuency, the culture medium was replaced by fresh DMEM containing 50 μM cisplatin (Beijing Ouhe Technology, China), and the cells were incubated at 37 °C for 24 h. Then the cells were harvested, and genomic DNA was extracted by whole genome extraction kit (Tian Gen Biotech). The DNA extracts were crushed into fragments between 250 to 750 bp with an ultrasonic breaker (Scientz-IID ultrasonic homogenizer, SCIENTZ, Ningbo, China). At the same time, a similar amount of cells without exposure to cisplatin were harvested for genomic DNA extract and fragmentation. The DNA fragments from the cells treated with cisplatin were considered as a positive sample and those from the cells without cisplatin treatment were considered as a control blank for capture and sequencing of cisplatin damaged gene loci. An ICP-MS (Agilent 7700) instrument was applied to measure the Pt concentration of the two DNA fragment samples.

### Capture of cisplatin-bound DNA fragments

The same amount (74 μg) of DNA fragments gained from cells treated and untreated with cisplatin were individually re-dissolved in 10 mM Tris-HCl (pH 7.4, Beijing Biodee Biotechnology), and added into the HMGB1a-functionlized magnetic beads suspended in PBS, followed by rotation for 1 h at room temperature. The beads were then separated by magnetic force and washed twice with 1 mL 10 mM Tris-HCl, and the captured HMGB1a-DNA complexes were eluted by 1 mL buffer containing 20 mM sodium phosphate, 500 mM NaCl and 400 mM imidazole (pH 7.4) *via* rotation at room temperature for 40 min.

Each eluted HMGB1a-DNA complex sample was centrifuged and lyophilized, and then re-dissolved in 500 μL deionized water, followed by addition of 500 μL mixture of phenol (Solarbio) – chloroform (Beijing Chemical Works) – isopropanol (Concord Technology, Tianjin) (25: 24: 1). The resulting mixture was shaken firstly for 10 s, and kept at room temperature for 2 min, then centrifuged at 12000 rpm and 4 °C for 7 min. The supernatant was mixed with 500 μL mixture of chloroform-isopropanol (24: 1), shaken again for 10 s, kept at room temperature for 2 min, and then centrifuged at 12000 rpm and 4 °C for 7 min. The isolated supernatant was mixed with 1500 μL pre-chilled absolute ethanol (Concord Technology, Tianjin) and 50 μL sodium acetate solution (3M, Beijing Chemical Works), kept at −20 °C overnight, and then centrifuged at 12000 rpm and 4 °C for 7 min. The supernatant was carefully removed, and the residues were left at room temperature for drying. Finally, the purified DNA fragments were re-dissolved in appropriate amount of TE solution (Solarbio) for sequencing.

### Sequencing and gene mapping

Sequencing of captured and purified DNA fragments were performed by Shanghai Sangon Biotech Ltd. The DNA fragments were firstly characterized by 1% agarose gel electrophoresis for quality check, and then fragmented to about 500 bp by Covaris and recovered for DNA library construction. Library construction was performed following the instruction of NEB Next^®^ Ultra ™ DNA Library Prep Kit for Illumina^®^ (https://international.neb.com/products/e7370-nebnext-ultra-dna-library-prep-kit-for-illumina#Protocols,%20Manuals%20&%20Usage). Since cisplatin binding to DNA fragments inhibits DNA replication and synthesis,^3,9,10^ the PCR extension time was prolonged to 30 min. The constructed DNA library was characterized by a 2% agarose gel, and then each positive or control sample was sequenced by Illumina^®^ at least 20 G of data. The sequencing data of each pair of positive/control sample were simultaneously matched with the human reference genome (hg38) to map genome-widely the DNA fragments captured.

The softwares and databases used for gene screening and location are listed in Supplementary Table 1. Ingenuity Pathway Analysis (IPA, QIAGEN) was used to analyze the core pathways, diseases and functions associated with the identified genes damaged by cisplatin.^17^ IPA analysis maps each gene to the corresponding molecule in the Ingenuity Pathway Knowledge Base, which is available at the Ingenuity System’s web site (http://www.ingenuity.com).

### Western Blot assays

The A549 human non-small cell lung cancer cell line, NCCIT testicular cancer cell line, HeLa cervical cancer cell line and HepG2 hepatoma cell line (National Infrastructure of Cell Line Resource, Beijing, China) were individually grown in the presence and absence of cisplatin at various concentration (50 μM for A549 and HepG2, 12 μM for NCCIT, 25 μM for HeLa) for 24 h. The cells were then harvested for extracting whole cell proteins using the total protein extraction kit (BestBio, China). The concentration of each protein extract was determined by standard BCA assay (Beyotime, China). To determine the expressing level of a specific protein due to gene damage by cisplatin, the same amount of whole cell proteins extracted from a positive sample and a control sample were individually boiled with the gel-loading buffer (4×LDS, Genscript, Nanjing, China), and loaded onto a 4 – 12% gradient SDS-PAGE gel (Genscript) for electrophoresis. The separated proteins were transferred to a PVDF membrane (Millipore, 0.2 μm). The membrane was blocked in 5% nonfat dry milk (Biofroxx) dissolved in 0.1% TBST, of which 1 L contained 2.4 g Tris base (Innochem), 8.8 g NaCl and 0.1% (v/v) Tween 20 (Beijing Topbio Science&Technology), at 4 °C overnight and then incubated with respective primary antibodies (Abcam) in appropriate dilutions at room temperature for 2 h. Thereafter, the membranes were washed three times with 5% nonfat dry milk in 0.1% TBST for 30 min each time and incubated with horseradish peroxidase-conjugated secondary antibody (Abcam) at room temperature for 1 h. After washing with 5% nonfat dry milk in 0.1% TBST for twice and 0.1% TBST for one time, the protein bands were visualized by enhanced chemiluminescence (Beyotime, China), and the optical densities were quantified using an image analyzer (Image J).

## Results and Discussion

### The capture of DNA fragments and DNA library construction

The human high mobility group box 1 (HMGB1) protein is a non-specific DNA binding protein, and has been well known to selectively bind to cisplatin crosslinked double–stranded DNA in both cell free media^18^ and inside cells.^19^ The box a domain of HMGB1 was shown to have a higher affinity than box b to bind to 1,2-cisplatin crosslinked DNA.^13^ Thus, we functionalized Nickle-based magnetic microbeads by HMGB1a to assemble affinity microprobes for capturing cisplatin crosslinked DNA from genomic DNA fragments extracted from the A549 human non-small cell lung cancer cells treated with cisplatin. As HMGB1a can also bind to bend non-platinated DNA^3^, we also extracted and fragmentated genomic DNA from A549 cells without exposure to cisplatin as control samples for DNA sequencing. The ICP-MS analysis showed that platination rate of genomic DNA extracted from cisplatin treated A549 cells reached 0.034‰ (Supplementary Figure 3A), being in consistent with our previous report, where the platination rate of DNA in MCF7 cells was measured to be 0.064‰.^20^ This DNA platination rate is much lower than that determined by *in vitro* reaction of synthetic DNA with cisplatin^21^ as the in vitro reaction was free from the competition of intracellular proteins binding to cisplatin and DNA. Furthermore, the gel-electrophoresis results demonstrated that the size of the DNA fragments gained from both the positive sample and control sample ranged from 1000 to 250 pb, but the larger DNA fragments obtained from the cisplatin treated cells were less than those from the untreated cells (Supplementary Figure 3B). This is likely ascribed to the platination which made genomic DNA more fragile.^22^ As shown in Supplementary Figure 4, the absorbance at 260 nm of the DNA fragments from both positive sample and control sample significantly decreased subject to incubation with the affinity microprobes (Supplementary Figure 4). Since HMGB1a can bind to both non-platinated and platinated DNA, though the affinity is different, it is not surprising that the same amount of the microprobes captured similar amount of DNA fragments from both positive sample and control sample (Supplementary Table. 2). However, this did not mean that the captured DNA fragments originated from the same genes and have the same sequence. In fact, the subsequent sequencing revealed that there were significant difference between the genes mapped by sequencing the DNA fragments captured from the positive and control samples.

### Gene Sequencing

Firstly, We constructed DNA libraries based on the captured DNA fragments extracted from each positive and control sample. The results showed that the DNA sequences in DNA libraries of the positive samples have similar sizes to those of DNA libraries based on the control samples (Supplementary Figure 5). This means that the inhibition of cisplatin binding on DNA synthesis^23^ has been circumvented when we increased PCR extension time, which minimized the risk of false negative results of gene mapping.

Taking the protein kinase cGMP-dependent 1 (*PRKG1*) gene as a example (Figure 2A), we identified a cisplatin damaged gene by comparing the total read of all peaks mapped to a gene in the positive sample with that of all peaks mapped to the gene in the corresponding control sample. The enrichment fold (EF_P_) of a peak was calculated by using the equation 1.

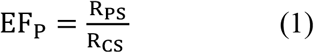

**Figure 2.**
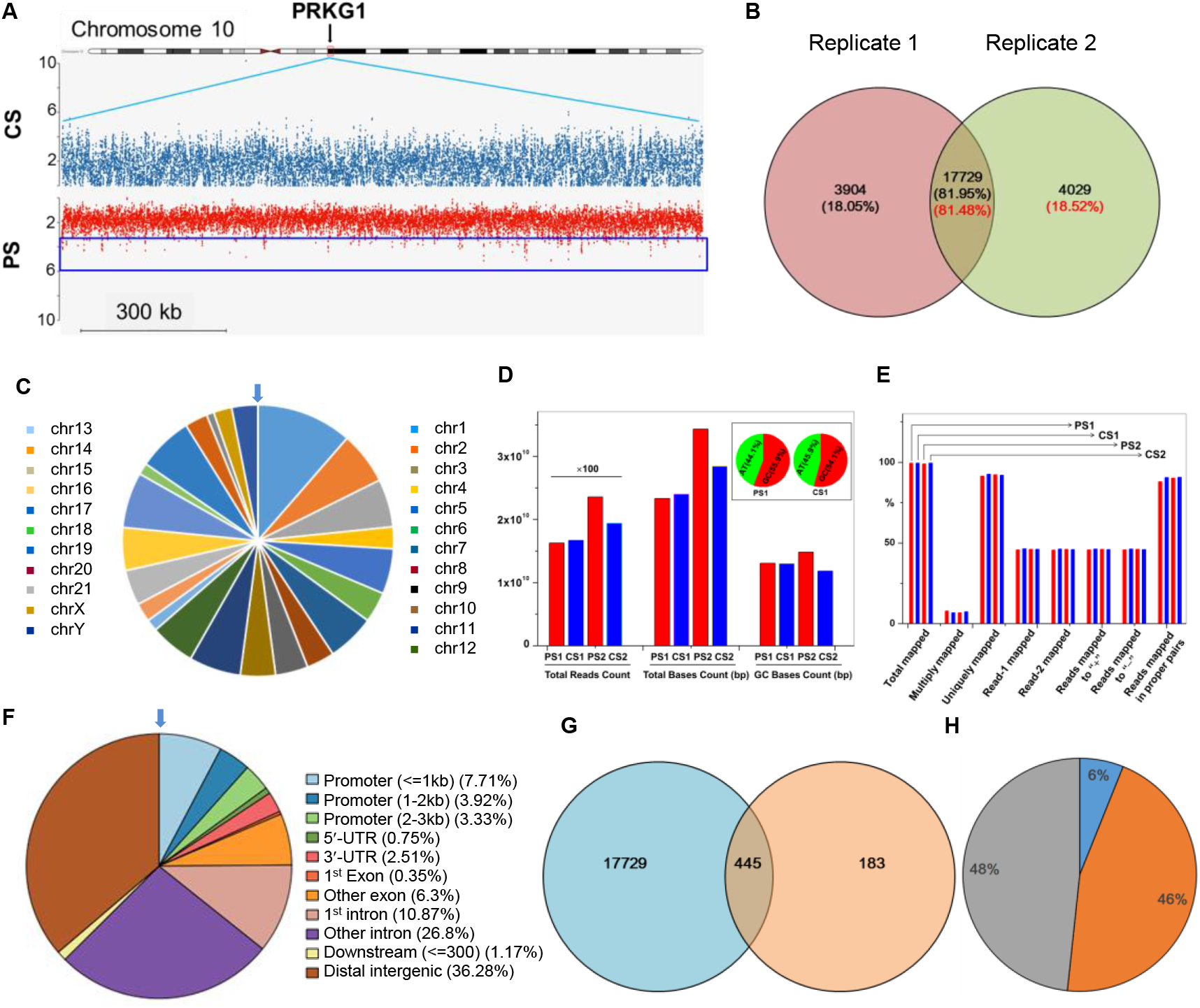
Mapping of cisplatin damaged genes. (A) Schematic diagram of mapped peaks for *PRKG1* gene in a pair of positive/control samples. (B) Venn Diagram of total cisplatin damaged genes mapped in two independent screenings with an enrichment fold of ≥2. (C) Distribution of cisplatin damaged genes over chromosomes. (D) Total reads, total bases and GC bases mapped, and (E) mapping rates counted in various ways of positive samples and control samples. (F) Distribution of cisplatin damaged DNA fragments in genetic regions. (G) Venn diagram of cisplatin damaged genes (navy) mapped in this work and the prior gene targets (light brown) for cancer treatment screened by the reference^25^. (H) The classification of commonly gene targets (445) for cancer treatments identified in this work and reference 25. Dark blue represents the genes which are targets of clinically used drugs or candidates in preclinical development, orange those which are not targets of drugs in clinical use but have evidences that support drugability, and grey those which have had no evidence or information supporting drugability.

Where R_PS_ and R_CS_ are the number of reads (or tags) mapped for a peak in a pair of positive sample (PS) and control sample (CS), respectively. It is worthy to point out that a peak was considered unreliable and then abandoned if the peak matched multiple genes, and that a peak with an enrichment fold (EF_p_) <1.5 was also taken as unreliable and then discarded.^24^ Thus, the enrichment fold of a mapped gene was the sum of enrichment folds of all peaks mapped to the gene, i.e.

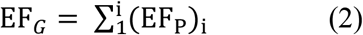

Where i represents the number of peaks mapped for a gene in a positive sample/control sample pair. Using equations 1 and 2, the enrichment fold of *PRKG1* was calculated to be 152, implicating that this gene was heavily damaged by cisplatin.

Using this method, we mapped 17729 common genes with a EF_G_ of ≥2 in two independent experiments (Figure 2B, Supplementary Table 5) by matched all DNA fragments in a positive/control sample pair with the human reference genome (hg38). They distributed over all 23 pairs of chromosomes, though various proportions can be seen from one to another (Figure 2C). The results showed that the total mapping rates of both positive samples and control samples achieved 99%, and uniquely mapping rates >91% (Figure 2D,2E, Supplementary Tables 3 and 4). As cisplatin prefers to crosslinking -GG- sites of DNA, the GC base pair count of the positive samples accounted for a higher ratio to the total base count than that of the control samples (Figure 2E, Supplementary Table 3). As a consequence, the mapping rates of total reads in proper base pairs of positive samples were slightly lower than those of control samples (Figure 2E, Supplementary Table 4). Moreover, we found that cisplatin mainly attacked the distal intergenic and intron regions, including the 1^st^ intron and other intron regions (Figure 2F). This is mainly ascribed to the relative large size of these genomic regions compared with others. However, given that the promoter and terminator sequences are generally short, they actually account for a quite higher platination proportion in cisplatin damaged genes (Figure 2F), being consistent with previous findings.^13^

Taking it into account that cisplatin attacks DNA to kill cells, one may expect that more cisplatin damaged genes could be detected in the death cells of positive samples. However, our sequencing data showed that there was no significant difference between the total numbers and enrichment folds of cisplatin damaged genes mapped from the DNA fragments collected from living cells alone and from living cells and death cells together after 24 h of incubation with 50 μM cisplatin (Data not shown).

Searching for potential gene targets for cancer therapy is always the most concern of drug discovery. Recently, Behan et. al. utilized the well-established CRISPR-Cas9 technique to screen the prior therapeutic gene targets for cancer treatment.^25^ They prioritized 628 gene targets for development of anticancer drugs, of which 445 (70.86%) were identified as cisplatin damaged genes in this work (Figure 2G, Supplementary Table 6), including 27 genes which are the targets of anticancer drugs in clinical use or candidates in preclinical research. For examples, *IGF1R*, *MTOR* and *ATR* for colorectal cancer treatment were identified as cisplatin damaged genes with an EF_G_ of 158, 66 and 40, respectively, and the EF_G_ values of *PIK3CB*, *ERBB2*/*B3*, *ESR1* and *TYMS* for breast cancer treatment were 101, 48/17, 27 and 6, respectively.^25^ The *WRN* gene, which expresses a RecQ DNA helicase, has been shown to be a promising drug target for cancers with microsatellite instability,^25,26^ and identified in this work as one of cisplatin targets with an EF_G_ of 32. At this term, cisplatin is really a multi-targeted chemotherapeutic drug.

### Bioinformatics Analysis

We applied Ingenuity Pathway Analysis (IPA) program to decipher the association and function of the cisplatin damaged genes mapped and described above. We input the commonly identified 17729 cisplatin damaged genes in two independent experiments with an EF_G_ of ≥ 2 into IPA data pool, and matched 14012 genes to human genome, including 2263 enzymes, 1308 transcription regulators, 768 transporters and 616 kinases (Figure 3A). Among the kinase genes, 427 are protein kinases, which account for 82% of putative protein kinase genes in human genome (Supplementary Table 7),^27^ suggesting that cisplatin appears to be a general protein kinase inhibitor.

**Figure 3.**
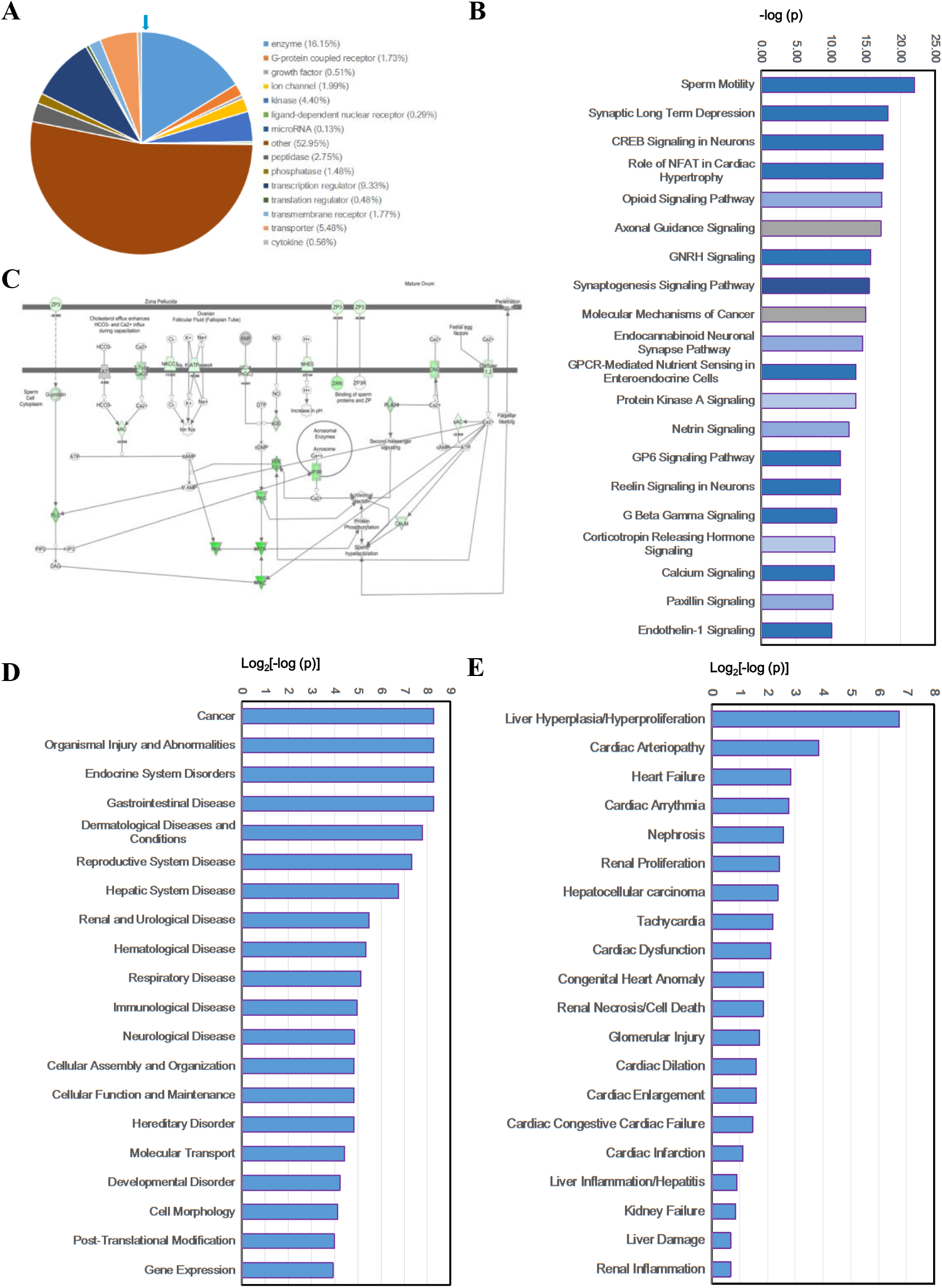
Bioinformatics analysis of cisplatin damaged genes. (A) Biological function classification of cisplatin damaged genes. (B) The core signaling pathways with which the cisplatin damaged genes are associated. The blue bar indicates that the signaling pathway is inhibited due to the gene damages. The deeper the blue color is, the stronger the inhibition is. The grey bar indicates that IPA cannot predict whether the core signaling pathway is activated or inhibited due to the gene damages. (C) The schematic diagram of the sperm motility signaling pathway with which the cisplatin damaged genes are the most associated. The green color indicates the gene (or gene group) was (were) damaged by cisplatin. The deeper the green color is, the higher the enrichment fold of genes mapped was. The number below the gene name represents the enrichment fold, the “−” indicates that the gene is damaged or its expression is inhibited. A more detailed schematic diagram of this signaling pathway is provided as Supplementary Figure 6. (D) The diseases and functions to which the cisplatin damaged genes are related. (E) The toxicity and functions to which the cisplatin damaged genes are associated. The *P* values represent associativity. The smaller the *P* values are, the more close the damaged genes are related to the pathway, disease, function or toxicity.

Limited by the data capacity of IPA, however, only 7578 genes, of which the EF_G_ is >12, could be analyzed further. The results showed that the 7578 cisplatin damaged genes are associated with 302 core signaling pathways with an association *P* value of < 0.05 (Supplementary Table 8). With a *P* value of 1.0×10^−22^, the most associated core signaling pathway of the 7578 genes is sperm motility, which was greatly inhibited with an inhibition index of −8.2 (*z*-score) (Figure 3B, Supplementary Table 8). Among the 7578 damaged genes, 146 genes are involved in the sperm motility pathway, which account for 68.5% of all genes in this signaling pathway (Figure 3C, Supplementary Table 9). Moreover, 87 of the 146 genes are members of kinase family, including *PKG*, *PKA*, *PTK* and *PKC* subfamilies (Supplementary Table 9 and Supplementary Figure 6). These results implicates that to a large extent cisplatin inhibits the sperm motility signaling by disturbing the kinase signaling. Indeed, we found that cisplatin damaged genes are also closely associated to protein kinase A signaling with an association *P* value of 2.5×10^−14^ (Figure 3B). As shown in Supplementary Figure 7 and Supplementary Table 10, 209 cisplatin damaged genes are associated with the protein kinase A signaling, including 38 kinases, 48 phosphatases and 17 transcription regulators, which account for 54.4% of all genes involved in this signaling pathway. As cisplatin damaged both kinases and phosphatases, e.g. protein kinase AMP-activated non-catalytic subunit gamma (*PRKAG2*), protein kinase C epsilon (*PRKCE*), protein tyrosine phosphatase receptor type D (*PTPRD*), protein tyrosine phosphatase receptor type T (*PTPRT*) of which the EF_G_ values were all >210 (Supplementary Table 5), IPA predicts that the net inhibition index (z-score) of cisplatin damages on the protein kinase A signaling is only −2.7 (Figure 3B).

It is notable that *PRKCA* gene is involved in both sperm motility and protein kinase A signaling pathways, and was heavily damaged by cisplatin with an EF_G_ of 180 (Supplementary Figures 6 and 7, Supplementary Table 5). Thus, we further analyzed its association with biological functions and diseases as well as its related up/downstream genes and molecules. We found that *PRKCA* is covered in many diseases and functions, e.g. apoptosis,^28^ differentiation of chondrocytes,^29^ tumorigenesis,^30^ migration of mammary tumor cells^31^ and triple-negative breast cancer,^32^ of course phosphorylation of tyrosine^33^ and threonine^34^ (Supplementary Figure 8). This gene is regulated by endo/exogenous molecules, such as tetradecanoylphorbol,^35^ bryostatin,^36^ VEGFA^37^ and insulin,^38^ etc. and regulates downstream signaling transducers, e.g. NFkB,^39^ Akt,^40^ p38,^41^ ERK1/2 ^42^ and RAF1^43^ (Supplementary Figure 9). This implicates that cisplatin damage on *PRKCA* may play a crucial role in action of this drug, deserving further investigation.

Protein phosphorylation is the most common and important protein posttranslational modification. It participates in and regulates many life activities and processes in the body. Through the dynamic phosphorylation and dephosphorylation of proteins, many cellular processes such as signal transduction, gene expression and cell cycle are finely controlled. For example, the deficiency of a fully developed and functioning initial segment, which is the most proximal region of the epididymis, leads to male infertility. Many protein kinases, e.g. SFK (SRC proto-oncogene family kinases), ERK pathway (known as the RAF/MEK/ERK pathway) components, and AMPK (AMP-activated protein kinases) pathway components, are involved in the development of initial segment.^44^ This explains why the cisplatin damage on kinase genes like *PRKCA*, *PRKC1* and *PRKAG2*, is the most related to sperm motility signaling pathways. More importantly, as abnormal protein phosphorylation is usually related to cancer, protein kinases has been considered to be the major drug targets for cancer therapy of this century.^45,46^ A number of small molecular protein kinase inhibitors (PKIs), e.g. gefitinib,^4^ imatinib^47^ and sorafenib,^48^ have gained approval for treatments of various cancers in clinic. Given the close association of cisplatin damaged genes identified herein with kinase signaling pathways, cisplatin appears to kill cancer cells by a similar way to that PKIs do, i.e. disrupting protein phosphorylation. However, on the one hand, cisplatin irreversibly blocks protein phosphorylation by damaging protein kinase genes but PKIs action are reversible. On the other hand, cisplatin damages many kinase genes, even phosphatases, as shown in Supplementary Figures 6 and 7. This non-specification makes cisplatin a Janus, causing toxic side effects while it kills cancer cells.

Another core signaling pathway with which the cisplatin damaged genes are highly associated (*P*=10^−15^) is the molecular mechanisms of cancer signaling (Figure 3B), and 213 genes damaged by cisplatin involve in this signaling pathway, accounting for 55.5% of all genes in this pathway (Supplementary Figure 10 and Supplementary Table 11). However, due to the complexity of the mechanisms of cancer, IPA could not predict how these gene lesions regulate the development of cancers, the *z*-scores of these gene damages on this signaling pathway were not available (Figure 3B). Apparently, at this term the ultimate fate of cancer cells upon cisplatin treatment requires further investigations.

Apart from the signaling pathways described above, unexpectedly, other signaling pathways with which the cisplatin damaged genes are closely associated (*P*<10^−10^) mainly gather in the neural system, including synaptic term depression (*P*=6.3×10^−19^, Supplementary Figure 11), GREB signaling in neurons (*P*=3.2×10^−18^, Supplementary Figure 12), axonal guidance signaling (*P*=6.3×10^−18^), GNRH signaling (*P*=2.0×10^−16^), synaptogenesis signaling pathway (*P*=3.2×10^−16^), endocannabinoid neuronal synapse pathway (*P*=2.5×10^−15^) and netrin signaling (*P*=2.5×10^−13^) (Figure 3B, Supplementary Table 8). Among them, the synaptogenesis signaling pathway is the most inhibited by cisplatin damages on genes with a *z*-score of −12.0. Due to the Blood Brain Barriers, cisplatin has been demonstrated to accumulate at a low level in human brain,^49^ thus we did not analyze the impact of gene damages by cisplatin on the neural system further. However, it does not exclude the necessity to explore further whether and how the cisplatin damages on these neural signaling related genes account for its neurotoxicity, such as peripheral sensory neuropathy and central nervous side effects, which is one of major toxic side effects of cisplatin in clinic use.^15,50^

Additionally, IPA revealed that the cisplatin damaged genes are highly related to the role of NFAT in cardiac hypertrophy signaling pathway with an association *P* value of 3.2×10^−18^ (Figure 3B). The 137 genes damaged by cisplatin account for 64.9% of all 211 genes involved in this signaling pathway (Supplementary Table 8). The nuclear factor of activated T cells (NFAT) as a transcription factor plays an important role in cardiac hypertrophic signaling which is mediated by the calcium-activated phosphatase calcineurin^51^ (Supplementary Figure 13). This indicates that the calcineurin-NFAT signaling is dominated by calcium signaling, with which the cisplatin damaged genes is also highly associated with an association *P* value of 3.2×10^−11^ (Figure 3B), and 116 out of 198 genes involved in this signaling pathway were damaged by cisplatin (Supplementary Figure 14 and Supplementary Table 8). The calcium signaling pathway is predicted to be strongly inhibited by cisplatin damages on these genes with a *z*-score of −8.5. We discovered that the calcium channel genes *CACNA1A/B/C* and *CACNA2D3* were severely damaged by cisplatin with an enrichment fold of 253, 179, 194 and 223, respectively (Supplementary Figure 14 and Supplementary Table 5). These implicate that gene damages imposed by cisplatin may lead to cardiac hypertrophy *via* disturbing calcium homeostasis.

The further IPA analysis unraveled that the most associated disease of the 7578 cisplatin damaged genes is cancer with an association *P* value of 1×10^−312^ (Figure 3D, Supplementary Table 12 and Supplementary Figure 15A). The four cancer-related diseases or functions, i.e. cancer (*P*=8.8×10^−154^), cancer of cells (*P*=2.27×10^−74^), cell transformation (*P*=9.5×10^−9^) and neoplasia of cells (*P*=5.4×10^−81^), are significantly inhibited by cisplatin damages on genes with a *z*-score of −3.15, −2.10, −3.56 and −2.48, respectively (Supplementary Figure 15B and Supplementary Table 13). It was notable that the *SPAG9* gene is involved in 147 of 361 cancers or functions with which the cisplatin damaged genes are highly associated (Supplementary Table 13). *SPAG9* abundantly expresses C-jun-amino-terminal kinase-interacting protein 4 (JIP-4) in testicular haploid germ cells, and is essential for the development, migration and invasion of cancer.^52^ JIP-4 is a scaffolding protein that knits mitogen-activated protein kinases (MAPKs) and their transcription factors to activate specific signaling pathways^53^. This implicates that cisplatin damage on *SPAG9* (EF_g_ = 61) may plays a crucial role in action of cisplatin as an anticancer drug.

With regard to this, we further analyzed the diseases and functions to which *SPAG9* gene is related. As shown in Supplementary Figure 16, *SPAG9* (JIP-4) activates JUN/MAP kinase signaling,^53,54^ and is associated with various types of cancers, e.g. anaplastic thyroid carcinoma,^55^ hairy cell leukemia^56^ and intestinal gastric adenocarcinoma^57^. It also plays a role in several important biological processes, such as fertility,^58^ differentiation of skeletal muscle cells,^59^ differentiation of neurons^60^. Moreover, *SPAG9* gene is regulated by ligand-dependent nuclear receptor ESR1,^61^ transmembrane receptor CAV1,^62^ transcription factor ZNE217^63^ and transporter BSG,^64^ etc. and regulate mainly ERK, JNK and P38 MAPK signaling^65^ (Supplementary Figure 17). MAPK complex consists of several highly conserved threonine/serine protein kinases, and its phosphorylation eventually lead to the activation of Jun N-terminal kinase (JNK), extracellular signal regulated kinase (ERK) and p38, which in turn induces apoptosis in human malignant testicular germ cell lines.^66–68^ Therefore, the damage on *SPAG9* by cisplatin may play an indispensable role in cisplatin-induced apoptosis in testicular cancer cells. Given that *SPAG9* specifically expresses JIP-4 in high abundance in testicular haploid germ cells, the cisplatin damage on it may account for the high cure rate of this drug to testicular cancer, also deserving further studies.

Other diseases or functions with which cisplatin damaged genes mainly include organismal injury and abnormalities, endocrine system disorders, gastrointestinal disease, and dermatological diseases and conditions, and etc. with an association *P* values of < 10^−222^. Again, due to the complexity of diseases, the IPA analysis cannot judge how the cisplatin damaged genes intervenes the disease development, which apparently request further experimental researches.

Tissue toxicity is one of major elements limiting clinic application of cisplatin. However, the molecular basis for its toxicity has remained unclear. We performed IPA analysis on the association of cisplatin damages on genomic DNA with its toxicity. The results demonstrated that the most related tissue toxicity is liver hyperplasia or hyperproliferation with an association P value of 6.80×10^−108^, and then cardiac arteriopathy, heart failure, cardiac arrhythmia and nephrosis with an P value of 4.83×10^−15^, 6.59×10^−8^, 1.76×10^−7^ and 1.12×10^−6^, respectively (Figure 3E, Supplementary Figure 18 and Supplementary Table 14. As many genes damaged by cisplatin are involved in a type of tissue toxicity, for example 3351 damaged genes are shown to be associated with the liver toxicity, the IPA analysis cannot judge whether the cisplatin lesions on these genes inhibit or activate hyperplasia/hyperproliferation to cause the side toxicity (Supplementary Figure 18A). In some cases, for example nephrotoxicity, IPA predicted that cisplatin may inhibit renal necrosis or cell death *via* damaging 156 genes involved in this process (Supplementary Figure 14B and Supplementary Table 15), but activate renal proliferation by attacking 96 genes related to this function (Supplementary Figure 14C and Supplementary Table 15). These results are not in well agreement with the clinic manifestations of the toxic side of cisplatin.^15^ With regard to this, the IPA results can only correlate the cisplatin gene damages with diseases, toxicity and biological functions, but not allow precisely prediction on the phenotypes of gene damages. To address these issues, further experimental researches are in great demand.

### Western Blot assays

In order to verify the genome-wide mapping results of cisplatin damaged genes, we performed traditional Western Blot assays to characterize the gene expression level of 20 functionally important genes with and without cisplatin exposure. Among the 20 genes, 18 including 13 protein kinase genes are closely associated with the sperm motility signaling pathway. Other two genes, *SPAG9* and *AKT3*, are the upstream mediator and middle transducer of MAPK signaling pathway, respectively (Supplementary Table 13 and Supplementary Figures 10 and 13).

Since these chosen genes express in different levels in different types of cells, we selected four human cancer cell lines, A549 (non-small cell lung cancer cell line), NCCIT (testicular cancer cell line), HepG2 (hepatocellular cancer cell line) and HeLa (cervical cancer cell line) to perform Western Blot assays. Among the four cell lines, NCCIT is the most sensitive to cisplatin with a IC_50_ value of 5.8 μM (24 h treatment). The IC_50_ of HeLa, A549 and HepG2 cells is 8.2 μM (24 h), 10.1 μM (48 h), 22.4 μM (24 h)^69^, seperately. Thus, we applied different dosage of cisplatin to treat the cell lines, i.e. 50 μM for A549 and HepG2, 25 μM for HeLa and 12 μM for NCCIT. As shown in Figure 4, the expression levels of all selected genes decreased subject to cisplatin treatment. The change folds (CFs) in expression levels of 12 genes, including 8 protein kinase genes and *SPAG9*, was significantly (*p* < 0.05) or very significantly (*p* < 0.01). These indicate that the genes were indeed damaged by cisplatin, strongly supporting the reliability and credibility of our gene mapping results.

**Figure 4.**
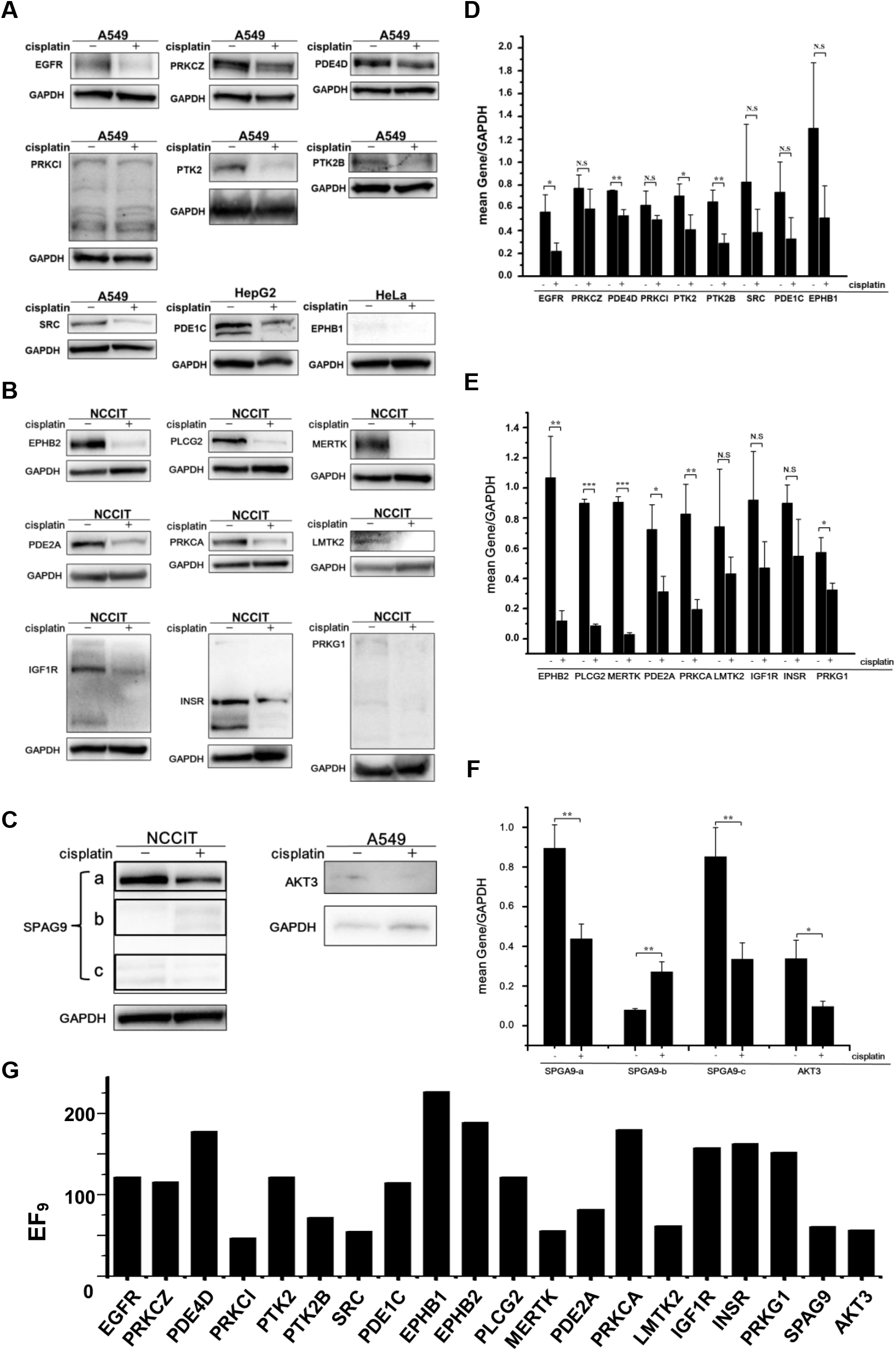
Western Blot assays. (A - C) Western Blot images of various proteins expressed in A549, HepG2, HeLa NCCIT cells with (+) and without (−) cisplatin treatment. GAPDH was used as the internal reference. (D – F) Changes in the expression levels of the proteins with and without cisplatin treatment. (G) The enrichment fold (EF_g_) of 20 genes damaged by cisplatin.

We noticed that there was no a simple correlation between the enrichment fold and change fold in the expression level of a specific gene. One of possible reasons is that the expression level of a selected gene is too low to be characterized, e.g. *LMTK2* in NCCIT. Another possible explanation for this phenomenon is that the gene expression is controlled by a complicated and sophisticated cellular system, including DNA replication and transcription machinery, and nuclear repair machinery, such that any alteration on this system could influence the expression of a specific gene addition to the direct damage imposed by cisplatin on the gene.

Interestingly, we found that the full length protein (JIP-4) product of *SPAG9* significantly decreased by 2.0-folds in abundance, but the two subunits visualized in band b of Figure 4D pronouncedly increased in intensity in NCCIT cells upon cisplatin treatment. This implies that the cisplatin lesion occurred more likely in the linkage region of *SPAG9* than in the core structural region of the subunits. As mentioned earlier, the cisplatin damage on *SPAG9* may account for the high sensitivity of testicular cancer to cisplatin, *SPAG9* may be a potential target for precise therapy of testicular cancer.

It is noted that the expression level of *MERTK* was the most strongly inhibited (CF = 32) by cisplatin damage on this gene (Figure 4B, E). The MERTK receptor tyrosine kinase phosphorylates Akt1-Y26, mediating Akt activation and survival signaling, which in turn drives oncogenesis and therapeutic resistance.^70^ Thus, cisplatin damages on *MERTK* gene may be implicated in mechanism of action of the drug. This result also support that MERTK is a potential therapeutic target.^70,71^

In addition to the phosphorylation-related genes mentioned above, another phosphorylation-related gene, *AKT3*, has also been identified as cisplatin gene target with an EG_g_ of 57 (Figure 4G). The protein encoded by this gene is a member of the AKT (or PKB) serine/threonine protein kinase family. As a middle-stream regulator of IGF1 signaling Akt3 plays a key role in regulating cell survival, insulin signaling, angiogenesis and tumor formation. The cisplatin damage to this gene reduced its expression by 3.5-folds (Figure 4C, F), which will trigger a series of downstream changes, as a consequence inducing apoptosis and death in cancer cells.^72^ Further research on *AKT3* will provide insights into better understanding the role of protein kinases in the mechanism of action of cisplatin.

## Conclusion

Combining affinity separation and high throughput sequencing, we mapped 17729 cisplatin damaged gene loci in human genome, which mainly function as enzymes, transcription regulators, transporters and kinases. Among them, 445 genes are prioritized gene targets for cancer therapy, implying that cisplatin is really a multi-targeting anticancer drug. Bioinformatic analysis demonstrated that 7578 genes damaged by cisplatin with a EF_G_ of >12 are the most closely related with the sperm motility signaling pathway, cancer diseases and hepatotoxicity with association *P* values of 1.0×10^−22^, 1×10^−312^ and 6.80×10^−108^, respectively. This provides novel insights into better understanding in the molecular mechanism of anticancer activity and toxicity of this drug. Cisplatin damaged protein kinase genes, 427 out of 518 putative human protein kinase genes, involved in all highly associated core signaling pathways, implicating that cisplatin may act as an irreversible protein kinase inhibitor to kill cancer cells, while it causes tissue toxicity like hepatotoxicity and nephrotoxicity. We selected 20 genes for Western Blot assays, the results showed that their expression level all reduced due to cisplatin damage. The expression of 8 protein kinases genes was significantly inhibited, that of *MERTK*, a receptor tyrosine kinase gene, was decreased by 32-fold subject to cisplatin damage. These intrigue further exploration in protein kinases as drugable targets for precise treatment of cancers. Interestingly, the kinase signaling mediating gene *SPAG9* was damaged by cisplatin, leading to 2-folds reduction in expression of full-length JIP-4 protein, but the subunits of JIP-4 increased in intensity, implicating that the cisplatin lesions mainly took place in linkage domain of the protein. Given *SPAG9* is involved in 147 out of 361 cancer diseases with which the cisplatin damaged genes are highly associated, and abundantly expresses JIP-4 in testicular haploid germ cells, this gene may account for the high sensitivity of testicular cancer to cisplatin, being a promising gene target for the precise treatment of testicular cancer.

## Acknowledgements

We thank the National Natural Science Foundation of China (Grant nos. 21575145, 21635008, 21790390 and 21790392) for support. Y.Z. also thanks the Youth Innovation Promotion Association of Chinese Academy of Sciences (grant no. 2017051).

