## Supplemental Tables and figures for "Genome-Wide Mapping of Cisplatin Damaged Gene Loci"

### Contents:

### Supplementary Chart 1

### Supplementary Figure 1 – 18

### Supplementary Table 1 - 16

ATGGGCAGCAGCCATCATCATCATCATCACAGCAGCGGCCTGGTGCCGCGC
GGCAGCCATATGGCTAGCATGACTGGTGGACAGCAAATGGGTCGCGGATCC
CCTCGTGGTAAAATGAGCAGCTATGCATTTTTTGTTCAGACCTGTCGTGAG
GAACACAAAAAAAAGCATCCTGATGCAAGCGTGAACTTTAGCGAATTCAG
CAAAAAATGTAGCGAACGCTGGAAAACCATGAGCGCAAAAGAAAAAGGC
AAATTTGAGGATATGGCCAAAGCAGATAAAGCCCGTTATGAACGTGAAATG
AAAACCTATTAA

**Supplementary Chart 1.** The cDNA sequence of human HMGB1 domain A (HMGB1a)

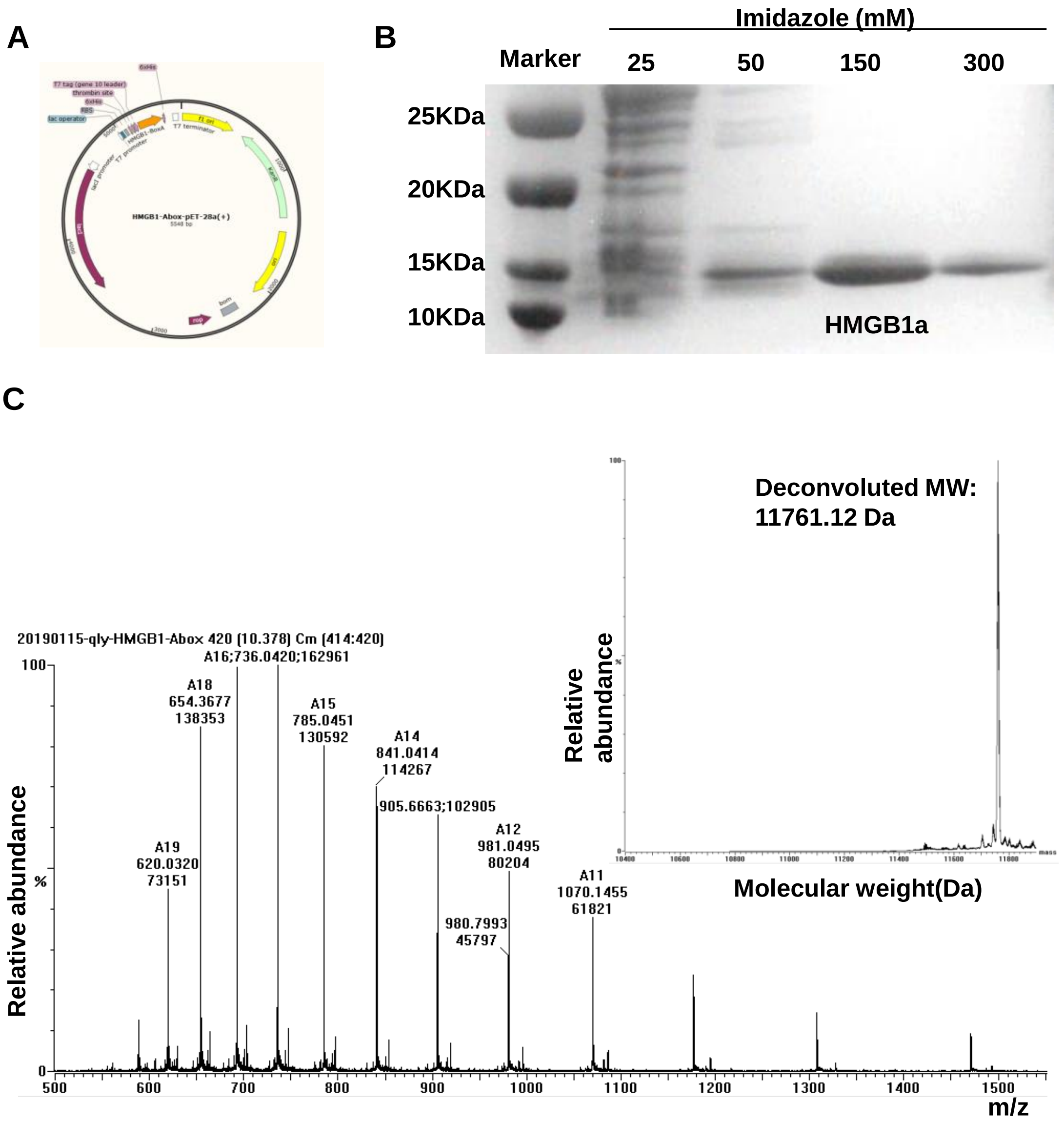

**Supplementary Figure 1.** Identification of HMGB1a. (A) Diagram of vector for HMGB1a expression in *E. coli*. (B) Coomassie-blue-stained gel of the wild type HMGB1a expressed in *E. coli*. (C) Mass spectrum of HMGB1a.

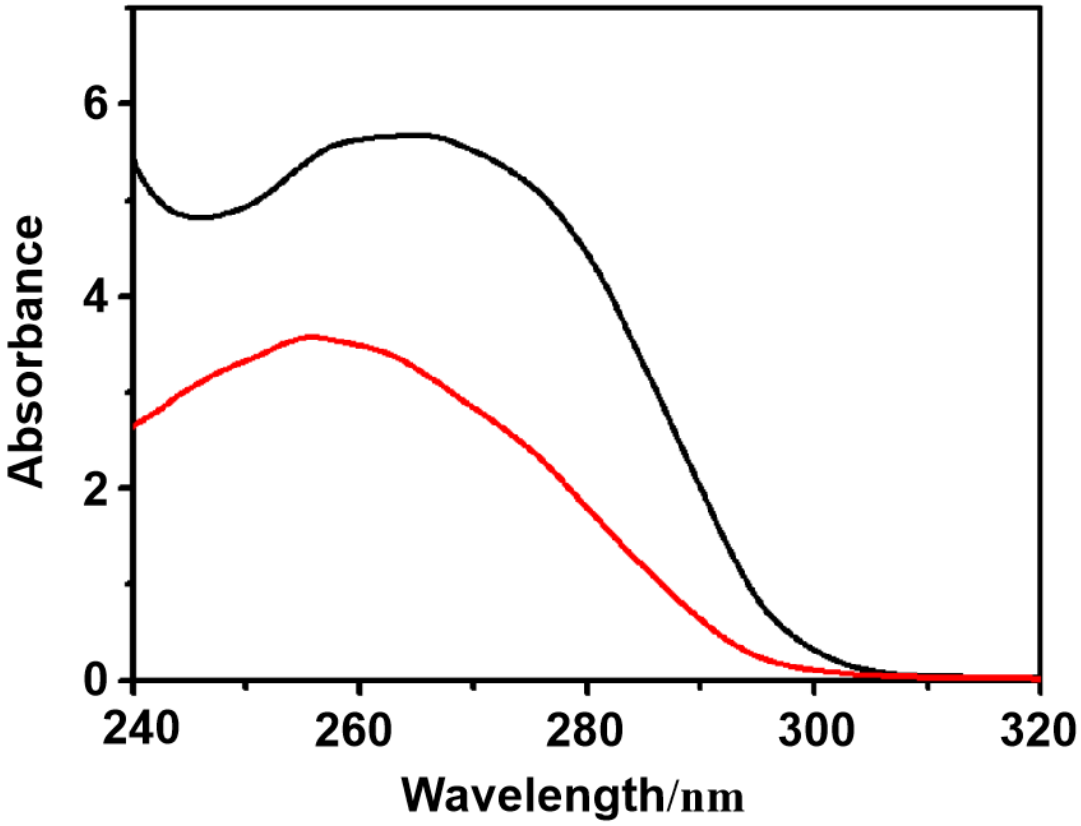

**Supplementary Figure 2.** UV absorption spectra of HMGB1a in the supernatant before (black) and after (red) incubated with nickel-modified magnetic beads suspended in PBS.

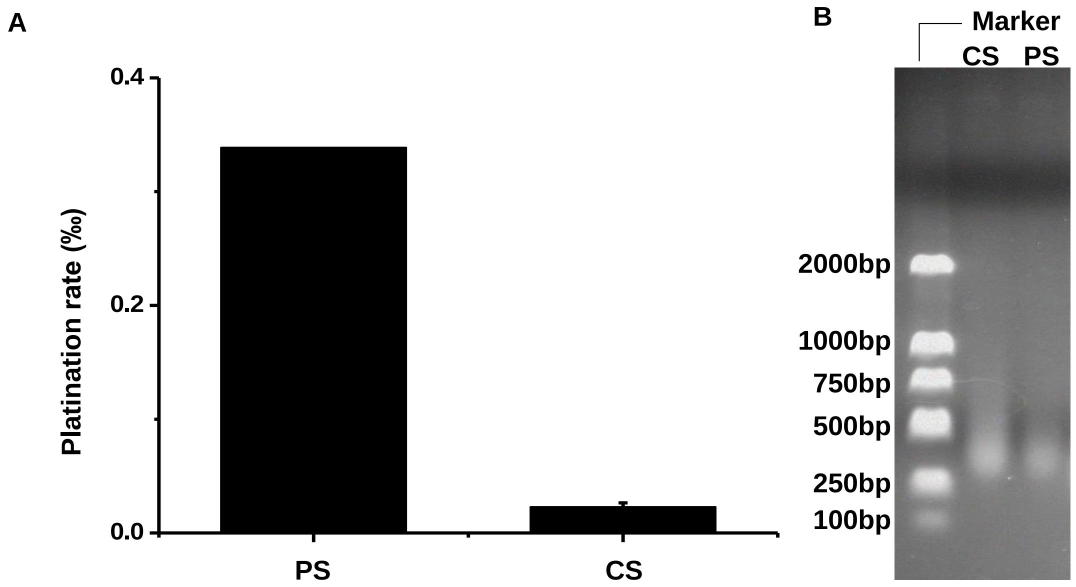

**Supplementary Figure 3.** Characterization of platinated/native DNA fragments and the capture of DNA fragments. (A) The platination rate of DNA fragments gained from positive sample (PS) and control sample (CS) determined by ICP-MS. (B) The size distribution of platinated/native DNA fragments characterized by 1% agarose gel electrophoresis.

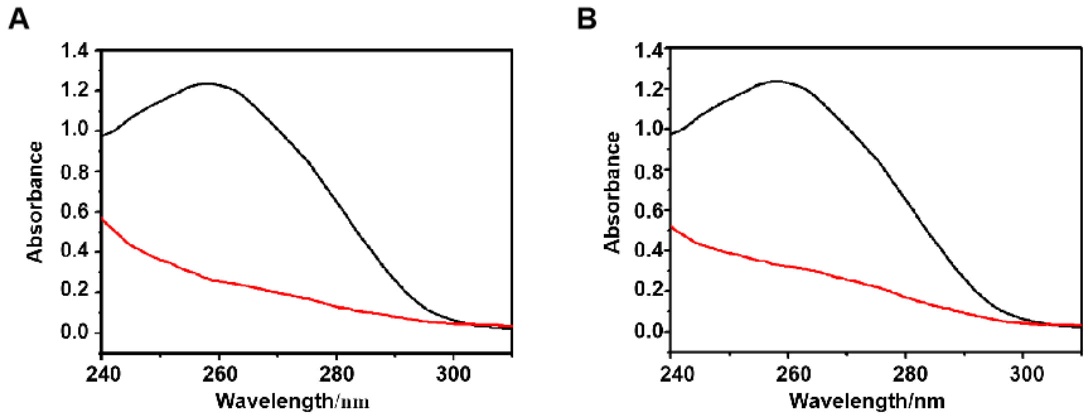

**Supplementary Figure 4.** UV absorption of the DNA fragments in the supernatant before (black) and after (red) incubation with HMGB1a-functionlized microprobes. (A) DNA fragments extracted from A549 cells treated with 50 μM cisplatin; (B) DNA fragments extracted from A549 without cisplatin treatment.

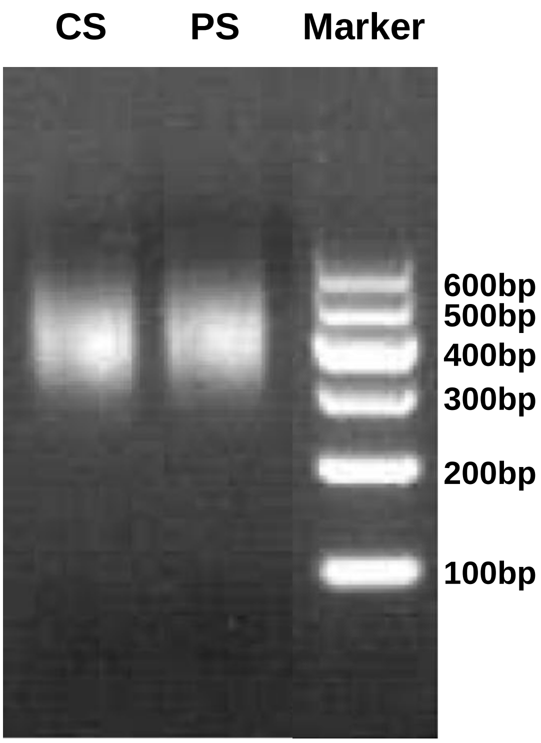

**Supplementary Figure 5**. DNA libraries constructed from captured DNA fragments.

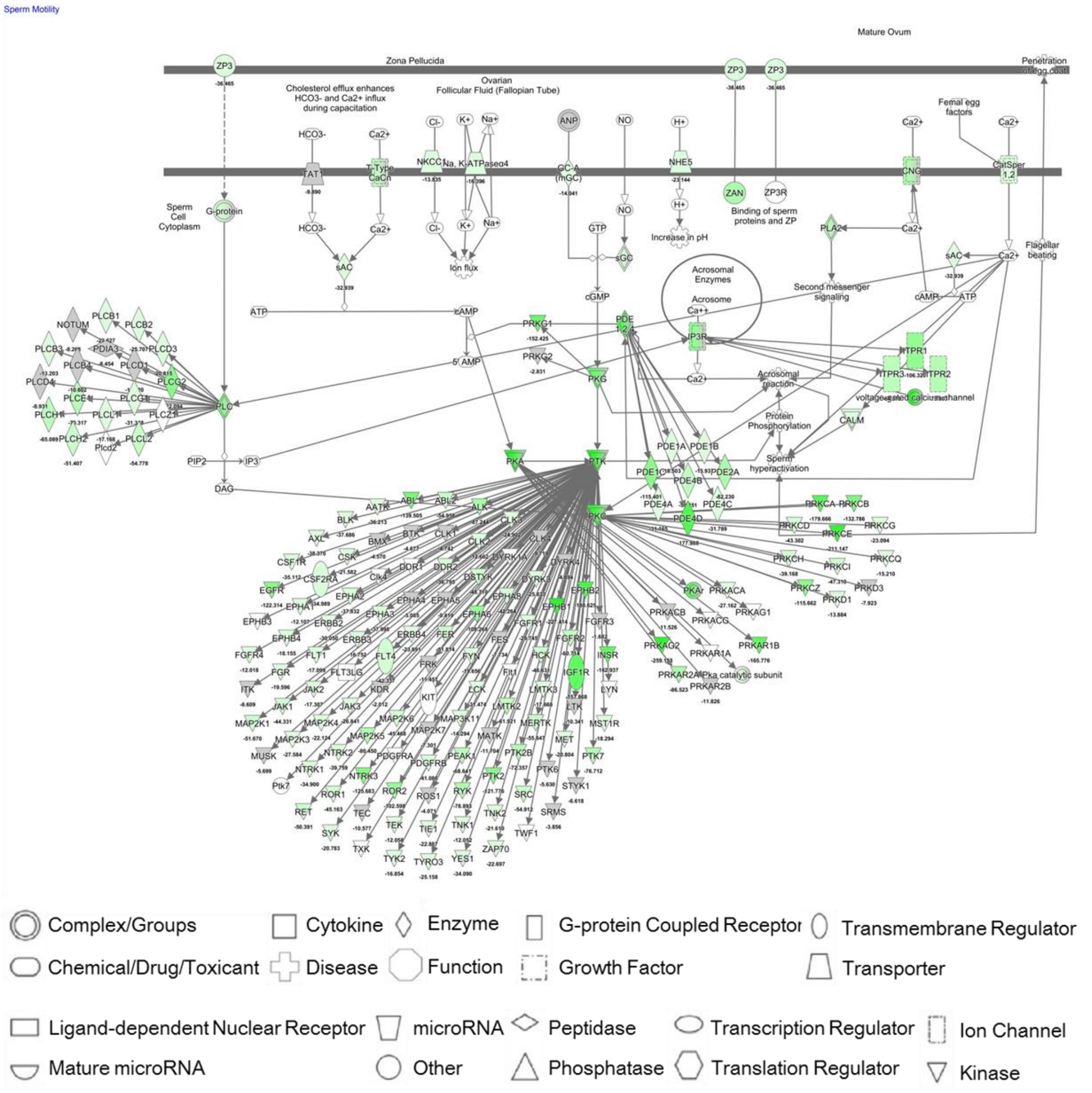

**Supplementary Figure 6.** The schematic diagram of sperm motility pathway with which the cisplatin damaged genes are the most associated. The green color indicates the gene (or gene group) was (were) damaged by cisplatin. The deeper the green color is, the higher the enrichment fold of genes mapped. The number below the gene name represents the enrichment fold, the minus symbol indicates that the gene is damaged or its expression is inhibited.

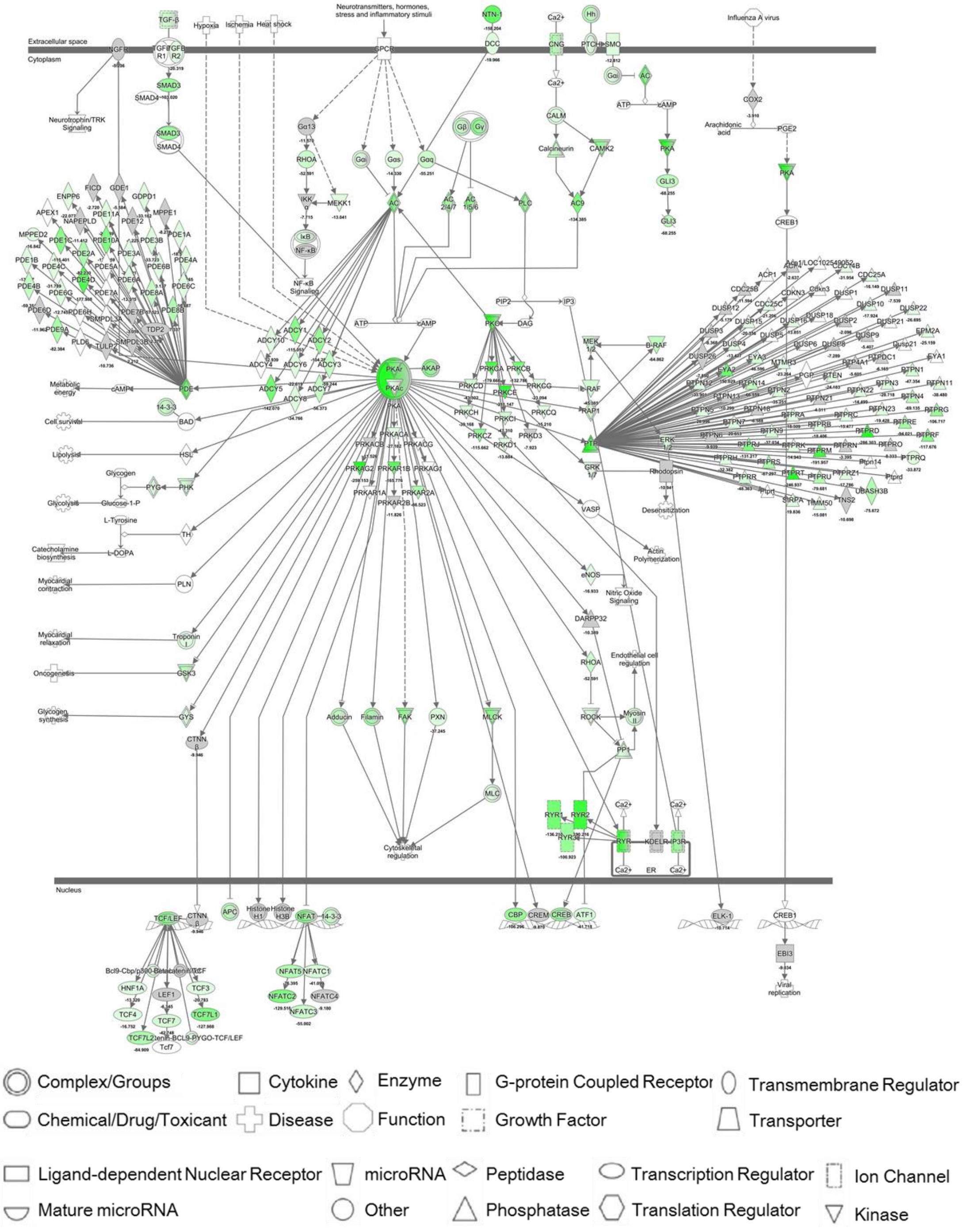

**Supplementary Figure 7.** Schematic diagram of protein kinase A signaling pathway with which the cisplatin damaged genes are highly associated. The green color indicates the gene (or gene group) was (were) damaged by cisplatin. The deeper the green color is, the higher the enrichment fold of genes mapped. The number below the gene name represents the enrichment fold, the minus symbol indicates that the gene is damaged or its expression is inhibited.

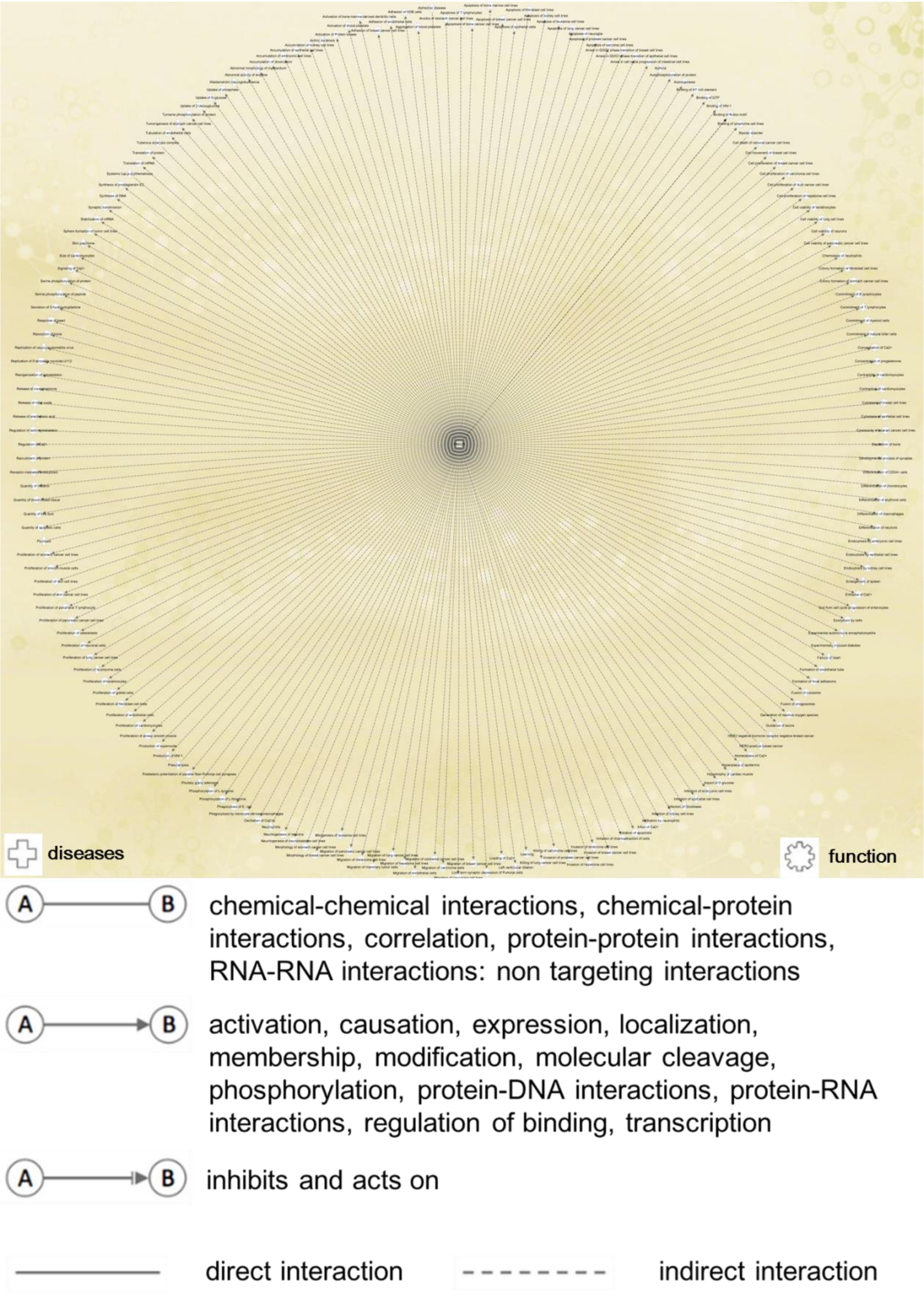

**Supplementary Figure 8.** The diseases and functions to which *PRKCA* is related.

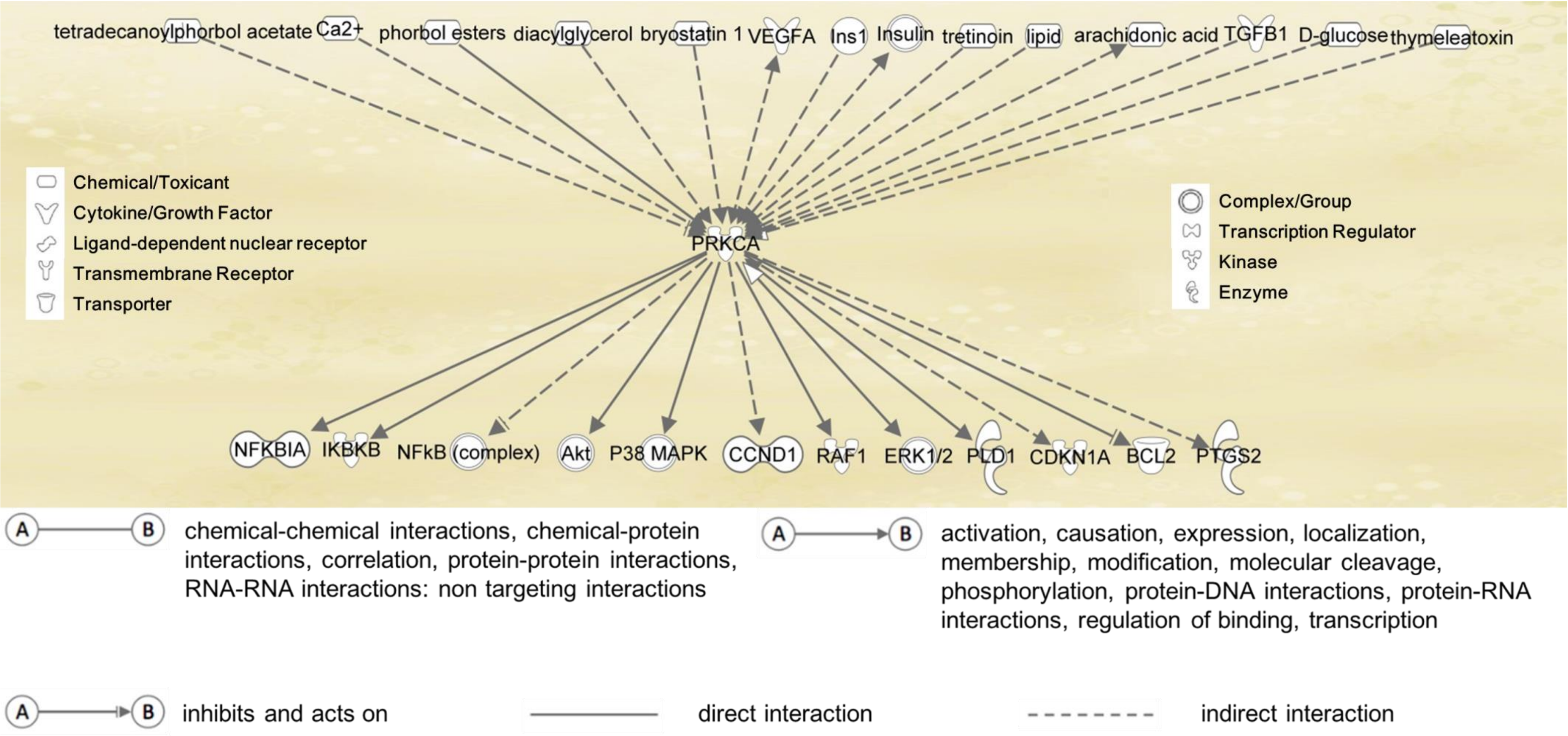

**Supplementary Figure 9.** The upstream and downstream genes or molecules which regulate *PRKCA* or are regulated by *PRKCA*.

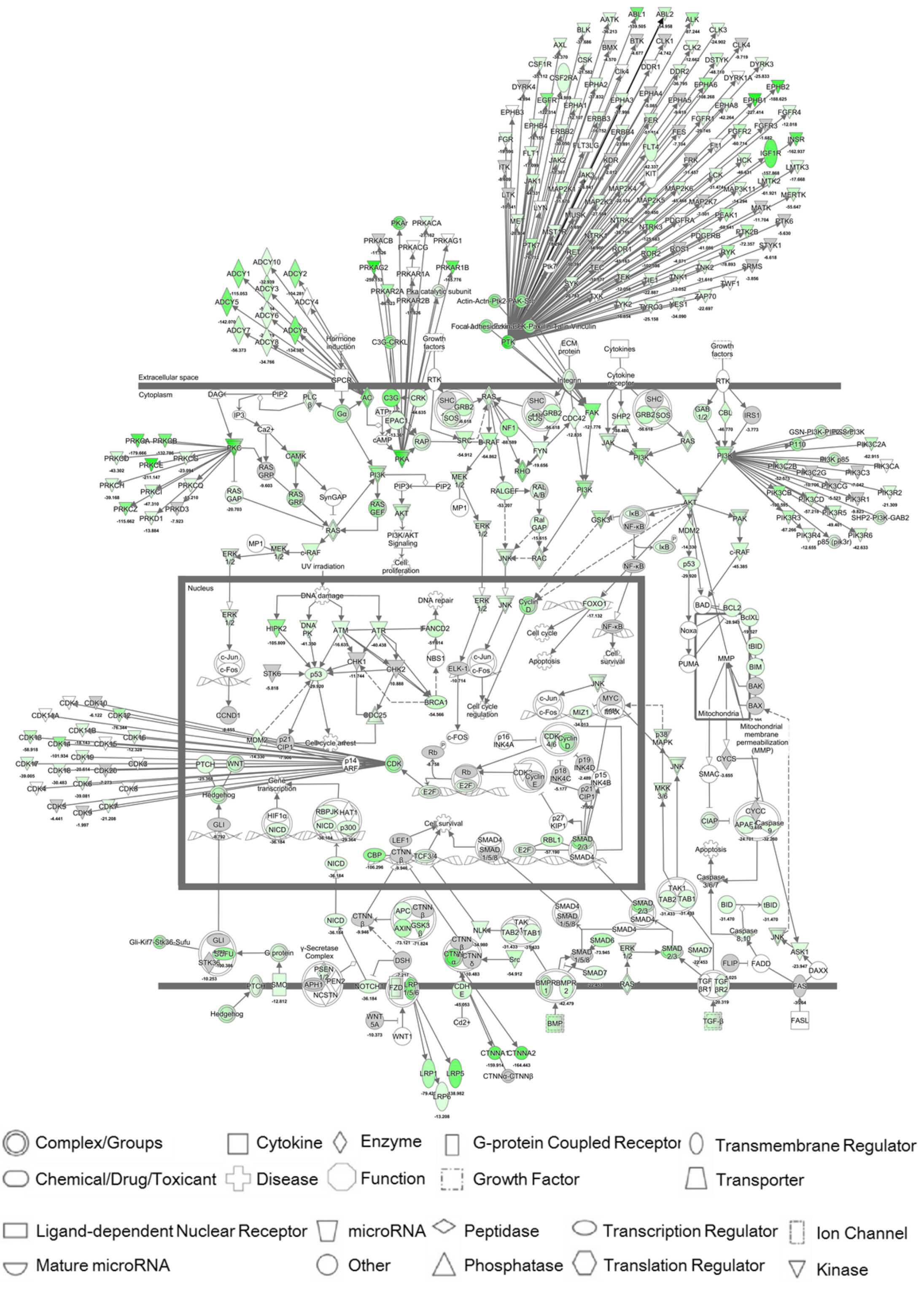

**Supplementary Figure 10.** Schematic diagram of molecular mechanism of cancer pathway with which the cisplatin damaged genes are highly associated. The green color indicates the gene (or gene group) was (were) damaged by cisplatin. The deeper the green color is, the higher the enrichment fold of genes mapped. The number below the gene name represents the enrichment fold, the minus symbol indicates that the gene is damaged or its expression is inhibited.

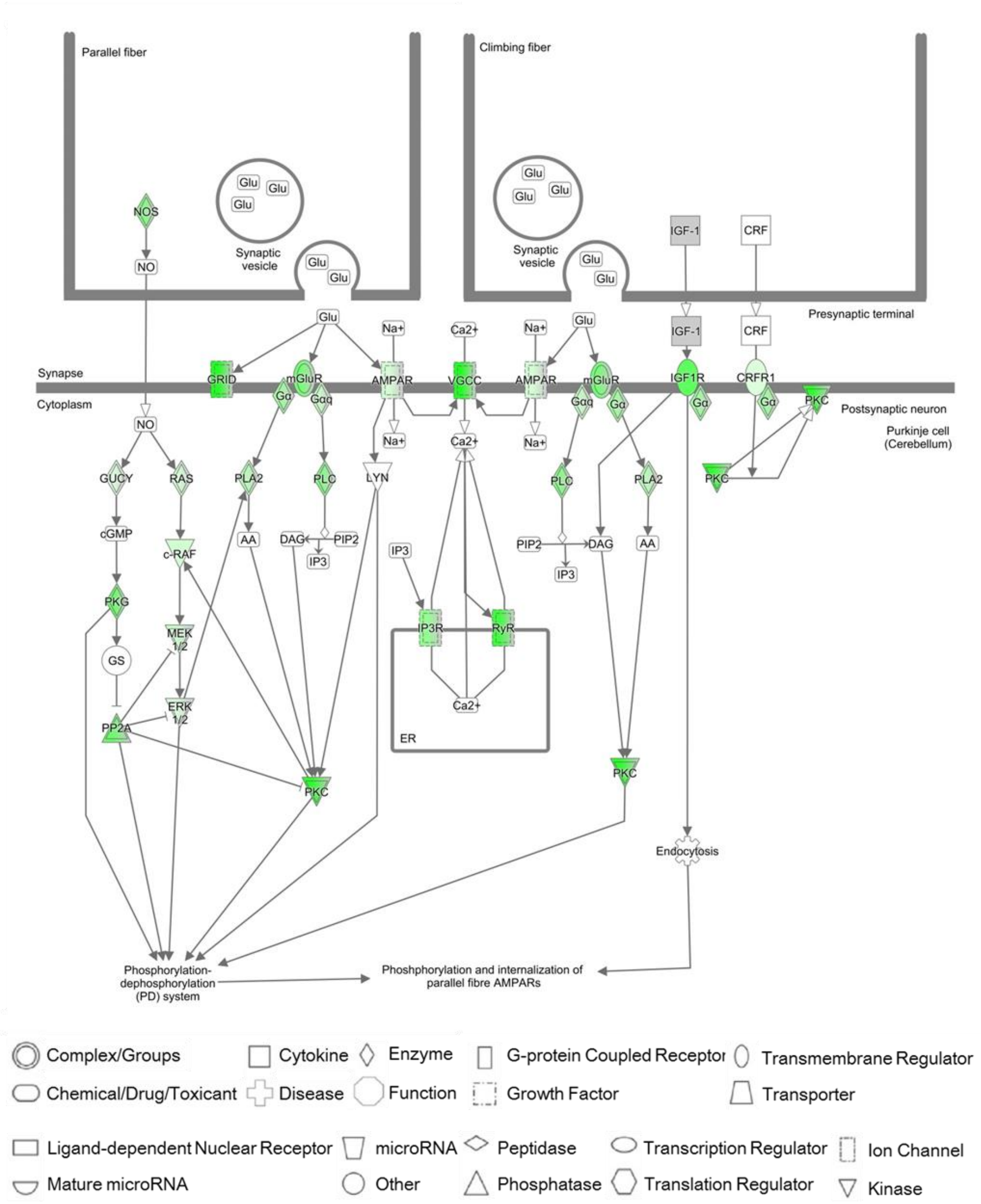

**Supplementary Figure 11**. Schematic diagram of synaptic long term depression pathway with which the cisplatin damaged genes are highly associated. The green color indicates the gene (or gene group) was (were) damaged by cisplatin. The deeper the green color is, the higher the enrichment fold of genes mapped.

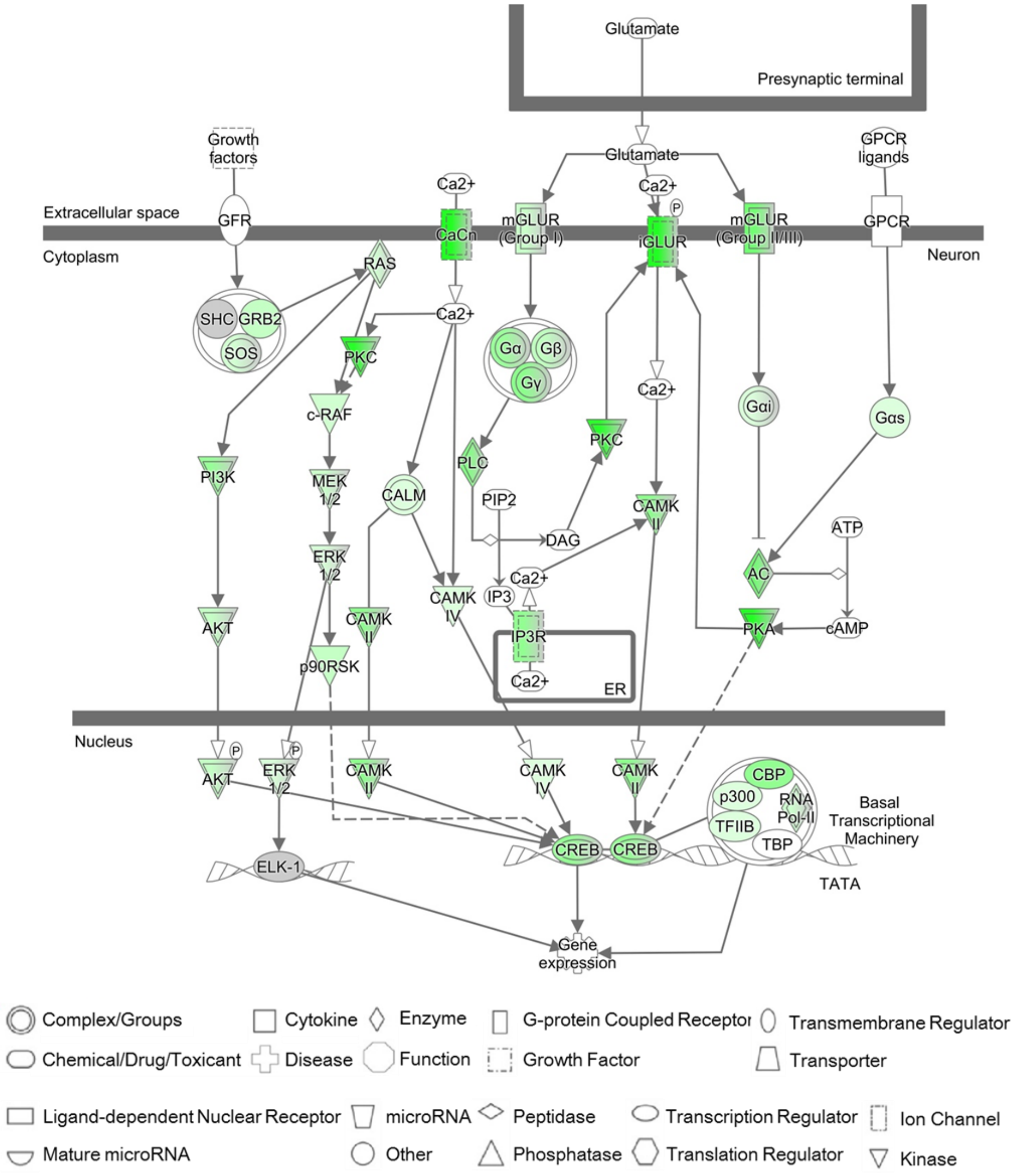

**Supplementary Figure 12**. Schematic diagram of CREB signaling in neurons pathway with which the cisplatin damaged genes are highly associated. The green color indicates the gene (or gene group) was (were) damaged by cisplatin. The deeper the green color is, the higher the enrichment fold of genes mapped.

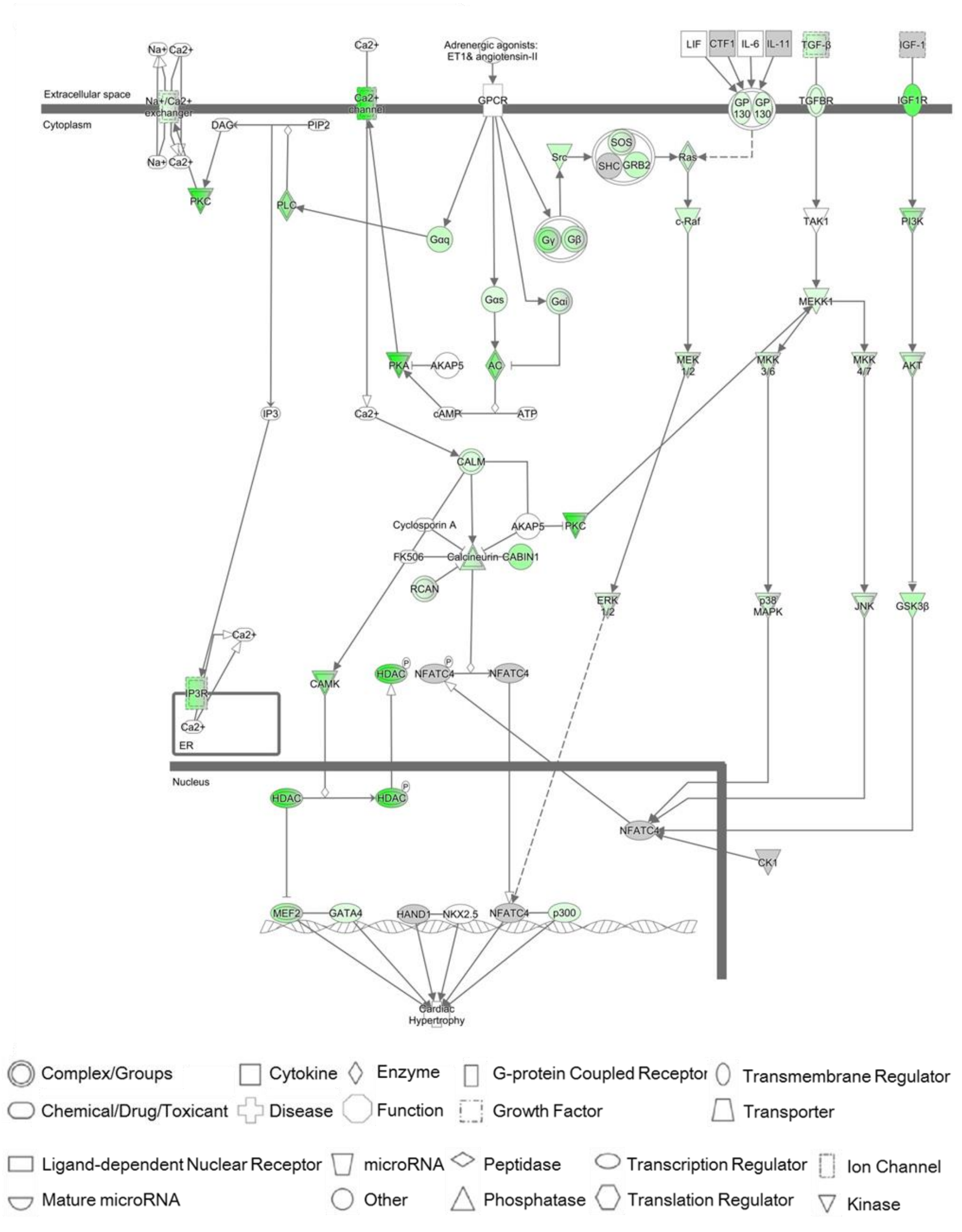

**Supplementary Figure 13**. Schematic diagram of role of NFAT in cardiac hypertrophy pathway with which the cisplatin damaged genes are highly associated. The green color indicates the gene (or gene group) was (were) damaged by cisplatin. The deeper the green color is, the higher the enrichment fold of genes mapped.

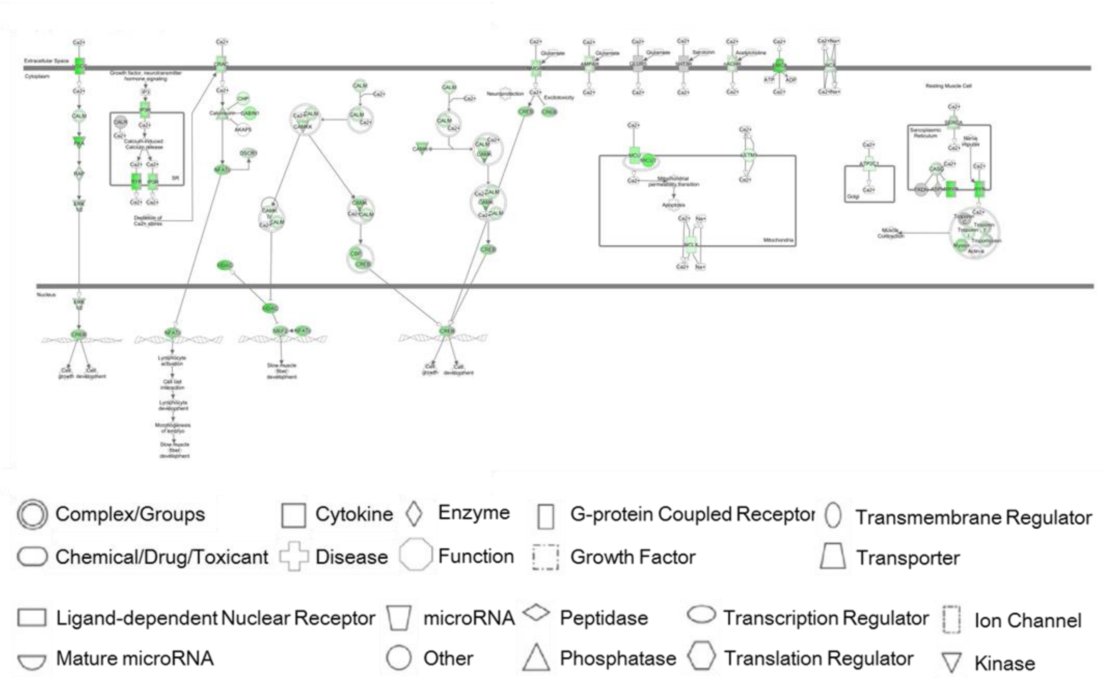

**Supplementary Figure 14**. Schematic diagram of calcium signaling pathway with which the cisplatin damaged genes are highly associated. The green color indicates the gene (or gene group) was (were) damaged by cisplatin. The deeper the green color is, the higher the enrichment fold of genes mapped.

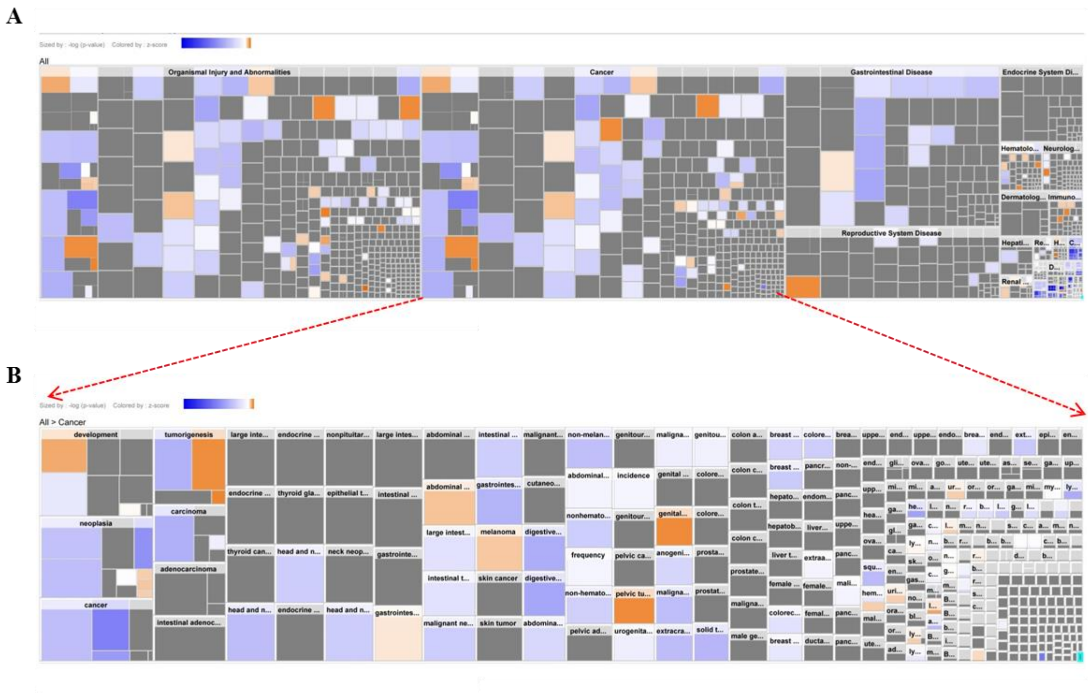

**Supplementary Figure 15.** (A) The heat map of diseases and functions which the cisplatin damaged genes are highly associated. (B) The enlarged view of heat map of cancer which the cisplatin damaged genes are highly associated. Orange indicates that the disease or function is activated, blue indicates inhibited, and gray indicates no prediction is available.

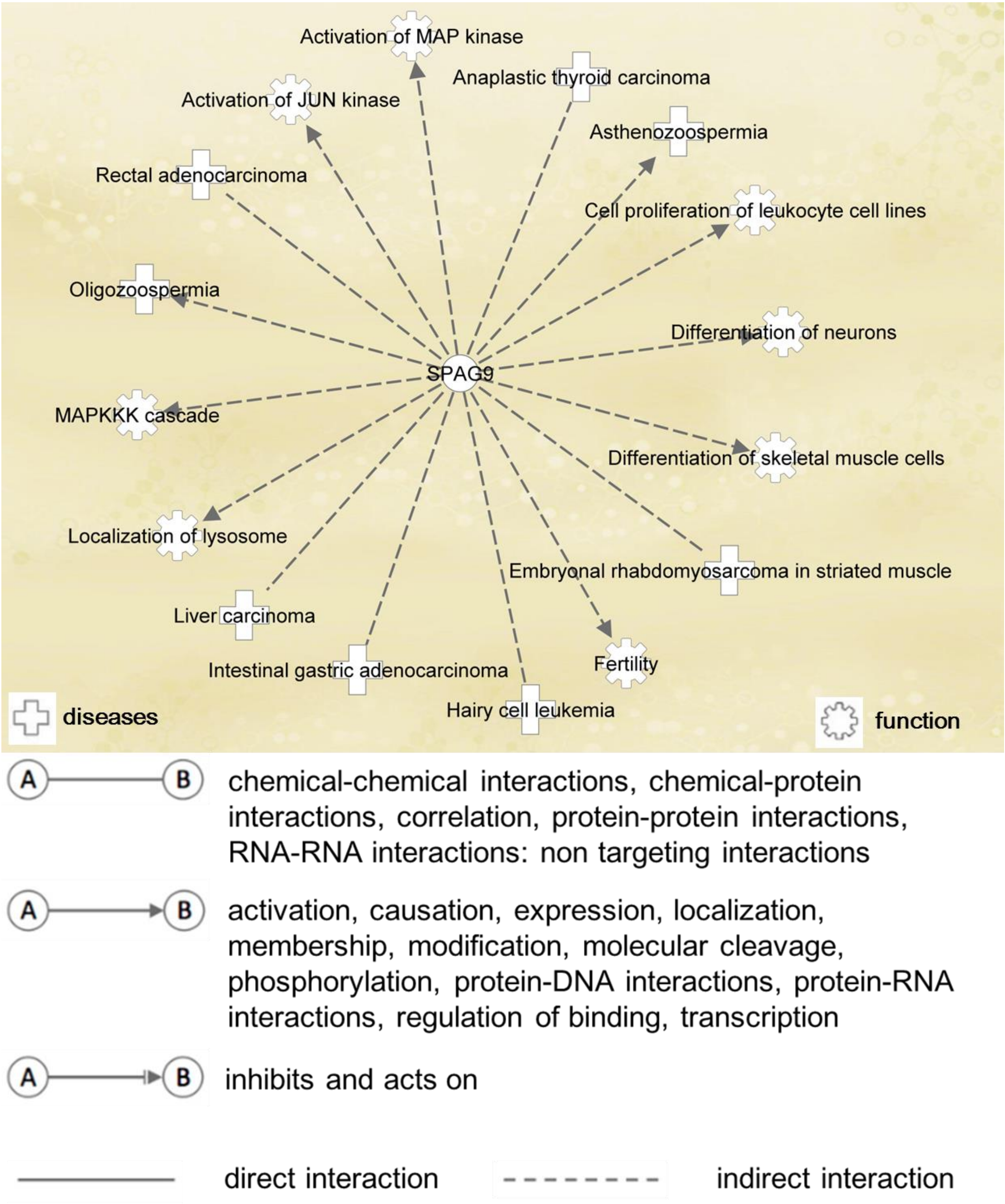

**Supplementary Figure 16**. Diseases and functions to which *SPAG9* gene is related.

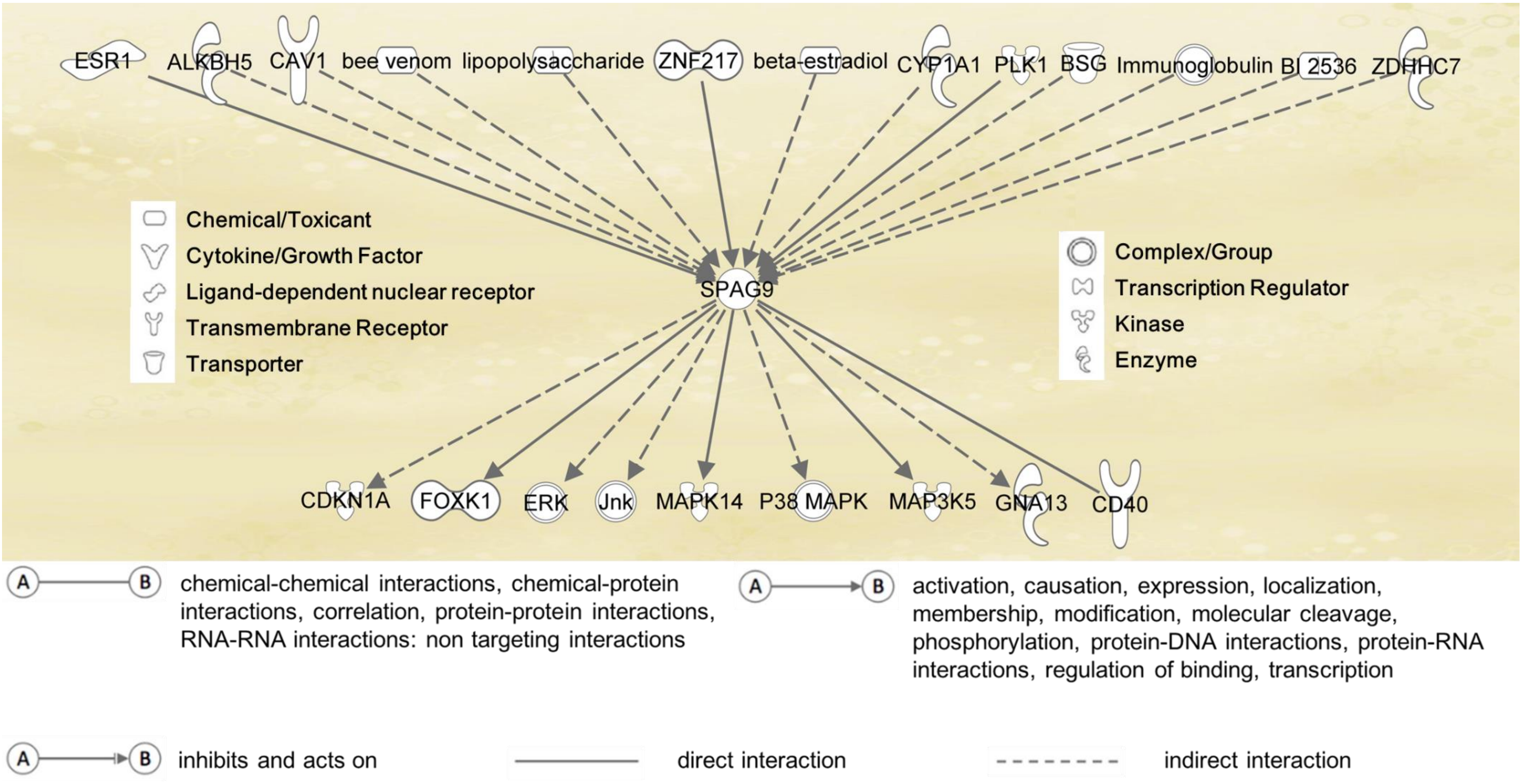

**Supplementary Figure 17**. The upstream and downstream genes or molecules which regulate *SPAG9* or are regulated by *SPAG9*.

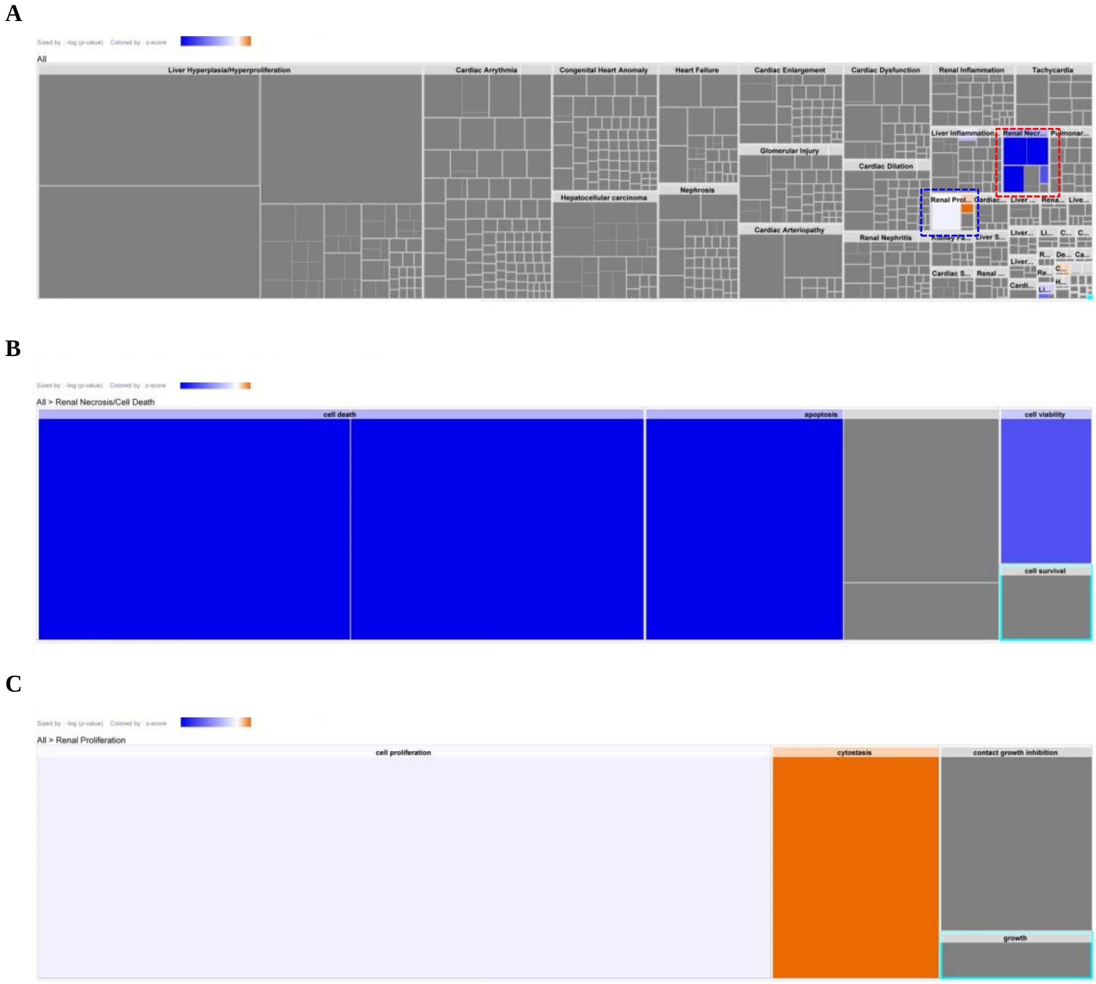

**Supplementary Figure 18.** (A) The heat map of toxicity and functions which the cisplatin damaged genes are highly associated. (B, C) The enlarged view of heat map of (B) renal necrosis and cell death, and (C) renal proliferation which the cisplatin damaged genes are highly associated. Orange indicates that the disease or function is activated, blue indicates inhibited, and gray indicates no prediction is available.

### Supplementary Tables

**Supplementary Table 1.** Softwares and databases used for gene screening and location.

| Software | Version | R Package | Version |
|---|---|---|---|
| BBTools | 37.48 | ggplot2 | 2.2.1 |
| R | 3.4.2 | reshape2 | 1.4.2 |
| BLAST+ | 2.2.28 | magrittr | 1.5 |
| Java | 1.8.0_151 | | |
| picard | 2.14.0 | | |
| htslib | 1.6 | | |
| samtools | 1.6 | | |
| macs14 | 1.4.2 | | |
| SnpEff | 4.3s | | |
| bedtools | 2.26.0 | | |

**Supplementary Table 2.** Information for sequencing samples prepared from genomic DNA extracts of A549 cells with cisplatin treatment (positive sample: PS) or without cisplatin treatment (control sample: CS).

| **Sample No.** | **PS-1** | **PS-2** | **CS-1** | **CS-2** |
|---|---|---|---|---|
| **DNA concentration (ng/μL)** | **17.1** | **15.1** | **15.7** | **17.1** |
| **Volume (μL)** | **260** | **200** | **260** | **200** |

**Supplementary Table 3.** Statistic analysis of Sequencing.

| Replicate | 1 | | 2 | |
|---|---|---|---|---|
| | Positive Sample | Negative Sample | Positive Sample | Negative Sample |
| Total Reads Count | 162704730 | 166832042 | 235408404 | 194048372 |
| Total Bases Count (bp) | 23334767257 | 23950671824 | 34314608427 | 28416732137 |
| Average Read Length (bp) | 143.42 | 143.56 | 145.77 | 146.44 |
| Q10 Bases Count (bp) | 23334513721 | 23950417040 | 34314572665 | 28416702861 |
| Q10 Bases Ratio (%) | 100.00% | 100.00% | 100.00% | 100.00% |
| Q20 Bases Count (bp) | 23043629520 | 23642092904 | 33800146069 | 27994785627 |
| Q20 Bases Ratio (%) | 98.75% | 98.71% | 98.50% | 98.52% |
| Q30 Bases Count (bp) | 22389034358 | 22959532355 | 32465733778 | 26882783529 |
| Q30 Bases Ratio (%) | 95.95% | 95.86% | 94.61% | 94.60% |
| N Bases Count (bp) | 57152 | 58518 | 35762 | 29276 |
| N Bases Ratio (%) | 0.00% | 0.00% | 0.00% | 0.00% |
| GC Bases Count (bp) | 13042819189 | 12964880382 | 14818176354 | 11829771086 |
| GC Bases Ratio (%) | 55.89% | 54.13% | 43.18% | 41.63% |

**Supplementary Table 4.** Rates of reads mapped to human genome.

| Replicate | 1 | | 2 | |
|---|---|---|---|---|
| | Positive Sample | Negative Sample | Positive Sample | Negative Sample |
| Total reads | 162704730(100.00%) | 166832042(100.00%) | 186590790(100.00%) | 199928074(100.00%) |
| Total mapped | 161812149(99.45%) | 165872115(99.42%) | 184756224(99.02%) | 198808893(99.44%) |
| Multiple mapped | 12953876(7.96%) | 11386662(6.83%) | 12622909(6.77%) | 14897960(7.45%) |
| Uniquely mapped | 148858273(91.49%) | 154485453(92.60%) | 172133315(92.25%) | 183910933(91.99%) |
| Read-1 mapped | 74517147(45.80%) | 77310628(46.34%) | 86135705(46.16%) | 92029339(46.03%) |
| Read-2 mapped | 74341126(45.69%) | 77174825(46.26%) | 85997610(46.09%) | 91881594(45.96%) |
| Reads map to '+' | 74429891(45.75%) | 77242602(46.30%) | 86075833(46.13%) | 91975255(46.00%) |
| Reads map to '-' | 74428382(45.74%) | 77242851(46.30%) | 86057482(46.12%) | 91935678(45.98%) |
| Reads mapped in proper pairs | 143215783(88.02%) | 151046514(90.54%) | 168403729(90.25%) | 181266998(90.67%) |

| Supplementary Table 5. Genes damaged by cisplatin. | | | | | | | | | |
|---|---|---|---|---|---|---|---|---|---|
| Chromosome | Gene | Average number of identified peaks (n=2) | Average number of reads (n=2) | Average enrichment fold (n=2) | Symbol in IPA | Entrez Gene Name | Location | Type(s) | Drug(s) |
| chr7 | *PTPRN2* | 361 | 17479 | 830 | PTPRN2 | protein tyrosine phosphatase receptor type N2 | Plasma Membrane | phosphatase | |
| chr1 | *CAMTA1* | 298 | 11871 | 721 | CAMTA1 | calmodulin binding transcription activator 1 | Other | other | |
| chr20 | *CDH4* | 278 | 17968 | 616 | CDH4 | cadherin 4 | Plasma Membrane | other | |
| chr8 | *TRAPPC9* | 227 | 8829 | 547 | TRAPPC9 | trafficking protein particle | Plasma Membrane | other | |
| chr11 | *SHANK2* | 228 | 9608 | 534 | SHANK2 | SH3 and multiple ankyrin repeat domains 2 | Plasma Membrane | other | |
| chr17 | *RBFOX3* | 198 | 9413 | 478 | RBFOX3 | RNA binding fox-1 homolog 3 | Nucleus | other | |
| chr7 | *CUX1* | 189 | 6788 | 452 | CUX1 | cut like homeobox 1 | Nucleus | transcription regulator | |
| chr7 | *SDK1* | 192 | 7788 | 441 | SDK1 | sidekick cell adhesion molecule 1 | Plasma Membrane | other | |
| chr4 | *SORCS2* | 150 | 6829 | 389 | SORCS2 | sortilin related VPS10 domain containing receptor 2 | Plasma Membrane | transporter | |
| chr1 | *KAZN* | 158 | 5457 | 369 | KAZN | kazrin, periplakin interacting | Plasma Membrane | other | |
| chr16 | *SNX29* | 156 | 4578 | 361 | SNX29 | sorting nexin 29 | Cytoplasm | other | |
| chr10 | *CDH23* | 152 | 10350 | 352 | CDH23 | cadherin related 23 | Plasma Membrane | transporter | |
| chr17 | *RPTOR* | 147 | 6474 | 351 | RPTOR | regulatory associated protein of MTOR complex 1 | Cytoplasm | other | |
| chr5 | *PDZD2* | 166 | 12122 | 347 | PDZD2 | PDZ domain containing 2 | Plasma Membrane | other | |
| chr7 | *MAD1L1* | 166 | 9975 | 342 | MAD1L1 | mitotic arrest deficient 1 like 1 | Nucleus | other | |
| chr1 | *PRDM16* | 153 | 7485 | 334 | PRDM16 | PR/SET domain 16 | Nucleus | transcription regulator | |
| chr10 | *GRID1* | 136 | 4640 | 317 | GRID1 | glutamate ionotropic receptor delta type subunit 1 | Plasma Membrane | ion channel | |
| chrX | *DMD* | 120 | 2795 | 312 | DMD | dystrophin | Plasma Membrane | other | |
| chr7 | *DPP6* | 139 | 3866 | 311 | DPP6 | dipeptidyl peptidase like 6 | Plasma Membrane | other | |
| chr18 | *DLGAP1* | 127 | 5196 | 306 | DLGAP1 | DLG associated protein 1 | Plasma Membrane | other | |
| chr6 | *RPS6KA2* | 122 | 5085 | 303 | RPS6KA2 | ribosomal protein S6 kinase A2 | Nucleus | kinase | PMD-026 |
| chr7 | *CNTNAP2* | 123 | 2699 | 298 | CNTNAP2 | contactin associated protein like | Plasma Membrane | other | |
| chr8 | *CSMD1* | 121 | 2754 | 290 | CSMD1 | CUB and Sushi multiple domains | Plasma Membrane | other | |
| chr17 | *RAP1GAP2* | 119 | 4658 | 287 | RAP1GAP2 | RAP1 GTPase activating protein | Cytoplasm | other | |
| chr9 | *PTPRD* | 112 | 3973 | 286 | PTPRD | protein tyrosine phosphatase receptor type D | Plasma Membrane | phosphatase | |
| chr7 | *TNS3* | 119 | 6409 | 286 | TNS3 | tensin 3 | Plasma Membrane | phosphatase | |
| chr5 | *COL23A1* | 120 | 5945 | 282 | COL23A1 | collagen type XXIII alpha 1 chain | Plasma Membrane | other | collagenase clostridium |
| chr11 | *DLG2* | 122 | 4844 | 282 | DLG2 | discs large MAGUK scaffold | Plasma Membrane | kinase | |
| chr17 | *MSI2* | 119 | 3877 | 282 | MSI2 | musashi RNA binding protein 2 | Cytoplasm | other | |
| chr2 | *SH3RF3* | 119 | 7857 | 280 | SH3RF3 | SH3 domain containing ring finger | Other | other | |
| chr16 | *CMIP* | 119 | 5984 | 279 | CMIP | c-Maf inducing protein | Cytoplasm | other | |
| chr1 | *HIVEP3* | 119 | 4160 | 278 | HIVEP3 | HIVEP zinc finger 3 | Nucleus | transcription regulator | |
| chr2 | *AGAP1* | 97 | 3589 | 277 | AGAP1 | ArfGAP with GTPase domain, ankyrin repeat and PH domain 1 | Cytoplasm | enzyme | |
| chr16 | *ZNF423* | 115 | 4278 | 270 | ZNF423 | zinc finger protein 423 | Nucleus | transcription regulator | |
| chr19 | *SIPA1L3* | 114 | 4026 | 267 | SIPA1L3 | signal induced proliferation associated 1 like 3 | Extracellular Space | other | |
| chr3 | *ATP2B2* | 114 | 4410 | 266 | ATP2B2 | ATPase plasma membrane Ca2+ transporting 2 | Plasma Membrane | transporter | |
| chr22 | *TBC1D22A* | 97 | 4291 | 263 | TBC1D22A | TBC1 domain family member 22A | Other | other | |
| chr17 | *SDK2* | 112 | 4768 | 262 | SDK2 | sidekick cell adhesion molecule 2 | Plasma Membrane | other | |
| chr10 | *CTNNA3* | 103 | 3684 | 260 | CTNNA3 | catenin alpha 3 | Plasma Membrane | other | |
| chr11 | *DSCAML1* | 111 | 4046 | 260 | DSCAML1 | DS cell adhesion molecule like 1 | Plasma Membrane | other | |
| chr16 | *RP11-420N3.2* | 121 | 2587 | 260 | | | | | |
| chr7 | *PRKAG2* | 106 | 4667 | 259 | PRKAG2 | protein kinase AMP-activated non-catalytic subunit gamma 2 | Cytoplasm | kinase | |
| chr10 | *ZMIZ1* | 110 | 5320 | 254 | ZMIZ1 | zinc finger MIZ-type containing 1 | Nucleus | other | |
| chr19 | *CACNA1A* | 107 | 2881 | 253 | CACNA1A | calcium voltage-gated channel subunit alpha1 A | Plasma Membrane | ion channel | pregabalin, bepridil, clevidipine butyrate |
| chr3 | *TBC1D5* | 101 | 3474 | 248 | TBC1D5 | TBC1 domain family member 5 | Extracellular Space | other | |
| chr20 | *PTPRT* | 117 | 3027 | 247 | PTPRT | protein tyrosine phosphatase receptor type T | Plasma Membrane | phosphatase | |
| chr2 | *GLI2* | 106 | 6355 | 245 | GLI2 | GLI family zinc finger 2 | Nucleus | transcription regulator | |
| chr1 | *CSMD2* | 108 | 2934 | 243 | CSMD2 | CUB and Sushi multiple domains | Other | other | |
| chr5 | *CTNND2* | 113 | 6139 | 242 | CTNND2 | catenin delta 2 | Plasma Membrane | other | |
| chr17 | *GAS7* | 100 | 3274 | 240 | GAS7 | growth arrest specific 7 | Cytoplasm | transcription regulator | |
| chr6 | *GMDS* | 93 | 2956 | 238 | GMDS | GDP-mannose 4,6-dehydratase | Cytoplasm | enzyme | |
| chr1 | *SMYD3* | 91 | 2500 | 238 | SMYD3 | SET and MYND domain | Nucleus | enzyme | |
| chr12 | *CUX2* | 101 | 3438 | 237 | CUX2 | cut like homeobox 2 | Nucleus | transcription regulator | |
| chr1 | *RERE* | 94 | 2532 | 236 | RERE | arginine-glutamic acid dipeptide repeats | Nucleus | transcription regulator | |
| chr9 | *VAV2* | 108 | 6301 | 232 | VAV2 | vav guanine nucleotide exchange factor 2 | Cytoplasm | transcription regulator | |
| chr12 | *NCOR2* | 107 | 5350 | 232 | NCOR2 | nuclear receptor corepressor 2 | Nucleus | transcription regulator | |
| chr17 | *ASIC2* | 107 | 2345 | 230 | ASIC2 | acid sensing ion channel subunit 2 | Plasma Membrane | ion channel | amiloride, amiloride/hydrochlorothiazide |
| chr3 | *EPHB1* | 109 | 3202 | 227 | EPHB1 | EPH receptor B1 | Plasma Membrane | kinase | |
| chr11 | *TENM4* | 108 | 3446 | 225 | TENM4 | teneurin transmembrane protein 4 | Plasma Membrane | other | |
| chr17 | *ABR* | 102 | 6183 | 224 | ABR | ABR activator of RhoGEF and GTPase | Cytoplasm | other | |
| chr17 | *BCAS3* | 89 | 2343 | 224 | BCAS3 | BCAS3 microtubule associated cell migration factor | Nucleus | other | |

| | | | | | | | | | |
|---|---|---|---|---|---|---|---|---|---|
| chr1 | *DAB1* | 101 | 2170 | 224 | DAB1 | DAB adaptor protein 1 | Cytoplasm | other | |
| chr7 | *AUTS2* | 95 | 2380 | 224 | AUTS2 | activator of transcription and developmental regulator AUTS2 | Nucleus | other | |
| chr15 | *ADAMTS17* | 101 | 4910 | 224 | ADAMTS17 | ADAM metallopeptidase with thrombospondin type 1 motif 17 | Extracellular Space | other | |
| chr3 | *CACNA2D3* | 99 | 2388 | 223 | CACNA2D3 | calcium voltage-gated channel auxiliary subunit alpha2delta 3 | Plasma Membrane | ion channel | pregabalin |
| chr5 | *TRIO* | 100 | 8450 | 223 | TRIO | trio Rho guanine nucleotide exchange factor | Cytoplasm | kinase | |
| chr10 | *INPP5A* | 103 | 5096 | 222 | INPP5A | inositol polyphosphate-5-phosphatase A | Plasma Membrane | phosphatase | |
| chr1 | *KIF26B* | 97 | 2591 | 222 | KIF26B | kinesin family member 26B | Other | other | |
| chr7 | *CALN1* | 100 | 2012 | 222 | CALN1 | calneuron 1 | Cytoplasm | other | |
| chr17 | *SEPT9* | 93 | 4349 | 221 | SEPTIN9 | septin 9 | Cytoplasm | enzyme | |
| chr5 | *ADAMTS2* | 99 | 5270 | 221 | ADAMTS2 | ADAM metallopeptidase with thrombospondin type 1 motif 2 | Extracellular Space | peptidase | |
| chr8 | *ERICH1-AS1* | 92 | 3651 | 220 | DLGAP2 | DLG associated protein 2 | Plasma Membrane | other | |
| chr10 | *ADK* | 89 | 3118 | 220 | ADK | adenosine kinase | Nucleus | kinase | pegintron/ribavirin, interferon alfacon-1/ribavirin, IFNA2B/ribavirin, ribavirin, nitazoxanide/peginterferon alfa-2a/ribavirin, adenosine kinase inhibitor, PEG- |
| chr10 | *ADARB2* | 80 | 3476 | 218 | ADARB2 | adenosine deaminase RNA specific B2 (inactive) | Nucleus | enzyme | |
| chr12 | *FAM101A* | 94 | 3152 | 217 | LNA/ZNF664-RFL | refilin A | Other | transporter | |
| chr7 | *PLXNA4* | 97 | 2610 | 216 | PLXNA4 | plexin A4 | Plasma Membrane | transmembrane receptor | |
| chr9 | *DENND1A* | 92 | 3620 | 215 | DENND1A | DENN domain containing 1A | Plasma Membrane | other | |
| chr10 | *GRK5* | 89 | 3590 | 212 | GRK5 | G protein-coupled receptor | Plasma Membrane | kinase | |
| chr1 | *RABGAP1L* | 85 | 2596 | 212 | RABGAP1L | RAB GTPase activating protein 1 | Cytoplasm | other | |
| chr2 | *PRKCE* | 99 | 3717 | 211 | PRKCE | protein kinase C epsilon | Cytoplasm | kinase | ingenol mebutate, protein kinase C-epsilon activator |
| chr11 | *GRIK4* | 92 | 2820 | 211 | GRIK4 | glutamate ionotropic receptor kainate type subunit 4 | Plasma Membrane | ion channel | |
| chr2 | *HDAC4* | 81 | 3919 | 207 | HDAC4 | histone deacetylase 4 | Nucleus | transcription regulator | OKI-179, tributyrin, belinostat, |
| chr15 | *THSD4* | 90 | 2351 | 207 | THSD4 | thrombospondin type 1 domain containing 4 | Cytoplasm | other | |
| chr11 | *NAV2* | 92 | 2496 | 206 | NAV2 | neuron navigator 2 | Nucleus | other | |
| chr19 | *ZNF536* | 90 | 2607 | 206 | ZNF536 | zinc finger protein 536 | Other | transcription regulator | |
| chr10 | *DIP2C* | 75 | 3183 | 205 | DIP2C | disco interacting protein 2 | Other | other | |
| chr5 | *FSTL4* | 85 | 2959 | 205 | FSTL4 | follistatin like 4 | Cytoplasm | other | |
| chr16 | *CDH13* | 95 | 2175 | 204 | CDH13 | cadherin 13 | Plasma Membrane | other | |
| chr11 | *CNTN5* | 82 | 4019 | 203 | CNTN5 | contactin 5 | Plasma Membrane | other | |
| chr5 | *SLIT3* | 95 | 3122 | 203 | SLIT3 | slit guidance ligand 3 | Extracellular Space | other | |
| chr2 | *THADA* | 88 | 5461 | 203 | THADA | THADA armadillo repeat | Cytoplasm | other | |
| chr10 | *PCDH15* | 85 | 2788 | 202 | PCDH15 | protocadherin related 15 | Plasma Membrane | other | |
| chr15 | *MEGF11* | 86 | 2845 | 202 | MEGF11 | multiple EGF like domains 11 | Other | other | |
| chr12 | *KSR2* | 92 | 2192 | 201 | KSR2 | kinase suppressor of ras 2 | Cytoplasm | kinase | |
| chr10 | *TACC2* | 85 | 3003 | 200 | TACC2 | transforming acidic coiled-coil containing protein 2 | Nucleus | other | |
| chr1 | *MACF1* | 78 | 1991 | 200 | MACF1 | microtubule actin crosslinking | Cytoplasm | enzyme | |
| chr9 | *EHMT1* | 84 | 4056 | 199 | EHMT1 | euchromatic histone lysine methyltransferase 1 | Nucleus | transcription regulator | |
| chr10 | *SH3PXD2A* | 86 | 3559 | 199 | SH3PXD2A | SH3 and PX domains 2A | Cytoplasm | other | |
| chr12 | *TMEM132D* | 84 | 2164 | 198 | TMEM132D | transmembrane protein 132D | Other | other | |
| chr2 | *DPP10* | 84 | 2160 | 197 | DPP10 | dipeptidyl peptidase like 10 | Extracellular Space | peptidase | |
| chr12 | *CACNA1C* | 72 | 2364 | 194 | CACNA1C | calcium voltage-gated channel subunit alpha1 C | Plasma Membrane | ion channel | amlodipine/perindopril, amlodipine/hydrochlorothiazide/olmesartan medoxomil, verapamil, amlodipine/hydrochlorothiazide/valsartan, dexamethasone/rituximab/verapamil, bepridil, mibefradil, clevidipine butyrate, nisoldipine, amlodipine/benazepril, nilvadipine, amlodipine/celecoxib, enalapril/felodipine, amlodipine/valsartan, amlodipine/ezetimibe/losartan/rosuvastatin, amlodipine/atorvastatin, [illegible] |
| chr3 | *LSAMP* | 80 | 2286 | 193 | LSAMP | limbic system associated membrane protein | Plasma Membrane | other | |

| | | | | | | | | | |
|---|---|---|---|---|---|---|---|---|---|
| chr18 | *PTPRM* | 81 | 2215 | 192 | PTPRM | protein tyrosine phosphatase receptor type M | Plasma Membrane | phosphatase | |
| chr1 | *RYR2* | 79 | 2268 | 190 | RYR2 | ryanodine receptor 2 | Plasma Membrane | ion channel | |
| chr10 | *DOCK1* | 82 | 2541 | 190 | DOCK1 | dedicator of cytokinesis 1 | Cytoplasm | other | |
| chr20 | *PREX1* | 82 | 4363 | 189 | PREX1 | phosphatidylinositol-3,4,5-trisphosphate dependent Rac exchange factor 1 | Cytoplasm | other | |
| chr3 | *KALRN* | 87 | 2677 | 189 | KALRN | kalirin RhoGEF kinase | Cytoplasm | kinase | |
| chr1 | *EPHB2* | 79 | 3304 | 189 | EPHB2 | EPH receptor B2 | Plasma Membrane | kinase | |
| chr17 | *STX8* | 72 | 1951 | 187 | STX8 | syntaxin 8 | Plasma Membrane | other | |
| chr22 | *FAM19A5* | 77 | 3654 | 186 | TAFA5 | TAFA chemokine like family member 5 | Extracellular Space | other | |
| chr9 | *COL5A1* | 86 | 5399 | 185 | COL5A1 | collagen type V alpha 1 chain | Extracellular Space | other | collagenase clostridium |
| chr11 | *NTM* | 83 | 2150 | 185 | NTM | neurotrimin | Plasma Membrane | other | |
| chr5 | *MYO10* | 85 | 5651 | 185 | MYO10 | myosin X | Cytoplasm | enzyme | |
| chr5 | *CDH18* | 81 | 3816 | 184 | CDH18 | cadherin 18 | Plasma Membrane | other | |
| chr17 | *PITPNC1* | 78 | 2009 | 184 | PITPNC1 | phosphatidylinositol transfer protein cytoplasmic 1 | Cytoplasm | transporter | |
| chr3 | *ERC2* | 82 | 1891 | 184 | ERC2 | ELKS/RAB6-interacting/CAST family member 2 | Cytoplasm | other | |
| chr7 | *GTF2IRD1* | 76 | 3283 | 183 | GTF2IRD1 | GTF2I repeat domain containing | Nucleus | transcription regulator | |
| chr7 | *MAGI2* | 79 | 2097 | 181 | MAGI2 | membrane associated guanylate kinase, WW and PDZ domain containing 2 | Plasma Membrane | kinase | |
| chr2 | *DYSF* | 73 | 3081 | 180 | DYSF | dysferlin | Plasma Membrane | other | |
| chr2 | *AFF3* | 79 | 2405 | 180 | AFF3 | AF4/FMR2 family member 3 | Nucleus | transcription regulator | |
| chr6 | *CASC15* | 71 | 1812 | 180 | CASC15 | cancer susceptibility 15 | Other | other | |
| chr17 | *PRKCA* | 78 | 3258 | 180 | PRKCA | protein kinase C alpha | Cytoplasm | kinase | ingenol mebutate, aprinocarsen, L-threo-safingol |
| chr9 | *CACNA1B* | 77 | 3754 | 179 | CACNA1B | calcium voltage-gated channel subunit alpha1 B | Plasma Membrane | ion channel | levetiracetam, gabapentin, ziconotide, bepridil, clevidipine butyrate, |
| chr7 | *COL26A1* | 78 | 2671 | 179 | COL26A1 | collagen type XXVI alpha 1 chain | Extracellular Space | other | |
| chr17 | *SMG6* | 73 | 2206 | 179 | SMG6 | SMG6 nonsense mediated mRNA decay factor | Nucleus | enzyme | |
| chr12 | *ANKS1B* | 77 | 1789 | 179 | ANKS1B | ankyrin repeat and sterile alpha motif domain containing 1B | Nucleus | other | |
| chr5 | *CDH12* | 77 | 3758 | 178 | CDH12 | cadherin 12 | Plasma Membrane | other | |
| chr5 | *PDE4D* | 73 | 1774 | 178 | PDE4D | phosphodiesterase 4D | Cytoplasm | enzyme | caffeine, pentoxifylline, arofylline, tetomilast, tolbutamide, anagrelide, theophylline, apremilast, milrinone, cilomilast, roflumilast, ketotifen, L-826,141, L 869298, |
| chr12 | *CHST11* | 79 | 2220 | 178 | CHST11 | carbohydrate sulfotransferase 11 | Cytoplasm | enzyme | |
| chr18 | *CTIF* | 68 | 2669 | 177 | CTIF | cap binding complex dependent translation initiation factor | Cytoplasm | other | |
| chr12 | *RIMBP2* | 74 | 2941 | 177 | RIMBP2 | RIMS binding protein 2 | Plasma Membrane | other | |
| chr16 | *GSG1L* | 78 | 2343 | 177 | GSG1L | GSG1 like | Plasma Membrane | other | |
| chr11 | *OPCML* | 79 | 1875 | 177 | OPCML | opioid binding protein/cell adhesion molecule like | Plasma Membrane | other | |
| chrX | *RP11-40F8.2* | 70 | 1646 | 177 | | | | | |
| chr13 | *MCF2L* | 80 | 4071 | 177 | MCF2L | MCF.2 cell line derived transforming sequence like | Cytoplasm | other | |
| chr3 | *IQSEC1* | 75 | 3665 | 175 | IQSEC1 | IQ motif and Sec7 domain | Cytoplasm | other | |
| chr1 | *GLIS1* | 74 | 3241 | 174 | GLIS1 | GLIS family zinc finger 1 | Nucleus | transcription regulator | |
| chr8 | *PVT1* | 74 | 2481 | 174 | PVT1 | Pvt1 oncogene | Other | other | |
| chr10 | *KCNMA1* | 78 | 2078 | 174 | KCNMA1 | potassium calcium-activated channel subfamily M alpha 1 | Plasma Membrane | ion channel | cromolyn, tedisamil |
| chr1 | *GRIK3* | 73 | 2562 | 173 | GRIK3 | glutamate ionotropic receptor kainate type subunit 3 | Plasma Membrane | ion channel | |
| chr3 | *STAG1* | 68 | 1921 | 173 | STAG1 | stromal antigen 1 | Nucleus | transcription regulator | |
| chr9 | *DAB2IP* | 76 | 4335 | 172 | DAB2IP | DAB2 interacting protein | Plasma Membrane | other | |
| chr10 | *C10orf11* | 79 | 2183 | 171 | LRMDA | leucine rich melanocyte differentiation associated | Other | other | |
| chr9 | *RALGPS1* | 73 | 3261 | 171 | RALGPS1 | Ral GEF with PH domain and SH3 binding motif 1 | Cytoplasm | other | |
| chr15 | *PCSK6* | 75 | 5213 | 171 | PCSK6 | proprotein convertase subtilisin/kexin type 6 | Extracellular Space | peptidase | |
| chr2 | *KLHL29* | 62 | 2543 | 171 | KLHL29 | kelch like family member 29 | Other | other | |
| chr7 | *C7orf10* | 73 | 1891 | 171 | SUGCT | succinyl-CoA:glutarate-CoA transferase | Cytoplasm | enzyme | |
| chr3 | *EEFSEC* | 72 | 3887 | 170 | EEFSEC | eukaryotic elongation factor, selenocysteine-tRNA specific | Cytoplasm | translation regulator | |
| chr3 | *DOCK3* | 74 | 1932 | 169 | DOCK3 | dedicator of cytokinesis 3 | Cytoplasm | other | |
| chr3 | *ROBO2* | 66 | 1624 | 169 | ROBO2 | roundabout guidance receptor 2 | Plasma Membrane | transmembrane receptor | |
| chr13 | *FARP1* | 68 | 2659 | 169 | FARP1 | FERM, ARH/RhoGEF and pleckstrin domain protein 1 | Plasma Membrane | other | |
| chr11 | *KIRREL3* | 72 | 2582 | 168 | KIRREL3 | kirre like nephrin family adhesion molecule 3 | Extracellular Space | other | |
| chr18 | *LDLRAD4* | 60 | 2210 | 167 | LDLRAD4 | low density lipoprotein receptor class A domain containing 4 | Cytoplasm | other | |

| | | | | | | | | | |
|---|---|---|---|---|---|---|---|---|---|
| chr11 | *KCNQ1* | 71 | 3625 | 167 | KCNQ1 | potassium voltage-gated channel subfamily Q member 1 | Plasma Membrane | ion channel | amlodipine/indapamide/perindopril, quinidine, indapamide, dextromethorphan/qu |
| chr11 | *SBF2* | 63 | 2093 | 167 | SBF2 | SET binding factor 2 | Cytoplasm | other | |
| chr10 | *MGMT* | 68 | 2763 | 167 | MGMT | O-6-methylguanine-DNA methyltransferase | Nucleus | enzyme | O6-alkylguanine-DNA alkyltransferase inhibitor, O6-benzylguanine |
| chr16 | *CLEC16A* | 68 | 2574 | 166 | CLEC16A | C-type lectin domain containing | Cytoplasm | other | |
| chr3 | *MAP4* | 68 | 1835 | 166 | MAP4 | microtubule associated protein 4 | Cytoplasm | other | |
| chr7 | *PRKAR1B* | 77 | 4290 | 166 | PRKAR1B | protein kinase cAMP-dependent type I regulatory subunit beta | Cytoplasm | kinase | |
| chr16 | *ABCC1* | 72 | 2331 | 166 | ABCC1 | ATP binding cassette subfamily C member 1 | Plasma Membrane | transporter | sulfinpyrazone |
| chr15 | *OTUD7A* | 61 | 2109 | 165 | OTUD7A | OTU deubiquitinase 7A | Cytoplasm | peptidase | |
| chr5 | *CTC-349C3.1* | 71 | 2667 | 165 | | | | | |
| chr1 | *AGBL4* | 73 | 1719 | 165 | AGBL4 | ATP/GTP binding protein like 4 | Cytoplasm | peptidase | |
| chr2 | *AC007682.1* | 68 | 1839 | 165 | | | | | |
| chr7 | *HIP1* | 74 | 2830 | 165 | HIP1 | huntingtin interacting protein 1 | Cytoplasm | other | |
| chr20 | *DLGAP4* | 73 | 3417 | 165 | DLGAP4 | DLG associated protein 4 | Plasma Membrane | other | |
| chr10 | *LINC00856* | 74 | 2153 | 165 | LINC00856 | long intergenic non-protein coding RNA 856 | Other | other | |
| chr1 | *IGSF21* | 73 | 2345 | 165 | IGSF21 | immunoglobin superfamily member 21 | Extracellular Space | other | |
| chr2 | *CTNNA2* | 71 | 1582 | 164 | CTNNA2 | catenin alpha 2 | Plasma Membrane | other | |
| chr10 | *COL13A1* | 73 | 4192 | 164 | COL13A1 | collagen type XIII alpha 1 chain | Plasma Membrane | other | collagenase clostridium |
| chr7 | *HECW1* | 71 | 2431 | 164 | HECW1 | HECT, C2 and WW domain containing E3 ubiquitin protein ligase 1 | Cytoplasm | enzyme | |
| chr17 | *WSCD1* | 70 | 1985 | 163 | WSCD1 | WSC domain containing 1 | Other | other | |
| chr17 | *TBCD* | 75 | 3672 | 163 | TBCD | tubulin folding cofactor D | Cytoplasm | other | |
| chr17 | *LINC00511* | 71 | 2212 | 163 | LINC00511 | long intergenic non-protein coding RNA 511 | Other | other | |
| chr19 | *INSR* | 68 | 1990 | 163 | INSR | insulin receptor | Plasma Membrane | kinase | canakinumab/INS, INS/pioglitazone, dapagliflozin/INS, insulin glargine/insulin lispro, insulin degludec, insulin aspart/insulin degludec, INS/sitagliptin, protamine zinc insulin, INS/protamine zinc insulin, subetta, insulin glargine, insulin detemir, insulin glulisine, alogliptin/INS, INS, INS/vildagliptin, insulin degludec/liraglutide, INS/saxagliptin, BMS-754807 |
| chr1 | *ARHGEF10L* | 68 | 3221 | 163 | ARHGEF10L | Rho guanine nucleotide exchange factor 10 like | Cytoplasm | enzyme | |
| chr2 | *MYT1L* | 58 | 2266 | 162 | MYT1L | myelin transcription factor 1 like | Nucleus | transcription regulator | |
| chr12 | *DIP2B* | 70 | 2651 | 162 | DIP2B | disco interacting protein 2 | Cytoplasm | other | |
| chr2 | *ACOXL* | 71 | 2653 | 160 | ACOXL | acyl-CoA oxidase like | Other | enzyme | |
| chr10 | *MICU1* | 66 | 2429 | 160 | MICU1 | mitochondrial calcium uptake 1 | Cytoplasm | other | |
| chr11 | *GALNT18* | 80 | 3827 | 160 | GALNT18 | polypeptide N-acetylgalactosaminyltransferase | Cytoplasm | enzyme | |
| chr13 | *ATP11A* | 70 | 3168 | 160 | ATP11A | ATPase phospholipid transporting 11A | Plasma Membrane | transporter | |
| chr5 | *CTNNA1* | 67 | 1831 | 160 | CTNNA1 | catenin alpha 1 | Plasma Membrane | other | |
| chr15 | *LINGO1* | 70 | 4062 | 159 | LINGO1 | leucine rich repeat and Ig domain containing 1 | Plasma Membrane | other | |
| chr12 | *TMEM132B* | 72 | 1871 | 159 | TMEM132B | transmembrane protein 132B | Other | other | |
| chr12 | *PITPNM2* | 69 | 2755 | 159 | PITPNM2 | phosphatidylinositol transfer protein membrane associated 2 | Cytoplasm | enzyme | |
| chr17 | *NTN1* | 70 | 2183 | 158 | NTN1 | netrin 1 | Extracellular Space | other | NP137 |

| | | | | | | | | | |
|---|---|---|---|---|---|---|---|---|---|
| chr15 | *IGF1R* | 68 | 3199 | 158 | IGF1R | insulin like growth factor 1 receptor | Plasma Membrane | transmembrane receptor | IGF-1R inhibitor, PL225B, IGF-methotrexate conjugate, cixutumumab, AVE1642, linsitinib, dalotuzumab, brigatinib, picropodophyllin, [225Ac]-FPI-1434, istiratumab, BMS-754807, ganitumab, KW-2450, ceritinib, [111In]-FPI-1547, XL228, BIIB022, [I- |
| chr5 | *TENM2* | 71 | 2148 | 157 | TENM2 | teneurin transmembrane protein 2 | Plasma Membrane | other | |
| chr3 | *CACNA1D* | 67 | 2047 | 157 | CACNA1D | calcium voltage-gated channel subunit alpha1 D | Plasma Membrane | ion channel | amlodipine/valsartan, amlodipine/ezetimibe/losartan/rosuvastatin, nicardipine, amlodipine/atorvastatin, aliskiren/amlodipine/hydrochlorothiazide, amlodipine/telmisartan, aliskiren/amlodipine, isradipine, MEM-1003, cinnarizine, nifedipine, amlodipine/indapamide/perindopril, amlodipine/perindopril, felodipine, verapamil, amlodipine/hydrochlorothiazide/valsartan, dexamethasone/rituximab/verapamil, [illegible] |
| chr1 | *ASH1L* | 63 | 1960 | 157 | ASH1L | ASH1 like histone lysine methyltransferase | Nucleus | transcription regulator | |
| chr16 | *ZFHX3* | 66 | 2203 | 157 | ZFHX3 | zinc finger homeobox 3 | Nucleus | transcription regulator | |
| chr20 | *PHACTR3* | 70 | 3915 | 155 | PHACTR3 | phosphatase and actin regulator | Nucleus | other | |
| chr16 | *RBFOX1* | 72 | 1352 | 155 | RBFOX1 | RNA binding fox-1 homolog 1 | Cytoplasm | other | |
| chr1 | *SSBP3* | 65 | 2540 | 155 | SSBP3 | single stranded DNA binding protein 3 | Nucleus | transcription regulator | |
| chr7 | *COBL* | 66 | 2920 | 154 | COBL | cordon-bleu WH2 repeat protein | Plasma Membrane | other | |
| chr4 | *ABLIM2* | 64 | 2750 | 154 | ABLIM2 | actin binding LIM protein family member 2 | Cytoplasm | other | |
| chr17 | *DNAH17* | 64 | 2606 | 154 | DNAH17 | dynein axonemal heavy chain 17 | Cytoplasm | other | |
| chr1 | *KCNN3* | 66 | 3105 | 154 | KCNN3 | potassium calcium-activated channel subfamily N member 3 | Plasma Membrane | ion channel | |
| chr5 | *STK10* | 63 | 3064 | 153 | STK10 | serine/threonine kinase 10 | Cytoplasm | kinase | |
| chr18 | *CELF4* | 61 | 2360 | 153 | CELF4 | CUGBP Elav-like family member | Nucleus | translation regulator | |
| chr16 | *TANGO6* | 60 | 1573 | 153 | TANGO6 | transport and golgi organization 6 homolog | Other | other | |
| chr10 | *PRKG1* | 67 | 2338 | 152 | PRKG1 | protein kinase cGMP-dependent | Cytoplasm | kinase | |
| chr10 | *TCERG1L* | 66 | 2843 | 152 | TCERG1L | transcription elongation regulator 1 like | Other | other | |
| chr10 | *SORCS1* | 63 | 1697 | 152 | SORCS1 | sortilin related VPS10 domain containing receptor 1 | Plasma Membrane | transporter | |
| chr20 | *PHF20* | 64 | 1956 | 150 | PHF20 | PHD finger protein 20 | Nucleus | transcription regulator | |
| chr13 | *PCCA* | 62 | 1766 | 150 | PCCA | propionyl-CoA carboxylase subunit alpha | Cytoplasm | enzyme | |
| chr20 | *EYA2* | 67 | 2374 | 150 | EYA2 | EYA transcriptional coactivator and phosphatase 2 | Nucleus | phosphatase | |
| chr16 | *CTD-3088G3.8* | 64 | 2692 | 150 | | | | | |
| chr1 | *EPB41* | 58 | 1558 | 149 | EPB41 | erythrocyte membrane protein band 4.1 | Plasma Membrane | other | |
| chr4 | *PPP2R2C* | 57 | 2523 | 149 | PPP2R2C | protein phosphatase 2 regulatory subunit Bgamma | Nucleus | phosphatase | |
| chr12 | *KDM2B* | 63 | 2155 | 149 | KDM2B | lysine demethylase 2B | Nucleus | enzyme | |
| chr2 | *FAM178B* | 64 | 3125 | 149 | FAM178B | family with sequence similarity 178 member B | Other | other | |
| chr20 | *ATP9A* | 62 | 2475 | 149 | ATP9A | ATPase phospholipid transporting 9A (putative) | Cytoplasm | transporter | |
| chr7 | *WBSCR17* | 69 | 1389 | 148 | GALNT17 | polypeptide N-acetylgalactosaminyltransferase | Cytoplasm | enzyme | |
| chr17 | *RAB11FIP4* | 62 | 2436 | 148 | RAB11FIP4 | RAB11 family interacting protein | Cytoplasm | other | |
| chr2 | *TBC1D8* | 62 | 2646 | 148 | TBC1D8 | TBC1 domain family member 8 | Plasma Membrane | other | |
| chrX | *IL1RAPL1* | 56 | 1289 | 148 | IL1RAPL1 | interleukin 1 receptor accessory protein like 1 | Plasma Membrane | transmembrane receptor | |
| chr1 | *ACOT7* | 65 | 3852 | 147 | ACOT7 | acyl-CoA thioesterase 7 | Cytoplasm | enzyme | |
| chr11 | *ABTB2* | 67 | 2715 | 147 | ABTB2 | ankyrin repeat and BTB domain containing 2 | Nucleus | other | |
| chr13 | *COL4A2* | 59 | 2518 | 147 | COL4A2 | collagen type IV alpha 2 chain | Extracellular Space | other | collagenase clostridium |
| chr14 | *RAD51B* | 54 | 1681 | 146 | RAD51B | RAD51 paralog B | Nucleus | enzyme | |

| | | | | | | | | | |
|---|---|---|---|---|---|---|---|---|---|
| chr16 | *SNX29P2* | 62 | 2421 | 146 | SNX29P2 | sorting nexin 29 pseudogene 2 | Other | other | |
| chr2 | *MTA3* | 62 | 2051 | 145 | MTA3 | metastasis associated 1 family member 3 | Nucleus | transcription regulator | |
| chr1 | *RGS7* | 60 | 1706 | 145 | RGS7 | regulator of G protein signaling 7 | Cytoplasm | enzyme | |
| chr11 | *PKNOX2* | 65 | 2041 | 145 | PKNOX2 | PBX/knotted 1 homeobox 2 | Nucleus | transcription regulator | |
| chr12 | *GPR133* | 64 | 3176 | 145 | ADGRD1 | adhesion G protein-coupled receptor D1 | Plasma Membrane | -protein coupled recept | |
| chr15 | *SCAPER* | 58 | 1917 | 144 | SCAPER | S-phase cyclin A associated protein in the ER | Extracellular Space | other | |
| chr7 | *TYW1B* | 55 | 1452 | 144 | TYW1B | tRNA-yW synthesizing protein 1 homolog B | Other | other | |
| chr15 | *TLN2* | 54 | 1504 | 143 | TLN2 | talin 2 | Nucleus | other | |
| chr1 | *RNF220* | 60 | 1973 | 143 | RNF220 | ring finger protein 220 | Cytoplasm | enzyme | |
| chr10 | *CTBP2* | 60 | 2432 | 143 | CTBP2 | C-terminal binding protein 2 | Nucleus | transcription regulator | |
| chr17 | *SLC39A11* | 62 | 1640 | 142 | SLC39A11 | solute carrier family 39 member | Other | transporter | |
| chr3 | *ADCY5* | 62 | 3041 | 142 | ADCY5 | adenylate cyclase 5 | Plasma Membrane | enzyme | |
| chr3 | *FOXP1* | 59 | 1396 | 142 | FOXP1 | forkhead box P1 | Nucleus | transcription regulator | |
| chr5 | *WDR70* | 62 | 4470 | 141 | WDR70 | WD repeat domain 70 | Other | other | |
| chr8 | *NRG1* | 58 | 1303 | 141 | NRG1 | neuregulin 1 | Plasma Membrane | growth factor | |
| chrX | *TBL1X* | 57 | 1862 | 141 | TBL1X | transducin beta like 1 X-linked | Nucleus | transcription regulator | |
| chr18 | *ZBTB7C* | 50 | 1754 | 140 | ZBTB7C | zinc finger and BTB domain containing 7C | Nucleus | transcription regulator | |
| chr13 | *GPC6* | 60 | 1650 | 140 | GPC6 | glypican 6 | Plasma Membrane | transmembrane recepto | |
| chr3 | *CACNA2D2* | 58 | 2872 | 140 | CACNA2D2 | calcium voltage-gated channel auxiliary subunit alpha2delta 2 | Plasma Membrane | ion channel | alpha 2 delta calcium channel agonist, amiodarone, pregabalin, gabapentin, bepridil, |
| chr1 | *BCAR3* | 63 | 2161 | 140 | BCAR3 | BCAR3 adaptor protein, NSP family member | Cytoplasm | other | |
| chr7 | *CLIP2* | 60 | 2082 | 140 | CLIP2 | CAP-Gly domain containing linker protein 2 | Cytoplasm | transcription regulator | |
| chr9 | *ABL1* | 57 | 2203 | 140 | ABL1 | ABL proto-oncogene 1, non-receptor tyrosine kinase | Nucleus | kinase | HQP1351, temozolomide, cytarabine/filgrastim/fludarabine phosphate/imatinib, clofarabine/cyclophosphamide/etoposide/imatinib, cyclophosphamide/daunorubicin/imatinib/prednisone/vincristine, regorafenib, dasatinib/nilotinib, ponatinib, imatinib, nilotinib, cytarabine/imatinib, saracatinib, imatinib/nilotinib/peginteron, L-asparaginase/dexamethasone/imatinib/vincristine, L-asparaginase/daunor[illegible] |
| chr13 | *CLYBL* | 60 | 1843 | 139 | CLYBL | citrate lyase beta like | Cytoplasm | enzyme | |
| chr2 | *EXOC6B* | 55 | 1426 | 139 | EXOC6B | exocyst complex component 6B | Other | other | |
| chr7 | *MKLN1* | 55 | 1288 | 139 | MKLN1 | muskelin 1 | Cytoplasm | other | |
| chr1 | *CASZ1* | 62 | 2993 | 139 | CASZ1 | castor zinc finger 1 | Nucleus | enzyme | |
| chr9 | *SCAI* | 56 | 1766 | 139 | SCAI | suppressor of cancer cell | Nucleus | transcription regulator | |
| chr11 | *LRP5* | 62 | 2914 | 139 | LRP5 | LDL receptor related protein 5 | Plasma Membrane | transmembrane recepto | |
| chr8 | *XKR6* | 59 | 1969 | 139 | XKR6 | XK related 6 | Other | other | |
| chr3 | *ROBO1* | 53 | 1264 | 138 | ROBO1 | roundabout guidance receptor 1 | Plasma Membrane | transmembrane recepto | |
| chr6 | *GRM4* | 58 | 2806 | 138 | GRM4 | glutamate metabotropic receptor | Plasma Membrane | -protein coupled recept | fasoracetam |
| chr8 | *TSNARE1* | 65 | 3348 | 138 | TSNARE1 | t-SNARE domain containing 1 | Other | other | |
| chr20 | *RP5-1043L13.1* | 67 | 2466 | 138 | | | | | |
| chr7 | *ELMO1* | 62 | 1705 | 137 | ELMO1 | engulfment and cell motility 1 | Cytoplasm | other | |
| chr11 | *FCHSD2* | 54 | 1819 | 137 | FCHSD2 | FCH and double SH3 domains 2 | Plasma Membrane | other | |
| chr9 | *RAPGEF1* | 60 | 2872 | 137 | RAPGEF1 | Rap guanine nucleotide exchange factor 1 | Cytoplasm | other | |
| chr19 | *RYR1* | 60 | 2279 | 136 | RYR1 | ryanodine receptor 1 | Cytoplasm | ion channel | dantrolene |
| chr22 | *MICAL3* | 50 | 1886 | 136 | MICAL3 | microtubule associated monooxygenase, calponin and LIM domain containing 3 | Cytoplasm | enzyme | |
| chr2 | *MIR4435-1HG* | 59 | 2478 | 136 | MIR4435-2HG | MIR4435-2 host gene | Other | other | |
| chr5 | *ERGIC1* | 59 | 4621 | 136 | ERGIC1 | endoplasmic reticulum-golgi intermediate compartment 1 | Cytoplasm | other | |
| chr1 | *PLXNA2* | 60 | 2532 | 135 | PLXNA2 | plexin A2 | Plasma Membrane | transmembrane recepto | |
| chr6 | *CDKAL1* | 55 | 1273 | 135 | CDKAL1 | CDK5 regulatory subunit associated protein 1 like 1 | Cytoplasm | enzyme | |
| chr14 | *ITPK1* | 53 | 2299 | 135 | ITPK1 | inositol-tetrakisphosphate 1- | Cytoplasm | kinase | |
| chr5 | *SIL1* | 60 | 1678 | 135 | SIL1 | SIL1 nucleotide exchange factor | Cytoplasm | transporter | |
| chr8 | *CSMD3* | 55 | 1425 | 135 | CSMD3 | CUB and Sushi multiple domains | Plasma Membrane | enzyme | |
| chr17 | *MPRIP* | 55 | 2333 | 134 | MPRIP | myosin phosphatase Rho interacting protein | Cytoplasm | other | |
| chr16 | *ADCY9* | 58 | 2058 | 134 | ADCY9 | adenylate cyclase 9 | Plasma Membrane | enzyme | |
| chr6 | *TRERF1* | 57 | 1702 | 134 | TRERF1 | transcriptional regulating factor 1 | Nucleus | transcription regulator | |

| | | | | | | | | | |
|---|---|---|---|---|---|---|---|---|---|
| chr15 | *FAM189A1* | 52 | 1747 | 133 | FAM189A1 | family with sequence similarity 189 member A1 | Other | other | |
| chr1 | *EIF4G3* | 52 | 1341 | 133 | EIF4G3 | eukaryotic translation initiation factor 4 gamma 3 | Cytoplasm | translation regulator | |
| chr16 | *PRKCB* | 61 | 1316 | 133 | PRKCB | protein kinase C beta | Cytoplasm | kinase | ingenol mebutate, sotrastaurin, enzastaurin, MS-553, ruboxistaurin |
| chr12 | *TSPAN9* | 48 | 2086 | 133 | TSPAN9 | tetraspanin 9 | Plasma Membrane | other | |
| chr15 | *RP11-266O8.1* | 59 | 2510 | 132 | | | | | |
| chr12 | *TMEM132C* | 61 | 1524 | 132 | TMEM132C | transmembrane protein 132C | Other | other | |
| chr7 | *HDAC9* | 54 | 1594 | 132 | HDAC9 | histone deacetylase 9 | Nucleus | transcription regulator | tributyrin, belinostat, pyroxamide |
| chr6 | *PHACTR1* | 60 | 1223 | 132 | PHACTR1 | phosphatase and actin regulator | Cytoplasm | other | |
| chr7 | *ATXN7L1* | 57 | 1674 | 132 | ATXN7L1 | ataxin 7 like 1 | Other | other | |
| chr1 | *INADL* | 53 | 1264 | 132 | PATJ | PATJ crumbs cell polarity complex component | Plasma Membrane | other | |
| chr10 | *JMJD1C* | 50 | 1761 | 132 | JMJD1C | jumonji domain containing 1C | Nucleus | enzyme | |
| chrX | *CASK* | 50 | 1280 | 131 | CASK | calcium/calmodulin dependent serine protein kinase | Plasma Membrane | kinase | |
| chr11 | *PTPRJ* | 60 | 3023 | 131 | PTPRJ | protein tyrosine phosphatase receptor type J | Plasma Membrane | phosphatase | |
| chr5 | *ARHGAP26* | 56 | 1359 | 131 | ARHGAP26 | Rho GTPase activating protein 26 | Cytoplasm | other | |
| chr10 | *CAMK1D* | 49 | 1492 | 131 | CAMK1D | calcium/calmodulin dependent protein kinase ID | Cytoplasm | kinase | |
| chr9 | *GLIS3* | 55 | 1425 | 130 | GLIS3 | GLIS family zinc finger 3 | Nucleus | transcription regulator | |
| chr1 | *FAF1* | 50 | 1207 | 130 | FAF1 | Fas associated factor 1 | Nucleus | other | |
| chr20 | *NFATC2* | 57 | 2511 | 130 | NFATC2 | nuclear factor of activated T cells | Nucleus | transcription regulator | |
| chr22 | *CELSR1* | 54 | 2428 | 129 | CELSR1 | cadherin EGF LAG seven-pass G-type receptor 1 | Plasma Membrane | G-protein coupled recepto | |
| chr8 | *MSRA* | 56 | 1704 | 129 | MSRA | methionine sulfoxide reductase A | Cytoplasm | enzyme | |
| chr10 | *ARHGAP22* | 57 | 3369 | 129 | ARHGAP22 | Rho GTPase activating protein 22 | Cytoplasm | other | |
| chr3 | *ITGA9* | 58 | 1444 | 129 | ITGA9 | integrin subunit alpha 9 | Plasma Membrane | other | |
| chr19 | *MYO9B* | 54 | 2202 | 129 | MYO9B | myosin IXB | Cytoplasm | enzyme | |
| chr3 | *RBM6* | 51 | 1457 | 129 | RBM6 | RNA binding motif protein 6 | Nucleus | other | |
| chr2 | *CLASP1* | 51 | 1889 | 128 | CLASP1 | cytoplasmic linker associated protein 1 | Cytoplasm | other | |
| chr1 | *ECE1* | 54 | 2397 | 128 | ECE1 | endothelin converting enzyme 1 | Plasma Membrane | peptidase | |
| chr17 | *JUP* | 55 | 2174 | 128 | JUP | junction plakoglobin | Plasma Membrane | other | |
| chr6 | *ZDHHC14* | 45 | 1555 | 128 | ZDHHC14 | zinc finger DHHC-type containing | Cytoplasm | enzyme | |
| chr2 | *TCF7L1* | 54 | 1705 | 128 | TCF7L1 | transcription factor 7 like 1 | Nucleus | transcription regulator | |
| chr16 | *C16orf95* | 53 | 1980 | 128 | C16orf95 | chromosome 16 open reading frame 95 | Other | other | |
| chr6 | *ANKS1A* | 54 | 1818 | 128 | ANKS1A | ankyrin repeat and sterile alpha motif domain containing 1A | Cytoplasm | other | |
| chr20 | *MACROD2* | 45 | 1225 | 127 | MACROD2 | mono-ADP ribosylhydrolase 2 | Nucleus | enzyme | |
| chr6 | *BTBD9* | 50 | 1408 | 127 | BTBD9 | BTB domain containing 9 | Other | other | |
| chr13 | *GPC5* | 54 | 1178 | 127 | GPC5 | glypican 5 | Plasma Membrane | other | |
| chr7 | *EXOC4* | 50 | 1237 | 127 | EXOC4 | exocyst complex component 4 | Cytoplasm | transporter | |
| chr13 | *RASA3* | 61 | 3224 | 127 | RASA3 | RAS p21 protein activator 3 | Plasma Membrane | ion channel | |
| chr16 | *PKD1L2* | 52 | 1843 | 127 | PKD1L2 | polycystin 1 like 2 (gene/pseudogene) | Other | ion channel | |
| chr1 | *NUP210L* | 54 | 1650 | 127 | NUP210L | nucleoporin 210 like | Other | other | |
| chr6 | *OFCC1* | 54 | 1307 | 127 | OFCC1 | orofacial cleft 1 candidate 1 | Cytoplasm | other | |
| chr1 | *VPS13D* | 48 | 1320 | 127 | VPS13D | vacuolar protein sorting 13 homolog D | Cytoplasm | other | |
| chr12 | *SCARB1* | 52 | 2291 | 127 | SCARB1 | scavenger receptor class B member 1 | Plasma Membrane | transporter | |
| chr8 | *PSD3* | 51 | 1231 | 126 | PSD3 | pleckstrin and Sec7 domain containing 3 | Cytoplasm | other | |
| chr17 | *ACACA* | 49 | 1319 | 126 | ACACA | acetyl-CoA carboxylase alpha | Cytoplasm | enzyme | |
| chr3 | *CLSTN2* | 61 | 1565 | 126 | CLSTN2 | calsyntenin 2 | Plasma Membrane | transporter | |
| chr16 | *ANKRD11* | 49 | 1914 | 126 | ANKRD11 | ankyrin repeat domain 11 | Nucleus | other | |
| chr15 | *APBA2* | 50 | 2025 | 126 | APBA2 | amyloid beta precursor protein binding family A member 2 | Cytoplasm | transporter | |
| chr12 | *NOS1* | 55 | 1566 | 126 | NOS1 | nitric oxide synthase 1 | Cytoplasm | enzyme | N(G)-monomethyl-D-arginine, GW 273629 |
| chr15 | *NTRK3* | 60 | 1913 | 126 | NTRK3 | neurotrophic receptor tyrosine kinase 3 | Plasma Membrane | kinase | larotrectinib, NOV1601, DS-6051b, belizatinib, lorlatinib, entrectinib, repotrectinib, |
| chr2 | *AC074391.1* | 55 | 1343 | 125 | | | | | |
| chr7 | *RNF216* | 53 | 1976 | 125 | RNF216 | ring finger protein 216 | Cytoplasm | enzyme | |
| chr6 | *FARS2* | 52 | 1445 | 125 | FARS2 | phenylalanyl-tRNA synthetase 2, mitochondrial | Cytoplasm | enzyme | |
| chr16 | *WWP2* | 52 | 1417 | 125 | WWP2 | WW domain containing E3 ubiquitin protein ligase 2 | Cytoplasm | enzyme | |
| chr6 | *SASH1* | 52 | 1344 | 124 | SASH1 | SAM and SH3 domain containing | Extracellular Space | other | |
| chr1 | *CACNA1E* | 59 | 1923 | 124 | CACNA1E | calcium voltage-gated channel subunit alpha1 E | Plasma Membrane | ion channel | |
| chr3 | *SRGAP3* | 57 | 2081 | 124 | SRGAP3 | SLIT-ROBO Rho GTPase activating protein 3 | Cytoplasm | other | |
| chr15 | *ADAMTSL3* | 52 | 1384 | 124 | ADAMTSL3 | ADAMTS like 3 | Other | other | |
| chr15 | *SLCO3A1* | 54 | 2631 | 124 | SLCO3A1 | solute carrier organic anion transporter family member 3A1 | Plasma Membrane | transporter | |
| chr19 | *PEPD* | 54 | 2605 | 124 | PEPD | peptidase D | Cytoplasm | peptidase | |
| chr1 | *ARID4B* | 44 | 1485 | 123 | ARID4B | AT-rich interaction domain 4B | Nucleus | transcription regulator | |
| chr10 | *HPSE2* | 49 | 1299 | 123 | HPSE2 | heparanase 2 (inactive) | Plasma Membrane | enzyme | |

| | | | | | | | | | |
|---|---|---|---|---|---|---|---|---|---|
| chr9 | *FNBP1* | 52 | 1745 | 123 | FNBP1 | formin binding protein 1 | Nucleus | enzyme | |
| chr17 | *RPH3AL* | 56 | 2699 | 123 | RPH3AL | rabphilin 3A like (without C2 | Plasma Membrane | other | |
| chr3 | *MYLK* | 56 | 2341 | 123 | MYLK | myosin light chain kinase | Cytoplasm | kinase | |
| chr1 | *NFASC* | 55 | 2361 | 123 | NFASC | neurofascin | Plasma Membrane | other | |
| chr16 | *KIAA0556* | 54 | 1785 | 123 | KIAA0556 | KIAA0556 | Extracellular Space | other | |
| chr16 | *BANP* | 52 | 2343 | 123 | BANP | BTG3 associated nuclear protein | Nucleus | other | |
| chr11 | *C11orf49* | 52 | 2837 | 123 | C11orf49 | chromosome 11 open reading frame 49 | Nucleus | other | |
| chr3 | *SUMF1* | 56 | 1325 | 122 | SUMF1 | sulfatase modifying factor 1 | Cytoplasm | enzyme | |
| chr7 | *EGFR* | 52 | 2523 | 122 | EGFR | epidermal growth factor receptor | Plasma Membrane | kinase | [illegible] cetuximab/erlotinib, CK-101, 18F-PEG6-IPQA, CLN-081, bis-tyrphostin, erlotinib/osimertinib, capecitabine/cetuximab/irinotecan, MCLA-158, neratinib, erlotinib, dasatinib/erlotinib, erlotinib/gemcitabine, BPI-15086, SCT200, BIBX 1382BS, docetaxel/erlotinib, ABBV-221, docetaxel/lapatinib, afatinib, afatinib/paclitaxel, cetuximab/vemurafenib, lazertinib, sym013, HLX07, BMS-690514, [illegible] |
| chr3 | *NUP210* | 51 | 2221 | 122 | NUP210 | nucleoporin 210 | Nucleus | transporter | |
| chr16 | *PLCG2* | 52 | 1602 | 122 | PLCG2 | phospholipase C gamma 2 | Cytoplasm | enzyme | |
| chr16 | *WWOX* | 53 | 1107 | 122 | WWOX | WW domain containing oxidoreductase | Cytoplasm | enzyme | |
| chr1 | *DNAH14* | 50 | 1528 | 122 | DNAH14 | dynein axonemal heavy chain 14 | Other | other | |
| chr6 | *PDE10A* | 49 | 1615 | 122 | PDE10A | phosphodiesterase 10A | Cytoplasm | enzyme | aspirin/dipyridamole/telmisartan, dipyridamole, aspirin/dipyridamole, |
| chr8 | *PTK2* | 50 | 1337 | 122 | PTK2 | protein tyrosine kinase 2 | Cytoplasm | kinase | conteltinib, lorlatinib, BI 853520, |
| chr16 | *HYDIN* | 52 | 1399 | 122 | HYDIN | HYDIN axonemal central pair apparatus protein | Cytoplasm | other | |
| chr5 | *AC005592.2* | 51 | 1345 | 122 | | | | | |
| chr1 | *HSPG2* | 54 | 2476 | 121 | HSPG2 | heparan sulfate proteoglycan 2 | Extracellular Space | enzyme | |
| chr10 | *ADAM12* | 51 | 1605 | 121 | ADAM12 | ADAM metallopeptidase domain | Plasma Membrane | peptidase | |
| chr3 | *ARHGEF3* | 55 | 1310 | 121 | ARHGEF3 | Rho guanine nucleotide exchange factor 3 | Cytoplasm | other | |
| chr19 | *CPAMD8* | 50 | 1699 | 121 | CPAMD8 | C3 and PZP like alpha-2-macroglobulin domain containing | Extracellular Space | other | |
| chr22 | *TTC28* | 48 | 1393 | 121 | TTC28 | tetratricopeptide repeat domain | Cytoplasm | other | |
| chr1 | *AJAP1* | 52 | 2053 | 121 | AJAP1 | adherens junctions associated protein 1 | Plasma Membrane | other | |
| chr17 | *KSR1* | 52 | 1961 | 121 | KSR1 | kinase suppressor of ras 1 | Cytoplasm | kinase | |
| chr13 | *HS6ST3* | 53 | 1424 | 120 | HS6ST3 | heparan sulfate 6-O-sulfotransferase 3 | Other | enzyme | |
| chr11 | *AMBRA1* | 49 | 1758 | 120 | AMBRA1 | autophagy and beclin 1 regulator | Cytoplasm | other | |
| chr15 | *OCA2* | 48 | 1963 | 120 | OCA2 | OCA2 melanosomal transmembrane protein | Plasma Membrane | transporter | |
| chr15 | *AKAP13* | 45 | 1555 | 120 | AKAP13 | A-kinase anchoring protein 13 | Cytoplasm | other | |
| chr2 | *USP34* | 47 | 1178 | 120 | USP34 | ubiquitin specific peptidase 34 | Extracellular Space | peptidase | |
| chr12 | *SPATS2* | 48 | 1944 | 119 | SPATS2 | spermatogenesis associated serine rich 2 | Cytoplasm | other | |
| chr9 | *C9orf171* | 53 | 2154 | 119 | CFAP77 | cilia and flagella associated | Other | other | |
| chr9 | *MAPKAP1* | 51 | 1656 | 119 | MAPKAP1 | MAPK associated protein 1 | Cytoplasm | other | |
| chr19 | *EPS15L1* | 50 | 1965 | 119 | EPS15L1 | epidermal growth factor receptor pathway substrate 15 like 1 | Plasma Membrane | other | |
| chr2 | *WDPCP* | 52 | 1270 | 118 | WDPCP | WD repeat containing planar cell polarity effector | Plasma Membrane | other | |
| chr1 | *SLC35F3* | 52 | 1658 | 118 | SLC35F3 | solute carrier family 35 member | Other | other | |
| chr8 | *ARHGAP39* | 53 | 2367 | 118 | ARHGAP39 | Rho GTPase activating protein 39 | Nucleus | other | |
| chr3 | *ATG7* | 51 | 1420 | 118 | ATG7 | autophagy related 7 | Cytoplasm | enzyme | |
| chr10 | *WDFY4* | 55 | 2584 | 118 | WDFY4 | WDFY family member 4 | Other | other | |
| chr1 | *PTPRF* | 48 | 2301 | 118 | PTPRF | protein tyrosine phosphatase receptor type F | Plasma Membrane | phosphatase | |
| chr11 | *TMEM135* | 44 | 1561 | 118 | TMEM135 | transmembrane protein 135 | Cytoplasm | other | |
| chr16 | *CRISPLD2* | 46 | 1916 | 117 | CRISPLD2 | cysteine rich secretory protein LCCL domain containing 2 | Cytoplasm | other | |
| chr1 | *CAPZB* | 50 | 1742 | 117 | CAPZB | capping actin protein of muscle Z-line subunit beta | Cytoplasm | other | |
| chr13 | *MYO16* | 50 | 1208 | 117 | MYO16 | myosin XVI | Nucleus | other | |
| chr1 | *KCNH1* | 51 | 1582 | 117 | KCNH1 | potassium voltage-gated channel subfamily H member 1 | Plasma Membrane | ion channel | |
| chr1 | *PLEKHA6* | 51 | 2606 | 117 | PLEKHA6 | pleckstrin homology domain containing A6 | Other | other | |
| chr12 | *DNAH10* | 48 | 1479 | 117 | DNAH10 | dynein axonemal heavy chain 10 | Cytoplasm | other | |

| | | | | | | | | | |
|---|---|---|---|---|---|---|---|---|---|
| chr1 | *NFIA* | 46 | 1186 | 117 | NFIA | nuclear factor I A | Nucleus | transcription regulator | |
| chr8 | *COL22A1* | 52 | 1484 | 117 | COL22A1 | collagen type XXII alpha 1 chain | Extracellular Space | other | collagenase clostridium |
| chr17 | *NXN* | 45 | 1714 | 116 | NXN | nucleoredoxin | Nucleus | enzyme | |
| chr1 | *NPHP4* | 49 | 2247 | 116 | NPHP4 | nephrocystin 4 | Cytoplasm | other | |
| chr9 | *MVB12B* | 50 | 2241 | 116 | MVB12B | multivesicular body subunit 12B | Cytoplasm | other | |
| chr1 | *PRKCZ* | 56 | 2601 | 116 | PRKCZ | protein kinase C zeta | Cytoplasm | kinase | ingenol mebutate, pkc-zeta inhibitor |
| chr7 | *PDE1C* | 52 | 1526 | 115 | PDE1C | phosphodiesterase 1C | Cytoplasm | enzyme | |
| chr15 | *ZNF609* | 44 | 1198 | 115 | ZNF609 | zinc finger protein 609 | Nucleus | other | |
| chr7 | *ADCY1* | 49 | 2474 | 115 | ADCY1 | adenylate cyclase 1 | Plasma Membrane | enzyme | |
| chr5 | *SH3PXD2B* | 50 | 2196 | 115 | SH3PXD2B | SH3 and PX domains 2B | Cytoplasm | other | |
| chr10 | *CPEB3* | 44 | 1082 | 115 | CPEB3 | cytoplasmic polyadenylation element binding protein 3 | Cytoplasm | translation regulator | |
| chr12 | *RBM19* | 47 | 1823 | 115 | RBM19 | RNA binding motif protein 19 | Nucleus | other | |
| chr11 | *LRRC4C* | 45 | 1087 | 114 | LRRC4C | leucine rich repeat containing 4C | Plasma Membrane | other | |
| chr17 | *RAI1* | 50 | 2454 | 114 | RAI1 | retinoic acid induced 1 | Cytoplasm | other | |
| chr6 | *PARK2* | 41 | 1187 | 114 | PRKN | parkin RBR E3 ubiquitin protein ligase | Cytoplasm | enzyme | |
| chr15 | *GABRG3* | 44 | 1433 | 113 | GABRG3 | gamma-aminobutyric acid type A receptor gamma3 subunit | Plasma Membrane | ion channel | [illegible] nitrazepam, sevoflurane, etomidate, pagoclone, zolpidem, phenobarbital, diazepam, isoflurane, halothane, pentobarbital, desflurane, adinazolam, SEP 174559, pregnenolone, zaleplon, tracazolate, enflurane, ezogabine, muscimol, flurazepam, metyrapone/oxazepam, isoniazid, midazolam, temazepam, diphenhydramine/midazolam, estazolam, [illegible] |
| chr12 | *HECTD4* | 46 | 1493 | 113 | HECTD4 | HECT domain E3 ubiquitin protein ligase 4 | Nucleus | other | |
| chr10 | *STK32C* | 53 | 2394 | 113 | STK32C | serine/threonine kinase 32C | Other | kinase | |
| chr19 | *CHST8* | 47 | 1823 | 113 | CHST8 | carbohydrate sulfotransferase 8 | Cytoplasm | enzyme | |
| chr3 | *CHCHD6* | 50 | 2217 | 113 | CHCHD6 | coiled-coil-helix-coiled-coil-helix domain containing 6 | Cytoplasm | other | |
| chr10 | *CNNM2* | 45 | 1223 | 113 | CNNM2 | cyclin and CBS domain divalent metal cation transport mediator 2 | Plasma Membrane | transporter | |
| chr3 | *MAGI1* | 50 | 1085 | 113 | MAGI1 | membrane associated guanylate kinase, WW and PDZ domain containing 1 | Plasma Membrane | kinase | |
| chr15 | *CCDC33* | 45 | 2042 | 113 | CCDC33 | coiled-coil domain containing 33 | Cytoplasm | other | |
| chr5 | *ARL15* | 42 | 1019 | 113 | ARL15 | ADP ribosylation factor like GTPase 15 | Cytoplasm | other | |
| chr11 | *PACS1* | 50 | 2252 | 113 | PACS1 | phosphofurin acidic cluster sorting protein 1 | Cytoplasm | other | |
| chr17 | *TAOK1* | 43 | 1095 | 113 | TAOK1 | TAO kinase 1 | Cytoplasm | kinase | |
| chr19 | *GNG7* | 42 | 1810 | 113 | GNG7 | G protein subunit gamma 7 | Plasma Membrane | enzyme | |
| chr7 | *CAMK2B* | 50 | 2965 | 113 | CAMK2B | calcium/calmodulin dependent protein kinase II beta | Cytoplasm | kinase | |
| chr20 | *SULF2* | 48 | 2391 | 113 | SULF2 | sulfatase 2 | Plasma Membrane | enzyme | |
| chr11 | *PC* | 52 | 3078 | 112 | PC | pyruvate carboxylase | Cytoplasm | enzyme | |
| chr6 | *DST* | 46 | 1094 | 112 | DST | dystonin | Plasma Membrane | other | |
| chr13 | *COL4A1* | 46 | 1763 | 112 | COL4A1 | collagen type IV alpha 1 chain | Extracellular Space | other | collagenase clostridium |
| chr1 | *PLD5* | 49 | 1422 | 112 | PLD5 | phospholipase D family member | Other | other | |
| chr3 | *ULK4* | 47 | 1206 | 112 | ULK4 | unc-51 like kinase 4 | Other | kinase | |
| chr3 | *MRPS22* | 48 | 1947 | 111 | MRPS22 | mitochondrial ribosomal protein | Cytoplasm | other | |
| chr7 | *SRPK2* | 45 | 1156 | 111 | SRPK2 | SRSF protein kinase 2 | Nucleus | kinase | |
| chr15 | *C15orf27* | 48 | 2241 | 111 | TMEM266 | transmembrane protein 266 | Plasma Membrane | other | |
| chrX | *SH3KBP1* | 46 | 1161 | 111 | SH3KBP1 | SH3 domain containing kinase binding protein 1 | Cytoplasm | other | |
| chr21 | *PCBP3* | 43 | 2045 | 111 | PCBP3 | poly(rC) binding protein 3 | Nucleus | transcription regulator | |
| chr17 | *DNAH9* | 48 | 1128 | 110 | DNAH9 | dynein axonemal heavy chain 9 | Cytoplasm | other | |
| chr3 | *MGLL* | 48 | 2285 | 110 | MGLL | monoglyceride lipase | Plasma Membrane | enzyme | |
| chr12 | *BTBD11* | 50 | 1224 | 110 | BTBD11 | BTB domain containing 11 | Other | transcription regulator | |
| chr12 | *FGD4* | 42 | 1127 | 110 | FGD4 | FYVE, RhoGEF and PH domain containing 4 | Cytoplasm | other | |
| chr7 | *SNX8* | 50 | 2400 | 110 | SNX8 | sorting nexin 8 | Cytoplasm | transporter | |
| chr7 | *AVL9* | 45 | 1430 | 110 | AVL9 | AVL9 cell migration associated | Other | other | |
| chr18 | *L3MBTL4* | 49 | 1974 | 110 | L3MBTL4 | L3MBTL histone methyl-lysine binding protein 4 | Other | other | |
| chr5 | *NSG2* | 48 | 1605 | 110 | NSG2 | neuronal vesicle trafficking associated 2 | Cytoplasm | other | |
| chr1 | *ST3GAL3* | 48 | 1391 | 110 | ST3GAL3 | ST3 beta-galactoside alpha-2,3-sialyltransferase 3 | Cytoplasm | enzyme | |

| | | | | | | | | | |
|---|---|---|---|---|---|---|---|---|---|
| chr8 | *DLC1* | 42 | 1022 | 110 | DLC1 | DLC1 Rho GTPase activating | Cytoplasm | other | |
| chr6 | *ATXN1* | 48 | 1049 | 109 | ATXN1 | ataxin 1 | Nucleus | transcription regulator | |
| chr1 | *MAST2* | 43 | 1202 | 109 | MAST2 | microtubule associated serine/threonine kinase 2 | Cytoplasm | kinase | |
| chr5 | *KCNIP1* | 50 | 2046 | 109 | KCNIP1 | potassium voltage-gated channel interacting protein 1 | Plasma Membrane | ion channel | |
| chr16 | *XYLT1* | 52 | 1103 | 109 | XYLT1 | xylosyltransferase 1 | Cytoplasm | enzyme | |
| chr19 | *KDM4B* | 52 | 2543 | 109 | KDM4B | lysine demethylase 4B | Nucleus | enzyme | |
| chr1 | *PEX14* | 45 | 1378 | 109 | PEX14 | peroxisomal biogenesis factor 14 | Cytoplasm | transcription regulator | |
| chr7 | *UPK3B* | 47 | 1727 | 109 | UPK3B | uroplakin 3B | Plasma Membrane | other | |
| chr7 | *TYW1* | 43 | 1106 | 109 | TYW1 | tRNA-yW synthesizing protein 1 homolog | Other | other | |
| chr7 | *IMMP2L* | 44 | 1060 | 109 | IMMP2L | inner mitochondrial membrane peptidase subunit 2 | Cytoplasm | peptidase | |
| chr15 | *RORA* | 47 | 989 | 108 | RORA | RAR related orphan receptor A | Nucleus | d-dependent nuclear rec | |
| chr3 | *MECOM* | 45 | 2051 | 108 | MECOM | MDS1 and EVI1 complex locus | Nucleus | transcription regulator | |
| chr7 | *FOXK1* | 48 | 2033 | 108 | FOXK1 | forkhead box K1 | Nucleus | transcription regulator | |
| chr20 | *EPB41L1* | 48 | 2164 | 108 | EPB41L1 | erythrocyte membrane protein band 4.1 like 1 | Plasma Membrane | other | |
| chr3 | *EPHA6* | 46 | 1172 | 108 | EPHA6 | EPH receptor A6 | Plasma Membrane | kinase | dacomitinib |
| chr2 | *SNTG2* | 38 | 1374 | 108 | SNTG2 | syntrophin gamma 2 | Plasma Membrane | other | |
| chr5 | *WWC1* | 47 | 1920 | 108 | WWC1 | WW and C2 domain containing 1 | Cytoplasm | transcription regulator | |
| chr22 | *C22orf34* | 43 | 1907 | 108 | C22orf34 | chromosome 22 open reading frame 34 | Other | other | |
| chr5 | *CTC-340A15.2* | 45 | 1065 | 108 | | | | | |
| chr19 | *MYH14* | 49 | 2068 | 108 | MYH14 | myosin heavy chain 14 | Extracellular Space | enzyme | |
| chr9 | *AK8* | 47 | 2156 | 108 | AK8 | adenylate kinase 8 | Cytoplasm | kinase | |
| chr7 | *ABCA13* | 45 | 1471 | 108 | ABCA13 | ATP binding cassette subfamily A member 13 | Extracellular Space | transporter | |
| chr5 | *NSD1* | 43 | 1132 | 108 | NSD1 | nuclear receptor binding SET domain protein 1 | Nucleus | transcription regulator | |
| chr2 | *TTC7A* | 44 | 2624 | 108 | TTC7A | tetratricopeptide repeat domain | Plasma Membrane | other | |
| chr8 | *CSGALNACT1* | 47 | 1213 | 108 | CSGALNACT1 | chondroitin sulfate N-acetylgalactosaminyltransferase | Cytoplasm | enzyme | |
| chr20 | *TOX2* | 45 | 2154 | 108 | TOX2 | TOX high mobility group box family member 2 | Nucleus | transcription regulator | |
| chr20 | *C20orf112* | 50 | 2632 | 107 | NOL4L | nucleolar protein 4 like | Other | other | |
| chr17 | *CCDC57* | 47 | 2043 | 107 | CCDC57 | coiled-coil domain containing 57 | Other | other | |
| chr17 | *DNAH2* | 45 | 1637 | 107 | DNAH2 | dynein axonemal heavy chain 2 | Cytoplasm | other | |
| chr11 | *STIM1* | 47 | 2221 | 107 | STIM1 | stromal interaction molecule 1 | Plasma Membrane | ion channel | |
| chr12 | *EP400* | 45 | 1893 | 107 | EP400 | E1A binding protein p400 | Nucleus | other | |
| chr7 | *SEMA3A* | 41 | 1397 | 107 | SEMA3A | semaphorin 3A | Extracellular Space | other | |
| chr3 | *PTPRG* | 46 | 1007 | 107 | PTPRG | protein tyrosine phosphatase receptor type G | Plasma Membrane | phosphatase | |
| chr17 | *CEP112* | 45 | 1104 | 107 | CEP112 | centrosomal protein 112 | Cytoplasm | other | |
| chr1 | *FMN2* | 40 | 1357 | 107 | FMN2 | formin 2 | Cytoplasm | other | |
| chr9 | *GABBR2* | 43 | 1281 | 107 | GABBR2 | gamma-aminobutyric acid type B receptor subunit 2 | Plasma Membrane | -protein coupled recepto | baclofen, arbaclofen, arbaclofen placarbil |
| chr12 | *CLIP1* | 44 | 1225 | 106 | CLIP1 | CAP-Gly domain containing linker protein 1 | Cytoplasm | other | |
| chr1 | *ZSWIM5* | 45 | 1062 | 106 | ZSWIM5 | zinc finger SWIM-type containing | Extracellular Space | other | |
| chr5 | *SEMA5A* | 49 | 2997 | 106 | SEMA5A | semaphorin 5A | Plasma Membrane | transmembrane recepto | |
| chr9 | *DAPK1* | 46 | 1488 | 106 | DAPK1 | death associated protein kinase 1 | Cytoplasm | kinase | |
| chr3 | *ITPR1* | 45 | 1262 | 106 | ITPR1 | inositol 1,4,5-trisphosphate receptor type 1 | Cytoplasm | ion channel | |
| chr16 | *CREBBP* | 42 | 1326 | 106 | CREBBP | CREB binding protein | Nucleus | transcription regulator | PRI-724, CCS1477 |
| chr17 | *TBC1D16* | 46 | 1994 | 106 | TBC1D16 | TBC1 domain family member 16 | Extracellular Space | other | |
| chr7 | *HIPK2* | 47 | 1916 | 106 | HIPK2 | homeodomain interacting protein kinase 2 | Nucleus | kinase | |
| chr7 | *TNRC18* | 49 | 2444 | 106 | TNRC18 | trinucleotide repeat containing 18 | Nucleus | other | |
| chr9 | *RXRA* | 54 | 3717 | 106 | RXRA | retinoid X receptor alpha | Nucleus | d-dependent nuclear rec | daunorubicin/tretinoin, alitretinoin, acitretin, bexarotene, doxorubicin/tretinoin, theophylline/tretinoin/valproic acid, etretinate, idarubicin/tretinoin, |
| chr15 | *CYFIP1* | 46 | 2138 | 105 | CYFIP1 | cytoplasmic FMR1 interacting protein 1 | Cytoplasm | translation regulator | |
| chr3 | *TFDP2* | 40 | 1271 | 105 | TFDP2 | transcription factor Dp-2 | Nucleus | transcription regulator | |
| chr1 | *MAN1C1* | 45 | 1584 | 105 | MAN1C1 | mannosidase alpha class 1C member 1 | Cytoplasm | enzyme | |
| chr12 | *DENND5B* | 41 | 1301 | 105 | DENND5B | DENN domain containing 5B | Cytoplasm | other | |
| chr8 | *ASAP1* | 46 | 1201 | 105 | ASAP1 | ArfGAP with SH3 domain, ankyrin repeat and PH domain 1 | Plasma Membrane | other | |
| chr7 | *GRB10* | 45 | 2017 | 105 | GRB10 | growth factor receptor bound protein 10 | Cytoplasm | other | |
| chr16 | *WDR59* | 43 | 1121 | 105 | WDR59 | WD repeat domain 59 | Cytoplasm | transporter | |
| chr10 | *C10orf76* | 40 | 999 | 105 | ARMH3 | armadillo like helical domain containing 3 | Other | other | |
| chr17 | *VPS53* | 45 | 1345 | 105 | VPS53 | VPS53 subunit of GARP complex | Cytoplasm | other | |
| chr9 | *LAMC3* | 47 | 2635 | 105 | LAMC3 | laminin subunit gamma 3 | Extracellular Space | other | |
| chr2 | *TNS1* | 39 | 1632 | 104 | TNS1 | tensin 1 | Plasma Membrane | other | |
| chr5 | *ADCY2* | 51 | 2872 | 104 | ADCY2 | adenylate cyclase 2 | Plasma Membrane | enzyme | |
| chr6 | *CCND3* | 41 | 1390 | 104 | CCND3 | cyclin D3 | Nucleus | kinase | |
| chr6 | *TIAM2* | 35 | 1008 | 104 | TIAM2 | T cell lymphoma invasion and metastasis 2 | Cytoplasm | enzyme | |
| chr11 | *UVRAG* | 39 | 1282 | 104 | UVRAG | UV radiation resistance | Nucleus | other | |

| | | | | | | | | | |
|---|---|---|---|---|---|---|---|---|---|
| chr12 | *ATXN2* | 39 | 1074 | 104 | ATXN2 | ataxin 2 | Nucleus | other | |
| chr16 | *JPH3* | 47 | 2257 | 104 | JPH3 | junctophilin 3 | Plasma Membrane | ion channel | |
| chr3 | *TMCC1* | 40 | 1207 | 104 | TMCC1 | transmembrane and coiled-coil domain family 1 | Cytoplasm | other | |
| chr1 | *CNIH3* | 48 | 1613 | 103 | CNIH3 | cornichon family AMPA receptor auxiliary protein 3 | Plasma Membrane | transporter | |
| chr8 | *AGO2* | 44 | 2190 | 103 | AGO2 | argonaute RISC catalytic component 2 | Cytoplasm | translation regulator | |
| chr15 | *AGBL1* | 50 | 1256 | 103 | AGBL1 | ATP/GTP binding protein like 1 | Other | peptidase | |
| chr10 | *DLG5* | 44 | 1799 | 103 | DLG5 | discs large MAGUK scaffold | Plasma Membrane | other | |
| chr15 | *SMAD3* | 43 | 1521 | 103 | SMAD3 | SMAD family member 3 | Nucleus | transcription regulator | |
| chr11 | *TEAD1* | 43 | 1427 | 103 | TEAD1 | TEA domain transcription factor 1 | Nucleus | transcription regulator | |
| chr12 | *GALNT9* | 49 | 2639 | 103 | GALNT9 | polypeptide N-acetylgalactosaminyltransferase | Cytoplasm | enzyme | |
| chr9 | *C9orf3* | 39 | 1137 | 103 | AOPEP | aminopeptidase O (putative) | Cytoplasm | peptidase | |
| chr11 | *PLEKHA7* | 45 | 1360 | 103 | PLEKHA7 | pleckstrin homology domain containing A7 | Cytoplasm | other | |
| chr2 | *CAMKMT* | 44 | 1202 | 103 | CAMKMT | calmodulin-lysine N-methyltransferase | Nucleus | enzyme | |
| chr7 | *TBXAS1* | 49 | 1652 | 103 | TBXAS1 | thromboxane A synthase 1 | Plasma Membrane | enzyme | ridogrel |
| chr11 | *TSPAN18* | 45 | 1614 | 103 | TSPAN18 | tetraspanin 18 | Other | other | |
| chr7 | *CARD11* | 46 | 2242 | 103 | CARD11 | caspase recruitment domain family member 11 | Cytoplasm | kinase | |
| chr19 | *NFIX* | 45 | 2139 | 103 | NFIX | nuclear factor I X | Nucleus | transcription regulator | |
| chr9 | *ROR2* | 36 | 1313 | 103 | ROR2 | receptor tyrosine kinase like orphan receptor 2 | Plasma Membrane | kinase | conditionally active biologic anti-ROR2 antibody drug conjugate |
| chr12 | *BICD1* | 38 | 999 | 103 | BICD1 | BICD cargo adaptor 1 | Cytoplasm | other | |
| chr20 | *CDH22* | 45 | 2155 | 102 | CDH22 | cadherin 22 | Plasma Membrane | other | |
| chr1 | *TRABD2B* | 45 | 1678 | 102 | TRABD2B | TraB domain containing 2B | Plasma Membrane | peptidase | |
| chr7 | *CDK14* | 43 | 1338 | 102 | CDK14 | cyclin dependent kinase 14 | Nucleus | kinase | |
| chr3 | *BSN* | 44 | 1461 | 102 | BSN | bassoon presynaptic cytomatrix protein | Plasma Membrane | other | |
| chr3 | *CNTN4* | 40 | 951 | 102 | CNTN4 | contactin 4 | Plasma Membrane | enzyme | |
| chr5 | *SPOCK1* | 49 | 1011 | 102 | SPOCK1 | SPARC (osteonectin), cwcv and kazal like domains proteoglycan 1 | Extracellular Space | other | |
| chr19 | *GATAD2A* | 45 | 1625 | 101 | GATAD2A | GATA zinc finger domain containing 2A | Nucleus | transcription regulator | |
| chr7 | *POU6F2* | 46 | 1260 | 101 | POU6F2 | POU class 6 homeobox 2 | Nucleus | transcription regulator | |
| chr10 | *LHPP* | 48 | 2145 | 101 | LHPP | phospholysine phosphohistidine inorganic pyrophosphate phosphatase | Cytoplasm | phosphatase | |
| chr15 | *ITGA11* | 41 | 1614 | 101 | ITGA11 | integrin subunit alpha 11 | Plasma Membrane | other | |
| chr2 | *CNTNAP5* | 46 | 1246 | 101 | CNTNAP5 | contactin associated protein like | Other | other | |
| chrX | *SHROOM2* | 43 | 1418 | 101 | SHROOM2 | shroom family member 2 | Plasma Membrane | ion channel | |
| chr6 | *CPNE5* | 42 | 1588 | 101 | CPNE5 | copine 5 | Plasma Membrane | other | |
| chr15 | *RYR3* | 41 | 947 | 101 | RYR3 | ryanodine receptor 3 | Plasma Membrane | ion channel | |
| chr20 | *CBFA2T2* | 41 | 1479 | 101 | CBFA2T2 | CBFA2/RUNX1 partner transcriptional co-repressor 2 | Nucleus | transcription regulator | |
| chr16 | *PARN* | 44 | 1074 | 101 | PARN | poly(A)-specific ribonuclease | Nucleus | enzyme | |
| chr13 | *STK24* | 41 | 1902 | 101 | STK24 | serine/threonine kinase 24 | Cytoplasm | kinase | bosutinib |
| chr11 | *RP11-179A16.1* | 44 | 1296 | 101 | | | | | |
| chr3 | *PIK3CB* | 39 | 1292 | 101 | PIK3CB | phosphatidylinositol-4,5-bisphosphate 3-kinase catalytic subunit beta | Cytoplasm | kinase | dactolisib, acalisib, BAY1082439, pilaralisib, KA2237, AZD8186, PX-866, SF 1126, pictilisib, CLR457, bimiralisib, SAR260301, GSK2636771, buparlisib |
| chr7 | *BBS9* | 40 | 1262 | 101 | BBS9 | Bardet-Biedl syndrome 9 | Extracellular Space | other | |
| chr1 | *KIF1B* | 39 | 1030 | 101 | KIF1B | kinesin family member 1B | Cytoplasm | transporter | |
| chr10 | *SUFU* | 42 | 1262 | 100 | SUFU | SUFU negative regulator of hedgehog signaling | Nucleus | transcription regulator | |
| chr20 | *TAF4* | 47 | 3195 | 100 | TAF4 | TATA-box binding protein associated factor 4 | Nucleus | transcription regulator | |
| chr12 | *C12orf55* | 42 | 958 | 100 | CFAP54 | cilia and flagella associated | Cytoplasm | other | |
| chr2 | *PCBP1-AS1* | 42 | 1348 | 100 | PCBP1-AS1 | PCBP1 antisense RNA 1 | Other | other | |
| chr3 | *SMARCC1* | 39 | 981 | 100 | SMARCC1 | SWI/SNF related, matrix associated, actin dependent regulator of chromatin subfamily c | Nucleus | transcription regulator | |
| chr2 | *SFXN5* | 42 | 1749 | 100 | SFXN5 | sideroflexin 5 | Cytoplasm | transporter | |
| chr19 | *FBN3* | 43 | 1630 | 100 | FBN3 | fibrillin 3 | Extracellular Space | other | |
| chr8 | *PEBP4* | 43 | 1265 | 100 | PEBP4 | phosphatidylethanolamine binding protein 4 | Cytoplasm | other | |
| chr17 | *USP32* | 39 | 1013 | 100 | USP32 | ubiquitin specific peptidase 32 | Cytoplasm | enzyme | |
| chr16 | *RAB11FIP3* | 43 | 1963 | 100 | RAB11FIP3 | RAB11 family interacting protein | Cytoplasm | other | |
| chr15 | *FAM174B* | 43 | 2186 | 100 | FAM174B | family with sequence similarity 174 member B | Other | other | |
| chr3 | *KCNAB1* | 44 | 1304 | 99 | KCNAB1 | potassium voltage-gated channel subfamily A member regulatory beta subunit 1 | Plasma Membrane | ion channel | |
| chr5 | *SGCD* | 43 | 1198 | 99 | SGCD | sarcoglycan delta | Cytoplasm | other | |
| chr5 | *FAM172A* | 38 | 992 | 99 | FAM172A | family with sequence similarity 172 member A | Cytoplasm | transcription regulator | |
| chr7 | *SND1* | 42 | 1211 | 99 | SND1 | staphylococcal nuclease and tudor domain containing 1 | Nucleus | enzyme | |

| | | | | | | | | | |
|---|---|---|---|---|---|---|---|---|---|
| chr9 | *SEMA4D* | 41 | 1874 | 99 | SEMA4D | semaphorin 4D | Plasma Membrane | transmembrane receptor | pepinemab |
| chr15 | *CORO2B* | 42 | 1496 | 99 | CORO2B | coronin 2B | Plasma Membrane | other | |
| chr17 | *MAPT* | 39 | 1464 | 99 | MAPT | microtubule associated protein | Plasma Membrane | other | |
| chr5 | *MCTP1* | 41 | 925 | 99 | MCTP1 | multiple C2 and transmembrane domain containing 1 | Cytoplasm | other | |
| chr7 | *TMEM178B* | 45 | 1142 | 99 | TMEM178B | transmembrane protein 178B | Other | other | |
| chr7 | *CHN2* | 44 | 1227 | 99 | CHN2 | chimerin 2 | Cytoplasm | other | |
| chr3 | *CADM2* | 42 | 911 | 99 | CADM2 | cell adhesion molecule 2 | Plasma Membrane | other | |
| chr9 | *KDM4C* | 41 | 1084 | 99 | KDM4C | lysine demethylase 4C | Nucleus | enzyme | |
| chr16 | *GNAO1* | 42 | 1672 | 98 | GNAO1 | G protein subunit alpha o1 | Plasma Membrane | enzyme | |
| chr10 | *HK1* | 44 | 2237 | 98 | HK1 | hexokinase 1 | Cytoplasm | kinase | |
| chr5 | *ZNF131* | 42 | 3530 | 98 | ZNF131 | zinc finger protein 131 | Nucleus | transcription regulator | |
| chr16 | *RP11-426C22.5* | 42 | 1914 | 98 | | | | | |
| chr7 | *JAZF1* | 41 | 1406 | 98 | JAZF1 | JAZF zinc finger 1 | Nucleus | transcription regulator | |
| chr9 | *HMCN2* | 44 | 2891 | 98 | HMCN2 | hemicentin 2 | Extracellular Space | other | |
| chr15 | *RP11-321F6.1* | 42 | 1439 | 98 | | | | | |
| chr1 | *PHACTR4* | 39 | 1030 | 98 | PHACTR4 | phosphatase and actin regulator | Plasma Membrane | other | |
| chr22 | *SCUBE1* | 39 | 1762 | 98 | SCUBE1 | signal peptide, CUB domain and EGF like domain containing 1 | Plasma Membrane | transmembrane receptor | |
| chr1 | *TTC34* | 42 | 13542 | 98 | TTC34 | tetratricopeptide repeat domain | Other | other | |
| chr1 | *OBSCN* | 48 | 2810 | 98 | OBSCN | obscurin, cytoskeletal calmodulin and titin-interacting RhoGEF | Cytoplasm | kinase | |
| chr2 | *NPAS2* | 42 | 2368 | 97 | NPAS2 | neuronal PAS domain protein 2 | Nucleus | transcription regulator | |
| chr7 | *KMT2C* | 39 | 1266 | 97 | KMT2C | lysine methyltransferase 2C | Nucleus | transcription regulator | |
| chr10 | *NRG3* | 43 | 897 | 97 | NRG3 | neuregulin 3 | Extracellular Space | growth factor | |
| chr6 | *SLC22A23* | 38 | 1409 | 97 | SLC22A23 | solute carrier family 22 member | Other | transporter | |
| chr10 | *ATRNL1* | 40 | 951 | 97 | ATRNL1 | attractin like 1 | Other | other | |
| chr8 | *DLGAP2* | 41 | 1706 | 97 | DLGAP2 | DLG associated protein 2 | Plasma Membrane | other | |
| chr3 | *XXYLT1* | 38 | 1516 | 97 | XXYLT1 | xyloside xylosyltransferase 1 | Cytoplasm | enzyme | |
| chr1 | *KCNAB2* | 44 | 2131 | 97 | KCNAB2 | potassium voltage-gated channel subfamily A regulatory beta | Plasma Membrane | ion channel | |
| chr11 | *ANO1* | 41 | 1980 | 97 | ANO1 | anoctamin 1 | Plasma Membrane | ion channel | crofelemer |
| chr18 | *LAMA1* | 36 | 1195 | 97 | LAMA1 | laminin subunit alpha 1 | Extracellular Space | other | |
| chr20 | *ITCH* | 39 | 1164 | 97 | ITCH | itchy E3 ubiquitin protein ligase | Nucleus | enzyme | |
| chr11 | *AMPD3* | 45 | 1746 | 97 | AMPD3 | adenosine monophosphate deaminase 3 | Cytoplasm | enzyme | |
| chr20 | *FAM65C* | 42 | 2146 | 97 | RIPOR3 | RIPOR family member 3 | Other | other | |
| chr17 | *PIGL* | 40 | 1043 | 97 | PIGL | phosphatidylinositol glycan anchor biosynthesis class L | Cytoplasm | enzyme | |
| chr19 | *CTC-525D6.1* | 43 | 1466 | 97 | | | | | |
| chr3 | *ZBTB20* | 40 | 1119 | 97 | ZBTB20 | zinc finger and BTB domain containing 20 | Nucleus | transcription regulator | |
| chr17 | *FBXL20* | 37 | 1005 | 97 | FBXL20 | F-box and leucine rich repeat protein 20 | Cytoplasm | other | |
| chr7 | *CACNA2D1* | 37 | 998 | 97 | CACNA2D1 | calcium voltage-gated channel auxiliary subunit alpha2delta 1 | Plasma Membrane | ion channel | nifedipine, amlodipine/indapamide/perindopril, amlodipine/perindopril, amlodipine/hydrochlorothiazide/olmesartan medoxomil, felodipine, amlodipine/hydrochlorothiazide/valsartan, gabapentin, bepridil, nitrendipine, amlodipine, nisoldipine, amlodipine/benazepril, nilvadipine, amlodipine/celecoxib, enalapril/felodipine, pregabalin, amlodipine/valsartan, amlodipine/ezetimibe/losartan/rosuvastatin [illegible] |
| chr7 | *TPST1* | 39 | 994 | 97 | TPST1 | tyrosylprotein sulfotransferase 1 | Cytoplasm | enzyme | |
| chr1 | *PCNXL2* | 42 | 1347 | 96 | PCNX2 | pecanex 2 | Other | other | |
| chr11 | *SIK3* | 37 | 927 | 96 | SIK3 | SIK family kinase 3 | Cytoplasm | kinase | |
| chr17 | *PEMT* | 41 | 1940 | 96 | PEMT | phosphatidylethanolamine N-methyltransferase | Cytoplasm | enzyme | |
| chr16 | *VAC14* | 40 | 1937 | 96 | VAC14 | VAC14 component of PIKFYVE complex | Cytoplasm | other | |
| chr9 | *BNC2* | 36 | 978 | 96 | BNC2 | basonuclin 2 | Nucleus | transcription regulator | |
| chr17 | *SSH2* | 37 | 918 | 96 | SSH2 | slingshot protein phosphatase 2 | Cytoplasm | phosphatase | |
| chr9 | *NR6A1* | 38 | 1352 | 96 | NR6A1 | nuclear receptor subfamily 6 group A member 1 | Nucleus | d-dependent nuclear rec | |
| chr8 | *SGCZ* | 39 | 831 | 96 | SGCZ | sarcoglycan zeta | Plasma Membrane | other | |
| chr11 | *AHNAK* | 40 | 1371 | 96 | AHNAK | AHNAK nucleoprotein | Nucleus | other | |
| chr10 | *TTC40* | 44 | 1903 | 96 | CFAP46 | cilia and flagella associated | Cytoplasm | other | |
| chr1 | *CROCC* | 45 | 2728 | 96 | CROCC | ciliary rootlet coiled-coil, rootletin | Plasma Membrane | other | |

| | | | | | | | | | |
|---|---|---|---|---|---|---|---|---|---|
| chr1 | *CHRM3* | 40 | 1117 | 96 | CHRM3 | cholinergic receptor muscarinic 3 | Plasma Membrane | G-protein coupled recepto | dexamethasone/olanzapine, dicyclomine, cevimeline, mirabegron/solifenacin, olanzapine, aclidinium/formoterol, acetylcholine, atropine/pralidoxime, brompheniramine, umeclidinium, mepenzolic acid, darifenacin, propantheline, orphenadrine, aspirin/caffeine/orphenadrine, tolterodine, chlorpheniramine/methscopolamine/phenylephrine, trihexyphenidyl, oxybutynin, formoterol/glycopyrrolate, scopolamine, [illegible] |
| chr10 | *MCU* | 39 | 1382 | 96 | MCU | mitochondrial calcium uniporter | Cytoplasm | ion channel | |
| chr9 | *EXD3* | 45 | 2175 | 96 | EXD3 | exonuclease 3'-5' domain containing 3 | Other | other | |
| chr1 | *GALNT2* | 41 | 1745 | 96 | GALNT2 | polypeptide N-acetylgalactosaminyltransferase | Cytoplasm | enzyme | |
| chr1 | *PAX7* | 41 | 1629 | 95 | PAX7 | paired box 7 | Nucleus | transcription regulator | |
| chr16 | *MYH11* | 44 | 1542 | 95 | MYH11 | myosin heavy chain 11 | Cytoplasm | other | |
| chr10 | *PALD1* | 42 | 2664 | 95 | PALD1 | phosphatase domain containing paladin 1 | Cytoplasm | phosphatase | |
| chr16 | *GLG1* | 36 | 1070 | 95 | GLG1 | golgi glycoprotein 1 | Cytoplasm | other | |
| chr3 | *SCN5A* | 40 | 1582 | 95 | SCN5A | sodium voltage-gated channel alpha subunit 5 | Plasma Membrane | ion channel | lidocaine/prilocaine, epinephrine/lidocaine, bupivacaine/lidocaine, encainide, epinephrine/prilocaine, tocainide, quinidine, aprindine, oxcarbazepine, mephenytoin, propafenone, flecainide, ajmaline, disopyramide, riluzole, moricizine, prilocaine, zonisamide, cocaine, procainamide, amiodarone, ranolazine, dibucaine, carbamazepine |
| chr11 | *ARRB1* | 41 | 2105 | 95 | ARRB1 | arrestin beta 1 | Cytoplasm | other | |
| chr5 | *GPR98* | 41 | 1158 | 95 | ADGRV1 | adhesion G protein-coupled receptor V1 | Plasma Membrane | G-protein coupled recepto | |
| chr1 | *LGR6* | 42 | 1808 | 95 | LGR6 | leucine rich repeat containing G protein-coupled receptor 6 | Plasma Membrane | G-protein coupled recepto | |
| chr2 | *AC131097.3* | 37 | 1878 | 95 | | | | | |
| chr2 | *NRXN1* | 41 | 1027 | 95 | NRXN1 | neurexin 1 | Plasma Membrane | transporter | |
| chr20 | *NCOA3* | 37 | 1185 | 95 | NCOA3 | nuclear receptor coactivator 3 | Nucleus | transcription regulator | |
| chr12 | *ANO2* | 38 | 963 | 94 | ANO2 | anoctamin 2 | Plasma Membrane | ion channel | |
| chr7 | *TRRAP* | 41 | 1979 | 94 | TRRAP | transformation/transcription domain associated protein | Nucleus | transcription regulator | |
| chr3 | *OSBPL10* | 43 | 989 | 94 | OSBPL10 | oxysterol binding protein like 10 | Cytoplasm | transporter | |
| chr20 | *ZMYND8* | 41 | 1485 | 94 | ZMYND8 | zinc finger MYND-type containing | Nucleus | transcription regulator | |
| chr11 | *KDM2A* | 38 | 1094 | 94 | KDM2A | lysine demethylase 2A | Nucleus | enzyme | |
| chr1 | *GATAD2B* | 40 | 1273 | 94 | GATAD2B | GATA zinc finger domain containing 2B | Nucleus | transcription regulator | |
| chr10 | *PTPRE* | 40 | 1555 | 94 | PTPRE | protein tyrosine phosphatase receptor type E | Plasma Membrane | phosphatase | |
| chr5 | *NRG2* | 40 | 1354 | 94 | NRG2 | neuregulin 2 | Extracellular Space | growth factor | |
| chr17 | *CYTH1* | 39 | 1226 | 94 | CYTH1 | cytohesin 1 | Cytoplasm | other | |
| chr17 | *ARHGAP23* | 39 | 1744 | 94 | ARHGAP23 | Rho GTPase activating protein 23 | Cytoplasm | other | |
| chr17 | *AC003051.1* | 36 | 920 | 94 | | | | | |
| chr12 | *CIT* | 40 | 1039 | 94 | CIT | citron rho-interacting serine/threonine kinase | Cytoplasm | kinase | |
| chr3 | *FBLN2* | 41 | 2060 | 94 | FBLN2 | fibulin 2 | Extracellular Space | other | |
| chr11 | *MICAL2* | 43 | 1923 | 94 | MICAL2 | microtubule associated monooxygenase, calponin and LIM domain containing 2 | Cytoplasm | enzyme | |
| chr17 | *KANSL1* | 34 | 977 | 94 | KANSL1 | KAT8 regulatory NSL complex subunit 1 | Nucleus | other | |
| chr1 | *FHAD1* | 39 | 1318 | 94 | FHAD1 | forkhead associated phosphopeptide binding domain 1 | Other | other | |

| | | | | | | | | | |
|---|---|---|---|---|---|---|---|---|---|
| chr5 | *MAST4* | 42 | 913 | 94 | MAST4 | microtubule associated serine/threonine kinase family member 4 | Other | kinase | |
| chr11 | *LDLRAD3* | 44 | 1424 | 93 | LDLRAD3 | low density lipoprotein receptor class A domain containing 3 | Plasma Membrane | other | |
| chr1 | *RPRD2* | 35 | 1278 | 93 | RPRD2 | regulation of nuclear pre-mRNA domain containing 2 | Nucleus | other | |
| chr7 | *CREB5* | 40 | 1206 | 93 | CREB5 | cAMP responsive element binding protein 5 | Nucleus | transcription regulator | |
| chr11 | *NUP98* | 37 | 1721 | 93 | NUP98 | nucleoporin 98 | Nucleus | transporter | |
| chr6 | *SMOC2* | 33 | 1340 | 93 | SMOC2 | SPARC related modular calcium binding 2 | Extracellular Space | other | |
| chr10 | *ANK3* | 40 | 1257 | 93 | ANK3 | ankyrin 3 | Plasma Membrane | other | |
| chr6 | *ZFAND3* | 35 | 912 | 93 | ZFAND3 | zinc finger AN1-type containing 3 | Other | other | |
| chr20 | *PIGU* | 38 | 1460 | 93 | PIGU | phosphatidylinositol glycan anchor biosynthesis class U | Cytoplasm | enzyme | |
| chr1 | *KCND3* | 43 | 1599 | 93 | KCND3 | potassium voltage-gated channel subfamily D member 3 | Plasma Membrane | ion channel | dalfampridine |
| chr1 | *MEGF6* | 48 | 2331 | 93 | MEGF6 | multiple EGF like domains 6 | Cytoplasm | other | |
| chr16 | *RANBP10* | 39 | 1313 | 93 | RANBP10 | RAN binding protein 10 | Cytoplasm | other | |
| chr15 | *MYO1E* | 37 | 948 | 93 | MYO1E | myosin IE | Cytoplasm | enzyme | |
| chr9 | *CNTLN* | 37 | 1001 | 93 | CNTLN | centlein | Cytoplasm | other | |
| chr5 | *KIAA0825* | 37 | 901 | 93 | KIAA0825 | KIAA0825 | Extracellular Space | other | |
| chrX | *BCOR* | 42 | 1435 | 92 | BCOR | BCL6 corepressor | Nucleus | transcription regulator | |
| chr13 | *DOCK9* | 37 | 1268 | 92 | DOCK9 | dedicator of cytokinesis 9 | Cytoplasm | other | |
| chr13 | *ABCC4* | 42 | 1300 | 92 | ABCC4 | ATP binding cassette subfamily C member 4 | Plasma Membrane | transporter | |
| chr12 | *RPH3A* | 43 | 957 | 92 | RPH3A | rabphilin 3A | Plasma Membrane | transporter | |
| chr7 | *AC011288.2* | 37 | 1068 | 92 | | | | | |
| chr22 | *CABIN1* | 37 | 1388 | 92 | CABIN1 | calcineurin binding protein 1 | Nucleus | other | |
| chr6 | *UST* | 36 | 1039 | 92 | UST | uronyl 2-sulfotransferase | Cytoplasm | enzyme | |
| chr17 | *PITPNM3* | 40 | 1680 | 92 | PITPNM3 | PITPNM family member 3 | Cytoplasm | enzyme | |
| chr17 | *PPM1E* | 35 | 841 | 92 | PPM1E | protein phosphatase, Mg2+/Mn2+ dependent 1E | Nucleus | phosphatase | |
| chr17 | *FOXK2* | 38 | 1770 | 92 | FOXK2 | forkhead box K2 | Nucleus | transcription regulator | |
| chr1 | *FGGY* | 40 | 1156 | 92 | FGGY | FGGY carbohydrate kinase domain containing | Other | other | |
| chr19 | *CRTC1* | 41 | 1774 | 92 | CRTC1 | CREB regulated transcription coactivator 1 | Nucleus | transcription regulator | sapanisertib, CC-115 |
| chr20 | *MYT1* | 40 | 2961 | 91 | MYT1 | myelin transcription factor 1 | Nucleus | transcription regulator | |
| chr3 | *FRMD4B* | 40 | 929 | 91 | FRMD4B | FERM domain containing 4B | Cytoplasm | other | |
| chr6 | *GMDS-AS1* | 37 | 1114 | 91 | GMDS-DT | GMDS divergent transcript | Other | other | |
| chr8 | *TG* | 41 | 1146 | 91 | TG | thyroglobulin | Extracellular Space | other | |
| chr3 | *FLNB* | 38 | 1279 | 91 | FLNB | filamin B | Cytoplasm | other | |
| chr8 | *MYOM2* | 38 | 1592 | 91 | MYOM2 | myomesin 2 | Cytoplasm | other | |
| chr1 | *GON4L* | 37 | 1245 | 91 | GON4L | gon-4 like | Cytoplasm | transcription regulator | |
| chr3 | *ZNF385D* | 41 | 996 | 91 | ZNF385D | zinc finger protein 385D | Nucleus | other | |
| chr7 | *FBXL18* | 40 | 2170 | 91 | FBXL18 | F-box and leucine rich repeat protein 18 | Cytoplasm | enzyme | |
| chr17 | *MGAT5B* | 38 | 1694 | 91 | MGAT5B | alpha-1,6-mannosylglycoprotein 6-beta-N-acetylglucosaminyltransferase B | Cytoplasm | enzyme | |
| chr11 | *ANO3* | 35 | 1474 | 91 | ANO3 | anoctamin 3 | Plasma Membrane | transporter | |
| chr2 | *TSSC1* | 35 | 1497 | 91 | EIPR1 | EARP complex and GARP complex interacting protein 1 | Cytoplasm | other | |
| chr7 | *CYTH3* | 38 | 1531 | 91 | CYTH3 | cytohesin 3 | Cytoplasm | other | |
| chr20 | *RP4-723E3.1* | 42 | 1157 | 91 | | | | | |
| chr21 | *RUNX1* | 36 | 892 | 91 | RUNX1 | RUNX family transcription factor | Nucleus | transcription regulator | |
| chr10 | *UNC5B* | 41 | 2940 | 91 | UNC5B | unc-5 netrin receptor B | Plasma Membrane | transmembrane receptor | |
| chr13 | *FAM155A* | 39 | 855 | 90 | FAM155A | family with sequence similarity 155 member A | Other | other | |
| chr3 | *EIF2B5* | 35 | 1139 | 90 | EIF2B5 | eukaryotic translation initiation factor 2B subunit epsilon | Cytoplasm | other | |
| chr8 | *ANK1* | 35 | 1271 | 90 | ANK1 | ankyrin 1 | Plasma Membrane | other | |
| chr19 | *SPTBN4* | 41 | 1327 | 90 | SPTBN4 | spectrin beta, non-erythrocytic 4 | Cytoplasm | other | |
| chr7 | *LINC01006* | 38 | 1236 | 90 | LINC01006 | long intergenic non-protein coding RNA 1006 | Other | other | |
| chr3 | *DNAH1* | 38 | 1776 | 90 | DNAH1 | dynein axonemal heavy chain 1 | Extracellular Space | other | |
| chr1 | *PPP1R12B* | 37 | 1162 | 90 | PPP1R12B | protein phosphatase 1 regulatory subunit 12B | Cytoplasm | phosphatase | |
| chr9 | *COL27A1* | 36 | 1477 | 90 | COL27A1 | collagen type XXVII alpha 1 chain | Extracellular Space | other | collagenase clostridium |
| chr15 | *MEF2A* | 36 | 1512 | 90 | MEF2A | myocyte enhancer factor 2A | Nucleus | transcription regulator | |
| chr1 | *ABCA4* | 39 | 1569 | 90 | ABCA4 | ATP binding cassette subfamily A member 4 | Plasma Membrane | transporter | |
| chr10 | *ASCC1* | 34 | 1378 | 90 | ASCC1 | activating signal cointegrator 1 complex subunit 1 | Nucleus | transcription regulator | |
| chr22 | *PACSIN2* | 34 | 1264 | 90 | PACSIN2 | protein kinase C and casein kinase substrate in neurons 2 | Cytoplasm | transporter | |
| chr10 | *ADAMTS14* | 41 | 2564 | 90 | ADAMTS14 | ADAM metallopeptidase with thrombospondin type 1 motif 14 | Extracellular Space | peptidase | |
| chr22 | *BCR* | 33 | 1400 | 89 | BCR | BCR activator of RhoGEF and GTPase | Cytoplasm | kinase | BCR targeting agent |
| chr5 | *C5orf42* | 39 | 2520 | 89 | CPLANE1 | ciliogenesis and planar polarity effector 1 | Other | other | |
| chr3 | *IQCJ-SCHIP1* | 38 | 1213 | 89 | IQCJ-SCHIP1 | IQCJ-SCHIP1 readthrough | Cytoplasm | other | |
| chr17 | *NPLOC4* | 37 | 1294 | 89 | NPLOC4 | NPL4 homolog, ubiquitin recognition factor | Nucleus | other | |
| chr8 | *GFRA2* | 36 | 1428 | 89 | GFRA2 | GDNF family receptor alpha 2 | Plasma Membrane | transmembrane receptor | |

| | | | | | | | | | |
|---|---|---|---|---|---|---|---|---|---|
| chr15 | *DAPK2* | 39 | 1088 | 89 | DAPK2 | death associated protein kinase 2 | Cytoplasm | kinase | |
| chr6 | *LRFN2* | 39 | 1146 | 89 | LRFN2 | leucine rich repeat and fibronectin type III domain containing 2 | Plasma Membrane | other | |
| chr1 | *CLSTN1* | 36 | 1178 | 89 | CLSTN1 | calsyntenin 1 | Plasma Membrane | other | |
| chr17 | *NF1* | 36 | 861 | 89 | NF1 | neurofibromin 1 | Cytoplasm | other | |
| chr7 | *DGKB* | 36 | 995 | 88 | DGKB | diacylglycerol kinase beta | Cytoplasm | kinase | |
| chr9 | *KANK1* | 37 | 986 | 88 | KANK1 | KN motif and ankyrin repeat domains 1 | Nucleus | transcription regulator | |
| chr12 | *VWF* | 31 | 1169 | 88 | VWF | von Willebrand factor | Extracellular Space | other | caplacizumab |
| chr2 | *ARHGEF4* | 34 | 1368 | 88 | ARHGEF4 | Rho guanine nucleotide exchange factor 4 | Cytoplasm | other | |
| chr5 | *RNF130* | 36 | 1470 | 88 | RNF130 | ring finger protein 130 | Cytoplasm | peptidase | |
| chr15 | *ALDH1A2* | 36 | 892 | 88 | ALDH1A2 | aldehyde dehydrogenase 1 family member A2 | Cytoplasm | enzyme | |
| chr12 | *TMEM117* | 37 | 1348 | 88 | TMEM117 | transmembrane protein 117 | Cytoplasm | other | |
| chr22 | *PHF21B* | 31 | 1407 | 88 | PHF21B | PHD finger protein 21B | Nucleus | other | |
| chr2 | *ACYP2* | 38 | 1118 | 88 | ACYP2 | acylphosphatase 2 | Cytoplasm | enzyme | |
| chr17 | *RNF213* | 37 | 1520 | 88 | RNF213 | ring finger protein 213 | Cytoplasm | enzyme | |
| chr15 | *HERC2* | 33 | 1208 | 88 | HERC2 | HECT and RLD domain containing E3 ubiquitin protein | Cytoplasm | enzyme | |
| chr1 | *XPR1* | 35 | 1037 | 88 | XPR1 | xenotropic and polytropic retrovirus receptor 1 | Plasma Membrane | G-protein coupled recepto | |
| chr1 | *CDC42BPA* | 34 | 973 | 88 | CDC42BPA | CDC42 binding protein kinase | Cytoplasm | kinase | |
| chr12 | *ACACB* | 38 | 1214 | 88 | ACACB | acetyl-CoA carboxylase beta | Cytoplasm | enzyme | |
| chr3 | *TRAK1* | 39 | 1115 | 88 | TRAK1 | trafficking kinesin protein 1 | Nucleus | other | |
| chr5 | *FBXL7* | 45 | 2369 | 88 | FBXL7 | F-box and leucine rich repeat protein 7 | Cytoplasm | enzyme | |
| chr10 | *ZMIZ1-AS1* | 37 | 1648 | 88 | ZMIZ1-AS1 | ZMIZ1 antisense RNA 1 | Other | other | |
| chr14 | *TTC7B* | 32 | 1225 | 88 | TTC7B | tetratricopeptide repeat domain | Other | other | |
| chr1 | *UBE4B* | 37 | 1022 | 87 | UBE4B | ubiquitination factor E4B | Cytoplasm | enzyme | |
| chr17 | *SKAP1* | 35 | 843 | 87 | SKAP1 | src kinase associated phosphoprotein 1 | Cytoplasm | kinase | |
| chr3 | *VGLL4* | 36 | 1204 | 87 | VGLL4 | vestigial like family member 4 | Nucleus | other | |
| chr19 | *PTPRS* | 36 | 1389 | 87 | PTPRS | protein tyrosine phosphatase receptor type S | Plasma Membrane | phosphatase | |
| chr17 | *ARSG* | 38 | 1079 | 87 | ARSG | arylsulfatase G | Cytoplasm | enzyme | |
| chrX | *DHRSX* | 30 | 886 | 87 | DHRSX | dehydrogenase/reductase X- | Extracellular Space | enzyme | |
| chr2 | *ALK* | 34 | 925 | 87 | ALK | ALK receptor tyrosine kinase | Plasma Membrane | kinase | gilteritinib, belizatinib, crizotinib/vemurafenib, TQ-B3139, ensartinib, alkotinib, WX-0593, crizotinib/prednisolone, brigatinib, CEP-37440, ASP3026, crizotinib/erlotinib, crizotinib/pazopanib, ALK inhibitor, PLB1003, ceritinib, crizotinib/gefitinib, conteltinib, lorlatinib, alectinib, entrectinib, [illegible] |
| chr7 | *UBE2H* | 34 | 852 | 87 | UBE2H | ubiquitin conjugating enzyme E2 | Cytoplasm | enzyme | |
| chr11 | *GDPD5* | 39 | 2325 | 87 | GDPD5 | glycerophosphodiester phosphodiesterase domain containing 5 | Plasma Membrane | enzyme | |
| chr17 | *MYO1D* | 37 | 990 | 87 | MYO1D | myosin ID | Cytoplasm | enzyme | |
| chr7 | *RP11-3P22.2* | 39 | 925 | 87 | | | | | |
| chr3 | *ITGB5* | 38 | 1552 | 87 | ITGB5 | integrin subunit beta 5 | Plasma Membrane | other | cilengitide |
| chr20 | *ZBTB46* | 41 | 2419 | 87 | ZBTB46 | zinc finger and BTB domain containing 46 | Nucleus | transcription regulator | |
| chr17 | *EPN2* | 36 | 1238 | 87 | EPN2 | epsin 2 | Cytoplasm | other | |
| chr10 | *CRTAC1* | 38 | 1398 | 87 | CRTAC1 | cartilage acidic protein 1 | Extracellular Space | other | |
| chr10 | *EBF3* | 37 | 1470 | 87 | EBF3 | EBF transcription factor 3 | Nucleus | transcription regulator | |
| chr5 | *UBE2D2* | 35 | 926 | 87 | UBE2D2 | ubiquitin conjugating enzyme E2 | Cytoplasm | enzyme | |
| chr3 | *PRKAR2A* | 34 | 912 | 87 | PRKAR2A | protein kinase cAMP-dependent type II regulatory subunit alpha | Cytoplasm | kinase | |
| chr15 | *PPCDC* | 34 | 1195 | 86 | PPCDC | phosphopantothenoylcysteine decarboxylase | Cytoplasm | enzyme | |

| | | | | | | | | | |
|---|---|---|---|---|---|---|---|---|---|
| chr7 | *POR* | 38 | 1738 | 86 | POR | cytochrome p450 oxidoreductase | Cytoplasm | enzyme | carboplatin/doxorubicin, nitazoxanide, cisplatin/doxorubicin/ifosfamide, cisplatin/doxorubicin, daunorubicin/tretinoin, cisplatin/doxorubicin/methotrexate, daunorubicin, doxorubicin/streptozocin, dacarbazine/doxorubicin, doxorubicin/ifosfamide, dacarbazine/doxorubicin/ifosfamide, cisplatin/doxorubicin/ifosfamide/methotrexate, doxorubicin, doxorubicin/tretinoin |
| chr8 | *DENND3* | 36 | 1756 | 86 | DENND3 | DENN domain containing 3 | Cytoplasm | other | |
| chr9 | *ADAMTSL1* | 35 | 953 | 86 | ADAMTSL1 | ADAMTS like 1 | Extracellular Space | other | |
| chr3 | *WWTR1* | 35 | 1164 | 86 | WWTR1 | WW domain containing transcription regulator 1 | Nucleus | transcription regulator | |
| chr16 | *NLRC5* | 35 | 1356 | 86 | NLRC5 | NLR family CARD domain containing 5 | Cytoplasm | transcription regulator | |
| chr7 | *KIAA1549* | 38 | 1464 | 86 | KIAA1549 | KIAA1549 | Plasma Membrane | other | |
| chr7 | *DENND2A* | 39 | 1514 | 86 | DENND2A | DENN domain containing 2A | Cytoplasm | other | |
| chr9 | *PRRC2B* | 38 | 1797 | 86 | PRRC2B | proline rich coiled-coil 2B | Other | other | |
| chr8 | *KCNK9* | 37 | 1420 | 86 | KCNK9 | potassium two pore domain channel subfamily K member 9 | Plasma Membrane | ion channel | doxapram, halothane |
| chr9 | *CCDC171* | 36 | 953 | 86 | CCDC171 | coiled-coil domain containing 171 | Other | other | |
| chr9 | *TRPM3* | 38 | 801 | 86 | TRPM3 | transient receptor potential cation channel subfamily M member 3 | Plasma Membrane | ion channel | |
| chr8 | *MROH1* | 37 | 1459 | 86 | MROH1 | maestro heat like repeat family member 1 | Other | other | |
| chr5 | *RP11-445O3.2* | 41 | 2389 | 86 | | | | | |
| chr9 | *CAMSAP1* | 38 | 1605 | 86 | CAMSAP1 | calmodulin regulated spectrin associated protein 1 | Cytoplasm | other | |
| chr17 | *RTN4RL1* | 37 | 1961 | 86 | RTN4RL1 | reticulon 4 receptor like 1 | Plasma Membrane | other | |
| chr2 | *INPP4A* | 38 | 1707 | 86 | INPP4A | inositol polyphosphate-4-phosphatase type I A | Cytoplasm | phosphatase | |
| chr14 | *CDC42BPB* | 32 | 1229 | 86 | CDC42BPB | CDC42 binding protein kinase | Cytoplasm | kinase | |
| chr16 | *FANCA* | 36 | 1452 | 86 | FANCA | FA complementation group A | Nucleus | other | |
| chr20 | *BCAS4* | 37 | 1893 | 85 | BCAS4 | breast carcinoma amplified sequence 4 | Cytoplasm | other | |
| chr5 | *LHFPL2* | 37 | 985 | 85 | LHFPL2 | LHFPL tetraspan subfamily member 2 | Other | enzyme | |
| chr11 | *DENND5A* | 33 | 1151 | 85 | DENND5A | DENN domain containing 5A | Cytoplasm | other | |
| chr4 | *JAKMIP1* | 33 | 1127 | 85 | JAKMIP1 | janus kinase and microtubule interacting protein 1 | Cytoplasm | other | |
| chr8 | *ZFAT* | 38 | 1105 | 85 | ZFAT | zinc finger and AT-hook domain containing | Nucleus | transcription regulator | |
| chr12 | *PPM1H* | 36 | 948 | 85 | PPM1H | protein phosphatase, Mg2+/Mn2+ dependent 1H | Nucleus | phosphatase | |
| chr11 | *NELL1* | 41 | 1091 | 85 | NELL1 | neural EGFL like 1 | Extracellular Space | growth factor | |
| chr15 | *RASGRF1* | 38 | 1864 | 85 | RASGRF1 | Ras protein specific guanine nucleotide releasing factor 1 | Cytoplasm | other | |
| chr10 | *TCF7L2* | 37 | 885 | 85 | TCF7L2 | transcription factor 7 like 2 | Nucleus | transcription regulator | |
| chr8 | *CTD-2281E23.2* | 35 | 1481 | 85 | | | | | |
| chr22 | *PARVB* | 33 | 1226 | 85 | PARVB | parvin beta | Cytoplasm | other | |
| chr17 | *BAIAP2* | 42 | 2153 | 85 | BAIAP2 | BAR/IMD domain containing adaptor protein 2 | Plasma Membrane | kinase | |
| chr1 | *SPSB1* | 34 | 1624 | 85 | SPSB1 | splA/ryanodine receptor domain and SOCS box containing 1 | Cytoplasm | other | |
| chr16 | *HS3ST4* | 42 | 728 | 85 | HS3ST4 | heparan sulfate-glucosamine 3-sulfotransferase 4 | Cytoplasm | enzyme | |
| chr17 | *SEC14L1* | 34 | 1177 | 85 | SEC14L1 | SEC14 like lipid binding 1 | Cytoplasm | transporter | |
| chr9 | *NEK6* | 38 | 2338 | 85 | NEK6 | NIMA related kinase 6 | Nucleus | kinase | |
| chrX | *FRMPD4* | 37 | 856 | 85 | FRMPD4 | FERM and PDZ domain | Plasma Membrane | other | |
| chr17 | *VMP1* | 31 | 855 | 84 | VMP1 | vacuole membrane protein 1 | Plasma Membrane | other | |
| chr7 | *AHCYL2* | 33 | 811 | 84 | AHCYL2 | adenosylhomocysteinase like 2 | Other | enzyme | |
| chr8 | *DOCK5* | 36 | 859 | 84 | DOCK5 | dedicator of cytokinesis 5 | Cytoplasm | other | |
| chr20 | *KCNB1* | 39 | 1502 | 84 | KCNB1 | potassium voltage-gated channel subfamily B member 1 | Plasma Membrane | ion channel | dalfampridine |
| chr10 | *VTI1A* | 35 | 1015 | 84 | VTI1A | vesicle transport through interaction with t-SNAREs 1A | Plasma Membrane | transporter | |
| chr16 | *PPL* | 37 | 1661 | 84 | PPL | periplakin | Cytoplasm | other | |
| chr3 | *RP11-23D24.2* | 32 | 951 | 84 | | | | | |
| chr11 | *MAML2* | 34 | 1668 | 84 | MAML2 | mastermind like transcriptional coactivator 2 | Nucleus | transcription regulator | |
| chr16 | *ZNRF1* | 34 | 1022 | 84 | ZNRF1 | zinc and ring finger 1 | Cytoplasm | enzyme | |
| chr8 | *SLC45A4* | 36 | 1747 | 84 | SLC45A4 | solute carrier family 45 member 4 | Other | transporter | |
| chr3 | *FGD5* | 35 | 1239 | 84 | FGD5 | FYVE, RhoGEF and PH domain containing 5 | Cytoplasm | other | |
| chr3 | *SLC9A9* | 37 | 1023 | 84 | SLC9A9 | solute carrier family 9 member A9 | Cytoplasm | transporter | |

| | | | | | | | | | |
|---|---|---|---|---|---|---|---|---|---|
| chr2 | *COMMD1* | 36 | 899 | 84 | COMMD1 | copper metabolism domain containing 1 | Nucleus | transporter | |
| chr1 | *C1orf222* | 38 | 1691 | 84 | CFAP74 | cilia and flagella associated | Cytoplasm | other | |
| chr8 | *RBPMS* | 33 | 1002 | 84 | RBPMS | RNA binding protein, mRNA processing factor | Cytoplasm | transcription regulator | |
| chr10 | *USP54* | 32 | 1307 | 84 | USP54 | ubiquitin specific peptidase 54 | Other | peptidase | |
| chr17 | *FAM222B* | 33 | 1029 | 84 | FAM222B | family with sequence similarity 222 member B | Nucleus | other | |
| chr20 | *SOGA1* | 37 | 1829 | 84 | SOGA1 | suppressor of glucose, autophagy associated 1 | Extracellular Space | other | |
| chr10 | *TET1* | 37 | 1354 | 84 | TET1 | tet methylcytosine dioxygenase 1 | Nucleus | enzyme | |
| chr2 | *KIAA1211L* | 37 | 1493 | 84 | KIAA1211L | KIAA1211 like | Other | other | |
| chr20 | *ZFP64* | 36 | 1285 | 84 | ZFP64 | ZFP64 zinc finger protein | Nucleus | transcription regulator | |
| chr8 | *KCNQ3* | 38 | 813 | 83 | KCNQ3 | potassium voltage-gated channel subfamily Q member 3 | Plasma Membrane | ion channel | ezogabine |
| chr4 | *ZFYVE28* | 34 | 1528 | 83 | ZFYVE28 | zinc finger FYVE-type containing | Cytoplasm | other | |
| chr9 | *PNPLA7* | 38 | 1988 | 83 | PNPLA7 | patatin like phospholipase domain containing 7 | Cytoplasm | enzyme | |
| chr1 | *NMNAT2* | 39 | 1281 | 83 | NMNAT2 | nicotinamide nucleotide adenylyltransferase 2 | Cytoplasm | enzyme | |
| chr15 | *ATP10A* | 33 | 1287 | 83 | ATP10A | ATPase phospholipid transporting 10A (putative) | Plasma Membrane | transporter | |
| chr3 | *SLMAP* | 32 | 827 | 83 | SLMAP | sarcolemma associated protein | Plasma Membrane | other | |
| chr8 | *EXTL3* | 36 | 949 | 83 | EXTL3 | exostosin like glycosyltransferase | Cytoplasm | enzyme | |
| chr9 | *FAM129B* | 37 | 2071 | 83 | NIBAN2 | niban apoptosis regulator 2 | Cytoplasm | transcription regulator | |
| chr19 | *ELL* | 34 | 1716 | 83 | ELL | elongation factor for RNA polymerase II | Nucleus | transcription regulator | |
| chr7 | *ZAN* | 34 | 1267 | 83 | ZAN | zonadhesin (gene/pseudogene) | Plasma Membrane | other | |
| chr17 | *B3GNTL1* | 37 | 1980 | 83 | B3GNTL1 | UDP-GlcNAc:betaGal beta-1,3-N-acetylglucosaminyltransferase | Other | enzyme | |
| chr3 | *SLC6A6* | 32 | 1542 | 83 | SLC6A6 | solute carrier family 6 member 6 | Plasma Membrane | transporter | |
| chr2 | *ARMC9* | 31 | 932 | 83 | ARMC9 | armadillo repeat containing 9 | Cytoplasm | other | |
| chr17 | *BPTF* | 33 | 804 | 82 | BPTF | bromodomain PHD finger transcription factor | Nucleus | transcription regulator | |
| chr13 | *ARHGEF7* | 33 | 1278 | 82 | ARHGEF7 | Rho guanine nucleotide exchange factor 7 | Cytoplasm | other | |
| chr21 | *PDE9A* | 33 | 1405 | 82 | PDE9A | phosphodiesterase 9A | Cytoplasm | enzyme | |
| chr7 | *VOPP1* | 35 | 1627 | 82 | VOPP1 | VOPP1 WW domain binding | Nucleus | other | |
| chr11 | *PDE2A* | 37 | 2165 | 82 | PDE2A | phosphodiesterase 2A | Cytoplasm | enzyme | |
| chr16 | *LITAF* | 34 | 1137 | 82 | LITAF | lipopolysaccharide induced TNF factor | Nucleus | transcription regulator | |
| chr16 | *RP11-525K10.3* | 36 | 825 | 82 | | | | | |
| chr14 | *WDR25* | 29 | 1085 | 82 | WDR25 | WD repeat domain 25 | Other | other | |
| chr17 | *TLK2* | 32 | 873 | 82 | TLK2 | tousled like kinase 2 | Cytoplasm | kinase | |
| chr17 | *SPECC1* | 35 | 915 | 82 | SPECC1 | sperm antigen with calponin homology and coiled-coil domains | Nucleus | other | |
| chr1 | *SYT2* | 37 | 1631 | 82 | SYT2 | synaptotagmin 2 | Cytoplasm | transporter | |
| chr22 | *MYH9* | 33 | 1390 | 82 | MYH9 | myosin heavy chain 9 | Cytoplasm | enzyme | |
| chr10 | *KAT6B* | 34 | 1243 | 82 | KAT6B | lysine acetyltransferase 6B | Nucleus | enzyme | |
| chr9 | *RP11-343J18.2* | 35 | 1483 | 82 | | | | | |
| chr9 | *GPR107* | 35 | 1240 | 82 | GPR107 | G protein-coupled receptor 107 | Plasma Membrane | -protein coupled recepto | |
| chr17 | *SRCIN1* | 36 | 1595 | 82 | SRCIN1 | SRC kinase signaling inhibitor 1 | Cytoplasm | other | |
| chr12 | *AACS* | 33 | 1528 | 82 | AACS | acetoacetyl-CoA synthetase | Cytoplasm | enzyme | |
| chr1 | *ZFYVE9* | 33 | 814 | 81 | ZFYVE9 | zinc finger FYVE-type containing | Cytoplasm | peptidase | |
| chr16 | *ATP2C2* | 33 | 1086 | 81 | ATP2C2 | ATPase secretory pathway Ca2+ transporting 2 | Cytoplasm | enzyme | |
| chr6 | *JARID2* | 33 | 1036 | 81 | JARID2 | jumonji and AT-rich interaction domain containing 2 | Nucleus | transcription regulator | |
| chr9 | *NTNG2* | 36 | 1980 | 81 | NTNG2 | netrin G2 | Plasma Membrane | other | |
| chr1 | *NOS1AP* | 35 | 1266 | 81 | NOS1AP | nitric oxide synthase 1 adaptor protein | Cytoplasm | other | |
| chr7 | *SRRM3* | 36 | 1660 | 81 | SRRM3 | serine/arginine repetitive matrix 3 | Other | other | |
| chr1 | *SKI* | 37 | 1871 | 81 | SKI | SKI proto-oncogene | Nucleus | transcription regulator | |
| chr15 | *FMN1* | 32 | 779 | 81 | FMN1 | formin 1 | Nucleus | other | |
| chr12 | *PPHLN1* | 33 | 1255 | 81 | PPHLN1 | periphilin 1 | Nucleus | other | |
| chr1 | *USH2A* | 36 | 906 | 81 | USH2A | usherin | Extracellular Space | other | |
| chr15 | *RP11-624L4.1* | 32 | 768 | 81 | | | | | |
| chr22 | *SYN3* | 30 | 884 | 81 | SYN3 | synapsin III | Plasma Membrane | other | |
| chr2 | *EML6* | 34 | 1026 | 81 | EML6 | EMAP like 6 | Other | other | |
| chr11 | *OTOG* | 34 | 1594 | 81 | OTOG | otogelin | Extracellular Space | enzyme | |
| chr10 | *RBM20* | 37 | 1403 | 81 | RBM20 | RNA binding motif protein 20 | Nucleus | other | |
| chr15 | *LRRK1* | 34 | 1949 | 81 | LRRK1 | leucine rich repeat kinase 1 | Cytoplasm | kinase | |
| chr8 | *KIF13B* | 32 | 942 | 80 | KIF13B | kinesin family member 13B | Cytoplasm | other | |
| chr15 | *MAP2K5* | 34 | 855 | 80 | MAP2K5 | mitogen-activated protein kinase kinase 5 | Cytoplasm | kinase | |
| chr1 | *EVI5* | 30 | 755 | 80 | EVI5 | ecotropic viral integration site 5 | Cytoplasm | other | |
| chr10 | *ABLIM1* | 35 | 826 | 80 | ABLIM1 | actin binding LIM protein 1 | Cytoplasm | other | |
| chr19 | *UNC13A* | 35 | 1267 | 80 | UNC13A | unc-13 homolog A | Plasma Membrane | other | |
| chr1 | *GNG4* | 35 | 1482 | 80 | GNG4 | G protein subunit gamma 4 | Plasma Membrane | enzyme | |
| chr20 | *NCOA6* | 34 | 1131 | 80 | NCOA6 | nuclear receptor coactivator 6 | Nucleus | transcription regulator | |
| chr3 | *CPNE4* | 37 | 1023 | 80 | CPNE4 | copine 4 | Cytoplasm | other | |
| chr14 | *CCDC88C* | 30 | 1354 | 80 | CCDC88C | coiled-coil domain containing 88C | Cytoplasm | other | |
| chr1 | *TMCO4* | 35 | 1203 | 80 | TMCO4 | transmembrane and coiled-coil domains 4 | Other | other | |
| chr2 | *HPCAL1* | 31 | 1469 | 80 | HPCAL1 | hippocalcin like 1 | Cytoplasm | other | |
| chr8 | *ARHGEF10* | 36 | 1545 | 80 | ARHGEF10 | Rho guanine nucleotide exchange factor 10 | Cytoplasm | enzyme | |
| chr9 | *TMC1* | 35 | 790 | 80 | TMC1 | transmembrane channel like 1 | Plasma Membrane | ion channel | |

| | | | | | | | | | |
|---|---|---|---|---|---|---|---|---|---|
| chr18 | *NEDD4L* | 29 | 953 | 80 | NEDD4L | NEDD4 like E3 ubiquitin protein ligase | Cytoplasm | enzyme | |
| chr7 | *SKAP2* | 32 | 931 | 80 | SKAP2 | src kinase associated phosphoprotein 2 | Cytoplasm | other | |
| chr2 | *AC073283.4* | 36 | 1322 | 80 | | | | | |
| chr3 | *DNAH12* | 33 | 761 | 80 | DNAH12 | dynein axonemal heavy chain 12 | Cytoplasm | other | |
| chr17 | *ZZEF1* | 34 | 993 | 80 | ZZEF1 | zinc finger ZZ-type and EF-hand domain containing 1 | Other | other | |
| chr9 | *MED27* | 34 | 1446 | 80 | MED27 | mediator complex subunit 27 | Nucleus | transcription regulator | |
| chr2 | *TET3* | 34 | 1139 | 80 | TET3 | tet methylcytosine dioxygenase 3 | Nucleus | enzyme | |
| chr1 | *PTPRU* | 36 | 1683 | 80 | PTPRU | protein tyrosine phosphatase receptor type U | Plasma Membrane | phosphatase | |
| chr20 | *EFCAB8* | 36 | 1563 | 80 | EFCAB8 | EF-hand calcium binding domain | Other | other | |
| chr16 | *CDH3* | 33 | 1152 | 80 | CDH3 | cadherin 3 | Plasma Membrane | other | PCA062, PF-06671008, FF21101, PF-3732010 |
| chr15 | *RP11-382A20.4* | 35 | 1182 | 80 | | | | | |
| chr5 | *IQGAP2* | 33 | 773 | 80 | IQGAP2 | IQ motif containing GTPase activating protein 2 | Cytoplasm | other | |
| chr13 | *PIBF1* | 31 | 805 | 79 | PIBF1 | progesterone immunomodulatory binding factor 1 | Nucleus | other | |
| chr12 | *LRP1* | 35 | 1800 | 79 | LRP1 | LDL receptor related protein 1 | Plasma Membrane | transmembrane receptor | paclitaxel-angiopep-2 conjugate, serpin peptide 16 |
| chr15 | *SLC28A1* | 35 | 1521 | 79 | SLC28A1 | solute carrier family 28 member 1 | Plasma Membrane | transporter | |
| chr3 | *POC1A* | 33 | 1429 | 79 | POC1A | POC1 centriolar protein A | Cytoplasm | peptidase | |
| chr17 | *TOM1L2* | 35 | 1122 | 79 | TOM1L2 | target of myb1 like 2 membrane trafficking protein | Cytoplasm | transporter | |
| chr1 | *RASAL2* | 32 | 1019 | 79 | RASAL2 | RAS protein activator like 2 | Extracellular Space | other | |
| chr8 | *RP11-317N12.1* | 36 | 736 | 79 | | | | | |
| chr21 | *AGPAT3* | 33 | 1480 | 79 | AGPAT3 | 1-acylglycerol-3-phosphate O-acyltransferase 3 | Cytoplasm | enzyme | |
| chr17 | *SHISA6* | 37 | 802 | 79 | SHISA6 | shisa family member 6 | Plasma Membrane | other | |
| chr2 | *NCK2* | 34 | 1197 | 79 | NCK2 | NCK adaptor protein 2 | Cytoplasm | kinase | |
| chr15 | *FAM81A* | 31 | 814 | 79 | FAM81A | family with sequence similarity 81 member A | Other | other | |
| chr9 | *GAPVD1* | 34 | 999 | 79 | GAPVD1 | GTPase activating protein and VPS9 domains 1 | Cytoplasm | other | |
| chr21 | *HSF2BP* | 33 | 995 | 79 | HSF2BP | heat shock transcription factor 2 binding protein | Cytoplasm | other | |
| chr6 | *KIF6* | 35 | 884 | 79 | KIF6 | kinesin family member 6 | Nucleus | other | |
| chr3 | *RYK* | 34 | 1230 | 79 | RYK | receptor like tyrosine kinase | Plasma Membrane | kinase | |
| chr13 | *KLF12* | 31 | 727 | 79 | KLF12 | Kruppel like factor 12 | Nucleus | transcription regulator | |
| chr7 | *UBE3C* | 30 | 856 | 79 | UBE3C | ubiquitin protein ligase E3C | Nucleus | enzyme | |
| chr10 | *CPXM2* | 34 | 1033 | 79 | CPXM2 | carboxypeptidase X, M14 family member 2 | Extracellular Space | peptidase | |
| chr1 | *TESK2* | 33 | 794 | 79 | TESK2 | testis associated actin remodelling kinase 2 | Nucleus | kinase | |
| chr1 | *SCMH1* | 32 | 885 | 79 | SCMH1 | Scm polycomb group protein homolog 1 | Nucleus | transcription regulator | |
| chr8 | *ZC3H3* | 39 | 1943 | 79 | ZC3H3 | zinc finger CCCH-type containing | Nucleus | transcription regulator | |
| chr17 | *TEX14* | 34 | 851 | 79 | TEX14 | testis expressed 14, intercellular bridge forming factor | Plasma Membrane | kinase | |
| chr17 | *TMEM104* | 33 | 1491 | 78 | TMEM104 | transmembrane protein 104 | Other | transporter | |
| chrX | *MID1* | 32 | 804 | 78 | MID1 | midline 1 | Nucleus | other | |
| chr3 | *SLC12A8* | 34 | 1316 | 78 | SLC12A8 | solute carrier family 12 member 8 | Other | transporter | |
| chr1 | *ALPL* | 33 | 1407 | 78 | ALPL | alkaline phosphatase, biomineralization associated | Plasma Membrane | phosphatase | zinc sulfate |
| chr8 | *SPIDR* | 26 | 758 | 78 | SPIDR | scaffold protein involved in DNA repair | Nucleus | other | |
| chr12 | *SPPL3* | 27 | 713 | 78 | SPPL3 | signal peptide peptidase like 3 | Cytoplasm | peptidase | |
| chr7 | *IGF2BP3* | 31 | 986 | 78 | IGF2BP3 | insulin like growth factor 2 mRNA binding protein 3 | Cytoplasm | translation regulator | |
| chr7 | *HUS1* | 33 | 1292 | 78 | HUS1 | HUS1 checkpoint clamp | Nucleus | kinase | |
| chr3 | *PDIA5* | 34 | 1487 | 78 | PDIA5 | protein disulfide isomerase family A member 5 | Cytoplasm | enzyme | |
| chr1 | *TNR* | 38 | 1151 | 78 | TNR | tenascin R | Plasma Membrane | other | |
| chr4 | *APBB2* | 29 | 750 | 78 | APBB2 | amyloid beta precursor protein binding family B member 2 | Cytoplasm | other | |
| chr2 | *VWA3B* | 33 | 1166 | 78 | VWA3B | von Willebrand factor A domain containing 3B | Other | other | |
| chr6 | *CDYL* | 33 | 942 | 78 | CDYL | chromodomain Y like | Nucleus | transcription regulator | |
| chr16 | *CNGB1* | 32 | 1337 | 78 | CNGB1 | cyclic nucleotide gated channel subunit beta 1 | Plasma Membrane | ion channel | |
| chr3 | *LIMD1* | 32 | 1122 | 78 | LIMD1 | LIM domains containing 1 | Nucleus | transcription regulator | |
| chr15 | *FRMD5* | 33 | 782 | 78 | FRMD5 | FERM domain containing 5 | Plasma Membrane | other | |
| chr6 | *FKBP5* | 31 | 809 | 78 | FKBP5 | FKBP prolyl isomerase 5 | Nucleus | enzyme | |
| chr6 | *KIF13A* | 31 | 759 | 77 | KIF13A | kinesin family member 13A | Cytoplasm | transporter | |
| chr7 | *C7orf50* | 38 | 2179 | 77 | C7orf50 | chromosome 7 open reading frame 50 | Other | other | |
| chr5 | *PDE8B* | 34 | 881 | 77 | PDE8B | phosphodiesterase 8B | Cytoplasm | enzyme | ketotifen |
| chr16 | *CPNE2* | 31 | 1390 | 77 | CPNE2 | copine 2 | Cytoplasm | other | |
| chrX | *TM4SF2* | 33 | 882 | 77 | TSPAN7 | tetraspanin 7 | Plasma Membrane | other | |
| chr12 | *SFSWAP* | 30 | 1258 | 77 | SFSWAP | splicing factor SWAP | Nucleus | other | |
| chr12 | *TAOK3* | 27 | 760 | 77 | TAOK3 | TAO kinase 3 | Cytoplasm | kinase | |
| chr8 | *SCARA5* | 31 | 1276 | 77 | SCARA5 | scavenger receptor class A member 5 | Plasma Membrane | transmembrane receptor | |
| chr3 | *MYRIP* | 36 | 903 | 77 | MYRIP | myosin VIIA and Rab interacting protein | Cytoplasm | other | |

| | | | | | | | | | |
|---|---|---|---|---|---|---|---|---|---|
| chr19 | *RHPN2* | 32 | 1049 | 77 | RHPN2 | rhophilin Rho GTPase binding protein 2 | Cytoplasm | other | |
| chr3 | *IRAK2* | 32 | 1235 | 77 | IRAK2 | interleukin 1 receptor associated kinase 2 | Plasma Membrane | kinase | |
| chr11 | *CLPB* | 34 | 1293 | 77 | CLPB | ClpB homolog, mitochondrial AAA ATPase chaperonin | Nucleus | transcription regulator | |
| chr5 | *ARHGEF28* | 32 | 788 | 77 | ARHGEF28 | Rho guanine nucleotide exchange factor 28 | Cytoplasm | other | |
| chr19 | *DNM2* | 29 | 1155 | 77 | DNM2 | dynamin 2 | Plasma Membrane | enzyme | |
| chr21 | *TSPEAR* | 32 | 1243 | 77 | TSPEAR | thrombospondin type laminin G domain and EAR repeats | Extracellular Space | other | |
| chr16 | *SMPD3* | 34 | 1337 | 77 | SMPD3 | sphingomyelin | Cytoplasm | enzyme | |
| chr5 | *RASGEF1C* | 33 | 1752 | 77 | RASGEF1C | RasGEF domain family member | Other | other | |
| chr16 | *DNAH3* | 36 | 711 | 77 | DNAH3 | dynein axonemal heavy chain 3 | Extracellular Space | enzyme | |
| chr6 | *PTK7* | 32 | 1157 | 77 | PTK7 | protein tyrosine kinase 7 | Plasma Membrane | kinase | cofetuzumab |
| chr11 | *C11orf80* | 33 | 1291 | 77 | C11orf80 | chromosome 11 open reading frame 80 | Other | other | |
| chr7 | *OGDH* | 34 | 1330 | 77 | OGDH | oxoglutarate dehydrogenase | Cytoplasm | enzyme | |
| chr5 | *ADAMTS6* | 28 | 743 | 77 | ADAMTS6 | ADAM metallopeptidase with thrombospondin type 1 motif 6 | Extracellular Space | peptidase | |
| chr2 | *LRRTM4* | 31 | 723 | 77 | LRRTM4 | leucine rich repeat transmembrane neuronal 4 | Extracellular Space | other | |
| chr1 | *COL24A1* | 32 | 816 | 76 | COL24A1 | collagen type XXIV alpha 1 chain | Extracellular Space | other | collagenase clostridium |
| chr12 | *RILPL1* | 31 | 1152 | 76 | RILPL1 | Rab interacting lysosomal protein like 1 | Cytoplasm | other | |
| chr10 | *LINC00842* | 38 | 3723 | 76 | LINC00842 | long intergenic non-protein coding RNA 842 | Other | other | |
| chr16 | *NFAT5* | 27 | 686 | 76 | NFAT5 | nuclear factor of activated T cells | Nucleus | transcription regulator | |
| chr11 | *PHF21A* | 29 | 1045 | 76 | PHF21A | PHD finger protein 21A | Nucleus | other | |
| chr17 | *CDK12* | 31 | 841 | 76 | CDK12 | cyclin dependent kinase 12 | Nucleus | kinase | |
| chr16 | *ZCCHC14* | 32 | 1365 | 76 | ZCCHC14 | zinc finger CCHC-type containing | Other | other | |
| chr16 | *CHD9* | 30 | 760 | 76 | CHD9 | chromodomain helicase DNA binding protein 9 | Cytoplasm | other | |
| chr11 | *USP2-AS1* | 33 | 1148 | 76 | USP2-AS1 | USP2 antisense RNA 1 (head to head) | Other | other | |
| chr6 | *EXOC2* | 29 | 949 | 76 | EXOC2 | exocyst complex component 2 | Cytoplasm | transporter | |
| chr16 | *SHISA9* | 36 | 683 | 76 | SHISA9 | shisa family member 9 | Plasma Membrane | other | |
| chr10 | *FRMD4A* | 29 | 823 | 76 | FRMD4A | FERM domain containing 4A | Plasma Membrane | other | |
| chr4 | *TENM3* | 25 | 696 | 76 | TENM3 | teneurin transmembrane protein 3 | Plasma Membrane | other | |
| chr3 | *SLCO2A1* | 34 | 1473 | 76 | SLCO2A1 | solute carrier organic anion transporter family member 2A1 | Plasma Membrane | transporter | |
| chr17 | *NCOR1* | 30 | 736 | 76 | NCOR1 | nuclear receptor corepressor 1 | Nucleus | transcription regulator | |
| chr11 | *MYO7A* | 33 | 2305 | 76 | MYO7A | myosin VIIA | Cytoplasm | enzyme | |
| chr22 | *KIAA1671* | 28 | 996 | 76 | KIAA1671 | KIAA1671 | Other | other | |
| chr11 | *UBASH3B* | 32 | 855 | 76 | UBASH3B | ubiquitin associated and SH3 domain containing B | Other | enzyme | |
| chr17 | *MYO15A* | 32 | 1595 | 76 | MYO15A | myosin XVA | Cytoplasm | other | |
| chr11 | *CELF1* | 31 | 1580 | 76 | CELF1 | CUGBP Elav-like family member | Cytoplasm | translation regulator | |
| chr11 | *CPT1A* | 34 | 1270 | 75 | CPT1A | carnitine palmitoyltransferase 1A | Cytoplasm | enzyme | perhexiline |
| chr11 | *DISC1FP1* | 30 | 1244 | 75 | DISC1FP1 | DISC1 fusion partner 1 | Other | other | |
| chr10 | *BTRC* | 27 | 716 | 75 | BTRC | beta-transducin repeat containing E3 ubiquitin protein ligase | Cytoplasm | enzyme | |
| chr1 | *CHD5* | 34 | 1921 | 75 | CHD5 | chromodomain helicase DNA binding protein 5 | Nucleus | enzyme | |
| chr3 | *LMCD1-AS1* | 35 | 887 | 75 | LMCD1-AS1 | LMCD1 antisense RNA 1 | Other | other | |
| chr11 | *CCDC73* | 28 | 963 | 75 | CCDC73 | coiled-coil domain containing 73 | Other | other | |
| chr9 | *PAX5* | 30 | 1091 | 75 | PAX5 | paired box 5 | Nucleus | transcription regulator | |
| chr17 | *ITGAE* | 32 | 1239 | 75 | ITGAE | integrin subunit alpha E | Plasma Membrane | other | |
| chr19 | *BRD4* | 32 | 1190 | 75 | BRD4 | bromodomain containing 4 | Nucleus | kinase | AZD5153, PLX51107, |
| chr16 | *LMF1* | 34 | 1610 | 75 | LMF1 | lipase maturation factor 1 | Cytoplasm | other | |
| chr5 | *EGFLAM* | 39 | 2564 | 75 | EGFLAM | EGF like, fibronectin type III and laminin G domains | Extracellular Space | other | |
| chr1 | *TNFRSF8* | 30 | 1186 | 75 | TNFRSF8 | TNF receptor superfamily member 8 | Plasma Membrane | transmembrane recepto | brentuximab vedotin/rituximab, monoclonal antibody SGN-30, HeFi-1 monoclonal antibody, MDX-1401, AFM13, iratumumab, anti-CD30 monoclonal antibody, |
| chr1 | *RAP1GAP* | 33 | 1563 | 75 | RAP1GAP | RAP1 GTPase activating protein | Cytoplasm | other | |
| chr1 | *TGFBR3* | 29 | 765 | 75 | TGFBR3 | transforming growth factor beta receptor 3 | Plasma Membrane | kinase | |
| chrX | *WNK3* | 30 | 862 | 75 | WNK3 | WNK lysine deficient protein | Plasma Membrane | kinase | |
| chr11 | *FAM168A* | 31 | 1031 | 75 | FAM168A | family with sequence similarity 168 member A | Other | other | |
| chr3 | *STXBP5L* | 33 | 856 | 75 | STXBP5L | syntaxin binding protein 5 like | Cytoplasm | other | |
| chr1 | *LPHN2* | 29 | 671 | 74 | ADGRL2 | adhesion G protein-coupled receptor L2 | Plasma Membrane | -protein coupled recepto | |
| chr11 | *PRDM11* | 32 | 1633 | 74 | PRDM11 | PR/SET domain 11 | Other | other | |
| chr1 | *EDARADD* | 33 | 952 | 74 | EDARADD | EDAR associated death domain | Cytoplasm | other | |
| chr9 | *KCNT1* | 38 | 2102 | 74 | KCNT1 | potassium sodium-activated channel subfamily T member 1 | Plasma Membrane | ion channel | |
| chr17 | *SLC43A2* | 32 | 1850 | 74 | SLC43A2 | solute carrier family 43 member 2 | Plasma Membrane | transporter | |
| chr15 | *SMAD6* | 28 | 1275 | 74 | SMAD6 | SMAD family member 6 | Nucleus | transcription regulator | |
| chr17 | *NLRP1* | 30 | 1012 | 74 | NLRP1 | NLR family pyrin domain | Cytoplasm | other | |

| | | | | | | | | | |
|---|---|---|---|---|---|---|---|---|---|
| chr15 | *ARNT2* | 33 | 1317 | 74 | ARNT2 | aryl hydrocarbon receptor nuclear translocator 2 | Nucleus | transcription regulator | |
| chr1 | *PDE4DIP* | 34 | 2801 | 74 | PDE4DIP | phosphodiesterase 4D interacting protein | Cytoplasm | enzyme | |
| chr16 | *RMI2* | 33 | 1120 | 74 | RMI2 | RecQ mediated genome | Nucleus | other | |
| chr12 | *ITPR2* | 30 | 907 | 74 | ITPR2 | inositol 1,4,5-trisphosphate receptor type 2 | Cytoplasm | ion channel | |
| chr3 | *RP11-384F7.1* | 32 | 773 | 74 | | | | | |
| chr7 | *DGKI* | 30 | 916 | 74 | DGKI | diacylglycerol kinase iota | Cytoplasm | kinase | |
| chr19 | *POU2F2* | 33 | 1159 | 74 | POU2F2 | POU class 2 homeobox 2 | Nucleus | transcription regulator | |
| chr9 | *PRUNE2* | 30 | 705 | 74 | PRUNE2 | prune homolog 2 with BCH | Cytoplasm | other | |
| chr8 | *ERICH1* | 33 | 1372 | 74 | ERICH1 | glutamate rich 1 | Other | other | |
| chr6 | *ARID1B* | 23 | 758 | 73 | ARID1B | AT-rich interaction domain 1B | Nucleus | transcription regulator | |
| chr17 | *CUEDC1* | 30 | 1208 | 73 | CUEDC1 | CUE domain containing 1 | Other | other | |
| chr1 | *CTNNBIP1* | 30 | 1156 | 73 | CTNNBIP1 | catenin beta interacting protein 1 | Cytoplasm | other | |
| chr2 | *INPP5D* | 27 | 1007 | 73 | INPP5D | inositol polyphosphate-5-phosphatase D | Cytoplasm | phosphatase | rosiptor |
| chr16 | *AXIN1* | 33 | 1576 | 73 | AXIN1 | axin 1 | Cytoplasm | other | |
| chr15 | *ZNF710* | 31 | 1866 | 73 | ZNF710 | zinc finger protein 710 | Nucleus | transcription regulator | |
| chrX | *POLA1* | 27 | 677 | 73 | POLA1 | DNA polymerase alpha 1, catalytic subunit | Nucleus | enzyme | entecavir, clofarabine/filgrastim/idarubicin, cladribine/cytarabine/filgrastim/idarubicin/plerixafor, cladribine, cytarabine/fludarabine phosphate/oxaliplatin/rituximab, cytarabine/fludarabine phosphate, cytarabine/IFNA2B, cladribine/cytarabine/filgrastim/idarubicin, cytarabine/etoposide/mitoxantrone, cytarabine/daunorubicin, cytarabine/dasatinib/filgrastim/fludarabine phosphate, cladribine/recombina[illegible] |
| chr6 | *UTRN* | 29 | 664 | 73 | UTRN | utrophin | Plasma Membrane | transmembrane receptor | |
| chr2 | *TMEM131* | 29 | 1074 | 73 | TMEM131 | transmembrane protein 131 | Extracellular Space | other | |
| chr10 | *GBF1* | 31 | 824 | 73 | GBF1 | golgi brefeldin A resistant guanine nucleotide exchange | Cytoplasm | other | |
| chr10 | *RP11-282I1.1* | 31 | 968 | 73 | | | | | |
| chr11 | *ZBTB16* | 33 | 870 | 73 | ZBTB16 | zinc finger and BTB domain containing 16 | Nucleus | transcription regulator | |
| chr20 | *DOK5* | 31 | 1219 | 73 | DOK5 | docking protein 5 | Plasma Membrane | other | |
| chr12 | *ERC1* | 25 | 641 | 73 | ERC1 | ELKS/RAB6-interacting/CAST family member 1 | Cytoplasm | other | |
| chrX | *PHF8* | 27 | 957 | 73 | PHF8 | PHD finger protein 8 | Nucleus | enzyme | |
| chr5 | *NUP155* | 32 | 2389 | 73 | NUP155 | nucleoporin 155 | Nucleus | other | |
| chr1 | *SLC9A1* | 31 | 1291 | 73 | SLC9A1 | solute carrier family 9 member A1 | Plasma Membrane | ion channel | amiloride, amiloride/hydrochlorothiazide |
| chr7 | *MKLN1-AS1* | 29 | 800 | 72 | LINC-PINT | long intergenic non-protein coding RNA, p53 induced transcript | Other | other | |
| chr12 | *TMTC2* | 30 | 707 | 72 | TMTC2 | transmembrane O-mannosyltransferase targeting cadherins 2 | Cytoplasm | enzyme | |
| chr5 | *CREBRF* | 29 | 1366 | 72 | CREBRF | CREB3 regulatory factor | Nucleus | transcription regulator | |
| chr8 | *PTK2B* | 32 | 1120 | 72 | PTK2B | protein tyrosine kinase 2 beta | Cytoplasm | kinase | conteltinib, lorlatinib, PF-562271 |
| chr8 | *SLC39A14* | 30 | 1101 | 72 | SLC39A14 | solute carrier family 39 member | Plasma Membrane | transporter | |
| chr3 | *RSRC1* | 30 | 888 | 72 | RSRC1 | arginine and serine rich coiled- | Nucleus | other | |
| chr9 | *NACC2* | 35 | 2105 | 72 | NACC2 | NACC family member 2 | Nucleus | transcription regulator | |
| chr22 | *OSBP2* | 30 | 1052 | 72 | OSBP2 | oxysterol binding protein 2 | Cytoplasm | other | |
| chr1 | *FBXO42* | 30 | 775 | 72 | FBXO42 | F-box protein 42 | Other | other | |
| chr17 | *CA10* | 34 | 661 | 72 | CA10 | carbonic anhydrase 10 | Other | enzyme | |
| chr1 | *ERI3* | 30 | 1051 | 72 | ERI3 | ERI1 exoribonuclease family member 3 | Plasma Membrane | other | |
| chr15 | *SNHG14* | 32 | 966 | 72 | SNHG14 | | Other | other | |
| chr9 | *SARDH* | 34 | 2048 | 72 | SARDH | sarcosine dehydrogenase | Cytoplasm | enzyme | |
| chr3 | *GSK3B* | 27 | 878 | 72 | GSK3B | glycogen synthase kinase 3 beta | Nucleus | kinase | glycogen synthase kinase-3beta inhibitor, enzastaurin |
| chr4 | *RNF4* | 26 | 830 | 72 | RNF4 | ring finger protein 4 | Nucleus | transcription regulator | |
| chr17 | *MYO18A* | 30 | 1462 | 72 | MYO18A | myosin XVIIIA | Cytoplasm | other | |
| chr11 | *OSBPL5* | 37 | 2875 | 72 | OSBPL5 | oxysterol binding protein like 5 | Cytoplasm | transporter | |
| chrX | *CDKL5* | 28 | 634 | 72 | CDKL5 | cyclin dependent kinase like 5 | Nucleus | kinase | |
| chr16 | *RP11-410D17.2* | 29 | 803 | 72 | | | | | |
| chr5 | *SEC24A* | 27 | 724 | 72 | SEC24A | SEC24 homolog A, COPII coat complex component | Cytoplasm | transporter | |
| chr17 | *RP11-101O21.1* | 32 | 963 | 72 | | | | | |
| chr16 | *CFDP1* | 28 | 763 | 71 | CFDP1 | craniofacial development protein | Extracellular Space | other | |
| chr11 | *DCDC1* | 31 | 843 | 71 | DCDC1 | doublecortin domain containing 1 | Other | other | |

| | | | | | | | | | |
|---|---|---|---|---|---|---|---|---|---|
| chr6 | *PACSIN1* | 30 | 1158 | 71 | PACSIN1 | protein kinase C and casein kinase substrate in neurons 1 | Cytoplasm | kinase | |
| chr11 | *TRPC6* | 33 | 1615 | 71 | TRPC6 | transient receptor potential cation channel subfamily C member 6 | Plasma Membrane | ion channel | |
| chr7 | *GRM8* | 32 | 855 | 71 | GRM8 | glutamate metabotropic receptor | Plasma Membrane | G-protein coupled recepto | fasoracetam |
| chr1 | *SPEN* | 28 | 937 | 71 | SPEN | spen family transcriptional | Nucleus | transcription regulator | |
| chr16 | *KIAA0513* | 32 | 1342 | 71 | KIAA0513 | KIAA0513 | Other | other | |
| chr7 | *EEPD1* | 32 | 1171 | 71 | EEPD1 | endonuclease/exonuclease/phosphatase family domain containing | Other | other | |
| chr8 | *ST3GAL1* | 32 | 1184 | 71 | ST3GAL1 | ST3 beta-galactoside alpha-2,3-sialyltransferase 1 | Cytoplasm | enzyme | |
| chr6 | *RREB1* | 31 | 967 | 71 | RREB1 | ras responsive element binding protein 1 | Nucleus | transcription regulator | |
| chr17 | *ABCC3* | 29 | 1252 | 71 | ABCC3 | ATP binding cassette subfamily C member 3 | Plasma Membrane | transporter | |
| chr13 | *PCDH9* | 27 | 705 | 71 | PCDH9 | protocadherin 9 | Plasma Membrane | other | |
| chr7 | *BAIAP2L1* | 29 | 1178 | 71 | BAIAP2L1 | BAR/IMD domain containing adaptor protein 2 like 1 | Cytoplasm | other | |
| chr18 | *MYOM1* | 29 | 1229 | 71 | MYOM1 | myomesin 1 | Cytoplasm | other | |
| chr5 | *ANKRD33B* | 34 | 2824 | 71 | ANKRD33B | ankyrin repeat domain 33B | Other | other | |
| chr5 | *BASP1* | 32 | 2035 | 71 | BASP1 | brain abundant membrane attached signal protein 1 | Nucleus | transcription regulator | |
| chr2 | *MYO7B* | 32 | 1749 | 71 | MYO7B | myosin VIIB | Cytoplasm | peptidase | |
| chr3 | *SLC6A11* | 30 | 966 | 71 | SLC6A11 | solute carrier family 6 member 11 | Plasma Membrane | transporter | tiagabine |
| chr12 | *GLT1D1* | 29 | 838 | 71 | GLT1D1 | glycosyltransferase 1 domain containing 1 | Extracellular Space | enzyme | |
| chr15 | *KLF13* | 28 | 1247 | 71 | KLF13 | Kruppel like factor 13 | Nucleus | transcription regulator | |
| chr10 | *MYOF* | 30 | 811 | 71 | MYOF | myoferlin | Nucleus | other | |
| chr6 | *GCNT2* | 28 | 746 | 71 | GCNT2 | glucosaminyl (N-acetyl) transferase 2 (I blood group) | Cytoplasm | enzyme | |
| chr1 | *EIF2B3* | 29 | 686 | 71 | EIF2B3 | eukaryotic translation initiation factor 2B subunit gamma | Cytoplasm | other | |
| chr2 | *EDAR* | 30 | 1603 | 71 | EDAR | ectodysplasin A receptor | Plasma Membrane | transmembrane receptor | |
| chr15 | *STARD9* | 27 | 691 | 70 | STARD9 | StAR related lipid transfer domain containing 9 | Cytoplasm | other | |
| chr17 | *SMURF2* | 27 | 691 | 70 | SMURF2 | SMAD specific E3 ubiquitin protein ligase 2 | Cytoplasm | enzyme | |
| chr16 | *GRIN2A* | 33 | 692 | 70 | GRIN2A | glutamate ionotropic receptor NMDA type subunit 2A | Plasma Membrane | ion channel | nebostinel, besonprodil, delucemine, 1-aminocyclopropane-1-carboxylic acid, orphenadrine, dextromethorphan/promethazine, neramexane, felbamate, chlorpheniramine/dextromethorphan/phenylephrine, memantine, donepezil/memantine, N-(2-indanyl)glycinamide, tenocyclidine, dextromethorphan, cycloserine, UK-240455, brompheniramine/dextromethorphan/pseudoephedrine, [illegible] |
| chr22 | *PLA2G6* | 27 | 1024 | 70 | PLA2G6 | phospholipase A2 group VI | Cytoplasm | enzyme | quinacrine |
| chr9 | *LMX1B* | 34 | 1841 | 70 | LMX1B | LIM homeobox transcription factor 1 beta | Nucleus | transcription regulator | |
| chr1 | *ENAH* | 28 | 838 | 70 | ENAH | ENAH actin regulator | Plasma Membrane | other | |
| chr11 | *NCAM1* | 31 | 735 | 70 | NCAM1 | neural cell adhesion molecule 1 | Plasma Membrane | other | BB-10901 |
| chr10 | *PLCE1* | 33 | 627 | 70 | PLCE1 | phospholipase C epsilon 1 | Cytoplasm | enzyme | |
| chr5 | *FBXL17* | 27 | 628 | 70 | FBXL17 | F-box and leucine rich repeat protein 17 | Nucleus | other | |
| chr3 | *LINC00882* | 30 | 922 | 70 | LINC00882 | long intergenic non-protein coding RNA 882 | Other | other | |
| chr4 | *TBC1D14* | 27 | 990 | 70 | TBC1D14 | TBC1 domain family member 14 | Extracellular Space | other | |
| chr10 | *HERC4* | 29 | 1005 | 70 | HERC4 | HECT and RLD domain containing E3 ubiquitin protein | Cytoplasm | enzyme | |
| chr12 | *ANAPC5* | 28 | 771 | 70 | ANAPC5 | anaphase promoting complex subunit 5 | Nucleus | other | |
| chr22 | *TNRC6B* | 24 | 656 | 70 | TNRC6B | trinucleotide repeat containing adaptor 6B | Cytoplasm | other | |
| chr9 | *GARNL3* | 31 | 1002 | 70 | GARNL3 | GTPase activating Rap/RanGAP domain like 3 | Other | other | |
| chr16 | *CDYL2* | 29 | 918 | 70 | CDYL2 | chromodomain Y like 2 | Nucleus | other | |
| chr2 | *MSH2* | 30 | 990 | 70 | MSH2 | mutS homolog 2 | Nucleus | enzyme | |
| chr7 | *LHFPL3* | 32 | 759 | 70 | LHFPL3 | LHFPL tetraspan subfamily member 3 | Other | other | |
| chr7 | *VIPR2* | 31 | 1502 | 70 | VIPR2 | vasoactive intestinal peptide receptor 2 | Plasma Membrane | G-protein coupled recepto | |
| chr1 | *ZMYM4* | 25 | 609 | 70 | ZMYM4 | zinc finger MYM-type containing | Nucleus | other | |
| chr16 | *KIFC3* | 31 | 1242 | 70 | KIFC3 | kinesin family member C3 | Cytoplasm | enzyme | |
| chr9 | *TTLL11* | 30 | 984 | 70 | TTLL11 | tubulin tyrosine ligase like 11 | Cytoplasm | enzyme | |

| | | | | | | | | | |
|---|---|---|---|---|---|---|---|---|---|
| chr8 | *DPYSL2* | 30 | 896 | 70 | DPYSL2 | dihydropyrimidinase like 2 | Cytoplasm | enzyme | |
| chr20 | *NDRG3* | 29 | 1045 | 70 | NDRG3 | NDRG family member 3 | Cytoplasm | other | |
| chr19 | *MUM1* | 33 | 1441 | 70 | | | | | |
| chr15 | *RP11-66B24.5* | 31 | 1805 | 70 | | | | | |
| chr16 | *LA16c-444G7.1* | 30 | 1555 | 70 | | | | | |
| chr10 | *SORCS3* | 34 | 689 | 70 | SORCS3 | sortilin related VPS10 domain containing receptor 3 | Nucleus | transporter | |
| chr7 | *COG5* | 26 | 732 | 70 | COG5 | component of oligomeric golgi complex 5 | Cytoplasm | transporter | |
| chr1 | *HMCN1* | 29 | 832 | 70 | HMCN1 | hemicentin 1 | Extracellular Space | other | |
| chr6 | *CAP2* | 28 | 699 | 70 | CAP2 | cyclase associated actin cytoskeleton regulatory protein 2 | Plasma Membrane | other | |
| chr7 | *NRF1* | 26 | 611 | 70 | NRF1 | nuclear respiratory factor 1 | Nucleus | transcription regulator | |
| chr11 | *DAGLA* | 33 | 2086 | 70 | DAGLA | diacylglycerol lipase alpha | Plasma Membrane | enzyme | |
| chr12 | *RP11-611E13.2* | 27 | 748 | 69 | | | | | |
| chr7 | *MYH16* | 31 | 1279 | 69 | MYH16 | myosin heavy chain 16 | Other | other | |
| chr15 | *MYO9A* | 28 | 719 | 69 | MYO9A | myosin IXA | Cytoplasm | enzyme | |
| chr18 | *ATP9B* | 25 | 969 | 69 | ATP9B | ATPase phospholipid transporting 9B (putative) | Cytoplasm | transporter | |
| chr10 | *C10orf90* | 31 | 861 | 69 | C10orf90 | chromosome 10 open reading frame 90 | Cytoplasm | enzyme | |
| chr3 | *DAG1* | 27 | 1214 | 69 | DAG1 | dystroglycan 1 | Plasma Membrane | transmembrane receptor | |
| chr1 | *WASF2* | 28 | 808 | 69 | WASF2 | WASP family member 2 | Plasma Membrane | other | |
| chr14 | *TECPR2* | 27 | 929 | 69 | TECPR2 | tectonin beta-propeller repeat containing 2 | Other | other | |
| chr2 | *PTPN4* | 25 | 876 | 69 | PTPN4 | protein tyrosine phosphatase non-receptor type 4 | Cytoplasm | phosphatase | |
| chr5 | *ADAMTS12* | 35 | 2260 | 69 | ADAMTS12 | ADAM metallopeptidase with thrombospondin type 1 motif 12 | Extracellular Space | peptidase | |
| chr22 | *CACNA1I* | 28 | 1374 | 69 | CACNA1I | calcium voltage-gated channel subunit alpha1 I | Plasma Membrane | ion channel | flunarizine, mibefradil, |
| chr16 | *ABAT* | 30 | 803 | 69 | ABAT | 4-aminobutyrate aminotransferase | Cytoplasm | enzyme | vigabatrin, theophylline/tretinoin/ valproic acid, |
| chr5 | *PPP2R2B* | 29 | 652 | 69 | PPP2R2B | protein phosphatase 2 regulatory subunit Bbeta | Cytoplasm | phosphatase | |
| chr12 | *CHFR* | 30 | 964 | 69 | CHFR | checkpoint with forkhead and ring finger domains | Nucleus | enzyme | |
| chr6 | *RP5-1120P11.1* | 28 | 1165 | 69 | | | | | |
| chr1 | *AHDC1* | 30 | 1354 | 69 | AHDC1 | AT-hook DNA binding motif containing 1 | Other | other | |
| chr14 | *TMEM179* | 30 | 1427 | 69 | TMEM179 | transmembrane protein 179 | Other | other | |
| chr15 | *BLM* | 30 | 1286 | 69 | BLM | BLM RecQ like helicase | Nucleus | enzyme | |
| chr20 | *PPP1R16B* | 32 | 1333 | 69 | PPP1R16B | protein phosphatase 1 regulatory subunit 16B | Plasma Membrane | phosphatase | |
| chr15 | *GABRB3* | 27 | 739 | 69 | GABRB3 | gamma-aminobutyric acid type A receptor beta3 subunit | Plasma Membrane | ion channel | [illegible] nitrazepam, methohexital, sevoflurane, etomidate, pagoclone, aspirin/butalbital/caffeine, zolpidem, atropine/hyoscyamine/phenobarbital/scopolamine, phenobarbital, diazepam, isoflurane, halothane, pentobarbital, acetaminophen/butalbital/caffeine/codeine, hyoscyamine/phenobarbital, desflurane, amobarbital, adinazolam, SEP 174559, [illegible] |
| chr9 | *WNK2* | 27 | 1257 | 69 | WNK2 | WNK lysine deficient protein | Cytoplasm | kinase | |
| chr15 | *PEAK1* | 27 | 776 | 69 | PEAK1 | pseudopodium enriched atypical kinase 1 | Plasma Membrane | kinase | |
| chr12 | *RP11-114G22.1* | 29 | 826 | 69 | | | | | |
| chr16 | *C16orf74* | 29 | 1234 | 69 | C16orf74 | chromosome 16 open reading frame 74 | Other | other | |
| chr8 | *HMBOX1* | 24 | 700 | 68 | HMBOX1 | homeobox containing 1 | Nucleus | transcription regulator | |
| chr1 | *KCNQ4* | 29 | 1417 | 68 | KCNQ4 | potassium voltage-gated channel subfamily Q member 4 | Plasma Membrane | ion channel | ezogabine |
| chr2 | *LRRFIP1* | 23 | 863 | 68 | LRRFIP1 | LRR binding FLII interacting | Cytoplasm | transcription regulator | |
| chr13 | *FNDC3A* | 25 | 743 | 68 | FNDC3A | fibronectin type III domain containing 3A | Cytoplasm | other | |
| chr7 | *GLI3* | 30 | 855 | 68 | GLI3 | GLI family zinc finger 3 | Nucleus | transcription regulator | |
| chr1 | *LINC01057* | 31 | 1015 | 68 | SLC44A3-AS1 | SLC44A3 antisense RNA 1 | Other | other | |
| chr12 | *MGAT4C* | 26 | 685 | 68 | MGAT4C | MGAT4 family member C | Cytoplasm | enzyme | |
| chr9 | *PCSK5* | 30 | 614 | 68 | PCSK5 | proprotein convertase subtilisin/kexin type 5 | Extracellular Space | peptidase | |
| chr12 | *MPHOSPH9* | 27 | 696 | 68 | MPHOSPH9 | M-phase phosphoprotein 9 | Cytoplasm | other | |
| chr19 | *NWD1* | 31 | 858 | 68 | NWD1 | NACHT and WD repeat domain containing 1 | Cytoplasm | other | |

| | | | | | | | | | |
|---|---|---|---|---|---|---|---|---|---|
| chr16 | *SCNN1B* | 29 | 939 | 68 | SCNN1B | sodium channel epithelial 1 beta subunit | Plasma Membrane | ion channel | hydrochlorothiazide/triamterene, amiloride, triamterene, amiloride/hydrochlorothiazide |
| chr1 | *ACBD6* | 25 | 812 | 68 | ACBD6 | acyl-CoA binding domain containing 6 | Cytoplasm | other | |
| chr11 | *ST5* | 32 | 1134 | 68 | ST5 | suppression of tumorigenicity 5 | Cytoplasm | enzyme | |
| chr5 | *RANBP17* | 28 | 1017 | 68 | RANBP17 | RAN binding protein 17 | Nucleus | transporter | |
| chr7 | *NUDCD3* | 31 | 1145 | 68 | NUDCD3 | NudC domain containing 3 | Cytoplasm | other | |
| chr11 | *RCN1* | 32 | 981 | 68 | RCN1 | reticulocalbin 1 | Cytoplasm | other | |
| chr19 | *NFIC* | 30 | 1233 | 68 | NFIC | nuclear factor I C | Nucleus | transcription regulator | |
| chr5 | *UNC5A* | 30 | 1624 | 68 | UNC5A | unc-5 netrin receptor A | Plasma Membrane | transmembrane recepto[illegible] | |
| chr6 | *SUPT3H* | 28 | 748 | 67 | SUPT3H | SPT3 homolog, SAGA and STAGA complex component | Nucleus | transcription regulator | |
| chr2 | *EML4* | 24 | 817 | 67 | EML4 | EMAP like 4 | Cytoplasm | other | |
| chr17 | *CCDC40* | 28 | 1150 | 67 | CCDC40 | coiled-coil domain containing 40 | Cytoplasm | other | |
| chr11 | *PPP6R3* | 26 | 746 | 67 | PPP6R3 | protein phosphatase 6 regulatory subunit 3 | Cytoplasm | other | |
| chr1 | *PIK3R3* | 27 | 749 | 67 | PIK3R3 | phosphoinositide-3-kinase regulatory subunit 3 | Cytoplasm | kinase | |
| chr9 | *NCS1* | 30 | 1691 | 67 | NCS1 | neuronal calcium sensor 1 | Plasma Membrane | other | |
| chr9 | *SPTAN1* | 29 | 1267 | 67 | SPTAN1 | spectrin alpha, non-erythrocytic 1 | Plasma Membrane | other | |
| chrX | *DIAPH2* | 23 | 615 | 67 | DIAPH2 | diaphanous related formin 2 | Cytoplasm | other | |
| chr15 | *MYO5A* | 26 | 699 | 67 | MYO5A | myosin VA | Cytoplasm | enzyme | |
| chr22 | *GRAMD4* | 30 | 1517 | 67 | GRAMD4 | GRAM domain containing 4 | Cytoplasm | other | |
| chr22 | *PI4KA* | 25 | 986 | 67 | PI4KA | phosphatidylinositol 4-kinase | Cytoplasm | kinase | |
| chr3 | *UBE2E2* | 29 | 847 | 67 | UBE2E2 | ubiquitin conjugating enzyme E2 | Cytoplasm | enzyme | |
| chr3 | *SETD2* | 26 | 638 | 67 | SETD2 | SET domain containing 2, histone lysine methyltransferase | Cytoplasm | enzyme | |
| chr16 | *VAT1L* | 30 | 656 | 67 | VAT1L | vesicle amine transport 1 like | Other | enzyme | |
| chr5 | *RAI14* | 33 | 2057 | 67 | RAI14 | retinoic acid induced 14 | Nucleus | transcription regulator | |
| chr2 | *ADD2* | 30 | 865 | 67 | ADD2 | adducin 2 | Cytoplasm | other | |
| chr5 | *MCC* | 31 | 749 | 67 | MCC | MCC regulator of WNT signaling pathway | Cytoplasm | other | |
| chr16 | *C16orf45* | 28 | 767 | 67 | BMERB1 | bMERB domain containing 1 | Other | other | |
| chr3 | *GRM7* | 31 | 649 | 67 | GRM7 | glutamate metabotropic receptor | Plasma Membrane | [illegible]-protein coupled recept[illegible] | fasoracetam |
| chr11 | *SOX6* | 29 | 702 | 67 | SOX6 | SRY-box transcription factor 6 | Nucleus | transcription regulator | |
| chr2 | *DIS3L2* | 28 | 916 | 67 | DIS3L2 | DIS3 like 3'-5' exoribonuclease 2 | Cytoplasm | enzyme | |
| chr8 | *FAM49B* | 27 | 830 | 67 | FAM49B | family with sequence similarity 49 member B | Extracellular Space | other | |
| chr1 | *NVL* | 28 | 904 | 67 | NVL | nuclear VCP like | Nucleus | other | |
| chr17 | *PRPSAP2* | 28 | 715 | 67 | PRPSAP2 | phosphoribosyl pyrophosphate synthetase associated protein 2 | Other | other | |
| chr3 | *SEMA5B* | 30 | 1332 | 67 | SEMA5B | semaphorin 5B | Plasma Membrane | other | |
| chr1 | *PBX1* | 29 | 902 | 67 | PBX1 | PBX homeobox 1 | Nucleus | transcription regulator | |
| chr22 | *FBLN1* | 27 | 1132 | 67 | FBLN1 | fibulin 1 | Extracellular Space | other | |
| chr1 | *PTPN14* | 28 | 926 | 67 | PTPN14 | protein tyrosine phosphatase non-receptor type 14 | Cytoplasm | phosphatase | |
| chr1 | *ITPKB* | 29 | 1419 | 67 | ITPKB | inositol-trisphosphate 3-kinase B | Cytoplasm | kinase | |
| chr15 | *TJP1* | 23 | 688 | 67 | TJP1 | tight junction protein 1 | Plasma Membrane | other | |
| chr5 | *CTC-535M15.2* | 27 | 737 | 66 | | | | | |
| chr12 | *CAMKK2* | 28 | 990 | 66 | CAMKK2 | calcium/calmodulin dependent protein kinase kinase 2 | Cytoplasm | kinase | |
| chr7 | *DNAH11* | 28 | 756 | 66 | DNAH11 | dynein axonemal heavy chain 11 | Cytoplasm | enzyme | |
| chr2 | *MAP4K3* | 24 | 795 | 66 | MAP4K3 | mitogen-activated protein kinase kinase kinase kinase 3 | Other | kinase | |
| chr1 | *MTOR* | 28 | 747 | 66 | MTOR | mechanistic target of rapamycin kinase | Nucleus | kinase | everolimus/fulvestrant, apitolisib, PF-4691502, AZD8055, CC-223, PKI-179, dactolisib, samotolisib, everolimus/letrozole, everolimus/gefitinib, everolimus/prednisone, ABI-009, methotrexate/sirolimus/tacrolimus, prednisone/tacrolimus, methylprednisolone/tacrolimus, cyclosporine A/sirolimus/tacrolimus, PWT33597, mTOR inhibitor, everolimus/sorafenib, everolimus, bimiralisib, [illegible] |
| chr2 | *AC019118.2* | 25 | 851 | 66 | | | | | |
| chr20 | *APCDD1L-AS1* | 30 | 1792 | 66 | APCDD1L-DT | APCDD1L divergent transcript | Other | other | |
| chr11 | *PRDM10* | 28 | 996 | 66 | PRDM10 | PR/SET domain 10 | Nucleus | transcription regulator | |
| chr1 | *RFWD2* | 24 | 785 | 66 | COP1 | COP1 E3 ubiquitin ligase | Cytoplasm | enzyme | |

| | | | | | | | | | |
|---|---|---|---|---|---|---|---|---|---|
| chr17 | *CACNA1G* | 29 | 1467 | 66 | CACNA1G | calcium voltage-gated channel subunit alpha1 G | Plasma Membrane | ion channel | flunarizine, ethosuximide, mibefradil, zonisamide, cinnarizine, |
| chr2 | *TFCP2L1* | 27 | 1396 | 66 | TFCP2L1 | transcription factor CP2 like 1 | Nucleus | transcription regulator | |
| chr16 | *C16orf62* | 31 | 791 | 66 | VPS35L | VPS35 endosomal protein sorting factor like | Cytoplasm | other | |
| chr2 | *ASAP2* | 24 | 878 | 66 | ASAP2 | ArfGAP with SH3 domain, ankyrin repeat and PH domain 2 | Nucleus | other | |
| chr7 | *DOCK4* | 26 | 600 | 66 | DOCK4 | dedicator of cytokinesis 4 | Plasma Membrane | other | |
| chr6 | *RP11-146I2.1* | 28 | 665 | 66 | | | | | |
| chr10 | *CHST15* | 28 | 1116 | 66 | CHST15 | carbohydrate sulfotransferase 15 | Plasma Membrane | enzyme | |
| chr5 | *DOCK2* | 30 | 953 | 66 | DOCK2 | dedicator of cytokinesis 2 | Cytoplasm | other | |
| chr18 | *GNAL* | 24 | 769 | 66 | GNAL | G protein subunit alpha L | Cytoplasm | enzyme | |
| chr5 | *RP11-454P21.1* | 29 | 1623 | 66 | | | | | |
| chr17 | *UBE2G1* | 25 | 702 | 66 | UBE2G1 | ubiquitin conjugating enzyme E2 | Cytoplasm | enzyme | |
| chr16 | *ABCA3* | 30 | 1245 | 66 | ABCA3 | ATP binding cassette subfamily A member 3 | Plasma Membrane | transporter | |
| chr1 | *ATP2B4* | 30 | 971 | 66 | ATP2B4 | ATPase plasma membrane Ca2+ transporting 4 | Plasma Membrane | transporter | |
| chr3 | *EGFEM1P* | 29 | 1102 | 66 | EGFEM1P | EGF like and EMI domain containing 1, pseudogene | Other | other | |
| chrX | *USP9X* | 23 | 624 | 66 | USP9X | ubiquitin specific peptidase 9 X- | Plasma Membrane | peptidase | |
| chr12 | *R3HDM2* | 26 | 679 | 66 | R3HDM2 | R3H domain containing 2 | Nucleus | other | |
| chr7 | *RADIL* | 30 | 1453 | 66 | RADIL | Rap associating with DIL domain | Cytoplasm | other | |
| chr22 | *TCF20* | 27 | 1172 | 66 | TCF20 | transcription factor 20 | Nucleus | transcription regulator | |
| chr2 | *KCNIP3* | 30 | 1691 | 66 | KCNIP3 | potassium voltage-gated channel interacting protein 3 | Nucleus | transcription regulator | |
| chr11 | *MYEOV* | 27 | 1070 | 65 | MYEOV | myeloma overexpressed | Nucleus | other | |
| chr1 | *HHAT* | 30 | 837 | 65 | HHAT | hedgehog acyltransferase | Cytoplasm | enzyme | |
| chr5 | *CPLX2* | 28 | 953 | 65 | CPLX2 | complexin 2 | Cytoplasm | other | |
| chr8 | *TUSC3* | 25 | 705 | 65 | TUSC3 | tumor suppressor candidate 3 | Plasma Membrane | transporter | |
| chr1 | *KIAA1324* | 29 | 952 | 65 | KIAA1324 | KIAA1324 | Plasma Membrane | other | |
| chr10 | *PARD3* | 24 | 623 | 65 | PARD3 | par-3 family cell polarity regulator | Plasma Membrane | other | |
| chr17 | *RPA1* | 27 | 925 | 65 | RPA1 | replication protein A1 | Nucleus | other | |
| chr10 | *TLL2* | 29 | 883 | 65 | TLL2 | tolloid like 2 | Extracellular Space | peptidase | |
| chr9 | *LRSAM1* | 27 | 1572 | 65 | LRSAM1 | leucine rich repeat and sterile alpha motif containing 1 | Cytoplasm | enzyme | |
| chr10 | *ARID5B* | 29 | 1131 | 65 | ARID5B | AT-rich interaction domain 5B | Nucleus | transcription regulator | |
| chr4 | *HTT* | 24 | 903 | 65 | HTT | huntingtin | Cytoplasm | transcription regulator | |
| chr2 | *NDUFA10* | 24 | 1036 | 65 | NDUFA10 | NADH:ubiquinone oxidoreductase subunit A10 | Cytoplasm | transporter | |
| chr3 | *PLCH1* | 28 | 784 | 65 | PLCH1 | phospholipase C eta 1 | Cytoplasm | enzyme | |
| chr1 | *CACNA1S* | 28 | 1478 | 65 | CACNA1S | calcium voltage-gated channel subunit alpha1 S | Plasma Membrane | ion channel | amlodipine/valsartan, amlodipine/ezetimibe/losartan/rosuvastatin, nicardipine, amlodipine/atorvastatin, aliskiren/amlodipine/hydrochlorothiazide, amlodipine/telmisartan, aliskiren/amlodipine, isradipine, MEM-1003, cinnarizine, nifedipine, amlodipine/indapamide/perindopril, amlodipine/perindopril, felodipine, verapamil, amlodipine/hydrochlorothiazide/valsartan, dexamethasone/rituximab/verapamil, [illegible] |
| chr2 | *TPO* | 25 | 998 | 65 | TPO | thyroid peroxidase | Plasma Membrane | enzyme | propylthiouracil, methimazole, carbimazole |
| chrX | *KDM6A* | 24 | 725 | 65 | KDM6A | lysine demethylase 6A | Nucleus | enzyme | |
| chr16 | *GTF3C1* | 28 | 966 | 65 | GTF3C1 | general transcription factor IIIC subunit 1 | Nucleus | transcription regulator | |
| chr10 | *RP11-170M17.1* | 29 | 1110 | 65 | | | | | |
| chr11 | *ZNF143* | 24 | 874 | 65 | ZNF143 | zinc finger protein 143 | Nucleus | transcription regulator | |
| chr15 | *TRPM1* | 26 | 898 | 65 | TRPM1 | transient receptor potential cation channel subfamily M member 1 | Plasma Membrane | ion channel | |
| chr10 | *TBC1D12* | 26 | 680 | 65 | TBC1D12 | TBC1 domain family member 12 | Extracellular Space | other | |
| chr13 | *RNF219-AS1* | 26 | 682 | 65 | OBI1-AS1 | OBI1 antisense RNA 1 | Other | other | |

| | | | | | | | | | |
|---|---|---|---|---|---|---|---|---|---|
| chr7 | *BRAF* | 24 | 627 | 65 | BRAF | B-Raf proto-oncogene, serine/threonine kinase | Cytoplasm | kinase | irinotecan/panitumumab/vemurafenib, sorafenib, dabrafenib/trametinib, sorafenib/sulindac/sunitinib, ARQ 736, vemurafenib, ipilimumab/vemurafenib, binimetinib/encorafenib/panitumumab, bevacizumab/sorafenib, dexamethasone/lenalidomide/sorafenib, regorafenib, BRAF inhibitor, 5-azacytidine/sorafenib, dabrafenib/ipilimumab/trametinib, belvarafenib, CEP-[illegible] |
| chr6 | *PPARD* | 27 | 931 | 65 | PPARD | peroxisome proliferator activated receptor delta | Nucleus | d-dependent nuclear rec | bezafibrate, icosapent, |
| chr19 | *GRIK5* | 29 | 1036 | 65 | GRIK5 | glutamate ionotropic receptor kainate type subunit 5 | Plasma Membrane | ion channel | |
| chr10 | *LRRC20* | 30 | 1531 | 65 | LRRC20 | leucine rich repeat containing 20 | Other | other | |
| chr10 | *ADD3* | 24 | 757 | 65 | ADD3 | adducin 3 | Cytoplasm | other | |
| chr14 | *RIN3* | 24 | 977 | 65 | RIN3 | Ras and Rab interactor 3 | Cytoplasm | other | |
| chr3 | *ZPLD1* | 27 | 768 | 64 | ZPLD1 | zona pellucida like domain containing 1 | Other | other | |
| chr2 | *LIMS1* | 25 | 945 | 64 | LIMS1 | LIM zinc finger domain containing | Plasma Membrane | other | |
| chr21 | *ADARB1* | 23 | 909 | 64 | ADARB1 | adenosine deaminase RNA specific B1 | Nucleus | enzyme | |
| chr5 | *FAM134B* | 29 | 2277 | 64 | RETREG1 | reticulophagy regulator 1 | Cytoplasm | other | |
| chr3 | *FHIT* | 28 | 629 | 64 | FHIT | fragile histidine triad diadenosine triphosphatase | Cytoplasm | enzyme | |
| chr6 | *EYS* | 23 | 626 | 64 | EYS | eyes shut homolog | Cytoplasm | other | |
| chr16 | *CBFA2T3* | 32 | 1541 | 64 | CBFA2T3 | CBFA2/RUNX1 partner transcriptional co-repressor 3 | Nucleus | transcription regulator | |
| chr6 | *ITPR3* | 29 | 1457 | 64 | ITPR3 | inositol 1,4,5-trisphosphate receptor type 3 | Cytoplasm | ion channel | |
| chr17 | *GPATCH8* | 24 | 767 | 64 | GPATCH8 | G-patch domain containing 8 | Other | other | |
| chr12 | *HCAR1* | 26 | 750 | 64 | HCAR1 | hydroxycarboxylic acid receptor 1 | Plasma Membrane | -protein coupled recepto | |
| chr10 | *BMPR1A* | 26 | 625 | 64 | BMPR1A | bone morphogenetic protein receptor type 1A | Plasma Membrane | kinase | |
| chr2 | *HS1BP3* | 24 | 1039 | 64 | HS1BP3 | HCLS1 binding protein 3 | Other | other | |
| chr11 | *SORL1* | 28 | 707 | 64 | SORL1 | sortilin related receptor 1 | Cytoplasm | transporter | |
| chr15 | *CTD-2544M6.1* | 31 | 1126 | 64 | | | | | |
| chr12 | *WDR66* | 28 | 811 | 64 | WDR66 | WD repeat domain 66 | Cytoplasm | other | |
| chr2 | *AC007246.3* | 27 | 848 | 64 | | | | | |
| chr8 | *TNKS* | 23 | 566 | 64 | TNKS | tankyrase | Nucleus | enzyme | |
| chr12 | *SCN8A* | 30 | 1075 | 64 | SCN8A | sodium voltage-gated channel alpha subunit 8 | Plasma Membrane | ion channel | riluzole |
| chr5 | *RP11-774D14.1* | 27 | 1391 | 64 | | | | | |
| chr16 | *CTCF* | 25 | 748 | 64 | CTCF | CCCTC-binding factor | Nucleus | transcription regulator | |
| chr20 | *OSBPL2* | 29 | 2151 | 64 | OSBPL2 | oxysterol binding protein like 2 | Cytoplasm | other | |
| chr15 | *SCAMP5* | 27 | 814 | 64 | SCAMP5 | secretory carrier membrane | Cytoplasm | other | |
| chr5 | *ANKH* | 31 | 2252 | 64 | ANKH | ANKH inorganic pyrophosphate transport regulator | Plasma Membrane | transporter | |
| chr15 | *HERC2P2* | 24 | 988 | 64 | HERC2P2 | hect domain and RLD 2 pseudogene 2 | Other | other | |
| chr16 | *ACSF3* | 30 | 1461 | 64 | ACSF3 | acyl-CoA synthetase family member 3 | Cytoplasm | enzyme | |
| chr3 | *THRB* | 27 | 815 | 64 | THRB | thyroid hormone receptor beta | Nucleus | d-dependent nuclear rec | dextrothyroxine, amiodarone, levothyroxine/l-triiodothyronine, denosumab/levothyroxine, L-triiodothyronine, 3,5-diiodothyropropionic |
| chr14 | *DPF3* | 25 | 766 | 64 | DPF3 | double PHD fingers 3 | Nucleus | other | |
| chr2 | *C2orf42* | 27 | 751 | 64 | C2orf42 | chromosome 2 open reading frame 42 | Nucleus | other | |
| chr15 | *SV2B* | 31 | 982 | 64 | SV2B | synaptic vesicle glycoprotein 2B | Plasma Membrane | transporter | |
| chr3 | *RBMS3* | 25 | 582 | 64 | RBMS3 | RNA binding motif single stranded interacting protein 3 | Cytoplasm | other | |
| chr5 | *DNAH5* | 30 | 1769 | 64 | DNAH5 | dynein axonemal heavy chain 5 | Cytoplasm | enzyme | |
| chr2 | *ANAPC1* | 23 | 758 | 63 | ANAPC1 | anaphase promoting complex subunit 1 | Nucleus | other | |
| chr16 | *FTO* | 28 | 588 | 63 | FTO | FTO alpha-ketoglutarate dependent dioxygenase | Nucleus | enzyme | |
| chr19 | *VAV1* | 28 | 761 | 63 | VAV1 | vav guanine nucleotide exchange factor 1 | Nucleus | transcription regulator | |
| chr5 | *CTD-2066L21.3* | 29 | 2044 | 63 | | | | | |
| chr11 | *MADD* | 28 | 1737 | 63 | MADD | MAP kinase activating death | Cytoplasm | other | |

| | | | | | | | | | |
|---|---|---|---|---|---|---|---|---|---|
| chr19 | *SMARCA4* | 27 | 1061 | 63 | SMARCA4 | SWI/SNF related, matrix associated, actin dependent regulator of chromatin, subfamily | Nucleus | transcription regulator | |
| chr20 | *BMP7* | 30 | 1628 | 63 | BMP7 | bone morphogenetic protein 7 | Extracellular Space | growth factor | |
| chr4 | *AFAP1* | 23 | 812 | 63 | AFAP1 | actin filament associated protein | Cytoplasm | other | |
| chr11 | *CLMP* | 27 | 702 | 63 | CLMP | CXADR like membrane protein | Plasma Membrane | other | |
| chr7 | *SUN1* | 30 | 1688 | 63 | SUN1 | Sad1 and UNC84 domain containing 1 | Nucleus | other | |
| chr2 | *AAK1* | 26 | 663 | 63 | AAK1 | AP2 associated kinase 1 | Cytoplasm | kinase | |
| chr20 | *ARFGEF2* | 27 | 837 | 63 | ARFGEF2 | ADP ribosylation factor guanine nucleotide exchange factor 2 | Cytoplasm | other | |
| chr15 | *DPH6* | 24 | 693 | 63 | DPH6 | diphthamine biosynthesis 6 | Cytoplasm | enzyme | |
| chr14 | *SPTB* | 24 | 907 | 63 | SPTB | spectrin beta, erythrocytic | Plasma Membrane | other | |
| chr2 | *ANXA4* | 26 | 609 | 63 | ANXA4 | annexin A4 | Plasma Membrane | other | |
| chr17 | *METTL16* | 25 | 693 | 63 | METTL16 | methyltransferase like 16 | Nucleus | enzyme | |
| chr19 | *FCGBP* | 30 | 1847 | 63 | FCGBP | Fc fragment of IgG binding | Extracellular Space | other | |
| chr9 | *STXBP1* | 28 | 1141 | 63 | STXBP1 | syntaxin binding protein 1 | Cytoplasm | transporter | |
| chr12 | *GRIP1* | 28 | 579 | 63 | GRIP1 | glutamate receptor interacting protein 1 | Plasma Membrane | transcription regulator | |
| chr6 | *SYNJ2* | 25 | 941 | 63 | SYNJ2 | synaptojanin 2 | Cytoplasm | phosphatase | |
| chr3 | *HRH1* | 27 | 843 | 63 | HRH1 | histamine receptor H1 | Plasma Membrane | G-protein coupled recepto | orphenadrine, clemizole, azatadine/pseudoephedrine, loxapine, diphenhydramine/ibuprofen, brompheniramine/codeine/phenylpropanolamine, pheniramine/phenyltoloxamine/pyrilamine, betahistine, loratadine/pseudoephedrine, chlorpheniramine/ibuprofen/pseudoephedrine, cimetidine/dexamethasone/diphenhydramine, chlorpheniramine/phenylephrine/pyrilamine, antazoline, [illegible] |
| chr22 | *CECR2* | 22 | 587 | 63 | CECR2 | CECR2 histone acetyl-lysine | Nucleus | other | |
| chr17 | *RP11-81K2.1* | 27 | 881 | 63 | | | | | |
| chr1 | *DPYD* | 24 | 699 | 63 | DPYD | dihydropyrimidine dehydrogenase | Cytoplasm | enzyme | gimeracil/irinotecan/oxonic acid/tegafur, uracil, bevacizumab/gimeracil/irinotecan/oxonic acid/tegafur, tegafur/uracil, eniluracil, gimeracil/oxonic acid/tegafur, cetuximab/gimeracil/irinotecan/oxonic acid/tegafur, gimeracil |
| chr17 | *UBE2O* | 26 | 1036 | 63 | UBE2O | ubiquitin conjugating enzyme E2 | Nucleus | enzyme | |
| chr10 | *ZNF365* | 28 | 1140 | 63 | ZNF365 | zinc finger protein 365 | Cytoplasm | other | |
| chr11 | *RP11-65M17.3* | 30 | 657 | 63 | | | | | |
| chr11 | *POLD3* | 29 | 826 | 63 | POLD3 | DNA polymerase delta 3, accessory subunit | Nucleus | transcription regulator | |
| chr11 | *PIK3C2A* | 25 | 613 | 63 | PIK3C2A | phosphatidylinositol-4-phosphate 3-kinase catalytic subunit type 2 | Cytoplasm | kinase | |
| chr3 | *AC133680.1* | 28 | 726 | 63 | | | | | |
| chr2 | *MAP4K4* | 26 | 756 | 63 | MAP4K4 | mitogen-activated protein kinase kinase kinase kinase 4 | Cytoplasm | kinase | |
| chr19 | *TDRD12* | 26 | 714 | 63 | TDRD12 | tudor domain containing 12 | Other | other | |
| chrX | *REPS2* | 26 | 586 | 63 | REPS2 | RALBP1 associated Eps domain containing 2 | Cytoplasm | other | |
| chr11 | *CEP164* | 26 | 826 | 63 | CEP164 | centrosomal protein 164 | Cytoplasm | other | |
| chr20 | *TSHZ2* | 29 | 963 | 63 | TSHZ2 | teashirt zinc finger homeobox 2 | Other | other | |
| chr1 | *SDCCAG8* | 27 | 819 | 63 | SDCCAG8 | serologically defined colon cancer antigen 8 | Cytoplasm | other | |
| chr13 | *NALCN-AS1* | 28 | 766 | 63 | NALCN-AS1 | NALCN antisense RNA 1 | Other | other | |
| chr12 | *LIMA1* | 27 | 1061 | 63 | LIMA1 | LIM domain and actin binding 1 | Cytoplasm | other | |
| chr11 | *MACROD1* | 28 | 1336 | 63 | MACROD1 | mono-ADP ribosylhydrolase 1 | Cytoplasm | enzyme | |
| chr16 | *CA5A* | 25 | 1126 | 63 | CA5A | carbonic anhydrase 5A | Cytoplasm | enzyme | hydrochlorothiazide, trichloromethiazide, sulfacetamide, benzthiazide, topiramate, chlorothiazide, chlorthalidone, acetazolamide, methazolamide |

| | | | | | | | | | |
|---|---|---|---|---|---|---|---|---|---|
| chr12 | *TPCN1* | 26 | 1087 | 63 | TPCN1 | two pore segment channel 1 | Plasma Membrane | ion channel | |
| chr1 | *EFCAB2* | 27 | 667 | 63 | EFCAB2 | EF-hand calcium binding domain | Other | other | |
| chr9 | *PBX3* | 25 | 768 | 63 | PBX3 | PBX homeobox 3 | Nucleus | transcription regulator | |
| chr10 | *JAKMIP3* | 29 | 1377 | 63 | JAKMIP3 | Janus kinase and microtubule interacting protein 3 | Other | other | |
| chr16 | *IL34* | 27 | 957 | 63 | IL34 | interleukin 34 | Extracellular Space | other | |
| chr22 | *LARGE* | 25 | 608 | 63 | LARGE1 | LARGE xylosyl- and glucuronyltransferase 1 | Cytoplasm | enzyme | |
| chr2 | *POLR1A* | 26 | 927 | 63 | POLR1A | RNA polymerase I subunit A | Nucleus | enzyme | |
| chr11 | *SYT12* | 28 | 1761 | 63 | SYT12 | synaptotagmin 12 | Plasma Membrane | transporter | |
| chr7 | *CALD1* | 26 | 729 | 62 | CALD1 | caldesmon 1 | Cytoplasm | other | |
| chr17 | *LLGL2* | 28 | 1363 | 62 | LLGL2 | LLGL scribble cell polarity complex component 2 | Cytoplasm | other | |
| chr3 | *ANO10* | 25 | 653 | 62 | ANO10 | anoctamin 10 | Plasma Membrane | ion channel | |
| chrX | *GPM6B* | 26 | 598 | 62 | GPM6B | glycoprotein M6B | Plasma Membrane | other | |
| chr11 | *RAB6A* | 25 | 796 | 62 | RAB6A | RAB6A, member RAS oncogene family | Cytoplasm | enzyme | |
| chr15 | *RP11-718O11.1* | 26 | 626 | 62 | | | | | |
| chr16 | *GSE1* | 28 | 1333 | 62 | GSE1 | Gse1 coiled-coil protein | Extracellular Space | other | |
| chrX | *MAP3K15* | 28 | 708 | 62 | MAP3K15 | mitogen-activated protein kinase kinase kinase 15 | Other | other | |
| chr9 | *CDK5RAP2* | 27 | 1025 | 62 | CDK5RAP2 | CDK5 regulatory subunit associated protein 2 | Cytoplasm | other | |
| chr9 | *PHF2* | 26 | 1073 | 62 | PHF2 | PHD finger protein 2 | Nucleus | enzyme | |
| chr3 | *CLASP2* | 23 | 588 | 62 | CLASP2 | cytoplasmic linker associated protein 2 | Cytoplasm | other | |
| chr1 | *ZCCHC11* | 23 | 554 | 62 | TUT4 | terminal uridylyl transferase 4 | Nucleus | enzyme | |
| chr11 | *LUZP2* | 23 | 570 | 62 | LUZP2 | leucine zipper protein 2 | Other | other | |
| chr15 | *TTBK2* | 23 | 626 | 62 | TTBK2 | tau tubulin kinase 2 | Cytoplasm | kinase | |
| chr9 | *GSN* | 27 | 1296 | 62 | GSN | gelsolin | Extracellular Space | other | |
| chrX | *PPEF1* | 24 | 692 | 62 | PPEF1 | protein phosphatase with EF-hand domain 1 | Extracellular Space | phosphatase | |
| chr7 | *AKAP9* | 23 | 713 | 62 | AKAP9 | A-kinase anchoring protein 9 | Cytoplasm | other | |
| chr1 | *RP11-431K24.1* | 26 | 706 | 62 | | | | | |
| chr10 | *GFRA1* | 28 | 766 | 62 | GFRA1 | GDNF family receptor alpha 1 | Plasma Membrane | transmembrane receptor | |
| chr1 | *ST6GALNAC3* | 27 | 532 | 62 | ST6GALNAC3 | ST6 N-acetylgalactosaminide alpha-2,6-sialyltransferase 3 | Cytoplasm | enzyme | |
| chr5 | *GHR* | 26 | 1954 | 62 | GHR | growth hormone receptor | Plasma Membrane | transmembrane receptor | GH1, prednisone/somatotropin, somatrem, pegvisomant |
| chr20 | *VSTM2L* | 26 | 1397 | 62 | VSTM2L | V-set and transmembrane domain containing 2 like | Extracellular Space | other | |
| chr17 | *WIPF2* | 26 | 659 | 62 | WIPF2 | WAS/WASL interacting protein family member 2 | Cytoplasm | other | |
| chr10 | *PAX2* | 28 | 972 | 62 | PAX2 | paired box 2 | Nucleus | transcription regulator | |
| chr1 | *NKAIN1* | 26 | 868 | 62 | NKAIN1 | sodium/potassium transporting ATPase interacting 1 | Plasma Membrane | other | |
| chr7 | *LMTK2* | 26 | 1170 | 62 | LMTK2 | lemur tyrosine kinase 2 | Cytoplasm | kinase | |
| chr20 | *JPH2* | 26 | 1225 | 62 | JPH2 | junctophilin 2 | Cytoplasm | enzyme | |
| chr11 | *MRPL48* | 27 | 941 | 62 | MRPL48 | mitochondrial ribosomal protein | Cytoplasm | other | |
| chr5 | *AP3B1* | 24 | 675 | 62 | AP3B1 | adaptor related protein complex 3 subunit beta 1 | Plasma Membrane | transporter | |
| chr16 | *TLDC1* | 26 | 999 | 62 | MEAK7 | MTOR associated protein, eak-7 homolog | Cytoplasm | other | |
| chr2 | *GREB1* | 24 | 878 | 62 | GREB1 | growth regulating estrogen receptor binding 1 | Cytoplasm | other | |
| chr10 | *ATE1* | 24 | 593 | 62 | ATE1 | arginyltransferase 1 | Nucleus | enzyme | |
| chr1 | *ADCK3* | 29 | 1392 | 62 | COQ8A | coenzyme Q8A | Cytoplasm | kinase | |
| chr1 | *LYST* | 27 | 709 | 62 | LYST | lysosomal trafficking regulator | Cytoplasm | transporter | |
| chr11 | *ARHGAP42* | 25 | 1334 | 62 | ARHGAP42 | Rho GTPase activating protein 42 | Other | other | |
| chr2 | *SPTBN1* | 28 | 984 | 62 | SPTBN1 | spectrin beta, non-erythrocytic 1 | Plasma Membrane | other | |
| chr2 | *AC016995.3* | 28 | 905 | 62 | | | | | |
| chrX | *APOO* | 24 | 660 | 62 | APOO | apolipoprotein O | Extracellular Space | other | |
| chr12 | *CNTN1* | 27 | 843 | 62 | CNTN1 | contactin 1 | Plasma Membrane | enzyme | |
| chr12 | *FAM19A2* | 24 | 604 | 62 | TAFA2 | TAFA chemokine like family member 2 | Cytoplasm | other | |
| chr20 | *SLC9A8* | 26 | 1147 | 62 | SLC9A8 | solute carrier family 9 member A8 | Cytoplasm | transporter | |
| chr1 | *GNB1* | 26 | 681 | 61 | GNB1 | G protein subunit beta 1 | Plasma Membrane | enzyme | |
| chr16 | *CMTM4* | 26 | 709 | 61 | CMTM4 | CKLF like MARVEL transmembrane domain | Extracellular Space | cytokine | |
| chr17 | *SPAG9* | 23 | 606 | 61 | SPAG9 | sperm associated antigen 9 | Cytoplasm | other | |
| chr2 | *DNMT3A* | 24 | 947 | 61 | DNMT3A | DNA methyltransferase 3 alpha | Nucleus | enzyme | |
| chr8 | *MCPH1* | 26 | 867 | 61 | MCPH1 | microcephalin 1 | Nucleus | other | |
| chr20 | *STAU1* | 23 | 938 | 61 | STAU1 | staufen double-stranded RNA binding protein 1 | Cytoplasm | transporter | |
| chr2 | *FER1L5* | 27 | 1230 | 61 | FER1L5 | fer-1 like family member 5 | Other | other | |
| chr15 | *SIN3A* | 24 | 722 | 61 | SIN3A | SIN3 transcription regulator family member A | Nucleus | transcription regulator | |
| chr12 | *RNFT2* | 28 | 727 | 61 | RNFT2 | ring finger protein, | Other | other | |
| chr21 | *ABCG1* | 24 | 906 | 61 | ABCG1 | ATP binding cassette subfamily G member 1 | Plasma Membrane | transporter | |
| chr12 | *MLXIP* | 26 | 1063 | 61 | MLXIP | MLX interacting protein | Nucleus | transcription regulator | |
| chr14 | *TRAF3* | 24 | 804 | 61 | TRAF3 | TNF receptor associated factor 3 | Cytoplasm | enzyme | |
| chr11 | *CAPN5* | 26 | 1743 | 61 | CAPN5 | calpain 5 | Cytoplasm | peptidase | |
| chr10 | *TSPAN15* | 27 | 1705 | 61 | TSPAN15 | tetraspanin 15 | Plasma Membrane | other | |
| chr10 | *AFAP1L2* | 26 | 1036 | 61 | AFAP1L2 | actin filament associated protein 1 like 2 | Cytoplasm | other | |
| chr2 | *AC009236.1* | 29 | 1273 | 61 | | | | | |

| | | | | | | | | | |
|---|---|---|---|---|---|---|---|---|---|
| chr21 | *HLCS* | 23 | 710 | 61 | HLCS | holocarboxylase synthetase | Cytoplasm | enzyme | |
| chr19 | *ACTN4* | 27 | 1207 | 61 | ACTN4 | actinin alpha 4 | Cytoplasm | transcription regulator | |
| chr10 | *SORBS1* | 27 | 788 | 61 | SORBS1 | sorbin and SH3 domain | Plasma Membrane | other | |
| chr6 | *DNAH8* | 27 | 548 | 61 | DNAH8 | dynein axonemal heavy chain 8 | Cytoplasm | enzyme | |
| chr8 | *RP11-124B13.1* | 27 | 650 | 61 | | | | | |
| chr1 | *LRRC7* | 25 | 599 | 61 | LRRC7 | leucine rich repeat containing 7 | Plasma Membrane | other | |
| chr16 | *ABCC6* | 26 | 945 | 61 | ABCC6 | ATP binding cassette subfamily C member 6 | Plasma Membrane | transporter | |
| chr5 | *DMGDH* | 26 | 572 | 61 | DMGDH | dimethylglycine dehydrogenase | Cytoplasm | enzyme | |
| chr5 | *GUSBP1* | 27 | 4678 | 61 | GUSBP1 | GUSB pseudogene 1 | Other | other | |
| chr8 | *EBF2* | 28 | 660 | 61 | EBF2 | EBF transcription factor 2 | Nucleus | transcription regulator | |
| chr16 | *SPIRE2* | 26 | 1200 | 61 | SPIRE2 | spire type actin nucleation factor | Cytoplasm | other | |
| chr2 | *REEP1* | 26 | 829 | 61 | REEP1 | receptor accessory protein 1 | Cytoplasm | other | |
| chr7 | *TPK1* | 23 | 578 | 61 | TPK1 | thiamin pyrophosphokinase 1 | Cytoplasm | kinase | |
| chr10 | *FGFR2* | 26 | 769 | 61 | FGFR2 | fibroblast growth factor receptor 2 | Plasma Membrane | kinase | BAY1179470, derazantinib, bortezomib/dexamethasone/thalidomide, thalidomide, rituximab/thalidomide, ASP5878, PRN1371, lenvatinib, 3D185, erdafitinib, debio 1347, prednisone/thalidomide, aprutumab ixadotin, regorafenib, lenvatinib/pembrolizumab, dexamethasone/thalidomide, pazopanib, docetaxel/nintedanib, everolimus/lenvatinib, FPA144, alofanib, pemigatinib, SKI-G-801, palifermin, lucitanib |
| chr5 | *RP11-423H2.1* | 25 | 806 | 61 | | | | | |
| chr1 | *MAGI3* | 24 | 989 | 61 | MAGI3 | membrane associated guanylate kinase, WW and PDZ domain containing 3 | Cytoplasm | kinase | |
| chr16 | *U91319.1* | 28 | 524 | 61 | | | | | |
| chr16 | *IQCK* | 25 | 592 | 61 | IQCK | IQ motif containing K | Other | other | |
| chrX | *FAM120C* | 24 | 897 | 61 | FAM120C | family with sequence similarity | Other | other | |
| chr5 | *AHRR* | 31 | 3080 | 61 | AHRR | aryl-hydrocarbon receptor | Nucleus | transcription regulator | |
| chr11 | *ARHGAP32* | 27 | 748 | 61 | ARHGAP32 | Rho GTPase activating protein 32 | Cytoplasm | other | |
| chr7 | *ZNF804B* | 28 | 641 | 60 | ZNF804B | zinc finger protein 804B | Other | other | |
| chr15 | *HERC1* | 22 | 570 | 60 | HERC1 | HECT and RLD domain containing E3 ubiquitin protein ligase family member 1 | Cytoplasm | other | |
| chr14 | *EML1* | 24 | 781 | 60 | EML1 | EMAP like 1 | Cytoplasm | other | |
| chr12 | *TXNRD1* | 23 | 710 | 60 | TXNRD1 | thioredoxin reductase 1 | Cytoplasm | enzyme | arsenic trioxide/cytarabine/methotrexate, arsenic trioxide/idarubicin/tretinoin, arsenic trioxide/tretinoin, arsenic trioxide, arsenic trioxide/daunorubicin/tretinoin, ethaselen, arsenic trioxide/gemtuzumab |
| chr8 | *MSR1* | 25 | 551 | 60 | MSR1 | macrophage scavenger receptor | Plasma Membrane | transmembrane receptor | |
| chrX | *WWC3* | 26 | 858 | 60 | WWC3 | WWC family member 3 | Cytoplasm | other | |
| chr12 | *LARP4* | 25 | 1008 | 60 | LARP4 | La ribonucleoprotein domain family member 4 | Cytoplasm | other | |
| chr10 | *VCL* | 27 | 1008 | 60 | VCL | vinculin | Plasma Membrane | enzyme | |
| chr6 | *LRRC16A* | 25 | 602 | 60 | CARMIL1 | capping protein regulator and myosin 1 linker 1 | Cytoplasm | enzyme | |
| chr2 | *ANTXR1* | 27 | 651 | 60 | ANTXR1 | ANTXR cell adhesion molecule 1 | Plasma Membrane | transmembrane receptor | |
| chr19 | *MYO1F* | 25 | 931 | 60 | MYO1F | myosin IF | Cytoplasm | other | |
| chr17 | *ARHGAP44* | 26 | 657 | 60 | ARHGAP44 | Rho GTPase activating protein 44 | Cytoplasm | other | |
| chr12 | *TFCP2* | 22 | 970 | 60 | TFCP2 | transcription factor CP2 | Nucleus | transcription regulator | |
| chr12 | *TMEM120B* | 26 | 1000 | 60 | TMEM120B | transmembrane protein 120B | Nucleus | other | |
| chr19 | *LIPE-AS1* | 26 | 629 | 60 | LIPE-AS1 | LIPE antisense RNA 1 | Other | other | |
| chr2 | *EHBP1* | 22 | 653 | 60 | EHBP1 | EH domain binding protein 1 | Cytoplasm | other | |
| chr17 | *USP43* | 26 | 707 | 60 | USP43 | ubiquitin specific peptidase 43 | Nucleus | peptidase | |
| chr17 | *RP11-1407O15.2* | 25 | 2017 | 60 | | | | | |
| chr16 | *MGRN1* | 27 | 1287 | 60 | MGRN1 | mahogunin ring finger 1 | Cytoplasm | enzyme | |
| chr13 | *FGF14* | 25 | 540 | 60 | FGF14 | fibroblast growth factor 14 | Extracellular Space | growth factor | |
| chr10 | *SVIL* | 21 | 632 | 60 | SVIL | supervillin | Plasma Membrane | other | |
| chr7 | *ADAM22* | 24 | 785 | 60 | ADAM22 | ADAM metallopeptidase domain | Plasma Membrane | peptidase | |
| chr20 | *UQCC1* | 25 | 900 | 60 | UQCC1 | ubiquinol-cytochrome c reductase complex assembly factor 1 | Cytoplasm | other | |
| chr1 | *RP4-799D16.1* | 25 | 1010 | 60 | | | | | |
| chr12 | *GCN1L1* | 24 | 987 | 60 | GCN1 | GCN1 activator of EIF2AK4 | Cytoplasm | translation regulator | |
| chr10 | *EXOC6* | 25 | 555 | 60 | EXOC6 | exocyst complex component 6 | Plasma Membrane | transporter | |

| | | | | | | | | | |
|---|---|---|---|---|---|---|---|---|---|
| chr16 | *ABCA17P* | 25 | 702 | 60 | ABCA17P | ATP binding cassette subfamily A member 17, pseudogene | Other | other | |
| chr12 | *CCDC64* | 24 | 800 | 60 | BICDL1 | BICD family like cargo adaptor 1 | Cytoplasm | other | |
| chr17 | *TNRC6C* | 22 | 694 | 59 | TNRC6C | trinucleotide repeat containing adaptor 6C | Cytoplasm | other | |
| chr11 | *RP1-65P5.1* | 26 | 782 | 59 | | | | | |
| chr5 | *SNX24* | 21 | 556 | 59 | SNX24 | sorting nexin 24 | Other | transporter | |
| chr1 | *RUNX3* | 26 | 1164 | 59 | RUNX3 | RUNX family transcription factor | Nucleus | transcription regulator | |
| chr12 | *SRGAP1* | 23 | 598 | 59 | SRGAP1 | SLIT-ROBO Rho GTPase activating protein 1 | Cytoplasm | other | |
| chr2 | *OTOF* | 23 | 1047 | 59 | OTOF | otoferlin | Plasma Membrane | other | |
| chr9 | *PRRX2* | 25 | 1587 | 59 | PRRX2 | paired related homeobox 2 | Nucleus | transcription regulator | |
| chr3 | *IL12A-AS1* | 30 | 759 | 59 | IL12A-AS1 | IL12A antisense RNA 1 | Other | other | |
| chr16 | *XPO6* | 26 | 779 | 59 | XPO6 | exportin 6 | Cytoplasm | other | |
| chr1 | *PDE4B* | 25 | 561 | 59 | PDE4B | phosphodiesterase 4B | Cytoplasm | enzyme | caffeine, pentoxifylline, arofylline, tetomilast, tolbutamide, anagrelide, theophylline, apremilast, milrinone, cilomilast, roflumilast, ketotifen, L-826,141, L 869298, enprofylline, ibudilast, papaverine, |
| chr1 | *MDS2* | 24 | 890 | 59 | MDS2 | myelodysplastic syndrome 2 translocation associated | Extracellular Space | other | |
| chr16 | *ITFG1* | 23 | 635 | 59 | ITFG1 | integrin alpha FG-GAP repeat containing 1 | Plasma Membrane | other | |
| chr12 | *SBNO1* | 24 | 665 | 59 | SBNO1 | strawberry notch homolog 1 | Other | enzyme | |
| chr18 | *RNF165* | 22 | 887 | 59 | RNF165 | ring finger protein 165 | Nucleus | enzyme | |
| chr2 | *LPIN1* | 20 | 746 | 59 | LPIN1 | lipin 1 | Nucleus | phosphatase | |
| chr7 | *HEATR2* | 27 | 1581 | 59 | DNAAF5 | dynein axonemal assembly factor | Cytoplasm | other | |
| chr15 | *PAQR5* | 26 | 798 | 59 | PAQR5 | progestin and adipoQ receptor family member 5 | Other | other | |
| chr2 | *ADCY3* | 22 | 906 | 59 | ADCY3 | adenylate cyclase 3 | Plasma Membrane | enzyme | |
| chr4 | *RGS12* | 26 | 1132 | 59 | RGS12 | regulator of G protein signaling | Nucleus | enzyme | |
| chr20 | *CTNNBL1* | 27 | 941 | 59 | CTNNBL1 | catenin beta like 1 | Nucleus | other | |
| chr22 | *TXNRD2* | 25 | 1058 | 59 | TXNRD2 | thioredoxin reductase 2 | Cytoplasm | enzyme | |
| chr7 | *GATS* | 25 | 1290 | 59 | CASTOR3 | CASTOR family member 3 | Other | other | |
| chr11 | *PARVA* | 28 | 927 | 59 | PARVA | parvin alpha | Cytoplasm | other | |
| chr1 | *PADI2* | 25 | 1061 | 59 | PADI2 | peptidyl arginine deiminase 2 | Cytoplasm | enzyme | |
| chr2 | *COX7A2L* | 25 | 1295 | 59 | COX7A2L | cytochrome c oxidase subunit 7A2 like | Cytoplasm | enzyme | |
| chr5 | *ANKRD55* | 24 | 619 | 59 | ANKRD55 | ankyrin repeat domain 55 | Nucleus | transcription regulator | |
| chr8 | *BAI1* | 32 | 1567 | 59 | ADGRB1 | adhesion G protein-coupled receptor B1 | Plasma Membrane | G-protein coupled recepto | |
| chr12 | *ABCB9* | 26 | 996 | 59 | ABCB9 | ATP binding cassette subfamily B member 9 | Cytoplasm | transporter | |
| chr17 | *RGS9* | 27 | 744 | 59 | RGS9 | regulator of G protein signaling 9 | Cytoplasm | enzyme | |
| chr3 | *LINC00620* | 26 | 1056 | 59 | LINC00620 | long intergenic non-protein coding RNA 620 | Other | other | |
| chr22 | *PPP6R2* | 23 | 861 | 59 | PPP6R2 | protein phosphatase 6 regulatory subunit 2 | Cytoplasm | other | |
| chr17 | *YWHAE* | 24 | 947 | 59 | YWHAE | tyrosine 3-monooxygenase/tryptophan 5-monooxygenase activation protein epsilon | Cytoplasm | other | |
| chr7 | *CDK13* | 23 | 694 | 59 | CDK13 | cyclin dependent kinase 13 | Nucleus | kinase | |
| chr6 | *COL21A1* | 25 | 595 | 59 | COL21A1 | collagen type XXI alpha 1 chain | Extracellular Space | other | collagenase clostridium |
| chr11 | *SLCO2B1* | 26 | 1333 | 59 | SLCO2B1 | solute carrier organic anion transporter family member 2B1 | Plasma Membrane | transporter | |
| chr22 | *MPPED1* | 21 | 941 | 59 | MPPED1 | metallophosphoesterase domain containing 1 | Other | other | |
| chr15 | *RP11-387D10.2* | 28 | 1463 | 59 | | | | | |
| chr1 | *NID1* | 26 | 940 | 59 | NID1 | nidogen 1 | Extracellular Space | other | |
| chr17 | *NPEPPS* | 23 | 665 | 59 | NPEPPS | aminopeptidase puromycin | Cytoplasm | peptidase | |
| chr7 | *NPSR1-AS1* | 27 | 775 | 59 | NPSR1-AS1 | NPSR1 antisense RNA 1 | Other | other | |
| chr8 | *PPP3CC* | 21 | 557 | 59 | PPP3CC | protein phosphatase 3 catalytic subunit gamma | Cytoplasm | phosphatase | voclosporin, cyclosporin A, pimecrolimus, prednisone/tacrolimus, cyclosporin A/methotrexate, methylprednisolone/t |
| chr11 | *SPON1* | 26 | 663 | 59 | SPON1 | spondin 1 | Extracellular Space | other | |
| chrX | *CTPS2* | 25 | 639 | 59 | CTPS2 | CTP synthase 2 | Cytoplasm | enzyme | |
| chr1 | *NR5A2* | 24 | 855 | 59 | NR5A2 | nuclear receptor subfamily 5 group A member 2 | Nucleus | d-dependent nuclear rec | |
| chr16 | *AC004158.2* | 24 | 565 | 59 | | | | | |
| chr2 | *PXDN* | 24 | 1037 | 58 | PXDN | peroxidasin | Extracellular Space | enzyme | |
| chr14 | *C14orf159* | 21 | 667 | 58 | DGLUCY | D-glutamate cyclase | Cytoplasm | enzyme | |
| chr17 | *AC005562.1* | 24 | 797 | 58 | | | | | |
| chr12 | *SOX5* | 24 | 566 | 58 | SOX5 | SRY-box transcription factor 5 | Nucleus | transcription regulator | |
| chr17 | *PAFAH1B1* | 24 | 611 | 58 | PAFAH1B1 | platelet activating factor acetylhydrolase 1b regulatory subunit 1 | Cytoplasm | enzyme | |

| | | | | | | | | | |
|---|---|---|---|---|---|---|---|---|---|
| chr2 | *PLB1* | 24 | 672 | 58 | PLB1 | phospholipase B1 | Cytoplasm | enzyme | |
| chr7 | *RP11-306G20.1* | 24 | 734 | 58 | | | | | |
| chr6 | *PRIM2* | 27 | 979 | 58 | PRIM2 | DNA primase subunit 2 | Nucleus | enzyme | alemtuzumab/cyclophosphamide/fludarabine phosphate, cytarabine/filgrastim/fludarabine phosphate/imatinib, fludarabine phosphate/rituximab, cytarabine/fludarabine phosphate/oxaliplatin/rituximab, dexamethasone/fludarabine phosphate/mitoxantrone/rituximab, cytarabine/fludarabine phosphate, cytarabine/filgrastim/fludarabine phosphate, cyclophosphamide/fludarabine [illegible] |
| chr13 | *NALCN* | 23 | 580 | 58 | NALCN | sodium leak channel, non- | Plasma Membrane | ion channel | |
| chr4 | *STK32B* | 22 | 647 | 58 | STK32B | serine/threonine kinase 32B | Other | kinase | |
| chr5 | *RP11-267A15.1* | 26 | 606 | 58 | | | | | |
| chr11 | *NUCB2* | 25 | 516 | 58 | NUCB2 | nucleobindin 2 | Nucleus | other | |
| chr3 | *ARIH2* | 23 | 691 | 58 | ARIH2 | ariadne RBR E3 ubiquitin protein ligase 2 | Nucleus | enzyme | |
| chr3 | *WNT7A* | 25 | 1024 | 58 | WNT7A | Wnt family member 7A | Extracellular Space | cytokine | |
| chr7 | *FBXL13* | 26 | 637 | 58 | FBXL13 | F-box and leucine rich repeat protein 13 | Cytoplasm | enzyme | |
| chr15 | *RP11-279F6.3* | 25 | 777 | 58 | | | | | |
| chr1 | *ACOT11* | 25 | 944 | 58 | ACOT11 | acyl-CoA thioesterase 11 | Cytoplasm | enzyme | |
| chr16 | *PDPK1* | 26 | 1308 | 58 | PDPK1 | 3-phosphoinositide dependent protein kinase 1 | Cytoplasm | kinase | |
| chr6 | *BMP6* | 23 | 637 | 58 | BMP6 | bone morphogenetic protein 6 | Extracellular Space | growth factor | |
| chr1 | *LUZP1* | 24 | 628 | 58 | LUZP1 | leucine zipper protein 1 | Nucleus | other | |
| chr8 | *INTS9* | 23 | 692 | 58 | INTS9 | integrator complex subunit 9 | Nucleus | other | |
| chr15 | *ACSBG1* | 25 | 1247 | 58 | ACSBG1 | acyl-CoA synthetase bubblegum family member 1 | Cytoplasm | enzyme | |
| chr2 | *BRE* | 21 | 603 | 58 | BABAM2 | BRISC and BRCA1 A complex member 2 | Cytoplasm | other | |
| chr12 | *RP11-114H23.1* | 26 | 550 | 58 | | | | | |
| chr18 | *RP11-161I6.2* | 27 | 912 | 58 | | | | | |
| chr3 | *RFTN1* | 28 | 961 | 58 | RFTN1 | raftlin, lipid raft linker 1 | Plasma Membrane | other | |
| chr11 | *PPFIBP2* | 27 | 1207 | 58 | PPFIBP2 | PPFIA binding protein 2 | Nucleus | phosphatase | |
| chr2 | *SH3BP4* | 22 | 875 | 58 | SH3BP4 | SH3 domain binding protein 4 | Cytoplasm | other | |
| chr7 | *LMBR1* | 23 | 601 | 58 | LMBR1 | limb development membrane protein 1 | Plasma Membrane | transmembrane receptor | |
| chr1 | *PGBD5* | 26 | 1230 | 58 | PGBD5 | piggyBac transposable element derived 5 | Nucleus | enzyme | |
| chr12 | *ARID2* | 23 | 782 | 58 | ARID2 | AT-rich interaction domain 2 | Nucleus | transcription regulator | |
| chr12 | *POLE* | 24 | 1079 | 58 | POLE | DNA polymerase epsilon, catalytic subunit | Nucleus | enzyme | cladribine/cytarabine/filgrastim/idarubicin/plerixafor, clofarabine/cyclophosphamide, nelarabine, cladribine, cladribine/cytarabine/daunorubicin, trifluridine, clofarabine/cyclophosphamide/etoposide/imatinib, cladribine/cytarabine/decitabine, clofarabine, cladribine/cytarabine/filgrastim/idarubicin, clofarabine/filgrastim, clofarabine/cytarabine/filgrastim, cladribine/recombinant interferon, [illegible] |
| chr7 | *OSBPL3* | 23 | 822 | 58 | OSBPL3 | oxysterol binding protein like 3 | Cytoplasm | other | |
| chr7 | *GTF2IRD2B* | 23 | 954 | 58 | F2IRD2/GTF2IRD | GTF2I repeat domain containing | Nucleus | other | |
| chr19 | *GPATCH1* | 23 | 716 | 58 | GPATCH1 | G-patch domain containing 1 | Extracellular Space | other | |
| chr15 | *PDE8A* | 26 | 789 | 58 | PDE8A | phosphodiesterase 8A | Cytoplasm | enzyme | pentoxifylline, tolbutamide, anagrelide, theophylline, milrinone, |
| chr2 | *KIF1A* | 26 | 1405 | 57 | KIF1A | kinesin family member 1A | Cytoplasm | other | |

| | | | | | | | | | |
|---|---|---|---|---|---|---|---|---|---|
| chrX | *IQSEC2* | 26 | 1193 | 57 | IQSEC2 | IQ motif and Sec7 domain | Cytoplasm | other | |
| chr1 | *SLC30A10* | 26 | 755 | 57 | SLC30A10 | solute carrier family 30 member | Plasma Membrane | transporter | |
| chr6 | *MDGA1* | 26 | 1020 | 57 | MDGA1 | MAM domain containing glycosylphosphatidylinositol | Plasma Membrane | other | |
| chr9 | *GLDC* | 24 | 589 | 57 | GLDC | glycine decarboxylase | Cytoplasm | enzyme | |
| chr2 | *WDR33* | 24 | 852 | 57 | WDR33 | WD repeat domain 33 | Nucleus | other | |
| chr5 | *UIMC1* | 21 | 552 | 57 | UIMC1 | ubiquitin interaction motif | Nucleus | other | |
| chr3 | *PLXND1* | 27 | 1644 | 57 | PLXND1 | plexin D1 | Plasma Membrane | transmembrane receptor | |
| chr19 | *KIAA0355* | 25 | 670 | 57 | KIAA0355 | KIAA0355 | Extracellular Space | other | |
| chr16 | *USP10* | 21 | 737 | 57 | USP10 | ubiquitin specific peptidase 10 | Cytoplasm | peptidase | |
| chr5 | *DDX46* | 23 | 597 | 57 | DDX46 | DEAD-box helicase 46 | Nucleus | enzyme | |
| chr1 | *ARID1A* | 25 | 751 | 57 | ARID1A | AT-rich interaction domain 1A | Nucleus | transcription regulator | |
| chr1 | *PIK3CD* | 25 | 898 | 57 | PIK3CD | phosphatidylinositol-4,5-bisphosphate 3-kinase catalytic subunit delta | Cytoplasm | kinase | buparlisib, SHC014748M, nemiralisib, GS-9901, parsaclisib, duvelisib, umbralisib, idelalisib/ofatumumab, PWT143, ACP-196/ACP-319, idelalisib, YY-20394, dezapelisib, bimiralisib, bendamustine/idelalisib/rituximab, copanlisib, dactolisib, acalisib, tenalisib, idelalisib/rituximab, pilaralisib, KA2237, PX-866, SF 1126 |
| chr1 | *TP73* | 28 | 1370 | 57 | TP73 | tumor protein p73 | Nucleus | transcription regulator | |
| chr11 | *RP11-430H10.4* | 27 | 923 | 57 | | | | | |
| chr20 | *RBL1* | 25 | 721 | 57 | RBL1 | RB transcriptional corepressor | Nucleus | transcription regulator | |
| chr15 | *RTF1* | 22 | 596 | 57 | RTF1 | RTF1 homolog, Paf1/RNA polymerase II complex | Nucleus | other | |
| chr1 | *AXDND1* | 24 | 675 | 57 | AXDND1 | axonemal dynein light chain domain containing 1 | Extracellular Space | other | |
| chr6 | *PACRG* | 20 | 717 | 57 | PACRG | parkin coregulated | Nucleus | other | |
| chr22 | *RP11-191L9.4* | 20 | 696 | 57 | | | | | |
| chr7 | *SMURF1* | 22 | 766 | 57 | SMURF1 | SMAD specific E3 ubiquitin protein ligase 1 | Cytoplasm | enzyme | |
| chr20 | *EDEM2* | 23 | 811 | 57 | EDEM2 | ER degradation enhancing alpha-mannosidase like protein 2 | Cytoplasm | enzyme | |
| chr11 | *KCNC1* | 24 | 1167 | 57 | KCNC1 | potassium voltage-gated channel subfamily C member 1 | Plasma Membrane | ion channel | dalfampridine |
| chr12 | *TSPAN11* | 26 | 1215 | 57 | TSPAN11 | tetraspanin 11 | Other | other | |
| chr9 | *DENND4C* | 22 | 562 | 57 | DENND4C | DENN domain containing 4C | Cytoplasm | other | |
| chr16 | *ZC3H18* | 24 | 1170 | 57 | ZC3H18 | zinc finger CCCH-type containing | Nucleus | other | |
| chr1 | *AKT3* | 22 | 654 | 57 | AKT3 | AKT serine/threonine kinase 3 | Cytoplasm | kinase | triciribine, triciribine phosphate, enzastaurin, GSK690693 |
| chr1 | *IL6R* | 25 | 1081 | 57 | IL6R | interleukin 6 receptor | Plasma Membrane | transmembrane receptor | methotrexate/tocilizumab, tocilizumab, methotrexate/sarilumab, sarilumab |
| chr20 | *PTGIS* | 25 | 1101 | 57 | PTGIS | prostaglandin I2 synthase | Cytoplasm | enzyme | phenylbutazone |
| chr17 | *RABEP1* | 19 | 539 | 57 | RABEP1 | rabaptin, RAB GTPase binding effector protein 1 | Cytoplasm | transporter | |
| chr15 | *TIPIN* | 20 | 623 | 57 | TIPIN | TIMELESS interacting protein | Nucleus | other | |
| chr1 | *KIF17* | 24 | 1104 | 57 | KIF17 | kinesin family member 17 | Cytoplasm | transporter | |
| chr7 | *STEAP1B* | 25 | 841 | 57 | STEAP1B | STEAP family member 1B | Other | other | |
| chr19 | *PAK4* | 24 | 1200 | 57 | PAK4 | p21 (RAC1) activated kinase 4 | Cytoplasm | kinase | PF-3758309, KPT-9274 |
| chr17 | *GRB2* | 22 | 674 | 57 | GRB2 | growth factor receptor bound protein 2 | Cytoplasm | other | liposome-incorporated Grb2 antisense |
| chr11 | *SYT7* | 27 | 1830 | 57 | SYT7 | synaptotagmin 7 | Cytoplasm | transporter | |
| chr3 | *GRIP2* | 22 | 1194 | 57 | GRIP2 | glutamate receptor interacting protein 2 | Plasma Membrane | other | |
| chr11 | *NRXN2* | 23 | 998 | 57 | NRXN2 | neurexin 2 | Plasma Membrane | transporter | |
| chr11 | *AP2A2* | 24 | 1222 | 57 | AP2A2 | adaptor related protein complex 2 subunit alpha 2 | Cytoplasm | transporter | |
| chr7 | *ATP6V0A4* | 26 | 804 | 57 | ATP6V0A4 | ATPase H+ transporting V0 subunit a4 | Cytoplasm | transporter | |
| chr2 | *EPB41L5* | 22 | 766 | 57 | EPB41L5 | erythrocyte membrane protein band 4.1 like 5 | Plasma Membrane | other | |
| chr9 | *DMRT1* | 25 | 574 | 57 | DMRT1 | doublesex and mab-3 related transcription factor 1 | Nucleus | transcription regulator | |
| chr13 | *LINC00393* | 23 | 568 | 56 | LINC00393 | long intergenic non-protein coding RNA 393 | Other | other | |
| chr19 | *CEP89* | 25 | 757 | 56 | CEP89 | centrosomal protein 89 | Cytoplasm | other | |
| chr1 | *INPP5B* | 25 | 621 | 56 | INPP5B | inositol polyphosphate-5-phosphatase B | Plasma Membrane | phosphatase | |
| chr16 | *ADCY7* | 24 | 1215 | 56 | ADCY7 | adenylate cyclase 7 | Plasma Membrane | enzyme | |
| chr9 | *SETX* | 20 | 680 | 56 | SETX | senataxin | Nucleus | enzyme | |
| chr15 | *SEMA4B* | 25 | 1434 | 56 | SEMA4B | semaphorin 4B | Plasma Membrane | other | |
| chr13 | *UBAC2* | 24 | 930 | 56 | UBAC2 | UBA domain containing 2 | Cytoplasm | enzyme | |
| chr1 | *WDTC1* | 22 | 701 | 56 | WDTC1 | WD and tetratricopeptide repeats | Nucleus | other | |

| | | | | | | | | | |
|---|---|---|---|---|---|---|---|---|---|
| chr16 | *SRL* | 24 | 935 | 56 | SRL | sarcalumenin | Cytoplasm | other | |
| chr17 | *ANKFY1* | 22 | 690 | 56 | ANKFY1 | ankyrin repeat and FYVE domain containing 1 | Cytoplasm | transcription regulator | |
| chr13 | *CARS2* | 23 | 1031 | 56 | CARS2 | cysteinyl-tRNA synthetase 2, mitochondrial | Cytoplasm | enzyme | |
| chr16 | *FA2H* | 23 | 996 | 56 | FA2H | fatty acid 2-hydroxylase | Cytoplasm | enzyme | |
| chr17 | *PRPSAP1* | 23 | 698 | 56 | PRPSAP1 | phosphoribosyl pyrophosphate synthetase associated protein 1 | Extracellular Space | other | |
| chr1 | *RPS6KA1* | 23 | 977 | 56 | RPS6KA1 | ribosomal protein S6 kinase A1 | Cytoplasm | kinase | PMD-026 |
| chr6 | *UHRF1BP1* | 22 | 584 | 56 | UHRF1BP1 | UHRF1 binding protein 1 | Other | other | |
| chr2 | *PLEKHH2* | 22 | 627 | 56 | PLEKHH2 | pleckstrin homology, MyTH4 and FERM domain containing H2 | Cytoplasm | other | |
| chr19 | *ANKRD27* | 23 | 843 | 56 | ANKRD27 | ankyrin repeat domain 27 | Cytoplasm | other | |
| chr3 | *SPSB4* | 26 | 1048 | 56 | SPSB4 | splA/ryanodine receptor domain and SOCS box containing 4 | Cytoplasm | other | |
| chr9 | *TRAF2* | 24 | 1570 | 56 | TRAF2 | TNF receptor associated factor 2 | Cytoplasm | enzyme | |
| chr18 | *TMEM241* | 21 | 633 | 56 | TMEM241 | transmembrane protein 241 | Other | other | |
| chr11 | *FAT3* | 28 | 1130 | 56 | FAT3 | FAT atypical cadherin 3 | Plasma Membrane | other | |
| chr1 | *RPS6KC1* | 23 | 693 | 56 | RPS6KC1 | ribosomal protein S6 kinase C1 | Cytoplasm | kinase | |
| chr19 | *KCNN1* | 23 | 918 | 56 | KCNN1 | potassium calcium-activated channel subfamily N member 1 | Plasma Membrane | ion channel | |
| chr16 | *KLHDC4* | 25 | 1060 | 56 | KLHDC4 | kelch domain containing 4 | Other | other | |
| chr15 | *MGA* | 20 | 550 | 56 | MGA | MAX dimerization protein MGA | Nucleus | transcription regulator | |
| chr7 | *PODXL* | 24 | 894 | 56 | PODXL | podocalyxin like | Plasma Membrane | kinase | |
| chr10 | *KIAA1217* | 23 | 572 | 56 | KIAA1217 | KIAA1217 | Cytoplasm | other | |
| chr15 | *INO80* | 22 | 570 | 56 | INO80 | INO80 complex ATPase subunit | Nucleus | enzyme | |
| chr16 | *NKD1* | 23 | 966 | 56 | NKD1 | NKD inhibitor of WNT signaling pathway 1 | Other | other | |
| chr2 | *NGEF* | 20 | 860 | 56 | NGEF | neuronal guanine nucleotide exchange factor | Cytoplasm | other | |
| chr6 | *PHACTR2* | 24 | 538 | 56 | PHACTR2 | phosphatase and actin regulator | Other | other | |
| chr2 | *KCMF1* | 20 | 571 | 56 | KCMF1 | potassium channel modulatory factor 1 | Extracellular Space | enzyme | |
| chr3 | *CD96* | 23 | 761 | 56 | CD96 | CD96 molecule | Plasma Membrane | other | |
| chr1 | *DISC1* | 23 | 706 | 56 | DISC1 | DISC1 scaffold protein | Cytoplasm | other | |
| chr5 | *RASGRF2* | 24 | 553 | 56 | RASGRF2 | Ras protein specific guanine nucleotide releasing factor 2 | Cytoplasm | other | |
| chr17 | *WNT3* | 22 | 1039 | 56 | WNT3 | Wnt family member 3 | Extracellular Space | other | |
| chr5 | *PHF15* | 22 | 1043 | 56 | JADE2 | jade family PHD finger 2 | Nucleus | enzyme | |
| chr5 | *HCN1* | 25 | 1570 | 56 | HCN1 | hyperpolarization activated cyclic nucleotide gated potassium channel 1 | Plasma Membrane | ion channel | |
| chr2 | *MERTK* | 24 | 801 | 56 | MERTK | MER proto-oncogene, tyrosine kinase | Plasma Membrane | kinase | MRX-2843, cabozantinib/nivolumab, cabozantinib/erlotinib, ningetinib, ONO-7475, sitravatinib, INCB081776, SKI-G-801, RXDX-106, |
| chr5 | *CTD-2337A12.1* | 25 | 529 | 56 | | | | | |
| chr11 | *ARHGEF17* | 26 | 1363 | 56 | ARHGEF17 | Rho guanine nucleotide exchange factor 17 | Cytoplasm | other | |
| chr5 | *NIPBL* | 23 | 1663 | 56 | NIPBL | NIPBL cohesin loading factor | Nucleus | transcription regulator | |
| chr1 | *GMEB1* | 22 | 680 | 56 | GMEB1 | glucocorticoid modulatory element binding protein 1 | Nucleus | transcription regulator | |
| chr7 | *CHCHD3* | 22 | 516 | 56 | CHCHD3 | coiled-coil-helix-coiled-coil-helix domain containing 3 | Cytoplasm | transcription regulator | |
| chr11 | *KIAA1549L* | 26 | 833 | 56 | KIAA1549L | KIAA1549 like | Cytoplasm | other | |
| chr12 | *IFT81* | 23 | 563 | 56 | IFT81 | intraflagellar transport 81 | Extracellular Space | other | |
| chr20 | *CHD6* | 24 | 740 | 55 | CHD6 | chromodomain helicase DNA binding protein 6 | Nucleus | enzyme | |
| chr8 | *RP11-624C23.1* | 25 | 503 | 55 | | | | | |
| chr22 | *MKL1* | 20 | 621 | 55 | MRTFA | myocardin related transcription factor A | Nucleus | transcription regulator | |
| chr10 | *P4HA1* | 23 | 808 | 55 | P4HA1 | prolyl 4-hydroxylase subunit | Cytoplasm | enzyme | |
| chr1 | *OSBPL9* | 23 | 569 | 55 | OSBPL9 | oxysterol binding protein like 9 | Cytoplasm | other | |
| chr9 | *GDA* | 21 | 562 | 55 | GDA | guanine deaminase | Cytoplasm | enzyme | |
| chr17 | *KIAA0195* | 24 | 981 | 55 | TMEM94 | transmembrane protein 94 | Extracellular Space | other | |
| chr16 | *DUS2* | 24 | 642 | 55 | DUS2 | dihydrouridine synthase 2 | Cytoplasm | enzyme | |
| chr15 | *GOLGA8A* | 23 | 988 | 55 | OLGA8A/GOLGA8 | golgin A8 family member A | Cytoplasm | other | |
| chr9 | *GNAQ* | 22 | 511 | 55 | GNAQ | G protein subunit alpha q | Plasma Membrane | enzyme | |
| chr2 | *HS6ST1* | 25 | 1519 | 55 | HS6ST1 | heparan sulfate 6-O-sulfotransferase 1 | Plasma Membrane | enzyme | |
| chr15 | *MORF4L1* | 23 | 750 | 55 | MORF4L1 | mortality factor 4 like 1 | Nucleus | other | |
| chr17 | *MRC2* | 25 | 1026 | 55 | MRC2 | mannose receptor C type 2 | Plasma Membrane | transmembrane receptor | |
| chr19 | *MUC16* | 24 | 603 | 55 | MUC16 | mucin 16, cell surface associated | Cytoplasm | other | REGN4018, oregovomab |
| chr1 | *PLEKHG5* | 27 | 1645 | 55 | PLEKHG5 | pleckstrin homology and RhoGEF domain containing G5 | Cytoplasm | other | |
| chr3 | *SCAP* | 23 | 718 | 55 | SCAP | SREBF chaperone | Cytoplasm | other | |
| chr7 | *STAG3L2* | 24 | 1072 | 55 | TAG3L2/STAG3L | stromal antigen 3-like 3 | Other | other | |
| chr11 | *SLC35F2* | 23 | 555 | 55 | SLC35F2 | solute carrier family 35 member | Other | other | |
| chr6 | *WDR27* | 21 | 731 | 55 | WDR27 | WD repeat domain 27 | Nucleus | other | |

| | | | | | | | | | |
|---|---|---|---|---|---|---|---|---|---|
| chr16 | *SLC12A3* | 23 | 992 | 55 | SLC12A3 | solute carrier family 12 member 3 | Plasma Membrane | transporter | hydrochlorothiazide/timolol, hydrochlorothiazide, hydrochlorothiazide/irbesartan, chlorothiazide/reserpine, hydrochlorothiazide/losartan, chlorothiazide, aliskiren/amlodipine/hydrochlorothiazide, chlorthalidone, enalapril/hydrochlorothiazide, fosinopril/hydrochlorothiazide, hydroflumethiazide/reserpine, candesartan cilexetil/hydrochlorothiazide, bendroflumethiazide/nadolol, [illegible] |
| chr7 | *RP4-724E13.2* | 23 | 1083 | 55 | | | | | |
| chr11 | *AGBL2* | 24 | 1102 | 55 | AGBL2 | ATP/GTP binding protein like 2 | Cytoplasm | enzyme | |
| chr3 | *LARS2* | 24 | 604 | 55 | LARS2 | leucyl-tRNA synthetase 2, mitochondrial | Cytoplasm | enzyme | |
| chr16 | *NFATC3* | 21 | 572 | 55 | NFATC3 | nuclear factor of activated T cells | Nucleus | transcription regulator | |
| chr7 | *ACTR3B* | 21 | 767 | 55 | ACTR3B | actin related protein 3B | Cytoplasm | other | |
| chr1 | *ABL2* | 24 | 756 | 55 | ABL2 | ABL proto-oncogene 2, non-receptor tyrosine kinase | Cytoplasm | kinase | nilotinib, dasatinib |
| chr20 | *SRC* | 26 | 1370 | 55 | SRC | SRC proto-oncogene, non-receptor tyrosine kinase | Cytoplasm | kinase | blinatumomab/dasatinib, KX-01, bosutinib/hydroxyurea/ponatinib, bosutinib, XL999, TPX-0046, docetaxel/nintedanib, dasatinib/nilotinib, dasatinib/erlotinib, AZD0424, AZM-475271, saracatinib, nintedanib, dasatinib/inotuzumab ozogamicin, repotrectinib |
| chr5 | *CTB-35F21.1* | 25 | 905 | 55 | | | | | |
| chr22 | *EFCAB6* | 19 | 715 | 55 | EFCAB6 | EF-hand calcium binding domain | Nucleus | other | |
| chr3 | *QRICH1* | 23 | 644 | 55 | QRICH1 | glutamine rich 1 | Nucleus | other | |
| chr17 | *CACNG4* | 24 | 904 | 55 | CACNG4 | calcium voltage-gated channel auxiliary subunit gamma 4 | Plasma Membrane | ion channel | |
| chr7 | *GRID2IP* | 25 | 1260 | 55 | GRID2IP | Grid2 interacting protein | Cytoplasm | other | |
| chr3 | *PLCL2* | 24 | 714 | 55 | PLCL2 | phospholipase C like 2 | Cytoplasm | enzyme | |
| chrX | *PCYT1B* | 23 | 516 | 55 | PCYT1B | phosphate cytidylyltransferase 1, choline, beta | Cytoplasm | enzyme | |
| chr17 | *ANKFN1* | 24 | 533 | 55 | ANKFN1 | ankyrin repeat and fibronectin type III domain containing 1 | Other | other | |
| chr16 | *NPRL3* | 23 | 856 | 55 | NPRL3 | NPR3 like, GATOR1 complex | Cytoplasm | other | |
| chr6 | *HIVEP1* | 25 | 634 | 55 | HIVEP1 | HIVEP zinc finger 1 | Nucleus | transcription regulator | |
| chr17 | *BRCA1* | 23 | 598 | 55 | BRCA1 | BRCA1 DNA repair associated | Nucleus | transcription regulator | |
| chr20 | *NTSR1* | 26 | 1868 | 55 | NTSR1 | neurotensin receptor 1 | Plasma Membrane | G-protein coupled receptor | contulakin-G, 177Lu-3BP-227, |
| chr12 | *CCDC91* | 23 | 556 | 55 | CCDC91 | coiled-coil domain containing 91 | Cytoplasm | other | |
| chr6 | *LYRM4* | 25 | 702 | 55 | LYRM4 | LYR motif containing 4 | Cytoplasm | other | |
| chr1 | *GABPB2* | 22 | 862 | 55 | GABPB2 | GA binding protein transcription factor subunit beta 2 | Nucleus | transcription regulator | |
| chr7 | *ISPD* | 20 | 703 | 54 | CRPPA | CDP-L-ribitol pyrophosphorylase | Cytoplasm | enzyme | |
| chr7 | *ABCB5* | 22 | 614 | 54 | ABCB5 | ATP binding cassette subfamily B member 5 | Plasma Membrane | transporter | |
| chr1 | *PLOD1* | 23 | 904 | 54 | PLOD1 | procollagen-lysine,2-oxoglutarate 5-dioxygenase 1 | Cytoplasm | enzyme | |
| chr19 | *SLC25A42* | 23 | 926 | 54 | SLC25A42 | solute carrier family 25 member | Cytoplasm | transporter | |
| chr3 | *CAND2* | 24 | 867 | 54 | CAND2 | cullin associated and neddylation dissociated 2 (putative) | Nucleus | transcription regulator | |
| chr1 | *USP48* | 24 | 578 | 54 | USP48 | ubiquitin specific peptidase 48 | Plasma Membrane | peptidase | |
| chr1 | *GPATCH2* | 19 | 608 | 54 | GPATCH2 | G-patch domain containing 2 | Nucleus | other | |
| chr4 | *SORBS2* | 20 | 539 | 54 | SORBS2 | sorbin and SH3 domain | Plasma Membrane | other | |
| chr11 | *PVRL1* | 23 | 1038 | 54 | NECTIN1 | nectin cell adhesion molecule 1 | Plasma Membrane | other | |
| chr20 | *RALY* | 25 | 872 | 54 | RALY | RALY heterogeneous nuclear ribonucleoprotein | Nucleus | transcription regulator | |
| chr15 | *CRTC3* | 24 | 1056 | 54 | CRTC3 | CREB regulated transcription coactivator 3 | Nucleus | other | |
| chr17 | *RP11-118B18.1* | 23 | 533 | 54 | | | | | |
| chr10 | *LDB3* | 24 | 928 | 54 | LDB3 | LIM domain binding 3 | Cytoplasm | transporter | |
| chr1 | *ZYG11B* | 20 | 541 | 54 | ZYG11B | zyg-11 family member B, cell cycle regulator | Other | other | |
| chr12 | *SSH1* | 23 | 894 | 54 | SSH1 | slingshot protein phosphatase 1 | Cytoplasm | phosphatase | |

| | | | | | | | | | |
|---|---|---|---|---|---|---|---|---|---|
| chr13 | *UGGT2* | 22 | 716 | 54 | UGGT2 | UDP-glucose glycoprotein glucosyltransferase 2 | Cytoplasm | enzyme | |
| chr7 | *MMD2* | 25 | 937 | 54 | MMD2 | monocyte to macrophage differentiation associated 2 | Other | kinase | |
| chr1 | *PUM1* | 22 | 617 | 54 | PUM1 | pumilio RNA binding family | Cytoplasm | other | |
| chr16 | *MAPK8IP3* | 24 | 1101 | 54 | MAPK8IP3 | mitogen-activated protein kinase 8 interacting protein 3 | Cytoplasm | other | |
| chr20 | *HNF4A* | 23 | 1399 | 54 | HNF4A | hepatocyte nuclear factor 4 alpha | Nucleus | transcription regulator | |
| chr5 | *CDC42SE2* | 19 | 595 | 54 | CDC42SE2 | CDC42 small effector 2 | Plasma Membrane | other | |
| chr20 | *SYNDIG1* | 20 | 669 | 54 | SYNDIG1 | synapse differentiation inducing 1 | Plasma Membrane | other | |
| chr16 | *SNTB2* | 20 | 566 | 54 | SNTB2 | syntrophin beta 2 | Plasma Membrane | other | |
| chr10 | *UBTD1* | 24 | 727 | 54 | UBTD1 | ubiquitin domain containing 1 | Other | other | |
| chr7 | *RP11-274B21.1* | 21 | 546 | 54 | | | | | |
| chr1 | *THRAP3* | 22 | 650 | 54 | THRAP3 | thyroid hormone receptor associated protein 3 | Nucleus | transcription regulator | |
| chr9 | *CCBL1* | 23 | 1070 | 54 | KYAT1 | kynurenine aminotransferase 1 | Cytoplasm | enzyme | |
| chr12 | *GOLGA3* | 24 | 1012 | 54 | GOLGA3 | golgin A3 | Cytoplasm | transporter | |
| chr5 | *CEP72* | 25 | 3679 | 54 | CEP72 | centrosomal protein 72 | Cytoplasm | other | |
| chr17 | *ACLY* | 23 | 747 | 54 | ACLY | ATP citrate lyase | Cytoplasm | enzyme | |
| chr14 | *SLC24A4* | 18 | 713 | 54 | SLC24A4 | solute carrier family 24 member 4 | Plasma Membrane | transporter | |
| chr11 | *CD82* | 23 | 1007 | 54 | CD82 | CD82 molecule | Plasma Membrane | other | |
| chr2 | *FAM179A* | 20 | 799 | 54 | TOGARAM2 | TOG array regulator of axonemal microtubules 2 | Other | other | |
| chr11 | *RP11-94P11.4* | 20 | 521 | 54 | | | | | |
| chr12 | *FAM186A* | 23 | 928 | 54 | FAM186A | family with sequence similarity 186 member A | Other | other | |
| chr21 | *HUNK* | 22 | 613 | 54 | HUNK | hormonally up-regulated Neu-associated kinase | Cytoplasm | kinase | |
| chr11 | *FLI1* | 25 | 908 | 54 | FLI1 | Fli-1 proto-oncogene, ETS transcription factor | Nucleus | transcription regulator | |
| chr3 | *RARB* | 26 | 577 | 54 | RARB | retinoic acid receptor beta | Nucleus | d-dependent nuclear rec | daunorubicin/tretinoin, alitretinoin, acitretin, AGN 190299, clindamycin phosphate/tretinoin, idarubicin/mitoxantrone/tretinoin, etretinate, fenretinide, idarubicin/tretinoin, tamibarotene, tretinoin, tazarotene, halobetasol propionate/tazarotene, adapalene, doxorubicin/tretinoin |
| chr13 | *TMCO3* | 23 | 1004 | 54 | TMCO3 | transmembrane and coiled-coil domains 3 | Other | transporter | |
| chr15 | *ATP8B4* | 23 | 524 | 54 | ATP8B4 | ATPase phospholipid transporting 8B4 (putative) | Plasma Membrane | transporter | |
| chr1 | *QSOX1* | 24 | 1429 | 54 | QSOX1 | quiescin sulfhydryl oxidase 1 | Cytoplasm | enzyme | |
| chr7 | *DNAJB6* | 24 | 905 | 54 | DNAJB6 | DnaJ heat shock protein family (Hsp40) member B6 | Nucleus | transcription regulator | |
| chr12 | *MED13L* | 22 | 473 | 54 | MED13L | mediator complex subunit 13L | Nucleus | other | |
| chr8 | *NDRG1* | 23 | 893 | 54 | NDRG1 | N-myc downstream regulated 1 | Nucleus | kinase | |
| chr2 | *TANC1* | 20 | 583 | 54 | TANC1 | tetratricopeptide repeat, ankyrin repeat and coiled-coil containing | Plasma Membrane | other | |
| chr21 | *ERG* | 20 | 618 | 54 | ERG | ETS transcription factor ERG | Nucleus | transcription regulator | |
| chr13 | *TFDP1* | 25 | 1334 | 54 | TFDP1 | transcription factor Dp-1 | Nucleus | transcription regulator | |
| chr12 | *HVCN1* | 23 | 736 | 53 | HVCN1 | hydrogen voltage gated channel | Plasma Membrane | ion channel | |
| chr7 | *CTB-111H14.1* | 24 | 552 | 53 | | | | | |
| chr1 | *TNFRSF1B* | 23 | 1052 | 53 | TNFRSF1B | TNF receptor superfamily | Plasma Membrane | transmembrane receptor | |
| chr8 | *CCDC26* | 23 | 524 | 53 | CCDC26 | CCDC26 long non-coding RNA | Other | other | |
| chr19 | *AKT2* | 22 | 1054 | 53 | AKT2 | AKT serine/threonine kinase 2 | Cytoplasm | kinase | triciribine, triciribine phosphate, enzastaurin, GSK690693, BAY1125976 |
| chr7 | *AMPH* | 24 | 651 | 53 | AMPH | amphiphysin | Plasma Membrane | other | |
| chrX | *PHEX* | 22 | 532 | 53 | PHEX | phosphate regulating endopeptidase homolog X-linked | Cytoplasm | peptidase | |
| chr9 | *RALGDS* | 26 | 1420 | 53 | RALGDS | ral guanine nucleotide dissociation stimulator | Cytoplasm | other | |
| chr6 | *SYTL3* | 21 | 622 | 53 | SYTL3 | synaptotagmin like 3 | Cytoplasm | other | |
| chr10 | *SGMS1* | 23 | 837 | 53 | SGMS1 | sphingomyelin synthase 1 | Cytoplasm | enzyme | |
| chr11 | *SLC1A2* | 26 | 716 | 53 | SLC1A2 | solute carrier family 1 member 2 | Plasma Membrane | transporter | riluzole |
| chr15 | *RNF111* | 19 | 557 | 53 | RNF111 | ring finger protein 111 | Nucleus | enzyme | |
| chr6 | *USP49* | 22 | 570 | 53 | USP49 | ubiquitin specific peptidase 49 | Nucleus | peptidase | |
| chr8 | *FAM135B* | 25 | 516 | 53 | FAM135B | family with sequence similarity 135 member B | Other | enzyme | |
| chr13 | *TUBGCP3* | 20 | 704 | 53 | TUBGCP3 | tubulin gamma complex associated protein 3 | Cytoplasm | other | |
| chr20 | *FER1L4* | 24 | 1111 | 53 | FER1L4 | fer-1 like family member 4 (pseudogene) | Other | other | |
| chr10 | *NT5C2* | 21 | 548 | 53 | NT5C2 | 5'-nucleotidase, cytosolic II | Cytoplasm | phosphatase | |
| chr3 | *IFT122* | 23 | 966 | 53 | IFT122 | intraflagellar transport 122 | Nucleus | other | |

| | | | | | | | | | |
|---|---|---|---|---|---|---|---|---|---|
| chr7 | *DDC* | 23 | 1152 | 53 | DDC | dopa decarboxylase | Cytoplasm | enzyme | S(-)-carbidopa, carbidopa/levodopa, DOPA decarboxylase inhibitor, levodopa, levodopa/pramipexole, carbidopa/entacapone/levodopa, carbidopa/levodopa/t |
| chr6 | *TULP4* | 19 | 545 | 53 | TULP4 | TUB like protein 4 | Cytoplasm | transcription regulator | |
| chr15 | *RP11-654A16.3* | 24 | 1011 | 53 | | | | | |
| chr3 | *VPRBP* | 23 | 588 | 53 | DCAF1 | DDB1 and CUL4 associated | Nucleus | kinase | |
| chr15 | *RP11-279F6.1* | 23 | 690 | 53 | | | | | |
| chr16 | *ATF7IP2* | 22 | 555 | 53 | ATF7IP2 | activating transcription factor 7 interacting protein 2 | Nucleus | other | |
| chr1 | *COL16A1* | 22 | 1141 | 53 | COL16A1 | collagen type XVI alpha 1 chain | Extracellular Space | other | collagenase clostridium |
| chr1 | *LMNA* | 24 | 1296 | 53 | LMNA | lamin A/C | Nucleus | other | |
| chr1 | *PRDM2* | 22 | 691 | 53 | PRDM2 | PR/SET domain 2 | Nucleus | transcription regulator | |
| chr20 | *PMEPA1* | 25 | 1754 | 53 | PMEPA1 | prostate transmembrane protein, androgen induced 1 | Plasma Membrane | other | |
| chr12 | *PCED1B* | 22 | 808 | 53 | PCED1B | PC-esterase domain containing | Other | other | |
| chr16 | *EEF2K* | 23 | 740 | 53 | EEF2K | eukaryotic elongation factor 2 | Cytoplasm | kinase | |
| chr14 | *PPP1R13B* | 19 | 561 | 53 | PPP1R13B | protein phosphatase 1 regulatory subunit 13B | Cytoplasm | phosphatase | |
| chr6 | *C6orf106* | 22 | 539 | 53 | ILRUN | inflammation and lipid regulator with UBA-like and NBR1-like | Other | other | |
| chr11 | *ABCC8* | 23 | 815 | 53 | ABCC8 | ATP binding cassette subfamily C member 8 | Plasma Membrane | transporter | nateglinide, chlorpropamide, acetohexamide, mitiglinide, tolazamide, gliquidone, gliclazide, repaglinide, glyburide, glipizide, |
| chr9 | *USP20* | 23 | 1103 | 53 | USP20 | ubiquitin specific peptidase 20 | Cytoplasm | peptidase | |
| chr6 | *FAM65B* | 23 | 607 | 53 | RIPOR2 | RHO family interacting cell polarization regulator 2 | Cytoplasm | other | |
| chr12 | *TESC* | 22 | 895 | 53 | TESC | tescalcin | Cytoplasm | other | |
| chr3 | *TRANK1* | 23 | 550 | 53 | TRANK1 | tetratricopeptide repeat and ankyrin repeat containing 1 | Nucleus | transcription regulator | |
| chr2 | *BIN1* | 27 | 1898 | 53 | BIN1 | bridging integrator 1 | Nucleus | other | |
| chr3 | *RP11-85M11.2* | 22 | 604 | 53 | | | | | |
| chr7 | *AGAP3* | 25 | 1216 | 53 | AGAP3 | ArfGAP with GTPase domain, ankyrin repeat and PH domain 3 | Nucleus | transcription regulator | |
| chr17 | *STAT3* | 22 | 580 | 53 | STAT3 | signal transducer and activator of transcription 3 | Nucleus | transcription regulator | danvatirsen, OPB-31121, OPB-51602, TTI-101, golotimod, STAT3 inhibitor |
| chr3 | *DGKG* | 20 | 541 | 53 | DGKG | diacylglycerol kinase gamma | Cytoplasm | kinase | |
| chr2 | *AC079807.4* | 23 | 680 | 53 | | | | | |
| chr9 | *FRMD3* | 23 | 536 | 53 | FRMD3 | FERM domain containing 3 | Other | other | |
| chr2 | *TRAPPC12* | 20 | 865 | 53 | TRAPPC12 | trafficking protein particle | Nucleus | other | |
| chr10 | *LCOR* | 19 | 501 | 53 | LCOR | ligand dependent nuclear receptor corepressor | Nucleus | transcription regulator | |
| chr11 | *GRM5* | 25 | 1044 | 53 | GRM5 | glutamate metabotropic receptor 5 | Plasma Membrane | G-protein coupled recepto | acamprosate, fasoracetam |
| chr4 | *CCSER1* | 18 | 527 | 53 | CCSER1 | coiled-coil serine rich protein 1 | Other | other | |
| chr17 | *RARA* | 22 | 1160 | 53 | RARA | retinoic acid receptor alpha | Nucleus | d-dependent nuclear rec | daunorubicin/tretinoin, alitretinoin, acitretin, AGN 190299, arsenic trioxide, NRX-195183, clindamycin phosphate/tretinoin, idarubicin/mitoxantrone/tretinoin, etretinate, idarubicin/tretinoin, tamibarotene, tretinoin, tazarotene, halobetasol propionate/tazarotene, adapalene, doxorubicin/tretinoin |
| chr1 | *AGO3* | 20 | 459 | 53 | AGO3 | argonaute RISC catalytic component 3 | Cytoplasm | translation regulator | |
| chr3 | *RHOA* | 22 | 681 | 53 | RHOA | ras homolog family member A | Cytoplasm | enzyme | |
| chr6 | *MTHFD1L* | 20 | 522 | 53 | MTHFD1L | methylenetetrahydrofolate dehydrogenase (NADP+ dependent) 1 like | Cytoplasm | enzyme | |
| chr16 | *RP11-347C12.2* | 22 | 689 | 53 | | | | | |
| chr10 | *CELF2* | 19 | 625 | 53 | CELF2 | CUGBP Elav-like family member | Nucleus | other | |
| chr1 | *PIK3C2B* | 24 | 1093 | 53 | PIK3C2B | phosphatidylinositol-4-phosphate 3-kinase catalytic subunit type 2 | Cytoplasm | kinase | |
| chr1 | *CNST* | 20 | 606 | 53 | CNST | consortin, connexin sorting | Cytoplasm | other | |
| chr3 | *PBRM1* | 19 | 554 | 53 | PBRM1 | polybromo 1 | Nucleus | other | |
| chr7 | *SLC37A3* | 23 | 696 | 53 | SLC37A3 | solute carrier family 37 member 3 | Cytoplasm | transporter | |
| chr8 | *LINC00536* | 23 | 550 | 52 | LINC00536 | long intergenic non-protein coding RNA 536 | Other | other | |

| | | | | | | | | | |
|---|---|---|---|---|---|---|---|---|---|
| chr21 | *PRDM15* | 22 | 1265 | 52 | PRDM15 | PR/SET domain 15 | Nucleus | transcription regulator | |
| chr9 | *SHC3* | 23 | 613 | 52 | SHC3 | SHC adaptor protein 3 | Cytoplasm | kinase | |
| chr2 | *CNNM4* | 22 | 1072 | 52 | CNNM4 | cyclin and CBS domain divalent metal cation transport mediator 4 | Plasma Membrane | transporter | |
| chr17 | *RHOT1* | 22 | 621 | 52 | RHOT1 | ras homolog family member T1 | Cytoplasm | enzyme | |
| chr18 | *RP11-172F10.1* | 23 | 826 | 52 | | | | | |
| chr11 | *CHKA* | 22 | 659 | 52 | CHKA | choline kinase alpha | Cytoplasm | kinase | TCD-717 |
| chr8 | *STK3* | 19 | 547 | 52 | STK3 | serine/threonine kinase 3 | Cytoplasm | kinase | |
| chr12 | *VSIG10* | 23 | 612 | 52 | VSIG10 | V-set and immunoglobulin domain containing 10 | Other | other | |
| chr10 | *CAMK2G* | 23 | 1455 | 52 | CAMK2G | calcium/calmodulin dependent protein kinase II gamma | Cytoplasm | kinase | |
| chr3 | *USP4* | 23 | 621 | 52 | USP4 | ubiquitin specific peptidase 4 | Nucleus | peptidase | |
| chr20 | *DIDO1* | 24 | 1475 | 52 | DIDO1 | death inducer-obliterator 1 | Nucleus | other | |
| chr19 | *CTC-548K16.2* | 22 | 731 | 52 | | | | | |
| chr15 | *LRRC49* | 21 | 576 | 52 | LRRC49 | leucine rich repeat containing 49 | Other | other | |
| chr2 | *EFR3B* | 19 | 646 | 52 | EFR3B | EFR3 homolog B | Plasma Membrane | other | |
| chr11 | *SERGEF* | 25 | 607 | 52 | SERGEF | secretion regulating guanine nucleotide exchange factor | Nucleus | other | |
| chr18 | *SPIRE1* | 19 | 463 | 52 | SPIRE1 | spire type actin nucleation factor | Cytoplasm | other | |
| chr8 | *MFHAS1* | 23 | 601 | 52 | MFHAS1 | malignant fibrous histiocytoma amplified sequence 1 | Cytoplasm | other | |
| chr14 | *BCL11B* | 19 | 781 | 52 | BCL11B | BAF chromatin remodeling complex subunit BCL11B | Nucleus | transcription regulator | |
| chr16 | *GPR56* | 22 | 1217 | 52 | ADGRG1 | adhesion G protein-coupled receptor G1 | Plasma Membrane | G-protein coupled recepto | |
| chr10 | *CHAT* | 24 | 1480 | 52 | CHAT | choline O-acetyltransferase | Nucleus | enzyme | |
| chr1 | *KCNC4* | 24 | 1065 | 52 | KCNC4 | potassium voltage-gated channel subfamily C member 4 | Plasma Membrane | ion channel | |
| chr10 | *WBP1L* | 21 | 660 | 52 | WBP1L | WW domain binding protein 1 like | Other | other | |
| chr7 | *GATSL1* | 20 | 828 | 52 | CASTOR2 | cytosolic arginine sensor for mTORC1 subunit 2 | Cytoplasm | other | |
| chr1 | *VAV3* | 22 | 638 | 52 | VAV3 | vav guanine nucleotide exchange factor 3 | Extracellular Space | cytokine | |
| chr1 | *LRP8* | 22 | 909 | 52 | LRP8 | LDL receptor related protein 8 | Plasma Membrane | transmembrane receptor | |
| chr2 | *TGFA* | 22 | 682 | 52 | TGFA | transforming growth factor alpha | Extracellular Space | growth factor | |
| chr22 | *MED15* | 21 | 755 | 52 | MED15 | mediator complex subunit 15 | Nucleus | transcription regulator | |
| chr5 | *ATP6V0E1* | 22 | 1183 | 52 | ATP6V0E1 | ATPase H+ transporting V0 subunit e1 | Cytoplasm | transporter | |
| chr5 | *ADAMTS16* | 25 | 2575 | 52 | ADAMTS16 | ADAM metallopeptidase with thrombospondin type 1 motif 16 | Extracellular Space | other | |
| chr5 | *LPCAT1* | 28 | 3118 | 52 | LPCAT1 | lysophosphatidylcholine acyltransferase 1 | Cytoplasm | enzyme | |
| chr3 | *RP11-649A16.1* | 24 | 544 | 52 | | | | | |
| chr10 | *VSTM4* | 24 | 1056 | 52 | VSTM4 | V-set and transmembrane domain containing 4 | Other | other | |
| chr14 | *NRXN3* | 20 | 566 | 52 | NRXN3 | neurexin 3 | Plasma Membrane | transporter | |
| chr5 | *FER* | 20 | 462 | 52 | FER | FER tyrosine kinase | Cytoplasm | kinase | lorlatinib |
| chr1 | *RGL1* | 24 | 753 | 52 | RGL1 | ral guanine nucleotide dissociation stimulator like 1 | Cytoplasm | other | |
| chr17 | *TEX2* | 21 | 617 | 52 | TEX2 | testis expressed 2 | Other | other | |
| chr12 | *NELL2* | 22 | 819 | 52 | NELL2 | neural EGFL like 2 | Extracellular Space | other | |
| chr10 | *RRP12* | 22 | 935 | 52 | RRP12 | ribosomal RNA processing 12 homolog | Nucleus | other | |
| chr17 | *PGS1* | 22 | 888 | 52 | PGS1 | phosphatidylglycerophosphate synthase 1 | Cytoplasm | enzyme | |
| chr20 | *SLC13A3* | 24 | 857 | 52 | SLC13A3 | solute carrier family 13 member 3 | Plasma Membrane | transporter | |
| chr5 | *ZFR* | 21 | 1428 | 52 | ZFR | zinc finger RNA binding protein | Nucleus | other | |
| chr18 | *LAMA3* | 19 | 563 | 52 | LAMA3 | laminin subunit alpha 3 | Extracellular Space | other | |
| chr15 | *MAP2K1* | 22 | 542 | 52 | MAP2K1 | mitogen-activated protein kinase kinase 1 | Cytoplasm | kinase | pimasertib, ARRY-424704, dabrafenib/panitumumab/trametinib, dabrafenib/trametinib, trametinib/vemurafenib, dabrafenib/trametinib/vemurafenib, AS703988, TAK 733, binimetinib/cetuximab/encorafenib, binimetinib/encorafenib/panitumumab, MKK1 inhibitor, refametinib, E 6201, PD184352, cetuximab/dabrafenib/trametinib, binimetinib, dabrafenib/ipilimumab/trametinib, trametinib |
| chr12 | *KNTC1* | 20 | 517 | 52 | KNTC1 | kinetochore associated 1 | Nucleus | other | |
| chr7 | *CCM2* | 21 | 812 | 52 | CCM2 | CCM2 scaffold protein | Cytoplasm | other | |
| chr3 | *FANCD2* | 20 | 512 | 52 | FANCD2 | FA complementation group D2 | Nucleus | other | |
| chr2 | *UGGT1* | 22 | 660 | 52 | UGGT1 | UDP-glucose glycoprotein glucosyltransferase 1 | Cytoplasm | enzyme | |

| | | | | | | | | | |
|---|---|---|---|---|---|---|---|---|---|
| chr4 | *LRBA* | 19 | 485 | 52 | LRBA | LPS responsive beige-like anchor protein | Cytoplasm | other | |
| chr15 | *EHD4* | 21 | 697 | 52 | EHD4 | EH domain containing 4 | Plasma Membrane | enzyme | |
| chr3 | *C3orf20* | 22 | 695 | 52 | C3orf20 | chromosome 3 open reading frame 20 | Cytoplasm | other | |
| chr18 | *RAB31* | 20 | 553 | 52 | RAB31 | RAB31, member RAS oncogene family | Cytoplasm | enzyme | |
| chr17 | *UNK* | 21 | 853 | 52 | UNK | unk zinc finger | Nucleus | transporter | |
| chr2 | *RAB11FIP5* | 23 | 787 | 51 | RAB11FIP5 | RAB11 family interacting protein | Cytoplasm | other | |
| chr1 | *NEGR1* | 20 | 535 | 51 | NEGR1 | neuronal growth regulator 1 | Plasma Membrane | other | |
| chr5 | *NDST1* | 19 | 729 | 51 | NDST1 | N-deacetylase and N-sulfotransferase 1 | Cytoplasm | enzyme | |
| chr5 | *RP11-541P9.3* | 23 | 624 | 51 | | | | | |
| chr17 | *MINK1* | 22 | 865 | 51 | MINK1 | misshapen like kinase 1 | Cytoplasm | kinase | |
| chr1 | *PLCH2* | 27 | 1122 | 51 | PLCH2 | phospholipase C eta 2 | Cytoplasm | enzyme | |
| chr11 | *GRAMD1B* | 23 | 791 | 51 | GRAMD1B | GRAM domain containing 1B | Other | other | |
| chr1 | *ESRRG* | 25 | 599 | 51 | ESRRG | estrogen related receptor gamma | Nucleus | d-dependent nuclear rec | diethylstilbestrol |
| chr6 | *GLTSCR1L* | 22 | 532 | 51 | BICRAL | BRD4 interacting chromatin remodeling complex associated protein like | Nucleus | other | |
| chr16 | *CACNG3* | 23 | 556 | 51 | CACNG3 | calcium voltage-gated channel auxiliary subunit gamma 3 | Plasma Membrane | ion channel | |
| chr9 | *ZNF618* | 20 | 812 | 51 | ZNF618 | zinc finger protein 618 | Other | other | |
| chr5 | *MAML1* | 24 | 989 | 51 | MAML1 | mastermind like transcriptional coactivator 1 | Nucleus | transcription regulator | |
| chr8 | *TOP1MT* | 23 | 1047 | 51 | TOP1MT | DNA topoisomerase I mitochondrial | Cytoplasm | enzyme | capecitabine/cetuximab/irinotecan, irinotecan/leucovorin, cetuximab/irinotecan, gemcitabine/irinotecan, aflibercept/irinotecan, capecitabine/irinotecan, irinotecan, cetuximab/irinotecan/ |
| chr12 | *CPNE8* | 21 | 774 | 51 | CPNE8 | copine 8 | Cytoplasm | other | |
| chr17 | *STAT5B* | 21 | 568 | 51 | STAT5B | signal transducer and activator of transcription 5B | Nucleus | transcription regulator | |
| chr16 | *PIEZO1* | 25 | 1368 | 51 | PIEZO1 | piezo type mechanosensitive ion channel component 1 | Cytoplasm | ion channel | |
| chr17 | *TANC2* | 19 | 519 | 51 | TANC2 | tetratricopeptide repeat, ankyrin repeat and coiled-coil containing | Nucleus | transcription regulator | |
| chr1 | *SZT2* | 22 | 740 | 51 | SZT2 | SZT2 subunit of KICSTOR | Cytoplasm | other | |
| chr3 | *CADPS* | 24 | 482 | 51 | CADPS | calcium dependent secretion | Plasma Membrane | other | |
| chr20 | *SAMHD1* | 22 | 654 | 51 | SAMHD1 | SAM and HD domain containing deoxynucleoside triphosphate triphosphohydrolase 1 | Nucleus | enzyme | |
| chr11 | *AP000797.3* | 22 | 574 | 51 | | | | | |
| chr3 | *CLDN11* | 23 | 957 | 51 | CLDN11 | claudin 11 | Plasma Membrane | other | |
| chr1 | *SPATA17* | 20 | 556 | 51 | SPATA17 | spermatogenesis associated 17 | Other | other | |
| chr8 | *ZHX2* | 18 | 592 | 51 | ZHX2 | zinc fingers and homeoboxes 2 | Nucleus | transcription regulator | |
| chr1 | *RP5-965F6.2* | 25 | 734 | 51 | | | | | |
| chr1 | *SCP2* | 21 | 499 | 51 | SCP2 | sterol carrier protein 2 | Cytoplasm | transporter | |
| chr1 | *MORN1* | 25 | 1060 | 51 | MORN1 | MORN repeat containing 1 | Other | other | |
| chr5 | *DIAPH1* | 18 | 488 | 51 | DIAPH1 | diaphanous related formin 1 | Plasma Membrane | other | |
| chr22 | *SGSM1* | 20 | 543 | 51 | SGSM1 | small G protein signaling | Nucleus | other | |
| chr1 | *CLIC4* | 21 | 484 | 51 | CLIC4 | chloride intracellular channel 4 | Plasma Membrane | ion channel | |
| chr11 | *YAP1* | 20 | 992 | 51 | YAP1 | Yes associated protein 1 | Nucleus | transcription regulator | |
| chr5 | *FBXW11* | 19 | 611 | 51 | FBXW11 | F-box and WD repeat domain containing 11 | Cytoplasm | enzyme | |
| chr2 | *SAP130* | 22 | 698 | 51 | SAP130 | Sin3A associated protein 130 | Nucleus | transcription regulator | |
| chr21 | *UMODL1* | 19 | 838 | 51 | UMODL1 | uromodulin like 1 | Cytoplasm | other | |
| chr9 | *RABGAP1* | 19 | 551 | 51 | RABGAP1 | RAB GTPase activating protein 1 | Cytoplasm | other | |
| chr2 | *AC007092.1* | 23 | 442 | 51 | | | | | |
| chr6 | *RUNX2* | 23 | 551 | 51 | RUNX2 | RUNX family transcription factor | Nucleus | transcription regulator | |
| chr17 | *SPNS3* | 22 | 938 | 51 | SPNS3 | sphingolipid transporter 3 | Other | other | |
| chr12 | *RP11-444D3.1* | 22 | 547 | 51 | | | | | |
| chr1 | *EPS15* | 22 | 462 | 51 | EPS15 | epidermal growth factor receptor pathway substrate 15 | Cytoplasm | other | |
| chr18 | *PIEZO2* | 18 | 530 | 51 | PIEZO2 | piezo type mechanosensitive ion channel component 2 | Other | ion channel | |
| chr7 | *PHTF2* | 21 | 565 | 51 | PHTF2 | putative homeodomain transcription factor 2 | Cytoplasm | other | |
| chr20 | *C20orf203* | 22 | 1010 | 51 | C20orf203 | chromosome 20 open reading frame 203 | Other | other | |
| chr5 | *ARSB* | 22 | 599 | 51 | ARSB | arylsulfatase B | Cytoplasm | enzyme | |
| chr11 | *HSD17B12* | 21 | 545 | 51 | HSD17B12 | hydroxysteroid 17-beta dehydrogenase 12 | Cytoplasm | enzyme | |
| chr3 | *SENP7* | 20 | 701 | 51 | SENP7 | SUMO specific peptidase 7 | Nucleus | peptidase | |
| chr1 | *TBCE* | 21 | 793 | 51 | TBCE | tubulin folding cofactor E | Cytoplasm | other | |
| chr12 | *FBRSL1* | 26 | 970 | 51 | FBRSL1 | fibrosin like 1 | Nucleus | other | |
| chr5 | *MARCH3;* | 20 | 553 | 51 | | | | | |
| chr6 | *UBR2* | 22 | 498 | 51 | UBR2 | ubiquitin protein ligase E3 component n-recognin 2 | Nucleus | enzyme | |
| chr10 | *STOX1* | 22 | 867 | 51 | STOX1 | storkhead box 1 | Cytoplasm | other | |
| chr10 | *MYPN* | 22 | 762 | 51 | MYPN | myopalladin | Cytoplasm | other | |
| chr15 | *NR2F2-AS1* | 23 | 852 | 51 | NR2F2-AS1 | NR2F2 antisense RNA 1 | Other | other | |
| chr10 | *TSPAN14* | 21 | 871 | 51 | TSPAN14 | tetraspanin 14 | Plasma Membrane | other | |

| | | | | | | | | | |
|---|---|---|---|---|---|---|---|---|---|
| chr5 | *CT49* | 20 | 1163 | 51 | LINC01194 | long intergenic non-protein coding RNA 1194 | Other | other | |
| chr18 | *EMILIN2* | 23 | 1134 | 51 | EMILIN2 | elastin microfibril interfacer 2 | Extracellular Space | other | |
| chr16 | *ZFPM1* | 26 | 1332 | 51 | ZFPM1 | zinc finger protein, FOG family member 1 | Nucleus | transcription regulator | |
| chr19 | *MEGF8* | 24 | 994 | 51 | MEGF8 | multiple EGF like domains 8 | Extracellular Space | other | |
| chr15 | *TTC23* | 22 | 984 | 51 | TTC23 | tetratricopeptide repeat domain | Extracellular Space | other | |
| chr20 | *ZHX3* | 19 | 682 | 51 | ZHX3 | zinc fingers and homeoboxes 3 | Nucleus | transcription regulator | |
| chr10 | *IDE* | 22 | 499 | 50 | IDE | insulin degrading enzyme | Extracellular Space | peptidase | bacitracin |
| chr2 | *LINC00152* | 21 | 771 | 50 | CYTOR | cytoskeleton regulator RNA | Other | other | |
| chr6 | *C6orf132* | 21 | 874 | 50 | C6orf132 | chromosome 6 open reading frame 132 | Other | other | |
| chr11 | *NUP160* | 21 | 934 | 50 | NUP160 | nucleoporin 160 | Nucleus | other | |
| chr19 | *FCHO1* | 22 | 928 | 50 | FCHO1 | FCH and mu domain containing endocytic adaptor 1 | Plasma Membrane | other | |
| chr10 | *RET* | 23 | 1227 | 50 | RET | ret proto-oncogene | Plasma Membrane | kinase | imatinib/sirolimus, imatinib/sorafenib, sorafenib, selpercatinib, lenvatinib, cytarabine/filgrastim/fludarabine phosphate/imatinib, sorafenib/sulindac/sunitinib, imatinib/sunitinib, vandetanib, cytarabine/filgrastim/fludarabine phosphate/idarubicin/imatinib, bevacizumab/sorafenib, dexamethasone/lenalidomide/sorafenib, sunitinib, BOS172738, cytarabine/imatinib/m[illegible] |
| chr1 | *LIN9* | 19 | 544 | 50 | LIN9 | lin-9 DREAM MuvB core complex component | Nucleus | other | |
| chr13 | *ATP8A2* | 19 | 520 | 50 | ATP8A2 | ATPase phospholipid transporting 8A2 | Plasma Membrane | transporter | |
| chr9 | *DNM1* | 22 | 1240 | 50 | DNM1 | dynamin 1 | Cytoplasm | enzyme | |
| chr1 | *SLC25A33* | 21 | 551 | 50 | SLC25A33 | solute carrier family 25 member | Cytoplasm | transporter | |
| chr5 | *RUFY1* | 22 | 931 | 50 | RUFY1 | RUN and FYVE domain | Cytoplasm | other | |
| chr8 | *LOXL2* | 22 | 926 | 50 | LOXL2 | lysyl oxidase like 2 | Extracellular Space | enzyme | simtuzumab |
| chr6 | *RP1-167F1.2* | 19 | 489 | 50 | | | | | |
| chr5 | *AACSP1* | 21 | 992 | 50 | AACSP1 | acetoacetyl-CoA synthetase pseudogene 1 | Other | other | |
| chr2 | *AC096559.1* | 19 | 490 | 50 | | | | | |
| chr11 | *ARAP1* | 22 | 1449 | 50 | ARAP1 | ArfGAP with RhoGAP domain, ankyrin repeat and PH domain 1 | Cytoplasm | other | |
| chr5 | *SV2C* | 22 | 490 | 50 | SV2C | synaptic vesicle glycoprotein 2C | Plasma Membrane | other | |
| chr14 | *EVL* | 19 | 752 | 50 | EVL | Enah/Vasp-like | Plasma Membrane | other | |
| chr3 | *HEG1* | 23 | 794 | 50 | HEG1 | heart development protein with EGF like domains 1 | Plasma Membrane | other | |
| chr2 | *ALMS1* | 20 | 552 | 50 | ALMS1 | ALMS1 centrosome and basal body associated protein | Cytoplasm | other | |
| chr11 | *RDX* | 20 | 530 | 50 | RDX | radixin | Cytoplasm | other | |
| chr5 | *NIM1* | 22 | 1554 | 50 | | | | | |
| chr20 | *ASXL1* | 22 | 673 | 50 | ASXL1 | ASXL transcriptional regulator 1 | Nucleus | transcription regulator | |
| chr6 | *SLC35F1* | 24 | 455 | 50 | SLC35F1 | solute carrier family 35 member | Other | other | |
| chrX | *NHS* | 25 | 529 | 50 | NHS | NHS actin remodeling regulator | Nucleus | other | |
| chr5 | *RP11-122F24.1* | 25 | 1439 | 50 | | | | | |
| chr3 | *MTMR14* | 21 | 977 | 50 | MTMR14 | myotubularin related protein 14 | Cytoplasm | phosphatase | |
| chr6 | *F13A1* | 22 | 624 | 50 | F13A1 | coagulation factor XIII A chain | Extracellular Space | enzyme | |
| chr15 | *ACAN* | 23 | 969 | 50 | ACAN | aggrecan | Extracellular Space | other | |
| chr13 | *CUL4A* | 21 | 993 | 50 | CUL4A | cullin 4A | Nucleus | other | |
| chr5 | *ZDHHC11* | 23 | 2967 | 50 | ZDHHC11 | zinc finger DHHC-type containing | Cytoplasm | other | |
| chr2 | *FARP2* | 19 | 606 | 50 | FARP2 | FERM, ARH/RhoGEF and pleckstrin domain protein 2 | Cytoplasm | other | |
| chr10 | *RUFY2* | 21 | 726 | 50 | RUFY2 | RUN and FYVE domain | Nucleus | other | |
| chr7 | *AC006481.1* | 19 | 528 | 50 | | | | | |
| chr12 | *CCDC62* | 20 | 578 | 50 | CCDC62 | coiled-coil domain containing 62 | Nucleus | transcription regulator | |
| chr11 | *RP11-115J23.1* | 19 | 654 | 50 | | | | | |
| chr11 | *FNBP4* | 22 | 1070 | 50 | FNBP4 | formin binding protein 4 | Nucleus | other | |

| | | | | | | | | | |
|---|---|---|---|---|---|---|---|---|---|
| chr19 | *DNMT1* | 19 | 681 | 50 | DNMT1 | DNA methyltransferase 1 | Nucleus | enzyme | cladribine/cytarabine/decitabine, 5-azacytidine/sorafenib, 5-azacytidine/decitabine, 4'-thio-2'-deoxycytidine, 5-azacytidine/lenalidomide, 5-azacytidine/cytarabine/decitabine, decitabine, decitabine/imatinib, fluorocyclopentenylcytosine, 5- |
| chr5 | *PRELID2* | 23 | 561 | 50 | PRELID2 | PRELI domain containing 2 | Cytoplasm | other | |
| chr17 | *USP36* | 22 | 762 | 50 | USP36 | ubiquitin specific peptidase 36 | Nucleus | peptidase | |
| chr5 | *RICTOR* | 20 | 1465 | 50 | RICTOR | RPTOR independent companion of MTOR complex 2 | Cytoplasm | other | |
| chr1 | *ASTN1* | 24 | 645 | 50 | ASTN1 | astrotactin 1 | Other | other | |
| chr11 | *DGKZ* | 24 | 1453 | 50 | DGKZ | diacylglycerol kinase zeta | Cytoplasm | kinase | |
| chr17 | *SHPK* | 20 | 830 | 50 | SHPK | sedoheptulokinase | Cytoplasm | kinase | |
| chrX | *ARHGAP6* | 25 | 545 | 50 | ARHGAP6 | Rho GTPase activating protein 6 | Cytoplasm | other | |
| chr11 | *HIPK3* | 17 | 603 | 50 | HIPK3 | homeodomain interacting protein kinase 3 | Nucleus | kinase | |
| chr9 | *AGTPBP1* | 19 | 476 | 50 | AGTPBP1 | ATP/GTP binding protein 1 | Nucleus | peptidase | |
| chr1 | *PTCHD2* | 23 | 933 | 50 | DISP3 | dispatched RND transporter family member 3 | Cytoplasm | other | |
| chr3 | *IGSF11* | 21 | 770 | 50 | IGSF11 | immunoglobulin superfamily member 11 | Plasma Membrane | other | |
| chr2 | *MSH6* | 23 | 651 | 50 | MSH6 | mutS homolog 6 | Nucleus | enzyme | |
| chr9 | *ASTN2* | 20 | 496 | 50 | ASTN2 | astrotactin 2 | Cytoplasm | other | |
| chr15 | *HERC2P3* | 20 | 890 | 50 | HERC2P3 | hect domain and RLD 2 pseudogene 3 | Other | other | |
| chr12 | *WSCD2* | 23 | 579 | 50 | WSCD2 | WSC domain containing 2 | Other | other | |
| chr17 | *RNF157* | 21 | 594 | 49 | RNF157 | ring finger protein 157 | Cytoplasm | enzyme | |
| chr15 | *MYO5C* | 21 | 615 | 49 | MYO5C | myosin VC | Cytoplasm | other | |
| chr15 | *RP11-152L20.3* | 21 | 796 | 49 | | | | | |
| chr7 | *DYNC1I1* | 21 | 651 | 49 | DYNC1I1 | dynein cytoplasmic 1 intermediate chain 1 | Cytoplasm | other | |
| chr19 | *C3* | 21 | 855 | 49 | C3 | complement C3 | Extracellular Space | peptidase | IgG, APL-2 |
| chr17 | *PIK3R5* | 22 | 696 | 49 | PIK3R5 | phosphoinositide-3-kinase regulatory subunit 5 | Cytoplasm | kinase | |
| chr21 | *SLC37A1* | 18 | 829 | 49 | SLC37A1 | solute carrier family 37 member 1 | Plasma Membrane | transporter | |
| chrX | *SLC9A7* | 21 | 532 | 49 | SLC9A7 | solute carrier family 9 member A7 | Cytoplasm | transporter | |
| chr12 | *SYT1* | 20 | 484 | 49 | SYT1 | synaptotagmin 1 | Cytoplasm | transporter | |
| chr12 | *PRMT8* | 18 | 631 | 49 | PRMT8 | protein arginine | Nucleus | enzyme | |
| chr17 | *SUZ12P* | 21 | 716 | 49 | SUZ12P1 | SUZ12 pseudogene 1 | Other | other | |
| chr1 | *PADI1* | 21 | 996 | 49 | PADI1 | peptidyl arginine deiminase 1 | Cytoplasm | enzyme | |
| chr7 | *RELN* | 21 | 421 | 49 | RELN | reelin | Extracellular Space | peptidase | |
| chr15 | *CGNL1* | 23 | 562 | 49 | CGNL1 | cingulin like 1 | Plasma Membrane | other | |
| chr20 | *FAM182B* | 20 | 1631 | 49 | FAM182B | family with sequence similarity 182 member B | Other | other | |
| chr5 | *SERINC5* | 21 | 486 | 49 | SERINC5 | serine incorporator 5 | Plasma Membrane | transporter | |
| chr1 | *POU2F1* | 19 | 470 | 49 | POU2F1 | POU class 2 homeobox 1 | Nucleus | transcription regulator | |
| chr2 | *EPAS1* | 23 | 930 | 49 | EPAS1 | endothelial PAS domain protein 1 | Nucleus | transcription regulator | PT-2385, PT2977 |
| chr17 | *WNT9B* | 21 | 708 | 49 | WNT9B | Wnt family member 9B | Extracellular Space | other | |
| chr20 | *MYBL2* | 22 | 920 | 49 | MYBL2 | MYB proto-oncogene like 2 | Nucleus | transcription regulator | |
| chrX | *EFHC2* | 21 | 449 | 49 | EFHC2 | EF-hand domain containing 2 | Other | other | |
| chr5 | *ADAM19* | 22 | 764 | 49 | ADAM19 | ADAM metallopeptidase domain | Plasma Membrane | peptidase | |
| chr12 | *OSBPL8* | 20 | 489 | 49 | OSBPL8 | oxysterol binding protein like 8 | Plasma Membrane | transporter | |
| chr15 | *CHD2* | 19 | 862 | 49 | CHD2 | chromodomain helicase DNA binding protein 2 | Nucleus | enzyme | |
| chr22 | *PPARA* | 18 | 568 | 49 | PPARA | peroxisome proliferator activated receptor alpha | Nucleus | d-dependent nuclear rec | bezafibrate, pemafibrate, ezetimibe/fenofibrate, TPST-1120, fenofibrate/simvastatin, gemfibrozil, fenofibrate, aleglitazar, NS-220, fenofibric acid/rosuvastatin, tesaglitazar, choline fenofibrate, docosahexaenoic acid, choline fenofibrate/simvastatin, clofibrate, atorvastatin/choline |

| | | | | | | | | | |
|---|---|---|---|---|---|---|---|---|---|
| chr12 | *VDR* | 23 | 1358 | 49 | VDR | vitamin D receptor | Nucleus | transcription regulator | cholecalciferol, alendronate/cholecalciferol, calcitriol, seocalcitol, 2-(3-hydroxypropoxy)calcitriol, doxercalciferol, calcifediol, paricalcitol, alfacalcidol, ILX-23-7553, inecalcitol, ergocalciferol, calcium carbonate/cholecalciferol, 1,25-dihydroxyergocalciferol, calcipotriene, betamethasone |
| chrX | *KAL1* | 21 | 438 | 49 | ANOS1 | anosmin 1 | Extracellular Space | other | |
| chr4 | *SPATA5* | 18 | 489 | 49 | SPATA5 | spermatogenesis associated 5 | Cytoplasm | other | |
| chr11 | *CARS* | 22 | 1772 | 49 | CARS1 | cysteinyl-tRNA synthetase 1 | Cytoplasm | enzyme | |
| chrX | *RP1-154K9.2* | 24 | 507 | 49 | | | | | |
| chr19 | *ELAVL1* | 21 | 912 | 49 | ELAVL1 | ELAV like RNA binding protein 1 | Cytoplasm | other | |
| chr3 | *LINC00877* | 24 | 555 | 49 | LINC00877 | long intergenic non-protein coding RNA 877 | Other | other | |
| chr7 | *GNA12* | 21 | 951 | 49 | GNA12 | G protein subunit alpha 12 | Plasma Membrane | enzyme | |
| chr1 | *CD247* | 21 | 700 | 49 | CD247 | CD247 molecule | Plasma Membrane | transmembrane receptor | blinatumomab |
| chr17 | *RP11-445F12.1* | 22 | 854 | 49 | | | | | |
| chr3 | *MUC4* | 20 | 795 | 49 | MUC4 | mucin 4, cell surface associated | Plasma Membrane | other | |
| chr18 | *SOGA2* | 18 | 630 | 49 | MTCL1 | microtubule crosslinking factor 1 | Cytoplasm | other | |
| chr1 | *ZBTB8OS* | 21 | 605 | 49 | ZBTB8OS | zinc finger and BTB domain containing 8 opposite strand | Cytoplasm | other | |
| chr1 | *TAF12* | 21 | 557 | 49 | TAF12 | TATA-box binding protein associated factor 12 | Nucleus | transcription regulator | |
| chr1 | *DSTYK* | 21 | 636 | 49 | DSTYK | dual serine/threonine and tyrosine protein kinase | Cytoplasm | kinase | |
| chr14 | *PPP2R5C* | 17 | 562 | 49 | PPP2R5C | protein phosphatase 2 regulatory subunit B'gamma | Nucleus | other | |
| chr18 | *FHOD3* | 18 | 474 | 49 | FHOD3 | formin homology 2 domain containing 3 | Nucleus | other | |
| chr13 | *DIAPH3* | 19 | 433 | 49 | DIAPH3 | diaphanous related formin 3 | Cytoplasm | enzyme | |
| chr22 | *CACNG2* | 21 | 614 | 49 | CACNG2 | calcium voltage-gated channel auxiliary subunit gamma 2 | Plasma Membrane | ion channel | |
| chr16 | *PHKB* | 20 | 527 | 49 | PHKB | phosphorylase kinase regulatory subunit beta | Cytoplasm | kinase | |
| chr12 | *FMNL3* | 22 | 1025 | 49 | FMNL3 | formin like 3 | Cytoplasm | other | |
| chr8 | *FBXO16* | 20 | 545 | 49 | FBXO16 | F-box protein 16 | Other | other | |
| chr20 | *ZNF341* | 22 | 862 | 49 | ZNF341 | zinc finger protein 341 | Nucleus | transcription regulator | |
| chr1 | *CNR2* | 19 | 565 | 49 | CNR2 | cannabinoid receptor 2 | Plasma Membrane | G-protein coupled receptor | BAY 38-7271, delta-9-tetrahydrocannabinol |
| chr11 | *PAK1* | 18 | 565 | 49 | PAK1 | p21 (RAC1) activated kinase 1 | Cytoplasm | kinase | |
| chr5 | *GFPT2* | 22 | 1120 | 49 | GFPT2 | glutamine-fructose-6-phosphate transaminase 2 | Cytoplasm | enzyme | |
| chr7 | *WDR60* | 22 | 855 | 49 | WDR60 | WD repeat domain 60 | Extracellular Space | other | |
| chr16 | *AC009120.4* | 22 | 462 | 49 | | | | | |
| chr5 | *CCNJL* | 22 | 937 | 49 | CCNJL | cyclin J like | Other | other | |
| chr10 | *SFMBT2* | 18 | 559 | 48 | SFMBT2 | Scm like with four mbt domains 2 | Nucleus | other | |
| chr7 | *CDHR3* | 22 | 532 | 48 | CDHR3 | cadherin related family member 3 | Plasma Membrane | other | |
| chr7 | *CNOT4* | 18 | 527 | 48 | CNOT4 | CCR4-NOT transcription complex subunit 4 | Cytoplasm | enzyme | |
| chr12 | *STAB2* | 22 | 522 | 48 | STAB2 | stabilin 2 | Plasma Membrane | transmembrane receptor | |
| chr1 | *SIPA1L2* | 21 | 732 | 48 | SIPA1L2 | signal induced proliferation associated 1 like 2 | Other | other | |
| chr7 | *ADCYAP1R1* | 21 | 1095 | 48 | ADCYAP1R1 | ADCYAP receptor type I | Plasma Membrane | G-protein coupled receptor | |
| chr5 | *SLC6A3* | 23 | 2641 | 48 | SLC6A3 | solute carrier family 6 member 3 | Plasma Membrane | transporter | venlafaxine, chlorpheniramine/hydrocodone/pseudoephedrine, methamphetamine, dexbrompheniramine/pseudoephedrine, azatadine/pseudoephedrine, codeine/pseudoephedrine/triprolidine, benzphetamine, phentermine/topiramate, phentermine, naproxen/pseudoephedrine, fenfluramine, brompheniramine/dextromethorphan/pseudoephedrine, benztropine, loratadine/pseudoephedrine, chlorpheniramine/ibu[illegible] |

| | | | | | | | | | |
|---|---|---|---|---|---|---|---|---|---|
| chr16 | *MKL2* | 20 | 475 | 48 | MRTFB | myocardin related transcription factor B | Nucleus | transcription regulator | |
| chr1 | *AHCTF1* | 19 | 489 | 48 | AHCTF1 | AT-hook containing transcription factor 1 | Nucleus | transcription regulator | |
| chr22 | *POLR2F* | 20 | 697 | 48 | POLR2F | RNA polymerase II subunit F | Nucleus | enzyme | |
| chr12 | *ANO4* | 22 | 461 | 48 | ANO4 | anoctamin 4 | Plasma Membrane | ion channel | |
| chr12 | *PTPRR* | 21 | 518 | 48 | PTPRR | protein tyrosine phosphatase receptor type R | Plasma Membrane | phosphatase | |
| chr12 | *C12orf75* | 21 | 485 | 48 | C12orf75 | chromosome 12 open reading frame 75 | Other | other | |
| chr20 | *ACSS1* | 21 | 984 | 48 | ACSS1 | acyl-CoA synthetase short chain family member 1 | Cytoplasm | enzyme | |
| chr11 | *RSF1* | 19 | 576 | 48 | RSF1 | remodeling and spacing factor 1 | Nucleus | transcription regulator | |
| chr5 | *MTMR12* | 23 | 1540 | 48 | MTMR12 | myotubularin related protein 12 | Plasma Membrane | other | |
| chr5 | *SLC38A9* | 18 | 453 | 48 | SLC38A9 | solute carrier family 38 member 9 | Extracellular Space | transporter | |
| chr7 | *GATSL2* | 19 | 738 | 48 | CASTOR2 | cytosolic arginine sensor for mTORC1 subunit 2 | Cytoplasm | other | |
| chr1 | *FAM129A* | 21 | 682 | 48 | NIBAN1 | niban apoptosis regulator 1 | Cytoplasm | other | |
| chr1 | *ASAP3* | 21 | 813 | 48 | ASAP3 | ArfGAP with SH3 domain, ankyrin repeat and PH domain 3 | Plasma Membrane | other | |
| chr1 | *AK5* | 22 | 441 | 48 | AK5 | adenylate kinase 5 | Cytoplasm | kinase | |
| chr16 | *TMC7* | 21 | 557 | 48 | TMC7 | transmembrane channel like 7 | Other | other | |
| chr19 | *MAST3* | 21 | 830 | 48 | MAST3 | microtubule associated serine/threonine kinase 3 | Other | kinase | |
| chr17 | *RHBDL3* | 20 | 769 | 48 | RHBDL3 | rhomboid like 3 | Plasma Membrane | peptidase | |
| chr17 | *CTD-2535L24.2* | 22 | 526 | 48 | | | | | |
| chr1 | *SOX13* | 20 | 1187 | 48 | SOX13 | SRY-box transcription factor 13 | Nucleus | transcription regulator | |
| chr12 | *SVOP* | 22 | 567 | 48 | SVOP | SV2 related protein | Cytoplasm | transporter | |
| chr9 | *STRBP* | 18 | 574 | 48 | STRBP | spermatid perinuclear RNA binding protein | Cytoplasm | other | |
| chr1 | *ST7L* | 16 | 503 | 48 | ST7L | suppression of tumorigenicity 7 | Other | other | |
| chr1 | *OPRD1* | 21 | 720 | 48 | OPRD1 | opioid receptor delta 1 | Plasma Membrane | G-protein coupled recepto | dextromethorphan/morphine, aspirin/caffeine/propoxyphene, chlorpheniramine/hydrocodone/pseudoephedrine, fentanyl, etorphine, acetaminophen/hydrocodone, nalmefene, morphine, hydromorphone, propoxyphene, acetaminophen/propoxyphene, naltrexone, naltrexone/oxycodone, codeine/pseudoephedrine/triprolidine, buprenorphine/naloxone, codeine, naldemedine, [illegible] |
| chr19 | *PKN1* | 22 | 845 | 48 | PKN1 | protein kinase N1 | Cytoplasm | kinase | |
| chr12 | *AEBP2* | 18 | 457 | 48 | AEBP2 | AE binding protein 2 | Nucleus | transcription regulator | |
| chr9 | *ASS1* | 21 | 1454 | 48 | ASS1 | argininosuccinate synthase 1 | Cytoplasm | enzyme | |
| chr10 | *NEURL* | 20 | 850 | 48 | NEURL1 | neuralized E3 ubiquitin protein ligase 1 | Cytoplasm | enzyme | |
| chr3 | *RPN1* | 20 | 756 | 48 | RPN1 | ribophorin I | Cytoplasm | enzyme | |
| chr7 | *SVOPL* | 21 | 617 | 48 | SVOPL | SVOP like | Other | other | |
| chr14 | *LTBP2* | 19 | 766 | 48 | LTBP2 | latent transforming growth factor beta binding protein 2 | Extracellular Space | other | |
| chr1 | *CAPN2* | 22 | 824 | 48 | CAPN2 | calpain 2 | Cytoplasm | peptidase | |
| chr19 | *IL12RB1* | 20 | 740 | 48 | IL12RB1 | interleukin 12 receptor subunit | Plasma Membrane | transmembrane recepto | |
| chr7 | *IQCE* | 23 | 1341 | 48 | IQCE | IQ motif containing E | Cytoplasm | other | |
| chr12 | *APOLD1* | 21 | 522 | 48 | APOLD1 | apolipoprotein L domain | Other | other | |
| chr5 | *DAP* | 23 | 2226 | 48 | DAP | death associated protein | Cytoplasm | transcription regulator | |
| chr7 | *WIPI2* | 21 | 943 | 48 | WIPI2 | WD repeat domain, phosphoinositide interacting 2 | Cytoplasm | other | |
| chr17 | *PITPNA* | 21 | 960 | 48 | PITPNA | phosphatidylinositol transfer protein alpha | Cytoplasm | transporter | |
| chr9 | *PIP5K1B* | 21 | 438 | 48 | PIP5K1B | phosphatidylinositol-4-phosphate 5-kinase type 1 beta | Cytoplasm | kinase | |
| chr5 | *ERBB2IP* | 19 | 487 | 48 | ERBIN | erbb2 interacting protein | Cytoplasm | other | |
| chr2 | *MGAT4A* | 20 | 671 | 48 | MGAT4A | alpha-1,3-mannosyl-glycoprotein 4-beta-N-acetylglucosaminyltransferase A | Cytoplasm | enzyme | |
| chr12 | *PRICKLE1* | 21 | 717 | 48 | PRICKLE1 | prickle planar cell polarity protein | Nucleus | other | |
| chr17 | *NMT1* | 20 | 879 | 48 | NMT1 | N-myristoyltransferase 1 | Cytoplasm | enzyme | |
| chr17 | *TMEM132E* | 21 | 715 | 48 | TMEM132E | transmembrane protein 132E | Other | other | |
| chr12 | *BCL7A* | 20 | 874 | 48 | BCL7A | BAF chromatin remodeling complex subunit BCL7A | Nucleus | other | |
| chr6 | *PKHD1* | 20 | 474 | 48 | PKHD1 | PKHD1 ciliary IPT domain containing fibrocystin/polyductin | Plasma Membrane | other | |
| chr10 | *DNMBP* | 20 | 490 | 48 | DNMBP | dynamin binding protein | Cytoplasm | other | |
| chr8 | *MTUS1* | 19 | 495 | 48 | MTUS1 | microtubule associated scaffold protein 1 | Extracellular Space | other | |

| | | | | | | | | | |
|---|---|---|---|---|---|---|---|---|---|
| chr11 | *RP11-627G23.1* | 21 | 717 | 48 | | | | | |
| chr15 | *NEO1* | 20 | 441 | 48 | NEO1 | neogenin 1 | Plasma Membrane | transcription regulator | |
| chr1 | *DCAF6* | 16 | 415 | 48 | DCAF6 | DDB1 and CUL4 associated | Nucleus | transcription regulator | |
| chr11 | *IPO7* | 19 | 740 | 48 | IPO7 | importin 7 | Nucleus | transporter | |
| chr5 | *KDM3B* | 18 | 532 | 48 | KDM3B | lysine demethylase 3B | Nucleus | enzyme | |
| chr1 | *SORT1* | 21 | 686 | 48 | SORT1 | sortilin 1 | Plasma Membrane | ;-protein coupled recepto | |
| chr1 | *TGFB2* | 19 | 667 | 48 | TGFB2 | transforming growth factor beta 2 | Extracellular Space | growth factor | fresolimumab, dalantercept, trabedersen |
| chr20 | *TM9SF4* | 21 | 893 | 48 | TM9SF4 | transmembrane 9 superfamily member 4 | Cytoplasm | transporter | |
| chr11 | *NADSYN1* | 21 | 877 | 48 | NADSYN1 | NAD synthetase 1 | Cytoplasm | enzyme | |
| chr3 | *ALDH1L1* | 20 | 1141 | 48 | ALDH1L1 | aldehyde dehydrogenase 1 family member L1 | Cytoplasm | enzyme | |
| chr12 | *ST8SIA1* | 20 | 587 | 48 | ST8SIA1 | ST8 alpha-N-acetyl-neuraminide alpha-2,8-sialyltransferase 1 | Cytoplasm | enzyme | |
| chr7 | *RHBDD2* | 20 | 811 | 48 | RHBDD2 | rhomboid domain containing 2 | Plasma Membrane | other | |
| chrX | *ACOT9* | 19 | 614 | 47 | ACOT9 | acyl-CoA thioesterase 9 | Cytoplasm | enzyme | |
| chr11 | *TRIM29* | 21 | 845 | 47 | TRIM29 | tripartite motif containing 29 | Cytoplasm | transcription regulator | |
| chr11 | *FADS2* | 19 | 986 | 47 | FADS2 | fatty acid desaturase 2 | Plasma Membrane | enzyme | |
| chr7 | *FOXP2* | 19 | 459 | 47 | FOXP2 | forkhead box P2 | Nucleus | transcription regulator | |
| chr10 | *RASGEF1A* | 23 | 1324 | 47 | RASGEF1A | RasGEF domain family member | Cytoplasm | other | |
| chr19 | *EMR2* | 19 | 696 | 47 | ADGRE2 | adhesion G protein-coupled receptor E2 | Plasma Membrane | other | |
| chr1 | *RP5-884C9.2* | 22 | 736 | 47 | | | | | |
| chr2 | *SPRED2* | 20 | 529 | 47 | SPRED2 | sprouty related EVH1 domain containing 2 | Extracellular Space | cytokine | |
| chr20 | *PTPN1* | 20 | 709 | 47 | PTPN1 | protein tyrosine phosphatase non-receptor type 1 | Cytoplasm | phosphatase | protein tyrosine phosphatase 1B inhibitor, |
| chr3 | *PRKCI* | 21 | 851 | 47 | PRKCI | protein kinase C iota | Cytoplasm | kinase | ingenol mebutate |
| chr15 | *LRRC28* | 20 | 678 | 47 | LRRC28 | leucine rich repeat containing 28 | Other | other | |
| chr20 | *TRPC4AP* | 20 | 880 | 47 | TRPC4AP | transient receptor potential cation channel subfamily C member 4 associated protein | Cytoplasm | transporter | |
| chr9 | *RGS3* | 18 | 723 | 47 | RGS3 | regulator of G protein signaling 3 | Nucleus | enzyme | |
| chr3 | *RP11-260O18.1* | 19 | 528 | 47 | | | | | |
| chr18 | *MBP* | 18 | 719 | 47 | MBP | myelin basic protein | Extracellular Space | other | |
| chr11 | *ST3GAL4* | 21 | 870 | 47 | ST3GAL4 | ST3 beta-galactoside alpha-2,3-sialyltransferase 4 | Cytoplasm | enzyme | |
| chr14 | *ACTN1* | 18 | 690 | 47 | ACTN1 | actinin alpha 1 | Cytoplasm | transcription regulator | |
| chr16 | *TXNDC11* | 20 | 515 | 47 | TXNDC11 | thioredoxin domain containing 11 | Cytoplasm | enzyme | |
| chr15 | *TRIP4* | 17 | 443 | 47 | TRIP4 | thyroid hormone receptor | Nucleus | transcription regulator | |
| chr9 | *DFNB31* | 20 | 595 | 47 | WHRN | whirlin | Plasma Membrane | other | |
| chr9 | *FGD3* | 18 | 804 | 47 | FGD3 | FYVE, RhoGEF and PH domain containing 3 | Cytoplasm | other | |
| chr17 | *ATP2A3* | 22 | 987 | 47 | ATP2A3 | ATPase sarcoplasmic/endoplasmic | Cytoplasm | transporter | |
| chr16 | *SMG1* | 19 | 515 | 47 | SMG1 | SMG1 nonsense mediated mRNA decay associated PI3K related kinase | Cytoplasm | kinase | |
| chr12 | *TMTC1* | 21 | 679 | 47 | TMTC1 | transmembrane O-mannosyltransferase targeting cadherins 1 | Cytoplasm | enzyme | |
| chr11 | *CADM1* | 18 | 434 | 47 | CADM1 | cell adhesion molecule 1 | Plasma Membrane | other | |
| chr8 | *PIWIL2* | 20 | 469 | 47 | PIWIL2 | piwi like RNA-mediated gene silencing 2 | Cytoplasm | enzyme | |
| chr3 | *SLC41A3* | 20 | 1065 | 47 | SLC41A3 | solute carrier family 41 member 3 | Plasma Membrane | transporter | |
| chr10 | *PFKFB3* | 18 | 733 | 47 | PFKFB3 | 6-phosphofructo-2-kinase/fructose-2,6- | Cytoplasm | kinase | PFK-158 |
| chr7 | *RSPH10B* | 19 | 913 | 47 | SPH10B/RSPH10 | radial spoke head 10 homolog B | Extracellular Space | other | |
| chr9 | *NUP188* | 20 | 775 | 47 | NUP188 | nucleoporin 188 | Nucleus | other | |
| chr16 | *CIRH1A* | 19 | 513 | 47 | UTP4 | UTP4 small subunit processome component | Nucleus | other | |
| chr8 | *BNIP3L* | 21 | 557 | 47 | BNIP3L | BCL2 interacting protein 3 like | Cytoplasm | other | |
| chr11 | *GAB2* | 21 | 637 | 47 | GAB2 | GRB2 associated binding protein | Cytoplasm | other | |
| chr2 | *DNAH6* | 20 | 474 | 47 | DNAH6 | dynein axonemal heavy chain 6 | Extracellular Space | other | |
| chr5 | *ATG10* | 19 | 435 | 47 | ATG10 | autophagy related 10 | Plasma Membrane | enzyme | |
| chr7 | *BAZ1B* | 18 | 530 | 47 | BAZ1B | bromodomain adjacent to zinc finger domain 1B | Nucleus | transcription regulator | |
| chr15 | *RP11-739G5.1* | 19 | 652 | 47 | | | | | |
| chr17 | *FAM117A* | 20 | 537 | 47 | FAM117A | family with sequence similarity 117 member A | Other | transporter | |
| chr11 | *CBL* | 20 | 465 | 47 | CBL | Cbl proto-oncogene | Nucleus | transcription regulator | |
| chr17 | *ERN1* | 19 | 685 | 47 | ERN1 | endoplasmic reticulum to nucleus signaling 1 | Cytoplasm | kinase | |
| chrX | *PHKA2* | 19 | 677 | 47 | PHKA2 | phosphorylase kinase regulatory subunit alpha 2 | Cytoplasm | kinase | |
| chr8 | *MROH5* | 20 | 1269 | 47 | MROH5 | maestro heat like repeat family member 5 (gene/pseudogene) | Other | other | |
| chr8 | *RP11-1080G15.1* | 20 | 488 | 47 | | | | | |
| chr12 | *CACNA2D4* | 17 | 763 | 47 | CACNA2D4 | calcium voltage-gated channel auxiliary subunit alpha2delta 4 | Plasma Membrane | ion channel | pregabalin |
| chr12 | *NAA25* | 18 | 463 | 47 | NAA25 | N(alpha)-acetyltransferase 25, NatB auxiliary subunit | Cytoplasm | other | |
| chr11 | *SPTBN2* | 23 | 1373 | 47 | SPTBN2 | spectrin beta, non-erythrocytic 2 | Cytoplasm | other | |
| chr7 | *AMZ1* | 21 | 1244 | 47 | AMZ1 | archaelysin family | Other | peptidase | |

| | | | | | | | | | |
|---|---|---|---|---|---|---|---|---|---|
| chr20 | *HCK* | 20 | 735 | 47 | HCK | HCK proto-oncogene, Src family tyrosine kinase | Cytoplasm | kinase | bosutinib/hydroxyurea/ponatinib, bosutinib, rebastinib |
| chr1 | *DHRS3* | 20 | 768 | 47 | DHRS3 | dehydrogenase/reductase 3 | Cytoplasm | enzyme | |
| chr1 | *EYA3* | 17 | 475 | 47 | EYA3 | EYA transcriptional coactivator and phosphatase 3 | Nucleus | phosphatase | |
| chr11 | *SWAP70* | 18 | 580 | 47 | SWAP70 | switching B cell complex subunit SWAP70 | Cytoplasm | other | |
| chr1 | *IQGAP3* | 21 | 869 | 47 | IQGAP3 | IQ motif containing GTPase activating protein 3 | Plasma Membrane | other | |
| chr10 | *CCAR1* | 19 | 765 | 47 | CCAR1 | cell division cycle and apoptosis regulator 1 | Nucleus | transcription regulator | |
| chr5 | *PPARGC1B* | 16 | 604 | 47 | PPARGC1B | PPARG coactivator 1 beta | Nucleus | transcription regulator | |
| chr6 | *LY86-AS1* | 21 | 475 | 46 | LY86-AS1 | LY86 antisense RNA 1 | Other | other | |
| chr9 | *FAM102A* | 20 | 1294 | 46 | FAM102A | family with sequence similarity 102 member A | Other | other | |
| chr9 | *ZER1* | 21 | 951 | 46 | ZER1 | zyg-11 related cell cycle regulator | Other | other | |
| chr12 | *FBXW8* | 19 | 678 | 46 | FBXW8 | F-box and WD repeat domain containing 8 | Cytoplasm | other | |
| chr19 | *UHRF1* | 19 | 815 | 46 | UHRF1 | ubiquitin like with PHD and ring finger domains 1 | Nucleus | transcription regulator | |
| chr9 | *VPS13A* | 16 | 430 | 46 | VPS13A | vacuolar protein sorting 13 homolog A | Cytoplasm | transporter | |
| chr1 | *VPS45* | 19 | 543 | 46 | VPS45 | vacuolar protein sorting 45 | Cytoplasm | transporter | |
| chr19 | *NCAN* | 20 | 692 | 46 | NCAN | neurocan | Extracellular Space | other | |
| chr7 | *AGMO* | 20 | 544 | 46 | AGMO | alkylglycerol monooxygenase | Cytoplasm | enzyme | |
| chr19 | *RFX1* | 21 | 887 | 46 | RFX1 | regulatory factor X1 | Nucleus | transcription regulator | |
| chr1 | *LMOD1* | 21 | 808 | 46 | LMOD1 | leiomodin 1 | Cytoplasm | other | |
| chr7 | *LIMK1* | 21 | 835 | 46 | LIMK1 | LIM domain kinase 1 | Cytoplasm | kinase | dabrafenib/ipilimumab/trametinib, dabrafenib, dabrafenib/panitumumab/trametinib, dabrafenib/trametinib, dabrafenib/trametinib/vemurafenib, cetuximab/encorafenib, binimetinib/encorafenib, encorafenib, binimetinib/cetuximab/encorafenib, binimetinib/encorafenib/panitumumab, cetuximab/dabrafenib [illegible] |
| chr15 | *CHSY1* | 19 | 975 | 46 | CHSY1 | chondroitin sulfate synthase 1 | Cytoplasm | enzyme | |
| chr7 | *ZNF398* | 19 | 591 | 46 | ZNF398 | zinc finger protein 398 | Nucleus | transcription regulator | |
| chr9 | *GOLGA1* | 19 | 811 | 46 | GOLGA1 | golgin A1 | Cytoplasm | other | |
| chr3 | *CCDC12* | 20 | 788 | 46 | CCDC12 | coiled-coil domain containing 12 | Other | other | |
| chr2 | *DHX57* | 19 | 609 | 46 | DHX57 | DExH-box helicase 57 | Extracellular Space | other | |
| chr5 | *BDP1* | 18 | 483 | 46 | BDP1 | B double prime 1, subunit of RNA polymerase III transcription initiation factor IIIB | Nucleus | transcription regulator | |
| chr11 | *ALX4* | 21 | 903 | 46 | ALX4 | ALX homeobox 4 | Nucleus | transcription regulator | |
| chr6 | *AIG1* | 19 | 476 | 46 | AIG1 | androgen induced 1 | Other | other | |
| chr9 | *UBAP2* | 15 | 483 | 46 | UBAP2 | ubiquitin associated protein 2 | Cytoplasm | other | |
| chr16 | *SEC14L5* | 20 | 742 | 46 | SEC14L5 | SEC14 like lipid binding 5 | Other | other | |
| chr3 | *PPM1L* | 19 | 553 | 46 | PPM1L | protein phosphatase, Mg2+/Mn2+ dependent 1L | Cytoplasm | phosphatase | |
| chr8 | *BMP1* | 21 | 914 | 46 | BMP1 | bone morphogenetic protein 1 | Extracellular Space | peptidase | |
| chr15 | *NRG4* | 19 | 617 | 46 | NRG4 | neuregulin 4 | Extracellular Space | growth factor | |
| chr20 | *PKIG* | 20 | 733 | 46 | PKIG | cAMP-dependent protein kinase inhibitor gamma | Other | other | |
| chr5 | *CTD-2116N24.1* | 23 | 1381 | 46 | | | | | |
| chr10 | *RP11-525A16.4* | 20 | 878 | 46 | | | | | |
| chr1 | *DISP1* | 18 | 627 | 46 | DISP1 | dispatched RND transporter family member 1 | Plasma Membrane | transporter | |
| chr3 | *CMTM7* | 21 | 559 | 46 | CMTM7 | CKLF like MARVEL transmembrane domain | Extracellular Space | cytokine | |
| chr5 | *EBF1* | 21 | 675 | 46 | EBF1 | EBF transcription factor 1 | Nucleus | transcription regulator | |
| chr1 | *KCNK2* | 19 | 563 | 46 | KCNK2 | potassium two pore domain channel subfamily K member 2 | Plasma Membrane | ion channel | |
| chr10 | *FAM35A* | 18 | 495 | 46 | SHLD2 | shieldin complex subunit 2 | Extracellular Space | other | |
| chr2 | *ZNF638* | 18 | 485 | 46 | ZNF638 | zinc finger protein 638 | Nucleus | other | |
| chr7 | *AOAH* | 20 | 674 | 46 | AOAH | acyloxyacyl hydrolase | Extracellular Space | enzyme | |
| chr1 | *CCT3* | 19 | 654 | 46 | CCT3 | chaperonin containing TCP1 subunit 3 | Cytoplasm | other | |
| chr1 | *WDR47* | 18 | 562 | 46 | WDR47 | WD repeat domain 47 | Other | other | |
| chr1 | *KDM4A* | 21 | 526 | 46 | KDM4A | lysine demethylase 4A | Nucleus | transcription regulator | |
| chr14 | *CCDC85C* | 19 | 947 | 46 | CCDC85C | coiled-coil domain containing 85C | Plasma Membrane | other | |
| chr1 | *ACTL8* | 20 | 716 | 46 | ACTL8 | actin like 8 | Cytoplasm | other | |
| chr11 | *CLNS1A* | 20 | 591 | 46 | CLNS1A | chloride nucleotide-sensitive channel 1A | Plasma Membrane | ion channel | |
| chr17 | *SLC38A10* | 22 | 1028 | 46 | SLC38A10 | solute carrier family 38 member | Cytoplasm | other | |
| chr21 | *DSCAM* | 19 | 481 | 46 | DSCAM | DS cell adhesion molecule | Plasma Membrane | other | |
| chr4 | *PALLD* | 17 | 411 | 46 | PALLD | palladin, cytoskeletal associated protein | Plasma Membrane | other | |
| chr15 | *RP11-798K3.2* | 19 | 592 | 46 | | | | | |

| | | | | | | | | | |
|---|---|---|---|---|---|---|---|---|---|
| chr15 | *MAPKBP1* | 19 | 648 | 46 | MAPKBP1 | mitogen-activated protein kinase binding protein 1 | Cytoplasm | other | |
| chr17 | *TADA2A* | 18 | 471 | 46 | TADA2A | transcriptional adaptor 2A | Nucleus | transcription regulator | |
| chr21 | *TIAM1* | 18 | 477 | 46 | TIAM1 | T cell lymphoma invasion and metastasis 1 | Cytoplasm | other | |
| chr14 | *RCOR1* | 17 | 453 | 46 | RCOR1 | REST corepressor 1 | Nucleus | transcription regulator | |
| chr1 | *SNX27* | 18 | 512 | 46 | SNX27 | sorting nexin 27 | Cytoplasm | other | |
| chr15 | *HERC2P9* | 18 | 542 | 46 | HERC2P9 | hect domain and RLD 2 pseudogene 9 | Other | other | |
| chr16 | *SRCAP* | 19 | 651 | 46 | SRCAP | Snf2 related CREBBP activator protein | Cytoplasm | transcription regulator | |
| chr1 | *LEMD1* | 20 | 1131 | 46 | LEMD1 | LEM domain containing 1 | Other | other | |
| chr1 | *ARHGEF2* | 20 | 886 | 46 | ARHGEF2 | Rho/Rac guanine nucleotide exchange factor 2 | Cytoplasm | other | |
| chr19 | *ZNF675* | 20 | 538 | 46 | ZNF675 | zinc finger protein 675 | Nucleus | transcription regulator | |
| chr17 | *SLC47A1* | 21 | 598 | 46 | SLC47A1 | solute carrier family 47 member 1 | Plasma Membrane | transporter | |
| chr13 | *MYCBP2* | 17 | 441 | 46 | MYCBP2 | MYC binding protein 2 | Nucleus | enzyme | |
| chr20 | *KCNQ2* | 24 | 1613 | 46 | KCNQ2 | potassium voltage-gated channel subfamily Q member 2 | Plasma Membrane | ion channel | ezogabine |
| chr3 | *LRIG1* | 20 | 669 | 46 | LRIG1 | leucine rich repeats and immunoglobulin like domains 1 | Extracellular Space | other | |
| chr16 | *FBXO31* | 20 | 966 | 46 | FBXO31 | F-box protein 31 | Extracellular Space | enzyme | |
| chr12 | *KRT8* | 21 | 1128 | 46 | KRT8 | keratin 8 | Cytoplasm | other | |
| chr15 | *HOMER2* | 20 | 829 | 46 | HOMER2 | homer scaffold protein 2 | Plasma Membrane | other | |
| chr2 | *IQCA1* | 16 | 645 | 46 | IQCA1 | IQ motif containing with AAA domain 1 | Other | other | |
| chr17 | *PYY* | 21 | 613 | 46 | PYY | peptide YY | Extracellular Space | other | |
| chr21 | *CHODL* | 19 | 471 | 46 | CHODL | chondrolectin | Plasma Membrane | other | |
| chr1 | *TTC39A* | 19 | 797 | 46 | TTC39A | tetratricopeptide repeat domain | Other | other | |
| chr3 | *CCDC36* | 20 | 564 | 46 | CCDC36 | coiled-coil domain containing 36 | Nucleus | other | |
| chr1 | *ARHGEF11* | 20 | 830 | 45 | ARHGEF11 | Rho guanine nucleotide exchange factor 11 | Cytoplasm | other | |
| chr16 | *C16orf96* | 19 | 763 | 45 | C16orf96 | chromosome 16 open reading frame 96 | Other | other | |
| chr1 | *GREM2* | 18 | 545 | 45 | GREM2 | gremlin 2, DAN family BMP antagonist | Extracellular Space | other | |
| chr17 | *MAP2K6* | 17 | 439 | 45 | MAP2K6 | mitogen-activated protein kinase kinase 6 | Cytoplasm | kinase | |
| chr20 | *HM13* | 21 | 1046 | 45 | HM13 | histocompatibility minor 13 | Cytoplasm | peptidase | |
| chr19 | *OPA3* | 18 | 594 | 45 | OPA3 | outer mitochondrial membrane lipid metabolism regulator OPA3 | Cytoplasm | other | |
| chr20 | *STK4* | 15 | 521 | 45 | STK4 | serine/threonine kinase 4 | Cytoplasm | kinase | |
| chr17 | *COX10-AS1* | 21 | 385 | 45 | COX10-AS1 | COX10 antisense RNA 1 | Other | other | |
| chr3 | *RAF1* | 19 | 554 | 45 | RAF1 | Raf-1 proto-oncogene, serine/threonine kinase | Cytoplasm | kinase | CHIR-265, irinotecan/panitumumab/vemurafenib, sorafenib, dabrafenib/trametinib, sorafenib/sulindac/sunitinib, vemurafenib, ipilimumab/vemurafenib, binimetinib/encorafenib/panitumumab, bevacizumab/sorafenib, dexamethasone/lenalidomide/sorafenib, regorafenib, 5-azacytidine/sorafenib, dabrafenib/ipilimumab/trametinib, belvarafenib, c-raf antisense [illegible] |
| chr15 | *NIPA1* | 19 | 777 | 45 | NIPA1 | NIPA magnesium transporter 1 | Plasma Membrane | transporter | |
| chr11 | *PICALM* | 17 | 716 | 45 | PICALM | phosphatidylinositol binding clathrin assembly protein | Cytoplasm | other | |
| chr1 | *PAPPA2* | 20 | 604 | 45 | PAPPA2 | pappalysin 2 | Extracellular Space | peptidase | |
| chr1 | *RP11-154H17.1* | 21 | 832 | 45 | | | | | |
| chr3 | *FBXL2* | 18 | 439 | 45 | FBXL2 | F-box and leucine rich repeat protein 2 | Cytoplasm | enzyme | |
| chr2 | *RFX8* | 19 | 782 | 45 | RFX8 | RFX family member 8, lacking RFX DNA binding domain | Other | other | |
| chr3 | *RAB7A* | 19 | 671 | 45 | RAB7A | RAB7A, member RAS oncogene family | Cytoplasm | enzyme | |
| chr20 | *RP11-305P22.9* | 23 | 1760 | 45 | | | | | |
| chr11 | *DIXDC1* | 18 | 537 | 45 | DIXDC1 | DIX domain containing 1 | Cytoplasm | other | |
| chr10 | *FBXW4* | 19 | 588 | 45 | FBXW4 | F-box and WD repeat domain containing 4 | Cytoplasm | other | |
| chr3 | *SLC25A26* | 21 | 481 | 45 | SLC25A26 | solute carrier family 25 member | Cytoplasm | transporter | |
| chr10 | *FRMPD2* | 21 | 933 | 45 | FRMPD2 | FERM and PDZ domain | Plasma Membrane | other | |
| chr17 | *MYH10* | 20 | 500 | 45 | MYH10 | myosin heavy chain 10 | Cytoplasm | enzyme | |
| chr7 | *RP11-396K3.1* | 18 | 678 | 45 | | | | | |
| chr1 | *ROR1* | 21 | 427 | 45 | ROR1 | receptor tyrosine kinase like orphan receptor 1 | Plasma Membrane | kinase | cirmtuzumab, VLS-101 |

| | | | | | | | | | |
|---|---|---|---|---|---|---|---|---|---|
| chr9 | *FUBP3* | 20 | 857 | 45 | FUBP3 | far upstream element binding protein 3 | Nucleus | transcription regulator | |
| chr3 | *RAD54L2* | 20 | 486 | 45 | RAD54L2 | RAD54 like 2 | Nucleus | transcription regulator | |
| chr11 | *ST14* | 20 | 1029 | 45 | ST14 | suppression of tumorigenicity 14 | Plasma Membrane | peptidase | |
| chr15 | *RP11-561C5.5* | 21 | 973 | 45 | | | | | |
| chr17 | *MED1* | 17 | 501 | 45 | MED1 | mediator complex subunit 1 | Nucleus | transcription regulator | |
| chr11 | *PHLDB1* | 20 | 788 | 45 | PHLDB1 | pleckstrin homology like domain family B member 1 | Cytoplasm | other | |
| chr10 | *ANXA11* | 18 | 766 | 45 | ANXA11 | annexin A11 | Nucleus | other | |
| chr9 | *MEGF9* | 16 | 519 | 45 | MEGF9 | multiple EGF like domains 9 | Extracellular Space | other | |
| chr16 | *CDH1* | 21 | 550 | 45 | CDH1 | cadherin 1 | Plasma Membrane | other | |
| chr11 | *RP11-356J5.12* | 20 | 592 | 45 | | | | | |
| chr1 | *ZNF678* | 19 | 618 | 45 | ZNF678 | zinc finger protein 678 | Nucleus | transcription regulator | |
| chr5 | *CTB-57H20.1* | 19 | 450 | 45 | | | | | |
| chr12 | *TEAD4* | 19 | 748 | 45 | TEAD4 | TEA domain transcription factor 4 | Nucleus | transcription regulator | |
| chr1 | *SLC1A7* | 20 | 899 | 45 | SLC1A7 | solute carrier family 1 member 7 | Plasma Membrane | transporter | riluzole |
| chr15 | *ARID3B* | 18 | 662 | 45 | ARID3B | AT-rich interaction domain 3B | Nucleus | transcription regulator | |
| chr22 | *ZC3H7B* | 18 | 772 | 45 | ZC3H7B | zinc finger CCCH-type containing | Nucleus | other | |
| chr1 | *DNM3* | 21 | 575 | 45 | DNM3 | dynamin 3 | Cytoplasm | enzyme | |
| chr1 | *PTAFR* | 20 | 587 | 45 | PTAFR | platelet activating factor receptor | Plasma Membrane | G-protein coupled recepto | lexipafant |
| chr17 | *EXOC7* | 19 | 634 | 45 | EXOC7 | exocyst complex component 7 | Cytoplasm | transporter | |
| chr5 | *OSMR* | 20 | 1679 | 45 | OSMR | oncostatin M receptor | Plasma Membrane | transmembrane receptor | |
| chr16 | *CPPED1* | 21 | 454 | 45 | CPPED1 | calcineurin like phosphoesterase domain containing 1 | Cytoplasm | enzyme | |
| chr3 | *FNDC3B* | 15 | 423 | 45 | FNDC3B | fibronectin type III domain containing 3B | Cytoplasm | other | |
| chr17 | *RP11-567L7.6* | 19 | 398 | 45 | | | | | |
| chr9 | *SPIN1* | 17 | 455 | 45 | SPIN1 | spindlin 1 | Nucleus | other | |
| chr17 | *AC005152.2* | 20 | 515 | 45 | | | | | |
| chr2 | *CCDC138* | 18 | 542 | 45 | CCDC138 | coiled-coil domain containing 138 | Other | other | |
| chr17 | *SPOP* | 16 | 430 | 45 | SPOP | speckle type BTB/POZ protein | Extracellular Space | other | |
| chr7 | *AC009784.3* | 20 | 646 | 45 | | | | | |
| chr12 | *IFLTD1* | 17 | 626 | 45 | LMNTD1 | lamin tail domain containing 1 | Nucleus | other | |
| chr15 | *ARHGAP11B* | 17 | 545 | 45 | ARHGAP11B | Rho GTPase activating protein | Cytoplasm | other | |
| chr6 | *TMEM181* | 16 | 657 | 45 | TMEM181 | transmembrane protein 181 | Other | other | |
| chr7 | *hsa-mir-490* | 18 | 542 | 45 | | | | | |
| chr11 | *TUB* | 21 | 919 | 45 | TUB | TUB bipartite transcription factor | Cytoplasm | transcription regulator | |
| chr20 | *TTLL9* | 20 | 848 | 45 | TTLL9 | tubulin tyrosine ligase like 9 | Other | other | |
| chr10 | *BICC1* | 21 | 783 | 45 | BICC1 | BicC family RNA binding protein | Other | other | |
| chr7 | *SSPO* | 22 | 1000 | 45 | SSPO | SCO-spondin | Cytoplasm | other | |
| chr10 | *CHST3* | 20 | 1187 | 45 | CHST3 | carbohydrate sulfotransferase 3 | Cytoplasm | enzyme | |
| chr10 | *PDCD11* | 19 | 537 | 45 | PDCD11 | programmed cell death 11 | Nucleus | other | |
| chr7 | *POM121C* | 20 | 729 | 45 | POM121/POM121C | POM121 transmembrane | Nucleus | other | |
| chr1 | *DNAJC11* | 19 | 778 | 45 | DNAJC11 | DnaJ heat shock protein family (Hsp40) member C11 | Cytoplasm | other | |
| chr17 | *CRK* | 19 | 831 | 45 | CRK | CRK proto-oncogene, adaptor protein | Cytoplasm | other | |
| chr1 | *SUSD4* | 22 | 729 | 45 | SUSD4 | sushi domain containing 4 | Other | other | |
| chr5 | *SLC12A7* | 25 | 2130 | 45 | SLC12A7 | solute carrier family 12 member 7 | Plasma Membrane | transporter | |
| chr15 | *UBR1* | 17 | 461 | 45 | UBR1 | ubiquitin protein ligase E3 component n-recognin 1 | Cytoplasm | enzyme | |
| chr1 | *INTS3* | 18 | 779 | 45 | INTS3 | integrator complex subunit 3 | Nucleus | other | |
| chr10 | *NUTM2A-AS1* | 18 | 587 | 45 | NUTM2A-AS1 | NUTM2A antisense RNA 1 | Other | other | |
| chr14 | *EGLN3* | 17 | 419 | 45 | EGLN3 | egl-9 family hypoxia inducible | Cytoplasm | enzyme | molidustat |
| chr21 | *PCNT* | 19 | 845 | 45 | PCNT | pericentrin | Cytoplasm | other | |
| chr3 | *DCP1A* | 17 | 459 | 45 | DCP1A | decapping mRNA 1A | Nucleus | other | |
| chr10 | *KIAA1598* | 17 | 479 | 45 | SHTN1 | shootin 1 | Plasma Membrane | other | |
| chr7 | *PPP1R9A* | 19 | 489 | 45 | PPP1R9A | protein phosphatase 1 regulatory subunit 9A | Plasma Membrane | other | |
| chrX | *ATP2B3* | 20 | 964 | 45 | ATP2B3 | ATPase plasma membrane Ca2+ transporting 3 | Plasma Membrane | transporter | |
| chr2 | *GALM* | 21 | 626 | 45 | GALM | galactose mutarotase | Cytoplasm | enzyme | |
| chr12 | *SLC4A8* | 20 | 754 | 44 | SLC4A8 | solute carrier family 4 member 8 | Plasma Membrane | transporter | |
| chr7 | *NXPH1* | 18 | 621 | 44 | NXPH1 | neurexophilin 1 | Extracellular Space | other | |
| chr3 | *NEK11* | 19 | 536 | 44 | NEK11 | NIMA related kinase 11 | Nucleus | kinase | dabrafenib/ipilimumab/trametinib, dabrafenib, dabrafenib/panitumumab/trametinib, dabrafenib/trametinib, dabrafenib/trametinib/vemurafenib, cetuximab/dabrafenib/trametinib. |
| chr19 | *RANBP3* | 18 | 712 | 44 | RANBP3 | RAN binding protein 3 | Nucleus | other | |
| chr4 | *ACOX3* | 17 | 810 | 44 | ACOX3 | acyl-CoA oxidase 3, pristanoyl | Cytoplasm | enzyme | |
| chr16 | *ST3GAL2* | 19 | 629 | 44 | ST3GAL2 | ST3 beta-galactoside alpha-2,3-sialyltransferase 2 | Cytoplasm | enzyme | |
| chr20 | *DNAJC5* | 20 | 1159 | 44 | DNAJC5 | DnaJ heat shock protein family (Hsp40) member C5 | Plasma Membrane | other | |
| chr21 | *KCNJ6* | 17 | 503 | 44 | KCNJ6 | potassium inwardly rectifying channel subfamily J member 6 | Plasma Membrane | ion channel | |
| chr16 | *GPR114* | 18 | 692 | 44 | ADGRG5 | adhesion G protein-coupled receptor G5 | Plasma Membrane | G-protein coupled recepto | |
| chr10 | *SAMD8* | 18 | 731 | 44 | SAMD8 | sterile alpha motif domain containing 8 | Cytoplasm | enzyme | |
| chr19 | *CELF5* | 17 | 659 | 44 | CELF5 | CUGBP Elav-like family member | Nucleus | other | |

| | | | | | | | | | |
|---|---|---|---|---|---|---|---|---|---|
| chr16 | *COG4* | 19 | 494 | 44 | COG4 | component of oligomeric golgi complex 4 | Cytoplasm | transporter | |
| chr1 | *JAK1* | 17 | 542 | 44 | JAK1 | Janus kinase 1 | Cytoplasm | kinase | erlotinib/ruxolitinib, baricitinib, ruxolitinib, tofacitinib, jaktinib, itacitinib, momelotinib, INCB052793, filgotinib, upadacitinib, methotrexate/tofacitinib, brepocitinib, |
| chr4 | *POLN* | 18 | 617 | 44 | POLN | DNA polymerase nu | Nucleus | enzyme | |
| chr7 | *PHF14* | 17 | 490 | 44 | PHF14 | PHD finger protein 14 | Nucleus | other | |
| chr6 | *PNPLA1* | 19 | 618 | 44 | PNPLA1 | patatin like phospholipase domain containing 1 | Cytoplasm | enzyme | |
| chr17 | *ULK2* | 18 | 467 | 44 | ULK2 | unc-51 like autophagy activating kinase 2 | Cytoplasm | kinase | |
| chr1 | *DENND2C* | 18 | 541 | 44 | DENND2C | DENN domain containing 2C | Other | other | |
| chr5 | *SSBP2* | 18 | 436 | 44 | SSBP2 | single stranded DNA binding protein 2 | Nucleus | transcription regulator | |
| chr14 | *WDR20* | 17 | 553 | 44 | WDR20 | WD repeat domain 20 | Nucleus | other | |
| chr7 | *LRGUK* | 20 | 492 | 44 | LRGUK | leucine rich repeats and guanylate kinase domain | Cytoplasm | other | |
| chr2 | *HDLBP* | 16 | 527 | 44 | HDLBP | high density lipoprotein binding protein | Nucleus | transporter | |
| chr7 | *GTF2IP1* | 18 | 623 | 44 | GTF2IP1 | general transcription factor IIi pseudogene 1 | Other | other | |
| chr2 | *PLEKHB2* | 16 | 653 | 44 | PLEKHB2 | pleckstrin homology domain containing B2 | Other | other | |
| chr11 | *QSER1* | 18 | 542 | 44 | QSER1 | glutamine and serine rich 1 | Other | other | |
| chr15 | *ADAMTS7* | 22 | 1120 | 44 | ADAMTS7 | ADAM metallopeptidase with thrombospondin type 1 motif 7 | Extracellular Space | peptidase | |
| chr16 | *SYT17* | 20 | 563 | 44 | SYT17 | synaptotagmin 17 | Plasma Membrane | other | |
| chr1 | *SYT14* | 19 | 525 | 44 | SYT14 | synaptotagmin 14 | Other | transporter | |
| chr10 | *PCGF6* | 18 | 498 | 44 | PCGF6 | polycomb group ring finger 6 | Nucleus | transcription regulator | |
| chr9 | *SLC24A2* | 21 | 457 | 44 | SLC24A2 | solute carrier family 24 member 2 | Plasma Membrane | transporter | |
| chrX | *MSN* | 20 | 486 | 44 | MSN | moesin | Plasma Membrane | other | |
| chr11 | *SIK2* | 17 | 416 | 44 | SIK2 | salt inducible kinase 2 | Cytoplasm | kinase | |
| chr4 | *RBM47* | 15 | 448 | 44 | RBM47 | RNA binding motif protein 47 | Nucleus | other | |
| chr1 | *PPFIA4* | 20 | 1079 | 44 | PPFIA4 | PTPRF interacting protein alpha 4 | Plasma Membrane | phosphatase | |
| chr4 | *GRK4* | 16 | 556 | 44 | GRK4 | G protein-coupled receptor | Plasma Membrane | kinase | |
| chr2 | *SLC8A1-AS1* | 20 | 587 | 44 | SLC8A1-AS1 | SLC8A1 antisense RNA 1 | Other | other | |
| chr1 | *SRGAP2* | 19 | 788 | 44 | SRGAP2 | SLIT-ROBO Rho GTPase activating protein 2 | Cytoplasm | other | |
| chr3 | *KIF15* | 19 | 459 | 44 | KIF15 | kinesin family member 15 | Nucleus | other | |
| chr14 | *TDRD9* | 14 | 427 | 44 | TDRD9 | tudor domain containing 9 | Cytoplasm | enzyme | |
| chr16 | *ATP6V0D1* | 18 | 874 | 44 | ATP6V0D1 | ATPase H+ transporting V0 subunit d1 | Cytoplasm | transporter | |
| chr17 | *RAB37* | 19 | 623 | 44 | RAB37 | RAB37, member RAS oncogene family | Cytoplasm | enzyme | |
| chr8 | *NSMCE2* | 16 | 375 | 44 | NSMCE2 | NSE2 (MMS21) homolog, SMC5-SMC6 complex SUMO ligase | Nucleus | enzyme | |
| chr2 | *FBLN7* | 20 | 986 | 44 | FBLN7 | fibulin 7 | Extracellular Space | other | |
| chr9 | *CORO2A* | 17 | 583 | 44 | CORO2A | coronin 2A | Cytoplasm | other | |
| chr16 | *OTOA* | 19 | 500 | 44 | OTOA | otoancorin | Extracellular Space | other | |
| chr19 | *WDR62* | 18 | 776 | 44 | WDR62 | WD repeat domain 62 | Nucleus | other | |
| chr5 | *ZNF346* | 18 | 527 | 44 | ZNF346 | zinc finger protein 346 | Nucleus | other | |
| chr18 | *COLEC12* | 20 | 465 | 44 | COLEC12 | collectin subfamily member 12 | Plasma Membrane | transmembrane receptor | |
| chr16 | *TERF2IP* | 18 | 415 | 44 | TERF2IP | TERF2 interacting protein | Nucleus | other | |
| chr6 | *NEDD9* | 20 | 524 | 44 | NEDD9 | neural precursor cell expressed, developmentally down-regulated | Nucleus | other | |
| chr4 | *SH3TC1* | 19 | 917 | 44 | SH3TC1 | SH3 domain and tetratricopeptide repeats 1 | Extracellular Space | other | |
| chr7 | *ESYT2* | 18 | 585 | 44 | ESYT2 | extended synaptotagmin 2 | Plasma Membrane | other | |
| chr17 | *ATP6V0A1* | 18 | 487 | 44 | ATP6V0A1 | ATPase H+ transporting V0 subunit a1 | Cytoplasm | transporter | |
| chr7 | *MTERF* | 18 | 656 | 44 | MTERF1 | mitochondrial transcription termination factor 1 | Cytoplasm | transcription regulator | |
| chr15 | *AQR* | 15 | 468 | 44 | AQR | aquarius intron-binding spliceosomal factor | Nucleus | other | |
| chr17 | *EFCAB5* | 19 | 407 | 44 | EFCAB5 | EF-hand calcium binding domain | Other | other | |
| chr20 | *MROH8* | 20 | 548 | 44 | MROH8 | maestro heat like repeat family member 8 | Other | other | |
| chr3 | *PFKFB4* | 18 | 750 | 44 | PFKFB4 | 6-phosphofructo-2-kinase/fructose-2,6- | Cytoplasm | kinase | |
| chr17 | *SUZ12* | 18 | 472 | 44 | SUZ12 | SUZ12 polycomb repressive complex 2 subunit | Nucleus | enzyme | |
| chr17 | *LASP1* | 19 | 795 | 44 | LASP1 | LIM and SH3 protein 1 | Cytoplasm | transporter | |
| chr5 | *DROSHA* | 19 | 1318 | 44 | DROSHA | drosha ribonuclease III | Nucleus | enzyme | |
| chr7 | *TRIM24* | 19 | 530 | 44 | TRIM24 | tripartite motif containing 24 | Nucleus | transcription regulator | |
| chr17 | *HDAC5* | 19 | 924 | 44 | HDAC5 | histone deacetylase 5 | Nucleus | transcription regulator | tributyrin, belinostat, pyroxamide |
| chr16 | *MYLK3* | 20 | 700 | 44 | MYLK3 | myosin light chain kinase 3 | Cytoplasm | kinase | |
| chr1 | *RP1-15D23.2* | 18 | 509 | 44 | | | | | |
| chr7 | *SEMA3E* | 18 | 474 | 44 | SEMA3E | semaphorin 3E | Extracellular Space | other | |
| chr3 | *RP11-88I21.2* | 19 | 960 | 44 | | | | | |
| chr11 | *DYNC2H1* | 18 | 444 | 44 | DYNC2H1 | dynein cytoplasmic 2 heavy chain | Cytoplasm | other | |

| | | | | | | | | | |
|---|---|---|---|---|---|---|---|---|---|
| chr5 | *ATP10B* | 19 | 563 | 44 | ATP10B | ATPase phospholipid transporting 10B (putative) | Plasma Membrane | transporter | |
| chr7 | *RPA3-AS1* | 18 | 541 | 44 | UMAD1 | UBAP1-MVB12-associated (UMA) domain containing 1 | Other | other | |
| chr12 | *TRPV4* | 20 | 1054 | 44 | TRPV4 | transient receptor potential cation channel subfamily V member 4 | Plasma Membrane | ion channel | |
| chr15 | *ABHD2* | 19 | 620 | 43 | ABHD2 | abhydrolase domain containing 2 | Cytoplasm | enzyme | |
| chr3 | *MRAS* | 20 | 800 | 43 | MRAS | muscle RAS oncogene homolog | Plasma Membrane | enzyme | |
| chr12 | *FGD6* | 18 | 405 | 43 | FGD6 | FYVE, RhoGEF and PH domain containing 6 | Cytoplasm | other | |
| chr5 | *NLN* | 19 | 454 | 43 | NLN | neurolysin | Cytoplasm | peptidase | |
| chr9 | *TJP2* | 18 | 455 | 43 | TJP2 | tight junction protein 2 | Plasma Membrane | kinase | |
| chr3 | *DHX30* | 19 | 633 | 43 | DHX30 | DExH-box helicase 30 | Nucleus | enzyme | |
| chr4 | *WHSC1* | 16 | 540 | 43 | NSD2 | nuclear receptor binding SET domain protein 2 | Nucleus | enzyme | |
| chr11 | *PRR5L* | 20 | 587 | 43 | PRR5L | proline rich 5 like | Cytoplasm | other | |
| chr18 | *OSBPL1A* | 15 | 361 | 43 | OSBPL1A | oxysterol binding protein like 1A | Cytoplasm | other | |
| chr15 | *MCTP2* | 20 | 651 | 43 | MCTP2 | multiple C2 and transmembrane domain containing 2 | Other | other | |
| chr7 | *PCLO* | 18 | 477 | 43 | PCLO | piccolo presynaptic cytomatrix | Cytoplasm | transporter | |
| chr3 | *IP6K2* | 18 | 554 | 43 | IP6K2 | inositol hexakisphosphate kinase | Cytoplasm | kinase | |
| chr3 | *PRKCD* | 18 | 857 | 43 | PRKCD | protein kinase C delta | Cytoplasm | kinase | ingenol mebutate |
| chr6 | *FOXP4* | 20 | 768 | 43 | FOXP4 | forkhead box P4 | Nucleus | transcription regulator | |
| chr19 | *LRRC4B* | 19 | 692 | 43 | LRRC4B | leucine rich repeat containing 4B | Plasma Membrane | other | |
| chr15 | *PSTPIP1* | 20 | 1021 | 43 | PSTPIP1 | proline-serine-threonine phosphatase interacting protein 1 | Cytoplasm | other | |
| chr1 | *RAP1A* | 17 | 534 | 43 | RAP1A | RAP1A, member of RAS oncogene family | Cytoplasm | enzyme | |
| chrX | *NLGN4X* | 20 | 430 | 43 | NLGN4X | neuroligin 4 X-linked | Plasma Membrane | enzyme | |
| chrX | *P2RY8* | 15 | 448 | 43 | P2RY8 | P2Y receptor family member 8 | Plasma Membrane | G-protein coupled recepto | |
| chr10 | *DMBT1* | 19 | 766 | 43 | DMBT1 | deleted in malignant brain tumors | Plasma Membrane | transmembrane receptor | |
| chr10 | *HTRA1* | 18 | 671 | 43 | HTRA1 | HtrA serine peptidase 1 | Extracellular Space | peptidase | |
| chr16 | *ITGAL* | 18 | 647 | 43 | ITGAL | integrin subunit alpha L | Plasma Membrane | transmembrane receptor | efalizumab |
| chr5 | *CAMK2A* | 18 | 659 | 43 | CAMK2A | calcium/calmodulin dependent protein kinase II alpha | Cytoplasm | kinase | |
| chr12 | *ACAD10* | 19 | 508 | 43 | ACAD10 | acyl-CoA dehydrogenase family member 10 | Cytoplasm | enzyme | |
| chr16 | *UNKL* | 20 | 908 | 43 | UNKL | unk like zinc finger | Cytoplasm | enzyme | |
| chr19 | *SAE1* | 16 | 476 | 43 | SAE1 | SUMO1 activating enzyme | Cytoplasm | enzyme | |
| chr20 | *TGM2* | 18 | 1163 | 43 | TGM2 | transglutaminase 2 | Cytoplasm | enzyme | |
| chr1 | *CAMSAP2* | 18 | 510 | 43 | CAMSAP2 | calmodulin regulated spectrin associated protein family member | Cytoplasm | other | |
| chr17 | *RAB40B* | 19 | 770 | 43 | RAB40B | RAB40B, member RAS oncogene family | Plasma Membrane | enzyme | |
| chr3 | *SH3BP5* | 20 | 524 | 43 | SH3BP5 | SH3 domain binding protein 5 | Cytoplasm | other | |
| chr6 | *GRM1* | 18 | 419 | 43 | GRM1 | glutamate metabotropic receptor | Plasma Membrane | G-protein coupled recepto | fasoracetam |
| chr9 | *RABEPK* | 19 | 610 | 43 | RABEPK | Rab9 effector protein with kelch motifs | Cytoplasm | other | |
| chr16 | *ARHGAP17* | 20 | 507 | 43 | ARHGAP17 | Rho GTPase activating protein 17 | Cytoplasm | other | |
| chr9 | *NUP214* | 19 | 598 | 43 | NUP214 | nucleoporin 214 | Nucleus | transporter | |
| chr1 | *PKP1* | 19 | 1089 | 43 | PKP1 | plakophilin 1 | Plasma Membrane | other | |
| chr3 | *XRN1* | 17 | 524 | 43 | XRN1 | 5'-3' exoribonuclease 1 | Cytoplasm | enzyme | |
| chr14 | *RGS6* | 16 | 448 | 43 | RGS6 | regulator of G protein signaling 6 | Cytoplasm | enzyme | |
| chr16 | *MEIOB* | 18 | 532 | 43 | MEIOB | meiosis specific with OB-fold | Cytoplasm | enzyme | |
| chr15 | *ST8SIA2* | 22 | 1044 | 43 | ST8SIA2 | ST8 alpha-N-acetyl-neuraminide alpha-2,8-sialyltransferase 2 | Cytoplasm | enzyme | |
| chr16 | *GINS3* | 18 | 495 | 43 | GINS3 | GINS complex subunit 3 | Nucleus | other | |
| chr3 | *IL17RD* | 18 | 479 | 43 | IL17RD | interleukin 17 receptor D | Cytoplasm | other | |
| chr10 | *HSPA12A* | 19 | 657 | 43 | HSPA12A | heat shock protein family A (Hsp70) member 12A | Cytoplasm | other | |
| chr16 | *ZDHHC7* | 18 | 655 | 43 | ZDHHC7 | zinc finger DHHC-type containing | Cytoplasm | enzyme | |
| chr9 | *GOLM1* | 18 | 610 | 43 | GOLM1 | golgi membrane protein 1 | Cytoplasm | other | |
| chr2 | *SRBD1* | 16 | 682 | 43 | SRBD1 | S1 RNA binding domain 1 | Extracellular Space | other | |
| chr2 | *FBXO11* | 17 | 526 | 43 | FBXO11 | F-box protein 11 | Cytoplasm | enzyme | |
| chr1 | *NUDC* | 18 | 567 | 43 | NUDC | nuclear distribution C, dynein complex regulator | Cytoplasm | other | |
| chr11 | *CKAP5* | 18 | 611 | 43 | CKAP5 | cytoskeleton associated protein 5 | Nucleus | transcription regulator | |
| chr12 | *TSPAN8* | 18 | 409 | 43 | TSPAN8 | tetraspanin 8 | Plasma Membrane | other | |
| chr16 | *MBTPS1* | 18 | 827 | 43 | MBTPS1 | membrane bound transcription factor peptidase, site 1 | Cytoplasm | peptidase | sonepcizumab |
| chr5 | *TCF7* | 18 | 845 | 43 | TCF7 | transcription factor 7 | Nucleus | transcription regulator | |
| chr3 | *NISCH* | 20 | 821 | 43 | NISCH | nischarin | Plasma Membrane | other | moxonidine, |
| chr16 | *AP1G1* | 16 | 477 | 43 | AP1G1 | adaptor related protein complex 1 subunit gamma 1 | Cytoplasm | other | |
| chr1 | *PHC2* | 17 | 593 | 43 | PHC2 | polyhomeotic homolog 2 | Nucleus | other | |
| chr19 | *MARK4* | 17 | 647 | 43 | MARK4 | microtubule affinity regulating kinase 4 | Cytoplasm | kinase | |
| chr16 | *TEKT5* | 19 | 593 | 43 | TEKT5 | tektin 5 | Nucleus | other | |
| chr20 | *NINL* | 15 | 686 | 43 | NINL | ninein like | Cytoplasm | other | |
| chr11 | *MRVI1* | 19 | 817 | 43 | MRVI1 | murine retrovirus integration site 1 homolog | Cytoplasm | other | |
| chr17 | *TTYH2* | 18 | 831 | 43 | TTYH2 | tweety family member 2 | Other | ion channel | |
| chr15 | *TRPM7* | 15 | 357 | 43 | TRPM7 | transient receptor potential cation channel subfamily M member 7 | Plasma Membrane | kinase | |
| chr16 | *NPIPB3* | 18 | 743 | 43 | PB4 (includes oth | nuclear pore complex interacting protein family member B4 | Nucleus | other | |
| chr2 | *SOS1* | 17 | 537 | 43 | SOS1 | SOS Ras/Rac guanine nucleotide exchange factor 1 | Cytoplasm | other | |
| chr21 | *DOPEY2* | 15 | 496 | 43 | DOP1B | DOP1 leucine zipper like protein | Cytoplasm | other | |

| | | | | | | | | | |
|---|---|---|---|---|---|---|---|---|---|
| chr15 | *CA12* | 18 | 537 | 43 | CA12 | carbonic anhydrase 12 | Plasma Membrane | enzyme | hydrochlorothiazide, trichloromethiazide, sulfacetamide, benzthiazide, topiramate, chlorothiazide, chlorthalidone, acetazolamide, methazolamide |
| chr3 | *RP11-493K19.3* | 18 | 923 | 43 | | | | | |
| chr1 | *FBXO28* | 16 | 538 | 43 | FBXO28 | F-box protein 28 | Nucleus | other | |
| chr2 | *AC144450.1* | 18 | 771 | 43 | | | | | |
| chr12 | *ATP2A2* | 16 | 544 | 43 | ATP2A2 | ATPase sarcoplasmic/endoplasmic | Cytoplasm | transporter | |
| chr5 | *MAPK9* | 17 | 785 | 43 | MAPK9 | mitogen-activated protein kinase 9 | Cytoplasm | kinase | cetuximab/encorafenib, binimetinib/encorafenib, encorafenib, binimetinib/cetuximab/encorafenib, binimetinib/encorafen |
| chr17 | *PIK3R6* | 18 | 627 | 43 | PIK3R6 | phosphoinositide-3-kinase regulatory subunit 6 | Cytoplasm | kinase | |
| chrX | *GPR64* | 17 | 432 | 43 | ADGRG2 | adhesion G protein-coupled receptor G2 | Plasma Membrane | i-protein coupled recepto | |
| chr1 | *FNBP1L* | 17 | 522 | 43 | FNBP1L | formin binding protein 1 like | Cytoplasm | other | |
| chr1 | *DOCK7* | 15 | 402 | 43 | DOCK7 | dedicator of cytokinesis 7 | Plasma Membrane | other | |
| chr9 | *SH2D3C* | 19 | 1040 | 43 | SH2D3C | SH2 domain containing 3C | Cytoplasm | other | |
| chr1 | *C1orf95* | 20 | 944 | 43 | STUM | stum, mechanosensory transduction mediator homolog | Other | other | |
| chr1 | *TMEM63A* | 18 | 968 | 43 | TMEM63A | transmembrane protein 63A | Cytoplasm | other | |
| chr5 | *NEURL1B* | 20 | 1342 | 43 | NEURL1B | neuralized E3 ubiquitin protein ligase 1B | Other | enzyme | |
| chr2 | *BMPR2* | 14 | 373 | 42 | BMPR2 | bone morphogenetic protein receptor type 2 | Plasma Membrane | kinase | |
| chr16 | *VWA3A* | 17 | 467 | 42 | VWA3A | von Willebrand factor A domain containing 3A | Other | other | |
| chr1 | *UBR4* | 17 | 496 | 42 | UBR4 | ubiquitin protein ligase E3 component n-recognin 4 | Nucleus | enzyme | |
| chr3 | *BOC* | 20 | 866 | 42 | BOC | BOC cell adhesion associated, oncogene regulated | Plasma Membrane | other | |
| chr15 | *FAM169B* | 19 | 976 | 42 | FAM169B | family with sequence similarity 169 member B | Other | other | |
| chr15 | *IL16* | 20 | 719 | 42 | IL16 | interleukin 16 | Extracellular Space | cytokine | |
| chr17 | *AC061992.2* | 17 | 703 | 42 | | | | | |
| chr19 | *SUGP1* | 20 | 746 | 42 | SUGP1 | SURP and G-patch domain containing 1 | Nucleus | other | |
| chr1 | *KPNA6* | 18 | 484 | 42 | KPNA6 | karyopherin subunit alpha 6 | Nucleus | transporter | |
| chr2 | *RGPD2* | 17 | 470 | 42 | PD4 (includes oth | RANBP2 like and GRIP domain containing 5 | Nucleus | enzyme | |
| chr10 | *TTC18* | 16 | 756 | 42 | CFAP70 | cilia and flagella associated | Cytoplasm | other | |
| chr2 | *MAP3K2* | 17 | 571 | 42 | MAP3K2 | mitogen-activated protein kinase kinase kinase 2 | Cytoplasm | kinase | |
| chr7 | *PUS7* | 18 | 422 | 42 | PUS7 | pseudouridine synthase 7 | Nucleus | enzyme | |
| chr5 | *FLT4* | 21 | 1279 | 42 | FLT4 | fms related tyrosine kinase 4 | Plasma Membrane | transmembrane receptor | imatinib/sorafenib, sorafenib, fruquintinib, lapatinib/pazopanib, lenvatinib, sorafenib/sulindac/sunitinib, IMC-3C5, imatinib/sunitinib, vandetanib, bevacizumab/sorafenib, dexamethasone/lenalidomide/sorafenib, sunitinib, JNJ-26483327, CEP 7055, axitinib, regorafenib, motesanib, 5-azacytidine/sorafenib, pazopanib, famitinib, cabozantinib/erlotinib, telatinib, [illegible] |
| chr7 | *RSPH10B2* | 17 | 757 | 42 | SPH10B/RSPH10 | radial spoke head 10 homolog B | Extracellular Space | other | |
| chr6 | *TTBK1* | 19 | 808 | 42 | TTBK1 | tau tubulin kinase 1 | Other | kinase | |
| chr3 | *EFCC1* | 18 | 989 | 42 | EFCC1 | EF-hand and coiled-coil domain containing 1 | Other | other | |
| chr1 | *KIF21B* | 18 | 1192 | 42 | KIF21B | kinesin family member 21B | Cytoplasm | other | |
| chr2 | *SCTR* | 18 | 805 | 42 | SCTR | secretin receptor | Plasma Membrane | i-protein coupled recepto | synthetic human secretin |
| chr1 | *EPHA8* | 20 | 965 | 42 | EPHA8 | EPH receptor A8 | Plasma Membrane | kinase | |

| | | | | | | | | | |
|---|---|---|---|---|---|---|---|---|---|
| chr1 | *HDAC1* | 18 | 500 | 42 | HDAC1 | histone deacetylase 1 | Nucleus | transcription regulator | R 306465, pyroxamide, bortezomib/vorinostat, OKI-179, tributyrin, trametinib/vorinostat, mocetinostat, belinostat, vorinostat, domatinostat, HDAC |
| chr17 | *FBXW10* | 16 | 436 | 42 | FBXW10 | F-box and WD repeat domain containing 10 | Cytoplasm | other | |
| chrX | *PHF16* | 16 | 410 | 42 | JADE3 | jade family PHD finger 3 | Nucleus | other | |
| chr10 | *BTBD16* | 18 | 641 | 42 | BTBD16 | BTB domain containing 16 | Other | other | |
| chr5 | *CDH9* | 16 | 899 | 42 | CDH9 | cadherin 9 | Plasma Membrane | other | |
| chr1 | *LMX1A* | 20 | 704 | 42 | LMX1A | LIM homeobox transcription factor 1 alpha | Nucleus | transcription regulator | |
| chr7 | *CAPZA2* | 16 | 387 | 42 | CAPZA2 | capping actin protein of muscle Z-line subunit alpha 2 | Cytoplasm | other | |
| chr12 | *DHX37* | 18 | 876 | 42 | DHX37 | DEAH-box helicase 37 | Nucleus | enzyme | |
| chr2 | *PPM1B* | 15 | 485 | 42 | PPM1B | protein phosphatase, Mg2+/Mn2+ dependent 1B | Cytoplasm | phosphatase | |
| chr16 | *IFT140* | 19 | 883 | 42 | IFT140 | intraflagellar transport 140 | Extracellular Space | other | |
| chr20 | *C20orf26* | 15 | 425 | 42 | CFAP61 | cilia and flagella associated | Cytoplasm | other | |
| chr14 | *ELMSAN1* | 18 | 596 | 42 | ELMSAN1 | ELM2 and Myb/SANT domain containing 1 | Nucleus | transcription regulator | |
| chr1 | *RP5-1180C18.1* | 17 | 555 | 42 | | | | | |
| chr22 | *KIAA1644* | 17 | 761 | 42 | SHISAL1 | shisa like 1 | Other | other | |
| chr16 | *LRRC36* | 16 | 398 | 42 | LRRC36 | leucine rich repeat containing 36 | Other | other | |
| chr5 | *CTD-2127H9.1* | 20 | 1649 | 42 | | | | | |
| chr15 | *SH3GL3* | 19 | 534 | 42 | SH3GL3 | SH3 domain containing GRB2 like 3, endophilin A3 | Cytoplasm | other | |
| chr11 | *LRP4* | 17 | 836 | 42 | LRP4 | LDL receptor related protein 4 | Extracellular Space | other | |
| chr15 | *RP11-307C19.2* | 18 | 977 | 42 | | | | | |
| chr20 | *KIAA1755* | 18 | 931 | 42 | KIAA1755 | KIAA1755 | Other | other | |
| chr15 | *RGMA* | 19 | 1401 | 42 | RGMA | repulsive guidance molecule BMP co-receptor a | Plasma Membrane | other | |
| chr7 | *DTX2* | 18 | 1240 | 42 | DTX2 | deltex E3 ubiquitin ligase 2 | Nucleus | other | |
| chr3 | *XYLB* | 19 | 540 | 42 | XYLB | xylulokinase | Cytoplasm | kinase | |
| chr16 | *AARS* | 17 | 599 | 42 | AARS1 | alanyl-tRNA synthetase 1 | Cytoplasm | enzyme | |
| chr3 | *SHISA5* | 17 | 702 | 42 | SHISA5 | shisa family member 5 | Nucleus | other | |
| chr11 | *USH1C* | 19 | 758 | 42 | USH1C | USH1 protein network component harmonin | Plasma Membrane | other | |
| chr1 | *SLC44A5* | 17 | 416 | 42 | SLC44A5 | solute carrier family 44 member 5 | Plasma Membrane | transporter | |
| chr8 | *RP1L1* | 18 | 783 | 42 | RP1L1 | RP1 like 1 | Cytoplasm | other | |
| chr5 | *PSD2* | 18 | 737 | 42 | PSD2 | pleckstrin and Sec7 domain containing 2 | Plasma Membrane | other | |
| chr18 | *TGIF1* | 17 | 801 | 42 | TGIF1 | TGFB induced factor homeobox 1 | Nucleus | transcription regulator | |
| chr2 | *AC007319.1* | 16 | 423 | 42 | | | | | |
| chr17 | *NT5M* | 17 | 588 | 42 | NT5M | 5',3'-nucleotidase, mitochondrial | Cytoplasm | phosphatase | |
| chr19 | *ARHGAP35* | 16 | 525 | 42 | ARHGAP35 | Rho GTPase activating protein 35 | Nucleus | transcription regulator | |
| chr17 | *SCN4A* | 18 | 861 | 42 | SCN4A | sodium voltage-gated channel alpha subunit 4 | Plasma Membrane | ion channel | riluzole, diclofenac/omeprazole, flecainide, diclofenac/misoprostol, diclofenac, |
| chr9 | *APBA1* | 19 | 464 | 42 | APBA1 | amyloid beta precursor protein binding family A member 1 | Cytoplasm | transporter | |
| chr14 | *MARK3* | 14 | 372 | 42 | MARK3 | microtubule affinity regulating kinase 3 | Cytoplasm | kinase | cdc25C phosphatase (211-221) |
| chr3 | *SFMBT1* | 16 | 404 | 42 | SFMBT1 | Scm like with four mbt domains 1 | Nucleus | transcription regulator | |
| chr7 | *CADPS2* | 19 | 399 | 42 | CADPS2 | calcium dependent secretion activator 2 | Plasma Membrane | other | |
| chr16 | *TERF2* | 18 | 644 | 42 | TERF2 | telomeric repeat binding factor 2 | Nucleus | transcription regulator | |
| chr9 | *CNTRL* | 17 | 637 | 42 | CNTRL | centriolin | Nucleus | transcription regulator | |
| chr12 | *ATF1* | 18 | 624 | 42 | ATF1 | activating transcription factor 1 | Nucleus | transcription regulator | |
| chr16 | *CNOT1* | 17 | 423 | 42 | CNOT1 | CCR4-NOT transcription complex subunit 1 | Cytoplasm | other | |
| chr16 | *NAA60* | 17 | 524 | 42 | NAA60 | N(alpha)-acetyltransferase 60, NatF catalytic subunit | Cytoplasm | enzyme | |
| chr20 | *TASP1* | 16 | 382 | 42 | TASP1 | taspase 1 | Nucleus | peptidase | |
| chr12 | *SH2B3* | 19 | 794 | 42 | SH2B3 | SH2B adaptor protein 3 | Plasma Membrane | other | |
| chr7 | *WBSCR16* | 17 | 631 | 42 | RCC1L | RCC1 like | Cytoplasm | other | |
| chr2 | *DGKD* | 16 | 768 | 42 | DGKD | diacylglycerol kinase delta | Cytoplasm | kinase | |
| chr10 | *CCNY* | 15 | 436 | 42 | CCNY | cyclin Y | Nucleus | other | |
| chr1 | *CEP350* | 16 | 473 | 42 | CEP350 | centrosomal protein 350 | Cytoplasm | other | |
| chr15 | *RP11-1084I9.1* | 18 | 514 | 42 | | | | | |
| chr10 | *SLC29A3* | 17 | 1293 | 42 | SLC29A3 | solute carrier family 29 member 3 | Plasma Membrane | transporter | |
| chr3 | *IP6K1* | 17 | 508 | 42 | IP6K1 | inositol hexakisphosphate kinase | Cytoplasm | kinase | |
| chr17 | *PRPF8* | 17 | 761 | 41 | PRPF8 | pre-mRNA processing factor 8 | Nucleus | other | |
| chr9 | *OLFM1* | 21 | 1082 | 41 | OLFM1 | olfactomedin 1 | Cytoplasm | other | |
| chr7 | *ZNF767* | 18 | 517 | 41 | ZNF767P | zinc finger family member 767, pseudogene | Other | other | |
| chr6 | *MYLK4* | 17 | 433 | 41 | MYLK4 | myosin light chain kinase family member 4 | Cytoplasm | kinase | |
| chr7 | *FAM20C* | 21 | 1211 | 41 | FAM20C | FAM20C golgi associated secretory pathway kinase | Extracellular Space | enzyme | |
| chr9 | *RP11-247A12.2* | 16 | 874 | 41 | | | | | |
| chrX | *HUWE1* | 16 | 586 | 41 | HUWE1 | HECT, UBA and WWE domain containing E3 ubiquitin protein ligase 1 | Nucleus | transcription regulator | |

| | | | | | | | | | |
|---|---|---|---|---|---|---|---|---|---|
| chr14 | *SMOC1* | 17 | 523 | 41 | SMOC1 | SPARC related modular calcium binding 1 | Extracellular Space | other | |
| chr8 | *PRKDC* | 13 | 414 | 41 | PRKDC | protein kinase, DNA-activated, catalytic subunit | Nucleus | kinase | panulisib, MSC2490484A, CC-115 |
| chr7 | *ARPC1A* | 18 | 725 | 41 | ARPC1A | actin related protein 2/3 complex subunit 1A | Extracellular Space | other | |
| chr5 | *PRLR* | 21 | 1177 | 41 | PRLR | prolactin receptor | Plasma Membrane | transmembrane receptor | fluoxymesterone, fluoxymesterone/tamoxifen |
| chr1 | *C1orf101* | 19 | 466 | 41 | CATSPERE | catsper channel auxiliary subunit epsilon | Extracellular Space | other | |
| chr18 | *ZNF516* | 16 | 698 | 41 | ZNF516 | zinc finger protein 516 | Nucleus | transcription regulator | |
| chr7 | *SEPT14* | 16 | 450 | 41 | SEPTIN14 | septin 14 | Cytoplasm | other | |
| chr1 | *CCDC30* | 18 | 415 | 41 | CCDC30 | coiled-coil domain containing 30 | Cytoplasm | other | |
| chr3 | *TMEM108* | 20 | 577 | 41 | TMEM108 | transmembrane protein 108 | Plasma Membrane | other | |
| chr1 | *SNAP47* | 18 | 1149 | 41 | SNAP47 | synaptosome associated protein | Plasma Membrane | other | |
| chr9 | *PPP2R4* | 17 | 906 | 41 | PTPA | protein phosphatase 2 phosphatase activator | Cytoplasm | phosphatase | |
| chr11 | *ATG13* | 17 | 601 | 41 | ATG13 | autophagy related 13 | Cytoplasm | other | |
| chr1 | *POGZ* | 15 | 513 | 41 | POGZ | pogo transposable element derived with ZNF domain | Nucleus | enzyme | |
| chr7 | *VPS41* | 17 | 506 | 41 | VPS41 | VPS41 subunit of HOPS complex | Cytoplasm | transporter | |
| chr5 | *RP11-6N13.1* | 17 | 458 | 41 | | | | | |
| chr1 | *FAM163A* | 20 | 642 | 41 | FAM163A | family with sequence similarity 163 member A | Other | other | |
| chr7 | *ELFN1* | 20 | 1141 | 41 | ELFN1 | extracellular leucine rich repeat and fibronectin type III domain containing 1 | Plasma Membrane | other | |
| chr14 | *UNC79* | 15 | 373 | 41 | UNC79 | unc-79 homolog, NALCN channel complex subunit | Other | other | |
| chr18 | *EPB41L3* | 18 | 581 | 41 | EPB41L3 | erythrocyte membrane protein band 4.1 like 3 | Plasma Membrane | other | |
| chr18 | *NFATC1* | 20 | 1005 | 41 | NFATC1 | nuclear factor of activated T cells | Nucleus | transcription regulator | |
| chr10 | *MARCH8;* | 17 | 618 | 41 | | | | | |
| chr5 | *PDGFRB* | 18 | 768 | 41 | PDGFRB | platelet derived growth factor receptor beta | Plasma Membrane | kinase | midostaurin, sorafenib, imatinib/inotuzumab ozogamicin, lapatinib/pazopanib, sorafenib/sulindac/sunitinib, imatinib/sunitinib, bevacizumab/sorafenib, dexamethasone/lenalidomide/sorafenib, sunitinib, 5-azacytidine/midostaurin, axitinib, regorafenib, 5-azacytidine/sorafenib, tandutinib, pazopanib, JI 101, dasatinib/nilotinib, dasatinib/erlotinib, telatinib, imatinib, becaplermin, [illegible] |
| chr3 | *AC018359.1* | 20 | 408 | 41 | | | | | |
| chr4 | *FAM193A* | 17 | 481 | 41 | FAM193A | family with sequence similarity 193 member A | Other | other | |
| chr1 | *LAMC1* | 16 | 494 | 41 | LAMC1 | laminin subunit gamma 1 | Extracellular Space | other | |
| chr17 | *MAP3K14* | 17 | 838 | 41 | MAP3K14 | mitogen-activated protein kinase kinase kinase 14 | Cytoplasm | kinase | |
| chr5 | *SIMC1* | 18 | 467 | 41 | SIMC1 | SUMO interacting motifs | Other | other | |
| chr20 | *RIMS4* | 19 | 874 | 41 | RIMS4 | regulating synaptic membrane exocytosis 4 | Plasma Membrane | other | |
| chr19 | *ZNF573* | 15 | 388 | 41 | ZNF573 | zinc finger protein 573 | Nucleus | other | |
| chr16 | *BCMO1* | 18 | 489 | 41 | BCO1 | beta-carotene oxygenase 1 | Cytoplasm | enzyme | |
| chr3 | *PXK* | 18 | 408 | 41 | PXK | PX domain containing serine/threonine kinase like | Cytoplasm | kinase | |
| chr12 | *PPFIBP1* | 16 | 499 | 41 | PPFIBP1 | PPFIA binding protein 1 | Plasma Membrane | other | |
| chr15 | *PRTG* | 15 | 399 | 41 | PRTG | protogenin | Extracellular Space | other | |
| chr5 | *MEF2C-AS1* | 18 | 377 | 41 | MEF2C-AS1 | MEF2C antisense RNA 1 | Other | other | |
| chr19 | *AP1M1* | 17 | 727 | 41 | AP1M1 | adaptor related protein complex 1 subunit mu 1 | Cytoplasm | transporter | |
| chr14 | *SLC7A8* | 15 | 442 | 41 | SLC7A8 | solute carrier family 7 member 8 | Plasma Membrane | transporter | |
| chr1 | *C1orf132* | 18 | 841 | 41 | MIR29B2CHG | MIR29B2 and MIR29C host gene | Other | other | |
| chr3 | *RAB6B* | 18 | 873 | 41 | RAB6B | RAB6B, member RAS oncogene family | Cytoplasm | enzyme | |
| chr14 | *ESRRB* | 16 | 502 | 41 | ESRRB | estrogen related receptor beta | Nucleus | d-dependent nuclear rec | |
| chr17 | *MARCH10;* | 17 | 542 | 41 | | | | | |
| chr10 | *MLLT10* | 14 | 380 | 41 | MLLT10 | MLLT10 histone lysine methyltransferase DOT1L | Nucleus | transcription regulator | |
| chr19 | *MAU2* | 19 | 770 | 41 | MAU2 | MAU2 sister chromatid cohesion factor | Nucleus | other | |
| chr5 | *AFF4* | 16 | 378 | 41 | AFF4 | AF4/FMR2 family member 4 | Nucleus | transcription regulator | |
| chr7 | *RBM33* | 18 | 501 | 41 | RBM33 | RNA binding motif protein 33 | Other | other | |

| | | | | | | | | | |
|---|---|---|---|---|---|---|---|---|---|
| chr7 | *MUC12* | 18 | 1121 | 41 | MUC12 | mucin 12, cell surface associated | Cytoplasm | other | |
| chr11 | *TPCN2* | 19 | 1260 | 41 | TPCN2 | two pore segment channel 2 | Cytoplasm | ion channel | |
| chr9 | *TMEM2* | 17 | 431 | 41 | CEMIP2 | cell migration inducing hyaluronidase 2 | Cytoplasm | enzyme | |
| chr19 | *OLFM2* | 16 | 523 | 41 | OLFM2 | olfactomedin 2 | Cytoplasm | other | |
| chr9 | *GFI1B* | 18 | 800 | 41 | GFI1B | growth factor independent 1B transcriptional repressor | Nucleus | transcription regulator | |
| chr1 | *SLC2A5* | 18 | 553 | 41 | SLC2A5 | solute carrier family 2 member 5 | Plasma Membrane | transporter | |
| chr1 | *PADI3* | 16 | 735 | 41 | PADI3 | peptidyl arginine deiminase 3 | Cytoplasm | enzyme | |
| chr22 | *BCL2L13* | 14 | 372 | 41 | BCL2L13 | BCL2 like 13 | Cytoplasm | other | |
| chr19 | *FAM129C* | 18 | 634 | 41 | NIBAN3 | niban apoptosis regulator 3 | Other | other | |
| chr15 | *NEDD4* | 16 | 344 | 41 | NEDD4 | NEDD4 E3 ubiquitin protein | Cytoplasm | enzyme | |
| chr9 | *LRRC8A* | 18 | 961 | 41 | LRRC8A | leucine rich repeat containing 8 VRAC subunit A | Plasma Membrane | ion channel | |
| chr17 | *PLXDC1* | 18 | 591 | 41 | PLXDC1 | plexin domain containing 1 | Plasma Membrane | other | |
| chr20 | *SLA2* | 17 | 750 | 41 | SLA2 | Src like adaptor 2 | Plasma Membrane | other | |
| chr7 | *POM121* | 19 | 534 | 41 | POM121/POM121 | POM121 transmembrane | Nucleus | other | |
| chr1 | *ZCCHC17* | 16 | 472 | 41 | ZCCHC17 | zinc finger CCHC-type containing | Nucleus | other | |
| chr22 | *KIAA0930* | 16 | 701 | 41 | KIAA0930 | KIAA0930 | Other | other | |
| chr17 | *SGSM2* | 18 | 753 | 41 | SGSM2 | small G protein signaling | Cytoplasm | other | |
| chr19 | *PBX4* | 17 | 635 | 41 | PBX4 | PBX homeobox 4 | Nucleus | transcription regulator | |
| chr8 | *KB-1047C11.2* | 17 | 394 | 41 | | | | | |
| chr7 | *LANCL2* | 16 | 560 | 40 | LANCL2 | LanC like 2 | Plasma Membrane | other | |
| chr17 | *DHRS7B* | 17 | 558 | 40 | DHRS7B | dehydrogenase/reductase 7B | Other | other | |
| chr7 | *RP11-313P13.3* | 18 | 464 | 40 | | | | | |
| chr16 | *NUP93* | 18 | 423 | 40 | NUP93 | nucleoporin 93 | Nucleus | other | |
| chr3 | *ATR* | 15 | 474 | 40 | ATR | ATR serine/threonine kinase | Nucleus | kinase | VX-803, berzosertib, BAY1895344, AZD6738 |
| chr6 | *PDSS2* | 14 | 367 | 40 | PDSS2 | decaprenyl diphosphate synthase subunit 2 | Cytoplasm | enzyme | |
| chr3 | *RASA2* | 16 | 446 | 40 | RASA2 | RAS p21 protein activator 2 | Cytoplasm | other | |
| chr16 | *CDH8* | 16 | 457 | 40 | CDH8 | cadherin 8 | Plasma Membrane | other | |
| chr15 | *SLC12A6* | 15 | 486 | 40 | SLC12A6 | solute carrier family 12 member 6 | Plasma Membrane | transporter | |
| chr1 | *SYCP1* | 17 | 478 | 40 | SYCP1 | synaptonemal complex protein 1 | Nucleus | other | |
| chr19 | *CTD-2189E23.1* | 17 | 499 | 40 | | | | | |
| chrX | *SCML2* | 16 | 389 | 40 | SCML2 | Scm polycomb group protein like | Nucleus | transcription regulator | |
| chr19 | *SIN3B* | 18 | 791 | 40 | SIN3B | SIN3 transcription regulator family member B | Nucleus | transcription regulator | |
| chr22 | *NFAM1* | 15 | 703 | 40 | NFAM1 | NFAT activating protein with ITAM motif 1 | Plasma Membrane | transmembrane receptor | |
| chr12 | *ATP5G2* | 17 | 659 | 40 | ATP5MC2 | ATP synthase membrane subunit c locus 2 | Cytoplasm | transporter | |
| chr19 | *ZFR2* | 18 | 645 | 40 | ZFR2 | zinc finger RNA binding protein 2 | Other | other | |
| chrX | *GPR143* | 18 | 480 | 40 | GPR143 | G protein-coupled receptor 143 | Plasma Membrane | G-protein coupled receptor | |
| chr1 | *WNT3A* | 18 | 1095 | 40 | WNT3A | Wnt family member 3A | Extracellular Space | cytokine | |
| chr9 | *OLFML2A* | 18 | 955 | 40 | OLFML2A | olfactomedin like 2A | Extracellular Space | other | |
| chr1 | *RNF115* | 17 | 607 | 40 | RNF115 | ring finger protein 115 | Cytoplasm | enzyme | |
| chr1 | *KIRREL* | 19 | 869 | 40 | KIRREL1 | kirre like nephrin family adhesion molecule 1 | Plasma Membrane | other | |
| chr2 | *MBD5* | 15 | 425 | 40 | MBD5 | methyl-CpG binding domain | Nucleus | other | |
| chr17 | *TRIM37* | 15 | 385 | 40 | TRIM37 | tripartite motif containing 37 | Cytoplasm | enzyme | |
| chr21 | *DIP2A* | 14 | 589 | 40 | DIP2A | disco interacting protein 2 | Nucleus | transcription regulator | |
| chr16 | *LUC7L* | 16 | 489 | 40 | LUC7L | LUC7 like | Nucleus | other | |
| chr20 | *TPX2* | 17 | 721 | 40 | TPX2 | TPX2 microtubule nucleation | Nucleus | other | |
| chr16 | *GALNS* | 18 | 922 | 40 | GALNS | galactosamine (N-acetyl)-6- | Cytoplasm | enzyme | |
| chr16 | *BCAR1* | 19 | 874 | 40 | BCAR1 | BCAR1 scaffold protein, Cas family member | Plasma Membrane | enzyme | |
| chr1 | *ANKRD13C* | 15 | 364 | 40 | ANKRD13C | ankyrin repeat domain 13C | Cytoplasm | other | |
| chr9 | *ADAMTS13* | 18 | 1053 | 40 | ADAMTS13 | ADAM metallopeptidase with thrombospondin type 1 motif 13 | Extracellular Space | peptidase | |
| chr5 | *LARP1* | 18 | 519 | 40 | LARP1 | La ribonucleoprotein domain family member 1 | Cytoplasm | translation regulator | |
| chr1 | *KIAA0319L* | 16 | 443 | 40 | KIAA0319L | KIAA0319 like | Cytoplasm | other | |
| chr15 | *TBC1D2B* | 16 | 885 | 40 | TBC1D2B | TBC1 domain family member 2B | Cytoplasm | other | |
| chr1 | *GPR161* | 17 | 585 | 40 | GPR161 | G protein-coupled receptor 161 | Plasma Membrane | G-protein coupled receptor | |
| chr5 | *EDIL3* | 17 | 421 | 40 | EDIL3 | EGF like repeats and discoidin domains 3 | Extracellular Space | other | |
| chr1 | *ARNT* | 16 | 525 | 40 | ARNT | aryl hydrocarbon receptor nuclear translocator | Nucleus | transcription regulator | |
| chr3 | *GXYLT2* | 18 | 382 | 40 | GXYLT2 | glucoside xylosyltransferase 2 | Other | other | |
| chr12 | *RBMS2* | 16 | 435 | 40 | RBMS2 | RNA binding motif single stranded interacting protein 2 | Nucleus | other | |
| chr22 | *PI4KAP2* | 14 | 622 | 40 | PI4KAP2 | phosphatidylinositol 4-kinase alpha pseudogene 2 | Other | other | |
| chr12 | *NDUFA12* | 16 | 368 | 40 | NDUFA12 | NADH:ubiquinone oxidoreductase subunit A12 | Cytoplasm | enzyme | |
| chr20 | *SEL1L2* | 17 | 430 | 40 | SEL1L2 | SEL1L2 adaptor subunit of ERAD E3 ligase | Other | other | |
| chr2 | *SERTAD2* | 16 | 447 | 40 | SERTAD2 | SERTA domain containing 2 | Cytoplasm | transcription regulator | |
| chr7 | *ACTB* | 17 | 1068 | 40 | ACTB | actin beta | Cytoplasm | other | |
| chr7 | *CTTNBP2* | 17 | 393 | 40 | CTTNBP2 | cortactin binding protein 2 | Cytoplasm | other | |
| chr1 | *IGSF3* | 19 | 728 | 40 | IGSF3 | immunoglobulin superfamily member 3 | Plasma Membrane | other | |
| chr8 | *PLEC* | 19 | 783 | 40 | PLEC | plectin | Cytoplasm | other | |

| | | | | | | | | | |
|---|---|---|---|---|---|---|---|---|---|
| chr2 | *C2orf48* | 17 | 683 | 40 | RRM2 | ribonucleotide reductase regulatory subunit M2 | Nucleus | enzyme | docetaxel/gemcitabine/vincristine, cladribine/cytarabine/filgrastim/idarubicin/plerixafor, cytarabine/filgrastim/fludarabine phosphate/imatinib, fludarabine phosphate/rituximab, cladribine, cytarabine/fludarabine phosphate/oxaliplatin/rituximab, gemcitabine/paclitaxel, cytarabine/fludarabine phosphate, cytarabine/filgrastim/fludarabine phosphate, gallium nitrate, hydroxyurea/6 |
| chr5 | *CDHR2* | 18 | 900 | 40 | CDHR2 | cadherin related family member 2 | Plasma Membrane | other | |
| chr2 | *AC018737.1* | 18 | 650 | 40 | | | | | |
| chr5 | *RP11-348J24.2* | 19 | 503 | 40 | | | | | |
| chr7 | *USP42* | 17 | 719 | 40 | USP42 | ubiquitin specific peptidase 42 | Nucleus | peptidase | |
| chr9 | *CNTNAP3B* | 15 | 435 | 40 | CNTNAP3B | contactin associated protein like | Other | other | |
| chr16 | *PDPR* | 17 | 425 | 40 | PDPR | pyruvate dehydrogenase phosphatase regulatory subunit | Cytoplasm | enzyme | |
| chr7 | *CD36* | 17 | 522 | 40 | CD36 | CD36 molecule | Plasma Membrane | transmembrane recepto | |
| chr3 | *FAM19A1* | 19 | 354 | 40 | TAFA1 | TAFA chemokine like family member 1 | Extracellular Space | other | |
| chr15 | *SEMA6D* | 17 | 321 | 40 | SEMA6D | semaphorin 6D | Plasma Membrane | other | |
| chr5 | *NDUFAF2* | 15 | 401 | 40 | NDUFAF2 | NADH:ubiquinone oxidoreductase complex | Cytoplasm | enzyme | |
| chr1 | *SRGAP2B* | 18 | 1070 | 40 | SRGAP2B | SLIT-ROBO Rho GTPase activating protein 2B | Other | other | |
| chr9 | *NTRK2* | 19 | 392 | 40 | NTRK2 | neurotrophic receptor tyrosine kinase 2 | Plasma Membrane | kinase | larotrectinib, NOV1601, DS-6051b, belizatinib, cabozantinib/nivolumab, lorlatinib, cabozantinib/erlotinib, entrectinib, repotrectinib, PLX7486, LOXO- |
| chr16 | *USP7* | 17 | 507 | 40 | USP7 | ubiquitin specific peptidase 7 | Nucleus | peptidase | |
| chr17 | *ACOX1* | 16 | 424 | 40 | ACOX1 | acyl-CoA oxidase 1 | Cytoplasm | enzyme | |
| chr7 | *GCK* | 19 | 819 | 40 | GCK | glucokinase | Cytoplasm | kinase | |
| chr7 | *COA1* | 15 | 574 | 40 | COA1 | cytochrome c oxidase assembly factor 1 homolog | Cytoplasm | other | |
| chr17 | *ASPSCR1* | 19 | 767 | 40 | ASPSCR1 | ASPSCR1 tether for SLC2A4, UBX domain containing | Cytoplasm | other | |
| chr16 | *C16orf70* | 17 | 498 | 40 | C16orf70 | chromosome 16 open reading frame 70 | Cytoplasm | other | |
| chr3 | *NEK10* | 16 | 439 | 40 | NEK10 | NIMA related kinase 10 | Other | kinase | |
| chr7 | *CRHR2* | 18 | 946 | 40 | CRHR2 | corticotropin releasing hormone receptor 2 | Plasma Membrane | -protein coupled recepto | |
| chr19 | *CTB-133G6.1* | 16 | 522 | 40 | | | | | |
| chr17 | *PLEKHM1* | 16 | 664 | 40 | PLEKHM1 | pleckstrin homology and RUN domain containing M1 | Cytoplasm | other | |
| chr19 | *MLLT1* | 18 | 867 | 40 | MLLT1 | MLLT1 super elongation complex subunit | Nucleus | transcription regulator | |
| chr1 | *LPGAT1* | 15 | 510 | 40 | LPGAT1 | lysophosphatidylglycerol acyltransferase 1 | Cytoplasm | enzyme | |
| chr15 | *FBN1* | 16 | 348 | 40 | FBN1 | fibrillin 1 | Extracellular Space | other | |
| chr17 | *YPEL2* | 18 | 453 | 40 | YPEL2 | yippee like 2 | Nucleus | other | |
| chr11 | *RP11-121M22.1* | 17 | 536 | 40 | | | | | |
| chr2 | *ANKMY1* | 16 | 679 | 40 | ANKMY1 | ankyrin repeat and MYND domain containing 1 | Other | other | |
| chr7 | *ELN* | 17 | 778 | 40 | ELN | elastin | Extracellular Space | other | |
| chr1 | *RP4-749H3.1* | 18 | 616 | 40 | | | | | |
| chr22 | *DEPDC5* | 16 | 380 | 40 | DEPDC5 | DEP domain containing 5 | Cytoplasm | other | |
| chr14 | *BRF1* | 15 | 723 | 40 | BRF1 | BRF1 RNA polymerase III transcription initiation factor | Nucleus | transcription regulator | |
| chr18 | *LPIN2* | 16 | 729 | 39 | LPIN2 | lipin 2 | Nucleus | phosphatase | |
| chr13 | *CRYL1* | 14 | 468 | 39 | CRYL1 | crystallin lambda 1 | Cytoplasm | enzyme | |
| chr2 | *UXS1* | 18 | 721 | 39 | UXS1 | UDP-glucuronate decarboxylase | Cytoplasm | enzyme | |
| chr4 | *MAML3* | 16 | 385 | 39 | MAML3 | mastermind like transcriptional coactivator 3 | Nucleus | transcription regulator | |
| chr12 | *ANO6* | 17 | 686 | 39 | ANO6 | anoctamin 6 | Plasma Membrane | ion channel | |
| chr5 | *CTB-43E15.3* | 17 | 816 | 39 | | | | | |
| chr1 | *FOXJ3* | 15 | 383 | 39 | FOXJ3 | forkhead box J3 | Nucleus | transcription regulator | |
| chr19 | *TLE2* | 15 | 620 | 39 | TLE2 | TLE family member 2, transcriptional corepressor | Nucleus | transcription regulator | |
| chr5 | *CNOT6* | 15 | 477 | 39 | CNOT6 | CCR4-NOT transcription complex subunit 6 | Nucleus | enzyme | |

| | | | | | | | | | |
|---|---|---|---|---|---|---|---|---|---|
| chr1 | *SDHC* | 16 | 546 | 39 | SDHC | succinate dehydrogenase complex subunit C | Cytoplasm | enzyme | |
| chrX | *CLCN4* | 17 | 611 | 39 | CLCN4 | chloride voltage-gated channel 4 | Plasma Membrane | ion channel | |
| chr12 | *RP11-438N16.1* | 18 | 547 | 39 | | | | | |
| chr3 | *NEK4* | 15 | 452 | 39 | NEK4 | NIMA related kinase 4 | Nucleus | kinase | |
| chr10 | *TMEM72-AS1* | 18 | 976 | 39 | TMEM72-AS1 | TMEM72 antisense RNA 1 | Other | other | |
| chr21 | *COL18A1* | 19 | 955 | 39 | COL18A1 | collagen type XVIII alpha 1 chain | Extracellular Space | other | collagenase clostridium |
| chr14 | *CEP128* | 13 | 331 | 39 | CEP128 | centrosomal protein 128 | Cytoplasm | other | |
| chr1 | *AKNAD1* | 17 | 581 | 39 | AKNAD1 | AKNA domain containing 1 | Cytoplasm | other | |
| chr12 | *FAR2* | 16 | 522 | 39 | FAR2 | fatty acyl-CoA reductase 2 | Cytoplasm | enzyme | |
| chr1 | *ATF6* | 16 | 483 | 39 | ATF6 | activating transcription factor 6 | Cytoplasm | transcription regulator | |
| chr16 | *COTL1* | 16 | 670 | 39 | COTL1 | coactosin like F-actin binding protein 1 | Cytoplasm | other | |
| chr7 | *IKZF1* | 17 | 686 | 39 | IKZF1 | IKAROS family zinc finger 1 | Nucleus | transcription regulator | |
| chr1 | *ALG14* | 15 | 477 | 39 | ALG14 | ALG14 UDP-N-acetylglucosaminyltransferase subunit | Cytoplasm | enzyme | |
| chr5 | *P4HA2* | 18 | 535 | 39 | P4HA2 | prolyl 4-hydroxylase subunit | Cytoplasm | transporter | |
| chr10 | *GPR123* | 19 | 1085 | 39 | ADGRA1 | adhesion G protein-coupled receptor A1 | Plasma Membrane | G-protein coupled recepto | |
| chr7 | *RSBN1L* | 15 | 437 | 39 | RSBN1L | round spermatid basic protein 1 | Other | other | |
| chr20 | *IFT52* | 18 | 553 | 39 | IFT52 | intraflagellar transport 52 | Cytoplasm | other | |
| chr14 | *PRKCH* | 15 | 383 | 39 | PRKCH | protein kinase C eta | Cytoplasm | kinase | ingenol mebutate |
| chr6 | *TBC1D22B* | 17 | 436 | 39 | TBC1D22B | TBC1 domain family member 22B | Other | other | |
| chr1 | *LRRN2* | 18 | 890 | 39 | LRRN2 | leucine rich repeat neuronal 2 | Plasma Membrane | other | |
| chr1 | *SYT6* | 17 | 731 | 39 | SYT6 | synaptotagmin 6 | Cytoplasm | transporter | |
| chr17 | *TRPV3* | 17 | 686 | 39 | TRPV3 | transient receptor potential cation channel subfamily V member 3 | Plasma Membrane | ion channel | |
| chr7 | *CDK6* | 17 | 539 | 39 | CDK6 | cyclin dependent kinase 6 | Nucleus | kinase | letrozole/ribociclib, abemaciclib/aromatase inhibitor/GnRH analog, abemaciclib/fulvestrant, ribociclib/tamoxifen, lerociclib, letrozole/palbociclib, anastrozole/palbociclib, abemaciclib/letrozole, palbociclib, alvocidib, exemestane/ribociclib, fulvestrant/palbociclib, HS-10342, abemaciclib/anastrozole, fulvestrant/ribociclib, FCN-437, abemaciclib/exemest[illegible] |
| chr3 | *CNOT10* | 15 | 368 | 39 | CNOT10 | CCR4-NOT transcription complex subunit 10 | Cytoplasm | other | |
| chr2 | *AC113607.1* | 14 | 523 | 39 | | | | | |
| chr20 | *CNBD2* | 17 | 608 | 39 | CNBD2 | cyclic nucleotide binding domain containing 2 | Cytoplasm | other | |
| chr7 | *SNX10* | 16 | 524 | 39 | SNX10 | sorting nexin 10 | Cytoplasm | transporter | |
| chr5 | *RBM27* | 15 | 389 | 39 | RBM27 | RNA binding motif protein 27 | Nucleus | other | |
| chr4 | *MAN2B2* | 14 | 735 | 39 | MAN2B2 | mannosidase alpha class 2B member 2 | Cytoplasm | enzyme | |
| chr12 | *CDK17* | 15 | 361 | 39 | CDK17 | cyclin dependent kinase 17 | Cytoplasm | kinase | |
| chr2 | *TRABD2A* | 19 | 486 | 39 | TRABD2A | TraB domain containing 2A | Plasma Membrane | peptidase | |
| chr6 | *ADTRP* | 17 | 457 | 39 | ADTRP | androgen dependent TFPI regulating protein | Plasma Membrane | enzyme | |
| chr20 | *BCAS1* | 17 | 739 | 39 | BCAS1 | breast carcinoma amplified sequence 1 | Plasma Membrane | other | |
| chr19 | *UPF1* | 18 | 902 | 39 | UPF1 | UPF1 RNA helicase and ATPase | Nucleus | enzyme | |
| chr5 | *GPBP1* | 16 | 413 | 39 | GPBP1 | GC-rich promoter binding protein | Nucleus | transcription regulator | |
| chr1 | *TMCC2* | 18 | 787 | 39 | TMCC2 | transmembrane and coiled-coil domain family 2 | Extracellular Space | other | |
| chr2 | *PSD4* | 18 | 835 | 39 | PSD4 | pleckstrin and Sec7 domain containing 4 | Cytoplasm | other | |
| chr17 | *TTLL6* | 17 | 430 | 39 | TTLL6 | tubulin tyrosine ligase like 6 | Cytoplasm | enzyme | |
| chr7 | *SEMA3C* | 15 | 420 | 39 | SEMA3C | semaphorin 3C | Extracellular Space | other | |
| chr10 | *DNA2* | 17 | 598 | 39 | DNA2 | DNA replication | Cytoplasm | enzyme | |
| chr5 | *UBTD2* | 16 | 492 | 39 | UBTD2 | ubiquitin domain containing 2 | Other | other | |
| chr16 | *NAE1* | 17 | 500 | 39 | NAE1 | NEDD8 activating enzyme E1 subunit 1 | Cytoplasm | enzyme | |
| chr22 | *MAPK1* | 13 | 390 | 39 | MAPK1 | mitogen-activated protein kinase 1 | Cytoplasm | kinase | LTT462, pexmetinib, ulixertinib, LY3214996, ASN007, binimetinib, ASTX029, KO-947, LY-3007113, MAP |
| chr7 | *RP5-945F2.3* | 19 | 375 | 39 | | | | | |
| chr5 | *FYB* | 17 | 1129 | 39 | FYB1 | FYN binding protein 1 | Nucleus | other | |

| | | | | | | | | | |
|---|---|---|---|---|---|---|---|---|---|
| chr7 | *GNGT1* | 15 | 471 | 39 | GNGT1 | G protein subunit gamma transducin 1 | Plasma Membrane | enzyme | |
| chr10 | *LRRC27* | 18 | 669 | 39 | LRRC27 | leucine rich repeat containing 27 | Other | other | |
| chr1 | *RCC1* | 16 | 568 | 39 | RCC1 | regulator of chromosome condensation 1 | Cytoplasm | other | |
| chr8 | *SGK223* | 15 | 535 | 39 | | | | | |
| chr1 | *RAB3B* | 16 | 417 | 39 | RAB3B | RAB3B, member RAS oncogene family | Cytoplasm | enzyme | |
| chr7 | *DFNA5* | 16 | 723 | 39 | GSDME | gasdermin E | Cytoplasm | other | |
| chr19 | *CTC-260E6.6* | 17 | 370 | 39 | | | | | |
| chr2 | *DTNB* | 15 | 440 | 39 | DTNB | dystrobrevin beta | Plasma Membrane | other | |
| chr15 | *KIAA1199* | 19 | 653 | 39 | CEMIP | cell migration inducing hyaluronidase 1 | Cytoplasm | enzyme | |
| chr5 | *TBC1D9B* | 19 | 1141 | 39 | TBC1D9B | TBC1 domain family member 9B | Other | other | |
| chr10 | *WAPAL* | 13 | 361 | 39 | WAPL | WAPL cohesin release factor | Nucleus | other | |
| chr16 | *NDRG4* | 17 | 661 | 39 | NDRG4 | NDRG family member 4 | Plasma Membrane | other | |
| chr5 | *RP11-114J13.1* | 17 | 486 | 39 | | | | | |
| chr2 | *ARHGAP25* | 17 | 488 | 39 | ARHGAP25 | Rho GTPase activating protein 25 | Cytoplasm | other | |
| chr1 | *SMIM12* | 18 | 554 | 39 | SMIM12 | small integral membrane protein | Other | other | |
| chr11 | *SCUBE2* | 16 | 556 | 39 | SCUBE2 | signal peptide, CUB domain and EGF like domain containing 2 | Extracellular Space | other | |
| chr4 | *GALNTL6* | 14 | 427 | 39 | GALNTL6 | polypeptide N-acetylgalactosaminyltransferase | Other | other | |
| chr3 | *CTDSPL* | 17 | 440 | 39 | CTDSPL | CTD small phosphatase like | Nucleus | other | |
| chr9 | *ODF2* | 17 | 634 | 39 | ODF2 | outer dense fiber of sperm tails 2 | Cytoplasm | other | |
| chr1 | *NAV1* | 18 | 728 | 39 | NAV1 | neuron navigator 1 | Other | enzyme | |
| chr17 | *HELZ* | 15 | 330 | 39 | HELZ | helicase with zinc finger | Nucleus | enzyme | |
| chr5 | *GRIA1* | 17 | 369 | 39 | GRIA1 | glutamate ionotropic receptor AMPA type subunit 1 | Plasma Membrane | ion channel | methoxyflurane, sevoflurane, isoflurane, perampanel, tezampanel, farampator, LY451395, |
| chr5 | *RP11-155G15.2* | 17 | 436 | 39 | | | | | |
| chr11 | *NARS2* | 15 | 450 | 39 | NARS2 | asparaginyl-tRNA synthetase 2, mitochondrial | Cytoplasm | enzyme | |
| chr15 | *FAM63B* | 12 | 340 | 39 | MINDY2 | MINDY lysine 48 deubiquitinase 2 | Nucleus | transporter | |
| chr19 | *HNRNPM* | 16 | 458 | 39 | HNRNPM | heterogeneous nuclear ribonucleoprotein M | Nucleus | other | |
| chr3 | *MED12L* | 15 | 411 | 39 | MED12L | mediator complex subunit 12L | Extracellular Space | other | |
| chr1 | *IL12RB2* | 17 | 392 | 39 | IL12RB2 | interleukin 12 receptor subunit | Plasma Membrane | transmembrane receptor | |
| chr12 | *SLC8B1* | 16 | 585 | 39 | SLC8B1 | solute carrier family 8 member B1 | Plasma Membrane | transporter | |
| chr19 | *AP3D1* | 17 | 694 | 39 | AP3D1 | adaptor related protein complex 3 subunit delta 1 | Cytoplasm | other | |
| chr20 | *ARHGAP40* | 17 | 715 | 39 | ARHGAP40 | Rho GTPase activating protein 40 | Cytoplasm | other | |
| chr19 | *VRK3* | 14 | 547 | 39 | VRK3 | VRK serine/threonine kinase 3 | Nucleus | kinase | |
| chr1 | *GS1-122H1.2* | 19 | 486 | 39 | | | | | |
| chr17 | *ADAP2* | 17 | 449 | 39 | ADAP2 | ArfGAP with dual PH domains 2 | Cytoplasm | other | |
| chr1 | *CEP104* | 17 | 634 | 39 | CEP104 | centrosomal protein 104 | Cytoplasm | other | |
| chr1 | *ZNF362* | 16 | 627 | 39 | ZNF362 | zinc finger protein 362 | Other | other | |
| chr6 | *SLC26A8* | 16 | 376 | 38 | SLC26A8 | solute carrier family 26 member 8 | Plasma Membrane | transporter | |
| chr8 | *MTMR7* | 17 | 393 | 38 | MTMR7 | myotubularin related protein 7 | Cytoplasm | phosphatase | |
| chr12 | *PTPN11* | 17 | 386 | 38 | PTPN11 | protein tyrosine phosphatase non-receptor type 11 | Cytoplasm | phosphatase | TNO155, RMC-4630, JAB-3068 |
| chr12 | *NCKAP5L* | 17 | 1158 | 38 | NCKAP5L | NCK associated protein 5 like | Cytoplasm | other | |
| chr1 | *SDC3* | 16 | 799 | 38 | SDC3 | syndecan 3 | Plasma Membrane | other | |
| chr18 | *ARHGAP28* | 16 | 385 | 38 | ARHGAP28 | Rho GTPase activating protein 28 | Cytoplasm | other | |
| chr3 | *PVRL3* | 16 | 425 | 38 | NECTIN3 | nectin cell adhesion molecule 3 | Plasma Membrane | other | |
| chr15 | *TCF12* | 13 | 345 | 38 | TCF12 | transcription factor 12 | Nucleus | transcription regulator | |
| chr16 | *RAB40C* | 18 | 906 | 38 | RAB40C | RAB40C, member RAS oncogene family | Plasma Membrane | enzyme | |
| chr12 | *BRI3BP* | 16 | 494 | 38 | BRI3BP | BRI3 binding protein | Extracellular Space | other | |
| chr19 | *AXL* | 17 | 609 | 38 | AXL | AXL receptor tyrosine kinase | Plasma Membrane | kinase | gilteritinib, cabozantinib/nivolumab, INCB081776, bosutinib, cabozantinib, DS-1205c, humax-AXL-ADC, ADCT-601, BPI-9016M, cabozantinib/erlotinib, ningetinib, ONO-7475, TP-0903, sitravatinib, SKI-G-801, bemcentinib, |
| chrY | *IL3RA* | 15 | 437 | 38 | IL3RA | interleukin 3 receptor subunit alpha | Plasma Membrane | transmembrane receptor | rituximab/sargramostim, EPO/sargramostim, IMGN632, flotetuzumab, sargramostim, DT388IL3, XmAb14045, |
| chr1 | *PIP5K1A* | 16 | 586 | 38 | PIP5K1A | phosphatidylinositol-4-phosphate 5-kinase type 1 alpha | Cytoplasm | kinase | |
| chr16 | *RP11-166B2.1* | 17 | 463 | 38 | | | | | |
| chr1 | *TARBP1* | 16 | 744 | 38 | TARBP1 | TAR (HIV-1) RNA binding protein | Nucleus | transcription regulator | |

| | | | | | | | | | |
|---|---|---|---|---|---|---|---|---|---|
| chr6 | *GFOD1* | 17 | 431 | 38 | GFOD1 | glucose-fructose oxidoreductase domain containing 1 | Other | enzyme | |
| chr1 | *KANK4* | 19 | 505 | 38 | KANK4 | KN motif and ankyrin repeat domains 4 | Cytoplasm | other | |
| chr17 | *MAP3K3* | 16 | 495 | 38 | MAP3K3 | mitogen-activated protein kinase kinase kinase 3 | Cytoplasm | kinase | |
| chr22 | *AP1B1* | 15 | 584 | 38 | AP1B1 | adaptor related protein complex 1 subunit beta 1 | Cytoplasm | other | |
| chr21 | *TRAPPC10* | 13 | 481 | 38 | TRAPPC10 | trafficking protein particle | Cytoplasm | other | |
| chr17 | *PLEKHM1P* | 16 | 665 | 38 | PLEKHM1P1 | pleckstrin homology and RUN domain containing M1 | Other | other | |
| chr9 | *RP11-154D17.1* | 16 | 411 | 38 | | | | | |
| chr11 | *PSMA1* | 16 | 421 | 38 | PSMA1 | proteasome subunit alpha 1 | Cytoplasm | peptidase | |
| chr15 | *PARP16* | 16 | 455 | 38 | PARP16 | poly(ADP-ribose) polymerase family member 16 | Cytoplasm | enzyme | |
| chr3 | *GOLGA4* | 15 | 369 | 38 | GOLGA4 | golgin A4 | Cytoplasm | other | |
| chr5 | *DTWD2* | 17 | 411 | 38 | DTWD2 | DTW domain containing 2 | Other | other | |
| chr1 | *NCMAP* | 16 | 553 | 38 | NCMAP | non-compact myelin associated protein | Plasma Membrane | other | |
| chr4 | *ARHGAP10* | 14 | 377 | 38 | ARHGAP10 | Rho GTPase activating protein 10 | Cytoplasm | other | |
| chr12 | *CCDC63* | 17 | 452 | 38 | CCDC63 | coiled-coil domain containing 63 | Other | other | |
| chr1 | *RP11-418J17.1* | 16 | 598 | 38 | | | | | |
| chr17 | *RP11-848P1.9* | 17 | 381 | 38 | | | | | |
| chr20 | *DNMT3B* | 17 | 792 | 38 | DNMT3B | DNA methyltransferase 3 beta | Nucleus | enzyme | |
| chr17 | *TOP3A* | 16 | 537 | 38 | TOP3A | DNA topoisomerase III alpha | Nucleus | enzyme | |
| chr3 | *TRIM71* | 17 | 507 | 38 | TRIM71 | tripartite motif containing 71 | Cytoplasm | enzyme | |
| chr20 | *SNTA1* | 17 | 703 | 38 | SNTA1 | syntrophin alpha 1 | Plasma Membrane | other | |
| chr1 | *TMEM51* | 16 | 606 | 38 | TMEM51 | transmembrane protein 51 | Other | other | |
| chr15 | *MAPK6* | 14 | 404 | 38 | MAPK6 | mitogen-activated protein kinase | Cytoplasm | kinase | |
| chr11 | *STK33* | 17 | 478 | 38 | STK33 | serine/threonine kinase 33 | Cytoplasm | kinase | |
| chr11 | *SLC22A6* | 16 | 710 | 38 | SLC22A6 | solute carrier family 22 member 6 | Plasma Membrane | transporter | allopurinol/lesinurad, lesinurad, colchicine/probenecid, probenecid |
| chr1 | *CDC14A* | 16 | 484 | 38 | CDC14A | cell division cycle 14A | Nucleus | phosphatase | |
| chr15 | *EIF2AK4* | 14 | 372 | 38 | EIF2AK4 | eukaryotic translation initiation factor 2 alpha kinase 4 | Cytoplasm | kinase | |
| chr8 | *NUGGC* | 17 | 416 | 38 | NUGGC | nuclear GTPase, germinal center associated | Nucleus | enzyme | |
| chr1 | *ESPNP* | 19 | 1539 | 38 | ESPNP | espin pseudogene | Other | other | |
| chr2 | *NCKAP5* | 15 | 344 | 38 | NCKAP5 | NCK associated protein 5 | Other | other | |
| chr12 | *PPTC7* | 15 | 421 | 38 | PPTC7 | PTC7 protein phosphatase | Cytoplasm | phosphatase | |
| chr8 | *RP11-13N12.1* | 14 | 352 | 38 | | | | | |
| chr3 | *EPHA3* | 15 | 421 | 38 | EPHA3 | EPH receptor A3 | Plasma Membrane | kinase | anti-EphA3 monoclonal antibody |
| chr3 | *PHC3* | 15 | 678 | 38 | PHC3 | polyhomeotic homolog 3 | Nucleus | transcription regulator | |
| chr5 | *BRD9* | 20 | 2131 | 38 | BRD9 | bromodomain containing 9 | Nucleus | other | |
| chr12 | *PPP1R12A* | 15 | 336 | 38 | PPP1R12A | protein phosphatase 1 regulatory subunit 12A | Cytoplasm | phosphatase | |
| chr19 | *TJP3* | 16 | 557 | 38 | TJP3 | tight junction protein 3 | Cytoplasm | other | |
| chr1 | *BRDT* | 14 | 370 | 38 | BRDT | bromodomain testis associated | Nucleus | kinase | BI 894999 |
| chr1 | *FAM69A* | 15 | 366 | 38 | DIPK1A | divergent protein kinase domain | Other | other | |
| chr1 | *CRB1* | 15 | 431 | 38 | CRB1 | crumbs cell polarity complex component 1 | Plasma Membrane | other | |
| chr12 | *CRADD* | 17 | 415 | 38 | CRADD | CASP2 and RIPK1 domain containing adaptor with death | Cytoplasm | other | |
| chr6 | *LAMA2* | 12 | 397 | 38 | LAMA2 | laminin subunit alpha 2 | Extracellular Space | other | |
| chr3 | *KCNH8* | 15 | 421 | 38 | KCNH8 | potassium voltage-gated channel subfamily H member 8 | Plasma Membrane | ion channel | |
| chr2 | *ATOH8* | 17 | 793 | 38 | ATOH8 | atonal bHLH transcription factor 8 | Nucleus | transcription regulator | |
| chr21 | *TMPRSS2* | 14 | 478 | 38 | TMPRSS2 | transmembrane serine protease 2 | Plasma Membrane | peptidase | |
| chr1 | *SLC6A17* | 18 | 812 | 38 | SLC6A17 | solute carrier family 6 member 17 | Cytoplasm | transporter | |
| chr15 | *ALPK3* | 19 | 703 | 38 | ALPK3 | alpha kinase 3 | Nucleus | kinase | |
| chr21 | *CXADR* | 15 | 365 | 38 | CXADR | CXADR Ig-like cell adhesion molecule | Plasma Membrane | transmembrane receptor | rAd-p53 |
| chr1 | *EPHA2* | 17 | 831 | 38 | EPHA2 | EPH receptor A2 | Plasma Membrane | kinase | blinatumomab/dasatinib, dasatinib/dexamethasone, regorafenib, cytarabine/dasatinib/dexamethasone/methotrexate, cytarabine/dasatinib/filgrastim/fludarabine phosphate, dasatinib/6-mercaptopurine/methotrexate, dasatinib/nilotinib, DS-8895a, cytarabine/dasatinib/filgrastim/fludarabine phosphate/idarubicin, dasatinib/prednisone, dasatinib/erlotinib, BT5528, cytarabine/dasatinib, dasatinib/inotuzumab [illegible] |

| | | | | | | | | | |
|---|---|---|---|---|---|---|---|---|---|
| chr19 | *BRSK1* | 16 | 728 | 38 | BRSK1 | BR serine/threonine kinase 1 | Cytoplasm | kinase | |
| chr5 | *CTD-2143L24.1* | 20 | 1126 | 38 | | | | | |
| chr12 | *EP400NL* | 16 | 664 | 38 | EP400P1 | EP400 pseudogene 1 | Other | other | |
| chrX | *HDHD1* | 15 | 435 | 38 | PUDP | pseudouridine 5'-phosphatase | Cytoplasm | enzyme | |
| chr22 | *CERK* | 16 | 663 | 38 | CERK | ceramide kinase | Plasma Membrane | kinase | |
| chr1 | *BRINP3* | 15 | 375 | 38 | BRINP3 | BMP/retinoic acid inducible neural specific 3 | Cytoplasm | other | |
| chr1 | *GNG12-AS1* | 17 | 403 | 38 | GNG12-AS1 | GNG12, DIRAS3 and WLS antisense RNA 1 | Other | other | |
| chr16 | *GFOD2* | 15 | 445 | 38 | GFOD2 | glucose-fructose oxidoreductase domain containing 2 | Extracellular Space | enzyme | |
| chr1 | *LINC00466* | 15 | 386 | 38 | LINC00466 | long intergenic non-protein coding RNA 466 | Other | other | |
| chr9 | *CIZ1* | 16 | 810 | 38 | CIZ1 | CDKN1A interacting zinc finger protein 1 | Nucleus | transporter | |
| chr7 | *THSD7A* | 17 | 390 | 38 | THSD7A | thrombospondin type 1 domain containing 7A | Cytoplasm | other | |
| chr8 | *BLK* | 16 | 574 | 38 | BLK | BLK proto-oncogene, Src family tyrosine kinase | Cytoplasm | kinase | afatinib/osimertinib, gefitinib/osimertinib, erlotinib/osimertinib, osimertinib |
| chr1 | *RASSF5* | 17 | 687 | 38 | RASSF5 | Ras association domain family member 5 | Plasma Membrane | other | |
| chr12 | *ADAMTS20* | 15 | 517 | 38 | ADAMTS20 | ADAM metallopeptidase with thrombospondin type 1 motif 20 | Extracellular Space | peptidase | |
| chr1 | *RLF* | 13 | 375 | 38 | RLF | rearranged L-myc fusion | Nucleus | transcription regulator | |
| chr19 | *KLHL26* | 17 | 713 | 38 | KLHL26 | kelch like family member 26 | Other | other | |
| chr3 | *NCK1* | 15 | 472 | 38 | NCK1 | NCK adaptor protein 1 | Cytoplasm | kinase | |
| chr20 | *PPP4R1L* | 15 | 679 | 38 | PPP4R1L | protein phosphatase 4 regulatory subunit 1 like (pseudogene) | Other | other | |
| chr16 | *TRAP1* | 15 | 494 | 38 | TRAP1 | TNF receptor associated protein | Cytoplasm | enzyme | |
| chr2 | *CCDC88A* | 15 | 398 | 38 | CCDC88A | coiled-coil domain containing 88A | Cytoplasm | other | |
| chr16 | *CTD-2547E10.2* | 16 | 828 | 38 | | | | | |
| chr12 | *PKP2* | 15 | 386 | 38 | PKP2 | plakophilin 2 | Plasma Membrane | other | |
| chr5 | *FBN2* | 15 | 380 | 38 | FBN2 | fibrillin 2 | Extracellular Space | other | |
| chr16 | *CACNA1H* | 19 | 681 | 38 | CACNA1H | calcium voltage-gated channel subunit alpha1 H | Plasma Membrane | ion channel | flunarizine, bepridil, mibefradil, isradipine, zonisamide, |
| chr17 | *SPNS2* | 16 | 835 | 38 | SPNS2 | sphingolipid transporter 2 | Other | transporter | |
| chr19 | *RFX2* | 15 | 573 | 38 | RFX2 | regulatory factor X2 | Nucleus | transcription regulator | |
| chr16 | *NPIPB4* | 16 | 681 | 38 | PB4 (includes oth | nuclear pore complex interacting protein family member B4 | Nucleus | other | |
| chr8 | *ADRA1A* | 17 | 392 | 37 | ADRA1A | adrenoceptor alpha 1A | Plasma Membrane | G-protein coupled recepto | chlorpheniramine/met hscopolamine/phenyl ephrine, apraclonidine, epinephrine/lidocaine , azatadine/pseudoeph edrine, dobutamine, benzphetamine, terazosine, naproxen/pseudoeph edrine, quinidine, doxazosin, dihydroergotamine, guanethidine, droperidol, tolazoline, brompheniramine/co deine/phenylpropanol amine, loratadine/pseudoep hedrine, chlorpheniramine/ibu profen/pseudoephedr [illegible] |
| chr7 | *DAGLB* | 16 | 758 | 37 | DAGLB | diacylglycerol lipase beta | Plasma Membrane | enzyme | |
| chr21 | *RRP1B* | 16 | 570 | 37 | RRP1B | ribosomal RNA processing 1B | Nucleus | transcription regulator | |
| chr1 | *RALGPS2* | 16 | 473 | 37 | RALGPS2 | Ral GEF with PH domain and SH3 binding motif 2 | Other | other | |
| chr2 | *GALNT14* | 14 | 427 | 37 | GALNT14 | polypeptide N-acetylgalactosaminyltransferase | Cytoplasm | enzyme | |
| chr8 | *CLVS1* | 13 | 371 | 37 | CLVS1 | clavesin 1 | Cytoplasm | other | |
| chr4 | *FAM53A* | 18 | 739 | 37 | FAM53A | family with sequence similarity 53 member A | Nucleus | other | |
| chr19 | *WDR88* | 16 | 523 | 37 | WDR88 | WD repeat domain 88 | Other | other | |
| chr1 | *FBLIM1* | 16 | 622 | 37 | FBLIM1 | filamin binding LIM protein 1 | Plasma Membrane | other | |
| chr15 | *TP53BP1* | 15 | 376 | 37 | TP53BP1 | tumor protein p53 binding protein | Nucleus | transcription regulator | |
| chr11 | *RP11-849H4.2* | 15 | 385 | 37 | | | | | |
| chr3 | *PPP2R3A* | 15 | 500 | 37 | PPP2R3A | protein phosphatase 2 regulatory subunit B''alpha | Nucleus | phosphatase | |
| chrX | *FGD1* | 17 | 608 | 37 | FGD1 | FYVE, RhoGEF and PH domain containing 1 | Cytoplasm | other | |
| chr2 | *PNPT1* | 15 | 507 | 37 | PNPT1 | polyribonucleotide nucleotidyltransferase 1 | Cytoplasm | enzyme | |
| chr10 | *WDR37* | 15 | 506 | 37 | WDR37 | WD repeat domain 37 | Other | other | |
| chr15 | *RP11-661P17.1* | 18 | 536 | 37 | | | | | |
| chr18 | *CABLES1* | 14 | 477 | 37 | CABLES1 | Cdk5 and Abl enzyme substrate 1 | Nucleus | other | |
| chr22 | *CLTCL1* | 14 | 498 | 37 | CLTCL1 | clathrin heavy chain like 1 | Plasma Membrane | other | |

| | | | | | | | | | |
|---|---|---|---|---|---|---|---|---|---|
| chr15 | *CASC4* | 12 | 353 | 37 | CASC4 | cancer susceptibility 4 | Cytoplasm | other | |
| chr16 | *SEZ6L2* | 16 | 569 | 37 | SEZ6L2 | seizure related 6 homolog like 2 | Cytoplasm | other | |
| chr22 | *MYO18B* | 15 | 445 | 37 | MYO18B | myosin XVIIIB | Cytoplasm | other | |
| chr12 | *HDAC7* | 18 | 1512 | 37 | HDAC7 | histone deacetylase 7 | Nucleus | transcription regulator | tributyrin, belinostat, pyroxamide |
| chr11 | *C2CD3* | 15 | 487 | 37 | C2CD3 | C2 domain containing 3 centriole elongation regulator | Cytoplasm | other | |
| chr1 | *RP1-35C21.2* | 19 | 441 | 37 | | | | | |
| chr12 | *CMKLR1* | 16 | 548 | 37 | CMKLR1 | chemerin chemokine-like receptor | Plasma Membrane | G-protein coupled recepto | |
| chr15 | *RP11-108K3.1* | 15 | 443 | 37 | | | | | |
| chr12 | *PXN* | 16 | 601 | 37 | PXN | paxillin | Cytoplasm | other | |
| chr20 | *GMEB2* | 17 | 1071 | 37 | GMEB2 | glucocorticoid modulatory element binding protein 2 | Nucleus | transcription regulator | |
| chr20 | *SLC23A2* | 13 | 425 | 37 | SLC23A2 | solute carrier family 23 member 2 | Plasma Membrane | transporter | |
| chr18 | *ESCO1* | 14 | 387 | 37 | ESCO1 | establishment of sister chromatid cohesion N-acetyltransferase 1 | Nucleus | enzyme | |
| chr14 | *DYNC1H1* | 13 | 417 | 37 | DYNC1H1 | dynein cytoplasmic 1 heavy chain | Cytoplasm | peptidase | |
| chr20 | *AAR2* | 15 | 722 | 37 | AAR2 | AAR2 splicing factor | Other | other | |
| chr8 | *UNC5D* | 17 | 329 | 37 | UNC5D | unc-5 netrin receptor D | Plasma Membrane | other | |
| chr10 | *SLC25A16* | 15 | 548 | 37 | SLC25A16 | solute carrier family 25 member | Cytoplasm | transporter | |
| chr11 | *TRIM5* | 16 | 576 | 37 | TRIM5 | tripartite motif containing 5 | Cytoplasm | enzyme | |
| chr7 | *BMPER* | 18 | 556 | 37 | BMPER | BMP binding endothelial regulator | Extracellular Space | other | |
| chr1 | *RP11-473A10.2* | 15 | 544 | 37 | | | | | |
| chr12 | *NTN4* | 17 | 424 | 37 | NTN4 | netrin 4 | Extracellular Space | other | |
| chr1 | *TRIM67* | 17 | 745 | 37 | TRIM67 | tripartite motif containing 67 | Cytoplasm | other | |
| chr9 | *UBE2R2* | 13 | 331 | 37 | UBE2R2 | ubiquitin conjugating enzyme E2 | Cytoplasm | enzyme | |
| chr12 | *EEA1* | 13 | 358 | 37 | EEA1 | early endosome antigen 1 | Cytoplasm | other | |
| chr16 | *NDE1* | 17 | 461 | 37 | NDE1 | nudE neurodevelopment protein 1 | Nucleus | other | |
| chr2 | *DPYSL5* | 13 | 472 | 37 | DPYSL5 | dihydropyrimidinase like 5 | Cytoplasm | enzyme | |
| chr1 | *IPP* | 13 | 354 | 37 | IPP | intracisternal A particle-promoted polypeptide | Cytoplasm | other | |
| chr17 | *DBF4B* | 15 | 499 | 37 | DBF4B | DBF4 zinc finger B | Nucleus | other | |
| chr15 | *PTPN9* | 14 | 386 | 37 | PTPN9 | protein tyrosine phosphatase non-receptor type 9 | Cytoplasm | phosphatase | |
| chr6 | *DAAM2* | 18 | 521 | 37 | DAAM2 | dishevelled associated activator of morphogenesis 2 | Cytoplasm | other | |
| chr1 | *LINC01036* | 15 | 460 | 37 | LINC01036 | long intergenic non-protein coding RNA 1036 | Other | other | |
| chr5 | *SLC27A6* | 16 | 400 | 37 | SLC27A6 | solute carrier family 27 member 6 | Plasma Membrane | transporter | |
| chr16 | *NOMO2* | 17 | 566 | 37 | MO1 (includes oth | NODAL modulator 1 | Plasma Membrane | other | |
| chr19 | *LPHN1* | 15 | 622 | 37 | ADGRL1 | adhesion G protein-coupled receptor L1 | Plasma Membrane | G-protein coupled recepto | |
| chr5 | *ZSWIM6* | 13 | 341 | 37 | ZSWIM6 | zinc finger SWIM-type containing | Extracellular Space | other | |
| chr12 | *APPL2* | 16 | 409 | 37 | APPL2 | adaptor protein, phosphotyrosine interacting with PH domain and leucine zipper 2 | Cytoplasm | other | |
| chr3 | *POLQ* | 16 | 492 | 37 | POLQ | DNA polymerase theta | Nucleus | enzyme | |
| chr9 | *SEC16A* | 18 | 1151 | 37 | SEC16A | SEC16 homolog A, endoplasmic reticulum export factor | Cytoplasm | phosphatase | |
| chr3 | *TEX264* | 16 | 665 | 37 | TEX264 | testis expressed 264, ER-phagy receptor | Extracellular Space | other | |
| chr6 | *PLEKHG1* | 14 | 368 | 37 | PLEKHG1 | pleckstrin homology and RhoGEF domain containing G1 | Extracellular Space | other | |
| chr3 | *PPARG* | 16 | 353 | 37 | PPARG | peroxisome proliferator activated receptor gamma | Nucleus | d-dependent nuclear rec | glimepiride/pioglitazone, bezafibrate, glimepiride/rosiglitazone, hydrochlorothiazide/telmisartan, pioglitazone, clopidogrel/telmisartan, balsalazide, icosapent, aspirin/dipyridamole/telmisartan, troglitazone, alogliptin/pioglitazone, telmisartan, inolitazone, farglitazar, sulfasalazine, rosiglitazone, nicotinic acid/pioglitazone |
| chr10 | *PIK3AP1* | 17 | 365 | 37 | PIK3AP1 | phosphoinositide-3-kinase adaptor protein 1 | Cytoplasm | other | |
| chr12 | *DRAM1* | 16 | 348 | 37 | DRAM1 | DNA damage regulated autophagy modulator 1 | Cytoplasm | other | |
| chr1 | *CEP170* | 13 | 406 | 37 | CEP170 | centrosomal protein 170 | Nucleus | other | |
| chr17 | *CACNG5* | 15 | 489 | 37 | CACNG5 | calcium voltage-gated channel auxiliary subunit gamma 5 | Plasma Membrane | ion channel | |
| chr16 | *RP11-345J4.8* | 15 | 413 | 37 | | | | | |
| chr12 | *PRPF40B* | 16 | 760 | 37 | PRPF40B | pre-mRNA processing factor 40 homolog B | Nucleus | other | |
| chr9 | *ENG* | 16 | 915 | 37 | ENG | endoglin | Plasma Membrane | transmembrane recepto | carotuximab |
| chr6 | *NUP153* | 14 | 366 | 37 | NUP153 | nucleoporin 153 | Nucleus | other | |
| chr4 | *RNF212* | 13 | 564 | 37 | RNF212 | ring finger protein 212 | Nucleus | enzyme | |
| chr9 | *BRINP1* | 16 | 509 | 37 | BRINP1 | BMP/retinoic acid inducible neural specific 1 | Nucleus | peptidase | |

| | | | | | | | | | |
|---|---|---|---|---|---|---|---|---|---|
| chrX | *ZNF81* | 15 | 404 | 37 | ZNF81 | zinc finger protein 81 | Nucleus | transcription regulator | |
| chr10 | *PPP3CB* | 14 | 620 | 37 | PPP3CB | protein phosphatase 3 catalytic subunit beta | Plasma Membrane | phosphatase | voclosporin, cyclosporin A, pimecrolimus, prednisone/tacrolimus, cyclosporin A/methotrexate, methylprednisolone/t |
| chr7 | *KCND2* | 15 | 347 | 37 | KCND2 | potassium voltage-gated channel subfamily D member 2 | Plasma Membrane | ion channel | dalfampridine |
| chr16 | *ZC3H7A* | 15 | 402 | 37 | ZC3H7A | zinc finger CCCH-type containing | Extracellular Space | other | |
| chr20 | *SLCO4A1* | 18 | 1712 | 37 | SLCO4A1 | solute carrier organic anion transporter family member 4A1 | Plasma Membrane | transporter | |
| chr1 | *DDR2* | 17 | 454 | 37 | DDR2 | discoidin domain receptor tyrosine kinase 2 | Plasma Membrane | kinase | blinatumomab/imatinib, imatinib/nilotinib, imatinib/inotuzumab ozogamicin, decitabine/imatinib, imatinib/sunitinib, imatinib, sitravatinib, imatinib/sirolimus, regorafenib, imatinib/sorafenib, dacomitinib |
| chr1 | *AGO1* | 16 | 417 | 37 | AGO1 | argonaute RISC component 1 | Cytoplasm | translation regulator | |
| chr15 | *ZNF592* | 16 | 636 | 37 | ZNF592 | zinc finger protein 592 | Nucleus | transcription regulator | |
| chr8 | *LZTS1* | 16 | 595 | 37 | LZTS1 | leucine zipper tumor suppressor | Nucleus | transcription regulator | |
| chr3 | *SNX4* | 14 | 408 | 37 | SNX4 | sorting nexin 4 | Cytoplasm | transporter | |
| chr15 | *ARIH1* | 14 | 348 | 37 | ARIH1 | ariadne RBR E3 ubiquitin protein ligase 1 | Cytoplasm | enzyme | |
| chr16 | *CES5A* | 16 | 378 | 37 | CES5A | carboxylesterase 5A | Other | enzyme | |
| chr1 | *PLEKHM2* | 16 | 529 | 37 | PLEKHM2 | pleckstrin homology and RUN domain containing M2 | Extracellular Space | other | |
| chr5 | *TNIP1* | 14 | 527 | 37 | TNIP1 | TNFAIP3 interacting protein 1 | Nucleus | other | |
| chr17 | *EZH1* | 16 | 382 | 37 | EZH1 | enhancer of zeste 1 polycomb repressive complex 2 subunit | Nucleus | enzyme | tazemetostat, valemetostat |
| chr8 | *TRPS1* | 18 | 462 | 37 | TRPS1 | transcriptional repressor GATA binding 1 | Nucleus | transcription regulator | |
| chr12 | *RACGAP1* | 16 | 736 | 37 | RACGAP1 | Rac GTPase activating protein 1 | Cytoplasm | transporter | |
| chr3 | *LRRFIP2* | 15 | 359 | 37 | LRRFIP2 | LRR binding FLII interacting | Cytoplasm | other | |
| chr17 | *SMYD4* | 16 | 502 | 37 | SMYD4 | SET and MYND domain | Other | other | |
| chr16 | *GPT2* | 16 | 645 | 37 | GPT2 | glutamic--pyruvic transaminase 2 | Cytoplasm | enzyme | |
| chr13 | *MTUS2* | 15 | 414 | 37 | MTUS2 | microtubule associated scaffold protein 2 | Nucleus | other | |
| chr15 | *CHP1* | 13 | 374 | 37 | CHP1 | calcineurin like EF-hand protein 1 | Cytoplasm | other | |
| chr12 | *TRHDE* | 15 | 396 | 37 | TRHDE | thyrotropin releasing hormone degrading enzyme | Plasma Membrane | peptidase | |
| chr5 | *MSH3* | 16 | 341 | 37 | MSH3 | mutS homolog 3 | Nucleus | enzyme | |
| chr19 | *TPM4* | 15 | 526 | 37 | TPM4 | tropomyosin 4 | Cytoplasm | other | |
| chr3 | *TKT* | 15 | 716 | 37 | TKT | transketolase | Cytoplasm | enzyme | |
| chr1 | *OTUD7B* | 15 | 537 | 37 | OTUD7B | OTU deubiquitinase 7B | Cytoplasm | peptidase | |
| chr8 | *RP11-30J20.1* | 17 | 314 | 37 | | | | | |
| chr7 | *GTF2I* | 15 | 555 | 37 | GTF2I | general transcription factor IIi | Nucleus | transcription regulator | |
| chr1 | *MFN2* | 15 | 568 | 37 | MFN2 | mitofusin 2 | Cytoplasm | enzyme | |
| chr8 | *RIMS2* | 13 | 369 | 37 | RIMS2 | regulating synaptic membrane exocytosis 2 | Plasma Membrane | other | |
| chr14 | *NUMB* | 13 | 357 | 37 | NUMB | NUMB endocytic adaptor protein | Plasma Membrane | other | |
| chr19 | *CERS4* | 15 | 651 | 37 | CERS4 | ceramide synthase 4 | Cytoplasm | transcription regulator | |
| chr15 | *CHRNA7* | 15 | 461 | 37 | CHRNA7 | cholinergic receptor nicotinic alpha 7 subunit | Plasma Membrane | transmembrane receptor | succinylcholine, lobeline, JNJ-39393406, rapacuronium, doxacurium, vecuronium, pipecuronium, metocurine, mivacurium, D-tubocurarine, nicotine, arecoline, pancuronium, atracurium, isoflurane, cisatracurium, rocuronium, amobarbital, mecamylamine, enflurane, ABT-089 |
| chr18 | *GREB1L* | 13 | 341 | 37 | GREB1L | GREB1 like retinoic acid receptor coactivator | Extracellular Space | other | |
| chr9 | *TLE1* | 15 | 390 | 37 | TLE1 | TLE family member 1, transcriptional corepressor | Nucleus | transcription regulator | |
| chr7 | *SCRN1* | 16 | 613 | 37 | SCRN1 | secernin 1 | Cytoplasm | other | |
| chr15 | *AVEN* | 13 | 332 | 36 | AVEN | apoptosis and caspase activation inhibitor | Nucleus | other | |
| chr15 | *CTDSPL2* | 13 | 343 | 36 | CTDSPL2 | CTD small phosphatase like 2 | Nucleus | phosphatase | |
| chr17 | *SYNRG* | 15 | 393 | 36 | SYNRG | synergin gamma | Cytoplasm | other | |
| chr19 | *AC006115.3* | 17 | 363 | 36 | | | | | |
| chr3 | *KIF9-AS1* | 16 | 376 | 36 | KIF9-AS1 | KIF9 antisense RNA 1 | Other | other | |
| chr7 | *ZP3* | 16 | 701 | 36 | ZP3 | zona pellucida glycoprotein 3 | Extracellular Space | other | |

| | | | | | | | | | |
|---|---|---|---|---|---|---|---|---|---|
| chr8 | *SLC20A2* | 13 | 405 | 36 | SLC20A2 | solute carrier family 20 member 2 | Plasma Membrane | transporter | |
| chr14 | *GALNT16* | 13 | 482 | 36 | GALNT16 | polypeptide N-acetylgalactosaminyltransferase | Cytoplasm | enzyme | |
| chr10 | *SH2D4B* | 15 | 459 | 36 | SH2D4B | SH2 domain containing 4B | Other | other | |
| chr1 | *ZYG11A* | 15 | 402 | 36 | ZYG11A | zyg-11 family member A, cell cycle regulator | Other | other | |
| chr2 | *LTBP1* | 12 | 353 | 36 | LTBP1 | latent transforming growth factor beta binding protein 1 | Extracellular Space | other | |
| chr19 | *AKAP8L* | 16 | 501 | 36 | AKAP8L | A-kinase anchoring protein 8 like | Nucleus | other | |
| chr1 | *USP24* | 15 | 348 | 36 | USP24 | ubiquitin specific peptidase 24 | Nucleus | peptidase | |
| chr7 | *RAPGEF5* | 17 | 430 | 36 | RAPGEF5 | Rap guanine nucleotide exchange factor 5 | Nucleus | other | |
| chr6 | *AKAP12* | 15 | 402 | 36 | AKAP12 | A-kinase anchoring protein 12 | Cytoplasm | transporter | |
| chr1 | *SLC6A9* | 17 | 743 | 36 | SLC6A9 | solute carrier family 6 member 9 | Plasma Membrane | transporter | |
| chr7 | *TNPO3* | 14 | 369 | 36 | TNPO3 | transportin 3 | Cytoplasm | other | |
| chr4 | *TBC1D1* | 14 | 395 | 36 | TBC1D1 | TBC1 domain family member 1 | Nucleus | other | |
| chr20 | *SMOX* | 16 | 521 | 36 | SMOX | spermine oxidase | Cytoplasm | enzyme | |
| chr10 | *RP11-556E13.1* | 16 | 552 | 36 | | | | | |
| chrX | *IL1RAPL2* | 14 | 370 | 36 | IL1RAPL2 | interleukin 1 receptor accessory protein like 2 | Plasma Membrane | transmembrane recepto | |
| chr3 | *AC144521.1* | 16 | 471 | 36 | | | | | |
| chr19 | *NLRP5* | 16 | 377 | 36 | NLRP5 | NLR family pyrin domain | Cytoplasm | other | |
| chr11 | *SLC22A10* | 17 | 407 | 36 | SLC22A10 | solute carrier family 22 member | Other | other | |
| chr17 | *HNF1B* | 15 | 436 | 36 | HNF1B | HNF1 homeobox B | Nucleus | transcription regulator | |
| chr11 | *INSC* | 16 | 401 | 36 | INSC | INSC spindle orientation adaptor protein | Plasma Membrane | other | |
| chr18 | *LOXHD1* | 15 | 450 | 36 | LOXHD1 | lipoxygenase homology domains | Extracellular Space | other | |
| chr15 | *DNAJC17* | 15 | 489 | 36 | DNAJC17 | DnaJ heat shock protein family (Hsp40) member C17 | Other | other | |
| chr6 | *IGF2R* | 13 | 482 | 36 | IGF2R | insulin like growth factor 2 | Plasma Membrane | transmembrane recepto | |
| chr17 | *AATK* | 18 | 650 | 36 | AATK | apoptosis associated tyrosine | Cytoplasm | kinase | |
| chr3 | *RAD18* | 14 | 469 | 36 | RAD18 | RAD18 E3 ubiquitin protein ligase | Nucleus | other | |
| chr17 | *RP11-60A24.3* | 16 | 418 | 36 | | | | | |
| chr9 | *NOTCH1* | 18 | 721 | 36 | NOTCH1 | notch receptor 1 | Plasma Membrane | transcription regulator | OMP-52M51 |
| chr11 | *NAP1L4* | 15 | 940 | 36 | NAP1L4 | nucleosome assembly protein 1 | Cytoplasm | other | |
| chr12 | *FAIM2* | 15 | 1174 | 36 | FAIM2 | Fas apoptotic inhibitory molecule | Plasma Membrane | other | |
| chr10 | *FAM107B* | 14 | 355 | 36 | FAM107B | family with sequence similarity 107 member B | Other | other | |
| chr4 | *RP11-371F15.3* | 14 | 308 | 36 | | | | | |
| chr1 | *TNN* | 16 | 548 | 36 | TNN | tenascin N | Plasma Membrane | other | |
| chr14 | *PRIMA1* | 14 | 533 | 36 | PRIMA1 | proline rich membrane anchor 1 | Plasma Membrane | other | |
| chr4 | *EVC2* | 13 | 440 | 36 | EVC2 | EvC ciliary complex subunit 2 | Extracellular Space | other | |
| chr11 | *RP11-707M1.1* | 16 | 698 | 36 | | | | | |
| chr10 | *CXCL12* | 18 | 759 | 36 | CXCL12 | C-X-C motif chemokine ligand 12 | Extracellular Space | cytokine | NOX-A12 |
| chr7 | *TTC26* | 13 | 472 | 36 | TTC26 | tetratricopeptide repeat domain | Extracellular Space | other | |
| chr20 | *RPN2* | 16 | 520 | 36 | RPN2 | ribophorin II | Cytoplasm | enzyme | |
| chr17 | *RP11-120M18.2* | 15 | 344 | 36 | | | | | |
| chr12 | *RP11-81H14.2* | 16 | 381 | 36 | | | | | |
| chr12 | *PLXNC1* | 16 | 355 | 36 | PLXNC1 | plexin C1 | Plasma Membrane | transmembrane recepto | |
| chr9 | *FAM120A* | 12 | 368 | 36 | FAM120A | family with sequence similarity | Cytoplasm | other | |
| chr2 | *GFPT1* | 13 | 329 | 36 | GFPT1 | glutamine--fructose-6-phosphate transaminase 1 | Cytoplasm | enzyme | |
| chr22 | *SMARCB1* | 13 | 488 | 36 | SMARCB1 | SWI/SNF related, matrix associated, actin dependent regulator of chromatin, subfamily | Nucleus | transcription regulator | |
| chr19 | *NOTCH3* | 17 | 599 | 36 | NOTCH3 | notch receptor 3 | Plasma Membrane | transcription regulator | |
| chr3 | *ARHGAP31* | 16 | 542 | 36 | ARHGAP31 | Rho GTPase activating protein 31 | Cytoplasm | other | |
| chr9 | *FAM73B* | 16 | 908 | 36 | MIGA2 | mitoguardin 2 | Plasma Membrane | other | |
| chr15 | *EPB42* | 14 | 432 | 36 | EPB42 | erythrocyte membrane protein band 4.2 | Plasma Membrane | transporter | |
| chr1 | *ZDHHC18* | 14 | 601 | 36 | ZDHHC18 | zinc finger DHHC-type containing | Cytoplasm | enzyme | |
| chr10 | *RGS10* | 15 | 572 | 36 | RGS10 | regulator of G protein signaling | Cytoplasm | enzyme | |
| chr10 | *ALOX5* | 17 | 1008 | 36 | ALOX5 | arachidonate 5-lipoxygenase | Cytoplasm | enzyme | diclofenac/omeprazole, diethylcarbamazine, benoxaprofen, balsalazide, diclofenac/misoprostol, ebselen, diclofenac, masoprocol, meclofenamic acid, mesalamine, nordihydroguaiaretic |
| chr19 | *ZNF331* | 15 | 486 | 36 | ZNF331 | zinc finger protein 331 | Nucleus | transcription regulator | |
| chr1 | *C1orf21* | 17 | 425 | 36 | C1orf21 | chromosome 1 open reading frame 21 | Other | other | |
| chr3 | *GNAI2* | 15 | 780 | 36 | GNAI2 | G protein subunit alpha i2 | Plasma Membrane | enzyme | |
| chr2 | *RTN4* | 14 | 489 | 36 | RTN4 | reticulon 4 | Cytoplasm | other | |
| chr10 | *REEP3* | 14 | 485 | 36 | REEP3 | receptor accessory protein 3 | Other | other | |
| chr17 | *MXRA7* | 15 | 661 | 36 | MXRA7 | matrix remodeling associated 7 | Extracellular Space | other | |
| chr3 | *GRAMD1C* | 16 | 484 | 36 | GRAMD1C | GRAM domain containing 1C | Other | other | |
| chr19 | *EMR3* | 17 | 302 | 36 | ADGRE3 | adhesion G protein-coupled receptor E3 | Plasma Membrane | -protein coupled recepto | |
| chr20 | *SLC12A5* | 16 | 804 | 36 | SLC12A5 | solute carrier family 12 member 5 | Plasma Membrane | transporter | bumetanide |
| chr15 | *RBPMS2* | 15 | 576 | 36 | RBPMS2 | RNA binding protein, mRNA processing factor 2 | Cytoplasm | other | |
| chr17 | *ZNF385C* | 15 | 598 | 36 | ZNF385C | zinc finger protein 385C | Other | other | |

| | | | | | | | | | |
|---|---|---|---|---|---|---|---|---|---|
| chr19 | *PIP5K1C* | 18 | 794 | 36 | PIP5K1C | phosphatidylinositol-4-phosphate 5-kinase type 1 gamma | Plasma Membrane | kinase | |
| chr1 | *NOTCH2* | 16 | 650 | 36 | NOTCH2 | notch receptor 2 | Plasma Membrane | transcription regulator | |
| chr10 | *KNDC1* | 18 | 743 | 36 | KNDC1 | kinase non-catalytic C-lobe domain containing 1 | Plasma Membrane | other | |
| chr3 | *LHFPL4* | 17 | 417 | 36 | LHFPL4 | LHFPL tetraspan subfamily member 4 | Plasma Membrane | other | |
| chr1 | *IGFN1* | 16 | 842 | 36 | IGFN1 | immunoglobulin like and fibronectin type III domain | Nucleus | other | |
| chr22 | *TMEM184B* | 13 | 601 | 36 | TMEM184B | transmembrane protein 184B | Other | other | |
| chr17 | *B9D1* | 15 | 705 | 36 | B9D1 | B9 domain containing 1 | Plasma Membrane | other | |
| chr17 | *CDC27* | 14 | 373 | 36 | CDC27 | cell division cycle 27 | Nucleus | other | |
| chr8 | *ELP3* | 14 | 370 | 36 | ELP3 | elongator acetyltransferase complex subunit 3 | Nucleus | enzyme | |
| chr3 | *RUVBL1* | 16 | 695 | 36 | RUVBL1 | RuvB like AAA ATPase 1 | Nucleus | transcription regulator | |
| chr1 | *CAPN8* | 16 | 603 | 36 | CAPN8 | calpain 8 | Cytoplasm | peptidase | |
| chr1 | *KCNT2* | 15 | 406 | 36 | KCNT2 | potassium sodium-activated channel subfamily T member 2 | Plasma Membrane | ion channel | |
| chr5 | *TBCA* | 15 | 408 | 36 | TBCA | tubulin folding cofactor A | Cytoplasm | other | |
| chr1 | *ZRANB2-AS2* | 16 | 345 | 36 | ZRANB2-AS2 | ZRANB2 antisense RNA 2 (head to head) | Other | other | |
| chr7 | *GTF2IRD2P1* | 14 | 481 | 36 | GTF2IRD2P1 | GTF2I repeat domain containing 2 pseudogene 1 | Other | other | |
| chr21 | *C2CD2* | 13 | 497 | 36 | C2CD2 | C2 calcium dependent domain containing 2 | Cytoplasm | other | |
| chrX | *PCDH11X* | 12 | 353 | 36 | PCDH11X | protocadherin 11 X-linked | Plasma Membrane | other | |
| chr15 | *C15orf26* | 18 | 375 | 36 | CFAP161 | cilia and flagella associated protein 161 | Other | other | |
| chr1 | *MDM4* | 16 | 510 | 36 | MDM4 | MDM4 regulator of p53 | Nucleus | transcription regulator | ALRN-6924 |
| chr2 | *GYPC* | 16 | 700 | 36 | GYPC | glycophorin C (Gerbich blood | Plasma Membrane | other | |
| chr13 | *MBNL2* | 17 | 481 | 36 | MBNL2 | muscleblind like splicing regulator | Nucleus | other | |
| chr1 | *PTBP2* | 13 | 348 | 36 | PTBP2 | polypyrimidine tract binding | Nucleus | other | |
| chr1 | *RCC2* | 15 | 580 | 36 | RCC2 | regulator of chromosome condensation 2 | Nucleus | other | |
| chr17 | *IKZF3* | 14 | 339 | 36 | IKZF3 | IKAROS family zinc finger 3 | Nucleus | transcription regulator | |
| chr3 | *ANKRD28* | 15 | 332 | 36 | ANKRD28 | ankyrin repeat domain 28 | Cytoplasm | other | |
| chr11 | *P2RY6* | 16 | 906 | 36 | P2RY6 | pyrimidinergic receptor P2Y6 | Plasma Membrane | G-protein coupled recepto | |
| chr3 | *ZDHHC3* | 15 | 503 | 36 | ZDHHC3 | zinc finger DHHC-type containing | Cytoplasm | peptidase | |
| chr1 | *IPO9* | 14 | 456 | 36 | IPO9 | importin 9 | Nucleus | transporter | |
| chrX | *CA5B* | 14 | 407 | 36 | CA5B | carbonic anhydrase 5B | Cytoplasm | enzyme | hydrochlorothiazide, trichloromethiazide, sulfacetamide, benzthiazide, topiramate, chlorothiazide, chlorthalidone, acetazolamide, methazolamide |
| chr13 | *TBC1D4* | 13 | 363 | 36 | TBC1D4 | TBC1 domain family member 4 | Cytoplasm | other | |
| chrX | *ZFX* | 13 | 356 | 36 | ZFX | zinc finger protein X-linked | Nucleus | transcription regulator | |
| chr11 | *ETS1* | 16 | 562 | 36 | ETS1 | ETS proto-oncogene 1, transcription factor | Nucleus | transcription regulator | |
| chr3 | *PCCB* | 15 | 420 | 36 | PCCB | propionyl-CoA carboxylase subunit beta | Cytoplasm | enzyme | |
| chr11 | *MARK2* | 15 | 450 | 36 | MARK2 | microtubule affinity regulating kinase 2 | Cytoplasm | kinase | |
| chr19 | *SUPT5H* | 15 | 559 | 36 | SUPT5H | SPT5 homolog, DSIF elongation factor subunit | Nucleus | transcription regulator | |
| chr22 | *GNB1L* | 15 | 657 | 36 | GNB1L | G protein subunit beta 1 like | Cytoplasm | other | |
| chr1 | *C1orf228* | 15 | 426 | 36 | ARMH1 | armadillo like helical domain containing 1 | Other | other | |
| chr17 | *NOS2* | 14 | 571 | 35 | NOS2 | nitric oxide synthase 2 | Cytoplasm | enzyme | pimagedine, N(G)-monomethyl-D-arginine, GW |
| chr11 | *ASRGL1* | 15 | 557 | 35 | ASRGL1 | asparaginase and isoaspartyl peptidase 1 | Cytoplasm | enzyme | |
| chr7 | *INTS4L1* | 15 | 332 | 35 | INTS4P1 | integrator complex subunit 4 pseudogene 1 | Other | other | |
| chr22 | *MEI1* | 13 | 318 | 35 | MEI1 | meiotic double-stranded break formation protein 1 | Other | other | |
| chr15 | *KLHL25* | 16 | 748 | 35 | KLHL25 | kelch like family member 25 | Cytoplasm | other | |
| chr17 | *CAMKK1* | 16 | 767 | 35 | CAMKK1 | calcium/calmodulin dependent protein kinase kinase 1 | Cytoplasm | kinase | |
| chr13 | *DNAJC3* | 14 | 496 | 35 | DNAJC3 | DnaJ heat shock protein family (Hsp40) member C3 | Cytoplasm | other | |
| chr5 | *CYSTM1* | 15 | 435 | 35 | CYSTM1 | cysteine rich transmembrane module containing 1 | Cytoplasm | other | |
| chr16 | *NLRC3* | 15 | 597 | 35 | NLRC3 | NLR family CARD domain containing 3 | Cytoplasm | other | |
| chr17 | *RECQL5* | 15 | 570 | 35 | RECQL5 | RecQ like helicase 5 | Nucleus | enzyme | |
| chr12 | *SLC2A13* | 15 | 510 | 35 | SLC2A13 | solute carrier family 2 member 13 | Plasma Membrane | transporter | |
| chr1 | *SSU72* | 16 | 662 | 35 | SSU72 | SSU72 homolog, RNA polymerase II CTD phosphatase | Other | phosphatase | |
| chr17 | *RHBDF2* | 15 | 706 | 35 | RHBDF2 | rhomboid 5 homolog 2 | Cytoplasm | other | |
| chr19 | *ERCC1* | 16 | 494 | 35 | ERCC1 | ERCC excision repair 1, endonuclease non-catalytic | Nucleus | enzyme | |
| chr19 | *COLGALT1* | 15 | 604 | 35 | COLGALT1 | collagen beta(1-O)galactosyltransferase 1 | Cytoplasm | enzyme | |
| chr17 | *WDR16* | 14 | 403 | 35 | CFAP52 | cilia and flagella associated | Cytoplasm | other | |

| | | | | | | | | | |
|---|---|---|---|---|---|---|---|---|---|
| chr9 | *FANCC* | 13 | 399 | 35 | FANCC | FA complementation group C | Nucleus | other | |
| chr15 | *IVD* | 15 | 526 | 35 | IVD | isovaleryl-CoA dehydrogenase | Cytoplasm | enzyme | |
| chr7 | *KCP* | 17 | 837 | 35 | KCP | kielin cysteine rich BMP regulator | Extracellular Space | other | |
| chr17 | *RP11-1055B8.7* | 18 | 840 | 35 | | | | | |
| chr16 | *ITFG3* | 16 | 608 | 35 | FAM234A | family with sequence similarity 234 member A | Plasma Membrane | other | |
| chr6 | *HIVEP2* | 15 | 401 | 35 | HIVEP2 | HIVEP zinc finger 2 | Nucleus | transcription regulator | |
| chr5 | *CTC-228N24.1* | 15 | 353 | 35 | | | | | |
| chr20 | *RALGAPB* | 14 | 415 | 35 | RALGAPB | Ral GTPase activating protein non-catalytic beta subunit | Other | other | |
| chr19 | *SYMPK* | 14 | 516 | 35 | SYMPK | symplekin | Cytoplasm | other | |
| chr2 | *MLPH* | 13 | 496 | 35 | MLPH | melanophilin | Cytoplasm | other | |
| chr14 | *CLMN* | 13 | 442 | 35 | CLMN | calmin | Cytoplasm | other | |
| chr6 | *STXBP5-AS1* | 16 | 344 | 35 | STXBP5-AS1 | STXBP5 antisense RNA 1 | Other | other | |
| chr2 | *REV1* | 12 | 389 | 35 | REV1 | REV1 DNA directed polymerase | Nucleus | enzyme | |
| chr18 | *PTPN2* | 14 | 386 | 35 | PTPN2 | protein tyrosine phosphatase non-receptor type 2 | Cytoplasm | phosphatase | |
| chr15 | *RP11-492D6.3* | 17 | 1792 | 35 | | | | | |
| chr12 | *SP1* | 15 | 747 | 35 | SP1 | Sp1 transcription factor | Nucleus | transcription regulator | |
| chr7 | *STYXL1* | 17 | 587 | 35 | STYXL1 | serine/threonine/tyrosine interacting like 1 | Cytoplasm | phosphatase | |
| chr2 | *TSGA10* | 15 | 416 | 35 | TSGA10 | testis specific 10 | Cytoplasm | other | |
| chr12 | *FRS2* | 12 | 371 | 35 | FRS2 | fibroblast growth factor receptor substrate 2 | Plasma Membrane | other | |
| chr9 | *SH3GL2* | 15 | 355 | 35 | SH3GL2 | SH3 domain containing GRB2 like 2, endophilin A1 | Plasma Membrane | enzyme | |
| chr7 | *KLHDC10* | 13 | 336 | 35 | KLHDC10 | kelch domain containing 10 | Nucleus | other | |
| chr16 | *BEAN1* | 15 | 496 | 35 | BEAN1 | brain expressed associated with NEDD4 1 | Other | other | |
| chr5 | *CSF1R* | 14 | 519 | 35 | CSF1R | colony stimulating factor 1 receptor | Plasma Membrane | kinase | [illegible] imatinib/inotuzumab ozogamicin, lapatinib/pazopanib, imatinib/sunitinib, sunitinib, regorafenib, pazopanib, quizartinib, dasatinib/nilotinib, SNDX-6352, ARRY-382, imatinib, nilotinib, SKI-G-801, pexidartinib, imatinib/nilotinib, NMS-03592088, c-FMS inhibitor, 3D185, emactuzumab, erdafitinib, DCC-3014, IMC-CS4, JNJ-40346527, bosutinib, BLZ-945, blinatumomab/imatini[illegible] |
| chr11 | *TMEM41B* | 14 | 438 | 35 | TMEM41B | transmembrane protein 41B | Cytoplasm | other | |
| chr10 | *LIPA* | 15 | 446 | 35 | LIPA | lipase A, lysosomal acid type | Cytoplasm | enzyme | |
| chr19 | *ZSWIM4* | 16 | 631 | 35 | ZSWIM4 | zinc finger SWIM-type containing | Other | other | |
| chr11 | *DNHD1* | 16 | 896 | 35 | DNHD1 | dynein heavy chain domain 1 | Cytoplasm | other | |
| chr16 | *SLC5A11* | 16 | 420 | 35 | SLC5A11 | solute carrier family 5 member 11 | Plasma Membrane | transporter | |
| chr16 | *ABCC12* | 15 | 507 | 35 | ABCC12 | ATP binding cassette subfamily C member 12 | Cytoplasm | transporter | |
| chr5 | *CWC27* | 14 | 321 | 35 | CWC27 | CWC27 spliceosome associated cyclophilin | Nucleus | enzyme | |
| chr11 | *POLA2* | 16 | 872 | 35 | POLA2 | DNA polymerase alpha 2, accessory subunit | Nucleus | enzyme | |
| chr1 | *DDAH1* | 16 | 556 | 35 | DDAH1 | dimethylarginine dimethylaminohydrolase 1 | Cytoplasm | enzyme | |
| chr1 | *DENND1B* | 15 | 405 | 35 | DENND1B | DENN domain containing 1B | Other | other | |
| chr12 | *ESPL1* | 16 | 936 | 35 | ESPL1 | extra spindle pole bodies like 1, separase | Nucleus | peptidase | |
| chr6 | *KDM1B* | 15 | 378 | 35 | KDM1B | lysine demethylase 1B | Nucleus | enzyme | |
| chr15 | *GRAMD2* | 15 | 556 | 35 | GRAMD2A | GRAM domain containing 2A | Other | other | |
| chr5 | *EPB41L4A* | 14 | 334 | 35 | EPB41L4A | erythrocyte membrane protein band 4.1 like 4A | Extracellular Space | other | |
| chr19 | *BTBD2* | 15 | 641 | 35 | BTBD2 | BTB domain containing 2 | Cytoplasm | other | |
| chr1 | *TRIM33* | 14 | 401 | 35 | TRIM33 | tripartite motif containing 33 | Nucleus | transcription regulator | |
| chrX | *CSF2RA* | 14 | 450 | 35 | CSF2RA | colony stimulating factor 2 receptor alpha subunit | Plasma Membrane | transmembrane receptor | rituximab/sargramostim, EPO/sargramostim, sargramostim, |
| chr10 | *PBLD* | 15 | 559 | 35 | PBLD | phenazine biosynthesis like protein domain containing | Cytoplasm | enzyme | |
| chr17 | *NLK* | 13 | 348 | 35 | NLK | nemo like kinase | Nucleus | kinase | |
| chr14 | *PACS2* | 17 | 776 | 35 | PACS2 | phosphofurin acidic cluster sorting protein 2 | Cytoplasm | other | |
| chr19 | *SULT2B1* | 14 | 466 | 35 | SULT2B1 | sulfotransferase family 2B | Cytoplasm | enzyme | |
| chr1 | *MEF2D* | 17 | 919 | 35 | MEF2D | myocyte enhancer factor 2D | Nucleus | transcription regulator | |
| chr16 | *RP11-505K9.4* | 16 | 859 | 35 | | | | | |
| chr7 | *GALNTL5* | 15 | 360 | 35 | GALNTL5 | polypeptide N-acetylgalactosaminyltransferase | Cytoplasm | enzyme | |

| | | | | | | | | | |
|---|---|---|---|---|---|---|---|---|---|
| chr9 | *AIF1L* | 14 | 805 | 35 | AIF1L | allograft inflammatory factor 1 like | Plasma Membrane | other | |
| chr17 | *AP2B1* | 11 | 329 | 35 | AP2B1 | adaptor related protein complex 2 subunit beta 1 | Plasma Membrane | transporter | |
| chr15 | *EDC3* | 14 | 372 | 35 | EDC3 | enhancer of mRNA decapping 3 | Cytoplasm | enzyme | |
| chr3 | *LPP* | 14 | 316 | 35 | LPP | LIM domain containing preferred translocation partner in lipoma | Nucleus | other | |
| chr5 | *WDR41* | 15 | 361 | 35 | WDR41 | WD repeat domain 41 | Cytoplasm | other | |
| chr1 | *NTRK1* | 16 | 831 | 35 | NTRK1 | neurotrophic receptor tyrosine kinase 1 | Plasma Membrane | kinase | larotrectinib, VMD-928, NOV1601, DS-6051b, belizatinib, lorlatinib, entrectinib, TRK tyrosine kinase inhibitor, repotrectinib, PLX7486, milciclib, |
| chr12 | *SLC35E3* | 13 | 321 | 35 | SLC35E3 | solute carrier family 35 member | Other | other | |
| chr17 | *RPS6KB1* | 15 | 383 | 35 | RPS6KB1 | ribosomal protein S6 kinase B1 | Cytoplasm | kinase | |
| chr12 | *MYO1H* | 14 | 360 | 35 | MYO1H | myosin IH | Other | other | |
| chr19 | *CATSPERD* | 12 | 442 | 35 | CATSPERD | cation channel sperm associated auxiliary subunit delta | Extracellular Space | other | |
| chr17 | *MYO15B* | 17 | 827 | 35 | MYO15B | myosin XVB | Other | other | |
| chr7 | *SH2B2* | 15 | 653 | 35 | SH2B2 | SH2B adaptor protein 2 | Cytoplasm | other | |
| chr14 | *C14orf166B* | 14 | 446 | 35 | LRRC74A | leucine rich repeat containing | Other | other | |
| chr13 | *ENOX1* | 13 | 293 | 35 | ENOX1 | ecto-NOX disulfide-thiol | Plasma Membrane | other | |
| chr2 | *EPCAM* | 14 | 480 | 35 | EPCAM | epithelial cell adhesion molecule | Plasma Membrane | other | catumaxomab, tucotuzumab celmoleukin, edrecolomab, solitomab, vicinium, adecatumumab |
| chr8 | *ADCY8* | 16 | 351 | 35 | ADCY8 | adenylate cyclase 8 | Plasma Membrane | enzyme | |
| chr16 | *ITGAM* | 16 | 481 | 35 | ITGAM | integrin subunit alpha M | Plasma Membrane | transmembrane recepto | abciximab, GB1275 |
| chr5 | *EFNA5* | 15 | 339 | 35 | EFNA5 | ephrin A5 | Plasma Membrane | kinase | |
| chr7 | *RP5-921G16.1* | 16 | 301 | 35 | | | | | |
| chr21 | *URB1* | 15 | 459 | 35 | URB1 | URB1 ribosome biogenesis | Nucleus | other | |
| chr17 | *TNS4* | 14 | 693 | 35 | TNS4 | tensin 4 | Cytoplasm | other | |
| chr1 | *RP5-1198O20.4* | 15 | 390 | 35 | | | | | |
| chr3 | *PRICKLE2* | 16 | 386 | 35 | PRICKLE2 | prickle planar cell polarity protein | Nucleus | other | |
| chr3 | *NR2C2* | 15 | 373 | 35 | NR2C2 | nuclear receptor subfamily 2 group C member 2 | Nucleus | d-dependent nuclear rec | |
| chr10 | *PPAPDC1A* | 15 | 362 | 35 | PLPP4 | phospholipid phosphatase 4 | Plasma Membrane | phosphatase | |
| chr17 | *GJC1* | 14 | 379 | 35 | GJC1 | gap junction protein gamma 1 | Plasma Membrane | ion channel | |
| chr5 | *FAM153A* | 14 | 541 | 35 | AM153A/FAM153 | family with sequence similarity 153 member B | Other | other | |
| chr1 | *CNTN2* | 15 | 872 | 35 | CNTN2 | contactin 2 | Plasma Membrane | other | |
| chr7 | *CREB3L2* | 14 | 481 | 35 | CREB3L2 | cAMP responsive element binding protein 3 like 2 | Nucleus | transcription regulator | |
| chr13 | *ADPRHL1* | 15 | 863 | 35 | ADPRHL1 | ADP-ribosylhydrolase like 1 | Other | enzyme | |
| chr1 | *BAI2* | 16 | 813 | 35 | ADGRB2 | adhesion G protein-coupled receptor B2 | Plasma Membrane | G-protein coupled recepto | |
| chr20 | *SLC24A3* | 14 | 331 | 35 | SLC24A3 | solute carrier family 24 member 3 | Plasma Membrane | transporter | |
| chr2 | *KCNG3* | 14 | 400 | 35 | KCNG3 | potassium voltage-gated channel modifier subfamily G member 3 | Cytoplasm | ion channel | |
| chr19 | *TGFB1* | 15 | 481 | 35 | TGFB1 | transforming growth factor beta 1 | Extracellular Space | growth factor | NIS793, MSB0011359C, fresolimumab, LY3200882, AVID200, YL-13027, |
| chr1 | *STK40* | 15 | 697 | 35 | STK40 | serine/threonine kinase 40 | Cytoplasm | kinase | |
| chr16 | *ANKS3* | 15 | 571 | 35 | ANKS3 | ankyrin repeat and sterile alpha motif domain containing 3 | Nucleus | transcription regulator | |
| chr19 | *DPP9* | 14 | 685 | 35 | DPP9 | dipeptidyl peptidase 9 | Cytoplasm | peptidase | |
| chr3 | *PLXNA1* | 19 | 1141 | 35 | PLXNA1 | plexin A1 | Plasma Membrane | transmembrane recepto | |
| chr11 | *CREB3L1* | 16 | 939 | 35 | CREB3L1 | cAMP responsive element binding protein 3 like 1 | Nucleus | transcription regulator | |
| chr17 | *SHMT1* | 14 | 515 | 35 | SHMT1 | serine hydroxymethyltransferase | Cytoplasm | enzyme | mimosine |
| chr7 | *ANKIB1* | 15 | 368 | 35 | ANKIB1 | ankyrin repeat and IBR domain containing 1 | Nucleus | transcription regulator | |
| chr11 | *METTL15* | 14 | 443 | 35 | METTL15 | methyltransferase like 15 | Other | other | |
| chr11 | *MAP6* | 15 | 522 | 35 | MAP6 | microtubule associated protein 6 | Cytoplasm | other | |
| chr5 | *DPYSL3* | 16 | 358 | 35 | DPYSL3 | dihydropyrimidinase like 3 | Cytoplasm | enzyme | |
| chr12 | *LINC00935* | 15 | 580 | 35 | TEX49 | testis expressed 49 | Other | other | |
| chr11 | *APLP2* | 14 | 525 | 35 | APLP2 | amyloid beta precursor like | Cytoplasm | other | |
| chr3 | *COLQ* | 15 | 406 | 35 | COLQ | collagen like tail subunit of asymmetric acetylcholinesterase | Extracellular Space | other | |
| chr1 | *RP11-84A19.4* | 14 | 329 | 35 | | | | | |
| chr19 | *LTBP4* | 17 | 786 | 35 | LTBP4 | latent transforming growth factor beta binding protein 4 | Extracellular Space | growth factor | |
| chr6 | *RP11-307P5.1* | 15 | 309 | 34 | | | | | |
| chr2 | *SMEK2* | 14 | 469 | 34 | PPP4R3B | protein phosphatase 4 regulatory subunit 3B | Nucleus | other | |
| chr16 | *LONP2* | 13 | 333 | 34 | LONP2 | lon peptidase 2, peroxisomal | Cytoplasm | peptidase | |
| chr17 | *LINC00469* | 14 | 513 | 34 | LINC00469 | long intergenic non-protein coding RNA 469 | Other | other | |
| chr18 | *MYO5B* | 14 | 385 | 34 | MYO5B | myosin VB | Cytoplasm | enzyme | |
| chr4 | *LIMCH1* | 13 | 340 | 34 | LIMCH1 | LIM and calponin homology domains 1 | Cytoplasm | other | |
| chr5 | *LINC01019* | 17 | 1157 | 34 | LINC01019 | long intergenic non-protein coding RNA 1019 | Other | other | |

| | | | | | | | | | |
|---|---|---|---|---|---|---|---|---|---|
| chr4 | *GRID2* | 13 | 318 | 34 | GRID2 | glutamate ionotropic receptor delta type subunit 2 | Plasma Membrane | ion channel | |
| chr7 | *TECPR1* | 16 | 829 | 34 | TECPR1 | tectonin beta-propeller repeat containing 1 | Cytoplasm | other | |
| chr14 | *MOK* | 13 | 466 | 34 | MOK | MOK protein kinase | Plasma Membrane | kinase | |
| chr9 | *SLC25A25* | 15 | 578 | 34 | SLC25A25 | solute carrier family 25 member | Cytoplasm | transporter | |
| chr17 | *MED24* | 15 | 600 | 34 | MED24 | mediator complex subunit 24 | Nucleus | transcription regulator | |
| chr9 | *RP11-65J3.1* | 14 | 839 | 34 | | | | | |
| chr22 | *TOM1* | 13 | 619 | 34 | TOM1 | target of myb1 membrane trafficking protein | Cytoplasm | transporter | |
| chr22 | *CSNK1E* | 14 | 531 | 34 | CSNK1E | casein kinase 1 epsilon | Cytoplasm | kinase | |
| chr6 | *KIAA0319* | 14 | 397 | 34 | KIAA0319 | KIAA0319 | Plasma Membrane | other | |
| chr15 | *RHCG* | 15 | 813 | 34 | RHCG | Rh family C glycoprotein | Plasma Membrane | transporter | |
| chr1 | *SOAT1* | 13 | 408 | 34 | SOAT1 | sterol O-acyltransferase 1 | Cytoplasm | enzyme | ezetimibe/simvastatin, pactimibe, ezetimibe, ezetimibe/fenofibrate, hesperetin, amlodipine/ezetimibe/losartan/rosuvastatin, ezetimibe/fluvastatin, |
| chr1 | *BTBD8* | 14 | 371 | 34 | BTBD8 | BTB domain containing 8 | Nucleus | other | |
| chr15 | *PLA2G4E* | 13 | 573 | 34 | PLA2G4E | phospholipase A2 group IVE | Cytoplasm | enzyme | |
| chr12 | *NR4A1* | 16 | 1068 | 34 | NR4A1 | nuclear receptor subfamily 4 group A member 1 | Nucleus | d-dependent nuclear rec | |
| chr1 | *SEMA4A* | 15 | 734 | 34 | SEMA4A | semaphorin 4A | Plasma Membrane | other | |
| chr6 | *PXDC1* | 15 | 678 | 34 | PXDC1 | PX domain containing 1 | Other | other | |
| chr17 | *CARD14* | 16 | 692 | 34 | CARD14 | caspase recruitment domain family member 14 | Cytoplasm | other | |
| chr1 | *GPR137B* | 15 | 687 | 34 | GPR137B | G protein-coupled receptor 137B | Plasma Membrane | other | |
| chr3 | *ZXDC* | 14 | 782 | 34 | ZXDC | ZXD family zinc finger C | Other | transcription regulator | |
| chr17 | *MBTD1* | 14 | 323 | 34 | MBTD1 | mbt domain containing 1 | Other | other | |
| chr22 | *EMID1* | 14 | 548 | 34 | EMID1 | EMI domain containing 1 | Extracellular Space | other | |
| chr18 | *CTDP1* | 15 | 567 | 34 | CTDP1 | CTD phosphatase subunit 1 | Nucleus | phosphatase | |
| chr7 | *SLC25A13* | 13 | 420 | 34 | SLC25A13 | solute carrier family 25 member | Cytoplasm | transporter | |
| chr12 | *MON2* | 12 | 327 | 34 | MON2 | MON2 homolog, regulator of endosome-to-Golgi trafficking | Cytoplasm | other | |
| chr4 | *LINC01060* | 12 | 315 | 34 | LINC01060 | long intergenic non-protein coding RNA 1060 | Other | other | |
| chr1 | *RC3H1* | 13 | 363 | 34 | RC3H1 | ring finger and CCCH-type | Cytoplasm | enzyme | |
| chr5 | *NR2F1-AS1* | 14 | 392 | 34 | NR2F1-AS1 | NR2F1 antisense RNA 1 | Other | other | |
| chr17 | *DNAI2* | 15 | 545 | 34 | DNAI2 | dynein axonemal intermediate chain 2 | Cytoplasm | other | |
| chr17 | *STAT5A* | 15 | 620 | 34 | STAT5A | signal transducer and activator of transcription 5A | Nucleus | transcription regulator | |
| chr16 | *PRMT7* | 16 | 589 | 34 | PRMT7 | protein arginine | Cytoplasm | enzyme | |
| chrX | *FAM156A* | 15 | 623 | 34 | AM156A/FAM156 | family with sequence similarity 156 member A | Nucleus | other | |
| chr18 | *RP11-838N2.5* | 13 | 576 | 34 | | | | | |
| chr5 | *GUSBP3* | 12 | 710 | 34 | GUSBP3 | GUSB pseudogene 3 | Other | other | |
| chr15 | *CASC5* | 13 | 377 | 34 | KNL1 | kinetochore scaffold 1 | Nucleus | other | |
| chr1 | *TCEA3* | 16 | 506 | 34 | TCEA3 | transcription elongation factor A3 | Nucleus | transcription regulator | |
| chr5 | *MARCH6;* | 16 | 1062 | 34 | | | | | |
| chr3 | *TFP1* | 14 | 526 | 34 | INHCAP | inhibitor of carbonic anhydrase pseudogene | Other | other | |
| chr18 | *YES1* | 13 | 296 | 34 | YES1 | YES proto-oncogene 1, Src family tyrosine kinase | Cytoplasm | kinase | cytarabine/dasatinib/dexamethasone/methotrexate, blinatumomab/dasatinib, cytarabine/dasatinib/filgrastim/fludarabine phosphate, dasatinib/6-mercaptopurine/methotrexate, dasatinib/nilotinib, cytarabine/dasatinib/filgrastim/fludarabine phosphate/idarubicin, dasatinib/prednisone, dasatinib/erlotinib, cytarabine/dasatinib, dasatinib/dexamethasone, dasatinib/inotuzumab ozogamicin, [illegible] |
| chr22 | *RANGAP1* | 14 | 509 | 34 | RANGAP1 | Ran GTPase activating protein 1 | Nucleus | other | |
| chr1 | *SPAG17* | 14 | 437 | 34 | SPAG17 | sperm associated antigen 17 | Cytoplasm | other | |
| chr1 | *LHX4* | 16 | 855 | 34 | LHX4 | LIM homeobox 4 | Nucleus | transcription regulator | |
| chr10 | *ARL3* | 14 | 426 | 34 | ARL3 | ADP ribosylation factor like | Cytoplasm | enzyme | |
| chr3 | *PIGX* | 12 | 352 | 34 | PIGX | phosphatidylinositol glycan anchor biosynthesis class X | Cytoplasm | other | |
| chr3 | *NAALADL2* | 13 | 337 | 34 | NAALADL2 | N-acetylated alpha-linked acidic dipeptidase like 2 | Nucleus | other | |
| chr10 | *LARP4B* | 12 | 381 | 34 | LARP4B | La ribonucleoprotein domain family member 4B | Cytoplasm | other | |

| | | | | | | | | | |
|---|---|---|---|---|---|---|---|---|---|
| chr16 | *PHLPP2* | 12 | 364 | 34 | PHLPP2 | PH domain and leucine rich repeat protein phosphatase 2 | Cytoplasm | enzyme | |
| chr3 | *PLS1* | 14 | 341 | 34 | PLS1 | plastin 1 | Plasma Membrane | other | |
| chr1 | *NFYC* | 15 | 413 | 34 | NFYC | nuclear transcription factor Y subunit gamma | Nucleus | transcription regulator | |
| chr1 | *ZBTB17* | 14 | 600 | 34 | ZBTB17 | zinc finger and BTB domain containing 17 | Nucleus | transcription regulator | |
| chr19 | *ATCAY* | 13 | 479 | 34 | ATCAY | ATCAY kinesin light chain interacting caytaxin | Plasma Membrane | other | |
| chr12 | *P2RX7* | 16 | 400 | 34 | P2RX7 | purinergic receptor P2X 7 | Plasma Membrane | ion channel | |
| chr2 | *SLC8A1* | 15 | 414 | 34 | SLC8A1 | solute carrier family 8 member A1 | Plasma Membrane | transporter | icosapent |
| chr10 | *SPOCK2* | 15 | 1092 | 34 | SPOCK2 | SPARC (osteonectin), cwcv and kazal like domains proteoglycan 2 | Extracellular Space | other | |
| chr5 | *CTD-2532K18.1* | 15 | 445 | 34 | | | | | |
| chr7 | *CUL1* | 12 | 418 | 34 | CUL1 | cullin 1 | Nucleus | enzyme | |
| chr12 | *DTX1* | 15 | 675 | 34 | DTX1 | deltex E3 ubiquitin ligase 1 | Nucleus | transcription regulator | |
| chr12 | *PLEKHA5* | 13 | 296 | 34 | PLEKHA5 | pleckstrin homology domain containing A5 | Cytoplasm | other | |
| chr5 | *FAF2* | 14 | 412 | 34 | FAF2 | Fas associated factor family member 2 | Cytoplasm | other | |
| chr2 | *RP11-444A22.1* | 15 | 321 | 34 | | | | | |
| chr22 | *WNT7B* | 16 | 835 | 34 | WNT7B | Wnt family member 7B | Extracellular Space | other | |
| chr9 | *PPP6C* | 13 | 459 | 34 | PPP6C | protein phosphatase 6 catalytic subunit | Nucleus | phosphatase | |
| chr17 | *KIF1C* | 15 | 587 | 34 | KIF1C | kinesin family member 1C | Cytoplasm | other | |
| chr20 | *COMMD7* | 14 | 503 | 34 | COMMD7 | COMM domain containing 7 | Cytoplasm | other | |
| chr7 | *PTPN12* | 13 | 369 | 34 | PTPN12 | protein tyrosine phosphatase non-receptor type 12 | Cytoplasm | phosphatase | |
| chr8 | *ZNF251* | 14 | 500 | 34 | ZNF251 | zinc finger protein 251 | Nucleus | transcription regulator | |
| chr12 | *PTPRQ* | 13 | 393 | 34 | PTPRQ | protein tyrosine phosphatase receptor type Q | Extracellular Space | other | |
| chr2 | *HECW2* | 11 | 312 | 34 | HECW2 | HECT, C2 and WW domain containing E3 ubiquitin protein ligase 2 | Extracellular Space | enzyme | |
| chr1 | *TMEM39B* | 14 | 449 | 34 | TMEM39B | transmembrane protein 39B | Cytoplasm | other | |
| chr19 | *PHLDB3* | 15 | 485 | 34 | PHLDB3 | pleckstrin homology like domain family B member 3 | Other | other | |
| chr19 | *EML2* | 14 | 450 | 34 | EML2 | EMAP like 2 | Cytoplasm | other | |
| chr1 | *NOL9* | 15 | 597 | 34 | NOL9 | nucleolar protein 9 | Nucleus | kinase | |
| chr15 | *GALK2* | 12 | 326 | 34 | GALK2 | galactokinase 2 | Cytoplasm | kinase | |
| chr3 | *CMTM8* | 16 | 338 | 34 | CMTM8 | CKLF like MARVEL transmembrane domain | Extracellular Space | cytokine | |
| chr1 | *RP11-475O6.1* | 15 | 327 | 34 | | | | | |
| chr15 | *LIPC* | 14 | 423 | 34 | LIPC | lipase C, hepatic type | Extracellular Space | enzyme | |
| chr9 | *DDX31* | 15 | 568 | 34 | DDX31 | DEAD-box helicase 31 | Nucleus | enzyme | |
| chr1 | *MYOM3* | 14 | 682 | 34 | MYOM3 | myomesin 3 | Other | other | |
| chr16 | *UBN1* | 13 | 489 | 34 | UBN1 | ubinuclein 1 | Nucleus | transcription regulator | |
| chr3 | *GK5* | 13 | 404 | 34 | GK5 | glycerol kinase 5 | Other | kinase | |
| chr7 | *NCAPG2* | 12 | 537 | 34 | NCAPG2 | non-SMC condensin II complex subunit G2 | Nucleus | other | |
| chr1 | *CEP85* | 14 | 373 | 34 | CEP85 | centrosomal protein 85 | Nucleus | other | |
| chr6 | *CAPN11* | 13 | 507 | 34 | CAPN11 | calpain 11 | Cytoplasm | peptidase | |
| chr15 | *ETFA* | 13 | 435 | 34 | ETFA | electron transfer flavoprotein subunit alpha | Cytoplasm | transporter | |
| chr17 | *IGF2BP1* | 15 | 448 | 34 | IGF2BP1 | insulin like growth factor 2 mRNA binding protein 1 | Cytoplasm | translation regulator | |
| chr1 | *COL11A1* | 13 | 378 | 34 | COL11A1 | collagen type XI alpha 1 chain | Extracellular Space | other | collagenase clostridium |
| chr12 | *B4GALNT3* | 12 | 521 | 34 | B4GALNT3 | beta-1,4-N-acetyl-galactosaminyltransferase 3 | Cytoplasm | enzyme | |
| chr11 | *PDE3B* | 15 | 389 | 34 | PDE3B | phosphodiesterase 3B | Cytoplasm | enzyme | cilostazol, pentoxifylline, tolbutamide, amrinone, theophylline, nitroglycerin, |
| chr2 | *PASK* | 14 | 589 | 34 | PASK | PAS domain containing serine/threonine kinase | Cytoplasm | kinase | |
| chr7 | *LAT2* | 15 | 616 | 34 | LAT2 | linker for activation of T cells family member 2 | Plasma Membrane | other | |
| chr22 | *GGT3P* | 14 | 523 | 34 | GGT3P | gamma-glutamyltransferase 3 pseudogene | Extracellular Space | other | |
| chr8 | *BIN3* | 16 | 607 | 34 | BIN3 | bridging integrator 3 | Cytoplasm | other | |
| chr19 | *TECR* | 15 | 639 | 34 | TECR | trans-2,3-enoyl-CoA reductase | Plasma Membrane | enzyme | |
| chr11 | *ACER3* | 14 | 379 | 34 | ACER3 | alkaline ceramidase 3 | Cytoplasm | enzyme | |
| chr10 | *DNAJC12* | 14 | 548 | 34 | DNAJC12 | DnaJ heat shock protein family (Hsp40) member C12 | Cytoplasm | other | |
| chr11 | *RP11-820L6.1* | 14 | 356 | 34 | | | | | |
| chr16 | *ABCC11* | 15 | 412 | 34 | ABCC11 | ATP binding cassette subfamily C member 11 | Plasma Membrane | transporter | |
| chr12 | *UTP20* | 12 | 310 | 34 | UTP20 | UTP20 small subunit processome component | Nucleus | other | |
| chr16 | *SLC38A8* | 15 | 791 | 34 | SLC38A8 | solute carrier family 38 member 8 | Other | other | |
| chr5 | *C5orf17* | 14 | 715 | 34 | C5orf17 | chromosome 5 open reading frame 17 (putative) | Other | other | |
| chr7 | *PMS2CL* | 16 | 603 | 34 | PMS2CL | PMS2 C-terminal like | Other | other | |
| chr7 | *STEAP2-AS1* | 15 | 422 | 34 | STEAP2-AS1 | STEAP2 antisense RNA 1 | Other | other | |
| chr17 | *GLP2R* | 16 | 435 | 34 | GLP2R | glucagon like peptide 2 receptor | Plasma Membrane | ;-protein coupled recepto | teduglutide, |
| chr11 | *RNF214* | 13 | 361 | 34 | RNF214 | ring finger protein 214 | Other | other | |

| | | | | | | | | | |
|---|---|---|---|---|---|---|---|---|---|
| chr11 | *UEVLD* | 13 | 298 | 34 | UEVLD | UEV and lactate/malate dehyrogenase domains | Cytoplasm | enzyme | |
| chr11 | *CDON* | 15 | 325 | 34 | CDON | cell adhesion associated, oncogene regulated | Plasma Membrane | other | |
| chr9 | *RP11-509J21.1* | 14 | 344 | 34 | | | | | |
| chr10 | *CACUL1* | 13 | 404 | 34 | CACUL1 | CDK2 associated cullin domain 1 | Other | other | |
| chr3 | *TPRXL* | 16 | 399 | 34 | TPRXL | tetrapeptide repeat homeobox | Other | other | |
| chr20 | *CSE1L* | 13 | 433 | 34 | CSE1L | chromosome segregation 1 like | Nucleus | transporter | |
| chr1 | *LRIG2* | 14 | 403 | 34 | LRIG2 | leucine rich repeats and immunoglobulin like domains 2 | Cytoplasm | other | |
| chr3 | *BBX* | 14 | 372 | 34 | BBX | BBX high mobility group box domain containing | Nucleus | transcription regulator | |
| chr7 | *EIF3B* | 13 | 994 | 34 | EIF3B | eukaryotic translation initiation factor 3 subunit B | Cytoplasm | translation regulator | |
| chr1 | *EPHX1* | 15 | 711 | 34 | EPHX1 | epoxide hydrolase 1 | Cytoplasm | peptidase | |
| chr7 | *WDR86* | 15 | 696 | 34 | WDR86 | WD repeat domain 86 | Other | other | |
| chr16 | *AFG3L1P* | 16 | 619 | 34 | AFG3L1P | AFG3 like matrix AAA peptidase subunit 1, pseudogene | Cytoplasm | other | |
| chr5 | *CTC-419K13.1* | 16 | 326 | 34 | | | | | |
| chr1 | *DRAXIN* | 15 | 651 | 34 | DRAXIN | dorsal inhibitory axon guidance protein | Extracellular Space | other | |
| chr17 | *PIPOX* | 15 | 409 | 34 | PIPOX | pipecolic acid and sarcosine | Cytoplasm | enzyme | |
| chr2 | *AC017099.3* | 15 | 530 | 34 | | | | | |
| chr12 | *ANKLE2* | 14 | 670 | 34 | ANKLE2 | ankyrin repeat and LEM domain containing 2 | Nucleus | transcription regulator | |
| chr7 | *HIBADH* | 12 | 363 | 34 | HIBADH | 3-hydroxyisobutyrate | Cytoplasm | enzyme | |
| chr15 | *RAB27A* | 13 | 332 | 33 | RAB27A | RAB27A, member RAS oncogene family | Cytoplasm | enzyme | |
| chr1 | *IFFO2* | 14 | 634 | 33 | IFFO2 | intermediate filament family | Other | other | |
| chr5 | *NDUFS4* | 11 | 345 | 33 | NDUFS4 | NADH:ubiquinone oxidoreductase subunit S4 | Cytoplasm | enzyme | |
| chr11 | *NUMA1* | 14 | 430 | 33 | NUMA1 | nuclear mitotic apparatus protein | Nucleus | other | |
| chr19 | *DOT1L* | 16 | 706 | 33 | DOT1L | DOT1 like histone lysine methyltransferase | Nucleus | phosphatase | pinometostat |
| chrX | *RP11-622K12.1* | 17 | 1369 | 33 | | | | | |
| chr17 | *ACSF2* | 15 | 525 | 33 | ACSF2 | acyl-CoA synthetase family member 2 | Cytoplasm | enzyme | |
| chr1 | *TMEM206* | 14 | 449 | 33 | PACC1 | proton activated chloride channel | Plasma Membrane | ion channel | |
| chr6 | *AGPAT4* | 12 | 508 | 33 | AGPAT4 | 1-acylglycerol-3-phosphate O-acyltransferase 4 | Cytoplasm | enzyme | |
| chr12 | *ATP6V0A2* | 14 | 410 | 33 | ATP6V0A2 | ATPase H+ transporting V0 subunit a2 | Cytoplasm | transporter | |
| chr6 | *TFEB* | 14 | 660 | 33 | TFEB | transcription factor EB | Nucleus | transcription regulator | |
| chr1 | *H6PD* | 14 | 622 | 33 | H6PD | hexose-6-phosphate dehydrogenase/glucose 1-dehydrogenase | Cytoplasm | enzyme | |
| chr17 | *SMTNL2* | 14 | 638 | 33 | SMTNL2 | smoothelin like 2 | Other | other | |
| chr5 | *SAR1B* | 14 | 391 | 33 | SAR1B | secretion associated Ras related GTPase 1B | Cytoplasm | enzyme | |
| chr3 | *SSUH2* | 16 | 673 | 33 | SSUH2 | ssu-2 homolog | Cytoplasm | other | |
| chr8 | *XPO7* | 12 | 334 | 33 | XPO7 | exportin 7 | Nucleus | transporter | |
| chr19 | *NLRP12* | 14 | 458 | 33 | NLRP12 | NLR family pyrin domain containing 12 | Cytoplasm | other | |
| chr15 | *SPPL2A* | 13 | 300 | 33 | SPPL2A | signal peptide peptidase like 2A | Plasma Membrane | peptidase | |
| chr9 | *RNF38* | 12 | 275 | 33 | RNF38 | ring finger protein 38 | Nucleus | enzyme | |
| chr14 | *SAMD4A* | 13 | 355 | 33 | SAMD4A | sterile alpha motif domain containing 4A | Cytoplasm | translation regulator | |
| chr16 | *CCDC113* | 13 | 324 | 33 | CCDC113 | coiled-coil domain containing 113 | Cytoplasm | other | |
| chr7 | *CFTR* | 12 | 337 | 33 | CFTR | CF transmembrane conductance regulator | Plasma Membrane | ion channel | ivacaftor/tezacaftor, tezacaftor, lumacaftor, ivacaftor/lumacaftor, |
| chr3 | *COPG1* | 15 | 788 | 33 | COPG1 | coatomer protein complex subunit gamma 1 | Cytoplasm | transporter | |
| chr9 | *GNA14* | 14 | 403 | 33 | GNA14 | G protein subunit alpha 14 | Plasma Membrane | enzyme | |
| chr19 | *HOOK2* | 15 | 611 | 33 | HOOK2 | hook microtubule tethering | Cytoplasm | other | |
| chr11 | *C11orf73* | 13 | 487 | 33 | HIKESHI | heat shock protein nuclear import factor hikeshi | Cytoplasm | transporter | |
| chr8 | *GATA4* | 14 | 688 | 33 | GATA4 | GATA binding protein 4 | Nucleus | transcription regulator | |
| chr5 | *RAB3C* | 15 | 324 | 33 | RAB3C | RAB3C, member RAS oncogene family | Cytoplasm | enzyme | |
| chr12 | *CCDC41* | 14 | 317 | 33 | CEP83 | centrosomal protein 83 | Extracellular Space | other | |
| chr11 | *AP002954.4* | 14 | 401 | 33 | | | | | |
| chr5 | *DMXL1* | 13 | 310 | 33 | DMXL1 | Dmx like 1 | Extracellular Space | other | |
| chr4 | *WWC2* | 11 | 288 | 33 | WWC2 | WW and C2 domain containing 2 | Cytoplasm | other | |
| chr11 | *ENPP7P8* | 15 | 373 | 33 | ENPP7P8 | ectonucleotide pyrophosphatase/phosphodiesterase 7 pseudogene 8 | Other | other | |
| chr12 | *LRRC43* | 15 | 508 | 33 | LRRC43 | leucine rich repeat containing 43 | Other | other | |
| chr3 | *CEP70* | 13 | 564 | 33 | CEP70 | centrosomal protein 70 | Cytoplasm | other | |
| chr7 | *MLXIPL* | 14 | 525 | 33 | MLXIPL | MLX interacting protein like | Nucleus | transcription regulator | |
| chr11 | *C11orf30* | 15 | 508 | 33 | EMSY | EMSY transcriptional repressor, BRCA2 interacting | Nucleus | other | |
| chr17 | *GDPD1* | 12 | 337 | 33 | GDPD1 | glycerophosphodiester phosphodiesterase domain containing 1 | Other | enzyme | |
| chr2 | *THSD7B* | 13 | 327 | 33 | THSD7B | thrombospondin type 1 domain containing 7B | Other | other | |
| chrX | *MECP2* | 12 | 422 | 33 | MECP2 | methyl-CpG binding protein 2 | Nucleus | transcription regulator | |

| | | | | | | | | | |
|---|---|---|---|---|---|---|---|---|---|
| chr2 | *TMEM163* | 13 | 375 | 33 | TMEM163 | transmembrane protein 163 | Cytoplasm | other | |
| chr11 | *CD5* | 14 | 777 | 33 | CD5 | CD5 molecule | Plasma Membrane | transmembrane receptor | |
| chr4 | *ADD1* | 12 | 443 | 33 | ADD1 | adducin 1 | Cytoplasm | other | |
| chr11 | *SYT9* | 16 | 509 | 33 | SYT9 | synaptotagmin 9 | Plasma Membrane | transporter | |
| chr4 | *KIAA0922* | 12 | 344 | 33 | TMEM131L | transmembrane 131 like | Other | other | |
| chr5 | *C5orf56* | 13 | 515 | 33 | IRF1-AS1 | IRF1 antisense RNA 1 | Other | other | |
| chr10 | *FAM21A* | 13 | 468 | 33 | ASHC2A/WASHC | WASH complex subunit 2A | Cytoplasm | other | |
| chr20 | *POFUT1* | 14 | 557 | 33 | POFUT1 | protein O-fucosyltransferase 1 | Cytoplasm | enzyme | |
| chr2 | *EFHD1* | 13 | 378 | 33 | EFHD1 | EF-hand domain family member | Cytoplasm | other | |
| chr11 | *EHD1* | 15 | 756 | 33 | EHD1 | EH domain containing 1 | Cytoplasm | other | |
| chr11 | *IGSF9B* | 16 | 673 | 33 | IGSF9B | immunoglobulin superfamily member 9B | Plasma Membrane | other | |
| chr16 | *SPG7* | 15 | 602 | 33 | SPG7 | SPG7 matrix AAA peptidase subunit, paraplegin | Cytoplasm | peptidase | |
| chr21 | *EVA1C* | 14 | 351 | 33 | EVA1C | eva-1 homolog C | Other | other | |
| chr11 | *AP002856.5* | 15 | 541 | 33 | | | | | |
| chr10 | *PI4K2A* | 13 | 362 | 33 | PI4K2A | phosphatidylinositol 4-kinase type 2 alpha | Cytoplasm | kinase | |
| chr12 | *RASAL1* | 15 | 542 | 33 | RASAL1 | RAS protein activator like 1 | Cytoplasm | other | |
| chr16 | *SLC7A5* | 15 | 748 | 33 | SLC7A5 | solute carrier family 7 member 5 | Plasma Membrane | transporter | |
| chr1 | *ADCY10* | 14 | 333 | 33 | ADCY10 | adenylate cyclase 10 | Cytoplasm | enzyme | |
| chr16 | *NPIPB5* | 15 | 619 | 33 | PB4 (includes oth | nuclear pore complex interacting protein family member B4 | Nucleus | other | |
| chr20 | *KIF3B* | 16 | 418 | 33 | KIF3B | kinesin family member 3B | Cytoplasm | transporter | |
| chr10 | *MXI1* | 13 | 496 | 33 | MXI1 | MAX interactor 1, dimerization | Nucleus | transcription regulator | |
| chr17 | *NSF* | 12 | 319 | 33 | NSF | N-ethylmaleimide sensitive factor, vesicle fusing ATPase | Cytoplasm | transporter | |
| chr10 | *FAM171A1* | 12 | 346 | 33 | FAM171A1 | family with sequence similarity 171 member A1 | Cytoplasm | other | |
| chr3 | *ARMC8* | 11 | 372 | 33 | ARMC8 | armadillo repeat containing 8 | Nucleus | other | |
| chr22 | *IL2RB* | 12 | 582 | 33 | IL2RB | interleukin 2 receptor subunit beta | Plasma Membrane | transmembrane receptor | denileukin diftitox, aldesleukin, bempegaldesleukin, monoclonal antibody Mik-Beta-1, |
| chr2 | *KIAA1841* | 13 | 345 | 33 | KIAA1841 | KIAA1841 | Other | other | |
| chr7 | *SNX13* | 13 | 455 | 33 | SNX13 | sorting nexin 13 | Cytoplasm | transporter | |
| chr1 | *KDM5B* | 13 | 388 | 33 | KDM5B | lysine demethylase 5B | Nucleus | transcription regulator | |
| chr15 | *IQGAP1* | 14 | 587 | 33 | IQGAP1 | IQ motif containing GTPase activating protein 1 | Cytoplasm | other | |
| chr3 | *SYN2* | 13 | 393 | 33 | SYN2 | synapsin II | Plasma Membrane | other | |
| chr6 | *C6orf89* | 12 | 371 | 33 | C6orf89 | chromosome 6 open reading frame 89 | Cytoplasm | other | |
| chr9 | *ADAMTSL2* | 15 | 830 | 33 | ADAMTSL2 | ADAMTS like 2 | Extracellular Space | other | |
| chr2 | *XPO1* | 13 | 378 | 33 | XPO1 | exportin 1 | Nucleus | transporter | eltanexor, leptomycin B, selinexor, SL-801 |
| chr14 | *RP11-260M19.2* | 14 | 711 | 33 | | | | | |
| chr19 | *ZNF83* | 14 | 301 | 33 | ZNF83 | zinc finger protein 83 | Nucleus | transcription regulator | |
| chr12 | *BAZ2A* | 14 | 414 | 33 | BAZ2A | bromodomain adjacent to zinc finger domain 2A | Nucleus | transcription regulator | |
| chr20 | *SS18L1* | 16 | 1214 | 33 | SS18L1 | SS18L1 subunit of BAF chromatin remodeling complex | Nucleus | transcription regulator | |
| chr20 | *RRBP1* | 14 | 558 | 33 | RRBP1 | ribosome binding protein 1 | Cytoplasm | other | |
| chr8 | *CASC8* | 14 | 399 | 33 | CASC8 | cancer susceptibility 8 | Other | other | |
| chr12 | *COL2A1* | 15 | 1141 | 33 | COL2A1 | collagen type II alpha 1 chain | Extracellular Space | other | collagenase clostridium |
| chr2 | *GPR55* | 12 | 531 | 33 | GPR55 | G protein-coupled receptor 55 | Plasma Membrane | G-protein coupled recepto | |
| chr12 | *BCAT1* | 15 | 439 | 33 | BCAT1 | branched chain amino acid transaminase 1 | Cytoplasm | enzyme | |
| chrX | *STAG2* | 12 | 316 | 33 | STAG2 | stromal antigen 2 | Nucleus | transcription regulator | |
| chr17 | *TIMP2* | 14 | 499 | 33 | TIMP2 | TIMP metallopeptidase inhibitor 2 | Extracellular Space | other | |
| chr3 | *TMEM44* | 12 | 451 | 33 | TMEM44 | transmembrane protein 44 | Other | other | |
| chr15 | *LINC00923* | 15 | 482 | 33 | LINC00923 | long intergenic non-protein coding RNA 923 | Other | other | |
| chr15 | *AC069368.3* | 14 | 495 | 33 | | | | | |
| chr10 | *EGR2* | 16 | 505 | 33 | EGR2 | early growth response 2 | Nucleus | transcription regulator | |
| chr9 | *NXNL2* | 14 | 408 | 33 | NXNL2 | nucleoredoxin like 2 | Other | transporter | |
| chr12 | *KMT2D* | 17 | 1141 | 33 | KMT2D | lysine methyltransferase 2D | Nucleus | transcription regulator | |
| chr12 | *RP11-511B23.2* | 15 | 305 | 33 | | | | | |
| chr9 | *MELK* | 12 | 292 | 33 | MELK | maternal embryonic leucine zipper kinase | Cytoplasm | kinase | OTS167 |
| chr19 | *TSHZ3* | 13 | 475 | 33 | TSHZ3 | teashirt zinc finger homeobox 3 | Nucleus | transcription regulator | |
| chr8 | *MATN2* | 12 | 373 | 33 | MATN2 | matrilin 2 | Extracellular Space | other | |
| chr1 | *HMGCL* | 14 | 379 | 33 | HMGCL | 3-hydroxy-3-methylglutaryl-CoA | Cytoplasm | enzyme | |
| chr6 | *PXT1* | 12 | 380 | 33 | PXT1 | peroxisomal testis enriched | Cytoplasm | other | |
| chrX | *RP11-258C19.5* | 14 | 429 | 33 | | | | | |
| chr19 | *CATSPERG* | 14 | 539 | 33 | CATSPERG | cation channel sperm associated auxiliary subunit gamma | Extracellular Space | other | |
| chr5 | *GUSBP9* | 14 | 705 | 33 | GUSBP9 | GUSB pseudogene 9 | Other | other | |
| chr10 | *PFKP* | 13 | 526 | 33 | PFKP | phosphofructokinase, platelet | Cytoplasm | kinase | |
| chr10 | *VWA2* | 15 | 440 | 33 | VWA2 | von Willebrand factor A domain containing 2 | Extracellular Space | other | |
| chr12 | *NCAPD2* | 12 | 415 | 33 | NCAPD2 | non-SMC condensin I complex subunit D2 | Nucleus | other | |
| chr13 | *TMEM255B* | 16 | 724 | 33 | TMEM255B | transmembrane protein 255B | Other | other | |
| chr16 | *USP31* | 15 | 364 | 33 | USP31 | ubiquitin specific peptidase 31 | Nucleus | peptidase | |
| chr13 | *GRTP1* | 16 | 586 | 33 | GRTP1 | growth hormone regulated TBC protein 1 | Other | other | |
| chr12 | *BIN2* | 14 | 633 | 33 | BIN2 | bridging integrator 2 | Plasma Membrane | other | |

| | | | | | | | | | |
|---|---|---|---|---|---|---|---|---|---|
| chr4 | *GAK* | 16 | 650 | 33 | GAK | cyclin G associated kinase | Nucleus | kinase | |
| chr1 | *MAPKAPK2* | 13 | 695 | 33 | MAPKAPK2 | MAPK activated protein kinase 2 | Nucleus | kinase | cdc25C phosphatase (211-221) |
| chr22 | *LRP5L* | 14 | 579 | 33 | LRP5L | LDL receptor related protein 5 | Other | other | |
| chr8 | *TNFRSF10A* | 13 | 443 | 33 | TNFRSF10A | TNF receptor superfamily member 10a | Plasma Membrane | transmembrane recepto | dulanermin, trm-1, mapatumumab |
| chr11 | *ELP4* | 14 | 416 | 33 | ELP4 | elongator acetyltransferase complex subunit 4 | Cytoplasm | other | |
| chr5 | *SDHA* | 15 | 1288 | 33 | SDHA | succinate dehydrogenase complex flavoprotein subunit A | Cytoplasm | enzyme | |
| chr1 | *SLC35E2B* | 15 | 735 | 33 | SLC35E2B | solute carrier family 35 member | Other | other | |
| chr3 | *SUCLG2* | 14 | 331 | 33 | SUCLG2 | succinate-CoA ligase GDP-forming beta subunit | Cytoplasm | enzyme | |
| chr3 | *PTPLB* | 13 | 423 | 33 | HACD2 | 3-hydroxyacyl-CoA dehydratase | Cytoplasm | phosphatase | |
| chr5 | *CAST* | 13 | 328 | 33 | CAST | calpastatin | Cytoplasm | peptidase | |
| chr3 | *SLC25A20* | 14 | 375 | 33 | SLC25A20 | solute carrier family 25 member | Cytoplasm | transporter | |
| chr15 | *ADAM10* | 11 | 278 | 33 | ADAM10 | ADAM metallopeptidase domain | Plasma Membrane | peptidase | |
| chr7 | *GALNT11* | 14 | 351 | 33 | GALNT11 | polypeptide N-acetylgalactosaminyltransferase | Cytoplasm | enzyme | |
| chr2 | *RALB* | 15 | 637 | 32 | RALB | RAS like proto-oncogene B | Cytoplasm | enzyme | |
| chr11 | *DRD2* | 14 | 492 | 32 | DRD2 | dopamine receptor D2 | Plasma Membrane | -protein coupled recepto | thiethylperazine, SLV-314, remoxipride, haloperidol/octreotide, haloperidol decanoate, loxapine, clozapine, sulpiride, brexpiprazole, dihydroergotamine, droperidol, benperidol, meloxicam, fluspirilene, carbidopa/entacapone/levodopa, apomorphine, fluphenazine decanoate, rotigotine, paliperidone, levodopa, risperidone, droperidol/fentanyl, [illegible] |
| chr19 | *SHANK1* | 15 | 597 | 32 | SHANK1 | SH3 and multiple ankyrin repeat domains 1 | Cytoplasm | other | |
| chr1 | *RCOR3* | 12 | 378 | 32 | RCOR3 | REST corepressor 3 | Nucleus | transcription regulator | |
| chr17 | *NUP85* | 13 | 442 | 32 | NUP85 | nucleoporin 85 | Cytoplasm | other | |
| chr5 | *AC022431.2* | 15 | 360 | 32 | | | | | |
| chr11 | *GLB1L2* | 15 | 628 | 32 | GLB1L2 | galactosidase beta 1 like 2 | Other | other | |
| chr11 | *B3GAT1* | 14 | 691 | 32 | B3GAT1 | beta-1,3-glucuronyltransferase 1 | Cytoplasm | enzyme | |
| chr19 | *PTPRH* | 13 | 593 | 32 | PTPRH | protein tyrosine phosphatase receptor type H | Plasma Membrane | phosphatase | |
| chr20 | *B4GALT5* | 15 | 439 | 32 | B4GALT5 | beta-1,4-galactosyltransferase 5 | Cytoplasm | enzyme | |
| chrX | *TEX11* | 12 | 342 | 32 | TEX11 | testis expressed 11 | Nucleus | other | |
| chr7 | *PMS2P5* | 14 | 604 | 32 | PMS2P5/PMS2P | PMS1 homolog 2, mismatch repair system component | Other | other | |
| chr17 | *AIPL1* | 14 | 553 | 32 | AIPL1 | aryl hydrocarbon receptor interacting protein like 1 | Nucleus | other | |
| chr4 | *UBE2K* | 13 | 372 | 32 | UBE2K | ubiquitin conjugating enzyme E2 | Cytoplasm | transcription regulator | |
| chr11 | *USP35* | 15 | 815 | 32 | USP35 | ubiquitin specific peptidase 35 | Extracellular Space | peptidase | |
| chr1 | *EPHA10* | 15 | 531 | 32 | EPHA10 | EPH receptor A10 | Plasma Membrane | transmembrane recepto | |
| chr10 | *CCSER2* | 12 | 312 | 32 | CCSER2 | coiled-coil serine rich protein 2 | Other | other | |
| chr3 | *MITF* | 13 | 334 | 32 | MITF | melanocyte inducing transcription factor | Nucleus | transcription regulator | |
| chr11 | *CTTN* | 15 | 652 | 32 | CTTN | cortactin | Plasma Membrane | other | |
| chr11 | *MICALCL* | 14 | 553 | 32 | MICALCL | MICAL C-terminal like | Cytoplasm | other | |
| chr11 | *ALG8* | 14 | 455 | 32 | ALG8 | ALG8 alpha-1,3- | Cytoplasm | enzyme | |
| chr1 | *CASP9* | 14 | 407 | 32 | CASP9 | caspase 9 | Cytoplasm | peptidase | caspase-9 inhibitor |
| chr6 | *RCAN2* | 15 | 319 | 32 | RCAN2 | regulator of calcineurin 2 | Other | other | |
| chr15 | *HCN4* | 15 | 561 | 32 | HCN4 | hyperpolarization activated cyclic nucleotide gated potassium channel 4 | Plasma Membrane | ion channel | |
| chr5 | *JAKMIP2* | 14 | 355 | 32 | JAKMIP2 | janus kinase and microtubule interacting protein 2 | Extracellular Space | other | |
| chr19 | *CTD-2562J15.6* | 14 | 480 | 32 | | | | | |
| chr12 | *DDX11* | 14 | 798 | 32 | DDX11 | DEAD/H-box helicase 11 | Cytoplasm | enzyme | |
| chr7 | *NRCAM* | 14 | 360 | 32 | NRCAM | neuronal cell adhesion molecule | Plasma Membrane | other | |
| chrX | *CXorf58* | 13 | 334 | 32 | CXorf58 | chromosome X open reading frame 58 | Other | other | |
| chr1 | *RIMS3* | 14 | 470 | 32 | RIMS3 | regulating synaptic membrane exocytosis 3 | Plasma Membrane | other | |
| chr21 | *BACE2* | 12 | 384 | 32 | BACE2 | beta-secretase 2 | Cytoplasm | peptidase | |
| chr10 | *VPS26A* | 12 | 510 | 32 | VPS26A | VPS26, retromer complex component A | Cytoplasm | transporter | |
| chr10 | *SFXN2* | 13 | 403 | 32 | SFXN2 | sideroflexin 2 | Cytoplasm | transporter | |
| chr11 | *TECTA* | 15 | 462 | 32 | TECTA | tectorin alpha | Extracellular Space | enzyme | |
| chr1 | *GOLPH3L* | 13 | 367 | 32 | GOLPH3L | golgi phosphoprotein 3 like | Cytoplasm | other | |
| chr1 | *UBXN11* | 13 | 566 | 32 | UBXN11 | UBX domain protein 11 | Cytoplasm | other | |

| | | | | | | | | | |
|---|---|---|---|---|---|---|---|---|---|
| chr4 | *FSTL5* | 10 | 288 | 32 | FSTL5 | follistatin like 5 | Extracellular Space | other | |
| chr11 | *DPP3* | 14 | 937 | 32 | DPP3 | dipeptidyl peptidase 3 | Cytoplasm | peptidase | |
| chr2 | *PSME4* | 12 | 444 | 32 | PSME4 | proteasome activator subunit 4 | Cytoplasm | other | |
| chr22 | *KLHL22* | 14 | 463 | 32 | KLHL22 | kelch like family member 22 | Cytoplasm | other | |
| chr12 | *NINJ2* | 11 | 430 | 32 | NINJ2 | ninjurin 2 | Plasma Membrane | other | |
| chr2 | *ELMOD3* | 14 | 377 | 32 | ELMOD3 | ELMO domain containing 3 | Other | other | |
| chr4 | *SHROOM3* | 12 | 326 | 32 | SHROOM3 | shroom family member 3 | Cytoplasm | other | |
| chr2 | *TTL* | 13 | 529 | 32 | TTL | tubulin tyrosine ligase | Cytoplasm | enzyme | |
| chr11 | *INTS4* | 15 | 473 | 32 | INTS4 | integrator complex subunit 4 | Nucleus | other | |
| chr2 | *DIRC3* | 13 | 370 | 32 | DIRC3 | disrupted in renal carcinoma 3 | Other | other | |
| chr6 | *RP1-209A6.1* | 14 | 298 | 32 | | | | | |
| chr17 | *KCNH4* | 14 | 658 | 32 | KCNH4 | potassium voltage-gated channel subfamily H member 4 | Plasma Membrane | ion channel | |
| chr9 | *RP11-399H11.3* | 14 | 783 | 32 | | | | | |
| chr1 | *PADI4* | 13 | 475 | 32 | PADI4 | peptidyl arginine deiminase 4 | Cytoplasm | enzyme | |
| chr5 | *HOMER1* | 11 | 306 | 32 | HOMER1 | homer scaffold protein 1 | Plasma Membrane | other | |
| chr7 | *COL28A1* | 14 | 408 | 32 | COL28A1 | collagen type XXVIII alpha 1 | Extracellular Space | other | |
| chr22 | *DGCR2* | 12 | 502 | 32 | DGCR2 | DiGeorge syndrome critical region gene 2 | Plasma Membrane | other | |
| chr2 | *CNGA3* | 14 | 638 | 32 | CNGA3 | cyclic nucleotide gated channel subunit alpha 3 | Plasma Membrane | ion channel | |
| chr16 | *CBFB* | 13 | 326 | 32 | CBFB | core-binding factor subunit beta | Nucleus | transcription regulator | |
| chr9 | *CDC14B* | 12 | 315 | 32 | CDC14B | cell division cycle 14B | Nucleus | phosphatase | |
| chr7 | *ZNF815P* | 15 | 549 | 32 | ZNF815P | zinc finger protein 815, | Other | other | |
| chr17 | *RP11-115K3.2* | 14 | 408 | 32 | | | | | |
| chr19 | *SPINT2* | 14 | 498 | 32 | SPINT2 | serine peptidase inhibitor, Kunitz type 2 | Extracellular Space | other | |
| chr13 | *SLC7A1* | 11 | 410 | 32 | SLC7A1 | solute carrier family 7 member 1 | Plasma Membrane | transporter | |
| chr1 | *SZRD1* | 13 | 391 | 32 | SZRD1 | SUZ RNA binding domain containing 1 | Other | other | |
| chr1 | *NMNAT1* | 13 | 336 | 32 | NMNAT1 | nicotinamide nucleotide adenylyltransferase 1 | Nucleus | enzyme | |
| chr13 | *RAB20* | 15 | 558 | 32 | RAB20 | RAB20, member RAS oncogene family | Cytoplasm | enzyme | |
| chr8 | *RP11-115J16.1* | 14 | 383 | 32 | | | | | |
| chr16 | *GAN* | 14 | 364 | 32 | GAN | gigaxonin | Cytoplasm | other | |
| chr8 | *PARP10* | 14 | 583 | 32 | PARP10 | poly(ADP-ribose) polymerase family member 10 | Cytoplasm | enzyme | |
| chr7 | *GTF2IRD1P1* | 13 | 549 | 32 | GTF2IRD1P1 | GTF2I repeat domain containing 1 pseudogene 1 | Other | other | |
| chr17 | *QRICH2* | 13 | 514 | 32 | QRICH2 | glutamine rich 2 | Extracellular Space | other | |
| chr12 | *ASIC1* | 14 | 850 | 32 | ASIC1 | acid sensing ion channel subunit 1 | Plasma Membrane | ion channel | diclofenac/omeprazole, amiloride, diclofenac/misoprostol, diclofenac, amiloride/hydrochlorothiazide |
| chr12 | *TUBA1C* | 15 | 535 | 32 | TUBA1C | tubulin alpha 1c | Cytoplasm | other | docetaxel/gemcitabine/vincristine, cyclophosphamide/epirubicin/5-fluorouracil/paclitaxel, bevacizumab/paclitaxel/topotecan, colchicine/probenecid, bevacizumab/docetaxel/prednisone/thalidomide/zoledronic acid, gemcitabine/paclitaxel, L-asparaginase/daunorubicin/dexamethasone/vincristine, prednisone/vincristine, docetaxel/5-fluorouracil, vinflunine, L-[illegible] |
| chr10 | *ZCCHC24* | 14 | 587 | 32 | ZCCHC24 | zinc finger CCHC-type containing | Other | other | |
| chr7 | *PMS2* | 13 | 579 | 32 | PMS2 | PMS1 homolog 2, mismatch repair system component | Nucleus | enzyme | |
| chr15 | *RP11-648K4.2* | 15 | 409 | 32 | | | | | |
| chr5 | *COMMD10* | 15 | 297 | 32 | COMMD10 | COMM domain containing 10 | Nucleus | other | |
| chr15 | *ANPEP* | 14 | 859 | 32 | ANPEP | alanyl aminopeptidase, membrane | Plasma Membrane | peptidase | CNGRC peptide-TNF alpha conjugate |
| chr22 | *UBE2L3* | 13 | 427 | 32 | UBE2L3 | ubiquitin conjugating enzyme E2 | Nucleus | enzyme | |
| chr14 | *RP11-187O7.3* | 13 | 303 | 32 | | | | | |
| chr12 | *TMBIM6* | 16 | 583 | 32 | TMBIM6 | transmembrane BAX inhibitor motif containing 6 | Nucleus | other | |
| chr1 | *ADC* | 13 | 458 | 32 | AZIN2 | antizyme inhibitor 2 | Cytoplasm | enzyme | |
| chr19 | *HNRNPUL1* | 14 | 404 | 32 | HNRNPUL1 | heterogeneous nuclear ribonucleoprotein U like 1 | Nucleus | other | |
| chr16 | *RP11-1166P10.1* | 14 | 565 | 32 | | | | | |
| chr1 | *TMCO1* | 12 | 376 | 32 | TMCO1 | transmembrane and coiled-coil domains 1 | Cytoplasm | ion channel | |
| chr16 | *PSKH1* | 14 | 515 | 32 | PSKH1 | protein serine kinase H1 | Nucleus | kinase | |

| | | | | | | | | | |
|---|---|---|---|---|---|---|---|---|---|
| chr16 | *SIAH1* | 14 | 329 | 32 | SIAH1 | siah E3 ubiquitin protein ligase 1 | Nucleus | enzyme | |
| chr5 | *SYNPO* | 14 | 485 | 32 | SYNPO | synaptopodin | Cytoplasm | other | |
| chr17 | *EFTUD2* | 14 | 397 | 32 | EFTUD2 | elongation factor Tu GTP binding domain containing 2 | Nucleus | enzyme | |
| chr6 | *NHSL1* | 11 | 298 | 32 | NHSL1 | NHS like 1 | Other | other | |
| chr19 | *PDE4C* | 13 | 555 | 32 | PDE4C | phosphodiesterase 4C | Cytoplasm | enzyme | caffeine, pentoxifylline, arofylline, tetomilast, tolbutamide, anagrelide, theophylline, milrinone, cilomilast, roflumilast, ketotifen, L-826,141, L 869298, ibudilast, |
| chr17 | *SLC26A11* | 14 | 655 | 32 | SLC26A11 | solute carrier family 26 member | Cytoplasm | transporter | |
| chr17 | *DCAKD* | 13 | 691 | 32 | DCAKD | dephospho-CoA kinase domain containing | Cytoplasm | other | |
| chr15 | *RP13-98N21.2* | 14 | 637 | 32 | | | | | |
| chr11 | *MMP26* | 15 | 440 | 32 | MMP26 | matrix metallopeptidase 26 | Extracellular Space | peptidase | marimastat |
| chr5 | *SUB1* | 16 | 983 | 32 | SUB1 | SUB1 regulator of transcription | Nucleus | transcription regulator | |
| chr22 | *ATXN10* | 12 | 328 | 32 | ATXN10 | ataxin 10 | Cytoplasm | other | |
| chr16 | *CCNF* | 14 | 580 | 32 | CCNF | cyclin F | Nucleus | other | |
| chr6 | *LRRC1* | 12 | 298 | 32 | LRRC1 | leucine rich repeat containing 1 | Cytoplasm | other | |
| chr17 | *TTC25* | 13 | 454 | 32 | TTC25 | tetratricopeptide repeat domain | Cytoplasm | other | |
| chr7 | *ACTR3C* | 12 | 412 | 32 | ACTR3C | actin related protein 3C | Cytoplasm | other | |
| chr5 | *PLCXD3* | 14 | 921 | 32 | PLCXD3 | phosphatidylinositol specific phospholipase C X domain containing 3 | Plasma Membrane | other | |
| chr1 | *CACHD1* | 14 | 310 | 32 | CACHD1 | cache domain containing 1 | Other | other | |
| chr22 | *XPNPEP3* | 10 | 293 | 32 | XPNPEP3 | X-prolyl aminopeptidase 3 | Cytoplasm | peptidase | |
| chr5 | *PLEKHG4B* | 17 | 2431 | 32 | PLEKHG4B | pleckstrin homology and RhoGEF domain containing G4B | Other | other | |
| chrX | *PDK3* | 12 | 288 | 32 | PDK3 | pyruvate dehydrogenase kinase 3 | Cytoplasm | kinase | |
| chr2 | *LRP1B* | 11 | 292 | 32 | LRP1B | LDL receptor related protein 1B | Plasma Membrane | transmembrane receptor | |
| chr22 | *FLJ27365* | 15 | 838 | 32 | | | | | |
| chr19 | *HPN-AS1* | 15 | 662 | 32 | HPN-AS1 | HPN antisense RNA 1 | Other | other | |
| chr12 | *WNT5B* | 12 | 361 | 32 | WNT5B | Wnt family member 5B | Extracellular Space | other | |
| chr1 | *MFSD4* | 14 | 861 | 32 | MFSD4A | major facilitator superfamily domain containing 4A | Other | other | |
| chr7 | *BRI3* | 14 | 626 | 32 | BRI3 | brain protein I3 | Cytoplasm | other | |
| chr14 | *SYNE3* | 12 | 479 | 32 | SYNE3 | spectrin repeat containing nuclear envelope family member 3 | Nucleus | other | |
| chr7 | *CCDC146* | 14 | 408 | 32 | CCDC146 | coiled-coil domain containing 146 | Cytoplasm | other | |
| chr17 | *THRA* | 13 | 641 | 32 | THRA | thyroid hormone receptor alpha | Nucleus | d-dependent nuclear rec | dextrothyroxine, amiodarone, levothyroxine/l-triiodothyronine, denosumab/levothyroxine, L-triiodothyronine, 3,5-diiodothyropropionic |
| chr3 | *ABHD6* | 14 | 340 | 32 | ABHD6 | abhydrolase domain containing 6 | Cytoplasm | enzyme | |
| chr3 | *C3orf67* | 14 | 345 | 32 | C3orf67 | chromosome 3 open reading frame 67 | Other | other | |
| chr15 | *AAGAB* | 11 | 296 | 32 | AAGAB | alpha and gamma adaptin binding protein | Cytoplasm | other | |
| chr8 | *WRN* | 11 | 292 | 32 | WRN | WRN RecQ like helicase | Nucleus | enzyme | |
| chr5 | *S100Z* | 14 | 381 | 32 | S100Z | S100 calcium binding protein Z | Other | other | |
| chr20 | *TPD52L2* | 15 | 870 | 32 | TPD52L2 | TPD52 like 2 | Cytoplasm | other | |
| chr15 | *UNC13C* | 12 | 334 | 32 | UNC13C | unc-13 homolog C | Cytoplasm | other | |
| chr16 | *GLYR1* | 14 | 404 | 32 | GLYR1 | glyoxylate reductase 1 homolog | Nucleus | other | |
| chr3 | *ZNF589* | 14 | 372 | 32 | ZNF589 | zinc finger protein 589 | Nucleus | transcription regulator | |
| chr5 | *SLC25A48* | 15 | 517 | 32 | SLC25A48 | solute carrier family 25 member | Other | other | |
| chr19 | *SLC7A9* | 13 | 454 | 32 | SLC7A9 | solute carrier family 7 member 9 | Plasma Membrane | transporter | |
| chr19 | *ANKRD24* | 13 | 512 | 32 | ANKRD24 | ankyrin repeat domain 24 | Other | other | |
| chr19 | *CTC-490E21.12* | 14 | 333 | 31 | | | | | |
| chr4 | *SCD5* | 12 | 334 | 31 | SCD5 | stearoyl-CoA desaturase 5 | Cytoplasm | enzyme | |
| chr2 | *MROH2A* | 12 | 503 | 31 | MROH2A | maestro heat like repeat family member 2A | Other | other | |
| chr11 | *CHID1* | 15 | 734 | 31 | CHID1 | chitinase domain containing 1 | Extracellular Space | other | |
| chr15 | *EXD1* | 12 | 310 | 31 | EXD1 | exonuclease 3'-5' domain containing 1 | Other | other | |
| chr1 | *LCK* | 14 | 365 | 31 | LCK | LCK proto-oncogene, Src family tyrosine kinase | Cytoplasm | kinase | nintedanib, dasatinib/inotuzumab ozogamicin, JNJ-26483327, dasatinib, dacomitinib, pazopanib, blinatumomab/dasatinib, crizotinib/pazopanib, docetaxel/nintedanib, lapatinib/pazopanib, |
| chr16 | *POLR3E* | 14 | 568 | 31 | POLR3E | RNA polymerase III subunit E | Nucleus | transcription regulator | |
| chr22 | *BID* | 12 | 477 | 31 | BID | BH3 interacting domain death | Cytoplasm | other | |
| chr3 | *PLXNB1* | 13 | 638 | 31 | PLXNB1 | plexin B1 | Plasma Membrane | transmembrane receptor | |
| chr3 | *GPR160* | 14 | 648 | 31 | GPR160 | G protein-coupled receptor 160 | Plasma Membrane | G-protein coupled receptor | |

| | | | | | | | | | |
|---|---|---|---|---|---|---|---|---|---|
| chr1 | *HS2ST1* | 12 | 281 | 31 | HS2ST1 | heparan sulfate 2-O-sulfotransferase 1 | Cytoplasm | enzyme | |
| chr22 | *ARVCF* | 13 | 680 | 31 | ARVCF | ARVCF delta catenin family | Plasma Membrane | other | |
| chr15 | *CTD-3049M7.1* | 15 | 600 | 31 | | | | | |
| chr6 | *TAB2* | 13 | 305 | 31 | TAB2 | TGF-beta activated kinase 1 (MAP3K7) binding protein 2 | Cytoplasm | other | |
| chr7 | *ZC3HAV1* | 14 | 469 | 31 | ZC3HAV1 | zinc finger CCCH-type containing, antiviral 1 | Plasma Membrane | other | |
| chr15 | *DMXL2* | 11 | 359 | 31 | DMXL2 | Dmx like 2 | Cytoplasm | other | |
| chr1 | *TNNI1* | 13 | 845 | 31 | TNNI1 | troponin I1, slow skeletal type | Cytoplasm | other | |
| chr21 | *LINC00478* | 10 | 263 | 31 | MIR99AHG | mir-99a-let-7c cluster host gene | Other | other | |
| chr10 | *COL17A1* | 13 | 499 | 31 | COL17A1 | collagen type XVII alpha 1 chain | Extracellular Space | other | collagenase clostridium |
| chr15 | *RPAP1* | 13 | 475 | 31 | RPAP1 | RNA polymerase II associated protein 1 | Other | other | |
| chr20 | *PLCG1* | 14 | 780 | 31 | PLCG1 | phospholipase C gamma 1 | Cytoplasm | enzyme | |
| chr3 | *ZNF148* | 13 | 381 | 31 | ZNF148 | zinc finger protein 148 | Nucleus | transcription regulator | |
| chr1 | *KIAA1614* | 15 | 824 | 31 | KIAA1614 | KIAA1614 | Other | other | |
| chr6 | *STK38* | 12 | 332 | 31 | STK38 | serine/threonine kinase 38 | Nucleus | kinase | |
| chr19 | *PGPEP1* | 13 | 484 | 31 | PGPEP1 | pyroglutamyl-peptidase I | Cytoplasm | peptidase | |
| chr3 | *AC026188.1* | 14 | 360 | 31 | | | | | |
| chr1 | *MTF2* | 12 | 363 | 31 | MTF2 | metal response element binding transcription factor 2 | Nucleus | transcription regulator | |
| chr16 | *KCTD19* | 14 | 473 | 31 | KCTD19 | potassium channel tetramerization domain containing | Other | other | |
| chr18 | *PHLPP1* | 11 | 324 | 31 | PHLPP1 | PH domain and leucine rich repeat protein phosphatase 1 | Cytoplasm | enzyme | |
| chr6 | *CNKSR3* | 12 | 338 | 31 | CNKSR3 | CNKSR family member 3 | Plasma Membrane | kinase | |
| chr1 | *CMPK1* | 13 | 345 | 31 | CMPK1 | cytidine/uridine monophosphate kinase 1 | Nucleus | kinase | |
| chr16 | *IL21R* | 14 | 493 | 31 | IL21R | interleukin 21 receptor | Plasma Membrane | transmembrane recepto | denenicokin |
| chr8 | *WISP1* | 14 | 480 | 31 | CCN4 | cellular communication network factor 4 | Extracellular Space | other | |
| chr1 | *PTGFRN* | 15 | 481 | 31 | PTGFRN | prostaglandin F2 receptor | Plasma Membrane | other | |
| chr5 | *PAPD7* | 13 | 1341 | 31 | TENT4A | terminal nucleotidyltransferase | Nucleus | enzyme | |
| chr1 | *RP1-27O5.3* | 14 | 341 | 31 | | | | | |
| chr7 | *COPG2* | 13 | 354 | 31 | COPG2 | coatomer protein complex subunit gamma 2 | Cytoplasm | transporter | |
| chr17 | *SLFN12L* | 13 | 493 | 31 | SLFN12L | schlafen family member 12 like | Other | enzyme | |
| chr12 | *MAPKAPK5* | 14 | 341 | 31 | MAPKAPK5 | MAPK activated protein kinase 5 | Cytoplasm | kinase | |
| chr19 | *ARMC6* | 13 | 487 | 31 | ARMC6 | armadillo repeat containing 6 | Cytoplasm | other | |
| chr9 | *B4GALT1* | 14 | 412 | 31 | B4GALT1 | beta-1,4-galactosyltransferase 1 | Cytoplasm | enzyme | |
| chr7 | *AQP1* | 13 | 940 | 31 | AQP1 | aquaporin 1 (Colton blood group) | Plasma Membrane | transporter | |
| chr15 | *RP11-499F3.2* | 15 | 456 | 31 | | | | | |
| chr3 | *CBLB* | 12 | 387 | 31 | CBLB | Cbl proto-oncogene B | Nucleus | enzyme | |
| chr2 | *ATP6V1C2* | 12 | 499 | 31 | ATP6V1C2 | ATPase H+ transporting V1 subunit C2 | Cytoplasm | transporter | |
| chr22 | *PRR14L* | 12 | 318 | 31 | PRR14L | proline rich 14 like | Other | enzyme | |
| chr5 | *SKIV2L2* | 12 | 264 | 31 | MTREX | Mtr4 exosome RNA helicase | Nucleus | enzyme | |
| chr12 | *DDX12P* | 11 | 600 | 31 | DDX12P | DEAD/H-box helicase 12, pseudogene | Other | other | |
| chr3 | *OSBPL11* | 13 | 340 | 31 | OSBPL11 | oxysterol binding protein like 11 | Cytoplasm | other | |
| chrX | *RBM10* | 13 | 567 | 31 | RBM10 | RNA binding motif protein 10 | Nucleus | other | |
| chr5 | *NPR3* | 16 | 1277 | 31 | NPR3 | natriuretic peptide receptor 3 | Plasma Membrane | ì-protein coupled recepto | nesiritide |
| chr6 | *PSMG4* | 13 | 473 | 31 | PSMG4 | proteasome assembly chaperone | Other | transcription regulator | |
| chr1 | *RORC* | 13 | 773 | 31 | RORC | RAR related orphan receptor C | Nucleus | d-dependent nuclear rec | ROR gamma t inhibitor, LYC-55716, |
| chr3 | *CEP63* | 12 | 421 | 31 | CEP63 | centrosomal protein 63 | Cytoplasm | other | |
| chr19 | *PDE4A* | 14 | 461 | 31 | PDE4A | phosphodiesterase 4A | Cytoplasm | enzyme | caffeine, pentoxifylline, arofylline, tetomilast, amrinone, anagrelide, aspirin/dipyridamole/t elmisartan, theophylline, apremilast, milrinone, cilomilast, drotaverin, roflumilast, ketotifen, aspirin/dipyridamole, L-826,141, L 869298, enprofylline, |
| chr15 | *TLE3* | 15 | 522 | 31 | TLE3 | TLE family member 3, transcriptional corepressor | Nucleus | other | |
| chr20 | *DYNLRB1* | 14 | 654 | 31 | DYNLRB1 | dynein light chain roadblock-type | Cytoplasm | other | |
| chr5 | *HSD17B4* | 12 | 293 | 31 | HSD17B4 | hydroxysteroid 17-beta dehydrogenase 4 | Cytoplasm | enzyme | |
| chr6 | *DCDC2* | 14 | 258 | 31 | DCDC2 | doublecortin domain containing 2 | Extracellular Space | other | |
| chr14 | *SIPA1L1* | 13 | 322 | 31 | SIPA1L1 | signal induced proliferation associated 1 like 1 | Cytoplasm | other | |
| chr1 | *RPAP2* | 12 | 308 | 31 | RPAP2 | RNA polymerase II associated protein 2 | Nucleus | enzyme | |
| chr8 | *DMTN* | 14 | 646 | 31 | DMTN | dematin actin binding protein | Plasma Membrane | other | |
| chr3 | *UROC1* | 14 | 832 | 31 | UROC1 | urocanate hydratase 1 | Cytoplasm | enzyme | |
| chr20 | *LAMA5* | 16 | 530 | 31 | LAMA5 | laminin subunit alpha 5 | Extracellular Space | other | |
| chr22 | *SHANK3* | 15 | 762 | 31 | SHANK3 | SH3 and multiple ankyrin repeat domains 3 | Plasma Membrane | other | |
| chr7 | *STK31* | 13 | 455 | 31 | STK31 | serine/threonine kinase 31 | Cytoplasm | kinase | |
| chr1 | *ZNF670* | 13 | 358 | 31 | ZNF670 | zinc finger protein 670 | Cytoplasm | transcription regulator | |

| | | | | | | | | | |
|---|---|---|---|---|---|---|---|---|---|
| chr17 | *C17orf85* | 12 | 373 | 31 | NCBP3 | nuclear cap binding subunit 3 | Nucleus | other | |
| chr19 | *ARHGEF1* | 15 | 669 | 31 | ARHGEF1 | Rho guanine nucleotide exchange factor 1 | Cytoplasm | other | |
| chr17 | *APPBP2* | 13 | 272 | 31 | APPBP2 | amyloid beta precursor protein binding protein 2 | Cytoplasm | other | |
| chr2 | *M1AP* | 13 | 378 | 31 | M1AP | meiosis 1 associated protein | Cytoplasm | other | |
| chr7 | *NPC1L1* | 12 | 715 | 31 | NPC1L1 | NPC1 like intracellular cholesterol transporter 1 | Cytoplasm | transporter | ezetimibe/simvastatin, ezetimibe, ezetimibe/fenofibrate, amlodipine/ezetimibe/losartan/rosuvastatin, ezetimibe/fluvastatin, ezetimibe/rosuvastati |
| chr17 | *RAD51C* | 12 | 336 | 31 | RAD51C | RAD51 paralog C | Nucleus | enzyme | |
| chr1 | *CCDC18* | 11 | 347 | 31 | CCDC18 | coiled-coil domain containing 18 | Other | other | |
| chr8 | *FUT10* | 14 | 302 | 31 | FUT10 | fucosyltransferase 10 | Cytoplasm | enzyme | |
| chr5 | *MFAP3* | 14 | 318 | 31 | MFAP3 | microfibril associated protein 3 | Extracellular Space | other | |
| chr16 | *TCF25* | 15 | 694 | 31 | TCF25 | transcription factor 25 | Nucleus | transcription regulator | |
| chr20 | *DHX35* | 13 | 414 | 31 | DHX35 | DEAH-box helicase 35 | Other | enzyme | |
| chr12 | *RASSF3* | 13 | 320 | 31 | RASSF3 | Ras association domain family member 3 | Other | other | |
| chr9 | *TTC39B* | 12 | 309 | 31 | TTC39B | tetratricopeptide repeat domain | Other | other | |
| chr7 | *SMARCD3* | 13 | 618 | 31 | SMARCD3 | SWI/SNF related, matrix associated, actin dependent regulator of chromatin, subfamily | Nucleus | transcription regulator | |
| chr11 | *IFT46* | 13 | 315 | 31 | IFT46 | intraflagellar transport 46 | Cytoplasm | other | |
| chrX | *KLHL15* | 12 | 358 | 31 | KLHL15 | kelch like family member 15 | Nucleus | other | |
| chr16 | *EMP2* | 14 | 401 | 31 | EMP2 | epithelial membrane protein 2 | Plasma Membrane | other | |
| chr15 | *CSPG4* | 15 | 1069 | 31 | CSPG4 | chondroitin sulfate proteoglycan 4 | Plasma Membrane | other | |
| chr1 | *KHDRBS1* | 12 | 326 | 31 | KHDRBS1 | KH RNA binding domain containing, signal transduction | Nucleus | transcription regulator | CWP232291 |
| chr17 | *POLR2A* | 13 | 445 | 31 | POLR2A | RNA polymerase II subunit A | Nucleus | enzyme | |
| chr11 | *DDX10* | 13 | 321 | 31 | DDX10 | DEAD-box helicase 10 | Nucleus | enzyme | |
| chr17 | *OGFOD3* | 14 | 617 | 31 | OGFOD3 | 2-oxoglutarate and iron dependent oxygenase domain | Cytoplasm | enzyme | |
| chr19 | *DOCK6* | 12 | 469 | 31 | DOCK6 | dedicator of cytokinesis 6 | Cytoplasm | other | |
| chr11 | *RTN3* | 11 | 259 | 31 | RTN3 | reticulon 3 | Cytoplasm | other | |
| chr20 | *ASIP* | 12 | 490 | 31 | ASIP | agouti signaling protein | Extracellular Space | other | |
| chr1 | *NIPAL3* | 13 | 353 | 31 | NIPAL3 | NIPA like domain containing 3 | Other | other | |
| chr16 | *COG7* | 13 | 315 | 31 | COG7 | component of oligomeric golgi complex 7 | Cytoplasm | transporter | |
| chr20 | *APCDD1L* | 15 | 860 | 31 | APCDD1L | APC down-regulated 1 like | Other | other | |
| chrX | *SMC1A* | 14 | 419 | 31 | SMC1A | structural maintenance of chromosomes 1A | Nucleus | transporter | |
| chr2 | *AC068718.1* | 11 | 295 | 31 | | | | | |
| chr7 | *AC006372.4* | 14 | 674 | 31 | | | | | |
| chr4 | *STPG2* | 10 | 318 | 31 | STPG2 | sperm tail PG-rich repeat | Other | other | |
| chr14 | *SNX6* | 11 | 282 | 31 | SNX6 | sorting nexin 6 | Cytoplasm | transporter | |
| chr3 | *FYCO1* | 14 | 407 | 31 | FYCO1 | FYVE and coiled-coil domain containing 1 | Cytoplasm | other | |
| chr3 | *TNIK* | 11 | 273 | 31 | TNIK | TRAF2 and NCK interacting | Plasma Membrane | kinase | |
| chr16 | *HS3ST2* | 15 | 276 | 31 | HS3ST2 | heparan sulfate-glucosamine 3-sulfotransferase 2 | Cytoplasm | enzyme | |
| chr17 | *SAP30BP* | 13 | 606 | 31 | SAP30BP | SAP30 binding protein | Nucleus | transcription regulator | |
| chr19 | *SH3GL1* | 14 | 623 | 31 | SH3GL1 | SH3 domain containing GRB2 like 1, endophilin A2 | Cytoplasm | other | |
| chr15 | *STRA6* | 12 | 627 | 31 | STRA6 | stimulated by retinoic acid 6 | Plasma Membrane | transporter | |
| chrX | *RP11-120D5.1* | 14 | 305 | 31 | | | | | |
| chr1 | *PI4KB* | 13 | 539 | 31 | PI4KB | phosphatidylinositol 4-kinase beta | Cytoplasm | kinase | |
| chr1 | *STX12* | 13 | 307 | 31 | STX12 | syntaxin 12 | Cytoplasm | other | |
| chr16 | *CCDC102A* | 14 | 582 | 31 | CCDC102A | coiled-coil domain containing | Other | other | |
| chr3 | *TBL1XR1* | 10 | 245 | 31 | TBL1XR1 | transducin beta like 1 X-linked receptor 1 | Nucleus | transcription regulator | |
| chr3 | *UBE2E1* | 13 | 376 | 31 | UBE2E1 | ubiquitin conjugating enzyme E2 | Cytoplasm | enzyme | |
| chr2 | *HK2* | 14 | 423 | 31 | HK2 | hexokinase 2 | Cytoplasm | kinase | |
| chr11 | *SLC22A20* | 14 | 692 | 31 | SLC22A20P | solute carrier family 22 member 20, pseudogene | Other | other | |
| chr1 | *CHIT1* | 15 | 534 | 31 | CHIT1 | chitinase 1 | Extracellular Space | enzyme | |
| chr6 | *MLIP* | 11 | 276 | 31 | MLIP | muscular LMNA interacting | Nucleus | transcription regulator | |
| chr19 | *LDLR* | 11 | 412 | 31 | LDLR | low density lipoprotein receptor | Plasma Membrane | transporter | |
| chr4 | *TRIM2* | 11 | 301 | 31 | TRIM2 | tripartite motif containing 2 | Cytoplasm | enzyme | |
| chr1 | *COL8A2* | 14 | 586 | 31 | COL8A2 | collagen type VIII alpha 2 chain | Extracellular Space | other | collagenase clostridium |
| chr9 | *C9orf85* | 12 | 323 | 31 | C9orf85 | chromosome 9 open reading frame 85 | Other | other | |
| chr1 | *GPBP1L1* | 12 | 303 | 31 | GPBP1L1 | GC-rich promoter binding protein 1 like 1 | Other | other | |
| chr1 | *WNT2B* | 14 | 536 | 31 | WNT2B | Wnt family member 2B | Extracellular Space | other | |
| chr21 | *BACH1* | 14 | 281 | 31 | BACH1 | BTB domain and CNC homolog 1 | Nucleus | transcription regulator | |
| chr14 | *ADCK1* | 12 | 375 | 31 | ADCK1 | aarF domain containing kinase 1 | Cytoplasm | kinase | |
| chr3 | *KLHL18* | 13 | 342 | 31 | KLHL18 | kelch like family member 18 | Other | other | |
| chr11 | *VWCE* | 14 | 925 | 31 | VWCE | von Willebrand factor C and EGF domains | Cytoplasm | other | |
| chr6 | *TMEM63B* | 14 | 581 | 31 | TMEM63B | transmembrane protein 63B | Plasma Membrane | other | |
| chr10 | *ANTXRL* | 13 | 1004 | 31 | ANTXRL | ANTXR like | Other | other | |
| chr21 | *AP000282.2* | 14 | 323 | 31 | | | | | |
| chr19 | *HKR1* | 13 | 350 | 31 | ZNF875 | zinc finger protein 875 | Nucleus | transcription regulator | |
| chr12 | *TIMELESS* | 13 | 339 | 31 | TIMELESS | timeless circadian regulator | Nucleus | other | |

| | | | | | | | | | |
|---|---|---|---|---|---|---|---|---|---|
| chr3 | *GALNT15* | 13 | 555 | 31 | GALNT15 | polypeptide N-acetylgalactosaminyltransferase | Cytoplasm | enzyme | |
| chr17 | *ARHGAP27* | 13 | 621 | 31 | ARHGAP27 | Rho GTPase activating protein 27 | Cytoplasm | other | |
| chr19 | *ARID3A* | 15 | 675 | 31 | ARID3A | AT-rich interaction domain 3A | Nucleus | transcription regulator | |
| chr13 | *KLHL1* | 12 | 325 | 31 | KLHL1 | kelch like family member 1 | Cytoplasm | other | |
| chr16 | *TSC2* | 14 | 452 | 31 | TSC2 | TSC complex subunit 2 | Cytoplasm | other | |
| chr6 | *CUL9* | 13 | 509 | 31 | CUL9 | cullin 9 | Cytoplasm | other | |
| chr16 | *NSMCE1* | 13 | 508 | 31 | NSMCE1 | NSE1 homolog, SMC5-SMC6 complex component | Nucleus | transporter | |
| chr8 | *EIF3H* | 12 | 383 | 31 | EIF3H | eukaryotic translation initiation factor 3 subunit H | Cytoplasm | other | |
| chr9 | *RCL1* | 14 | 284 | 31 | RCL1 | RNA terminal phosphate cyclase like 1 | Nucleus | enzyme | |
| chr1 | *CLCN6* | 12 | 499 | 31 | CLCN6 | chloride voltage-gated channel 6 | Plasma Membrane | ion channel | |
| chr2 | *ATL2* | 11 | 351 | 31 | ATL2 | atlastin GTPase 2 | Cytoplasm | other | |
| chr12 | *BRAP* | 12 | 314 | 31 | BRAP | BRCA1 associated protein | Cytoplasm | enzyme | |
| chr3 | *BDH1* | 12 | 423 | 31 | BDH1 | 3-hydroxybutyrate | Cytoplasm | enzyme | |
| chr6 | *DTNBP1* | 13 | 357 | 31 | DTNBP1 | dystrobrevin binding protein 1 | Plasma Membrane | other | |
| chr5 | *FGF18* | 14 | 695 | 30 | FGF18 | fibroblast growth factor 18 | Extracellular Space | growth factor | |
| chr5 | *RP11-434D9.1* | 14 | 284 | 30 | | | | | |
| chr1 | *TRIM62* | 14 | 504 | 30 | TRIM62 | tripartite motif containing 62 | Cytoplasm | enzyme | |
| chr1 | *CDK18* | 14 | 956 | 30 | CDK18 | cyclin dependent kinase 18 | Cytoplasm | kinase | |
| chr5 | *ACSL6* | 12 | 495 | 30 | ACSL6 | acyl-CoA synthetase long chain family member 6 | Cytoplasm | enzyme | |
| chr5 | *TMEM232* | 15 | 282 | 30 | TMEM232 | transmembrane protein 232 | Other | other | |
| chr3 | *PDZRN3* | 13 | 310 | 30 | PDZRN3 | PDZ domain containing ring | Extracellular Space | enzyme | |
| chr9 | *FIBCD1* | 15 | 978 | 30 | FIBCD1 | fibrinogen C domain containing 1 | Other | other | |
| chr6 | *CLIC5* | 14 | 279 | 30 | CLIC5 | chloride intracellular channel 5 | Cytoplasm | ion channel | |
| chr20 | *CEP250* | 13 | 662 | 30 | CEP250 | centrosomal protein 250 | Nucleus | other | |
| chr10 | *PYROXD2* | 13 | 507 | 30 | PYROXD2 | pyridine nucleotide-disulphide oxidoreductase domain 2 | Other | other | |
| chr22 | *TANGO2* | 11 | 580 | 30 | TANGO2 | transport and golgi organization 2 homolog | Cytoplasm | other | |
| chr20 | *PABPC1L* | 14 | 632 | 30 | PABPC1L | poly(A) binding protein cytoplasmic 1 like | Cytoplasm | other | |
| chr7 | *AC007091.1* | 14 | 358 | 30 | | | | | |
| chr1 | *ARHGAP29* | 13 | 371 | 30 | ARHGAP29 | Rho GTPase activating protein 29 | Cytoplasm | other | |
| chr5 | *TNFAIP8* | 14 | 362 | 30 | TNFAIP8 | TNF alpha induced protein 8 | Cytoplasm | other | |
| chr15 | *ZNF280D* | 13 | 251 | 30 | ZNF280D | zinc finger protein 280D | Other | other | |
| chr14 | *AP4S1* | 10 | 240 | 30 | AP4S1 | adaptor related protein complex 4 subunit sigma 1 | Cytoplasm | other | |
| chr3 | *ADAMTS9-AS2* | 14 | 251 | 30 | ADAMTS9-AS2 | ADAMTS9 antisense RNA 2 | Other | other | |
| chr6 | *MBOAT1* | 14 | 292 | 30 | MBOAT1 | membrane bound O-acyltransferase domain | Cytoplasm | enzyme | |
| chr22 | *BRD1* | 14 | 635 | 30 | BRD1 | bromodomain containing 1 | Nucleus | other | |
| chr20 | *BPIFB1* | 13 | 663 | 30 | BPIFB1 | BPI fold containing family B member 1 | Extracellular Space | other | |
| chr6 | *KCTD20* | 12 | 332 | 30 | KCTD20 | potassium channel tetramerization domain containing | Cytoplasm | other | |
| chr2 | *AC007392.3* | 13 | 380 | 30 | | | | | |
| chr16 | *PDXDC1* | 14 | 690 | 30 | PDXDC1 | pyridoxal dependent decarboxylase domain containing | Cytoplasm | other | |
| chr1 | *F11R* | 14 | 456 | 30 | F11R | F11 receptor | Plasma Membrane | other | |
| chrX | *NHSL2* | 12 | 336 | 30 | NHSL2 | NHS like 2 | Other | other | |
| chr1 | *RHCE* | 13 | 340 | 30 | RHCE/RHD | Rh blood group D antigen | Plasma Membrane | transporter | |
| chr4 | *LETM1* | 13 | 490 | 30 | LETM1 | leucine zipper and EF-hand containing transmembrane | Cytoplasm | transporter | |
| chr5 | *HEXB* | 12 | 332 | 30 | HEXB | hexosaminidase subunit beta | Cytoplasm | enzyme | |
| chr18 | *ZNF407* | 11 | 273 | 30 | ZNF407 | zinc finger protein 407 | Nucleus | transcription regulator | |
| chr1 | *E2F2* | 12 | 445 | 30 | E2F2 | E2F transcription factor 2 | Nucleus | transcription regulator | |
| chr3 | *SKIL* | 13 | 537 | 30 | SKIL | SKI like proto-oncogene | Nucleus | transcription regulator | |
| chr9 | *CENPP* | 10 | 321 | 30 | CENPP | centromere protein P | Nucleus | other | |
| chr10 | *SLC16A12* | 13 | 285 | 30 | SLC16A12 | solute carrier family 16 member | Plasma Membrane | transporter | |
| chrX | *CACNA1F* | 13 | 502 | 30 | CACNA1F | calcium voltage-gated channel subunit alpha1 F | Plasma Membrane | ion channel | verapamil, dexamethasone/rituximab/verapamil, nicardipine, bepridil, mibefradil, isradipine, MEM-1003, nimodipine, cinnarizine, trandolapril/verapamil |
| chr1 | *C1orf127* | 14 | 499 | 30 | C1orf127 | chromosome 1 open reading frame 127 | Other | other | |
| chr1 | *INTS7* | 10 | 393 | 30 | INTS7 | integrator complex subunit 7 | Nucleus | other | |
| chr19 | *FBXO27* | 13 | 325 | 30 | FBXO27 | F-box protein 27 | Cytoplasm | enzyme | |
| chr5 | *CXXC5* | 14 | 735 | 30 | CXXC5 | CXXC finger protein 5 | Nucleus | other | |
| chr15 | *NUSAP1* | 11 | 344 | 30 | NUSAP1 | nucleolar and spindle associated protein 1 | Nucleus | other | |
| chr17 | *KCTD2* | 13 | 485 | 30 | KCTD2 | potassium channel tetramerization domain containing | Other | ion channel | |
| chr3 | *FAM19A4* | 15 | 335 | 30 | TAFA4 | TAFA chemokine like family member 4 | Extracellular Space | other | |
| chr2 | *TGFBRAP1* | 12 | 437 | 30 | TGFBRAP1 | transforming growth factor beta receptor associated protein 1 | Cytoplasm | other | |
| chr16 | *CCDC101* | 12 | 429 | 30 | SGF29 | SAGA complex associated factor | Nucleus | other | |
| chr16 | *TMC5* | 15 | 326 | 30 | TMC5 | transmembrane channel like 5 | Cytoplasm | other | |
| chr11 | *ARHGEF12* | 11 | 283 | 30 | ARHGEF12 | Rho guanine nucleotide exchange factor 12 | Cytoplasm | other | |

| | | | | | | | | | |
|---|---|---|---|---|---|---|---|---|---|
| chr3 | *RP11-894J14.5* | 13 | 447 | 30 | | | | | |
| chr12 | *TMEM116* | 13 | 349 | 30 | TMEM116 | transmembrane protein 116 | Other | other | |
| chr15 | *DPP8* | 12 | 274 | 30 | DPP8 | dipeptidyl peptidase 8 | Cytoplasm | peptidase | |
| chr6 | *GPLD1* | 14 | 342 | 30 | GPLD1 | glycosylphosphatidylinositol specific phospholipase D1 | Cytoplasm | enzyme | |
| chr9 | *KIAA1432* | 12 | 309 | 30 | RIC1 | RIC1 homolog, RAB6A GEF complex partner 1 | Cytoplasm | other | |
| chr15 | *GPR176* | 13 | 253 | 30 | GPR176 | G protein-coupled receptor 176 | Plasma Membrane | G-protein coupled recepto | |
| chr8 | *HNF4G* | 11 | 273 | 30 | HNF4G | hepatocyte nuclear factor 4 | Nucleus | transcription regulator | |
| chr1 | *TRNAU1AP* | 13 | 354 | 30 | TRNAU1AP | tRNA selenocysteine 1 associated protein 1 | Nucleus | other | |
| chr19 | *PLD3* | 13 | 524 | 30 | PLD3 | phospholipase D family member | Cytoplasm | enzyme | |
| chr12 | *IQSEC3* | 12 | 475 | 30 | IQSEC3 | IQ motif and Sec7 domain | Cytoplasm | other | |
| chr22 | *ACO2* | 12 | 417 | 30 | ACO2 | aconitase 2 | Cytoplasm | enzyme | |
| chr2 | *IL1R1* | 14 | 400 | 30 | IL1R1 | interleukin 1 receptor type 1 | Plasma Membrane | transmembrane recepto | anakinra |
| chr16 | *IL4R* | 12 | 510 | 30 | IL4R | interleukin 4 receptor | Plasma Membrane | transmembrane recepto | dupilumab, MDNA55 |
| chr1 | *RP11-286B14.1* | 14 | 361 | 30 | | | | | |
| chr7 | *PRKRIP1* | 13 | 377 | 30 | PRKRIP1 | PRKR interacting protein 1 | Nucleus | other | |
| chr12 | *ELK3* | 14 | 469 | 30 | ELK3 | ETS transcription factor ELK3 | Nucleus | transcription regulator | |
| chr17 | *ERBB2* | 12 | 535 | 30 | ERBB2 | erb-b2 receptor tyrosine kinase 2 | Plasma Membrane | kinase | paclitaxel/pertuzuma b/trastuzumab, ADCT-502, tucatinib, erlotinib/osimertinib, docetaxel/pertuzuma b/trastuzumab, neratinib, 89Zr-trastuzumab, erlotinib, A166, anti-HER2 antibody, HER2 inhibitor, docetaxel/lapatinib, afatinib, afatinib/paclitaxel, cyclophosphamide/docetaxel/epirubicin/5-fluorouracil/trastuzumab, sym013, trastuzumab, MM-111, BMS-690514, cyclophosphamide/docetaxel/trastuzumab, MBS301, selatinib, [illegible] |
| chr15 | *DPH6-AS1* | 13 | 291 | 30 | DPH6-DT | DPH6 divergent transcript | Other | other | |
| chr9 | *COL15A1* | 12 | 453 | 30 | COL15A1 | collagen type XV alpha 1 chain | Extracellular Space | other | collagenase clostridium |
| chr15 | *SPTBN5* | 15 | 688 | 30 | SPTBN5 | spectrin beta, non-erythrocytic 5 | Plasma Membrane | other | |
| chr1 | *RP11-435B5.4* | 13 | 751 | 30 | | | | | |
| chr17 | *KCNJ12* | 14 | 1335 | 30 | KCNJ12 | potassium inwardly rectifying channel subfamily J member 12 | Plasma Membrane | ion channel | amiodarone, nicorandil, dofetilide |
| chrX | *RP11-305F18.1* | 13 | 315 | 30 | | | | | |
| chr1 | *YARS* | 14 | 317 | 30 | YARS1 | tyrosyl-tRNA synthetase 1 | Cytoplasm | enzyme | |
| chr11 | *RP11-727A23.10* | 13 | 354 | 30 | | | | | |
| chr4 | *WDR1* | 11 | 479 | 30 | WDR1 | WD repeat domain 1 | Extracellular Space | other | |
| chr1 | *CDC73* | 10 | 369 | 30 | CDC73 | cell division cycle 73 | Nucleus | other | |
| chr19 | *ZC3H4* | 12 | 443 | 30 | ZC3H4 | zinc finger CCCH-type containing | Nucleus | transcription regulator | |
| chr17 | *TP53* | 12 | 395 | 30 | TP53 | tumor protein p53 | Nucleus | transcription regulator | BI 907828, kevetrin, cenersen, azurin 50-77, ALT-801, APR-246, COTI-2, |
| chr5 | *CMBL* | 14 | 1107 | 30 | CMBL | carboxymethylenebutenolidase homolog | Cytoplasm | enzyme | |
| chr11 | *AP002387.1* | 13 | 481 | 30 | | | | | |
| chr13 | *PHF2P2* | 11 | 465 | 30 | PHF2P2 | PHD finger protein 2 pseudogene | Other | other | |
| chr2 | *ZRANB3* | 11 | 302 | 30 | ZRANB3 | zinc finger RANBP2-type containing 3 | Nucleus | enzyme | |
| chr14 | *RALGAPA1* | 10 | 231 | 30 | RALGAPA1 | Ral GTPase activating protein catalytic alpha subunit 1 | Cytoplasm | other | |
| chrX | *RP11-545D19.1* | 11 | 324 | 30 | | | | | |
| chr1 | *VANGL1* | 13 | 467 | 30 | VANGL1 | VANGL planar cell polarity | Cytoplasm | other | |
| chr3 | *AC090044.1* | 12 | 308 | 30 | | | | | |
| chr7 | *SLC12A9* | 13 | 544 | 30 | SLC12A9 | solute carrier family 12 member 9 | Plasma Membrane | transporter | |
| chr2 | *PARD3B* | 12 | 287 | 30 | PARD3B | par-3 family cell polarity regulator | Plasma Membrane | other | |
| chr2 | *NBEAL1* | 10 | 252 | 30 | NBEAL1 | neurobeachin like 1 | Other | other | |
| chr15 | *PIAS1* | 12 | 245 | 30 | PIAS1 | protein inhibitor of activated | Nucleus | transcription regulator | |
| chr14 | *BEGAIN* | 13 | 659 | 30 | BEGAIN | brain enriched guanylate kinase associated | Nucleus | other | |
| chr2 | *AC009499.1* | 12 | 296 | 30 | | | | | |
| chr1 | *SLC44A3* | 13 | 444 | 30 | SLC44A3 | solute carrier family 44 member 3 | Plasma Membrane | transporter | |
| chr8 | *SCARA3* | 13 | 477 | 30 | SCARA3 | scavenger receptor class A member 3 | Plasma Membrane | transmembrane recepto | |
| chr15 | *TMED3* | 13 | 427 | 30 | TMED3 | transmembrane p24 trafficking protein 3 | Cytoplasm | transporter | |
| chr7 | *PILRA* | 14 | 513 | 30 | PILRA | paired immunoglobin like type 2 receptor alpha | Plasma Membrane | other | |
| chr15 | *SPESP1* | 12 | 343 | 30 | SPESP1 | sperm equatorial segment protein | Cytoplasm | other | |
| chr15 | *IGDCC4* | 13 | 538 | 30 | IGDCC4 | immunoglobulin superfamily DCC subclass member 4 | Plasma Membrane | other | |

| | | | | | | | | | |
|---|---|---|---|---|---|---|---|---|---|
| chrY | *ASMTL* | 11 | 471 | 30 | ASMTL | acetylserotonin O-methyltransferase like | Cytoplasm | enzyme | |
| chr8 | *FGFR1* | 11 | 406 | 30 | FGFR1 | fibroblast growth factor receptor 1 | Plasma Membrane | kinase | sorafenib, ASP5878, lapatinib/pazopanib, PRN1371, lenvatinib, sorafenib/sulindac/sunitinib, debio 1347, dexamethasone/lenalidomide/sorafenib, regorafenib, 5-azacytidine/sorafenib, pazopanib, SAR103168, SKI-G-801, derazantinib, orantinib, everolimus/sorafenib, sulfatinib, bortezomib/sorafenib, 3D185, erdafitinib, decitabine/sorafenib, clofarabine/cytarabine/sorafenib, lenvatinib/pembrolizumab, cytarabine/idarubicin/[illegible] |
| chr11 | *PPP2R1B* | 11 | 420 | 30 | PPP2R1B | protein phosphatase 2 scaffold subunit Abeta | Plasma Membrane | phosphatase | |
| chr17 | *GRAP* | 12 | 561 | 30 | GRAP | GRB2 related adaptor protein | Cytoplasm | other | |
| chr1 | *LEPR* | 12 | 386 | 30 | LEPR | leptin receptor | Plasma Membrane | transmembrane receptor | recombinant-methionyl human |
| chr17 | *TNFRSF13B* | 12 | 477 | 30 | TNFRSF13B | TNF receptor superfamily member 13B | Plasma Membrane | transmembrane receptor | |
| chr9 | *TBC1D13* | 13 | 583 | 30 | TBC1D13 | TBC1 domain family member 13 | Cytoplasm | other | |
| chr17 | *GOSR1* | 11 | 265 | 30 | GOSR1 | golgi SNAP receptor complex member 1 | Cytoplasm | transporter | |
| chr7 | *AC005062.2* | 11 | 354 | 30 | | | | | |
| chr1 | *FAM20B* | 11 | 466 | 30 | FAM20B | FAM20B glycosaminoglycan xylosylkinase | Extracellular Space | enzyme | |
| chr7 | *VKORC1L1* | 12 | 298 | 30 | VKORC1L1 | vitamin K epoxide reductase complex subunit 1 like 1 | Cytoplasm | enzyme | vitamin K3 |
| chr15 | *RAB11A* | 12 | 311 | 30 | RAB11A | RAB11A, member RAS oncogene family | Cytoplasm | enzyme | |
| chr19 | *COL5A3* | 12 | 393 | 30 | COL5A3 | collagen type V alpha 3 chain | Extracellular Space | other | collagenase clostridium |
| chr1 | *SPATA21* | 13 | 487 | 30 | SPATA21 | spermatogenesis associated 21 | Other | other | |
| chr13 | *DACH1* | 11 | 234 | 30 | DACH1 | dachshund family transcription factor 1 | Nucleus | transcription regulator | |
| chr7 | *LAMB4* | 13 | 288 | 30 | LAMB4 | laminin subunit beta 4 | Other | other | |
| chr4 | *PDS5A* | 10 | 242 | 30 | PDS5A | PDS5 cohesin associated factor | Nucleus | other | |
| chr4 | *STOX2* | 11 | 350 | 30 | STOX2 | storkhead box 2 | Other | other | |
| chr17 | *HSF5* | 14 | 293 | 30 | HSF5 | heat shock transcription factor 5 | Other | other | |
| chr2 | *LMAN2L* | 12 | 447 | 30 | LMAN2L | lectin, mannose binding 2 like | Cytoplasm | other | |
| chr1 | *RP11-353N4.5* | 15 | 846 | 30 | | | | | |
| chr1 | *SLC25A24P1* | 11 | 345 | 30 | SLC25A24P1 | SLC25A24 pseudogene 1 | Other | other | |
| chr6 | *RP3-468B3.2* | 13 | 484 | 30 | | | | | |
| chr1 | *RP11-278H7.1* | 13 | 392 | 30 | | | | | |
| chr8 | *LINC00535* | 12 | 312 | 30 | LINC00535 | long intergenic non-protein coding RNA 535 | Other | other | |
| chr17 | *MYO1C* | 15 | 961 | 30 | MYO1C | myosin IC | Cytoplasm | enzyme | |
| chr17 | *SLC13A5* | 13 | 512 | 30 | SLC13A5 | solute carrier family 13 member 5 | Plasma Membrane | transporter | |
| chr20 | *LBP* | 12 | 526 | 30 | LBP | lipopolysaccharide binding protein | Plasma Membrane | transporter | |
| chr22 | *KCNJ4* | 12 | 517 | 30 | KCNJ4 | potassium inwardly rectifying channel subfamily J member 4 | Plasma Membrane | ion channel | |
| chr1 | *KLHL20* | 12 | 376 | 29 | KLHL20 | kelch like family member 20 | Plasma Membrane | enzyme | |
| chr6 | *SAMD5* | 12 | 248 | 29 | SAMD5 | sterile alpha motif domain containing 5 | Cytoplasm | other | |
| chr15 | *ZFAND6* | 11 | 317 | 29 | ZFAND6 | zinc finger AN1-type containing 6 | Cytoplasm | other | |
| chr11 | *POU2AF1* | 13 | 393 | 29 | POU2AF1 | POU class 2 homeobox associating factor 1 | Nucleus | transcription regulator | |
| chr19 | *CAPN12* | 13 | 473 | 29 | CAPN12 | calpain 12 | Other | peptidase | |
| chr5 | *PPAP2A* | 12 | 304 | 29 | PLPP1 | phospholipid phosphatase 1 | Plasma Membrane | phosphatase | |
| chr1 | *NME7* | 13 | 439 | 29 | NME7 | NME/NM23 family member 7 | Cytoplasm | kinase | |
| chr8 | *XKR4* | 11 | 308 | 29 | XKR4 | XK related 4 | Other | other | |
| chr11 | *POU2F3* | 13 | 363 | 29 | POU2F3 | POU class 2 homeobox 3 | Nucleus | transcription regulator | |
| chr2 | *GCC2* | 10 | 396 | 29 | GCC2 | GRIP and coiled-coil domain containing 2 | Cytoplasm | other | |
| chr16 | *CDIP1* | 12 | 466 | 29 | CDIP1 | cell death inducing p53 target 1 | Nucleus | other | |
| chr1 | *CTTNBP2NL* | 12 | 374 | 29 | CTTNBP2NL | CTTNBP2 N-terminal like | Cytoplasm | other | |
| chr6 | *ETV7* | 13 | 422 | 29 | ETV7 | ETS variant transcription factor 7 | Nucleus | transcription regulator | |
| chr1 | *WDR64* | 12 | 344 | 29 | WDR64 | WD repeat domain 64 | Other | other | |
| chr3 | *RP11-536I6.2* | 13 | 538 | 29 | | | | | |
| chr11 | *CD6* | 13 | 661 | 29 | CD6 | CD6 molecule | Plasma Membrane | transmembrane receptor | |
| chr13 | *NBEA* | 10 | 278 | 29 | NBEA | neurobeachin | Cytoplasm | other | |
| chr1 | *NDC1* | 11 | 297 | 29 | NDC1 | NDC1 transmembrane | Nucleus | other | |
| chr19 | *ZNF98* | 13 | 331 | 29 | ZNF492/ZNF98 | zinc finger protein 492 | Nucleus | transcription regulator | |
| chr19 | *ZSCAN5A* | 12 | 392 | 29 | ZSCAN5A | zinc finger and SCAN domain containing 5A | Nucleus | transcription regulator | |

| | | | | | | | | | |
|---|---|---|---|---|---|---|---|---|---|
| chr3 | *GPD1L* | 13 | 291 | 29 | GPD1L | glycerol-3-phosphate dehydrogenase 1 like | Cytoplasm | enzyme | |
| chr16 | *RP11-396B14.2* | 13 | 530 | 29 | | | | | |
| chr2 | *ITSN2* | 11 | 346 | 29 | ITSN2 | intersectin 2 | Cytoplasm | other | |
| chr10 | *C10orf128* | 13 | 701 | 29 | TMEM273 | transmembrane protein 273 | Other | other | |
| chr22 | *EP300* | 9 | 250 | 29 | EP300 | E1A binding protein p300 | Nucleus | transcription regulator | CCS1477 |
| chr16 | *DNASE1* | 11 | 398 | 29 | DNASE1 | deoxyribonuclease 1 | Extracellular Space | enzyme | |
| chr15 | *GABPB1* | 11 | 274 | 29 | GABPB1 | GA binding protein transcription factor subunit beta 1 | Nucleus | transcription regulator | |
| chr19 | *SCGB2B2* | 13 | 470 | 29 | SCGB2B2 | secretoglobin family 2B member | Other | other | |
| chr11 | *DEAF1* | 12 | 540 | 29 | DEAF1 | DEAF1 transcription factor | Nucleus | transcription regulator | |
| chr2 | *NFU1* | 12 | 297 | 29 | NFU1 | NFU1 iron-sulfur cluster scaffold | Cytoplasm | other | |
| chr9 | *SLC27A4* | 13 | 764 | 29 | SLC27A4 | solute carrier family 27 member 4 | Plasma Membrane | transporter | |
| chr9 | *C9orf96* | 13 | 703 | 29 | STKLD1 | serine/threonine kinase like domain containing 1 | Other | kinase | |
| chr14 | *KIAA0391* | 11 | 248 | 29 | PRORP | protein only RNase P catalytic subunit | Cytoplasm | enzyme | |
| chr12 | *STX2* | 12 | 483 | 29 | STX2 | syntaxin 2 | Cytoplasm | transporter | |
| chr9 | *SNX30* | 11 | 333 | 29 | SNX30 | sorting nexin family member 30 | Other | other | |
| chr15 | *PML* | 13 | 476 | 29 | PML | promyelocytic leukemia | Nucleus | transcription regulator | arsenic trioxide |
| chr12 | *RAP1B* | 10 | 261 | 29 | RAP1B | RAP1B, member of RAS oncogene family | Cytoplasm | enzyme | |
| chr3 | *SEMA3F* | 13 | 689 | 29 | SEMA3F | semaphorin 3F | Extracellular Space | other | |
| chr11 | *RNF121* | 11 | 306 | 29 | RNF121 | ring finger protein 121 | Other | other | |
| chr22 | *RTN4R* | 13 | 668 | 29 | RTN4R | reticulon 4 receptor | Plasma Membrane | transmembrane receptor | |
| chr2 | *C2orf88* | 12 | 282 | 29 | C2orf88 | chromosome 2 open reading frame 88 | Other | other | |
| chr3 | *KIAA0226* | 10 | 325 | 29 | RUBCN | rubicon autophagy regulator | Cytoplasm | other | |
| chr9 | *TMOD1* | 12 | 401 | 29 | TMOD1 | tropomodulin 1 | Cytoplasm | enzyme | |
| chr16 | *PDXDC2P* | 13 | 359 | 29 | PDXDC2P | | Other | other | |
| chr3 | *SCN10A* | 13 | 372 | 29 | SCN10A | sodium voltage-gated channel alpha subunit 10 | Plasma Membrane | ion channel | procaine, riluzole, mepivacaine, benzocaine, lidocaine/prilocaine, bupivacaine/sufentanil, epinephrine/lidocaine, bupivacaine/lidocaine, ropivacaine, lacosamide, bupivacaine/epinephrine, dibucaine, dyclonine, proxymetacaine |
| chr5 | *RP11-124N3.3* | 13 | 691 | 29 | | | | | |
| chr20 | *RNF24* | 9 | 249 | 29 | RNF24 | ring finger protein 24 | Cytoplasm | other | |
| chr1 | *ACTN2* | 12 | 443 | 29 | ACTN2 | actinin alpha 2 | Nucleus | transcription regulator | |
| chr22 | *ZNRF3* | 11 | 290 | 29 | ZNRF3 | zinc and ring finger 3 | Plasma Membrane | enzyme | |
| chr20 | *NKAIN4* | 15 | 1064 | 29 | NKAIN4 | sodium/potassium transporting ATPase interacting 4 | Other | other | |
| chr9 | *PSMB7* | 13 | 519 | 29 | PSMB7 | proteasome subunit beta 7 | Cytoplasm | peptidase | |
| chr19 | *ZNF420* | 13 | 444 | 29 | ZNF420 | zinc finger protein 420 | Cytoplasm | transcription regulator | |
| chr19 | *EVI5L* | 14 | 707 | 29 | EVI5L | ecotropic viral integration site 5 | Other | other | |
| chr2 | *ERCC3* | 12 | 464 | 29 | ERCC3 | ERCC excision repair 3, TFIIH core complex helicase subunit | Nucleus | enzyme | |
| chr2 | *BCL11A* | 14 | 329 | 29 | BCL11A | BAF chromatin remodeling complex subunit BCL11A | Nucleus | transcription regulator | |
| chr2 | *RAMP1* | 11 | 532 | 29 | RAMP1 | receptor activity modifying protein | Plasma Membrane | other | pramlintide |
| chr15 | *SRP14-AS1* | 12 | 420 | 29 | SRP14-AS1 | SRP14 antisense RNA1 (head to head) | Other | other | |
| chr16 | *C16orf52* | 11 | 274 | 29 | MOSMO | modulator of smoothened | Other | other | |
| chr17 | *NBR2* | 12 | 395 | 29 | NBR2 | neighbor of BRCA1 lncRNA 2 | Other | other | |
| chr8 | *GPR124* | 12 | 512 | 29 | ADGRA2 | adhesion G protein-coupled receptor A2 | Plasma Membrane | G-protein coupled receptor | |
| chr19 | *SSC5D* | 13 | 543 | 29 | SSC5D | scavenger receptor cysteine rich family member with 5 domains | Plasma Membrane | transmembrane receptor | |
| chr2 | *HEATR5B* | 10 | 249 | 29 | HEATR5B | HEAT repeat containing 5B | Cytoplasm | other | |
| chr1 | *DHCR24* | 12 | 426 | 29 | DHCR24 | 24-dehydrocholesterol reductase | Cytoplasm | enzyme | |
| chr11 | *ARNTL* | 13 | 401 | 29 | ARNTL | aryl hydrocarbon receptor nuclear translocator like | Nucleus | transcription regulator | |
| chr8 | *PXDNL* | 11 | 276 | 29 | PXDNL | peroxidasin like | Extracellular Space | enzyme | |
| chr1 | *SLC35A3* | 11 | 340 | 29 | SLC35A3 | solute carrier family 35 member | Cytoplasm | transporter | |
| chr15 | *WDR93* | 13 | 416 | 29 | WDR93 | WD repeat domain 93 | Other | other | |
| chr17 | *CCDC144B* | 11 | 435 | 29 | CCDC144B | coiled-coil domain containing 144B (pseudogene) | Other | other | |
| chr19 | *EMR1* | 13 | 286 | 29 | ADGRE1 | adhesion G protein-coupled receptor E1 | Plasma Membrane | G-protein coupled receptor | |
| chr5 | *LIFR* | 12 | 961 | 29 | LIFR | LIF receptor subunit alpha | Plasma Membrane | transmembrane receptor | |
| chr1 | *ST6GALNAC5* | 14 | 331 | 29 | ST6GALNAC5 | ST6 N-acetylgalactosaminide alpha-2,6-sialyltransferase 5 | Cytoplasm | enzyme | |
| chr6 | *CD2AP* | 11 | 283 | 29 | CD2AP | CD2 associated protein | Cytoplasm | other | |
| chr1 | *IARS2* | 13 | 379 | 29 | IARS2 | isoleucyl-tRNA synthetase 2, mitochondrial | Cytoplasm | enzyme | |
| chr5 | *CANX* | 12 | 452 | 29 | CANX | calnexin | Cytoplasm | other | |
| chr9 | *ZBTB43* | 12 | 368 | 29 | ZBTB43 | zinc finger and BTB domain containing 43 | Nucleus | transcription regulator | |
| chr19 | *DPY19L3* | 12 | 277 | 29 | DPY19L3 | dpy-19 like C- | Other | other | |

| | | | | | | | | | |
|---|---|---|---|---|---|---|---|---|---|
| chr4 | *FRAS1* | 12 | 336 | 29 | FRAS1 | Fraser extracellular matrix complex subunit 1 | Extracellular Space | other | |
| chr1 | *VWA5B1* | 13 | 457 | 29 | VWA5B1 | von Willebrand factor A domain containing 5B1 | Extracellular Space | other | |
| chr7 | *ZNF273* | 13 | 498 | 29 | ZNF273 | zinc finger protein 273 | Nucleus | transcription regulator | |
| chr11 | *PTPN5* | 13 | 387 | 29 | PTPN5 | protein tyrosine phosphatase non-receptor type 5 | Plasma Membrane | phosphatase | |
| chr16 | *HERC2P4* | 11 | 468 | 29 | HERC2P4 | hect domain and RLD 2 pseudogene 4 | Other | other | |
| chr17 | *RP11-64J4.2* | 13 | 354 | 29 | | | | | |
| chr16 | *NECAB2* | 12 | 737 | 29 | NECAB2 | N-terminal EF-hand calcium binding protein 2 | Cytoplasm | other | |
| chr19 | *TRPM4* | 11 | 374 | 29 | TRPM4 | transient receptor potential cation channel subfamily M member 4 | Plasma Membrane | ion channel | |
| chr5 | *LMNB1* | 12 | 321 | 29 | LMNB1 | lamin B1 | Nucleus | other | |
| chr22 | *TTLL1* | 11 | 367 | 29 | TTLL1 | tubulin tyrosine ligase like 1 | Extracellular Space | enzyme | |
| chr3 | *CNTN6* | 12 | 309 | 29 | CNTN6 | contactin 6 | Plasma Membrane | enzyme | |
| chr3 | *NMD3* | 13 | 312 | 29 | NMD3 | NMD3 ribosome export adaptor | Nucleus | other | |
| chr19 | *LIG1* | 11 | 389 | 29 | LIG1 | DNA ligase 1 | Nucleus | enzyme | |
| chr2 | *BIRC6* | 9 | 234 | 29 | BIRC6 | baculoviral IAP repeat containing | Cytoplasm | enzyme | |
| chr12 | *CHD4* | 13 | 429 | 29 | CHD4 | chromodomain helicase DNA binding protein 4 | Nucleus | enzyme | |
| chr1 | *HFM1* | 12 | 536 | 29 | HFM1 | helicase for meiosis 1 | Other | other | |
| chr2 | *GPR75-ASB3* | 13 | 445 | 29 | SB3/GPR75-ASB | ankyrin repeat and SOCS box containing 3 | Cytoplasm | transcription regulator | |
| chr18 | *BCL2* | 10 | 295 | 29 | BCL2 | BCL2 apoptosis regulator | Cytoplasm | transporter | lapatinib/paclitaxel, paclitaxel/pertuzumab/trastuzumab, venetoclax, bortezomib/paclitaxel, gemcitabine/paclitaxel, APG-1252, decitabine/venetoclax, rituximab/venetoclax, chelerythrine, doxorubicin/paclitaxel, doxorubicin/lapatinib/paclitaxel/trastuzumab, afatinib/paclitaxel, SPC2996, BCL-2 blocker, paclitaxel/rituximab, paclitaxel/pembrolizumab/ramucirumab, paclitaxel/ramucirum [illegible] |
| chr3 | *MLH1* | 12 | 306 | 29 | MLH1 | mutL homolog 1 | Nucleus | enzyme | |
| chr22 | *MN1* | 11 | 418 | 29 | MN1 | MN1 proto-oncogene, transcriptional regulator | Nucleus | other | |
| chr6 | *TMEM217* | 13 | 308 | 29 | TMEM217 | transmembrane protein 217 | Nucleus | other | |
| chr16 | *SH2B1* | 12 | 508 | 29 | SH2B1 | SH2B adaptor protein 1 | Cytoplasm | other | |
| chr7 | *CHST12* | 13 | 561 | 29 | CHST12 | carbohydrate sulfotransferase 12 | Cytoplasm | enzyme | |
| chr5 | *PCBD2* | 12 | 299 | 29 | PCBD2 | pterin-4 alpha-carbinolamine dehydratase 2 | Nucleus | enzyme | |
| chr12 | *USP44* | 10 | 282 | 29 | USP44 | ubiquitin specific peptidase 44 | Nucleus | peptidase | |
| chr17 | *ACE* | 13 | 569 | 29 | ACE | angiotensin I converting enzyme | Plasma Membrane | peptidase | [illegible]sinopril, fosinopril, amlodipine/perindopril, trandolaprilat, trandolapril, perindopril, captopril, captopril/hydrochlorothiazide, enalapril, quinaprilat, amlodipine/benazepril, ramiprilat, enalapril/felodipine, benazeprilat, benazepril, lisinopril, angiotensin-converting enzyme inhibitor, quinapril, hydrochlorothiazide/moexipril, amlodipine/indapamide/perindopril, aspirin/lisinopril, perindoprilat, [illegible] |
| chr12 | *STK38L* | 11 | 340 | 29 | STK38L | serine/threonine kinase 38 like | Cytoplasm | kinase | |
| chr19 | *CARM1* | 12 | 498 | 29 | CARM1 | coactivator associated arginine methyltransferase 1 | Nucleus | transcription regulator | |

| | | | | | | | | | |
|---|---|---|---|---|---|---|---|---|---|
| chr20 | *ADA* | 13 | 683 | 29 | ADA | adenosine deaminase | Cytoplasm | enzyme | cyclophosphamide/pentostatin/rituximab, pentostatin, alemtuzumab/pentostatin, vidarabine, pentostatin/recombinant interferon, aspirin/dipyridamole/telmisartan, pentostatin/rituximab, dipyridamole, aspirin/dipyridamole, adenosine deaminase inhibitor |
| chr2 | *PCDP1* | 13 | 434 | 29 | | | | | |
| chr22 | *TMPRSS6* | 12 | 546 | 29 | TMPRSS6 | transmembrane serine protease 6 | Plasma Membrane | peptidase | |
| chr1 | *RP11-400N13.1* | 14 | 320 | 29 | | | | | |
| chr10 | *ANTXRLP1* | 12 | 889 | 29 | ANTXRLP1 | ANTXR like pseudogene 1 | Other | other | |
| chr16 | *CETP* | 12 | 513 | 29 | CETP | cholesteryl ester transfer protein | Extracellular Space | enzyme | torcetrapib |
| chr1 | *AK4* | 12 | 286 | 29 | AK4 | adenylate kinase 4 | Cytoplasm | kinase | |
| chr19 | *SHC2* | 14 | 712 | 29 | SHC2 | SHC adaptor protein 2 | Cytoplasm | other | |
| chr1 | *TPM3* | 13 | 443 | 29 | TPM3 | tropomyosin 3 | Cytoplasm | other | AZD-7451 |
| chr16 | *KCTD5* | 13 | 635 | 29 | KCTD5 | potassium channel tetramerization domain containing | Cytoplasm | other | |
| chr16 | *SBK1* | 13 | 558 | 29 | SBK1 | SH3 domain binding kinase 1 | Other | kinase | |
| chr2 | *RANBP2* | 11 | 381 | 29 | RANBP2 | RAN binding protein 2 | Nucleus | enzyme | |
| chr19 | *ZNF787* | 13 | 597 | 29 | ZNF787 | zinc finger protein 787 | Other | other | |
| chr2 | *USP39* | 13 | 331 | 29 | USP39 | ubiquitin specific peptidase 39 | Nucleus | peptidase | |
| chr11 | *ADAMTS15* | 13 | 716 | 29 | ADAMTS15 | ADAM metallopeptidase with thrombospondin type 1 motif 15 | Extracellular Space | peptidase | |
| chr17 | *CHD3* | 13 | 474 | 29 | CHD3 | chromodomain helicase DNA binding protein 3 | Nucleus | enzyme | |
| chr19 | *PPP6R1* | 14 | 622 | 29 | PPP6R1 | protein phosphatase 6 regulatory subunit 1 | Cytoplasm | other | |
| chr7 | *SLC29A4* | 14 | 716 | 29 | SLC29A4 | solute carrier family 29 member 4 | Plasma Membrane | transporter | |
| chr15 | *CHRNB4* | 13 | 593 | 29 | CHRNB4 | cholinergic receptor nicotinic beta 4 subunit | Plasma Membrane | transmembrane receptor | succinylcholine, rapacuronium, doxacurium, vecuronium, pipecuronium, metocurine, mivacurium, D-tubocurarine, nicotine, arecoline, pancuronium, atracurium, isoflurane, cisatracurium, rocuronium, ibogaine |
| chr7 | *ZNRF2* | 11 | 301 | 29 | ZNRF2 | zinc and ring finger 2 | Cytoplasm | enzyme | |
| chr1 | *PRRC2C* | 12 | 355 | 29 | PRRC2C | proline rich coiled-coil 2C | Cytoplasm | other | |
| chr9 | *LINGO2* | 10 | 275 | 29 | LINGO2 | leucine rich repeat and Ig domain containing 2 | Extracellular Space | other | |
| chr15 | *RP11-1007O24.3* | 12 | 270 | 29 | | | | | |
| chr3 | *HDAC11* | 12 | 536 | 29 | HDAC11 | histone deacetylase 11 | Nucleus | transcription regulator | tributyrin, belinostat, pyroxamide |
| chr3 | *PCOLCE2* | 13 | 342 | 29 | PCOLCE2 | procollagen C-endopeptidase enhancer 2 | Extracellular Space | other | |
| chr22 | *MOV10L1* | 11 | 459 | 29 | MOV10L1 | Mov10 like RISC complex RNA helicase 1 | Nucleus | enzyme | |
| chr10 | *C10orf68* | 10 | 320 | 29 | CCDC7 | coiled-coil domain containing 7 | Other | other | |
| chr3 | *PODXL2* | 13 | 609 | 29 | PODXL2 | podocalyxin like 2 | Plasma Membrane | other | |
| chr3 | *MAP3K13* | 11 | 241 | 29 | MAP3K13 | mitogen-activated protein kinase kinase kinase 13 | Cytoplasm | kinase | |
| chr16 | *PMM2* | 11 | 351 | 29 | PMM2 | phosphomannomutase 2 | Cytoplasm | enzyme | |
| chr4 | *HTRA3* | 11 | 524 | 29 | HTRA3 | HtrA serine peptidase 3 | Extracellular Space | peptidase | |
| chr19 | *AP2A1* | 12 | 496 | 29 | AP2A1 | adaptor related protein complex 2 subunit alpha 1 | Cytoplasm | transporter | |
| chr6 | *IP6K3* | 12 | 561 | 29 | IP6K3 | inositol hexakisphosphate kinase | Cytoplasm | kinase | |
| chr3 | *NBEAL2* | 14 | 729 | 29 | NBEAL2 | neurobeachin like 2 | Cytoplasm | other | |
| chr17 | *AATF* | 12 | 307 | 29 | AATF | apoptosis antagonizing transcription factor | Nucleus | transcription regulator | |
| chr9 | *PTPN3* | 12 | 298 | 29 | PTPN3 | protein tyrosine phosphatase non-receptor type 3 | Cytoplasm | phosphatase | |
| chr16 | *RP4-536B24.3* | 13 | 517 | 29 | | | | | |
| chr2 | *NCAPH* | 12 | 469 | 29 | NCAPH | non-SMC condensin I complex subunit H | Nucleus | other | |
| chr5 | *FCHO2* | 11 | 257 | 29 | FCHO2 | FCH and mu domain containing endocytic adaptor 2 | Cytoplasm | other | |
| chr12 | *SLC41A2* | 11 | 323 | 29 | SLC41A2 | solute carrier family 41 member 2 | Plasma Membrane | transporter | |
| chr8 | *FAM167A* | 14 | 653 | 29 | FAM167A | family with sequence similarity 167 member A | Cytoplasm | other | |
| chr1 | *LINC01035* | 12 | 346 | 29 | LINC01035 | | Other | other | |
| chr12 | *UHRF1BP1L* | 13 | 255 | 29 | UHRF1BP1L | UHRF1 binding protein 1 like | Cytoplasm | other | |
| chr21 | *TRPM2* | 12 | 599 | 29 | TRPM2 | transient receptor potential cation channel subfamily M member 2 | Plasma Membrane | ion channel | |
| chr17 | *MFSD11* | 12 | 257 | 29 | MFSD11 | major facilitator superfamily domain containing 11 | Other | other | |

| | | | | | | | | | |
|---|---|---|---|---|---|---|---|---|---|
| chr11 | *C11orf65* | 12 | 317 | 29 | C11orf65 | chromosome 11 open reading frame 65 | Other | other | |
| chr1 | *CDA* | 12 | 395 | 29 | CDA | cytidine deaminase | Nucleus | enzyme | cytidine deaminase inhibitor, |
| chr22 | *NF2* | 11 | 355 | 29 | NF2 | neurofibromin 2 | Plasma Membrane | other | |
| chr15 | *FSD2* | 12 | 398 | 29 | FSD2 | fibronectin type III and SPRY domain containing 2 | Other | other | |
| chr5 | *FAM13B* | 11 | 301 | 29 | FAM13B | family with sequence similarity 13 member B | Cytoplasm | other | |
| chr10 | *PKD2L1* | 12 | 311 | 29 | PKD2L1 | polycystin 2 like 1, transient receptor potential cation channel | Plasma Membrane | ion channel | |
| chr9 | *NR5A1* | 14 | 844 | 29 | NR5A1 | nuclear receptor subfamily 5 group A member 1 | Nucleus | d-dependent nuclear rec | |
| chr6 | *RIPK1* | 12 | 320 | 29 | RIPK1 | receptor interacting serine/threonine kinase 1 | Plasma Membrane | kinase | GSK3145095, GSK2982772 |
| chrX | *FAM9B* | 13 | 319 | 29 | FAM9B | family with sequence similarity 9 member B | Nucleus | other | |
| chr20 | *COL20A1* | 15 | 1132 | 29 | COL20A1 | collagen type XX alpha 1 chain | Extracellular Space | other | |
| chr22 | *DGCR8* | 12 | 431 | 29 | DGCR8 | DGCR8 microprocessor complex subunit | Nucleus | enzyme | |
| chr10 | *POLR3A* | 13 | 354 | 29 | POLR3A | RNA polymerase III subunit A | Nucleus | enzyme | |
| chr9 | *CERCAM* | 13 | 767 | 29 | CERCAM | cerebral endothelial cell adhesion molecule | Other | other | |
| chr8 | *PLEKHA2* | 10 | 353 | 29 | PLEKHA2 | pleckstrin homology domain containing A2 | Cytoplasm | other | |
| chr19 | *SLC1A6* | 13 | 371 | 29 | SLC1A6 | solute carrier family 1 member 6 | Plasma Membrane | transporter | riluzole |
| chr19 | *AC011524.1* | 13 | 370 | 29 | | | | | |
| chr5 | *CYFIP2* | 14 | 434 | 29 | CYFIP2 | cytoplasmic FMR1 interacting protein 2 | Cytoplasm | other | |
| chr1 | *DLGAP3* | 13 | 465 | 29 | DLGAP3 | DLG associated protein 3 | Cytoplasm | other | |
| chr17 | *NDEL1* | 11 | 375 | 29 | NDEL1 | nudE neurodevelopment protein 1 like 1 | Nucleus | peptidase | |
| chr18 | *MAPK4* | 12 | 297 | 29 | MAPK4 | mitogen-activated protein kinase | Cytoplasm | kinase | |
| chr11 | *LMO1* | 13 | 654 | 29 | LMO1 | LIM domain only 1 | Nucleus | transcription regulator | |
| chr12 | *NTF3* | 11 | 318 | 29 | NTF3 | neurotrophin 3 | Extracellular Space | growth factor | |
| chr4 | *CCDC149* | 12 | 272 | 29 | CCDC149 | coiled-coil domain containing 149 | Other | other | |
| chr12 | *RNF10* | 12 | 324 | 29 | RNF10 | ring finger protein 10 | Cytoplasm | enzyme | |
| chr16 | *TMEM219* | 12 | 399 | 29 | TMEM219 | transmembrane protein 219 | Plasma Membrane | other | |
| chr19 | *PALM* | 14 | 534 | 29 | PALM | paralemmin | Plasma Membrane | other | |
| chr5 | *TCOF1* | 11 | 442 | 29 | TCOF1 | treacle ribosome biogenesis | Nucleus | transporter | |
| chr3 | *PPP4R2* | 11 | 261 | 29 | PPP4R2 | protein phosphatase 4 regulatory subunit 2 | Cytoplasm | other | |
| chr7 | *BLVRA* | 13 | 424 | 28 | BLVRA | biliverdin reductase A | Cytoplasm | enzyme | |
| chr20 | *RP5-1112F19.2* | 13 | 417 | 28 | | | | | |
| chr1 | *RP11-382E9.1* | 12 | 358 | 28 | | | | | |
| chr1 | *SWT1* | 11 | 348 | 28 | SWT1 | SWT1 RNA endoribonuclease homolog | Extracellular Space | other | |
| chr8 | *ST18* | 11 | 283 | 28 | ST18 | ST18 C2H2C-type zinc finger transcription factor | Nucleus | transcription regulator | |
| chr6 | *UNC93A* | 12 | 764 | 28 | UNC93A | unc-93 homolog A | Plasma Membrane | other | |
| chr19 | *ADAMTS10* | 13 | 529 | 28 | ADAMTS10 | ADAM metallopeptidase with thrombospondin type 1 motif 10 | Extracellular Space | peptidase | |
| chrX | *RP11-342D14.1* | 11 | 328 | 28 | | | | | |
| chr3 | *DLEC1* | 13 | 375 | 28 | DLEC1 | DLEC1 cilia and flagella associated protein | Cytoplasm | other | |
| chr2 | *RGPD6* | 10 | 391 | 28 | PD4 (includes oth | RANBP2 like and GRIP domain containing 5 | Nucleus | enzyme | |
| chr19 | *SUGP2* | 12 | 421 | 28 | SUGP2 | SURP and G-patch domain containing 2 | Nucleus | other | |
| chr19 | *TMEM143* | 12 | 396 | 28 | TMEM143 | transmembrane protein 143 | Cytoplasm | other | |
| chr9 | *UBAC1* | 14 | 817 | 28 | UBAC1 | UBA domain containing 1 | Nucleus | other | |
| chr10 | *PARG* | 12 | 352 | 28 | PARG | poly(ADP-ribose) glycohydrolase | Cytoplasm | enzyme | |
| chr5 | *MRPS27* | 11 | 282 | 28 | MRPS27 | mitochondrial ribosomal protein | Cytoplasm | other | |
| chr12 | *PDZRN4* | 13 | 419 | 28 | PDZRN4 | PDZ domain containing ring | Other | other | |
| chr6 | *SGK1* | 10 | 265 | 28 | SGK1 | serum/glucocorticoid regulated kinase 1 | Cytoplasm | kinase | |
| chr1 | *TTC13* | 11 | 364 | 28 | TTC13 | tetratricopeptide repeat domain | Other | other | |
| chr1 | *RP11-31F15.1* | 12 | 381 | 28 | | | | | |
| chr1 | *EPRS* | 10 | 311 | 28 | EPRS1 | glutamyl-prolyl-tRNA synthetase | Cytoplasm | enzyme | |
| chr7 | *MGAM* | 13 | 355 | 28 | MGAM | maltase-glucoamylase | Plasma Membrane | enzyme | celgosivir, voglibose, acarbose, miglitol |
| chr1 | *PTPRVP* | 11 | 834 | 28 | PTPRVP | protein tyrosine phosphatase receptor type V, pseudogene | Other | other | |
| chr9 | *TRPM6* | 12 | 309 | 28 | TRPM6 | transient receptor potential cation channel subfamily M member 6 | Plasma Membrane | kinase | |
| chr8 | *PTP4A3* | 14 | 838 | 28 | PTP4A3 | protein tyrosine phosphatase 4A3 | Plasma Membrane | phosphatase | |
| chr21 | *TMPRSS15* | 12 | 261 | 28 | TMPRSS15 | transmembrane serine protease | Extracellular Space | peptidase | |
| chr13 | *TMTC4* | 12 | 472 | 28 | TMTC4 | transmembrane O-mannosyltransferase targeting cadherins 4 | Other | other | |
| chr5 | *PWWP2A* | 13 | 352 | 28 | PWWP2A | PWWP domain containing 2A | Nucleus | other | |
| chr6 | *FGD2* | 13 | 601 | 28 | FGD2 | FYVE, RhoGEF and PH domain containing 2 | Cytoplasm | other | |
| chr16 | *PLLP* | 12 | 518 | 28 | PLLP | plasmolipin | Plasma Membrane | transporter | |
| chr15 | *RFX7* | 10 | 267 | 28 | RFX7 | regulatory factor X7 | Other | other | |
| chr14 | *HHIPL1* | 12 | 525 | 28 | HHIPL1 | HHIP like 1 | Other | other | |
| chr12 | *TCTN2* | 12 | 339 | 28 | TCTN2 | tectonic family member 2 | Extracellular Space | other | |
| chr2 | *IMMT* | 12 | 283 | 28 | IMMT | inner membrane mitochondrial | Cytoplasm | other | |

| | | | | | | | | | |
|---|---|---|---|---|---|---|---|---|---|
| chr16 | *DNAJA3* | 12 | 445 | 28 | DNAJA3 | DnaJ heat shock protein family (Hsp40) member A3 | Cytoplasm | other | |
| chr1 | *SLC2A7* | 12 | 466 | 28 | SLC2A7 | solute carrier family 2 member 7 | Plasma Membrane | transporter | |
| chr2 | *SPAG16* | 9 | 277 | 28 | SPAG16 | sperm associated antigen 16 | Cytoplasm | other | |
| chr20 | *RPRD1B* | 12 | 368 | 28 | RPRD1B | regulation of nuclear pre-mRNA domain containing 1B | Nucleus | other | |
| chr19 | *PLVAP* | 12 | 390 | 28 | PLVAP | plasmalemma vesicle associated protein | Plasma Membrane | other | |
| chr13 | *SLC15A1* | 13 | 458 | 28 | SLC15A1 | solute carrier family 15 member 1 | Plasma Membrane | transporter | |
| chr12 | *RP11-638F5.1* | 12 | 519 | 28 | | | | | |
| chr18 | *IMPA2* | 10 | 359 | 28 | IMPA2 | inositol monophosphatase 2 | Cytoplasm | phosphatase | |
| chr9 | *RFX3* | 12 | 254 | 28 | RFX3 | regulatory factor X3 | Nucleus | transcription regulator | |
| chr9 | *GGTA1P* | 13 | 438 | 28 | GGTA1P | glycoprotein alpha-galactosyltransferase 1, | Cytoplasm | other | |
| chr10 | *ECD* | 10 | 382 | 28 | ECD | ecdysoneless cell cycle regulator | Nucleus | transcription regulator | |
| chr1 | *IKBKE* | 11 | 700 | 28 | IKBKE | inhibitor of nuclear factor kappa B kinase subunit epsilon | Cytoplasm | kinase | |
| chr15 | *UACA* | 10 | 259 | 28 | UACA | uveal autoantigen with coiled-coil domains and ankyrin repeats | Cytoplasm | other | |
| chr17 | *USP22* | 12 | 426 | 28 | USP22 | ubiquitin specific peptidase 22 | Nucleus | peptidase | |
| chr11 | *TMEM45B* | 12 | 451 | 28 | TMEM45B | transmembrane protein 45B | Extracellular Space | other | |
| chr3 | *SLC9C1* | 12 | 356 | 28 | SLC9C1 | solute carrier family 9 member C1 | Extracellular Space | other | |
| chr12 | *RP11-144F15.1* | 11 | 335 | 28 | | | | | |
| chr16 | *BFAR* | 13 | 372 | 28 | BFAR | bifunctional apoptosis regulator | Cytoplasm | enzyme | |
| chr3 | *ESYT3* | 13 | 510 | 28 | ESYT3 | extended synaptotagmin 3 | Plasma Membrane | other | |
| chr15 | *SPG11* | 10 | 282 | 28 | SPG11 | SPG11 vesicle trafficking associated, spatacsin | Cytoplasm | other | |
| chr7 | *CLDN4* | 13 | 432 | 28 | CLDN4 | claudin 4 | Plasma Membrane | transmembrane receptor | |
| chr8 | *KHDRBS3* | 12 | 258 | 28 | KHDRBS3 | KH RNA binding domain containing, signal transduction | Nucleus | other | |
| chr17 | *NARF* | 11 | 482 | 28 | NARF | nuclear prelamin A recognition | Nucleus | enzyme | |
| chr3 | *HMCES* | 11 | 515 | 28 | HMCES | 5-hydroxymethylcytosine binding, ES cell specific | Nucleus | other | |
| chr20 | *CHMP4B* | 13 | 486 | 28 | CHMP4B | charged multivesicular body protein 4B | Cytoplasm | other | |
| chr12 | *RP4-601P9.2* | 12 | 315 | 28 | | | | | |
| chr17 | *SLC47A2* | 12 | 451 | 28 | SLC47A2 | solute carrier family 47 member 2 | Plasma Membrane | transporter | |
| chr1 | *RP5-896L10.1* | 12 | 363 | 28 | | | | | |
| chr16 | *CIITA* | 13 | 371 | 28 | CIITA | class II major histocompatibility complex transactivator | Nucleus | transcription regulator | |
| chr10 | *BTAF1* | 10 | 253 | 28 | BTAF1 | B-TFIID TATA-box binding protein associated factor 1 | Nucleus | transcription regulator | |
| chr6 | *BRPF3* | 11 | 423 | 28 | BRPF3 | bromodomain and PHD finger containing 3 | Cytoplasm | other | |
| chr3 | *ABI3BP* | 11 | 281 | 28 | ABI3BP | ABI family member 3 binding | Extracellular Space | other | |
| chr11 | *PAAF1* | 11 | 352 | 28 | PAAF1 | proteasomal ATPase associated factor 1 | Other | other | |
| chr3 | *KIAA1257* | 13 | 453 | 28 | KIAA1257 | KIAA1257 | Other | other | |
| chr17 | *TOB1-AS1* | 13 | 419 | 28 | TOB1-AS1 | TOB1 antisense RNA 1 | Other | other | |
| chrX | *RP11-357C3.3* | 11 | 296 | 28 | | | | | |
| chr3 | *PAK2* | 9 | 255 | 28 | PAK2 | p21 (RAC1) activated kinase 2 | Cytoplasm | kinase | |
| chr3 | *STAB1* | 13 | 625 | 28 | STAB1 | stabilin 1 | Plasma Membrane | transporter | FP-1305 |
| chr17 | *MPP2* | 12 | 549 | 28 | MPP2 | membrane palmitoylated protein | Plasma Membrane | kinase | |
| chr16 | *CTA-363E6.2* | 12 | 296 | 28 | | | | | |
| chr12 | *ETV6* | 11 | 255 | 28 | ETV6 | ETS variant transcription factor 6 | Nucleus | transcription regulator | |
| chr16 | *PKD1* | 13 | 388 | 28 | PKD1 | polycystin 1, transient receptor potential channel interacting | Plasma Membrane | ion channel | |
| chr11 | *C11orf63* | 11 | 265 | 28 | JHY | junctional cadherin complex | Other | other | |
| chr13 | *LINC00351* | 12 | 315 | 28 | LINC00351 | long intergenic non-protein coding RNA 351 | Other | other | |
| chr11 | *DDB1* | 12 | 414 | 28 | DDB1 | damage specific DNA binding protein 1 | Nucleus | other | |
| chr12 | *SLC11A2* | 13 | 490 | 28 | SLC11A2 | solute carrier family 11 member 2 | Plasma Membrane | transporter | |
| chr11 | *CRY2* | 14 | 513 | 28 | CRY2 | cryptochrome circadian regulator | Nucleus | enzyme | |
| chr17 | *KIF18B* | 13 | 658 | 28 | KIF18B | kinesin family member 18B | Cytoplasm | other | |
| chr1 | *CR1L* | 13 | 513 | 28 | CR1L | complement C3b/C4b receptor 1 | Plasma Membrane | other | |
| chr5 | *ANKRD32* | 9 | 300 | 28 | SLF1 | SMC5-SMC6 complex localization factor 1 | Nucleus | transcription regulator | |
| chr17 | *AKAP10* | 11 | 306 | 28 | AKAP10 | A-kinase anchoring protein 10 | Cytoplasm | other | |
| chr17 | *ITGA3* | 11 | 542 | 28 | ITGA3 | integrin subunit alpha 3 | Plasma Membrane | other | |
| chr2 | *MGAT5* | 11 | 341 | 28 | MGAT5 | alpha-1,6-mannosylglycoprotein 6-beta-N- | Cytoplasm | enzyme | |
| chr5 | *KCTD16* | 11 | 263 | 28 | KCTD16 | potassium channel tetramerization domain containing | Plasma Membrane | other | |
| chr1 | *OLFML2B* | 13 | 676 | 28 | OLFML2B | olfactomedin like 2B | Extracellular Space | other | |
| chr9 | *SNAPC4* | 14 | 842 | 28 | SNAPC4 | small nuclear RNA activating complex polypeptide 4 | Nucleus | transcription regulator | |
| chr16 | *ITGAD* | 12 | 447 | 28 | ITGAD | integrin subunit alpha D | Plasma Membrane | other | |
| chr7 | *AC008060.8* | 13 | 491 | 28 | | | | | |
| chr16 | *GGA2* | 14 | 348 | 28 | GGA2 | golgi associated, gamma adaptin ear containing, ARF binding | Cytoplasm | transporter | |
| chr22 | *RBFOX2* | 10 | 246 | 28 | RBFOX2 | RNA binding fox-1 homolog 2 | Nucleus | transcription regulator | |
| chr7 | *ORAI2* | 13 | 466 | 28 | ORAI2 | ORAI calcium release-activated calcium modulator 2 | Plasma Membrane | other | |
| chr1 | *SLC45A1* | 13 | 642 | 28 | SLC45A1 | solute carrier family 45 member 1 | Other | transporter | |
| chr18 | *ST8SIA5* | 10 | 293 | 28 | ST8SIA5 | ST8 alpha-N-acetyl-neuraminide alpha-2,8-sialyltransferase 5 | Cytoplasm | enzyme | |

| | | | | | | | | | |
|---|---|---|---|---|---|---|---|---|---|
| chr1 | *PARP1* | 12 | 520 | 28 | PARP1 | poly(ADP-ribose) polymerase 1 | Nucleus | enzyme | SC10914, talazoparib, simmiparib, olaparib, ABT-767, rucaparib, veliparib, poly ADP ribose polymerase 1 inhibitor, CEP-9722, niraparib, [18F]fluorthanatrace, 2X-121, INO-1001, |
| chrX | *SPANXA2-OT1* | 10 | 299 | 28 | SPANXA2-OT1 | SPANXA2 overlapping transcript | Other | other | |
| chr19 | *SLC5A5* | 13 | 453 | 28 | SLC5A5 | solute carrier family 5 member 5 | Plasma Membrane | transporter | |
| chrX | *ZNF630* | 12 | 377 | 28 | ZNF630 | zinc finger protein 630 | Other | other | |
| chr2 | *SPEG* | 13 | 563 | 28 | SPEG | striated muscle enriched protein kinase | Nucleus | kinase | |
| chr2 | *LRPPRC* | 10 | 327 | 28 | LRPPRC | leucine rich pentatricopeptide repeat containing | Cytoplasm | other | |
| chr3 | *CHDH* | 11 | 486 | 28 | CHDH | choline dehydrogenase | Cytoplasm | enzyme | |
| chr7 | *RAC1* | 12 | 400 | 28 | RAC1 | Rac family small GTPase 1 | Plasma Membrane | enzyme | |
| chr16 | *CCDC79* | 11 | 282 | 28 | TERB1 | telomere repeat binding bouquet formation protein 1 | Nucleus | peptidase | |
| chr2 | *RAB1A* | 11 | 261 | 28 | RAB1A | RAB1A, member RAS oncogene family | Cytoplasm | enzyme | |
| chr19 | *ZNF586* | 12 | 320 | 28 | ZNF586 | zinc finger protein 586 | Other | other | |
| chr19 | *CD97* | 12 | 460 | 28 | ADGRE5 | adhesion G protein-coupled receptor E5 | Plasma Membrane | G-protein coupled recepto | |
| chr12 | *POLR3B* | 11 | 287 | 28 | POLR3B | RNA polymerase III subunit B | Nucleus | enzyme | |
| chr17 | *TRIM16L* | 10 | 333 | 28 | TRIM16L | tripartite motif containing 16 like | Other | other | |
| chr12 | *HIP1R* | 13 | 715 | 28 | HIP1R | huntingtin interacting protein 1 | Cytoplasm | other | |
| chr7 | *INTS1* | 14 | 554 | 28 | INTS1 | integrator complex subunit 1 | Nucleus | other | |
| chr5 | *TERT* | 15 | 1469 | 28 | TERT | telomerase reverse transcriptase | Nucleus | enzyme | imetelstat, telomerase inhibitor |
| chr12 | *RAD9B* | 11 | 283 | 28 | RAD9B | RAD9 checkpoint clamp component B | Nucleus | other | |
| chr12 | *FAM216A* | 11 | 291 | 28 | FAM216A | family with sequence similarity 216 member A | Other | other | |
| chr16 | *SF3B3* | 11 | 324 | 28 | SF3B3 | splicing factor 3b subunit 3 | Nucleus | other | |
| chr17 | *AZI1* | 13 | 564 | 28 | CEP131 | centrosomal protein 131 | Cytoplasm | other | |
| chr1 | *TDRD10* | 13 | 460 | 28 | TDRD10 | tudor domain containing 10 | Other | other | |
| chr7 | *GS1-179L18.1* | 12 | 376 | 28 | | | | | |
| chr19 | *ZNF578* | 14 | 304 | 28 | ZNF578 | zinc finger protein 578 | Nucleus | other | |
| chr18 | *CTD-2130O13.1* | 11 | 296 | 28 | | | | | |
| chr20 | *BPIFB4* | 12 | 573 | 28 | BPIFB4 | BPI fold containing family B member 4 | Extracellular Space | other | |
| chrX | *ZNF674* | 12 | 286 | 28 | ZNF674 | zinc finger protein 674 | Other | other | |
| chr19 | *ZNF333* | 12 | 340 | 28 | ZNF333 | zinc finger protein 333 | Nucleus | transcription regulator | |
| chr3 | *RP11-319G6.1* | 12 | 381 | 28 | | | | | |
| chr5 | *RP11-844P9.2* | 11 | 511 | 28 | | | | | |
| chr7 | *KLRG2* | 12 | 517 | 28 | KLRG2 | killer cell lectin like receptor G2 | Other | other | |
| chr12 | *RP11-103B5.2* | 12 | 379 | 28 | | | | | |
| chr2 | *TRPM8* | 9 | 284 | 28 | TRPM8 | transient receptor potential cation channel subfamily M member 8 | Plasma Membrane | ion channel | D-3263 |
| chr3 | *GMPS* | 11 | 276 | 28 | GMPS | guanine monophosphate synthase | Nucleus | enzyme | glutamine amidotransferase inhibitor |
| chr9 | *BICD2* | 10 | 450 | 28 | BICD2 | BICD cargo adaptor 2 | Cytoplasm | other | |
| chr7 | *GLCCI1* | 10 | 334 | 28 | GLCCI1 | glucocorticoid induced 1 | Cytoplasm | other | |
| chr12 | *NAV3* | 12 | 263 | 28 | NAV3 | neuron navigator 3 | Nucleus | other | |
| chr5 | *TRPC7* | 14 | 302 | 28 | TRPC7 | transient receptor potential cation channel subfamily C member 7 | Plasma Membrane | ion channel | |
| chr11 | *SMCO4* | 13 | 775 | 28 | SMCO4 | single-pass membrane protein with coiled-coil domains 4 | Other | other | |
| chr5 | *RP11-60A8.1* | 12 | 321 | 28 | | | | | |
| chr9 | *C5* | 11 | 413 | 28 | C5 | complement C5 | Extracellular Space | cytokine | IgG, eculizumab, ravulizumab, IFX-1 |
| chr9 | *QSOX2* | 13 | 789 | 28 | QSOX2 | quiescin sulfhydryl oxidase 2 | Other | enzyme | |
| chr17 | *DLG4* | 12 | 565 | 28 | DLG4 | discs large MAGUK scaffold | Plasma Membrane | kinase | |
| chr2 | *RNF144A* | 10 | 346 | 28 | RNF144A | ring finger protein 144A | Nucleus | enzyme | |
| chr15 | *FSIP1* | 12 | 252 | 28 | FSIP1 | fibrous sheath interacting protein | Other | other | |
| chr4 | *RP11-420A23.1* | 10 | 261 | 28 | | | | | |
| chr5 | *UBE2QL1* | 13 | 1258 | 28 | UBE2QL1 | ubiquitin conjugating enzyme E2 Q family like 1 | Nucleus | enzyme | |
| chr17 | *MAP2K3* | 13 | 1112 | 28 | MAP2K3 | mitogen-activated protein kinase kinase 3 | Cytoplasm | kinase | |
| chr7 | *MUC17* | 13 | 368 | 28 | MUC17 | mucin 17, cell surface associated | Plasma Membrane | other | |
| chr11 | *DDX6* | 11 | 291 | 28 | DDX6 | DEAD-box helicase 6 | Nucleus | enzyme | |
| chr22 | *FAM83F* | 11 | 422 | 28 | FAM83F | family with sequence similarity 83 member F | Other | other | |
| chr12 | *C12orf76* | 11 | 321 | 28 | C12orf76 | chromosome 12 open reading frame 76 | Other | other | |
| chr15 | *USP8* | 11 | 269 | 28 | USP8 | ubiquitin specific peptidase 8 | Cytoplasm | peptidase | |
| chr8 | *NCALD* | 12 | 265 | 28 | NCALD | neurocalcin delta | Cytoplasm | other | |
| chr16 | *PKD1L3* | 12 | 317 | 28 | PKD1L3 | polycystin 1 like 3, transient receptor potential channel | Plasma Membrane | ion channel | |
| chr13 | *LATS2* | 10 | 307 | 28 | LATS2 | large tumor suppressor kinase 2 | Nucleus | kinase | |
| chr2 | *LIMS2* | 13 | 699 | 28 | LIMS2 | LIM zinc finger domain containing | Cytoplasm | other | |
| chr5 | *CTD-2023N9.1* | 14 | 276 | 28 | | | | | |
| chr20 | *ZNF335* | 13 | 701 | 28 | ZNF335 | zinc finger protein 335 | Nucleus | transcription regulator | |

| | | | | | | | | | |
|---|---|---|---|---|---|---|---|---|---|
| chr11 | *SDHD* | 12 | 264 | 28 | SDHD | succinate dehydrogenase complex subunit D | Cytoplasm | enzyme | hexachlorophene |
| chr19 | *CCDC130* | 12 | 375 | 28 | CCDC130 | coiled-coil domain containing 130 | Other | other | |
| chr7 | *INMT* | 13 | 566 | 28 | INMT | indolethylamine N- | Cytoplasm | enzyme | |
| chr7 | *TMEM130* | 12 | 588 | 28 | TMEM130 | transmembrane protein 130 | Cytoplasm | other | |
| chr7 | *JHDM1D* | 11 | 310 | 28 | KDM7A | lysine demethylase 7A | Nucleus | enzyme | |
| chr17 | *PIP4K2B* | 12 | 377 | 28 | PIP4K2B | phosphatidylinositol-5-phosphate 4-kinase type 2 beta | Cytoplasm | kinase | |
| chr1 | *DNAJC16* | 12 | 301 | 27 | DNAJC16 | DnaJ heat shock protein family (Hsp40) member C16 | Other | other | |
| chr12 | *RP11-486A14.2* | 11 | 252 | 27 | | | | | |
| chr11 | *NAT10* | 11 | 439 | 27 | NAT10 | N-acetyltransferase 10 | Nucleus | enzyme | |
| chr12 | *RP11-690J15.1* | 13 | 245 | 27 | | | | | |
| chr1 | *KIAA1522* | 13 | 428 | 27 | KIAA1522 | KIAA1522 | Other | other | |
| chr17 | *GPRC5C* | 12 | 525 | 27 | GPRC5C | G protein-coupled receptor class C group 5 member C | Plasma Membrane | G-protein coupled recepto | |
| chr15 | *RP11-507B12.2* | 11 | 259 | 27 | | | | | |
| chr17 | *CDR2L* | 12 | 492 | 27 | CDR2L | cerebellar degeneration related protein 2 like | Other | other | |
| chr16 | *ZFP90* | 12 | 275 | 27 | ZFP90 | ZFP90 zinc finger protein | Nucleus | transcription regulator | |
| chr1 | *OLFM3* | 11 | 316 | 27 | OLFM3 | olfactomedin 3 | Cytoplasm | other | |
| chr6 | *CMTR1* | 12 | 322 | 27 | CMTR1 | cap methyltransferase 1 | Nucleus | enzyme | |
| chr10 | *RP11-540N6.1* | 13 | 451 | 27 | | | | | |
| chr10 | *ZNF487* | 12 | 375 | 27 | ZNF487 | zinc finger protein 487 | Other | other | |
| chr8 | *EXT1* | 11 | 236 | 27 | EXT1 | exostosin glycosyltransferase 1 | Cytoplasm | enzyme | |
| chr7 | *KCNH2* | 13 | 700 | 27 | KCNH2 | potassium voltage-gated channel subfamily H member 2 | Plasma Membrane | ion channel | dofetilide, sotalol, pimozide, quinidine, procainamide, amiodarone, amitriptyline, propafenone, ibutilide, dextromethorphan/qu |
| chr7 | *RP11-514P8.6* | 12 | 836 | 27 | | | | | |
| chr18 | *ZNF532* | 10 | 264 | 27 | ZNF532 | zinc finger protein 532 | Nucleus | transcription regulator | |
| chr1 | *ZC3H11A* | 11 | 347 | 27 | ZC3H11A | zinc finger CCCH-type containing 11A | Nucleus | other | |
| chr16 | *TNRC6A* | 11 | 292 | 27 | TNRC6A | trinucleotide repeat containing adaptor 6A | Nucleus | other | |
| chr1 | *CYP4B1* | 12 | 319 | 27 | CYP4B1 | cytochrome P450 family 4 subfamily B member 1 | Cytoplasm | enzyme | |
| chr10 | *XPNPEP1* | 13 | 373 | 27 | XPNPEP1 | X-prolyl aminopeptidase 1 | Cytoplasm | peptidase | |
| chr3 | *COL7A1* | 12 | 707 | 27 | COL7A1 | collagen type VII alpha 1 chain | Extracellular Space | other | collagenase clostridium |
| chr12 | *RP1-34H18.1* | 12 | 288 | 27 | | | | | |
| chr1 | *ZMYM1* | 12 | 275 | 27 | ZMYM1 | zinc finger MYM-type containing | Other | other | |
| chr5 | *GALNT10* | 13 | 282 | 27 | GALNT10 | polypeptide N-acetylgalactosaminyltransferase | Cytoplasm | enzyme | |
| chr11 | *ARHGAP1* | 13 | 570 | 27 | ARHGAP1 | Rho GTPase activating protein 1 | Cytoplasm | other | |
| chr12 | *RP11-478C19.2* | 11 | 264 | 27 | | | | | |
| chr3 | *RNF123* | 12 | 541 | 27 | RNF123 | ring finger protein 123 | Cytoplasm | other | |
| chr21 | *PDXK* | 13 | 550 | 27 | PDXK | pyridoxal kinase | Cytoplasm | kinase | |
| chr13 | *IPO5* | 11 | 312 | 27 | IPO5 | importin 5 | Nucleus | transporter | |
| chr16 | *CNTNAP4* | 12 | 224 | 27 | CNTNAP4 | contactin associated protein like | Plasma Membrane | other | |
| chr19 | *MIER2* | 12 | 476 | 27 | MIER2 | MIER family member 2 | Nucleus | transcription regulator | |
| chr12 | *CORO1C* | 12 | 300 | 27 | CORO1C | coronin 1C | Cytoplasm | other | |
| chr2 | *HAAO* | 11 | 582 | 27 | HAAO | 3-hydroxyanthranilate 3,4-dioxygenase | Cytoplasm | enzyme | |
| chr1 | *DNAJC8* | 12 | 336 | 27 | DNAJC8 | DnaJ heat shock protein family (Hsp40) member C8 | Nucleus | other | |
| chr1 | *PRPF3* | 11 | 412 | 27 | PRPF3 | pre-mRNA processing factor 3 | Nucleus | other | |
| chr6 | *RSPH9* | 11 | 445 | 27 | RSPH9 | radial spoke head 9 homolog | Cytoplasm | other | |
| chr19 | *SLC44A2* | 12 | 389 | 27 | SLC44A2 | solute carrier family 44 member 2 | Extracellular Space | transporter | |
| chr5 | *SLC6A7* | 10 | 436 | 27 | SLC6A7 | solute carrier family 6 member 7 | Plasma Membrane | transporter | |
| chr16 | *HAGH* | 12 | 514 | 27 | HAGH | hydroxyacylglutathione hydrolase | Cytoplasm | enzyme | |
| chr3 | *IGF2BP2* | 10 | 325 | 27 | IGF2BP2 | insulin like growth factor 2 mRNA binding protein 2 | Cytoplasm | translation regulator | |
| chr10 | *DNAJC1* | 10 | 308 | 27 | DNAJC1 | DnaJ heat shock protein family (Hsp40) member C1 | Cytoplasm | other | |
| chr11 | *INCENP* | 12 | 831 | 27 | INCENP | inner centromere protein | Nucleus | other | |
| chr1 | *PADI6* | 12 | 436 | 27 | PADI6 | peptidyl arginine deiminase 6 | Cytoplasm | enzyme | |
| chr21 | *AL109763.2* | 10 | 275 | 27 | | | | | |
| chr3 | *KAT2B* | 10 | 342 | 27 | KAT2B | lysine acetyltransferase 2B | Nucleus | transcription regulator | |
| chr17 | *GRAPL* | 11 | 459 | 27 | GRAPL | GRB2 related adaptor protein like | Other | other | |
| chr16 | *LINC01081* | 11 | 423 | 27 | LINC01081 | long intergenic non-protein coding RNA 1081 | Other | other | |
| chr9 | *HAUS6* | 12 | 307 | 27 | HAUS6 | HAUS augmin like complex | Cytoplasm | other | |
| chr2 | *ASXL2* | 9 | 255 | 27 | ASXL2 | ASXL transcriptional regulator 2 | Extracellular Space | other | |
| chr15 | *IGDCC3* | 11 | 501 | 27 | IGDCC3 | immunoglobulin superfamily DCC subclass member 3 | Other | other | |
| chr20 | *DDX27* | 11 | 507 | 27 | DDX27 | DEAD-box helicase 27 | Nucleus | enzyme | |
| chrX | *ZNF41* | 12 | 330 | 27 | ZNF41 | zinc finger protein 41 | Nucleus | transcription regulator | |
| chr19 | *TMEM150B* | 11 | 404 | 27 | TMEM150B | transmembrane protein 150B | Plasma Membrane | other | |
| chr6 | *LHFPL5* | 11 | 312 | 27 | LHFPL5 | LHFPL tetraspan subfamily member 5 | Plasma Membrane | other | |
| chr4 | *SH3D19* | 9 | 254 | 27 | SH3D19 | SH3 domain containing 19 | Plasma Membrane | other | |
| chr2 | *THAP4* | 10 | 533 | 27 | THAP4 | THAP domain containing 4 | Nucleus | transcription regulator | |
| chr19 | *PRKACA* | 11 | 460 | 27 | PRKACA | protein kinase cAMP-activated catalytic subunit alpha | Cytoplasm | kinase | |

| | | | | | | | | | |
|---|---|---|---|---|---|---|---|---|---|
| chr14 | *LIN52* | 10 | 241 | 27 | LIN52 | lin-52 DREAM MuvB core complex component | Nucleus | other | |
| chr12 | *HPD* | 12 | 443 | 27 | HPD | 4-hydroxyphenylpyruvate dioxygenase | Cytoplasm | enzyme | nitisinone |
| chr5 | *GABRA6* | 11 | 331 | 27 | GABRA6 | gamma-aminobutyric acid type A receptor alpha6 subunit | Plasma Membrane | ion channel | flunitrazepam, sevoflurane, butobarbital, atropine/hyoscyamine/phenobarbital/scopolamine, phenobarbital, diazepam, barbital, acetaminophen/butalbital/caffeine/codeine, hyoscyamine/phenobarbital, desflurane, SEP 174559, zaleplon, mephobarbital, ezogabine, flurazepam, butabarbital, midazolam, temazepam, diphenhydramine/midazolam, alphadolone, [illegible] |
| chr12 | *KIF5A* | 12 | 335 | 27 | KIF5A | kinesin family member 5A | Cytoplasm | transporter | |
| chr15 | *FAM227B* | 10 | 301 | 27 | FAM227B | family with sequence similarity 227 member B | Other | other | |
| chr11 | *DGAT2* | 13 | 546 | 27 | DGAT2 | diacylglycerol O-acyltransferase 2 | Cytoplasm | enzyme | |
| chr17 | *MPP3* | 11 | 433 | 27 | MPP3 | membrane palmitoylated protein | Plasma Membrane | kinase | |
| chr11 | *PLA2G16* | 11 | 358 | 27 | PLAAT3 | phospholipase A and acyltransferase 3 | Nucleus | enzyme | |
| chr22 | *SFI1* | 11 | 357 | 27 | SFI1 | SFI1 centrin binding protein | Cytoplasm | other | |
| chr14 | *NUBPL* | 11 | 282 | 27 | NUBPL | nucleotide binding protein like | Cytoplasm | other | |
| chr14 | *JDP2* | 11 | 398 | 27 | JDP2 | Jun dimerization protein 2 | Nucleus | transcription regulator | |
| chrX | *RP3-410B11.1* | 12 | 266 | 27 | | | | | |
| chr17 | *FBXO47* | 9 | 273 | 27 | FBXO47 | F-box protein 47 | Other | other | |
| chr3 | *MFI2* | 11 | 537 | 27 | MELTF | melanotransferrin | Plasma Membrane | other | SGN-CD228A |
| chr6 | *ESR1* | 11 | 273 | 27 | ESR1 | estrogen receptor 1 | Nucleus | d-dependent nuclear rec | desogestrel, clomiphene, estrone sulfate, progesterone, bazedoxifene, exemestane/fulvestrant/goserelin/letrozole/tamoxifen, SRN-927, CC8490, sulindac/toremifene, synthetic conjugated estrogens, B, esterified estrogens, exemestane/fulvestrant/tamoxifen, exemestane/fulvestrant, MITO-4509, dienestrol, estradiol cypionate/testosterone cypionate, estradiol cypionate/medroxyprogesterone acetate, acolbifene, D-0502, [illegible] |
| chr3 | *MBNL1* | 11 | 293 | 27 | MBNL1 | muscleblind like splicing regulator | Nucleus | other | |
| chr15 | *SNX1* | 10 | 336 | 27 | SNX1 | sorting nexin 1 | Cytoplasm | transporter | |
| chr2 | *TRAF3IP1* | 10 | 331 | 27 | TRAF3IP1 | TRAF3 interacting protein 1 | Cytoplasm | other | |
| chr1 | *C1orf177* | 12 | 438 | 27 | LEXM | lymphocyte expansion molecule | Cytoplasm | other | |
| chr22 | *SREBF2* | 10 | 353 | 27 | SREBF2 | sterol regulatory element binding transcription factor 2 | Nucleus | transcription regulator | |
| chr19 | *ZFP30* | 10 | 293 | 27 | ZFP30 | ZFP30 zinc finger protein | Nucleus | transcription regulator | |
| chr1 | *SLC26A9* | 13 | 689 | 27 | SLC26A9 | solute carrier family 26 member 9 | Plasma Membrane | transporter | |
| chr11 | *PDGFD* | 12 | 244 | 27 | PDGFD | platelet derived growth factor D | Extracellular Space | growth factor | |

| | | | | | | | | | |
|---|---|---|---|---|---|---|---|---|---|
| chr19 | *MAP2K2* | 12 | 479 | 27 | MAP2K2 | mitogen-activated protein kinase kinase 2 | Cytoplasm | kinase | dabrafenib/panitumumab/trametinib, dabrafenib/trametinib, trametinib/vemurafenib, dabrafenib/trametinib/vemurafenib, AS703988, TAK 733, binimetinib/cetuximab/encorafenib, binimetinib/encorafenib/panitumumab, PD184352, cetuximab/dabrafenib/trametinib, binimetinib, dabrafenib/ipilimumab/trametinib, trametinib, cobimetinib/vemurafenib, binimetinib/encorafen |
| chr19 | *NLRP2* | 12 | 372 | 27 | NLRP2 | NLR family pyrin domain | Nucleus | other | |
| chr11 | *SLC3A2* | 13 | 405 | 27 | SLC3A2 | solute carrier family 3 member 2 | Plasma Membrane | transporter | |
| chr16 | *QPRT* | 11 | 416 | 27 | QPRT | quinolinate phosphoribosyltransferase | Cytoplasm | enzyme | lovastatin/niacin, nicotinic acid/pioglitazone, atorvastatin/niacin, nicotinic acid |
| chr6 | *SYNE1* | 10 | 246 | 27 | SYNE1 | spectrin repeat containing nuclear envelope protein 1 | Nucleus | other | |
| chr17 | *ENDOV* | 13 | 601 | 27 | ENDOV | endonuclease V | Cytoplasm | enzyme | |
| chr12 | *TMPRSS12* | 12 | 412 | 27 | TMPRSS12 | transmembrane serine protease | Other | peptidase | |
| chr11 | *DKK3* | 12 | 697 | 27 | DKK3 | dickkopf WNT signaling pathway inhibitor 3 | Extracellular Space | cytokine | |
| chr20 | *FAM182A* | 11 | 570 | 27 | FAM182A | family with sequence similarity 182 member A | Other | other | |
| chrX | *SHROOM4* | 13 | 242 | 27 | SHROOM4 | shroom family member 4 | Plasma Membrane | other | |
| chr8 | *ERI1* | 13 | 277 | 27 | ERI1 | exoribonuclease 1 | Nucleus | enzyme | |
| chr15 | *DNM1P47* | 11 | 2183 | 27 | DNM1P47 | dynamin 1 pseudogene 47 | Other | other | |
| chr18 | *RP11-78F17.1* | 13 | 237 | 27 | | | | | |
| chr16 | *CYB5B* | 9 | 299 | 27 | CYB5B | cytochrome b5 type B | Cytoplasm | enzyme | |
| chr20 | *KIF16B* | 9 | 279 | 27 | KIF16B | kinesin family member 16B | Cytoplasm | enzyme | |
| chr3 | *SUSD5* | 10 | 292 | 27 | SUSD5 | sushi domain containing 5 | Other | other | |
| chr11 | *TMPRSS4-AS1* | 11 | 350 | 27 | SMIM35 | small integral membrane protein | Other | other | |
| chr1 | *COLGALT2* | 13 | 398 | 27 | COLGALT2 | collagen beta(1-O)galactosyltransferase 2 | Cytoplasm | enzyme | |
| chr7 | *RAMP3* | 12 | 599 | 27 | RAMP3 | receptor activity modifying protein | Plasma Membrane | other | pramlintide |
| chrX | *BCORL1* | 11 | 357 | 27 | BCORL1 | BCL6 corepressor like 1 | Nucleus | transcription regulator | |
| chr1 | *ZZZ3* | 11 | 266 | 27 | ZZZ3 | zinc finger ZZ-type containing 3 | Nucleus | transcription regulator | |
| chr19 | *C19orf47* | 11 | 463 | 27 | C19orf47 | chromosome 19 open reading frame 47 | Nucleus | other | |
| chr12 | *RP11-510I5.4* | 11 | 280 | 27 | | | | | |
| chr20 | *RP11-445H22.4* | 11 | 607 | 27 | | | | | |
| chr5 | *SLC9A3* | 15 | 1108 | 27 | SLC9A3 | solute carrier family 9 member A3 | Plasma Membrane | ion channel | tenapanor, AVE0657, squalamine |
| chr10 | *RP11-222A11.1* | 11 | 382 | 27 | | | | | |
| chr17 | *DHX8* | 12 | 332 | 27 | DHX8 | DEAH-box helicase 8 | Nucleus | enzyme | |
| chr19 | *CLPTM1* | 11 | 383 | 27 | CLPTM1 | CLPTM1 regulator of GABA type A receptor forward trafficking | Plasma Membrane | other | |
| chr5 | *CTC-340D7.1* | 11 | 279 | 27 | | | | | |
| chr11 | *PRSS23* | 13 | 509 | 27 | PRSS23 | serine protease 23 | Extracellular Space | peptidase | |
| chr3 | *MME* | 12 | 309 | 27 | MME | membrane metalloendopeptidase | Plasma Membrane | peptidase | sacubitril, LBQ657, sacubitril/valsartan |
| chr3 | *ALS2CL* | 12 | 570 | 27 | ALS2CL | ALS2 C-terminal like | Cytoplasm | other | |
| chr1 | *LAD1* | 12 | 664 | 27 | LAD1 | ladinin 1 | Extracellular Space | other | |
| chr6 | *TAF8* | 12 | 345 | 27 | TAF8 | TATA-box binding protein associated factor 8 | Nucleus | transcription regulator | |
| chr19 | *JAK3* | 12 | 498 | 27 | JAK3 | Janus kinase 3 | Cytoplasm | kinase | PF-06651600, tofacitinib, R-348, methotrexate/tofaciti |
| chr15 | *LINC00927* | 11 | 475 | 27 | LINC00927 | long intergenic non-protein coding RNA 927 | Other | other | |
| chr19 | *MYBPC2* | 12 | 426 | 27 | MYBPC2 | myosin binding protein C, fast | Cytoplasm | other | |
| chr6 | *PPP1R14C* | 9 | 267 | 27 | PPP1R14C | protein phosphatase 1 regulatory inhibitor subunit 14C | Cytoplasm | other | |
| chr3 | *CELSR3* | 12 | 491 | 27 | CELSR3 | cadherin EGF LAG seven-pass G-type receptor 3 | Plasma Membrane | G-protein coupled recepto | |
| chr19 | *AC005307.3* | 13 | 284 | 27 | | | | | |
| chr10 | *FAM160B1* | 11 | 267 | 27 | FAM160B1 | family with sequence similarity 160 member B1 | Other | other | |
| chr8 | *RP11-136O12.2* | 12 | 283 | 27 | | | | | |
| chr1 | *DNAJC6* | 11 | 319 | 27 | DNAJC6 | DnaJ heat shock protein family (Hsp40) member C6 | Cytoplasm | other | |
| chr8 | *EPHX2* | 12 | 344 | 27 | EPHX2 | epoxide hydrolase 2 | Cytoplasm | enzyme | |
| chr2 | *AC074289.1* | 11 | 256 | 27 | | | | | |
| chr12 | *KRT80* | 11 | 983 | 27 | KRT80 | keratin 80 | Cytoplasm | other | |
| chr12 | *USP30* | 11 | 294 | 27 | USP30 | ubiquitin specific peptidase 30 | Cytoplasm | peptidase | |

| | | | | | | | | | |
|---|---|---|---|---|---|---|---|---|---|
| chr3 | *HLTF* | 10 | 272 | 27 | HLTF | helicase like transcription factor | Nucleus | transcription regulator | |
| chr1 | *MROH3P* | 13 | 357 | 27 | MROH3P | maestro heat like repeat family member 3, pseudogene | Other | other | |
| chr6 | *SRPK1* | 10 | 276 | 27 | SRPK1 | SRSF protein kinase 1 | Nucleus | kinase | |
| chr9 | *DBH* | 12 | 749 | 27 | DBH | dopamine beta-hydroxylase | Cytoplasm | enzyme | disulfiram, disulfiram/gemcitabin |
| chr22 | *CTA-714B7.5* | 11 | 254 | 27 | | | | | |
| chr9 | *FREM1* | 11 | 275 | 27 | FREM1 | FRAS1 related extracellular | Extracellular Space | other | |
| chr11 | *TRIM66* | 12 | 445 | 27 | TRIM66 | tripartite motif containing 66 | Nucleus | transcription regulator | |
| chr7 | *AC004540.5* | 12 | 333 | 27 | | | | | |
| chr12 | *PAWR* | 11 | 257 | 27 | PAWR | pro-apoptotic WT1 regulator | Nucleus | transcription regulator | |
| chr1 | *PPIE* | 11 | 349 | 27 | PPIE | peptidylprolyl isomerase E | Nucleus | enzyme | |
| chr15 | *RP11-10O17.3* | 11 | 291 | 27 | | | | | |
| chr16 | *CCDC135* | 11 | 424 | 27 | DRC7 | dynein regulatory complex | Cytoplasm | other | |
| chr11 | *NOX4* | 13 | 540 | 27 | NOX4 | NADPH oxidase 4 | Cytoplasm | enzyme | imipramine blue |
| chr3 | *SHQ1* | 12 | 264 | 27 | SHQ1 | SHQ1, H/ACA ribonucleoprotein assembly factor | Nucleus | other | |
| chr15 | *SEMA7A* | 11 | 589 | 27 | SEMA7A | semaphorin 7A (John Milton Hagen blood group) | Plasma Membrane | transmembrane recepto | |
| chr6 | *DUSP22* | 13 | 422 | 27 | DUSP22 | dual specificity phosphatase 22 | Cytoplasm | phosphatase | |
| chr2 | *STEAP3* | 12 | 637 | 27 | STEAP3 | STEAP3 metalloreductase | Cytoplasm | transporter | |
| chr1 | *STPG1* | 11 | 325 | 27 | STPG1 | sperm tail PG-rich repeat | Cytoplasm | other | |
| chr2 | *CD8B* | 11 | 387 | 27 | CD8B | CD8b molecule | Plasma Membrane | other | |
| chr14 | *TMEM229B* | 10 | 320 | 27 | TMEM229B | transmembrane protein 229B | Other | other | |
| chr1 | *ZBTB40* | 11 | 360 | 27 | ZBTB40 | zinc finger and BTB domain containing 40 | Nucleus | transcription regulator | |
| chr2 | *CIB4* | 11 | 420 | 27 | CIB4 | calcium and integrin binding family member 4 | Other | other | |
| chr10 | *DHTKD1* | 9 | 280 | 27 | DHTKD1 | dehydrogenase E1 and transketolase domain containing | Cytoplasm | enzyme | |
| chr12 | *SETD1B* | 13 | 540 | 27 | SETD1B | SET domain containing 1B, histone lysine methyltransferase | Nucleus | enzyme | |
| chr4 | *WFS1* | 11 | 436 | 27 | WFS1 | wolframin ER transmembrane glycoprotein | Cytoplasm | enzyme | |
| chr3 | *LRRC2* | 12 | 280 | 27 | LRRC2 | leucine rich repeat containing 2 | Other | other | |
| chr11 | *ARCN1* | 9 | 286 | 27 | ARCN1 | archain 1 | Cytoplasm | other | |
| chrX | *STS* | 13 | 246 | 27 | STS | steroid sulfatase | Cytoplasm | enzyme | STX64 compound, coumate |
| chr3 | *WDR52* | 11 | 360 | 27 | CFAP44 | cilia and flagella associated | Extracellular Space | other | |
| chr2 | *RGPD1* | 11 | 271 | 27 | PD4 (includes oth | RANBP2 like and GRIP domain containing 5 | Nucleus | enzyme | |
| chr15 | *SCAMP2* | 11 | 341 | 27 | SCAMP2 | secretory carrier membrane | Cytoplasm | other | |
| chr17 | *KCNH6* | 12 | 504 | 27 | KCNH6 | potassium voltage-gated channel subfamily H member 6 | Plasma Membrane | ion channel | |
| chr4 | *SCFD2* | 10 | 299 | 27 | SCFD2 | sec1 family domain containing 2 | Other | transporter | |
| chr22 | *PI4KAP1* | 9 | 353 | 27 | PI4KAP1 | phosphatidylinositol 4-kinase alpha pseudogene 1 | Other | other | |
| chr19 | *LSM14A* | 10 | 261 | 27 | LSM14A | LSM14A mRNA processing body assembly factor | Cytoplasm | other | |
| chr11 | *BCL9L* | 11 | 628 | 27 | BCL9L | BCL9 like | Cytoplasm | other | |
| chr11 | *KMT2A* | 12 | 351 | 27 | KMT2A | lysine methyltransferase 2A | Nucleus | transcription regulator | KO-539 |
| chr1 | *SLC2A1* | 12 | 446 | 27 | SLC2A1 | solute carrier family 2 member 1 | Plasma Membrane | transporter | |
| chr1 | *SESN2* | 11 | 361 | 27 | SESN2 | sestrin 2 | Cytoplasm | enzyme | |
| chr17 | *SKA2* | 10 | 299 | 27 | SKA2 | spindle and kinetochore associated complex subunit 2 | Nucleus | other | |
| chr17 | *ZNF652* | 11 | 250 | 27 | ZNF652 | zinc finger protein 652 | Other | other | |
| chr6 | *RP11-239H6.2* | 12 | 260 | 27 | | | | | |
| chr1 | *SGIP1* | 11 | 264 | 27 | SGIP1 | SH3GL interacting endocytic adaptor 1 | Cytoplasm | other | |
| chr11 | *CHRDL2* | 13 | 558 | 27 | CHRDL2 | chordin like 2 | Cytoplasm | other | |
| chr6 | *KCNK5* | 12 | 434 | 27 | KCNK5 | potassium two pore domain channel subfamily K member 5 | Plasma Membrane | ion channel | |
| chr20 | *C20orf194* | 10 | 279 | 27 | C20orf194 | chromosome 20 open reading frame 194 | Nucleus | other | |
| chr4 | *FIP1L1* | 10 | 282 | 27 | FIP1L1 | factor interacting with PAPOLA and CPSF1 | Nucleus | other | |
| chr16 | *FLYWCH1* | 12 | 469 | 27 | FLYWCH1 | FLYWCH-type zinc finger 1 | Nucleus | other | |
| chr12 | *IFNG-AS1* | 13 | 268 | 27 | IFNG-AS1 | IFNG antisense RNA 1 | Other | other | |
| chr1 | *FAM159A* | 12 | 414 | 27 | SHISAL2A | shisa like 2A | Other | other | |
| chr1 | *CROCCP2* | 14 | 1746 | 27 | CROCCP2 | CROCC pseudogene 2 | Other | other | |

| | | | | | | | | | |
|---|---|---|---|---|---|---|---|---|---|
| chr1 | *ADORA1* | 11 | 605 | 26 | ADORA1 | adenosine A1 receptor | Plasma Membrane | G-protein coupled recepto | clofarabine, acetaminophen/aspirin/caffeine, acetaminophen/butalbital/caffeine/codeine, caffeine/ethanol, theobromine, acetaminophen/butalbital/caffeine, aminophylline, aspirin/butalbital/caffeine/codeine, theophylline/tretinoin/valproic acid, aspirin/caffeine/phenacetin, dyphylline, caffeine/ergotamine, adenosine, tecadenoson, PBF-680, acetaminophen/caffeine/chlorpheniramine/hydrocodone/phenylephrine, caffeine |
| chr17 | *PFAS* | 11 | 516 | 26 | PFAS | phosphoribosylformylglycinamidine synthase | Cytoplasm | enzyme | |
| chr7 | *EIF2AK1* | 12 | 405 | 26 | EIF2AK1 | eukaryotic translation initiation factor 2 alpha kinase 1 | Cytoplasm | kinase | |
| chr15 | *RP11-17L5.4* | 13 | 355 | 26 | | | | | |
| chr1 | *MMEL1* | 12 | 536 | 26 | MMEL1 | membrane metalloendopeptidase like 1 | Cytoplasm | peptidase | |
| chrX | *MAP7D2* | 12 | 265 | 26 | MAP7D2 | MAP7 domain containing 2 | Other | other | |
| chr16 | *SDR42E2* | 12 | 398 | 26 | SDR42E2 | short chain dehydrogenase/reductase family 42E, member 2 | Other | other | |
| chr7 | *GS1-124K5.12* | 11 | 358 | 26 | | | | | |
| chr22 | *CARD10* | 11 | 492 | 26 | CARD10 | caspase recruitment domain family member 10 | Cytoplasm | other | |
| chr22 | *PPIL2* | 12 | 485 | 26 | PPIL2 | peptidylprolyl isomerase like 2 | Nucleus | enzyme | |
| chr17 | *GGNBP2* | 10 | 270 | 26 | GGNBP2 | gametogenetin binding protein 2 | Other | other | |
| chr7 | *CPVL* | 12 | 319 | 26 | CPVL | carboxypeptidase vitellogenic like | Cytoplasm | peptidase | |
| chr1 | *DFFB* | 13 | 573 | 26 | DFFB | DNA fragmentation factor subunit beta | Nucleus | enzyme | |
| chr4 | *MARCH1;* | 9 | 299 | 26 | | | | | |
| chr19 | *RELB* | 10 | 354 | 26 | RELB | RELB proto-oncogene, NF-kB subunit | Nucleus | transcription regulator | |
| chr1 | *ADAR* | 11 | 420 | 26 | ADAR | adenosine deaminase RNA | Nucleus | enzyme | |
| chr4 | *SH3BP2* | 12 | 596 | 26 | SH3BP2 | SH3 domain binding protein 2 | Cytoplasm | other | |
| chr20 | *WFDC3* | 12 | 376 | 26 | WFDC3 | WAP four-disulfide core domain 3 | Extracellular Space | other | |
| chr4 | *RBPJ* | 10 | 238 | 26 | RBPJ | recombination signal binding protein for immunoglobulin kappa | Nucleus | transcription regulator | |
| chr19 | *ZNF565* | 12 | 294 | 26 | ZNF565 | zinc finger protein 565 | Nucleus | transcription regulator | |
| chr22 | *CBX7* | 11 | 445 | 26 | CBX7 | chromobox 7 | Nucleus | other | |
| chr9 | *WDR34* | 12 | 587 | 26 | WDR34 | WD repeat domain 34 | Cytoplasm | other | |
| chr11 | *TRIM44* | 12 | 413 | 26 | TRIM44 | tripartite motif containing 44 | Cytoplasm | other | |
| chr1 | *TAS1R1* | 12 | 649 | 26 | TAS1R1 | taste 1 receptor member 1 | Plasma Membrane | G-protein coupled recepto | |
| chr19 | *PSG8* | 11 | 310 | 26 | PSG8 | pregnancy specific beta-1-glycoprotein 8 | Other | other | |
| chr1 | *SMYD2* | 11 | 405 | 26 | SMYD2 | SET and MYND domain | Cytoplasm | enzyme | |
| chr17 | *RP11-392O1.4* | 12 | 354 | 26 | | | | | |
| chr3 | *MAPKAPK3* | 12 | 425 | 26 | MAPKAPK3 | MAPK activated protein kinase 3 | Nucleus | kinase | |
| chr11 | *RP11-613D13.4* | 12 | 319 | 26 | | | | | |
| chr10 | *TLX1NB* | 11 | 448 | 26 | TLX1NB | TLX1 neighbor | Other | other | |
| chr5 | *ROPN1L* | 12 | 1199 | 26 | ROPN1L | rhophilin associated tail protein 1 | Cytoplasm | kinase | |
| chr16 | *ADAMTS18* | 11 | 260 | 26 | ADAMTS18 | ADAM metallopeptidase with thrombospondin type 1 motif 18 | Extracellular Space | peptidase | |
| chr17 | *SRR* | 10 | 298 | 26 | SRR | serine racemase | Cytoplasm | enzyme | |
| chr11 | *P2RX3* | 12 | 425 | 26 | P2RX3 | purinergic receptor P2X 3 | Plasma Membrane | ion channel | |
| chr16 | *ARHGAP23P1* | 10 | 473 | 26 | ARHGAP23P1 | Rho GTPase activating protein 23 pseudogene 1 | Other | other | |
| chr9 | *TBC1D2* | 11 | 401 | 26 | TBC1D2 | TBC1 domain family member 2 | Cytoplasm | other | |
| chr18 | *DYM* | 10 | 289 | 26 | DYM | dymeclin | Cytoplasm | other | |
| chr19 | *CC2D1A* | 12 | 435 | 26 | CC2D1A | coiled-coil and C2 domain containing 1A | Cytoplasm | transcription regulator | |
| chr1 | *PROX1-AS1* | 13 | 322 | 26 | PROX1-AS1 | PROX1 antisense RNA 1 | Other | other | |
| chr16 | *RFWD3* | 11 | 308 | 26 | RFWD3 | ring finger and WD repeat domain | Nucleus | enzyme | |
| chr2 | *ITGB6* | 11 | 262 | 26 | ITGB6 | integrin subunit beta 6 | Plasma Membrane | other | |
| chr17 | *RP11-463M16.4* | 11 | 325 | 26 | | | | | |
| chr12 | *RASSF8* | 10 | 284 | 26 | RASSF8 | Ras association domain family member 8 | Extracellular Space | other | |
| chr1 | *TMEM201* | 12 | 602 | 26 | TMEM201 | transmembrane protein 201 | Nucleus | other | |
| chrX | *LINC00894* | 10 | 310 | 26 | LINC00894 | long intergenic non-protein coding RNA 894 | Other | other | |
| chr7 | *STAG3L1* | 11 | 434 | 26 | STAG3L1 | stromal antigen 3-like 1 | Other | other | |
| chr8 | *RP11-380I10.4* | 13 | 281 | 26 | | | | | |
| chr17 | *COPS3* | 11 | 314 | 26 | COPS3 | COP9 signalosome subunit 3 | Cytoplasm | other | |

| | | | | | | | | | |
|---|---|---|---|---|---|---|---|---|---|
| chr11 | *GUCY1A2* | 10 | 243 | 26 | GUCY1A2 | guanylate cyclase 1 soluble subunit alpha 2 | Cytoplasm | enzyme | nitroprusside, nitroglycerin, isosorbide dinitrate, isosorbide-5- |
| chr13 | *EPSTI1* | 11 | 275 | 26 | EPSTI1 | epithelial stromal interaction 1 | Other | other | |
| chr7 | *AC073325.1* | 13 | 393 | 26 | | | | | |
| chr7 | *UBN2* | 10 | 347 | 26 | UBN2 | ubinuclein 2 | Nucleus | other | |
| chr12 | *CPM* | 12 | 272 | 26 | CPM | carboxypeptidase M | Plasma Membrane | peptidase | |
| chr7 | *GBAS* | 11 | 330 | 26 | NIPSNAP2 | nipsnap homolog 2 | Plasma Membrane | other | |
| chr15 | *ZNF106* | 9 | 249 | 26 | ZNF106 | zinc finger protein 106 | Cytoplasm | other | |
| chr12 | *SRRM4* | 11 | 307 | 26 | SRRM4 | serine/arginine repetitive matrix 4 | Nucleus | other | |
| chr16 | *RSPRY1* | 10 | 266 | 26 | RSPRY1 | ring finger and SPRY domain containing 1 | Other | other | |
| chr12 | *TMEM194A* | 11 | 326 | 26 | NEMP1 | nuclear envelope integral membrane protein 1 | Nucleus | other | |
| chr20 | *GGT7* | 12 | 486 | 26 | GGT7 | gamma-glutamyltransferase 7 | Plasma Membrane | enzyme | |
| chr17 | *LSM12* | 10 | 315 | 26 | LSM12 | LSM12 homolog | Other | other | |
| chr3 | *COL6A6* | 12 | 387 | 26 | COL6A6 | collagen type VI alpha 6 chain | Extracellular Space | other | |
| chr16 | *WFDC1* | 12 | 383 | 26 | WFDC1 | WAP four-disulfide core domain 1 | Extracellular Space | other | |
| chr12 | *FOXN4* | 11 | 521 | 26 | FOXN4 | forkhead box N4 | Nucleus | transcription regulator | |
| chr4 | *PET112* | 10 | 285 | 26 | GATB | glutamyl-tRNA amidotransferase subunit B | Cytoplasm | enzyme | |
| chr21 | *MAP3K7CL* | 8 | 212 | 26 | MAP3K7CL | MAP3K7 C-terminal like | Other | other | |
| chr9 | *NFX1* | 10 | 228 | 26 | NFX1 | nuclear transcription factor, X-box binding 1 | Nucleus | transcription regulator | |
| chr11 | *SLC22A8* | 11 | 544 | 26 | SLC22A8 | solute carrier family 22 member 8 | Plasma Membrane | transporter | colchicine/probenecid, probenecid |
| chr14 | *RP11-1029J19.5* | 9 | 327 | 26 | | | | | |
| chr11 | *GRIA4* | 10 | 251 | 26 | GRIA4 | glutamate ionotropic receptor AMPA type subunit 4 | Plasma Membrane | ion channel | tezampanel, farampator, LY451395, |
| chr1 | *LZIC* | 10 | 292 | 26 | LZIC | leucine zipper and CTNNBIP1 domain containing | Other | other | |
| chr22 | *KREMEN1* | 12 | 318 | 26 | KREMEN1 | kringle containing transmembrane protein 1 | Plasma Membrane | other | |
| chr17 | *ITGB4* | 13 | 750 | 26 | ITGB4 | integrin subunit beta 4 | Plasma Membrane | transmembrane recepto | |
| chr5 | *FAM193B* | 11 | 446 | 26 | FAM193B | family with sequence similarity 193 member B | Nucleus | other | |
| chr12 | *VEZT* | 11 | 278 | 26 | VEZT | vezatin, adherens junctions transmembrane protein | Plasma Membrane | other | |
| chr11 | *AC129929.5* | 10 | 373 | 26 | | | | | |
| chr17 | *ITGB3* | 11 | 324 | 26 | ITGB3 | integrin subunit beta 3 | Plasma Membrane | transmembrane recepto | TP 9201, eptifibatide, cilengitide, abciximab, tirofiban, |
| chr19 | *SAMD4B* | 12 | 441 | 26 | SAMD4B | sterile alpha motif domain containing 4B | Cytoplasm | other | |
| chr7 | *PMS2P2* | 12 | 282 | 26 | PMS2P2 | PMS1 homolog 2, mismatch repair system component | Other | other | |
| chr21 | *MX2* | 10 | 353 | 26 | MX2 | MX dynamin like GTPase 2 | Nucleus | enzyme | |
| chr5 | *CTC-321K16.1* | 12 | 349 | 26 | | | | | |
| chr8 | *PDGFRL* | 11 | 331 | 26 | PDGFRL | platelet derived growth factor receptor like | Plasma Membrane | kinase | |
| chr17 | *MYH3* | 10 | 346 | 26 | MYH3 | myosin heavy chain 3 | Cytoplasm | enzyme | |
| chr13 | *RP11-65D24.2* | 12 | 434 | 26 | | | | | |
| chr9 | *DPH7* | 12 | 712 | 26 | DPH7 | diphthamide biosynthesis 7 | Cytoplasm | other | |
| chr17 | *INPP5K* | 12 | 609 | 26 | INPP5K | inositol polyphosphate-5-phosphatase K | Cytoplasm | phosphatase | |
| chr4 | *LPHN3* | 9 | 287 | 26 | ADGRL3 | adhesion G protein-coupled receptor L3 | Plasma Membrane | -protein coupled recepto | |
| chr7 | *LINC00265* | 13 | 375 | 26 | LINC00265 | long intergenic non-protein coding RNA 265 | Other | other | |
| chr10 | *AP3M1* | 11 | 498 | 26 | AP3M1 | adaptor related protein complex 3 subunit mu 1 | Cytoplasm | transporter | |
| chr1 | *ZNF124* | 11 | 278 | 26 | ZNF124 | zinc finger protein 124 | Nucleus | transcription regulator | |
| chr2 | *PUS10* | 10 | 292 | 26 | PUS10 | pseudouridine synthase 10 | Other | other | |
| chr1 | *NTNG1* | 11 | 282 | 26 | NTNG1 | netrin G1 | Extracellular Space | other | |
| chr2 | *KANSL3* | 12 | 444 | 26 | KANSL3 | KAT8 regulatory NSL complex subunit 3 | Nucleus | other | |
| chr7 | *RALA* | 11 | 338 | 26 | RALA | RAS like proto-oncogene A | Cytoplasm | enzyme | |
| chr10 | *FANK1* | 10 | 323 | 26 | FANK1 | fibronectin type III and ankyrin repeat domains 1 | Nucleus | transcription regulator | |
| chr5 | *IL31RA* | 11 | 300 | 26 | IL31RA | interleukin 31 receptor A | Plasma Membrane | transmembrane recepto | |
| chr11 | *CALCB* | 14 | 382 | 26 | CALCB | calcitonin related polypeptide | Extracellular Space | other | |
| chr7 | *CPED1* | 10 | 257 | 26 | CPED1 | cadherin like and PC-esterase domain containing 1 | Cytoplasm | other | |
| chr6 | *MAPK14* | 11 | 266 | 26 | MAPK14 | mitogen-activated protein kinase 14 | Cytoplasm | kinase | RO-3201195, p38 MAP kinase inhibitor, talmapimod, |
| chr1 | *IL22RA1* | 10 | 424 | 26 | IL22RA1 | interleukin 22 receptor subunit alpha 1 | Plasma Membrane | transmembrane recepto | |
| chr16 | *AJ003147.9* | 12 | 425 | 26 | | | | | |
| chr14 | *MEG3* | 11 | 398 | 26 | MEG3 | maternally expressed 3 | Other | other | |
| chr3 | *PVRL3-AS1* | 13 | 386 | 26 | NECTIN3-AS1 | NECTIN3 antisense RNA 1 | Other | other | |
| chr1 | *PHTF1* | 11 | 304 | 26 | PHTF1 | putative homeodomain transcription factor 1 | Nucleus | transcription regulator | |
| chr20 | *AC005808.3* | 10 | 463 | 26 | | | | | |
| chr1 | *RP11-145A3.1* | 11 | 445 | 26 | | | | | |
| chr4 | *KCNIP4* | 10 | 305 | 26 | KCNIP4 | potassium voltage-gated channel interacting protein 4 | Plasma Membrane | ion channel | |

| | | | | | | | | | |
|---|---|---|---|---|---|---|---|---|---|
| chr22 | *SMPD4P1* | 10 | 347 | 26 | SMPD4P1 | sphingomyelin phosphodiesterase 4 pseudogene | Other | other | |
| chr10 | *ABCC2* | 12 | 285 | 26 | ABCC2 | ATP binding cassette subfamily C member 2 | Plasma Membrane | transporter | sulfinpyrazone |
| chr12 | *RP11-370I10.6* | 10 | 405 | 26 | | | | | |
| chr9 | *RC3H2* | 10 | 315 | 26 | RC3H2 | ring finger and CCCH-type | Plasma Membrane | enzyme | |
| chr6 | *GLP1R* | 11 | 492 | 26 | GLP1R | glucagon like peptide 1 receptor | Plasma Membrane | -protein coupled recepto | insulin degludec/liraglutide, exenatide, GLP-1 (7-36) amide, dulaglutide, dulaglutide/INS, liraglutide, semaglutide, |
| chr16 | *TSNAXIP1* | 11 | 355 | 26 | TSNAXIP1 | translin associated factor X interacting protein 1 | Cytoplasm | other | |
| chr15 | *TARSL2* | 11 | 499 | 26 | TARS3 | threonyl-tRNA synthetase 3 | Other | enzyme | |
| chr10 | *OGDHL* | 12 | 930 | 26 | OGDHL | oxoglutarate dehydrogenase like | Other | enzyme | |
| chr3 | *GLB1* | 12 | 297 | 26 | GLB1 | galactosidase beta 1 | Cytoplasm | enzyme | |
| chr5 | *COL4A3BP* | 10 | 235 | 26 | CERT1 | ceramide transporter 1 | Cytoplasm | kinase | |
| chrX | *TBC1D25* | 11 | 316 | 26 | TBC1D25 | TBC1 domain family member 25 | Other | other | |
| chr12 | *C12orf42* | 13 | 295 | 26 | C12orf42 | chromosome 12 open reading frame 42 | Other | other | |
| chr1 | *DYRK3* | 12 | 350 | 26 | DYRK3 | dual specificity tyrosine phosphorylation regulated kinase | Nucleus | kinase | |
| chr8 | *ZNF250* | 12 | 379 | 26 | ZNF250 | zinc finger protein 250 | Nucleus | transcription regulator | |
| chr20 | *MYH7B* | 12 | 664 | 26 | MYH7B | myosin heavy chain 7B | Other | other | |
| chr1 | *SNRNP40* | 8 | 258 | 26 | SNRNP40 | small nuclear ribonucleoprotein U5 subunit 40 | Nucleus | other | |
| chr15 | *TMOD3* | 10 | 253 | 26 | TMOD3 | tropomodulin 3 | Cytoplasm | other | |
| chr17 | *STARD3* | 11 | 533 | 26 | STARD3 | StAR related lipid transfer domain containing 3 | Cytoplasm | transporter | |
| chr15 | *RP11-429B14.1* | 13 | 405 | 26 | | | | | |
| chr3 | *LTF* | 12 | 395 | 26 | LTF | lactotransferrin | Extracellular Space | peptidase | |
| chr22 | *MGAT3* | 10 | 475 | 26 | MGAT3 | mannosyl (beta-1,4-)-glycoprotein beta-1,4-N-acetylglucosaminyltransferase | Cytoplasm | enzyme | |
| chr17 | *SLC5A10* | 12 | 577 | 26 | SLC5A10 | solute carrier family 5 member 10 | Cytoplasm | transporter | |
| chr3 | *CCDC37* | 12 | 550 | 26 | CFAP100 | cilia and flagella associated protein 100 | Cytoplasm | other | |
| chr1 | *DDI2* | 10 | 300 | 26 | DDI2 | DNA damage inducible 1 | Plasma Membrane | transporter | |
| chr3 | *BHLHE40-AS1* | 9 | 250 | 26 | BHLHE40-AS1 | BHLHE40 antisense RNA 1 | Other | other | |
| chr2 | *ATP6V1E2* | 11 | 483 | 26 | ATP6V1E2 | ATPase H+ transporting V1 subunit E2 | Cytoplasm | transporter | |
| chr10 | *PDZD7* | 13 | 506 | 26 | PDZD7 | PDZ domain containing 7 | Extracellular Space | other | |
| chr20 | *ZNF831* | 12 | 571 | 26 | ZNF831 | zinc finger protein 831 | Other | other | |
| chr1 | *DDX59* | 11 | 361 | 26 | DDX59 | DEAD-box helicase 59 | Other | enzyme | |
| chr12 | *RP11-642P15.1* | 10 | 270 | 26 | | | | | |
| chr1 | *TDRD5* | 11 | 313 | 26 | TDRD5 | tudor domain containing 5 | Nucleus | other | |
| chr1 | *NBPF13P* | 13 | 396 | 26 | NBPF13P | NBPF member 13, pseudogene | Other | other | |
| chr10 | *GPR158* | 10 | 257 | 26 | GPR158 | G protein-coupled receptor 158 | Plasma Membrane | -protein coupled recepto | |
| chr10 | *KIF11* | 10 | 271 | 26 | KIF11 | kinesin family member 11 | Nucleus | other | 4SC-205, ispinesib, ARRY-520, AZD4877, litronesib, SB-743921, ALN- |
| chr6 | *NKAIN2* | 10 | 262 | 26 | NKAIN2 | sodium/potassium transporting ATPase interacting 2 | Plasma Membrane | other | |
| chr19 | *ATP1A3* | 12 | 491 | 26 | ATP1A3 | ATPase Na+/K+ transporting subunit alpha 3 | Plasma Membrane | transporter | perphenazine, ethacrynic acid, |
| chr1 | *NEXN* | 10 | 234 | 26 | NEXN | nexilin F-actin binding protein | Plasma Membrane | other | |
| chr11 | *IGHMBP2* | 12 | 468 | 26 | IGHMBP2 | immunoglobulin mu DNA binding protein 2 | Nucleus | enzyme | |
| chr10 | *UPF2* | 9 | 245 | 26 | UPF2 | UPF2 regulator of nonsense mediated mRNA decay | Cytoplasm | other | |
| chr17 | *CWC25* | 11 | 325 | 26 | CWC25 | CWC25 spliceosome associated protein homolog | Nucleus | other | |
| chr3 | *FAM3D* | 11 | 463 | 26 | FAM3D | family with sequence similarity 3 member D | Extracellular Space | cytokine | |
| chr15 | *PLCB2* | 11 | 465 | 26 | PLCB2 | phospholipase C beta 2 | Cytoplasm | enzyme | |
| chr1 | *COPA* | 10 | 315 | 26 | COPA | coatomer protein complex subunit alpha | Cytoplasm | transporter | |
| chr9 | *AKNA* | 10 | 399 | 26 | AKNA | AT-hook transcription factor | Nucleus | transcription regulator | |
| chr12 | *CSAD* | 10 | 563 | 26 | CSAD | cysteine sulfinic acid | Cytoplasm | enzyme | |
| chr17 | *CBX1* | 10 | 305 | 26 | CBX1 | chromobox 1 | Nucleus | transcription regulator | |
| chr1 | *CR1* | 11 | 319 | 26 | CR1 | complement C3b/C4b receptor 1 (Knops blood group) | Plasma Membrane | transmembrane recepto | |
| chr9 | *RP11-395D3.1* | 11 | 231 | 26 | | | | | |
| chr3 | *CCR3* | 12 | 255 | 26 | CCR3 | C-C motif chemokine receptor 3 | Plasma Membrane | -protein coupled recepto | |
| chr21 | *PKNOX1* | 10 | 329 | 26 | PKNOX1 | PBX/knotted 1 homeobox 1 | Nucleus | transcription regulator | |
| chr3 | *ZBTB38* | 12 | 408 | 26 | ZBTB38 | zinc finger and BTB domain containing 38 | Nucleus | transcription regulator | |
| chr9 | *GCNT1* | 11 | 243 | 26 | GCNT1 | glucosaminyl (N-acetyl) | Cytoplasm | enzyme | |
| chr19 | *PVRL2* | 10 | 383 | 26 | NECTIN2 | nectin cell adhesion molecule 2 | Plasma Membrane | transmembrane recepto | |
| chr5 | *LARS* | 9 | 285 | 26 | LARS1 | leucyl-tRNA synthetase 1 | Cytoplasm | enzyme | |
| chr5 | *JMY* | 11 | 279 | 26 | JMY | junction mediating and regulatory protein, p53 cofactor | Nucleus | transcription regulator | |
| chr1 | *PTCH2* | 12 | 484 | 26 | PTCH2 | patched 2 | Plasma Membrane | transmembrane recepto | |
| chr19 | *WTIP* | 11 | 554 | 26 | WTIP | WT1 interacting protein | Nucleus | transcription regulator | |

| | | | | | | | | | |
|---|---|---|---|---|---|---|---|---|---|
| chr9 | *SH3GLB2* | 12 | 566 | 26 | SH3GLB2 | SH3 domain containing GRB2 like, endophilin B2 | Cytoplasm | other | |
| chr3 | *ATXN7* | 11 | 246 | 26 | ATXN7 | ataxin 7 | Nucleus | other | |
| chr5 | *NR3C1* | 10 | 235 | 26 | NR3C1 | nuclear receptor subfamily 3 group C member 1 | Nucleus | d-dependent nuclear rec | dexamethasone/netu pitant/palonosetron, bevacizumab/docetaxel/prednisone/thalidomide/zoledronic acid, beclomethasone, prednisone/thalidomide, methylprednisolone, aprepitant/dexamethasone/palonosetron, methylprednisolone/mycophenolate mofetil, carfilzomib/dexamethasone/rituximab, alclometasone, prednisone/tacrolimus, cimetidine/dexamethasone/diphenhydramine, miconazole, [illegible] |
| chr17 | *CA4* | 11 | 427 | 26 | CA4 | carbonic anhydrase 4 | Plasma Membrane | enzyme | chlorthalidone, acetazolamide, methazolamide, hydrochlorothiazide, trichloromethiazide, phentermine/topiramate, sulfacetamide, benzthiazide, ethoxyzolamide, topiramate, chlorothiazide/reserpine, chlorothiazide, dichlorphenamide |
| chr4 | *ELF2* | 9 | 232 | 26 | ELF2 | E74 like ETS transcription factor | Nucleus | transcription regulator | |
| chr1 | *RP4-717I23.3* | 10 | 266 | 26 | | | | | |
| chr3 | *RP11-59J16.1* | 11 | 538 | 26 | | | | | |
| chr19 | *AKAP8* | 13 | 469 | 26 | AKAP8 | A-kinase anchoring protein 8 | Nucleus | other | |
| chr18 | *ZNF236* | 9 | 301 | 26 | ZNF236 | zinc finger protein 236 | Nucleus | transcription regulator | |
| chr14 | *FUT8* | 9 | 274 | 26 | FUT8 | fucosyltransferase 8 | Cytoplasm | enzyme | |
| chr1 | *WNT4* | 10 | 632 | 26 | WNT4 | Wnt family member 4 | Extracellular Space | cytokine | |
| chrX | *GRIPAP1* | 11 | 412 | 26 | GRIPAP1 | GRIP1 associated protein 1 | Plasma Membrane | other | |
| chr9 | *GRIN1* | 12 | 710 | 26 | GRIN1 | glutamate ionotropic receptor NMDA type subunit 1 | Plasma Membrane | ion channel | ketamine, carbinoxamine/dextromethorphan/pseudoephedrine, dextromethorphan/morphine, ifenprodil, nebostinel, besonprodil, delucemine, 1-aminocyclopropane-1-carboxylic acid, orphenadrine, dextromethorphan/promethazine, aspirin/caffeine/orphenadrine, neramexane, felbamate, chlorpheniramine/dextromethorphan/phenylephrine, memantine, agmatine, donepezil/memantine, [illegible] |
| chr8 | *TNFRSF10D* | 10 | 416 | 26 | TNFRSF10D | TNF receptor superfamily member 10d | Plasma Membrane | transmembrane receptor | |
| chr12 | *ACVR1B* | 10 | 537 | 26 | ACVR1B | activin A receptor type 1B | Plasma Membrane | kinase | |
| chr5 | *SCAMP1* | 11 | 310 | 26 | SCAMP1 | secretory carrier membrane | Cytoplasm | transporter | |
| chr3 | *SYNPR* | 11 | 262 | 26 | SYNPR | synaptoporin | Plasma Membrane | transporter | |
| chr3 | *ST6GAL1* | 10 | 259 | 26 | ST6GAL1 | ST6 beta-galactoside alpha-2,6-sialyltransferase 1 | Cytoplasm | enzyme | |
| chr9 | *HIATL2* | 9 | 253 | 26 | MFSD14C | major facilitator superfamily domain containing 14C | Other | other | |
| chr15 | *IQCH* | 10 | 213 | 26 | IQCH | IQ motif containing H | Other | other | |
| chr1 | *S100PBP* | 9 | 246 | 26 | S100PBP | S100P binding protein | Nucleus | other | |
| chr1 | *MECR* | 11 | 443 | 26 | MECR | mitochondrial trans-2-enoyl-CoA reductase | Cytoplasm | enzyme | |
| chr17 | *SUPT6H* | 10 | 347 | 26 | SUPT6H | SPT6 homolog, histone chaperone and transcription | Nucleus | transcription regulator | |
| chr11 | *RP11-736K20.6* | 12 | 483 | 26 | | | | | |
| chr5 | *SH3RF2* | 12 | 298 | 25 | SH3RF2 | SH3 domain containing ring finger | Extracellular Space | enzyme | |

| | | | | | | | | | |
|---|---|---|---|---|---|---|---|---|---|
| chr16 | *ZDHHC1* | 12 | 474 | 25 | ZDHHC1 | zinc finger DHHC-type containing | Nucleus | enzyme | |
| chr11 | *ATG16L2* | 12 | 630 | 25 | ATG16L2 | autophagy related 16 like 2 | Extracellular Space | other | |
| chr12 | *FAM222A* | 11 | 451 | 25 | FAM222A | family with sequence similarity 222 member A | Other | other | |
| chr3 | *GPR156* | 12 | 397 | 25 | GPR156 | G protein-coupled receptor 156 | Plasma Membrane | [illegible]-protein coupled recept[illegible] | |
| chr16 | *LINC00922* | 11 | 248 | 25 | LINC00922 | long intergenic non-protein coding RNA 922 | Other | other | |
| chr17 | *CACNB1* | 10 | 396 | 25 | CACNB1 | calcium voltage-gated channel auxiliary subunit beta 1 | Plasma Membrane | ion channel | verapamil, ibutilide, dexamethasone/rituximab/verapamil, mibefradil, nimodipine, |
| chr19 | *ZNF826P* | 10 | 251 | 25 | ZNF826P | zinc finger protein 826, | Other | other | |
| chr8 | *RP11-273G15.2* | 11 | 476 | 25 | | | | | |
| chr11 | *KIAA1377* | 11 | 548 | 25 | CEP126 | centrosomal protein 126 | Cytoplasm | other | |
| chr19 | *EPN1* | 12 | 529 | 25 | EPN1 | epsin 1 | Plasma Membrane | other | |
| chr11 | *SLC43A1* | 12 | 391 | 25 | SLC43A1 | solute carrier family 43 member 1 | Plasma Membrane | transporter | |
| chr1 | *SMG5* | 11 | 483 | 25 | SMG5 | SMG5 nonsense mediated mRNA decay factor | Nucleus | other | |
| chr17 | *RP11-333E1.1* | 11 | 297 | 25 | | | | | |
| chr1 | *HNRNPR* | 9 | 283 | 25 | HNRNPR | heterogeneous nuclear ribonucleoprotein R | Nucleus | other | |
| chr19 | *DHX34* | 11 | 471 | 25 | DHX34 | DExH-box helicase 34 | Other | enzyme | |
| chr16 | *PAPD5* | 11 | 236 | 25 | TENT4B | terminal nucleotidyltransferase | Nucleus | enzyme | |
| chr20 | *CDK5RAP1* | 13 | 284 | 25 | CDK5RAP1 | CDK5 regulatory subunit associated protein 1 | Extracellular Space | enzyme | |
| chr10 | *DDX50* | 12 | 378 | 25 | DDX50 | DExD-box helicase 50 | Nucleus | enzyme | |
| chr1 | *KIFAP3* | 10 | 376 | 25 | KIFAP3 | kinesin associated protein 3 | Cytoplasm | other | |
| chr11 | *TRPC2* | 13 | 762 | 25 | TRPC2 | transient receptor potential cation channel subfamily C member 2 (pseudogene) | Other | other | |
| chr16 | *RP11-19N8.2* | 11 | 661 | 25 | | | | | |
| chr6 | *KIAA1244* | 10 | 303 | 25 | ARFGEF3 | ARFGEF family member 3 | Other | other | |
| chr19 | *AC007796.1* | 12 | 264 | 25 | | | | | |
| chr1 | *LAPTM5* | 10 | 458 | 25 | LAPTM5 | lysosomal protein transmembrane | Plasma Membrane | other | |
| chr9 | *PTCH1* | 10 | 319 | 25 | PTCH1 | patched 1 | Plasma Membrane | transmembrane recepto[illegible] | |
| chrX | *CLCN5* | 11 | 325 | 25 | CLCN5 | chloride voltage-gated channel 5 | Plasma Membrane | ion channel | |
| chr13 | *LAMP1* | 12 | 398 | 25 | LAMP1 | lysosomal associated membrane protein 1 | Plasma Membrane | other | SAR428926 |
| chr1 | *MTF1* | 10 | 257 | 25 | MTF1 | metal regulatory transcription | Nucleus | transcription regulator | APTO-253 |
| chr15 | *ANP32A* | 10 | 357 | 25 | ANP32A | acidic nuclear phosphoprotein 32 family member A | Other | other | |
| chr17 | *CALCOCO2* | 10 | 246 | 25 | CALCOCO2 | calcium binding and coiled-coil domain 2 | Nucleus | other | |
| chr9 | *NFIB* | 11 | 239 | 25 | NFIB | nuclear factor I B | Nucleus | transcription regulator | |
| chr3 | *SLC4A7* | 10 | 285 | 25 | SLC4A7 | solute carrier family 4 member 7 | Plasma Membrane | transporter | |
| chr4 | *FAM13A* | 9 | 246 | 25 | FAM13A | family with sequence similarity 13 member A | Cytoplasm | other | |
| chr1 | *RP11-439E19.8* | 12 | 393 | 25 | | | | | |
| chr3 | *TRAIP* | 11 | 367 | 25 | TRAIP | TRAF interacting protein | Cytoplasm | enzyme | |
| chr5 | *GABRB2* | 11 | 270 | 25 | GABRB2 | gamma-aminobutyric acid type A receptor beta2 subunit | Plasma Membrane | ion channel | [illegible] nitrazepam, methohexital, sevoflurane, etomidate, pagoclone, aspirin/butalbital/caffeine, zolpidem, atropine/hyoscyamine/phenobarbital/scopolamine, phenobarbital, diazepam, isoflurane, halothane, pentobarbital, acetaminophen/butalbital/caffeine/codeine, hyoscyamine/phenobarbital, desflurane, amobarbital, adinazolam, SEP 174559, [illegible] |
| chrX | *ATRX* | 9 | 265 | 25 | ATRX | ATRX chromatin remodeler | Nucleus | transcription regulator | |
| chr10 | *PSAP* | 12 | 697 | 25 | PSAP | prosaposin | Extracellular Space | enzyme | |
| chr11 | *CCDC81* | 11 | 451 | 25 | CCDC81 | coiled-coil domain containing 81 | Cytoplasm | other | |
| chrX | *SYAP1* | 11 | 258 | 25 | SYAP1 | synapse associated protein 1 | Nucleus | other | |
| chr2 | *UBR3* | 8 | 234 | 25 | UBR3 | ubiquitin protein ligase E3 component n-recognin 3 | Other | enzyme | |
| chr3 | *ATP6V1A* | 10 | 281 | 25 | ATP6V1A | ATPase H+ transporting V1 subunit A | Plasma Membrane | transporter | bafilomycin A1, bafilomycin b1 |
| chr2 | *VPS54* | 10 | 250 | 25 | VPS54 | VPS54 subunit of GARP complex | Extracellular Space | other | |
| chr7 | *CCDC129* | 10 | 274 | 25 | ITPRID1 | ITPR interacting domain | Other | other | |
| chr5 | *IL6ST* | 9 | 221 | 25 | IL6ST | interleukin 6 signal transducer | Plasma Membrane | transmembrane recepto[illegible] | |
| chr1 | *FAM73A* | 10 | 234 | 25 | MIGA1 | mitoguardin 1 | Plasma Membrane | other | |
| chr3 | *STAC* | 11 | 249 | 25 | STAC | SH3 and cysteine rich domain | Cytoplasm | other | |
| chr5 | *NNT* | 10 | 729 | 25 | NNT | nicotinamide nucleotide transhydrogenase | Cytoplasm | enzyme | |

| | | | | | | | | | |
|---|---|---|---|---|---|---|---|---|---|
| chr19 | *POLD1* | 11 | 493 | 25 | POLD1 | DNA polymerase delta 1, catalytic subunit | Nucleus | enzyme | entecavir, nelarabine, cytarabine/filgrastim/fludarabine phosphate/imatinib, cytarabine/fludarabine phosphate/oxaliplatin/rituximab, cytarabine/fludarabine phosphate, cytarabine/filgrastim/fludarabine phosphate, trifluridine, clofarabine/cyclophosphamide/etoposide/imatinib, cytarabine/dexamethasone, cytarabine/gemtuzumab ozogamicin, cytarabine/IFNA2B, clofarabine, cytarabine/etoposide/ |
| chr17 | *FADS6* | 11 | 452 | 25 | FADS6 | fatty acid desaturase 6 | Other | enzyme | |
| chr8 | *SORBS3* | 11 | 575 | 25 | SORBS3 | sorbin and SH3 domain | Cytoplasm | other | |
| chr12 | *CCNT1* | 10 | 433 | 25 | CCNT1 | cyclin T1 | Nucleus | transcription regulator | |
| chr11 | *Y_RNA* | 11 | 295 | 25 | | | | | |
| chr12 | *CD4* | 11 | 325 | 25 | CD4 | CD4 molecule | Plasma Membrane | transmembrane receptor | ibalizumab, zanolimumab |
| chr4 | *NEK1* | 9 | 269 | 25 | NEK1 | NIMA related kinase 1 | Nucleus | kinase | |
| chr2 | *CYP27C1* | 10 | 483 | 25 | CYP27C1 | cytochrome P450 family 27 subfamily C member 1 | Other | other | |
| chr1 | *TRIT1* | 10 | 308 | 25 | TRIT1 | tRNA isopentenyltransferase 1 | Cytoplasm | enzyme | |
| chr14 | *L2HGDH* | 8 | 220 | 25 | L2HGDH | L-2-hydroxyglutarate | Cytoplasm | enzyme | |
| chr6 | *EPM2A* | 11 | 269 | 25 | EPM2A | EPM2A glucan phosphatase, | Cytoplasm | phosphatase | |
| chr15 | *TYRO3* | 10 | 368 | 25 | TYRO3 | TYRO3 protein tyrosine kinase | Plasma Membrane | kinase | cabozantinib/nivolumab, cabozantinib/erlotinib, RXDX-106, |
| chr10 | *MMRN2* | 11 | 479 | 25 | MMRN2 | multimerin 2 | Extracellular Space | other | |
| chr1 | *LDLRAP1* | 10 | 580 | 25 | LDLRAP1 | low density lipoprotein receptor adaptor protein 1 | Cytoplasm | transporter | |
| chr1 | *GNG12* | 12 | 280 | 25 | GNG12 | G protein subunit gamma 12 | Plasma Membrane | enzyme | |
| chr10 | *TEX36* | 12 | 331 | 25 | TEX36 | testis expressed 36 | Other | other | |
| chr2 | *ST3GAL5* | 11 | 347 | 25 | ST3GAL5 | ST3 beta-galactoside alpha-2,3-sialyltransferase 5 | Cytoplasm | enzyme | |
| chr5 | *GRK6* | 11 | 423 | 25 | GRK6 | G protein-coupled receptor | Plasma Membrane | kinase | |
| chr15 | *USP50* | 10 | 232 | 25 | USP50 | ubiquitin specific peptidase 50 | Other | other | |
| chr2 | *AC010967.2* | 8 | 287 | 25 | | | | | |
| chr17 | *PRR11* | 10 | 248 | 25 | PRR11 | proline rich 11 | Other | other | |
| chr7 | *STAG3L3* | 11 | 499 | 25 | STAG3L2/STAG3L | stromal antigen 3-like 3 | Other | other | |
| chr12 | *HSD17B6* | 11 | 258 | 25 | HSD17B6 | hydroxysteroid 17-beta dehydrogenase 6 | Other | enzyme | 3-hydroxysteroid dehydrogenase inhibitor |
| chr5 | *STARD4-AS1* | 12 | 272 | 25 | STARD4-AS1 | STARD4 antisense RNA 1 | Other | other | |
| chr19 | *IL27RA* | 10 | 361 | 25 | IL27RA | interleukin 27 receptor subunit | Plasma Membrane | transmembrane receptor | |
| chr1 | *RCSD1* | 11 | 311 | 25 | RCSD1 | RCSD domain containing 1 | Other | other | |
| chr7 | *ZNF282* | 11 | 370 | 25 | ZNF282 | zinc finger protein 282 | Nucleus | transcription regulator | |
| chr17 | *MYO19* | 11 | 388 | 25 | MYO19 | myosin XIX | Cytoplasm | enzyme | |
| chr1 | *SLC22A15* | 10 | 292 | 25 | SLC22A15 | solute carrier family 22 member | Other | transporter | |
| chr2 | *IWS1* | 10 | 423 | 25 | IWS1 | interacts with SUPT6H, CTD assembly factor 1 | Nucleus | enzyme | |
| chr5 | *ATP6AP1L* | 12 | 249 | 25 | ATP6AP1L | ATPase H+ transporting accessory protein 1 like | Other | other | |
| chr2 | *TMEM87B* | 11 | 320 | 25 | TMEM87B | transmembrane protein 87B | Cytoplasm | other | |
| chr19 | *CHAF1A* | 11 | 414 | 25 | CHAF1A | chromatin assembly factor 1 subunit A | Nucleus | other | |
| chr18 | *C18orf25* | 8 | 237 | 25 | C18orf25 | chromosome 18 open reading frame 25 | Other | other | |
| chr20 | *CTCFL* | 11 | 491 | 25 | CTCFL | CCCTC-binding factor like | Nucleus | transcription regulator | |
| chr19 | *NPAS1* | 11 | 406 | 25 | NPAS1 | neuronal PAS domain protein 1 | Nucleus | transcription regulator | |
| chr1 | *RP11-541F9.2* | 9 | 249 | 25 | | | | | |
| chr17 | *MED13* | 9 | 252 | 25 | MED13 | mediator complex subunit 13 | Nucleus | transcription regulator | |
| chrX | *RP11-268G12.3* | 9 | 245 | 25 | | | | | |
| chr1 | *SHCBP1L* | 11 | 300 | 25 | SHCBP1L | SHC binding and spindle associated 1 like | Other | other | |
| chr3 | *LINC00879* | 11 | 297 | 25 | LINC00879 | long intergenic non-protein coding RNA 879 | Other | other | |
| chr2 | *DOCK10* | 9 | 190 | 25 | DOCK10 | dedicator of cytokinesis 10 | Cytoplasm | other | |
| chr1 | *RP11-463J7.2* | 10 | 251 | 25 | | | | | |
| chr19 | *PPP5C* | 10 | 394 | 25 | PPP5C | protein phosphatase 5 catalytic subunit | Nucleus | phosphatase | |
| chrX | *RP11-268G12.1* | 10 | 271 | 25 | | | | | |
| chr19 | *MAP4K1* | 10 | 380 | 25 | MAP4K1 | mitogen-activated protein kinase kinase kinase kinase 1 | Cytoplasm | kinase | |
| chr3 | *RP11-340E6.1* | 12 | 352 | 25 | | | | | |
| chr11 | *RP11-750H9.5* | 12 | 809 | 25 | | | | | |

| | | | | | | | | | |
|---|---|---|---|---|---|---|---|---|---|
| chr2 | *DNER* | 10 | 232 | 25 | DNER | delta/notch like EGF repeat containing | Plasma Membrane | transmembrane receptor | |
| chr4 | *ARHGAP24* | 9 | 243 | 25 | ARHGAP24 | Rho GTPase activating protein 24 | Cytoplasm | other | |
| chr1 | *BCL9* | 10 | 429 | 25 | BCL9 | BCL9 transcription coactivator | Nucleus | other | |
| chr15 | *CLK3* | 10 | 506 | 25 | CLK3 | CDC like kinase 3 | Nucleus | kinase | |
| chr2 | *MRPS5* | 9 | 314 | 25 | MRPS5 | mitochondrial ribosomal protein | Cytoplasm | other | |
| chr6 | *XPO5* | 11 | 251 | 25 | XPO5 | exportin 5 | Nucleus | other | |
| chr17 | *RP11-580I16.2* | 10 | 265 | 25 | | | | | |
| chr1 | *DHDDS* | 11 | 303 | 25 | DHDDS | dehydrodolichyl diphosphate synthase subunit | Cytoplasm | enzyme | |
| chr12 | *RP11-530C5.1* | 10 | 239 | 25 | | | | | |
| chr12 | *ITGA7* | 11 | 790 | 25 | ITGA7 | integrin subunit alpha 7 | Plasma Membrane | other | |
| chr15 | *SYNM* | 12 | 671 | 25 | SYNM | synemin | Cytoplasm | other | |
| chr17 | *TRPV2* | 10 | 389 | 25 | TRPV2 | transient receptor potential cation channel subfamily V member 2 | Plasma Membrane | ion channel | |
| chrX | *GPR173* | 10 | 434 | 25 | GPR173 | G protein-coupled receptor 173 | Plasma Membrane | G-protein coupled receptor | |
| chr16 | *CTD-2349B8.1* | 12 | 275 | 25 | | | | | |
| chr20 | *EBF4* | 10 | 363 | 25 | EBF4 | EBF family member 4 | Nucleus | transcription regulator | |
| chr1 | *TMEM51-AS1* | 10 | 368 | 25 | TMEM51-AS1 | TMEM51 antisense RNA 1 | Other | other | |
| chr8 | *R3HCC1* | 11 | 444 | 25 | R3HCC1 | R3H domain and coiled-coil containing 1 | Other | other | |
| chr16 | *SETD1A* | 10 | 512 | 25 | SETD1A | SET domain containing 1A, histone lysine methyltransferase | Nucleus | ion channel | |
| chr15 | *CTSH* | 11 | 580 | 25 | CTSH | cathepsin H | Cytoplasm | peptidase | |
| chr3 | *SLC6A20* | 11 | 425 | 25 | SLC6A20 | solute carrier family 6 member 20 | Plasma Membrane | transporter | |
| chr14 | *RP11-85K15.2* | 9 | 207 | 25 | | | | | |
| chr17 | *LGALS9C* | 10 | 539 | 25 | LGALS9C | galectin 9C | Cytoplasm | other | |
| chr5 | *AC005355.1* | 10 | 305 | 25 | | | | | |
| chr13 | *C13orf35* | 11 | 408 | 25 | ATP11AUN | ATP11A upstream neighbor | Other | other | |
| chr7 | *AC005537.2* | 12 | 315 | 25 | | | | | |
| chr17 | *HID1* | 11 | 570 | 25 | HID1 | HID1 domain containing | Plasma Membrane | other | |
| chr22 | *CYB5R3* | 10 | 458 | 25 | CYB5R3 | cytochrome b5 reductase 3 | Cytoplasm | enzyme | |
| chr7 | *CNPY1* | 11 | 466 | 25 | CNPY1 | canopy FGF signaling regulator 1 | Other | other | |
| chr9 | *GTF3C5* | 11 | 609 | 25 | GTF3C5 | general transcription factor IIIC subunit 5 | Nucleus | transcription regulator | |
| chr12 | *PPFIA2* | 10 | 249 | 25 | PPFIA2 | PTPRF interacting protein alpha 2 | Plasma Membrane | phosphatase | |
| chr19 | *GPI* | 11 | 453 | 25 | GPI | glucose-6-phosphate isomerase | Extracellular Space | enzyme | |
| chr11 | *DCPS* | 10 | 379 | 25 | DCPS | decapping enzyme, scavenger | Nucleus | enzyme | |
| chrX | *EDA* | 10 | 261 | 25 | EDA | ectodysplasin A | Plasma Membrane | cytokine | |
| chr17 | *RP11-156P1.3* | 10 | 283 | 25 | | | | | |
| chr19 | *ZSCAN18* | 11 | 345 | 25 | ZSCAN18 | zinc finger and SCAN domain containing 18 | Other | other | |
| chr5 | *CCDC125* | 10 | 264 | 25 | CCDC125 | coiled-coil domain containing 125 | Cytoplasm | other | |
| chr19 | *CYP4F22* | 12 | 338 | 25 | CYP4F22 | cytochrome P450 family 4 subfamily F member 22 | Extracellular Space | enzyme | |
| chr1 | *SF3A3* | 10 | 283 | 25 | SF3A3 | splicing factor 3a subunit 3 | Nucleus | other | |
| chr19 | *PAPL* | 10 | 326 | 25 | | | | | |
| chr1 | *CTSS* | 10 | 282 | 25 | CTSS | cathepsin S | Cytoplasm | peptidase | |
| chr15 | *LOXL1* | 11 | 463 | 25 | LOXL1 | lysyl oxidase like 1 | Extracellular Space | enzyme | |
| chr15 | *LINC00925* | 12 | 662 | 25 | MIR9-3HG | MIR9-3 host gene | Other | other | |
| chr8 | *ENPP7P6* | 12 | 417 | 25 | ENPP7P6 | | Other | other | |
| chr11 | *ORAOV1* | 10 | 456 | 25 | LTO1 | LTO1 maturation factor of ABCE1 | Nucleus | other | |
| chr12 | *STAT6* | 10 | 404 | 25 | STAT6 | signal transducer and activator of transcription 6 | Nucleus | transcription regulator | |
| chr4 | *INPP4B* | 9 | 234 | 25 | INPP4B | inositol polyphosphate-4-phosphatase type II B | Cytoplasm | phosphatase | |
| chr17 | *PPM1D* | 10 | 250 | 25 | PPM1D | protein phosphatase, Mg2+/Mn2+ dependent 1D | Cytoplasm | phosphatase | |
| chr22 | *PISD* | 10 | 391 | 25 | PISD | phosphatidylserine | Cytoplasm | enzyme | |
| chr19 | *ZNF702P* | 11 | 249 | 25 | ZNF702P | zinc finger protein 702, | Nucleus | other | |
| chr1 | *CA6* | 11 | 303 | 25 | CA6 | carbonic anhydrase 6 | Extracellular Space | enzyme | hydrochlorothiazide, trichloromethiazide, sulfacetamide, benzthiazide, topiramate, chlorothiazide, SAR566658, chlorthalidone, acetazolamide, methazolamide |
| chr1 | *SETDB1* | 12 | 329 | 25 | SETDB1 | SET domain bifurcated histone lysine methyltransferase 1 | Nucleus | enzyme | |
| chr12 | *APAF1* | 10 | 249 | 25 | APAF1 | apoptotic peptidase activating | Cytoplasm | other | |
| chr16 | *BCL7C* | 10 | 261 | 25 | BCL7C | BAF chromatin remodeling complex subunit BCL7C | Nucleus | other | |
| chr10 | *MGEA5* | 10 | 278 | 25 | OGA | O-GlcNAcase | Cytoplasm | enzyme | |
| chr1 | *ZMYM6* | 10 | 211 | 25 | ZMYM6 | zinc finger MYM-type containing | Nucleus | other | |
| chr16 | *MMP2* | 12 | 325 | 25 | MMP2 | matrix metallopeptidase 2 | Extracellular Space | peptidase | prinomastat, rebimastat, marimastat, MMP2 |
| chr15 | *RP11-111A22.1* | 11 | 271 | 25 | | | | | |
| chr20 | *AHCY* | 11 | 540 | 25 | AHCY | adenosylhomocysteinase | Cytoplasm | enzyme | 3-deazaaristeromycin, cyanovirin-N, neplanocin A, 6'-C-methylneplanocin A, 9-(2',3'-dihydroxycyclopent-4'-enyl)adenine, 3- |

| | | | | | | | | | |
|---|---|---|---|---|---|---|---|---|---|
| chr7 | *AGFG2* | 12 | 515 | 25 | AGFG2 | ArfGAP with FG repeats 2 | Other | other | |
| chr1 | *CGN* | 11 | 511 | 25 | CGN | cingulin | Plasma Membrane | other | |
| chr19 | *TMEM38A* | 11 | 290 | 25 | TMEM38A | transmembrane protein 38A | Cytoplasm | ion channel | |
| chr17 | *MEOX1* | 11 | 479 | 25 | MEOX1 | mesenchyme homeobox 1 | Nucleus | transcription regulator | |
| chr2 | *PPP1R21* | 8 | 257 | 25 | PPP1R21 | protein phosphatase 1 regulatory subunit 21 | Other | other | |
| chr10 | *FAM13C* | 10 | 379 | 25 | FAM13C | family with sequence similarity 13 member C | Other | other | |
| chr12 | *UBE3B* | 10 | 393 | 25 | UBE3B | ubiquitin protein ligase E3B | Extracellular Space | enzyme | |
| chr8 | *SAMD12* | 8 | 240 | 25 | SAMD12 | sterile alpha motif domain containing 12 | Other | other | |
| chr12 | *SETD8* | 11 | 386 | 25 | KMT5A | lysine methyltransferase 5A | Nucleus | enzyme | |
| chr3 | *MSL2* | 9 | 325 | 25 | MSL2 | MSL complex subunit 2 | Nucleus | other | |
| chr12 | *SDS* | 11 | 430 | 25 | SDS | serine dehydratase | Cytoplasm | enzyme | |
| chr15 | *RAB8B* | 9 | 223 | 25 | RAB8B | RAB8B, member RAS oncogene family | Cytoplasm | enzyme | |
| chr5 | *APBB3* | 10 | 392 | 25 | APBB3 | amyloid beta precursor protein binding family B member 3 | Cytoplasm | other | |
| chr17 | *LLGL1* | 12 | 585 | 25 | LLGL1 | LLGL scribble cell polarity complex component 1 | Cytoplasm | other | |
| chr12 | *CD9* | 10 | 321 | 25 | CD9 | CD9 molecule | Plasma Membrane | other | |
| chr8 | *MTSS1* | 10 | 263 | 25 | MTSS1 | MTSS I-BAR domain containing 1 | Cytoplasm | other | |
| chr10 | *IPMK* | 11 | 415 | 25 | IPMK | inositol polyphosphate | Nucleus | kinase | |
| chr7 | *AC003090.1* | 13 | 370 | 25 | | | | | |
| chr1 | *AGRN* | 13 | 617 | 25 | AGRN | agrin | Plasma Membrane | other | |
| chr3 | *RP11-81N13.1* | 12 | 236 | 25 | | | | | |
| chr9 | *TTF1* | 10 | 388 | 25 | TTF1 | transcription termination factor 1 | Nucleus | transcription regulator | |
| chr2 | *TACR1* | 11 | 272 | 25 | TACR1 | tachykinin receptor 1 | Plasma Membrane | G-protein coupled recepto | SLV317, dexamethasone/netupitant/palonosetron, dexamethasone/netupitant, aprepitant/dexamethasone, aprepitant/granisetron, aprepitant/dexamethasone/granisetron, aprepitant/dexamethasone/palonosetron, fosaprepitant dimeglumine, casopitant, aprepitant, aprepitant/palonosetron, netupitant, aprepitant/ondansetron, aprepitant/dexameth |
| chr12 | *AMN1* | 10 | 352 | 25 | AMN1 | antagonist of mitotic exit network 1 homolog | Plasma Membrane | other | |
| chr3 | *NR1I2* | 11 | 424 | 25 | NR1I2 | nuclear receptor subfamily 1 group I member 2 | Nucleus | d-dependent nuclear rec | |
| chr11 | *WT1* | 12 | 409 | 25 | WT1 | WT1 transcription factor | Nucleus | transcription regulator | |
| chr2 | *CLHC1* | 10 | 281 | 25 | CLHC1 | clathrin heavy chain linker domain containing 1 | Other | other | |
| chr9 | *PPAPDC3* | 11 | 608 | 25 | PLPP7 | phospholipid phosphatase 7 (inactive) | Nucleus | phosphatase | |
| chr1 | *VASH2* | 12 | 412 | 25 | VASH2 | vasohibin 2 | Cytoplasm | peptidase | |
| chr7 | *ETV1* | 9 | 265 | 25 | ETV1 | ETS variant transcription factor 1 | Nucleus | transcription regulator | |
| chr7 | *WIPF3* | 11 | 332 | 25 | WIPF3 | WAS/WASL interacting protein family member 3 | Plasma Membrane | other | |
| chr10 | *ACTR1A* | 10 | 391 | 25 | ACTR1A | actin related protein 1A | Cytoplasm | other | |
| chr19 | *ZFP14* | 10 | 268 | 25 | ZFP14 | ZFP14 zinc finger protein | Nucleus | transcription regulator | |
| chr17 | *RP11-285E9.6* | 10 | 354 | 25 | | | | | |
| chr5 | *THOC3* | 11 | 371 | 25 | THOC3 | THO complex 3 | Nucleus | other | |
| chr11 | *CD44* | 12 | 356 | 24 | CD44 | CD44 molecule (Indian blood group) | Plasma Membrane | other | A6 peptide, anti-CD44v7 antibody, AMC303 |
| chr1 | *PHGDH* | 11 | 539 | 24 | PHGDH | phosphoglycerate dehydrogenase | Cytoplasm | enzyme | |
| chr16 | *RP11-10K17.6* | 11 | 286 | 24 | | | | | |
| chr1 | *PRUNE* | 10 | 390 | 24 | PRUNE1 | prune exopolyphosphatase 1 | Nucleus | enzyme | |
| chr16 | *NOMO1* | 11 | 399 | 24 | MO1 (includes oth | NODAL modulator 1 | Plasma Membrane | other | |
| chr22 | *FAM227A* | 10 | 234 | 24 | FAM227A | family with sequence similarity 227 member A | Other | other | |
| chr2 | *AC010884.1* | 10 | 313 | 24 | | | | | |
| chr1 | *WDR63* | 10 | 219 | 24 | WDR63 | WD repeat domain 63 | Other | other | |
| chr10 | *LINC00840* | 11 | 451 | 24 | LINC00840 | long intergenic non-protein coding RNA 840 | Other | other | |
| chr19 | *ZNF529* | 10 | 206 | 24 | ZNF529 | zinc finger protein 529 | Other | other | |
| chr12 | *LRMP* | 10 | 262 | 24 | LRMP | lymphoid restricted membrane | Cytoplasm | other | |
| chr12 | *GALNT6* | 11 | 545 | 24 | GALNT6 | polypeptide N-acetylgalactosaminyltransferase | Cytoplasm | enzyme | |
| chr19 | *HSH2D* | 10 | 386 | 24 | HSH2D | hematopoietic SH2 domain containing | Cytoplasm | other | |
| chr6 | *PLAGL1* | 11 | 272 | 24 | PLAGL1 | PLAG1 like zinc finger 1 | Nucleus | transcription regulator | |
| chr22 | *MICALL1* | 11 | 435 | 24 | MICALL1 | MICAL like 1 | Cytoplasm | other | |
| chr19 | *PPP1R12C* | 11 | 525 | 24 | PPP1R12C | protein phosphatase 1 regulatory subunit 12C | Cytoplasm | phosphatase | |

| | | | | | | | | | |
|---|---|---|---|---|---|---|---|---|---|
| chr11 | *AMOTL1* | 12 | 644 | 24 | AMOTL1 | angiomotin like 1 | Plasma Membrane | other | |
| chr5 | *TNPO1* | 9 | 217 | 24 | TNPO1 | transportin 1 | Nucleus | transporter | |
| chr7 | *ZKSCAN5* | 10 | 332 | 24 | ZKSCAN5 | zinc finger with KRAB and SCAN domains 5 | Nucleus | transcription regulator | |
| chr22 | *FAM118A* | 9 | 323 | 24 | FAM118A | family with sequence similarity 118 member A | Other | other | |
| chr5 | *SGTB* | 10 | 235 | 24 | SGTB | small glutamine rich tetratricopeptide repeat | Other | other | |
| chr19 | *VSTM2B* | 11 | 341 | 24 | VSTM2B | V-set and transmembrane domain containing 2B | Other | other | |
| chr3 | *PHLDB2* | 11 | 295 | 24 | PHLDB2 | pleckstrin homology like domain family B member 2 | Cytoplasm | other | |
| chr17 | *NUP88* | 9 | 247 | 24 | NUP88 | nucleoporin 88 | Nucleus | transporter | |
| chr9 | *KIAA2026* | 10 | 231 | 24 | KIAA2026 | KIAA2026 | Other | other | |
| chr5 | *RP11-589F5.4* | 11 | 617 | 24 | | | | | |
| chr17 | *RP11-524F11.2* | 10 | 348 | 24 | | | | | |
| chr1 | *YTHDF2* | 9 | 266 | 24 | YTHDF2 | YTH N6-methyladenosine RNA binding protein 2 | Cytoplasm | other | |
| chr5 | *BRD8* | 9 | 235 | 24 | BRD8 | bromodomain containing 8 | Nucleus | transcription regulator | |
| chr11 | *RAB3IL1* | 11 | 699 | 24 | RAB3IL1 | RAB3A interacting protein like 1 | Cytoplasm | other | |
| chr4 | *SEC31A* | 8 | 214 | 24 | SEC31A | SEC31 homolog A, COPII coat complex component | Cytoplasm | other | |
| chr7 | *RP11-514P8.8* | 10 | 751 | 24 | | | | | |
| chr19 | *SLC27A1* | 11 | 410 | 24 | SLC27A1 | solute carrier family 27 member 1 | Plasma Membrane | transporter | |
| chr16 | *CTD-2540M10.1* | 10 | 284 | 24 | | | | | |
| chr11 | *PTDSS2* | 12 | 462 | 24 | PTDSS2 | phosphatidylserine synthase 2 | Cytoplasm | enzyme | |
| chr13 | *HMGB1* | 9 | 220 | 24 | HMGB1 | high mobility group box 1 | Nucleus | transcription regulator | |
| chr14 | *C14orf132* | 10 | 292 | 24 | C14orf132 | chromosome 14 open reading frame 132 | Other | other | |
| chr9 | *SMC5* | 10 | 224 | 24 | SMC5 | structural maintenance of chromosomes 5 | Nucleus | other | |
| chr1 | *DENND4B* | 11 | 626 | 24 | DENND4B | DENN domain containing 4B | Extracellular Space | other | |
| chr15 | *ZSCAN2* | 11 | 423 | 24 | ZSCAN2 | zinc finger and SCAN domain containing 2 | Nucleus | transcription regulator | |
| chr22 | *SULT4A1* | 10 | 439 | 24 | SULT4A1 | sulfotransferase family 4A | Cytoplasm | enzyme | |
| chr1 | *SRRM1* | 9 | 292 | 24 | SRRM1 | serine and arginine repetitive | Nucleus | other | |
| chr6 | *TCP10L2* | 11 | 405 | 24 | TCP10/TCP10L2 | t-complex 10 | Cytoplasm | other | |
| chr16 | *ATP2A1* | 11 | 484 | 24 | ATP2A1 | ATPase sarcoplasmic/endoplasmic | Cytoplasm | transporter | |
| chr3 | *ACPL2* | 11 | 417 | 24 | PXYLP1 | 2-phosphoxylose phosphatase 1 | Cytoplasm | phosphatase | |
| chr7 | *ZKSCAN1* | 10 | 343 | 24 | ZKSCAN1 | zinc finger with KRAB and SCAN domains 1 | Nucleus | transcription regulator | |
| chr3 | *AC026167.1* | 11 | 248 | 24 | | | | | |
| chr6 | *GMPR* | 11 | 290 | 24 | GMPR | guanosine monophosphate reductase | Cytoplasm | enzyme | |
| chr5 | *FAXDC2* | 11 | 270 | 24 | FAXDC2 | fatty acid hydroxylase domain containing 2 | Other | other | |
| chr17 | *B4GALNT2* | 10 | 273 | 24 | B4GALNT2 | beta-1,4-N-acetyl-galactosaminyltransferase 2 | Cytoplasm | enzyme | |
| chr4 | *BANK1* | 9 | 236 | 24 | BANK1 | B cell scaffold protein with ankyrin repeats 1 | Extracellular Space | other | |
| chr1 | *RP11-206L10.11* | 10 | 481 | 24 | | | | | |
| chr2 | *FBXO36* | 8 | 237 | 24 | FBXO36 | F-box protein 36 | Other | other | |
| chr6 | *FNDC1* | 9 | 270 | 24 | FNDC1 | fibronectin type III domain containing 1 | Plasma Membrane | other | AGS-8M4 |
| chr12 | *PLBD2* | 11 | 408 | 24 | PLBD2 | phospholipase B domain | Extracellular Space | other | |
| chr10 | *ARHGAP19-SLIT1* | 11 | 358 | 24 | ARHGAP19-SLIT1 | ARHGAP19-SLIT1 readthrough (NMD candidate) | Other | other | |
| chr3 | *LINC00578* | 9 | 223 | 24 | LINC00578 | long intergenic non-protein coding RNA 578 | Other | other | |
| chr11 | *SYTL2* | 11 | 394 | 24 | SYTL2 | synaptotagmin like 2 | Cytoplasm | other | |
| chr9 | *RORB* | 9 | 214 | 24 | RORB | RAR related orphan receptor B | Nucleus | d-dependent nuclear rec | |
| chr11 | *ADRBK1* | 11 | 524 | 24 | GRK2 | G protein-coupled receptor | Cytoplasm | kinase | |
| chr21 | *APP* | 9 | 208 | 24 | APP | amyloid beta precursor protein | Plasma Membrane | other | florbetaben F, florbetapir F18, bapineuzumab |
| chr19 | *NUMBL* | 11 | 429 | 24 | NUMBL | NUMB like endocytic adaptor | Cytoplasm | other | |
| chr1 | *USP33* | 10 | 223 | 24 | USP33 | ubiquitin specific peptidase 33 | Cytoplasm | peptidase | |
| chr1 | *GIPC2* | 11 | 221 | 24 | GIPC2 | GIPC PDZ domain containing family member 2 | Cytoplasm | other | |
| chr5 | *MAP1B* | 12 | 268 | 24 | MAP1B | microtubule associated protein | Cytoplasm | other | |
| chr14 | *ASB2* | 9 | 342 | 24 | ASB2 | ankyrin repeat and SOCS box containing 2 | Nucleus | transcription regulator | |
| chr5 | *HRH2* | 11 | 292 | 24 | HRH2 | histamine receptor H2 | Plasma Membrane | G-protein coupled recepto | nizatidine, epinastine, ranitidine, methantheline, dexamethasone/diphenhydramine/ranitidine, famotidine/ibuprofen, metiamide, tesmilifene, famotidine, doxepin, cimetidine, asenapine, buclizine, |
| chr3 | *ACOX2* | 10 | 378 | 24 | ACOX2 | acyl-CoA oxidase 2 | Cytoplasm | enzyme | |
| chrX | *TXLNG* | 10 | 341 | 24 | TXLNG | taxilin gamma | Nucleus | other | |
| chrX | *PIR* | 9 | 217 | 24 | PIR | pirin | Nucleus | transcription regulator | |
| chr13 | *CLDN10* | 12 | 330 | 24 | CLDN10 | claudin 10 | Plasma Membrane | other | |

| | | | | | | | | | |
|---|---|---|---|---|---|---|---|---|---|
| chr12 | *RP11-13A1.3* | 11 | 233 | 24 | | | | | |
| chr1 | *SLC16A1* | 10 | 306 | 24 | SLC16A1 | solute carrier family 16 member 1 | Plasma Membrane | transporter | AZD-3965 |
| chr1 | *PPAP2B* | 11 | 266 | 24 | PLPP3 | phospholipid phosphatase 3 | Plasma Membrane | phosphatase | |
| chr12 | *TMCC3* | 10 | 218 | 24 | TMCC3 | transmembrane and coiled-coil domain family 3 | Cytoplasm | other | |
| chr10 | *PTEN* | 10 | 229 | 24 | PTEN | phosphatase and tensin homolog | Cytoplasm | phosphatase | |
| chr16 | *ITGAX* | 11 | 441 | 24 | ITGAX | integrin subunit alpha X | Plasma Membrane | transmembrane receptor | |
| chr1 | *ECHDC2* | 10 | 330 | 24 | ECHDC2 | enoyl-CoA hydratase domain containing 2 | Other | other | |
| chr15 | *GLCE* | 10 | 251 | 24 | GLCE | glucuronic acid epimerase | Cytoplasm | enzyme | |
| chr1 | *EFHD2* | 10 | 485 | 24 | EFHD2 | EF-hand domain family member | Other | other | |
| chr7 | *KMT2E* | 9 | 207 | 24 | KMT2E | lysine methyltransferase 2E | Nucleus | enzyme | |
| chr1 | *EXO1* | 9 | 329 | 24 | EXO1 | exonuclease 1 | Nucleus | enzyme | |
| chr20 | *ZNFX1* | 10 | 329 | 24 | ZNFX1 | zinc finger NFX1-type containing | Nucleus | transcription regulator | |
| chr17 | *HN1* | 10 | 330 | 24 | JPT1 | Jupiter microtubule associated homolog 1 | Nucleus | other | |
| chr11 | *CCDC67* | 10 | 430 | 24 | DEUP1 | deuterosome assembly protein 1 | Other | other | |
| chr17 | *ADAM11* | 11 | 505 | 24 | ADAM11 | ADAM metallopeptidase domain | Plasma Membrane | peptidase | |
| chrX | *RP2* | 9 | 271 | 24 | RP2 | RP2 activator of ARL3 GTPase | Cytoplasm | enzyme | |
| chr17 | *RP5-1029K10.2* | 10 | 386 | 24 | | | | | |
| chr1 | *RNF19B* | 9 | 271 | 24 | RNF19B | ring finger protein 19B | Cytoplasm | enzyme | |
| chr3 | *CNTN3* | 11 | 202 | 24 | CNTN3 | contactin 3 | Plasma Membrane | other | |
| chr3 | *RP11-454C18.2* | 9 | 329 | 24 | | | | | |
| chr11 | *VPS37C* | 10 | 580 | 24 | VPS37C | VPS37C subunit of ESCRT-I | Cytoplasm | other | |
| chr12 | *HCFC2* | 8 | 216 | 24 | HCFC2 | host cell factor C2 | Nucleus | transcription regulator | |
| chrX | *DOCK11* | 8 | 197 | 24 | DOCK11 | dedicator of cytokinesis 11 | Cytoplasm | other | |
| chr8 | *LINC00589* | 10 | 216 | 24 | LINC00589 | long intergenic non-protein coding RNA 589 | Other | other | |
| chr17 | *USP6* | 11 | 390 | 24 | USP6 | ubiquitin specific peptidase 6 | Cytoplasm | peptidase | |
| chr5 | *PDCD6* | 11 | 1022 | 24 | PDCD6 | programmed cell death 6 | Cytoplasm | other | |
| chr9 | *SMARCA2* | 10 | 270 | 24 | SMARCA2 | SWI/SNF related, matrix associated, actin dependent regulator of chromatin, subfamily | Nucleus | transcription regulator | |
| chr8 | *IKBKB* | 9 | 370 | 24 | IKBKB | inhibitor of nuclear factor kappa B kinase subunit beta | Cytoplasm | kinase | auranofin, IKKbeta inhibitor |
| chr3 | *TMEM40* | 10 | 358 | 24 | TMEM40 | transmembrane protein 40 | Other | other | |
| chr19 | *CRLF1* | 11 | 411 | 24 | CRLF1 | cytokine receptor like factor 1 | Extracellular Space | other | |
| chr5 | *CTD-2587M2.1* | 11 | 333 | 24 | | | | | |
| chr4 | *BMPR1B* | 9 | 215 | 24 | BMPR1B | bone morphogenetic protein receptor type 1B | Plasma Membrane | kinase | |
| chr17 | *ABCA8* | 9 | 254 | 24 | ABCA8 | ATP binding cassette subfamily A member 8 | Plasma Membrane | transporter | |
| chr14 | *FOXN3* | 9 | 243 | 24 | FOXN3 | forkhead box N3 | Nucleus | transcription regulator | |
| chr12 | *COQ5* | 11 | 283 | 24 | COQ5 | coenzyme Q5, methyltransferase | Cytoplasm | enzyme | |
| chr17 | *RP11-344E13.3* | 10 | 427 | 24 | | | | | |
| chr19 | *MARCH2;* | 10 | 329 | 24 | | | | | |
| chr17 | *SLC52A1* | 10 | 305 | 24 | SLC52A1 | solute carrier family 52 member 1 | Plasma Membrane | transporter | |
| chr1 | *PEAR1* | 10 | 751 | 24 | PEAR1 | platelet endothelial aggregation receptor 1 | Plasma Membrane | other | |
| chr17 | *TMEM105* | 11 | 509 | 24 | TMEM105 | TMEM105 long non-coding RNA | Other | other | |
| chr4 | *LARP1B* | 9 | 254 | 24 | LARP1B | La ribonucleoprotein domain family member 1B | Extracellular Space | other | |
| chr19 | *PLEKHA4* | 10 | 347 | 24 | PLEKHA4 | pleckstrin homology domain containing A4 | Cytoplasm | other | |
| chr7 | *RUNDC3B* | 10 | 297 | 24 | RUNDC3B | RUN domain containing 3B | Other | other | |
| chr5 | *RP11-804N13.1* | 10 | 541 | 24 | | | | | |
| chr8 | *C8orf12* | 11 | 382 | 24 | FAM167A-AS1 | FAM167A antisense RNA 1 | Other | other | |
| chr15 | *SHC4* | 10 | 247 | 24 | SHC4 | SHC adaptor protein 4 | Cytoplasm | other | |
| chr11 | *RRM1* | 8 | 432 | 24 | RRM1 | ribonucleotide reductase catalytic subunit M1 | Nucleus | enzyme | L-asparaginase/gemcitabine/oxaliplatin, cyclophosphamide/fludarabine phosphate, alemtuzumab/cyclophosphamide/fludarabine phosphate/rituximab, ribonucleotide reductase inhibitor, cetuximab/gemcitabine, clofarabine/filgrastim/idarubicin, docetaxel/gemcitabine/vincristine, cladribine/cytarabine/filgrastim/idarubicin/plerixafor, cladribine, cytarabine/fludarabine [illegible] |
| chr1 | *NEK7* | 11 | 360 | 24 | NEK7 | NIMA related kinase 7 | Nucleus | kinase | |
| chr7 | *RP11-745C15.2* | 9 | 388 | 24 | | | | | |
| chr6 | *TEAD3* | 11 | 503 | 24 | TEAD3 | TEA domain transcription factor 3 | Nucleus | transcription regulator | |
| chr6 | *MAP3K5* | 8 | 181 | 24 | MAP3K5 | mitogen-activated protein kinase kinase kinase 5 | Cytoplasm | kinase | selonsertib |
| chr1 | *HP1BP3* | 9 | 228 | 24 | HP1BP3 | heterochromatin protein 1 binding protein 3 | Nucleus | other | |

| | | | | | | | | | |
|---|---|---|---|---|---|---|---|---|---|
| chr9 | *NAA35* | 9 | 199 | 24 | NAA35 | N(alpha)-acetyltransferase 35, NatC auxiliary subunit | Cytoplasm | other | |
| chr13 | *RP11-141M1.3* | 9 | 219 | 24 | | | | | |
| chr12 | *NR1H4* | 10 | 234 | 24 | NR1H4 | nuclear receptor subfamily 1 group H member 4 | Nucleus | d-dependent nuclear rec | apomine, obeticholic acid |
| chr12 | *CASC1* | 9 | 267 | 24 | CASC1 | cancer susceptibility 1 | Cytoplasm | other | |
| chr7 | *RSBN1L-AS1* | 10 | 314 | 24 | APTR | Alu-mediated CDKN1A/p21 transcriptional regulator | Other | other | |
| chr17 | *TUSC5* | 10 | 548 | 24 | TRARG1 | trafficking regulator of GLUT4 (SLC2A4) 1 | Plasma Membrane | other | |
| chr22 | *TBX1* | 11 | 501 | 24 | TBX1 | T-box transcription factor 1 | Nucleus | transcription regulator | |
| chr2 | *ERBB4* | 9 | 242 | 24 | ERBB4 | erb-b2 receptor tyrosine kinase 4 | Plasma Membrane | kinase | afatinib, afatinib/paclitaxel, pirotinib, afatinib/cetuximab, neratinib/paclitaxel, erlotinib/osimertinib, JNJ-26483327, poziotinib, dacomitinib, neratinib, afatinib/osimertinib, BMS-599626, allitinib, |
| chr20 | *RP11-445H22.3* | 11 | 412 | 24 | | | | | |
| chr17 | *ARMC7* | 10 | 429 | 24 | ARMC7 | armadillo repeat containing 7 | Cytoplasm | other | |
| chr1 | *MAP7D1* | 10 | 523 | 24 | MAP7D1 | MAP7 domain containing 1 | Cytoplasm | other | |
| chr20 | *RASSF2* | 9 | 294 | 24 | RASSF2 | Ras association domain family member 2 | Nucleus | kinase | |
| chr10 | *MAPK8* | 9 | 374 | 24 | MAPK8 | mitogen-activated protein kinase 8 | Cytoplasm | kinase | aplidine, cetuximab/encorafenib, binimetinib/encorafenib, encorafenib, binimetinib/cetuximab/encorafenib, binimetinib/encorafen |
| chr17 | *CDC42EP4* | 10 | 401 | 24 | CDC42EP4 | CDC42 effector protein 4 | Cytoplasm | other | |
| chr12 | *RP11-446N19.1* | 11 | 452 | 24 | | | | | |
| chr10 | *PPP2R2D* | 9 | 431 | 24 | PPP2R2D | protein phosphatase 2 regulatory subunit Bdelta | Nucleus | other | |
| chr9 | *SUSD1* | 10 | 216 | 24 | SUSD1 | sushi domain containing 1 | Other | other | |
| chr20 | *VAPB* | 9 | 359 | 24 | VAPB | VAMP associated protein B and | Plasma Membrane | other | |
| chr19 | *C5AR1* | 9 | 308 | 24 | C5AR1 | complement C5a receptor 1 | Plasma Membrane | G-protein coupled recepto | IPH5401 |
| chr22 | *SUN2* | 10 | 462 | 24 | SUN2 | Sad1 and UNC84 domain containing 2 | Nucleus | other | |
| chr19 | *ZNF444* | 11 | 424 | 24 | ZNF444 | zinc finger protein 444 | Nucleus | transcription regulator | |
| chr16 | *DNAAF1* | 10 | 335 | 24 | DNAAF1 | dynein axonemal assembly factor | Cytoplasm | other | |
| chr4 | *KIAA0232* | 10 | 193 | 24 | KIAA0232 | KIAA0232 | Extracellular Space | other | |
| chr17 | *GID4* | 11 | 290 | 24 | GID4 | GID complex subunit 4 homolog | Nucleus | other | |
| chr4 | *PROM1* | 9 | 265 | 24 | PROM1 | prominin 1 | Plasma Membrane | other | anti-CD133-PE38-KDEL fusion protein |
| chr11 | *NR1H3* | 10 | 729 | 24 | NR1H3 | nuclear receptor subfamily 1 group H member 3 | Nucleus | d-dependent nuclear rec | |
| chr5 | *MEGF10* | 10 | 255 | 24 | MEGF10 | multiple EGF like domains 10 | Plasma Membrane | transmembrane receptor | |
| chr19 | *CD22* | 9 | 439 | 24 | CD22 | CD22 molecule | Plasma Membrane | transmembrane receptor | combotox, DT2219ARL, yttrium y 90 epratuzumab tetraxetan, dasatinib/inotuzumab ozogamicin, RFB4-dgRTA, inotuzumab ozogamicin, BL22 immunotoxin, TRPH-222, immunotoxin HA22, imatinib/inotuzumab ozogamicin, epratuzumab-Cys-tesirine, |
| chr11 | *MTCH2* | 10 | 529 | 24 | MTCH2 | mitochondrial carrier 2 | Cytoplasm | other | |
| chr1 | *RP11-76N22.2* | 8 | 205 | 24 | | | | | |
| chr2 | *GEMIN6* | 10 | 316 | 24 | GEMIN6 | gem nuclear organelle associated protein 6 | Nucleus | other | |
| chr2 | *AC010733.4* | 10 | 263 | 24 | | | | | |
| chr12 | *MSRB3* | 10 | 236 | 24 | MSRB3 | methionine sulfoxide reductase | Cytoplasm | enzyme | |
| chr10 | *BEND7* | 9 | 251 | 24 | BEND7 | BEN domain containing 7 | Cytoplasm | other | |
| chr12 | *TCHP* | 10 | 436 | 24 | TCHP | trichoplein keratin filament | Cytoplasm | other | |
| chr1 | *NSL1* | 10 | 324 | 24 | NSL1 | NSL1 component of MIS12 kinetochore complex | Nucleus | other | |
| chr17 | *MLLT6* | 10 | 455 | 24 | MLLT6 | MLLT6, PHD finger containing | Nucleus | transcription regulator | |
| chrX | *ASB9* | 10 | 258 | 24 | ASB9 | ankyrin repeat and SOCS box containing 9 | Nucleus | transcription regulator | |
| chr10 | *HELLS* | 9 | 297 | 24 | HELLS | helicase, lymphoid specific | Nucleus | enzyme | |
| chr16 | *SLC12A4* | 10 | 462 | 24 | SLC12A4 | solute carrier family 12 member 4 | Plasma Membrane | transporter | bumetanide |
| chr22 | *RFPL1S* | 9 | 284 | 24 | RFPL1S | RFPL1 antisense RNA 1 | Other | other | |
| chr7 | *CCZ1* | 11 | 493 | 24 | CCZ1/CCZ1B | CCZ1 homolog B, vacuolar protein trafficking and biogenesis | Cytoplasm | other | |
| chr12 | *LPAR5* | 10 | 406 | 24 | LPAR5 | lysophosphatidic acid receptor 5 | Plasma Membrane | G-protein coupled recepto | |

| | | | | | | | | | |
|---|---|---|---|---|---|---|---|---|---|
| chr5 | *SEPT8* | 11 | 366 | 24 | SEPTIN8 | septin 8 | Extracellular Space | other | |
| chr14 | *STRN3* | 9 | 202 | 24 | STRN3 | striatin 3 | Nucleus | transcription regulator | |
| chr15 | *GLDN* | 11 | 253 | 24 | GLDN | gliomedin | Cytoplasm | other | |
| chr8 | *TPD52* | 9 | 215 | 24 | TPD52 | tumor protein D52 | Cytoplasm | other | |
| chr18 | *TSHZ1* | 8 | 296 | 24 | TSHZ1 | teashirt zinc finger homeobox 1 | Nucleus | transcription regulator | |
| chr1 | *LINC00970* | 11 | 299 | 24 | LINC00970 | long intergenic non-protein coding RNA 970 | Other | other | |
| chr19 | *ZNF682* | 10 | 240 | 24 | ZNF682 | zinc finger protein 682 | Nucleus | transcription regulator | |
| chr1 | *NSUN4* | 9 | 307 | 24 | NSUN4 | NOP2/Sun RNA | Cytoplasm | enzyme | |
| chr1 | *RPA2* | 9 | 302 | 24 | RPA2 | replication protein A2 | Nucleus | other | |
| chr5 | *EXOC3* | 11 | 1185 | 24 | EXOC3 | exocyst complex component 3 | Plasma Membrane | transporter | |
| chr1 | *FLVCR1* | 9 | 270 | 24 | FLVCR1 | feline leukemia virus subgroup C cellular receptor 1 | Plasma Membrane | transporter | |
| chr7 | *GS1-124K5.2* | 11 | 239 | 24 | | | | | |
| chr11 | *FRMD8* | 11 | 878 | 24 | FRMD8 | FERM domain containing 8 | Other | other | |
| chr15 | *TGM5* | 11 | 315 | 24 | TGM5 | transglutaminase 5 | Cytoplasm | enzyme | |
| chr15 | *RP11-184D12.1* | 10 | 286 | 24 | | | | | |
| chr5 | *C9* | 12 | 833 | 24 | C9 | complement C9 | Extracellular Space | other | |
| chrX | *FAAH2* | 11 | 267 | 24 | FAAH2 | fatty acid amide hydrolase 2 | Cytoplasm | enzyme | |
| chr10 | *MARCH5;* | 10 | 225 | 24 | | | | | |
| chr7 | *FKBP6* | 11 | 306 | 24 | FKBP6 | FKBP prolyl isomerase 6 | Nucleus | enzyme | |
| chr17 | *SRP68* | 10 | 248 | 24 | SRP68 | signal recognition particle 68 | Nucleus | other | |
| chr2 | *PDE11A* | 9 | 273 | 24 | PDE11A | phosphodiesterase 11A | Cytoplasm | enzyme | pentoxifylline, tolbutamide, tadalafil, theophylline, nitroglycerin, dyphylline, |
| chr16 | *PMFBP1* | 11 | 291 | 24 | PMFBP1 | polyamine modulated factor 1 binding protein 1 | Cytoplasm | other | |
| chr16 | *TVP23A* | 11 | 255 | 24 | TVP23A | trans-golgi network vesicle protein 23 homolog A | Other | other | |
| chr1 | *SYT11* | 10 | 371 | 24 | SYT11 | synaptotagmin 11 | Cytoplasm | transporter | |
| chr5 | *ARHGEF37* | 9 | 241 | 24 | ARHGEF37 | Rho guanine nucleotide exchange factor 37 | Other | other | |
| chr16 | *FUK* | 11 | 415 | 24 | FCSK | fucose kinase | Cytoplasm | kinase | |
| chr6 | *PRPH2* | 10 | 337 | 24 | PRPH2 | peripherin 2 | Plasma Membrane | transmembrane recepto | |
| chr2 | *FHL2* | 11 | 395 | 23 | FHL2 | four and a half LIM domains 2 | Nucleus | transcription regulator | |
| chr12 | *CABP1* | 10 | 376 | 23 | CABP1 | calcium binding protein 1 | Plasma Membrane | other | |
| chr19 | *CTC-513N18.6* | 12 | 217 | 23 | | | | | |
| chr11 | *ARHGAP20* | 9 | 276 | 23 | ARHGAP20 | Rho GTPase activating protein 20 | Cytoplasm | other | |
| chr19 | *PLAUR* | 10 | 321 | 23 | PLAUR | plasminogen activator, urokinase receptor | Plasma Membrane | transmembrane recepto | |
| chr5 | *RAD17* | 9 | 215 | 23 | RAD17 | RAD17 checkpoint clamp loader component | Nucleus | enzyme | |
| chr9 | *TLN1* | 9 | 348 | 23 | TLN1 | talin 1 | Plasma Membrane | other | |
| chr5 | *CTC-228N24.3* | 9 | 217 | 23 | | | | | |
| chr1 | *GPR157* | 10 | 400 | 23 | GPR157 | G protein-coupled receptor 157 | Plasma Membrane | -protein coupled recepto | |
| chr17 | *DHX40* | 9 | 268 | 23 | DHX40 | DEAH-box helicase 40 | Other | enzyme | |
| chr1 | *C1orf159* | 12 | 454 | 23 | C1orf159 | chromosome 1 open reading frame 159 | Other | other | |
| chr9 | *WDR5* | 11 | 681 | 23 | WDR5 | WD repeat domain 5 | Nucleus | transcription regulator | |
| chr7 | *RFC2* | 10 | 334 | 23 | RFC2 | replication factor C subunit 2 | Nucleus | other | |
| chr15 | *RP11-34F13.3* | 10 | 298 | 23 | | | | | |
| chr16 | *CRAMP1L* | 9 | 337 | 23 | CRAMP1 | cramped chromatin regulator homolog 1 | Other | other | |
| chr11 | *NCAPD3* | 10 | 286 | 23 | NCAPD3 | non-SMC condensin II complex subunit D3 | Nucleus | other | |
| chr17 | *TUBD1* | 10 | 225 | 23 | TUBD1 | tubulin delta 1 | Cytoplasm | other | docetaxel/gemcitabin e/vincristine, cyclophosphamide/epirubicin/5-fluorouracil/paclitaxel, bevacizumab/paclitaxel/topotecan, colchicine/probenecid, bevacizumab/docetaxel/prednisone/thalidomide/zoledronic acid, gemcitabine/paclitaxel, L-asparaginase/daunorubicin/dexamethasone/vincristine, prednisone/vincristine, docetaxel/5-fluorouracil, vinflunine, L-[illegible] |
| chr1 | *RP11-466L17.1* | 12 | 227 | 23 | | | | | |
| chr1 | *CELSR2* | 12 | 650 | 23 | CELSR2 | cadherin EGF LAG seven-pass G-type receptor 2 | Plasma Membrane | -protein coupled recepto | |
| chr2 | *CDKL4* | 9 | 295 | 23 | CDKL4 | cyclin dependent kinase like 4 | Nucleus | kinase | |
| chr5 | *RP11-1415C14.4* | 9 | 454 | 23 | | | | | |
| chr2 | *AC104782.3* | 10 | 402 | 23 | | | | | |
| chr9 | *LINC00963* | 10 | 673 | 23 | LINC00963 | long intergenic non-protein coding RNA 963 | Other | other | |

| | | | | | | | | | |
|---|---|---|---|---|---|---|---|---|---|
| chr5 | *KLHL3* | 11 | 226 | 23 | KLHL3 | kelch like family member 3 | Cytoplasm | other | |
| chr6 | *STXBP5* | 8 | 213 | 23 | STXBP5 | syntaxin binding protein 5 | Plasma Membrane | other | |
| chr12 | *RP13-507P19.2* | 10 | 434 | 23 | | | | | |
| chr7 | *GHRHR* | 11 | 624 | 23 | GHRHR | growth hormone releasing hormone receptor | Plasma Membrane | G-protein coupled recepto | sermorelin, tesamorelin |
| chr15 | *SEC11A* | 9 | 289 | 23 | SEC11A | SEC11 homolog A, signal peptidase complex subunit | Cytoplasm | peptidase | |
| chr9 | *CRB2* | 12 | 709 | 23 | CRB2 | crumbs cell polarity complex component 2 | Extracellular Space | other | |
| chr12 | *POU6F1* | 11 | 697 | 23 | POU6F1 | POU class 6 homeobox 1 | Nucleus | transcription regulator | |
| chr16 | *GPRC5B* | 11 | 334 | 23 | GPRC5B | G protein-coupled receptor class C group 5 member B | Plasma Membrane | G-protein coupled recepto | |
| chr22 | *ATP6V1E1* | 8 | 229 | 23 | ATP6V1E1 | ATPase H+ transporting V1 subunit E1 | Cytoplasm | transporter | |
| chr8 | *GML* | 11 | 295 | 23 | GML | glycosylphosphatidylinositol anchored molecule like | Plasma Membrane | other | |
| chr9 | *FAM157B* | 10 | 633 | 23 | AM157A/FAM157 | family with sequence similarity 157 member A | Other | other | |
| chr12 | *RP11-405A12.2* | 9 | 307 | 23 | | | | | |
| chr2 | *ANKRD44* | 10 | 214 | 23 | ANKRD44 | ankyrin repeat domain 44 | Other | other | |
| chr7 | *DPY19L1* | 10 | 284 | 23 | DPY19L1 | dpy-19 like C- | Other | other | |
| chr1 | *MPZL1* | 9 | 268 | 23 | MPZL1 | myelin protein zero like 1 | Plasma Membrane | other | |
| chr12 | *GTF2H3* | 9 | 242 | 23 | GTF2H3 | general transcription factor IIH subunit 3 | Nucleus | transcription regulator | |
| chr2 | *RAPGEF4* | 9 | 207 | 23 | RAPGEF4 | Rap guanine nucleotide exchange factor 4 | Cytoplasm | other | |
| chr20 | *TCEA2* | 12 | 1065 | 23 | TCEA2 | transcription elongation factor A2 | Nucleus | transcription regulator | |
| chr9 | *RP11-65J3.3* | 11 | 569 | 23 | | | | | |
| chr1 | *SPOCD1* | 10 | 383 | 23 | SPOCD1 | SPOC domain containing 1 | Other | other | |
| chr16 | *RP11-77K12.8* | 10 | 269 | 23 | | | | | |
| chr11 | *TNKS1BP1* | 10 | 424 | 23 | TNKS1BP1 | tankyrase 1 binding protein 1 | Nucleus | other | |
| chr6 | *SPDEF* | 10 | 504 | 23 | SPDEF | SAM pointed domain containing ETS transcription factor | Nucleus | transcription regulator | |
| chr17 | *SERPINF1* | 10 | 400 | 23 | SERPINF1 | serpin family F member 1 | Extracellular Space | other | |
| chr3 | *COL8A1* | 10 | 334 | 23 | COL8A1 | collagen type VIII alpha 1 chain | Extracellular Space | other | collagenase clostridium |
| chr3 | *DLG1* | 7 | 188 | 23 | DLG1 | discs large MAGUK scaffold | Plasma Membrane | kinase | |
| chr1 | *FAAH* | 10 | 469 | 23 | FAAH | fatty acid amide hydrolase | Plasma Membrane | enzyme | |
| chr6 | *TJAP1* | 9 | 473 | 23 | TJAP1 | tight junction associated protein 1 | Plasma Membrane | other | |
| chr20 | *LPIN3* | 10 | 644 | 23 | LPIN3 | lipin 3 | Nucleus | phosphatase | |
| chr1 | *URB2* | 10 | 382 | 23 | URB2 | URB2 ribosome biogenesis | Nucleus | other | |
| chr2 | *MALL* | 11 | 582 | 23 | MALL | mal, T cell differentiation protein | Plasma Membrane | other | |
| chr7 | *NUP205* | 9 | 280 | 23 | NUP205 | nucleoporin 205 | Nucleus | other | |
| chr22 | *MTMR3* | 8 | 193 | 23 | MTMR3 | myotubularin related protein 3 | Cytoplasm | phosphatase | |
| chr1 | *TCEANC2* | 10 | 255 | 23 | TCEANC2 | transcription elongation factor A N-terminal and central domain containing 2 | Other | other | |
| chr7 | *FAM185A* | 9 | 259 | 23 | FAM185A | family with sequence similarity 185 member A | Cytoplasm | other | |
| chr7 | *TFEC* | 10 | 245 | 23 | TFEC | transcription factor EC | Nucleus | transcription regulator | |
| chr1 | *BPNT1* | 9 | 285 | 23 | BPNT1 | 3'(2'), 5'-bisphosphate | Nucleus | phosphatase | |
| chr2 | *ZC3H6* | 9 | 252 | 23 | ZC3H6 | zinc finger CCCH-type containing | Other | other | |
| chr7 | *SHFM1* | 9 | 330 | 23 | SEM1 | SEM1 26S proteasome complex subunit | Nucleus | peptidase | |
| chr1 | *MARK1* | 9 | 291 | 23 | MARK1 | microtubule affinity regulating kinase 1 | Cytoplasm | kinase | |
| chr12 | *RP11-768F21.1* | 9 | 218 | 23 | | | | | |
| chr1 | *C1orf61* | 11 | 446 | 23 | C1orf61 | chromosome 1 open reading frame 61 | Cytoplasm | other | |
| chr2 | *ANKRD36C* | 10 | 362 | 23 | ANKRD36C | ankyrin repeat domain 36C | Other | other | |
| chr11 | *SLC37A2* | 9 | 454 | 23 | SLC37A2 | solute carrier family 37 member 2 | Cytoplasm | transporter | |
| chr8 | *RP11-238K6.1* | 10 | 206 | 23 | | | | | |
| chr12 | *CNOT2* | 8 | 236 | 23 | CNOT2 | CCR4-NOT transcription complex subunit 2 | Nucleus | transcription regulator | |
| chr3 | *MKRN2* | 10 | 248 | 23 | MKRN2 | makorin ring finger protein 2 | Other | other | |
| chr16 | *CDH11* | 11 | 289 | 23 | CDH11 | cadherin 11 | Plasma Membrane | other | |
| chr16 | *RP11-523L20.2* | 11 | 231 | 23 | | | | | |
| chr1 | *ATP8B2* | 10 | 461 | 23 | ATP8B2 | ATPase phospholipid transporting 8B2 | Plasma Membrane | transporter | |
| chr10 | *FAM178A* | 9 | 234 | 23 | SLF2 | SMC5-SMC6 complex localization factor 2 | Extracellular Space | other | |
| chr3 | *VPS8* | 9 | 209 | 23 | VPS8 | VPS8 subunit of CORVET | Cytoplasm | other | |
| chr18 | *ABHD3* | 9 | 218 | 23 | ABHD3 | abhydrolase domain containing 3 | Plasma Membrane | enzyme | |
| chr19 | *WIZ* | 11 | 486 | 23 | WIZ | WIZ zinc finger | Nucleus | transcription regulator | |
| chr20 | *PCSK2* | 9 | 307 | 23 | PCSK2 | proprotein convertase subtilisin/kexin type 2 | Extracellular Space | peptidase | |
| chr10 | *ZFAND4* | 9 | 358 | 23 | ZFAND4 | zinc finger AN1-type containing 4 | Other | other | |
| chr5 | *RNF180* | 9 | 261 | 23 | RNF180 | ring finger protein 180 | Cytoplasm | enzyme | |
| chr16 | *SLC9A5* | 10 | 392 | 23 | SLC9A5 | solute carrier family 9 member A5 | Plasma Membrane | transporter | |
| chr1 | *PLA2G5* | 10 | 277 | 23 | PLA2G5 | phospholipase A2 group V | Extracellular Space | enzyme | varespladib, varespladib methyl |
| chr1 | *STIL* | 9 | 223 | 23 | STIL | STIL centriolar assembly protein | Nucleus | other | |
| chr7 | *KCTD7* | 10 | 329 | 23 | KCTD7 | potassium channel tetramerization domain containing | Plasma Membrane | ion channel | |
| chr5 | *SQSTM1* | 11 | 538 | 23 | SQSTM1 | sequestosome 1 | Cytoplasm | transcription regulator | |
| chr19 | *MOB3A* | 10 | 395 | 23 | MOB3A | MOB kinase activator 3A | Other | other | |
| chr17 | *DRG2* | 10 | 417 | 23 | DRG2 | developmentally regulated GTP binding protein 2 | Cytoplasm | enzyme | |

| | | | | | | | | | |
|---|---|---|---|---|---|---|---|---|---|
| chr9 | *MAN1B1* | 11 | 952 | 23 | MAN1B1 | mannosidase alpha class 1B member 1 | Cytoplasm | enzyme | |
| chr12 | *ANKRD52* | 10 | 409 | 23 | ANKRD52 | ankyrin repeat domain 52 | Nucleus | transcription regulator | |
| chr1 | *B3GALNT2* | 11 | 262 | 23 | B3GALNT2 | beta-1,3-N-acetylgalactosaminyltransferase | Cytoplasm | enzyme | |
| chrX | *TAF1* | 8 | 217 | 23 | TAF1 | TATA-box binding protein associated factor 1 | Nucleus | transcription regulator | |
| chr19 | *PRKCG* | 11 | 399 | 23 | PRKCG | protein kinase C gamma | Cytoplasm | kinase | ingenol mebutate |
| chr17 | *NUFIP2* | 9 | 256 | 23 | NUFIP2 | nuclear FMR1 interacting protein | Cytoplasm | other | |
| chr2 | *EPC2* | 8 | 221 | 23 | EPC2 | enhancer of polycomb homolog 2 | Other | other | |
| chr22 | *ASCC2* | 9 | 290 | 23 | ASCC2 | activating signal cointegrator 1 complex subunit 2 | Nucleus | other | |
| chr1 | *RP4-669L17.10* | 11 | 460 | 23 | | | | | |
| chr6 | *RNF8* | 9 | 234 | 23 | RNF8 | ring finger protein 8 | Nucleus | enzyme | |
| chr1 | *ESPN* | 11 | 647 | 23 | ESPN | espin | Cytoplasm | other | |
| chr19 | *GRIN2D* | 10 | 366 | 23 | GRIN2D | glutamate ionotropic receptor NMDA type subunit 2D | Plasma Membrane | ion channel | dextromethorphan/morphine, nebostinel, besonprodil, delucemine, 1-aminocyclopropane-1-carboxylic acid, orphenadrine, dextromethorphan/promethazine, aspirin/caffeine/orphenadrine, neramexane, felbamate, chlorpheniramine/dextromethorphan/phenylephrine, memantine, donepezil/memantine, N-(2-indanyl)glycinamide, dextromethorphan, cycloserine, UK-240455, brompheniramine/de[illegible] |
| chr15 | *RP11-519G16.3* | 10 | 353 | 23 | | | | | |
| chr15 | *PKM* | 10 | 305 | 23 | PKM | pyruvate kinase M1/2 | Cytoplasm | kinase | CAP-232 |
| chr8 | *SLC30A8* | 10 | 340 | 23 | SLC30A8 | solute carrier family 30 member 8 | Cytoplasm | transporter | |
| chr5 | *PAPD4* | 9 | 221 | 23 | TENT2 | terminal nucleotidyltransferase 2 | Nucleus | enzyme | |
| chr11 | *STX3* | 10 | 246 | 23 | STX3 | syntaxin 3 | Plasma Membrane | transporter | |
| chr17 | *ADORA2B* | 10 | 355 | 23 | ADORA2B | adenosine A2b receptor | Plasma Membrane | G-protein coupled recepto | aspirin/caffeine/propoxyphene, aspirin/caffeine/orphenadrine, theophylline, acetaminophen/caffeine/dihydrocodeine, aspirin/butalbital/caffeine, aspirin/caffeine/dihydrocodeine, acetaminophen/aspirin/caffeine, enprofylline, acetaminophen/butalbital/caffeine/codeine, acetaminophen/butalbital/caffeine, aspirin/butalbital/caffeine/codeine, AB928, theophylline/tretinoin/valproic acid, PBF-1129, [illegible] |
| chr19 | *RPL28* | 10 | 376 | 23 | RPL28 | ribosomal protein L28 | Cytoplasm | other | |
| chr8 | *RALYL* | 9 | 256 | 23 | RALYL | RALY RNA binding protein like | Nucleus | other | |
| chr17 | *CANT1* | 10 | 456 | 23 | CANT1 | calcium activated nucleotidase 1 | Extracellular Space | enzyme | |
| chr6 | *E2F3* | 9 | 230 | 23 | E2F3 | E2F transcription factor 3 | Nucleus | transcription regulator | |
| chr1 | *KIF14* | 9 | 232 | 23 | KIF14 | kinesin family member 14 | Cytoplasm | enzyme | |
| chr16 | *KATNB1* | 10 | 469 | 23 | KATNB1 | katanin regulatory subunit B1 | Cytoplasm | enzyme | |
| chr5 | *CTD-3179P9.1* | 11 | 210 | 23 | | | | | |
| chr10 | *ZNF33B* | 10 | 297 | 23 | ZNF33B | zinc finger protein 33B | Nucleus | transcription regulator | |
| chr19 | *LYPD5* | 9 | 320 | 23 | LYPD5 | LY6/PLAUR domain containing 5 | Other | other | |
| chr4 | *RXFP1* | 8 | 209 | 23 | RXFP1 | relaxin family peptide receptor 1 | Plasma Membrane | G-protein coupled recepto | relaxin |
| chr2 | *RUFY4* | 8 | 319 | 23 | RUFY4 | RUN and FYVE domain | Other | other | |
| chr16 | *SMG1P1* | 9 | 346 | 23 | SMG1P1 | SMG1 pseudogene 1 | Other | other | |
| chr2 | *AC007040.8* | 9 | 321 | 23 | | | | | |
| chr6 | *TTLL2* | 10 | 542 | 23 | TTLL2 | tubulin tyrosine ligase like 2 | Other | other | |
| chr19 | *LRP3* | 11 | 406 | 23 | LRP3 | LDL receptor related protein 3 | Plasma Membrane | transmembrane recepto | |
| chr2 | *RP11-13J10.1* | 11 | 324 | 23 | | | | | |
| chr14 | *AHNAK2* | 11 | 517 | 23 | AHNAK2 | AHNAK nucleoprotein 2 | Cytoplasm | other | |
| chr3 | *LRCH3* | 7 | 219 | 23 | LRCH3 | leucine rich repeats and calponin homology domain containing 3 | Cytoplasm | other | |
| chr17 | *DDX42* | 9 | 247 | 23 | DDX42 | DEAD-box helicase 42 | Cytoplasm | enzyme | |
| chr1 | *RP11-399E6.1* | 11 | 316 | 23 | | | | | |

| | | | | | | | | | |
|---|---|---|---|---|---|---|---|---|---|
| chr21 | *UBE2G2* | 7 | 229 | 23 | UBE2G2 | ubiquitin conjugating enzyme E2 | Cytoplasm | enzyme | |
| chr11 | *RP11-347E10.1* | 9 | 230 | 23 | | | | | |
| chr12 | *KRT86* | 11 | 634 | 23 | KRT86 | keratin 86 | Cytoplasm | other | |
| chr1 | *TIE1* | 9 | 415 | 23 | TIE1 | tyrosine kinase with immunoglobulin like and EGF like | Plasma Membrane | kinase | |
| chr16 | *CTC-786C10.1* | 11 | 532 | 23 | | | | | |
| chr7 | *MPP6* | 9 | 271 | 23 | MPP6 | membrane palmitoylated protein | Plasma Membrane | kinase | |
| chr21 | *UBASH3A* | 9 | 346 | 23 | UBASH3A | ubiquitin associated and SH3 domain containing A | Cytoplasm | enzyme | |
| chr5 | *FAM151B* | 9 | 226 | 23 | FAM151B | family with sequence similarity 151 member B | Other | other | |
| chr3 | *ACAD9* | 10 | 440 | 23 | ACAD9 | acyl-CoA dehydrogenase family member 9 | Cytoplasm | enzyme | |
| chr16 | *LCMT1* | 10 | 305 | 23 | LCMT1 | leucine carboxyl | Cytoplasm | enzyme | |
| chr11 | *RP13-631K18.2* | 10 | 447 | 23 | | | | | |
| chr11 | *ADAMTS8* | 11 | 472 | 23 | ADAMTS8 | ADAM metallopeptidase with thrombospondin type 1 motif 8 | Extracellular Space | peptidase | |
| chr17 | *NSRP1* | 8 | 242 | 23 | NSRP1 | nuclear speckle splicing regulatory protein 1 | Nucleus | other | |
| chr11 | *RP11-672A2.3* | 10 | 576 | 23 | | | | | |
| chr15 | *COX5A* | 9 | 297 | 23 | COX5A | cytochrome c oxidase subunit 5A | Cytoplasm | enzyme | |
| chr11 | *OR9Q1* | 11 | 244 | 23 | OR9Q1 | olfactory receptor family 9 subfamily Q member 1 | Plasma Membrane | G-protein coupled recepto | |
| chr18 | *SLC14A2* | 10 | 225 | 23 | SLC14A2 | solute carrier family 14 member 2 | Plasma Membrane | transporter | |
| chr9 | *RP11-125B21.2* | 10 | 214 | 23 | | | | | |
| chr17 | *UNC45B* | 11 | 395 | 23 | UNC45B | unc-45 myosin chaperone B | Cytoplasm | other | |
| chr19 | *PLIN3* | 10 | 356 | 23 | PLIN3 | perilipin 3 | Cytoplasm | other | |
| chr2 | *MEIS1* | 9 | 219 | 23 | MEIS1 | Meis homeobox 1 | Nucleus | transcription regulator | |
| chr17 | *RP1-41C23.1* | 10 | 344 | 23 | | | | | |
| chr4 | *PDLIM5* | 7 | 171 | 23 | PDLIM5 | PDZ and LIM domain 5 | Cytoplasm | other | |
| chr15 | *KATNBL1* | 10 | 308 | 23 | KATNBL1 | katanin regulatory subunit B1 like | Nucleus | other | |
| chr19 | *ILF3* | 9 | 358 | 23 | ILF3 | interleukin enhancer binding | Nucleus | transcription regulator | |
| chr3 | *ACAP2* | 7 | 187 | 23 | ACAP2 | ArfGAP with coiled-coil, ankyrin repeat and PH domains 2 | Nucleus | other | |
| chr5 | *ZNF454* | 10 | 315 | 23 | ZNF454 | zinc finger protein 454 | Nucleus | transcription regulator | |
| chr17 | *ZBTB4* | 11 | 408 | 23 | ZBTB4 | zinc finger and BTB domain containing 4 | Nucleus | transcription regulator | |
| chr15 | *PLA2G4D* | 9 | 421 | 23 | PLA2G4D | phospholipase A2 group IVD | Cytoplasm | enzyme | |
| chr17 | *ATAD5* | 9 | 218 | 23 | ATAD5 | ATPase family AAA domain containing 5 | Nucleus | enzyme | |
| chr5 | *NAIP* | 9 | 244 | 23 | NAIP | NLR family apoptosis inhibitory protein | Cytoplasm | other | |
| chr11 | *ZNF195* | 10 | 501 | 23 | ZNF195 | zinc finger protein 195 | Nucleus | transcription regulator | |
| chr9 | *SPATA6L* | 10 | 246 | 23 | SPATA6L | spermatogenesis associated 6 | Other | other | |
| chr12 | *RP11-110A12.2* | 11 | 372 | 23 | | | | | |
| chr5 | *CTB-43E15.2* | 10 | 477 | 23 | | | | | |
| chr8 | *PPP1R16A* | 11 | 488 | 23 | PPP1R16A | protein phosphatase 1 regulatory subunit 16A | Plasma Membrane | other | |
| chr2 | *ZAP70* | 11 | 613 | 23 | ZAP70 | zeta chain of T cell receptor associated protein kinase 70 | Plasma Membrane | kinase | |
| chr15 | *FANCI* | 10 | 298 | 23 | FANCI | FA complementation group I | Nucleus | other | |
| chr17 | *ATPAF2* | 10 | 312 | 23 | ATPAF2 | ATP synthase mitochondrial F1 complex assembly factor 2 | Cytoplasm | other | |
| chr6 | *SIRT5* | 9 | 308 | 23 | SIRT5 | sirtuin 5 | Cytoplasm | enzyme | suramin |
| chr10 | *CNNM1* | 10 | 231 | 23 | CNNM1 | cyclin and CBS domain divalent metal cation transport mediator 1 | Plasma Membrane | other | |
| chr1 | *GNPAT* | 9 | 293 | 23 | GNPAT | glyceronephosphate O-acyltransferase | Cytoplasm | enzyme | |
| chr1 | *C1orf112* | 10 | 271 | 23 | C1orf112 | chromosome 1 open reading frame 112 | Other | other | |
| chr11 | *SCGB1A1* | 9 | 454 | 23 | SCGB1A1 | secretoglobin family 1A member | Extracellular Space | cytokine | |
| chr21 | *FAM207A* | 11 | 511 | 23 | FAM207A | family with sequence similarity 207 member A | Other | other | |
| chr15 | *RP13-98N21.3* | 9 | 464 | 23 | | | | | |
| chr4 | *STX18-AS1* | 9 | 228 | 23 | STX18-AS1 | STX18 antisense RNA 1 (head to head) | Other | other | |
| chr13 | *PSPC1* | 8 | 195 | 23 | PSPC1 | paraspeckle component 1 | Nucleus | transcription regulator | |
| chr3 | *CDCP1* | 10 | 282 | 23 | CDCP1 | CUB domain containing protein 1 | Plasma Membrane | other | |
| chr20 | *PLCB1* | 8 | 238 | 23 | PLCB1 | phospholipase C beta 1 | Cytoplasm | enzyme | |
| chr12 | *MSI1* | 10 | 420 | 23 | MSI1 | musashi RNA binding protein 1 | Cytoplasm | other | |
| chr1 | *RP11-495P10.2* | 10 | 304 | 23 | | | | | |
| chr6 | *ZNF76* | 10 | 352 | 23 | ZNF76 | zinc finger protein 76 | Nucleus | transcription regulator | |
| chr11 | *RP11-867G2.8* | 10 | 466 | 23 | | | | | |
| chr5 | *RP11-1E3.1* | 10 | 233 | 23 | | | | | |
| chr12 | *ADCY6* | 10 | 590 | 23 | ADCY6 | adenylate cyclase 6 | Plasma Membrane | enzyme | |
| chr1 | *PKN2* | 10 | 204 | 23 | PKN2 | protein kinase N2 | Cytoplasm | kinase | |
| chr19 | *MAP3K10* | 11 | 359 | 23 | MAP3K10 | mitogen-activated protein kinase kinase kinase 10 | Cytoplasm | kinase | |
| chr3 | *METTL6* | 10 | 235 | 23 | METTL6 | methyltransferase like 6 | Other | enzyme | |
| chr1 | *AIM1L* | 10 | 340 | 23 | CRYBG2 | crystallin beta-gamma domain containing 2 | Extracellular Space | other | |
| chr12 | *RAB35* | 11 | 508 | 23 | RAB35 | RAB35, member RAS oncogene family | Cytoplasm | enzyme | |
| chr6 | *SMIM13* | 9 | 229 | 23 | SMIM13 | small integral membrane protein | Other | other | |
| chr7 | *MICALL2* | 12 | 627 | 23 | MICALL2 | MICAL like 2 | Cytoplasm | other | |
| chr3 | *CASR* | 11 | 401 | 23 | CASR | calcium sensing receptor | Plasma Membrane | G-protein coupled recepto | cinacalcet, |
| chr1 | *RP11-550P17.5* | 10 | 277 | 23 | | | | | |
| chr10 | *BMS1* | 9 | 305 | 23 | BMS1 | BMS1 ribosome biogenesis factor | Nucleus | other | |
| chr17 | *STXBP4* | 7 | 187 | 23 | STXBP4 | syntaxin binding protein 4 | Cytoplasm | other | |

| | | | | | | | | | |
|---|---|---|---|---|---|---|---|---|---|
| chr5 | *HSPA4* | 8 | 252 | 23 | HSPA4 | heat shock protein family A (Hsp70) member 4 | Cytoplasm | other | |
| chr16 | *RP11-57A19.4* | 10 | 286 | 23 | | | | | |
| chr16 | *ZNF263* | 10 | 344 | 23 | ZNF263 | zinc finger protein 263 | Nucleus | transcription regulator | |
| chr16 | *RP11-293B20.2* | 10 | 275 | 23 | | | | | |
| chr5 | *KIF2A* | 8 | 215 | 23 | KIF2A | kinesin family member 2A | Cytoplasm | other | |
| chr1 | *RP4-773N10.6* | 10 | 459 | 23 | | | | | |
| chr1 | *CELA3B* | 10 | 292 | 23 | CELA3B | chymotrypsin like elastase 3B | Other | peptidase | |
| chr17 | *SLC16A5* | 10 | 498 | 23 | SLC16A5 | solute carrier family 16 member 5 | Plasma Membrane | transporter | |
| chr8 | *TNFRSF10B* | 10 | 306 | 23 | TNFRSF10B | TNF receptor superfamily member 10b | Plasma Membrane | transmembrane receptor | dulanermin, DS-8273a, RO6874813, drozitumab, conatumumab, GEN1029, |
| chr19 | *ATP8B3* | 10 | 417 | 22 | ATP8B3 | ATPase phospholipid transporting 8B3 | Cytoplasm | transporter | |
| chr2 | *TAF1B* | 8 | 244 | 22 | TAF1B | TATA-box binding protein associated factor, RNA | Nucleus | transcription regulator | |
| chrX | *OTUD5* | 10 | 366 | 22 | OTUD5 | OTU deubiquitinase 5 | Cytoplasm | enzyme | |
| chr12 | *RP11-575F12.1* | 9 | 206 | 22 | | | | | |
| chr1 | *ALG6* | 8 | 204 | 22 | ALG6 | ALG6 alpha-1,3- | Cytoplasm | enzyme | |
| chr17 | *EVPL* | 11 | 568 | 22 | EVPL | envoplakin | Plasma Membrane | other | |
| chr20 | *GSS* | 9 | 379 | 22 | GSS | glutathione synthetase | Cytoplasm | enzyme | N-acetyl-L-cysteine |
| chr9 | *RP11-54D18.3* | 10 | 214 | 22 | | | | | |
| chr11 | *RNF169* | 8 | 325 | 22 | RNF169 | ring finger protein 169 | Nucleus | other | |
| chr19 | *ITPKC* | 10 | 356 | 22 | ITPKC | inositol-trisphosphate 3-kinase C | Cytoplasm | kinase | |
| chr18 | *SMAD7* | 8 | 343 | 22 | SMAD7 | SMAD family member 7 | Nucleus | transcription regulator | |
| chr1 | *KIAA0907* | 9 | 342 | 22 | KHDC4 | KH domain containing 4, pre-mRNA splicing factor | Nucleus | other | |
| chr1 | *PSMB2* | 9 | 242 | 22 | PSMB2 | proteasome subunit beta 2 | Cytoplasm | peptidase | bortezomib/cladribine/rituximab, bortezomib/dexamethasone/pomalidomide, bortezomib/doxorubicin, bortezomib/dexamethasone/thalidomide, bortezomib/paclitaxel, bortezomib/fulvestrant, bortezomib/rituximab, bortezomib/sorafenib, carfilzomib, bortezomib/lenalidomide, carfilzomib/dexamethasone/lenalidomide, bortezomib/dexamethasone/lenalidomide, bortezomib/dexamet [illegible] |
| chr10 | *RP11-215N21.1* | 9 | 257 | 22 | | | | | |
| chr2 | *CHST10* | 10 | 448 | 22 | CHST10 | carbohydrate sulfotransferase 10 | Cytoplasm | enzyme | |
| chr7 | *ICA1* | 11 | 291 | 22 | ICA1 | islet cell autoantigen 1 | Cytoplasm | other | |
| chr6 | *SLC29A1* | 10 | 407 | 22 | SLC29A1 | solute carrier family 29 member 1 (Augustine blood group) | Plasma Membrane | transporter | |
| chr12 | *PDE3A* | 8 | 200 | 22 | PDE3A | phosphodiesterase 3A | Cytoplasm | enzyme | cilostazol, pentoxifylline, tolbutamide, amrinone, anagrelide, enoximone, theophylline, |
| chr11 | *TMEM123* | 10 | 247 | 22 | TMEM123 | transmembrane protein 123 | Plasma Membrane | other | |
| chr3 | *ATP1B3* | 10 | 323 | 22 | ATP1B3 | ATPase Na+/K+ transporting subunit beta 3 | Plasma Membrane | transporter | |
| chr16 | *FAM57B* | 9 | 368 | 22 | TLCD3B | TLC domain containing 3B | Cytoplasm | enzyme | |
| chr1 | *RP11-296A18.3* | 10 | 209 | 22 | | | | | |
| chr1 | *TMEM9* | 9 | 556 | 22 | TMEM9 | transmembrane protein 9 | Cytoplasm | other | |
| chr9 | *NTMT1* | 10 | 600 | 22 | NTMT1 | N-terminal Xaa-Pro-Lys N-methyltransferase 1 | Nucleus | enzyme | |
| chr9 | *PHYHD1* | 10 | 426 | 22 | PHYHD1 | phytanoyl-CoA dioxygenase domain containing 1 | Other | other | |
| chr1 | *PIGK* | 9 | 220 | 22 | PIGK | phosphatidylinositol glycan anchor biosynthesis class K | Cytoplasm | peptidase | |
| chr12 | *BCL2L14* | 9 | 215 | 22 | BCL2L14 | BCL2 like 14 | Cytoplasm | other | |
| chr4 | *RP11-526A4.1* | 9 | 228 | 22 | | | | | |
| chr19 | *NLRP4* | 10 | 267 | 22 | NLRP4 | NLR family pyrin domain | Cytoplasm | other | |
| chr8 | *ZNF623* | 10 | 322 | 22 | ZNF623 | zinc finger protein 623 | Nucleus | transcription regulator | |
| chr20 | *CDH26* | 11 | 417 | 22 | CDH26 | cadherin 26 | Plasma Membrane | other | |
| chr11 | *JAM3* | 9 | 238 | 22 | JAM3 | junctional adhesion molecule 3 | Plasma Membrane | other | |
| chr9 | *DOCK8* | 9 | 236 | 22 | DOCK8 | dedicator of cytokinesis 8 | Cytoplasm | other | |
| chr4 | *FAT1* | 8 | 253 | 22 | FAT1 | FAT atypical cadherin 1 | Plasma Membrane | other | |
| chr1 | *LRRC38* | 10 | 258 | 22 | LRRC38 | leucine rich repeat containing 38 | Plasma Membrane | ion channel | |
| chr17 | *UTP6* | 9 | 210 | 22 | UTP6 | UTP6 small subunit processome component | Nucleus | other | |
| chr5 | *GOLPH3* | 10 | 727 | 22 | GOLPH3 | golgi phosphoprotein 3 | Cytoplasm | other | |

| | | | | | | | | | |
|---|---|---|---|---|---|---|---|---|---|
| chr16 | *FENDRR* | 10 | 388 | 22 | FENDRR | FOXF1 adjacent non-coding developmental regulatory RNA | Other | other | |
| chr17 | *CCDC144CP* | 9 | 259 | 22 | CCDC144CP | coiled-coil domain containing 144C, pseudogene | Other | other | |
| chr7 | *TFR2* | 10 | 366 | 22 | TFR2 | transferrin receptor 2 | Plasma Membrane | transporter | |
| chr2 | *SATB2* | 10 | 195 | 22 | SATB2 | SATB homeobox 2 | Nucleus | transcription regulator | |
| chr1 | *RPL22* | 10 | 447 | 22 | RPL22 | ribosomal protein L22 | Cytoplasm | translation regulator | |
| chr8 | *GTF2E2* | 9 | 235 | 22 | GTF2E2 | general transcription factor IIE subunit 2 | Nucleus | transcription regulator | |
| chr17 | *MRPL45P2* | 9 | 282 | 22 | MRPL45P2 | mitochondrial ribosomal protein L45 pseudogene 2 | Other | other | |
| chr1 | *ELAVL4* | 10 | 189 | 22 | ELAVL4 | ELAV like RNA binding protein 4 | Cytoplasm | translation regulator | |
| chr15 | *CIB2* | 10 | 467 | 22 | CIB2 | calcium and integrin binding family member 2 | Cytoplasm | kinase | |
| chr8 | *RP11-175E9.1* | 10 | 242 | 22 | | | | | |
| chr17 | *CD300LG* | 9 | 416 | 22 | CD300LG | CD300 molecule like family | Plasma Membrane | other | |
| chr2 | *UBXN2A* | 8 | 194 | 22 | UBXN2A | UBX domain protein 2A | Other | other | |
| chr19 | *PGLYRP2* | 10 | 285 | 22 | PGLYRP2 | peptidoglycan recognition protein | Plasma Membrane | transmembrane receptor | |
| chr8 | *RP11-1101K5.1* | 11 | 234 | 22 | | | | | |
| chr10 | *HKDC1* | 11 | 589 | 22 | HKDC1 | hexokinase domain containing 1 | Cytoplasm | kinase | |
| chr11 | *PGM2L1* | 10 | 286 | 22 | PGM2L1 | phosphoglucomutase 2 like 1 | Cytoplasm | enzyme | |
| chr11 | *TMPRSS13* | 9 | 431 | 22 | TMPRSS13 | transmembrane serine protease | Other | peptidase | |
| chr17 | *KIF19* | 10 | 454 | 22 | KIF19 | kinesin family member 19 | Extracellular Space | enzyme | |
| chr12 | *MORN3* | 9 | 346 | 22 | MORN3 | MORN repeat containing 3 | Nucleus | other | |
| chr1 | *RP4-543J13.1* | 11 | 289 | 22 | | | | | |
| chr19 | *MAG* | 10 | 451 | 22 | MAG | myelin associated glycoprotein | Plasma Membrane | other | |
| chr3 | *LEKR1* | 10 | 300 | 22 | LEKR1 | leucine, glutamate and lysine rich | Other | other | |
| chr8 | *SLC7A2* | 10 | 238 | 22 | SLC7A2 | solute carrier family 7 member 2 | Plasma Membrane | transporter | |
| chr19 | *EIF3K* | 9 | 328 | 22 | EIF3K | eukaryotic translation initiation factor 3 subunit K | Cytoplasm | translation regulator | |
| chr12 | *NT5DC3* | 10 | 255 | 22 | NT5DC3 | 5'-nucleotidase domain | Cytoplasm | other | |
| chr2 | *COL6A3* | 8 | 297 | 22 | COL6A3 | collagen type VI alpha 3 chain | Extracellular Space | other | collagenase clostridium |
| chr10 | *USP6NL* | 8 | 215 | 22 | USP6NL | USP6 N-terminal like | Plasma Membrane | other | |
| chr6 | *SCUBE3* | 10 | 352 | 22 | SCUBE3 | signal peptide, CUB domain and EGF like domain containing 3 | Plasma Membrane | other | |
| chr1 | *DTL* | 8 | 309 | 22 | DTL | denticleless E3 ubiquitin protein ligase homolog | Nucleus | other | |
| chr11 | *SUV420H1* | 9 | 244 | 22 | KMT5B | lysine methyltransferase 5B | Nucleus | enzyme | |
| chr7 | *GSAP* | 11 | 229 | 22 | GSAP | gamma-secretase activating | Cytoplasm | peptidase | |
| chr2 | *SLC9A2* | 9 | 253 | 22 | SLC9A2 | solute carrier family 9 member A2 | Plasma Membrane | transporter | |
| chr7 | *RP11-467H10.2* | 9 | 366 | 22 | | | | | |
| chr11 | *EXPH5* | 10 | 176 | 22 | EXPH5 | exophilin 5 | Cytoplasm | other | |
| chr11 | *PAMR1* | 11 | 370 | 22 | PAMR1 | peptidase domain containing associated with muscle regeneration 1 | Extracellular Space | peptidase | |
| chr11 | *RP11-286N22.8* | 11 | 390 | 22 | | | | | |
| chr7 | *EPHA1-AS1* | 10 | 235 | 22 | EPHA1-AS1 | EPHA1 antisense RNA 1 | Other | other | |
| chr6 | *SYCP2L* | 8 | 207 | 22 | SYCP2L | synaptonemal complex protein 2 | Nucleus | other | |
| chr5 | *CTD-2503O16.4* | 10 | 193 | 22 | | | | | |
| chr17 | *LINC00671* | 9 | 279 | 22 | LINC00671 | long intergenic non-protein coding RNA 671 | Other | other | |
| chr7 | *ZNF775* | 10 | 470 | 22 | ZNF775 | zinc finger protein 775 | Other | other | |
| chrX | *ELF4* | 9 | 326 | 22 | ELF4 | E74 like ETS transcription factor | Nucleus | transcription regulator | |
| chr5 | *NKD2* | 12 | 1062 | 22 | NKD2 | NKD inhibitor of WNT signaling pathway 2 | Nucleus | other | |
| chr12 | *GLT8D2* | 9 | 286 | 22 | GLT8D2 | glycosyltransferase 8 domain containing 2 | Other | enzyme | |
| chr10 | *RNLS* | 9 | 226 | 22 | RNLS | renalase, FAD dependent amine oxidase | Extracellular Space | enzyme | |
| chr22 | *CDC45* | 8 | 271 | 22 | CDC45 | cell division cycle 45 | Nucleus | other | |
| chr3 | *CCDC13* | 9 | 397 | 22 | CCDC13 | coiled-coil domain containing 13 | Cytoplasm | other | |
| chr1 | *RP4-660H19.1* | 9 | 265 | 22 | | | | | |
| chr10 | *CPN1* | 10 | 246 | 22 | CPN1 | carboxypeptidase N subunit 1 | Extracellular Space | peptidase | |
| chr3 | *LZTFL1* | 9 | 224 | 22 | LZTFL1 | leucine zipper transcription factor like 1 | Cytoplasm | other | |
| chr17 | *MAP2K4* | 8 | 206 | 22 | MAP2K4 | mitogen-activated protein kinase kinase 4 | Cytoplasm | kinase | cetuximab/encorafenib, binimetinib/encorafenib, encorafenib, binimetinib/cetuximab/encorafenib, binimetinib/encorafen |
| chr18 | *ENOSF1* | 9 | 290 | 22 | ENOSF1 | enolase superfamily member 1 | Other | enzyme | |
| chr16 | *PRDM7* | 9 | 273 | 22 | PRDM7 | PR/SET domain 7 | Nucleus | other | |
| chr2 | *ACTR3* | 8 | 273 | 22 | ACTR3 | actin related protein 3 | Plasma Membrane | other | |
| chr10 | *PAPSS2* | 9 | 245 | 22 | PAPSS2 | 3'-phosphoadenosine 5'-phosphosulfate synthase 2 | Cytoplasm | enzyme | |
| chr17 | *MAPT-AS1* | 9 | 226 | 22 | MAPT-AS1 | MAPT antisense RNA 1 | Other | other | |
| chr10 | *RHOBTB1* | 11 | 321 | 22 | RHOBTB1 | Rho related BTB domain | Other | enzyme | |
| chr12 | *FBXO21* | 10 | 314 | 22 | FBXO21 | F-box protein 21 | Extracellular Space | enzyme | |
| chr4 | *MAEA* | 10 | 496 | 22 | MAEA | macrophage erythroblast attacher | Plasma Membrane | other | |
| chr6 | *RP1-230L10.1* | 8 | 281 | 22 | | | | | |
| chr12 | *AAAS* | 10 | 496 | 22 | AAAS | aladin WD repeat nucleoporin | Nucleus | other | |
| chr7 | *CCDC126* | 9 | 238 | 22 | CCDC126 | coiled-coil domain containing 126 | Other | other | |
| chr19 | *CEACAM21* | 10 | 339 | 22 | CEACAM21 | CEA cell adhesion molecule 21 | Other | other | |
| chr12 | *SOAT2* | 10 | 557 | 22 | SOAT2 | sterol O-acyltransferase 2 | Cytoplasm | enzyme | |

| | | | | | | | | | |
|---|---|---|---|---|---|---|---|---|---|
| chr4 | *ENPP6* | 9 | 323 | 22 | ENPP6 | ectonucleotide pyrophosphatase/phosphodiesterase 6 | Cytoplasm | enzyme | |
| chr10 | *SFXN4* | 9 | 295 | 22 | SFXN4 | sideroflexin 4 | Cytoplasm | transporter | |
| chr10 | *NPFFR1* | 11 | 387 | 22 | NPFFR1 | neuropeptide FF receptor 1 | Plasma Membrane | G-protein coupled recepto | |
| chr15 | *TUBGCP5* | 10 | 281 | 22 | TUBGCP5 | tubulin gamma complex associated protein 5 | Cytoplasm | other | |
| chr1 | *TEX35* | 9 | 573 | 22 | TEX35 | testis expressed 35 | Nucleus | other | |
| chr19 | *ZNF813* | 10 | 260 | 22 | ZNF813 | zinc finger protein 813 | Other | other | |
| chr1 | *ZNF496* | 10 | 353 | 22 | ZNF496 | zinc finger protein 496 | Nucleus | transcription regulator | |
| chr6 | *GUSBP2* | 9 | 329 | 22 | GUSBP2 | GUSB pseudogene 2 | Other | other | |
| chr12 | *RP11-370I10.2* | 9 | 259 | 22 | | | | | |
| chr3 | *SLC22A14* | 9 | 357 | 22 | SLC22A14 | solute carrier family 22 member | Plasma Membrane | transporter | |
| chr1 | *MRPL37* | 10 | 347 | 22 | MRPL37 | mitochondrial ribosomal protein | Cytoplasm | enzyme | |
| chr11 | *MTL5* | 10 | 291 | 22 | TESMIN | testis expressed metallothionein like protein | Cytoplasm | other | |
| chr12 | *PTGES3* | 9 | 245 | 22 | PTGES3 | prostaglandin E synthase 3 | Cytoplasm | enzyme | |
| chr15 | *RP11-798K3.3* | 9 | 384 | 22 | | | | | |
| chr1 | *ATF3* | 11 | 306 | 22 | ATF3 | activating transcription factor 3 | Nucleus | transcription regulator | |
| chr1 | *DCAF8* | 9 | 296 | 22 | DCAF8 | DDB1 and CUL4 associated | Nucleus | other | |
| chr2 | *RGPD5* | 8 | 296 | 22 | PD4 (includes oth | RANBP2 like and GRIP domain containing 5 | Nucleus | enzyme | |
| chr7 | *AC073343.13* | 8 | 243 | 22 | | | | | |
| chr5 | *RP11-72L22.1* | 10 | 199 | 22 | | | | | |
| chr3 | *RP11-457K10.1* | 10 | 284 | 22 | | | | | |
| chr10 | *HPS1* | 10 | 473 | 22 | HPS1 | HPS1 biogenesis of lysosomal organelles complex 3 subunit 1 | Cytoplasm | other | |
| chr15 | *TGM7* | 9 | 315 | 22 | TGM7 | transglutaminase 7 | Other | enzyme | |
| chr9 | *ST6GALNAC6* | 10 | 509 | 22 | ST6GALNAC6 | ST6 N-acetylgalactosaminide alpha-2,6-sialyltransferase 6 | Cytoplasm | enzyme | |
| chr12 | *RP11-551L14.1* | 8 | 265 | 22 | | | | | |
| chr1 | *BMP8A* | 11 | 500 | 22 | BMP8A | bone morphogenetic protein 8a | Extracellular Space | cytokine | |
| chr8 | *VPS13B* | 9 | 238 | 22 | VPS13B | vacuolar protein sorting 13 homolog B | Nucleus | transporter | |
| chr21 | *AP000320.7* | 8 | 272 | 22 | | | | | |
| chr2 | *ATG4B* | 9 | 372 | 22 | ATG4B | autophagy related 4B cysteine peptidase | Cytoplasm | peptidase | |
| chr12 | *RARG* | 10 | 718 | 22 | RARG | retinoic acid receptor gamma | Nucleus | d-dependent nuclear rec | 6-mercaptopurine/methotrexate/tretinoin, daunorubicin/tretinoin, alitretinoin, cytarabine/daunorubicin/tretinoin, acitretin, AGN 190299, clindamycin phosphate/tretinoin, palovarotene, arsenic trioxide/daunorubicin/tretinoin, idarubicin/mitoxantrone/tretinoin, etretinate, fenretinide, idarubicin/tretinoin, tretinoin, tazarotene, arsenic trioxide/gemtuzumab ozogamicin/tretinoin, halobetasol [illegible] |
| chr14 | *PPP2R5E* | 8 | 220 | 22 | PPP2R5E | protein phosphatase 2 regulatory subunit B'epsilon | Cytoplasm | phosphatase | |
| chr11 | *ANKRD42* | 9 | 256 | 22 | ANKRD42 | ankyrin repeat domain 42 | Nucleus | transcription regulator | |
| chr11 | *DDB2* | 10 | 580 | 22 | DDB2 | damage specific DNA binding protein 2 | Nucleus | other | |
| chr2 | *EIF5B* | 8 | 232 | 22 | EIF5B | eukaryotic translation initiation factor 5B | Cytoplasm | translation regulator | |
| chr1 | *PARK7* | 9 | 354 | 22 | PARK7 | Parkinsonism associated | Nucleus | enzyme | |
| chr13 | *ITGBL1* | 9 | 221 | 22 | ITGBL1 | integrin subunit beta like 1 | Extracellular Space | other | |
| chr12 | *RP11-392P7.6* | 10 | 212 | 22 | | | | | |
| chrX | *OPHN1* | 8 | 225 | 22 | OPHN1 | oligophrenin 1 | Cytoplasm | other | |
| chr22 | *TTLL8* | 10 | 430 | 22 | TTLL8 | tubulin tyrosine ligase like 8 | Extracellular Space | enzyme | |
| chr16 | *CLN3* | 10 | 345 | 22 | CLN3 | CLN3 lysosomal/endosomal transmembrane protein, battenin | Cytoplasm | other | |
| chr10 | *ITIH5* | 10 | 271 | 22 | ITIH5 | inter-alpha-trypsin inhibitor heavy chain 5 | Plasma Membrane | other | |
| chr1 | *IFNLR1* | 9 | 314 | 22 | IFNLR1 | interferon lambda receptor 1 | Plasma Membrane | transmembrane recepto | |
| chr15 | *ISLR2* | 10 | 287 | 22 | ISLR2 | immunoglobulin superfamily containing leucine rich repeat 2 | Plasma Membrane | other | |
| chr8 | *NKAIN3* | 8 | 193 | 22 | NKAIN3 | sodium/potassium transporting ATPase interacting 3 | Other | other | |
| chr3 | *ZNF717* | 9 | 289 | 22 | ZNF717 | zinc finger protein 717 | Nucleus | transcription regulator | |
| chr1 | *BSDC1* | 9 | 296 | 22 | BSDC1 | BSD domain containing 1 | Other | other | |
| chr14 | *SYNE2* | 8 | 188 | 22 | SYNE2 | spectrin repeat containing nuclear envelope protein 2 | Nucleus | other | |
| chr16 | *PLA2G10* | 10 | 298 | 22 | PLA2G10 | phospholipase A2 group X | Cytoplasm | enzyme | varespladib, varespladib methyl |
| chr1 | *GPR89A* | 8 | 276 | 22 | GPR89A/GPR89E | G protein-coupled receptor 89B | Cytoplasm | ion channel | |

| | | | | | | | | | |
|---|---|---|---|---|---|---|---|---|---|
| chr16 | *SLX4* | 10 | 400 | 22 | SLX4 | SLX4 structure-specific endonuclease subunit | Nucleus | other | |
| chr22 | *COMT* | 9 | 340 | 22 | COMT | catechol-O-methyltransferase | Cytoplasm | enzyme | entacapone, carbidopa/entacapone/levodopa, carbidopa/levodopa/tolcapone, tolcapone, BIA-3-202 |
| chr16 | *RP11-368L12.1* | 9 | 212 | 22 | | | | | |
| chr2 | *PAPOLG* | 9 | 187 | 22 | PAPOLG | poly(A) polymerase gamma | Nucleus | enzyme | |
| chr4 | *RP11-665G4.1* | 9 | 217 | 22 | | | | | |
| chr6 | *MTCH1* | 8 | 367 | 22 | MTCH1 | mitochondrial carrier 1 | Cytoplasm | other | |
| chr1 | *RAB3GAP2* | 9 | 300 | 22 | RAB3GAP2 | RAB3 GTPase activating non-catalytic protein subunit 2 | Cytoplasm | enzyme | |
| chr19 | *BLVRB* | 10 | 373 | 22 | BLVRB | biliverdin reductase B | Cytoplasm | enzyme | |
| chr21 | *SIM2* | 9 | 406 | 22 | SIM2 | SIM bHLH transcription factor 2 | Nucleus | transcription regulator | |
| chr2 | *SAG* | 8 | 275 | 22 | SAG | S-antigen visual arrestin | Cytoplasm | other | |
| chr5 | *SRD5A1* | 10 | 683 | 22 | SRD5A1 | steroid 5 alpha-reductase 1 | Cytoplasm | enzyme | estradiol/levonorgestrel, ethinyl estradiol/levonorgestrel, levonorgestrel, dutasteride/tamsulosin, dutasteride, finasteride, 5-alpha-reductase inhibitor |
| chr12 | *RP11-70F11.8* | 10 | 479 | 22 | | | | | |
| chr1 | *TXLNA* | 9 | 322 | 22 | TXLNA | taxilin alpha | Extracellular Space | cytokine | |
| chr3 | *PDCD6IP* | 8 | 208 | 22 | PDCD6IP | programmed cell death 6 interacting protein | Cytoplasm | other | |
| chr5 | *ARAP3* | 9 | 375 | 22 | ARAP3 | ArfGAP with RhoGAP domain, ankyrin repeat and PH domain 3 | Cytoplasm | other | |
| chr2 | *SH2D6* | 9 | 400 | 22 | SH2D6 | SH2 domain containing 6 | Other | other | |
| chr12 | *RP11-22B23.1* | 9 | 357 | 22 | | | | | |
| chr5 | *PAM* | 8 | 230 | 22 | PAM | peptidylglycine alpha-amidating monooxygenase | Plasma Membrane | enzyme | |
| chr11 | *CDCA5* | 9 | 352 | 22 | CDCA5 | cell division cycle associated 5 | Cytoplasm | other | |
| chr19 | *TMPRSS9* | 9 | 323 | 22 | TMPRSS9 | transmembrane serine protease 9 | Plasma Membrane | peptidase | |
| chr22 | *TEF* | 9 | 286 | 22 | TEF | TEF transcription factor, PAR bZIP family member | Nucleus | transcription regulator | |
| chr15 | *RAD51* | 8 | 222 | 22 | RAD51 | RAD51 recombinase | Nucleus | enzyme | CYT-0851 |
| chr8 | *NDUFAF6* | 8 | 204 | 22 | NDUFAF6 | NADH:ubiquinone oxidoreductase complex | Cytoplasm | other | |
| chr5 | *SRFBP1* | 8 | 247 | 22 | SRFBP1 | serum response factor binding protein 1 | Nucleus | other | |
| chr5 | *SMN1* | 9 | 254 | 22 | SMN1/SMN2 | survival of motor neuron 1, | Nucleus | other | nusinersen |
| chr11 | *CCDC83* | 11 | 381 | 22 | CCDC83 | coiled-coil domain containing 83 | Other | other | |
| chr19 | *CACNG7* | 10 | 289 | 22 | CACNG7 | calcium voltage-gated channel auxiliary subunit gamma 7 | Plasma Membrane | ion channel | |
| chr7 | *LINC01004* | 8 | 211 | 22 | LINC01004 | long intergenic non-protein coding RNA 1004 | Other | other | |
| chr10 | *SHOC2* | 8 | 251 | 22 | SHOC2 | SHOC2 leucine rich repeat scaffold protein | Cytoplasm | other | |
| chr7 | *BCL7B* | 9 | 304 | 22 | BCL7B | BAF chromatin remodeling complex subunit BCL7B | Nucleus | other | |
| chr10 | *OIT3* | 10 | 334 | 22 | OIT3 | oncoprotein induced transcript 3 | Nucleus | other | |
| chr5 | *SLC1A3* | 10 | 654 | 22 | SLC1A3 | solute carrier family 1 member 3 | Plasma Membrane | transporter | riluzole |
| chr17 | *CLTC* | 8 | 240 | 22 | CLTC | clathrin heavy chain | Plasma Membrane | other | |
| chr12 | *DDX54* | 10 | 423 | 22 | DDX54 | DEAD-box helicase 54 | Nucleus | transcription regulator | |
| chr1 | *RCAN3* | 9 | 238 | 22 | RCAN3 | RCAN family member 3 | Other | other | |
| chr5 | *SH3TC2* | 9 | 239 | 22 | SH3TC2 | SH3 domain and tetratricopeptide repeats 2 | Cytoplasm | other | |
| chr12 | *SLC38A1* | 10 | 454 | 22 | SLC38A1 | solute carrier family 38 member 1 | Plasma Membrane | transporter | |
| chr22 | *WBP2NL* | 8 | 204 | 22 | WBP2NL | WBP2 N-terminal like | Cytoplasm | other | |
| chr1 | *STRIP1* | 10 | 427 | 22 | STRIP1 | striatin interacting protein 1 | Nucleus | other | |
| chr17 | *AKAP1* | 10 | 381 | 22 | AKAP1 | A-kinase anchoring protein 1 | Cytoplasm | other | |
| chr17 | *CTD-2020K17.1* | 10 | 341 | 22 | | | | | |
| chr7 | *ZNF277* | 9 | 226 | 22 | ZNF277 | zinc finger protein 277 | Nucleus | transcription regulator | |
| chr10 | *NEBL* | 9 | 203 | 22 | NEBL | nebulette | Plasma Membrane | other | |
| chr7 | *ZC3HC1* | 8 | 271 | 22 | ZC3HC1 | zinc finger C3HC-type containing | Nucleus | other | |
| chr17 | *CYB5D2* | 9 | 367 | 22 | CYB5D2 | cytochrome b5 domain containing | Extracellular Space | other | |
| chr3 | *RP11-167H9.4* | 11 | 300 | 22 | | | | | |
| chr8 | *RP11-89M16.1* | 10 | 228 | 22 | | | | | |
| chr7 | *CPA4* | 9 | 332 | 22 | CPA4 | carboxypeptidase A4 | Extracellular Space | peptidase | |
| chr11 | *STX5* | 10 | 261 | 22 | STX5 | syntaxin 5 | Cytoplasm | transporter | |
| chr11 | *C11orf85* | 9 | 254 | 22 | MAJIN | membrane anchored junction | Nucleus | other | |
| chr10 | *ZFYVE27* | 9 | 391 | 22 | ZFYVE27 | zinc finger FYVE-type containing | Plasma Membrane | other | |
| chr1 | *ATAD3B* | 11 | 364 | 22 | ATAD3B | ATPase family AAA domain containing 3B | Nucleus | other | |
| chr11 | *RBM4* | 9 | 391 | 22 | RBM4 | RNA binding motif protein 4 | Nucleus | other | |
| chr7 | *AGBL3* | 9 | 312 | 22 | AGBL3 | ATP/GTP binding protein like 3 | Other | peptidase | |

| | | | | | | | | | |
|---|---|---|---|---|---|---|---|---|---|
| chr3 | *TNK2* | 9 | 474 | 22 | TNK2 | tyrosine kinase non receptor 2 | Cytoplasm | kinase | cetuximab/vemurafenib, irinotecan/panitumumab/vemurafenib, trametinib/vemurafenib, dabrafenib/trametinib/vemurafenib, vemurafenib, crizotinib/vemurafenib, ipilimumab/vemurafenib, erlotinib/osimertinib, cobimetinib/vemurafenib, afatinib/osimertinib |
| chr10 | *H2AFY2* | 10 | 402 | 22 | MACROH2A2 | macroH2A.2 histone | Nucleus | other | |
| chr5 | *RP11-1023L17.1* | 10 | 1404 | 22 | | | | | |
| chr2 | *FOXN2* | 9 | 215 | 22 | FOXN2 | forkhead box N2 | Nucleus | transcription regulator | |
| chr5 | *LIFR-AS1* | 11 | 776 | 22 | LIFR-AS1 | LIFR antisense RNA 1 | Other | other | |
| chr1 | *PIGR* | 10 | 451 | 22 | PIGR | polymeric immunoglobulin | Plasma Membrane | transporter | |
| chr1 | *KIF2C* | 9 | 275 | 22 | KIF2C | kinesin family member 2C | Nucleus | other | |
| chr19 | *PPP5D1* | 9 | 298 | 22 | PPP5D1 | PPP5 tetratricopeptide repeat domain containing 1 | Other | other | |
| chr10 | *FAM21C* | 9 | 343 | 22 | ASHC2A/WASHC | WASH complex subunit 2A | Cytoplasm | other | |
| chr15 | *CSK* | 9 | 350 | 22 | CSK | C-terminal Src kinase | Cytoplasm | kinase | bosutinib |
| chr11 | *ZBTB44* | 9 | 262 | 22 | ZBTB44 | zinc finger and BTB domain containing 44 | Other | other | |
| chr11 | *MAP4K2* | 9 | 417 | 22 | MAP4K2 | mitogen-activated protein kinase kinase kinase kinase 2 | Cytoplasm | kinase | |
| chr1 | *C1orf167* | 11 | 485 | 22 | C1orf167 | chromosome 1 open reading frame 167 | Other | other | |
| chr20 | *RP11-290F20.1* | 9 | 591 | 22 | | | | | |
| chr18 | *MIB1* | 7 | 200 | 22 | MIB1 | mindbomb E3 ubiquitin protein ligase 1 | Cytoplasm | enzyme | |
| chr1 | *DCST1* | 9 | 522 | 22 | DCST1 | DC-STAMP domain containing 1 | Extracellular Space | enzyme | |
| chr4 | *F11-AS1* | 8 | 285 | 22 | F11-AS1 | F11 antisense RNA 1 | Other | other | |
| chr12 | *NUP107* | 9 | 198 | 22 | NUP107 | nucleoporin 107 | Nucleus | other | |
| chr2 | *CAPG* | 10 | 385 | 22 | CAPG | capping actin protein, gelsolin like | Nucleus | other | |
| chr1 | *AGO4* | 8 | 217 | 22 | AGO4 | argonaute RISC component 4 | Cytoplasm | translation regulator | |
| chr1 | *RP11-815M8.1* | 10 | 265 | 22 | | | | | |
| chr5 | *PHAX* | 8 | 228 | 22 | PHAX | phosphorylated adaptor for RNA export | Cytoplasm | other | |
| chr7 | *TMEM248* | 9 | 271 | 22 | TMEM248 | transmembrane protein 248 | Other | other | |
| chr22 | *TAB1* | 8 | 343 | 22 | TAB1 | TGF-beta activated kinase 1 (MAP3K7) binding protein 1 | Cytoplasm | enzyme | |
| chr8 | *GSR* | 10 | 190 | 22 | GSR | glutathione-disulfide reductase | Cytoplasm | enzyme | carmustine/prednisone, carmustine |
| chr11 | *CAPRIN1* | 9 | 332 | 22 | CAPRIN1 | cell cycle associated protein 1 | Plasma Membrane | other | |
| chr2 | *NBAS* | 8 | 224 | 22 | NBAS | NBAS subunit of NRZ tethering complex | Nucleus | other | |
| chr22 | *MCM5* | 8 | 340 | 22 | MCM5 | minichromosome maintenance complex component 5 | Nucleus | enzyme | |
| chr17 | *COX10* | 9 | 219 | 22 | COX10 | cytochrome c oxidase assembly factor heme A:farnesyltransferase COX10 | Cytoplasm | enzyme | |
| chr11 | *TSSC2* | 10 | 590 | 21 | TSSC2 | tumor suppressing subtransferable candidate 2 | Other | other | |
| chr17 | *SLC25A19* | 9 | 357 | 21 | SLC25A19 | solute carrier family 25 member | Cytoplasm | transporter | |
| chr1 | *EGLN1* | 8 | 232 | 21 | EGLN1 | egl-9 family hypoxia inducible | Cytoplasm | enzyme | |
| chr7 | *RHEB* | 8 | 261 | 21 | RHEB | Ras homolog, mTORC1 binding | Cytoplasm | enzyme | |
| chr5 | *BTNL8* | 9 | 358 | 21 | BTNL8 | butyrophilin like 8 | Plasma Membrane | other | |
| chr19 | *GDF15* | 9 | 371 | 21 | GDF15 | growth differentiation factor 15 | Extracellular Space | growth factor | |
| chr1 | *PTGER3* | 9 | 204 | 21 | PTGER3 | prostaglandin E receptor 3 | Plasma Membrane | i-protein coupled recepto | prostaglandin E2, bimatoprost/timolol, bimatoprost, diclofenac/misoprostol, misoprostol, prostaglandin E1 |
| chr8 | *CTSB* | 10 | 366 | 21 | CTSB | cathepsin B | Cytoplasm | peptidase | |
| chr3 | *ACVR2B* | 10 | 301 | 21 | ACVR2B | activin A receptor type 2B | Plasma Membrane | kinase | |
| chr7 | *TAX1BP1* | 8 | 224 | 21 | TAX1BP1 | Tax1 binding protein 1 | Cytoplasm | other | |
| chr19 | *CIC* | 11 | 479 | 21 | CIC | capicua transcriptional repressor | Nucleus | transcription regulator | |
| chr16 | *FBXL19* | 10 | 383 | 21 | FBXL19 | F-box and leucine rich repeat protein 19 | Cytoplasm | enzyme | |
| chr2 | *AC104809.3* | 10 | 538 | 21 | | | | | |
| chr9 | *FAM78A* | 10 | 512 | 21 | FAM78A | family with sequence similarity 78 member A | Other | other | |
| chr12 | *A2ML1* | 7 | 168 | 21 | A2ML1 | alpha-2-macroglobulin like 1 | Cytoplasm | other | |
| chr17 | *TMEM11* | 10 | 381 | 21 | TMEM11 | transmembrane protein 11 | Plasma Membrane | i-protein coupled recepto | |
| chr1 | *CROCCP3* | 10 | 362 | 21 | CROCCP3 | CROCC pseudogene 3 | Other | other | |
| chr16 | *ACSM3* | 8 | 282 | 21 | ACSM3 | acyl-CoA synthetase medium chain family member 3 | Cytoplasm | enzyme | |
| chr10 | *DPCD* | 10 | 319 | 21 | DPCD | deleted in primary ciliary dyskinesia homolog (mouse) | Nucleus | other | |
| chr11 | *PGAP2* | 10 | 611 | 21 | PGAP2 | post-GPI attachment to proteins 2 | Other | other | |
| chr8 | *SNX31* | 8 | 210 | 21 | SNX31 | sorting nexin 31 | Other | other | |
| chr7 | *ZNF777* | 8 | 263 | 21 | ZNF777 | zinc finger protein 777 | Other | other | |

| | | | | | | | | | |
|---|---|---|---|---|---|---|---|---|---|
| chr15 | *SNRPN* | 9 | 193 | 21 | SNRPN | small nuclear ribonucleoprotein polypeptide N | Nucleus | other | |
| chr20 | *SDC4* | 10 | 375 | 21 | SDC4 | syndecan 4 | Plasma Membrane | other | |
| chr20 | *GDAP1L1* | 10 | 418 | 21 | GDAP1L1 | ganglioside induced differentiation associated protein | Other | other | |
| chr17 | *CTC-501O10.1* | 9 | 351 | 21 | | | | | |
| chr10 | *GLRX3* | 8 | 218 | 21 | GLRX3 | glutaredoxin 3 | Cytoplasm | enzyme | |
| chr3 | *YEATS2* | 7 | 196 | 21 | YEATS2 | YEATS domain containing 2 | Nucleus | transcription regulator | |
| chr12 | *RNF34* | 8 | 245 | 21 | RNF34 | ring finger protein 34 | Cytoplasm | enzyme | |
| chr19 | *ZNF627* | 9 | 256 | 21 | ZNF627 | zinc finger protein 627 | Other | other | |
| chr22 | *GGA1* | 9 | 408 | 21 | GGA1 | golgi associated, gamma adaptin ear containing, ARF binding | Cytoplasm | transporter | |
| chr18 | *MAPRE2* | 8 | 188 | 21 | MAPRE2 | microtubule associated protein RP/EB family member 2 | Cytoplasm | other | |
| chr19 | *PRX* | 9 | 374 | 21 | PRX | periaxin | Nucleus | other | |
| chr12 | *RAD52* | 8 | 212 | 21 | RAD52 | RAD52 homolog, DNA repair | Nucleus | other | |
| chr1 | *CD244* | 10 | 288 | 21 | CD244 | CD244 molecule | Plasma Membrane | transmembrane receptor | |
| chr3 | *TPRG1* | 8 | 205 | 21 | TPRG1 | tumor protein p63 regulated 1 | Cytoplasm | other | |
| chr7 | *SEMA3D* | 11 | 346 | 21 | SEMA3D | semaphorin 3D | Extracellular Space | other | |
| chr14 | *NPAS3* | 8 | 193 | 21 | NPAS3 | neuronal PAS domain protein 3 | Nucleus | transcription regulator | |
| chr14 | *RPS6KA5* | 8 | 172 | 21 | RPS6KA5 | ribosomal protein S6 kinase A5 | Nucleus | kinase | |
| chr5 | *PCDH1* | 10 | 444 | 21 | PCDH1 | protocadherin 1 | Plasma Membrane | other | |
| chr7 | *PLEKHA8* | 10 | 278 | 21 | PLEKHA8 | pleckstrin homology domain containing A8 | Extracellular Space | transporter | |
| chr10 | *WDR96* | 10 | 192 | 21 | CFAP43 | cilia and flagella associated | Cytoplasm | other | |
| chr17 | *LINC00670* | 8 | 182 | 21 | LINC00670 | long intergenic non-protein coding RNA 670 | Other | other | |
| chr19 | *PIK3R2* | 10 | 437 | 21 | PIK3R2 | phosphoinositide-3-kinase regulatory subunit 2 | Cytoplasm | kinase | |
| chr15 | *SNAP23* | 8 | 228 | 21 | SNAP23 | synaptosome associated protein | Plasma Membrane | transporter | |
| chr4 | *FAM114A1* | 9 | 198 | 21 | FAM114A1 | family with sequence similarity 114 member A1 | Extracellular Space | other | |
| chr18 | *SNRPD1* | 8 | 247 | 21 | SNRPD1 | small nuclear ribonucleoprotein D1 polypeptide | Nucleus | other | |
| chr8 | *SLC25A37* | 9 | 306 | 21 | SLC25A37 | solute carrier family 25 member | Cytoplasm | transporter | |
| chr7 | *ZPBP* | 10 | 257 | 21 | ZPBP | zona pellucida binding protein | Nucleus | other | |
| chr12 | *NR2C1* | 7 | 191 | 21 | NR2C1 | nuclear receptor subfamily 2 group C member 1 | Nucleus | transcription regulator | |
| chr2 | *TLK1* | 7 | 189 | 21 | TLK1 | tousled like kinase 1 | Nucleus | kinase | |
| chr4 | *AFF1* | 8 | 183 | 21 | AFF1 | AF4/FMR2 family member 1 | Nucleus | transcription regulator | |
| chr17 | *P2RX1* | 10 | 421 | 21 | P2RX1 | purinergic receptor P2X 1 | Plasma Membrane | ion channel | |
| chr1 | *PSEN2* | 10 | 493 | 21 | PSEN2 | presenilin 2 | Cytoplasm | peptidase | tarenflurbil |
| chr16 | *IRF8* | 9 | 351 | 21 | IRF8 | interferon regulatory factor 8 | Nucleus | transcription regulator | |
| chr1 | *RBP7* | 10 | 201 | 21 | RBP7 | retinol binding protein 7 | Cytoplasm | other | |
| chr7 | *GNAI1* | 8 | 242 | 21 | GNAI1 | G protein subunit alpha i1 | Plasma Membrane | enzyme | |
| chr1 | *COL9A2* | 10 | 416 | 21 | COL9A2 | collagen type IX alpha 2 chain | Extracellular Space | other | collagenase clostridium |
| chr5 | *CTC-347C20.2* | 10 | 220 | 21 | | | | | |
| chr7 | *MYL10* | 9 | 446 | 21 | MYL10 | myosin light chain 10 | Cytoplasm | other | |
| chr19 | *LMNB2* | 10 | 478 | 21 | LMNB2 | lamin B2 | Nucleus | other | |
| chr10 | *EIF3A* | 9 | 250 | 21 | EIF3A | eukaryotic translation initiation factor 3 subunit A | Cytoplasm | other | |
| chr3 | *LINC00693* | 10 | 191 | 21 | LINC00693 | long intergenic non-protein coding RNA 693 | Other | other | |
| chr16 | *CDH5* | 9 | 429 | 21 | CDH5 | cadherin 5 | Plasma Membrane | other | |
| chr19 | *KANK2* | 9 | 350 | 21 | KANK2 | KN motif and ankyrin repeat domains 2 | Nucleus | transcription regulator | |
| chr5 | *CDK7* | 9 | 216 | 21 | CDK7 | cyclin dependent kinase 7 | Nucleus | kinase | SB-1317, BMS-387032, roscovitine, alvocidib, CT7001, SY-1365 |
| chr5 | *CDC25C* | 8 | 259 | 21 | CDC25C | cell division cycle 25C | Nucleus | phosphatase | |
| chr2 | *ACTR2* | 8 | 194 | 21 | ACTR2 | actin related protein 2 | Plasma Membrane | other | |
| chr1 | *RP5-857K21.4* | 9 | 299 | 21 | | | | | |
| chr3 | *ARHGEF26* | 9 | 236 | 21 | ARHGEF26 | Rho guanine nucleotide exchange factor 26 | Plasma Membrane | other | |
| chr8 | *CPQ* | 8 | 223 | 21 | CPQ | carboxypeptidase Q | Extracellular Space | peptidase | |
| chr22 | *PDGFB* | 9 | 420 | 21 | PDGFB | platelet derived growth factor subunit B | Extracellular Space | growth factor | pegpleranib |
| chr19 | *C19orf57* | 10 | 378 | 21 | C19orf57 | chromosome 19 open reading frame 57 | Other | other | |
| chr17 | *MMP28* | 10 | 268 | 21 | MMP28 | matrix metallopeptidase 28 | Extracellular Space | peptidase | marimastat |
| chr16 | *AMFR* | 8 | 306 | 21 | AMFR | autocrine motility factor receptor | Plasma Membrane | transmembrane receptor | |
| chr1 | *CLSPN* | 8 | 216 | 21 | CLSPN | claspin | Nucleus | other | |

| | | | | | | | | | |
|---|---|---|---|---|---|---|---|---|---|
| chr11 | *PGR* | 11 | 494 | 21 | PGR | progesterone receptor | Nucleus | d-dependent nuclear rec | levonorgestrel, dydrogesterone, norethindrone, ethinyl estradiol/norgestrel, ulipristal acetate, drospirenone, ethinyl estradiol/norgestimate, megestrol acetate/tamoxifen, asoprisnil, beta-estradiol/progesterone, ulipristal, ethynodiol diacetate, dienogest, danazol, estradiol/norethindrone acetate, fluorine F 18 fluoro furanyl norprogesterone, norelgestromin, desogestrel, leuprolide acetate/norethindrone acetate |
| chr8 | *ZFP41* | 10 | 482 | 21 | ZFP41 | ZFP41 zinc finger protein | Nucleus | transcription regulator | |
| chr1 | *ORC1* | 10 | 211 | 21 | ORC1 | origin recognition complex | Nucleus | other | |
| chr12 | *NUAK1* | 10 | 285 | 21 | NUAK1 | NUAK family kinase 1 | Nucleus | kinase | |
| chr9 | *GPR144* | 11 | 559 | 21 | ADGRD2 | adhesion G protein-coupled receptor D2 | Plasma Membrane | G-protein coupled recepto | |
| chr13 | *TEX29* | 10 | 491 | 21 | TEX29 | testis expressed 29 | Other | other | |
| chr16 | *FAM65A* | 9 | 341 | 21 | RIPOR1 | RHO family interacting cell polarization regulator 1 | Cytoplasm | other | |
| chr7 | *C7orf57* | 9 | 357 | 21 | C7orf57 | chromosome 7 open reading frame 57 | Other | other | |
| chr4 | *NSG1* | 9 | 321 | 21 | NSG1 | neuronal vesicle trafficking associated 1 | Cytoplasm | other | |
| chr20 | *ADNP* | 9 | 273 | 21 | ADNP | activity dependent neuroprotector homeobox | Nucleus | transcription regulator | |
| chrX | *PDZD4* | 9 | 369 | 21 | PDZD4 | PDZ domain containing 4 | Cytoplasm | other | |
| chr3 | *ITIH3* | 9 | 373 | 21 | ITIH3 | inter-alpha-trypsin inhibitor heavy chain 3 | Extracellular Space | other | |
| chr2 | *GIGYF2* | 8 | 178 | 21 | GIGYF2 | GRB10 interacting GYF protein 2 | Cytoplasm | other | |
| chr9 | *DNAI1* | 8 | 257 | 21 | DNAI1 | dynein axonemal intermediate chain 1 | Extracellular Space | other | |
| chr5 | *ALDH7A1* | 9 | 237 | 21 | ALDH7A1 | aldehyde dehydrogenase 7 family member A1 | Cytoplasm | enzyme | |
| chr18 | *NPC1* | 8 | 234 | 21 | NPC1 | NPC intracellular cholesterol transporter 1 | Cytoplasm | transporter | |
| chr1 | *RAVER2* | 7 | 232 | 21 | RAVER2 | ribonucleoprotein, PTB binding 2 | Nucleus | other | |
| chr22 | *SYNGR1* | 8 | 412 | 21 | SYNGR1 | synaptogyrin 1 | Plasma Membrane | transporter | |
| chr12 | *ULK1* | 11 | 382 | 21 | ULK1 | unc-51 like autophagy activating kinase 1 | Cytoplasm | kinase | |
| chr18 | *CEP192* | 8 | 228 | 21 | CEP192 | centrosomal protein 192 | Cytoplasm | other | |
| chr15 | *AP4E1* | 8 | 198 | 21 | AP4E1 | adaptor related protein complex 4 subunit epsilon 1 | Cytoplasm | other | |
| chr19 | *GIPC1* | 9 | 366 | 21 | GIPC1 | GIPC PDZ domain containing family member 1 | Cytoplasm | other | |
| chr1 | *ENSA* | 9 | 331 | 21 | ENSA | endosulfine alpha | Cytoplasm | transporter | |
| chr2 | *RP11-481J13.1* | 10 | 327 | 21 | | | | | |
| chr16 | *STX1B* | 10 | 461 | 21 | STX1B | syntaxin 1B | Plasma Membrane | other | |
| chr11 | *EXT2* | 9 | 205 | 21 | EXT2 | exostosin glycosyltransferase 2 | Cytoplasm | enzyme | |
| chr8 | *C8orf34* | 7 | 190 | 21 | C8orf34 | chromosome 8 open reading frame 34 | Other | other | |
| chr14 | *NIN* | 8 | 201 | 21 | NIN | ninein | Cytoplasm | other | |
| chr14 | *ANGEL1* | 8 | 232 | 21 | ANGEL1 | angel homolog 1 | Extracellular Space | other | |
| chr15 | *SH2D7* | 9 | 434 | 21 | SH2D7 | SH2 domain containing 7 | Other | other | |
| chr8 | *GULOP* | 9 | 399 | 21 | GULOP | gulonolactone (L-) oxidase, pseudogene | Other | other | |
| chr11 | *RP11-702H23.4* | 10 | 263 | 21 | | | | | |
| chr11 | *TMPRSS4* | 9 | 239 | 21 | TMPRSS4 | transmembrane serine protease 4 | Cytoplasm | peptidase | |
| chr6 | *NQO2* | 10 | 312 | 21 | NQO2 | N-ribosyldihydronicotinamide:quinon | Cytoplasm | enzyme | caricotamide |
| chr1 | *CASQ2* | 9 | 316 | 21 | CASQ2 | calsequestrin 2 | Cytoplasm | other | |
| chr15 | *MEIS2* | 8 | 223 | 21 | MEIS2 | Meis homeobox 2 | Nucleus | transcription regulator | |
| chr3 | *MUC13* | 9 | 245 | 21 | MUC13 | mucin 13, cell surface associated | Extracellular Space | other | |
| chr8 | *PCM1* | 7 | 215 | 21 | PCM1 | pericentriolar material 1 | Cytoplasm | other | |
| chr20 | *BPIFB3* | 10 | 526 | 21 | BPIFB3 | BPI fold containing family B member 3 | Extracellular Space | other | |
| chr7 | *AC009518.4* | 10 | 247 | 21 | | | | | |
| chr7 | *ZNF890P* | 10 | 497 | 21 | ZNF890P | zinc finger protein 890, | Other | other | |
| chr19 | *KMT2B* | 10 | 514 | 21 | KMT2B | lysine methyltransferase 2B | Nucleus | transcription regulator | |
| chr17 | *TRIM65* | 9 | 397 | 21 | TRIM65 | tripartite motif containing 65 | Nucleus | other | |
| chr9 | *C9orf92* | 10 | 252 | 21 | C9orf92 | chromosome 9 open reading frame 92 | Other | other | |
| chr11 | *HNRNPKP3* | 10 | 206 | 21 | HNRNPKP3 | heterogeneous nuclear ribonucleoprotein K pseudogene | Other | other | |
| chr17 | *TBX4* | 10 | 345 | 21 | TBX4 | T-box transcription factor 4 | Nucleus | transcription regulator | |
| chr11 | *PANX1* | 9 | 489 | 21 | PANX1 | pannexin 1 | Plasma Membrane | transporter | |

| | | | | | | | | | |
|---|---|---|---|---|---|---|---|---|---|
| chr4 | *SPOCK3* | 8 | 222 | 21 | SPOCK3 | SPARC (osteonectin), cwcv and kazal like domains proteoglycan 3 | Extracellular Space | other | |
| chr17 | *SLC6A4* | 10 | 272 | 21 | SLC6A4 | solute carrier family 6 member 4 | Plasma Membrane | transporter | SLV-314, escitalopram, methamphetamine, milnacipran, amoxapine, azatadine/pseudoephedrine, trazodone, fluoxetine, phentermine/topiramate, naproxen/pseudoephedrine, loratadine/pseudoephedrine, chlorpheniramine/ibuprofen/pseudoephedrine, amitriptyline/melatonin, mazindol, clomipramine, fenfluramine/phentermine, sibutramine, protriptyline, [illegible] |
| chr3 | *TAMM41* | 8 | 220 | 21 | TAMM41 | TAM41 mitochondrial translocator assembly and maintenance | Cytoplasm | enzyme | |
| chr7 | *NCF1C* | 8 | 463 | 21 | NCF1C | neutrophil cytosolic factor 1C pseudogene | Cytoplasm | other | |
| chr11 | *NFRKB* | 9 | 389 | 21 | NFRKB | nuclear factor related to kappaB binding protein | Nucleus | transcription regulator | |
| chr20 | *OPRL1* | 11 | 1223 | 21 | OPRL1 | opioid related nociceptin receptor 1 | Plasma Membrane | G-protein coupled recepto | cebranopadol, buprenorphine/naloxone, buprenorphine, ZP120 |
| chr15 | *KIF7* | 10 | 478 | 21 | KIF7 | kinesin family member 7 | Extracellular Space | other | |
| chr18 | *RP11-476K15.1* | 9 | 229 | 21 | | | | | |
| chr16 | *HSDL1* | 8 | 369 | 21 | HSDL1 | hydroxysteroid dehydrogenase | Cytoplasm | enzyme | |
| chr17 | *LPO* | 9 | 261 | 21 | LPO | lactoperoxidase | Extracellular Space | enzyme | |
| chr7 | *C7orf26* | 10 | 477 | 21 | C7orf26 | chromosome 7 open reading frame 26 | Nucleus | other | |
| chr5 | *CTD-2231H16.1* | 10 | 1021 | 21 | | | | | |
| chr2 | *KIF3C* | 8 | 310 | 21 | KIF3C | kinesin family member 3C | Cytoplasm | other | |
| chr9 | *PHF19* | 9 | 520 | 21 | PHF19 | PHD finger protein 19 | Nucleus | other | |
| chr3 | *SDHAP1* | 8 | 341 | 21 | SDHAP1 | succinate dehydrogenase complex flavoprotein subunit A | Other | other | |
| chr11 | *BARX2* | 10 | 313 | 21 | BARX2 | BARX homeobox 2 | Nucleus | transcription regulator | |
| chr18 | *ALPK2* | 7 | 203 | 21 | ALPK2 | alpha kinase 2 | Nucleus | kinase | |
| chr7 | *SLC26A5* | 10 | 184 | 21 | SLC26A5 | solute carrier family 26 member 5 | Plasma Membrane | transporter | |
| chr22 | *DGCR5* | 9 | 384 | 21 | DGCR5 | DiGeorge syndrome critical region gene 5 | Other | other | |
| chr1 | *LAMC2* | 8 | 286 | 21 | LAMC2 | laminin subunit gamma 2 | Extracellular Space | other | |
| chr14 | *MNAT1* | 8 | 214 | 21 | MNAT1 | MNAT1 component of CDK activating kinase | Nucleus | other | |
| chr5 | *TTC37* | 9 | 221 | 21 | TTC37 | tetratricopeptide repeat domain | Nucleus | other | |
| chrX | *CXXC1P1* | 9 | 233 | 21 | CXXC1P1 | CXXC finger protein 1 | Other | other | |
| chr3 | *KIF9* | 10 | 255 | 21 | KIF9 | kinesin family member 9 | Cytoplasm | enzyme | |
| chr17 | *STAC2* | 9 | 460 | 21 | STAC2 | SH3 and cysteine rich domain 2 | Plasma Membrane | other | |
| chr1 | *C1orf86* | 11 | 575 | 21 | FAAP20 | FA core complex associated protein 20 | Nucleus | other | |
| chr11 | *RP11-460B17.3* | 9 | 312 | 21 | | | | | |
| chr16 | *RABEP2* | 10 | 293 | 21 | RABEP2 | rabaptin, RAB GTPase binding effector protein 2 | Cytoplasm | other | |
| chr5 | *TXNDC15* | 10 | 282 | 21 | TXNDC15 | thioredoxin domain containing 15 | Extracellular Space | enzyme | |
| chr17 | *TOP2A* | 8 | 214 | 21 | TOP2A | DNA topoisomerase II alpha | Nucleus | enzyme | bortezomib/doxorubicin, daunorubicin/tretinoin, idarubicin, cladribine/cytarabine/filgrastim/idarubicin/plerixafor, cisplatin/doxorubicin/methotrexate, cytarabine/idarubicin, cytarabine/daunorubicin, cytarabine/doxorubicin, cyclophosphamide/epirubicin, doxorubicin/paclitaxel, fludarabine phosphate/mitoxantrone/rituximab, novobiocin, bevacizumab/etoposide, [illegible] |

| | | | | | | | | | |
|---|---|---|---|---|---|---|---|---|---|
| chr12 | *LIN7A* | 8 | 234 | 21 | LIN7A | lin-7 homolog A, crumbs cell polarity complex component | Plasma Membrane | other | |
| chr14 | *HSP90AA1* | 9 | 233 | 21 | HSP90AA1 | heat shock protein 90 alpha family class A member 1 | Cytoplasm | enzyme | luminespib, cisplatin, retaspimycin, alvespimycin, TAS- |
| chr12 | *IKZF4* | 9 | 476 | 21 | IKZF4 | IKAROS family zinc finger 4 | Nucleus | transcription regulator | |
| chr17 | *C1QTNF1* | 9 | 400 | 21 | C1QTNF1 | C1q and TNF related 1 | Extracellular Space | other | |
| chr12 | *TRAFD1* | 9 | 263 | 21 | TRAFD1 | TRAF-type zinc finger domain containing 1 | Other | other | |
| chr16 | *CTB-31N19.2* | 10 | 653 | 21 | | | | | |
| chr15 | *EFTUD1* | 9 | 237 | 21 | EFL1 | elongation factor like GTPase 1 | Cytoplasm | translation regulator | |
| chr7 | *CLCN1* | 9 | 296 | 21 | CLCN1 | chloride voltage-gated channel 1 | Plasma Membrane | ion channel | |
| chr17 | *PLCD3* | 9 | 574 | 21 | PLCD3 | phospholipase C delta 3 | Cytoplasm | enzyme | |
| chr5 | *ZCCHC10* | 9 | 224 | 21 | ZCCHC10 | zinc finger CCHC-type containing | Other | other | |
| chr5 | *BNIP1* | 8 | 337 | 21 | BNIP1 | BCL2 interacting protein 1 | Cytoplasm | transporter | |
| chr7 | *MET* | 10 | 232 | 21 | MET | MET proto-oncogene, receptor tyrosine kinase | Plasma Membrane | kinase | EMD 1204831, volitinib, TQ-B3139, bozitinib, onartuzumab, TR1801, LY3164530, ARGX-111, glumetinib, OMO-1, AL2846, MK2461, EMI-137, cabozantinib/erlotinib, crizotinib, tivantinib, ABT-700, JNJ-38877605, AMG 208, EMB-01, amuvatinib, c-MET inhibitor, LY2875358, SAR125844, crizotinib/vemurafenib, cabozantinib/nivolumab, merestinib, foretinib, glesatinib, SAIT301, [illegible] |
| chr15 | *NIPA2* | 8 | 304 | 21 | NIPA2 | NIPA magnesium transporter 2 | Cytoplasm | other | |
| chr7 | *DPY19L2P2* | 8 | 210 | 21 | DPY19L2P2 | DPY19L2 pseudogene 2 | Other | other | |
| chr3 | *SIDT1* | 8 | 240 | 21 | SIDT1 | SID1 transmembrane family member 1 | Other | other | |
| chr7 | *PARP12* | 9 | 402 | 21 | PARP12 | poly(ADP-ribose) polymerase family member 12 | Nucleus | enzyme | |
| chr15 | *SPG21* | 8 | 220 | 21 | SPG21 | SPG21 abhydrolase domain containing, maspardin | Plasma Membrane | enzyme | |
| chr1 | *WDR78* | 8 | 172 | 21 | WDR78 | WD repeat domain 78 | Other | other | |
| chr19 | *TCF3* | 11 | 516 | 21 | TCF3 | transcription factor 3 | Nucleus | transcription regulator | |
| chr9 | *SYK* | 7 | 252 | 21 | SYK | spleen associated tyrosine kinase | Cytoplasm | kinase | SKI-O-592, cerdulatinib, fostamatinib, HMPL-523, GSK2646264, R-348, entospletinib, |
| chr1 | *LINC00505* | 10 | 464 | 21 | FAAHP1 | fatty acid amide hydrolase pseudogene 1 | Other | other | |
| chr3 | *CPNE9* | 9 | 341 | 21 | CPNE9 | copine family member 9 | Extracellular Space | other | |
| chr2 | *NCOA1* | 7 | 177 | 21 | NCOA1 | nuclear receptor coactivator 1 | Nucleus | transcription regulator | |
| chr2 | *AC008074.3* | 9 | 200 | 21 | | | | | |
| chr11 | *DPF2* | 10 | 588 | 21 | DPF2 | double PHD fingers 2 | Nucleus | transcription regulator | |
| chr22 | *HIC2* | 9 | 488 | 21 | HIC2 | HIC ZBTB transcriptional | Nucleus | transcription regulator | |
| chr1 | *PCSK9* | 9 | 482 | 21 | PCSK9 | proprotein convertase subtilisin/kexin type 9 | Extracellular Space | peptidase | PCSK9 inhibitor, BMS-844421, evinacumab, evolocumab, |
| chr16 | *ZNF720* | 10 | 241 | 21 | ZNF720 | zinc finger protein 720 | Other | other | |
| chr11 | *TOLLIP* | 9 | 532 | 21 | TOLLIP | toll interacting protein | Cytoplasm | other | |
| chr7 | *TTYH3* | 12 | 561 | 21 | TTYH3 | tweety family member 3 | Plasma Membrane | ion channel | |
| chr17 | *DNAJC7* | 9 | 224 | 21 | DNAJC7 | DnaJ heat shock protein family (Hsp40) member C7 | Cytoplasm | other | |
| chr16 | *METTL9* | 8 | 224 | 21 | METTL9 | methyltransferase like 9 | Other | other | |
| chr15 | *RP11-321G12.1* | 9 | 208 | 21 | | | | | |
| chr6 | *DEK* | 7 | 209 | 21 | DEK | DEK proto-oncogene | Nucleus | transcription regulator | |
| chr7 | *AC011294.3* | 10 | 259 | 21 | | | | | |
| chr17 | *LRRC37B* | 8 | 220 | 21 | LRRC37B | leucine rich repeat containing | Cytoplasm | other | |
| chr1 | *OSCP1* | 8 | 260 | 21 | OSCP1 | organic solute carrier partner 1 | Plasma Membrane | transporter | |
| chr14 | *BAZ1A* | 7 | 177 | 21 | BAZ1A | bromodomain adjacent to zinc finger domain 1A | Nucleus | other | |
| chr12 | *SUDS3* | 9 | 200 | 21 | SUDS3 | SDS3 homolog, SIN3A corepressor complex component | Nucleus | transcription regulator | |
| chr1 | *PDZK1* | 10 | 292 | 21 | PDZK1 | PDZ domain containing 1 | Plasma Membrane | other | |
| chr12 | *DDX11-AS1* | 10 | 326 | 21 | DDX11-AS1 | DDX11 antisense RNA 1 | Other | other | |
| chr4 | *MAPK10* | 8 | 209 | 21 | MAPK10 | mitogen-activated protein kinase 10 | Cytoplasm | kinase | cetuximab/encorafenib, binimetinib/encorafenib, encorafenib, binimetinib/cetuximab/encorafenib, binimetinib/encorafen |

| | | | | | | | | | |
|---|---|---|---|---|---|---|---|---|---|
| chr5 | *RASA1* | 9 | 228 | 21 | RASA1 | RAS p21 protein activator 1 | Cytoplasm | transporter | |
| chr5 | *CDC23* | 8 | 211 | 21 | CDC23 | cell division cycle 23 | Nucleus | enzyme | |
| chr3 | *TPRA1* | 10 | 617 | 21 | TPRA1 | transmembrane protein adipocyte associated 1 | Plasma Membrane | ;-protein coupled recepto | |
| chr17 | *MYCBPAP* | 9 | 331 | 21 | MYCBPAP | MYCBP associated protein | Nucleus | other | |
| chr7 | *ABCB4* | 9 | 249 | 21 | ABCB4 | ATP binding cassette subfamily B member 4 | Plasma Membrane | transporter | |
| chr12 | *RP11-352M15.1* | 10 | 317 | 21 | | | | | |
| chr5 | *ELOVL7* | 9 | 209 | 21 | ELOVL7 | ELOVL fatty acid elongase 7 | Cytoplasm | enzyme | |
| chr6 | *C6orf222* | 9 | 304 | 21 | BNIP5 | BCL2 interacting protein 5 | Other | other | |
| chr17 | *COL1A1* | 10 | 457 | 21 | COL1A1 | collagen type I alpha 1 chain | Extracellular Space | other | collagenase clostridium |
| chr1 | *PTPN7* | 9 | 512 | 21 | PTPN7 | protein tyrosine phosphatase non-receptor type 7 | Cytoplasm | phosphatase | |
| chr22 | *GTSE1* | 8 | 311 | 21 | GTSE1 | G2 and S-phase expressed 1 | Cytoplasm | other | |
| chr5 | *MARCH11;* | 10 | 435 | 21 | | | | | |
| chr19 | *ZNF91* | 9 | 195 | 21 | ZNF91 | zinc finger protein 91 | Nucleus | transcription regulator | |
| chr2 | *PKP4* | 8 | 209 | 21 | PKP4 | plakophilin 4 | Plasma Membrane | other | |
| chr17 | *HEXDC* | 9 | 328 | 21 | HEXD | hexosaminidase D | Extracellular Space | enzyme | |
| chr6 | *CDK19* | 7 | 187 | 21 | CDK19 | cyclin dependent kinase 19 | Nucleus | kinase | SEL120, BCD-115, alvocidib |
| chr13 | *LINC00452* | 9 | 312 | 21 | LINC00452 | long intergenic non-protein coding RNA 452 | Other | other | |
| chr11 | *RPS3* | 9 | 333 | 21 | RPS3 | ribosomal protein S3 | Cytoplasm | enzyme | |
| chr2 | *AC062028.1* | 9 | 311 | 21 | | | | | |
| chr7 | *SEPT7* | 8 | 234 | 21 | SEPTIN7 | septin 7 | Cytoplasm | other | |
| chr14 | *MDGA2* | 7 | 204 | 21 | MDGA2 | MAM domain containing glycosylphosphatidylinositol | Plasma Membrane | other | |
| chr10 | *TAF3* | 7 | 182 | 21 | TAF3 | TATA-box binding protein associated factor 3 | Nucleus | transcription regulator | |
| chr17 | *APOH* | 9 | 188 | 21 | APOH | apolipoprotein H | Extracellular Space | transporter | |
| chr2 | *MAL* | 10 | 499 | 21 | MAL | mal, T cell differentiation protein | Plasma Membrane | other | |
| chr7 | *SP4* | 9 | 222 | 21 | SP4 | Sp4 transcription factor | Nucleus | transcription regulator | |
| chr17 | *DUSP14* | 8 | 290 | 21 | DUSP14 | dual specificity phosphatase 14 | Other | phosphatase | |
| chr2 | *SLC4A10* | 7 | 209 | 21 | SLC4A10 | solute carrier family 4 member 10 | Plasma Membrane | transporter | |
| chr20 | *CASS4* | 10 | 456 | 21 | CASS4 | Cas scaffold protein family | Plasma Membrane | other | |
| chr11 | *FAM118B* | 8 | 252 | 21 | FAM118B | family with sequence similarity 118 member B | Other | other | |
| chr14 | *MIPOL1* | 7 | 193 | 21 | MIPOL1 | mirror-image polydactyly 1 | Nucleus | other | |
| chr3 | *TMEM45A* | 10 | 283 | 21 | TMEM45A | transmembrane protein 45A | Plasma Membrane | other | |
| chr5 | *RP11-434D11.4* | 7 | 231 | 21 | | | | | |
| chr12 | *TBX5* | 10 | 240 | 21 | TBX5 | T-box transcription factor 5 | Nucleus | transcription regulator | |
| chr14 | *PLEKHG3* | 9 | 385 | 21 | PLEKHG3 | pleckstrin homology and RhoGEF domain containing G3 | Other | other | |
| chr12 | *CRY1* | 8 | 170 | 21 | CRY1 | cryptochrome circadian regulator | Nucleus | enzyme | |
| chr10 | *HTR7* | 9 | 221 | 21 | HTR7 | 5-hydroxytryptamine receptor 7 | Plasma Membrane | ;-protein coupled recepto | methysergide, almotriptan, brexpiprazole, fenfluramine, vortioxetine, iloperidone, ergotamine, eletriptan, lurasidone, sultopride, |
| chr20 | *RP1-148H17.1* | 9 | 295 | 21 | | | | | |
| chr11 | *RP13-726E6.2* | 10 | 435 | 21 | | | | | |
| chr6 | *ABCC10* | 9 | 361 | 21 | ABCC10 | ATP binding cassette subfamily C member 10 | Plasma Membrane | transporter | |
| chr2 | *PTH2R* | 8 | 217 | 21 | PTH2R | parathyroid hormone 2 receptor | Plasma Membrane | ;-protein coupled recepto | PTH, raloxifene/teriparatid |
| chr15 | *C15orf60* | 9 | 231 | 21 | REC114 | REC114 meiotic recombination protein | Other | other | |
| chr5 | *XRCC4* | 8 | 217 | 21 | XRCC4 | X-ray repair cross complementing | Nucleus | other | |
| chr3 | *PLD1* | 7 | 181 | 21 | PLD1 | phospholipase D1 | Cytoplasm | enzyme | |
| chr20 | *BPIFB2* | 9 | 455 | 21 | BPIFB2 | BPI fold containing family B member 2 | Extracellular Space | other | |
| chr6 | *PPP2R5D* | 9 | 268 | 21 | PPP2R5D | protein phosphatase 2 regulatory subunit B'delta | Nucleus | phosphatase | |
| chr1 | *DARS2* | 7 | 238 | 21 | DARS2 | aspartyl-tRNA synthetase 2, mitochondrial | Cytoplasm | enzyme | |
| chr1 | *LAMTOR5-AS1* | 9 | 308 | 21 | LAMTOR5-AS1 | LAMTOR5 antisense RNA 1 | Other | other | |
| chr5 | *TRIP13* | 9 | 942 | 21 | TRIP13 | thyroid hormone receptor interactor 13 | Cytoplasm | transcription regulator | |
| chr15 | *NPTN* | 8 | 193 | 21 | NPTN | neuroplastin | Plasma Membrane | other | |
| chr22 | *CYTH4* | 9 | 383 | 21 | CYTH4 | cytohesin 4 | Cytoplasm | other | |
| chr1 | *PLA2G4A* | 8 | 236 | 21 | PLA2G4A | phospholipase A2 group IVA | Cytoplasm | enzyme | cytosolic phospholipase A2 |
| chr15 | *RP11-2E17.1* | 11 | 262 | 20 | | | | | |
| chr5 | *TRAPPC13* | 8 | 179 | 20 | TRAPPC13 | trafficking protein particle | Cytoplasm | other | |
| chrX | *BEND2* | 9 | 266 | 20 | BEND2 | BEN domain containing 2 | Other | other | |
| chr19 | *HSPBP1* | 9 | 395 | 20 | HSPBP1 | HSPA (Hsp70) binding protein 1 | Cytoplasm | other | |
| chr21 | *BAGE2* | 10 | 973 | 20 | BAGE2 | BAGE family member 2 | Other | other | |
| chr8 | *ZMAT4* | 8 | 222 | 20 | ZMAT4 | zinc finger matrin-type 4 | Nucleus | other | |
| chr5 | *SEPP1* | 8 | 570 | 20 | SELENOP | selenoprotein P | Extracellular Space | other | |
| chr6 | *MOCS1* | 10 | 278 | 20 | MOCS1 | molybdenum cofactor synthesis 1 | Cytoplasm | other | |
| chr12 | *VPS37B* | 10 | 284 | 20 | VPS37B | VPS37B subunit of ESCRT-I | Cytoplasm | other | |
| chr22 | *RP1-63G5.5* | 8 | 412 | 20 | | | | | |
| chr11 | *GLB1L3* | 10 | 356 | 20 | GLB1L3 | galactosidase beta 1 like 3 | Other | enzyme | |
| chr9 | *ZNF658* | 7 | 173 | 20 | ZNF658 | zinc finger protein 658 | Other | other | |

| | | | | | | | | | |
|---|---|---|---|---|---|---|---|---|---|
| chr2 | *CAB39* | 7 | 162 | 20 | CAB39 | calcium binding protein 39 | Cytoplasm | enzyme | |
| chr12 | *RP11-110L15.2* | 9 | 213 | 20 | | | | | |
| chr19 | *TTYH1* | 10 | 528 | 20 | TTYH1 | tweety family member 1 | Plasma Membrane | ion channel | |
| chr19 | *NLRP7* | 9 | 298 | 20 | NLRP7 | NLR family pyrin domain | Extracellular Space | other | |
| chr6 | *RAB44* | 8 | 379 | 20 | RAB44 | RAB44, member RAS oncogene family | Cytoplasm | other | |
| chr3 | *ITIH1* | 8 | 375 | 20 | ITIH1 | inter-alpha-trypsin inhibitor heavy chain 1 | Extracellular Space | other | |
| chr2 | *ANKRD36* | 8 | 376 | 20 | ANKRD36 | ankyrin repeat domain 36 | Other | other | |
| chr15 | *CHRNA5* | 6 | 211 | 20 | CHRNA5 | cholinergic receptor nicotinic alpha 5 subunit | Plasma Membrane | transmembrane receptor | succinylcholine, rapacuronium, doxacurium, vecuronium, pipecuronium, metocurine, mivacurium, D-tubocurarine, nicotine, arecoline, pancuronium, atracurium, isoflurane, cisatracurium, rocuronium, amobarbital, mecamylamine |
| chr8 | *CLU* | 8 | 303 | 20 | CLU | clusterin | Cytoplasm | other | clusterin antisense oligonucleotide |
| chr1 | *CTRC* | 8 | 371 | 20 | CTRC | chymotrypsin C | Extracellular Space | peptidase | |
| chr11 | *NLRX1* | 10 | 387 | 20 | NLRX1 | NLR family member X1 | Cytoplasm | other | |
| chr12 | *CELA1* | 10 | 390 | 20 | CELA1 | chymotrypsin like elastase 1 | Extracellular Space | peptidase | |
| chr8 | *RP11-513O17.2* | 9 | 581 | 20 | | | | | |
| chr18 | *RP11-865B13.1* | 10 | 358 | 20 | | | | | |
| chr20 | *OVOL2* | 8 | 243 | 20 | OVOL2 | ovo like zinc finger 2 | Nucleus | transcription regulator | |
| chr2 | *TWIST2* | 8 | 354 | 20 | TWIST2 | twist family bHLH transcription factor 2 | Nucleus | transcription regulator | |
| chrX | *WDR44* | 8 | 197 | 20 | WDR44 | WD repeat domain 44 | Cytoplasm | other | |
| chr6 | *RP1-153P14.8* | 8 | 293 | 20 | | | | | |
| chr3 | *RP11-572M11.4* | 10 | 250 | 20 | | | | | |
| chr16 | *NOD2* | 9 | 345 | 20 | NOD2 | nucleotide binding oligomerization domain containing 2 | Cytoplasm | other | |
| chr19 | *PPFIA3* | 9 | 352 | 20 | PPFIA3 | PTPRF interacting protein alpha 3 | Plasma Membrane | phosphatase | |
| chr3 | *AC027119.1* | 9 | 176 | 20 | | | | | |
| chr19 | *NLRP13* | 9 | 197 | 20 | NLRP13 | NLR family pyrin domain containing 13 | Other | other | |
| chr17 | *CTC-264K15.6* | 9 | 373 | 20 | | | | | |
| chr1 | *MCOLN2* | 8 | 204 | 20 | MCOLN2 | mucolipin 2 | Plasma Membrane | ion channel | |
| chr16 | *RP11-482M8.1* | 9 | 377 | 20 | | | | | |
| chr16 | *TAOK2* | 9 | 372 | 20 | TAOK2 | TAO kinase 2 | Cytoplasm | kinase | |
| chr16 | *NOMO3* | 8 | 235 | 20 | MO1 (includes oth | NODAL modulator 1 | Plasma Membrane | other | |
| chr10 | *SVILP1* | 8 | 258 | 20 | SVIL2P | supervillin family member 2, pseudogene | Other | other | |
| chr20 | *DUSP15* | 10 | 595 | 20 | DUSP15 | dual specificity phosphatase 15 | Cytoplasm | phosphatase | |
| chr10 | *ANXA7* | 8 | 316 | 20 | ANXA7 | annexin A7 | Plasma Membrane | ion channel | |
| chr5 | *CTC-329D1.2* | 9 | 239 | 20 | | | | | |
| chr22 | *PARVG* | 8 | 387 | 20 | PARVG | parvin gamma | Cytoplasm | other | |
| chr2 | *ZC3H8* | 8 | 219 | 20 | ZC3H8 | zinc finger CCCH-type containing | Nucleus | transcription regulator | |
| chr16 | *RP11-252A24.2* | 8 | 231 | 20 | | | | | |
| chr4 | *LDB2* | 7 | 187 | 20 | LDB2 | LIM domain binding 2 | Nucleus | transcription regulator | |
| chr9 | *ZCCHC7* | 7 | 161 | 20 | ZCCHC7 | zinc finger CCHC-type containing | Nucleus | other | |
| chr22 | *ZMAT5* | 8 | 302 | 20 | ZMAT5 | zinc finger matrin-type 5 | Nucleus | other | |
| chr2 | *AC013271.3* | 9 | 294 | 20 | | | | | |
| chr12 | *RP11-74M13.4* | 9 | 196 | 20 | | | | | |
| chr15 | *CTD-2315E11.1* | 10 | 359 | 20 | | | | | |
| chr17 | *RP11-74E22.4* | 9 | 311 | 20 | | | | | |
| chr3 | *TGFBR2* | 9 | 241 | 20 | TGFBR2 | transforming growth factor beta receptor 2 | Plasma Membrane | kinase | IMC-TR1 |
| chr20 | *XKR7* | 9 | 395 | 20 | XKR7 | XK related 7 | Other | other | |
| chr7 | *WDR91* | 9 | 427 | 20 | WDR91 | WD repeat domain 91 | Cytoplasm | other | |
| chr2 | *AC007563.5* | 8 | 216 | 20 | | | | | |
| chr11 | *NPAT* | 9 | 220 | 20 | NPAT | nuclear protein, coactivator of histone transcription | Nucleus | transcription regulator | |
| chr9 | *GLE1* | 9 | 369 | 20 | GLE1 | GLE1 RNA export mediator | Nucleus | other | |
| chr20 | *RAB22A* | 9 | 323 | 20 | RAB22A | RAB22A, member RAS oncogene family | Cytoplasm | enzyme | |
| chr20 | *APMAP* | 9 | 370 | 20 | APMAP | adipocyte plasma membrane associated protein | Plasma Membrane | enzyme | |
| chr2 | *VIL1* | 8 | 276 | 20 | VIL1 | villin 1 | Cytoplasm | other | |
| chr9 | *MORN5* | 9 | 387 | 20 | MORN5 | MORN repeat containing 5 | Other | other | |
| chr10 | *BMS1P5* | 8 | 372 | 20 | BMS1P1 | BMS1 pseudogene 1 | Other | other | |
| chr12 | *SLC16A7* | 9 | 267 | 20 | SLC16A7 | solute carrier family 16 member 7 | Plasma Membrane | transporter | |
| chr7 | *BRAT1* | 10 | 463 | 20 | BRAT1 | BRCA1 associated ATM activator | Nucleus | other | |
| chr12 | *RP11-173P15.7* | 9 | 289 | 20 | | | | | |
| chr19 | *ZNF274* | 9 | 254 | 20 | ZNF274 | zinc finger protein 274 | Nucleus | transcription regulator | |
| chr6 | *CMAHP* | 9 | 201 | 20 | CMAHP | cytidine monophospho-N-acetylneuraminic acid hydroxylase, pseudogene | Cytoplasm | other | |
| chr1 | *MROH9* | 8 | 247 | 20 | MROH9 | maestro heat like repeat family member 9 | Other | other | |
| chr2 | *CBWD2* | 9 | 287 | 20 | CBWD2 | COBW domain containing 2 | Other | other | |

| | | | | | | | | | |
|---|---|---|---|---|---|---|---|---|---|
| chr17 | *CPD* | 8 | 223 | 20 | CPD | carboxypeptidase D | Extracellular Space | peptidase | |
| chr3 | *XPC* | 9 | 290 | 20 | XPC | XPC complex subunit, DNA damage recognition and repair | Nucleus | other | |
| chr12 | *MMP17* | 10 | 560 | 20 | MMP17 | matrix metallopeptidase 17 | Extracellular Space | peptidase | marimastat |
| chr11 | *ZDHHC5* | 8 | 300 | 20 | ZDHHC5 | zinc finger DHHC-type containing | Nucleus | enzyme | |
| chr22 | *TTC38* | 8 | 329 | 20 | TTC38 | tetratricopeptide repeat domain | Cytoplasm | other | |
| chr1 | *TMEM57* | 8 | 197 | 20 | MACO1 | macoilin 1 | Nucleus | other | |
| chr18 | *LINC00669* | 7 | 238 | 20 | MIR924HG | MIR924 host gene | Other | other | |
| chr3 | *ZFYVE20* | 8 | 233 | 20 | RBSN | rabenosyn, RAB effector | Cytoplasm | other | |
| chr7 | *COX19* | 10 | 558 | 20 | COX19 | cytochrome c oxidase assembly factor COX19 | Cytoplasm | other | |
| chr11 | *RP11-360K13.1* | 9 | 373 | 20 | | | | | |
| chr3 | *MCCC1* | 8 | 188 | 20 | MCCC1 | methylcrotonoyl-CoA carboxylase | Cytoplasm | enzyme | |
| chr12 | *LRRIQ1* | 8 | 174 | 20 | LRRIQ1 | leucine rich repeats and IQ motif containing 1 | Other | other | |
| chr17 | *ALOXE3* | 10 | 315 | 20 | ALOXE3 | arachidonate lipoxygenase 3 | Extracellular Space | enzyme | |
| chr19 | *CLEC17A* | 9 | 249 | 20 | CLEC17A | C-type lectin domain containing | Plasma Membrane | other | |
| chr7 | *VWC2* | 8 | 267 | 20 | VWC2 | von Willebrand factor C domain containing 2 | Extracellular Space | other | |
| chr16 | *RP11-314O13.1* | 8 | 186 | 20 | | | | | |
| chr11 | *DLAT* | 8 | 195 | 20 | DLAT | dihydrolipoamide S- | Cytoplasm | enzyme | |
| chr5 | *MCCC2* | 9 | 214 | 20 | MCCC2 | methylcrotonoyl-CoA carboxylase | Cytoplasm | enzyme | |
| chr1 | *SMG7* | 8 | 271 | 20 | SMG7 | SMG7 nonsense mediated mRNA decay factor | Nucleus | other | |
| chr19 | *SLC27A5* | 9 | 346 | 20 | SLC27A5 | solute carrier family 27 member 5 | Cytoplasm | transporter | |
| chr12 | *RP3-473L9.4* | 9 | 303 | 20 | | | | | |
| chr2 | *CCDC108* | 7 | 316 | 20 | CFAP65 | cilia and flagella associated | Other | other | |
| chr19 | *PLA2G4C* | 8 | 218 | 20 | PLA2G4C | phospholipase A2 group IVC | Plasma Membrane | enzyme | |
| chr16 | *TBC1D24* | 10 | 393 | 20 | TBC1D24 | TBC1 domain family member 24 | Cytoplasm | other | |
| chr1 | *CAPZA1* | 8 | 251 | 20 | CAPZA1 | capping actin protein of muscle Z-line subunit alpha 1 | Cytoplasm | other | |
| chr10 | *RTKN2* | 8 | 272 | 20 | RTKN2 | rhotekin 2 | Plasma Membrane | other | |
| chr12 | *SCAF11* | 7 | 306 | 20 | SCAF11 | SR-related CTD associated factor | Nucleus | other | |
| chr3 | *DAZL* | 9 | 253 | 20 | DAZL | deleted in azoospermia like | Cytoplasm | translation regulator | |
| chr7 | *LAMB1* | 9 | 245 | 20 | LAMB1 | laminin subunit beta 1 | Extracellular Space | other | |
| chr15 | *CPEB1* | 9 | 304 | 20 | CPEB1 | cytoplasmic polyadenylation element binding protein 1 | Cytoplasm | translation regulator | |
| chr22 | *DMC1* | 7 | 184 | 20 | DMC1 | DNA meiotic recombinase 1 | Nucleus | enzyme | |
| chr5 | *NSUN2* | 9 | 799 | 20 | NSUN2 | NOP2/Sun RNA | Nucleus | enzyme | |
| chr9 | *LINC00476* | 7 | 219 | 20 | LINC00476 | long intergenic non-protein coding RNA 476 | Other | other | |
| chr11 | *MRPL11* | 10 | 362 | 20 | MRPL11 | mitochondrial ribosomal protein | Cytoplasm | other | |
| chr16 | *MTHFSD* | 9 | 380 | 20 | MTHFSD | methenyltetrahydrofolate synthetase domain containing | Other | other | |
| chr12 | *KCNMB4* | 9 | 180 | 20 | KCNMB4 | potassium calcium-activated channel subfamily M regulatory | Plasma Membrane | ion channel | tedisamil |
| chrX | *MED14* | 9 | 227 | 20 | MED14 | mediator complex subunit 14 | Nucleus | transcription regulator | |
| chr5 | *RP11-259O2.1* | 9 | 1056 | 20 | | | | | |
| chr1 | *RP11-634B7.4* | 9 | 234 | 20 | | | | | |
| chr19 | *CLASRP* | 8 | 336 | 20 | CLASRP | CLK4 associating serine/arginine rich protein | Nucleus | other | |
| chr15 | *PTPLAD1* | 7 | 173 | 20 | HACD3 | 3-hydroxyacyl-CoA dehydratase | Cytoplasm | enzyme | |
| chr3 | *ARL8B* | 8 | 182 | 20 | ARL8B | ADP ribosylation factor like GTPase 8B | Plasma Membrane | enzyme | |
| chr11 | *LDHC* | 8 | 197 | 20 | LDHC | lactate dehydrogenase C | Cytoplasm | enzyme | |
| chr1 | *CCDC27* | 9 | 388 | 20 | CCDC27 | coiled-coil domain containing 27 | Other | other | |
| chr2 | *SLC1A4* | 8 | 259 | 20 | SLC1A4 | solute carrier family 1 member 4 | Plasma Membrane | transporter | riluzole |
| chr1 | *RP5-1074L1.1* | 10 | 310 | 20 | | | | | |
| chr10 | *MCMBP* | 7 | 196 | 20 | MCMBP | minichromosome maintenance complex binding protein | Nucleus | other | |
| chr22 | *ZBED4* | 8 | 309 | 20 | ZBED4 | zinc finger BED-type containing 4 | Nucleus | transcription regulator | |
| chr2 | *PLEKHM3* | 7 | 182 | 20 | PLEKHM3 | pleckstrin homology domain containing M3 | Other | other | |
| chr1 | *NASP* | 9 | 215 | 20 | NASP | nuclear autoantigenic sperm | Nucleus | other | |
| chr4 | *FAM184B* | 8 | 238 | 20 | FAM184B | family with sequence similarity 184 member B | Other | other | |
| chr10 | *ZNF438* | 8 | 222 | 20 | ZNF438 | zinc finger protein 438 | Nucleus | transcription regulator | |
| chr5 | *DEPDC1B* | 8 | 225 | 20 | DEPDC1B | DEP domain containing 1B | Cytoplasm | other | |
| chr9 | *PLIN2* | 9 | 184 | 20 | PLIN2 | perilipin 2 | Plasma Membrane | other | |
| chrX | *EIF2S3* | 8 | 234 | 20 | EIF2S3 | eukaryotic translation initiation factor 2 subunit gamma | Cytoplasm | translation regulator | |
| chr11 | *CAPN1* | 9 | 474 | 20 | CAPN1 | calpain 1 | Cytoplasm | peptidase | |
| chr18 | *DCC* | 7 | 207 | 20 | DCC | DCC netrin 1 receptor | Plasma Membrane | transmembrane receptor | |
| chr3 | *DNASE1L3* | 9 | 269 | 20 | DNASE1L3 | deoxyribonuclease 1 like 3 | Nucleus | enzyme | |
| chr10 | *AGAP9* | 7 | 232 | 20 | AP5 (includes oth | ArfGAP with GTPase domain, ankyrin repeat and PH domain 5 | Other | other | |
| chr12 | *DENR* | 7 | 241 | 20 | DENR | density regulated re-initiation and release factor | Other | other | |
| chr3 | *RP11-680B3.2* | 9 | 219 | 20 | | | | | |
| chr17 | *FAM134C* | 9 | 217 | 20 | RETREG3 | reticulophagy regulator family member 3 | Other | other | |
| chr11 | *PHRF1* | 9 | 464 | 20 | PHRF1 | PHD and ring finger domains 1 | Nucleus | other | |
| chr16 | *NPIPB8* | 8 | 231 | 20 | PB15 (includes oth | nuclear pore complex interacting protein family member B15 | Nucleus | other | |
| chr3 | *TDGF1* | 9 | 214 | 20 | TDGF1 | teratocarcinoma-derived growth factor 1 | Extracellular Space | growth factor | huB3F6-SPDB-DM4 immunoconjugate |
| chrX | *MAGT1* | 7 | 179 | 20 | MAGT1 | magnesium transporter 1 | Plasma Membrane | enzyme | |
| chr11 | *CUL5* | 8 | 197 | 20 | CUL5 | cullin 5 | Nucleus | ion channel | |

| | | | | | | | | | |
|---|---|---|---|---|---|---|---|---|---|
| chr18 | *ROCK1* | 7 | 170 | 20 | ROCK1 | Rho associated coiled-coil containing protein kinase 1 | Cytoplasm | kinase | |
| chr13 | *ATP7B* | 7 | 229 | 20 | ATP7B | ATPase copper transporting beta | Cytoplasm | transporter | |
| chr15 | *ANKDD1A* | 7 | 284 | 20 | ANKDD1A | ankyrin repeat and death domain containing 1A | Other | other | |
| chr9 | *CLTA* | 7 | 193 | 20 | CLTA | clathrin light chain A | Plasma Membrane | other | |
| chr5 | *AFAP1L1* | 7 | 259 | 20 | AFAP1L1 | actin filament associated protein 1 like 1 | Other | other | |
| chr12 | *PAH* | 10 | 213 | 20 | PAH | phenylalanine hydroxylase | Cytoplasm | enzyme | sapropterin, 5,6,7,8-tetrahydrobiopterin |
| chrX | *TKTL1* | 8 | 271 | 20 | TKTL1 | transketolase like 1 | Cytoplasm | enzyme | |
| chr21 | *DSCR4* | 7 | 178 | 20 | DSCR4 | Down syndrome critical region 4 | Other | other | |
| chr9 | *TLE4* | 9 | 203 | 20 | TLE4 | TLE family member 4, transcriptional corepressor | Nucleus | transcription regulator | |
| chr3 | *IQCB1* | 9 | 202 | 20 | IQCB1 | IQ motif containing B1 | Extracellular Space | other | |
| chr8 | *RP11-706C16.8* | 9 | 427 | 20 | | | | | |
| chr14 | *ZNF839* | 8 | 256 | 20 | ZNF839 | zinc finger protein 839 | Other | other | |
| chr2 | *ST6GAL2* | 8 | 203 | 20 | ST6GAL2 | ST6 beta-galactoside alpha-2,6-sialyltransferase 2 | Cytoplasm | enzyme | |
| chr3 | *FAM208A* | 7 | 178 | 20 | TASOR | transcription activation | Nucleus | other | |
| chr20 | *NFS1* | 9 | 302 | 20 | NFS1 | NFS1 cysteine desulfurase | Cytoplasm | enzyme | |
| chr6 | *LINC01016* | 9 | 307 | 20 | LINC01016 | long intergenic non-protein coding RNA 1016 | Other | other | |
| chr17 | *NBR1* | 8 | 272 | 20 | NBR1 | NBR1 autophagy cargo receptor | Cytoplasm | other | |
| chr11 | *CDC42BPG* | 10 | 525 | 20 | CDC42BPG | CDC42 binding protein kinase gamma | Cytoplasm | kinase | |
| chr7 | *NUB1* | 8 | 295 | 20 | NUB1 | negative regulator of ubiquitin like proteins 1 | Nucleus | other | |
| chr15 | *MAN2A2* | 9 | 717 | 20 | MAN2A2 | mannosidase alpha class 2A member 2 | Cytoplasm | enzyme | |
| chr1 | *GRHL3* | 9 | 378 | 20 | GRHL3 | grainyhead like transcription | Nucleus | transcription regulator | |
| chr12 | *CCDC92* | 8 | 296 | 20 | CCDC92 | coiled-coil domain containing 92 | Cytoplasm | other | |
| chr1 | *NADK* | 9 | 467 | 20 | NADK | NAD kinase | Cytoplasm | kinase | |
| chr12 | *RILPL2* | 9 | 257 | 20 | RILPL2 | Rab interacting lysosomal protein like 2 | Cytoplasm | other | |
| chr21 | *TTC3* | 7 | 219 | 20 | TTC3 | tetratricopeptide repeat domain 3 | Cytoplasm | enzyme | |
| chr20 | *SIRPA* | 8 | 271 | 20 | SIRPA | signal regulatory protein alpha | Plasma Membrane | phosphatase | BI 765063, CC- |
| chr7 | *INTS4L2* | 9 | 201 | 20 | INTS4P2 | integrator complex subunit 4 pseudogene 2 | Other | other | |
| chr8 | *TACC1* | 8 | 215 | 20 | TACC1 | transforming acidic coiled-coil containing protein 1 | Nucleus | other | |
| chr9 | *FBP1* | 8 | 294 | 20 | FBP1 | fructose-bisphosphatase 1 | Cytoplasm | phosphatase | |
| chr15 | *MKRN3* | 8 | 243 | 20 | MKRN3 | makorin ring finger protein 3 | Nucleus | other | |
| chr5 | *RP11-98J23.2* | 8 | 772 | 20 | | | | | |
| chr2 | *AC079354.1* | 8 | 200 | 20 | | | | | |
| chrX | *SYN1* | 9 | 300 | 20 | SYN1 | synapsin I | Plasma Membrane | transporter | |
| chr7 | *AC016831.7* | 8 | 198 | 20 | | | | | |
| chr1 | *KCNK1* | 9 | 331 | 20 | KCNK1 | potassium two pore domain channel subfamily K member 1 | Plasma Membrane | ion channel | |
| chr1 | *TOR1AIP1* | 9 | 220 | 20 | TOR1AIP1 | torsin 1A interacting protein 1 | Nucleus | other | |
| chr9 | *GLIPR2* | 8 | 272 | 20 | GLIPR2 | GLI pathogenesis related 2 | Cytoplasm | other | |
| chr15 | *NR2E3* | 8 | 369 | 20 | NR2E3 | nuclear receptor subfamily 2 group E member 3 | Nucleus | d-dependent nuclear rec | |
| chr9 | *ZNF79* | 8 | 293 | 20 | ZNF79 | zinc finger protein 79 | Nucleus | transcription regulator | |
| chr8 | *ZNF707* | 9 | 365 | 20 | ZNF707 | zinc finger protein 707 | Other | other | |
| chr18 | *SETBP1* | 9 | 184 | 20 | SETBP1 | SET binding protein 1 | Nucleus | transcription regulator | |
| chr1 | *SARS* | 8 | 253 | 20 | SARS1 | seryl-tRNA synthetase 1 | Cytoplasm | enzyme | |
| chr5 | *APC* | 7 | 183 | 20 | APC | APC regulator of WNT signaling pathway | Nucleus | enzyme | |
| chr10 | *CCDC3* | 7 | 231 | 20 | CCDC3 | coiled-coil domain containing 3 | Cytoplasm | other | |
| chr3 | *SENP2* | 7 | 190 | 20 | SENP2 | SUMO specific peptidase 2 | Nucleus | peptidase | |
| chr17 | *CTNS* | 9 | 320 | 20 | CTNS | cystinosin, lysosomal cystine transporter | Cytoplasm | transporter | |
| chr6 | *GPR116* | 9 | 185 | 20 | ADGRF5 | adhesion G protein-coupled receptor F5 | Plasma Membrane | i-protein coupled recepto | |
| chr20 | *TTI1* | 9 | 316 | 20 | TTI1 | TELO2 interacting protein 1 | Cytoplasm | other | |
| chr9 | *UBAP1* | 7 | 194 | 20 | UBAP1 | ubiquitin associated protein 1 | Cytoplasm | other | |
| chr6 | *RIMS1* | 7 | 191 | 20 | RIMS1 | regulating synaptic membrane exocytosis 1 | Plasma Membrane | other | |
| chr19 | *CAMSAP3* | 10 | 461 | 20 | CAMSAP3 | calmodulin regulated spectrin associated protein family member | Cytoplasm | other | |
| chr6 | *RANBP9* | 7 | 206 | 20 | RANBP9 | RAN binding protein 9 | Nucleus | other | |
| chr14 | *SPTLC2* | 7 | 169 | 20 | SPTLC2 | serine palmitoyltransferase long chain base subunit 2 | Cytoplasm | enzyme | |
| chr10 | *INPP5F* | 8 | 197 | 20 | INPP5F | inositol polyphosphate-5-phosphatase F | Plasma Membrane | phosphatase | |
| chr19 | *FBXO17* | 9 | 280 | 20 | FBXO17 | F-box protein 17 | Cytoplasm | enzyme | |
| chr11 | *RP11-563P16.1* | 8 | 235 | 20 | | | | | |
| chr1 | *RP4-651E10.4* | 8 | 253 | 20 | | | | | |
| chr4 | *DHX15* | 7 | 160 | 20 | DHX15 | DEAH-box helicase 15 | Nucleus | enzyme | |
| chr11 | *CWF19L2* | 8 | 213 | 20 | CWF19L2 | CWF19 like cell cycle control | Other | other | |
| chr2 | *WDFY1* | 7 | 191 | 20 | WDFY1 | WD repeat and FYVE domain containing 1 | Cytoplasm | other | |
| chrX | *RP6-1O2.1* | 10 | 211 | 20 | | | | | |
| chr11 | *FADS3* | 9 | 536 | 20 | FADS3 | fatty acid desaturase 3 | Plasma Membrane | enzyme | |
| chr11 | *TREH* | 9 | 271 | 20 | TREH | trehalase | Plasma Membrane | enzyme | |
| chr9 | *SLC44A1* | 7 | 171 | 20 | SLC44A1 | solute carrier family 44 member 1 | Plasma Membrane | transporter | |
| chr10 | *RP11-318C4.2* | 9 | 245 | 20 | | | | | |

| | | | | | | | | | |
|---|---|---|---|---|---|---|---|---|---|
| chr15 | *FAM214A* | 7 | 163 | 20 | FAM214A | family with sequence similarity 214 member A | Other | other | |
| chr5 | *RAD50* | 8 | 194 | 20 | RAD50 | RAD50 double strand break repair protein | Nucleus | enzyme | |
| chr16 | *TK2* | 8 | 313 | 20 | TK2 | thymidine kinase 2 | Cytoplasm | kinase | |
| chr8 | *RP11-150O12.1* | 8 | 281 | 20 | | | | | |
| chr16 | *SLC6A2* | 9 | 272 | 20 | SLC6A2 | solute carrier family 6 member 2 | Plasma Membrane | transporter | orphenadrine, aspirin/caffeine/orphenadrine, methamphetamine, milnacipran, amoxapine, azatadine/pseudoephedrine, benzphetamine, phentermine/topiramate, naproxen/pseudoephedrine, loratadine/pseudoephedrine, chlorpheniramine/ibuprofen/pseudoephedrine, amitriptyline/melatonin, D-methylphenidate, bupropion/naltrexone, mazindol, [illegible] |
| chr11 | *C11orf70* | 7 | 418 | 20 | CFAP300 | cilia and flagella associated protein 300 | Cytoplasm | other | |
| chr2 | *AC093802.1* | 7 | 316 | 20 | | | | | |
| chr6 | *FYN* | 8 | 183 | 20 | FYN | FYN proto-oncogene, Src family tyrosine kinase | Plasma Membrane | kinase | blinatumomab/dasatinib, dasatinib/nilotinib, dasatinib/erlotinib, dasatinib/inotuzumab ozogamicin, JNJ- [illegible] |
| chr1 | *GTF2B* | 8 | 204 | 20 | GTF2B | general transcription factor IIB | Nucleus | transcription regulator | |
| chr5 | *SLC34A1* | 9 | 433 | 20 | SLC34A1 | solute carrier family 34 member 1 | Plasma Membrane | transporter | |
| chr17 | *CLUH* | 10 | 339 | 20 | CLUH | clustered mitochondria homolog | Cytoplasm | translation regulator | |
| chr17 | *ALKBH5* | 9 | 253 | 20 | ALKBH5 | alkB homolog 5, RNA | Nucleus | enzyme | |
| chr5 | *RP11-166A12.1* | 9 | 190 | 20 | | | | | |
| chr3 | *FAM157A* | 9 | 360 | 20 | AM157A/FAM157 | family with sequence similarity 157 member A | Other | other | |
| chr13 | *SPACA7* | 9 | 317 | 20 | SPACA7 | sperm acrosome associated 7 | Cytoplasm | other | |
| chr15 | *HDC* | 8 | 238 | 20 | HDC | histidine decarboxylase | Cytoplasm | enzyme | |
| chr16 | *HERC2P8* | 7 | 176 | 20 | HERC2P8 | hect domain and RLD 2 pseudogene 8 | Other | other | |
| chr12 | *GLTP* | 9 | 328 | 20 | GLTP | glycolipid transfer protein | Cytoplasm | transporter | |
| chr3 | *CMSS1* | 8 | 232 | 20 | CMSS1 | cms1 ribosomal small subunit homolog | Other | other | |
| chr2 | *AC018866.1* | 9 | 413 | 20 | | | | | |
| chr20 | *ACSS2* | 9 | 269 | 20 | ACSS2 | acyl-CoA synthetase short chain family member 2 | Cytoplasm | enzyme | |
| chr19 | *ZNF568* | 8 | 165 | 20 | ZNF568 | zinc finger protein 568 | Other | other | |
| chr10 | *CTA-109P11.4* | 9 | 221 | 20 | | | | | |
| chr10 | *FAM45A* | 8 | 222 | 20 | DENND10 | DENN domain containing 10 | Other | other | |
| chr9 | *RP11-379C10.1* | 9 | 299 | 20 | | | | | |
| chr6 | *GUCA1A* | 10 | 294 | 20 | GUCA1A | guanylate cyclase activator 1A | Cytoplasm | other | |
| chr1 | *TMEM50A* | 7 | 188 | 20 | TMEM50A | transmembrane protein 50A | Plasma Membrane | other | |
| chr2 | *SLC25A12* | 8 | 190 | 20 | SLC25A12 | solute carrier family 25 member | Cytoplasm | transporter | |
| chr19 | *RDH13* | 8 | 296 | 20 | RDH13 | retinol dehydrogenase 13 | Cytoplasm | enzyme | |
| chr20 | *TTPAL* | 8 | 343 | 20 | TTPAL | alpha tocopherol transfer protein | Other | other | |
| chr5 | *CLPTM1L* | 10 | 1068 | 20 | CLPTM1L | CLPTM1 like | Other | other | |
| chr1 | *FGR* | 8 | 353 | 20 | FGR | FGR proto-oncogene, Src family tyrosine kinase | Nucleus | kinase | cetuximab/vemurafenib, irinotecan/panitumumab/vemurafenib, cobimetinib/vemurafenib, trametinib/vemurafenib, dabrafenib/trametinib/vemurafenib, vemurafenib, crizotinib/vemurafenib, ipilimumab/vemurafe [illegible] |
| chr10 | *CEP164P1* | 9 | 312 | 20 | CEP164P1 | centrosomal protein 164 pseudogene 1 | Other | other | |
| chr2 | *RGPD8* | 8 | 331 | 20 | PD4 (includes oth | RANBP2 like and GRIP domain containing 5 | Nucleus | enzyme | |
| chr13 | *DLEU1* | 8 | 235 | 20 | DLEU1 | deleted in lymphocytic leukemia 1 | Other | other | |
| chr3 | *BFSP2* | 9 | 325 | 20 | BFSP2 | beaded filament structural protein | Cytoplasm | other | |
| chr1 | *UBAP2L* | 7 | 250 | 20 | UBAP2L | ubiquitin associated protein 2 like | Nucleus | other | |

| | | | | | | | | | |
|---|---|---|---|---|---|---|---|---|---|
| chr2 | *CACNB4* | 8 | 188 | 20 | CACNB4 | calcium voltage-gated channel auxiliary subunit beta 4 | Plasma Membrane | ion channel | verapamil, dexamethasone/rituximab/verapamil, mibefradil, nimodipine, |
| chr9 | *FAM154A* | 9 | 228 | 20 | SAXO1 | stabilizer of axonemal | Cytoplasm | other | |
| chr9 | *PTGES* | 9 | 442 | 20 | PTGES | prostaglandin E synthase | Cytoplasm | enzyme | |
| chr16 | *RP11-481J2.2* | 10 | 219 | 20 | | | | | |
| chr20 | *PIEZO1P2* | 10 | 386 | 20 | PIEZO1P2 | piezo type mechanosensitive ion channel component 1 | Other | other | |
| chr2 | *APLF* | 9 | 239 | 20 | APLF | aprataxin and PNKP like factor | Cytoplasm | enzyme | |
| chr1 | *SYNC* | 8 | 271 | 20 | SYNC | syncoilin, intermediate filament | Cytoplasm | other | |
| chr14 | *TTC6* | 8 | 235 | 20 | TTC6 | tetratricopeptide repeat domain 6 | Other | other | |
| chr10 | *SEC61A2* | 7 | 199 | 20 | SEC61A2 | SEC61 translocon alpha 2 | Cytoplasm | transporter | |
| chr20 | *RBM39* | 7 | 313 | 20 | RBM39 | RNA binding motif protein 39 | Nucleus | transcription regulator | |
| chr20 | *BCL2L1* | 9 | 500 | 20 | BCL2L1 | BCL2 like 1 | Cytoplasm | other | |
| chr7 | *KIAA0895* | 7 | 208 | 20 | KIAA0895 | KIAA0895 | Other | other | |
| chr1 | *RP6-206I17.1* | 9 | 217 | 20 | | | | | |
| chr16 | *EMC8* | 8 | 318 | 20 | EMC8 | ER membrane protein complex subunit 8 | Cytoplasm | other | |
| chr12 | *SART3* | 8 | 264 | 20 | SART3 | spliceosome associated factor 3, U4/U6 recycling protein | Nucleus | other | |
| chr13 | *TDRD3* | 7 | 190 | 20 | TDRD3 | tudor domain containing 3 | Nucleus | transcription regulator | |
| chr12 | *ZCCHC8* | 8 | 225 | 20 | ZCCHC8 | zinc finger CCHC-type containing | Nucleus | other | |
| chr10 | *RP11-95I16.2* | 9 | 324 | 20 | | | | | |
| chr3 | *AC069277.2* | 8 | 195 | 20 | | | | | |
| chr17 | *ITGA2B* | 8 | 335 | 20 | ITGA2B | integrin subunit alpha 2b | Plasma Membrane | transmembrane receptor | TP 9201, eptifibatide, abciximab, tirofiban |
| chr16 | *CA5AP1* | 8 | 301 | 20 | CA5AP1 | carbonic anhydrase 5A pseudogene 1 | Other | other | |
| chrX | *SMS* | 9 | 220 | 20 | SMS | spermine synthase | Cytoplasm | enzyme | (±)-2-hydroxyoleic |
| chr10 | *DNAJB12* | 9 | 645 | 20 | DNAJB12 | DnaJ heat shock protein family (Hsp40) member B12 | Cytoplasm | other | |
| chr7 | *PPIA* | 9 | 285 | 20 | PPIA | peptidylprolyl isomerase A | Cytoplasm | enzyme | cyclosporine A/sirolimus/tacrolimus, N-methyl-4-Ile-cyclosporin, alemtuzumab/cyclosporin A, cyclosporin A, cyclosporine A/tacrolimus, cyclosporin A/methotrexate |
| chr11 | *PAFAH1B2* | 8 | 202 | 19 | PAFAH1B2 | platelet activating factor acetylhydrolase 1b catalytic | Cytoplasm | enzyme | |
| chr11 | *MYBPC3* | 10 | 808 | 19 | MYBPC3 | myosin binding protein C, cardiac | Cytoplasm | other | |
| chr2 | *SNRNP200* | 10 | 402 | 19 | SNRNP200 | small nuclear ribonucleoprotein U5 subunit 200 | Nucleus | enzyme | |
| chr5 | *CTC-359M8.1* | 9 | 198 | 19 | | | | | |
| chr22 | *PRR5* | 8 | 368 | 19 | PRR5 | proline rich 5 | Cytoplasm | other | |
| chr2 | *AC027612.4* | 9 | 302 | 19 | | | | | |
| chr20 | *ANGPT4* | 7 | 256 | 19 | ANGPT4 | angiopoietin 4 | Extracellular Space | growth factor | |
| chr7 | *YWHAG* | 8 | 312 | 19 | YWHAG | tyrosine 3-monooxygenase/tryptophan 5-monooxygenase activation protein gamma | Cytoplasm | other | |
| chr10 | *PHYHIPL* | 8 | 248 | 19 | PHYHIPL | phytanoyl-CoA 2-hydroxylase interacting protein like | Cytoplasm | other | |
| chr16 | *CES4A* | 8 | 283 | 19 | CES4A | carboxylesterase 4A | Extracellular Space | enzyme | |
| chr16 | *DEF8* | 9 | 377 | 19 | DEF8 | differentially expressed in FDCP 8 homolog | Other | other | |
| chr2 | *AC027612.6* | 11 | 1334 | 19 | | | | | |
| chr20 | *COL9A3* | 10 | 739 | 19 | COL9A3 | collagen type IX alpha 3 chain | Extracellular Space | other | collagenase clostridium |
| chr10 | *CASC2* | 9 | 170 | 19 | CASC2 | cancer susceptibility 2 | Other | other | |
| chr16 | *RP11-449H11.1* | 8 | 220 | 19 | | | | | |
| chr1 | *AK2* | 9 | 207 | 19 | AK2 | adenylate kinase 2 | Cytoplasm | kinase | |
| chr20 | *RP5-1121H13.4* | 10 | 281 | 19 | | | | | |
| chr17 | *AC139099.4* | 10 | 377 | 19 | | | | | |
| chrX | *MAOA* | 9 | 247 | 19 | MAOA | monoamine oxidase A | Cytoplasm | enzyme | fenfluramine/phentermine, moclobemide, methamphetamine, isocarboxazid, CX-1370, benzphetamine, phentermine/topiramate, N-(2-indanyl)glycinamide, iproniazid, phentermine, tranylcypromine, procainamide, MAO-A inhibitor, ladostigil, |
| chr19 | *ATP13A1* | 10 | 482 | 19 | ATP13A1 | ATPase 13A1 | Extracellular Space | transporter | |
| chr11 | *ALDH3B1* | 10 | 558 | 19 | ALDH3B1 | aldehyde dehydrogenase 3 family member B1 | Cytoplasm | enzyme | |
| chr9 | *ACER2* | 9 | 204 | 19 | ACER2 | alkaline ceramidase 2 | Cytoplasm | enzyme | |
| chr11 | *UPK2* | 9 | 300 | 19 | UPK2 | uroplakin 2 | Plasma Membrane | other | |

| | | | | | | | | | |
|---|---|---|---|---|---|---|---|---|---|
| chr3 | *PTPN23* | 9 | 272 | 19 | PTPN23 | protein tyrosine phosphatase non-receptor type 23 | Cytoplasm | phosphatase | |
| chr2 | *RAPH1* | 7 | 157 | 19 | RAPH1 | Ras association (RalGDS/AF-6) and pleckstrin homology domains | Plasma Membrane | other | |
| chr12 | *SPRYD3* | 8 | 551 | 19 | SPRYD3 | SPRY domain containing 3 | Other | other | |
| chr2 | *C2orf76* | 8 | 208 | 19 | C2orf76 | chromosome 2 open reading frame 76 | Other | other | |
| chr3 | *DENND6A* | 7 | 175 | 19 | DENND6A | DENN domain containing 6A | Cytoplasm | other | |
| chr22 | *SEZ6L* | 7 | 221 | 19 | SEZ6L | seizure related 6 homolog like | Plasma Membrane | other | |
| chr1 | *SRSF4* | 9 | 265 | 19 | SRSF4 | serine and arginine rich splicing factor 4 | Nucleus | other | |
| chr7 | *FLNC* | 10 | 482 | 19 | FLNC | filamin C | Cytoplasm | other | |
| chr3 | *LRRC3B* | 7 | 194 | 19 | LRRC3B | leucine rich repeat containing 3B | Other | other | |
| chr2 | *PER2* | 8 | 313 | 19 | PER2 | period circadian regulator 2 | Nucleus | transcription regulator | |
| chr4 | *AF146191.4* | 9 | 406 | 19 | | | | | |
| chr1 | *VTCN1* | 10 | 242 | 19 | VTCN1 | V-set domain containing T cell activation inhibitor 1 | Plasma Membrane | other | FPA150 |
| chr7 | *RP11-3B12.2* | 8 | 185 | 19 | | | | | |
| chr16 | *CALB2* | 9 | 266 | 19 | CALB2 | calbindin 2 | Cytoplasm | other | |
| chr19 | *ZNF90* | 9 | 197 | 19 | ZNF90 | zinc finger protein 90 | Nucleus | transcription regulator | |
| chr17 | *GGA3* | 8 | 316 | 19 | GGA3 | golgi associated, gamma adaptin ear containing, ARF binding | Cytoplasm | transporter | |
| chr2 | *USP37* | 7 | 167 | 19 | USP37 | ubiquitin specific peptidase 37 | Nucleus | peptidase | |
| chr8 | *HR* | 9 | 491 | 19 | HR | HR lysine demethylase and nuclear receptor corepressor | Nucleus | transcription regulator | |
| chr4 | *PIGG* | 8 | 326 | 19 | PIGG | phosphatidylinositol glycan anchor biosynthesis class G | Cytoplasm | enzyme | |
| chr6 | *DSP* | 8 | 214 | 19 | DSP | desmoplakin | Plasma Membrane | other | |
| chr2 | *PUM2* | 7 | 160 | 19 | PUM2 | pumilio RNA binding family | Cytoplasm | other | |
| chr20 | *TOP1* | 7 | 242 | 19 | TOP1 | DNA topoisomerase I | Nucleus | enzyme | cyclophosphamide/topotecan/vincristine, topoisomerase I inhibitor, bevacizumab/topotecan, aflibercept/irinotecan, bevacizumab/capecitabine/irinotecan, simmitecan, paclitaxel/topotecan, irinotecan, ifosfamide/topotecan, bevacizumab/irinotecan/oxaliplatin, SN-38, bevacizumab/paclitaxel/topotecan, T 0128, bevacizumab/irinotecan, belotecan, 5-fluorouracil/irinotecan/oxaliplatin, capecitabine/cetuximab/irinotecan, BAY 56-2798 |
| chr9 | *PTGS1* | 9 | 442 | 19 | PTGS1 | prostaglandin-endoperoxide synthase 1 | Cytoplasm | enzyme | sulindac/tamoxifen, esomeprazole/naproxen, acetaminophen/oxycodone, piroxicam, salicylic acid, raloxifene/sulindac, 8Z,11Z,14Z-eicosatrienoic acid, fenoprofen, acetaminophen, aspirin/rivaroxaban/ticlopidine, phenacetin, balsalazide, aspirin/caffeine/orphenadrine, acetaminophen/hydrocodone, aspirin/dabigatran etexilate, ketorolac/phenylephrine, [illegible] |
| chr9 | *UNC13B* | 8 | 175 | 19 | UNC13B | unc-13 homolog B | Cytoplasm | other | |
| chr18 | *LINC00908* | 8 | 262 | 19 | LINC00908 | long intergenic non-protein coding RNA 908 | Other | other | |
| chr1 | *C1orf146* | 7 | 187 | 19 | C1orf146 | chromosome 1 open reading frame 146 | Other | other | |
| chr7 | *RBM28* | 8 | 211 | 19 | RBM28 | RNA binding motif protein 28 | Nucleus | other | |
| chr4 | *SLIT2* | 7 | 154 | 19 | SLIT2 | slit guidance ligand 2 | Extracellular Space | other | |
| chr18 | *CCBE1* | 8 | 184 | 19 | CCBE1 | collagen and calcium binding EGF domains 1 | Extracellular Space | other | |
| chr16 | *CTD-2336H13.2* | 9 | 175 | 19 | | | | | |
| chr8 | *PPP2CB* | 6 | 189 | 19 | PPP2CB | protein phosphatase 2 catalytic subunit beta | Cytoplasm | phosphatase | |

| | | | | | | | | | |
|---|---|---|---|---|---|---|---|---|---|
| chr6 | *ELOVL2* | 7 | 162 | 19 | ELOVL2 | ELOVL fatty acid elongase 2 | Cytoplasm | enzyme | |
| chr10 | *SMC3* | 7 | 202 | 19 | SMC3 | structural maintenance of chromosomes 3 | Nucleus | other | |
| chr15 | *POLG* | 8 | 441 | 19 | POLG | DNA polymerase gamma, catalytic subunit | Cytoplasm | enzyme | lamivudine/stavudine, stavudine, vidarabine, lamivudine/nelfinavir/stavudine, zalcitabine, |
| chrX | *CD99L2* | 8 | 225 | 19 | CD99L2 | CD99 molecule like 2 | Plasma Membrane | other | |
| chr1 | *SLC45A3* | 9 | 652 | 19 | SLC45A3 | solute carrier family 45 member 3 | Cytoplasm | transporter | |
| chr16 | *SLC38A7* | 9 | 299 | 19 | SLC38A7 | solute carrier family 38 member 7 | Plasma Membrane | transporter | |
| chr10 | *HIF1AN* | 8 | 190 | 19 | HIF1AN | hypoxia inducible factor 1 subunit alpha inhibitor | Nucleus | enzyme | |
| chr17 | *ABCA10* | 8 | 224 | 19 | ABCA10 | ATP binding cassette subfamily A member 10 | Other | transporter | |
| chr17 | *TAF15* | 6 | 213 | 19 | TAF15 | TATA-box binding protein associated factor 15 | Nucleus | other | |
| chr11 | *TTC17* | 8 | 169 | 19 | TTC17 | tetratricopeptide repeat domain | Cytoplasm | other | |
| chr19 | *CCDC155* | 8 | 312 | 19 | CCDC155 | coiled-coil domain containing 155 | Nucleus | other | |
| chr9 | *TRAF1* | 8 | 432 | 19 | TRAF1 | TNF receptor associated factor 1 | Cytoplasm | other | |
| chr5 | *ADRA1B* | 9 | 322 | 19 | ADRA1B | adrenoceptor alpha 1B | Plasma Membrane | G-protein coupled recepto | chlorpheniramine/methscopolamine/phenylephrine, apraclonidine, epinephrine/lidocaine, azatadine/pseudoephedrine, dobutamine, terazosine, quinidine, doxazosin, brexpiprazole, dihydroergotamine, guanethidine, tolazoline, brompheniramine/codeine/phenylpropanolamine, loratadine/pseudoephedrine, chlorpheniramine/ibuprofen/pseudoephedrine, brimonidine, dipivefrin, guanadrel, [illegible] |
| chr1 | *RP13-15E13.1* | 9 | 377 | 19 | | | | | |
| chr3 | *SOX2-OT* | 7 | 193 | 19 | SOX2-OT | SOX2 overlapping transcript | Other | other | |
| chr7 | *LINC00689* | 10 | 384 | 19 | LINC00689 | long intergenic non-protein coding RNA 689 | Other | other | |
| chr20 | *L3MBTL1* | 9 | 377 | 19 | L3MBTL1 | L3MBTL histone methyl-lysine binding protein 1 | Nucleus | other | |
| chr19 | *CTC-360P9.4* | 8 | 190 | 19 | | | | | |
| chr1 | *NBPF20* | 10 | 399 | 19 | PF1 (includes othe | NBPF member 11 | Other | other | |
| chr10 | *TDRD1* | 8 | 200 | 19 | TDRD1 | tudor domain containing 1 | Cytoplasm | other | |
| chr11 | *RPLP0P2* | 8 | 474 | 19 | RPLP0P2 | ribosomal protein lateral stalk subunit P0 pseudogene 2 | Other | other | |
| chr17 | *FMNL1* | 9 | 537 | 19 | FMNL1 | formin like 1 | Cytoplasm | other | |
| chr15 | *WDR76* | 8 | 181 | 19 | WDR76 | WD repeat domain 76 | Nucleus | other | |
| chr1 | *SLC25A44* | 9 | 334 | 19 | SLC25A44 | solute carrier family 25 member | Cytoplasm | transporter | |
| chr12 | *CBX5* | 9 | 337 | 19 | CBX5 | chromobox 5 | Nucleus | transcription regulator | |
| chr20 | *MAVS* | 8 | 263 | 19 | MAVS | mitochondrial antiviral signaling protein | Cytoplasm | other | |
| chr10 | *RASSF4* | 9 | 633 | 19 | RASSF4 | Ras association domain family member 4 | Other | other | |
| chr5 | *POLK* | 8 | 193 | 19 | POLK | DNA polymerase kappa | Nucleus | enzyme | |
| chr21 | *LSS* | 8 | 370 | 19 | LSS | lanosterol synthase | Cytoplasm | enzyme | oxiconazole |

| | | | | | | | | | |
|---|---|---|---|---|---|---|---|---|---|
| chr14 | *ESR2* | 7 | 165 | 19 | ESR2 | estrogen receptor 2 | Nucleus | d-dependent nuclear rec | sulindac/tamoxifen, ethinyl estradiol/ethynodiol diacetate, lasofoxifene, raloxifene/sulindac, genistein, diethylstilbestrol, anastrozole/fulvestrant/goserelin/tamoxifen, estradiol valerate, beta-estradiol, anastrozole/fulvestrant/tamoxifen, trilostane, anastrozole/exemestane/fulvestrant/tamoxifen, goserelin/tamoxifen, exemestane/tamoxifen, ethinyl estradiol/norgestimate, 17-alpha- [illegible] |
| chrX | *PLAC1* | 8 | 171 | 19 | PLAC1 | placenta enriched 1 | Plasma Membrane | other | |
| chr17 | *C17orf67* | 8 | 229 | 19 | C17orf67 | chromosome 17 open reading frame 67 | Other | other | |
| chr1 | *RBBP4* | 7 | 226 | 19 | RBBP4 | RB binding protein 4, chromatin remodeling factor | Nucleus | enzyme | |
| chr11 | *IMMP1L* | 8 | 247 | 19 | IMMP1L | inner mitochondrial membrane peptidase subunit 1 | Cytoplasm | other | |
| chr19 | *NLRP11* | 8 | 216 | 19 | NLRP11 | NLR family pyrin domain containing 11 | Extracellular Space | other | |
| chrX | *ZNF157* | 8 | 182 | 19 | ZNF157 | zinc finger protein 157 | Nucleus | transcription regulator | |
| chr4 | *RP11-310I9.1* | 8 | 192 | 19 | | | | | |
| chr14 | *SLC25A21* | 7 | 172 | 19 | SLC25A21 | solute carrier family 25 member | Cytoplasm | transporter | |
| chr1 | *PTGFR* | 9 | 235 | 19 | PTGFR | prostaglandin F receptor | Plasma Membrane | G-protein coupled recepto | timolol/travoprost, bimatoprost/timolol, latanoprost/netarsudil, tafluprost, dinoprost, latanoprost, bimatoprost, latanoprost/timolol, isopropyl unoprostone, |
| chr9 | *URM1* | 9 | 442 | 19 | URM1 | ubiquitin related modifier 1 | Cytoplasm | other | |
| chr1 | *CELA2B* | 8 | 294 | 19 | CELA2B | chymotrypsin like elastase 2B | Extracellular Space | peptidase | |
| chr20 | *BPIFB5P* | 8 | 453 | 19 | BPIFB5P | BPI fold containing family B member 5, pseudogene | Other | other | |
| chr9 | *ANKS6* | 7 | 279 | 19 | ANKS6 | ankyrin repeat and sterile alpha motif domain containing 6 | Cytoplasm | other | |
| chr11 | *BACE1* | 9 | 228 | 19 | BACE1 | beta-secretase 1 | Cytoplasm | peptidase | atabecestat, MK-8931, elenbecestat, lanabecestat |
| chr19 | *SPC24* | 8 | 303 | 19 | SPC24 | SPC24 component of NDC80 kinetochore complex | Cytoplasm | other | |
| chrX | *CCNB3* | 9 | 203 | 19 | CCNB3 | cyclin B3 | Nucleus | other | |
| chr1 | *SCCPDH* | 8 | 203 | 19 | SCCPDH | saccharopine dehydrogenase (putative) | Cytoplasm | other | |
| chr17 | *EFCAB3* | 8 | 173 | 19 | EFCAB3 | EF-hand calcium binding domain | Other | other | |
| chr1 | *DAP3P1* | 8 | 306 | 19 | DAP3P1 | death associated protein 3 pseudogene 1 | Other | other | |
| chr1 | *C1orf106* | 9 | 442 | 19 | INAVA | innate immunity activator | Other | other | |
| chr7 | *AC073316.1* | 8 | 278 | 19 | | | | | |
| chrX | *PLCXD1* | 8 | 323 | 19 | PLCXD1 | phosphatidylinositol specific phospholipase C X domain containing 1 | Other | enzyme | |
| chr3 | *DZIP1L* | 10 | 392 | 19 | DZIP1L | DAZ interacting zinc finger protein 1 like | Extracellular Space | other | |
| chr1 | *ATAD3A* | 10 | 396 | 19 | ATAD3A | ATPase family AAA domain containing 3A | Cytoplasm | other | |
| chr12 | *LGR5* | 8 | 179 | 19 | LGR5 | leucine rich repeat containing G protein-coupled receptor 5 | Plasma Membrane | transmembrane receptor | MCLA-158, BNC101 |
| chr6 | *FAM120B* | 7 | 236 | 19 | FAM120B | family with sequence similarity | Nucleus | other | |
| chr9 | *MRRF* | 8 | 212 | 19 | MRRF | mitochondrial ribosome recycling factor | Cytoplasm | other | |
| chr1 | *RP11-15I11.2* | 8 | 303 | 19 | | | | | |
| chr2 | *AC098828.2* | 8 | 233 | 19 | | | | | |
| chr20 | *DSN1* | 9 | 258 | 19 | DSN1 | DSN1 component of MIS12 kinetochore complex | Nucleus | other | |
| chr20 | *TLDC2* | 8 | 439 | 19 | TLDC2 | TBC/LysM-associated domain containing 2 | Other | other | |
| chr15 | *PPIP5K1* | 8 | 190 | 19 | PPIP5K1 | diphosphoinositol pentakisphosphate kinase 1 | Nucleus | phosphatase | |
| chr7 | *AC005682.8* | 7 | 243 | 19 | | | | | |
| chr17 | *CRLF3* | 8 | 199 | 19 | CRLF3 | cytokine receptor like factor 3 | Cytoplasm | transcription regulator | |
| chr6 | *C6orf99* | 7 | 203 | 19 | C6orf99 | chromosome 6 open reading frame 99 | Other | other | |

| | | | | | | | | | |
|---|---|---|---|---|---|---|---|---|---|
| chr17 | *TBKBP1* | 9 | 457 | 19 | TBKBP1 | TBK1 binding protein 1 | Other | other | |
| chr7 | *AC004112.4* | 8 | 177 | 19 | | | | | |
| chr19 | *TNNT1* | 8 | 314 | 19 | TNNT1 | troponin T1, slow skeletal type | Cytoplasm | other | |
| chr16 | *SRRM2-AS1* | 8 | 312 | 19 | SRRM2-AS1 | SRRM2 antisense RNA 1 | Other | other | |
| chr11 | *CTC-497E21.5* | 10 | 261 | 19 | | | | | |
| chr9 | *TSC1* | 9 | 310 | 19 | TSC1 | TSC complex subunit 1 | Cytoplasm | other | |
| chr22 | *CITF22-49D8.1* | 7 | 279 | 19 | | | | | |
| chr7 | *FAM220A* | 8 | 310 | 19 | FAM220A | family with sequence similarity 220 member A | Nucleus | other | |
| chr12 | *LOH12CR1* | 7 | 169 | 19 | BORCS5 | BLOC-1 related complex subunit | Cytoplasm | other | |
| chr17 | *SOCS7* | 9 | 215 | 19 | SOCS7 | suppressor of cytokine signaling | Cytoplasm | other | |
| chr11 | *PCSK7* | 8 | 363 | 19 | PCSK7 | proprotein convertase subtilisin/kexin type 7 | Cytoplasm | peptidase | |
| chr20 | *BPI* | 9 | 291 | 19 | BPI | bactericidal permeability increasing protein | Plasma Membrane | transporter | |
| chr5 | *CTC-254B4.1* | 9 | 195 | 19 | | | | | |
| chr8 | *RP11-156K13.1* | 8 | 178 | 19 | | | | | |
| chrX | *HDAC6* | 8 | 404 | 19 | HDAC6 | histone deacetylase 6 | Nucleus | transcription regulator | ricolinostat, pyroxamide, bortezomib/vorinostat, tributyrin, trametinib/vorinostat, KA2507, belinostat, |
| chr7 | *MDH2* | 8 | 401 | 19 | MDH2 | malate dehydrogenase 2 | Cytoplasm | enzyme | |
| chr2 | *RP11-76I14.1* | 8 | 231 | 19 | | | | | |
| chr11 | *USP28* | 8 | 204 | 19 | USP28 | ubiquitin specific peptidase 28 | Nucleus | peptidase | |
| chr13 | *TM9SF2* | 8 | 232 | 19 | TM9SF2 | transmembrane 9 superfamily member 2 | Plasma Membrane | transporter | |
| chr8 | *CHMP7* | 8 | 320 | 19 | CHMP7 | charged multivesicular body | Cytoplasm | other | |
| chr19 | *ZFP82* | 8 | 160 | 19 | ZFP82 | ZFP82 zinc finger protein | Nucleus | transcription regulator | |
| chr3 | *EFCAB12* | 9 | 428 | 19 | EFCAB12 | EF-hand calcium binding domain | Other | other | |
| chr11 | *F2* | 8 | 416 | 19 | F2 | coagulation factor II, thrombin | Extracellular Space | peptidase | aspirin/bivalirudin, enoxaparin, dabigatran, Fibrinogen, dabigatran etexilate, bivalirudin, lepirudin, ximelagatran, argatroban, ulinastatin, thrombin inhibitor, |
| chr7 | *RP11-3B12.1* | 8 | 199 | 19 | | | | | |
| chr20 | *MANBAL* | 9 | 364 | 19 | MANBAL | mannosidase beta like | Other | other | |
| chr15 | *RP11-97O12.7* | 8 | 331 | 19 | | | | | |
| chr5 | *STK32A* | 8 | 160 | 19 | STK32A | serine/threonine kinase 32A | Other | kinase | |
| chr1 | *FRRS1* | 8 | 236 | 19 | FRRS1 | ferric chelate reductase 1 | Plasma Membrane | transmembrane recepto | |
| chr13 | *GRK1* | 8 | 560 | 19 | GRK1 | G protein-coupled receptor | Plasma Membrane | kinase | |
| chr7 | *KLHL7* | 8 | 273 | 19 | KLHL7 | kelch like family member 7 | Nucleus | other | |
| chr5 | *RARS* | 9 | 282 | 19 | RARS1 | arginyl-tRNA synthetase 1 | Cytoplasm | enzyme | |
| chr20 | *RP11-410N8.4* | 8 | 443 | 19 | | | | | |
| chr12 | *ACSS3* | 7 | 218 | 19 | ACSS3 | acyl-CoA synthetase short chain family member 3 | Cytoplasm | enzyme | |
| chr17 | *RP11-218F4.1* | 8 | 245 | 19 | | | | | |
| chr19 | *TMEM161A* | 8 | 294 | 19 | TMEM161A | transmembrane protein 161A | Other | other | |
| chr19 | *KCNN4* | 8 | 313 | 19 | KCNN4 | potassium calcium-activated channel subfamily N member 4 | Plasma Membrane | ion channel | clotrimazole, betamethasone/clotrimazole, senicapoc |
| chr3 | *CEP97* | 9 | 326 | 19 | CEP97 | centrosomal protein 97 | Cytoplasm | other | |
| chr10 | *OPN4* | 8 | 396 | 19 | OPN4 | opsin 4 | Plasma Membrane | -protein coupled recepto | |
| chr9 | *RP11-15B24.5* | 9 | 159 | 19 | | | | | |
| chr6 | *VEGFA* | 8 | 438 | 19 | VEGFA | vascular endothelial growth factor A | Extracellular Space | growth factor | bevacizumab/capecitabine, bevacizumab/paclitaxel/topotecan, bevacizumab/temozolomide, bevacizumab/docetaxel/prednisone/thalidomide/zoledronic acid, bevacizumab-IRDye 800CW, bevacizumab/5-fluorouracil/irinotecan/oxaliplatin, bevacizumab/irinotecan, bevacizumab/sorafenib, bevacizumab/erlotinib, brolucizumab, bevacizumab, bevacizumab/gemcitabine, [illegible] |
| chr2 | *DYNC2LI1* | 7 | 211 | 19 | DYNC2LI1 | dynein cytoplasmic 2 light intermediate chain 1 | Cytoplasm | other | |
| chr20 | *CABLES2* | 10 | 777 | 19 | CABLES2 | Cdk5 and Abl enzyme substrate 2 | Other | other | |
| chr1 | *SPATA6* | 9 | 230 | 19 | SPATA6 | spermatogenesis associated 6 | Cytoplasm | other | |

| | | | | | | | | | |
|---|---|---|---|---|---|---|---|---|---|
| chr8 | *TNFRSF10C* | 8 | 344 | 19 | TNFRSF10C | TNF receptor superfamily member 10c | Plasma Membrane | transmembrane receptor | |
| chr19 | *GNA11* | 9 | 417 | 19 | GNA11 | G protein subunit alpha 11 | Plasma Membrane | enzyme | |
| chr1 | *ZNF644* | 7 | 178 | 19 | ZNF644 | zinc finger protein 644 | Nucleus | transcription regulator | |
| chr3 | *SLC38A3* | 8 | 467 | 19 | SLC38A3 | solute carrier family 38 member 3 | Plasma Membrane | transporter | |
| chr19 | *NLRP9* | 8 | 217 | 19 | NLRP9 | NLR family pyrin domain | Cytoplasm | other | |
| chr16 | *RP5-991G20.1* | 8 | 178 | 19 | | | | | |
| chr9 | *BRD3* | 10 | 477 | 19 | BRD3 | bromodomain containing 3 | Nucleus | kinase | BI 894999 |
| chr1 | *LINC00623* | 7 | 250 | 19 | LINC00623 | long intergenic non-protein coding RNA 623 | Other | other | |
| chr19 | *FCAR* | 8 | 264 | 19 | FCAR | Fc fragment of IgA receptor | Plasma Membrane | transmembrane receptor | |
| chr8 | *STAU2* | 7 | 159 | 19 | STAU2 | staufen double-stranded RNA binding protein 2 | Cytoplasm | other | |
| chr10 | *DUPD1* | 9 | 417 | 19 | DUPD1 | dual specificity phosphatase and pro isomerase domain containing | Other | enzyme | |
| chr3 | *MYL3* | 8 | 392 | 19 | MYL3 | myosin light chain 3 | Cytoplasm | other | |
| chr3 | *P4HTM* | 9 | 335 | 19 | P4HTM | prolyl 4-hydroxylase, | Cytoplasm | enzyme | |
| chr12 | *DYNLL1* | 9 | 241 | 19 | DYNLL1 | dynein light chain LC8-type 1 | Cytoplasm | other | |
| chr5 | *SLC6A18* | 10 | 859 | 19 | SLC6A18 | solute carrier family 6 member 18 | Plasma Membrane | transporter | |
| chr16 | *RP11-1166P10.6* | 8 | 823 | 19 | | | | | |
| chr7 | *STAG3* | 8 | 301 | 19 | STAG3 | stromal antigen 3 | Nucleus | other | |
| chr15 | *CTD-2323K18.1* | 8 | 299 | 19 | | | | | |
| chr1 | *NUP133* | 9 | 219 | 19 | NUP133 | nucleoporin 133 | Nucleus | other | |
| chr22 | *RP1-272J12.1* | 7 | 192 | 19 | | | | | |
| chr5 | *DNAJC18* | 8 | 207 | 19 | DNAJC18 | DnaJ heat shock protein family (Hsp40) member C18 | Other | enzyme | |
| chr19 | *AC005780.1* | 9 | 174 | 19 | | | | | |
| chr7 | *ZMIZ2* | 9 | 466 | 19 | ZMIZ2 | zinc finger MIZ-type containing 2 | Nucleus | transcription regulator | |
| chr8 | *TOX* | 8 | 205 | 19 | TOX | thymocyte selection associated high mobility group box | Nucleus | transcription regulator | |
| chr3 | *MAATS1* | 8 | 247 | 19 | MAATS1 | MYCBP associated and testis expressed 1 | Cytoplasm | other | |
| chr11 | *RP11-744N12.3* | 9 | 240 | 19 | | | | | |
| chr7 | *SUN3* | 6 | 221 | 19 | SUN3 | Sad1 and UNC84 domain containing 3 | Nucleus | other | |
| chr6 | *RPS10-NUDT3* | 8 | 181 | 19 | RPS10-NUDT3 | RPS10-NUDT3 readthrough | Cytoplasm | other | |
| chr16 | *PDPK2* | 9 | 440 | 19 | | | | | |
| chr2 | *RAB10* | 7 | 173 | 19 | RAB10 | RAB10, member RAS oncogene family | Cytoplasm | enzyme | |
| chr1 | *TOR3A* | 8 | 402 | 19 | TOR3A | torsin family 3 member A | Cytoplasm | enzyme | |
| chr17 | *GPR179* | 9 | 385 | 19 | GPR179 | G protein-coupled receptor 179 | Plasma Membrane | G-protein coupled recepto | |
| chr20 | *ATRN* | 7 | 173 | 19 | ATRN | attractin | Extracellular Space | other | |
| chr6 | *SERPINB6* | 8 | 355 | 19 | SERPINB6 | serpin family B member 6 | Cytoplasm | other | |
| chr7 | *ABCB8* | 9 | 345 | 19 | ABCB8 | ATP binding cassette subfamily B member 8 | Cytoplasm | transporter | |
| chr20 | *ELMO2* | 9 | 322 | 19 | ELMO2 | engulfment and cell motility 2 | Cytoplasm | other | |
| chr14 | *KCNH5* | 7 | 152 | 19 | KCNH5 | potassium voltage-gated channel subfamily H member 5 | Plasma Membrane | ion channel | |
| chr17 | *FLOT2* | 8 | 310 | 19 | FLOT2 | flotillin 2 | Plasma Membrane | other | |
| chr3 | *CCDC66* | 7 | 226 | 19 | CCDC66 | coiled-coil domain containing 66 | Cytoplasm | other | |
| chr11 | *SIDT2* | 8 | 345 | 19 | SIDT2 | SID1 transmembrane family member 2 | Cytoplasm | transporter | |
| chr17 | *FN3KRP* | 8 | 301 | 19 | FN3KRP | fructosamine 3 kinase related | Cytoplasm | kinase | |
| chr19 | *KCTD15* | 8 | 336 | 19 | KCTD15 | potassium channel tetramerization domain containing | Other | ion channel | |
| chr16 | *TOX3* | 8 | 204 | 19 | TOX3 | TOX high mobility group box family member 3 | Nucleus | transcription regulator | |
| chr7 | *ZNF425* | 8 | 230 | 19 | ZNF425 | zinc finger protein 425 | Nucleus | other | |
| chr17 | *CTC-297N7.5* | 9 | 169 | 19 | | | | | |
| chr5 | *RP11-375B1.1* | 8 | 360 | 19 | | | | | |
| chr19 | *NR1H2* | 9 | 294 | 19 | NR1H2 | nuclear receptor subfamily 1 group H member 2 | Nucleus | d-dependent nuclear rec | RGX-104 |
| chr22 | *APOBEC3G* | 7 | 242 | 19 | APOBEC3G | apolipoprotein B mRNA editing enzyme catalytic subunit 3G | Nucleus | enzyme | |
| chr19 | *EMR4P* | 8 | 180 | 19 | ADGRE4P | adhesion G protein-coupled receptor E4, pseudogene | Plasma Membrane | other | |
| chr16 | *ALDOA* | 8 | 341 | 19 | ALDOA | aldolase, fructose-bisphosphate | Cytoplasm | enzyme | |
| chr11 | *KIAA1731* | 8 | 420 | 19 | CEP295 | centrosomal protein 295 | Cytoplasm | other | |
| chr2 | *COLEC11* | 8 | 365 | 19 | COLEC11 | collectin subfamily member 11 | Extracellular Space | other | |
| chrX | *GK* | 7 | 173 | 19 | GK | glycerol kinase | Cytoplasm | kinase | |
| chr11 | *GDPD4* | 8 | 250 | 19 | GDPD4 | glycerophosphodiester phosphodiesterase domain containing 4 | Other | kinase | |
| chr19 | *ZNF850* | 8 | 150 | 19 | ZNF850 | zinc finger protein 850 | Other | other | |
| chr15 | *FAN1* | 7 | 257 | 19 | FAN1 | FANCD2 and FANCI associated nuclease 1 | Nucleus | enzyme | |
| chr19 | *SAFB* | 7 | 233 | 19 | SAFB | scaffold attachment factor B | Nucleus | other | |
| chr19 | *CEACAM7* | 9 | 341 | 19 | CEACAM7 | CEA cell adhesion molecule 7 | Plasma Membrane | other | |
| chr3 | *PFN2* | 9 | 249 | 19 | PFN2 | profilin 2 | Cytoplasm | enzyme | |
| chr5 | *FABP6* | 9 | 262 | 19 | FABP6 | fatty acid binding protein 6 | Cytoplasm | transporter | |
| chr19 | *SIRT2* | 9 | 281 | 19 | SIRT2 | sirtuin 2 | Nucleus | transcription regulator | |
| chr21 | *TEKT4P2* | 6 | 343 | 19 | TEKT4P2 | tektin 4 pseudogene 2 | Other | other | |
| chr3 | *TGM4* | 8 | 256 | 19 | TGM4 | transglutaminase 4 | Extracellular Space | enzyme | |
| chr11 | *KCNJ5* | 8 | 369 | 19 | KCNJ5 | potassium inwardly rectifying channel subfamily J member 5 | Plasma Membrane | ion channel | minoxidil, amiodarone, |
| chr4 | *LIN54* | 7 | 154 | 19 | LIN54 | lin-54 DREAM MuvB core complex component | Nucleus | other | |
| chr7 | *AC072061.2* | 7 | 231 | 19 | | | | | |

| | | | | | | | | | |
|---|---|---|---|---|---|---|---|---|---|
| chr17 | *ST6GALNAC1* | 8 | 255 | 19 | ST6GALNAC1 | ST6 N-acetylgalactosaminide alpha-2,6-sialyltransferase 1 | Cytoplasm | enzyme | |
| chr17 | *C17orf99* | 8 | 344 | 19 | C17orf99 | chromosome 17 open reading frame 99 | Extracellular Space | other | |
| chr12 | *RNF41* | 8 | 281 | 19 | RNF41 | ring finger protein 41 | Cytoplasm | enzyme | |
| chr13 | *TPT1-AS1* | 9 | 208 | 19 | TPT1-AS1 | TPT1 antisense RNA 1 | Other | other | |
| chr10 | *C10orf107* | 9 | 277 | 19 | CABCOCO1 | ciliary associated calcium binding coiled-coil 1 | Cytoplasm | other | |
| chr8 | *WHSC1L1* | 7 | 204 | 19 | NSD3 | nuclear receptor binding SET domain protein 3 | Nucleus | enzyme | |
| chr19 | *ZNF793* | 8 | 187 | 19 | ZNF793 | zinc finger protein 793 | Other | other | |
| chr6 | *MDFI* | 8 | 400 | 19 | MDFI | MyoD family inhibitor | Cytoplasm | other | |
| chr5 | *ARSI* | 8 | 267 | 19 | ARSI | arylsulfatase family member I | Extracellular Space | enzyme | |
| chr16 | *FAM192A* | 7 | 181 | 19 | FAM192A | family with sequence similarity 192 member A | Nucleus | other | |
| chr17 | *RP11-304F15.6* | 9 | 384 | 19 | | | | | |
| chr18 | *KIAA1328* | 7 | 198 | 19 | KIAA1328 | KIAA1328 | Other | other | |
| chrX | *TAB3* | 7 | 171 | 19 | TAB3 | TGF-beta activated kinase 1 (MAP3K7) binding protein 3 | Cytoplasm | other | |
| chr2 | *VAMP8* | 8 | 250 | 19 | VAMP8 | vesicle associated membrane protein 8 | Plasma Membrane | transporter | |
| chr17 | *AC087499.7* | 7 | 186 | 19 | | | | | |
| chr5 | *FAM153B* | 8 | 315 | 19 | AM153A/FAM153 | family with sequence similarity 153 member B | Other | other | |
| chr6 | *BEND6* | 7 | 165 | 19 | BEND6 | BEN domain containing 6 | Nucleus | transcription regulator | |
| chr19 | *OR7A5* | 8 | 161 | 19 | OR7A5 | olfactory receptor family 7 subfamily A member 5 | Plasma Membrane | i-protein coupled recepto | |
| chr5 | *VDAC1* | 8 | 271 | 19 | VDAC1 | voltage dependent anion channel | Cytoplasm | ion channel | |
| chr14 | *VRTN* | 8 | 224 | 19 | VRTN | vertebrae development | Other | other | |
| chr13 | *STARD13* | 7 | 178 | 19 | STARD13 | StAR related lipid transfer domain containing 13 | Cytoplasm | other | |
| chr14 | *CDKL1* | 7 | 195 | 19 | CDKL1 | cyclin dependent kinase like 1 | Nucleus | kinase | |
| chr8 | *RP11-452N4.1* | 9 | 185 | 19 | | | | | |
| chr11 | *AP000487.5* | 7 | 264 | 19 | | | | | |
| chr4 | *NAA15* | 6 | 157 | 19 | NAA15 | N(alpha)-acetyltransferase 15, NatA auxiliary subunit | Nucleus | transcription regulator | |
| chr17 | *AFMID* | 8 | 267 | 19 | AFMID | arylformamidase | Nucleus | enzyme | |
| chr3 | *RP11-694I15.7* | 8 | 295 | 19 | | | | | |
| chr9 | *FXN* | 8 | 230 | 19 | FXN | frataxin | Cytoplasm | kinase | |
| chr2 | *AC012363.4* | 9 | 341 | 19 | | | | | |
| chr9 | *UHRF2* | 7 | 177 | 19 | UHRF2 | ubiquitin like with PHD and ring finger domains 2 | Nucleus | enzyme | |
| chr8 | *MICU3* | 8 | 163 | 19 | MICU3 | mitochondrial calcium uptake family member 3 | Cytoplasm | other | |
| chr7 | *HGF* | 7 | 189 | 19 | HGF | hepatocyte growth factor | Extracellular Space | growth factor | SCH 900105, MP0250, rilotumumab, bevacizumab/rilotum |
| chr5 | *SLC22A5* | 8 | 233 | 19 | SLC22A5 | solute carrier family 22 member 5 | Plasma Membrane | transporter | |
| chr3 | *AC090505.6* | 7 | 195 | 19 | | | | | |
| chr18 | *EPG5* | 7 | 192 | 19 | EPG5 | ectopic P-granules autophagy protein 5 homolog | Cytoplasm | other | |
| chr5 | *HK3* | 9 | 392 | 19 | HK3 | hexokinase 3 | Cytoplasm | kinase | |
| chr11 | *RP11-22P4.1* | 7 | 241 | 19 | | | | | |
| chr1 | *PER3* | 7 | 181 | 19 | PER3 | period circadian regulator 3 | Nucleus | other | |
| chr8 | *ADCK5* | 8 | 518 | 19 | ADCK5 | aarF domain containing kinase 5 | Cytoplasm | kinase | |
| chr17 | *ST6GALNAC2* | 8 | 344 | 19 | ST6GALNAC2 | ST6 N-acetylgalactosaminide alpha-2,6-sialyltransferase 2 | Cytoplasm | enzyme | |
| chr2 | *MARCO* | 9 | 336 | 19 | MARCO | macrophage receptor with collagenous structure | Plasma Membrane | transmembrane recepto | |
| chr6 | *GCLC* | 8 | 193 | 19 | GCLC | glutamate-cysteine ligase catalytic subunit | Cytoplasm | enzyme | |
| chr2 | *PDE1A* | 7 | 181 | 19 | PDE1A | phosphodiesterase 1A | Cytoplasm | enzyme | |
| chr3 | *WDR82* | 9 | 240 | 19 | WDR82 | WD repeat domain 82 | Nucleus | other | |
| chr7 | *GUSB* | 8 | 290 | 18 | GUSB | glucuronidase beta | Cytoplasm | enzyme | |
| chrX | *DDX3X* | 8 | 239 | 18 | DDX3X | DEAD-box helicase 3 X-linked | Cytoplasm | enzyme | |
| chr5 | *PRDM6* | 8 | 218 | 18 | PRDM6 | PR/SET domain 6 | Nucleus | transcription regulator | |
| chr1 | *MRPS21* | 8 | 293 | 18 | MRPS21 | mitochondrial ribosomal protein | Cytoplasm | other | |
| chr12 | *IPO8* | 6 | 244 | 18 | IPO8 | importin 8 | Nucleus | transporter | |
| chr2 | *ABCG8* | 9 | 390 | 18 | ABCG8 | ATP binding cassette subfamily G member 8 | Plasma Membrane | transporter | |
| chr2 | *AC092619.1* | 8 | 249 | 18 | | | | | |
| chr12 | *RP11-497G19.1* | 8 | 248 | 18 | | | | | |
| chr16 | *MMP15* | 8 | 478 | 18 | MMP15 | matrix metallopeptidase 15 | Extracellular Space | peptidase | marimastat |
| chr4 | *GRIA2* | 8 | 207 | 18 | GRIA2 | glutamate ionotropic receptor AMPA type subunit 2 | Plasma Membrane | ion channel | tezampanel, farampator, LY451395, |
| chr15 | *C15orf41* | 8 | 182 | 18 | C15orf41 | chromosome 15 open reading frame 41 | Other | other | |

| | | | | | | | | | |
|---|---|---|---|---|---|---|---|---|---|
| chr7 | *IMPDH1* | 8 | 355 | 18 | IMPDH1 | inosine monophosphate dehydrogenase 1 | Cytoplasm | enzyme | pegintron/ribavirin, interferon alfacon-1/ribavirin, cytarabine/daunorubicin/thioguanine, IFNA2B/ribavirin, ribavirin, mycophenolate mofetil, PEG-interferon alfa-2a/ribavirin, VX-944, mycophenolate mofetil/prednisone, methylprednisolone/mycophenolate mofetil, 6-mercaptopurine/prednisone/thioguanine, cytarabine/thioguanine, mycophenolic acid |
| chr12 | *GRIN2B* | 8 | 163 | 18 | GRIN2B | glutamate ionotropic receptor NMDA type subunit 2B | Plasma Membrane | ion channel | dextromethorphan/morphine, ifenprodil, nebostinel, besonprodil, delucemine, 1-aminocyclopropane-1-carboxylic acid, orphenadrine, dextromethorphan/promethazine, neramexane, felbamate, chlorpheniramine/dextromethorphan/phenylephrine, memantine, donepezil/memantine, N-(2-indanyl)glycinamide, tenocyclidine, dextromethorphan, cycloserine, UK-240455, brompheniramine/de[illegible] |
| chr13 | *UPF3A* | 8 | 349 | 18 | UPF3A | UPF3A regulator of nonsense mediated mRNA decay | Nucleus | transporter | |
| chr3 | *AC024560.3* | 7 | 209 | 18 | | | | | |
| chr5 | *CTD-2201E18.3* | 9 | 692 | 18 | | | | | |
| chr9 | *TRUB2* | 8 | 391 | 18 | TRUB2 | TruB pseudouridine synthase family member 2 | Cytoplasm | enzyme | |
| chr12 | *TBC1D30* | 9 | 187 | 18 | TBC1D30 | TBC1 domain family member 30 | Cytoplasm | other | |
| chr8 | *RP11-1134I14.2* | 6 | 221 | 18 | | | | | |
| chr16 | *METTL22* | 8 | 310 | 18 | METTL22 | methyltransferase like 22 | Nucleus | enzyme | |
| chrX | *CD99* | 7 | 214 | 18 | CD99 | CD99 molecule (Xg blood group) | Plasma Membrane | other | |
| chr20 | *MAP1LC3A* | 8 | 411 | 18 | MAP1LC3A | microtubule associated protein 1 light chain 3 alpha | Cytoplasm | other | |
| chr16 | *RP11-42I10.1* | 9 | 207 | 18 | | | | | |
| chr11 | *PRMT3* | 8 | 206 | 18 | PRMT3 | protein arginine | Nucleus | enzyme | |
| chr5 | *SMN2* | 7 | 210 | 18 | SMN1/SMN2 | survival of motor neuron 1, | Nucleus | other | nusinersen |
| chr12 | *RP11-161H23.9* | 8 | 262 | 18 | | | | | |
| chr9 | *RP11-180I4.2* | 7 | 206 | 18 | | | | | |
| chr10 | *MYO3A* | 7 | 169 | 18 | MYO3A | myosin IIIA | Cytoplasm | kinase | |
| chr12 | *PTPRB* | 9 | 170 | 18 | PTPRB | protein tyrosine phosphatase receptor type B | Plasma Membrane | phosphatase | |
| chr2 | *D2HGDH* | 9 | 412 | 18 | D2HGDH | D-2-hydroxyglutarate | Cytoplasm | enzyme | |
| chr16 | *NETO2* | 6 | 195 | 18 | NETO2 | neuropilin and tolloid like 2 | Plasma Membrane | other | |
| chr15 | *BMF* | 8 | 349 | 18 | BMF | Bcl2 modifying factor | Cytoplasm | other | |
| chr7 | *NOM1* | 8 | 291 | 18 | NOM1 | nucleolar protein with MIF4G domain 1 | Nucleus | other | |
| chr12 | *RIC8B* | 7 | 179 | 18 | RIC8B | RIC8 guanine nucleotide exchange factor B | Cytoplasm | other | |
| chr10 | *GUCY2GP* | 8 | 226 | 18 | GUCY2GP | guanylate cyclase 2G, | Other | other | |
| chr7 | *EZH2* | 8 | 175 | 18 | EZH2 | enhancer of zeste 2 polycomb repressive complex 2 subunit | Nucleus | transcription regulator | SHR2554, GSK2816126, tazemetostat, CPI-0209, PF-06821497, CPI-1205, |
| chrX | *L1CAM* | 9 | 336 | 18 | L1CAM | L1 cell adhesion molecule | Plasma Membrane | other | |
| chr22 | *MFNG* | 7 | 347 | 18 | MFNG | MFNG O-fucosylpeptide 3-beta-N-acetylglucosaminyltransferase | Cytoplasm | enzyme | |
| chr1 | *WDR26* | 7 | 219 | 18 | WDR26 | WD repeat domain 26 | Cytoplasm | other | |
| chr13 | *MIR4500HG* | 8 | 145 | 18 | MIR4500HG | MIR4500 host gene | Other | other | |
| chr1 | *UBE2J2* | 9 | 379 | 18 | UBE2J2 | ubiquitin conjugating enzyme E2 | Cytoplasm | enzyme | |
| chr6 | *RP11-444E17.6* | 9 | 201 | 18 | | | | | |
| chr9 | *PDCL* | 8 | 298 | 18 | PDCL | phosducin like | Cytoplasm | enzyme | |
| chr9 | *RP11-274B18.4* | 8 | 173 | 18 | | | | | |
| chr12 | *RP11-588H23.3* | 8 | 193 | 18 | | | | | |

| | | | | | | | | | |
|---|---|---|---|---|---|---|---|---|---|
| chr9 | *ZBTB34* | 8 | 226 | 18 | ZBTB34 | zinc finger and BTB domain containing 34 | Nucleus | transcription regulator | |
| chr20 | *CHRNA4* | 9 | 637 | 18 | CHRNA4 | cholinergic receptor nicotinic alpha 4 subunit | Plasma Membrane | transmembrane receptor | succinylcholine, lobeline, rapacuronium, doxacurium, vecuronium, pipecuronium, ABT 418, metocurine, mivacurium, D-tubocurarine, memantine, nicotine, arecoline, donepezil/memantine, pancuronium, atracurium, isoflurane, cisatracurium, rocuronium, amobarbital, mecamylamine, enflurane, ABT-089 |
| chr9 | *PRDM12* | 8 | 397 | 18 | PRDM12 | PR/SET domain 12 | Nucleus | transcription regulator | |
| chrX | *MAGED1* | 8 | 208 | 18 | MAGED1 | MAGE family member D1 | Plasma Membrane | transcription regulator | |
| chr3 | *IQCG* | 6 | 188 | 18 | IQCG | IQ motif containing G | Cytoplasm | other | |
| chr17 | *GRIN2C* | 9 | 409 | 18 | GRIN2C | glutamate ionotropic receptor NMDA type subunit 2C | Plasma Membrane | ion channel | nebostinel, besonprodil, delucemine, 1-aminocyclopropane-1-carboxylic acid, orphenadrine, dextromethorphan/promethazine, neramexane, felbamate, chlorpheniramine/dextromethorphan/phenylephrine, memantine, donepezil/memantine, N-(2-indanyl)glycinamide, tenocyclidine, dextromethorphan, cycloserine, UK-240455, brompheniramine/dextromethorphan/pseudoephedrine, [illegible] |
| chrX | *F8* | 7 | 200 | 18 | F8 | coagulation factor VIII | Extracellular Space | other | drotrecogin alfa, F9 |
| chr9 | *LHX2* | 9 | 349 | 18 | LHX2 | LIM homeobox 2 | Nucleus | transcription regulator | |
| chr6 | *ZC3H12D* | 8 | 237 | 18 | ZC3H12D | zinc finger CCCH-type containing 12D | Cytoplasm | other | |
| chr8 | *GRHL2* | 7 | 191 | 18 | GRHL2 | grainyhead like transcription | Nucleus | transcription regulator | |
| chr19 | *ADCK4* | 8 | 379 | 18 | COQ8B | coenzyme Q8B | Cytoplasm | kinase | |
| chrX | *RP11-320G24.1* | 9 | 242 | 18 | | | | | |
| chr19 | *ZNF329* | 8 | 204 | 18 | ZNF329 | zinc finger protein 329 | Other | other | |
| chr20 | *PTPRA* | 6 | 188 | 18 | PTPRA | protein tyrosine phosphatase receptor type A | Plasma Membrane | phosphatase | |
| chr9 | *RASEF* | 8 | 208 | 18 | RASEF | RAS and EF-hand domain | Cytoplasm | other | |
| chr22 | *TRMU* | 8 | 319 | 18 | TRMU | tRNA 5-methylaminomethyl-2-thiouridylate methyltransferase | Cytoplasm | enzyme | |
| chr3 | *MST1R* | 7 | 306 | 18 | MST1R | macrophage stimulating 1 receptor | Plasma Membrane | kinase | crizotinib/erlotinib, crizotinib/pazopanib, crizotinib/gefitinib, crizotinib/vemurafenib, narnatumab, glesatinib, crizotinib |
| chr3 | *SEC13* | 8 | 262 | 18 | SEC13 | SEC13 homolog, nuclear pore and COPII coat complex | Cytoplasm | transporter | |
| chr14 | *MMP14* | 7 | 338 | 18 | MMP14 | matrix metallopeptidase 14 | Extracellular Space | peptidase | prinomastat, rebimastat, |
| chr11 | *MAPK8IP1* | 9 | 501 | 18 | MAPK8IP1 | mitogen-activated protein kinase 8 interacting protein 1 | Cytoplasm | other | |
| chr2 | *PDIA6* | 7 | 234 | 18 | PDIA6 | protein disulfide isomerase family A member 6 | Cytoplasm | enzyme | |
| chr7 | *NCF1B* | 8 | 447 | 18 | NCF1B | neutrophil cytosolic factor 1B pseudogene | Other | other | |
| chr3 | *RP11-761N21.1* | 9 | 165 | 18 | | | | | |
| chr15 | *CYP11A1* | 8 | 325 | 18 | CYP11A1 | cytochrome P450 family 11 subfamily A member 1 | Cytoplasm | enzyme | ODM-208 |
| chr5 | *SPEF2* | 8 | 579 | 18 | SPEF2 | sperm flagellar 2 | Cytoplasm | other | |
| chr6 | *LPA* | 9 | 185 | 18 | LPA | lipoprotein(a) | Extracellular Space | other | |
| chr8 | *LRRC6* | 8 | 190 | 18 | LRRC6 | leucine rich repeat containing 6 | Cytoplasm | other | |
| chr14 | *AK7* | 7 | 157 | 18 | AK7 | adenylate kinase 7 | Cytoplasm | kinase | |
| chr16 | *CLUAP1* | 8 | 230 | 18 | CLUAP1 | clusterin associated protein 1 | Nucleus | other | |
| chr8 | *ZFHX4* | 8 | 175 | 18 | ZFHX4 | zinc finger homeobox 4 | Nucleus | transcription regulator | |
| chr5 | *C5orf22* | 7 | 466 | 18 | C5orf22 | chromosome 5 open reading frame 22 | Other | other | |

| | | | | | | | | | |
|---|---|---|---|---|---|---|---|---|---|
| chr14 | *PLEKHH1* | 7 | 237 | 18 | PLEKHH1 | pleckstrin homology, MyTH4 and FERM domain containing H1 | Cytoplasm | other | |
| chr3 | *NLGN1* | 6 | 188 | 18 | NLGN1 | neuroligin 1 | Plasma Membrane | enzyme | |
| chr15 | *POLR2M* | 7 | 205 | 18 | POLR2M | RNA polymerase II subunit M | Nucleus | other | |
| chr5 | *RP11-419C19.2* | 7 | 343 | 18 | | | | | |
| chr5 | *CCDC127* | 9 | 1430 | 18 | CCDC127 | coiled-coil domain containing 127 | Other | other | |
| chr12 | *RP11-1028N23.4* | 8 | 220 | 18 | | | | | |
| chr22 | *PEX26* | 7 | 197 | 18 | PEX26 | peroxisomal biogenesis factor 26 | Cytoplasm | other | |
| chr17 | *CD300A* | 8 | 307 | 18 | CD300A | CD300a molecule | Plasma Membrane | transmembrane receptor | |
| chr15 | *SCG5* | 7 | 162 | 18 | SCG5 | secretogranin V | Extracellular Space | other | |
| chr22 | *ADRBK2* | 6 | 158 | 18 | GRK3 | G protein-coupled receptor | Cytoplasm | kinase | |
| chr20 | *MYLK2* | 8 | 638 | 18 | MYLK2 | myosin light chain kinase 2 | Cytoplasm | kinase | |
| chr11 | *LTBP3* | 10 | 636 | 18 | LTBP3 | latent transforming growth factor beta binding protein 3 | Extracellular Space | other | |
| chrX | *PRKX* | 7 | 175 | 18 | PRKX | protein kinase X-linked | Cytoplasm | kinase | |
| chr10 | *LBX1-AS1* | 9 | 194 | 18 | LBX1-AS1 | LBX1 antisense RNA 1 | Other | other | |
| chr1 | *FAM78B* | 9 | 172 | 18 | FAM78B | family with sequence similarity 78 member B | Other | other | |
| chr20 | *RP11-290F20.3* | 8 | 518 | 18 | | | | | |
| chr6 | *RP11-517H2.6* | 8 | 262 | 18 | | | | | |
| chr17 | *MYBBP1A* | 9 | 308 | 18 | MYBBP1A | MYB binding protein 1a | Nucleus | transcription regulator | |
| chr11 | *GTF2H1* | 7 | 185 | 18 | GTF2H1 | general transcription factor IIH subunit 1 | Nucleus | transcription regulator | |
| chr14 | *NEMF* | 7 | 161 | 18 | NEMF | nuclear export mediator factor | Nucleus | other | |
| chr11 | *GANAB* | 8 | 239 | 18 | GANAB | glucosidase II alpha subunit | Cytoplasm | enzyme | miglitol |
| chr7 | *NOD1* | 8 | 489 | 18 | NOD1 | nucleotide binding oligomerization domain containing 1 | Cytoplasm | other | |
| chr11 | *RP11-855O10.2* | 9 | 508 | 18 | | | | | |
| chr7 | *EPHB4* | 8 | 378 | 18 | EPHB4 | EPH receptor B4 | Plasma Membrane | kinase | tesevatinib, JI 101 |
| chr1 | *ABCB10* | 7 | 278 | 18 | ABCB10 | ATP binding cassette subfamily B member 10 | Cytoplasm | transporter | |
| chr9 | *KIAA1958* | 7 | 176 | 18 | KIAA1958 | KIAA1958 | Other | other | |
| chr4 | *SLC2A9* | 7 | 227 | 18 | SLC2A9 | solute carrier family 2 member 9 | Plasma Membrane | transporter | |
| chr1 | *CDK11B* | 8 | 451 | 18 | CDK11B | cyclin dependent kinase 11B | Nucleus | kinase | |
| chr17 | *METTL2A* | 8 | 234 | 18 | METTL2A | methyltransferase like 2A | Other | enzyme | |
| chr22 | *CSF2RB* | 7 | 344 | 18 | CSF2RB | colony stimulating factor 2 receptor beta common subunit | Plasma Membrane | transmembrane receptor | rituximab/sargramostim, EPO/sargramostim, |
| chr5 | *CTB-32H22.1* | 9 | 246 | 18 | | | | | |
| chr11 | *RP11-708B6.2* | 9 | 148 | 18 | | | | | |
| chr8 | *INTS10* | 7 | 182 | 18 | INTS10 | integrator complex subunit 10 | Nucleus | other | |
| chr19 | *ZNF765* | 8 | 218 | 18 | ZNF765 | zinc finger protein 765 | Other | other | |
| chr10 | *CYP2C19* | 8 | 175 | 18 | CYP2C19 | cytochrome P450 family 2 subfamily C member 19 | Cytoplasm | enzyme | stiripentol |
| chr3 | *KBTBD12* | 8 | 301 | 18 | KBTBD12 | kelch repeat and BTB domain containing 12 | Other | other | |
| chr7 | *ZNF3* | 8 | 299 | 18 | ZNF3 | zinc finger protein 3 | Nucleus | transcription regulator | |
| chr1 | *CTPS1* | 9 | 258 | 18 | CTPS1 | CTP synthase 1 | Nucleus | enzyme | cytidine triphosphate synthetase inhibitor |
| chr17 | *RP11-219G17.4* | 8 | 398 | 18 | | | | | |
| chr2 | *LINC00486* | 7 | 169 | 18 | LINC00486 | long intergenic non-protein coding RNA 486 | Other | other | |
| chr9 | *SUSD3* | 8 | 275 | 18 | SUSD3 | sushi domain containing 3 | Plasma Membrane | other | |
| chr3 | *TMCC1-AS1* | 7 | 311 | 18 | TMCC1-AS1 | TMCC1 antisense RNA 1 (head to head) | Other | other | |
| chr11 | *WT1-AS* | 9 | 331 | 18 | WT1-AS | WT1 antisense RNA | Other | other | |
| chr1 | *UBIAD1* | 9 | 246 | 18 | UBIAD1 | UbiA prenyltransferase domain containing 1 | Nucleus | enzyme | |
| chr11 | *RP11-1L12.3* | 9 | 370 | 18 | | | | | |
| chr16 | *CDR2* | 8 | 191 | 18 | CDR2 | cerebellar degeneration related protein 2 | Cytoplasm | other | |
| chr5 | *TTC1* | 7 | 200 | 18 | TTC1 | tetratricopeptide repeat domain 1 | Cytoplasm | other | |
| chr20 | *TCFL5* | 8 | 493 | 18 | TCFL5 | transcription factor like 5 | Nucleus | transcription regulator | |
| chr17 | *EFCAB13* | 7 | 191 | 18 | EFCAB13 | EF-hand calcium binding domain | Nucleus | other | |
| chr17 | *DHRS7C* | 8 | 272 | 18 | DHRS7C | dehydrogenase/reductase 7C | Cytoplasm | enzyme | |
| chr2 | *ATAD2B* | 7 | 174 | 18 | ATAD2B | ATPase family AAA domain containing 2B | Nucleus | other | |
| chr19 | *RASGRP4* | 7 | 283 | 18 | RASGRP4 | RAS guanyl releasing protein 4 | Cytoplasm | other | |
| chr12 | *RP11-541G9.1* | 8 | 183 | 18 | | | | | |
| chr7 | *ZSCAN21* | 7 | 319 | 18 | ZSCAN21 | zinc finger and SCAN domain containing 21 | Nucleus | transcription regulator | |
| chr5 | *RP11-1319K7.1* | 8 | 168 | 18 | | | | | |
| chr4 | *RP11-138I17.1* | 7 | 224 | 18 | | | | | |
| chr18 | *RP11-703M24.5* | 8 | 294 | 18 | | | | | |
| chr3 | *LINC00971* | 8 | 197 | 18 | LINC00971 | long intergenic non-protein coding RNA 971 | Other | other | |
| chr22 | *PPM1F* | 8 | 395 | 18 | PPM1F | protein phosphatase, Mg2+/Mn2+ dependent 1F | Cytoplasm | phosphatase | |
| chr2 | *PEX13* | 7 | 201 | 18 | PEX13 | peroxisomal biogenesis factor 13 | Cytoplasm | transporter | |
| chr7 | *C1GALT1* | 8 | 202 | 18 | C1GALT1 | core 1 synthase, glycoprotein-N-acetylgalactosamine 3-beta-galactosyltransferase 1 | Plasma Membrane | enzyme | |
| chr15 | *CERS3* | 9 | 294 | 18 | CERS3 | ceramide synthase 3 | Nucleus | transcription regulator | |
| chr11 | *RP11-554A11.8* | 8 | 292 | 18 | | | | | |
| chr19 | *CACNG8* | 8 | 332 | 18 | CACNG8 | calcium voltage-gated channel auxiliary subunit gamma 8 | Plasma Membrane | ion channel | |
| chr6 | *RP1-261G23.5* | 8 | 268 | 18 | | | | | |
| chr1 | *OR2L13* | 7 | 167 | 18 | OR2L13 | olfactory receptor family 2 subfamily L member 13 | Plasma Membrane | G-protein coupled receptor | |

| | | | | | | | | | |
|---|---|---|---|---|---|---|---|---|---|
| chr1 | *RP3-522D1.1* | 7 | 274 | 18 | | | | | |
| chrX | *ZRSR2* | 7 | 218 | 18 | ZRSR2 | zinc finger CCCH-type, RNA binding motif and serine/arginine | Nucleus | other | |
| chr17 | *FXR2* | 8 | 227 | 18 | FXR2 | FMR1 autosomal homolog 2 | Cytoplasm | other | |
| chr1 | *MKNK1* | 8 | 206 | 18 | MKNK1 | MAPK interacting serine/threonine kinase 1 | Cytoplasm | kinase | ETC-1907206, BAY1143269, tomivosertib, dacomitinib |
| chr2 | *RGPD3* | 8 | 260 | 18 | PD4 (includes oth | RANBP2 like and GRIP domain containing 5 | Nucleus | enzyme | |
| chr1 | *TP53BP2* | 7 | 252 | 18 | TP53BP2 | tumor protein p53 binding protein | Nucleus | other | |
| chr22 | *SMC1B* | 6 | 163 | 18 | SMC1B | structural maintenance of chromosomes 1B | Nucleus | transporter | |
| chr22 | *TUBGCP6* | 8 | 430 | 18 | TUBGCP6 | tubulin gamma complex associated protein 6 | Cytoplasm | other | |
| chr19 | *SBNO2* | 10 | 411 | 18 | SBNO2 | strawberry notch homolog 2 | Other | transcription regulator | |
| chr2 | *AC097374.2* | 8 | 286 | 18 | | | | | |
| chr2 | *NPHP1* | 7 | 222 | 18 | NPHP1 | nephrocystin 1 | Nucleus | other | |
| chr17 | *LUC7L3* | 6 | 174 | 18 | LUC7L3 | LUC7 like 3 pre-mRNA splicing | Nucleus | other | |
| chr6 | *PTCHD4* | 9 | 167 | 18 | PTCHD4 | patched domain containing 4 | Other | other | |
| chr12 | *RP11-123O10.4* | 8 | 145 | 18 | | | | | |
| chr11 | *LGR4* | 8 | 307 | 18 | LGR4 | leucine rich repeat containing G protein-coupled receptor 4 | Plasma Membrane | transmembrane recepto | |
| chr2 | *CCDC104* | 8 | 258 | 18 | CFAP36 | cilia and flagella associated | Extracellular Space | other | |
| chr7 | *AC118138.2* | 8 | 265 | 18 | | | | | |
| chr12 | *C12orf79* | 8 | 176 | 18 | LINC01619 | long intergenic non-protein coding RNA 1619 | Other | other | |
| chr1 | *RP11-426L16.8* | 8 | 267 | 18 | | | | | |
| chr11 | *CSTF3-AS1* | 8 | 232 | 18 | CSTF3-DT | CSTF3 divergent transcript | Other | other | |
| chr7 | *TRA2A* | 7 | 292 | 18 | TRA2A | transformer 2 alpha homolog | Nucleus | other | |
| chr6 | *CCDC167* | 7 | 314 | 18 | CCDC167 | coiled-coil domain containing 167 | Other | other | |
| chr1 | *C1orf27* | 6 | 212 | 18 | ODR4 | odr-4 GPCR localization factor homolog | Other | other | |
| chr15 | *GNB5* | 8 | 228 | 18 | GNB5 | G protein subunit beta 5 | Plasma Membrane | enzyme | |
| chr17 | *GLOD4* | 8 | 249 | 18 | GLOD4 | glyoxalase domain containing 4 | Cytoplasm | enzyme | |
| chr18 | *ANKRD29* | 8 | 210 | 18 | ANKRD29 | ankyrin repeat domain 29 | Other | other | |
| chr5 | *SLCO6A1* | 8 | 160 | 18 | SLCO6A1 | solute carrier organic anion transporter family member 6A1 | Plasma Membrane | transporter | |
| chr16 | *RP11-467L24.1* | 7 | 162 | 18 | | | | | |
| chr7 | *DPY19L2P1* | 8 | 221 | 18 | DPY19L2P1 | DPY19L2 pseudogene 1 | Other | other | |
| chr16 | *AQP8* | 7 | 294 | 18 | AQP8 | aquaporin 8 | Plasma Membrane | transporter | |
| chr16 | *DOK4* | 7 | 383 | 18 | DOK4 | docking protein 4 | Plasma Membrane | other | |
| chr6 | *ZNF318* | 8 | 188 | 18 | ZNF318 | zinc finger protein 318 | Nucleus | other | |
| chr16 | *FAM92B* | 8 | 367 | 18 | FAM92B | family with sequence similarity 92 member B | Cytoplasm | other | |
| chr19 | *NACC1* | 8 | 305 | 18 | NACC1 | nucleus accumbens associated 1 | Nucleus | transcription regulator | |
| chr1 | *DUSP10* | 8 | 264 | 18 | DUSP10 | dual specificity phosphatase 10 | Nucleus | phosphatase | |
| chr5 | *LNPEP* | 8 | 153 | 18 | LNPEP | leucyl and cystinyl | Cytoplasm | peptidase | |
| chr12 | *MYRFL* | 7 | 193 | 18 | MYRFL | myelin regulatory factor like | Other | other | |
| chr12 | *LRRK2* | 8 | 208 | 18 | LRRK2 | leucine rich repeat kinase 2 | Cytoplasm | kinase | |
| chr16 | *MVP* | 8 | 291 | 18 | MVP | major vault protein | Nucleus | other | |
| chr11 | *CSTF3* | 7 | 212 | 18 | CSTF3 | cleavage stimulation factor | Nucleus | other | |
| chr11 | *P2RY2* | 9 | 381 | 18 | P2RY2 | purinergic receptor P2Y2 | Plasma Membrane | -protein coupled recepto | UTP, suramin |
| chr1 | *MYBPHL* | 8 | 340 | 18 | MYBPHL | myosin binding protein H like | Other | other | |
| chr1 | *CSF3R* | 7 | 369 | 18 | CSF3R | colony stimulating factor 3 receptor | Plasma Membrane | transmembrane recepto | clofarabine/cytarabine/filgrastim/idarubicin, clofarabine/filgrastim/idarubicin, benegrastim, cladribine/cytarabine/filgrastim/idarubicin/plerixafor, cytarabine/filgrastim/fludarabine phosphate/imatinib, cytarabine/filgrastim/fludarabine phosphate/idarubicin/imatinib, cytarabine/filgrastim/fludarabine phosphate, EPO/filgrastim, cladribine/cytarabine/filgrastim/idarubicin, cytarabine/filgrastim/f[illegible] |
| chr5 | *AC091969.1* | 7 | 215 | 18 | | | | | |
| chr6 | *RP1-223B1.1* | 7 | 311 | 18 | | | | | |
| chr1 | *HORMAD1* | 8 | 239 | 18 | HORMAD1 | HORMA domain containing 1 | Nucleus | other | |
| chr2 | *TSN* | 7 | 332 | 18 | TSN | translin | Nucleus | other | |
| chr5 | *ETF1* | 7 | 195 | 18 | ETF1 | eukaryotic translation termination factor 1 | Cytoplasm | translation regulator | |
| chr3 | *EAF2* | 6 | 222 | 18 | EAF2 | ELL associated factor 2 | Nucleus | transcription regulator | |
| chr3 | *ARF4* | 7 | 223 | 18 | ARF4 | ADP ribosylation factor 4 | Cytoplasm | enzyme | |
| chr10 | *KIF20B* | 7 | 192 | 18 | KIF20B | kinesin family member 20B | Nucleus | enzyme | |
| chr10 | *SGPL1* | 8 | 256 | 18 | SGPL1 | sphingosine-1-phosphate lyase 1 | Cytoplasm | enzyme | |

| | | | | | | | | |
|---|---|---|---|---|---|---|---|---|
| chr19 | *LSR* | 8 | 351 | 18 | LSR | lipolysis stimulated lipoprotein receptor | Nucleus | transcription regulator |
| chr1 | *NRD1* | 7 | 197 | 18 | NRDC | nardilysin convertase | Cytoplasm | peptidase |
| chr5 | *C5orf45* | 9 | 367 | 18 | MRNIP | MRN complex interacting protein | Nucleus | other |
| chr1 | *SHE* | 8 | 309 | 18 | SHE | Src homology 2 domain | Cytoplasm | other |
| chr19 | *CSNK1G2* | 10 | 538 | 18 | CSNK1G2 | casein kinase 1 gamma 2 | Cytoplasm | kinase |
| chr19 | *SGTA* | 9 | 474 | 18 | SGTA | small glutamine rich tetratricopeptide repeat | Cytoplasm | other |
| chr1 | *SDHB* | 8 | 219 | 18 | SDHB | succinate dehydrogenase complex iron sulfur subunit B | Cytoplasm | enzyme |
| chr19 | *CNOT3* | 8 | 414 | 18 | CNOT3 | CCR4-NOT transcription complex subunit 3 | Cytoplasm | other |
| chr10 | *FAM35DP* | 7 | 289 | 18 | SHLD2P3 | shieldin complex subunit 2 pseudogene 3 | Other | other |
| chr11 | *BRSK2* | 8 | 514 | 18 | BRSK2 | BR serine/threonine kinase 2 | Cytoplasm | kinase |
| chr19 | *DAZAP1* | 9 | 423 | 18 | DAZAP1 | DAZ associated protein 1 | Other | other |
| chr15 | *IDH3A* | 9 | 257 | 18 | IDH3A | isocitrate dehydrogenase (NAD(+)) 3 catalytic subunit alpha | Cytoplasm | enzyme |
| chr20 | *ESF1* | 6 | 158 | 18 | ESF1 | ESF1 nucleolar pre-rRNA processing protein homolog | Nucleus | other |
| chr10 | *BLNK* | 8 | 177 | 18 | BLNK | B cell linker | Cytoplasm | other |
| chr17 | *CTB-187M2.3* | 8 | 231 | 18 | | | | |
| chr7 | *ZNF316* | 8 | 480 | 18 | ZNF316 | zinc finger protein 316 | Other | other |
| chr11 | *TCP11L1* | 9 | 283 | 18 | TCP11L1 | t-complex 11 like 1 | Cytoplasm | other |
| chr12 | *SNRNP35* | 8 | 224 | 18 | SNRNP35 | small nuclear ribonucleoprotein U11/U12 subunit 35 | Nucleus | other |
| chr5 | *CTC-546K23.1* | 8 | 173 | 18 | | | | |
| chr19 | *ZNF490* | 7 | 221 | 18 | ZNF490 | zinc finger protein 490 | Nucleus | transcription regulator |
| chr18 | *RP13-270P17.2* | 7 | 296 | 18 | | | | |
| chr17 | *CTC-297N7.11* | 7 | 146 | 18 | | | | |
| chr12 | *C12orf40* | 8 | 393 | 18 | C12orf40 | chromosome 12 open reading frame 40 | Other | other |
| chr10 | *KIAA1462* | 7 | 200 | 18 | JCAD | junctional cadherin 5 associated | Plasma Membrane | other |
| chr22 | *SLC25A17* | 6 | 172 | 18 | SLC25A17 | solute carrier family 25 member | Cytoplasm | transporter |
| chr3 | *GOLGB1* | 7 | 217 | 18 | GOLGB1 | golgin B1 | Cytoplasm | other |
| chr7 | *PTPRZ1* | 7 | 149 | 18 | PTPRZ1 | protein tyrosine phosphatase receptor type Z1 | Plasma Membrane | phosphatase |
| chr7 | *LRRC72* | 8 | 247 | 18 | LRRC72 | leucine rich repeat containing 72 | Other | other |
| chr19 | *PRKD2* | 8 | 275 | 18 | PRKD2 | protein kinase D2 | Cytoplasm | kinase |
| chr11 | *SLC22A18* | 10 | 827 | 18 | SLC22A18 | solute carrier family 22 member | Plasma Membrane | transporter |
| chr8 | *GS1-57L11.1* | 8 | 198 | 18 | | | | |
| chr20 | *BLCAP* | 9 | 332 | 18 | BLCAP | BLCAP apoptosis inducing factor | Other | other |
| chr1 | *TARS2* | 8 | 289 | 18 | TARS2 | threonyl-tRNA synthetase 2, mitochondrial | Cytoplasm | enzyme |
| chr16 | *DHX38* | 8 | 317 | 18 | DHX38 | DEAH-box helicase 38 | Nucleus | enzyme |
| chr10 | *SEC24C* | 9 | 396 | 18 | SEC24C | SEC24 homolog C, COPII coat complex component | Cytoplasm | transporter |
| chr9 | *ABHD17B* | 7 | 161 | 18 | ABHD17B | abhydrolase domain containing | Plasma Membrane | peptidase |
| chr15 | *DUOX2* | 8 | 314 | 18 | DUOX2 | dual oxidase 2 | Plasma Membrane | enzyme |
| chr3 | *MCF2L2* | 7 | 194 | 18 | MCF2L2 | MCF.2 cell line derived transforming sequence-like 2 | Other | other |
| chr3 | *APPL1* | 6 | 150 | 18 | APPL1 | adaptor protein, phosphotyrosine interacting with PH domain and leucine zipper 1 | Cytoplasm | other |
| chr9 | *APTX* | 7 | 175 | 18 | APTX | aprataxin | Nucleus | phosphatase |
| chr1 | *FAM102B* | 8 | 207 | 18 | FAM102B | family with sequence similarity 102 member B | Other | other |
| chr7 | *POLR2J2* | 8 | 389 | 18 | 'OLR2J2/POLR2J | RNA polymerase II subunit J2 | Other | transcription regulator |
| chr3 | *NMNAT3* | 9 | 258 | 18 | NMNAT3 | nicotinamide nucleotide adenylyltransferase 3 | Cytoplasm | enzyme |
| chr22 | *TOP3B* | 7 | 352 | 18 | TOP3B | DNA topoisomerase III beta | Nucleus | enzyme |
| chr3 | *ALCAM* | 7 | 271 | 18 | ALCAM | activated leukocyte cell adhesion molecule | Plasma Membrane | other |
| chr20 | *ZSWIM3* | 8 | 297 | 18 | ZSWIM3 | zinc finger SWIM-type containing | Other | other |
| chr11 | *RHOD* | 8 | 413 | 18 | RHOD | ras homolog family member D | Cytoplasm | enzyme |
| chr12 | *MMAB* | 8 | 295 | 18 | MMAB | metabolism of cobalamin associated B | Cytoplasm | enzyme |
| chr1 | *FLG-AS1* | 9 | 289 | 18 | FLG-AS1 | FLG antisense RNA 1 | Other | other |
| chr12 | *RP11-630C16.2* | 8 | 273 | 18 | | | | |
| chr3 | *RP11-18H7.1* | 7 | 227 | 18 | | | | |
| chr1 | *SLC2A1-AS1* | 8 | 206 | 18 | SLC2A1-AS1 | SLC2A1 antisense RNA 1 | Other | other |
| chr3 | *FBXW12* | 7 | 214 | 18 | FBXW12 | F-box and WD repeat domain containing 12 | Cytoplasm | enzyme |
| chr19 | *CTC-523E23.4* | 8 | 189 | 18 | | | | |
| chr17 | *GUCY2D* | 8 | 306 | 18 | GUCY2D | guanylate cyclase 2D, retinal | Plasma Membrane | kinase |
| chr12 | *PUS1* | 8 | 303 | 18 | PUS1 | pseudouridine synthase 1 | Nucleus | enzyme |
| chr19 | *CEACAM22P* | 8 | 163 | 18 | CEACAM22P | CEA cell adhesion molecule 22, pseudogene | Other | other |
| chr1 | *BMP8B* | 8 | 401 | 18 | BMP8B | bone morphogenetic protein 8b | Extracellular Space | growth factor |
| chr3 | *CCDC80* | 7 | 226 | 18 | CCDC80 | coiled-coil domain containing 80 | Nucleus | other |
| chr3 | *ZBBX* | 8 | 210 | 18 | ZBBX | zinc finger B-box domain | Other | other |

| | | | | | | | | | |
|---|---|---|---|---|---|---|---|---|---|
| chr3 | *DRD3* | 8 | 258 | 18 | DRD3 | dopamine receptor D3 | Plasma Membrane | -protein coupled recepto | ziprasidone, pimozide, pramipexole, prochlorperazine, asenapine, haloperidol decanoate, chlorprothixene, dopamine, loxapine, clozapine, brexpiprazole, dihydroergotamine, carbidopa/entacapone/levodopa, carbidopa/levodopa/tolcapone, trifluoperazine, abaperidone, apomorphine, fluphenazine decanoate, lisuride, pergolide, mesoridazine, [illegible] |
| chr11 | *ALG9* | 7 | 154 | 18 | ALG9 | ALG9 alpha-1,2- | Cytoplasm | enzyme | |
| chr12 | *GNPTAB* | 8 | 159 | 18 | GNPTAB | N-acetylglucosamine-1-phosphate transferase subunits | Cytoplasm | enzyme | |
| chrX | *UBA1* | 8 | 294 | 18 | UBA1 | ubiquitin like modifier activating enzyme 1 | Cytoplasm | enzyme | MLN7243 |
| chr19 | *LMTK3* | 8 | 265 | 18 | LMTK3 | lemur tyrosine kinase 3 | Other | kinase | |
| chr20 | *PYGB* | 8 | 266 | 18 | PYGB | glycogen phosphorylase B | Cytoplasm | enzyme | |
| chr2 | *AC016722.2* | 8 | 444 | 18 | | | | | |
| chr20 | *SUN5* | 8 | 343 | 18 | SUN5 | Sad1 and UNC84 domain containing 5 | Nucleus | other | |
| chrX | *RP11-453F18__B.* | 6 | 160 | 18 | | | | | |
| chr9 | *RP11-292B8.1* | 6 | 166 | 18 | | | | | |
| chr22 | *NPTXR* | 7 | 306 | 18 | NPTXR | neuronal pentraxin receptor | Plasma Membrane | transmembrane receptor | |
| chr16 | *LINC00917* | 8 | 277 | 18 | LINC00917 | long intergenic non-protein coding RNA 917 | Other | other | |
| chr6 | *HCRTR2* | 7 | 173 | 18 | HCRTR2 | hypocretin receptor 2 | Plasma Membrane | -protein coupled recepto | seltorexant, |
| chr9 | *TMEM8C* | 9 | 544 | 18 | MYMK | myomaker, myoblast fusion factor | Plasma Membrane | other | |
| chr9 | *SECISBP2* | 7 | 174 | 18 | SECISBP2 | SECIS binding protein 2 | Cytoplasm | translation regulator | |
| chr11 | *APBB1* | 8 | 494 | 18 | APBB1 | amyloid beta precursor protein binding family B member 1 | Cytoplasm | transcription regulator | |
| chr5 | *PRKAA1* | 7 | 461 | 18 | PRKAA1 | protein kinase AMP-activated catalytic subunit alpha 1 | Cytoplasm | kinase | phenformin |
| chr1 | *PGD* | 8 | 211 | 18 | PGD | phosphogluconate | Cytoplasm | enzyme | |
| chr7 | *TRIM4* | 9 | 288 | 18 | TRIM4 | tripartite motif containing 4 | Cytoplasm | other | |
| chr10 | *BAG3* | 8 | 291 | 18 | BAG3 | BCL2 associated athanogene 3 | Cytoplasm | other | |
| chr7 | *AC024084.1* | 8 | 199 | 18 | | | | | |
| chr11 | *FAM86C2P* | 8 | 316 | 18 | FAM86C2P | family with sequence similarity 86, member A pseudogene | Other | other | |
| chr15 | *RP11-1069G10.1* | 7 | 220 | 18 | | | | | |
| chr4 | *LINC00504* | 7 | 193 | 18 | LINC00504 | | Other | other | |
| chr19 | *PIAS4* | 9 | 395 | 18 | PIAS4 | protein inhibitor of activated | Nucleus | transcription regulator | |
| chr3 | *CLRN1-AS1* | 6 | 190 | 18 | CLRN1-AS1 | CLRN1 antisense RNA 1 | Other | other | |
| chr5 | *UGT3A1* | 8 | 649 | 18 | UGT3A1 | UDP glycosyltransferase family 3 member A1 | Cytoplasm | enzyme | |
| chr19 | *UBA2* | 7 | 185 | 18 | UBA2 | ubiquitin like modifier activating enzyme 2 | Cytoplasm | enzyme | |
| chr7 | *GS1-124K5.10* | 7 | 296 | 18 | | | | | |
| chr7 | *C7orf31* | 8 | 256 | 18 | C7orf31 | chromosome 7 open reading frame 31 | Cytoplasm | transporter | |
| chr18 | *TTC39C* | 8 | 157 | 18 | TTC39C | tetratricopeptide repeat domain | Other | other | |
| chr6 | *DSE* | 7 | 195 | 18 | DSE | dermatan sulfate epimerase | Cytoplasm | enzyme | |
| chr19 | *EPS8L1* | 8 | 339 | 18 | EPS8L1 | EPS8 like 1 | Cytoplasm | other | |
| chr9 | *KIAA0020* | 7 | 187 | 18 | PUM3 | pumilio RNA binding family | Nucleus | other | |
| chr1 | *SPRR2B* | 9 | 247 | 18 | SPRR2B | small proline rich protein 2B | Cytoplasm | other | |
| chr15 | *SNUPN* | 8 | 332 | 18 | SNUPN | snurportin 1 | Nucleus | other | |
| chr9 | *HSD17B3* | 7 | 164 | 18 | HSD17B3 | hydroxysteroid 17-beta dehydrogenase 3 | Cytoplasm | enzyme | |
| chr8 | *CNBD1* | 6 | 207 | 18 | CNBD1 | cyclic nucleotide binding domain containing 1 | Other | other | |
| chr2 | *BCL2L11* | 8 | 251 | 18 | BCL2L11 | BCL2 like 11 | Cytoplasm | other | |
| chr11 | *P4HA3* | 9 | 214 | 18 | P4HA3 | prolyl 4-hydroxylase subunit | Cytoplasm | enzyme | |
| chr7 | *AC091801.1* | 8 | 283 | 18 | | | | | |
| chr17 | *SNX11* | 8 | 192 | 18 | SNX11 | sorting nexin 11 | Cytoplasm | transporter | |
| chr5 | *KIAA0947* | 6 | 446 | 18 | ICE1 | interactor of little elongation complex ELL subunit 1 | Nucleus | other | |
| chr19 | *GYS1* | 7 | 290 | 18 | GYS1 | glycogen synthase 1 | Cytoplasm | enzyme | |
| chr9 | *SWI5* | 8 | 364 | 18 | SWI5 | SWI5 homologous recombination repair protein | Nucleus | other | |
| chrX | *CNKSR2* | 9 | 151 | 18 | CNKSR2 | connector enhancer of kinase suppressor of Ras 2 | Plasma Membrane | other | |
| chr5 | *NDFIP1* | 6 | 141 | 18 | NDFIP1 | Nedd4 family interacting protein 1 | Cytoplasm | other | |
| chr10 | *MMS19* | 8 | 215 | 18 | MMS19 | MMS19 homolog, cytosolic iron-sulfur assembly component | Nucleus | transcription regulator | |
| chr9 | *SLC1A1* | 8 | 189 | 18 | SLC1A1 | solute carrier family 1 member 1 | Plasma Membrane | transporter | riluzole |
| chr19 | *AC016582.2* | 8 | 217 | 18 | | | | | |

| | | | | | | | | | |
|---|---|---|---|---|---|---|---|---|---|
| chr3 | *MB21D2* | 6 | 170 | 18 | MB21D2 | Mab-21 domain containing 2 | Other | other | |
| chr3 | *ZIC1* | 9 | 260 | 18 | ZIC1 | Zic family member 1 | Nucleus | transcription regulator | |
| chr11 | *PDHX* | 9 | 246 | 18 | PDHX | pyruvate dehydrogenase complex component X | Cytoplasm | enzyme | |
| chr1 | *HHIPL2* | 8 | 253 | 18 | HHIPL2 | HHIP like 2 | Other | other | |
| chr18 | *LINC00470* | 8 | 185 | 18 | LINC00470 | long intergenic non-protein coding RNA 470 | Other | other | |
| chr14 | *TRIM9* | 7 | 219 | 18 | TRIM9 | tripartite motif containing 9 | Cytoplasm | enzyme | |
| chr17 | *ZNF286A* | 6 | 210 | 17 | ZNF286A | zinc finger protein 286A | Nucleus | transcription regulator | |
| chr17 | *BZRAP1* | 8 | 396 | 17 | TSPOAP1 | TSPO associated protein 1 | Cytoplasm | other | |
| chr6 | *RP11-440G9.1* | 9 | 198 | 17 | | | | | |
| chr5 | *TGFBI* | 8 | 264 | 17 | TGFBI | transforming growth factor beta induced | Extracellular Space | other | |
| chr7 | *AC068533.7* | 8 | 215 | 17 | | | | | |
| chr7 | *PPP1R3A* | 7 | 207 | 17 | PPP1R3A | protein phosphatase 1 regulatory subunit 3A | Cytoplasm | phosphatase | |
| chr1 | *SV2A* | 8 | 345 | 17 | SV2A | synaptic vesicle glycoprotein 2A | Cytoplasm | other | brivaracetam, levetiracetam |
| chr2 | *PNKD* | 7 | 208 | 17 | PNKD | PNKD metallo-beta-lactamase domain containing | Nucleus | other | |
| chr19 | *KIAA1683* | 8 | 350 | 17 | IQCN | IQ motif containing N | Cytoplasm | other | |
| chr3 | *LRRC31* | 7 | 305 | 17 | LRRC31 | leucine rich repeat containing 31 | Other | other | |
| chr1 | *RP11-202K23.1* | 8 | 192 | 17 | | | | | |
| chr2 | *SPAST* | 6 | 147 | 17 | SPAST | spastin | Nucleus | enzyme | |
| chr17 | *LRRC37A11P* | 8 | 184 | 17 | LRRC37A11P | leucine rich repeat containing 37 member A11, pseudogene | Other | other | |
| chr16 | *CTA-481E9.4* | 7 | 181 | 17 | | | | | |
| chr10 | *NRAP* | 9 | 203 | 17 | NRAP | nebulin related anchoring protein | Cytoplasm | other | |
| chr10 | *RP13-137A17.4* | 8 | 340 | 17 | | | | | |
| chr1 | *TRAF5* | 8 | 233 | 17 | TRAF5 | TNF receptor associated factor 5 | Cytoplasm | transporter | |
| chr7 | *LFNG* | 9 | 500 | 17 | LFNG | LFNG O-fucosylpeptide 3-beta-N-acetylglucosaminyltransferase | Cytoplasm | enzyme | |
| chr17 | *RP1-66C13.4* | 8 | 220 | 17 | | | | | |
| chr19 | *AF038458.3* | 7 | 255 | 17 | | | | | |
| chr17 | *CTD-2309O5.3* | 8 | 220 | 17 | | | | | |
| chr17 | *C17orf70* | 9 | 404 | 17 | FAAP100 | FA core complex associated protein 100 | Nucleus | other | |
| chr14 | *MYH6* | 7 | 319 | 17 | MYH6 | myosin heavy chain 6 | Cytoplasm | enzyme | |
| chrX | *PRDX4* | 7 | 189 | 17 | PRDX4 | peroxiredoxin 4 | Cytoplasm | enzyme | |
| chr17 | *HS3ST3A1* | 8 | 209 | 17 | HS3ST3A1 | heparan sulfate-glucosamine 3-sulfotransferase 3A1 | Cytoplasm | enzyme | |
| chr2 | *DCDC2C* | 6 | 219 | 17 | DCDC2C | doublecortin domain containing | Cytoplasm | other | |
| chr5 | *AGXT2* | 9 | 533 | 17 | AGXT2 | alanine--glyoxylate aminotransferase 2 | Cytoplasm | enzyme | |
| chr17 | *LGALS9B* | 7 | 348 | 17 | LGALS9B | galectin 9B | Cytoplasm | other | |
| chr10 | *HABP2* | 8 | 212 | 17 | HABP2 | hyaluronan binding protein 2 | Extracellular Space | peptidase | |
| chr12 | *SMUG1* | 8 | 439 | 17 | SMUG1 | single-strand-selective monofunctional uracil-DNA glycosylase 1 | Nucleus | enzyme | |
| chr21 | *C21orf49* | 7 | 175 | 17 | C21orf62-AS1 | C21orf62 antisense RNA 1 | Other | other | |
| chr2 | *NOL10* | 7 | 143 | 17 | NOL10 | nucleolar protein 10 | Nucleus | other | |
| chr10 | *NUTM2D* | 7 | 341 | 17 | NUTM2D | NUT family member 2D | Other | other | |
| chr14 | *PSEN1* | 6 | 183 | 17 | PSEN1 | presenilin 1 | Plasma Membrane | peptidase | tarenflurbil |
| chr1 | *TRIM63* | 7 | 263 | 17 | TRIM63 | tripartite motif containing 63 | Nucleus | enzyme | |
| chr2 | *POLR1B* | 7 | 270 | 17 | POLR1B | RNA polymerase I subunit B | Nucleus | enzyme | |
| chrX | *ITIH6* | 8 | 303 | 17 | ITIH6 | inter-alpha-trypsin inhibitor heavy chain family member 6 | Other | other | |
| chr16 | *DOC2A* | 9 | 338 | 17 | DOC2A | double C2 domain alpha | Cytoplasm | transporter | |
| chr19 | *SNRPA* | 8 | 256 | 17 | SNRPA | small nuclear ribonucleoprotein polypeptide A | Nucleus | other | |
| chr3 | *UBP1* | 8 | 186 | 17 | UBP1 | upstream binding protein 1 | Cytoplasm | transcription regulator | |
| chr12 | *FAM60A* | 7 | 250 | 17 | SINHCAF | SIN3-HDAC complex associated factor | Extracellular Space | other | |
| chr4 | *CC2D2A* | 6 | 147 | 17 | CC2D2A | coiled-coil and C2 domain containing 2A | Cytoplasm | other | |
| chr20 | *SERINC3* | 7 | 359 | 17 | SERINC3 | serine incorporator 3 | Cytoplasm | transporter | |
| chr12 | *METTL25* | 7 | 156 | 17 | METTL25 | methyltransferase like 25 | Extracellular Space | other | |
| chrX | *STARD8* | 7 | 271 | 17 | STARD8 | StAR related lipid transfer domain containing 8 | Cytoplasm | other | |
| chr3 | *ZCWPW2* | 7 | 160 | 17 | ZCWPW2 | zinc finger CW-type and PWWP domain containing 2 | Other | other | |
| chr22 | *EIF3L* | 7 | 162 | 17 | EIF3L | eukaryotic translation initiation factor 3 subunit L | Cytoplasm | other | |
| chr19 | *CIB3* | 8 | 243 | 17 | CIB3 | calcium and integrin binding family member 3 | Other | other | |
| chr1 | *FNDC7* | 7 | 231 | 17 | FNDC7 | fibronectin type III domain containing 7 | Other | other | |
| chr11 | *MRE11A* | 7 | 323 | 17 | MRE11 | MRE11 homolog, double strand break repair nuclease | Nucleus | enzyme | |
| chr2 | *SMYD1* | 8 | 221 | 17 | SMYD1 | SET and MYND domain | Nucleus | transcription regulator | |
| chr9 | *SURF4* | 8 | 461 | 17 | SURF4 | surfeit 4 | Cytoplasm | other | |
| chr12 | *CCDC77* | 7 | 189 | 17 | CCDC77 | coiled-coil domain containing 77 | Cytoplasm | other | |
| chr6 | *RP3-323P13.2* | 7 | 169 | 17 | | | | | |
| chr5 | *ELL2* | 7 | 187 | 17 | ELL2 | elongation factor for RNA polymerase II 2 | Nucleus | transcription regulator | |
| chr12 | *KCNH3* | 9 | 545 | 17 | KCNH3 | potassium voltage-gated channel subfamily H member 3 | Plasma Membrane | ion channel | |
| chr14 | *CYP46A1* | 8 | 271 | 17 | CYP46A1 | cytochrome P450 family 46 subfamily A member 1 | Cytoplasm | enzyme | |

| | | | | | | | | | |
|---|---|---|---|---|---|---|---|---|---|
| chr20 | *SLC17A9* | 9 | 772 | 17 | SLC17A9 | solute carrier family 17 member 9 | Cytoplasm | other | |
| chr11 | *RP11-587D21.4* | 8 | 248 | 17 | | | | | |
| chr5 | *HAVCR2* | 8 | 218 | 17 | HAVCR2 | hepatitis A virus cellular receptor 2 | Plasma Membrane | other | LY3415244, Sym023, RO7121661, TSR-022, MBG453, BMS-986258, BGB-A425, |
| chr16 | *RP11-459F6.3* | 8 | 194 | 17 | | | | | |
| chr2 | *PROM2* | 8 | 492 | 17 | PROM2 | prominin 2 | Plasma Membrane | transmembrane recepto | |
| chr9 | *JAK2* | 7 | 197 | 17 | JAK2 | Janus kinase 2 | Cytoplasm | kinase | NS-018, erlotinib/ruxolitinib, baricitinib, ruxolitinib, jak2 inhibitor, SB-1317, tofacitinib, jaktinib, momelotinib, XL019, upadacitinib, methotrexate/tofacitinib, WP1066, BMS-911543, baricitinib/methotrexate, AZD1480, pacritinib, entrectinib, |
| chr15 | *CLN6* | 7 | 189 | 17 | CLN6 | CLN6 transmembrane ER protein | Cytoplasm | other | |
| chr1 | *PSMA5* | 7 | 282 | 17 | PSMA5 | proteasome subunit alpha 5 | Cytoplasm | peptidase | |
| chr17 | *CASC3* | 7 | 211 | 17 | CASC3 | CASC3 exon junction complex subunit | Nucleus | other | |
| chr14 | *GNG2* | 7 | 139 | 17 | GNG2 | G protein subunit gamma 2 | Plasma Membrane | enzyme | |
| chr5 | *CCL28* | 8 | 652 | 17 | CCL28 | C-C motif chemokine ligand 28 | Extracellular Space | cytokine | |
| chr6 | *ZC2HC1B* | 7 | 184 | 17 | ZC2HC1B | zinc finger C2HC-type containing | Other | other | |
| chrX | *ACSL4* | 6 | 175 | 17 | ACSL4 | acyl-CoA synthetase long chain family member 4 | Cytoplasm | enzyme | |
| chr17 | *PGAP3* | 8 | 271 | 17 | PGAP3 | post-GPI attachment to proteins 3 | Cytoplasm | enzyme | |
| chr1 | *CSRP1* | 8 | 327 | 17 | CSRP1 | cysteine and glycine rich protein | Nucleus | other | |
| chr2 | *SLC9A4* | 8 | 251 | 17 | SLC9A4 | solute carrier family 9 member A4 | Plasma Membrane | transporter | |
| chr11 | *OAF* | 8 | 361 | 17 | OAF | out at first homolog | Cytoplasm | other | |
| chr1 | *AGL* | 8 | 201 | 17 | AGL | amylo-alpha-1, 6-glucosidase, 4-alpha-glucanotransferase | Cytoplasm | enzyme | |
| chr19 | *ETHE1* | 8 | 214 | 17 | ETHE1 | ETHE1 persulfide dioxygenase | Cytoplasm | enzyme | |
| chr12 | *ANKRD13A* | 8 | 187 | 17 | ANKRD13A | ankyrin repeat domain 13A | Plasma Membrane | other | |
| chr20 | *ABHD12* | 6 | 169 | 17 | ABHD12 | abhydrolase domain containing | Plasma Membrane | enzyme | |
| chr1 | *RNPEP* | 8 | 226 | 17 | RNPEP | arginyl aminopeptidase | Cytoplasm | peptidase | |
| chr19 | *HAUS8* | 7 | 283 | 17 | HAUS8 | HAUS augmin like complex | Cytoplasm | other | |
| chr3 | *PROS1* | 7 | 224 | 17 | PROS1 | protein S | Extracellular Space | other | |
| chr1 | *TINAGL1* | 8 | 358 | 17 | TINAGL1 | tubulointerstitial nephritis antigen like 1 | Extracellular Space | transporter | |
| chr7 | *KPNA7* | 9 | 205 | 17 | KPNA7 | karyopherin subunit alpha 7 | Other | other | |
| chr19 | *NANOS3* | 8 | 289 | 17 | NANOS3 | nanos C2HC-type zinc finger 3 | Cytoplasm | other | |
| chr1 | *MLK4* | 6 | 232 | 17 | | | | | |
| chr3 | *VEPH1* | 8 | 212 | 17 | VEPH1 | ventricular zone expressed PH domain containing 1 | Nucleus | other | |
| chr7 | *PDGFA* | 9 | 501 | 17 | PDGFA | platelet derived growth factor subunit A | Extracellular Space | growth factor | |
| chr7 | *AC009365.3* | 7 | 206 | 17 | | | | | |
| chr11 | *ME3* | 8 | 314 | 17 | ME3 | malic enzyme 3 | Cytoplasm | enzyme | |
| chr19 | *THOP1* | 8 | 414 | 17 | THOP1 | thimet oligopeptidase 1 | Cytoplasm | peptidase | |
| chr19 | *DEDD2* | 7 | 321 | 17 | DEDD2 | death effector domain containing | Nucleus | other | |
| chr8 | *RP11-32K4.1* | 6 | 206 | 17 | | | | | |
| chr3 | *KPNA4* | 7 | 206 | 17 | KPNA4 | karyopherin subunit alpha 4 | Nucleus | transporter | |
| chr12 | *IFFO1* | 8 | 353 | 17 | IFFO1 | intermediate filament family | Other | other | |
| chr17 | *DHX58* | 7 | 326 | 17 | DHX58 | DExH-box helicase 58 | Cytoplasm | enzyme | |
| chr12 | *P2RX4* | 8 | 341 | 17 | P2RX4 | purinergic receptor P2X 4 | Plasma Membrane | ion channel | |
| chr17 | *MRPL45* | 7 | 224 | 17 | MRPL45 | mitochondrial ribosomal protein | Cytoplasm | other | |
| chr1 | *SLC41A1* | 8 | 422 | 17 | SLC41A1 | solute carrier family 41 member 1 | Plasma Membrane | transporter | |
| chr15 | *KIAA1024* | 8 | 296 | 17 | MINAR1 | membrane integral NOTCH2 associated receptor 1 | Plasma Membrane | other | |
| chr3 | *UQCRC1* | 8 | 337 | 17 | UQCRC1 | ubiquinol-cytochrome c reductase core protein 1 | Cytoplasm | enzyme | |
| chr7 | *FAM126A* | 6 | 169 | 17 | FAM126A | family with sequence similarity 126 member A | Cytoplasm | other | |
| chr1 | *CSF1* | 8 | 391 | 17 | CSF1 | colony stimulating factor 1 | Extracellular Space | cytokine | lacnotuzumab, PD-360324 |
| chr2 | *PLCL1* | 8 | 143 | 17 | PLCL1 | phospholipase C like 1 (inactive) | Cytoplasm | enzyme | quinacrine |
| chr7 | *AGK* | 7 | 156 | 17 | AGK | acylglycerol kinase | Cytoplasm | kinase | |
| chrX | *PPP1R3F* | 8 | 270 | 17 | PPP1R3F | protein phosphatase 1 regulatory subunit 3F | Other | other | |
| chr17 | *XYLT2* | 8 | 340 | 17 | XYLT2 | xylosyltransferase 2 | Cytoplasm | enzyme | |
| chr20 | *ZNF217* | 8 | 345 | 17 | ZNF217 | zinc finger protein 217 | Nucleus | transcription regulator | |
| chr7 | *CALU* | 6 | 183 | 17 | CALU | calumenin | Cytoplasm | other | |
| chr2 | *PIGF* | 6 | 214 | 17 | PIGF | phosphatidylinositol glycan anchor biosynthesis class F | Cytoplasm | enzyme | |
| chr3 | *COL6A5* | 8 | 182 | 17 | COL6A5 | collagen type VI alpha 5 chain | Extracellular Space | other | |
| chr3 | *SPICE1* | 8 | 250 | 17 | SPICE1 | spindle and centriole associated protein 1 | Cytoplasm | other | |
| chr13 | *FOXO1* | 7 | 162 | 17 | FOXO1 | forkhead box O1 | Nucleus | transcription regulator | |
| chr1 | *CYP4X1* | 8 | 203 | 17 | CYP4X1 | cytochrome P450 family 4 subfamily X member 1 | Cytoplasm | enzyme | |
| chr6 | *TCP10* | 7 | 251 | 17 | TCP10/TCP10L2 | t-complex 10 | Cytoplasm | other | |
| chr6 | *EZR* | 5 | 215 | 17 | EZR | ezrin | Plasma Membrane | other | |

| | | | | | | | | | |
|---|---|---|---|---|---|---|---|---|---|
| chr7 | *ABCF2* | 8 | 330 | 17 | ABCF2 | ATP binding cassette subfamily F member 2 | Cytoplasm | transporter | |
| chrX | *HTR2C* | 6 | 179 | 17 | HTR2C | 5-hydroxytryptamine receptor 2C | Plasma Membrane | G-protein coupled recepto | pimavanserin, methysergide, ziprasidone, lorcaserin, agomelatine, ergotamine, doxepin, asenapine, chlorprothixene, trazodone, dihydroergotamine, fenfluramine, quetiapine, opipramol, apomorphine, lisuride, epinastine, clomipramine, fenfluramine/phentermine, paliperidone, sertindole, risperidone, cyproheptadine, blonanserin, vabicaserin, [illegible] |
| chr2 | *RGPD4* | 6 | 220 | 17 | PD4 (includes oth | RANBP2 like and GRIP domain containing 5 | Nucleus | enzyme | |
| chr1 | *RP5-898J17.1* | 8 | 211 | 17 | | | | | |
| chr10 | *TAF5* | 7 | 185 | 17 | TAF5 | TATA-box binding protein associated factor 5 | Nucleus | transcription regulator | |
| chr1 | *LAMB3* | 8 | 311 | 17 | LAMB3 | laminin subunit beta 3 | Extracellular Space | transporter | |
| chr1 | *NECAP2* | 8 | 286 | 17 | NECAP2 | NECAP endocytosis associated 2 | Cytoplasm | other | |
| chr4 | *DDX60L* | 7 | 204 | 17 | DDX60L | DExD/H-box 60 like | Other | other | |
| chr3 | *ENTPD3-AS1* | 8 | 193 | 17 | ENTPD3-AS1 | ENTPD3 antisense RNA 1 | Other | other | |
| chr4 | *COL25A1* | 6 | 180 | 17 | COL25A1 | collagen type XXV alpha 1 chain | Cytoplasm | other | collagenase clostridium |
| chrX | *PTCHD1* | 8 | 176 | 17 | PTCHD1 | patched domain containing 1 | Plasma Membrane | other | |
| chr6 | *ADGB* | 8 | 138 | 17 | ADGB | androglobin | Other | other | |
| chr13 | *FLT1* | 7 | 186 | 17 | FLT1 | fms related tyrosine kinase 1 | Plasma Membrane | kinase | sorafenib, fruquintinib, lapatinib/pazopanib, lenvatinib, sorafenib/sulindac/sunitinib, imatinib/sunitinib, vandetanib, bevacizumab/sorafenib, dexamethasone/lenalidomide/sorafenib, sunitinib, CEP 7055, axitinib, regorafenib, motesanib, icrucumab, 5-azacytidine/sorafenib, pazopanib, famitinib, cabozantinib/erlotinib, SAR103168, SKI-G-801, everolimus/sorafenib, [illegible] |
| chr3 | *ABCC5* | 7 | 160 | 17 | ABCC5 | ATP binding cassette subfamily C member 5 | Plasma Membrane | transporter | |
| chr5 | *TMEM161B-AS1* | 7 | 175 | 17 | TMEM161B-AS1 | TMEM161B antisense RNA 1 | Other | other | |
| chr15 | *VPS13C* | 7 | 200 | 17 | VPS13C | vacuolar protein sorting 13 homolog C | Cytoplasm | other | |
| chr2 | *COL4A4* | 7 | 192 | 17 | COL4A4 | collagen type IV alpha 4 chain | Extracellular Space | other | collagenase clostridium |
| chr9 | *AQP7* | 7 | 365 | 17 | AQP7 | aquaporin 7 | Plasma Membrane | transporter | |
| chr1 | *RNF11* | 6 | 170 | 17 | RNF11 | ring finger protein 11 | Cytoplasm | enzyme | |
| chr3 | *ADAMTS9* | 8 | 157 | 17 | ADAMTS9 | ADAM metallopeptidase with thrombospondin type 1 motif 9 | Extracellular Space | peptidase | |
| chr1 | *ABCD3* | 7 | 183 | 17 | ABCD3 | ATP binding cassette subfamily D member 3 | Cytoplasm | transporter | |
| chr11 | *CTD-3138F19.1* | 7 | 242 | 17 | | | | | |
| chr2 | *BAZ2B* | 6 | 143 | 17 | BAZ2B | bromodomain adjacent to zinc finger domain 2B | Extracellular Space | other | |
| chr11 | *RELT* | 8 | 431 | 17 | RELT | RELT TNF receptor | Plasma Membrane | transmembrane receptor | |
| chr3 | *C3orf18* | 7 | 339 | 17 | C3orf18 | chromosome 3 open reading frame 18 | Other | other | |
| chr15 | *BAHD1* | 8 | 315 | 17 | BAHD1 | bromo adjacent homology domain containing 1 | Nucleus | other | |
| chr12 | *ALDH1L2* | 7 | 186 | 17 | ALDH1L2 | aldehyde dehydrogenase 1 family member L2 | Cytoplasm | enzyme | |
| chr3 | *RP11-407B7.1* | 7 | 186 | 17 | | | | | |
| chr2 | *KCNK3* | 7 | 282 | 17 | KCNK3 | potassium two pore domain channel subfamily K member 3 | Plasma Membrane | ion channel | doxapram, halothane, |

| | | | | | | | | | |
|---|---|---|---|---|---|---|---|---|---|
| chr17 | *FAM104A* | 6 | 196 | 17 | FAM104A | family with sequence similarity 104 member A | Other | other | |
| chr8 | *CPA6* | 6 | 190 | 17 | CPA6 | carboxypeptidase A6 | Extracellular Space | peptidase | |
| chr16 | *CDH15* | 9 | 409 | 17 | CDH15 | cadherin 15 | Plasma Membrane | other | |
| chr9 | *GOLGA2* | 8 | 426 | 17 | GOLGA2 | golgin A2 | Cytoplasm | other | |
| chr9 | *ALDH1A1* | 7 | 168 | 17 | ALDH1A1 | aldehyde dehydrogenase 1 family member A1 | Cytoplasm | enzyme | disulfiram, chlorpropamide |
| chr6 | *POLH* | 7 | 155 | 17 | POLH | DNA polymerase eta | Nucleus | enzyme | |
| chr11 | *RAG1* | 7 | 221 | 17 | RAG1 | recombination activating 1 | Nucleus | enzyme | |
| chr8 | *EEF1D* | 9 | 395 | 17 | EEF1D | eukaryotic translation elongation factor 1 delta | Cytoplasm | translation regulator | |
| chr17 | *KRT16P4* | 7 | 260 | 17 | KRT16P4 | keratin 16 pseudogene 4 | Other | other | |
| chr17 | *RP11-192H23.4* | 8 | 290 | 17 | | | | | |
| chr3 | *AC093415.2* | 7 | 239 | 17 | | | | | |
| chr7 | *AC004009.3* | 8 | 201 | 17 | | | | | |
| chr2 | *ANO7* | 8 | 394 | 17 | ANO7 | anoctamin 7 | Plasma Membrane | transporter | |
| chr3 | *UBXN7* | 6 | 171 | 17 | UBXN7 | UBX domain protein 7 | Nucleus | other | |
| chr5 | *STC2* | 7 | 375 | 17 | STC2 | stanniocalcin 2 | Extracellular Space | other | |
| chr2 | *AC092594.1* | 7 | 173 | 17 | | | | | |
| chr2 | *PID1* | 6 | 179 | 17 | PID1 | phosphotyrosine interaction domain containing 1 | Cytoplasm | other | |
| chr4 | *UNC5C* | 7 | 188 | 17 | UNC5C | unc-5 netrin receptor C | Plasma Membrane | transmembrane receptor | |
| chr19 | *VSTM1* | 8 | 257 | 17 | VSTM1 | V-set and transmembrane domain containing 1 | Other | other | |
| chr1 | *TFB2M* | 7 | 193 | 17 | TFB2M | transcription factor B2, | Cytoplasm | enzyme | |
| chr9 | *SVEP1* | 6 | 175 | 17 | SVEP1 | sushi, von Willebrand factor type A, EGF and pentraxin domain containing 1 | Cytoplasm | other | |
| chr5 | *CPEB4* | 7 | 256 | 17 | CPEB4 | cytoplasmic polyadenylation element binding protein 4 | Plasma Membrane | other | |
| chr20 | *SLC2A10* | 8 | 290 | 17 | SLC2A10 | solute carrier family 2 member 10 | Plasma Membrane | transporter | |
| chr6 | *TNFRSF21* | 7 | 183 | 17 | TNFRSF21 | TNF receptor superfamily | Plasma Membrane | transmembrane receptor | |
| chr10 | *RP11-500G22.2* | 7 | 178 | 17 | | | | | |
| chr12 | *POC1B* | 7 | 172 | 17 | POC1B | POC1 centriolar protein B | Cytoplasm | other | |
| chr9 | *FPGS* | 8 | 466 | 17 | FPGS | folylpolyglutamate synthase | Cytoplasm | enzyme | pralatrexate |
| chr16 | *ALG1* | 7 | 342 | 17 | ALG1 | ALG1 chitobiosyldiphosphodolichol | Cytoplasm | enzyme | |
| chr14 | *TMEM63C* | 6 | 294 | 17 | TMEM63C | transmembrane protein 63C | Other | ion channel | |
| chr1 | *RNF2* | 8 | 234 | 17 | RNF2 | ring finger protein 2 | Nucleus | transcription regulator | |
| chr7 | *AC004538.3* | 8 | 196 | 17 | | | | | |
| chr13 | *GTF2F2* | 7 | 149 | 17 | GTF2F2 | general transcription factor IIF subunit 2 | Nucleus | transcription regulator | |
| chr17 | *RP11-296K13.4* | 7 | 201 | 17 | | | | | |
| chr12 | *KIF21A* | 7 | 260 | 17 | KIF21A | kinesin family member 21A | Cytoplasm | other | |
| chr8 | *UTP23* | 8 | 193 | 17 | UTP23 | UTP23 small subunit processome component | Other | other | |
| chrX | *AF196970.3* | 7 | 190 | 17 | | | | | |
| chr19 | *PNPLA6* | 8 | 267 | 17 | PNPLA6 | patatin like phospholipase domain containing 6 | Cytoplasm | enzyme | |
| chrX | *HS6ST2* | 7 | 151 | 17 | HS6ST2 | heparan sulfate 6-O-sulfotransferase 2 | Plasma Membrane | enzyme | |
| chr2 | *ROCK2* | 6 | 138 | 17 | ROCK2 | Rho associated coiled-coil containing protein kinase 2 | Cytoplasm | kinase | KD025 |
| chr1 | *GNL2* | 6 | 180 | 17 | GNL2 | G protein nucleolar 2 | Nucleus | enzyme | |
| chr1 | *EBNA1BP2* | 7 | 205 | 17 | EBNA1BP2 | EBNA1 binding protein 2 | Nucleus | other | |
| chr11 | *CCDC15* | 7 | 203 | 17 | CCDC15 | coiled-coil domain containing 15 | Cytoplasm | other | |
| chr1 | *ANGEL2* | 6 | 227 | 17 | ANGEL2 | angel homolog 2 | Nucleus | other | |
| chr1 | *ATP1A2* | 7 | 403 | 17 | ATP1A2 | ATPase Na+/K+ transporting subunit alpha 2 | Plasma Membrane | transporter | perphenazine, ethacrynic acid, |
| chr17 | *TEKT1* | 7 | 201 | 17 | TEKT1 | tektin 1 | Cytoplasm | other | |
| chr4 | *RP11-503N18.1* | 7 | 353 | 17 | | | | | |
| chr16 | *GINS2* | 7 | 262 | 17 | GINS2 | GINS complex subunit 2 | Nucleus | other | |
| chr7 | *NOS3* | 8 | 373 | 17 | NOS3 | nitric oxide synthase 3 | Cytoplasm | enzyme | 5,6,7,8-tetrahydrobiopterin, N(G)-monomethyl-D-arginine, GW 273629 |
| chr2 | *AC140481.2* | 6 | 277 | 17 | | | | | |
| chr10 | *SUPV3L1* | 8 | 358 | 17 | SUPV3L1 | Suv3 like RNA helicase | Cytoplasm | enzyme | |
| chr15 | *CTD-2184D3.5* | 8 | 211 | 17 | | | | | |
| chr4 | *CLNK* | 7 | 134 | 17 | CLNK | cytokine dependent hematopoietic cell linker | Cytoplasm | other | |
| chr14 | *RP11-638I2.4* | 7 | 189 | 17 | | | | | |
| chr7 | *PDIA4* | 7 | 317 | 17 | PDIA4 | protein disulfide isomerase family A member 4 | Cytoplasm | enzyme | |
| chr5 | *CAMK4* | 8 | 165 | 17 | CAMK4 | calcium/calmodulin dependent protein kinase IV | Nucleus | kinase | |
| chr8 | *SNTG1* | 7 | 183 | 17 | SNTG1 | syntrophin gamma 1 | Nucleus | other | |
| chrX | *FOXP3* | 7 | 329 | 17 | FOXP3 | forkhead box P3 | Nucleus | transcription regulator | |
| chr12 | *ASB8* | 7 | 266 | 17 | ASB8 | ankyrin repeat and SOCS box containing 8 | Nucleus | transcription regulator | |
| chr12 | *DNM1L* | 7 | 177 | 17 | DNM1L | dynamin 1 like | Cytoplasm | enzyme | |
| chr5 | *ANKRD31* | 9 | 184 | 17 | ANKRD31 | ankyrin repeat domain 31 | Extracellular Space | other | |
| chr11 | *COPB1* | 6 | 161 | 17 | COPB1 | coatomer protein complex subunit beta 1 | Cytoplasm | transporter | |
| chr15 | *ABHD17C* | 7 | 171 | 17 | ABHD17C | abhydrolase domain containing | Plasma Membrane | enzyme | |
| chr19 | *NPHS1* | 8 | 276 | 17 | NPHS1 | NPHS1 adhesion molecule, | Plasma Membrane | other | |
| chr22 | *DRG1* | 6 | 138 | 17 | DRG1 | developmentally regulated GTP binding protein 1 | Cytoplasm | enzyme | |
| chr7 | *HERPUD2* | 7 | 252 | 17 | HERPUD2 | HERPUD family member 2 | Other | other | |

| | | | | | | | | | |
|---|---|---|---|---|---|---|---|---|---|
| chr2 | *ATG16L1* | 7 | 187 | 17 | ATG16L1 | autophagy related 16 like 1 | Cytoplasm | enzyme | |
| chr15 | *ISG20* | 8 | 389 | 17 | ISG20 | interferon stimulated exonuclease gene 20 | Nucleus | enzyme | |
| chr13 | *DZIP1* | 9 | 292 | 17 | DZIP1 | DAZ interacting zinc finger | Cytoplasm | other | |
| chr6 | *DEF6* | 8 | 254 | 17 | DEF6 | DEF6 guanine nucleotide exchange factor | Extracellular Space | other | |
| chr3 | *EMC3* | 7 | 191 | 17 | EMC3 | ER membrane protein complex subunit 3 | Nucleus | other | |
| chr20 | *CDS2* | 6 | 169 | 17 | CDS2 | CDP-diacylglycerol synthase 2 | Cytoplasm | enzyme | |
| chr22 | *CCDC157* | 7 | 293 | 17 | CCDC157 | coiled-coil domain containing 157 | Other | other | |
| chr16 | *RP11-65J21.3* | 7 | 267 | 17 | | | | | |
| chr3 | *HESX1* | 7 | 171 | 17 | HESX1 | HESX homeobox 1 | Nucleus | transcription regulator | |
| chr7 | *PNPLA8* | 7 | 149 | 17 | PNPLA8 | patatin like phospholipase domain containing 8 | Cytoplasm | enzyme | |
| chr2 | *AC093616.4* | 7 | 163 | 17 | | | | | |
| chr2 | *CYS1* | 7 | 349 | 17 | CYS1 | cystin 1 | Cytoplasm | transcription regulator | |
| chr2 | *AP1S3* | 7 | 143 | 17 | AP1S3 | adaptor related protein complex 1 subunit sigma 3 | Cytoplasm | transporter | |
| chr14 | *ATG14* | 7 | 170 | 17 | ATG14 | autophagy related 14 | Cytoplasm | other | |
| chr19 | *TYK2* | 7 | 282 | 17 | TYK2 | tyrosine kinase 2 | Plasma Membrane | kinase | brepocitinib, |
| chr7 | *ACN9* | 7 | 262 | 17 | SDHAF3 | succinate dehydrogenase complex assembly factor 3 | Cytoplasm | other | |
| chr11 | *MPPED2* | 7 | 197 | 17 | MPPED2 | metallophosphoesterase domain containing 2 | Other | other | |
| chrX | *RPS6KA3* | 7 | 150 | 17 | RPS6KA3 | ribosomal protein S6 kinase A3 | Cytoplasm | kinase | PMD-026 |
| chr1 | *SERINC2* | 8 | 271 | 17 | SERINC2 | serine incorporator 2 | Plasma Membrane | transporter | |
| chr11 | *SCYL1* | 8 | 501 | 17 | SCYL1 | SCY1 like pseudokinase 1 | Cytoplasm | kinase | |
| chr12 | *KRT7* | 8 | 453 | 17 | KRT7 | keratin 7 | Cytoplasm | other | |
| chr17 | *RP11-849N15.4* | 7 | 173 | 17 | | | | | |
| chr19 | *PRAM1* | 8 | 318 | 17 | PRAM1 | PML-RARA regulated adaptor molecule 1 | Other | other | |
| chr5 | *ZNF366* | 8 | 205 | 17 | ZNF366 | zinc finger protein 366 | Nucleus | transcription regulator | |
| chr6 | *ARMC2* | 6 | 174 | 17 | ARMC2 | armadillo repeat containing 2 | Extracellular Space | other | |
| chr17 | *WNK4* | 8 | 290 | 17 | WNK4 | WNK lysine deficient protein | Plasma Membrane | kinase | |
| chr5 | *NREP* | 9 | 172 | 17 | NREP | neuronal regeneration related | Cytoplasm | other | |
| chr2 | *AC006227.1* | 8 | 218 | 17 | | | | | |
| chrX | *ARMCX4* | 6 | 152 | 17 | ARMCX4 | armadillo repeat containing X- | Other | other | |
| chr7 | *AC007285.6* | 6 | 192 | 17 | | | | | |
| chr17 | *FLII* | 9 | 434 | 17 | FLII | FLII actin remodeling protein | Nucleus | other | |
| chr2 | *SLC39A10* | 6 | 161 | 17 | SLC39A10 | solute carrier family 39 member | Extracellular Space | transporter | |
| chr22 | *LIMK2* | 6 | 200 | 17 | LIMK2 | LIM domain kinase 2 | Cytoplasm | kinase | cetuximab/encorafenib, binimetinib/encorafenib, encorafenib, binimetinib/cetuximab/encorafenib, binimetinib/encorafen |
| chr2 | *AC012307.3* | 9 | 252 | 17 | | | | | |
| chr5 | *AGGF1* | 6 | 173 | 17 | AGGF1 | angiogenic factor with G-patch and FHA domains 1 | Cytoplasm | other | |
| chr1 | *KLHL21* | 8 | 479 | 17 | KLHL21 | kelch like family member 21 | Cytoplasm | other | |
| chr21 | *AF064858.8* | 6 | 202 | 17 | | | | | |
| chr15 | *CHRFAM7A* | 7 | 234 | 17 | CHRFAM7A | CHRNA7 (exons 5-10) and FAM7A (exons A-E) fusion | Other | other | |
| chr2 | *CREG2* | 7 | 311 | 17 | CREG2 | cellular repressor of E1A stimulated genes 2 | Extracellular Space | enzyme | |
| chr3 | *NPHP3* | 7 | 174 | 17 | NPHP3 | nephrocystin 3 | Extracellular Space | other | |
| chr17 | *RP11-141J13.5* | 6 | 226 | 17 | | | | | |
| chr22 | *AC008132.13* | 6 | 446 | 17 | | | | | |
| chr12 | *ERBB3* | 8 | 254 | 17 | ERBB3 | erb-b2 receptor tyrosine kinase 3 | Plasma Membrane | kinase | afatinib, afatinib/paclitaxel, afatinib/cetuximab, GSK2849330, sym013, REGN1400, MM-111, KTN3379, zirconium-89-GSK2849330, zirconium-89-labeled RO5479599, patritumab, BDTX-189, erlotinib/osimertinib, lumretuzumab, seribantumab, CDX-3379, istiratumab, elgemtumab, sapitinib, MEHD7945A, afatinib/osimertinib |
| chr18 | *TCF4* | 6 | 137 | 17 | TCF4 | transcription factor 4 | Nucleus | transcription regulator | |
| chr4 | *TRMT44* | 6 | 290 | 17 | TRMT44 | tRNA methyltransferase 44 | Other | other | |
| chr16 | *RP11-368J21.3* | 8 | 286 | 17 | | | | | |
| chr21 | *WDR4* | 7 | 319 | 17 | WDR4 | WD repeat domain 4 | Nucleus | other | |
| chr8 | *RP11-149P24.1* | 7 | 157 | 17 | | | | | |
| chr14 | *LINC00617* | 6 | 205 | 17 | TUNAR | TCL1 upstream neural differentiation-associated RNA | Other | other | |
| chr19 | *SLC8A2* | 7 | 255 | 17 | SLC8A2 | solute carrier family 8 member A2 | Cytoplasm | transporter | |
| chr1 | *RP11-231P20.2* | 8 | 215 | 17 | | | | | |
| chr11 | *USP47* | 6 | 294 | 17 | USP47 | ubiquitin specific peptidase 47 | Cytoplasm | peptidase | |

| | | | | | | | | | |
|---|---|---|---|---|---|---|---|---|---|
| chr17 | *ALOX15* | 7 | 377 | 17 | ALOX15 | arachidonate 15-lipoxygenase | Cytoplasm | enzyme | |
| chr11 | *RP11-807H22.7* | 8 | 269 | 17 | | | | | |
| chr4 | *RP11-742B18.1* | 7 | 172 | 17 | | | | | |
| chr2 | *AC083899.3* | 6 | 155 | 17 | | | | | |
| chr15 | *PLA2G4F* | 6 | 304 | 17 | PLA2G4F | phospholipase A2 group IVF | Cytoplasm | enzyme | |
| chr17 | *CASC17* | 8 | 168 | 17 | CASC17 | cancer susceptibility 17 | Other | other | |
| chr1 | *FOXO6* | 8 | 422 | 17 | FOXO6 | forkhead box O6 | Nucleus | transcription regulator | |
| chr21 | *PCP4* | 6 | 171 | 17 | PCP4 | Purkinje cell protein 4 | Cytoplasm | other | |
| chr17 | *KAT7* | 7 | 176 | 17 | KAT7 | lysine acetyltransferase 7 | Nucleus | enzyme | |
| chr7 | *KRBA1* | 8 | 400 | 17 | KRBA1 | KRAB-A domain containing 1 | Other | other | |
| chr2 | *SPHKAP* | 7 | 178 | 17 | SPHKAP | SPHK1 interactor, AKAP domain containing | Cytoplasm | other | |
| chr9 | *GKAP1* | 7 | 193 | 17 | GKAP1 | G kinase anchoring protein 1 | Cytoplasm | other | |
| chr5 | *RP1-167G20.1* | 9 | 459 | 17 | | | | | |
| chr1 | *LRRC41* | 8 | 264 | 17 | LRRC41 | leucine rich repeat containing 41 | Cytoplasm | other | |
| chr21 | *PTTG1IP* | 7 | 278 | 17 | PTTG1IP | PTTG1 interacting protein | Nucleus | other | |
| chr10 | *PDE6C* | 8 | 161 | 17 | PDE6C | phosphodiesterase 6C | Cytoplasm | enzyme | |
| chr13 | *LINC00539* | 6 | 241 | 17 | LINC00539 | long intergenic non-protein coding RNA 539 | Other | other | |
| chr8 | *CTC-756D1.2* | 7 | 251 | 17 | | | | | |
| chr19 | *NLRP8* | 7 | 176 | 17 | NLRP8 | NLR family pyrin domain | Other | other | |
| chr17 | *ASGR2* | 7 | 260 | 17 | ASGR2 | asialoglycoprotein receptor 2 | Plasma Membrane | transmembrane receptor | |
| chr1 | *NBPF1* | 9 | 811 | 17 | PF1 (includes othe | NBPF member 11 | Other | other | |
| chr17 | *TBC1D3P5* | 7 | 319 | 17 | TBC1D3P5 | TBC1 domain family member 3 pseudogene 5 | Other | other | |
| chr2 | *DNAH7* | 5 | 154 | 17 | DNAH7 | dynein axonemal heavy chain 7 | Cytoplasm | other | |
| chr10 | *TMEM180* | 8 | 356 | 17 | MFSD13A | major facilitator superfamily domain containing 13A | Other | other | |
| chr11 | *ATM* | 7 | 176 | 17 | ATM | ATM serine/threonine kinase | Nucleus | kinase | AZD1390, AZD0156, M3541 |
| chr22 | *FOXRED2* | 7 | 255 | 17 | FOXRED2 | FAD dependent oxidoreductase domain containing 2 | Cytoplasm | other | |
| chr7 | *ZNF786* | 7 | 240 | 17 | ZNF786 | zinc finger protein 786 | Nucleus | transcription regulator | |
| chr7 | *CCL24* | 7 | 246 | 17 | CCL24 | C-C motif chemokine ligand 24 | Extracellular Space | cytokine | |
| chr20 | *MTG2* | 8 | 489 | 17 | MTG2 | mitochondrial ribosome associated GTPase 2 | Cytoplasm | enzyme | |
| chr13 | *FAM194B* | 7 | 200 | 17 | ERICH6B | glutamate rich 6B | Other | other | |
| chr1 | *RBBP5* | 7 | 193 | 17 | RBBP5 | RB binding protein 5, histone lysine methyltransferase complex | Nucleus | transcription regulator | |
| chr20 | *DTD1* | 6 | 185 | 17 | DTD1 | D-aminoacyl-tRNA deacylase 1 | Cytoplasm | enzyme | |
| chr10 | *DRGX* | 8 | 505 | 17 | DRGX | dorsal root ganglia homeobox | Nucleus | transcription regulator | |
| chr5 | *ZNF608* | 8 | 186 | 17 | ZNF608 | zinc finger protein 608 | Other | other | |
| chr2 | *SDC1* | 6 | 318 | 17 | SDC1 | syndecan 1 | Plasma Membrane | enzyme | indatuximab |
| chr8 | *ZNF34* | 7 | 237 | 17 | ZNF34 | zinc finger protein 34 | Nucleus | transcription regulator | |
| chr2 | *HADHB* | 5 | 156 | 17 | HADHB | hydroxyacyl-CoA dehydrogenase trifunctional multienzyme complex subunit beta | Cytoplasm | enzyme | |
| chr5 | *DBN1* | 8 | 393 | 17 | DBN1 | drebrin 1 | Cytoplasm | other | |
| chr19 | *C19orf38* | 7 | 189 | 17 | C19orf38 | chromosome 19 open reading frame 38 | Other | other | |
| chr12 | *PAN2* | 8 | 190 | 17 | PAN2 | poly(A) specific ribonuclease subunit PAN2 | Cytoplasm | peptidase | |
| chr12 | *ACVRL1* | 8 | 622 | 17 | ACVRL1 | activin A receptor like type 1 | Plasma Membrane | kinase | PF-3446962, |
| chr14 | *RP11-99E15.2* | 8 | 175 | 17 | | | | | |
| chr14 | *POMT2* | 7 | 183 | 17 | POMT2 | protein O-mannosyltransferase 2 | Cytoplasm | enzyme | |
| chr9 | *PKN3* | 8 | 488 | 17 | PKN3 | protein kinase N3 | Nucleus | kinase | ATU-027 |
| chr14 | *TTLL5* | 6 | 141 | 17 | TTLL5 | tubulin tyrosine ligase like 5 | Cytoplasm | enzyme | |
| chr1 | *ANO7P1* | 8 | 301 | 17 | ANO7L1 | anoctamin 7 like 1 (pseudogene) | Other | other | |
| chr19 | *DMKN* | 7 | 330 | 17 | DMKN | dermokine | Extracellular Space | other | |
| chr11 | *INPPL1* | 8 | 514 | 17 | INPPL1 | inositol polyphosphate phosphatase like 1 | Cytoplasm | phosphatase | |
| chr4 | *C4orf22* | 6 | 206 | 17 | CFAP299 | cilia and flagella associated protein 299 | Cytoplasm | other | |
| chr4 | *CCDC109B* | 6 | 141 | 17 | MCUB | mitochondrial calcium uniporter dominant negative beta subunit | Cytoplasm | other | |
| chr16 | *RP11-488I20.8* | 7 | 203 | 17 | | | | | |
| chr4 | *KIAA1211* | 7 | 180 | 17 | CRACD | capping protein inhibiting regulator of actin dynamics | Other | other | |
| chr2 | *RNF149* | 7 | 214 | 17 | RNF149 | ring finger protein 149 | Cytoplasm | enzyme | |
| chr19 | *TIMM44* | 8 | 314 | 17 | TIMM44 | translocase of inner mitochondrial membrane 44 | Cytoplasm | transporter | |
| chr17 | *AC015922.5* | 7 | 198 | 17 | | | | | |
| chr11 | *AP000770.1* | 8 | 260 | 17 | | | | | |
| chr5 | *PAIP2* | 7 | 197 | 17 | PAIP2 | poly(A) binding protein interacting protein 2 | Cytoplasm | translation regulator | |
| chr1 | *PPP1R8* | 7 | 201 | 17 | PPP1R8 | protein phosphatase 1 regulatory subunit 8 | Nucleus | phosphatase | |
| chr2 | *CUL3* | 5 | 134 | 17 | CUL3 | cullin 3 | Nucleus | enzyme | |
| chr1 | *ACADM* | 7 | 186 | 17 | ACADM | acyl-CoA dehydrogenase medium chain | Cytoplasm | enzyme | |
| chr17 | *CCDC42* | 8 | 286 | 17 | CCDC42 | coiled-coil domain containing 42 | Other | other | |
| chr14 | *AL163953.3* | 7 | 169 | 17 | | | | | |
| chr4 | *FAM149A* | 6 | 179 | 17 | FAM149A | family with sequence similarity 149 member A | Other | other | |
| chr10 | *AIFM2* | 7 | 478 | 17 | AIFM2 | apoptosis inducing factor mitochondria associated 2 | Cytoplasm | enzyme | |
| chr13 | *RBM26* | 6 | 154 | 17 | RBM26 | RNA binding motif protein 26 | Other | other | |
| chr15 | *FAH* | 8 | 393 | 17 | FAH | fumarylacetoacetate hydrolase | Cytoplasm | enzyme | |

| | | | | | | | | | |
|---|---|---|---|---|---|---|---|---|---|
| chr10 | *ASB13* | 7 | 294 | 17 | ASB13 | ankyrin repeat and SOCS box containing 13 | Cytoplasm | other | |
| chrX | *WAS* | 7 | 284 | 17 | WAS | WASP actin nucleation promoting factor | Cytoplasm | other | |
| chr1 | *CATSPER4* | 7 | 270 | 17 | CATSPER4 | cation channel sperm associated | Cytoplasm | transporter | |
| chr3 | *IL17RC* | 7 | 396 | 17 | IL17RC | interleukin 17 receptor C | Plasma Membrane | transmembrane receptor | |
| chr17 | *CASKIN2* | 8 | 335 | 16 | CASKIN2 | CASK interacting protein 2 | Cytoplasm | other | |
| chr6 | *SLC17A5* | 6 | 142 | 16 | SLC17A5 | solute carrier family 17 member 5 | Plasma Membrane | transporter | |
| chr11 | *ZC3H12C* | 8 | 174 | 16 | ZC3H12C | zinc finger CCCH-type containing 12C | Other | other | |
| chr15 | *RP11-1000B6.3* | 6 | 188 | 16 | | | | | |
| chr6 | *SNX9* | 7 | 175 | 16 | SNX9 | sorting nexin 9 | Cytoplasm | transporter | |
| chr20 | *RNF114* | 8 | 278 | 16 | RNF114 | ring finger protein 114 | Cytoplasm | transcription regulator | |
| chr10 | *NUDT13* | 6 | 260 | 16 | NUDT13 | nudix hydrolase 13 | Cytoplasm | enzyme | |
| chr3 | *SAMD7* | 7 | 303 | 16 | SAMD7 | sterile alpha motif domain containing 7 | Other | other | |
| chr19 | *TRMT1* | 8 | 311 | 16 | TRMT1 | tRNA methyltransferase 1 | Extracellular Space | enzyme | |
| chr2 | *NOP58* | 6 | 160 | 16 | NOP58 | NOP58 ribonucleoprotein | Nucleus | enzyme | |
| chr1 | *TIMM17A* | 6 | 267 | 16 | TIMM17A | translocase of inner mitochondrial membrane 17A | Cytoplasm | transporter | |
| chr8 | *RP11-566H8.3* | 8 | 148 | 16 | | | | | |
| chr19 | *ZNF254* | 7 | 174 | 16 | ZNF254 | zinc finger protein 254 | Nucleus | transcription regulator | |
| chr1 | *ARHGAP30* | 8 | 336 | 16 | ARHGAP30 | Rho GTPase activating protein 30 | Cytoplasm | other | |
| chr13 | *GAS6* | 8 | 428 | 16 | GAS6 | growth arrest specific 6 | Extracellular Space | growth factor | AVB-S6-500 |
| chr12 | *EFCAB4B* | 7 | 200 | 16 | CRACR2A | calcium release activated channel regulator 2A | Cytoplasm | other | |
| chr7 | *ADCK2* | 8 | 310 | 16 | ADCK2 | aarF domain containing kinase 2 | Cytoplasm | kinase | |
| chr19 | *ZNF546* | 7 | 177 | 16 | ZNF546 | zinc finger protein 546 | Nucleus | transcription regulator | |
| chr11 | *KCTD21-AS1* | 8 | 289 | 16 | KCTD21-AS1 | KCTD21 antisense RNA 1 | Other | other | |
| chr7 | *AKR1D1* | 7 | 201 | 16 | AKR1D1 | aldo-keto reductase family 1 member D1 | Cytoplasm | enzyme | azelaic acid |
| chr1 | *ANP32E* | 7 | 202 | 16 | ANP32E | acidic nuclear phosphoprotein 32 family member E | Nucleus | other | |
| chr10 | *SEC23IP* | 6 | 159 | 16 | SEC23IP | SEC23 interacting protein | Cytoplasm | other | |
| chr1 | *HEYL* | 7 | 231 | 16 | HEYL | hes related family bHLH transcription factor with YRPW | Nucleus | transcription regulator | |
| chr3 | *SNRK* | 7 | 176 | 16 | SNRK | SNF related kinase | Cytoplasm | kinase | |
| chr17 | *PTRF* | 7 | 201 | 16 | CAVIN1 | caveolae associated protein 1 | Nucleus | transcription regulator | |
| chr8 | *KIAA1456* | 7 | 184 | 16 | TRMT9B | tRNA methyltransferase 9B | Cytoplasm | enzyme | |
| chr11 | *DHCR7* | 6 | 321 | 16 | DHCR7 | 7-dehydrocholesterol reductase | Cytoplasm | enzyme | |
| chr12 | *PFKM* | 8 | 263 | 16 | PFKM | phosphofructokinase, muscle | Cytoplasm | kinase | |
| chr17 | *P4HB* | 8 | 400 | 16 | P4HB | prolyl 4-hydroxylase subunit beta | Cytoplasm | enzyme | |
| chr8 | *CTD-2530N21.4* | 7 | 198 | 16 | | | | | |
| chr5 | *PARP8* | 6 | 134 | 16 | PARP8 | poly(ADP-ribose) polymerase family member 8 | Other | other | |
| chr12 | *UBE2N* | 6 | 158 | 16 | UBE2N | ubiquitin conjugating enzyme E2 | Cytoplasm | enzyme | |
| chr22 | *LZTR1* | 8 | 337 | 16 | LZTR1 | leucine zipper like transcription regulator 1 | Cytoplasm | transcription regulator | |
| chr1 | *ETNK2* | 7 | 302 | 16 | ETNK2 | ethanolamine kinase 2 | Cytoplasm | kinase | |
| chr15 | *NOX5* | 7 | 264 | 16 | NOX5 | NADPH oxidase 5 | Cytoplasm | ion channel | |
| chr1 | *ATP1A4* | 8 | 216 | 16 | ATP1A4 | ATPase Na+/K+ transporting subunit alpha 4 | Plasma Membrane | transporter | perphenazine, ethacrynic acid, |
| chr15 | *GOLGA6C* | 7 | 468 | 16 | GA6A (includes ot | golgin A6 family member A | Other | other | |
| chr10 | *RSU1* | 6 | 167 | 16 | RSU1 | Ras suppressor protein 1 | Cytoplasm | other | |
| chr15 | *CSPG4P8* | 6 | 317 | 16 | | | | | |
| chr17 | *CTC1* | 8 | 235 | 16 | CTC1 | CST telomere replication complex component 1 | Nucleus | other | |
| chr3 | *SPATA16* | 7 | 139 | 16 | SPATA16 | spermatogenesis associated 16 | Cytoplasm | other | |
| chr15 | *RP11-82L7.4* | 6 | 164 | 16 | | | | | |
| chr8 | *UBXN8* | 6 | 216 | 16 | UBXN8 | UBX domain protein 8 | Cytoplasm | other | |
| chr3 | *BTLA* | 6 | 210 | 16 | BTLA | B and T lymphocyte associated | Plasma Membrane | other | TAB004 |
| chr12 | *MAP3K12* | 7 | 352 | 16 | MAP3K12 | mitogen-activated protein kinase kinase kinase 12 | Cytoplasm | kinase | |
| chr19 | *KANK3* | 8 | 222 | 16 | KANK3 | KN motif and ankyrin repeat domains 3 | Cytoplasm | other | |
| chr16 | *AC141586.5* | 8 | 357 | 16 | | | | | |
| chr2 | *SOCS5* | 7 | 191 | 16 | SOCS5 | suppressor of cytokine signaling | Extracellular Space | cytokine | |
| chr16 | *N4BP1* | 7 | 215 | 16 | N4BP1 | NEDD4 binding protein 1 | Cytoplasm | other | |
| chr12 | *ORAI1* | 7 | 253 | 16 | ORAI1 | ORAI calcium release-activated calcium modulator 1 | Plasma Membrane | ion channel | |
| chr10 | *NRP1* | 6 | 157 | 16 | NRP1 | neuropilin 1 | Plasma Membrane | transmembrane receptor | vesencumab |
| chr3 | *TOPBP1* | 7 | 184 | 16 | TOPBP1 | DNA topoisomerase II binding protein 1 | Nucleus | other | |
| chr5 | *IPO11* | 7 | 143 | 16 | IPO11 | importin 11 | Nucleus | transporter | |
| chr11 | *PCNXL3* | 8 | 631 | 16 | PCNX3 | pecanex 3 | Other | other | |
| chr16 | *NMRAL1* | 7 | 328 | 16 | NMRAL1 | NmrA like redox sensor 1 | Nucleus | other | |
| chr11 | *SIAE* | 7 | 181 | 16 | SIAE | sialic acid acetylesterase | Cytoplasm | enzyme | |
| chr12 | *RPSAP52* | 7 | 178 | 16 | RPSAP52 | ribosomal protein SA pseudogene | Other | other | |
| chr17 | *CRHR1* | 7 | 336 | 16 | CRHR1 | corticotropin releasing hormone receptor 1 | Plasma Membrane | G-protein coupled recepto | SSR125543A, CRA1001, CRA0165 |
| chr8 | *SH2D4A* | 8 | 152 | 16 | SH2D4A | SH2 domain containing 4A | Cytoplasm | other | |
| chr11 | *PRRG4* | 7 | 190 | 16 | PRRG4 | proline rich and Gla domain 4 | Plasma Membrane | other | |
| chr8 | *KBTBD11* | 7 | 336 | 16 | KBTBD11 | kelch repeat and BTB domain containing 11 | Other | other | |
| chr22 | *NOL12* | 7 | 245 | 16 | NOL12 | nucleolar protein 12 | Nucleus | other | |
| chr17 | *hsa-mir-6080* | 8 | 328 | 16 | | | | | |
| chr17 | *OSBPL7* | 7 | 398 | 16 | OSBPL7 | oxysterol binding protein like 7 | Cytoplasm | other | |
| chr1 | *FAM212B* | 8 | 296 | 16 | INKA2 | inka box actin regulator 2 | Nucleus | other | |

| | | | | | | | | | |
|---|---|---|---|---|---|---|---|---|---|
| chr7 | *PSMG3-AS1* | 8 | 435 | 16 | PSMG3-AS1 | PSMG3 antisense RNA 1 (head to head) | Other | other | |
| chr4 | *CRMP1* | 6 | 212 | 16 | CRMP1 | collapsin response mediator | Cytoplasm | enzyme | |
| chr3 | *SLC6A1* | 7 | 262 | 16 | SLC6A1 | solute carrier family 6 member 1 | Plasma Membrane | transporter | tiagabine |
| chr12 | *LMBR1L* | 8 | 408 | 16 | LMBR1L | limb development membrane protein 1 like | Plasma Membrane | transmembrane recepto | |
| chr9 | *MPDZ* | 7 | 134 | 16 | MPDZ | multiple PDZ domain crumbs cell polarity complex component | Plasma Membrane | other | |
| chr3 | *PTH1R* | 8 | 421 | 16 | PTH1R | parathyroid hormone 1 receptor | Plasma Membrane | -protein coupled recepto | PTH, abaloparatide, raloxifene/teriparatid |
| chr16 | *RHBDF1* | 8 | 410 | 16 | RHBDF1 | rhomboid 5 homolog 1 | Cytoplasm | other | |
| chr8 | *ATAD2* | 5 | 128 | 16 | ATAD2 | ATPase family AAA domain containing 2 | Nucleus | enzyme | |
| chr9 | *ABO* | 7 | 333 | 16 | ABO | ABO, alpha 1-3-N-acetylgalactosaminyltransferase and alpha 1-3- | Plasma Membrane | enzyme | |
| chr2 | *RABL2A* | 7 | 227 | 16 | RABL2A | RAB, member of RAS oncogene family like 2A | Cytoplasm | enzyme | |
| chr17 | *SPACA3* | 7 | 229 | 16 | SPACA3 | sperm acrosome associated 3 | Cytoplasm | other | |
| chr3 | *ARL13B* | 6 | 197 | 16 | ARL13B | ADP ribosylation factor like GTPase 13B | Extracellular Space | other | |
| chr9 | *CARM1P1* | 8 | 166 | 16 | CARM1P1 | coactivator associated arginine methyltransferase 1 pseudogene | Other | other | |
| chr12 | *ANHX* | 8 | 317 | 16 | ANHX | anomalous homeobox | Other | other | |
| chr7 | *CALCR* | 7 | 194 | 16 | CALCR | calcitonin receptor | Plasma Membrane | -protein coupled recepto | pramlintide |
| chr3 | *SEC22A* | 6 | 177 | 16 | SEC22A | SEC22 homolog A, vesicle trafficking protein | Cytoplasm | transporter | |
| chr1 | *C1orf141* | 7 | 156 | 16 | C1orf141 | chromosome 1 open reading frame 141 | Other | other | |
| chr6 | *TRAM2* | 7 | 208 | 16 | TRAM2 | translocation associated membrane protein 2 | Other | other | |
| chr6 | *RRP36* | 8 | 191 | 16 | RRP36 | ribosomal RNA processing 36 | Nucleus | other | |
| chr16 | *TFAP4* | 7 | 288 | 16 | TFAP4 | transcription factor AP-4 | Nucleus | transcription regulator | |
| chr17 | *MTMR4* | 7 | 211 | 16 | MTMR4 | myotubularin related protein 4 | Cytoplasm | phosphatase | |
| chr10 | *SIRT1* | 7 | 255 | 16 | SIRT1 | sirtuin 1 | Nucleus | transcription regulator | SIRT1 inhibitor III |
| chr15 | *MFAP1* | 6 | 179 | 16 | MFAP1 | microfibril associated protein 1 | Extracellular Space | other | |
| chr10 | *TBATA* | 8 | 444 | 16 | TBATA | thymus, brain and testes | Cytoplasm | other | |
| chr12 | *OTOGL* | 8 | 179 | 16 | OTOGL | otogelin like | Other | other | |
| chr1 | *CC2D1B* | 7 | 269 | 16 | CC2D1B | coiled-coil and C2 domain containing 1B | Nucleus | transcription regulator | |
| chr10 | *FAM175B* | 7 | 172 | 16 | ABRAXAS2 | abraxas 2, BRISC complex | Nucleus | other | |
| chr12 | *CEP290* | 7 | 143 | 16 | CEP290 | centrosomal protein 290 | Cytoplasm | transcription regulator | |
| chrX | *ZNF185* | 6 | 262 | 16 | ZNF185 | zinc finger protein 185 with LIM domain | Plasma Membrane | other | |
| chr3 | *LRRC15* | 6 | 252 | 16 | LRRC15 | leucine rich repeat containing 15 | Plasma Membrane | other | |
| chr4 | *RP11-130F10.1* | 6 | 161 | 16 | | | | | |
| chr12 | *HELB* | 8 | 166 | 16 | HELB | DNA helicase B | Nucleus | enzyme | cyclophosphamide/docetaxel/epirubicin/5-fluorouracil/trastuzumab, chlorambucil/epirubicin, cyclophosphamide/epirubicin/prednisone/rituximab/vincristine, cyclophosphamide/epirubicin/5-fluorouracil/paclitaxel, cyclophosphamide/epirubicin/paclitaxel, cyclophosphamide/docetaxel/epirubicin/5-fluorouracil, cyclophosphamide/epirubicin/prednisone/vincristine, cyclophosphamide/epirubicin/5-[illegible] |
| chr1 | *EXTL1* | 8 | 278 | 16 | EXTL1 | exostosin like glycosyltransferase | Cytoplasm | enzyme | |
| chr11 | *SPI1* | 8 | 588 | 16 | SPI1 | Spi-1 proto-oncogene | Nucleus | transcription regulator | |
| chr14 | *RBM25* | 7 | 163 | 16 | RBM25 | RNA binding motif protein 25 | Nucleus | other | |
| chr17 | *SREBF1* | 7 | 464 | 16 | SREBF1 | sterol regulatory element binding transcription factor 1 | Nucleus | transcription regulator | |
| chr22 | *MB* | 6 | 229 | 16 | MB | myoglobin | Cytoplasm | transporter | |
| chr20 | *CSNK2A1* | 5 | 135 | 16 | CSNK2A1 | casein kinase 2 alpha 1 | Nucleus | kinase | |
| chr22 | *BIK* | 7 | 282 | 16 | BIK | BCL2 interacting killer | Cytoplasm | other | |
| chr3 | *CPB1* | 7 | 205 | 16 | CPB1 | carboxypeptidase B1 | Extracellular Space | peptidase | |
| chr7 | *RAB19* | 8 | 237 | 16 | RAB19 | RAB19, member RAS oncogene family | Cytoplasm | enzyme | |
| chr17 | *DCAF7* | 8 | 145 | 16 | DCAF7 | DDB1 and CUL4 associated | Cytoplasm | other | |
| chr17 | *ARL17B* | 7 | 159 | 16 | ARL17A/ARL17B | ADP ribosylation factor like GTPase 17A | Other | other | |
| chrX | *SEPT6* | 7 | 175 | 16 | SEPTIN6 | septin 6 | Cytoplasm | other | |
| chr16 | *RPL3L* | 8 | 334 | 16 | RPL3L | ribosomal protein L3 like | Cytoplasm | other | |
| chr5 | *ANXA6* | 7 | 194 | 16 | ANXA6 | annexin A6 | Plasma Membrane | ion channel | |
| chr7 | *ANLN* | 7 | 200 | 16 | ANLN | anillin actin binding protein | Cytoplasm | other | |

| | | | | | | | | | |
|---|---|---|---|---|---|---|---|---|---|
| chr1 | *AGTRAP* | 7 | 281 | 16 | AGTRAP | angiotensin II receptor associated protein | Plasma Membrane | G-protein coupled recepto | |
| chr18 | *PSTPIP2* | 7 | 172 | 16 | PSTPIP2 | proline-serine-threonine phosphatase interacting protein 2 | Cytoplasm | other | |
| chr4 | *MTHFD2L* | 6 | 137 | 16 | MTHFD2L | methylenetetrahydrofolate dehydrogenase (NADP+ dependent) 2 like | Cytoplasm | enzyme | |
| chr12 | *RP11-386G11.10* | 7 | 292 | 16 | | | | | |
| chr19 | *ZBTB45* | 8 | 239 | 16 | ZBTB45 | zinc finger and BTB domain containing 45 | Nucleus | transcription regulator | |
| chr12 | *KRT79* | 7 | 459 | 16 | KRT79 | keratin 79 | Extracellular Space | other | |
| chr17 | *SLC9A3R1* | 7 | 335 | 16 | SLC9A3R1 | SLC9A3 regulator 1 | Plasma Membrane | other | |
| chr7 | *CCDC132* | 6 | 194 | 16 | VPS50 | VPS50 subunit of EARP/GARPII complex | Cytoplasm | other | |
| chr13 | *DCUN1D2* | 7 | 276 | 16 | DCUN1D2 | defective in cullin neddylation 1 domain containing 2 | Other | other | |
| chr15 | *STRC* | 7 | 258 | 16 | STRC | stereocilin | Extracellular Space | other | |
| chr14 | *ENTPD5* | 6 | 177 | 16 | ENTPD5 | ectonucleoside triphosphate diphosphohydrolase 5 (inactive) | Cytoplasm | enzyme | |
| chr10 | *SPRN* | 7 | 335 | 16 | SPRN | shadow of prion protein | Nucleus | other | |
| chr20 | *PARD6B* | 6 | 239 | 16 | PARD6B | par-6 family cell polarity regulator | Plasma Membrane | other | |
| chr5 | *CTD-2260A17.2* | 8 | 157 | 16 | | | | | |
| chr16 | *RP11-345M22.1* | 7 | 143 | 16 | | | | | |
| chr6 | *GABRR2* | 5 | 176 | 16 | GABRR2 | gamma-aminobutyric acid type A receptor rho2 subunit | Plasma Membrane | ion channel | triazolam, flurazepam, metyrapone/oxazepam, nitrazepam, diazepam, midazolam, temazepam, quazepam, diphenhydramine/mid |
| chr11 | *OR7E14P* | 7 | 204 | 16 | OR7E14P | olfactory receptor family 7 subfamily E member 14 | Other | other | |
| chr8 | *DPY19L4* | 6 | 140 | 16 | DPY19L4 | dpy-19 like 4 | Other | other | |
| chr3 | *BRPF1* | 7 | 385 | 16 | BRPF1 | bromodomain and PHD finger containing 1 | Nucleus | transporter | |
| chr16 | *CLCN7* | 8 | 241 | 16 | CLCN7 | chloride voltage-gated channel 7 | Plasma Membrane | ion channel | |
| chr8 | *PHYHIP* | 7 | 369 | 16 | PHYHIP | phytanoyl-CoA 2-hydroxylase interacting protein | Cytoplasm | other | |
| chr3 | *CDC25A* | 7 | 197 | 16 | CDC25A | cell division cycle 25A | Nucleus | phosphatase | |
| chr4 | *SETD7* | 7 | 137 | 16 | SETD7 | SET domain containing 7, histone lysine methyltransferase | Nucleus | enzyme | |
| chr14 | *HEATR4* | 6 | 153 | 16 | HEATR4 | HEAT repeat containing 4 | Other | other | |
| chr17 | *WRAP53* | 7 | 238 | 16 | WRAP53 | WD repeat containing antisense to TP53 | Nucleus | other | |
| chr8 | *SLC18A1* | 7 | 175 | 16 | SLC18A1 | solute carrier family 18 member A1 | Plasma Membrane | transporter | reserpine, hydrochlorothiazide/reserpine, reserpine/trichloromethiazide, deserpidine/methyclothiazide, chlorothiazide/reserpine, chlorthalidone/reserpine, polythiazide/reserpine, methamphetamine, deserpidine |
| chr7 | *ITGB8* | 6 | 194 | 16 | ITGB8 | integrin subunit beta 8 | Plasma Membrane | other | |
| chr6 | *AHI1* | 6 | 132 | 16 | AHI1 | Abelson helper integration site 1 | Cytoplasm | other | |
| chr8 | *SCRIB* | 8 | 327 | 16 | SCRIB | scribble planar cell polarity | Cytoplasm | other | |
| chr1 | *TM2D1* | 6 | 168 | 16 | TM2D1 | TM2 domain containing 1 | Plasma Membrane | G-protein coupled recepto | |
| chr19 | *AC074212.3* | 7 | 256 | 16 | | | | | |
| chr19 | *RPS19* | 7 | 305 | 16 | RPS19 | ribosomal protein S19 | Cytoplasm | other | |
| chr17 | *RAPGEFL1* | 7 | 249 | 16 | RAPGEFL1 | Rap guanine nucleotide exchange factor like 1 | Other | other | |
| chr17 | *RP11-806H10.4* | 7 | 319 | 16 | | | | | |
| chr17 | *CTD-2357A8.3* | 7 | 291 | 16 | | | | | |
| chr17 | *C17orf53* | 7 | 278 | 16 | C17orf53 | chromosome 17 open reading frame 53 | Other | other | |
| chr1 | *LIN28A* | 7 | 161 | 16 | LIN28A | lin-28 homolog A | Cytoplasm | other | |
| chr12 | *SLC15A4* | 7 | 231 | 16 | SLC15A4 | solute carrier family 15 member 4 | Cytoplasm | transporter | |
| chr12 | *TBK1* | 6 | 168 | 16 | TBK1 | TANK binding kinase 1 | Cytoplasm | kinase | |
| chr14 | *C14orf79* | 7 | 298 | 16 | CLBA1 | clathrin binding box of aftiphilin containing 1 | Other | other | |
| chr1 | *TTLL10* | 8 | 249 | 16 | TTLL10 | tubulin tyrosine ligase like 10 | Cytoplasm | enzyme | |
| chr1 | *NLRP3* | 8 | 195 | 16 | NLRP3 | NLR family pyrin domain | Cytoplasm | other | BMS-986299 |
| chr5 | *RP11-1277A3.2* | 7 | 153 | 16 | | | | | |
| chr15 | *IDH2* | 7 | 450 | 16 | IDH2 | isocitrate dehydrogenase (NADP(+)) 2 | Cytoplasm | enzyme | enasidenib |
| chr2 | *AC092155.4* | 7 | 216 | 16 | | | | | |
| chr15 | *TMOD2* | 7 | 150 | 16 | TMOD2 | tropomodulin 2 | Cytoplasm | other | |
| chr15 | *CTD-3092A11.1* | 5 | 152 | 16 | | | | | |
| chr7 | *RINT1* | 7 | 198 | 16 | RINT1 | RAD50 interactor 1 | Nucleus | other | |
| chr1 | *RP11-422J8.1* | 7 | 176 | 16 | | | | | |
| chr11 | *ATG2A* | 8 | 433 | 16 | ATG2A | autophagy related 2A | Plasma Membrane | other | |

| | | | | | | | | | |
|---|---|---|---|---|---|---|---|---|---|
| chr1 | *AIDA* | 6 | 190 | 16 | AIDA | axin interactor, dorsalization associated | Cytoplasm | other | |
| chr22 | *ARFGAP3* | 6 | 174 | 16 | ARFGAP3 | ADP ribosylation factor GTPase activating protein 3 | Cytoplasm | transporter | |
| chr1 | *PRCC* | 7 | 277 | 16 | PRCC | proline rich mitotic checkpoint control factor | Nucleus | other | |
| chr4 | *PCGF3* | 7 | 324 | 16 | PCGF3 | polycomb group ring finger 3 | Nucleus | other | |
| chr9 | *RP11-390F4.6* | 6 | 196 | 16 | | | | | |
| chr1 | *GNAI3* | 6 | 257 | 16 | GNAI3 | G protein subunit alpha i3 | Cytoplasm | enzyme | |
| chr5 | *CDH10* | 7 | 338 | 16 | CDH10 | cadherin 10 | Plasma Membrane | other | |
| chr1 | *DESI2* | 6 | 138 | 16 | DESI2 | desumoylating isopeptidase 2 | Cytoplasm | enzyme | |
| chr1 | *EXOSC10* | 7 | 216 | 16 | EXOSC10 | exosome component 10 | Nucleus | kinase | |
| chr8 | *RP11-946L20.4* | 6 | 159 | 16 | | | | | |
| chr1 | *PAFAH2* | 8 | 191 | 16 | PAFAH2 | platelet activating factor acetylhydrolase 2 | Cytoplasm | enzyme | |
| chr2 | *LONRF2* | 8 | 258 | 16 | LONRF2 | LON peptidase N-terminal domain and ring finger 2 | Other | other | |
| chr16 | *MTSS1L* | 8 | 309 | 16 | MTSS2 | MTSS I-BAR domain containing 2 | Plasma Membrane | other | |
| chr7 | *AOC1* | 7 | 265 | 16 | AOC1 | amine oxidase copper containing | Extracellular Space | enzyme | |
| chr1 | *TBX15* | 8 | 371 | 16 | TBX15 | T-box transcription factor 15 | Nucleus | transcription regulator | |
| chr2 | *MCEE* | 6 | 190 | 16 | MCEE | methylmalonyl-CoA epimerase | Cytoplasm | enzyme | |
| chr7 | *AC005154.6* | 7 | 232 | 16 | | | | | |
| chr7 | *SLC13A4* | 8 | 330 | 16 | SLC13A4 | solute carrier family 13 member 4 | Plasma Membrane | transporter | |
| chr6 | *ELOVL2-AS1* | 6 | 196 | 16 | ELOVL2-AS1 | ELOVL2 antisense RNA 1 | Other | other | |
| chr5 | *PKD2L2* | 8 | 183 | 16 | PKD2L2 | polycystin 2 like 2, transient receptor potential cation channel | Plasma Membrane | ion channel | |
| chr9 | *RP11-473E2.4* | 7 | 412 | 16 | | | | | |
| chr5 | *CENPH* | 6 | 154 | 16 | CENPH | centromere protein H | Nucleus | other | |
| chr19 | *ZNF541* | 6 | 212 | 16 | ZNF541 | zinc finger protein 541 | Nucleus | transcription regulator | |
| chr20 | *RSPO4* | 7 | 238 | 16 | RSPO4 | R-spondin 4 | Plasma Membrane | other | |
| chr22 | *A4GALT* | 7 | 268 | 16 | A4GALT | alpha 1,4-galactosyltransferase (P blood group) | Cytoplasm | enzyme | |
| chr2 | *SLC11A1* | 7 | 263 | 16 | SLC11A1 | solute carrier family 11 member 1 | Plasma Membrane | transporter | |
| chr19 | *ERCC2* | 7 | 275 | 16 | ERCC2 | ERCC excision repair 2, TFIIH core complex helicase subunit | Nucleus | enzyme | |
| chr7 | *TMEM213* | 7 | 209 | 16 | TMEM213 | transmembrane protein 213 | Other | other | |
| chr1 | *ARHGEF16* | 8 | 519 | 16 | ARHGEF16 | Rho guanine nucleotide exchange factor 16 | Cytoplasm | other | |
| chr1 | *LINC00862* | 7 | 210 | 16 | LINC00862 | long intergenic non-protein coding RNA 862 | Other | other | |
| chrX | *TENM1* | 7 | 155 | 16 | TENM1 | teneurin transmembrane protein 1 | Plasma Membrane | transmembrane receptor | |
| chr12 | *RP11-81H3.2* | 7 | 162 | 16 | | | | | |
| chr10 | *PTPN20B* | 7 | 171 | 16 | PTPN20 | protein tyrosine phosphatase non-receptor type 20 | Extracellular Space | phosphatase | |
| chr10 | *CDHR1* | 8 | 258 | 16 | CDHR1 | cadherin related family member 1 | Plasma Membrane | other | |
| chr2 | *CYP20A1* | 6 | 160 | 16 | CYP20A1 | cytochrome P450 family 20 subfamily A member 1 | Other | enzyme | |
| chr20 | *SGK2* | 8 | 325 | 16 | SGK2 | serum/glucocorticoid regulated kinase 2 | Cytoplasm | kinase | |
| chr20 | *PXMP4* | 7 | 350 | 16 | PXMP4 | peroxisomal membrane protein 4 | Cytoplasm | other | |
| chr12 | *RSRC2* | 6 | 156 | 16 | RSRC2 | arginine and serine rich coiled- | Other | other | |
| chr2 | *ASIC4* | 7 | 259 | 16 | ASIC4 | acid sensing ion channel subunit family member 4 | Plasma Membrane | ion channel | amiloride, amiloride/hydrochlorothiazide |
| chr9 | *AGPAT2* | 8 | 589 | 16 | AGPAT2 | 1-acylglycerol-3-phosphate O-acyltransferase 2 | Cytoplasm | enzyme | |
| chr2 | *MRPS9* | 6 | 180 | 16 | MRPS9 | mitochondrial ribosomal protein | Cytoplasm | other | |
| chr14 | *LINC00639* | 5 | 155 | 16 | LINC00639 | long intergenic non-protein coding RNA 639 | Other | other | |
| chr1 | *RP11-470E16.1* | 7 | 199 | 16 | | | | | |
| chr11 | *TP53I11* | 8 | 440 | 16 | TP53I11 | tumor protein p53 inducible | Other | other | |
| chr12 | *SLC6A12* | 6 | 248 | 16 | SLC6A12 | solute carrier family 6 member 12 | Plasma Membrane | transporter | tiagabine |
| chr19 | *PRR12* | 7 | 257 | 16 | PRR12 | proline rich 12 | Extracellular Space | other | |
| chr22 | *MORC2* | 6 | 163 | 16 | MORC2 | MORC family CW-type zinc finger | Nucleus | enzyme | |
| chr11 | *PPP2R5B* | 8 | 328 | 16 | PPP2R5B | protein phosphatase 2 regulatory subunit B'beta | Cytoplasm | phosphatase | |
| chr15 | *IREB2* | 7 | 247 | 16 | IREB2 | iron responsive element binding protein 2 | Cytoplasm | translation regulator | |
| chr14 | *ISM2* | 6 | 219 | 16 | ISM2 | isthmin 2 | Other | other | |
| chr10 | *CCDC147* | 7 | 157 | 16 | CFAP58 | cilia and flagella associated | Extracellular Space | other | |
| chr7 | *KIAA1324L* | 7 | 261 | 16 | KIAA1324L | KIAA1324 like | Other | other | |
| chr21 | *BRWD1* | 6 | 150 | 16 | BRWD1 | bromodomain and WD repeat domain containing 1 | Nucleus | transcription regulator | |
| chr1 | *UBQLN4* | 7 | 312 | 16 | UBQLN4 | ubiquilin 4 | Cytoplasm | other | |
| chr14 | *GPHN* | 5 | 116 | 16 | GPHN | gephyrin | Plasma Membrane | enzyme | |
| chr8 | *ADAM32* | 5 | 153 | 16 | ADAM32 | ADAM metallopeptidase domain | Other | peptidase | |
| chr2 | *ANAPC1P1* | 6 | 135 | 16 | ANAPC1P1 | anaphase promoting complex subunit 1 pseudogene 1 | Other | other | |
| chr6 | *GUSBP4* | 7 | 245 | 16 | GUSBP4 | GUSB pseudogene 4 | Other | other | |
| chr14 | *CATSPERB* | 6 | 169 | 16 | CATSPERB | cation channel sperm associated auxiliary subunit beta | Plasma Membrane | other | |
| chr9 | *SNAPC3* | 7 | 171 | 16 | SNAPC3 | small nuclear RNA activating complex polypeptide 3 | Nucleus | other | |
| chr12 | *METTL20* | 7 | 242 | 16 | ETFBKMT | electron transfer flavoprotein subunit beta lysine | Cytoplasm | enzyme | |
| chr17 | *ACAP1* | 7 | 325 | 16 | ACAP1 | ArfGAP with coiled-coil, ankyrin repeat and PH domains 1 | Plasma Membrane | other | |
| chr17 | *AANAT* | 7 | 295 | 16 | AANAT | aralkylamine N-acetyltransferase | Cytoplasm | enzyme | |

| | | | | | | | | | |
|---|---|---|---|---|---|---|---|---|---|
| chr11 | *LINC00301* | 8 | 231 | 16 | LINC00301 | long intergenic non-protein coding RNA 301 | Other | other | |
| chr4 | *GPR78* | 7 | 296 | 16 | GPR78 | G protein-coupled receptor 78 | Plasma Membrane | G-protein coupled recepto | |
| chr7 | *AC017060.1* | 6 | 169 | 16 | | | | | |
| chr11 | *RP11-867G23.8* | 7 | 347 | 16 | | | | | |
| chr16 | *NQO1* | 7 | 204 | 16 | NQO1 | NAD(P)H quinone | Cytoplasm | enzyme | |
| chr3 | *AMOTL2* | 8 | 394 | 16 | AMOTL2 | angiomotin like 2 | Plasma Membrane | other | |
| chr2 | *MBOAT2* | 7 | 144 | 16 | MBOAT2 | membrane bound O-acyltransferase domain | Cytoplasm | enzyme | |
| chr3 | *STT3B* | 6 | 160 | 16 | STT3B | STT3 oligosaccharyltransferase complex catalytic subunit B | Cytoplasm | enzyme | |
| chr15 | *BUB1B* | 5 | 173 | 16 | BUB1B | BUB1 mitotic checkpoint serine/threonine kinase B | Nucleus | kinase | |
| chr12 | *ABCD2* | 7 | 220 | 16 | ABCD2 | ATP binding cassette subfamily D member 2 | Cytoplasm | transporter | |
| chr10 | *SLC16A9* | 7 | 215 | 16 | SLC16A9 | solute carrier family 16 member 9 | Other | other | |
| chr11 | *RRAS2* | 6 | 172 | 16 | RRAS2 | RAS related 2 | Plasma Membrane | enzyme | |
| chr19 | *B3GNT3* | 7 | 231 | 16 | B3GNT3 | UDP-GlcNAc:betaGal beta-1,3-N-acetylglucosaminyltransferase 3 | Cytoplasm | enzyme | |
| chr17 | *NFE2L1* | 7 | 273 | 16 | NFE2L1 | nuclear factor, erythroid 2 like 1 | Nucleus | transcription regulator | |
| chr6 | *BPHL* | 6 | 243 | 16 | BPHL | biphenyl hydrolase like | Cytoplasm | enzyme | |
| chr16 | *AC002310.13* | 7 | 230 | 16 | | | | | |
| chr3 | *MYH15* | 7 | 171 | 16 | MYH15 | myosin heavy chain 15 | Extracellular Space | other | |
| chr12 | *HOXC4* | 8 | 407 | 16 | HOXC4 | homeobox C4 | Nucleus | transcription regulator | |
| chr8 | *PLAT* | 6 | 228 | 16 | PLAT | plasminogen activator, tissue | Extracellular Space | peptidase | 6-aminocaproic acid |
| chr14 | *RPGRIP1* | 6 | 151 | 16 | RPGRIP1 | RPGR interacting protein 1 | Extracellular Space | other | |
| chr17 | *UNC13D* | 8 | 380 | 16 | UNC13D | unc-13 homolog D | Cytoplasm | other | |
| chr11 | *UNC93B1* | 7 | 354 | 16 | UNC93B1 | unc-93 homolog B1, TLR signaling regulator | Cytoplasm | other | |
| chr11 | *RP11-480C22.1* | 6 | 147 | 16 | | | | | |
| chr12 | *KIAA1551* | 6 | 160 | 16 | RESF1 | retroelement silencing factor 1 | Nucleus | other | |
| chr19 | *ZNF320* | 8 | 165 | 16 | ZNF320 | zinc finger protein 320 | Other | other | |
| chr1 | *SEMA6C* | 7 | 430 | 16 | SEMA6C | semaphorin 6C | Plasma Membrane | other | |
| chr12 | *KANSL2* | 7 | 258 | 16 | KANSL2 | KAT8 regulatory NSL complex subunit 2 | Nucleus | other | |
| chr20 | *RP4-553F4.6* | 6 | 245 | 16 | | | | | |
| chr2 | *KIF5C* | 7 | 187 | 16 | KIF5C | kinesin family member 5C | Cytoplasm | enzyme | |
| chr19 | *ZNF808* | 8 | 173 | 16 | ZNF808 | zinc finger protein 808 | Other | other | |
| chr16 | *EIF3CL* | 6 | 208 | 16 | EIF3CL | eukaryotic translation initiation factor 3 subunit C like | Other | other | |
| chr1 | *WNT9A* | 9 | 574 | 16 | WNT9A | Wnt family member 9A | Extracellular Space | other | |
| chr10 | *GPR26* | 7 | 232 | 16 | GPR26 | G protein-coupled receptor 26 | Plasma Membrane | G-protein coupled recepto | |
| chr5 | *RP11-497H16.5* | 6 | 385 | 16 | | | | | |
| chr1 | *CLCNKA* | 7 | 345 | 16 | CLCNKA | chloride voltage-gated channel | Plasma Membrane | ion channel | |
| chr7 | *EFCAB10* | 7 | 180 | 16 | EFCAB10 | EF-hand calcium binding domain | Other | other | |
| chr2 | *FAM168B* | 5 | 163 | 16 | FAM168B | family with sequence similarity 168 member B | Cytoplasm | other | |
| chr1 | *TLR5* | 7 | 281 | 16 | TLR5 | toll like receptor 5 | Plasma Membrane | transmembrane recepto | entolimod |
| chr5 | *C5orf64* | 7 | 155 | 16 | C5orf64 | chromosome 5 open reading frame 64 | Other | other | |
| chr19 | *GRAMD1A* | 7 | 290 | 16 | GRAMD1A | GRAM domain containing 1A | Cytoplasm | transporter | |
| chr16 | *EARS2* | 7 | 202 | 16 | EARS2 | glutamyl-tRNA synthetase 2, mitochondrial | Cytoplasm | enzyme | |
| chr14 | *SLC7A7* | 6 | 153 | 16 | SLC7A7 | solute carrier family 7 member 7 | Plasma Membrane | transporter | |
| chr19 | *CACNG6* | 7 | 199 | 16 | CACNG6 | calcium voltage-gated channel auxiliary subunit gamma 6 | Plasma Membrane | ion channel | |
| chr7 | *RP11-479O9.2* | 7 | 219 | 16 | | | | | |
| chr3 | *TCAIM* | 7 | 166 | 16 | TCAIM | T cell activation inhibitor, mitochondrial | Cytoplasm | other | |
| chr19 | *PEX11G* | 8 | 317 | 16 | PEX11G | peroxisomal biogenesis factor 11 gamma | Cytoplasm | other | |
| chr7 | *IQCA1P1* | 7 | 321 | 16 | IQCA1L | IQ motif containing with AAA domain 1 like | Other | other | |
| chr5 | *CTD-2324F15.2* | 9 | 477 | 16 | | | | | |
| chr11 | *RP11-469N6.1* | 7 | 451 | 16 | | | | | |
| chr7 | *SPDYE3* | 7 | 332 | 16 | SPDYE3 | speedy/RINGO cell cycle regulator family member E3 | Other | other | |
| chr16 | *CSNK2A2* | 7 | 151 | 16 | CSNK2A2 | casein kinase 2 alpha 2 | Cytoplasm | kinase | |
| chr16 | *UBE2I* | 8 | 383 | 16 | UBE2I | ubiquitin conjugating enzyme E2 I | Nucleus | enzyme | |
| chr17 | *SERPINF2* | 8 | 464 | 16 | SERPINF2 | serpin family F member 2 | Extracellular Space | other | |
| chr10 | *MAT1A* | 7 | 262 | 16 | MAT1A | methionine adenosyltransferase | Cytoplasm | enzyme | |
| chr1 | *CLCNKB* | 8 | 391 | 16 | CLCNKB | chloride voltage-gated channel | Plasma Membrane | ion channel | |
| chr19 | *RPS9* | 8 | 338 | 16 | RPS9 | ribosomal protein S9 | Cytoplasm | translation regulator | |
| chr1 | *FLAD1* | 7 | 352 | 16 | FLAD1 | flavin adenine dinucleotide synthetase 1 | Cytoplasm | enzyme | |
| chr7 | *AKR1B1* | 7 | 325 | 16 | AKR1B1 | aldo-keto reductase family 1 member B | Cytoplasm | enzyme | sorbinil |
| chr5 | *ZNF354B* | 7 | 272 | 16 | ZNF354B | zinc finger protein 354B | Other | other | |
| chr17 | *PHF12* | 7 | 250 | 16 | PHF12 | PHD finger protein 12 | Nucleus | transcription regulator | |
| chr16 | *ZNF19* | 7 | 148 | 16 | ZNF19 | zinc finger protein 19 | Nucleus | transcription regulator | |
| chr5 | *LMAN2* | 7 | 336 | 16 | LMAN2 | lectin, mannose binding 2 | Cytoplasm | transporter | |
| chr1 | *HEATR1* | 6 | 215 | 16 | HEATR1 | HEAT repeat containing 1 | Nucleus | other | |
| chr18 | *THEMIS3P* | 7 | 170 | 16 | THEMIS3P | thymocyte selection associated family member 3, pseudogene | Other | other | |
| chr1 | *MAP3K6* | 7 | 412 | 16 | MAP3K6 | mitogen-activated protein kinase kinase kinase 6 | Other | kinase | |
| chr7 | *CCL26* | 7 | 196 | 16 | CCL26 | C-C motif chemokine ligand 26 | Extracellular Space | cytokine | |
| chr12 | *ESYT1* | 8 | 324 | 16 | ESYT1 | extended synaptotagmin 1 | Cytoplasm | other | |

| | | | | | | | | | |
|---|---|---|---|---|---|---|---|---|---|
| chr2 | *C2orf78* | 5 | 135 | 16 | C2orf78 | chromosome 2 open reading frame 78 | Other | other | |
| chrX | *RP4-537K23.4* | 6 | 170 | 16 | | | | | |
| chr16 | *RP11-430C1.1* | 7 | 163 | 16 | | | | | |
| chr1 | *LHX9* | 8 | 367 | 16 | LHX9 | LIM homeobox 9 | Nucleus | transcription regulator | |
| chr18 | *ZNF521* | 5 | 126 | 16 | ZNF521 | zinc finger protein 521 | Nucleus | transcription regulator | |
| chr1 | *LRRC8B* | 7 | 139 | 16 | LRRC8B | leucine rich repeat containing 8 VRAC subunit B | Plasma Membrane | transporter | |
| chr8 | *VPS37A* | 5 | 150 | 16 | VPS37A | VPS37A subunit of ESCRT-I | Cytoplasm | other | |
| chr16 | *CCNYL3* | 7 | 272 | 16 | CCNYL3 | cyclin Y like 3 (pseudogene) | Other | other | |
| chr1 | *MYSM1* | 6 | 159 | 16 | MYSM1 | Myb like, SWIRM and MPN domains 1 | Nucleus | enzyme | |
| chr12 | *KRT78* | 7 | 405 | 16 | KRT78 | keratin 78 | Cytoplasm | other | |
| chr13 | *RNF219* | 5 | 142 | 16 | OBI1 | ORC ubiquitin ligase 1 | Nucleus | enzyme | |
| chr19 | *C3P1* | 7 | 201 | 16 | C3P1 | complement component 3 precursor pseudogene | Other | other | |
| chr17 | *CSNK1D* | 7 | 287 | 16 | CSNK1D | casein kinase 1 delta | Cytoplasm | kinase | |
| chr15 | *HMGN2P46* | 7 | 168 | 16 | HMGN2P46 | high mobility group nucleosomal binding domain 2 pseudogene 46 | Other | other | |
| chr10 | *PWWP2B* | 8 | 408 | 16 | PWWP2B | PWWP domain containing 2B | Nucleus | other | |
| chr15 | *SPINT1* | 6 | 224 | 16 | SPINT1 | serine peptidase inhibitor, Kunitz type 1 | Extracellular Space | other | |
| chr12 | *SENP1* | 7 | 230 | 16 | SENP1 | SUMO specific peptidase 1 | Nucleus | peptidase | |
| chr3 | *DNAJC13* | 6 | 220 | 16 | DNAJC13 | DnaJ heat shock protein family (Hsp40) member C13 | Cytoplasm | other | |
| chr20 | *C20orf197* | 7 | 375 | 16 | C20orf197 | chromosome 20 open reading frame 197 | Other | other | |
| chr12 | *RP11-493L12.3* | 7 | 229 | 16 | | | | | |
| chr1 | *FCRL5* | 7 | 251 | 16 | FCRL5 | Fc receptor like 5 | Plasma Membrane | other | |
| chr12 | *RP11-20E24.1* | 7 | 170 | 16 | | | | | |
| chr1 | *LINC00467* | 7 | 228 | 16 | LINC00467 | long intergenic non-protein coding RNA 467 | Other | other | |
| chr17 | *RP11-453A12.1* | 7 | 164 | 16 | | | | | |
| chr1 | *AMPD2* | 7 | 392 | 16 | AMPD2 | adenosine monophosphate deaminase 2 | Cytoplasm | enzyme | |
| chr21 | *LINC00334* | 7 | 246 | 16 | LINC00334 | long intergenic non-protein coding RNA 334 | Other | other | |
| chr6 | *RP1-292B18.4* | 7 | 146 | 16 | | | | | |
| chrX | *DACH2* | 7 | 180 | 16 | DACH2 | dachshund family transcription factor 2 | Nucleus | transcription regulator | |
| chr6 | *GPR126* | 7 | 145 | 16 | ADGRG6 | adhesion G protein-coupled receptor G6 | Plasma Membrane | -protein coupled recepto | |
| chr15 | *CLPX* | 6 | 154 | 16 | CLPX | caseinolytic mitochondrial matrix peptidase chaperone subunit | Cytoplasm | enzyme | |
| chr18 | *RALBP1* | 6 | 136 | 16 | RALBP1 | ralA binding protein 1 | Cytoplasm | enzyme | |
| chr1 | *TATDN3* | 7 | 190 | 16 | TATDN3 | TatD DNase domain containing 3 | Cytoplasm | other | |
| chr14 | *TGM1* | 6 | 270 | 16 | TGM1 | transglutaminase 1 | Plasma Membrane | enzyme | |
| chr1 | *MOV10* | 7 | 309 | 16 | MOV10 | Mov10 RISC complex RNA | Nucleus | enzyme | |
| chr11 | *VPS51* | 8 | 248 | 16 | VPS51 | VPS51 subunit of GARP complex | Cytoplasm | other | |
| chrX | *COL4A6* | 6 | 200 | 16 | COL4A6 | collagen type IV alpha 6 chain | Extracellular Space | other | collagenase clostridium |
| chr19 | *ZNF101* | 6 | 171 | 16 | ZNF101 | zinc finger protein 101 | Nucleus | transcription regulator | |
| chr3 | *PLA1A* | 7 | 234 | 16 | PLA1A | phospholipase A1 member A | Extracellular Space | enzyme | |
| chr17 | *RP11-697E22.2* | 6 | 151 | 16 | | | | | |
| chr12 | *CSRNP2* | 7 | 326 | 16 | CSRNP2 | cysteine and serine rich nuclear protein 2 | Nucleus | transcription regulator | |
| chr19 | *ZNF345* | 6 | 138 | 16 | ZNF345 | zinc finger protein 345 | Nucleus | transcription regulator | |
| chr4 | *C4orf29* | 5 | 122 | 16 | ABHD18 | abhydrolase domain containing | Other | other | |
| chr19 | *CREB3L3* | 7 | 224 | 16 | CREB3L3 | cAMP responsive element binding protein 3 like 3 | Cytoplasm | transcription regulator | |
| chr1 | *BRINP2* | 7 | 207 | 16 | BRINP2 | BMP/retinoic acid inducible neural specific 2 | Cytoplasm | other | |
| chr19 | *STRN4* | 7 | 338 | 16 | STRN4 | striatin 4 | Cytoplasm | other | |
| chr5 | *RP11-325I22.2* | 7 | 631 | 16 | | | | | |
| chr1 | *MSH4* | 6 | 152 | 16 | MSH4 | mutS homolog 4 | Nucleus | enzyme | |
| chr11 | *ABCG4* | 7 | 287 | 16 | ABCG4 | ATP binding cassette subfamily G member 4 | Plasma Membrane | transporter | |
| chr3 | *MASP1* | 6 | 198 | 16 | MASP1 | mannan binding lectin serine peptidase 1 | Extracellular Space | peptidase | |
| chr16 | *MEFV* | 7 | 245 | 16 | MEFV | MEFV innate immuity regulator, | Nucleus | other | |
| chr15 | *CD276* | 8 | 300 | 16 | CD276 | CD276 molecule | Plasma Membrane | other | MGC018, MGA271, MGD009 |
| chr12 | *ARNTL2* | 6 | 186 | 16 | ARNTL2 | aryl hydrocarbon receptor nuclear translocator like 2 | Nucleus | transcription regulator | |
| chr15 | *RP11-24M17.4* | 7 | 351 | 16 | | | | | |
| chr4 | *PLAC8* | 6 | 173 | 16 | PLAC8 | placenta associated 8 | Nucleus | other | |
| chr17 | *PSMD11* | 7 | 165 | 16 | PSMD11 | proteasome 26S subunit, non-ATPase 11 | Cytoplasm | other | |
| chr9 | *LHX6* | 8 | 331 | 16 | LHX6 | LIM homeobox 6 | Nucleus | transcription regulator | |
| chr1 | *BTF3L4* | 7 | 153 | 16 | BTF3L4 | basic transcription factor 3 like 4 | Other | other | |
| chr10 | *ANAPC16* | 7 | 269 | 16 | ANAPC16 | anaphase promoting complex subunit 16 | Nucleus | other | |
| chr10 | *CWF19L1* | 6 | 179 | 16 | CWF19L1 | CWF19 like cell cycle control | Other | other | |
| chr2 | *SMPD4* | 7 | 309 | 16 | SMPD4 | sphingomyelin | Cytoplasm | enzyme | |
| chr3 | *NR1D2* | 6 | 195 | 16 | NR1D2 | nuclear receptor subfamily 1 group D member 2 | Nucleus | d-dependent nuclear rec | |
| chr12 | *RP11-167N24.3* | 7 | 149 | 16 | | | | | |
| chr5 | *TTC23L* | 7 | 427 | 16 | TTC23L | tetratricopeptide repeat domain 23 like | Other | other | |

| | | | | | | | | | |
|---|---|---|---|---|---|---|---|---|---|
| chr5 | *CLTB* | 7 | 268 | 16 | CLTB | clathrin light chain B | Plasma Membrane | other | |
| chr4 | *ABCG2* | 5 | 123 | 16 | ABCG2 | ATP binding cassette subfamily G member 2 (Junior blood group) | Plasma Membrane | transporter | |
| chrX | *FIGF* | 6 | 144 | 16 | VEGFD | vascular endothelial growth factor | Extracellular Space | growth factor | |
| chr8 | *RGS20* | 6 | 170 | 16 | RGS20 | regulator of G protein signaling | Cytoplasm | enzyme | |
| chr5 | *CTD-2281M20.1* | 8 | 175 | 15 | | | | | |
| chr3 | *CMC1* | 6 | 137 | 15 | CMC1 | C-X9-C motif containing 1 | Cytoplasm | other | |
| chr20 | *YTHDF1* | 7 | 463 | 15 | YTHDF1 | YTH N6-methyladenosine RNA binding protein 1 | Other | other | |
| chr5 | *ITGA2* | 8 | 172 | 15 | ITGA2 | integrin subunit alpha 2 | Plasma Membrane | transmembrane recepto | |
| chr1 | *PTPRC* | 6 | 146 | 15 | PTPRC | protein tyrosine phosphatase receptor type C | Plasma Membrane | phosphatase | At 211 MAb BC8-B10, 111 In-BC8 |
| chr3 | *KPNA1* | 6 | 187 | 15 | KPNA1 | karyopherin subunit alpha 1 | Nucleus | transporter | |
| chr8 | *MTFR1* | 6 | 144 | 15 | MTFR1 | mitochondrial fission regulator 1 | Cytoplasm | other | |
| chr6 | *MYO6* | 5 | 142 | 15 | MYO6 | myosin VI | Cytoplasm | other | |
| chr11 | *TRIM3* | 8 | 272 | 15 | TRIM3 | tripartite motif containing 3 | Cytoplasm | enzyme | |
| chr10 | *PLA2G12B* | 7 | 225 | 15 | PLA2G12B | phospholipase A2 group XIIB | Extracellular Space | enzyme | |
| chr14 | *FBLN5* | 6 | 199 | 15 | FBLN5 | fibulin 5 | Extracellular Space | other | |
| chr14 | *SLC35F4* | 6 | 143 | 15 | SLC35F4 | solute carrier family 35 member | Other | other | |
| chr19 | *ZNF665* | 7 | 227 | 15 | ZNF665 | zinc finger protein 665 | Other | other | |
| chr20 | *SALL4* | 8 | 309 | 15 | SALL4 | spalt like transcription factor 4 | Nucleus | transcription regulator | |
| chr3 | *GUCA1C* | 6 | 160 | 15 | GUCA1C | guanylate cyclase activator 1C | Cytoplasm | other | |
| chr5 | *RP11-152K4.2* | 8 | 431 | 15 | | | | | |
| chr15 | *CTD-2034I4.1* | 8 | 189 | 15 | | | | | |
| chr14 | *LRRC16B* | 6 | 293 | 15 | CARMIL3 | capping protein regulator and myosin 1 linker 3 | Plasma Membrane | other | |
| chr11 | *RIC3* | 6 | 208 | 15 | RIC3 | RIC3 acetylcholine receptor chaperone | Cytoplasm | other | |
| chr10 | *C10orf71* | 7 | 436 | 15 | C10orf71 | chromosome 10 open reading frame 71 | Other | other | |
| chr13 | *MPHOSPH8* | 6 | 183 | 15 | MPHOSPH8 | M-phase phosphoprotein 8 | Nucleus | transcription regulator | |
| chr10 | *UROS* | 7 | 178 | 15 | UROS | uroporphyrinogen III synthase | Cytoplasm | enzyme | |
| chr1 | *RP11-99J16__A.2* | 7 | 222 | 15 | | | | | |
| chr7 | *CCT6P3* | 7 | 128 | 15 | CCT6P3 | chaperonin containing TCP1 subunit 6 pseudogene 3 | Other | other | |
| chr1 | *SMAP2* | 7 | 168 | 15 | SMAP2 | small ArfGAP2 | Cytoplasm | other | |
| chr3 | *SETD5* | 6 | 135 | 15 | SETD5 | SET domain containing 5 | Nucleus | enzyme | |
| chr11 | *ATL3* | 6 | 155 | 15 | ATL3 | atlastin GTPase 3 | Cytoplasm | other | |
| chr1 | *DIO1* | 6 | 195 | 15 | DIO1 | iodothyronine deiodinase 1 | Cytoplasm | enzyme | propylthiouracil |
| chrX | *DCAF8L2* | 6 | 159 | 15 | DCAF8L2 | DDB1 and CUL4 associated factor 8 like 2 | Other | other | |
| chr15 | *TEX9* | 7 | 180 | 15 | TEX9 | testis expressed 9 | Other | other | |
| chr2 | *ALS2CR11* | 6 | 190 | 15 | C2CD6 | C2 calcium dependent domain containing 6 | Other | other | |
| chr15 | *RMDN3* | 7 | 214 | 15 | RMDN3 | regulator of microtubule dynamics | Cytoplasm | other | |
| chr1 | *MTR* | 7 | 241 | 15 | MTR | 5-methyltetrahydrofolate-homocysteine methyltransferase | Cytoplasm | enzyme | mecobalamin, hydroxocobalamin, cyanocobalamin |
| chr11 | *CCS* | 7 | 389 | 15 | CCS | copper chaperone for superoxide dismutase | Cytoplasm | enzyme | |
| chr10 | *EPC1* | 5 | 137 | 15 | EPC1 | enhancer of polycomb homolog 1 | Nucleus | transcription regulator | |
| chr19 | *LINC00661* | 7 | 316 | 15 | LINC00661 | long intergenic non-protein coding RNA 661 | Other | other | |
| chr11 | *TTC12* | 7 | 196 | 15 | TTC12 | tetratricopeptide repeat domain | Cytoplasm | other | |
| chr2 | *INO80D* | 5 | 143 | 15 | INO80D | INO80 complex subunit D | Nucleus | other | |
| chr3 | *EFHB* | 6 | 190 | 15 | EFHB | EF-hand domain family member | Other | other | |
| chr19 | *KEAP1* | 7 | 183 | 15 | KEAP1 | kelch like ECH associated protein | Cytoplasm | transcription regulator | dimethyl fumarate |
| chr3 | *FAM107A* | 7 | 214 | 15 | FAM107A | family with sequence similarity 107 member A | Nucleus | other | |
| chr16 | *TEPP* | 7 | 164 | 15 | TEPP | testis, prostate and placenta expressed | Other | other | |
| chr10 | *RP11-672F9.1* | 8 | 320 | 15 | | | | | |
| chr3 | *KRBOX1* | 7 | 143 | 15 | KRBOX1 | KRAB box domain containing 1 | Other | other | |
| chr3 | *CCDC14* | 6 | 193 | 15 | CCDC14 | coiled-coil domain containing 14 | Cytoplasm | other | |
| chr16 | *BBS2* | 5 | 134 | 15 | BBS2 | Bardet-Biedl syndrome 2 | Cytoplasm | other | |
| chr18 | *RP11-595B24.1* | 6 | 142 | 15 | | | | | |
| chr22 | *SMTN* | 8 | 323 | 15 | SMTN | smoothelin | Extracellular Space | other | |
| chr12 | *MVK* | 6 | 358 | 15 | MVK | mevalonate kinase | Cytoplasm | kinase | |
| chr11 | *PRCP* | 6 | 184 | 15 | PRCP | prolylcarboxypeptidase | Cytoplasm | peptidase | |
| chr19 | *ZNF160* | 7 | 148 | 15 | ZNF160 | zinc finger protein 160 | Nucleus | transcription regulator | |
| chr4 | *G3BP2* | 5 | 149 | 15 | G3BP2 | G3BP stress granule assembly factor 2 | Cytoplasm | enzyme | |
| chr22 | *NDUFA6-AS1* | 7 | 200 | 15 | NDUFA6-DT | NDUFA6 divergent transcript | Other | other | |
| chr8 | *MTDH* | 6 | 125 | 15 | MTDH | metadherin | Cytoplasm | transcription regulator | |
| chr9 | *NDUFA8* | 6 | 234 | 15 | NDUFA8 | NADH:ubiquinone oxidoreductase subunit A8 | Cytoplasm | enzyme | |
| chr2 | *KCNS3* | 5 | 177 | 15 | KCNS3 | potassium voltage-gated channel modifier subfamily S member 3 | Plasma Membrane | ion channel | |
| chr14 | *HECTD1* | 7 | 136 | 15 | HECTD1 | HECT domain E3 ubiquitin protein ligase 1 | Other | enzyme | |
| chr16 | *CTB-134H23.3* | 7 | 252 | 15 | | | | | |
| chrX | *RAI2* | 7 | 188 | 15 | RAI2 | retinoic acid induced 2 | Other | other | |
| chr3 | *RP11-702L6.4* | 7 | 188 | 15 | | | | | |
| chr3 | *TOPAZ1* | 6 | 141 | 15 | TOPAZ1 | testis and ovary specific PAZ domain containing 1 | Other | other | |
| chr13 | *LHFP* | 5 | 140 | 15 | LHFPL6 | LHFPL tetraspan subfamily member 6 | Other | other | |
| chr8 | *CASC11* | 8 | 150 | 15 | CASC11 | cancer susceptibility 11 | Other | other | |

| | | | | | | | | | |
|---|---|---|---|---|---|---|---|---|---|
| chr1 | *ADORA3* | 7 | 277 | 15 | ADORA3 | adenosine A3 receptor | Plasma Membrane | G-protein coupled recepto | caffeine, clofarabine, aminothiadiazole, theophylline, theophylline/tretinoin/valproic acid, dyphylline, CF102, |
| chr17 | *TRIM25* | 7 | 235 | 15 | TRIM25 | tripartite motif containing 25 | Cytoplasm | transcription regulator | |
| chr10 | *TUBGCP2* | 7 | 337 | 15 | TUBGCP2 | tubulin gamma complex associated protein 2 | Cytoplasm | peptidase | |
| chr2 | *C2orf15* | 7 | 177 | 15 | C2orf15 | chromosome 2 open reading frame 15 | Other | other | |
| chr8 | *LPL* | 7 | 151 | 15 | LPL | lipoprotein lipase | Cytoplasm | enzyme | lovastatin/niacin, nicotinic acid/pioglitazone, atorvastatin/niacin, nicotinic acid, |
| chr1 | *WRAP73* | 8 | 429 | 15 | WRAP73 | WD repeat containing, antisense to TP73 | Nucleus | other | |
| chr19 | *SH2D3A* | 8 | 320 | 15 | SH2D3A | SH2 domain containing 3A | Cytoplasm | other | |
| chr18 | *RBBP8* | 5 | 126 | 15 | RBBP8 | RB binding protein 8, | Nucleus | enzyme | |
| chr2 | *CYP26B1* | 7 | 419 | 15 | CYP26B1 | cytochrome P450 family 26 subfamily B member 1 | Cytoplasm | enzyme | |
| chr3 | *EXOG* | 6 | 198 | 15 | EXOG | exo/endonuclease G | Cytoplasm | enzyme | |
| chr1 | *GPR153* | 8 | 522 | 15 | GPR153 | G protein-coupled receptor 153 | Plasma Membrane | G-protein coupled recepto | |
| chr3 | *OXSR1* | 7 | 206 | 15 | OXSR1 | oxidative stress responsive | Nucleus | kinase | |
| chr19 | *MED25* | 6 | 278 | 15 | MED25 | mediator complex subunit 25 | Nucleus | other | |
| chr11 | *MS4A15* | 7 | 351 | 15 | MS4A15 | membrane spanning 4-domains | Other | other | |
| chr19 | *SHKBP1* | 7 | 317 | 15 | SHKBP1 | SH3KBP1 binding protein 1 | Other | other | |
| chr16 | *RP11-140I24.1* | 7 | 169 | 15 | | | | | |
| chr18 | *RP11-176N18.2* | 6 | 172 | 15 | | | | | |
| chrX | *NYX* | 6 | 163 | 15 | NYX | nyctalopin | Extracellular Space | other | |
| chrX | *FTX* | 6 | 166 | 15 | FTX | FTX transcript, XIST regulator | Other | other | |
| chrX | *XIAP* | 5 | 139 | 15 | XIAP | X-linked inhibitor of apoptosis | Cytoplasm | enzyme | GDC-0152, idronoxil, BI-891065, birinapant, ASTX660, XIAP antisense oligonucleotide AEG35156, GDC- |
| chr1 | *RP11-131L23.1* | 7 | 213 | 15 | | | | | |
| chr20 | *RBBP8NL* | 8 | 691 | 15 | RBBP8NL | RBBP8 N-terminal like | Extracellular Space | other | |
| chr16 | *KCNG4* | 8 | 314 | 15 | KCNG4 | potassium voltage-gated channel modifier subfamily G member 4 | Plasma Membrane | ion channel | |
| chr7 | *AC004840.9* | 7 | 389 | 15 | | | | | |
| chrX | *MBTPS2* | 6 | 147 | 15 | MBTPS2 | membrane bound transcription factor peptidase, site 2 | Cytoplasm | peptidase | |
| chr10 | *PIP4K2A* | 6 | 176 | 15 | PIP4K2A | phosphatidylinositol-5-phosphate 4-kinase type 2 alpha | Cytoplasm | kinase | |
| chr15 | *SNX33* | 7 | 621 | 15 | SNX33 | sorting nexin 33 | Cytoplasm | other | |
| chr17 | *AATK-AS1* | 8 | 329 | 15 | PVALEF | parvalbumin like EF-hand | Other | other | |
| chr6 | *REV3L* | 5 | 116 | 15 | REV3L | REV3 like, DNA directed polymerase zeta catalytic subunit | Nucleus | enzyme | |
| chr6 | *RP11-716O23.1* | 7 | 170 | 15 | | | | | |
| chr17 | *RP11-1055B8.2* | 7 | 344 | 15 | | | | | |
| chr14 | *ZFYVE1* | 6 | 163 | 15 | ZFYVE1 | zinc finger FYVE-type containing | Cytoplasm | other | |
| chr6 | *RP11-439L18.3* | 7 | 156 | 15 | | | | | |
| chr2 | *PPP1R7* | 6 | 222 | 15 | PPP1R7 | protein phosphatase 1 regulatory subunit 7 | Nucleus | phosphatase | |
| chr1 | *EFNA3* | 7 | 466 | 15 | EFNA3 | ephrin A3 | Plasma Membrane | kinase | |
| chr4 | *WDR19* | 7 | 173 | 15 | WDR19 | WD repeat domain 19 | Extracellular Space | other | |
| chr15 | *NUTM1* | 7 | 213 | 15 | NUTM1 | NUT midline carcinoma family member 1 | Other | other | |
| chr4 | *RP11-341G5.1* | 6 | 152 | 15 | | | | | |
| chr1 | *FAM131C* | 6 | 312 | 15 | FAM131C | family with sequence similarity 131 member C | Other | other | |
| chr16 | *MLKL* | 7 | 229 | 15 | MLKL | mixed lineage kinase domain like pseudokinase | Cytoplasm | kinase | |
| chr11 | *MOGAT2* | 6 | 340 | 15 | MOGAT2 | monoacylglycerol O- | Cytoplasm | enzyme | |
| chr9 | *RP11-63P12.6* | 6 | 152 | 15 | | | | | |
| chr22 | *RAC2* | 6 | 241 | 15 | RAC2 | Rac family small GTPase 2 | Cytoplasm | enzyme | |
| chr10 | *PRKCQ* | 7 | 165 | 15 | PRKCQ | protein kinase C theta | Cytoplasm | kinase | ingenol mebutate, sotrastaurin |
| chr6 | *FAM135A* | 5 | 128 | 15 | FAM135A | family with sequence similarity 135 member A | Other | enzyme | |
| chr2 | *NRP2* | 6 | 188 | 15 | NRP2 | neuropilin 2 | Plasma Membrane | kinase | |
| chr9 | *PMPCA* | 7 | 461 | 15 | PMPCA | peptidase, mitochondrial processing alpha subunit | Cytoplasm | peptidase | |
| chr19 | *AC008686.1* | 6 | 231 | 15 | | | | | |
| chr9 | *FBXW2* | 6 | 204 | 15 | FBXW2 | F-box and WD repeat domain containing 2 | Cytoplasm | enzyme | |
| chrX | *OFD1* | 6 | 168 | 15 | OFD1 | OFD1 centriole and centriolar satellite protein | Plasma Membrane | other | |
| chr19 | *SEMA6B* | 7 | 272 | 15 | SEMA6B | semaphorin 6B | Plasma Membrane | other | |
| chr7 | *EIF4H* | 7 | 208 | 15 | EIF4H | eukaryotic translation initiation factor 4H | Cytoplasm | translation regulator | |
| chr11 | *ATP5L* | 6 | 162 | 15 | ATP5MG | ATP synthase membrane subunit | Cytoplasm | enzyme | |
| chr11 | *DCHS1* | 8 | 474 | 15 | DCHS1 | dachsous cadherin-related 1 | Plasma Membrane | other | |
| chr1 | *FAM76A* | 7 | 159 | 15 | FAM76A | family with sequence similarity 76 member A | Nucleus | other | |
| chr9 | *FOCAD* | 6 | 132 | 15 | FOCAD | focadhesin | Plasma Membrane | other | |

| | | | | | | | | | |
|---|---|---|---|---|---|---|---|---|---|
| chr18 | *SMCHD1* | 5 | 221 | 15 | SMCHD1 | structural maintenance of chromosomes flexible hinge domain containing 1 | Nucleus | enzyme | |
| chr3 | *GRK7* | 8 | 267 | 15 | GRK7 | G protein-coupled receptor | Cytoplasm | kinase | |
| chr3 | *GBE1* | 7 | 145 | 15 | GBE1 | 1,4-alpha-glucan branching enzyme 1 | Cytoplasm | enzyme | |
| chr10 | *AGAP6* | 6 | 358 | 15 | AP5 (includes oth | ArfGAP with GTPase domain, ankyrin repeat and PH domain 5 | Other | other | |
| chr19 | *CKM* | 7 | 248 | 15 | CKM | creatine kinase, M-type | Cytoplasm | kinase | |
| chr3 | *MORC1* | 7 | 175 | 15 | MORC1 | MORC family CW-type zinc finger | Nucleus | other | |
| chr16 | *CTD-3037G24.3* | 7 | 197 | 15 | | | | | |
| chr1 | *FUCA1* | 6 | 166 | 15 | FUCA1 | alpha-L-fucosidase 1 | Cytoplasm | enzyme | |
| chr16 | *CAPN15* | 8 | 359 | 15 | CAPN15 | calpain 15 | Other | peptidase | |
| chr9 | *SPTLC1* | 6 | 183 | 15 | SPTLC1 | serine palmitoyltransferase long chain base subunit 1 | Cytoplasm | enzyme | |
| chr1 | *DCST2* | 7 | 416 | 15 | DCST2 | DC-STAMP domain containing 2 | Other | other | |
| chr7 | *SUMF2* | 7 | 294 | 15 | SUMF2 | sulfatase modifying factor 2 | Cytoplasm | other | |
| chr5 | *RP11-43F13.1* | 7 | 673 | 15 | | | | | |
| chr11 | *PLEKHB1* | 7 | 391 | 15 | PLEKHB1 | pleckstrin homology domain containing B1 | Cytoplasm | other | |
| chr3 | *RP11-438D8.2* | 6 | 278 | 15 | | | | | |
| chr17 | *FN3K* | 7 | 265 | 15 | FN3K | fructosamine 3 kinase | Cytoplasm | kinase | |
| chr6 | *LY86* | 7 | 172 | 15 | LY86 | lymphocyte antigen 86 | Plasma Membrane | other | |
| chr15 | *GCNT3* | 7 | 131 | 15 | GCNT3 | glucosaminyl (N-acetyl) transferase 3, mucin type | Plasma Membrane | enzyme | |
| chr2 | *TMEM247* | 8 | 276 | 15 | TMEM247 | transmembrane protein 247 | Cytoplasm | other | |
| chr12 | *SP7* | 7 | 437 | 15 | SP7 | Sp7 transcription factor | Nucleus | transcription regulator | |
| chr2 | *LINC00299* | 6 | 179 | 15 | LINC00299 | long intergenic non-protein coding RNA 299 | Other | other | |
| chr5 | *BTNL3* | 6 | 216 | 15 | BTNL3 | butyrophilin like 3 | Other | other | |
| chr1 | *KCNA2* | 7 | 235 | 15 | KCNA2 | potassium voltage-gated channel subfamily A member 2 | Plasma Membrane | ion channel | dalfampridine |
| chr2 | *FBXO41* | 7 | 344 | 15 | FBXO41 | F-box protein 41 | Cytoplasm | other | |
| chr16 | *RP11-462G12.2* | 6 | 169 | 15 | | | | | |
| chr16 | *RP11-615I2.1* | 6 | 217 | 15 | | | | | |
| chr20 | *C20orf196* | 6 | 156 | 15 | SHLD1 | shieldin complex subunit 1 | Nucleus | other | |
| chr2 | *CRIM1* | 6 | 147 | 15 | CRIM1 | cysteine rich transmembrane BMP regulator 1 | Extracellular Space | kinase | |
| chr7 | *RP13-492C18.2* | 6 | 181 | 15 | | | | | |
| chr22 | *GGT5* | 6 | 279 | 15 | GGT5 | gamma-glutamyltransferase 5 | Plasma Membrane | enzyme | |
| chr3 | *SERPINI1* | 6 | 330 | 15 | SERPINI1 | serpin family I member 1 | Extracellular Space | other | |
| chr3 | *NCKIPSD* | 7 | 261 | 15 | NCKIPSD | NCK interacting protein with SH3 domain | Other | other | |
| chr1 | *THEMIS2* | 7 | 294 | 15 | THEMIS2 | thymocyte selection associated family member 2 | Other | other | |
| chr17 | *ICAM2* | 6 | 210 | 15 | ICAM2 | intercellular adhesion molecule 2 | Plasma Membrane | other | |
| chr17 | *MED9* | 6 | 251 | 15 | MED9 | mediator complex subunit 9 | Nucleus | other | |
| chr5 | *SPINK5* | 6 | 145 | 15 | SPINK5 | serine peptidase inhibitor, Kazal type 5 | Extracellular Space | other | |
| chr2 | *VIT* | 7 | 151 | 15 | VIT | vitrin | Extracellular Space | other | |
| chr13 | *ZMYM2* | 5 | 140 | 15 | ZMYM2 | zinc finger MYM-type containing | Nucleus | transcription regulator | |
| chr19 | *TIMM50* | 6 | 237 | 15 | TIMM50 | translocase of inner mitochondrial membrane 50 | Cytoplasm | phosphatase | |
| chr11 | *RP11-124G5.3* | 7 | 124 | 15 | | | | | |
| chr16 | *RP11-252E2.2* | 7 | 174 | 15 | | | | | |
| chr15 | *CDAN1* | 6 | 235 | 15 | CDAN1 | codanin 1 | Nucleus | other | |
| chr7 | *PSPH* | 8 | 212 | 15 | PSPH | phosphoserine phosphatase | Cytoplasm | phosphatase | |
| chr10 | *RP11-225H22.5* | 6 | 297 | 15 | | | | | |
| chr10 | *SLC18A2* | 7 | 180 | 15 | SLC18A2 | solute carrier family 18 member A2 | Plasma Membrane | transporter | N-methyl-3,4-methylenedioxyamphetamine, reserpine, hydrochlorothiazide/reserpine, amphetamine, deserpidine/methyclothiazide, polythiazide/reserpine, methamphetamine, deserpidine, tetrabenazine, valbenazine, secologanin tryptamine alkaloid derivative, dihydrotetrabenazine, hydralazine/hydrochlorothiazide/reserpine, reserpine/trichlorome[illegible] |
| chr2 | *AC010987.5* | 6 | 164 | 15 | | | | | |
| chr19 | *MVB12A* | 7 | 270 | 15 | MVB12A | multivesicular body subunit 12A | Cytoplasm | other | |
| chr14 | *ADSSL1* | 7 | 344 | 15 | ADSS1 | adenylosuccinate synthase 1 | Cytoplasm | enzyme | |
| chr4 | *WDFY3* | 5 | 128 | 15 | WDFY3 | WD repeat and FYVE domain containing 3 | Cytoplasm | enzyme | |
| chr9 | *MOB3B* | 6 | 153 | 15 | MOB3B | MOB kinase activator 3B | Other | other | |
| chr1 | *PLA2G2C* | 6 | 231 | 15 | PLA2G2C | phospholipase A2 group IIC | Extracellular Space | enzyme | |

| | | | | | | | | | |
|---|---|---|---|---|---|---|---|---|---|
| chr8 | *FZD3* | 6 | 148 | 15 | FZD3 | frizzled class receptor 3 | Plasma Membrane | i-protein coupled recepto | |
| chr20 | *STK35* | 6 | 186 | 15 | STK35 | serine/threonine kinase 35 | Cytoplasm | kinase | |
| chr5 | *FAM173B* | 7 | 506 | 15 | ATPSCKMT | ATP synthase c subunit lysine N-methyltransferase | Cytoplasm | enzyme | |
| chr22 | *BAIAP2L2* | 7 | 274 | 15 | BAIAP2L2 | BAR/IMD domain containing adaptor protein 2 like 2 | Cytoplasm | other | |
| chr2 | *TRIP12* | 5 | 135 | 15 | TRIP12 | thyroid hormone receptor interactor 12 | Cytoplasm | enzyme | |
| chr5 | *CTD-2152M20.2* | 6 | 426 | 15 | | | | | |
| chr1 | *SYTL1* | 8 | 326 | 15 | SYTL1 | synaptotagmin like 1 | Cytoplasm | enzyme | |
| chr9 | *DCAF12* | 6 | 156 | 15 | DCAF12 | DDB1 and CUL4 associated factor 12 | Cytoplasm | other | |
| chr7 | *STX1A* | 6 | 255 | 15 | STX1A | syntaxin 1A | Cytoplasm | transporter | |
| chrX | *RP11-245M24.1* | 6 | 190 | 15 | | | | | |
| chr17 | *AC144836.1* | 7 | 288 | 15 | | | | | |
| chr5 | *RGS14* | 7 | 329 | 15 | RGS14 | regulator of G protein signaling | Cytoplasm | enzyme | |
| chr13 | *XPO4* | 5 | 143 | 15 | XPO4 | exportin 4 | Nucleus | transporter | |
| chr2 | *ACTG2* | 6 | 168 | 15 | ACTG2 | actin gamma 2, smooth muscle | Cytoplasm | other | |
| chr18 | *RP11-638L3.1* | 5 | 143 | 15 | | | | | |
| chr6 | *IPCEF1* | 6 | 147 | 15 | IPCEF1 | interaction protein for cytohesin exchange factors 1 | Cytoplasm | enzyme | |
| chr9 | *RP11-613M10.9* | 6 | 174 | 15 | | | | | |
| chr2 | *GMCL1* | 7 | 152 | 15 | GMCL1 | germ cell-less 1, spermatogenesis associated | Nucleus | other | |
| chr9 | *ERMP1* | 5 | 181 | 15 | ERMP1 | endoplasmic reticulum metallopeptidase 1 | Cytoplasm | other | |
| chr1 | *C1orf198* | 6 | 388 | 15 | C1orf198 | chromosome 1 open reading frame 198 | Cytoplasm | other | |
| chr19 | *ASF1B* | 7 | 168 | 15 | ASF1B | anti-silencing function 1B histone chaperone | Nucleus | other | |
| chr16 | *TBC1D10B* | 7 | 344 | 15 | TBC1D10B | TBC1 domain family member 10B | Plasma Membrane | enzyme | |
| chr9 | *CD72* | 6 | 174 | 15 | CD72 | CD72 molecule | Plasma Membrane | transmembrane recepto | |
| chr11 | *BDNF-AS* | 8 | 193 | 15 | BDNF-AS | BDNF antisense RNA | Other | other | |
| chr9 | *ARPC5L* | 7 | 320 | 15 | ARPC5L | actin related protein 2/3 complex subunit 5 like | Cytoplasm | other | |
| chr12 | *IQCD* | 6 | 207 | 15 | IQCD | IQ motif containing D | Other | other | |
| chr1 | *VPS72* | 7 | 287 | 15 | VPS72 | vacuolar protein sorting 72 | Nucleus | transcription regulator | |
| chr5 | *FAM169A* | 6 | 121 | 15 | FAM169A | family with sequence similarity 169 member A | Other | other | |
| chr2 | *FANCL* | 5 | 156 | 15 | FANCL | FA complementation group L | Nucleus | enzyme | |
| chr20 | *GNAS-AS1* | 7 | 447 | 15 | GNAS-AS1 | GNAS antisense RNA 1 | Other | other | |
| chr15 | *BBS4* | 6 | 150 | 15 | BBS4 | Bardet-Biedl syndrome 4 | Cytoplasm | other | |
| chr1 | *ITGB3BP* | 6 | 140 | 15 | ITGB3BP | integrin subunit beta 3 binding | Nucleus | other | |
| chr5 | *CHSY3* | 6 | 117 | 15 | CHSY3 | chondroitin sulfate synthase 3 | Cytoplasm | enzyme | |
| chr16 | *USB1* | 6 | 186 | 15 | USB1 | U6 snRNA biogenesis phosphodiesterase 1 | Nucleus | enzyme | |
| chr5 | *MATR3* | 6 | 139 | 15 | MATR3 | matrin 3 | Nucleus | other | |
| chr12 | *CALCOCO1* | 7 | 344 | 15 | CALCOCO1 | calcium binding and coiled-coil domain 1 | Nucleus | transcription regulator | |
| chr3 | *CP* | 6 | 178 | 15 | CP | ceruloplasmin | Extracellular Space | enzyme | |
| chr3 | *C3orf83* | 6 | 186 | 15 | MKRN2OS | MKRN2 opposite strand | Other | other | |
| chr18 | *AFG3L2* | 6 | 151 | 15 | AFG3L2 | AFG3 like matrix AAA peptidase subunit 2 | Cytoplasm | peptidase | |
| chr10 | *LGI1* | 6 | 151 | 15 | LGI1 | leucine rich glioma inactivated 1 | Plasma Membrane | other | |
| chr19 | *NOVA2* | 6 | 210 | 15 | NOVA2 | NOVA alternative splicing | Nucleus | other | |
| chr6 | *PTPRK* | 5 | 136 | 15 | PTPRK | protein tyrosine phosphatase receptor type K | Plasma Membrane | phosphatase | |
| chr5 | *LMBRD2* | 7 | 687 | 15 | LMBRD2 | LMBR1 domain containing 2 | Other | other | |
| chr3 | *RFT1* | 7 | 164 | 15 | RFT1 | RFT1 homolog | Other | other | |
| chr1 | *RP1-91G5.3* | 7 | 201 | 15 | | | | | |
| chr3 | *CIDECP* | 6 | 218 | 15 | CIDECP1 | cell death inducing DFFA like effector c pseudogene 1 | Other | other | |
| chr7 | *RNF32* | 5 | 165 | 15 | RNF32 | ring finger protein 32 | Cytoplasm | other | |
| chr1 | *CD46* | 5 | 197 | 15 | CD46 | CD46 molecule | Plasma Membrane | transmembrane recepto | FOR46 |
| chrX | *SLC38A5* | 7 | 259 | 15 | SLC38A5 | solute carrier family 38 member 5 | Plasma Membrane | transporter | |
| chr8 | *HSF1* | 7 | 277 | 15 | HSF1 | heat shock transcription factor 1 | Nucleus | transcription regulator | |
| chr9 | *AQP7P4* | 6 | 239 | 15 | AQP7P4 | aquaporin 7 pseudogene 4 | Other | other | |
| chr7 | *CPA5* | 6 | 263 | 15 | CPA5 | carboxypeptidase A5 | Extracellular Space | peptidase | |
| chr22 | *PIWIL3* | 6 | 170 | 15 | PIWIL3 | piwi like RNA-mediated gene silencing 3 | Other | other | |
| chr3 | *RP11-24F11.2* | 7 | 137 | 15 | | | | | |
| chr16 | *RP11-461O7.1* | 7 | 161 | 15 | | | | | |
| chr21 | *PWP2* | 7 | 378 | 15 | )C102724159/PW | PWP2 small subunit processome component | Nucleus | other | |
| chr2 | *SLC35F5* | 6 | 192 | 15 | SLC35F5 | solute carrier family 35 member | Other | other | |
| chr13 | *CDC16* | 6 | 195 | 15 | CDC16 | cell division cycle 16 | Nucleus | other | |
| chr15 | *FAM96A* | 6 | 178 | 15 | CIAO2A | cytosolic iron-sulfur assembly component 2A | Extracellular Space | other | |
| chr1 | *RP11-46A10.5* | 6 | 274 | 15 | | | | | |
| chr6 | *RP11-554D15.1* | 6 | 152 | 15 | | | | | |
| chr3 | *CSPG5* | 7 | 229 | 15 | CSPG5 | chondroitin sulfate proteoglycan 5 | Extracellular Space | growth factor | |
| chrX | *FLNA* | 8 | 424 | 15 | FLNA | filamin A | Cytoplasm | other | |
| chr14 | *TMED10* | 5 | 157 | 15 | TMED10 | transmembrane p24 trafficking protein 10 | Cytoplasm | transporter | |
| chr4 | *RP11-17E2.2* | 6 | 139 | 15 | | | | | |
| chr5 | *FAM153C* | 6 | 263 | 15 | FAM153CP | family with sequence similarity 153 member C | Other | other | |
| chr2 | *RHOQ* | 8 | 248 | 15 | RHOQ | ras homolog family member Q | Plasma Membrane | enzyme | |

| | | | | | | | | | |
|---|---|---|---|---|---|---|---|---|---|
| chr16 | *NPIPB6* | 6 | 170 | 15 | PB15 (includes oth | nuclear pore complex interacting protein family member B15 | Nucleus | other | |
| chr4 | *STX18* | 5 | 178 | 15 | STX18 | syntaxin 18 | Cytoplasm | transporter | |
| chr13 | *SMPD4P2* | 5 | 211 | 15 | SMPD4P2 | sphingomyelin phosphodiesterase 4 pseudogene | Other | other | |
| chr3 | *C3orf22* | 7 | 376 | 15 | C3orf22 | chromosome 3 open reading frame 22 | Other | other | |
| chr18 | *RP11-945C19.1* | 8 | 270 | 15 | | | | | |
| chr12 | *SIRT4* | 6 | 190 | 15 | SIRT4 | sirtuin 4 | Cytoplasm | enzyme | |
| chr10 | *TNKS2* | 6 | 150 | 15 | TNKS2 | tankyrase 2 | Nucleus | enzyme | |
| chr5 | *CTC-347C20.1* | 7 | 203 | 15 | | | | | |
| chr7 | *RP11-1220K2.2* | 7 | 130 | 15 | | | | | |
| chr19 | *DDA1* | 7 | 313 | 15 | DDA1 | DET1 and DDB1 associated 1 | Nucleus | other | |
| chr8 | *RP11-120I21.2* | 5 | 152 | 15 | | | | | |
| chr5 | *RP11-1C1.5* | 8 | 565 | 15 | | | | | |
| chr7 | *ACTL6B* | 6 | 199 | 15 | ACTL6B | actin like 6B | Nucleus | other | |
| chr3 | *OGG1* | 7 | 317 | 15 | OGG1 | 8-oxoguanine DNA glycosylase | Nucleus | enzyme | |
| chr2 | *LBH* | 6 | 157 | 15 | LBH | LBH regulator of WNT signaling pathway | Nucleus | transcription regulator | |
| chr4 | *ANK2* | 5 | 137 | 15 | ANK2 | ankyrin 2 | Plasma Membrane | other | |
| chr9 | *PGM5* | 6 | 152 | 15 | PGM5 | phosphoglucomutase 5 | Cytoplasm | enzyme | |
| chr1 | *CCNL2* | 7 | 259 | 15 | CCNL2 | cyclin L2 | Nucleus | other | |
| chr1 | *OPTC* | 6 | 298 | 15 | OPTC | opticin | Extracellular Space | other | |
| chr1 | *C1orf186* | 7 | 210 | 15 | RHEX | regulator of hemoglobinization and erythroid cell expansion | Plasma Membrane | other | |
| chr14 | *RP11-666E17.1* | 6 | 160 | 15 | | | | | |
| chr2 | *PCYOX1* | 7 | 175 | 15 | PCYOX1 | prenylcysteine oxidase 1 | Cytoplasm | enzyme | |
| chr11 | *HEPHL1* | 8 | 388 | 15 | HEPHL1 | hephaestin like 1 | Other | enzyme | |
| chr15 | *ASB7* | 6 | 252 | 15 | ASB7 | ankyrin repeat and SOCS box containing 7 | Cytoplasm | other | |
| chr2 | *VAX2* | 7 | 238 | 15 | VAX2 | ventral anterior homeobox 2 | Nucleus | transcription regulator | |
| chr15 | *GOLGA8EP* | 5 | 385 | 15 | OLGA8EP/GOLGA | golgin A8 family member G | Other | other | |
| chr10 | *GDI2* | 7 | 174 | 15 | GDI2 | GDP dissociation inhibitor 2 | Cytoplasm | other | |
| chr10 | *FFAR4* | 7 | 181 | 15 | FFAR4 | free fatty acid receptor 4 | Plasma Membrane | -protein coupled recepto | |
| chr12 | *WIBG* | 7 | 289 | 15 | PYM1 | PYM homolog 1, exon junction complex associated factor | Nucleus | other | |
| chr1 | *ZNHIT6* | 6 | 139 | 15 | ZNHIT6 | zinc finger HIT-type containing 6 | Cytoplasm | other | |
| chr17 | *PMP22* | 6 | 165 | 15 | PMP22 | peripheral myelin protein 22 | Plasma Membrane | other | |
| chr3 | *TBC1D23* | 6 | 154 | 15 | TBC1D23 | TBC1 domain family member 23 | Cytoplasm | other | |
| chr14 | *DNAL1* | 5 | 147 | 15 | DNAL1 | dynein axonemal light chain 1 | Cytoplasm | other | |
| chr16 | *CLEC18C* | 7 | 343 | 15 | LEC18A/CLEC18 | C-type lectin domain family 18 member A | Other | other | |
| chr17 | *RP11-87G24.3* | 7 | 225 | 15 | | | | | |
| chr16 | *ZNF276* | 7 | 305 | 15 | ZNF276 | zinc finger protein 276 | Nucleus | transcription regulator | |
| chr1 | *TNNT2* | 7 | 374 | 15 | TNNT2 | troponin T2, cardiac type | Cytoplasm | other | |
| chr12 | *YAF2* | 7 | 185 | 15 | YAF2 | YY1 associated factor 2 | Nucleus | transcription regulator | |
| chr2 | *AC092684.1* | 5 | 126 | 15 | | | | | |
| chr4 | *NR3C2* | 6 | 162 | 15 | NR3C2 | nuclear receptor subfamily 3 group C member 2 | Nucleus | d-dependent nuclear rec | fludrocortisone acetate, eplerenone, spironolactone, aldosterone receptor antagonist, fludrocortisone, progesterone, drospirenone, hydrochlorothiazide/spironolactone, AZD9977, beta-estradiol/progesterone, drospirenone/ethinyl |
| chr17 | *NOS2P1* | 6 | 276 | 15 | NOS2P1 | nitric oxide synthase 2 | Other | other | |
| chr10 | *CTGLF10P* | 7 | 358 | 15 | AGAP10P | ArfGAP with GTPase domain, ankyrin repeat and PH domain 10, pseudogene | Other | other | |
| chr17 | *RP11-6N17.10* | 6 | 291 | 15 | | | | | |
| chr12 | *RAB5B* | 6 | 338 | 15 | RAB5B | RAB5B, member RAS oncogene family | Cytoplasm | enzyme | |
| chr10 | *CCDC6* | 7 | 304 | 15 | CCDC6 | coiled-coil domain containing 6 | Cytoplasm | other | |
| chr21 | *ITGB2* | 8 | 426 | 15 | ITGB2 | integrin subunit beta 2 | Plasma Membrane | transmembrane receptor | |
| chr22 | *IL17RA* | 6 | 218 | 15 | IL17RA | interleukin 17 receptor A | Plasma Membrane | transmembrane receptor | brodalumab |
| chrX | *ASB11* | 7 | 135 | 15 | ASB11 | ankyrin repeat and SOCS box containing 11 | Nucleus | transcription regulator | |
| chr6 | *CDC5L* | 7 | 158 | 15 | CDC5L | cell division cycle 5 like | Nucleus | transcription regulator | |
| chr1 | *LRRC40* | 5 | 144 | 15 | LRRC40 | leucine rich repeat containing 40 | Nucleus | other | |
| chr19 | *FSD1* | 6 | 219 | 15 | FSD1 | fibronectin type III and SPRY domain containing 1 | Cytoplasm | other | |
| chrX | *RP4-755D9.1* | 6 | 157 | 15 | | | | | |
| chr1 | *ZNF683* | 7 | 256 | 15 | ZNF683 | zinc finger protein 683 | Other | other | |
| chr11 | *RTN4RL2* | 7 | 281 | 15 | RTN4RL2 | reticulon 4 receptor like 2 | Plasma Membrane | other | |
| chr15 | *RP11-210M15.2* | 7 | 174 | 15 | | | | | |
| chr19 | *IL4I1* | 6 | 273 | 15 | IL4I1 | interleukin 4 induced 1 | Cytoplasm | enzyme | |
| chr1 | *VAV3-AS1* | 7 | 229 | 15 | VAV3-AS1 | VAV3 antisense RNA 1 | Other | other | |
| chr12 | *STAT2* | 6 | 165 | 15 | STAT2 | signal transducer and activator of transcription 2 | Nucleus | transcription regulator | |
| chrX | *WDR45* | 7 | 224 | 15 | WDR45 | WD repeat domain 45 | Cytoplasm | other | |
| chr7 | *GAL3ST4* | 6 | 308 | 15 | GAL3ST4 | galactose-3-O-sulfotransferase 4 | Cytoplasm | enzyme | |
| chr6 | *TBC1D7* | 7 | 188 | 15 | TBC1D7 | TBC1 domain family member 7 | Cytoplasm | other | |
| chr5 | *C7* | 7 | 408 | 15 | C7 | complement C7 | Extracellular Space | other | |

| | | | | | | | | | |
|---|---|---|---|---|---|---|---|---|---|
| chr10 | *GLUD1* | 6 | 134 | 15 | GLUD1 | glutamate dehydrogenase 1 | Cytoplasm | enzyme | |
| chr19 | *ZNF833P* | 6 | 167 | 15 | ZNF833P | zinc finger protein 833, | Other | other | |
| chr6 | *GOPC* | 6 | 167 | 15 | GOPC | golgi associated PDZ and coiled-coil motif containing | Cytoplasm | transporter | |
| chr19 | *CADM4* | 6 | 304 | 15 | CADM4 | cell adhesion molecule 4 | Plasma Membrane | other | |
| chr1 | *PBXIP1* | 7 | 392 | 15 | PBXIP1 | PBX homeobox interacting | Nucleus | transcription regulator | |
| chr4 | *IRF2* | 6 | 159 | 15 | IRF2 | interferon regulatory factor 2 | Nucleus | transcription regulator | |
| chr7 | *RASA4DP* | 5 | 327 | 15 | RASA4DP | RAS p21 protein activator 4CD, pseudogene | Other | other | |
| chr8 | *PPP2R2A* | 6 | 170 | 15 | PPP2R2A | protein phosphatase 2 regulatory subunit Balpha | Cytoplasm | phosphatase | |
| chr5 | *CCDC152* | 5 | 290 | 15 | CCDC152 | coiled-coil domain containing 152 | Other | other | |
| chr1 | *IGSF9* | 7 | 398 | 15 | IGSF9 | immunoglobulin superfamily member 9 | Plasma Membrane | other | |
| chr20 | *C20orf166* | 8 | 499 | 15 | MIR1-1HG | MIR1-1 host gene | Other | other | |
| chr3 | *GPR128* | 6 | 206 | 15 | ADGRG7 | adhesion G protein-coupled receptor G7 | Plasma Membrane | -protein coupled recepto | |
| chr19 | *C19orf10* | 6 | 222 | 15 | MYDGF | myeloid derived growth factor | Extracellular Space | cytokine | |
| chr6 | *CCDC170* | 6 | 149 | 15 | CCDC170 | coiled-coil domain containing 170 | Extracellular Space | other | |
| chr4 | *RP11-386B13.3* | 5 | 153 | 15 | | | | | |
| chr7 | *TSPAN13* | 6 | 200 | 15 | TSPAN13 | tetraspanin 13 | Plasma Membrane | other | |
| chr12 | *KIAA1033* | 5 | 137 | 15 | WASHC4 | WASH complex subunit 4 | Cytoplasm | other | |
| chr1 | *BSND* | 6 | 235 | 15 | BSND | barttin CLCNK type accessory beta subunit | Plasma Membrane | ion channel | |
| chr12 | *BEST3* | 6 | 141 | 15 | BEST3 | bestrophin 3 | Nucleus | ion channel | |
| chr1 | *CD5L* | 7 | 182 | 15 | CD5L | CD5 molecule like | Plasma Membrane | transmembrane recepto | |
| chr5 | *DAB2* | 5 | 409 | 15 | DAB2 | DAB adaptor protein 2 | Plasma Membrane | other | |
| chr4 | *FAM160A1* | 5 | 145 | 15 | FAM160A1 | family with sequence similarity 160 member A1 | Other | other | |
| chr7 | *ZCWPW1* | 7 | 246 | 15 | ZCWPW1 | zinc finger CW-type and PWWP domain containing 1 | Other | other | |
| chr10 | *ENTPD7* | 6 | 118 | 15 | ENTPD7 | ectonucleoside triphosphate diphosphohydrolase 7 | Cytoplasm | enzyme | |
| chr2 | *SLC3A1* | 6 | 204 | 15 | SLC3A1 | solute carrier family 3 member 1 | Plasma Membrane | transporter | |
| chr11 | *RP11-290F24.3* | 7 | 288 | 15 | | | | | |
| chr4 | *SEL1L3* | 6 | 123 | 15 | SEL1L3 | SEL1L family member 3 | Nucleus | other | |
| chr17 | *GSDMB* | 7 | 166 | 15 | GSDMB | gasdermin B | Cytoplasm | other | |
| chr19 | *ZNF28* | 7 | 148 | 15 | ZNF28 | zinc finger protein 28 | Nucleus | transcription regulator | |
| chr1 | *ERO1LB* | 7 | 193 | 15 | ERO1B | endoplasmic reticulum oxidoreductase 1 beta | Cytoplasm | enzyme | |
| chr21 | *POFUT2* | 6 | 318 | 15 | POFUT2 | protein O-fucosyltransferase 2 | Cytoplasm | enzyme | |
| chrX | *KRBOX4* | 6 | 189 | 15 | KRBOX4 | KRAB box domain containing 4 | Other | other | |
| chr17 | *RP1-66C13.3* | 6 | 259 | 15 | | | | | |
| chr15 | *HYKK* | 7 | 209 | 15 | HYKK | hydroxylysine kinase | Cytoplasm | kinase | |
| chr18 | *RTTN* | 5 | 168 | 15 | RTTN | rotatin | Cytoplasm | other | |
| chr11 | *HRASLS5* | 6 | 166 | 15 | PLAAT5 | phospholipase A and acyltransferase 5 | Other | enzyme | |
| chr6 | *RP11-328M4.2* | 7 | 206 | 15 | | | | | |
| chr7 | *SCIN* | 7 | 220 | 15 | SCIN | scinderin | Cytoplasm | other | |
| chr1 | *STXBP3* | 7 | 155 | 15 | STXBP3 | syntaxin binding protein 3 | Plasma Membrane | transporter | |
| chr4 | *ANKRD17* | 5 | 125 | 15 | ANKRD17 | ankyrin repeat domain 17 | Nucleus | other | |
| chr13 | *TSC22D1* | 6 | 156 | 15 | TSC22D1 | TSC22 domain family member 1 | Nucleus | transcription regulator | |
| chr10 | *EMX2OS* | 7 | 157 | 15 | EMX2OS | EMX2 opposite strand/antisense RNA | Other | other | |
| chr2 | *AC083949.1* | 6 | 184 | 15 | | | | | |
| chr6 | *KIF25* | 6 | 219 | 15 | KIF25 | kinesin family member 25 | Cytoplasm | enzyme | |
| chr4 | *LINC00955* | 7 | 288 | 15 | LINC00955 | long intergenic non-protein coding RNA 955 | Other | other | |
| chr14 | *SETD3* | 5 | 146 | 15 | SETD3 | SET domain containing 3, actin histidine methyltransferase | Nucleus | enzyme | |
| chr7 | *KIAA1147* | 6 | 182 | 15 | DENND11 | DENN domain containing 11 | Other | other | |
| chr19 | *CYP2B7P1* | 7 | 157 | 15 | CYP2B7P | cytochrome P450 family 2 subfamily B member 7, | Other | other | |
| chr10 | *LINC00866* | 7 | 132 | 15 | LINC00866 | long intergenic non-protein coding RNA 866 | Other | other | |
| chr17 | *DPH1* | 7 | 401 | 15 | DPH1 | diphthamide biosynthesis 1 | Cytoplasm | other | |
| chr11 | *TSGA10IP* | 7 | 403 | 15 | TSGA10IP | testis specific 10 interacting | Other | other | |
| chr13 | *PCID2* | 6 | 218 | 15 | PCID2 | PCI domain containing 2 | Nucleus | transcription regulator | |
| chr3 | *FAM194A* | 6 | 180 | 15 | ERICH6 | glutamate rich 6 | Other | other | |
| chr3 | *ATG3* | 5 | 184 | 15 | ATG3 | autophagy related 3 | Cytoplasm | enzyme | |
| chr10 | *HECTD2* | 6 | 201 | 15 | HECTD2 | HECT domain E3 ubiquitin protein ligase 2 | Cytoplasm | enzyme | |
| chr16 | *HMOX2* | 6 | 206 | 15 | HMOX2 | heme oxygenase 2 | Cytoplasm | enzyme | tin mesoporphyrin |
| chr3 | *RP11-654C22.2* | 7 | 153 | 15 | | | | | |
| chr9 | *ABCA1* | 7 | 156 | 15 | ABCA1 | ATP binding cassette subfamily A member 1 | Plasma Membrane | transporter | probucol |
| chr17 | *AC005772.2* | 6 | 125 | 15 | | | | | |
| chr1 | *RNF207* | 7 | 361 | 15 | RNF207 | ring finger protein 207 | Other | other | |
| chr11 | *WNT11* | 8 | 467 | 15 | WNT11 | Wnt family member 11 | Extracellular Space | other | |
| chr9 | *KIF24* | 6 | 160 | 15 | KIF24 | kinesin family member 24 | Cytoplasm | other | |
| chr1 | *GPA33* | 7 | 239 | 15 | GPA33 | glycoprotein A33 | Plasma Membrane | other | MGD007 |
| chr2 | *RASGRP3* | 5 | 138 | 15 | RASGRP3 | RAS guanyl releasing protein 3 | Cytoplasm | other | |
| chr22 | *ANKRD54* | 6 | 181 | 15 | ANKRD54 | ankyrin repeat domain 54 | Nucleus | transcription regulator | |
| chr14 | *KCNK13* | 7 | 134 | 15 | KCNK13 | potassium two pore domain channel subfamily K member 13 | Plasma Membrane | ion channel | |
| chr3 | *IL1RAP* | 6 | 148 | 15 | IL1RAP | interleukin 1 receptor accessory protein | Plasma Membrane | transmembrane recepto | CAN04 |
| chr16 | *IST1* | 6 | 143 | 15 | IST1 | IST1 factor associated with ESCRT-III | Cytoplasm | other | |

| | | | | | | | | | |
|---|---|---|---|---|---|---|---|---|---|
| chr15 | RP11-522B15.3 | 8 | 360 | 15 | | | | | |
| chr17 | XAF1 | 5 | 194 | 15 | XAF1 | XIAP associated factor 1 | Nucleus | other | |
| chr1 | SEP15; | 6 | 172 | 15 | | | | | |
| chr16 | CES1 | 7 | 167 | 15 | CES1 | carboxylesterase 1 | Cytoplasm | enzyme | |
| chr13 | SCEL | 7 | 125 | 15 | SCEL | sciellin | Cytoplasm | other | |
| chr8 | PCAT1 | 5 | 122 | 15 | PCAT1 | prostate cancer associated transcript 1 | Other | other | |
| chr11 | BEST1 | 7 | 368 | 15 | BEST1 | bestrophin 1 | Plasma Membrane | ion channel | |
| chr14 | FBXO34 | 5 | 124 | 15 | FBXO34 | F-box protein 34 | Other | other | |
| chrX | ENOX2 | 5 | 131 | 15 | ENOX2 | ecto-NOX disulfide-thiol | Plasma Membrane | enzyme | |
| chr5 | CTB-12O2.1 | 6 | 171 | 15 | | | | | |
| chr10 | RP11-324H6.5 | 7 | 260 | 15 | | | | | |
| chr6 | KCNQ5 | 6 | 151 | 15 | KCNQ5 | potassium voltage-gated channel subfamily Q member 5 | Plasma Membrane | ion channel | ezogabine |
| chr7 | ZSCAN25 | 7 | 333 | 15 | ZSCAN25 | zinc finger and SCAN domain containing 25 | Other | other | |
| chr10 | TACR2 | 6 | 443 | 15 | TACR2 | tachykinin receptor 2 | Plasma Membrane | G-protein coupled recepto | SR 48968 |
| chr6 | MCUR1 | 5 | 131 | 15 | MCUR1 | mitochondrial calcium uniporter regulator 1 | Cytoplasm | other | |
| chr17 | ENGASE | 7 | 375 | 15 | ENGASE | endo-beta-N- | Cytoplasm | enzyme | |
| chr1 | SLC25A24 | 6 | 209 | 15 | SLC25A24 | solute carrier family 25 member | Cytoplasm | transporter | |
| chr1 | DFFA | 6 | 210 | 15 | DFFA | DNA fragmentation factor subunit alpha | Nucleus | enzyme | |
| chr9 | POMT1 | 6 | 331 | 15 | POMT1 | protein O-mannosyltransferase 1 | Cytoplasm | enzyme | |
| chr10 | PDLIM1 | 6 | 152 | 15 | PDLIM1 | PDZ and LIM domain 1 | Cytoplasm | transcription regulator | |
| chr16 | RP11-58C22.1 | 5 | 131 | 15 | | | | | |
| chr5 | TSPAN17 | 7 | 310 | 15 | TSPAN17 | tetraspanin 17 | Other | enzyme | |
| chr16 | KAT8 | 7 | 273 | 15 | KAT8 | lysine acetyltransferase 8 | Nucleus | enzyme | |
| chr4 | RAPGEF2 | 5 | 116 | 15 | RAPGEF2 | Rap guanine nucleotide exchange factor 2 | Cytoplasm | other | |
| chr13 | PROZ | 7 | 404 | 15 | PROZ | protein Z, vitamin K dependent plasma glycoprotein | Extracellular Space | peptidase | |
| chr12 | ATP2B1 | 5 | 122 | 15 | ATP2B1 | ATPase plasma membrane Ca2+ transporting 1 | Plasma Membrane | transporter | |
| chr6 | SCML4 | 5 | 144 | 15 | SCML4 | Scm polycomb group protein like | Nucleus | other | |
| chr8 | CDCA2 | 7 | 125 | 15 | CDCA2 | cell division cycle associated 2 | Nucleus | other | |
| chr1 | PTPN22 | 6 | 191 | 14 | PTPN22 | protein tyrosine phosphatase non-receptor type 22 | Cytoplasm | phosphatase | |
| chr17 | ATP1B2 | 7 | 230 | 14 | ATP1B2 | ATPase Na+/K+ transporting subunit beta 2 | Plasma Membrane | transporter | |
| chr1 | PRAMEF6 | 8 | 170 | 14 | MEF4 (includes ot | PRAME family member 4 | Other | other | |
| chrX | CXorf30 | 6 | 169 | 14 | CFAP47 | cilia and flagella associated | Other | other | |
| chr8 | RP11-89K10.1 | 6 | 119 | 14 | | | | | |
| chr13 | USP12 | 5 | 118 | 14 | USP12 | ubiquitin specific peptidase 12 | Cytoplasm | peptidase | |
| chr4 | C4orf19 | 6 | 159 | 14 | C4orf19 | chromosome 4 open reading frame 19 | Nucleus | other | |
| chrX | MAOB | 6 | 133 | 14 | MAOB | monoamine oxidase B | Cytoplasm | enzyme | fenfluramine/phentermine, rasagiline, methamphetamine, isocarboxazid, MAO-B inhibitor, benzphetamine, phentermine/topiramate, safinamide, phentermine, tranylcypromine, procainamide, pargyline, levodopa/rasagiline, selegiline, ladostigil, phenelzine, dextroamphetamine |
| chr12 | RP11-897M7.1 | 6 | 341 | 14 | | | | | |
| chr12 | MLF2 | 7 | 229 | 14 | MLF2 | myeloid leukemia factor 2 | Nucleus | other | |
| chr12 | LTA4H | 6 | 163 | 14 | LTA4H | leukotriene A4 hydrolase | Cytoplasm | enzyme | |
| chr11 | PYGM | 6 | 296 | 14 | PYGM | glycogen phosphorylase, muscle associated | Cytoplasm | enzyme | |
| chr13 | IRS2 | 7 | 174 | 14 | IRS2 | insulin receptor substrate 2 | Cytoplasm | enzyme | |
| chr17 | PLD2 | 6 | 267 | 14 | PLD2 | phospholipase D2 | Cytoplasm | enzyme | |
| chr12 | ZDHHC17 | 5 | 127 | 14 | ZDHHC17 | zinc finger DHHC-type containing | Cytoplasm | enzyme | |
| chr1 | TTLL7 | 5 | 121 | 14 | TTLL7 | tubulin tyrosine ligase like 7 | Cytoplasm | enzyme | |
| chr20 | ARFGAP1 | 7 | 420 | 14 | ARFGAP1 | ADP ribosylation factor GTPase activating protein 1 | Cytoplasm | transporter | |
| chr22 | SBF1 | 7 | 377 | 14 | SBF1 | SET binding factor 1 | Plasma Membrane | phosphatase | |
| chr22 | RIBC2 | 6 | 238 | 14 | RIBC2 | RIB43A domain with coiled-coils | Extracellular Space | other | |
| chr3 | FSTL1 | 8 | 181 | 14 | FSTL1 | follistatin like 1 | Extracellular Space | other | |
| chr9 | AQP7P3 | 7 | 251 | 14 | AQP7P3 | aquaporin 7 pseudogene 3 | Cytoplasm | other | |
| chr5 | RP11-1252I4.2 | 6 | 265 | 14 | | | | | |
| chr17 | RP11-260A9.6 | 5 | 346 | 14 | | | | | |
| chr1 | VANGL2 | 7 | 343 | 14 | VANGL2 | VANGL planar cell polarity | Plasma Membrane | other | |
| chr1 | C1orf63 | 6 | 160 | 14 | RSRP1 | arginine and serine rich protein 1 | Other | other | |
| chr8 | TDRP | 6 | 179 | 14 | TDRP | testis development related protein | Cytoplasm | other | |
| chr12 | MIP | 6 | 178 | 14 | MIP | major intrinsic protein of lens fiber | Plasma Membrane | transporter | |
| chr16 | KLHL36 | 7 | 261 | 14 | KLHL36 | kelch like family member 36 | Cytoplasm | other | |
| chr1 | ZBTB7B | 7 | 519 | 14 | ZBTB7B | zinc finger and BTB domain containing 7B | Nucleus | transcription regulator | |
| chr17 | DDX52 | 5 | 136 | 14 | DDX52 | DExD-box helicase 52 | Nucleus | enzyme | |
| chr6 | NT5DC1 | 6 | 127 | 14 | NT5DC1 | 5'-nucleotidase domain | Other | other | |

| | | | | | | | | | |
|---|---|---|---|---|---|---|---|---|---|
| chr12 | *GPRC5A* | 7 | 214 | 14 | GPRC5A | G protein-coupled receptor class C group 5 member A | Plasma Membrane | -protein coupled recepto | |
| chr6 | *AK9* | 6 | 167 | 14 | AK9 | adenylate kinase 9 | Nucleus | kinase | |
| chr3 | *ABHD5* | 6 | 128 | 14 | ABHD5 | abhydrolase domain containing 5 | Cytoplasm | enzyme | |
| chr1 | *PFKFB2* | 7 | 268 | 14 | PFKFB2 | 6-phosphofructo-2-kinase/fructose-2,6- | Cytoplasm | kinase | |
| chr4 | *PHF17* | 5 | 151 | 14 | JADE1 | jade family PHD finger 1 | Nucleus | transcription regulator | |
| chr1 | *MYOC* | 7 | 214 | 14 | MYOC | myocilin | Cytoplasm | other | |
| chr5 | *RP11-480D4.1* | 6 | 363 | 14 | | | | | |
| chr5 | *ZFP2* | 6 | 170 | 14 | ZFP2 | ZFP2 zinc finger protein | Nucleus | transcription regulator | |
| chr15 | *DUOX1* | 6 | 228 | 14 | DUOX1 | dual oxidase 1 | Plasma Membrane | enzyme | |
| chr17 | *JMJD6* | 7 | 199 | 14 | JMJD6 | jumonji domain containing 6, arginine demethylase and lysine | Plasma Membrane | transmembrane recepto | |
| chr7 | *SRRT* | 6 | 265 | 14 | SRRT | serrate, RNA effector molecule | Nucleus | other | |
| chr19 | *ZNF71* | 7 | 174 | 14 | ZNF71 | zinc finger protein 71 | Nucleus | transcription regulator | |
| chr19 | *RCN3* | 7 | 227 | 14 | RCN3 | reticulocalbin 3 | Cytoplasm | other | |
| chr5 | *PPIP5K2* | 5 | 155 | 14 | PPIP5K2 | diphosphoinositol pentakisphosphate kinase 2 | Cytoplasm | kinase | |
| chr9 | *HABP4* | 6 | 133 | 14 | HABP4 | hyaluronan binding protein 4 | Cytoplasm | other | |
| chr12 | *ALDH2* | 7 | 177 | 14 | ALDH2 | aldehyde dehydrogenase 2 family member | Cytoplasm | enzyme | disulfiram, chlorpropamide, disulfiram/gemcitabine, ANS-6637 |
| chr3 | *C3orf84* | 6 | 187 | 14 | C3orf84 | chromosome 3 open reading frame 84 | Other | other | |
| chr2 | *AC010148.1* | 6 | 169 | 14 | | | | | |
| chr10 | *CACNB2* | 6 | 116 | 14 | CACNB2 | calcium voltage-gated channel auxiliary subunit beta 2 | Plasma Membrane | ion channel | nifedipine, amlodipine/indapamide/perindopril, amlodipine/perindopril, felodipine, verapamil, amlodipine/hydrochlorothiazide/valsartan, dexamethasone/rituximab/verapamil, mibefradil, nimodipine, nitrendipine, trandolapril/verapamil, amlodipine, nisoldipine, amlodipine/benazepril, nilvadipine, amlodipine/celecoxib, enalapril/felodipine, amlodipine/valsartan, amlodipine/ezetimibe/losartan/rosuvastatin [illegible] |
| chr5 | *CTD-2288O8.1* | 6 | 148 | 14 | | | | | |
| chr2 | *AC016735.2* | 6 | 317 | 14 | | | | | |
| chr4 | *SCLT1* | 5 | 150 | 14 | SCLT1 | sodium channel and clathrin | Plasma Membrane | transporter | |
| chr8 | *CHD7* | 5 | 155 | 14 | CHD7 | chromodomain helicase DNA binding protein 7 | Nucleus | enzyme | |
| chr2 | *KLHL23* | 5 | 128 | 14 | 23/PHOSPHO2-K | kelch like family member 23 | Extracellular Space | other | |
| chr2 | *SFTPB* | 6 | 301 | 14 | SFTPB | surfactant protein B | Extracellular Space | other | |
| chr22 | *RP1-149A16.12* | 6 | 219 | 14 | | | | | |
| chr1 | *SRSF11* | 6 | 152 | 14 | SRSF11 | serine and arginine rich splicing factor 11 | Nucleus | other | |
| chr16 | *PDILT* | 6 | 177 | 14 | PDILT | protein disulfide isomerase like, testis expressed | Extracellular Space | enzyme | |
| chr19 | *HDGFRP2* | 7 | 333 | 14 | | | | | |
| chr1 | *PTP4A2* | 6 | 214 | 14 | PTP4A2 | protein tyrosine phosphatase 4A2 | Cytoplasm | phosphatase | |
| chr1 | *SLC30A7* | 6 | 151 | 14 | SLC30A7 | solute carrier family 30 member 7 | Cytoplasm | transporter | |
| chr2 | *HNRNPLL* | 5 | 203 | 14 | HNRNPLL | heterogeneous nuclear ribonucleoprotein L like | Other | other | |
| chr22 | *CTA-292E10.6* | 6 | 143 | 14 | | | | | |
| chr6 | *EYA4* | 5 | 145 | 14 | EYA4 | EYA transcriptional coactivator and phosphatase 4 | Cytoplasm | phosphatase | |
| chr17 | *CHMP6* | 7 | 348 | 14 | CHMP6 | charged multivesicular body | Cytoplasm | other | |
| chr13 | *DCLK1* | 7 | 169 | 14 | DCLK1 | doublecortin like kinase 1 | Plasma Membrane | kinase | |
| chr12 | *CS* | 6 | 158 | 14 | CS | citrate synthase | Cytoplasm | enzyme | |
| chr12 | *MDM2* | 6 | 149 | 14 | MDM2 | MDM2 proto-oncogene | Nucleus | transcription regulator | ASTX295, MK-8242, RO6839921, APG-115, CGM097, HDM201, idasanutlin, milademetan, BI 907828, AMG-232, ALRN-6924, RO-5045337, KRT-232, MI-773, serdemetan |
| chr20 | *GNAS* | 7 | 325 | 14 | GNAS | GNAS complex locus | Plasma Membrane | enzyme | |
| chr17 | *CCR7* | 6 | 279 | 14 | CCR7 | C-C motif chemokine receptor 7 | Plasma Membrane | -protein coupled recepto | |
| chr1 | *C1orf94* | 7 | 188 | 14 | C1orf94 | chromosome 1 open reading frame 94 | Extracellular Space | other | |
| chr12 | *OASL* | 7 | 164 | 14 | OASL | 2'-5'-oligoadenylate synthetase | Cytoplasm | enzyme | |
| chr11 | *LDHA* | 5 | 169 | 14 | LDHA | lactate dehydrogenase A | Cytoplasm | enzyme | |
| chr3 | *RP11-71H9.1* | 7 | 163 | 14 | | | | | |

| | | | | | | | | | |
|---|---|---|---|---|---|---|---|---|---|
| chr10 | *RP11-119F19.2* | 7 | 215 | 14 | | | | | |
| chr19 | *NUCB1* | 6 | 228 | 14 | NUCB1 | nucleobindin 1 | Cytoplasm | other | |
| chr7 | *RP11-310H4.6* | 5 | 196 | 14 | | | | | |
| chr7 | *GUSBP6* | 7 | 161 | 14 | GUSBP6 | GUSB pseudogene 6 | Other | other | |
| chr4 | *FRYL* | 6 | 142 | 14 | FRYL | FRY like transcription coactivator | Other | other | |
| chr12 | *SLC48A1* | 6 | 551 | 14 | SLC48A1 | solute carrier family 48 member 1 | Cytoplasm | transporter | |
| chr7 | *WBSCR22* | 6 | 229 | 14 | BUD23 | BUD23 rRNA methyltransferase and ribosome maturation factor | Nucleus | enzyme | |
| chr13 | *SUCLA2* | 5 | 114 | 14 | SUCLA2 | succinate-CoA ligase ADP-forming beta subunit | Cytoplasm | enzyme | |
| chr15 | *SLC24A1* | 6 | 192 | 14 | SLC24A1 | solute carrier family 24 member 1 | Plasma Membrane | transporter | |
| chr7 | *TRIM73* | 7 | 332 | 14 | TRIM73/TRIM74 | tripartite motif containing 74 | Cytoplasm | other | |
| chr1 | *TOR1AIP2* | 6 | 181 | 14 | TOR1AIP2 | torsin 1A interacting protein 2 | Cytoplasm | other | |
| chr7 | *PHKG1* | 7 | 332 | 14 | PHKG1 | phosphorylase kinase catalytic subunit gamma 1 | Cytoplasm | kinase | |
| chr9 | *AQP7P2* | 7 | 296 | 14 | AQP7P2 | aquaporin 7 pseudogene 2 | Other | other | |
| chr10 | *IL2RA* | 5 | 164 | 14 | IL2RA | interleukin 2 receptor subunit alpha | Plasma Membrane | transmembrane receptor | denileukin diftitox, LMB-2, basiliximab, camidanlumab tesirine, aldesleukin, yttrium Y 90 basiliximab, |
| chr4 | *SH3RF1* | 5 | 109 | 14 | SH3RF1 | SH3 domain containing ring finger | Cytoplasm | enzyme | |
| chr11 | *MAP3K11* | 7 | 533 | 14 | MAP3K11 | mitogen-activated protein kinase kinase kinase 11 | Cytoplasm | kinase | |
| chr17 | *snoU13* | 6 | 190 | 14 | | | | | |
| chr8 | *KB-1930G5.4* | 6 | 170 | 14 | | | | | |
| chr2 | *ZNF385B* | 5 | 132 | 14 | ZNF385B | zinc finger protein 385B | Nucleus | other | |
| chr15 | *RP11-484P15.1* | 7 | 133 | 14 | | | | | |
| chr1 | *MARC1* | 6 | 248 | 14 | MARC1 | mitochondrial amidoxime reducing component 1 | Cytoplasm | enzyme | |
| chr1 | *GLMN* | 6 | 155 | 14 | GLMN | glomulin, FKBP associated | Cytoplasm | other | |
| chr21 | *TMPRSS3* | 6 | 200 | 14 | TMPRSS3 | transmembrane serine protease 3 | Plasma Membrane | peptidase | |
| chr1 | *NBPF4* | 7 | 235 | 14 | NBPF4/NBPF6 | NBPF member 6 | Other | other | |
| chr12 | *RP11-87C12.5* | 7 | 209 | 14 | | | | | |
| chr14 | *KIAA0247* | 5 | 145 | 14 | SUSD6 | sushi domain containing 6 | Other | other | |
| chr11 | *NEU3* | 7 | 188 | 14 | NEU3 | neuraminidase 3 | Plasma Membrane | enzyme | |
| chr10 | *WDR11-AS1* | 7 | 176 | 14 | WDR11-AS1 | WDR11 antisense RNA 1 | Other | other | |
| chr10 | *PTPN20A* | 7 | 169 | 14 | PTPN20 | protein tyrosine phosphatase non-receptor type 20 | Extracellular Space | phosphatase | |
| chr16 | *TUBB3* | 7 | 311 | 14 | TUBB3 | tubulin beta 3 class III | Cytoplasm | other | docetaxel/gemcitabine/vincristine, cyclophosphamide/epirubicin/5-fluorouracil/paclitaxel, bevacizumab/paclitaxel/topotecan, colchicine/probenecid, bevacizumab/docetaxel/prednisone/thalidomide/zoledronic acid, gemcitabine/paclitaxel, simotaxel, L-asparaginase/daunorubicin/dexamethasone/vincristine, prednisone/vincristine, docetaxel/5-fluorouracil, vinflunine, L-[illegible] |
| chr3 | *NPHP3-AS1* | 6 | 180 | 14 | NPHP3-AS1 | NPHP3 antisense RNA 1 | Other | other | |
| chr1 | *THEM4* | 7 | 187 | 14 | THEM4 | thioesterase superfamily member | Plasma Membrane | enzyme | |
| chr17 | *KIAA0100* | 6 | 155 | 14 | KIAA0100 | KIAA0100 | Extracellular Space | other | |
| chr2 | *CAD* | 6 | 245 | 14 | CAD | carbamoyl-phosphate synthetase 2, aspartate transcarbamylase, and dihydroorotase | Cytoplasm | enzyme | |
| chr19 | *LGALS4* | 6 | 229 | 14 | LGALS4 | galectin 4 | Extracellular Space | other | |
| chr1 | *CNKSR1* | 6 | 304 | 14 | CNKSR1 | connector enhancer of kinase suppressor of Ras 1 | Cytoplasm | other | |
| chr15 | *TSPAN3* | 6 | 175 | 14 | TSPAN3 | tetraspanin 3 | Plasma Membrane | other | |
| chr2 | *AGPS* | 5 | 148 | 14 | AGPS | alkylglycerone phosphate | Cytoplasm | enzyme | |
| chr1 | *SLC5A9* | 6 | 228 | 14 | SLC5A9 | solute carrier family 5 member 9 | Plasma Membrane | transporter | |
| chr19 | *CYP2F1* | 6 | 246 | 14 | CYP2F1 | cytochrome P450 family 2 subfamily F member 1 | Cytoplasm | enzyme | |
| chr7 | *MEOX2* | 7 | 196 | 14 | MEOX2 | mesenchyme homeobox 2 | Nucleus | transcription regulator | |
| chr2 | *ABCG5* | 6 | 189 | 14 | ABCG5 | ATP binding cassette subfamily G member 5 | Plasma Membrane | transporter | |
| chr17 | *LIG3* | 6 | 234 | 14 | LIG3 | DNA ligase 3 | Nucleus | enzyme | |
| chr16 | *UMOD* | 6 | 148 | 14 | UMOD | uromodulin | Extracellular Space | other | |
| chr19 | *LGI4* | 7 | 282 | 14 | LGI4 | leucine rich repeat LGI family member 4 | Extracellular Space | other | |
| chr1 | *ARF1* | 6 | 320 | 14 | ARF1 | ADP ribosylation factor 1 | Cytoplasm | enzyme | |
| chr12 | *RP11-536G4.2* | 6 | 131 | 14 | | | | | |
| chr8 | *GSDMC* | 7 | 135 | 14 | GSDMC | gasdermin C | Cytoplasm | other | |
| chr2 | *USP40* | 6 | 212 | 14 | USP40 | ubiquitin specific peptidase 40 | Other | peptidase | |

| | | | | | | | | | |
|---|---|---|---|---|---|---|---|---|---|
| chr19 | *MISP* | 7 | 248 | 14 | MISP | mitotic spindle positioning | Plasma Membrane | other | |
| chr11 | *FOLR3* | 6 | 213 | 14 | FOLR3 | folate receptor gamma | Extracellular Space | other | |
| chr4 | *N4BP2* | 5 | 116 | 14 | N4BP2 | NEDD4 binding protein 2 | Cytoplasm | enzyme | |
| chr8 | *RP11-586K2.1* | 6 | 152 | 14 | | | | | |
| chr6 | *C6orf201* | 6 | 134 | 14 | C6orf201 | chromosome 6 open reading frame 201 | Other | other | |
| chr17 | *CTB-91J4.1* | 6 | 254 | 14 | | | | | |
| chr3 | *MCM2* | 7 | 388 | 14 | MCM2 | minichromosome maintenance complex component 2 | Nucleus | enzyme | |
| chr1 | *ADAM15* | 7 | 401 | 14 | ADAM15 | ADAM metallopeptidase domain | Plasma Membrane | peptidase | |
| chr7 | *ZNF789* | 7 | 220 | 14 | ZNF789 | zinc finger protein 789 | Nucleus | transcription regulator | |
| chr11 | *VPS11* | 7 | 186 | 14 | VPS11 | VPS11 core subunit of CORVET and HOPS complexes | Cytoplasm | transporter | |
| chr16 | *PLEKHG4* | 7 | 342 | 14 | PLEKHG4 | pleckstrin homology and RhoGEF domain containing G4 | Other | other | |
| chr1 | *RGS5* | 6 | 159 | 14 | RGS5 | regulator of G protein signaling 5 | Plasma Membrane | enzyme | |
| chr15 | *VPS39* | 6 | 171 | 14 | VPS39 | VPS39 subunit of HOPS complex | Cytoplasm | transporter | |
| chr8 | *UBR5* | 5 | 122 | 14 | UBR5 | ubiquitin protein ligase E3 component n-recognin 5 | Nucleus | enzyme | |
| chr12 | *PMEL* | 6 | 363 | 14 | PMEL | premelanosome protein | Plasma Membrane | enzyme | IMCgp100 |
| chr17 | *TBX21* | 6 | 256 | 14 | TBX21 | T-box transcription factor 21 | Nucleus | transcription regulator | |
| chr10 | *KIAA1279* | 7 | 257 | 14 | KIFBP | kinesin family binding protein | Cytoplasm | enzyme | |
| chr1 | *CEPT1* | 5 | 142 | 14 | CEPT1 | choline/ethanolamine phosphotransferase 1 | Cytoplasm | enzyme | |
| chr5 | *AC011343.1* | 7 | 208 | 14 | | | | | |
| chr15 | *RP11-578F21.12* | 5 | 146 | 14 | | | | | |
| chr2 | *FAM117B* | 5 | 120 | 14 | FAM117B | family with sequence similarity 117 member B | Other | other | |
| chr22 | *USP18* | 7 | 160 | 14 | USP18 | ubiquitin specific peptidase 18 | Cytoplasm | peptidase | |
| chr22 | *UFD1L* | 5 | 209 | 14 | UFD1 | ubiquitin recognition factor in ER associated degradation 1 | Cytoplasm | peptidase | |
| chr5 | *RP11-321E2.4* | 6 | 484 | 14 | | | | | |
| chr1 | *TCEB3* | 6 | 145 | 14 | ELOA | elongin A | Nucleus | transcription regulator | |
| chr19 | *CYP2S1* | 6 | 276 | 14 | CYP2S1 | cytochrome P450 family 2 subfamily S member 1 | Cytoplasm | enzyme | |
| chr12 | *GNS* | 6 | 146 | 14 | GNS | glucosamine (N-acetyl)-6- | Cytoplasm | enzyme | |
| chr20 | *TMC2* | 5 | 163 | 14 | TMC2 | transmembrane channel like 2 | Plasma Membrane | ion channel | |
| chr12 | *RP11-620J15.2* | 6 | 195 | 14 | | | | | |
| chr8 | *ZNF517* | 6 | 293 | 14 | ZNF517 | zinc finger protein 517 | Nucleus | transcription regulator | |
| chr15 | *CSPG4P12* | 6 | 592 | 14 | CSPG4P12 | chondroitin sulfate proteoglycan 4 pseudogene 12 | Other | other | |
| chr20 | *GDF5* | 7 | 249 | 14 | GDF5 | growth differentiation factor 5 | Extracellular Space | growth factor | |
| chr17 | *KRT15* | 6 | 286 | 14 | KRT15 | keratin 15 | Cytoplasm | other | |
| chr10 | *TMEM254-AS1* | 5 | 128 | 14 | TMEM254-AS1 | TMEM254 antisense RNA 1 | Other | other | |
| chr3 | *USP13* | 6 | 136 | 14 | USP13 | ubiquitin specific peptidase 13 | Cytoplasm | peptidase | |
| chr8 | *XKR5* | 7 | 223 | 14 | XKR5 | XK related 5 | Other | other | |
| chr17 | *AC003043.1* | 6 | 208 | 14 | | | | | |
| chr16 | *RP11-645C24.2* | 5 | 190 | 14 | | | | | |
| chr8 | *ENTPD4* | 6 | 170 | 14 | ENTPD4 | ectonucleoside triphosphate diphosphohydrolase 4 | Cytoplasm | enzyme | |
| chr17 | *PSMD3* | 6 | 214 | 14 | PSMD3 | proteasome 26S subunit, non-ATPase 3 | Cytoplasm | other | |
| chr5 | *SOX30* | 7 | 196 | 14 | SOX30 | SRY-box transcription factor 30 | Nucleus | transcription regulator | |
| chr16 | *RP11-510J16.5* | 6 | 188 | 14 | | | | | |
| chr19 | *ZNF606* | 6 | 166 | 14 | ZNF606 | zinc finger protein 606 | Nucleus | transcription regulator | |
| chr2 | *AC079305.11* | 6 | 125 | 14 | | | | | |
| chr17 | *GIP* | 6 | 200 | 14 | GIP | gastric inhibitory polypeptide | Extracellular Space | other | |
| chr17 | *FAM211A* | 6 | 270 | 14 | LRRC75A | leucine rich repeat containing | Other | other | |
| chr10 | *ITPRIP* | 7 | 253 | 14 | ITPRIP | inositol 1,4,5-trisphosphate receptor interacting protein | Extracellular Space | other | |
| chr10 | *RPS24* | 7 | 186 | 14 | RPS24 | ribosomal protein S24 | Cytoplasm | other | |
| chr7 | *UPP1* | 7 | 334 | 14 | UPP1 | uridine phosphorylase 1 | Cytoplasm | enzyme | |
| chr16 | *CDH16* | 6 | 291 | 14 | CDH16 | cadherin 16 | Plasma Membrane | enzyme | |
| chr10 | *KCNIP2* | 7 | 215 | 14 | KCNIP2 | potassium voltage-gated channel interacting protein 2 | Plasma Membrane | other | |
| chr14 | *C14orf164* | 5 | 125 | 14 | RNF212B | ring finger protein 212B | Other | other | |
| chr17 | *G6PC* | 6 | 152 | 14 | G6PC | glucose-6-phosphatase catalytic subunit | Cytoplasm | phosphatase | |
| chr7 | *ST7* | 6 | 176 | 14 | ST7 | suppression of tumorigenicity 7 | Other | other | |
| chr11 | *CTD-2562J17.4* | 7 | 226 | 14 | | | | | |

| | | | | | | | | | |
|---|---|---|---|---|---|---|---|---|---|
| chr11 | *HTR3B* | 6 | 137 | 14 | HTR3B | 5-hydroxytryptamine receptor 3B | Plasma Membrane | ion channel | dexamethasone/granisetron, dolasetron, ergotamine, alosetron, cisapride, granisetron, D-tubocurarine, aprepitant/granisetron, dexamethasone/ondansetron, dexamethasone/palonosetron, ondansetron, aprepitant/dexamethasone/granisetron, mirtazapine, aprepitant/dexamethasone/palonosetron, fenfluramine, aprepitant/palonosetron, palonosetron |
| chr10 | *ZNF32-AS3* | 7 | 231 | 14 | ZNF32-AS3 | ZNF32 antisense RNA 3 | Other | other | |
| chr7 | *ANKMY2* | 5 | 211 | 14 | ANKMY2 | ankyrin repeat and MYND domain containing 2 | Plasma Membrane | other | |
| chr3 | *EDEM1* | 7 | 147 | 14 | EDEM1 | ER degradation enhancing alpha-mannosidase like protein 1 | Cytoplasm | enzyme | |
| chr2 | *AFTPH* | 6 | 126 | 14 | AFTPH | aftiphilin | Cytoplasm | other | |
| chr7 | *STRA8* | 7 | 246 | 14 | STRA8 | stimulated by retinoic acid 8 | Nucleus | other | |
| chr2 | *LINC00276* | 5 | 162 | 14 | LINC00276 | long intergenic non-protein coding RNA 276 | Other | other | |
| chr7 | *FKBP9L* | 7 | 244 | 14 | FKBP9P1 | FKBP prolyl isomerase 9 pseudogene 1 | Other | other | |
| chr17 | *TVP23B* | 6 | 137 | 14 | TVP23B | trans-golgi network vesicle protein 23 homolog B | Other | other | |
| chr7 | *MKRN1* | 6 | 207 | 14 | MKRN1 | makorin ring finger protein 1 | Cytoplasm | enzyme | |
| chr7 | *ZNF862* | 7 | 231 | 14 | ZNF862 | zinc finger protein 862 | Other | other | |
| chr2 | *DAPL1* | 5 | 119 | 14 | DAPL1 | death associated protein like 1 | Other | other | |
| chr6 | *ATG5* | 6 | 141 | 14 | ATG5 | autophagy related 5 | Cytoplasm | other | |
| chr15 | *C15orf39* | 7 | 385 | 14 | C15orf39 | chromosome 15 open reading frame 39 | Cytoplasm | other | |
| chr14 | *C14orf37* | 5 | 146 | 14 | ARMH4 | armadillo like helical domain containing 4 | Extracellular Space | other | |
| chr3 | *LAMB2* | 6 | 236 | 14 | LAMB2 | laminin subunit beta 2 | Extracellular Space | enzyme | |
| chr11 | *MCAM* | 7 | 364 | 14 | MCAM | melanoma cell adhesion molecule | Plasma Membrane | other | |
| chr11 | *BIRC2* | 6 | 186 | 14 | BIRC2 | baculoviral IAP repeat containing 2 | Cytoplasm | enzyme | GDC-0152, BI-891065, birinapant, ASTX660, GDC- |
| chr1 | *NPR1* | 6 | 361 | 14 | NPR1 | natriuretic peptide receptor 1 | Plasma Membrane | enzyme | nesiritide, nitroprusside, nitroglycerin, hydralazine/isosorbide dinitrate, isosorbide |
| chr14 | *NRDE2* | 5 | 172 | 14 | NRDE2 | NRDE-2, necessary for RNA interference, domain containing | Nucleus | other | |
| chr2 | *ARID5A* | 7 | 437 | 14 | ARID5A | AT-rich interaction domain 5A | Nucleus | transcription regulator | |
| chr3 | *TOMM70A* | 5 | 175 | 14 | TOMM70 | translocase of outer mitochondrial membrane 70 | Cytoplasm | transporter | |
| chr7 | *DBNL* | 6 | 315 | 14 | DBNL | drebrin like | Cytoplasm | other | |
| chr1 | *LRRC71* | 6 | 351 | 14 | LRRC71 | leucine rich repeat containing 71 | Other | other | |
| chr12 | *KCNC2* | 6 | 180 | 14 | KCNC2 | potassium voltage-gated channel subfamily C member 2 | Plasma Membrane | ion channel | dalfampridine |
| chr10 | *CEP55* | 6 | 158 | 14 | CEP55 | centrosomal protein 55 | Cytoplasm | other | |
| chr1 | *AIM2* | 7 | 233 | 14 | AIM2 | absent in melanoma 2 | Cytoplasm | other | |
| chr17 | *GAA* | 7 | 348 | 14 | GAA | glucosidase alpha, acid | Cytoplasm | enzyme | acarbose, miglitol |
| chr18 | *RP11-138E9.2* | 6 | 167 | 14 | | | | | |
| chr12 | *SLC2A14* | 5 | 135 | 14 | SLC2A14 | solute carrier family 2 member 14 | Nucleus | transporter | |
| chr11 | *FOLH1* | 6 | 246 | 14 | FOLH1 | folate hydrolase 1 | Plasma Membrane | peptidase | HPN424, 131I-MIP-1095, AMG-160, 177Lu-PSMA617, monoclonal antibody J591, 123I-MIP-1095, capromab pendetide, REGN5678, fluorine |
| chr6 | *SERAC1* | 5 | 121 | 14 | SERAC1 | serine active site containing 1 | Extracellular Space | other | |
| chr17 | *CAMTA2* | 6 | 185 | 14 | CAMTA2 | calmodulin binding transcription activator 2 | Nucleus | transcription regulator | |
| chr6 | *CD109* | 5 | 118 | 14 | CD109 | CD109 molecule | Plasma Membrane | other | |
| chr15 | *CILP* | 6 | 212 | 14 | CILP | cartilage intermediate layer | Extracellular Space | phosphatase | |
| chr2 | *CPSF3* | 6 | 134 | 14 | CPSF3 | cleavage and polyadenylation specific factor 3 | Nucleus | enzyme | |
| chr1 | *ZBTB37* | 5 | 158 | 14 | ZBTB37 | zinc finger and BTB domain containing 37 | Nucleus | transcription regulator | |
| chr7 | *FAM180A* | 6 | 250 | 14 | FAM180A | family with sequence similarity 180 member A | Other | other | |
| chr1 | *SH2D5* | 7 | 356 | 14 | SH2D5 | SH2 domain containing 5 | Plasma Membrane | other | |
| chr17 | *CDC6* | 6 | 191 | 14 | CDC6 | cell division cycle 6 | Nucleus | other | |
| chr1 | *NOC2L* | 7 | 351 | 14 | NOC2L | NOC2 like nucleolar associated transcriptional repressor | Nucleus | transcription regulator | |

| | | | | | | | | | |
|---|---|---|---|---|---|---|---|---|---|
| chr3 | *TSEN2* | 6 | 135 | 14 | TSEN2 | tRNA splicing endonuclease subunit 2 | Cytoplasm | enzyme | |
| chr22 | *NCAPH2* | 6 | 258 | 14 | NCAPH2 | non-SMC condensin II complex subunit H2 | Nucleus | other | |
| chr13 | *KATNAL1* | 5 | 120 | 14 | KATNAL1 | katanin catalytic subunit A1 like 1 | Cytoplasm | enzyme | |
| chr16 | *RP11-169E6.4* | 7 | 126 | 14 | | | | | |
| chr1 | *ARHGEF19* | 6 | 350 | 14 | ARHGEF19 | Rho guanine nucleotide exchange factor 19 | Cytoplasm | other | |
| chr5 | *FGF1* | 6 | 170 | 14 | FGF1 | fibroblast growth factor 1 | Extracellular Space | growth factor | pentosan polysulfate |
| chrX | *TMEM164* | 6 | 147 | 14 | TMEM164 | transmembrane protein 164 | Other | other | |
| chr10 | *PSD* | 7 | 293 | 14 | PSD | pleckstrin and Sec7 domain containing | Cytoplasm | transporter | |
| chr10 | *PARGP1* | 6 | 167 | 14 | PARGP1 | poly(ADP-ribose) glycohydrolase pseudogene 1 | Other | other | |
| chr6 | *GUCA1B* | 6 | 189 | 14 | GUCA1B | guanylate cyclase activator 1B | Cytoplasm | other | |
| chr3 | *TRPC1* | 6 | 136 | 14 | TRPC1 | transient receptor potential cation channel subfamily C member 1 | Plasma Membrane | ion channel | |
| chr6 | *TCP11* | 7 | 163 | 14 | TCP11 | t-complex 11 | Cytoplasm | other | |
| chr19 | *ZNF557* | 7 | 175 | 14 | ZNF557 | zinc finger protein 557 | Nucleus | transcription regulator | |
| chr9 | *RP11-318K12.2* | 5 | 125 | 14 | | | | | |
| chr4 | *CPLX1* | 7 | 286 | 14 | CPLX1 | complexin 1 | Plasma Membrane | transporter | |
| chr11 | *GLYAT* | 5 | 113 | 14 | GLYAT | glycine-N-acyltransferase | Cytoplasm | enzyme | |
| chr8 | *LEPROTL1* | 6 | 142 | 14 | LEPROTL1 | leptin receptor overlapping transcript like 1 | Plasma Membrane | other | |
| chr3 | *LRRIQ4* | 7 | 279 | 14 | LRRIQ4 | leucine rich repeats and IQ motif containing 4 | Other | other | |
| chr16 | *FAM157C* | 6 | 319 | 14 | FAM157C | family with sequence similarity 157 member C | Other | other | |
| chr1 | *FAM63A* | 7 | 285 | 14 | MINDY1 | MINDY lysine 48 deubiquitinase 1 | Cytoplasm | peptidase | |
| chr3 | *RP11-231I13.2* | 7 | 158 | 14 | | | | | |
| chr12 | *PDE1B* | 7 | 298 | 14 | PDE1B | phosphodiesterase 1B | Cytoplasm | enzyme | |
| chr1 | *MFSD2A* | 6 | 300 | 14 | MFSD2A | major facilitator superfamily domain containing 2A | Plasma Membrane | transporter | |
| chr6 | *BEND3* | 6 | 180 | 14 | BEND3 | BEN domain containing 3 | Nucleus | other | |
| chr15 | *VPS33B* | 7 | 211 | 14 | VPS33B | VPS33B late endosome and lysosome associated | Cytoplasm | transporter | |
| chr8 | *RP11-513H8.1* | 7 | 175 | 14 | | | | | |
| chr18 | *TNFRSF11A* | 5 | 130 | 14 | TNFRSF11A | TNF receptor superfamily member 11a | Plasma Membrane | transmembrane receptor | |
| chr11 | *ZBED5-AS1* | 6 | 183 | 14 | ZBED5-AS1 | ZBED5 antisense RNA 1 | Other | other | |
| chr18 | *RP11-567M16.3* | 5 | 168 | 14 | | | | | |
| chr8 | *SFRP1* | 5 | 152 | 14 | SFRP1 | secreted frizzled related protein 1 | Plasma Membrane | transmembrane receptor | |
| chr14 | *SEC23A* | 4 | 114 | 14 | SEC23A | Sec23 homolog A, coat complex II component | Cytoplasm | transporter | |
| chr7 | *NFE2L3* | 7 | 172 | 14 | NFE2L3 | nuclear factor, erythroid 2 like 3 | Nucleus | transcription regulator | |
| chr20 | *ENTPD6* | 6 | 245 | 14 | ENTPD6 | ectonucleoside triphosphate diphosphohydrolase 6 | Cytoplasm | enzyme | |
| chr19 | *REXO1* | 7 | 325 | 14 | REXO1 | RNA exonuclease 1 homolog | Nucleus | transcription regulator | |
| chr3 | *RP11-6B4.1* | 6 | 166 | 14 | | | | | |
| chr1 | *GCSAML* | 7 | 221 | 14 | GCSAML | germinal center associated signaling and motility like | Other | other | |
| chr7 | *PAXIP1* | 6 | 196 | 14 | PAXIP1 | PAX interacting protein 1 | Nucleus | other | |
| chr19 | *FCER2* | 6 | 220 | 14 | FCER2 | Fc fragment of IgE receptor II | Plasma Membrane | transmembrane receptor | lumiliximab |
| chr19 | *ZNF100* | 6 | 143 | 14 | ZNF100 | zinc finger protein 100 | Nucleus | transcription regulator | |
| chr1 | *RP4-680D5.2* | 6 | 193 | 14 | | | | | |
| chr15 | *RP11-173D3.1* | 6 | 358 | 14 | | | | | |
| chr3 | *C3orf35* | 7 | 150 | 14 | C3orf35 | chromosome 3 open reading frame 35 | Other | other | |
| chr14 | *SLC38A6* | 5 | 151 | 14 | SLC38A6 | solute carrier family 38 member 6 | Plasma Membrane | transporter | |
| chr2 | *GRHL1* | 5 | 141 | 14 | GRHL1 | grainyhead like transcription | Nucleus | transcription regulator | |
| chr7 | *MYO1G* | 7 | 437 | 14 | MYO1G | myosin IG | Cytoplasm | other | |
| chr11 | *RP11-179A10.1* | 7 | 331 | 14 | | | | | |
| chr20 | *DPM1* | 6 | 223 | 14 | DPM1 | dolichyl-phosphate mannosyltransferase subunit 1, catalytic | Cytoplasm | enzyme | |
| chr5 | *HAVCR1* | 6 | 187 | 14 | HAVCR1 | hepatitis A virus cellular receptor | Plasma Membrane | other | |
| chr10 | *CDC123* | 5 | 129 | 14 | CDC123 | cell division cycle 123 | Cytoplasm | other | |
| chr5 | *SFXN1* | 6 | 150 | 14 | SFXN1 | sideroflexin 1 | Cytoplasm | transporter | |
| chr15 | *WHAMMP3* | 5 | 147 | 14 | WHAMMP3 | WAS protein homolog associated with actin, golgi membranes and microtubules pseudogene 3 | Other | other | |
| chr1 | *RP4-601K24.1* | 7 | 120 | 14 | | | | | |
| chr14 | *PRKD1* | 5 | 138 | 14 | PRKD1 | protein kinase D1 | Cytoplasm | kinase | |
| chr7 | *C7orf60* | 5 | 174 | 14 | BMT2 | base methyltransferase of 25S rRNA 2 homolog | Other | other | |
| chr3 | *ALAS1* | 6 | 206 | 14 | ALAS1 | 5'-aminolevulinate synthase 1 | Cytoplasm | enzyme | |
| chr16 | *RP11-418I22.2* | 6 | 122 | 14 | | | | | |
| chr8 | *ZDHHC2* | 6 | 144 | 14 | ZDHHC2 | zinc finger DHHC-type containing | Nucleus | enzyme | |
| chr2 | *PTCD3* | 5 | 132 | 14 | PTCD3 | pentatricopeptide repeat domain | Cytoplasm | other | |
| chr16 | *ANKRD26P1* | 6 | 213 | 14 | ANKRD26P1 | ankyrin repeat domain 26 pseudogene 1 | Other | other | |
| chr9 | *CRAT* | 6 | 420 | 14 | CRAT | carnitine O-acetyltransferase | Cytoplasm | enzyme | |
| chr11 | *C11orf24* | 6 | 295 | 14 | C11orf24 | chromosome 11 open reading frame 24 | Extracellular Space | other | |
| chr14 | *TEP1* | 6 | 179 | 14 | TEP1 | telomerase associated protein 1 | Nucleus | enzyme | |
| chr15 | *VWA9* | 5 | 131 | 14 | INTS14 | integrator complex subunit 14 | Cytoplasm | other | |
| chr17 | *CD300E* | 6 | 215 | 14 | CD300E | CD300e molecule | Plasma Membrane | other | |
| chr17 | *SHBG* | 6 | 200 | 14 | SHBG | sex hormone binding globulin | Extracellular Space | other | |
| chr16 | *CCL17* | 6 | 249 | 14 | CCL17 | C-C motif chemokine ligand 17 | Extracellular Space | cytokine | |

| | | | | | | | | | |
|---|---|---|---|---|---|---|---|---|---|
| chrX | *CXorf36* | 6 | 185 | 14 | DIPK2B | divergent protein kinase domain | Other | other | |
| chr12 | *DUSP16* | 5 | 139 | 14 | DUSP16 | dual specificity phosphatase 16 | Nucleus | phosphatase | |
| chr17 | *TMC6* | 7 | 309 | 14 | TMC6 | transmembrane channel like 6 | Cytoplasm | transporter | |
| chr5 | *LINC01033* | 6 | 159 | 14 | LINC01033 | | Other | other | |
| chr2 | *TTLL4* | 6 | 128 | 14 | TTLL4 | tubulin tyrosine ligase like 4 | Cytoplasm | enzyme | |
| chr8 | *RGS22* | 5 | 117 | 14 | RGS22 | regulator of G protein signaling | Cytoplasm | other | |
| chr5 | *SLC12A2* | 6 | 132 | 14 | SLC12A2 | solute carrier family 12 member 2 | Plasma Membrane | transporter | bumetanide, quinethazone |
| chr11 | *VPS26B* | 6 | 184 | 14 | VPS26B | VPS26, retromer complex component B | Cytoplasm | transporter | |
| chr19 | *ZNF155* | 5 | 139 | 14 | ZNF155 | zinc finger protein 155 | Nucleus | transcription regulator | |
| chr12 | *SLC6A13* | 6 | 214 | 14 | SLC6A13 | solute carrier family 6 member 13 | Plasma Membrane | transporter | tiagabine |
| chr19 | *ZSCAN1* | 7 | 220 | 14 | ZSCAN1 | zinc finger and SCAN domain containing 1 | Nucleus | transcription regulator | |
| chr16 | *CLEC18B* | 7 | 461 | 14 | CLEC18B | C-type lectin domain family 18 member B | Cytoplasm | other | |
| chr13 | *SACS* | 5 | 122 | 14 | SACS | sacsin molecular chaperone | Plasma Membrane | other | |
| chr1 | *PROX1* | 5 | 239 | 14 | PROX1 | prospero homeobox 1 | Nucleus | transcription regulator | |
| chr2 | *GACAT1* | 7 | 167 | 14 | GACAT1 | gastric cancer associated | Other | other | |
| chr1 | *ILDR2* | 6 | 159 | 14 | ILDR2 | immunoglobulin like domain containing receptor 2 | Other | other | BAY1905254 |
| chr9 | *RP11-162D16.2* | 6 | 181 | 14 | | | | | |
| chr3 | *FAIM* | 6 | 160 | 14 | FAIM | Fas apoptotic inhibitory molecule | Plasma Membrane | other | |
| chr18 | *MOCOS* | 5 | 161 | 14 | MOCOS | molybdenum cofactor sulfurase | Cytoplasm | enzyme | |
| chr14 | *VASH1* | 6 | 264 | 14 | VASH1 | vasohibin 1 | Extracellular Space | peptidase | |
| chr15 | *LINC00924* | 6 | 194 | 14 | LINC00924 | long intergenic non-protein coding RNA 924 | Other | other | |
| chr2 | *MTIF2* | 5 | 179 | 14 | MTIF2 | mitochondrial translational initiation factor 2 | Cytoplasm | translation regulator | |
| chr5 | *SLC6A19* | 8 | 676 | 14 | SLC6A19 | solute carrier family 6 member 19 | Plasma Membrane | transporter | |
| chr17 | *C17orf104* | 5 | 137 | 14 | MEIOC | meiosis specific with coiled-coil domain | Cytoplasm | other | |
| chr12 | *DENND5B-AS1* | 7 | 176 | 14 | DENND5B-AS1 | DENND5B antisense RNA 1 | Other | other | |
| chr19 | *TSKS* | 5 | 226 | 14 | TSKS | testis specific serine kinase | Cytoplasm | kinase | |
| chr16 | *CASKIN1* | 7 | 193 | 14 | CASKIN1 | CASK interacting protein 1 | Nucleus | transcription regulator | |
| chr1 | *RD3* | 6 | 305 | 14 | RD3 | retinal degeneration 3, GUCY2D regulator | Cytoplasm | other | |
| chr1 | *ACP6* | 6 | 223 | 14 | ACP6 | acid phosphatase 6, | Cytoplasm | phosphatase | |
| chr1 | *MIR3916* | 6 | 138 | 14 | MIR3916 | microRNA 3916 | Cytoplasm | microRNA | |
| chr2 | *GTDC1* | 5 | 124 | 14 | GTDC1 | glycosyltransferase like domain containing 1 | Other | other | |
| chr15 | *HMG20A* | 4 | 167 | 14 | HMG20A | high mobility group 20A | Nucleus | transcription regulator | |
| chr6 | *C6orf203* | 5 | 129 | 14 | MTRES1 | mitochondrial transcription rescue factor 1 | Cytoplasm | other | |
| chr13 | *FRY* | 5 | 111 | 14 | FRY | FRY microtubule binding protein | Extracellular Space | other | |
| chr16 | *DHODH* | 7 | 169 | 14 | DHODH | dihydroorotate dehydrogenase (quinone) | Cytoplasm | enzyme | ASLAN003, teriflunomide, brequinar, BAY2402234, leflunomide, |
| chr21 | *TPTE* | 7 | 216 | 14 | TPTE | transmembrane phosphatase with tensin homology | Plasma Membrane | phosphatase | |
| chr3 | *HACL1* | 5 | 142 | 14 | HACL1 | 2-hydroxyacyl-CoA lyase 1 | Cytoplasm | enzyme | |
| chr2 | *RBMS1* | 5 | 111 | 14 | RBMS1 | RNA binding motif single stranded interacting protein 1 | Nucleus | other | |
| chr10 | *MPP7* | 6 | 116 | 14 | MPP7 | membrane palmitoylated protein | Plasma Membrane | other | |
| chr19 | *ACER1* | 5 | 192 | 14 | ACER1 | alkaline ceramidase 1 | Cytoplasm | enzyme | |
| chr19 | *ZNF667* | 7 | 136 | 14 | ZNF667 | zinc finger protein 667 | Nucleus | transcription regulator | |
| chr19 | *SNRNP70* | 6 | 225 | 14 | SNRNP70 | small nuclear ribonucleoprotein U1 subunit 70 | Nucleus | other | |
| chr7 | *DDX56* | 5 | 387 | 14 | DDX56 | DEAD-box helicase 56 | Nucleus | enzyme | |
| chr16 | *CES3* | 6 | 219 | 14 | CES3 | carboxylesterase 3 | Cytoplasm | enzyme | |
| chr3 | *KLHL6* | 6 | 133 | 14 | KLHL6 | kelch like family member 6 | Other | other | |
| chr2 | *DCTN1* | 7 | 264 | 14 | DCTN1 | dynactin subunit 1 | Cytoplasm | other | |
| chr19 | *ZNF136* | 5 | 136 | 14 | ZNF136 | zinc finger protein 136 | Nucleus | transcription regulator | |
| chr19 | *ARRDC2* | 7 | 260 | 14 | ARRDC2 | arrestin domain containing 2 | Other | other | |
| chr6 | *ECT2L* | 5 | 113 | 14 | ECT2L | epithelial cell transforming 2 like | Other | other | |
| chr2 | *PPP1R1C* | 6 | 132 | 14 | PPP1R1C | protein phosphatase 1 regulatory inhibitor subunit 1C | Cytoplasm | phosphatase | |
| chr7 | *PMS2P1* | 6 | 184 | 14 | PMS2P1 | PMS1 homolog 2, mismatch repair system component | Other | other | |
| chr17 | *HS3ST3B1* | 7 | 161 | 14 | HS3ST3B1 | heparan sulfate-glucosamine 3-sulfotransferase 3B1 | Cytoplasm | enzyme | |
| chr19 | *SMG9* | 7 | 226 | 14 | SMG9 | SMG9 nonsense mediated mRNA decay factor | Cytoplasm | other | |
| chr19 | *NFKBID* | 7 | 268 | 14 | NFKBID | NFKB inhibitor delta | Nucleus | transcription regulator | |
| chr7 | *PMS2P3* | 6 | 207 | 14 | PMS2P3 | PMS1 homolog 2, mismatch repair system component | Other | other | |
| chr5 | *C5orf28* | 6 | 379 | 14 | TMEM267 | transmembrane protein 267 | Other | other | |
| chr20 | *RIN2* | 6 | 157 | 14 | RIN2 | Ras and Rab interactor 2 | Cytoplasm | other | |
| chr17 | *TRIM16* | 6 | 155 | 14 | TRIM16 | tripartite motif containing 16 | Cytoplasm | transcription regulator | |
| chr14 | *DCAF4* | 6 | 187 | 14 | DCAF4 | DDB1 and CUL4 associated | Nucleus | other | |
| chr4 | *RP11-539G18.2* | 5 | 120 | 14 | | | | | |
| chr16 | *KDM8* | 6 | 206 | 14 | KDM8 | lysine demethylase 8 | Nucleus | enzyme | |
| chr20 | *DNTTIP1* | 6 | 179 | 14 | DNTTIP1 | deoxynucleotidyltransferase terminal interacting protein 1 | Nucleus | d-dependent nuclear rec | |
| chr17 | *FBF1* | 6 | 199 | 14 | FBF1 | Fas binding factor 1 | Nucleus | other | |
| chr19 | *ZNF600* | 6 | 122 | 14 | ZNF600 | zinc finger protein 600 | Other | other | |

| | | | | | | | | | |
|---|---|---|---|---|---|---|---|---|---|
| chr3 | *TIGIT* | 6 | 231 | 14 | TIGIT | T cell immunoreceptor with Ig and ITIM domains | Plasma Membrane | other | OMP-313M32, BGB-A1217, AB154, tiragolumab, MK- |
| chr6 | *ICK* | 6 | 136 | 14 | ICK | intestinal cell kinase | Cytoplasm | kinase | |
| chr17 | *PCGF2* | 6 | 200 | 14 | PCGF2 | polycomb group ring finger 2 | Nucleus | transcription regulator | |
| chr1 | *ITGA10* | 7 | 222 | 14 | ITGA10 | integrin subunit alpha 10 | Plasma Membrane | other | |
| chr19 | *YIF1B* | 7 | 228 | 14 | YIF1B | Yip1 interacting factor homolog B, membrane trafficking protein | Cytoplasm | other | |
| chr15 | *RP11-386M24.9* | 6 | 385 | 14 | | | | | |
| chr15 | *NDUFAF1* | 5 | 158 | 14 | NDUFAF1 | NADH:ubiquinone oxidoreductase complex | Cytoplasm | other | |
| chr12 | *WNK1* | 5 | 117 | 14 | WNK1 | WNK lysine deficient protein | Cytoplasm | kinase | |
| chr19 | *ABCA7* | 7 | 324 | 14 | ABCA7 | ATP binding cassette subfamily A member 7 | Plasma Membrane | transporter | |
| chr1 | *PANK4* | 8 | 373 | 14 | PANK4 | pantothenate kinase 4 | Cytoplasm | kinase | |
| chr19 | *LILRB4* | 6 | 250 | 14 | LILRB4 | leukocyte immunoglobulin like receptor B4 | Plasma Membrane | other | |
| chr1 | *MAN1A2* | 6 | 163 | 14 | MAN1A2 | mannosidase alpha class 1A member 2 | Cytoplasm | enzyme | |
| chr11 | *ELF5* | 6 | 256 | 14 | ELF5 | E74 like ETS transcription factor | Other | transcription regulator | |
| chr12 | *KRAS* | 5 | 172 | 14 | KRAS | KRAS proto-oncogene, GTPase | Cytoplasm | enzyme | JNJ-74699157, AZD4785, chelerythrine, MRTX849 |
| chr7 | *ZNRF2P2* | 7 | 174 | 14 | ZNRF2P2 | zinc and ring finger 2 | Other | other | |
| chr16 | *RP11-805I24.1* | 6 | 237 | 14 | | | | | |
| chr20 | *PCMTD2* | 6 | 353 | 14 | PCMTD2 | protein-L-isoaspartate (D-aspartate) O-methyltransferase domain containing 2 | Cytoplasm | enzyme | |
| chr10 | *NRBF2* | 6 | 260 | 14 | NRBF2 | nuclear receptor binding factor 2 | Nucleus | transcription regulator | |
| chr11 | *DTX4* | 7 | 185 | 14 | DTX4 | deltex E3 ubiquitin ligase 4 | Cytoplasm | enzyme | |
| chr13 | *GUCY1B2* | 5 | 142 | 14 | GUCY1B2 | guanylate cyclase 1 soluble subunit beta 2 (pseudogene) | Cytoplasm | other | |
| chr1 | *RHBG* | 6 | 336 | 14 | RHBG | Rh family B glycoprotein (gene/pseudogene) | Plasma Membrane | transporter | |
| chr11 | *SPTY2D1* | 6 | 188 | 14 | SPTY2D1 | SPT2 chromatin protein domain containing 1 | Nucleus | other | |
| chr21 | *CBS* | 6 | 320 | 14 | CBS/CBSL | cystathionine beta-synthase | Cytoplasm | enzyme | |
| chrX | *KDM5C* | 6 | 254 | 14 | KDM5C | lysine demethylase 5C | Nucleus | enzyme | |
| chr4 | *DCTD* | 5 | 157 | 14 | DCTD | dCMP deaminase | Cytoplasm | enzyme | tetrahydrouridine |
| chr10 | *ZNF511* | 7 | 187 | 14 | ZNF511 | zinc finger protein 511 | Nucleus | transcription regulator | |
| chr6 | *TBC1D32* | 5 | 190 | 14 | TBC1D32 | TBC1 domain family member 32 | Other | other | |
| chr1 | *RP11-436K8.1* | 6 | 121 | 14 | | | | | |
| chr5 | *HARS* | 5 | 131 | 14 | HARS1 | histidyl-tRNA synthetase 1 | Cytoplasm | enzyme | |
| chr16 | *TRAF7* | 7 | 348 | 14 | TRAF7 | TNF receptor associated factor 7 | Cytoplasm | enzyme | |
| chr1 | *TBX19* | 6 | 244 | 14 | TBX19 | T-box transcription factor 19 | Nucleus | transcription regulator | |
| chr22 | *CTA-211A9.5* | 6 | 166 | 14 | | | | | |
| chrX | *ARHGEF9* | 6 | 124 | 14 | ARHGEF9 | Cdc42 guanine nucleotide exchange factor 9 | Cytoplasm | other | |
| chr7 | *SOSTDC1* | 7 | 198 | 14 | SOSTDC1 | sclerostin domain containing 1 | Extracellular Space | growth factor | |
| chr9 | *RP11-203I2.1* | 5 | 128 | 14 | | | | | |
| chr5 | *TARS* | 5 | 326 | 14 | TARS1 | threonyl-tRNA synthetase 1 | Nucleus | enzyme | |
| chr2 | *MYO1B* | 6 | 145 | 14 | MYO1B | myosin IB | Cytoplasm | enzyme | |
| chr16 | *NPIPA1* | 7 | 295 | 14 | PA1 (includes oth | nuclear pore complex interacting protein family member A3 | Nucleus | other | |
| chr12 | *SLC17A8* | 6 | 114 | 14 | SLC17A8 | solute carrier family 17 member 8 | Plasma Membrane | transporter | |
| chr7 | *ZNF746* | 6 | 254 | 14 | ZNF746 | zinc finger protein 746 | Cytoplasm | transcription regulator | |
| chr6 | *LINC00680* | 6 | 172 | 14 | LINC00680 | | Other | other | |
| chr2 | *CAPN13* | 6 | 156 | 14 | CAPN13 | calpain 13 | Other | other | |
| chr1 | *C1orf50* | 6 | 195 | 14 | C1orf50 | chromosome 1 open reading frame 50 | Other | other | |
| chr1 | *RP1-117O3.2* | 6 | 202 | 14 | | | | | |
| chr17 | *POLG2* | 5 | 166 | 14 | POLG2 | DNA polymerase gamma 2, accessory subunit | Cytoplasm | enzyme | |
| chr19 | *RGL3* | 6 | 227 | 14 | RGL3 | ral guanine nucleotide dissociation stimulator like 3 | Cytoplasm | other | |
| chr4 | *SLC39A8* | 5 | 119 | 14 | SLC39A8 | solute carrier family 39 member 8 | Extracellular Space | transporter | |
| chr17 | *LSMD1* | 6 | 168 | 14 | NAA38 | N(alpha)-acetyltransferase 38, NatC auxiliary subunit | Nucleus | other | |
| chr10 | *MTPAP* | 5 | 119 | 14 | MTPAP | mitochondrial poly(A) polymerase | Cytoplasm | enzyme | |
| chr8 | *LINC00534* | 5 | 130 | 14 | LINC00534 | long intergenic non-protein coding RNA 534 | Other | other | |
| chr9 | *C9orf89* | 6 | 225 | 14 | CARD19 | caspase recruitment domain family member 19 | Cytoplasm | other | |
| chr16 | *ERN2* | 6 | 176 | 14 | ERN2 | endoplasmic reticulum to nucleus signaling 2 | Cytoplasm | kinase | |
| chr19 | *CCDC94* | 5 | 261 | 14 | YJU2 | YJU2 splicing factor homolog | Nucleus | other | |
| chr2 | *METAP1D* | 5 | 148 | 14 | METAP1D | methionyl aminopeptidase type 1D, mitochondrial | Cytoplasm | peptidase | |
| chr8 | *RP11-770E5.1* | 5 | 170 | 14 | | | | | |
| chr11 | *XRRA1* | 6 | 241 | 14 | XRRA1 | X-ray radiation resistance associated 1 | Cytoplasm | other | |
| chr19 | *ZNF709* | 5 | 140 | 14 | ZNF709 | zinc finger protein 709 | Other | other | |
| chr4 | *CDKL2* | 5 | 125 | 14 | CDKL2 | cyclin dependent kinase like 2 | Nucleus | kinase | |
| chr16 | *RP11-525K10.1* | 7 | 145 | 14 | | | | | |
| chr9 | *HIATL1* | 5 | 113 | 14 | MFSD14B | major facilitator superfamily domain containing 14B | Other | other | |
| chr16 | *NOB1* | 6 | 216 | 14 | NOB1 | NIN1 (RPN12) binding protein 1 homolog | Nucleus | enzyme | |

| | | | | | | | | | |
|---|---|---|---|---|---|---|---|---|---|
| chr1 | *B4GALT2* | 7 | 318 | 14 | B4GALT2 | beta-1,4-galactosyltransferase 2 | Cytoplasm | enzyme | |
| chr19 | *URI1* | 7 | 150 | 14 | URI1 | URI1 prefoldin like chaperone | Nucleus | transcription regulator | |
| chr9 | *RP11-65N13.8* | 7 | 221 | 14 | | | | | |
| chr1 | *ATG4C* | 5 | 135 | 14 | ATG4C | autophagy related 4C cysteine peptidase | Cytoplasm | peptidase | |
| chr5 | *CTC-575N7.1* | 5 | 125 | 14 | | | | | |
| chr7 | *ABCB1* | 6 | 198 | 14 | ABCB1 | ATP binding cassette subfamily B member 1 | Plasma Membrane | transporter | OC 144-093, dofequidar, valspodar, eicosapentenoic |
| chr15 | *GOLGA8F* | 5 | 350 | 14 | GOLGA8F | golgin A8 family member F | Other | other | |
| chr1 | *RP5-998N21.7* | 6 | 193 | 14 | | | | | |
| chr1 | *PQLC2* | 7 | 298 | 14 | SLC66A1 | solute carrier family 66 member 1 | Cytoplasm | transporter | |
| chr15 | *CSPG4P9* | 5 | 284 | 14 | | | | | |
| chr3 | *U2SURP* | 4 | 138 | 14 | U2SURP | U2 snRNP associated SURP domain containing | Nucleus | other | |
| chr2 | *AC125232.1* | 5 | 115 | 14 | | | | | |
| chr2 | *AC018730.1* | 6 | 139 | 14 | | | | | |
| chr2 | *SPDYA* | 5 | 118 | 14 | SPDYA | speedy/RINGO cell cycle regulator family member A | Nucleus | other | |
| chr11 | *ZDHHC13* | 5 | 169 | 14 | ZDHHC13 | zinc finger DHHC-type containing | Nucleus | transcription regulator | |
| chr16 | *DBNDD1* | 7 | 381 | 14 | DBNDD1 | dysbindin domain containing 1 | Other | other | |
| chr14 | *RP11-1017G21.4* | 6 | 238 | 14 | | | | | |
| chr1 | *MTHFR* | 5 | 248 | 14 | MTHFR | methylenetetrahydrofolate | Cytoplasm | enzyme | |
| chr12 | *RPAP3* | 6 | 240 | 14 | RPAP3 | RNA polymerase II associated protein 3 | Cytoplasm | enzyme | |
| chr9 | *ABCA2* | 7 | 203 | 14 | ABCA2 | ATP binding cassette subfamily A member 2 | Plasma Membrane | transporter | |
| chr1 | *SH3D21* | 6 | 191 | 14 | SH3D21 | SH3 domain containing 21 | Cytoplasm | other | |
| chr22 | *PNPLA3* | 6 | 178 | 14 | PNPLA3 | patatin like phospholipase domain containing 3 | Cytoplasm | enzyme | |
| chr8 | *RP11-459E5.1* | 7 | 171 | 14 | | | | | |
| chr17 | *LINC00868* | 6 | 175 | 14 | LINC00868 | long intergenic non-protein coding RNA 868 | Other | other | |
| chr20 | *PIGT* | 6 | 337 | 14 | PIGT | phosphatidylinositol glycan anchor biosynthesis class T | Cytoplasm | enzyme | |
| chr19 | *LAIR1* | 6 | 270 | 14 | LAIR1 | leukocyte associated immunoglobulin like receptor 1 | Plasma Membrane | transmembrane receptor | |
| chr17 | *RP11-477N12.3* | 6 | 201 | 14 | | | | | |
| chr12 | *LETMD1* | 6 | 240 | 14 | LETMD1 | LETM1 domain containing 1 | Plasma Membrane | other | |
| chr19 | *SLC25A23* | 6 | 189 | 14 | SLC25A23 | solute carrier family 25 member | Cytoplasm | transporter | |
| chr10 | *FAM149B1* | 5 | 214 | 14 | FAM149B1 | family with sequence similarity 149 member B1 | Other | other | |
| chr16 | *RP11-1348G14.4* | 6 | 227 | 14 | | | | | |
| chr12 | *PLEKHA8P1* | 6 | 202 | 14 | PLEKHA8P1 | pleckstrin homology domain containing A8 pseudogene 1 | Other | other | |
| chr16 | *SLC9A3R2* | 7 | 312 | 14 | SLC9A3R2 | SLC9A3 regulator 2 | Plasma Membrane | transporter | |
| chr2 | *SEPT10* | 6 | 181 | 14 | SEPTIN10 | septin 10 | Cytoplasm | transcription regulator | |
| chr8 | *COL14A1* | 6 | 135 | 13 | COL14A1 | collagen type XIV alpha 1 chain | Extracellular Space | other | collagenase clostridium |
| chr9 | *RP11-146D12.2* | 6 | 162 | 13 | | | | | |
| chr11 | *SCN4B* | 6 | 208 | 13 | SCN4B | sodium voltage-gated channel beta subunit 4 | Plasma Membrane | ion channel | riluzole, zonisamide |
| chr19 | *ZNF699* | 5 | 188 | 13 | ZNF699 | zinc finger protein 699 | Other | other | |
| chr1 | *PRSS38* | 7 | 249 | 13 | PRSS38 | serine protease 38 | Other | other | |
| chr22 | *CSDC2* | 6 | 254 | 13 | CSDC2 | cold shock domain containing C2 | Cytoplasm | other | |
| chr7 | *CEP41* | 5 | 133 | 13 | CEP41 | centrosomal protein 41 | Cytoplasm | other | |
| chr22 | *YPEL1* | 6 | 188 | 13 | YPEL1 | yippee like 1 | Nucleus | enzyme | |
| chr4 | *NELFA* | 6 | 176 | 13 | NELFA | negative elongation factor complex member A | Nucleus | other | |
| chr6 | *TULP1* | 6 | 254 | 13 | TULP1 | TUB like protein 1 | Nucleus | transcription regulator | |
| chr1 | *SNIP1* | 5 | 170 | 13 | SNIP1 | Smad nuclear interacting protein | Nucleus | other | |
| chr13 | *KPNA3* | 5 | 109 | 13 | KPNA3 | karyopherin subunit alpha 3 | Nucleus | transporter | |
| chrX | *TSC22D3* | 6 | 144 | 13 | TSC22D3 | TSC22 domain family member 3 | Nucleus | transcription regulator | |
| chr14 | *PAPLN* | 6 | 311 | 13 | PAPLN | papilin, proteoglycan like sulfated glycoprotein | Extracellular Space | other | |
| chr8 | *RP11-211C9.1* | 7 | 145 | 13 | | | | | |
| chr9 | *IARS* | 5 | 129 | 13 | IARS1 | isoleucyl-tRNA synthetase 1 | Cytoplasm | enzyme | |
| chr3 | *NKIRAS1* | 6 | 198 | 13 | NKIRAS1 | NFKB inhibitor interacting Ras | Other | enzyme | |
| chr7 | *KRIT1* | 5 | 197 | 13 | KRIT1 | KRIT1 ankyrin repeat containing | Plasma Membrane | other | |
| chr9 | *TRIM14* | 6 | 177 | 13 | TRIM14 | tripartite motif containing 14 | Cytoplasm | other | |
| chr19 | *USHBP1* | 6 | 265 | 13 | USHBP1 | USH1 protein network component harmonin binding protein 1 | Other | other | |
| chr18 | *RP11-449D8.5* | 6 | 119 | 13 | | | | | |
| chr2 | *LRP2* | 6 | 152 | 13 | LRP2 | LDL receptor related protein 2 | Plasma Membrane | transporter | |
| chr5 | *GEMIN5* | 6 | 127 | 13 | GEMIN5 | gem nuclear organelle associated protein 5 | Nucleus | other | |
| chr1 | *TMEM222* | 5 | 251 | 13 | TMEM222 | transmembrane protein 222 | Other | other | |
| chr7 | *TSPAN33* | 6 | 163 | 13 | TSPAN33 | tetraspanin 33 | Plasma Membrane | other | |
| chr17 | *ALOX12B* | 6 | 313 | 13 | ALOX12B | arachidonate 12-lipoxygenase, 12R type | Cytoplasm | enzyme | |
| chr2 | *CHN1* | 5 | 100 | 13 | CHN1 | chimerin 1 | Cytoplasm | other | |
| chr1 | *IBA57* | 7 | 367 | 13 | IBA57 | iron-sulfur cluster assembly factor IBA57 | Cytoplasm | other | |
| chr9 | *TUBBP5* | 7 | 355 | 13 | TUBBP5 | tubulin beta pseudogene 5 | Other | other | |
| chr5 | *CTD-2353F22.1* | 7 | 392 | 13 | | | | | |
| chr15 | *ANXA2* | 6 | 135 | 13 | ANXA2 | annexin A2 | Plasma Membrane | other | |
| chr20 | *FAM210B* | 6 | 312 | 13 | FAM210B | family with sequence similarity 210 member B | Extracellular Space | other | |

| | | | | | | | | | |
|---|---|---|---|---|---|---|---|---|---|
| chr6 | *ELOVL5* | 5 | 154 | 13 | ELOVL5 | ELOVL fatty acid elongase 5 | Cytoplasm | enzyme | |
| chr1 | *CHRNB2* | 6 | 390 | 13 | CHRNB2 | cholinergic receptor nicotinic beta 2 subunit | Plasma Membrane | transmembrane recepto | succinylcholine, lobeline, rapacuronium, doxacurium, vecuronium, pipecuronium, ABT 418, metocurine, mivacurium, D-tubocurarine, nicotine, arecoline, pancuronium, atracurium, isoflurane, cisatracurium, rocuronium, amobarbital, mecamylamine |
| chr17 | *SCARF1* | 6 | 363 | 13 | SCARF1 | scavenger receptor class F member 1 | Plasma Membrane | transmembrane recepto | |
| chr5 | *PHYKPL* | 5 | 219 | 13 | PHYKPL | 5-phosphohydroxy-L-lysine phospho-lyase | Cytoplasm | enzyme | |
| chr22 | *TTLL12* | 6 | 291 | 13 | TTLL12 | tubulin tyrosine ligase like 12 | Cytoplasm | other | |
| chr15 | *SORD* | 6 | 216 | 13 | SORD | sorbitol dehydrogenase | Cytoplasm | enzyme | |
| chr2 | *IFT172* | 6 | 153 | 13 | IFT172 | intraflagellar transport 172 | Extracellular Space | other | |
| chr8 | *DUSP4* | 6 | 247 | 13 | DUSP4 | dual specificity phosphatase 4 | Nucleus | phosphatase | |
| chr10 | *CTGLF12P* | 5 | 411 | 13 | AGAP12P | ArfGAP with GTPase domain, ankyrin repeat and PH domain 12, pseudogene | Other | other | |
| chr3 | *LINC00886* | 5 | 172 | 13 | LINC00886 | long intergenic non-protein coding RNA 886 | Other | other | |
| chr7 | *AP5Z1* | 7 | 212 | 13 | AP5Z1 | adaptor related protein complex 5 subunit zeta 1 | Nucleus | other | |
| chr5 | *SDHAP3* | 7 | 479 | 13 | SDHAP3 | succinate dehydrogenase complex flavoprotein subunit A | Other | other | |
| chr9 | *C9orf57* | 5 | 111 | 13 | C9orf57 | chromosome 9 open reading frame 57 | Other | other | |
| chr1 | *RRP15* | 7 | 219 | 13 | RRP15 | ribosomal RNA processing 15 homolog | Nucleus | other | |
| chr7 | *GRM3* | 7 | 163 | 13 | GRM3 | glutamate metabotropic receptor | Plasma Membrane | G-protein coupled recepto | fasoracetam |
| chr16 | *TELO2* | 7 | 242 | 13 | TELO2 | telomere maintenance 2 | Cytoplasm | other | |
| chr9 | *RP11-213G2.1* | 5 | 132 | 13 | | | | | |
| chr9 | *PIP5KL1* | 6 | 437 | 13 | PIP5KL1 | phosphatidylinositol-4-phosphate 5-kinase like 1 | Plasma Membrane | kinase | |
| chr22 | *GTPBP1* | 6 | 206 | 13 | GTPBP1 | GTP binding protein 1 | Cytoplasm | enzyme | |
| chr1 | *PLA2G2F* | 6 | 261 | 13 | PLA2G2F | phospholipase A2 group IIF | Extracellular Space | enzyme | |
| chr22 | *XRCC6* | 5 | 121 | 13 | XRCC6 | X-ray repair cross complementing | Nucleus | enzyme | |
| chr17 | *ABCA9* | 6 | 128 | 13 | ABCA9 | ATP binding cassette subfamily A member 9 | Cytoplasm | transporter | |
| chr5 | *PDE6A* | 6 | 118 | 13 | PDE6A | phosphodiesterase 6A | Plasma Membrane | enzyme | |
| chr4 | *RP11-8L2.1* | 6 | 130 | 13 | | | | | |
| chr18 | *SIGLEC15* | 5 | 189 | 13 | SIGLEC15 | sialic acid binding Ig like lectin 15 | Plasma Membrane | other | NC318 |
| chr6 | *IRF4* | 6 | 179 | 13 | IRF4 | interferon regulatory factor 4 | Nucleus | transcription regulator | |
| chrX | *CHM* | 5 | 119 | 13 | CHM | CHM Rab escort protein | Cytoplasm | enzyme | |
| chr19 | *TMC4* | 7 | 289 | 13 | TMC4 | transmembrane channel like 4 | Cytoplasm | other | |
| chr16 | *GAS8* | 7 | 219 | 13 | GAS8 | growth arrest specific 8 | Cytoplasm | other | |
| chr3 | *MYLK-AS1* | 6 | 173 | 13 | MYLK-AS1 | MYLK antisense RNA 1 | Other | other | |
| chr1 | *RGS8* | 7 | 190 | 13 | RGS8 | regulator of G protein signaling 8 | Cytoplasm | enzyme | |
| chr2 | *VRK2* | 6 | 164 | 13 | VRK2 | VRK serine/threonine kinase 2 | Nucleus | kinase | |
| chr17 | *CTD-2532D12.4* | 6 | 215 | 13 | | | | | |
| chr15 | *UNC45A* | 7 | 385 | 13 | UNC45A | unc-45 myosin chaperone A | Plasma Membrane | other | |
| chr11 | *RP11-626H12.1* | 6 | 152 | 13 | | | | | |
| chr12 | *RAPGEF3* | 6 | 590 | 13 | RAPGEF3 | Rap guanine nucleotide exchange factor 3 | Nucleus | other | |
| chr8 | *CASC9* | 5 | 183 | 13 | CASC9 | cancer susceptibility 9 | Other | other | |
| chr14 | *IFT43* | 5 | 135 | 13 | IFT43 | intraflagellar transport 43 | Cytoplasm | other | |
| chr12 | *DAO* | 6 | 174 | 13 | DAO | D-amino acid oxidase | Cytoplasm | enzyme | |
| chr8 | *CLN8* | 6 | 234 | 13 | CLN8 | CLN8 transmembrane ER and ERGIC protein | Cytoplasm | other | |
| chr2 | *CEP68* | 6 | 185 | 13 | CEP68 | centrosomal protein 68 | Cytoplasm | other | |
| chr12 | *C12orf43* | 5 | 202 | 13 | C12orf43 | chromosome 12 open reading frame 43 | Other | other | |
| chr9 | *PPP1R26-AS1* | 6 | 310 | 13 | PPP1R26-AS1 | PPP1R26 antisense RNA 1 | Other | other | |
| chr11 | *RP11-886D15.1* | 6 | 130 | 13 | | | | | |
| chr20 | *RP5-843L14.1* | 6 | 242 | 13 | | | | | |
| chr19 | *SSBP4* | 6 | 365 | 13 | SSBP4 | single stranded DNA binding protein 4 | Nucleus | transcription regulator | |
| chr15 | *CSPG4P10* | 5 | 269 | 13 | CSPG4P10 | chondroitin sulfate proteoglycan 4 pseudogene 10 | Other | other | |
| chr17 | *KIAA0753* | 6 | 182 | 13 | KIAA0753 | KIAA0753 | Cytoplasm | other | |
| chr3 | *FGF12* | 5 | 147 | 13 | FGF12 | fibroblast growth factor 12 | Extracellular Space | other | |
| chr3 | *RPP14* | 6 | 176 | 13 | RPP14 | ribonuclease P/MRP subunit p14 | Nucleus | enzyme | |
| chr12 | *HNF1A* | 5 | 239 | 13 | HNF1A | HNF1 homeobox A | Nucleus | transcription regulator | |
| chr16 | *KIF22* | 6 | 182 | 13 | KIF22 | kinesin family member 22 | Nucleus | other | |
| chr3 | *NXPE3* | 6 | 170 | 13 | NXPE3 | neurexophilin and PC-esterase domain family member 3 | Other | other | |
| chr1 | *CYP4Z2P* | 6 | 154 | 13 | CYP4Z2P | cytochrome P450 family 4 subfamily Z member 2, | Other | other | |
| chr8 | *SPAG1* | 5 | 118 | 13 | SPAG1 | sperm associated antigen 1 | Cytoplasm | other | |

| | | | | | | | | | |
|---|---|---|---|---|---|---|---|---|---|
| chr1 | *DPT* | 7 | 213 | 13 | DPT | dermatopontin | Extracellular Space | other | |
| chr1 | *RP5-886K2.3* | 6 | 137 | 13 | | | | | |
| chr17 | *GFAP* | 6 | 457 | 13 | GFAP | glial fibrillary acidic protein | Cytoplasm | other | |
| chr14 | *RP11-112J1.2* | 5 | 168 | 13 | | | | | |
| chr16 | *RAB26* | 6 | 306 | 13 | RAB26 | RAB26, member RAS oncogene family | Plasma Membrane | enzyme | |
| chrX | *FUNDC1* | 5 | 142 | 13 | FUNDC1 | FUN14 domain containing 1 | Cytoplasm | other | |
| chr8 | *RP11-150O12.4* | 5 | 140 | 13 | | | | | |
| chr10 | *C10orf112* | 5 | 117 | 13 | MALRD1 | MAM and LDL receptor class A domain containing 1 | Cytoplasm | other | |
| chr5 | *TIMD4* | 6 | 145 | 13 | TIMD4 | T cell immunoglobulin and mucin domain containing 4 | Plasma Membrane | other | |
| chr2 | *AC096579.13* | 6 | 206 | 13 | | | | | |
| chr17 | *RP11-15E18.1* | 6 | 126 | 13 | | | | | |
| chr11 | *PITPNM1* | 7 | 306 | 13 | PITPNM1 | phosphatidylinositol transfer protein membrane associated 1 | Cytoplasm | transporter | |
| chr20 | *SEC23B* | 5 | 128 | 13 | SEC23B | SEC23 homolog B, coat complex II component | Extracellular Space | transporter | |
| chr10 | *DPYSL4* | 7 | 337 | 13 | DPYSL4 | dihydropyrimidinase like 4 | Cytoplasm | enzyme | |
| chr3 | *TSC22D2* | 6 | 194 | 13 | TSC22D2 | TSC22 domain family member 2 | Nucleus | transcription regulator | |
| chr2 | *ZNF804A* | 4 | 136 | 13 | ZNF804A | zinc finger protein 804A | Plasma Membrane | other | |
| chr10 | *AGAP7* | 5 | 249 | 13 | AP5 (includes oth | ArfGAP with GTPase domain, ankyrin repeat and PH domain 5 | Other | other | |
| chr9 | *NDOR1* | 7 | 354 | 13 | NDOR1 | NADPH dependent diflavin oxidoreductase 1 | Cytoplasm | enzyme | |
| chr5 | *SNX2* | 5 | 134 | 13 | SNX2 | sorting nexin 2 | Cytoplasm | transporter | |
| chr11 | *PPME1* | 6 | 169 | 13 | PPME1 | protein phosphatase methylesterase 1 | Nucleus | enzyme | |
| chr19 | *CTB-171A8.1* | 5 | 175 | 13 | | | | | |
| chr15 | *RLBP1* | 6 | 306 | 13 | RLBP1 | retinaldehyde binding protein 1 | Cytoplasm | transporter | |
| chr17 | *LRRC37A16P* | 6 | 137 | 13 | LRRC37A16P | leucine rich repeat containing 37 member A16, pseudogene | Other | other | |
| chr8 | *AGPAT5* | 6 | 138 | 13 | AGPAT5 | 1-acylglycerol-3-phosphate O-acyltransferase 5 | Cytoplasm | enzyme | |
| chr10 | *LOXL4* | 6 | 272 | 13 | LOXL4 | lysyl oxidase like 4 | Extracellular Space | enzyme | |
| chr1 | *TDRKH* | 6 | 154 | 13 | TDRKH | tudor and KH domain containing | Cytoplasm | other | |
| chr8 | *CHRNA2* | 6 | 254 | 13 | CHRNA2 | cholinergic receptor nicotinic alpha 2 subunit | Plasma Membrane | transmembrane recepto | carbamylcholine, succinylcholine, rapacuronium, doxacurium, vecuronium, pipecuronium, biperiden, ABT 418, metocurine, mivacurium, D-tubocurarine, nicotine, arecoline, pancuronium, atracurium, isoflurane, cisatracurium, rocuronium, decamethonium, mecamylamine |
| chr17 | *TBC1D28* | 6 | 179 | 13 | TBC1D28 | TBC1 domain family member 28 | Other | other | |
| chrX | *PFKFB1* | 6 | 255 | 13 | PFKFB1 | 6-phosphofructo-2-kinase/fructose-2,6- | Cytoplasm | kinase | |
| chr3 | *KLF15* | 7 | 351 | 13 | KLF15 | Kruppel like factor 15 | Nucleus | transcription regulator | |
| chr1 | *LEPRE1* | 6 | 219 | 13 | P3H1 | prolyl 3-hydroxylase 1 | Nucleus | enzyme | |
| chr11 | *CTD-2005H7.2* | 6 | 199 | 13 | | | | | |
| chr2 | *RFTN2* | 6 | 155 | 13 | RFTN2 | raftlin family member 2 | Other | other | |
| chr1 | *RP11-34P13.13* | 6 | 200 | 13 | | | | | |
| chr19 | *CYP4F12* | 7 | 161 | 13 | CYP4F12 | cytochrome P450 family 4 subfamily F member 12 | Cytoplasm | enzyme | |
| chr6 | *BMP5* | 5 | 104 | 13 | BMP5 | bone morphogenetic protein 5 | Extracellular Space | growth factor | |
| chr1 | *HCRTR1* | 5 | 312 | 13 | HCRTR1 | hypocretin receptor 1 | Plasma Membrane | -protein coupled recept | suvorexant |
| chr1 | *DBT* | 6 | 180 | 13 | DBT | dihydrolipoamide branched chain transacylase E2 | Cytoplasm | enzyme | |
| chr4 | *AC108142.1* | 6 | 186 | 13 | | | | | |
| chr3 | *CDHR4* | 5 | 221 | 13 | CDHR4 | cadherin related family member 4 | Other | other | |
| chr19 | *SEC1P* | 6 | 203 | 13 | SEC1P | secretory blood group 1, | Other | other | |
| chr16 | *GPR139* | 7 | 141 | 13 | GPR139 | G protein-coupled receptor 139 | Plasma Membrane | -protein coupled recept | |
| chr15 | *PPP1R14D* | 5 | 163 | 13 | PPP1R14D | protein phosphatase 1 regulatory inhibitor subunit 14D | Cytoplasm | other | |
| chr10 | *RP11-313J2.1* | 6 | 154 | 13 | | | | | |
| chr19 | *KCNC3* | 7 | 310 | 13 | KCNC3 | potassium voltage-gated channel subfamily C member 3 | Plasma Membrane | ion channel | dalfampridine |
| chr19 | *TARM1* | 6 | 188 | 13 | TARM1 | T cell-interacting, activating receptor on myeloid cells 1 | Plasma Membrane | other | |
| chr13 | *CARKD* | 6 | 292 | 13 | NAXD | NAD(P)HX dehydratase | Cytoplasm | enzyme | |
| chr3 | *MOBP* | 5 | 114 | 13 | MOBP | myelin associated oligodendrocyte basic protein | Cytoplasm | other | |
| chr10 | *CSGALNACT2* | 6 | 144 | 13 | CSGALNACT2 | chondroitin sulfate N-acetylgalactosaminyltransferase | Cytoplasm | enzyme | |
| chr19 | *LSM4* | 6 | 277 | 13 | LSM4 | LSM4 homolog, U6 small nuclear RNA and mRNA degradation associated | Nucleus | other | |
| chr2 | *CASP8* | 5 | 114 | 13 | CASP8 | caspase 8 | Nucleus | peptidase | |
| chr21 | *CLDN14* | 5 | 154 | 13 | CLDN14 | claudin 14 | Plasma Membrane | other | |

| | | | | | | | | | |
|---|---|---|---|---|---|---|---|---|---|
| chr7 | *AC083864.3* | 7 | 217 | 13 | | | | | |
| chr12 | *METAP2* | 5 | 156 | 13 | METAP2 | methionyl aminopeptidase 2 | Cytoplasm | peptidase | XMT-1107, O-(chloroacetylcarbamoyl)fumagillol, beloranib, PPI-2458, SDX-7320, |
| chr1 | *LACTBL1* | 5 | 211 | 13 | LACTBL1 | lactamase beta like 1 | Other | other | |
| chr12 | *RP11-230G5.2* | 6 | 170 | 13 | | | | | |
| chr9 | *L09NC01-254D11.* | 6 | 352 | 13 | | | | | |
| chr19 | *ZNF253* | 4 | 128 | 13 | ZNF253 | zinc finger protein 253 | Nucleus | transcription regulator | |
| chr11 | *SLC22A11* | 6 | 314 | 13 | SLC22A11 | solute carrier family 22 member 11 | Plasma Membrane | transporter | colchicine/probenecid, probenecid |
| chr12 | *LRP6* | 4 | 104 | 13 | LRP6 | LDL receptor related protein 6 | Plasma Membrane | transmembrane receptor | |
| chr1 | *PKLR* | 6 | 372 | 13 | PKLR | pyruvate kinase L/R | Cytoplasm | kinase | |
| chr14 | *RP11-298I3.1* | 4 | 126 | 13 | | | | | |
| chr1 | *RP4-735C1.6* | 6 | 181 | 13 | | | | | |
| chr11 | *PLCB3* | 7 | 374 | 13 | PLCB3 | phospholipase C beta 3 | Cytoplasm | enzyme | |
| chr20 | *ZNF663P* | 6 | 255 | 13 | ZNF663P | zinc finger protein 663, | Other | other | |
| chr17 | *BRIP1* | 5 | 115 | 13 | BRIP1 | BRCA1 interacting protein C-terminal helicase 1 | Nucleus | enzyme | |
| chr19 | *HSD17B14* | 6 | 206 | 13 | HSD17B14 | hydroxysteroid 17-beta dehydrogenase 14 | Cytoplasm | enzyme | |
| chrX | *CCDC120* | 6 | 248 | 13 | CCDC120 | coiled-coil domain containing 120 | Plasma Membrane | other | |
| chr12 | *RP11-542B15.1* | 6 | 116 | 13 | | | | | |
| chr3 | *RP11-451B8.1* | 5 | 124 | 13 | | | | | |
| chr8 | *CTD-2541M15.1* | 6 | 129 | 13 | | | | | |
| chr4 | *SEC24B* | 6 | 114 | 13 | SEC24B | SEC24 homolog B, COPII coat complex component | Cytoplasm | transporter | |
| chr15 | *AC068831.10* | 5 | 270 | 13 | | | | | |
| chr5 | *OXCT1* | 6 | 464 | 13 | OXCT1 | 3-oxoacid CoA-transferase 1 | Cytoplasm | enzyme | |
| chr16 | *PPP4C* | 6 | 215 | 13 | PPP4C | protein phosphatase 4 catalytic subunit | Cytoplasm | phosphatase | |
| chr9 | *FAM189A2* | 6 | 115 | 13 | FAM189A2 | family with sequence similarity 189 member A2 | Other | other | |
| chr19 | *SYT3* | 6 | 223 | 13 | SYT3 | synaptotagmin 3 | Cytoplasm | transporter | |
| chr8 | *FGL1* | 5 | 129 | 13 | FGL1 | fibrinogen like 1 | Extracellular Space | other | |
| chr3 | *RYBP* | 5 | 106 | 13 | RYBP | RING1 and YY1 binding protein | Nucleus | transcription regulator | |
| chr10 | *RP11-523O18.1* | 6 | 335 | 13 | | | | | |
| chrX | *BCAP31* | 6 | 277 | 13 | BCAP31 | B cell receptor associated protein | Cytoplasm | transporter | |
| chr9 | *CNTFR* | 6 | 294 | 13 | CNTFR | ciliary neurotrophic factor | Plasma Membrane | transmembrane receptor | |
| chr17 | *SECTM1* | 7 | 275 | 13 | SECTM1 | secreted and transmembrane 1 | Extracellular Space | other | |
| chr4 | *PDE6B* | 6 | 274 | 13 | PDE6B | phosphodiesterase 6B | Cytoplasm | enzyme | |
| chr1 | *SDF4* | 7 | 206 | 13 | SDF4 | stromal cell derived factor 4 | Cytoplasm | other | |
| chr6 | *HMGCLL1* | 5 | 145 | 13 | HMGCLL1 | 3-hydroxymethyl-3-methylglutaryl-CoA lyase like 1 | Cytoplasm | enzyme | |
| chr12 | *TDG* | 4 | 132 | 13 | TDG | thymine DNA glycosylase | Nucleus | enzyme | |
| chr1 | *KMO* | 6 | 146 | 13 | KMO | kynurenine 3-monooxygenase | Cytoplasm | enzyme | |
| chr3 | *TTC21A* | 6 | 222 | 13 | TTC21A | tetratricopeptide repeat domain | Extracellular Space | other | |
| chr1 | *CELA3A* | 5 | 220 | 13 | CELA3A | chymotrypsin like elastase 3A | Other | peptidase | |
| chr10 | *PPRC1* | 6 | 205 | 13 | PPRC1 | PPARG related coactivator 1 | Nucleus | transcription regulator | |
| chr4 | *RFC1* | 4 | 100 | 13 | RFC1 | replication factor C subunit 1 | Nucleus | transcription regulator | |
| chr11 | *TMPRSS5* | 6 | 243 | 13 | TMPRSS5 | transmembrane serine protease 5 | Plasma Membrane | peptidase | |
| chr5 | *LCP2* | 7 | 185 | 13 | LCP2 | lymphocyte cytosolic protein 2 | Cytoplasm | other | |
| chr2 | *STAMBP* | 5 | 138 | 13 | STAMBP | STAM binding protein | Nucleus | enzyme | |
| chr1 | *MASP2* | 6 | 204 | 13 | MASP2 | mannan binding lectin serine peptidase 2 | Extracellular Space | peptidase | |
| chr12 | *SCYL2* | 5 | 105 | 13 | SCYL2 | SCY1 like pseudokinase 2 | Cytoplasm | other | |
| chr20 | *ZNF512B* | 7 | 379 | 13 | ZNF512B | zinc finger protein 512B | Nucleus | transcription regulator | |
| chr2 | *MYO3B* | 6 | 116 | 13 | MYO3B | myosin IIIB | Plasma Membrane | kinase | |
| chr8 | *FBXO25* | 5 | 173 | 13 | FBXO25 | F-box protein 25 | Nucleus | enzyme | |
| chr1 | *MKNK1-AS1* | 5 | 214 | 13 | MKNK1-AS1 | MKNK1 antisense RNA 1 | Other | other | |
| chr3 | *RP11-20B7.1* | 6 | 107 | 13 | | | | | |
| chrX | *SLC25A43* | 5 | 137 | 13 | SLC25A43 | solute carrier family 25 member | Cytoplasm | transporter | |
| chr1 | *RP5-952N6.1* | 6 | 172 | 13 | | | | | |
| chr19 | *MEIS3* | 6 | 226 | 13 | MEIS3 | Meis homeobox 3 | Nucleus | transcription regulator | |
| chr5 | *RP11-549K20.1* | 5 | 335 | 13 | | | | | |
| chr14 | *RP11-298I3.5* | 4 | 157 | 13 | | | | | |
| chr11 | *TSG101* | 6 | 131 | 13 | TSG101 | tumor susceptibility 101 | Cytoplasm | transcription regulator | |
| chr5 | *H2AFY* | 5 | 211 | 13 | MACROH2A1 | macroH2A.1 histone | Nucleus | other | |
| chr5 | *SERF1A* | 5 | 140 | 13 | SERF1A/SERF1B | small EDRK-rich factor 1A | Other | other | |
| chr12 | *NACA* | 6 | 169 | 13 | NACA | nascent polypeptide associated complex subunit alpha | Cytoplasm | transcription regulator | |
| chr1 | *IPO13* | 6 | 257 | 13 | IPO13 | importin 13 | Nucleus | transporter | |
| chr20 | *RAE1* | 6 | 329 | 13 | RAE1 | ribonucleic acid export 1 | Nucleus | other | |
| chr20 | *CST7* | 5 | 274 | 13 | CST7 | cystatin F | Extracellular Space | other | |
| chr19 | *RAB11B* | 6 | 212 | 13 | RAB11B | RAB11B, member RAS oncogene family | Cytoplasm | enzyme | |
| chr15 | *DENND4A* | 5 | 137 | 13 | DENND4A | DENN domain containing 4A | Nucleus | other | |
| chr19 | *PLEKHG2* | 6 | 281 | 13 | PLEKHG2 | pleckstrin homology and RhoGEF domain containing G2 | Cytoplasm | other | |
| chr7 | *AC003984.1* | 5 | 138 | 13 | | | | | |
| chr5 | *ST8SIA4* | 5 | 149 | 13 | ST8SIA4 | ST8 alpha-N-acetyl-neuraminide alpha-2,8-sialyltransferase 4 | Cytoplasm | enzyme | |
| chr7 | *SEPT7P2* | 5 | 188 | 13 | SEPTIN7P2 | septin 7 pseudogene 2 | Other | other | |
| chr8 | *NAT1* | 6 | 146 | 13 | NAT1 | N-acetyltransferase 1 | Cytoplasm | enzyme | |
| chr7 | *YAE1D1* | 5 | 172 | 13 | YAE1 | YAE1 maturation factor of ABCE1 | Other | other | |
| chr16 | *RP11-46C24.5* | 6 | 289 | 13 | | | | | |
| chr2 | *ING5* | 6 | 231 | 13 | ING5 | inhibitor of growth family member | Nucleus | other | |
| chr9 | *RP5-1050E16.2* | 6 | 186 | 13 | | | | | |

| | | | | | | | | | |
|---|---|---|---|---|---|---|---|---|---|
| chr22 | *TPST2* | 5 | 169 | 13 | TPST2 | tyrosylprotein sulfotransferase 2 | Cytoplasm | enzyme | |
| chr5 | *MAP3K1* | 6 | 116 | 13 | MAP3K1 | mitogen-activated protein kinase kinase kinase 1 | Cytoplasm | kinase | E 6201 |
| chr11 | *SERPINH1* | 6 | 320 | 13 | SERPINH1 | serpin family H member 1 | Extracellular Space | other | |
| chr1 | *NUAK2* | 6 | 308 | 13 | NUAK2 | NUAK family kinase 2 | Other | kinase | |
| chr2 | *CKAP2L* | 6 | 187 | 13 | CKAP2L | cytoskeleton associated protein 2 | Cytoplasm | other | |
| chr2 | *AMMECR1L* | 6 | 168 | 13 | AMMECR1L | AMMECR1 like | Other | other | |
| chr19 | *ZNF180* | 5 | 167 | 13 | ZNF180 | zinc finger protein 180 | Nucleus | transcription regulator | |
| chr17 | *KRT32* | 6 | 192 | 13 | KRT32 | keratin 32 | Cytoplasm | other | |
| chr12 | *TM7SF3* | 5 | 170 | 13 | TM7SF3 | transmembrane 7 superfamily member 3 | Plasma Membrane | other | |
| chr5 | *TAF9* | 5 | 140 | 13 | TAF9 | TATA-box binding protein associated factor 9 | Nucleus | transcription regulator | |
| chr15 | *GOLGA8R* | 4 | 177 | 13 | GA8K (includes ot | golgin A8 family member K | Other | other | |
| chr22 | *ENTHD1* | 5 | 129 | 13 | ENTHD1 | ENTH domain containing 1 | Other | other | |
| chrX | *TRO* | 6 | 245 | 13 | TRO | trophinin | Plasma Membrane | other | |
| chr7 | *FAM221A* | 5 | 170 | 13 | FAM221A | family with sequence similarity 221 member A | Other | other | |
| chr1 | *MR1* | 6 | 264 | 13 | MR1 | major histocompatibility complex, class I-related | Plasma Membrane | transmembrane receptor | anti-thymocyte globulin (rabbit) |
| chr8 | *ZNF704* | 6 | 137 | 13 | ZNF704 | zinc finger protein 704 | Nucleus | other | |
| chr1 | *C1orf227* | 5 | 195 | 13 | SPATA45 | spermatogenesis associated 45 | Other | other | |
| chr7 | *CCZ1B* | 5 | 199 | 13 | CCZ1/CCZ1B | CCZ1 homolog B, vacuolar protein trafficking and biogenesis | Cytoplasm | other | |
| chr18 | *KATNAL2* | 6 | 498 | 13 | KATNAL2 | katanin catalytic subunit A1 like 2 | Cytoplasm | other | |
| chr6 | *RP11-514O12.4* | 6 | 163 | 13 | | | | | |
| chr7 | *TRIM50* | 6 | 201 | 13 | TRIM50 | tripartite motif containing 50 | Cytoplasm | other | |
| chr19 | *LILRB1* | 6 | 250 | 13 | LILRB1 | leukocyte immunoglobulin like receptor B1 | Plasma Membrane | transmembrane receptor | |
| chr16 | *CHTF8* | 5 | 188 | 13 | CHTF8 | chromosome transmission fidelity factor 8 | Cytoplasm | other | |
| chrX | *PORCN* | 5 | 260 | 13 | PORCN | porcupine O-acyltransferase | Cytoplasm | enzyme | CGX1321, ETC-1922159, LGK-974 |
| chr7 | *TSC22D4* | 7 | 393 | 13 | TSC22D4 | TSC22 domain family member 4 | Nucleus | transcription regulator | |
| chr12 | *RIMKLB* | 5 | 118 | 13 | RIMKLB | ribosomal modification protein rimK like family member B | Cytoplasm | enzyme | |
| chr7 | *AC008060.7* | 6 | 235 | 13 | | | | | |
| chr12 | *YEATS4* | 6 | 154 | 13 | YEATS4 | YEATS domain containing 4 | Nucleus | transcription regulator | |
| chr22 | *GGT2* | 5 | 228 | 13 | GT2/LOC1027241 | gamma-glutamyltransferase 2 | Plasma Membrane | enzyme | |
| chr5 | *CTD-2044J15.1* | 7 | 348 | 13 | | | | | |
| chr16 | *CPNE7* | 7 | 276 | 13 | CPNE7 | copine 7 | Cytoplasm | transporter | |
| chr14 | *CINP* | 5 | 162 | 13 | CINP | cyclin dependent kinase 2 interacting protein | Nucleus | other | |
| chr5 | *TPPP* | 7 | 839 | 13 | TPPP | tubulin polymerization promoting protein | Cytoplasm | other | |
| chr22 | *GAB4* | 5 | 196 | 13 | GAB4 | GRB2 associated binding protein family member 4 | Other | other | |
| chr14 | *CTD-3006G17.2* | 5 | 134 | 13 | | | | | |
| chr5 | *RP11-756H20.1* | 6 | 113 | 13 | | | | | |
| chr19 | *CDC37* | 6 | 222 | 13 | CDC37 | cell division cycle 37 | Cytoplasm | other | |
| chr1 | *MIA3* | 6 | 189 | 13 | MIA3 | MIA SH3 domain ER export | Cytoplasm | other | |
| chr11 | *CATSPER1* | 6 | 454 | 13 | CATSPER1 | cation channel sperm associated | Plasma Membrane | ion channel | |
| chr8 | *RP11-134O21.1* | 5 | 133 | 13 | | | | | |
| chr20 | *HELZ2* | 7 | 362 | 13 | HELZ2 | helicase with zinc finger 2 | Nucleus | transcription regulator | |
| chr12 | *RPS11P6* | 6 | 141 | 13 | RPS11P6 | ribosomal protein S11 | Other | other | |
| chr5 | *CTC-207P7.1* | 6 | 145 | 13 | | | | | |
| chr7 | *SERPINE1* | 6 | 203 | 13 | SERPINE1 | serpin family E member 1 | Extracellular Space | other | drotrecogin alfa |
| chr2 | *ASTL* | 6 | 319 | 13 | ASTL | astacin like metalloendopeptidase | Cytoplasm | peptidase | |
| chr19 | *ANGPTL4* | 6 | 224 | 13 | ANGPTL4 | angiopoietin like 4 | Extracellular Space | other | |
| chr1 | *RSPO1* | 6 | 187 | 13 | RSPO1 | R-spondin 1 | Plasma Membrane | other | |
| chr4 | *RP11-63A11.1* | 5 | 127 | 13 | | | | | |
| chr12 | *SUOX* | 6 | 355 | 13 | SUOX | sulfite oxidase | Cytoplasm | enzyme | |
| chr16 | *CRYM* | 6 | 138 | 13 | CRYM | crystallin mu | Cytoplasm | enzyme | |
| chr8 | *BOP1* | 6 | 293 | 13 | BOP1 | BOP1 ribosomal biogenesis | Nucleus | other | |
| chr2 | *SEMA4F* | 6 | 159 | 13 | SEMA4F | ssemaphorin 4F | Plasma Membrane | other | |
| chr16 | *RP11-510M2.1* | 6 | 156 | 13 | | | | | |
| chr15 | *MAP1A* | 6 | 201 | 13 | MAP1A | microtubule associated protein | Cytoplasm | other | estramustine |
| chr16 | *GOT2* | 6 | 136 | 13 | GOT2 | glutamic-oxaloacetic | Cytoplasm | enzyme | |
| chr19 | *CTC-513N18.7* | 6 | 111 | 13 | | | | | |
| chr10 | *PROSER2-AS1* | 5 | 197 | 13 | PROSER2-AS1 | PROSER2 antisense RNA 1 | Other | other | |
| chr10 | *ACBD5* | 6 | 125 | 13 | ACBD5 | acyl-CoA binding domain containing 5 | Nucleus | other | |
| chr11 | *EHBP1L1* | 6 | 558 | 13 | EHBP1L1 | EH domain binding protein 1 like | Other | other | |
| chr17 | *PELP1* | 6 | 197 | 13 | PELP1 | proline, glutamate and leucine rich protein 1 | Nucleus | other | |
| chr3 | *LMLN* | 4 | 103 | 13 | LMLN | leishmanolysin like peptidase | Plasma Membrane | peptidase | |
| chr12 | *RP11-834C11.3* | 5 | 289 | 13 | | | | | |
| chrX | *PRICKLE3* | 6 | 249 | 13 | PRICKLE3 | prickle planar cell polarity protein | Other | other | |
| chr14 | *PLEKHD1* | 5 | 163 | 13 | PLEKHD1 | pleckstrin homology and coiled-coil domain containing D1 | Other | other | |
| chr14 | *MEG8* | 5 | 161 | 13 | MEG8 | maternally expressed 8, small nucleolar RNA host gene | Other | other | |
| chr10 | *CUBN* | 5 | 115 | 13 | CUBN | cubilin | Plasma Membrane | transmembrane receptor | |
| chr3 | *TFRC* | 4 | 124 | 13 | TFRC | transferrin receptor | Plasma Membrane | transporter | CX-2029, CALAA-01 |
| chr6 | *MAPK13* | 6 | 202 | 13 | MAPK13 | mitogen-activated protein kinase | Cytoplasm | kinase | talmapimod |
| chrX | *ZNF280C* | 5 | 132 | 13 | ZNF280C | zinc finger protein 280C | Nucleus | transcription regulator | |
| chr2 | *REL* | 5 | 121 | 13 | REL | REL proto-oncogene, NF-kB | Nucleus | transcription regulator | |
| chr19 | *NOSIP* | 5 | 160 | 13 | NOSIP | nitric oxide synthase interacting protein | Cytoplasm | other | |

| | | | | | | | | | |
|---|---|---|---|---|---|---|---|---|---|
| chr19 | *AC008984.2* | 5 | 241 | 13 | | | | | |
| chr3 | *SENP5* | 5 | 109 | 13 | SENP5 | SUMO specific peptidase 5 | Nucleus | peptidase | |
| chr1 | *LPAR3* | 6 | 136 | 13 | LPAR3 | lysophosphatidic acid receptor 3 | Plasma Membrane | i-protein coupled recepto | |
| chr11 | *RP11-856F16.2* | 6 | 232 | 13 | | | | | |
| chr19 | *TICAM1* | 6 | 171 | 13 | TICAM1 | toll like receptor adaptor molecule | Nucleus | other | |
| chr2 | *LCLAT1* | 5 | 125 | 13 | LCLAT1 | lysocardiolipin acyltransferase 1 | Cytoplasm | enzyme | |
| chr1 | *ANKRD45* | 6 | 181 | 13 | ANKRD45 | ankyrin repeat domain 45 | Nucleus | transcription regulator | |
| chr5 | *HTR4* | 5 | 134 | 13 | HTR4 | 5-hydroxytryptamine receptor 4 | Plasma Membrane | i-protein coupled recepto | fenfluramine, ergotamine, cisapride, renzapride, prucalopride, PRX-03140, tegaserod |
| chr20 | *RALGAPA2* | 5 | 164 | 13 | RALGAPA2 | Ral GTPase activating protein catalytic alpha subunit 2 | Cytoplasm | other | |
| chr16 | *PRSS36* | 6 | 296 | 13 | PRSS36 | serine protease 36 | Extracellular Space | peptidase | |
| chr6 | *FOXO3* | 5 | 138 | 13 | FOXO3 | forkhead box O3 | Nucleus | transcription regulator | |
| chr10 | *FBXO18* | 5 | 138 | 13 | FBH1 | F-box DNA helicase 1 | Nucleus | enzyme | |
| chr1 | *MPL* | 5 | 180 | 13 | MPL | MPL proto-oncogene, thrombopoietin receptor | Plasma Membrane | transmembrane recepto | avatrombopag, eltrombopag, lusutrombopag, hetrombopag, romiplostim |
| chr1 | *RP11-24P14.1* | 5 | 114 | 13 | | | | | |
| chr1 | *POLR3C* | 6 | 159 | 13 | POLR3C | RNA polymerase III subunit C | Nucleus | enzyme | |
| chr16 | *IL27* | 6 | 268 | 13 | IL27 | interleukin 27 | Extracellular Space | cytokine | |
| chr13 | *WASF3* | 4 | 126 | 13 | WASF3 | WASP family member 3 | Cytoplasm | other | |
| chr20 | *RP4-724E16.2* | 6 | 197 | 13 | | | | | |
| chr4 | *ELOVL6* | 5 | 121 | 13 | ELOVL6 | ELOVL fatty acid elongase 6 | Cytoplasm | enzyme | |
| chr1 | *LPPR5* | 6 | 134 | 13 | | | | | |
| chr13 | *EBPL* | 5 | 121 | 13 | EBPL | EBP like | Cytoplasm | enzyme | |
| chr7 | *PTN* | 6 | 153 | 13 | PTN | pleiotrophin | Extracellular Space | growth factor | |
| chr12 | *TBC1D15* | 5 | 137 | 13 | TBC1D15 | TBC1 domain family member 15 | Cytoplasm | other | |
| chr11 | *RP11-770G2.2* | 5 | 209 | 13 | | | | | |
| chr7 | *SSBP1* | 5 | 160 | 13 | SSBP1 | single stranded DNA binding protein 1 | Cytoplasm | other | |
| chr1 | *CDC42* | 6 | 131 | 13 | CDC42 | cell division cycle 42 | Cytoplasm | enzyme | |
| chr19 | *NUDT19* | 6 | 112 | 13 | NUDT19 | nudix hydrolase 19 | Cytoplasm | enzyme | |
| chr5 | *GRAMD3* | 5 | 120 | 13 | GRAMD2B | GRAM domain containing 2B | Extracellular Space | other | |
| chr15 | *OIP5-AS1* | 4 | 130 | 13 | OIP5-AS1 | OIP5 antisense RNA 1 | Other | other | |
| chr12 | *WSB2* | 5 | 157 | 13 | WSB2 | WD repeat and SOCS box containing 2 | Cytoplasm | other | |
| chr11 | *ACAT1* | 5 | 135 | 13 | ACAT1 | acetyl-CoA acetyltransferase 1 | Cytoplasm | enzyme | |
| chr8 | *PUF60* | 7 | 295 | 13 | PUF60 | poly(U) binding splicing factor 60 | Nucleus | other | |
| chr7 | *AC018720.10* | 6 | 250 | 13 | | | | | |
| chr16 | *RP11-266L9.4* | 5 | 159 | 13 | | | | | |
| chr6 | *LACE1* | 4 | 125 | 13 | AFG1L | AFG1 like ATPase | Cytoplasm | other | |
| chr19 | *MFSD12* | 5 | 224 | 13 | MFSD12 | major facilitator superfamily domain containing 12 | Cytoplasm | other | |
| chr7 | *SMO* | 5 | 164 | 13 | SMO | smoothened, frizzled class receptor | Plasma Membrane | i-protein coupled recepto | gemcitabine/vismodegib, BMS-833923, vismodegib, erlotinib/vismodegib, TAK-441, ZSP1602, NVP-LEQ-506, glasdegib, saridegib, taladegib, |
| chr8 | *RP11-941H19.3* | 4 | 118 | 13 | | | | | |
| chrX | *CHST7* | 6 | 117 | 13 | CHST7 | carbohydrate sulfotransferase 7 | Cytoplasm | enzyme | |
| chr1 | *CFH* | 6 | 138 | 13 | CFH | complement factor H | Extracellular Space | other | |
| chr7 | *NDUFB2* | 6 | 194 | 13 | NDUFB2 | NADH:ubiquinone oxidoreductase subunit B2 | Cytoplasm | enzyme | |
| chr1 | *AUNIP* | 5 | 132 | 13 | AUNIP | aurora kinase A and ninein interacting protein | Cytoplasm | other | |
| chr6 | *NCR2* | 6 | 192 | 13 | NCR2 | natural cytotoxicity triggering receptor 2 | Plasma Membrane | transmembrane recepto | |
| chr7 | *TMEM184A* | 7 | 192 | 13 | TMEM184A | transmembrane protein 184A | Cytoplasm | other | |
| chr9 | *TOR1A* | 6 | 254 | 13 | TOR1A | torsin family 1 member A | Cytoplasm | enzyme | |
| chr1 | *ATAD3C* | 7 | 282 | 13 | ATAD3C | ATPase family AAA domain containing 3C | Cytoplasm | other | |
| chr18 | *RP11-838N2.4* | 5 | 227 | 13 | | | | | |
| chr1 | *RP4-794H19.2* | 7 | 284 | 13 | | | | | |
| chr1 | *ANXA9* | 6 | 225 | 13 | ANXA9 | annexin A9 | Plasma Membrane | transmembrane recepto | |
| chr3 | *GADL1* | 6 | 138 | 13 | GADL1 | glutamate decarboxylase like 1 | Cytoplasm | enzyme | |
| chr10 | *ERCC6* | 5 | 227 | 13 | ERCC6 | ERCC excision repair 6, chromatin remodeling factor | Nucleus | transcription regulator | |
| chr21 | *IFNGR2* | 5 | 144 | 13 | IFNGR2 | interferon gamma receptor 2 | Plasma Membrane | transmembrane recepto | interferon gamma-1b |
| chr9 | *PDCD1LG2* | 6 | 149 | 13 | PDCD1LG2 | programmed cell death 1 ligand 2 | Plasma Membrane | enzyme | CA-170, anti PD-L2 antibody |
| chr7 | *AC023669.1* | 6 | 152 | 13 | | | | | |

| | | | | | | | | | |
|---|---|---|---|---|---|---|---|---|---|
| chr17 | *CHRNB1* | 6 | 196 | 13 | CHRNB1 | cholinergic receptor nicotinic beta 1 subunit | Plasma Membrane | transmembrane receptor | succinylcholine, rapacuronium, doxacurium, vecuronium, pipecuronium, metocurine, mivacurium, D-tubocurarine, nicotine, arecoline, pancuronium, atracurium, isoflurane, cisatracurium, rocuronium, amobarbital, mecamylamine |
| chr8 | *NEIL2* | 6 | 177 | 13 | NEIL2 | nei like DNA glycosylase 2 | Nucleus | enzyme | |
| chr19 | *ZNF177* | 6 | 116 | 13 | ZNF177 | zinc finger protein 177 | Nucleus | transcription regulator | |
| chr12 | *RP11-923I11.4* | 6 | 305 | 13 | | | | | |
| chr9 | *RP11-98L5.2* | 6 | 353 | 13 | | | | | |
| chr7 | *AC073115.6* | 6 | 252 | 13 | | | | | |
| chr15 | *MFGE8* | 5 | 336 | 13 | MFGE8 | milk fat globule-EGF factor 8 | Extracellular Space | other | |
| chr6 | *MAD2L1BP* | 5 | 206 | 13 | MAD2L1BP | MAD2L1 binding protein | Nucleus | other | |
| chrX | *GABRA3* | 5 | 160 | 13 | GABRA3 | gamma-aminobutyric acid type A receptor alpha3 subunit | Plasma Membrane | ion channel | flunitrazepam, sevoflurane, alprazolam, butobarbital, atropine/hyoscyamine/phenobarbital/scopolamine, phenobarbital, diazepam, barbital, acetaminophen/butalbital/caffeine/codeine, hyoscyamine/phenobarbital, desflurane, adinazolam, SEP 174559, zaleplon, mephobarbital, ezogabine, flurazepam, butabarbital, midazolam, temazepam, diphenhydramine/midazolam, [illegible] |
| chr17 | *LGALS9* | 5 | 242 | 13 | LGALS9 | galectin 9 | Plasma Membrane | other | |
| chr2 | *FAM126B* | 4 | 101 | 13 | FAM126B | family with sequence similarity 126 member B | Other | other | |
| chr1 | *CD58* | 6 | 249 | 13 | CD58 | CD58 molecule | Plasma Membrane | transmembrane receptor | |
| chr1 | *EIF2D* | 6 | 176 | 13 | EIF2D | eukaryotic translation initiation factor 2D | Cytoplasm | transporter | |
| chr14 | *RTN1* | 5 | 152 | 13 | RTN1 | reticulon 1 | Cytoplasm | other | |
| chr10 | *VDAC2* | 6 | 289 | 13 | VDAC2 | voltage dependent anion channel | Cytoplasm | ion channel | |
| chr17 | *PCTP* | 6 | 134 | 13 | PCTP | phosphatidylcholine transfer | Cytoplasm | transporter | |
| chr2 | *AC012442.5* | 6 | 160 | 13 | | | | | |
| chr22 | *PLXNB2* | 7 | 372 | 13 | PLXNB2 | plexin B2 | Plasma Membrane | transmembrane receptor | |
| chr20 | *TGM3* | 5 | 151 | 13 | TGM3 | transglutaminase 3 | Cytoplasm | enzyme | |
| chr4 | *TBCK* | 5 | 172 | 13 | TBCK | TBC1 domain containing kinase | Cytoplasm | kinase | |
| chr10 | *AL450307.1* | 6 | 272 | 13 | | | | | |
| chr16 | *CTD-2012K14.6* | 6 | 131 | 13 | | | | | |
| chr19 | *MBOAT7* | 7 | 299 | 13 | MBOAT7 | membrane bound O-acyltransferase domain | Plasma Membrane | enzyme | |
| chr1 | *AKR1A1* | 6 | 182 | 13 | AKR1A1 | aldo-keto reductase family 1 member A1 | Cytoplasm | enzyme | aldose reductase inhibitor |
| chr12 | *HSPB8* | 6 | 145 | 13 | HSPB8 | heat shock protein family B (small) member 8 | Cytoplasm | kinase | |
| chrX | *IKBKG* | 6 | 252 | 13 | IKBKG | inhibitor of nuclear factor kappa B kinase regulatory subunit gamma | Nucleus | kinase | |
| chr8 | *HGSNAT* | 5 | 114 | 13 | HGSNAT | heparan-alpha-glucosaminide N-acetyltransferase | Cytoplasm | enzyme | |
| chr7 | *AC006960.7* | 6 | 210 | 13 | | | | | |
| chr17 | *PDE6G* | 6 | 234 | 13 | PDE6G | phosphodiesterase 6G | Cytoplasm | enzyme | |
| chrX | *TEX28* | 5 | 171 | 13 | TEX28 | testis expressed 28 | Other | other | |
| chr15 | *TMEM87A* | 5 | 132 | 13 | TMEM87A | transmembrane protein 87A | Cytoplasm | other | |
| chr12 | *RP11-148B3.2* | 6 | 159 | 13 | | | | | |
| chr11 | *RARRES3* | 6 | 205 | 13 | PLAAT4 | phospholipase A and acyltransferase 4 | Cytoplasm | enzyme | |
| chr17 | *ENO3* | 5 | 248 | 13 | ENO3 | enolase 3 | Cytoplasm | enzyme | |
| chr2 | *AC067959.1* | 5 | 125 | 13 | | | | | |
| chr5 | *RP11-798K23.4* | 6 | 259 | 13 | | | | | |
| chr3 | *DYNC1LI1* | 5 | 102 | 13 | DYNC1LI1 | dynein cytoplasmic 1 light intermediate chain 1 | Cytoplasm | other | |
| chr3 | *FAM86DP* | 6 | 231 | 13 | FAM86DP | family with sequence similarity 86 member D, pseudogene | Other | other | |
| chr10 | *PPA1* | 5 | 212 | 13 | PPA1 | inorganic pyrophosphatase 1 | Cytoplasm | enzyme | |
| chr5 | *N4BP3* | 6 | 278 | 13 | N4BP3 | NEDD4 binding protein 3 | Cytoplasm | other | |
| chr17 | *KLHL10* | 6 | 178 | 13 | KLHL10 | kelch like family member 10 | Nucleus | other | |

| | | | | | | | | | |
|---|---|---|---|---|---|---|---|---|---|
| chr22 | *POLDIP3* | 6 | 159 | 13 | POLDIP3 | DNA polymerase delta interacting protein 3 | Nucleus | other | |
| chr5 | *SNCAIP* | 6 | 113 | 13 | SNCAIP | synuclein alpha interacting | Cytoplasm | transcription regulator | |
| chr16 | *RP11-254F19.3* | 6 | 282 | 13 | | | | | |
| chr16 | *TRIM72* | 5 | 207 | 13 | TRIM72 | tripartite motif containing 72 | Cytoplasm | enzyme | |
| chr15 | *RP11-168G16.2* | 6 | 202 | 13 | | | | | |
| chr8 | *RP11-122C21.1* | 4 | 183 | 13 | | | | | |
| chr20 | *STMN3* | 6 | 497 | 13 | STMN3 | stathmin 3 | Nucleus | other | |
| chr8 | *FAM160B2* | 6 | 254 | 13 | FAM160B2 | family with sequence similarity 160 member B2 | Other | other | |
| chr5 | *RP11-133F8.2* | 5 | 134 | 13 | | | | | |
| chr17 | *AC004231.2* | 6 | 118 | 13 | | | | | |
| chr19 | *CCL25* | 6 | 171 | 13 | CCL25 | C-C motif chemokine ligand 25 | Extracellular Space | cytokine | |
| chr5 | *CTC-281M20.1* | 5 | 121 | 13 | | | | | |
| chr1 | *TSNAX-DISC1* | 6 | 139 | 13 | TSNAX-DISC1 | TSNAX-DISC1 readthrough (NMD candidate) | Other | other | |
| chr2 | *AC093838.4* | 6 | 240 | 13 | | | | | |
| chr3 | *EIF4E3* | 6 | 118 | 13 | EIF4E3 | eukaryotic translation initiation factor 4E family member 3 | Cytoplasm | translation regulator | |
| chr11 | *SYT13* | 7 | 222 | 13 | SYT13 | synaptotagmin 13 | Plasma Membrane | transporter | |
| chr22 | *PRODH* | 5 | 277 | 13 | C102724788/PRC | proline dehydrogenase 1 | Cytoplasm | enzyme | |
| chrX | *MAMLD1* | 6 | 219 | 13 | MAMLD1 | mastermind like domain | Nucleus | other | |
| chr1 | *DHX9* | 5 | 141 | 13 | DHX9 | DExH-box helicase 9 | Nucleus | enzyme | |
| chr16 | *NUTF2* | 6 | 159 | 13 | NUTF2 | nuclear transport factor 2 | Nucleus | transporter | |
| chr16 | *SULT1A1* | 5 | 233 | 13 | SULT1A1 | sulfotransferase family 1A | Cytoplasm | enzyme | |
| chr19 | *QTRT1* | 6 | 175 | 13 | QTRT1 | queuine tRNA-ribosyltransferase catalytic subunit 1 | Cytoplasm | enzyme | |
| chr8 | *TBC1D31* | 5 | 110 | 13 | TBC1D31 | TBC1 domain family member 31 | Cytoplasm | other | |
| chr16 | *CTD-2033A16.3* | 6 | 155 | 13 | | | | | |
| chr11 | *ZDHHC24* | 6 | 416 | 13 | ZDHHC24 | zinc finger DHHC-type containing | Plasma Membrane | other | |
| chr7 | *H2AFV* | 5 | 189 | 13 | H2AZ2 | H2A.Z variant histone 2 | Nucleus | other | |
| chr8 | *PRSS51* | 6 | 180 | 13 | PRSS51 | serine protease 51 | Other | other | |
| chr19 | *ZNF317* | 5 | 171 | 13 | ZNF317 | zinc finger protein 317 | Nucleus | transcription regulator | |
| chr7 | *AC007566.11* | 5 | 197 | 13 | | | | | |
| chr17 | *SAMD14* | 6 | 259 | 13 | SAMD14 | sterile alpha motif domain containing 14 | Other | other | |
| chr12 | *MDM1* | 5 | 113 | 13 | MDM1 | Mdm1 nuclear protein | Cytoplasm | other | |
| chr1 | *KLHDC8A* | 6 | 319 | 13 | KLHDC8A | kelch domain containing 8A | Other | other | |
| chr1 | *FCAMR* | 6 | 236 | 13 | FCAMR | Fc fragment of IgA and IgM | Plasma Membrane | transmembrane receptor | |
| chr16 | *E2F4* | 6 | 270 | 13 | E2F4 | E2F transcription factor 4 | Nucleus | transcription regulator | |
| chr1 | *CLK2* | 6 | 270 | 13 | CLK2 | CDC like kinase 2 | Nucleus | kinase | |
| chr19 | *SPPL2B* | 6 | 236 | 13 | SPPL2B | signal peptide peptidase like 2B | Plasma Membrane | peptidase | |
| chr16 | *HEATR3* | 5 | 148 | 13 | HEATR3 | HEAT repeat containing 3 | Other | other | |
| chr19 | *ATP4A* | 6 | 306 | 13 | ATP4A | ATPase H+/K+ transporting subunit alpha | Plasma Membrane | transporter | ilaprazole, esomeprazole magnesium, aspirin/omeprazole, tenatoprazole, clarithromycin/omeprazole, lansoprazole, AR-H047108, diclofenac/omeprazole, dexlansoprazole, AGN 201904, magnesium hydroxide/omeprazole/sodium bicarbonate, omeprazole/sodium bicarbonate, pantoprazole, esomeprazole/naproxen, rabeprazole |
| chr9 | *FSD1L* | 4 | 134 | 13 | FSD1L | fibronectin type III and SPRY domain containing 1 like | Other | other | |
| chr3 | *PIK3R4* | 5 | 174 | 13 | PIK3R4 | phosphoinositide-3-kinase regulatory subunit 4 | Cytoplasm | kinase | |
| chr15 | *EMC7* | 4 | 150 | 13 | EMC7 | ER membrane protein complex subunit 7 | Cytoplasm | other | |
| chr6 | *SSR1* | 5 | 239 | 13 | SSR1 | signal sequence receptor subunit | Cytoplasm | other | |
| chr11 | *NDUFC2-KCTD14* | 6 | 198 | 13 | NDUFC2-KCTD14 | NDUFC2-KCTD14 readthrough | Cytoplasm | other | |
| chr17 | *C17orf103* | 6 | 251 | 13 | NATD1 | N-acetyltransferase domain containing 1 | Other | other | |
| chr6 | *SERPINB9* | 6 | 150 | 13 | SERPINB9 | serpin family B member 9 | Cytoplasm | other | |
| chr11 | *SLC6A5* | 6 | 181 | 13 | SLC6A5 | solute carrier family 6 member 5 | Plasma Membrane | transporter | VVZ-149 |
| chr2 | *STRN* | 4 | 95 | 13 | STRN | striatin | Cytoplasm | other | |
| chr17 | *GSDMA* | 6 | 167 | 13 | GSDMA | gasdermin A | Cytoplasm | other | |
| chr5 | *RGS7BP* | 6 | 122 | 13 | RGS7BP | regulator of G protein signaling 7 binding protein | Cytoplasm | other | |
| chr13 | *RP11-321C24.1* | 5 | 110 | 13 | | | | | |
| chr1 | *RP11-277A4.4* | 5 | 165 | 13 | | | | | |
| chr8 | *C8orf31* | 6 | 196 | 13 | C8orf31 | chromosome 8 open reading frame 31 | Other | other | |
| chr8 | *RP11-661A12.5* | 5 | 241 | 13 | | | | | |
| chr1 | *RAB25* | 5 | 304 | 13 | RAB25 | RAB25, member RAS oncogene family | Cytoplasm | enzyme | |
| chrX | *DMRTC1* | 5 | 150 | 13 | MRTC1/DMRTC1 | DMRT like family C1 | Other | other | |
| chr19 | *ZNF701* | 5 | 154 | 13 | ZNF701 | zinc finger protein 701 | Nucleus | transcription regulator | |
| chr9 | *FCN1* | 6 | 319 | 13 | FCN1 | ficolin 1 | Extracellular Space | other | |

| | | | | | | | | |
|---|---|---|---|---|---|---|---|---|
| chr10 | RP11-298J20.4 | 5 | 131 | 13 | | | | |
| chr3 | RP11-290K4.2 | 6 | 173 | 13 | | | | |
| chr1 | SLC25A3P1 | 6 | 143 | 13 | SLC25A3P1 | solute carrier family 25 member 3 pseudogene 1 | Other | other |
| chr16 | CTD-2555A7.1 | 6 | 267 | 13 | | | | |
| chr1 | COG2 | 6 | 168 | 13 | COG2 | component of oligomeric golgi complex 2 | Cytoplasm | other |
| chr15 | RP11-208K4.1 | 6 | 106 | 13 | | | | |
| chr5 | CTB-43E15.4 | 6 | 245 | 13 | | | | |
| chr11 | RP11-788M5.3 | 6 | 227 | 13 | | | | |
| chr3 | GAP43 | 5 | 174 | 13 | GAP43 | growth associated protein 43 | Plasma Membrane | other |
| chr10 | NCOA4 | 5 | 186 | 13 | NCOA4 | nuclear receptor coactivator 4 | Nucleus | transcription regulator |
| chr3 | GOLIM4 | 5 | 267 | 13 | GOLIM4 | golgi integral membrane protein 4 | Cytoplasm | other |
| chr1 | RP11-261C10.3 | 6 | 241 | 13 | | | | |
| chr1 | TFAP2E | 6 | 190 | 13 | TFAP2E | transcription factor AP-2 epsilon | Nucleus | transcription regulator |
| chr16 | SLC7A6 | 6 | 158 | 13 | SLC7A6 | solute carrier family 7 member 6 | Plasma Membrane | transporter |
| chr1 | ARL8A | 5 | 295 | 13 | ARL8A | ADP ribosylation factor like GTPase 8A | Cytoplasm | enzyme |
| chr19 | SHISA7 | 6 | 281 | 13 | SHISA7 | shisa family member 7 | Plasma Membrane | other |
| chr12 | DDX23 | 6 | 224 | 13 | DDX23 | DEAD-box helicase 23 | Nucleus | enzyme |
| chr15 | C15orf32 | 6 | 235 | 13 | C15orf32 | chromosome 15 open reading frame 32 | Other | other |
| chr2 | SLC30A6 | 4 | 99 | 13 | SLC30A6 | solute carrier family 30 member 6 | Cytoplasm | transporter |
| chr18 | LINC00907 | 5 | 127 | 13 | LINC00907 | long intergenic non-protein coding RNA 907 | Other | other |
| chr1 | RP11-739N20.2 | 5 | 168 | 13 | | | | |
| chr7 | TNRC18P2 | 7 | 340 | 13 | TNRC18P2 | trinucleotide repeat containing 18 pseudogene 2 | Other | other |
| chr19 | ARRDC5 | 5 | 157 | 13 | ARRDC5 | arrestin domain containing 5 | Other | other |
| chr17 | SMCR8 | 5 | 248 | 13 | SMCR8 | SMCR8-C9orf72 complex subunit | Nucleus | other |
| chr22 | EWSR1 | 5 | 128 | 13 | EWSR1 | EWS RNA binding protein 1 | Nucleus | other |
| chr2 | METTL8 | 5 | 128 | 13 | METTL8 | methyltransferase like 8 | Other | enzyme |
| chr11 | UBE4A | 5 | 148 | 13 | UBE4A | ubiquitination factor E4A | Cytoplasm | enzyme |
| chr17 | HGS | 7 | 365 | 13 | HGS | hepatocyte growth factor-regulated tyrosine kinase | Cytoplasm | other |
| chr17 | ZFP3 | 5 | 140 | 13 | ZFP3 | ZFP3 zinc finger protein | Nucleus | transcription regulator |
| chr22 | CECR1 | 5 | 171 | 13 | ADA2 | adenosine deaminase 2 | Extracellular Space | enzyme |
| chr15 | RASL12 | 6 | 198 | 13 | RASL12 | RAS like family 12 | Other | enzyme |
| chr22 | RTDR1 | 6 | 209 | 13 | RSPH14 | radial spoke head 14 homolog | Other | other |
| chr12 | CLSTN3 | 5 | 202 | 13 | CLSTN3 | calsyntenin 3 | Plasma Membrane | other |
| chr11 | NRIP3 | 4 | 168 | 13 | NRIP3 | nuclear receptor interacting | Other | other |
| chr1 | CD34 | 6 | 226 | 13 | CD34 | CD34 molecule | Plasma Membrane | other |
| chr1 | HHLA3 | 5 | 138 | 13 | HHLA3 | HERV-H LTR-associating 3 | Other | other |
| chr3 | CRELD1 | 6 | 317 | 13 | CRELD1 | cysteine rich with EGF like | Other | other |
| chr22 | UPK3A | 5 | 203 | 13 | UPK3A | uroplakin 3A | Cytoplasm | other |
| chr1 | MIER1 | 4 | 112 | 13 | MIER1 | MIER1 transcriptional regulator | Nucleus | transcription regulator |
| chr10 | ACSL5 | 6 | 173 | 13 | ACSL5 | acyl-CoA synthetase long chain family member 5 | Cytoplasm | enzyme |
| chr17 | TMEM101 | 6 | 234 | 13 | TMEM101 | transmembrane protein 101 | Extracellular Space | other |
| chr20 | RP5-963E22.4 | 7 | 517 | 13 | | | | |
| chr10 | PTPN20CP | 5 | 138 | 13 | PTPN20CP | protein tyrosine phosphatase non-receptor type 20C, | Other | other |
| chr11 | CHST1 | 6 | 297 | 13 | CHST1 | carbohydrate sulfotransferase 1 | Cytoplasm | enzyme |
| chr5 | CMYA5 | 5 | 148 | 13 | CMYA5 | cardiomyopathy associated 5 | Other | other |
| chr10 | NUTM2HP | 6 | 394 | 13 | NUTM2HP | NUT family member 2H, | Other | other |
| chr12 | ETNK1 | 5 | 118 | 13 | ETNK1 | ethanolamine kinase 1 | Cytoplasm | kinase |
| chr9 | RECK | 5 | 114 | 13 | RECK | reversion inducing cysteine rich protein with kazal motifs | Plasma Membrane | other |
| chr5 | TTC33 | 6 | 424 | 13 | TTC33 | tetratricopeptide repeat domain | Other | other |
| chr15 | CTD-2008A1.2 | 6 | 153 | 13 | | | | |
| chr19 | ZNF708 | 5 | 133 | 13 | ZNF708 | zinc finger protein 708 | Nucleus | other |
| chr7 | MACC1 | 5 | 149 | 12 | MACC1 | MET transcriptional regulator | Nucleus | transcription regulator |
| chr1 | SNX7 | 5 | 138 | 12 | SNX7 | sorting nexin 7 | Other | transporter |
| chr14 | LINC00523 | 5 | 230 | 12 | LINC00523 | long intergenic non-protein coding RNA 523 | Other | other |
| chr20 | HSPA12B | 6 | 266 | 12 | HSPA12B | heat shock protein family A (Hsp70) member 12B | Other | other |
| chr7 | RASA4CP | 6 | 352 | 12 | RASA4CP | RAS p21 protein activator 4C, pseudogene | Other | other |
| chr5 | CTD-2194D22.4 | 6 | 610 | 12 | CTD-2194D22.4 | uncharacterized LOC101929081 | Other | other |
| chr20 | C20orf27 | 5 | 207 | 12 | C20orf27 | chromosome 20 open reading frame 27 | Other | other |
| chr20 | BPIFA2 | 6 | 203 | 12 | BPIFA2 | BPI fold containing family A member 2 | Extracellular Space | other |
| chr11 | CTD-2589M5.4 | 6 | 256 | 12 | | | | |
| chr15 | GOLGA6L10 | 5 | 177 | 12 | GA6L9 (includes o | golgin A6 family like 9 | Other | other |
| chr19 | PODNL1 | 6 | 152 | 12 | PODNL1 | podocan like 1 | Extracellular Space | other |
| chr10 | IL15RA | 5 | 182 | 12 | IL15RA | interleukin 15 receptor subunit | Plasma Membrane | transmembrane receptor |
| chr7 | GTPBP10 | 5 | 180 | 12 | GTPBP10 | GTP binding protein 10 | Cytoplasm | other |
| chr1 | DUSP27 | 5 | 225 | 12 | DUSP27 | dual specificity phosphatase 27, atypical | Other | phosphatase |
| chr5 | LINC00992 | 6 | 150 | 12 | LINC00992 | long intergenic non-protein coding RNA 992 | Other | other |
| chr2 | DNPEP | 5 | 165 | 12 | DNPEP | aspartyl aminopeptidase | Cytoplasm | peptidase |
| chr9 | NELFB | 7 | 305 | 12 | NELFB | negative elongation factor complex member B | Nucleus | other |
| chr17 | RP11-104H15.7 | 6 | 219 | 12 | | | | |
| chr19 | MED29 | 6 | 202 | 12 | MED29 | mediator complex subunit 29 | Nucleus | other |

| | | | | | | | | | |
|---|---|---|---|---|---|---|---|---|---|
| chr1 | *NUCKS1* | 5 | 195 | 12 | NUCKS1 | nuclear casein kinase and cyclin dependent kinase substrate 1 | Nucleus | kinase | |
| chr14 | *CTD-2134A5.4* | 5 | 220 | 12 | | | | | |
| chr1 | *REN* | 6 | 310 | 12 | REN | renin | Extracellular Space | peptidase | aliskiren, aliskiren/amlodipine/hydrochlorothiazide, aliskiren/amlodipine, remikiren, aliskiren/valsartan, aliskiren/hydrochlorothiazide |
| chr1 | *PRKAA2* | 5 | 112 | 12 | PRKAA2 | protein kinase AMP-activated catalytic subunit alpha 2 | Cytoplasm | kinase | |
| chr3 | *ATP2C1* | 6 | 147 | 12 | ATP2C1 | ATPase secretory pathway Ca2+ transporting 1 | Cytoplasm | transporter | Ca2+, sevoflurane, isoflurane, desflurane, enflurane |
| chr17 | *ALDH3A2* | 6 | 171 | 12 | ALDH3A2 | aldehyde dehydrogenase 3 family member A2 | Cytoplasm | enzyme | |
| chr10 | *MMP21* | 5 | 154 | 12 | MMP21 | matrix metallopeptidase 21 | Extracellular Space | peptidase | marimastat |
| chr15 | *RP11-1129I3.1* | 6 | 102 | 12 | | | | | |
| chr1 | *RP11-280O1.2* | 6 | 146 | 12 | | | | | |
| chr17 | *AC015849.16* | 5 | 123 | 12 | | | | | |
| chr2 | *AC084193.1* | 6 | 107 | 12 | | | | | |
| chr16 | *IL32* | 5 | 258 | 12 | IL32 | interleukin 32 | Extracellular Space | cytokine | |
| chr19 | *R3HDM4* | 6 | 281 | 12 | R3HDM4 | R3H domain containing 4 | Other | other | |
| chr10 | *PANK1* | 5 | 135 | 12 | PANK1 | pantothenate kinase 1 | Cytoplasm | kinase | |
| chr13 | *F10* | 6 | 333 | 12 | F10 | coagulation factor X | Extracellular Space | peptidase | aspirin/enoxaparin, heparin, rivaroxaban, enoxaparin/warfarin, RPR 120844, emicizumab, edoxaban, idraparinux, aspirin/rivaroxaban/ticlopidine, dalteparin, nematode anticoagulant protein c2, DPC 423, aspirin/rivaroxaban, betrixaban, tifacogin, aspirin/clopidogrel/rivaroxaban, F9, activated recombinant human factor VII, apixaban, fondaparinux/warfarin, fondaparinux |
| chr8 | *RP11-63E5.6* | 5 | 170 | 12 | | | | | |
| chr2 | *ALMS1P* | 6 | 150 | 12 | ALMS1P1 | ALMS1 pseudogene 1 | Other | other | |
| chr12 | *FGFR1OP2* | 5 | 183 | 12 | FGFR1OP2 | FGFR1 oncogene partner 2 | Cytoplasm | other | |
| chr3 | *RP11-10O22.1* | 5 | 125 | 12 | | | | | |
| chr17 | *PDK2* | 5 | 216 | 12 | PDK2 | pyruvate dehydrogenase kinase 2 | Cytoplasm | kinase | |
| chr1 | *RP11-31F15.2* | 6 | 193 | 12 | | | | | |
| chr1 | *RHBDL2* | 6 | 155 | 12 | RHBDL2 | rhomboid like 2 | Plasma Membrane | peptidase | |
| chr2 | *MAP2* | 4 | 120 | 12 | MAP2 | microtubule associated protein 2 | Plasma Membrane | other | estramustine |
| chr7 | *AC004901.1* | 6 | 174 | 12 | | | | | |
| chr7 | *AC002454.1* | 6 | 196 | 12 | | | | | |
| chr12 | *C1S* | 5 | 171 | 12 | C1S | complement C1s | Extracellular Space | peptidase | sutimlimab, |
| chr12 | *NOP2* | 6 | 156 | 12 | NOP2 | NOP2 nucleolar protein | Nucleus | other | |
| chr10 | *KCNK18* | 5 | 152 | 12 | KCNK18 | potassium two pore domain channel subfamily K member 18 | Plasma Membrane | ion channel | |
| chr3 | *PROSP* | 5 | 164 | 12 | PROS2P | protein S (beta) pseudogene | Other | other | |
| chr19 | *RUVBL2* | 6 | 221 | 12 | RUVBL2 | RuvB like AAA ATPase 2 | Nucleus | transcription regulator | |
| chr19 | *DUXA* | 5 | 122 | 12 | DUXA | double homeobox A | Other | other | |
| chrX | *OPN1MW* | 4 | 186 | 12 | 1MW (includes ot | opsin 1, medium wave sensitive | Plasma Membrane | i-protein coupled recepto | |
| chr15 | *RP11-566K19.3* | 5 | 208 | 12 | | | | | |
| chr2 | *FOSL2* | 6 | 193 | 12 | FOSL2 | FOS like 2, AP-1 transcription factor subunit | Nucleus | transcription regulator | |
| chr17 | *CCT6B* | 5 | 145 | 12 | CCT6B | chaperonin containing TCP1 subunit 6B | Cytoplasm | other | |
| chr19 | *CCDC114* | 6 | 204 | 12 | CCDC114 | coiled-coil domain containing 114 | Extracellular Space | other | |
| chr15 | *FOXB1* | 6 | 181 | 12 | FOXB1 | forkhead box B1 | Nucleus | transcription regulator | |
| chr19 | *DYRK1B* | 6 | 231 | 12 | DYRK1B | dual specificity tyrosine phosphorylation regulated kinase | Nucleus | kinase | |
| chr17 | *ALDH3A1* | 6 | 223 | 12 | ALDH3A1 | aldehyde dehydrogenase 3 family member A1 | Cytoplasm | enzyme | |
| chrX | *CDK16* | 5 | 232 | 12 | CDK16 | cyclin dependent kinase 16 | Cytoplasm | kinase | |
| chr19 | *ZNF790* | 5 | 158 | 12 | ZNF790 | zinc finger protein 790 | Other | other | |
| chr2 | *ESPNL* | 6 | 296 | 12 | ESPNL | espin like | Extracellular Space | other | |
| chrX | *USP11* | 5 | 201 | 12 | USP11 | ubiquitin specific peptidase 11 | Nucleus | peptidase | |
| chr14 | *MYH7* | 5 | 175 | 12 | MYH7 | myosin heavy chain 7 | Cytoplasm | enzyme | |
| chr15 | *GATM* | 5 | 120 | 12 | GATM | glycine amidinotransferase | Cytoplasm | enzyme | |
| chr15 | *GOLGA6A* | 5 | 249 | 12 | GA6A (includes ot | golgin A6 family member A | Other | other | |
| chr6 | *RP11-459O1.2* | 4 | 130 | 12 | | | | | |
| chr9 | *AQP7P1* | 6 | 333 | 12 | AQP7P1 | aquaporin 7 pseudogene 1 | Other | other | |
| chr16 | *RPGRIP1L* | 6 | 108 | 12 | RPGRIP1L | RPGRIP1 like | Cytoplasm | other | |
| chr14 | *CEP170B* | 7 | 392 | 12 | CEP170B | centrosomal protein 170B | Other | other | |
| chrX | *CCDC22* | 5 | 213 | 12 | CCDC22 | coiled-coil domain containing 22 | Cytoplasm | other | |
| chr4 | *SNX25* | 4 | 120 | 12 | SNX25 | sorting nexin 25 | Cytoplasm | other | |

| | | | | | | | | | |
|---|---|---|---|---|---|---|---|---|---|
| chr11 | *MED17* | 5 | 246 | 12 | MED17 | mediator complex subunit 17 | Nucleus | transcription regulator | |
| chr2 | *LIMS3L* | 5 | 170 | 12 | LIMS3/LIMS4 | LIM zinc finger domain containing | Other | other | |
| chr12 | *DDX55* | 6 | 139 | 12 | DDX55 | DEAD-box helicase 55 | Nucleus | enzyme | |
| chr12 | *MAP1LC3B2* | 5 | 141 | 12 | MAP1LC3B2 | microtubule associated protein 1 light chain 3 beta 2 | Other | other | |
| chr1 | *PDPN* | 5 | 108 | 12 | PDPN | podoplanin | Plasma Membrane | other | |
| chr17 | *ARHGEF15* | 5 | 242 | 12 | ARHGEF15 | Rho guanine nucleotide exchange factor 15 | Cytoplasm | other | |
| chr7 | *PLOD3* | 6 | 290 | 12 | PLOD3 | procollagen-lysine,2-oxoglutarate 5-dioxygenase 3 | Cytoplasm | enzyme | |
| chr1 | *SLAMF1* | 5 | 140 | 12 | SLAMF1 | signaling lymphocytic activation molecule family member 1 | Plasma Membrane | transmembrane receptor | |
| chr10 | *ZNF518A* | 5 | 149 | 12 | ZNF518A | zinc finger protein 518A | Other | other | |
| chr15 | *UBE2Q2* | 5 | 131 | 12 | UBE2Q2 | ubiquitin conjugating enzyme E2 | Cytoplasm | enzyme | |
| chr2 | *RETSAT* | 6 | 185 | 12 | RETSAT | retinol saturase | Cytoplasm | enzyme | |
| chr2 | *NT5DC4* | 5 | 257 | 12 | NT5DC4 | 5'-nucleotidase domain | Other | other | |
| chr15 | *OIP5* | 5 | 160 | 12 | OIP5 | Opa interacting protein 5 | Nucleus | other | |
| chr17 | *KPNB1* | 5 | 123 | 12 | KPNB1 | karyopherin subunit beta 1 | Nucleus | transporter | |
| chr9 | *C9orf9* | 5 | 266 | 12 | SPACA9 | sperm acrosome associated 9 | Cytoplasm | other | |
| chr6 | *VTA1* | 5 | 131 | 12 | VTA1 | vesicle trafficking 1 | Cytoplasm | other | |
| chr3 | *RNF168* | 4 | 120 | 12 | RNF168 | ring finger protein 168 | Nucleus | enzyme | |
| chr8 | *AC100802.3* | 6 | 107 | 12 | | | | | |
| chr2 | *PLA2R1* | 6 | 121 | 12 | PLA2R1 | phospholipase A2 receptor 1 | Plasma Membrane | transmembrane receptor | |
| chr11 | *PGA3* | 5 | 472 | 12 | GA5 (includes othe | pepsinogen A3 | Extracellular Space | peptidase | sucralfate |
| chr12 | *SMARCD1* | 6 | 242 | 12 | SMARCD1 | SWI/SNF related, matrix associated, actin dependent regulator of chromatin, subfamily | Nucleus | transcription regulator | |
| chr8 | *GPR20* | 6 | 392 | 12 | GPR20 | G protein-coupled receptor 20 | Plasma Membrane | G-protein coupled recepto | |
| chr12 | *ITGB7* | 6 | 267 | 12 | ITGB7 | integrin subunit beta 7 | Plasma Membrane | transmembrane receptor | vedolizumab |
| chr3 | *TM4SF19* | 4 | 139 | 12 | TM4SF19 | transmembrane 4 L six family member 19 | Other | other | |
| chr2 | *PDCL3* | 6 | 204 | 12 | PDCL3 | phosducin like 3 | Cytoplasm | other | |
| chrX | *LAS1L* | 5 | 217 | 12 | LAS1L | LAS1 like ribosome biogenesis | Nucleus | other | |
| chr2 | *PROC* | 6 | 315 | 12 | PROC | protein C, inactivator of coagulation factors Va and VIIIa | Extracellular Space | peptidase | |
| chr1 | *TTF2* | 5 | 149 | 12 | TTF2 | transcription termination factor 2 | Cytoplasm | transcription regulator | |
| chr5 | *ACOT12* | 6 | 147 | 12 | ACOT12 | acyl-CoA thioesterase 12 | Cytoplasm | enzyme | |
| chr19 | *IGLON5* | 6 | 206 | 12 | IGLON5 | IgLON family member 5 | Other | other | |
| chr12 | *IKBIP* | 5 | 127 | 12 | IKBIP | IKBKB interacting protein | Cytoplasm | other | |
| chr1 | *HYDIN2* | 4 | 134 | 12 | HYDIN2 | HYDIN axonemal central pair apparatus protein 2 (pseudogene) | Other | other | |
| chr10 | *WDR11* | 6 | 136 | 12 | WDR11 | WD repeat domain 11 | Cytoplasm | other | |
| chr16 | *RP11-324D17.1* | 5 | 155 | 12 | | | | | |
| chr8 | *TMEM71* | 5 | 124 | 12 | TMEM71 | transmembrane protein 71 | Cytoplasm | other | |
| chr19 | *ONECUT3* | 5 | 249 | 12 | ONECUT3 | one cut homeobox 3 | Other | other | |
| chr1 | *WARS2* | 6 | 178 | 12 | WARS2 | tryptophanyl tRNA synthetase 2, mitochondrial | Cytoplasm | enzyme | |
| chr11 | *C2CD2L* | 5 | 233 | 12 | C2CD2L | C2CD2 like | Cytoplasm | transporter | |
| chr1 | *PSMD4* | 5 | 165 | 12 | PSMD4 | proteasome 26S subunit, non-ATPase 4 | Cytoplasm | other | |
| chr12 | *RMST* | 6 | 105 | 12 | RMST | rhabdomyosarcoma 2 associated transcript | Other | other | |
| chr3 | *NME9* | 6 | 219 | 12 | NME9 | NME/NM23 family member 9 | Cytoplasm | enzyme | |
| chr3 | *CIDEC* | 5 | 212 | 12 | CIDEC | cell death inducing DFFA like effector c | Cytoplasm | other | |
| chr1 | *AKR7L* | 5 | 196 | 12 | AKR7L | aldo-keto reductase family 7 like (gene/pseudogene) | Cytoplasm | enzyme | |
| chr1 | *LRRC47* | 6 | 263 | 12 | LRRC47 | leucine rich repeat containing 47 | Other | other | |
| chr11 | *ELMOD1* | 6 | 107 | 12 | ELMOD1 | ELMO domain containing 1 | Other | other | |
| chr9 | *RP11-389K14.3* | 4 | 100 | 12 | | | | | |
| chr18 | *FAM69C* | 5 | 154 | 12 | DIPK1C | divergent protein kinase domain | Other | other | |
| chr2 | *C2orf43* | 5 | 170 | 12 | LDAH | lipid droplet associated hydrolase | Cytoplasm | enzyme | |
| chr6 | *LINC00271* | 4 | 137 | 12 | LINC00271 | long intergenic non-protein coding RNA 271 | Other | other | |
| chr19 | *MED16* | 6 | 219 | 12 | MED16 | mediator complex subunit 16 | Nucleus | transcription regulator | |
| chr21 | *SLC19A1* | 6 | 296 | 12 | SLC19A1 | solute carrier family 19 member 1 | Plasma Membrane | transporter | |
| chr5 | *MARVELD2* | 5 | 113 | 12 | MARVELD2 | MARVEL domain containing 2 | Plasma Membrane | other | |
| chr2 | *WNT10A* | 6 | 229 | 12 | WNT10A | Wnt family member 10A | Extracellular Space | other | |
| chr6 | *PGC* | 5 | 222 | 12 | PGC | progastricsin | Extracellular Space | peptidase | |
| chr21 | *RRP1* | 6 | 234 | 12 | RRP1 | ribosomal RNA processing 1 | Nucleus | other | |
| chr19 | *CD177* | 6 | 219 | 12 | CD177 | CD177 molecule | Cytoplasm | other | |
| chr4 | *MANBA* | 5 | 111 | 12 | MANBA | mannosidase beta | Cytoplasm | enzyme | |
| chr11 | *FAR1* | 4 | 102 | 12 | FAR1 | fatty acyl-CoA reductase 1 | Cytoplasm | enzyme | |
| chr21 | *KCNJ15* | 4 | 100 | 12 | KCNJ15 | potassium inwardly rectifying channel subfamily J member 15 | Plasma Membrane | ion channel | |
| chr20 | *MAPRE1* | 5 | 194 | 12 | MAPRE1 | microtubule associated protein RP/EB family member 1 | Cytoplasm | other | |
| chr4 | *TACC3* | 5 | 252 | 12 | TACC3 | transforming acidic coiled-coil containing protein 3 | Nucleus | other | |
| chr1 | *RSBN1* | 5 | 144 | 12 | RSBN1 | round spermatid basic protein 1 | Nucleus | other | |
| chr19 | *RASIP1* | 6 | 203 | 12 | RASIP1 | Ras interacting protein 1 | Cytoplasm | other | |
| chr17 | *RP11-13K12.1* | 5 | 197 | 12 | | | | | |
| chr11 | *SART1* | 6 | 430 | 12 | SART1 | spliceosome associated factor 1, recruiter of U4/U6.U5 tri-snRNP | Nucleus | other | |
| chr4 | *SPON2* | 6 | 323 | 12 | SPON2 | spondin 2 | Extracellular Space | other | |
| chr12 | *MUC19* | 6 | 173 | 12 | MUC19 | mucin 19, oligomeric | Cytoplasm | other | |
| chr19 | *ZNF766* | 5 | 145 | 12 | ZNF766 | zinc finger protein 766 | Nucleus | transcription regulator | |
| chr20 | *TGM6* | 5 | 174 | 12 | TGM6 | transglutaminase 6 | Cytoplasm | enzyme | |
| chr17 | *DOC2B* | 6 | 281 | 12 | DOC2B | double C2 domain beta | Cytoplasm | transporter | |

| | | | | | | | | | |
|---|---|---|---|---|---|---|---|---|---|
| chr2 | *AC073987.1* | 5 | 126 | 12 | | | | | |
| chr5 | *FAT2* | 6 | 187 | 12 | FAT2 | FAT atypical cadherin 2 | Plasma Membrane | other | |
| chr10 | *AGAP10* | 4 | 125 | 12 | AGAP10P | ArfGAP with GTPase domain, ankyrin repeat and PH domain 10, pseudogene | Other | other | |
| chr22 | *RNF215* | 5 | 183 | 12 | RNF215 | ring finger protein 215 | Other | other | |
| chr1 | *RXRG* | 6 | 189 | 12 | RXRG | retinoid X receptor gamma | Nucleus | d-dependent nuclear rec | 6-mercaptopurine/methotrexate/tretinoin, daunorubicin/tretinoin, alitretinoin, cytarabine/daunorubicin/tretinoin, acitretin, arsenic trioxide/daunorubicin/tretinoin, idarubicin/mitoxantrone/tretinoin, etretinate, idarubicin/tretinoin, tretinoin, arsenic trioxide/gemtuzumab ozogamicin/tretinoin, cytarabine/daunorubicin/gemtuzumab ozogamicin/tretinoin, adapalene, arsenic trioxide/idarubicin/tretinoin, bexarotene, [illegible] |
| chr20 | *E2F1* | 6 | 341 | 12 | E2F1 | E2F transcription factor 1 | Nucleus | transcription regulator | |
| chr19 | *RASAL3* | 6 | 202 | 12 | RASAL3 | RAS protein activator like 3 | Cytoplasm | other | |
| chr15 | *GOLGA8M* | 5 | 156 | 12 | GA8K (includes ot | golgin A8 family member K | Other | other | |
| chr17 | *UTP18* | 4 | 129 | 12 | UTP18 | UTP18 small subunit processome component | Nucleus | other | |
| chr2 | *WASH2P* | 6 | 443 | 12 | WASH2P | WASP family homolog 2, pseudogene | Other | other | |
| chr10 | *ZNF37BP* | 5 | 223 | 12 | ZNF37BP | zinc finger protein 37B, | Other | other | |
| chrX | *GNL3L* | 7 | 176 | 12 | GNL3L | G protein nucleolar 3 like | Nucleus | other | |
| chr9 | *RP11-213G2.3* | 4 | 110 | 12 | | | | | |
| chr5 | *CTD-2247C11.1* | 7 | 383 | 12 | | | | | |
| chr20 | *RTFDC1* | 5 | 216 | 12 | RTF2 | replication termination factor 2 | Nucleus | other | |
| chr7 | *EPHA1* | 5 | 238 | 12 | EPHA1 | EPH receptor A1 | Plasma Membrane | kinase | |
| chr12 | *RAB3IP* | 5 | 128 | 12 | RAB3IP | RAB3A interacting protein | Cytoplasm | other | |
| chr19 | *STAP2* | 6 | 180 | 12 | STAP2 | signal transducing adaptor family member 2 | Cytoplasm | other | |
| chr11 | *TAF1D* | 6 | 387 | 12 | TAF1D | TATA-box binding protein associated factor, RNA | Nucleus | other | |
| chr6 | *CYP39A1* | 6 | 112 | 12 | CYP39A1 | cytochrome P450 family 39 subfamily A member 1 | Cytoplasm | enzyme | |
| chr20 | *ZBP1* | 6 | 410 | 12 | ZBP1 | Z-DNA binding protein 1 | Cytoplasm | other | |
| chr8 | *PTDSS1* | 5 | 115 | 12 | PTDSS1 | phosphatidylserine synthase 1 | Cytoplasm | enzyme | |
| chr17 | *BLMH* | 5 | 129 | 12 | BLMH | bleomycin hydrolase | Cytoplasm | peptidase | |
| chr11 | *SCN3B* | 5 | 132 | 12 | SCN3B | sodium voltage-gated channel beta subunit 3 | Plasma Membrane | ion channel | riluzole, zonisamide |
| chr1 | *TMEM53* | 6 | 219 | 12 | TMEM53 | transmembrane protein 53 | Nucleus | other | |
| chr1 | *TIPRL* | 5 | 141 | 12 | TIPRL | TOR signaling pathway regulator | Cytoplasm | other | |
| chr1 | *HECTD3* | 6 | 227 | 12 | HECTD3 | HECT domain E3 ubiquitin protein ligase 3 | Cytoplasm | enzyme | |
| chr2 | *FAM49A* | 6 | 150 | 12 | FAM49A | family with sequence similarity 49 member A | Other | other | |
| chr1 | *SSR2* | 5 | 209 | 12 | SSR2 | signal sequence receptor subunit | Cytoplasm | other | |
| chr12 | *GPD1* | 5 | 387 | 12 | GPD1 | glycerol-3-phosphate dehydrogenase 1 | Cytoplasm | enzyme | |
| chr8 | *ZNF7* | 6 | 197 | 12 | ZNF7 | zinc finger protein 7 | Nucleus | transcription regulator | |
| chr1 | *SDE2* | 5 | 137 | 12 | SDE2 | SDE2 telomere maintenance homolog | Nucleus | other | |
| chr17 | *AC002117.1* | 5 | 188 | 12 | | | | | |
| chr10 | *R3HCC1L* | 5 | 154 | 12 | R3HCC1L | R3H domain and coiled-coil containing 1 like | Other | other | |
| chr17 | *VAT1* | 5 | 199 | 12 | VAT1 | vesicle amine transport 1 | Plasma Membrane | transporter | |
| chr6 | *CASP8AP2* | 4 | 111 | 12 | CASP8AP2 | caspase 8 associated protein 2 | Nucleus | transcription regulator | |
| chr10 | *NSUN6* | 4 | 104 | 12 | NSUN6 | NOP2/Sun RNA | Cytoplasm | enzyme | |
| chr15 | *VPS18* | 6 | 235 | 12 | VPS18 | VPS18 core subunit of CORVET and HOPS complexes | Cytoplasm | transporter | |
| chrX | *KCND1* | 5 | 289 | 12 | KCND1 | potassium voltage-gated channel subfamily D member 1 | Plasma Membrane | ion channel | dalfampridine |
| chr9 | *TEK* | 4 | 97 | 12 | TEK | TEK receptor tyrosine kinase | Plasma Membrane | kinase | everolimus/vandetanib, CEP-11981, cabozantinib/nivolumab, vandetanib, glesatinib, bosutinib/hydroxyurea/ponatinib, altiratinib, regorafenib, cabozantinib, ponatinib, cabozantinib/erlotinib |

| | | | | | | | | | |
|---|---|---|---|---|---|---|---|---|---|
| chr6 | *RNF144B* | 5 | 138 | 12 | RNF144B | ring finger protein 144B | Cytoplasm | enzyme | |
| chr22 | *APOBEC3D* | 5 | 206 | 12 | APOBEC3D | apolipoprotein B mRNA editing enzyme catalytic subunit 3D | Cytoplasm | other | |
| chr17 | *TNK1* | 6 | 197 | 12 | TNK1 | tyrosine kinase non receptor 1 | Cytoplasm | kinase | |
| chr11 | *ACCS* | 5 | 187 | 12 | ACCS | 1-aminocyclopropane-1-carboxylate synthase homolog | Other | enzyme | |
| chr15 | *UBAP1L* | 5 | 169 | 12 | UBAP1L | ubiquitin associated protein 1 like | Other | other | |
| chr17 | *TBC1D3P4* | 5 | 274 | 12 | TBC1D3P4 | TBC1 domain family member 3 pseudogene 4 | Other | other | |
| chr3 | *LRRC34* | 5 | 225 | 12 | LRRC34 | leucine rich repeat containing 34 | Other | other | |
| chr1 | *WI2-3658N16.1* | 6 | 173 | 12 | | | | | |
| chr1 | *GPR37L1* | 6 | 370 | 12 | GPR37L1 | G protein-coupled receptor 37 | Plasma Membrane | G-protein coupled recepto | |
| chr22 | *AC002472.13* | 5 | 177 | 12 | | | | | |
| chr1 | *BTBD19* | 5 | 219 | 12 | BTBD19 | BTB domain containing 19 | Other | other | |
| chr3 | *SCN11A* | 6 | 136 | 12 | SCN11A | sodium voltage-gated channel alpha subunit 11 | Plasma Membrane | ion channel | riluzole, zonisamide |
| chr19 | *AC011513.4* | 5 | 134 | 12 | | | | | |
| chr9 | *CNTNAP3* | 4 | 116 | 12 | CNTNAP3 | contactin associated protein like | Extracellular Space | other | |
| chr9 | *RP11-132E11.2* | 4 | 97 | 12 | | | | | |
| chr1 | *GCLM* | 4 | 114 | 12 | GCLM | glutamate-cysteine ligase modifier subunit | Cytoplasm | enzyme | |
| chr19 | *ZNF43* | 6 | 122 | 12 | ZNF43 | zinc finger protein 43 | Nucleus | transcription regulator | |
| chr14 | *SLC39A9* | 4 | 110 | 12 | SLC39A9 | solute carrier family 39 member 9 | Other | transporter | |
| chr1 | *RP11-296O14.3* | 5 | 142 | 12 | | | | | |
| chr20 | *SIGLEC1* | 5 | 254 | 12 | SIGLEC1 | sialic acid binding Ig like lectin 1 | Plasma Membrane | other | |
| chr5 | *B4GALT7* | 6 | 239 | 12 | B4GALT7 | beta-1,4-galactosyltransferase 7 | Cytoplasm | enzyme | |
| chr3 | *C3orf55* | 6 | 156 | 12 | SLC66A1L | solute carrier family 66 member 1 | Other | other | |
| chr5 | *FGFR4* | 6 | 296 | 12 | FGFR4 | fibroblast growth factor receptor 4 | Plasma Membrane | kinase | roblitinib, ASP5878, PRN1371, lenvatinib, everolimus/lenvatinib, U3-1784, erdafitinib, BLU-554, LY2874455, INCB062079, lenvatinib/pembrolizu |
| chr20 | *ACOT8* | 5 | 256 | 12 | ACOT8 | acyl-CoA thioesterase 8 | Cytoplasm | enzyme | |
| chr19 | *ATP5SL* | 5 | 217 | 12 | DMAC2 | distal membrane arm assembly complex 2 | Cytoplasm | other | |
| chr19 | *TEX101* | 5 | 163 | 12 | TEX101 | testis expressed 101 | Plasma Membrane | other | |
| chr1 | *RP11-14N7.2* | 6 | 734 | 12 | | | | | |
| chr17 | *IGFBP4* | 6 | 200 | 12 | IGFBP4 | insulin like growth factor binding protein 4 | Extracellular Space | other | |
| chr11 | *MS4A6E* | 5 | 164 | 12 | MS4A6E | membrane spanning 4-domains | Other | other | |
| chr9 | *PAPPA* | 5 | 169 | 12 | PAPPA | pappalysin 1 | Extracellular Space | peptidase | |
| chr19 | *AC003006.7* | 4 | 114 | 12 | | | | | |
| chr19 | *VN2R19P* | 5 | 183 | 12 | VN2R19P | vomeronasal 2 receptor 19, pseudogene | Other | other | |
| chr10 | *SLC35G1* | 6 | 111 | 12 | SLC35G1 | solute carrier family 35 member | Plasma Membrane | other | |
| chr11 | *TSKU* | 6 | 312 | 12 | TSKU | tsukushi, small leucine rich proteoglycan | Extracellular Space | other | |
| chr4 | *ATP8A1* | 5 | 104 | 12 | ATP8A1 | ATPase phospholipid transporting 8A1 | Cytoplasm | transporter | |
| chr16 | *RP11-863P13.4* | 6 | 253 | 12 | | | | | |
| chr7 | *PEX1* | 4 | 124 | 12 | PEX1 | peroxisomal biogenesis factor 1 | Cytoplasm | enzyme | |
| chr17 | *KRT16P1* | 5 | 290 | 12 | KRT16P1 | keratin 16 pseudogene 1 | Other | other | |
| chr7 | *CPA2* | 6 | 162 | 12 | CPA2 | carboxypeptidase A2 | Extracellular Space | peptidase | |
| chr11 | *CCDC88B* | 6 | 305 | 12 | CCDC88B | coiled-coil domain containing 88B | Nucleus | enzyme | |
| chr4 | *FNIP2* | 4 | 109 | 12 | FNIP2 | folliculin interacting protein 2 | Cytoplasm | other | |
| chr3 | *DCLK3* | 5 | 158 | 12 | DCLK3 | doublecortin like kinase 3 | Cytoplasm | kinase | |
| chr3 | *HEMK1* | 5 | 246 | 12 | HEMK1 | HemK methyltransferase family member 1 | Nucleus | enzyme | |
| chr21 | *DNMT3L* | 5 | 242 | 12 | DNMT3L | DNA methyltransferase 3 like | Nucleus | transcription regulator | |
| chr4 | *C4orf45* | 4 | 102 | 12 | C4orf45 | chromosome 4 open reading frame 45 | Other | other | |
| chr19 | *AC007204.2* | 6 | 126 | 12 | | | | | |
| chr8 | *DNAJC5B* | 4 | 110 | 12 | DNAJC5B | DnaJ heat shock protein family (Hsp40) member C5 beta | Other | transporter | |
| chr5 | *ANKDD1B* | 5 | 167 | 12 | ANKDD1B | ankyrin repeat and death domain containing 1B | Other | other | |
| chr13 | *NEK5* | 5 | 108 | 12 | NEK5 | NIMA related kinase 5 | Other | kinase | |
| chrX | *FAM122C* | 5 | 108 | 12 | FAM122C | family with sequence similarity | Other | other | |
| chr2 | *IL1RL2* | 6 | 210 | 12 | IL1RL2 | interleukin 1 receptor like 2 | Plasma Membrane | transmembrane receptor | |
| chr1 | *UHMK1* | 4 | 136 | 12 | UHMK1 | U2AF homology motif kinase 1 | Nucleus | kinase | |
| chr15 | *GOLGA6L6* | 4 | 181 | 12 | GA6L1 (includes o | golgin A6 family like 1 | Other | other | |
| chr5 | *RP11-116A1.1* | 5 | 122 | 12 | | | | | |
| chr3 | *TLR9* | 6 | 292 | 12 | TLR9 | toll like receptor 9 | Plasma Membrane | transmembrane recepto | AST-008, MGN1703, ISS-1018, CPG 10104, agatolimod, DV281, IMO-2125, GNKG168, DUK-CPG-001, hydroxychloroquine, |
| chr2 | *PDE6D* | 4 | 109 | 12 | PDE6D | phosphodiesterase 6D | Cytoplasm | enzyme | |
| chr21 | *RCAN1* | 5 | 142 | 12 | RCAN1 | regulator of calcineurin 1 | Nucleus | transcription regulator | |
| chr1 | *ZNF684* | 4 | 119 | 12 | ZNF684 | zinc finger protein 684 | Nucleus | transcription regulator | |
| chr19 | *ZNF837* | 5 | 224 | 12 | ZNF837 | zinc finger protein 837 | Other | other | |
| chr3 | *PARP9* | 5 | 153 | 12 | PARP9 | poly(ADP-ribose) polymerase family member 9 | Nucleus | enzyme | |

| | | | | | | | | | |
|---|---|---|---|---|---|---|---|---|---|
| chr16 | *C16orf91* | 6 | 212 | 12 | C16orf91 | chromosome 16 open reading frame 91 | Other | other | |
| chr12 | *TMPO* | 5 | 120 | 12 | TMPO | thymopoietin | Nucleus | other | |
| chr16 | *MAPK3* | 5 | 220 | 12 | MAPK3 | mitogen-activated protein kinase 3 | Cytoplasm | kinase | ASTX029, KO-947, ulixertinib, LY3214996, ASN007 |
| chr5 | *GDNF-AS1* | 6 | 385 | 12 | GDNF-AS1 | GDNF antisense RNA 1 | Other | other | |
| chr1 | *TRIM11* | 6 | 297 | 12 | TRIM11 | tripartite motif containing 11 | Cytoplasm | enzyme | |
| chr14 | *KIAA0586* | 4 | 97 | 12 | KIAA0586 | KIAA0586 | Cytoplasm | other | |
| chrX | *DMRTC1B* | 5 | 143 | 12 | MRTC1/DMRTC1 | DMRT like family C1 | Other | other | |
| chr12 | *LRRC23* | 4 | 122 | 12 | LRRC23 | leucine rich repeat containing 23 | Plasma Membrane | other | |
| chr4 | *RP11-15B17.1* | 5 | 142 | 12 | | | | | |
| chr8 | *SPAG11A* | 5 | 261 | 12 | SPAG11A | sperm associated antigen 11A | Other | other | |
| chr3 | *IL17RE* | 6 | 276 | 12 | IL17RE | interleukin 17 receptor E | Plasma Membrane | other | |
| chr16 | *GPR97* | 5 | 240 | 12 | ADGRG3 | adhesion G protein-coupled receptor G3 | Plasma Membrane | ;-protein coupled recepto | |
| chrX | *CXorf23* | 5 | 150 | 12 | BCLAF3 | BCLAF1 and THRAP3 family member 3 | Other | other | |
| chr19 | *GAPDHS* | 6 | 215 | 12 | GAPDHS | glyceraldehyde-3-phosphate dehydrogenase, spermatogenic | Cytoplasm | enzyme | |
| chr12 | *PARPBP* | 5 | 115 | 12 | PARPBP | PARP1 binding protein | Nucleus | other | |
| chr15 | *HERC2P10* | 5 | 170 | 12 | HERC2P10 | hect domain and RLD 2 pseudogene 10 | Other | other | |
| chr2 | *ALLC* | 5 | 138 | 12 | ALLC | allantoicase | Other | enzyme | |
| chrX | *KLHL13* | 4 | 118 | 12 | KLHL13 | kelch like family member 13 | Cytoplasm | other | |
| chr3 | *ZBTB11* | 5 | 141 | 12 | ZBTB11 | zinc finger and BTB domain containing 11 | Nucleus | transcription regulator | |
| chr12 | *RP11-13G14.4* | 6 | 112 | 12 | | | | | |
| chr3 | *LINC00969* | 5 | 196 | 12 | MIR570HG | MIR570 host gene | Other | other | |
| chr22 | *UPB1* | 5 | 188 | 12 | UPB1 | beta-ureidopropionase 1 | Cytoplasm | enzyme | |
| chr17 | *KDM6B* | 6 | 276 | 12 | KDM6B | lysine demethylase 6B | Extracellular Space | enzyme | |
| chr1 | *ZMPSTE24* | 4 | 107 | 12 | ZMPSTE24 | zinc metallopeptidase STE24 | Nucleus | peptidase | |
| chr7 | *POT1* | 5 | 115 | 12 | POT1 | protection of telomeres 1 | Nucleus | other | |
| chr3 | *FBXO40* | 6 | 144 | 12 | FBXO40 | F-box protein 40 | Cytoplasm | other | |
| chr5 | *MYOZ3* | 5 | 143 | 12 | MYOZ3 | myozenin 3 | Cytoplasm | other | |
| chr19 | *ZNF611* | 6 | 148 | 12 | ZNF611 | zinc finger protein 611 | Other | other | |
| chr1 | *TP73-AS1* | 6 | 212 | 12 | TP73-AS1 | TP73 antisense RNA 1 | Other | other | |
| chr16 | *RNF40* | 5 | 255 | 12 | RNF40 | ring finger protein 40 | Cytoplasm | enzyme | |
| chr17 | *AC022816.2* | 5 | 101 | 12 | | | | | |
| chr15 | *CTD-2054N24.2* | 6 | 201 | 12 | | | | | |
| chr1 | *PRELP* | 6 | 218 | 12 | PRELP | proline and arginine rich end leucine rich repeat protein | Extracellular Space | other | |
| chr7 | *TWIST1* | 6 | 210 | 12 | TWIST1 | twist family bHLH transcription factor 1 | Nucleus | transcription regulator | |
| chr11 | *MS4A10* | 5 | 266 | 12 | MS4A10 | membrane spanning 4-domains | Other | other | |
| chr1 | *FNDC5* | 5 | 270 | 12 | FNDC5 | fibronectin type III domain containing 5 | Extracellular Space | other | |
| chr11 | *ALDH3B2* | 6 | 246 | 12 | ALDH3B2 | aldehyde dehydrogenase 3 family member B2 | Cytoplasm | enzyme | |
| chr20 | *RBPJL* | 6 | 249 | 12 | RBPJL | recombination signal binding protein for immunoglobulin kappa J region like | Nucleus | transcription regulator | |
| chr21 | *RIPK4* | 5 | 246 | 12 | RIPK4 | receptor interacting serine/threonine kinase 4 | Nucleus | kinase | |
| chr2 | *FSHR* | 6 | 140 | 12 | FSHR | follicle stimulating hormone receptor | Plasma Membrane | ;-protein coupled recepto | FSH, menotropin, suramin |
| chr7 | *IQUB* | 4 | 135 | 12 | IQUB | IQ motif and ubiquitin domain containing | Cytoplasm | other | |
| chr4 | *TSPAN5* | 4 | 148 | 12 | TSPAN5 | tetraspanin 5 | Plasma Membrane | other | |
| chr2 | *GPR39* | 5 | 129 | 12 | GPR39 | G protein-coupled receptor 39 | Plasma Membrane | ;-protein coupled recepto | |
| chr9 | *FAM219A* | 6 | 127 | 12 | FAM219A | family with sequence similarity 219 member A | Other | other | |
| chr1 | *PCP4L1* | 6 | 134 | 12 | PCP4L1 | Purkinje cell protein 4 like 1 | Other | other | |
| chr1 | *GOLT1A* | 6 | 265 | 12 | GOLT1A | golgi transport 1A | Cytoplasm | other | |
| chr1 | *RP4-580N22.2* | 5 | 152 | 12 | | | | | |
| chr2 | *AC027612.3* | 5 | 294 | 12 | | | | | |
| chr2 | *AC007880.1* | 6 | 161 | 12 | | | | | |
| chr17 | *OTOP3* | 6 | 238 | 12 | OTOP3 | otopetrin 3 | Other | other | |
| chr4 | *BEND4* | 5 | 148 | 12 | BEND4 | BEN domain containing 4 | Other | other | |
| chr16 | *KARS* | 6 | 121 | 12 | KARS1 | lysyl-tRNA synthetase 1 | Cytoplasm | enzyme | |
| chr7 | *VWDE* | 4 | 150 | 12 | VWDE | von Willebrand factor D and EGF domains | Other | other | |
| chr3 | *CCDC174* | 5 | 155 | 12 | CCDC174 | coiled-coil domain containing 174 | Nucleus | other | |
| chrX | *MTMR1* | 5 | 137 | 12 | MTMR1 | myotubularin related protein 1 | Cytoplasm | phosphatase | |
| chr19 | *DPF1* | 6 | 232 | 12 | DPF1 | double PHD fingers 1 | Nucleus | other | |
| chr10 | *ZEB1* | 4 | 135 | 12 | ZEB1 | zinc finger E-box binding | Nucleus | transcription regulator | |
| chr2 | *MARCH4;* | 5 | 141 | 12 | | | | | |
| chr16 | *CMC2* | 5 | 108 | 12 | CMC2 | C-X9-C motif containing 2 | Cytoplasm | other | |
| chr9 | *PTBP3* | 4 | 94 | 12 | PTBP3 | polypyrimidine tract binding | Nucleus | other | |
| chr15 | *UBL7* | 5 | 164 | 12 | UBL7 | ubiquitin like 7 | Other | other | |
| chr19 | *TLE6* | 5 | 216 | 12 | TLE6 | TLE family member 6, subcortical maternal complex member | Nucleus | other | |
| chr10 | *ACADSB* | 6 | 145 | 12 | ACADSB | acyl-CoA dehydrogenase short/branched chain | Cytoplasm | enzyme | |
| chr7 | *HBP1* | 4 | 115 | 12 | HBP1 | HMG-box transcription factor 1 | Nucleus | transcription regulator | |
| chr1 | *NBPF3* | 6 | 135 | 12 | NBPF3 | NBPF member 3 | Other | other | |
| chr1 | *TSPAN2* | 5 | 167 | 12 | TSPAN2 | tetraspanin 2 | Extracellular Space | other | |
| chr19 | *PSMC4* | 5 | 193 | 12 | PSMC4 | proteasome 26S subunit, ATPase | Nucleus | peptidase | |
| chr7 | *AC018647.3* | 5 | 162 | 12 | | | | | |

| | | | | | | | | | |
|---|---|---|---|---|---|---|---|---|---|
| chr7 | *PRKAR2B* | 5 | 106 | 12 | PRKAR2B | protein kinase cAMP-dependent type II regulatory subunit beta | Cytoplasm | kinase | |
| chr2 | *CD8A* | 5 | 165 | 12 | CD8A | CD8a molecule | Plasma Membrane | other | |
| chr1 | *POMGNT1* | 6 | 200 | 12 | POMGNT1 | protein O-linked mannose N-acetylglucosaminyltransferase 1 (beta 1,2-) | Cytoplasm | enzyme | |
| chr19 | *PPP2R1A* | 6 | 144 | 12 | PPP2R1A | protein phosphatase 2 scaffold subunit Aalpha | Cytoplasm | phosphatase | |
| chr19 | *CYP4F2* | 5 | 182 | 12 | CYP4F2 | cytochrome P450 family 4 subfamily F member 2 | Cytoplasm | enzyme | |
| chr17 | *CTD-2033D24.2* | 5 | 161 | 12 | | | | | |
| chr5 | *ZNF354C* | 5 | 157 | 12 | ZNF354C | zinc finger protein 354C | Nucleus | transcription regulator | |
| chr4 | *HERC3* | 4 | 120 | 12 | HERC3 | HECT and RLD domain containing E3 ubiquitin protein | Cytoplasm | enzyme | |
| chr1 | *SELENBP1* | 5 | 272 | 12 | SELENBP1 | selenium binding protein 1 | Cytoplasm | other | |
| chr16 | *RP11-429P3.5* | 5 | 101 | 12 | | | | | |
| chr9 | *RP11-276H19.1* | 5 | 122 | 12 | | | | | |
| chr5 | *CENPK* | 5 | 110 | 12 | CENPK | centromere protein K | Nucleus | other | |
| chr4 | *SEC24B-AS1* | 4 | 127 | 12 | SEC24B-AS1 | SEC24B antisense RNA 1 | Other | other | |
| chr3 | *UBA7* | 5 | 216 | 12 | UBA7 | ubiquitin like modifier activating enzyme 7 | Cytoplasm | enzyme | |
| chr11 | *RP11-713P17.3* | 6 | 276 | 12 | | | | | |
| chr1 | *RP5-1109J22.1* | 6 | 174 | 12 | | | | | |
| chr12 | *HRK* | 6 | 172 | 12 | HRK | harakiri, BCL2 interacting protein | Cytoplasm | other | |
| chr12 | *IRAK3* | 4 | 116 | 12 | IRAK3 | interleukin 1 receptor associated kinase 3 | Cytoplasm | kinase | |
| chr6 | *LEMD2* | 5 | 235 | 12 | LEMD2 | LEM domain containing 2 | Nucleus | other | |
| chr2 | *NLRC4* | 4 | 113 | 12 | NLRC4 | NLR family CARD domain containing 4 | Cytoplasm | other | |
| chr16 | *TMEM8A* | 6 | 142 | 12 | TMEM8A | transmembrane protein 8A | Plasma Membrane | other | |
| chr3 | *HSPBAP1* | 6 | 156 | 12 | HSPBAP1 | HSPB1 associated protein 1 | Other | other | |
| chr2 | *MXD1* | 6 | 139 | 12 | MXD1 | MAX dimerization protein 1 | Nucleus | transcription regulator | |
| chr9 | *SLC2A8* | 6 | 296 | 12 | SLC2A8 | solute carrier family 2 member 8 | Plasma Membrane | transporter | |
| chr12 | *RP11-439H13.2* | 5 | 111 | 12 | | | | | |
| chr1 | *RP11-177F15.1* | 5 | 167 | 12 | | | | | |
| chr2 | *SMARCAL1* | 4 | 130 | 12 | SMARCAL1 | SWI/SNF related, matrix associated, actin dependent regulator of chromatin, subfamily | Nucleus | enzyme | |
| chr1 | *LY9* | 6 | 188 | 12 | LY9 | lymphocyte antigen 9 | Plasma Membrane | other | |
| chr3 | *B4GALT4-AS1* | 6 | 151 | 12 | B4GALT4-AS1 | B4GALT4 antisense RNA 1 | Other | other | |
| chr2 | *AC105760.2* | 4 | 126 | 12 | | | | | |
| chr8 | *ARMC1* | 4 | 115 | 12 | ARMC1 | armadillo repeat containing 1 | Cytoplasm | other | |
| chr13 | *PDS5B* | 4 | 96 | 12 | PDS5B | PDS5 cohesin associated factor | Nucleus | other | |
| chr7 | *GPNMB* | 5 | 175 | 12 | GPNMB | glycoprotein nmb | Plasma Membrane | enzyme | glembatumumab, glembatumumab vedotin |
| chr2 | *RMND5A* | 5 | 114 | 12 | RMND5A | required for meiotic nuclear division 5 homolog A | Nucleus | enzyme | |
| chr8 | *LINC00051* | 5 | 284 | 12 | LINC00051 | long intergenic non-protein coding RNA 51 | Other | other | |
| chr20 | *PLTP* | 5 | 284 | 12 | PLTP | phospholipid transfer protein | Extracellular Space | enzyme | |
| chr19 | *LINC00085* | 6 | 189 | 12 | SPACA6 | sperm acrosome associated 6 | Other | other | |
| chr17 | *EPX* | 5 | 212 | 12 | EPX | eosinophil peroxidase | Cytoplasm | enzyme | |
| chr1 | *ACBD3* | 5 | 156 | 12 | ACBD3 | acyl-CoA binding domain containing 3 | Cytoplasm | other | |
| chr20 | *SPATA2* | 6 | 280 | 12 | SPATA2 | spermatogenesis associated 2 | Extracellular Space | other | |
| chr3 | *PARL* | 4 | 93 | 12 | PARL | presenilin associated rhomboid | Cytoplasm | peptidase | |
| chr17 | *FDXR* | 6 | 260 | 12 | FDXR | ferredoxin reductase | Cytoplasm | enzyme | |
| chr19 | *HPN* | 5 | 238 | 12 | HPN | hepsin | Plasma Membrane | peptidase | |
| chr15 | *RP11-707P17.1* | 5 | 174 | 12 | | | | | |
| chr15 | *GANC* | 5 | 122 | 12 | GANC | glucosidase alpha, neutral C | Extracellular Space | enzyme | miglitol |
| chr13 | *CAB39L* | 5 | 111 | 12 | CAB39L | calcium binding protein 39 like | Cytoplasm | kinase | |
| chr14 | *RP11-596D21.1* | 5 | 128 | 12 | | | | | |
| chr2 | *PKDCC* | 5 | 264 | 12 | PKDCC | protein kinase domain containing, cytoplasmic | Cytoplasm | kinase | |
| chr1 | *RP11-511I2.2* | 5 | 140 | 12 | | | | | |
| chr5 | *CTD-2308B18.3* | 5 | 124 | 12 | | | | | |
| chr11 | *SLC15A3* | 6 | 331 | 12 | SLC15A3 | solute carrier family 15 member 3 | Cytoplasm | transporter | |
| chr17 | *LRRC37A4P* | 4 | 199 | 12 | LRRC37A4P | leucine rich repeat containing 37 member A4, pseudogene | Other | other | |
| chr16 | *MVD* | 6 | 271 | 12 | MVD | mevalonate diphosphate decarboxylase | Cytoplasm | enzyme | |
| chr16 | *RP11-95P2.1* | 5 | 182 | 12 | | | | | |
| chr17 | *WBP2* | 5 | 209 | 12 | WBP2 | WW domain binding protein 2 | Cytoplasm | transcription regulator | |
| chr9 | *AUH* | 4 | 110 | 12 | AUH | AU RNA binding methylglutaconyl-CoA hydratase | Cytoplasm | enzyme | |
| chr6 | *SOD2* | 4 | 128 | 12 | SOD2 | superoxide dismutase 2 | Cytoplasm | enzyme | |
| chr2 | *CD8BP* | 5 | 202 | 12 | CD8B2 | CD8b2 molecule | Plasma Membrane | other | |
| chr1 | *RP11-446E24.4* | 5 | 213 | 12 | | | | | |
| chr11 | *CHEK1* | 5 | 131 | 12 | CHEK1 | checkpoint kinase 1 | Nucleus | kinase | UCN-01, prexasertib, CCT245737, GDC-0575, PF 00477736, MK-8776, CHK1 inhibitor, XL844, GDC-0425, cdc25C phosphatase (211- |
| chr8 | *POLR3D* | 5 | 167 | 12 | POLR3D | RNA polymerase III subunit D | Nucleus | transcription regulator | |
| chr1 | *CTH* | 4 | 133 | 12 | CTH | cystathionine gamma-lyase | Cytoplasm | enzyme | |
| chr9 | *SPATA31C2* | 5 | 210 | 12 | SPATA31C2 | SPATA31 subfamily C member 2 | Other | other | |

| | | | | | | | | | |
|---|---|---|---|---|---|---|---|---|---|
| chr16 | *PRR25* | 6 | 306 | 12 | PRR25 | proline rich 25 | Other | other | |
| chr5 | *ITGA1* | 5 | 96 | 12 | ITGA1 | integrin subunit alpha 1 | Plasma Membrane | other | |
| chr7 | *LRCH4* | 6 | 240 | 12 | LRCH4 | leucine rich repeats and calponin homology domain containing 4 | Cytoplasm | transcription regulator | |
| chr19 | *TPRX2P* | 4 | 146 | 12 | TPRX2P | tetrapeptide repeat homeobox 2, pseudogene | Other | other | |
| chr5 | *LINC01024* | 5 | 113 | 12 | MALINC1 | mitosis associated long intergenic non-coding RNA 1 | Other | other | |
| chr15 | *hsa-mir-7162* | 5 | 172 | 12 | | | | | |
| chr6 | *FRMD1* | 6 | 267 | 12 | FRMD1 | FERM domain containing 1 | Other | other | |
| chrX | *DGKK* | 6 | 107 | 12 | DGKK | diacylglycerol kinase kappa | Cytoplasm | kinase | |
| chr12 | *PPP1CC* | 4 | 135 | 12 | PPP1CC | protein phosphatase 1 catalytic subunit gamma | Nucleus | phosphatase | |
| chr3 | *PTPRG-AS1* | 6 | 127 | 12 | PTPRG-AS1 | PTPRG antisense RNA 1 | Other | other | |
| chr12 | *PLEKHG7* | 5 | 113 | 12 | PLEKHG7 | pleckstrin homology and RhoGEF domain containing G7 | Other | other | |
| chr12 | *PHC1* | 4 | 153 | 12 | PHC1 | polyhomeotic homolog 1 | Nucleus | transcription regulator | |
| chr7 | *FAM115A* | 5 | 180 | 12 | TCAF1 | TRPM8 channel associated factor | Plasma Membrane | other | |
| chr19 | *KLC3* | 6 | 232 | 12 | KLC3 | kinesin light chain 3 | Cytoplasm | other | |
| chr5 | *TRIM36* | 5 | 153 | 12 | TRIM36 | tripartite motif containing 36 | Cytoplasm | enzyme | |
| chr11 | *JRKL* | 5 | 240 | 12 | JRKL | JRK like | Nucleus | other | |
| chr5 | *NNT-AS1* | 6 | 314 | 12 | NNT-AS1 | NNT antisense RNA 1 | Other | other | |
| chr12 | *DGKA* | 6 | 351 | 12 | DGKA | diacylglycerol kinase alpha | Cytoplasm | kinase | |
| chr15 | *RP11-69H14.6* | 6 | 342 | 12 | | | | | |
| chr2 | *GALNT13* | 4 | 136 | 12 | GALNT13 | polypeptide N-acetylgalactosaminyltransferase | Cytoplasm | enzyme | |
| chr10 | *FAM35BP* | 5 | 204 | 12 | SHLD2P1 | shieldin complex subunit 2 pseudogene 1 | Other | other | |
| chr3 | *WDR49* | 6 | 149 | 12 | WDR49 | WD repeat domain 49 | Other | other | |
| chr9 | *INSL6* | 6 | 162 | 12 | INSL6 | insulin like 6 | Extracellular Space | other | |
| chr19 | *CYP4F11* | 6 | 199 | 12 | CYP4F11 | cytochrome P450 family 4 subfamily F member 11 | Cytoplasm | enzyme | |
| chr15 | *MESP2* | 5 | 236 | 12 | MESP2 | mesoderm posterior bHLH transcription factor 2 | Nucleus | transcription regulator | |
| chr20 | *RP5-977B1.7* | 5 | 194 | 12 | | | | | |
| chr21 | *LINC00111* | 5 | 191 | 12 | LINC00111 | long intergenic non-protein coding RNA 111 | Other | other | |
| chr3 | *ATP13A3* | 4 | 86 | 12 | ATP13A3 | ATPase 13A3 | Extracellular Space | transporter | |
| chr3 | *SLC35G2* | 4 | 166 | 12 | SLC35G2 | solute carrier family 35 member | Cytoplasm | other | |
| chr2 | *FMNL2* | 5 | 95 | 12 | FMNL2 | formin like 2 | Cytoplasm | other | |
| chr19 | *MATK* | 6 | 216 | 12 | MATK | megakaryocyte-associated tyrosine kinase | Cytoplasm | kinase | |
| chr15 | *RP11-276M12.1* | 5 | 168 | 12 | | | | | |
| chr1 | *HPCAL4* | 6 | 189 | 12 | HPCAL4 | hippocalcin like 4 | Other | transporter | |
| chr17 | *RP11-304F15.4* | 5 | 229 | 12 | | | | | |
| chr3 | *VIPR1* | 6 | 143 | 12 | VIPR1 | vasoactive intestinal peptide receptor 1 | Plasma Membrane | -protein coupled recepto | |
| chr15 | *RP11-326A19.4* | 6 | 183 | 12 | | | | | |
| chr3 | *TMIE* | 5 | 247 | 12 | TMIE | transmembrane inner ear | Other | other | |
| chr18 | *NDC80* | 5 | 151 | 12 | NDC80 | NDC80 kinetochore complex component | Nucleus | other | |
| chr17 | *RP11-143K11.5* | 5 | 139 | 12 | | | | | |
| chr8 | *CYHR1* | 6 | 277 | 12 | CYHR1 | cysteine and histidine rich 1 | Nucleus | other | |
| chrX | *ZC4H2* | 6 | 122 | 12 | ZC4H2 | zinc finger C4H2-type containing | Nucleus | other | |
| chr2 | *NEURL3* | 6 | 261 | 12 | NEURL3 | neuralized E3 ubiquitin protein ligase 3 | Other | enzyme | |
| chr3 | *RABL3* | 5 | 192 | 12 | RABL3 | RAB, member of RAS oncogene family like 3 | Other | enzyme | |
| chr2 | *ITGAV* | 5 | 93 | 12 | ITGAV | integrin subunit alpha V | Plasma Membrane | transmembrane recepto | cilengitide, fluciclatide F 18, abciximab, 68Ga-NOTA-BBN-RGD, abituzumab, intetumumab, |
| chr8 | *TRIM35* | 5 | 166 | 12 | TRIM35 | tripartite motif containing 35 | Cytoplasm | other | |
| chr4 | *CCNG2* | 5 | 88 | 12 | CCNG2 | cyclin G2 | Nucleus | other | |
| chr17 | *ZSWIM7* | 5 | 113 | 12 | ZSWIM7 | zinc finger SWIM-type containing | Other | other | |
| chr4 | *RP11-93L9.1* | 5 | 105 | 12 | | | | | |
| chr16 | *CLEC18A* | 6 | 258 | 12 | LEC18A/CLEC18 | C-type lectin domain family 18 member A | Other | other | |
| chr17 | *SPATA22* | 5 | 142 | 12 | SPATA22 | spermatogenesis associated 22 | Other | other | |
| chr17 | *ZNF624* | 4 | 92 | 12 | ZNF624 | zinc finger protein 624 | Nucleus | transcription regulator | |
| chr6 | *OSTM1* | 5 | 172 | 12 | OSTM1 | osteoclastogenesis associated transmembrane protein 1 | Cytoplasm | other | |
| chr3 | *CHL1* | 5 | 98 | 12 | CHL1 | cell adhesion molecule L1 like | Plasma Membrane | other | |
| chr12 | *SDSL* | 5 | 214 | 12 | SDSL | serine dehydratase like | Cytoplasm | enzyme | |
| chr19 | *FZR1* | 6 | 277 | 12 | FZR1 | fizzy and cell division cycle 20 related 1 | Nucleus | kinase | |
| chr22 | *GAL3ST1* | 5 | 199 | 12 | GAL3ST1 | galactose-3-O-sulfotransferase 1 | Cytoplasm | enzyme | |
| chr5 | *NDUFS6* | 5 | 526 | 12 | NDUFS6 | NADH:ubiquinone oxidoreductase subunit S6 | Cytoplasm | enzyme | |
| chr6 | *FAM83B* | 5 | 108 | 12 | FAM83B | family with sequence similarity 83 member B | Cytoplasm | other | |
| chr10 | *EIF4EBP2* | 5 | 186 | 12 | EIF4EBP2 | eukaryotic translation initiation factor 4E binding protein 2 | Cytoplasm | translation regulator | |
| chr17 | *BECN1* | 5 | 142 | 12 | BECN1 | beclin 1 | Cytoplasm | other | |
| chr19 | *PRKCSH* | 5 | 163 | 12 | PRKCSH | protein kinase C substrate 80K-H | Cytoplasm | enzyme | |
| chr10 | *CUL2* | 5 | 127 | 12 | CUL2 | cullin 2 | Nucleus | enzyme | |
| chr6 | *PI16* | 5 | 258 | 12 | PI16 | peptidase inhibitor 16 | Extracellular Space | other | |

| | | | | | | | | |
|---|---|---|---|---|---|---|---|---|
| chr5 | *CTD-2001C12.1* | 5 | 125 | 12 | | | | |
| chr11 | *ROBO3* | 5 | 205 | 12 | ROBO3 | roundabout guidance receptor 3 | Plasma Membrane | transmembrane recepto |
| chr6 | *TFB1M* | 4 | 115 | 12 | TFB1M | transcription factor B1, | Cytoplasm | transcription regulator |
| chr19 | *FCGRT* | 5 | 158 | 12 | FCGRT | Fc fragment of IgG receptor and transporter | Plasma Membrane | transmembrane recepto |
| chr7 | *FIS1* | 6 | 197 | 12 | FIS1 | fission, mitochondrial 1 | Cytoplasm | other |
| chr11 | *SERPING1* | 4 | 132 | 12 | SERPING1 | serpin family G member 1 | Extracellular Space | other |
| chr5 | *RP11-1415C14.3* | 5 | 351 | 12 | | | | |
| chr5 | *CRHBP* | 6 | 176 | 12 | CRHBP | corticotropin releasing hormone binding protein | Extracellular Space | other |
| chr17 | *ABCA5* | 5 | 97 | 12 | ABCA5 | ATP binding cassette subfamily A member 5 | Plasma Membrane | transporter |
| chr17 | *ANAPC11* | 6 | 258 | 12 | ANAPC11 | anaphase promoting complex subunit 11 | Cytoplasm | enzyme |
| chr10 | *ALDH18A1* | 5 | 105 | 12 | ALDH18A1 | aldehyde dehydrogenase 18 family member A1 | Cytoplasm | kinase |
| chr8 | *GS1-24F4.2* | 6 | 177 | 12 | GS1-24F4.2 | uncharacterized LOC100652791 | Other | other |
| chr7 | *ZNF736* | 5 | 158 | 12 | ZNF736 | zinc finger protein 736 | Other | other |
| chr3 | *SLC26A6* | 5 | 198 | 12 | SLC26A6 | solute carrier family 26 member 6 | Cytoplasm | transporter |
| chr22 | *CTA-85E5.10* | 5 | 137 | 12 | | | | |
| chr1 | *ICMT* | 5 | 222 | 12 | ICMT | isoprenylcysteine carboxyl methyltransferase | Cytoplasm | enzyme |
| chr17 | *RP11-690G19.4* | 5 | 207 | 12 | | | | |
| chr6 | *SNRPC* | 5 | 140 | 12 | SNRPC | small nuclear ribonucleoprotein polypeptide C | Nucleus | other |
| chr4 | *ENPEP* | 5 | 137 | 12 | ENPEP | glutamyl aminopeptidase | Plasma Membrane | peptidase |
| chr12 | *CCDC53* | 4 | 128 | 12 | WASHC3 | WASH complex subunit 3 | Cytoplasm | other |
| chr3 | *CD200R1L* | 5 | 201 | 12 | CD200R1L | CD200 receptor 1 like | Plasma Membrane | other |
| chr11 | *NAALAD2* | 5 | 205 | 12 | NAALAD2 | N-acetylated alpha-linked acidic dipeptidase 2 | Plasma Membrane | peptidase |
| chr16 | *WDR90* | 6 | 259 | 12 | WDR90 | WD repeat domain 90 | Cytoplasm | other |
| chr17 | *CNP* | 5 | 212 | 12 | CNP | 2',3'-cyclic nucleotide 3' phosphodiesterase | Cytoplasm | enzyme |
| chr16 | *RRN3P1* | 5 | 136 | 12 | RRN3P1 | RRN3 homolog, RNA polymerase I transcription factor pseudogene | Other | other |
| chr20 | *CDC25B* | 4 | 213 | 12 | CDC25B | cell division cycle 25B | Nucleus | phosphatase |
| chr1 | *SCYL3* | 4 | 166 | 12 | SCYL3 | SCY1 like pseudokinase 3 | Cytoplasm | kinase |
| chr4 | *SLC10A7* | 4 | 112 | 12 | SLC10A7 | solute carrier family 10 member 7 | Cytoplasm | transporter |
| chr11 | *RP11-958J22.1* | 6 | 152 | 12 | | | | |
| chr14 | *KLHDC1* | 4 | 108 | 12 | KLHDC1 | kelch domain containing 1 | Cytoplasm | other |
| chr14 | *DAAM1* | 4 | 96 | 12 | DAAM1 | dishevelled associated activator of morphogenesis 1 | Cytoplasm | other |
| chr1 | *KLHL12* | 5 | 195 | 12 | KLHL12 | kelch like family member 12 | Cytoplasm | other |
| chr6 | *PEX6* | 5 | 214 | 12 | PEX6 | peroxisomal biogenesis factor 6 | Cytoplasm | enzyme |
| chr1 | *SRP9* | 5 | 151 | 12 | SRP9 | signal recognition particle 9 | Cytoplasm | other |
| chr12 | *LINC00592* | 5 | 308 | 12 | LINC00592 | long intergenic non-protein coding RNA 592 | Other | other |
| chr19 | *CTD-2043I16.1* | 5 | 139 | 12 | | | | |
| chr12 | *CCND2* | 4 | 133 | 12 | CCND2 | cyclin D2 | Nucleus | other |
| chr15 | *AC124997.1* | 5 | 211 | 12 | | | | |
| chr2 | *NEB* | 5 | 152 | 12 | NEB | nebulin | Cytoplasm | other |
| chr17 | *GNA13* | 5 | 123 | 12 | GNA13 | G protein subunit alpha 13 | Plasma Membrane | enzyme |
| chr15 | *MPI* | 5 | 196 | 12 | MPI | mannose phosphate isomerase | Cytoplasm | enzyme |
| chr3 | *RP11-657O9.1* | 5 | 165 | 12 | | | | |
| chr19 | *CCDC124* | 6 | 251 | 12 | CCDC124 | coiled-coil domain containing 124 | Other | other |
| chr20 | *BANF2* | 5 | 145 | 12 | BANF2 | barrier to autointegration factor 2 | Cytoplasm | other |
| chr17 | *TBC1D3F* | 5 | 801 | 12 | C1D3 (includes oth | TBC1 domain family member 3F | Extracellular Space | other |
| chr6 | *EFHC1* | 5 | 146 | 12 | EFHC1 | EF-hand domain containing 1 | Cytoplasm | other |
| chr12 | *CCDC60* | 6 | 116 | 12 | CCDC60 | coiled-coil domain containing 60 | Other | other |
| chr14 | *ATP6V1D* | 5 | 107 | 12 | ATP6V1D | ATPase H+ transporting V1 subunit D | Cytoplasm | transporter |
| chr4 | *ZNF827* | 4 | 103 | 12 | ZNF827 | zinc finger protein 827 | Other | other |
| chr1 | *SUCO* | 6 | 142 | 12 | SUCO | SUN domain containing ossification factor | Other | other |
| chr1 | *FAM89A* | 5 | 197 | 12 | FAM89A | family with sequence similarity 89 member A | Other | other |
| chr6 | *MAP3K4* | 4 | 105 | 12 | MAP3K4 | mitogen-activated protein kinase kinase kinase 4 | Cytoplasm | kinase |
| chr4 | *RNF150* | 5 | 100 | 12 | RNF150 | ring finger protein 150 | Other | peptidase |
| chr2 | *KYNU* | 5 | 139 | 12 | KYNU | kynureninase | Cytoplasm | enzyme |
| chr1 | *CAP1* | 5 | 115 | 12 | CAP1 | cyclase associated actin cytoskeleton regulatory protein 1 | Plasma Membrane | other |
| chr15 | *PWRN1* | 5 | 113 | 12 | PWRN1 | Prader-Willi region non-protein coding RNA 1 | Other | other |
| chr20 | *GHRH* | 5 | 263 | 12 | GHRH | growth hormone releasing | Extracellular Space | other |
| chr5 | *C6* | 5 | 336 | 12 | C6 | complement C6 | Extracellular Space | other |
| chr1 | *ANKRD35* | 6 | 212 | 12 | ANKRD35 | ankyrin repeat domain 35 | Other | other |
| chr9 | *ST6GALNAC4* | 6 | 324 | 12 | ST6GALNAC4 | ST6 N-acetylgalactosaminide alpha-2,6-sialyltransferase 4 | Cytoplasm | enzyme |
| chr18 | *CIDEA* | 5 | 171 | 12 | CIDEA | cell death inducing DFFA like effector a | Cytoplasm | other |
| chr12 | *RP11-96H19.1* | 5 | 229 | 12 | | | | |
| chr14 | *PPP4R4* | 5 | 123 | 12 | PPP4R4 | protein phosphatase 4 regulatory subunit 4 | Cytoplasm | other |
| chr2 | *RP11-521D12.2* | 5 | 140 | 12 | | | | |
| chr2 | *DRC1* | 4 | 133 | 12 | DRC1 | dynein regulatory complex | Extracellular Space | other |
| chr11 | *EHF* | 6 | 160 | 12 | EHF | ETS homologous factor | Nucleus | transcription regulator |
| chr1 | *PRKACB* | 5 | 142 | 12 | PRKACB | protein kinase cAMP-activated catalytic subunit beta | Cytoplasm | kinase |

| | | | | | | | | | |
|---|---|---|---|---|---|---|---|---|---|
| chr7 | *SLC25A40* | 5 | 138 | 12 | SLC25A40 | solute carrier family 25 member | Cytoplasm | other | |
| chr19 | *RSPH6A* | 5 | 167 | 12 | RSPH6A | radial spoke head 6 homolog A | Extracellular Space | other | |
| chr3 | *HYAL1* | 5 | 177 | 12 | HYAL1 | hyaluronidase 1 | Cytoplasm | enzyme | diclofenac/hyaluronic acid, hyaluronic acid, hylan |
| chr11 | *SIRT3* | 5 | 166 | 12 | SIRT3 | sirtuin 3 | Cytoplasm | enzyme | |
| chr5 | *ZFYVE16* | 6 | 109 | 12 | ZFYVE16 | zinc finger FYVE-type containing | Nucleus | other | |
| chr15 | *DNM1P35* | 6 | 316 | 12 | DNM1P35 | dynamin 1 pseudogene 35 | Other | other | |
| chr9 | *GALNT12* | 5 | 114 | 12 | GALNT12 | polypeptide N-acetylgalactosaminyltransferase | Cytoplasm | enzyme | |
| chr7 | *CPA1* | 5 | 245 | 12 | CPA1 | carboxypeptidase A1 | Extracellular Space | peptidase | |
| chrX | *XPNPEP2* | 5 | 169 | 12 | XPNPEP2 | X-prolyl aminopeptidase 2 | Plasma Membrane | peptidase | |
| chr3 | *APEH* | 5 | 276 | 12 | APEH | acylaminoacyl-peptide hydrolase | Cytoplasm | peptidase | |
| chr1 | *ENO1* | 5 | 172 | 12 | ENO1 | enolase 1 | Cytoplasm | enzyme | |
| chr19 | *ZNF556* | 5 | 122 | 12 | ZNF556 | zinc finger protein 556 | Other | other | |
| chr10 | *GTPBP4* | 5 | 172 | 12 | GTPBP4 | GTP binding protein 4 | Nucleus | enzyme | |
| chr5 | *GDNF* | 6 | 496 | 12 | GDNF | glial cell derived neurotrophic | Extracellular Space | growth factor | |
| chr1 | *TAF5L* | 6 | 204 | 11 | TAF5L | TATA-box binding protein associated factor 5 like | Nucleus | transcription regulator | |
| chr6 | *RNGTT* | 4 | 116 | 11 | RNGTT | RNA guanylyltransferase and 5'-phosphatase | Nucleus | phosphatase | |
| chr19 | *GCDH* | 5 | 229 | 11 | GCDH | glutaryl-CoA dehydrogenase | Cytoplasm | enzyme | |
| chr6 | *ARHGAP18* | 5 | 104 | 11 | ARHGAP18 | Rho GTPase activating protein 18 | Cytoplasm | other | |
| chr22 | *RP1-90G24.10* | 4 | 141 | 11 | | | | | |
| chr16 | *RP11-166B2.8* | 5 | 120 | 11 | | | | | |
| chr16 | *NUDT21* | 5 | 111 | 11 | NUDT21 | nudix hydrolase 21 | Nucleus | other | |
| chr12 | *RP11-175P13.3* | 5 | 101 | 11 | | | | | |
| chr2 | *NYAP2* | 5 | 106 | 11 | NYAP2 | neuronal tyrosine-phosphorylated phosphoinositide-3-kinase | Other | other | |
| chr11 | *LGALS12* | 5 | 230 | 11 | LGALS12 | galectin 12 | Extracellular Space | other | |
| chr1 | *RP11-160H22.5* | 5 | 157 | 11 | | | | | |
| chr5 | *RP11-321E2.3* | 5 | 346 | 11 | | | | | |
| chr2 | *IL1R2* | 5 | 198 | 11 | IL1R2 | interleukin 1 receptor type 2 | Plasma Membrane | transmembrane receptor | |
| chr19 | *SAFB2* | 5 | 195 | 11 | SAFB2 | scaffold attachment factor B2 | Cytoplasm | other | |
| chr10 | *RP11-452K12.7* | 5 | 137 | 11 | | | | | |
| chr8 | *RP11-127H5.1* | 5 | 131 | 11 | | | | | |
| chr14 | *DEGS2* | 5 | 237 | 11 | DEGS2 | delta 4-desaturase, sphingolipid 2 | Cytoplasm | enzyme | |
| chr20 | *EDN3* | 6 | 283 | 11 | EDN3 | endothelin 3 | Extracellular Space | other | |
| chr19 | *ZNF114* | 4 | 122 | 11 | ZNF114 | zinc finger protein 114 | Cytoplasm | transcription regulator | |
| chr2 | *PGAP1* | 5 | 145 | 11 | PGAP1 | post-GPI attachment to proteins 1 | Cytoplasm | enzyme | |
| chr6 | *FRK* | 5 | 149 | 11 | FRK | fyn related Src family tyrosine | Nucleus | kinase | regorafenib |
| chr7 | *DLX6-AS1* | 5 | 258 | 11 | DLX6-AS1 | DLX6 antisense RNA 1 | Other | other | |
| chr19 | *AC004791.2* | 5 | 131 | 11 | | | | | |
| chr1 | *RP11-101C11.1* | 5 | 100 | 11 | | | | | |
| chr2 | *AC124861.1* | 5 | 195 | 11 | | | | | |
| chr6 | *TMEM170B* | 5 | 91 | 11 | TMEM170B | transmembrane protein 170B | Plasma Membrane | other | |
| chr8 | *RHPN1* | 6 | 236 | 11 | RHPN1 | rhophilin Rho GTPase binding protein 1 | Cytoplasm | other | |
| chr20 | *RP4-610C12.1* | 6 | 347 | 11 | | | | | |
| chr15 | *SENP8* | 4 | 112 | 11 | SENP8 | SUMO peptidase family member, NEDD8 specific | Cytoplasm | peptidase | |
| chr15 | *KNSTRN* | 5 | 152 | 11 | KNSTRN | kinetochore localized astrin (SPAG5) binding protein | Cytoplasm | other | |
| chr1 | *KLF17* | 5 | 151 | 11 | KLF17 | Kruppel like factor 17 | Nucleus | transcription regulator | |
| chr17 | *ARRB2* | 5 | 182 | 11 | ARRB2 | arrestin beta 2 | Cytoplasm | other | |
| chr15 | *RP11-386M24.3* | 6 | 260 | 11 | | | | | |
| chr11 | *MS4A4E* | 5 | 130 | 11 | MS4A4E | membrane spanning 4-domains | Other | other | |
| chr1 | *RP11-57H12.5* | 4 | 110 | 11 | | | | | |
| chr16 | *DPEP1* | 6 | 219 | 11 | DPEP1 | dipeptidase 1 | Cytoplasm | peptidase | cilastatin/imipenem/relebactam, cilastatin, imipenem-cilastatin |
| chr7 | *AC147651.1* | 5 | 276 | 11 | | | | | |
| chr6 | *RNF182* | 5 | 104 | 11 | RNF182 | ring finger protein 182 | Cytoplasm | enzyme | |
| chr17 | *SLC4A1* | 5 | 213 | 11 | SLC4A1 | solute carrier family 4 member 1 (Diego blood group) | Plasma Membrane | transporter | |
| chr2 | *AC079305.8* | 5 | 125 | 11 | | | | | |
| chr1 | *RP4-639F20.1* | 6 | 135 | 11 | | | | | |
| chr20 | *PANK2* | 5 | 120 | 11 | PANK2 | pantothenate kinase 2 | Cytoplasm | kinase | |
| chr5 | *IL5* | 4 | 117 | 11 | IL5 | interleukin 5 | Extracellular Space | cytokine | mepolizumab, reslizumab |
| chr3 | *DIRC2* | 5 | 173 | 11 | SLC49A4 | solute carrier family 49 member 4 | Other | other | |
| chr5 | *MTRR* | 5 | 245 | 11 | MTRR | 5-methyltetrahydrofolate-homocysteine methyltransferase reductase | Cytoplasm | enzyme | |
| chr11 | *ENDOD1* | 5 | 235 | 11 | ENDOD1 | endonuclease domain containing | Extracellular Space | enzyme | |
| chr1 | *LRRC42* | 4 | 110 | 11 | LRRC42 | leucine rich repeat containing 42 | Other | other | |
| chr7 | *NAPEPLD* | 5 | 120 | 11 | NAPEPLD | N-acyl phosphatidylethanolamine phospholipase D | Cytoplasm | enzyme | |
| chr9 | *OSTF1* | 5 | 108 | 11 | OSTF1 | osteoclast stimulating factor 1 | Nucleus | transcription regulator | |
| chr11 | *TALDO1* | 5 | 171 | 11 | TALDO1 | transaldolase 1 | Cytoplasm | enzyme | |
| chr20 | *ERGIC3* | 6 | 249 | 11 | ERGIC3 | ERGIC and golgi 3 | Cytoplasm | other | |
| chr21 | *C21orf58* | 5 | 190 | 11 | C21orf58 | chromosome 21 open reading frame 58 | Other | other | |
| chr9 | *COQ4* | 6 | 240 | 11 | COQ4 | coenzyme Q4 | Cytoplasm | other | |
| chr15 | *RPL9P8* | 4 | 174 | 11 | | | | | |
| chr10 | *RP11-15K3.1* | 5 | 111 | 11 | | | | | |
| chr7 | *AC073842.19* | 4 | 153 | 11 | | | | | |
| chr10 | *PAOX* | 5 | 199 | 11 | PAOX | polyamine oxidase | Cytoplasm | enzyme | |
| chr9 | *RP11-18B16.2* | 5 | 139 | 11 | | | | | |

| | | | | | | | | | |
|---|---|---|---|---|---|---|---|---|---|
| chr19 | *USF2* | 6 | 226 | 11 | USF2 | upstream transcription factor 2, c-fos interacting | Nucleus | transcription regulator | |
| chr7 | *HYAL4* | 5 | 109 | 11 | HYAL4 | hyaluronidase 4 | Plasma Membrane | enzyme | diclofenac/hyaluronic acid, hyaluronic acid, hylan |
| chr10 | *ECHDC3* | 5 | 181 | 11 | ECHDC3 | enoyl-CoA hydratase domain containing 3 | Cytoplasm | enzyme | |
| chr15 | *RP11-323I15.5* | 4 | 106 | 11 | | | | | |
| chr15 | *CT60* | 5 | 183 | 11 | LINC01193 | long intergenic non-protein coding RNA 1193 | Other | other | |
| chr15 | *CEP152* | 4 | 97 | 11 | CEP152 | centrosomal protein 152 | Cytoplasm | other | |
| chr19 | *KPTN* | 5 | 163 | 11 | KPTN | kaptin, actin binding protein | Plasma Membrane | other | |
| chr5 | *CTD-2410N18.5* | 5 | 110 | 11 | | | | | |
| chr11 | *TMEM25* | 5 | 224 | 11 | TMEM25 | transmembrane protein 25 | Extracellular Space | other | |
| chr17 | *LEPREL4* | 6 | 259 | 11 | P3H4 | prolyl 3-hydroxylase family member 4 (inactive) | Nucleus | other | |
| chr11 | *RP11-867G23.10* | 5 | 297 | 11 | | | | | |
| chr1 | *CCSAP* | 5 | 190 | 11 | CCSAP | centriole, cilia and spindle associated protein | Cytoplasm | other | |
| chr6 | *GFRAL* | 5 | 138 | 11 | GFRAL | GDNF family receptor alpha like | Plasma Membrane | transmembrane receptor | |
| chrX | *MTMR8* | 4 | 194 | 11 | MTMR8 | myotubularin related protein 8 | Nucleus | phosphatase | |
| chr7 | *PDAP1* | 5 | 314 | 11 | PDAP1 | PDGFA associated protein 1 | Cytoplasm | other | |
| chr2 | *TMEM182* | 5 | 133 | 11 | TMEM182 | transmembrane protein 182 | Other | other | |
| chr5 | *RP11-574H6.1* | 5 | 129 | 11 | | | | | |
| chr10 | *LDB1* | 6 | 226 | 11 | LDB1 | LIM domain binding 1 | Nucleus | transcription regulator | |
| chr14 | *POLE2* | 3 | 97 | 11 | POLE2 | DNA polymerase epsilon 2, accessory subunit | Nucleus | enzyme | bortezomib/cladribine/rituximab, cladribine/cytarabine/decitabine, cladribine/cytarabine/filgrastim/idarubicin, cladribine/cytarabine/filgrastim/idarubicin/plerixafor, cladribine/recombinant interferon, cladribine, cladribine/cytarabine/filgrastim, cladribine/cytarabine/filgrastim/mitoxantrone, cladribine/rituximab, cladribine/cytarabine/daunorubicin |
| chr11 | *VSTM5* | 6 | 273 | 11 | VSTM5 | V-set and transmembrane domain containing 5 | Plasma Membrane | other | |
| chr15 | *DISP2* | 5 | 211 | 11 | DISP2 | dispatched RND transporter family member 2 | Plasma Membrane | other | |
| chr7 | *SRCRB4D* | 5 | 266 | 11 | SSC4D | scavenger receptor cysteine rich family member with 4 domains | Plasma Membrane | transmembrane receptor | |
| chr4 | *UVSSA* | 6 | 240 | 11 | UVSSA | UV stimulated scaffold protein A | Nucleus | other | |
| chr12 | *PWP1* | 4 | 108 | 11 | PWP1 | PWP1 homolog, endonuclein | Nucleus | other | |
| chr15 | *RP13-996F3.3* | 5 | 147 | 11 | | | | | |
| chr10 | *NUDT5* | 4 | 101 | 11 | NUDT5 | nudix hydrolase 5 | Cytoplasm | phosphatase | |
| chr9 | *DOLPP1* | 5 | 372 | 11 | DOLPP1 | dolichyldiphosphatase 1 | Cytoplasm | enzyme | |
| chr19 | *BCAT2* | 5 | 188 | 11 | BCAT2 | branched chain amino acid transaminase 2 | Cytoplasm | enzyme | |
| chr8 | *FDFT1* | 5 | 118 | 11 | FDFT1 | farnesyl-diphosphate farnesyltransferase 1 | Cytoplasm | enzyme | zoledronic acid, lapaquistat |
| chr8 | *XKR9* | 4 | 136 | 11 | XKR9 | XK related 9 | Plasma Membrane | other | |
| chr3 | *SEMA3G* | 6 | 330 | 11 | SEMA3G | semaphorin 3G | Cytoplasm | other | |
| chr19 | *RBM42* | 5 | 288 | 11 | RBM42 | RNA binding motif protein 42 | Other | other | |
| chr19 | *ZNF14* | 5 | 114 | 11 | ZNF14 | zinc finger protein 14 | Nucleus | transcription regulator | |
| chr2 | *CHRND* | 5 | 215 | 11 | CHRND | cholinergic receptor nicotinic delta subunit | Plasma Membrane | transmembrane receptor | succinylcholine, rapacuronium, doxacurium, vecuronium, pipecuronium, metocurine, mivacurium, D-tubocurarine, nicotine, arecoline, pancuronium, atracurium, isoflurane, cisatracurium, rocuronium, amobarbital, mecamylamine |
| chr18 | *ZCCHC2* | 4 | 123 | 11 | ZCCHC2 | zinc finger CCHC-type containing | Cytoplasm | other | |
| chr1 | *CCBL2* | 4 | 103 | 11 | KYAT3 | kynurenine aminotransferase 3 | Cytoplasm | enzyme | |
| chr12 | *RND1* | 5 | 277 | 11 | RND1 | Rho family GTPase 1 | Cytoplasm | enzyme | |
| chr19 | *ZNF714* | 5 | 111 | 11 | ZNF714 | zinc finger protein 714 | Other | other | |
| chr10 | *RP11-242G20.1* | 5 | 232 | 11 | | | | | |
| chr1 | *RP11-776H12.1* | 6 | 120 | 11 | | | | | |
| chr1 | *EPHX4* | 4 | 105 | 11 | EPHX4 | epoxide hydrolase 4 | Other | enzyme | |
| chr19 | *LRRC8E* | 5 | 192 | 11 | LRRC8E | leucine rich repeat containing 8 VRAC subunit E | Plasma Membrane | ion channel | |

| | | | | | | | | | |
|---|---|---|---|---|---|---|---|---|---|
| chr15 | *WHAMM* | 4 | 130 | 11 | WHAMM | WASP homolog associated with actin, golgi membranes and microtubules | Cytoplasm | other | |
| chr17 | *TM4SF5* | 5 | 206 | 11 | TM4SF5 | transmembrane 4 L six family member 5 | Plasma Membrane | other | |
| chr12 | *MYBPC1* | 5 | 113 | 11 | MYBPC1 | myosin binding protein C, slow | Cytoplasm | other | |
| chr10 | *TRIM8* | 6 | 317 | 11 | TRIM8 | tripartite motif containing 8 | Nucleus | other | |
| chr5 | *DCTN4* | 4 | 100 | 11 | DCTN4 | dynactin subunit 4 | Nucleus | other | |
| chr12 | *KRT71* | 5 | 359 | 11 | KRT71 | keratin 71 | Cytoplasm | other | |
| chr10 | *PLEKHA1* | 5 | 106 | 11 | PLEKHA1 | pleckstrin homology domain containing A1 | Cytoplasm | other | |
| chr8 | *LGI3* | 5 | 253 | 11 | LGI3 | leucine rich repeat LGI family member 3 | Extracellular Space | other | |
| chr5 | *AP3S1* | 5 | 110 | 11 | AP3S1 | adaptor related protein complex 3 subunit sigma 1 | Cytoplasm | transporter | |
| chr2 | *AC073218.1* | 4 | 115 | 11 | | | | | |
| chr13 | *TPTE2P5* | 5 | 120 | 11 | TPTE2P5 | transmembrane phosphoinositide 3-phosphatase and tensin homolog 2 pseudogene 5 | Other | other | |
| chr6 | *C6orf183* | 4 | 122 | 11 | CCDC162P | coiled-coil domain containing 162, pseudogene | Other | other | |
| chr1 | *IL19* | 5 | 150 | 11 | IL19 | interleukin 19 | Extracellular Space | cytokine | |
| chr12 | *KCTD10* | 5 | 129 | 11 | KCTD10 | potassium channel tetramerization domain containing | Nucleus | ion channel | |
| chr3 | *VILL* | 5 | 297 | 11 | VILL | villin like | Cytoplasm | other | |
| chr12 | *KRT74* | 5 | 337 | 11 | KRT74 | keratin 74 | Cytoplasm | other | |
| chr6 | *RP1-40E16.9* | 6 | 127 | 11 | | | | | |
| chr7 | *POLM* | 4 | 341 | 11 | POLM | DNA polymerase mu | Nucleus | enzyme | |
| chr14 | *CDH24* | 5 | 194 | 11 | CDH24 | cadherin 24 | Plasma Membrane | other | |
| chr1 | *GSTM2* | 5 | 200 | 11 | GSTM2 | glutathione S-transferase mu 2 | Cytoplasm | enzyme | |
| chr16 | *ZG16B* | 5 | 272 | 11 | ZG16B | zymogen granule protein 16B | Cytoplasm | other | |
| chr4 | *RP11-380P13.1* | 4 | 102 | 11 | | | | | |
| chr1 | *HIPK1* | 6 | 165 | 11 | HIPK1 | homeodomain interacting protein kinase 1 | Nucleus | kinase | |
| chr6 | *TDRG1* | 5 | 171 | 11 | TDRG1 | testis development related 1 | Other | other | |
| chr3 | *ELP6* | 5 | 157 | 11 | ELP6 | elongator acetyltransferase complex subunit 6 | Other | other | |
| chr14 | *ZBTB25* | 4 | 98 | 11 | ZBTB25 | zinc finger and BTB domain containing 25 | Nucleus | transcription regulator | |
| chr11 | *RP11-881M11.8* | 5 | 203 | 11 | | | | | |
| chr5 | *RP11-61G23.1* | 5 | 125 | 11 | | | | | |
| chr19 | *CYP4F23P* | 6 | 122 | 11 | CYP4F23P | cytochrome P450 family 4 subfamily F member 23, | Other | other | |
| chr19 | *ARHGAP33* | 5 | 209 | 11 | ARHGAP33 | Rho GTPase activating protein 33 | Plasma Membrane | transporter | |
| chr1 | *MPC2* | 3 | 86 | 11 | MPC2 | mitochondrial pyruvate carrier 2 | Plasma Membrane | other | |
| chr7 | *AC007276.5* | 4 | 127 | 11 | | | | | |
| chr11 | *RP11-166D19.1* | 5 | 90 | 11 | | | | | |
| chr2 | *RP11-1223D19.1* | 5 | 144 | 11 | | | | | |
| chr18 | *WDR7* | 4 | 116 | 11 | WDR7 | WD repeat domain 7 | Cytoplasm | other | |
| chr9 | *CEP78* | 4 | 118 | 11 | CEP78 | centrosomal protein 78 | Cytoplasm | other | |
| chrX | *TFE3* | 5 | 156 | 11 | TFE3 | transcription factor binding to IGHM enhancer 3 | Nucleus | transcription regulator | |
| chr5 | *HSPA9* | 4 | 128 | 11 | HSPA9 | heat shock protein family A (Hsp70) member 9 | Cytoplasm | other | |
| chr16 | *CX3CL1* | 5 | 183 | 11 | CX3CL1 | C-X3-C motif chemokine ligand 1 | Extracellular Space | cytokine | |
| chr1 | *MRTO4* | 5 | 198 | 11 | MRTO4 | MRT4 homolog, ribosome maturation factor | Cytoplasm | other | |
| chr8 | *FAM86B2* | 5 | 277 | 11 | AM86B2/FAM86K | family with sequence similarity 86 member B2 | Other | other | |
| chr5 | *RP11-497H16.6* | 3 | 270 | 11 | | | | | |
| chr12 | *SLC6A15* | 5 | 120 | 11 | SLC6A15 | solute carrier family 6 member 15 | Plasma Membrane | transporter | |
| chr6 | *KHDRBS2* | 4 | 129 | 11 | KHDRBS2 | KH RNA binding domain containing, signal transduction | Nucleus | other | |
| chr15 | *GOLGA8S* | 4 | 209 | 11 | GOLGA8S | golgin A8 family member S | Other | other | |
| chr6 | *ALDH5A1* | 5 | 120 | 11 | ALDH5A1 | aldehyde dehydrogenase 5 family member A1 | Cytoplasm | enzyme | theophylline/tretinoin/ valproic acid, |
| chr19 | *TMIGD2* | 5 | 156 | 11 | TMIGD2 | transmembrane and immunoglobulin domain | Other | other | |
| chr18 | *CNDP2* | 5 | 191 | 11 | CNDP2 | carnosine dipeptidase 2 | Cytoplasm | peptidase | |
| chr17 | *RCVRN* | 5 | 172 | 11 | RCVRN | recoverin | Cytoplasm | other | |
| chr19 | *CTC-457E21.9* | 5 | 115 | 11 | | | | | |
| chr11 | *KIF18A* | 4 | 127 | 11 | KIF18A | kinesin family member 18A | Cytoplasm | enzyme | |
| chr2 | *FAM161A* | 5 | 105 | 11 | FAM161A | FAM161 centrosomal protein A | Cytoplasm | other | |
| chr15 | *COPS2* | 4 | 110 | 11 | COPS2 | COP9 signalosome subunit 2 | Cytoplasm | other | |
| chr22 | *MCAT* | 5 | 174 | 11 | MCAT | malonyl-CoA-acyl carrier protein transacylase | Cytoplasm | enzyme | |
| chr2 | *SLC30A3* | 5 | 157 | 11 | SLC30A3 | solute carrier family 30 member 3 | Plasma Membrane | transporter | |
| chr19 | *ZBTB7A* | 6 | 243 | 11 | ZBTB7A | zinc finger and BTB domain containing 7A | Nucleus | transcription regulator | |
| chr2 | *FARSB* | 4 | 116 | 11 | FARSB | phenylalanyl-tRNA synthetase subunit beta | Cytoplasm | enzyme | |
| chr11 | *TTC9C* | 5 | 258 | 11 | TTC9C | tetratricopeptide repeat domain | Other | other | |
| chr1 | *CTD-2090I13.1* | 5 | 154 | 11 | | | | | |
| chr19 | *NDUFA7* | 5 | 215 | 11 | NDUFA7 | NADH:ubiquinone oxidoreductase subunit A7 | Cytoplasm | enzyme | |
| chr16 | *TMEM159* | 5 | 107 | 11 | TMEM159 | transmembrane protein 159 | Other | other | |
| chr8 | *AC133633.2* | 5 | 114 | 11 | | | | | |
| chr1 | *OR2W3* | 5 | 168 | 11 | OR2W3 | olfactory receptor family 2 subfamily W member 3 | Plasma Membrane | -protein coupled recepto | |

| | | | | | | | | | |
|---|---|---|---|---|---|---|---|---|---|
| chr1 | *AADACL4* | 5 | 116 | 11 | AADACL4 | arylacetamide deacetylase like 4 | Other | other | |
| chr1 | *RP11-10N16.2* | 5 | 163 | 11 | | | | | |
| chr1 | *KCTD3* | 3 | 121 | 11 | KCTD3 | potassium channel tetramerization domain containing | Plasma Membrane | ion channel | |
| chr15 | *CTD-2643K12.3* | 6 | 158 | 11 | | | | | |
| chrX | *PGK1* | 4 | 83 | 11 | PGK1 | phosphoglycerate kinase 1 | Cytoplasm | kinase | |
| chr12 | *CTDSP2* | 6 | 143 | 11 | CTDSP2 | CTD small phosphatase 2 | Nucleus | phosphatase | |
| chr20 | *KCNG1* | 6 | 274 | 11 | KCNG1 | potassium voltage-gated channel modifier subfamily G member 1 | Plasma Membrane | ion channel | |
| chr11 | *PSMC3* | 5 | 420 | 11 | PSMC3 | proteasome 26S subunit, ATPase | Nucleus | enzyme | |
| chr1 | *SPRTN* | 4 | 168 | 11 | SPRTN | SprT-like N-terminal domain | Nucleus | other | |
| chr7 | *DNAJC2* | 4 | 106 | 11 | DNAJC2 | DnaJ heat shock protein family (Hsp40) member C2 | Nucleus | transcription regulator | |
| chr17 | *EPN3* | 6 | 247 | 11 | EPN3 | epsin 3 | Cytoplasm | other | |
| chr10 | *CASP7* | 5 | 150 | 11 | CASP7 | caspase 7 | Cytoplasm | peptidase | |
| chr9 | *SLC35D2* | 4 | 117 | 11 | SLC35D2 | solute carrier family 35 member | Cytoplasm | transporter | |
| chr8 | *RP11-1102P16.1* | 4 | 142 | 11 | | | | | |
| chr1 | *TXNDC12* | 4 | 121 | 11 | TXNDC12 | thioredoxin domain containing 12 | Cytoplasm | enzyme | |
| chr3 | *EHHADH* | 4 | 95 | 11 | EHHADH | enoyl-CoA hydratase and 3-hydroxyacyl CoA dehydrogenase | Cytoplasm | enzyme | |
| chr5 | *EIF4E1B* | 5 | 191 | 11 | EIF4E1B | eukaryotic translation initiation factor 4E family member 1B | Other | other | |
| chr9 | *RP11-492E3.2* | 6 | 247 | 11 | | | | | |
| chr11 | *C11orf44* | 5 | 168 | 11 | C11orf44 | chromosome 11 open reading frame 44 | Other | other | |
| chr19 | *CTD-2245F17.3* | 5 | 158 | 11 | | | | | |
| chr11 | *MUC5AC* | 6 | 346 | 11 | MUC5AC | mucin 5AC, oligomeric mucus/gel-forming | Cytoplasm | peptidase | |
| chr15 | *PLIN1* | 5 | 275 | 11 | PLIN1 | perilipin 1 | Cytoplasm | other | |
| chr18 | *TXNL4A* | 5 | 115 | 11 | TXNL4A | thioredoxin like 4A | Nucleus | enzyme | |
| chr19 | *AC008753.6* | 5 | 131 | 11 | | | | | |
| chr19 | *HNRNPL* | 5 | 204 | 11 | HNRNPL | heterogeneous nuclear ribonucleoprotein L | Nucleus | other | |
| chr11 | *HYOU1* | 5 | 206 | 11 | HYOU1 | hypoxia up-regulated 1 | Cytoplasm | other | |
| chr18 | *THOC1* | 4 | 99 | 11 | THOC1 | THO complex 1 | Nucleus | transcription regulator | |
| chr18 | *TWSG1* | 4 | 86 | 11 | TWSG1 | twisted gastrulation BMP signaling modulator 1 | Extracellular Space | other | |
| chr17 | *RP11-700H6.1* | 5 | 235 | 11 | | | | | |
| chr6 | *NCOA7* | 4 | 88 | 11 | NCOA7 | nuclear receptor coactivator 7 | Nucleus | transcription regulator | |
| chr7 | *ZNF716* | 5 | 123 | 11 | ZNF716 | zinc finger protein 716 | Other | other | |
| chr1 | *RP11-193H5.1* | 5 | 175 | 11 | | | | | |
| chr6 | *OPRM1* | 4 | 110 | 11 | OPRM1 | opioid receptor mu 1 | Plasma Membrane | G-protein coupled receptor | O-demethyltramadol, fentanyl, etorphine, acetaminophen/hydrocodone, nalmefene, morphine, hydromorphone, acetaminophen/caffeine/dihydrocodeine, naltrexone/oxycodone, pentazocine, levo-alpha-acetylmethadol, codeine, acetaminophen/butalbital/caffeine/codeine, piritramide, brompheniramine/codeine/phenylpropanolamine, ethylmorphine, aspirin/hydrocodone, bupropion/naltrexone, oxycodone, [illegible] |
| chr5 | *SLC4A9* | 5 | 188 | 11 | SLC4A9 | solute carrier family 4 member 9 | Plasma Membrane | transporter | |
| chr11 | *DNAJB13* | 6 | 200 | 11 | DNAJB13 | DnaJ heat shock protein family (Hsp40) member B13 | Cytoplasm | other | |
| chr18 | *DTNA* | 5 | 97 | 11 | DTNA | dystrobrevin alpha | Plasma Membrane | other | |
| chr22 | *NIPSNAP1* | 5 | 133 | 11 | NIPSNAP1 | nipsnap homolog 1 | Cytoplasm | enzyme | |
| chr22 | *GUCD1* | 4 | 190 | 11 | GUCD1 | guanylyl cyclase domain | Other | other | |
| chr13 | *TGDS* | 5 | 109 | 11 | TGDS | TDP-glucose 4,6-dehydratase | Extracellular Space | enzyme | |
| chr1 | *TOMM20* | 4 | 145 | 11 | TOMM20 | translocase of outer mitochondrial membrane 20 | Cytoplasm | transporter | |
| chr3 | *RP11-501O2.5* | 5 | 138 | 11 | | | | | |
| chr1 | *HCN3* | 5 | 271 | 11 | HCN3 | hyperpolarization activated cyclic nucleotide gated potassium channel 3 | Plasma Membrane | ion channel | |
| chr12 | *RP11-1060G2.1* | 5 | 116 | 11 | | | | | |
| chr3 | *CYP2AB1P* | 4 | 167 | 11 | CYP2AB1P | cytochrome P450 family 2 subfamily AB member 1, | Other | other | |
| chr5 | *FAM105A* | 5 | 364 | 11 | OTULINL | OTU deubiquitinase with linear linkage specificity like | Other | other | |
| chr6 | *CEP85L* | 4 | 96 | 11 | CEP85L | centrosomal protein 85 like | Cytoplasm | other | |
| chr1 | *RP4-665J23.1* | 6 | 115 | 11 | | | | | |
| chr18 | *APCDD1* | 4 | 130 | 11 | APCDD1 | APC down-regulated 1 | Plasma Membrane | other | |
| chr1 | *LINC01031* | 4 | 131 | 11 | LINC01031 | long intergenic non-protein coding RNA 1031 | Other | other | |
| chr22 | *MIEF1* | 5 | 156 | 11 | MIEF1 | mitochondrial elongation factor 1 | Cytoplasm | other | |

| | | | | | | | | | |
|---|---|---|---|---|---|---|---|---|---|
| chr1 | *SLFNL1* | 5 | 241 | 11 | SLFNL1 | schlafen like 1 | Other | other | |
| chr20 | *SNAP25-AS1* | 5 | 138 | 11 | SNAP25-AS1 | SNAP25 antisense RNA 1 | Other | other | |
| chr15 | *LARP6* | 5 | 140 | 11 | LARP6 | La ribonucleoprotein domain family member 6 | Cytoplasm | other | |
| chr6 | *SLC17A3* | 5 | 88 | 11 | SLC17A3 | solute carrier family 17 member 3 | Plasma Membrane | transporter | |
| chr19 | *CTD-2561J22.3* | 5 | 124 | 11 | | | | | |
| chr4 | *SMIM14* | 4 | 95 | 11 | SMIM14 | small integral membrane protein | Cytoplasm | other | |
| chr3 | *AC098973.2* | 5 | 108 | 11 | | | | | |
| chr19 | *LAIR2* | 5 | 208 | 11 | LAIR2 | leukocyte associated immunoglobulin like receptor 2 | Plasma Membrane | other | |
| chr2 | *CCDC93* | 5 | 125 | 11 | CCDC93 | coiled-coil domain containing 93 | Extracellular Space | other | |
| chr16 | *RP11-598D12.4* | 5 | 219 | 11 | | | | | |
| chr2 | *HTR5BP* | 6 | 139 | 11 | HTR5BP | 5-hydroxytryptamine receptor 5B, pseudogene | Other | other | |
| chr18 | *USP14* | 5 | 106 | 11 | USP14 | ubiquitin specific peptidase 14 | Cytoplasm | peptidase | VLX1570 |
| chr11 | *MTMR2* | 4 | 156 | 11 | MTMR2 | myotubularin related protein 2 | Cytoplasm | phosphatase | |
| chr2 | *AOX2P* | 5 | 122 | 11 | AOX2P | aldehyde oxidase 2, pseudogene | Other | other | |
| chr11 | *RAPSN* | 5 | 470 | 11 | RAPSN | receptor associated protein of the synapse | Plasma Membrane | other | |
| chr3 | *RP11-299J3.8* | 3 | 141 | 11 | | | | | |
| chr5 | *F2R* | 5 | 108 | 11 | F2R | coagulation factor II thrombin receptor | Plasma Membrane | G-protein coupled recepto | chrysalin, bivalirudin, argatroban, vorapaxar, PAR1 |
| chr5 | *RMND5B* | 5 | 198 | 11 | RMND5B | required for meiotic nuclear division 5 homolog B | Cytoplasm | other | |
| chr2 | *PQLC3* | 5 | 171 | 11 | SLC66A3 | solute carrier family 66 member 3 | Other | other | |
| chr3 | *AC124944.5* | 4 | 181 | 11 | | | | | |
| chr3 | *RP11-325B23.2* | 4 | 104 | 11 | | | | | |
| chr5 | *KIAA0141* | 5 | 143 | 11 | DELE1 | DAP3 binding cell death | Cytoplasm | other | |
| chr4 | *HMX1* | 5 | 222 | 11 | HMX1 | H6 family homeobox 1 | Nucleus | transcription regulator | |
| chr10 | *SEC31B* | 5 | 119 | 11 | SEC31B | SEC31 homolog B, COPII coat complex component | Cytoplasm | other | |
| chr7 | *FAM71F1* | 6 | 117 | 11 | FAM71F1 | family with sequence similarity 71 member F1 | Other | other | |
| chr3 | *SRPRB* | 5 | 168 | 11 | SRPRB | SRP receptor subunit beta | Cytoplasm | other | |
| chr7 | *SPDYE6* | 5 | 307 | 11 | DYE2 (includes oth | speedy/RINGO cell cycle regulator family member E2 | Other | other | |
| chr2 | *XDH* | 4 | 119 | 11 | XDH | xanthine dehydrogenase | Cytoplasm | enzyme | xanthine oxidase inhibitor, allopurinol/lesinurad, allopurinol/colchicine, oxypurinol, allopurinol, |
| chr7 | *BZW2* | 5 | 122 | 11 | BZW2 | basic leucine zipper and W2 domains 2 | Cytoplasm | translation regulator | |
| chr5 | *NME5* | 4 | 112 | 11 | NME5 | NME/NM23 family member 5 | Other | kinase | |
| chr12 | *C2CD5* | 5 | 142 | 11 | C2CD5 | C2 calcium dependent domain containing 5 | Cytoplasm | other | |
| chr3 | *ACPP* | 5 | 119 | 11 | ACPP | acid phosphatase, prostate | Extracellular Space | phosphatase | |
| chr5 | *SERF1B* | 4 | 160 | 11 | SERF1A/SERF1B | small EDRK-rich factor 1A | Other | other | |
| chr2 | *FAM171B* | 4 | 125 | 11 | FAM171B | family with sequence similarity 171 member B | Other | other | |
| chr10 | *GPRIN2* | 6 | 700 | 11 | GPRIN2 | G protein regulated inducer of neurite outgrowth 2 | Other | other | |
| chr15 | *RP11-798K3.4* | 5 | 128 | 11 | | | | | |
| chr2 | *C2orf70* | 5 | 164 | 11 | FAM166C | family with sequence similarity 166 member C | Nucleus | other | |
| chr9 | *PRSS3* | 4 | 98 | 11 | PRSS3 | serine protease 3 | Extracellular Space | peptidase | |
| chr18 | *CD226* | 4 | 93 | 11 | CD226 | CD226 molecule | Plasma Membrane | other | |
| chr15 | *ADPGK* | 5 | 86 | 11 | ADPGK | ADP dependent glucokinase | Plasma Membrane | kinase | |
| chr2 | *ALS2CR12* | 5 | 120 | 11 | FLACC1 | flagellum associated containing coiled-coil domains 1 | Cytoplasm | other | |
| chr20 | *DDRGK1* | 5 | 193 | 11 | DDRGK1 | DDRGK domain containing 1 | Extracellular Space | other | |
| chr8 | *ESCO2* | 5 | 121 | 11 | ESCO2 | establishment of sister chromatid cohesion N-acetyltransferase 2 | Nucleus | enzyme | |
| chr5 | *CTD-2081C10.1* | 6 | 114 | 11 | | | | | |
| chr3 | *DPPA2* | 5 | 157 | 11 | DPPA2 | developmental pluripotency associated 2 | Nucleus | other | |
| chr16 | *ZFP1* | 5 | 109 | 11 | ZFP1 | ZFP1 zinc finger protein | Nucleus | transcription regulator | |
| chr14 | *ZFYVE26* | 4 | 131 | 11 | ZFYVE26 | zinc finger FYVE-type containing | Cytoplasm | other | |
| chrX | *AKAP14* | 4 | 94 | 11 | AKAP14 | A-kinase anchoring protein 14 | Nucleus | other | |
| chr5 | *CHD1* | 4 | 88 | 11 | CHD1 | chromodomain helicase DNA binding protein 1 | Nucleus | enzyme | epirubicin, epirubicin/paclitaxel |
| chr11 | *ACTN3* | 5 | 429 | 11 | ACTN3 | actinin alpha 3 | Plasma Membrane | other | |
| chr5 | *LYRM7* | 5 | 106 | 11 | LYRM7 | LYR motif containing 7 | Cytoplasm | other | |
| chr6 | *ME1* | 4 | 128 | 11 | ME1 | malic enzyme 1 | Cytoplasm | enzyme | |
| chr15 | *SLC27A2* | 5 | 117 | 11 | SLC27A2 | solute carrier family 27 member 2 | Cytoplasm | transporter | |
| chr5 | *PDLIM7* | 5 | 208 | 11 | PDLIM7 | PDZ and LIM domain 7 | Cytoplasm | other | |
| chr20 | *C20orf62* | 5 | 189 | 11 | LINC01620 | long intergenic non-protein coding RNA 1620 | Other | other | |
| chr17 | *RP11-15E18.4* | 4 | 123 | 11 | | | | | |
| chr14 | *PGF* | 5 | 199 | 11 | PGF | placental growth factor | Extracellular Space | growth factor | aflibercept, aflibercept/bevacizumab, |
| chr12 | *TAC3* | 4 | 162 | 11 | TAC3 | tachykinin precursor 3 | Extracellular Space | other | |
| chr16 | *FBRS* | 5 | 195 | 11 | FBRS | fibrosin | Extracellular Space | cytokine | |
| chr15 | *LCTL* | 5 | 169 | 11 | LCTL | lactase like | Cytoplasm | enzyme | |
| chr3 | *PLCD1* | 5 | 220 | 11 | PLCD1 | phospholipase C delta 1 | Cytoplasm | enzyme | |
| chr18 | *SMAD2* | 4 | 98 | 11 | SMAD2 | SMAD family member 2 | Nucleus | transcription regulator | |

| | | | | | | | | | |
|---|---|---|---|---|---|---|---|---|---|
| chr14 | *PCNX* | 4 | 98 | 11 | PCNX1 | pecanex 1 | Plasma Membrane | other | |
| chr1 | *UCK2* | 6 | 147 | 11 | UCK2 | uridine-cytidine kinase 2 | Cytoplasm | kinase | |
| chr3 | *RP11-329B9.1* | 4 | 166 | 11 | | | | | |
| chr1 | *NDUFS5* | 5 | 155 | 11 | NDUFS5 | NADH:ubiquinone oxidoreductase subunit S5 | Cytoplasm | enzyme | |
| chr11 | *BTBD10* | 4 | 121 | 11 | BTBD10 | BTB domain containing 10 | Cytoplasm | other | |
| chr5 | *GTF2H2B* | 4 | 95 | 11 | GTF2H2B | general transcription factor IIH subunit 2B (pseudogene) | Other | other | |
| chr16 | *CLEC19A* | 6 | 108 | 11 | CLEC19A | C-type lectin domain containing | Other | other | |
| chr18 | *PPP4R1* | 4 | 91 | 11 | PPP4R1 | protein phosphatase 4 regulatory subunit 1 | Other | phosphatase | |
| chr19 | *ZNF430* | 5 | 122 | 11 | ZNF430 | zinc finger protein 430 | Nucleus | transcription regulator | |
| chr4 | *RP11-384K6.6* | 5 | 126 | 11 | | | | | |
| chr18 | *CCDC178* | 4 | 133 | 11 | CCDC178 | coiled-coil domain containing 178 | Other | other | |
| chr16 | *RP11-958N24.2* | 5 | 159 | 11 | | | | | |
| chr15 | *GOLGA8CP* | 4 | 291 | 11 | GOLGA8CP | golgin A8 family member C, pseudogene | Other | other | |
| chr3 | *KIAA1143* | 5 | 127 | 11 | KIAA1143 | KIAA1143 | Other | other | |
| chr10 | *ITGA8* | 5 | 120 | 11 | ITGA8 | integrin subunit alpha 8 | Plasma Membrane | other | |
| chr15 | *TRIM69* | 4 | 129 | 11 | TRIM69 | tripartite motif containing 69 | Nucleus | enzyme | |
| chr19 | *CCDC61* | 4 | 165 | 11 | CCDC61 | coiled-coil domain containing 61 | Cytoplasm | other | |
| chr2 | *ITM2C* | 4 | 212 | 11 | ITM2C | integral membrane protein 2C | Cytoplasm | other | |
| chr1 | *UBE2T* | 5 | 186 | 11 | UBE2T | ubiquitin conjugating enzyme E2 | Nucleus | enzyme | |
| chr10 | *DDX21* | 4 | 202 | 11 | DDX21 | DExD-box helicase 21 | Nucleus | enzyme | |
| chr11 | *RP11-148I19.1* | 6 | 111 | 11 | | | | | |
| chr10 | *FAM21B* | 5 | 101 | 11 | ASHC2A/WASHC | WASH complex subunit 2A | Cytoplasm | other | |
| chr13 | *CDADC1* | 5 | 135 | 11 | CDADC1 | cytidine and dCMP deaminase domain containing 1 | Other | kinase | |
| chr18 | *CNDP1* | 4 | 136 | 11 | CNDP1 | carnosine dipeptidase 1 | Cytoplasm | peptidase | |
| chr11 | *ZNF215* | 5 | 233 | 11 | ZNF215 | zinc finger protein 215 | Nucleus | transcription regulator | |
| chr8 | *RP11-94H18.1* | 5 | 150 | 11 | | | | | |
| chr6 | *POM121L2* | 5 | 123 | 11 | POM121L2 | POM121 transmembrane nucleoporin like 2 | Extracellular Space | other | |
| chrX | *DLG3* | 4 | 123 | 11 | DLG3 | discs large MAGUK scaffold | Plasma Membrane | kinase | |
| chrX | *TCEANC* | 5 | 135 | 11 | TCEANC | transcription elongation factor A N-terminal and central domain containing | Other | other | |
| chr14 | *SOS2* | 5 | 86 | 11 | SOS2 | SOS Ras/Rho guanine nucleotide exchange factor 2 | Cytoplasm | other | |
| chr22 | *APOL3* | 4 | 139 | 11 | APOL3 | apolipoprotein L3 | Cytoplasm | transporter | |
| chr14 | *RP11-594C13.1* | 4 | 115 | 11 | | | | | |
| chr4 | *CAMK2D* | 4 | 92 | 11 | CAMK2D | calcium/calmodulin dependent protein kinase II delta | Cytoplasm | kinase | |
| chr5 | *ERCC8* | 5 | 115 | 11 | ERCC8 | ERCC excision repair 8, CSA ubiquitin ligase complex subunit | Nucleus | transcription regulator | |
| chr11 | *HARBI1* | 5 | 147 | 11 | HARBI1 | harbinger transposase derived 1 | Cytoplasm | other | |
| chr8 | *TMEM67* | 4 | 113 | 11 | TMEM67 | transmembrane protein 67 | Plasma Membrane | other | |
| chr19 | *CPT1C* | 5 | 194 | 11 | CPT1C | carnitine palmitoyltransferase 1C | Cytoplasm | enzyme | |
| chr11 | *CLP1* | 4 | 164 | 11 | CLP1 | cleavage and polyadenylation factor I subunit 1 | Nucleus | kinase | |
| chr16 | *7SK* | 5 | 96 | 11 | | | | | |
| chr3 | *RHO* | 5 | 269 | 11 | RHO | rhodopsin | Plasma Membrane | -protein coupled recepto | |
| chr5 | *RP11-1101H11.1* | 5 | 138 | 11 | | | | | |
| chr19 | *ZNF208* | 6 | 109 | 11 | ZNF208 | zinc finger protein 208 | Nucleus | transcription regulator | |
| chr1 | *GPN2* | 5 | 130 | 11 | GPN2 | GPN-loop GTPase 2 | Other | other | |
| chr1 | *RP13-279N23.2* | 5 | 223 | 11 | | | | | |
| chr5 | *HNRNPH1* | 5 | 220 | 11 | HNRNPH1 | heterogeneous nuclear ribonucleoprotein H1 | Nucleus | other | |
| chr10 | *GAD2* | 4 | 97 | 11 | GAD2 | glutamate decarboxylase 2 | Cytoplasm | enzyme | theophylline/tretinoin/ valproic acid, |
| chr11 | *EED* | 5 | 140 | 11 | EED | embryonic ectoderm development | Nucleus | transcription regulator | MAK683 |
| chr1 | *RP11-39H13.1* | 4 | 119 | 11 | | | | | |
| chr14 | *INF2* | 6 | 310 | 11 | INF2 | inverted formin, FH2 and WH2 domain containing | Cytoplasm | other | |
| chr1 | *RP5-887A10.1* | 4 | 106 | 11 | | | | | |
| chr10 | *RP11-123B3.2* | 5 | 221 | 11 | | | | | |
| chr12 | *RP11-328J6.1* | 5 | 88 | 11 | | | | | |
| chr9 | *CDC37L1* | 4 | 108 | 11 | CDC37L1 | cell division cycle 37 like 1 | Cytoplasm | other | |
| chr21 | *USP25* | 4 | 105 | 11 | USP25 | ubiquitin specific peptidase 25 | Cytoplasm | peptidase | |
| chr12 | *RP11-377D9.3* | 5 | 101 | 11 | | | | | |
| chr3 | *LINC00698* | 5 | 107 | 11 | LINC00698 | long intergenic non-protein coding RNA 698 | Other | other | |
| chr1 | *PHF13* | 5 | 266 | 11 | PHF13 | PHD finger protein 13 | Nucleus | other | |
| chr12 | *TESPA1* | 5 | 162 | 11 | TESPA1 | thymocyte expressed, positive selection associated 1 | Cytoplasm | other | |
| chr2 | *TMEM127* | 5 | 211 | 11 | TMEM127 | transmembrane protein 127 | Other | other | |
| chrX | *RP11-157D23.2* | 5 | 99 | 11 | | | | | |
| chrX | *HDAC8* | 4 | 99 | 11 | HDAC8 | histone deacetylase 8 | Nucleus | transcription regulator | tributyrin, belinostat, R 306465, |
| chr19 | *AP1M2* | 4 | 150 | 11 | AP1M2 | adaptor related protein complex 1 subunit mu 2 | Cytoplasm | transporter | |
| chr3 | *BTD* | 5 | 113 | 11 | BTD | biotinidase | Extracellular Space | enzyme | |
| chr2 | *AC012354.6* | 6 | 224 | 11 | | | | | |
| chr2 | *ARHGAP15* | 4 | 94 | 11 | ARHGAP15 | Rho GTPase activating protein 15 | Cytoplasm | other | |
| chr9 | *SPINK4* | 5 | 142 | 11 | SPINK4 | serine peptidase inhibitor, Kazal type 4 | Extracellular Space | other | |
| chr1 | *CNIH4* | 5 | 126 | 11 | CNIH4 | cornichon family AMPA receptor auxiliary protein 4 | Plasma Membrane | other | |

| | | | | | | | | | |
|---|---|---|---|---|---|---|---|---|---|
| chr7 | *FAM86LP* | 5 | 244 | 11 | FAM86LP | family with sequence similarity 86, member A pseudogene | Other | other | |
| chr22 | *CHEK2* | 4 | 100 | 11 | CHEK2 | checkpoint kinase 2 | Nucleus | kinase | XL844 |
| chr2 | *CDC42EP3* | 5 | 138 | 11 | CDC42EP3 | CDC42 effector protein 3 | Cytoplasm | other | |
| chr9 | *SMU1* | 4 | 141 | 11 | SMU1 | SMU1 DNA replication regulator and spliceosomal factor | Nucleus | other | |
| chr1 | *NPL* | 5 | 165 | 11 | NPL | N-acetylneuraminate pyruvate | Cytoplasm | enzyme | |
| chr6 | *LAMA4* | 4 | 84 | 11 | LAMA4 | laminin subunit alpha 4 | Extracellular Space | enzyme | |
| chr17 | *ALOX15B* | 5 | 172 | 11 | ALOX15B | arachidonate 15-lipoxygenase | Cytoplasm | enzyme | |
| chr22 | *POLR3H* | 5 | 158 | 11 | POLR3H | RNA polymerase III subunit H | Nucleus | enzyme | |
| chr1 | *CD53* | 5 | 114 | 11 | CD53 | CD53 molecule | Plasma Membrane | other | |
| chr2 | *EVA1A* | 6 | 138 | 11 | EVA1A | eva-1 homolog A, regulator of programmed cell death | Plasma Membrane | other | |
| chr19 | *CYP4F3* | 5 | 149 | 11 | CYP4F3 | cytochrome P450 family 4 subfamily F member 3 | Cytoplasm | enzyme | |
| chr19 | *CTD-2568A17.1* | 5 | 164 | 11 | | | | | |
| chr1 | *NT5C1A* | 5 | 154 | 11 | NT5C1A | 5'-nucleotidase, cytosolic IA | Cytoplasm | phosphatase | |
| chr2 | *RAB17* | 4 | 194 | 11 | RAB17 | RAB17, member RAS oncogene family | Cytoplasm | enzyme | |
| chr2 | *SPATS2L* | 4 | 97 | 11 | SPATS2L | spermatogenesis associated serine rich 2 like | Nucleus | other | |
| chr2 | *CERS6* | 5 | 112 | 11 | CERS6 | ceramide synthase 6 | Nucleus | transcription regulator | |
| chr12 | *ACRBP* | 5 | 197 | 11 | ACRBP | acrosin binding protein | Extracellular Space | other | |
| chr7 | *INHBA-AS1* | 5 | 124 | 11 | INHBA-AS1 | INHBA antisense RNA 1 | Other | other | |
| chr9 | *RUSC2* | 5 | 187 | 11 | RUSC2 | RUN and SH3 domain containing | Cytoplasm | other | |
| chr22 | *ALG12* | 5 | 236 | 11 | ALG12 | ALG12 alpha-1,6-mannosyltransferase | Cytoplasm | enzyme | |
| chr19 | *ZNF260* | 4 | 103 | 11 | ZNF260 | zinc finger protein 260 | Nucleus | transcription regulator | |
| chr12 | *SLC39A5* | 5 | 168 | 11 | SLC39A5 | solute carrier family 39 member 5 | Plasma Membrane | transporter | |
| chr16 | *RP11-20I23.1* | 5 | 266 | 11 | | | | | |
| chr1 | *RP1-272L16.1* | 6 | 131 | 11 | | | | | |
| chr12 | *MYL2* | 5 | 135 | 11 | MYL2 | myosin light chain 2 | Cytoplasm | other | |
| chr19 | *LONP1* | 6 | 321 | 11 | LONP1 | lon peptidase 1, mitochondrial | Cytoplasm | peptidase | |
| chr19 | *PSG3* | 4 | 126 | 11 | PSG3 | pregnancy specific beta-1-glycoprotein 3 | Extracellular Space | other | |
| chr19 | *KIR3DX1* | 5 | 169 | 11 | KIR3DX1 | killer cell immunoglobulin like receptor, three Ig domains X1 (pseudogene) | Other | other | |
| chr19 | *JSRP1* | 5 | 214 | 11 | JSRP1 | junctional sarcoplasmic reticulum protein 1 | Cytoplasm | other | |
| chr19 | *CBLC* | 4 | 171 | 11 | CBLC | Cbl proto-oncogene C | Nucleus | enzyme | |
| chr3 | *FANCD2P2* | 5 | 126 | 11 | FANCD2P2 | FANCD2 pseudogene 2 | Other | other | |
| chr11 | *RAB38* | 5 | 166 | 11 | RAB38 | RAB38, member RAS oncogene family | Cytoplasm | enzyme | |
| chr1 | *RP11-169K16.9* | 6 | 178 | 11 | | | | | |
| chr7 | *C7orf33* | 5 | 112 | 11 | C7orf33 | chromosome 7 open reading frame 33 | Other | other | |
| chr1 | *SYF2* | 5 | 142 | 11 | SYF2 | SYF2 pre-mRNA splicing factor | Nucleus | other | |
| chr9 | *PLGRKT* | 5 | 128 | 11 | PLGRKT | plasminogen receptor with a C-terminal lysine | Cytoplasm | other | |
| chr8 | *FAM83A* | 4 | 142 | 11 | FAM83A | family with sequence similarity 83 member A | Other | other | |
| chr7 | *URGCP* | 5 | 144 | 11 | URGCP | upregulator of cell proliferation | Cytoplasm | other | |
| chr17 | *RP11-1109M24.5* | 5 | 174 | 11 | | | | | |
| chr2 | *CSRNP3* | 4 | 160 | 11 | CSRNP3 | cysteine and serine rich nuclear protein 3 | Nucleus | transcription regulator | |
| chr7 | *AC004485.3* | 5 | 117 | 11 | | | | | |
| chr1 | *PLA2G2D* | 5 | 127 | 11 | PLA2G2D | phospholipase A2 group IID | Extracellular Space | enzyme | |
| chr14 | *EAPP* | 4 | 113 | 11 | EAPP | E2F associated phosphoprotein | Cytoplasm | other | |
| chr5 | *RP11-492A10.1* | 5 | 121 | 11 | | | | | |
| chr9 | *SDCCAG3* | 5 | 357 | 11 | ENTR1 | endosome associated trafficking regulator 1 | Cytoplasm | other | |
| chr5 | *UGT3A2* | 6 | 439 | 11 | UGT3A2 | UDP glycosyltransferase family 3 member A2 | Other | enzyme | |
| chr14 | *NYNRIN* | 4 | 204 | 11 | NYNRIN | NYN domain and retroviral integrase containing | Other | other | |
| chr11 | *RP11-50B3.2* | 5 | 203 | 11 | | | | | |
| chr10 | *NOC3L* | 3 | 97 | 11 | NOC3L | NOC3 like DNA replication | Nucleus | other | |
| chr17 | *RP11-180P8.1* | 5 | 101 | 11 | | | | | |
| chr4 | *CD38* | 5 | 107 | 11 | CD38 | CD38 molecule | Plasma Membrane | enzyme | TAK-079, anti-CD38 antibody, isatuximab, daratumumab, MOR03087 |
| chr11 | *RP11-119D9.1* | 5 | 102 | 11 | | | | | |
| chr11 | *RP11-396O20.1* | 5 | 124 | 11 | | | | | |
| chr21 | *KCNE1* | 4 | 160 | 11 | KCNE1 | potassium voltage-gated channel subfamily E regulatory subunit 1 | Plasma Membrane | ion channel | amlodipine/indapamide/perindopril, amiodarone, indapamide, azimilide, nicorandil |
| chr2 | *GPAT2* | 6 | 298 | 11 | GPAT2 | glycerol-3-phosphate acyltransferase 2, mitochondrial | Cytoplasm | enzyme | |
| chr2 | *MFSD6* | 5 | 103 | 11 | MFSD6 | major facilitator superfamily domain containing 6 | Plasma Membrane | transmembrane receptor | |
| chr4 | *KIAA1109* | 4 | 90 | 11 | KIAA1109 | KIAA1109 | Other | other | |
| chrX | *GLRA2* | 4 | 104 | 11 | GLRA2 | glycine receptor alpha 2 | Plasma Membrane | ion channel | caffeine/ethanol, ethanol |
| chr6 | *GLO1* | 4 | 95 | 11 | GLO1 | glyoxalase I | Cytoplasm | enzyme | |
| chr6 | *RP1-80N2.2* | 5 | 122 | 11 | | | | | |

| | | | | | | | | | |
|---|---|---|---|---|---|---|---|---|---|
| chr8 | *PAG1* | 4 | 97 | 11 | PAG1 | phosphoprotein membrane anchor with glycosphingolipid microdomains 1 | Plasma Membrane | other | |
| chr15 | *SCG3* | 5 | 127 | 11 | SCG3 | secretogranin III | Extracellular Space | other | |
| chr14 | *ARHGAP5* | 4 | 88 | 11 | ARHGAP5 | Rho GTPase activating protein 5 | Cytoplasm | enzyme | |
| chr14 | *RP11-486O13.4* | 4 | 147 | 11 | | | | | |
| chr2 | *AC019118.3* | 4 | 155 | 11 | | | | | |
| chr19 | *CTD-2192J16.20* | 4 | 98 | 11 | | | | | |
| chr12 | *NXPH4* | 5 | 245 | 11 | NXPH4 | neurexophilin 4 | Extracellular Space | other | |
| chr6 | *FANCE* | 5 | 182 | 11 | FANCE | FA complementation group E | Nucleus | other | |
| chr22 | *RP1-213J1P__B.1* | 5 | 131 | 11 | | | | | |
| chr19 | *CD320* | 5 | 264 | 11 | CD320 | CD320 molecule | Plasma Membrane | transmembrane receptor | |
| chr9 | *TNC* | 4 | 115 | 11 | TNC | tenascin C | Extracellular Space | other | F16-IL2 fusion |
| chr12 | *3C7-42389800N1* | 5 | 862 | 11 | | | | | |
| chr14 | *RPS6KL1* | 5 | 212 | 11 | RPS6KL1 | ribosomal protein S6 kinase like 1 | Other | kinase | |
| chr4 | *FAM175A* | 5 | 110 | 11 | ABRAXAS1 | abraxas 1, BRCA1 A complex subunit | Nucleus | other | |
| chr17 | *NLGN2* | 5 | 220 | 11 | NLGN2 | neuroligin 2 | Plasma Membrane | enzyme | |
| chr15 | *ZNF774* | 6 | 235 | 11 | ZNF774 | zinc finger protein 774 | Other | other | |
| chr21 | *NDUFV3* | 4 | 118 | 11 | NDUFV3 | NADH:ubiquinone oxidoreductase subunit V3 | Cytoplasm | enzyme | |
| chr12 | *USP15* | 5 | 104 | 11 | USP15 | ubiquitin specific peptidase 15 | Cytoplasm | peptidase | |
| chr15 | *RP11-505E24.2* | 5 | 306 | 11 | | | | | |
| chr3 | *RP11-91K9.1* | 4 | 102 | 11 | | | | | |
| chr1 | *LRRC39* | 5 | 131 | 11 | LRRC39 | leucine rich repeat containing 39 | Other | other | |
| chr1 | *NBPF6* | 5 | 129 | 11 | NBPF4/NBPF6 | NBPF member 6 | Other | other | |
| chr13 | *LINC00371* | 5 | 141 | 11 | C13orf42 | chromosome 13 open reading frame 42 | Other | other | |
| chr19 | *TULP2* | 4 | 156 | 11 | TULP2 | TUB like protein 2 | Other | enzyme | |
| chr5 | *RNF44* | 5 | 284 | 11 | RNF44 | ring finger protein 44 | Other | other | |
| chr11 | *GPR83* | 6 | 225 | 11 | GPR83 | G protein-coupled receptor 83 | Plasma Membrane | G-protein coupled recepto | |
| chr13 | *F7* | 5 | 225 | 11 | F7 | coagulation factor VII | Plasma Membrane | peptidase | PCI-27483, nematode |
| chr11 | *FDX1* | 5 | 116 | 11 | FDX1 | ferredoxin 1 | Cytoplasm | transporter | |
| chr2 | *GNLY* | 4 | 220 | 11 | GNLY | granulysin | Cytoplasm | other | |
| chr8 | *ADAM5* | 4 | 96 | 11 | ADAM5 | ADAM metallopeptidase domain 5 (pseudogene) | Other | other | |
| chr11 | *SSH3* | 5 | 251 | 11 | SSH3 | slingshot protein phosphatase 3 | Cytoplasm | phosphatase | |
| chr15 | *WASH3P* | 5 | 558 | 11 | WASH3P | WASP family homolog 3, pseudogene | Plasma Membrane | other | |
| chr8 | *CPSF1* | 5 | 188 | 11 | CPSF1 | cleavage and polyadenylation specific factor 1 | Nucleus | other | |
| chr20 | *CTSZ* | 5 | 301 | 11 | CTSZ | cathepsin Z | Cytoplasm | peptidase | |
| chr2 | *ARPC2* | 4 | 99 | 11 | ARPC2 | actin related protein 2/3 complex subunit 2 | Cytoplasm | other | |
| chr1 | *GABRD* | 6 | 244 | 11 | GABRD | gamma-aminobutyric acid type A receptor delta subunit | Plasma Membrane | ion channel | nitrazepam, sevoflurane, etomidate, pagoclone, zolpidem, phenobarbital, diazepam, isoflurane, halothane, pentobarbital, desflurane, adinazolam, SEP 174559, pregnenolone, zaleplon, tracazolate, enflurane, ezogabine, muscimol, flurazepam, metyrapone/oxazepam, midazolam, temazepam, diphenhydramine/midazolam, estazolam, felbamate, [illegible] |
| chr15 | *RP11-24M17.5* | 4 | 214 | 11 | | | | | |
| chrX | *ELK1* | 5 | 164 | 11 | ELK1 | ETS transcription factor ELK1 | Nucleus | transcription regulator | |
| chrX | *LINC00630* | 4 | 94 | 11 | LINC00630 | long intergenic non-protein coding RNA 630 | Other | other | |
| chr12 | *AQP6* | 5 | 465 | 11 | AQP6 | aquaporin 6 | Cytoplasm | transporter | |
| chr12 | *PIK3C2G* | 4 | 104 | 11 | PIK3C2G | phosphatidylinositol-4-phosphate 3-kinase catalytic subunit type 2 gamma | Cytoplasm | kinase | |
| chr1 | *AKIRIN1* | 4 | 124 | 11 | AKIRIN1 | akirin 1 | Cytoplasm | enzyme | |
| chr3 | *LIPH* | 4 | 85 | 11 | LIPH | lipase H | Extracellular Space | enzyme | |
| chr4 | *RP11-689P11.2* | 4 | 189 | 11 | | | | | |
| chr1 | *PGLYRP3* | 5 | 192 | 11 | PGLYRP3 | peptidoglycan recognition protein | Plasma Membrane | transmembrane receptor | |
| chr12 | *TENC1* | 5 | 374 | 11 | TNS2 | tensin 2 | Plasma Membrane | other | |
| chr21 | *AP001347.6* | 5 | 235 | 11 | | | | | |
| chr20 | *COX4I2* | 5 | 324 | 11 | COX4I2 | cytochrome c oxidase subunit 4I2 | Cytoplasm | enzyme | |
| chr16 | *RP11-296I10.3* | 5 | 183 | 11 | | | | | |
| chr8 | *HOOK3* | 4 | 106 | 11 | HOOK3 | hook microtubule tethering | Cytoplasm | other | |
| chr20 | *PFDN4* | 4 | 160 | 11 | PFDN4 | prefoldin subunit 4 | Cytoplasm | other | |
| chr2 | *AC011747.3* | 5 | 167 | 11 | | | | | |
| chr17 | *TEKT3* | 5 | 107 | 11 | TEKT3 | tektin 3 | Cytoplasm | other | |

| | | | | | | | | | |
|---|---|---|---|---|---|---|---|---|---|
| chr1 | *CREG1* | 5 | 137 | 11 | CREG1 | cellular repressor of E1A stimulated genes 1 | Nucleus | transcription regulator | |
| chr9 | *FCN2* | 5 | 321 | 11 | FCN2 | ficolin 2 | Extracellular Space | other | |
| chr5 | *FAM105B* | 5 | 294 | 11 | OTULIN | OTU deubiquitinase with linear linkage specificity | Cytoplasm | peptidase | |
| chr9 | *C9orf135* | 5 | 92 | 11 | C9orf135 | chromosome 9 open reading frame 135 | Other | other | |
| chr12 | *GOLGA2B* | 4 | 158 | 11 | GOLGA2P5 | GOLGA2 pseudogene 5 | Other | other | |
| chr3 | *AGTR1* | 5 | 149 | 11 | AGTR1 | angiotensin II receptor type 1 | Plasma Membrane | G-protein coupled recepto | candesartan cilexetil, eprosartan, amlodipine/hydrochlorothiazide/olmesartan medoxomil, clopidogrel/telmisartan, amlodipine/hydrochlorothiazide/valsartan, azilsartan, telmisartan, hydrochlorothiazide/olmesartan medoxomil, nebivolol/valsartan, sacubitril/valsartan, aliskiren/valsartan, azilsartan medoxomil/chlorthalidone, amlodipine/valsartan, amlodipine/ezetimibe/losartan/rosuvastatin, [illegible] |
| chr16 | *CTB-134H23.2* | 4 | 188 | 11 | | | | | |
| chr16 | *C16orf72* | 4 | 97 | 11 | C16orf72 | chromosome 16 open reading frame 72 | Other | other | |
| chr2 | *PROKR1* | 5 | 145 | 11 | PROKR1 | prokineticin receptor 1 | Plasma Membrane | G-protein coupled recepto | |
| chr15 | *CSPG4P11* | 5 | 522 | 11 | CSPG4P11 | chondroitin sulfate proteoglycan 4 pseudogene 11 | Other | other | |
| chr1 | *PM20D1* | 6 | 221 | 11 | PM20D1 | peptidase M20 domain containing | Cytoplasm | peptidase | |
| chr17 | *ABI3* | 5 | 186 | 11 | ABI3 | ABI family member 3 | Plasma Membrane | other | |
| chr2 | *PPM1G* | 5 | 116 | 11 | PPM1G | protein phosphatase, Mg2+/Mn2+ dependent 1G | Nucleus | phosphatase | |
| chr15 | *SKOR1* | 6 | 194 | 11 | SKOR1 | SKI family transcriptional corepressor 1 | Nucleus | transcription regulator | |
| chr7 | *GBX1* | 5 | 159 | 11 | GBX1 | gastrulation brain homeobox 1 | Nucleus | transcription regulator | |
| chr20 | *NECAB3* | 5 | 356 | 11 | NECAB3 | N-terminal EF-hand calcium binding protein 3 | Cytoplasm | other | |
| chr15 | *CHRNA3* | 5 | 214 | 11 | CHRNA3 | cholinergic receptor nicotinic alpha 3 subunit | Plasma Membrane | transmembrane recepto | succinylcholine, rapacuronium, doxacurium, vecuronium, pipecuronium, metocurine, mivacurium, levamisole, D-tubocurarine, nicotine, arecoline, pancuronium, atracurium, isoflurane, cisatracurium, rocuronium, ibogaine, amobarbital |
| chr7 | *AC003088.1* | 5 | 153 | 11 | | | | | |
| chr20 | *CSTF1* | 4 | 171 | 11 | CSTF1 | cleavage stimulation factor | Nucleus | other | |
| chr19 | *HAUS5* | 5 | 228 | 11 | HAUS5 | HAUS augmin like complex | Cytoplasm | other | |
| chr13 | *NDFIP2* | 5 | 96 | 11 | NDFIP2 | Nedd4 family interacting protein 2 | Cytoplasm | other | |
| chr1 | *MAD2L2* | 5 | 186 | 11 | MAD2L2 | mitotic arrest deficient 2 like 2 | Nucleus | enzyme | |
| chr22 | *CCDC134* | 4 | 146 | 11 | CCDC134 | coiled-coil domain containing 134 | Other | other | |
| chr7 | *TBX20* | 5 | 120 | 11 | TBX20 | T-box transcription factor 20 | Nucleus | transcription regulator | |
| chr2 | *AC023347.1* | 5 | 131 | 11 | | | | | |
| chr15 | *RP11-307C19.1* | 5 | 232 | 11 | | | | | |
| chr18 | *CLUL1* | 5 | 118 | 11 | CLUL1 | clusterin like 1 | Extracellular Space | other | |
| chr19 | *GNA15* | 5 | 143 | 11 | GNA15 | G protein subunit alpha 15 | Plasma Membrane | enzyme | |
| chr6 | *MTO1* | 5 | 99 | 11 | MTO1 | mitochondrial tRNA translation optimization 1 | Cytoplasm | enzyme | |
| chr13 | *B3GALTL* | 4 | 142 | 11 | B3GLCT | beta 3-glucosyltransferase | Cytoplasm | enzyme | |
| chr2 | *HJURP* | 4 | 153 | 11 | HJURP | Holliday junction recognition | Nucleus | other | |
| chr4 | *KIAA1239* | 5 | 107 | 11 | NWD2 | NACHT and WD repeat domain containing 2 | Other | other | |
| chr15 | *NBEAP1* | 4 | 159 | 11 | NBEAP1 | neurobeachin pseudogene 1 | Other | other | |
| chr10 | *MSMB* | 5 | 180 | 11 | MSMB | microseminoprotein beta | Extracellular Space | other | |
| chr16 | *HAS3* | 4 | 166 | 11 | HAS3 | hyaluronan synthase 3 | Plasma Membrane | enzyme | |
| chr7 | *AC092657.2* | 5 | 132 | 11 | | | | | |

| | | | | | | | | | |
|---|---|---|---|---|---|---|---|---|---|
| chr17 | *AOC3* | 5 | 192 | 11 | AOC3 | amine oxidase copper containing 3 | Plasma Membrane | enzyme | ASP8232, hydralazine/hydrochlorothiazide/reserpine , hydralazine/hydrochlorothiazide, hydralazine, |
| chr1 | *HSPB11* | 5 | 111 | 11 | HSPB11 | heat shock protein family B (small) member 11 | Extracellular Space | other | |
| chr5 | *CTB-164N12.1* | 5 | 171 | 11 | | | | | |
| chrX | *FAM156B* | 5 | 260 | 11 | AM156A/FAM156 | family with sequence similarity 156 member A | Nucleus | other | |
| chr3 | *CAMKV* | 5 | 210 | 11 | CAMKV | CaM kinase like vesicle | Plasma Membrane | kinase | |
| chr1 | *SYDE2* | 4 | 104 | 11 | SYDE2 | synapse defective Rho GTPase homolog 2 | Cytoplasm | other | |
| chr9 | *FBP2* | 4 | 113 | 11 | FBP2 | fructose-bisphosphatase 2 | Cytoplasm | phosphatase | |
| chr3 | *GPR149* | 5 | 176 | 11 | GPR149 | G protein-coupled receptor 149 | Plasma Membrane | G-protein coupled recepto | |
| chr2 | *NFE2L2* | 4 | 87 | 11 | NFE2L2 | nuclear factor, erythroid 2 like 2 | Nucleus | transcription regulator | RTA 408 |
| chr20 | *PLCB4* | 4 | 92 | 11 | PLCB4 | phospholipase C beta 4 | Cytoplasm | enzyme | |
| chr6 | *COL19A1* | 4 | 86 | 11 | COL19A1 | collagen type XIX alpha 1 chain | Extracellular Space | other | collagenase clostridium |
| chr19 | *DAND5* | 5 | 198 | 11 | DAND5 | DAN domain BMP antagonist family member 5 | Extracellular Space | other | |
| chr12 | *KRT72* | 5 | 284 | 11 | KRT72 | keratin 72 | Cytoplasm | other | |
| chr5 | *RP11-119J18.1* | 4 | 182 | 11 | | | | | |
| chrX | *GPC3* | 4 | 104 | 11 | GPC3 | glypican 3 | Plasma Membrane | other | ERY974, |
| chr5 | *LINC01020* | 6 | 273 | 11 | LINC01020 | long intergenic non-protein coding RNA 1020 | Other | other | |
| chr12 | *KIAA1467* | 5 | 107 | 11 | FAM234B | family with sequence similarity 234 member B | Other | other | |
| chr16 | *MON1B* | 4 | 165 | 11 | MON1B | MON1 homolog B, secretory trafficking associated | Cytoplasm | other | |
| chr11 | *NAA40* | 4 | 149 | 11 | NAA40 | N(alpha)-acetyltransferase 40, NatD catalytic subunit | Other | other | |
| chrX | *RP11-761E20.1* | 4 | 133 | 11 | | | | | |
| chr4 | *TEC* | 4 | 97 | 11 | TEC | tec protein tyrosine kinase | Cytoplasm | kinase | |
| chr5 | *POC5* | 4 | 165 | 11 | POC5 | POC5 centriolar protein | Cytoplasm | other | |
| chr10 | *GOT1* | 4 | 114 | 11 | GOT1 | glutamic-oxaloacetic | Cytoplasm | enzyme | |
| chr6 | *RP11-420L9.4* | 5 | 135 | 11 | | | | | |
| chr20 | *GATA5* | 6 | 281 | 11 | GATA5 | GATA binding protein 5 | Nucleus | transcription regulator | |
| chr22 | *RAB36* | 5 | 214 | 11 | RAB36 | RAB36, member RAS oncogene family | Cytoplasm | enzyme | |
| chr20 | *EEF1A2* | 6 | 455 | 11 | EEF1A2 | eukaryotic translation elongation factor 1 alpha 2 | Cytoplasm | translation regulator | |
| chr3 | *CPHL1P* | 5 | 170 | 11 | CPHL1P | ceruloplasmin and hephaestin like 1, pseudogene | Other | other | |
| chr4 | *RELL1* | 4 | 88 | 11 | RELL1 | RELT like 1 | Plasma Membrane | other | |
| chr17 | *KRT18P55* | 4 | 145 | 11 | KRT18P55 | keratin 18 pseudogene 55 | Other | other | |
| chr3 | *SI* | 4 | 95 | 11 | SI | sucrase-isomaltase | Cytoplasm | enzyme | |
| chr11 | *PATL1* | 4 | 110 | 11 | PATL1 | PAT1 homolog 1, processing body mRNA decay factor | Cytoplasm | translation regulator | |
| chr10 | *FRMPD2P2* | 5 | 133 | 11 | FRMPD2B | FERM and PDZ domain containing 2B, pseudogene | Other | other | |
| chr5 | *VCAN* | 4 | 189 | 11 | VCAN | versican | Extracellular Space | other | |
| chr22 | *XXbac-B444P24.1* | 5 | 243 | 11 | | | | | |
| chr7 | *ZNF92* | 5 | 120 | 11 | ZNF92 | zinc finger protein 92 | Nucleus | transcription regulator | |
| chr19 | *CCDC151* | 5 | 198 | 11 | CCDC151 | coiled-coil domain containing 151 | Cytoplasm | other | |
| chr9 | *ZNF169* | 4 | 113 | 11 | ZNF169 | zinc finger protein 169 | Nucleus | transcription regulator | |
| chr11 | *RELA* | 5 | 386 | 11 | RELA | RELA proto-oncogene, NF-kB subunit | Nucleus | transcription regulator | NF-kappaB decoy |
| chr4 | *FAM86MP* | 4 | 138 | 11 | FAM86MP | family with sequence similarity 86, member A pseudogene | Other | other | |
| chr16 | *STX4* | 5 | 205 | 11 | STX4 | syntaxin 4 | Plasma Membrane | transporter | |
| chr5 | *CAPSL* | 5 | 495 | 11 | CAPSL | calcyphosine like | Other | other | |
| chr12 | *NCKAP1L* | 5 | 155 | 11 | NCKAP1L | NCK associated protein 1 like | Plasma Membrane | other | |
| chr1 | *L1TD1* | 5 | 128 | 11 | L1TD1 | LINE1 type transposase domain containing 1 | Other | other | |
| chr17 | *CYB561* | 5 | 226 | 11 | CYB561 | cytochrome b561 | Cytoplasm | enzyme | |
| chr17 | *MPO* | 5 | 193 | 11 | MPO | myeloperoxidase | Cytoplasm | enzyme | AZD4831 |
| chr10 | *ABI1* | 4 | 104 | 11 | ABI1 | abl interactor 1 | Cytoplasm | other | |
| chr9 | *MLLT3* | 5 | 116 | 11 | MLLT3 | MLLT3 super elongation complex subunit | Nucleus | transcription regulator | |
| chr15 | *AC100830.3* | 4 | 147 | 11 | | | | | |
| chr5 | *ARSK* | 4 | 98 | 11 | ARSK | arylsulfatase family member K | Extracellular Space | enzyme | |
| chr15 | *GOLGA8DP* | 5 | 265 | 11 | GOLGA8DP | golgin A8 family member D, pseudogene | Other | other | |
| chr10 | *RP11-34D15.2* | 5 | 165 | 11 | | | | | |
| chr16 | *RP11-26L20.3* | 6 | 99 | 11 | | | | | |
| chr4 | *UBA6-AS1* | 4 | 107 | 11 | UBA6-AS1 | UBA6 antisense RNA 1 (head to head) | Other | other | |
| chr20 | *PCIF1* | 5 | 282 | 11 | PCIF1 | PDX1 C-terminal inhibiting factor | Nucleus | enzyme | |
| chr16 | *GSPT1* | 5 | 95 | 11 | GSPT1 | G1 to S phase transition 1 | Cytoplasm | translation regulator | |
| chr6 | *SMAP1* | 4 | 105 | 11 | SMAP1 | small ArfGAP 1 | Cytoplasm | other | |
| chr2 | *AC097499.1* | 5 | 143 | 10 | | | | | |
| chr18 | *RP11-751H17.1* | 4 | 132 | 10 | | | | | |
| chr5 | *MEF2C* | 5 | 91 | 10 | MEF2C | myocyte enhancer factor 2C | Nucleus | transcription regulator | |
| chr11 | *RP11-25I9.2* | 4 | 86 | 10 | | | | | |
| chr19 | *DMRTC2* | 5 | 150 | 10 | DMRTC2 | DMRT like family C2 | Nucleus | transcription regulator | |
| chr19 | *GMIP* | 5 | 254 | 10 | GMIP | GEM interacting protein | Cytoplasm | enzyme | |

| | | | | | | | | | |
|---|---|---|---|---|---|---|---|---|---|
| chr2 | ARHGEF33 | 5 | 134 | 10 | ARHGEF33 | Rho guanine nucleotide exchange factor 33 | Other | other | |
| chr5 | CTD-2245E15.3 | 5 | 551 | 10 | | | | | |
| chr18 | RPRD1A | 4 | 83 | 10 | RPRD1A | regulation of nuclear pre-mRNA domain containing 1A | Extracellular Space | other | |
| chr7 | RP11-545G3.1 | 5 | 118 | 10 | | | | | |
| chr1 | RP11-38J22.6 | 5 | 201 | 10 | | | | | |
| chr11 | CTNND1 | 5 | 123 | 10 | CTNND1 | catenin delta 1 | Nucleus | other | |
| chr3 | C3orf33 | 5 | 117 | 10 | C3orf33 | chromosome 3 open reading frame 33 | Extracellular Space | other | |
| chr20 | LINC00494 | 4 | 215 | 10 | LINC00494 | long intergenic non-protein coding RNA 494 | Other | other | |
| chr7 | SLC4A2 | 5 | 258 | 10 | SLC4A2 | solute carrier family 4 member 2 | Plasma Membrane | transporter | |
| chr4 | PRDM5 | 4 | 131 | 10 | PRDM5 | PR/SET domain 5 | Nucleus | transcription regulator | |
| chr12 | PIWIL1 | 4 | 103 | 10 | PIWIL1 | piwi like RNA-mediated gene silencing 1 | Cytoplasm | enzyme | |
| chr19 | CCDC159 | 4 | 145 | 10 | CCDC159 | coiled-coil domain containing 159 | Other | other | |
| chr2 | AC016768.1 | 5 | 108 | 10 | | | | | |
| chr1 | UQCRH | 5 | 108 | 10 | UQCRH | ubiquinol-cytochrome c reductase hinge protein | Cytoplasm | enzyme | |
| chr14 | GCH1 | 4 | 106 | 10 | GCH1 | GTP cyclohydrolase 1 | Cytoplasm | enzyme | |
| chr12 | USP5 | 4 | 181 | 10 | USP5 | ubiquitin specific peptidase 5 | Cytoplasm | peptidase | |
| chr16 | C16orf58 | 5 | 211 | 10 | C16orf58 | chromosome 16 open reading frame 58 | Other | other | |
| chr1 | TAF13 | 5 | 148 | 10 | TAF13 | TATA-box binding protein associated factor 13 | Nucleus | transcription regulator | |
| chr22 | DENND6B | 5 | 212 | 10 | DENND6B | DENN domain containing 6B | Cytoplasm | other | |
| chr3 | ATP13A4 | 4 | 78 | 10 | ATP13A4 | ATPase 13A4 | Cytoplasm | transporter | |
| chr12 | RP11-776A13.3 | 5 | 124 | 10 | | | | | |
| chr11 | AP002856.7 | 5 | 164 | 10 | | | | | |
| chr3 | ABTB1 | 5 | 296 | 10 | ABTB1 | ankyrin repeat and BTB domain containing 1 | Cytoplasm | translation regulator | |
| chr19 | RAB8A | 4 | 152 | 10 | RAB8A | RAB8A, member RAS oncogene family | Plasma Membrane | enzyme | |
| chr7 | LEP | 5 | 172 | 10 | LEP | leptin | Extracellular Space | growth factor | |
| chr2 | MFSD2B | 5 | 183 | 10 | MFSD2B | major facilitator superfamily domain containing 2B | Plasma Membrane | transporter | |
| chr16 | BCAR4 | 5 | 119 | 10 | BCAR4 | breast cancer anti-estrogen resistance 4 | Other | other | |
| chr19 | ZNF558 | 5 | 97 | 10 | ZNF558 | zinc finger protein 558 | Nucleus | transcription regulator | |
| chr5 | CDX1 | 4 | 143 | 10 | CDX1 | caudal type homeobox 1 | Nucleus | transcription regulator | |
| chrX | HCFC1 | 5 | 254 | 10 | HCFC1 | host cell factor C1 | Nucleus | transcription regulator | |
| chr3 | TF | 5 | 168 | 10 | TF | transferrin | Extracellular Space | transporter | NKP-1339, ferric carboxymaltose |
| chr9 | MED22 | 5 | 318 | 10 | MED22 | mediator complex subunit 22 | Cytoplasm | other | |
| chr1 | TNFRSF9 | 5 | 102 | 10 | TNFRSF9 | TNF receptor superfamily member 9 | Plasma Membrane | transmembrane receptor | urelumab, INBRX 105, GEN1046, PRS-343, PF-05082566, anti-CD137 antibody, RO7227166, CTX-471, ADG106 |
| chrX | TEX28P2 | 4 | 128 | 10 | TEX28P2 | testis expressed 28 pseudogene | Other | other | |
| chr8 | ENPP7P1 | 5 | 317 | 10 | ENPP7P1 | ectonucleotide pyrophosphatase/phosphodiesterase 7 pseudogene 1 | Other | other | |
| chr6 | PRPF4B | 4 | 92 | 10 | PRPF4B | pre-mRNA processing factor 4B | Nucleus | kinase | |
| chr20 | PLAGL2 | 5 | 182 | 10 | PLAGL2 | PLAG1 like zinc finger 2 | Nucleus | transcription regulator | |
| chr15 | GOLGA6L1 | 5 | 329 | 10 | GA6L1 (includes o | golgin A6 family like 1 | Other | other | |
| chr14 | CTD-2058B24.2 | 4 | 92 | 10 | | | | | |
| chr1 | SLC27A3 | 4 | 264 | 10 | SLC27A3 | solute carrier family 27 member 3 | Cytoplasm | transporter | |
| chr1 | RP4-655J12.4 | 5 | 185 | 10 | | | | | |
| chr11 | SLC22A18AS | 5 | 493 | 10 | SLC22A18AS | solute carrier family 22 member 18 antisense | Other | other | |
| chrX | NSDHL | 4 | 126 | 10 | NSDHL | NAD(P) dependent steroid dehydrogenase-like | Cytoplasm | enzyme | |
| chr16 | RSL1D1 | 5 | 102 | 10 | RSL1D1 | ribosomal L1 domain containing 1 | Nucleus | other | |
| chr17 | LGALS3BP | 5 | 207 | 10 | LGALS3BP | galectin 3 binding protein | Plasma Membrane | transmembrane receptor | |
| chr7 | RP11-460N20.5 | 4 | 163 | 10 | | | | | |
| chr19 | ZNF461 | 5 | 116 | 10 | ZNF461 | zinc finger protein 461 | Nucleus | transcription regulator | |
| chr21 | IL10RB | 4 | 159 | 10 | IL10RB | interleukin 10 receptor subunit | Plasma Membrane | transmembrane receptor | |
| chr6 | MEI4 | 5 | 113 | 10 | MEI4 | meiotic double-stranded break formation protein 4 | Nucleus | other | |
| chr7 | PRRT4 | 5 | 172 | 10 | PRRT4 | proline rich transmembrane | Other | other | |
| chr4 | CTBP1-AS2 | 4 | 147 | 10 | CTBP1-DT | CTBP1 divergent transcript | Other | other | |
| chr14 | DAD1 | 4 | 111 | 10 | DAD1 | defender against cell death 1 | Cytoplasm | other | |
| chr6 | SLC17A1 | 4 | 87 | 10 | SLC17A1 | solute carrier family 17 member 1 | Plasma Membrane | transporter | |
| chr20 | RP5-823G15.5 | 5 | 196 | 10 | | | | | |
| chr10 | GDF10 | 5 | 276 | 10 | GDF10 | growth differentiation factor 10 | Extracellular Space | growth factor | |
| chr10 | ARHGAP21 | 4 | 85 | 10 | ARHGAP21 | Rho GTPase activating protein 21 | Cytoplasm | other | |
| chr3 | ZNF445 | 5 | 121 | 10 | ZNF445 | zinc finger protein 445 | Nucleus | transcription regulator | |
| chr17 | RUNDC1 | 5 | 140 | 10 | RUNDC1 | RUN domain containing 1 | Other | other | |
| chr17 | PPP1R1B | 5 | 228 | 10 | PPP1R1B | protein phosphatase 1 regulatory inhibitor subunit 1B | Cytoplasm | phosphatase | |
| chr3 | MON1A | 5 | 101 | 10 | MON1A | MON1 homolog A, secretory trafficking associated | Cytoplasm | other | |
| chr17 | CTD-2104P17.1 | 5 | 97 | 10 | | | | | |
| chr15 | HAPLN3 | 5 | 237 | 10 | HAPLN3 | hyaluronan and proteoglycan link protein 3 | Extracellular Space | other | |
| chr11 | AP000769.1 | 5 | 231 | 10 | | | | | |

| | | | | | | | | | |
|---|---|---|---|---|---|---|---|---|---|
| chr1 | *DEGS1* | 5 | 163 | 10 | DEGS1 | delta 4-desaturase, sphingolipid 1 | Plasma Membrane | enzyme | |
| chr15 | *CYP19A1* | 5 | 88 | 10 | CYP19A1 | cytochrome P450 family 19 subfamily A member 1 | Cytoplasm | enzyme | abemaciclib/anastrozole, lapatinib/letrozole/trastuzumab, exemestane/lapatinib, anastrozole/fulvestrant/goserelin/tamoxifen, steroidal aromatase inhibitor, anastrozole/fulvestrant/tamoxifen, anastrozole/exemestane/fulvestrant/tamoxifen, exemestane/tamoxifen, letrozole/tamoxifen, fulvestrant/letrozole, exemestane/palbociclib, fulvestrant/letrozole/tamoxifen, [illegible] |
| chr11 | *PRKRIR* | 5 | 156 | 10 | THAP12 | THAP domain containing 12 | Nucleus | transcription regulator | |
| chr10 | *NHLRC2* | 4 | 116 | 10 | NHLRC2 | NHL repeat containing 2 | Cytoplasm | enzyme | |
| chr3 | *EXOSC7* | 5 | 111 | 10 | EXOSC7 | exosome component 7 | Nucleus | enzyme | |
| chr4 | *DOK7* | 5 | 290 | 10 | DOK7 | docking protein 7 | Extracellular Space | other | |
| chr5 | *PAIP1* | 5 | 262 | 10 | PAIP1 | poly(A) binding protein interacting protein 1 | Cytoplasm | translation regulator | |
| chr3 | *WNT5A* | 5 | 209 | 10 | WNT5A | Wnt family member 5A | Extracellular Space | cytokine | |
| chr1 | *RP11-576I22.2* | 5 | 111 | 10 | | | | | |
| chr2 | *AC116050.1* | 6 | 852 | 10 | | | | | |
| chr6 | *TPMT* | 5 | 97 | 10 | TPMT | thiopurine S-methyltransferase | Cytoplasm | enzyme | olsalazine |
| chr3 | *CTD-2185K10.1* | 4 | 84 | 10 | | | | | |
| chr12 | *RP11-54A9.1* | 5 | 99 | 10 | | | | | |
| chr5 | *CTC-529P8.1* | 4 | 97 | 10 | | | | | |
| chr11 | *STIP1* | 5 | 146 | 10 | STIP1 | stress induced phosphoprotein 1 | Cytoplasm | other | lavendustin C |
| chr2 | *SF3B1* | 4 | 97 | 10 | SF3B1 | splicing factor 3b subunit 1 | Nucleus | other | H3B-8800 |
| chr2 | *POLR2D* | 4 | 156 | 10 | POLR2D | RNA polymerase II subunit D | Nucleus | enzyme | |
| chr14 | *ZFYVE21* | 5 | 233 | 10 | ZFYVE21 | zinc finger FYVE-type containing | Other | other | |
| chr15 | *RP11-516C1.1* | 4 | 102 | 10 | | | | | |
| chr1 | *RP11-978I15.10* | 5 | 140 | 10 | | | | | |
| chr15 | *LTK* | 5 | 208 | 10 | LTK | leukocyte receptor tyrosine | Plasma Membrane | kinase | lorlatinib |
| chr15 | *GOLGA6L4* | 4 | 341 | 10 | GA6L9 (includes o | golgin A6 family like 9 | Other | other | |
| chr4 | *GABRB1* | 4 | 102 | 10 | GABRB1 | gamma-aminobutyric acid type A receptor beta1 subunit | Plasma Membrane | ion channel | triazolam, nitrazepam, methohexital, sevoflurane, etomidate, lindane, pagoclone, aspirin/butalbital/caffeine, zolpidem, atropine/hyoscyamine/phenobarbital/scopolamine, phenobarbital, diazepam, isoflurane, halothane, pentobarbital, acetaminophen/butalbital/caffeine/codeine, hyoscyamine/phenobarbital, desflurane, amobarbital, adinazolam, SEP 174559, [illegible] |
| chr3 | *NUP210P1* | 5 | 226 | 10 | NUP210P1 | nucleoporin 210 pseudogene 1 | Other | other | |
| chr12 | *RP11-181C3.1* | 5 | 120 | 10 | | | | | |
| chr5 | *AC008834.1* | 5 | 109 | 10 | | | | | |
| chr3 | *DZIP3* | 5 | 140 | 10 | DZIP3 | DAZ interacting zinc finger | Cytoplasm | enzyme | |
| chr1 | *CHI3L1* | 5 | 249 | 10 | CHI3L1 | chitinase 3 like 1 | Extracellular Space | enzyme | |
| chr3 | *IL20RB* | 5 | 190 | 10 | IL20RB | interleukin 20 receptor subunit | Plasma Membrane | other | |
| chr1 | *PGLYRP4* | 5 | 166 | 10 | PGLYRP4 | peptidoglycan recognition protein | Plasma Membrane | transmembrane receptor | |
| chr16 | *U95743.1* | 5 | 86 | 10 | | | | | |
| chr19 | *FKRP* | 5 | 125 | 10 | FKRP | fukutin related protein | Cytoplasm | other | |
| chr11 | *LPXN* | 5 | 121 | 10 | LPXN | leupaxin | Cytoplasm | transcription regulator | |
| chr15 | *BNIP2* | 3 | 87 | 10 | BNIP2 | BCL2 interacting protein 2 | Cytoplasm | other | |
| chr6 | *MLLT4* | 4 | 119 | 10 | AFDN | afadin, adherens junction formation factor | Nucleus | other | |
| chr5 | *RP11-1084J3.4* | 4 | 228 | 10 | | | | | |
| chr15 | *FURIN* | 6 | 459 | 10 | FURIN | furin, paired basic amino acid cleaving enzyme | Cytoplasm | peptidase | nona-D-arginine amide, furin inhibitor, hexa-D-arginine |
| chr4 | *ART3* | 4 | 121 | 10 | ART3 | ADP-ribosyltransferase 3 | Plasma Membrane | enzyme | |
| chr6 | *SLC22A2* | 4 | 111 | 10 | SLC22A2 | solute carrier family 22 member 2 | Plasma Membrane | transporter | |

| | | | | | | | | | |
|---|---|---|---|---|---|---|---|---|---|
| chr8 | *RP11-429J17.7* | 5 | 123 | 10 | | | | | |
| chr5 | *MIR143HG* | 4 | 140 | 10 | CARMN | cardiac mesoderm enhancer-associated non-coding RNA | Other | other | |
| chr5 | *GM2A* | 4 | 106 | 10 | GM2A | GM2 ganglioside activator | Cytoplasm | enzyme | |
| chr14 | *CTD-2307P3.1* | 4 | 99 | 10 | | | | | |
| chr9 | *REXO4* | 5 | 281 | 10 | REXO4 | REX4 homolog, 3'-5' exonuclease | Nucleus | transcription regulator | |
| chr9 | *LPPR1* | 4 | 159 | 10 | | | | | |
| chr10 | *PDZD8* | 4 | 96 | 10 | PDZD8 | PDZ domain containing 8 | Extracellular Space | other | |
| chr5 | *CD74* | 4 | 143 | 10 | CD74 | CD74 molecule | Plasma Membrane | transmembrane recepto | milatuzumab |
| chr19 | *ZNF429* | 5 | 111 | 10 | ZNF429 | zinc finger protein 429 | Nucleus | transcription regulator | |
| chr19 | *FPR1* | 4 | 102 | 10 | FPR1 | formyl peptide receptor 1 | Plasma Membrane | G-protein coupled recepto | |
| chr4 | *PTPN13* | 5 | 108 | 10 | PTPN13 | protein tyrosine phosphatase non-receptor type 13 | Cytoplasm | phosphatase | |
| chr1 | *PITHD1* | 4 | 97 | 10 | PITHD1 | PITH domain containing 1 | Nucleus | other | |
| chr1 | *RP4-694A7.4* | 5 | 91 | 10 | | | | | |
| chr1 | *TMED5* | 5 | 113 | 10 | TMED5 | transmembrane p24 trafficking protein 5 | Cytoplasm | other | |
| chr7 | *ARHGEF5* | 5 | 199 | 10 | ARHGEF5 | Rho guanine nucleotide exchange factor 5 | Cytoplasm | other | |
| chr7 | *AC002463.3* | 5 | 122 | 10 | | | | | |
| chr1 | *BNIPL* | 5 | 149 | 10 | BNIPL | BCL2 interacting protein like | Cytoplasm | other | |
| chr19 | *AC011526.1* | 5 | 127 | 10 | | | | | |
| chr1 | *RIMKLA* | 5 | 92 | 10 | RIMKLA | ribosomal modification protein rimK like family member A | Cytoplasm | enzyme | |
| chr19 | *PLIN4* | 5 | 272 | 10 | PLIN4 | perilipin 4 | Cytoplasm | other | |
| chr1 | *ZFP69* | 4 | 94 | 10 | ZFP69 | ZFP69 zinc finger protein | Nucleus | transcription regulator | |
| chr1 | *RP11-278H7.4* | 5 | 114 | 10 | | | | | |
| chr20 | *SYCP2* | 4 | 113 | 10 | SYCP2 | synaptonemal complex protein 2 | Nucleus | other | |
| chr17 | *GHDC* | 5 | 239 | 10 | GHDC | GH3 domain containing | Cytoplasm | other | |
| chr13 | *LINC00571* | 3 | 106 | 10 | LINC00571 | long intergenic non-protein coding RNA 571 | Other | other | |
| chr14 | *YLPM1* | 4 | 106 | 10 | YLPM1 | YLP motif containing 1 | Nucleus | transcription regulator | |
| chr16 | *RP11-351A20.1* | 4 | 84 | 10 | | | | | |
| chr2 | *PREPL* | 5 | 138 | 10 | PREPL | prolyl endopeptidase like | Other | peptidase | |
| chr2 | *ICA1L* | 4 | 103 | 10 | ICA1L | islet cell autoantigen 1 like | Cytoplasm | other | |
| chr12 | *SCNN1A* | 4 | 166 | 10 | SCNN1A | sodium channel epithelial 1 alpha subunit | Plasma Membrane | ion channel | hydrochlorothiazide/triamterene, amiloride, triamterene, amiloride/hydrochlorothiazide |
| chr13 | *LINC00559* | 4 | 108 | 10 | LINC00559 | long intergenic non-protein coding RNA 559 | Other | other | |
| chr16 | *PKD1P6* | 5 | 228 | 10 | PKD1P6 | polycystin 1, transient receptor potential channel interacting pseudogene 6 | Other | other | |
| chr5 | *MRPL22* | 4 | 95 | 10 | MRPL22 | mitochondrial ribosomal protein | Cytoplasm | other | |
| chrX | *CHDC2* | 4 | 79 | 10 | CFAP47 | cilia and flagella associated | Other | other | |
| chr12 | *E2F7* | 5 | 101 | 10 | E2F7 | E2F transcription factor 7 | Nucleus | transcription regulator | |
| chr12 | *MYO1A* | 5 | 168 | 10 | MYO1A | myosin IA | Cytoplasm | peptidase | |
| chr17 | *PSMD12* | 3 | 99 | 10 | PSMD12 | proteasome 26S subunit, non-ATPase 12 | Cytoplasm | other | |
| chr2 | *STK36* | 4 | 111 | 10 | STK36 | serine/threonine kinase 36 | Cytoplasm | kinase | cetuximab/encorafenib, binimetinib/encorafenib, encorafenib, binimetinib/cetuximab/encorafenib, binimetinib/encorafen |
| chr2 | *AC007879.5* | 4 | 80 | 10 | | | | | |
| chr1 | *RP11-424D14.1* | 5 | 117 | 10 | | | | | |
| chr1 | *MST1P2* | 4 | 432 | 10 | MST1P2 | macrophage stimulating 1 pseudogene 2 | Other | other | |
| chr16 | *SEPT12* | 5 | 176 | 10 | SEPTIN12 | septin 12 | Cytoplasm | other | |
| chr15 | *GOLGA8O* | 4 | 152 | 10 | GA8K (includes ot | golgin A8 family member K | Other | other | |
| chr17 | *CTD-2377D24.8* | 5 | 89 | 10 | | | | | |
| chr11 | *RP11-867O8.5* | 5 | 275 | 10 | | | | | |
| chr15 | *RPS27L* | 3 | 85 | 10 | RPS27L | ribosomal protein S27 like | Cytoplasm | translation regulator | |
| chr15 | *RP11-665J16.1* | 4 | 145 | 10 | | | | | |
| chr13 | *SLC25A30* | 4 | 88 | 10 | SLC25A30 | solute carrier family 25 member | Cytoplasm | other | |
| chr15 | *TPM1* | 4 | 131 | 10 | TPM1 | tropomyosin 1 | Cytoplasm | other | |
| chr2 | *LIMS3* | 4 | 182 | 10 | LIMS3/LIMS4 | LIM zinc finger domain containing | Other | other | |
| chr16 | *ADAT1* | 5 | 154 | 10 | ADAT1 | adenosine deaminase tRNA specific 1 | Other | enzyme | |
| chr12 | *CCDC42B* | 5 | 156 | 10 | CFAP73 | cilia and flagella associated | Cytoplasm | other | |
| chr20 | *ACTR5* | 5 | 167 | 10 | ACTR5 | actin related protein 5 | Nucleus | other | |
| chr5 | *RP11-143A12.3* | 5 | 370 | 10 | | | | | |
| chr19 | *ZNF566* | 4 | 97 | 10 | ZNF566 | zinc finger protein 566 | Nucleus | transcription regulator | |
| chr17 | *NPTX1* | 5 | 253 | 10 | NPTX1 | neuronal pentraxin 1 | Extracellular Space | other | |
| chr14 | *RP11-159L20.2* | 4 | 97 | 10 | | | | | |
| chr19 | *GMFG* | 5 | 128 | 10 | GMFG | glia maturation factor gamma | Cytoplasm | growth factor | |
| chr12 | *TCTN1* | 4 | 105 | 10 | TCTN1 | tectonic family member 1 | Extracellular Space | other | |
| chr5 | *PRR16* | 5 | 121 | 10 | PRR16 | proline rich 16 | Other | other | |
| chr3 | *SACM1L* | 4 | 88 | 10 | SACM1L | SAC1 like phosphatidylinositide phosphatase | Cytoplasm | phosphatase | |
| chr12 | *MRPL42* | 4 | 119 | 10 | MRPL42 | mitochondrial ribosomal protein | Cytoplasm | other | |
| chr12 | *MCRS1* | 4 | 307 | 10 | MCRS1 | microspherule protein 1 | Nucleus | other | |
| chr3 | *MLF1* | 4 | 144 | 10 | MLF1 | myeloid leukemia factor 1 | Nucleus | other | |
| chr9 | *CACFD1* | 5 | 383 | 10 | CACFD1 | calcium channel flower domain containing 1 | Plasma Membrane | ion channel | |

| | | | | | | | | | |
|---|---|---|---|---|---|---|---|---|---|
| chr11 | *UBE2L6* | 5 | 107 | 10 | UBE2L6 | ubiquitin conjugating enzyme E2 | Cytoplasm | enzyme | |
| chr9 | *C9orf64* | 5 | 142 | 10 | C9orf64 | chromosome 9 open reading frame 64 | Other | other | |
| chr19 | *NAPA* | 4 | 172 | 10 | NAPA | NSF attachment protein alpha | Cytoplasm | transporter | |
| chr6 | *NFKBIE* | 4 | 195 | 10 | NFKBIE | NFKB inhibitor epsilon | Nucleus | transcription regulator | |
| chr6 | *GPR111* | 5 | 90 | 10 | ADGRF2 | adhesion G protein-coupled receptor F2 | Plasma Membrane | i-protein coupled recepto | |
| chr9 | *RP11-255A11.21* | 5 | 149 | 10 | | | | | |
| chr16 | *PDP2* | 5 | 199 | 10 | PDP2 | pyruvate dehyrogenase phosphatase catalytic subunit 2 | Cytoplasm | phosphatase | |
| chr1 | *RP11-216N14.5* | 5 | 283 | 10 | | | | | |
| chr12 | *MED21* | 5 | 155 | 10 | MED21 | mediator complex subunit 21 | Nucleus | transcription regulator | |
| chr6 | *BAI3* | 4 | 103 | 10 | ADGRB3 | adhesion G protein-coupled receptor B3 | Plasma Membrane | i-protein coupled recepto | |
| chr5 | *CNOT8* | 4 | 128 | 10 | CNOT8 | CCR4-NOT transcription complex subunit 8 | Nucleus | transcription regulator | |
| chr6 | *MRPL14* | 4 | 134 | 10 | MRPL14 | mitochondrial ribosomal protein | Cytoplasm | other | |
| chr20 | *RP11-560A15.3* | 5 | 266 | 10 | | | | | |
| chr22 | *L3MBTL2* | 4 | 154 | 10 | L3MBTL2 | L3MBTL histone methyl-lysine binding protein 2 | Nucleus | transcription regulator | |
| chr2 | *PAIP2B* | 5 | 125 | 10 | PAIP2B | poly(A) binding protein interacting protein 2B | Other | translation regulator | |
| chr5 | *CTD-2117L12.1* | 4 | 154 | 10 | | | | | |
| chr6 | *GCNT6* | 4 | 98 | 10 | GCNT2P1 | GCNT2 pseudogene 1 | Other | other | |
| chr3 | *CT64* | 5 | 131 | 10 | LINC01192 | long intergenic non-protein coding RNA 1192 | Other | other | |
| chr1 | *EMBP1* | 4 | 224 | 10 | EMBP1 | embigin pseudogene 1 | Other | other | |
| chr2 | *RP11-89K21.1* | 6 | 346 | 10 | | | | | |
| chr13 | *ANKRD10* | 4 | 126 | 10 | ANKRD10 | ankyrin repeat domain 10 | Nucleus | transcription regulator | |
| chr1 | *C1orf185* | 4 | 99 | 10 | C1orf185 | chromosome 1 open reading frame 185 | Other | other | |
| chr12 | *TNFRSF1A* | 4 | 146 | 10 | TNFRSF1A | TNF receptor superfamily | Plasma Membrane | transmembrane recepto | GSK2862277 |
| chr20 | *SLX4IP* | 4 | 114 | 10 | SLX4IP | SLX4 interacting protein | Other | other | |
| chr17 | *RP11-81A22.5* | 4 | 166 | 10 | | | | | |
| chr16 | *PIGQ* | 5 | 190 | 10 | PIGQ | phosphatidylinositol glycan anchor biosynthesis class Q | Cytoplasm | enzyme | |
| chr4 | *TNIP2* | 4 | 180 | 10 | TNIP2 | TNFAIP3 interacting protein 2 | Cytoplasm | other | |
| chr1 | *CENPL* | 4 | 143 | 10 | CENPL | centromere protein L | Cytoplasm | other | |
| chr16 | *HERC2P5* | 5 | 138 | 10 | HERC2P5 | hect domain and RLD 2 pseudogene 5 | Other | other | |
| chr16 | *CTD-2034I21.1* | 5 | 118 | 10 | | | | | |
| chr19 | *MCOLN1* | 5 | 193 | 10 | MCOLN1 | mucolipin 1 | Cytoplasm | ion channel | |
| chr3 | *FYTTD1* | 4 | 138 | 10 | FYTTD1 | forty-two-three domain containing | Nucleus | other | |
| chr16 | *MSLNL* | 5 | 187 | 10 | MSLNL | mesothelin like | Other | other | |
| chr8 | *EFR3A* | 4 | 103 | 10 | EFR3A | EFR3 homolog A | Plasma Membrane | other | |
| chr3 | *RP11-944L7.4* | 5 | 122 | 10 | | | | | |
| chr1 | *PEF1* | 4 | 116 | 10 | PEF1 | penta-EF-hand domain containing | Cytoplasm | other | |
| chr1 | *F5* | 5 | 140 | 10 | F5 | coagulation factor V | Extracellular Space | other | drotrecogin alfa, antithrombin alfa |
| chr1 | *PYHIN1* | 4 | 172 | 10 | PYHIN1 | pyrin and HIN domain family member 1 | Nucleus | other | |
| chr1 | *STX6* | 5 | 183 | 10 | STX6 | syntaxin 6 | Cytoplasm | transporter | |
| chr17 | *TTC19* | 5 | 104 | 10 | TTC19 | tetratricopeptide repeat domain | Cytoplasm | other | |
| chr1 | *GSTM4* | 4 | 216 | 10 | GSTM4 | glutathione S-transferase mu 4 | Cytoplasm | enzyme | |
| chr13 | *SIAH3* | 4 | 129 | 10 | SIAH3 | siah E3 ubiquitin protein ligase family member 3 | Cytoplasm | other | |
| chr12 | *TULP3* | 4 | 105 | 10 | TULP3 | TUB like protein 3 | Extracellular Space | other | |
| chr11 | *RASGRP2* | 4 | 175 | 10 | RASGRP2 | RAS guanyl releasing protein 2 | Cytoplasm | other | |
| chr1 | *C1orf189* | 5 | 153 | 10 | C1orf189 | chromosome 1 open reading frame 189 | Other | other | |
| chr19 | *CTB-102L5.4* | 4 | 152 | 10 | | | | | |
| chr2 | *STARD7* | 4 | 150 | 10 | STARD7 | StAR related lipid transfer domain containing 7 | Cytoplasm | other | |
| chr19 | *PDCD2L* | 5 | 111 | 10 | PDCD2L | programmed cell death 2 like | Other | other | |
| chr8 | *KB-1568E2.1* | 5 | 118 | 10 | | | | | |
| chr1 | *MUTYH* | 5 | 164 | 10 | MUTYH | mutY DNA glycosylase | Nucleus | enzyme | |
| chr2 | *KLF11* | 3 | 129 | 10 | KLF11 | Kruppel like factor 11 | Nucleus | transcription regulator | |
| chr11 | *CCDC153* | 4 | 138 | 10 | CCDC153 | coiled-coil domain containing 153 | Other | other | |
| chrX | *RP11-265P11.1* | 6 | 122 | 10 | | | | | |
| chr10 | *NODAL* | 6 | 224 | 10 | NODAL | nodal growth differentiation factor | Extracellular Space | growth factor | |
| chr10 | *MINPP1* | 5 | 106 | 10 | MINPP1 | multiple inositol-polyphosphate phosphatase 1 | Cytoplasm | phosphatase | |
| chr16 | *ECI1* | 5 | 292 | 10 | ECI1 | enoyl-CoA delta isomerase 1 | Cytoplasm | enzyme | |
| chr4 | *ANTXR2* | 4 | 138 | 10 | ANTXR2 | ANTXR cell adhesion molecule 2 | Plasma Membrane | transmembrane recepto | |
| chr12 | *A2ML1-AS1* | 4 | 84 | 10 | A2ML1-AS1 | A2ML1 antisense RNA 1 | Other | other | |
| chr5 | *CARD6* | 5 | 260 | 10 | CARD6 | caspase recruitment domain family member 6 | Cytoplasm | other | |
| chr11 | *C11orf16* | 4 | 205 | 10 | C11orf16 | chromosome 11 open reading frame 16 | Other | other | |
| chrX | *RP3-326I13.1* | 4 | 101 | 10 | | | | | |
| chr8 | *RP11-705O24.1* | 4 | 125 | 10 | | | | | |
| chr10 | *CCNYL2* | 5 | 184 | 10 | CCNYL2 | cyclin Y like 2 (pseudogene) | Other | other | |
| chr10 | *ECHS1* | 5 | 174 | 10 | ECHS1 | enoyl-CoA hydratase, short chain | Cytoplasm | enzyme | |
| chr12 | *RP11-417L19.2* | 5 | 213 | 10 | | | | | |

| | | | | | | | | | |
|---|---|---|---|---|---|---|---|---|---|
| chr8 | *CHRNA6* | 4 | 97 | 10 | CHRNA6 | cholinergic receptor nicotinic alpha 6 subunit | Plasma Membrane | transmembrane receptor | succinylcholine, rapacuronium, doxacurium, vecuronium, pipecuronium, metocurine, mivacurium, D-tubocurarine, nicotine, arecoline, pancuronium, atracurium, isoflurane, cisatracurium, rocuronium, mecamylamine |
| chr6 | *MCM3* | 4 | 121 | 10 | MCM3 | minichromosome maintenance complex component 3 | Nucleus | enzyme | |
| chr19 | *NCR1* | 4 | 137 | 10 | NCR1 | natural cytotoxicity triggering receptor 1 | Plasma Membrane | transmembrane receptor | |
| chr2 | *ILKAP* | 4 | 103 | 10 | ILKAP | ILK associated serine/threonine phosphatase | Cytoplasm | phosphatase | |
| chr8 | *FAM86B1* | 4 | 269 | 10 | FAM86B1 | family with sequence similarity 86 member B1 | Other | other | |
| chr4 | *CBR4* | 4 | 106 | 10 | CBR4 | carbonyl reductase 4 | Cytoplasm | enzyme | |
| chr3 | *POMGNT2* | 4 | 154 | 10 | POMGNT2 | protein O-linked mannose N-acetylglucosaminyltransferase 2 (beta 1,4-) | Cytoplasm | enzyme | |
| chr14 | *SLC25A29* | 5 | 184 | 10 | SLC25A29 | solute carrier family 25 member | Cytoplasm | transporter | |
| chr12 | *AMHR2* | 5 | 256 | 10 | AMHR2 | anti-Mullerian hormone receptor type 2 | Plasma Membrane | kinase | GM102 |
| chr7 | *DBF4* | 3 | 89 | 10 | DBF4 | DBF4 zinc finger | Nucleus | kinase | |
| chrX | *SYP* | 5 | 187 | 10 | SYP | synaptophysin | Cytoplasm | transporter | |
| chr17 | *PER1* | 4 | 202 | 10 | PER1 | period circadian regulator 1 | Nucleus | transcription regulator | avibactam |
| chr19 | *XAB2* | 5 | 242 | 10 | XAB2 | XPA binding protein 2 | Nucleus | other | |
| chr19 | *C19orf18* | 4 | 106 | 10 | C19orf18 | chromosome 19 open reading frame 18 | Cytoplasm | other | |
| chr2 | *C2orf73* | 5 | 124 | 10 | C2orf73 | chromosome 2 open reading frame 73 | Other | other | |
| chrX | *RP13-188A5.1* | 5 | 197 | 10 | | | | | |
| chr22 | *SLC35E4* | 5 | 159 | 10 | SLC35E4 | solute carrier family 35 member | Other | other | |
| chr16 | *RP11-82O18.2* | 5 | 121 | 10 | | | | | |
| chr1 | *RRAGC* | 4 | 97 | 10 | RRAGC | Ras related GTP binding C | Cytoplasm | enzyme | |
| chr1 | *RP5-936J12.1* | 5 | 129 | 10 | | | | | |
| chr16 | *RP11-161M6.3* | 5 | 192 | 10 | | | | | |
| chr11 | *RP11-111M22.2* | 5 | 121 | 10 | | | | | |
| chr2 | *C2orf71* | 4 | 144 | 10 | PCARE | photoreceptor cilium actin | Extracellular Space | other | |
| chr1 | *C1orf43* | 5 | 161 | 10 | C1orf43 | chromosome 1 open reading frame 43 | Other | other | |
| chr9 | *ARRDC1* | 5 | 295 | 10 | ARRDC1 | arrestin domain containing 1 | Cytoplasm | other | |
| chr12 | *ZFC3H1* | 4 | 92 | 10 | ZFC3H1 | zinc finger C3H1-type containing | Extracellular Space | other | |
| chr11 | *HPS5* | 4 | 100 | 10 | HPS5 | HPS5 biogenesis of lysosomal organelles complex 2 subunit 2 | Cytoplasm | other | |
| chr7 | *TRIP6* | 5 | 215 | 10 | TRIP6 | thyroid hormone receptor | Extracellular Space | cytokine | |
| chr4 | *MOB1B* | 4 | 89 | 10 | MOB1B | MOB kinase activator 1B | Cytoplasm | other | |
| chr12 | *GXYLT1* | 5 | 185 | 10 | GXYLT1 | glucoside xylosyltransferase 1 | Cytoplasm | enzyme | |
| chr1 | *RP11-354K1.1* | 4 | 199 | 10 | | | | | |
| chr9 | *RMI1* | 4 | 97 | 10 | RMI1 | RecQ mediated genome | Nucleus | other | |
| chr12 | *HOTAIR* | 5 | 342 | 10 | HOTAIR | HOX transcript antisense RNA | Other | other | |
| chr6 | *SCGN* | 4 | 117 | 10 | SCGN | secretagogin, EF-hand calcium binding protein | Cytoplasm | other | |
| chr4 | *RP11-775H9.2* | 4 | 144 | 10 | | | | | |
| chr11 | *C11orf84* | 4 | 172 | 10 | SPINDOC | spindlin interactor and repressor of chromatin binding | Other | other | |
| chr2 | *GAREML* | 5 | 181 | 10 | GAREM2 | GRB2 associated regulator of MAPK1 subtype 2 | Cytoplasm | other | |
| chr11 | *HRASLS2* | 4 | 138 | 10 | PLAAT2 | phospholipase A and acyltransferase 2 | Cytoplasm | enzyme | |
| chr7 | *ZNF680* | 4 | 122 | 10 | ZNF680 | zinc finger protein 680 | Nucleus | transcription regulator | |
| chr19 | *CNN1* | 5 | 172 | 10 | CNN1 | calponin 1 | Cytoplasm | other | |
| chr19 | *MAN2B1* | 5 | 177 | 10 | MAN2B1 | mannosidase alpha class 2B member 1 | Cytoplasm | enzyme | |
| chr19 | *AC004609.1* | 4 | 106 | 10 | | | | | |
| chr2 | *RTKN* | 5 | 178 | 10 | RTKN | rhotekin | Cytoplasm | other | |
| chr15 | *GOLGA8Q* | 4 | 149 | 10 | GOLGA8Q | golgin A8 family member Q | Other | other | |
| chr20 | *RP4-684O24.5* | 4 | 131 | 10 | | | | | |
| chr11 | *CEP57* | 4 | 195 | 10 | CEP57 | centrosomal protein 57 | Cytoplasm | other | |
| chr6 | *AL033381.1* | 4 | 143 | 10 | | | | | |
| chr3 | *CD80* | 5 | 130 | 10 | CD80 | CD80 molecule | Plasma Membrane | transmembrane receptor | belatacept, galiximab, abatacept, abatacept/methotrex |
| chr12 | *ENPP7P5* | 4 | 122 | 10 | ENPP7P5 | | Other | other | |
| chr20 | *LSM14B* | 5 | 342 | 10 | LSM14B | LSM family member 14B | Other | other | |

| | | | | | | | | | |
|---|---|---|---|---|---|---|---|---|---|
| chr3 | *TOP2B* | 4 | 104 | 10 | TOP2B | DNA topoisomerase II beta | Nucleus | enzyme | daunorubicin/tretinoin, chlorambucil/epirubicin, cyclophosphamide/epirubicin/5-fluorouracil/paclitaxel, cisplatin/doxorubicin/methotrexate, chlorambucil/mitoxantrone/prednisone/rituximab, AQ4N, cytarabine/etoposide/mitoxantrone, cytarabine/daunorubicin, clofarabine/cyclophosphamide/etoposide, cyclophosphamide/epirubicin, fludarabine phosphate/mitoxantrone/rituximab, [illegible] |
| chr19 | *CYP2B6* | 5 | 101 | 10 | CYP2B6 | cytochrome P450 family 2 subfamily B member 6 | Cytoplasm | enzyme | |
| chr5 | *RP11-93O7.5* | 4 | 90 | 10 | | | | | |
| chr16 | *DDX19A* | 5 | 166 | 10 | DDX19A | DEAD-box helicase 19A | Nucleus | enzyme | |
| chr6 | *FBXO9* | 4 | 120 | 10 | FBXO9 | F-box protein 9 | Cytoplasm | enzyme | |
| chr11 | *NEAT1* | 5 | 343 | 10 | NEAT1 | nuclear paraspeckle assembly transcript 1 | Other | other | |
| chr17 | *METRNL* | 5 | 201 | 10 | METRNL | meteorin like, glial cell differentiation regulator | Cytoplasm | other | |
| chr5 | *C1QTNF2* | 5 | 163 | 10 | C1QTNF2 | C1q and TNF related 2 | Extracellular Space | other | |
| chr19 | *SLC7A10* | 5 | 253 | 10 | SLC7A10 | solute carrier family 7 member 10 | Plasma Membrane | transporter | |
| chr6 | *TREM1* | 5 | 121 | 10 | TREM1 | triggering receptor expressed on myeloid cells 1 | Plasma Membrane | transmembrane recepto | anti-trem1 |
| chr19 | *THEG* | 5 | 163 | 10 | THEG | theg spermatid protein | Cytoplasm | other | |
| chrX | *SSX6* | 5 | 113 | 10 | SSX6P | SSX family member 6, | Other | other | |
| chr3 | *RP11-443P15.2* | 4 | 95 | 10 | | | | | |
| chr22 | *PNPLA5* | 4 | 168 | 10 | PNPLA5 | patatin like phospholipase domain containing 5 | Cytoplasm | enzyme | |
| chr11 | *RP11-466I1.1* | 4 | 162 | 10 | | | | | |
| chr2 | *CCDC148* | 4 | 147 | 10 | CCDC148 | coiled-coil domain containing 148 | Other | other | |
| chr15 | *GOLGA8I* | 4 | 188 | 10 | GOLGA8IP | golgin A8 family member I, pseudogene | Other | other | |
| chr17 | *TK1* | 5 | 179 | 10 | TK1 | thymidine kinase 1 | Cytoplasm | kinase | |
| chr5 | *AC004041.2* | 4 | 185 | 10 | | | | | |
| chr14 | *SIVA1* | 4 | 186 | 10 | SIVA1 | SIVA1 apoptosis inducing factor | Cytoplasm | other | |
| chr5 | *CSNK1A1* | 4 | 105 | 10 | CSNK1A1 | casein kinase 1 alpha 1 | Cytoplasm | kinase | |
| chr19 | *DDX39A* | 5 | 210 | 10 | DDX39A | DExD-box helicase 39A | Nucleus | enzyme | |
| chr11 | *ART1* | 5 | 233 | 10 | ART1 | ADP-ribosyltransferase 1 | Plasma Membrane | enzyme | |
| chr21 | *SYNJ1* | 3 | 77 | 10 | SYNJ1 | synaptojanin 1 | Cytoplasm | phosphatase | |
| chr2 | *MZT2A* | 4 | 148 | 10 | MZT2A | mitotic spindle organizing protein | Cytoplasm | other | |
| chr2 | *LOXL3* | 5 | 155 | 10 | LOXL3 | lysyl oxidase like 3 | Extracellular Space | enzyme | |
| chr1 | *RP11-139I14.2* | 4 | 280 | 10 | | | | | |
| chr3 | *GRM2* | 5 | 244 | 10 | GRM2 | glutamate metabotropic receptor 2 | Plasma Membrane | G-protein coupled recepto | talaglumetad, fasoracetam |
| chr10 | *NSMCE4A* | 4 | 113 | 10 | NSMCE4A | NSE4 homolog A, SMC5-SMC6 complex component | Nucleus | other | |
| chr1 | *MIIP* | 5 | 228 | 10 | MIIP | migration and invasion inhibitory protein | Other | other | |
| chr4 | *RP11-696N14.1* | 5 | 102 | 10 | | | | | |
| chr5 | *GPRIN1* | 5 | 199 | 10 | GPRIN1 | G protein regulated inducer of neurite outgrowth 1 | Plasma Membrane | other | |
| chr11 | *ZBED5* | 5 | 162 | 10 | ZBED5 | zinc finger BED-type containing 5 | Nucleus | transcription regulator | |
| chr2 | *AC012487.2* | 5 | 116 | 10 | | | | | |
| chr3 | *CTNNB1* | 4 | 91 | 10 | CTNNB1 | catenin beta 1 | Nucleus | transcription regulator | PRI-724 |
| chr19 | *HIF3A* | 4 | 140 | 10 | HIF3A | hypoxia inducible factor 3 subunit alpha | Nucleus | transcription regulator | |
| chr17 | *TNFAIP1* | 4 | 260 | 10 | TNFAIP1 | TNF alpha induced protein 1 | Plasma Membrane | ion channel | |
| chr6 | *MLN* | 4 | 210 | 10 | MLN | motilin | Extracellular Space | other | |
| chr5 | *RP11-1C1.4* | 5 | 429 | 10 | | | | | |
| chr12 | *PTPN6* | 5 | 194 | 10 | PTPN6 | protein tyrosine phosphatase non-receptor type 6 | Cytoplasm | phosphatase | SC-43 |
| chr19 | *ECSIT* | 4 | 166 | 10 | ECSIT | ECSIT signalling integrator | Nucleus | transcription regulator | |
| chr14 | *RP11-116N8.1* | 4 | 95 | 10 | | | | | |
| chr1 | *ZMYND12* | 4 | 135 | 10 | ZMYND12 | zinc finger MYND-type containing | Other | other | |
| chr1 | *FCGR2B* | 4 | 163 | 10 | FCGR2B | Fc fragment of IgG receptor IIb | Plasma Membrane | transmembrane recepto | BI-1206, IgG, XmAb5871 |
| chr18 | *CTD-2015H3.2* | 5 | 76 | 10 | | | | | |
| chr17 | *ARL5C* | 4 | 136 | 10 | ARL5C | ADP ribosylation factor like GTPase 5C | Other | other | |
| chr19 | *CTB-50L17.16* | 4 | 124 | 10 | | | | | |
| chr8 | *FAM90A18P* | 5 | 675 | 10 | | | | | |
| chr1 | *C1orf168* | 5 | 117 | 10 | FYB2 | FYN binding protein 2 | Plasma Membrane | other | |
| chr15 | *WDR73* | 5 | 151 | 10 | WDR73 | WD repeat domain 73 | Extracellular Space | other | |
| chr11 | *MRPL21* | 5 | 183 | 10 | MRPL21 | mitochondrial ribosomal protein | Cytoplasm | other | |
| chr3 | *VHL* | 5 | 135 | 10 | VHL | von Hippel-Lindau tumor | Nucleus | transcription regulator | |

| | | | | | | | | | |
|---|---|---|---|---|---|---|---|---|---|
| chr12 | *TMEM119* | 5 | 252 | 10 | TMEM119 | transmembrane protein 119 | Cytoplasm | other | |
| chr2 | *AC079610.1* | 4 | 102 | 10 | | | | | |
| chr9 | *XXyac-YRM2039.2* | 4 | 389 | 10 | | | | | |
| chr8 | *SULF1* | 3 | 88 | 10 | SULF1 | sulfatase 1 | Cytoplasm | enzyme | |
| chr20 | *PSMF1* | 5 | 116 | 10 | PSMF1 | proteasome inhibitor subunit 1 | Cytoplasm | other | |
| chr10 | *MBL1P* | 4 | 114 | 10 | MBL1P | mannose binding lectin 1, pseudogene | Other | other | |
| chr15 | *DLL4* | 4 | 226 | 10 | DLL4 | delta like canonical Notch ligand 4 | Extracellular Space | other | REGN421, OMP-305B83, OMP- |
| chr5 | *DCP2* | 4 | 117 | 10 | DCP2 | decapping mRNA 2 | Nucleus | enzyme | |
| chr12 | *RP11-1022B3.1* | 4 | 104 | 10 | | | | | |
| chr4 | *RP11-608O21.1* | 4 | 82 | 10 | | | | | |
| chr1 | *RP11-135J2.4* | 5 | 129 | 10 | | | | | |
| chr10 | *FAM25C* | 4 | 414 | 10 | M25A (includes oth | family with sequence similarity 25 member A | Other | other | |
| chr19 | *AC011523.2* | 4 | 111 | 10 | | | | | |
| chr10 | *PRDX3* | 5 | 127 | 10 | PRDX3 | peroxiredoxin 3 | Cytoplasm | enzyme | |
| chr7 | *AC004893.11* | 4 | 146 | 10 | | | | | |
| chr5 | *FAM81B* | 5 | 101 | 10 | FAM81B | family with sequence similarity 81 member B | Nucleus | other | |
| chr7 | *GPC2* | 5 | 205 | 10 | GPC2 | glypican 2 | Plasma Membrane | other | |
| chr6 | *SLC16A10* | 4 | 77 | 10 | SLC16A10 | solute carrier family 16 member | Plasma Membrane | transporter | |
| chr6 | *RP3-475N16.1* | 4 | 126 | 10 | | | | | |
| chr3 | *LINC00885* | 4 | 164 | 10 | LINC00885 | long intergenic non-protein coding RNA 885 | Other | other | |
| chr5 | *CTD-2235C13.2* | 5 | 94 | 10 | | | | | |
| chr1 | *SLC35E2* | 5 | 208 | 10 | SLC35E2A | solute carrier family 35 member | Other | other | |
| chr1 | *RNPC3* | 4 | 117 | 10 | RNPC3 | RNA binding region (RNP1, RRM) containing 3 | Nucleus | other | |
| chr10 | *CTGLF11P* | 4 | 207 | 10 | AGAP14P | ArfGAP with GTPase domain, ankyrin repeat and PH domain 14, pseudogene | Other | other | |
| chr19 | *FBL* | 5 | 144 | 10 | FBL | fibrillarin | Nucleus | enzyme | |
| chr5 | *TMEM161B* | 4 | 88 | 10 | TMEM161B | transmembrane protein 161B | Other | other | |
| chr11 | *MS4A13* | 5 | 152 | 10 | MS4A13 | membrane spanning 4-domains | Other | other | |
| chr16 | *RBL2* | 4 | 125 | 10 | RBL2 | RB transcriptional corepressor | Nucleus | other | |
| chr17 | *RP11-6N17.9* | 5 | 125 | 10 | | | | | |
| chr10 | *CREM* | 4 | 84 | 10 | CREM | cAMP responsive element | Nucleus | transcription regulator | |
| chr12 | *CACNB3* | 5 | 342 | 10 | CACNB3 | calcium voltage-gated channel auxiliary subunit beta 3 | Plasma Membrane | ion channel | verapamil, dexamethasone/rituximab/verapamil, mibefradil, nimodipine, |
| chr7 | *JAZF1-AS1* | 4 | 111 | 10 | JAZF1-AS1 | JAZF1 antisense RNA 1 | Other | other | |
| chr7 | *NYAP1* | 5 | 453 | 10 | NYAP1 | neuronal tyrosine phosphorylated phosphoinositide-3-kinase | Other | other | |
| chr5 | *CTD-2306M5.1* | 5 | 186 | 10 | | | | | |
| chr17 | *SMARCD2* | 5 | 231 | 10 | SMARCD2 | SWI/SNF related, matrix associated, actin dependent regulator of chromatin, subfamily | Nucleus | transcription regulator | |
| chr15 | *MESDC2* | 5 | 185 | 10 | MESD | mesoderm development LRP chaperone | Extracellular Space | other | |
| chr22 | *NAGA* | 4 | 123 | 10 | NAGA | alpha-N-acetylgalactosaminidase | Cytoplasm | enzyme | |
| chr12 | *RP11-726G1.1* | 3 | 84 | 10 | | | | | |
| chr10 | *NUTM2E* | 4 | 304 | 10 | NUTM2E | NUT family member 2E | Other | other | |
| chr1 | *TRAPPC3* | 4 | 133 | 10 | TRAPPC3 | trafficking protein particle | Cytoplasm | other | birabresib |
| chr8 | *ASAH1* | 4 | 110 | 10 | ASAH1 | N-acylsphingosine | Cytoplasm | enzyme | |
| chr3 | *RP11-135A1.3* | 4 | 132 | 10 | | | | | |
| chr2 | *AC016907.3* | 5 | 101 | 10 | | | | | |
| chr8 | *FGF17* | 5 | 232 | 10 | FGF17 | fibroblast growth factor 17 | Extracellular Space | growth factor | |
| chr2 | *AC105393.1* | 4 | 157 | 10 | | | | | |
| chr3 | *ZBTB47* | 5 | 293 | 10 | ZBTB47 | zinc finger and BTB domain containing 47 | Other | other | |
| chr5 | *AC016561.1* | 5 | 120 | 10 | | | | | |
| chr3 | *RBM5* | 4 | 121 | 10 | RBM5 | RNA binding motif protein 5 | Nucleus | other | |
| chr18 | *GAREM* | 4 | 88 | 10 | GAREM1 | GRB2 associated regulator of MAPK1 subtype 1 | Nucleus | other | |
| chr19 | *CEACAM18* | 5 | 143 | 10 | CEACAM18 | CEA cell adhesion molecule 18 | Other | other | |
| chrX | *U52111.14* | 4 | 198 | 10 | | | | | |
| chr10 | *APBB1IP* | 4 | 113 | 10 | APBB1IP | amyloid beta precursor protein binding family B member 1 interacting protein | Cytoplasm | other | |
| chr1 | *RP13-614K11.1* | 4 | 165 | 10 | | | | | |
| chr3 | *CRTAP* | 4 | 96 | 10 | CRTAP | cartilage associated protein | Extracellular Space | other | |
| chr10 | *PDSS1* | 4 | 98 | 10 | PDSS1 | decaprenyl diphosphate synthase subunit 1 | Cytoplasm | enzyme | |
| chr1 | *MEAF6* | 5 | 109 | 10 | MEAF6 | MYST/Esa1 associated factor 6 | Nucleus | other | |
| chr2 | *FAHD2B* | 5 | 184 | 10 | FAHD2B | fumarylacetoacetate hydrolase domain containing 2B | Cytoplasm | other | |
| chr8 | *PBK* | 3 | 125 | 10 | PBK | PDZ binding kinase | Cytoplasm | kinase | |
| chr10 | *TMEM26* | 4 | 134 | 10 | TMEM26 | transmembrane protein 26 | Other | other | |
| chr14 | *STON2* | 4 | 85 | 10 | STON2 | stonin 2 | Cytoplasm | other | |
| chr8 | *KB-1507C5.2* | 4 | 91 | 10 | | | | | |
| chr20 | *VPS16* | 4 | 112 | 10 | VPS16 | VPS16 core subunit of CORVET and HOPS complexes | Cytoplasm | transporter | |
| chr5 | *PIK3R1* | 5 | 109 | 10 | PIK3R1 | phosphoinositide-3-kinase regulatory subunit 1 | Cytoplasm | kinase | |
| chr5 | *CDC20B* | 4 | 90 | 10 | CDC20B | cell division cycle 20B | Other | other | |
| chr8 | *OC90* | 4 | 125 | 10 | OC90 | otoconin 90 | Extracellular Space | enzyme | |

| | | | | | | | | | |
|---|---|---|---|---|---|---|---|---|---|
| chr19 | *PSG2* | 5 | 120 | 10 | PSG2 | pregnancy specific beta-1-glycoprotein 2 | Extracellular Space | other | |
| chr1 | *C1orf100* | 4 | 92 | 10 | C1orf100 | chromosome 1 open reading frame 100 | Other | other | |
| chr3 | *SLC15A2* | 4 | 175 | 10 | SLC15A2 | solute carrier family 15 member 2 | Plasma Membrane | transporter | |
| chr1 | *MARC2* | 5 | 138 | 10 | MARC2 | mitochondrial amidoxime reducing component 2 | Cytoplasm | enzyme | |
| chr20 | *LINC00686* | 5 | 342 | 10 | LINC00686 | long intergenic non-protein coding RNA 686 | Other | other | |
| chr4 | *ALPK1* | 4 | 98 | 10 | ALPK1 | alpha kinase 1 | Cytoplasm | kinase | |
| chr1 | *GGPS1* | 4 | 163 | 10 | GGPS1 | geranylgeranyl diphosphate synthase 1 | Cytoplasm | enzyme | zoledronic acid |
| chr5 | *SLC23A1* | 4 | 215 | 10 | SLC23A1 | solute carrier family 23 member 1 | Plasma Membrane | transporter | |
| chr9 | *ZDHHC21* | 4 | 127 | 10 | ZDHHC21 | zinc finger DHHC-type containing | Plasma Membrane | enzyme | |
| chr16 | *ZSCAN10* | 5 | 218 | 10 | ZSCAN10 | zinc finger and SCAN domain containing 10 | Nucleus | transcription regulator | |
| chr6 | *MAP7* | 5 | 83 | 10 | MAP7 | microtubule associated protein 7 | Cytoplasm | other | |
| chr1 | *POLR3GL* | 4 | 140 | 10 | POLR3GL | RNA polymerase III subunit G | Nucleus | other | |
| chr2 | *AC022201.4* | 4 | 90 | 10 | | | | | |
| chr2 | *AC068282.3* | 4 | 148 | 10 | | | | | |
| chr7 | *CYP3A5* | 5 | 142 | 10 | CYP3A5 | cytochrome P450 family 3 subfamily A member 5 | Cytoplasm | enzyme | cobicistat, cobicistat/darunavir, cobicistat/elvitegravir/emtricitabine/tenofovir disoproxil, atazanavir/cobicistat |
| chr4 | *RP11-10L7.1* | 4 | 109 | 10 | | | | | |
| chr11 | *SLC22A24* | 4 | 97 | 10 | SLC22A24 | solute carrier family 22 member | Plasma Membrane | transporter | |
| chr18 | *RP11-15F12.1* | 3 | 151 | 10 | | | | | |
| chr19 | *APLP1* | 5 | 170 | 10 | APLP1 | amyloid beta precursor like | Extracellular Space | other | |
| chr1 | *AKR7A3* | 5 | 145 | 10 | AKR7A3 | aldo-keto reductase family 7 member A3 | Cytoplasm | enzyme | |
| chr8 | *C8orf44-SGK3* | 4 | 95 | 10 | 8orf44-SGK3/SGK | serum/glucocorticoid regulated kinase family member 3 | Cytoplasm | kinase | |
| chr1 | *ASPM* | 4 | 142 | 10 | ASPM | abnormal spindle microtubule assembly | Nucleus | other | |
| chr3 | *ATRIP* | 5 | 112 | 10 | ATRIP | ATR interacting protein | Nucleus | kinase | |
| chr5 | *SLU7* | 4 | 149 | 10 | SLU7 | SLU7 homolog, splicing factor | Nucleus | enzyme | |
| chr15 | *GOLGA8G* | 4 | 273 | 10 | OLGA8EP/GOLGA | golgin A8 family member G | Other | other | |
| chr7 | *RP11-667F9.1* | 5 | 120 | 10 | | | | | |
| chr15 | *C15orf59* | 5 | 144 | 10 | INSYN1 | inhibitory synaptic factor 1 | Plasma Membrane | other | |
| chr11 | *MOB2* | 5 | 225 | 10 | MOB2 | MOB kinase activator 2 | Nucleus | other | |
| chr17 | *ACBD4* | 5 | 195 | 10 | ACBD4 | acyl-CoA binding domain containing 4 | Other | other | |
| chr18 | *RP11-325K19.1* | 3 | 82 | 10 | | | | | |
| chr11 | *RP11-535A19.2* | 4 | 189 | 10 | | | | | |
| chr12 | *ATP5B* | 4 | 147 | 10 | ATP5F1B | ATP synthase F1 subunit beta | Cytoplasm | transporter | |
| chr7 | *CDCA7L* | 5 | 113 | 10 | CDCA7L | cell division cycle associated 7 | Nucleus | other | |
| chr9 | *RP11-307E17.8* | 4 | 92 | 10 | | | | | |
| chr12 | *SLC2A3* | 4 | 88 | 10 | SLC2A3 | solute carrier family 2 member 3 | Plasma Membrane | transporter | |
| chrX | *RP13-314C10.5* | 4 | 95 | 10 | | | | | |
| chr14 | *POTEG* | 4 | 156 | 10 | TEH (includes oth | POTE ankyrin domain family member G | Other | other | |
| chr14 | *BTBD7* | 4 | 85 | 10 | BTBD7 | BTB domain containing 7 | Nucleus | other | |
| chr19 | *KRTDAP* | 5 | 197 | 10 | KRTDAP | keratinocyte differentiation associated protein | Extracellular Space | other | |
| chr13 | *DCT* | 5 | 117 | 10 | DCT | dopachrome tautomerase | Cytoplasm | enzyme | |
| chr3 | *HGD* | 5 | 118 | 10 | HGD | homogentisate 1,2-dioxygenase | Cytoplasm | enzyme | |
| chr1 | *MATN1* | 4 | 201 | 10 | MATN1 | matrilin 1 | Extracellular Space | other | |
| chr21 | *ICOSLG* | 4 | 227 | 10 | OSLG/LOC102723 | inducible T cell costimulator | Plasma Membrane | other | |
| chr14 | *ARID4A* | 3 | 85 | 10 | ARID4A | AT-rich interaction domain 4A | Nucleus | transcription regulator | |
| chr1 | *BOLA1* | 4 | 133 | 10 | BOLA1 | bolA family member 1 | Cytoplasm | other | |
| chr5 | *UBE2B* | 4 | 103 | 10 | UBE2B | ubiquitin conjugating enzyme E2 | Cytoplasm | enzyme | |
| chr16 | *NUBP1* | 4 | 155 | 10 | NUBP1 | nucleotide binding protein 1 | Cytoplasm | other | |
| chr10 | *ASAH2B* | 4 | 144 | 10 | ASAH2B | N-acylsphingosine amidohydrolase 2B | Cytoplasm | other | |
| chr7 | *FAM71F2* | 4 | 91 | 10 | FAM71F2 | family with sequence similarity 71 member F2 | Other | other | |
| chr6 | *ACOT13* | 4 | 106 | 10 | ACOT13 | acyl-CoA thioesterase 13 | Cytoplasm | enzyme | |
| chr16 | *RLTPR* | 5 | 240 | 10 | CARMIL2 | capping protein regulator and myosin 1 linker 2 | Plasma Membrane | other | |
| chr16 | *SLC7A6OS* | 4 | 139 | 10 | SLC7A6OS | solute carrier family 7 member 6 opposite strand | Other | other | |

| | | | | | | | | | |
|---|---|---|---|---|---|---|---|---|---|
| chr15 | *SLC12A1* | 4 | 93 | 10 | SLC12A1 | solute carrier family 12 member 1 | Plasma Membrane | transporter | deserpidine/methyclothiazide, furosemide, bumetanide, hydroflumethiazide, ethacrynic acid, reserpine/trichloromethiazide, methyclothiazide, azilsartan medoxomil/chlorthalidone, chlorthalidone/reserpine, torsemide, chlorthalidone, hydroflumethiazide/reserpine, atenolol/chlorthalidone, quinethazone, chlorthalidone/clonidi |
| chr19 | *PRR19* | 5 | 207 | 10 | PRR19 | proline rich 19 | Other | other | |
| chr21 | *LINC00322* | 4 | 198 | 10 | LINC00322 | long intergenic non-protein coding RNA 322 | Other | other | |
| chr2 | *VAMP5* | 5 | 199 | 10 | VAMP5 | vesicle associated membrane protein 5 | Plasma Membrane | transporter | |
| chr11 | *HTR3A* | 4 | 146 | 10 | HTR3A | 5-hydroxytryptamine receptor 3A | Plasma Membrane | ion channel | netupitant/palonosetron, procaine, dexamethasone/granisetron, dexamethasone/netupitant/palonosetron, vortioxetine, dolasetron, ergotamine, alosetron, cisapride, granisetron, D-tubocurarine, aprepitant/granisetron, dexamethasone/granisetron/haloperidol, memantine, dexamethasone/ondansetron, dexamethasone/palonosetron, ondansetron, aprepitant/dexameth |
| chr2 | *SP140* | 5 | 132 | 10 | SP140 | SP140 nuclear body protein | Nucleus | transcription regulator | |
| chr19 | *ZNF283* | 5 | 91 | 10 | ZNF283 | zinc finger protein 283 | Nucleus | transcription regulator | |
| chr2 | *ADAM17* | 4 | 90 | 10 | ADAM17 | ADAM metallopeptidase domain | Plasma Membrane | peptidase | |
| chr4 | *RHOH* | 4 | 90 | 10 | RHOH | ras homolog family member H | Plasma Membrane | enzyme | |
| chr3 | *VGLL3* | 3 | 118 | 10 | VGLL3 | vestigial like family member 3 | Other | other | |
| chr12 | *CCDC38* | 5 | 132 | 10 | CCDC38 | coiled-coil domain containing 38 | Cytoplasm | other | |
| chr16 | *NPIPA7* | 4 | 117 | 10 | PA1 (includes oth | nuclear pore complex interacting protein family member A3 | Nucleus | other | |
| chr6 | *RP11-505P4.7* | 4 | 91 | 10 | | | | | |
| chr11 | *RP11-839D17.3* | 4 | 101 | 10 | | | | | |
| chr16 | *MYLPF* | 4 | 155 | 10 | MYLPF | myosin light chain, phosphorylatable, fast skeletal | Cytoplasm | other | |
| chr12 | *RERG* | 4 | 76 | 10 | RERG | RAS like estrogen regulated growth inhibitor | Nucleus | enzyme | |
| chr22 | *TCN2* | 4 | 123 | 10 | TCN2 | transcobalamin 2 | Extracellular Space | transporter | |
| chr5 | *CLK4* | 4 | 172 | 10 | CLK4 | CDC like kinase 4 | Nucleus | kinase | |
| chr7 | *LINC00972* | 5 | 145 | 10 | LINC00972 | | Other | other | |
| chr6 | *FIG4* | 4 | 110 | 10 | FIG4 | FIG4 phosphoinositide 5-phosphatase | Cytoplasm | enzyme | |
| chr13 | *VWA8* | 3 | 94 | 10 | VWA8 | von Willebrand factor A domain containing 8 | Cytoplasm | enzyme | |
| chr19 | *FXYD5* | 5 | 161 | 10 | FXYD5 | FXYD domain containing ion transport regulator 5 | Plasma Membrane | ion channel | |

| | | | | | | | | | |
|---|---|---|---|---|---|---|---|---|---|
| chr22 | *TUBA8* | 3 | 111 | 10 | TUBA8 | tubulin alpha 8 | Cytoplasm | other | docetaxel/gemcitabine/vincristine, cyclophosphamide/epirubicin/5-fluorouracil/paclitaxel, bevacizumab/paclitaxel/topotecan, colchicine/probenecid, bevacizumab/docetaxel/prednisone/thalidomide/zoledronic acid, gemcitabine/paclitaxel, L-asparaginase/daunorubicin/dexamethasone/vincristine, prednisone/vincristine, docetaxel/5-fluorouracil, vinflunine, L-[illegible] |
| chr3 | *NT5DC2* | 5 | 227 | 10 | NT5DC2 | 5'-nucleotidase domain | Cytoplasm | other | |
| chr2 | *AC067960.1* | 5 | 240 | 10 | | | | | |
| chr15 | *SPATA31E3P* | 4 | 365 | 10 | SPATA31E3P | SPATA31 subfamily E member 3, pseudogene | Other | other | |
| chr19 | *KRI1* | 5 | 175 | 10 | KRI1 | KRI1 homolog | Nucleus | other | |
| chr2 | *AC079613.1* | 3 | 88 | 10 | | | | | |
| chrX | *GPKOW* | 4 | 141 | 10 | GPKOW | G-patch domain and KOW motifs | Nucleus | other | |
| chr11 | *MRVI1-AS1* | 4 | 217 | 10 | MRVI1-AS1 | MRVI1 antisense RNA 1 | Other | other | |
| chr16 | *HN1L* | 4 | 154 | 10 | JPT2 | Jupiter microtubule associated homolog 2 | Cytoplasm | other | |
| chr1 | *SKINTL* | 5 | 152 | 10 | SKINT1L | Skint1 like (pseudogene) | Other | other | |
| chr22 | *BCRP4* | 4 | 185 | 10 | BCRP4 | BCR pseudogene 4 | Other | other | |
| chr16 | *RP11-598D12.2* | 5 | 150 | 10 | | | | | |
| chr17 | *KRT17P1* | 4 | 302 | 10 | KRT17P1 | keratin 17 pseudogene 1 | Other | other | |
| chr6 | *RP11-631F7.1* | 4 | 85 | 10 | | | | | |
| chr11 | *AIP* | 5 | 189 | 10 | AIP | aryl hydrocarbon receptor interacting protein | Nucleus | transcription regulator | |
| chrX | *PPP2R3B* | 5 | 209 | 10 | PPP2R3B | protein phosphatase 2 regulatory subunit B''beta | Nucleus | phosphatase | |
| chr7 | *STK17A* | 4 | 112 | 10 | STK17A | serine/threonine kinase 17a | Nucleus | kinase | |
| chr16 | *SPATA33* | 4 | 167 | 10 | SPATA33 | spermatogenesis associated 33 | Other | other | |
| chr11 | *SNX19* | 5 | 151 | 10 | SNX19 | sorting nexin 19 | Cytoplasm | transporter | |
| chr5 | *RP11-91H12.4* | 4 | 220 | 10 | | | | | |
| chr16 | *RP11-65J21.1* | 5 | 191 | 10 | | | | | |
| chr19 | *FAM83E* | 5 | 188 | 10 | FAM83E | family with sequence similarity 83 member E | Other | other | |
| chr1 | *ADAMTS4* | 5 | 187 | 10 | ADAMTS4 | ADAM metallopeptidase with thrombospondin type 1 motif 4 | Extracellular Space | peptidase | |
| chr8 | *FAM90A3P* | 4 | 361 | 10 | 90A26 (includes o | family with sequence similarity 90 member A26 | Other | other | |
| chr1 | *LDLRAD1* | 4 | 180 | 10 | LDLRAD1 | low density lipoprotein receptor class A domain containing 1 | Other | other | |
| chr2 | *C2orf91* | 5 | 144 | 10 | C2orf91 | chromosome 2 open reading frame 91 | Other | other | |
| chr5 | *IRF1* | 4 | 158 | 10 | IRF1 | interferon regulatory factor 1 | Nucleus | transcription regulator | |
| chr8 | *SFTPC* | 5 | 228 | 10 | SFTPC | surfactant protein C | Extracellular Space | other | |
| chr2 | *AC018717.1* | 4 | 106 | 10 | | | | | |
| chr12 | *CLEC4C* | 3 | 79 | 10 | CLEC4C | C-type lectin domain family 4 member C | Plasma Membrane | other | BIIB059 |
| chr1 | *CTSE* | 5 | 217 | 10 | CTSE | cathepsin E | Cytoplasm | peptidase | |
| chr3 | *RAB5A* | 3 | 111 | 10 | RAB5A | RAB5A, member RAS oncogene family | Cytoplasm | enzyme | |
| chr7 | *AC010091.1* | 4 | 93 | 10 | | | | | |
| chr21 | *SUMO3* | 5 | 159 | 10 | SUMO3 | small ubiquitin like modifier 3 | Nucleus | other | |
| chr19 | *ILVBL* | 4 | 199 | 10 | ILVBL | ilvB acetolactate synthase like | Cytoplasm | enzyme | |
| chr3 | *ZNF621* | 4 | 90 | 10 | ZNF621 | zinc finger protein 621 | Nucleus | transcription regulator | |
| chrX | *FTSJ1* | 4 | 198 | 10 | FTSJ1 | FtsJ RNA 2'-O-methyltransferase | Cytoplasm | enzyme | |
| chr10 | *YME1L1* | 3 | 85 | 10 | YME1L1 | YME1 like 1 ATPase | Cytoplasm | peptidase | |
| chr1 | *CRTC2* | 4 | 217 | 10 | CRTC2 | CREB regulated transcription coactivator 2 | Nucleus | other | |
| chr11 | *ARFGAP2* | 4 | 360 | 10 | ARFGAP2 | ADP ribosylation factor GTPase activating protein 2 | Nucleus | other | |
| chr19 | *INSL3* | 4 | 152 | 10 | INSL3 | insulin like 3 | Extracellular Space | growth factor | |
| chr20 | *SLC35C2* | 5 | 237 | 10 | SLC35C2 | solute carrier family 35 member | Cytoplasm | transporter | |
| chr17 | *FKBP10* | 5 | 238 | 10 | FKBP10 | FKBP prolyl isomerase 10 | Cytoplasm | enzyme | |
| chr6 | *KPNA5* | 4 | 113 | 10 | KPNA5 | karyopherin subunit alpha 5 | Cytoplasm | other | |
| chr19 | *TOMM40* | 4 | 176 | 10 | TOMM40 | translocase of outer mitochondrial membrane 40 | Cytoplasm | ion channel | |
| chr2 | *AC007403.2* | 5 | 93 | 10 | | | | | |
| chr2 | *AC099344.2* | 3 | 96 | 10 | | | | | |
| chr22 | *GGT1* | 4 | 177 | 10 | GGT1 | gamma-glutamyltransferase 1 | Plasma Membrane | enzyme | |
| chr1 | *RP11-134G8.7* | 4 | 165 | 10 | | | | | |
| chr19 | *RP11-420K14.8* | 4 | 82 | 10 | | | | | |
| chr5 | *SEMA6A* | 5 | 138 | 10 | SEMA6A | semaphorin 6A | Plasma Membrane | transmembrane receptor | |

| | | | | | | | | |
|---|---|---|---|---|---|---|---|---|
| chr8 | *FAM110B* | 4 | 97 | 10 | FAM110B | family with sequence similarity 110 member B | Cytoplasm | other |
| chr17 | *AC139099.6* | 4 | 215 | 10 | | | | |
| chr11 | *RP11-379J13.2* | 4 | 205 | 10 | | | | |
| chrX | *MOSPD1* | 3 | 91 | 10 | MOSPD1 | motile sperm domain containing 1 | Other | other |
| chr5 | *SLCO4C1* | 4 | 119 | 10 | SLCO4C1 | solute carrier organic anion transporter family member 4C1 | Plasma Membrane | transporter |
| chr13 | *RCBTB1* | 4 | 100 | 10 | RCBTB1 | RCC1 and BTB domain containing protein 1 | Nucleus | other |
| chr16 | *BAIAP3* | 5 | 172 | 10 | BAIAP3 | BAI1 associated protein 3 | Extracellular Space | other |
| chr15 | *DIS3L* | 4 | 96 | 10 | DIS3L | DIS3 like exosome 3'-5' exoribonuclease | Cytoplasm | enzyme |
| chr19 | *COMP* | 5 | 182 | 10 | COMP | cartilage oligomeric matrix protein | Extracellular Space | other |
| chr21 | *AF064858.6* | 4 | 135 | 10 | | | | |
| chr2 | *TRIM54* | 4 | 112 | 10 | TRIM54 | tripartite motif containing 54 | Cytoplasm | other |
| chr20 | *DEFB119* | 5 | 154 | 10 | DEFB119 | defensin beta 119 | Extracellular Space | other |
| chr10 | *RP11-432J9.6* | 5 | 254 | 10 | | | | |
| chr4 | *ANXA10* | 4 | 83 | 10 | ANXA10 | annexin A10 | Cytoplasm | other |
| chr9 | *DMRT3* | 4 | 124 | 10 | DMRT3 | doublesex and mab-3 related transcription factor 3 | Nucleus | transcription regulator |
| chr2 | *YIPF4* | 4 | 101 | 10 | YIPF4 | Yip1 domain family member 4 | Cytoplasm | other |
| chr2 | *ZNF142* | 4 | 161 | 10 | ZNF142 | zinc finger protein 142 | Nucleus | transcription regulator |
| chr7 | *POM121B* | 5 | 246 | 10 | POM121B | POM121 transmembrane nucleoporin B (pseudogene) | Other | other |
| chr19 | *ZNF677* | 4 | 90 | 10 | ZNF677 | zinc finger protein 677 | Other | other |
| chr17 | *SLC25A10* | 5 | 267 | 10 | SLC25A10 | solute carrier family 25 member | Cytoplasm | transporter |
| chr11 | *CCDC84* | 4 | 113 | 10 | CCDC84 | coiled-coil domain containing 84 | Other | other |
| chr1 | *RP11-70P17.1* | 4 | 163 | 10 | | | | |
| chr10 | *DHX32* | 4 | 134 | 10 | DHX32 | DEAH-box helicase 32 (putative) | Other | enzyme |
| chr16 | *PLA2G15* | 5 | 152 | 10 | PLA2G15 | phospholipase A2 group XV | Cytoplasm | enzyme |
| chr16 | *PKMYT1* | 5 | 257 | 10 | PKMYT1 | protein kinase, membrane associated tyrosine/threonine 1 | Cytoplasm | kinase |
| chr2 | *AC013269.5* | 3 | 174 | 10 | | | | |
| chr4 | *RP11-115D19.1* | 4 | 88 | 10 | | | | |
| chr12 | *RP11-609L23.1* | 3 | 126 | 10 | | | | |
| chr17 | *ZNF287* | 4 | 90 | 10 | ZNF287 | zinc finger protein 287 | Nucleus | transcription regulator |
| chr19 | *LILRA3* | 5 | 148 | 10 | LILRA3 | leukocyte immunoglobulin like receptor A3 | Extracellular Space | other |
| chr17 | *NLE1* | 4 | 230 | 10 | NLE1 | notchless homolog 1 | Nucleus | enzyme |
| chr16 | *CHMP1A* | 5 | 195 | 10 | CHMP1A | charged multivesicular body protein 1A | Extracellular Space | peptidase |
| chr15 | *RASGRP1* | 5 | 79 | 10 | RASGRP1 | RAS guanyl releasing protein 1 | Cytoplasm | other |
| chr17 | *NR1D1* | 4 | 240 | 10 | NR1D1 | nuclear receptor subfamily 1 group D member 1 | Nucleus | d-dependent nuclear rec |
| chr1 | *CFHR2* | 4 | 114 | 10 | CFHR2 | complement factor H related 2 | Extracellular Space | other |
| chr1 | *RP4-663N10.1* | 5 | 168 | 10 | | | | |
| chr6 | *BYSL* | 4 | 144 | 10 | BYSL | bystin like | Nucleus | other |
| chr3 | *LEPREL1* | 4 | 76 | 10 | P3H2 | prolyl 3-hydroxylase 2 | Cytoplasm | enzyme |
| chr2 | *AC104820.2* | 4 | 78 | 10 | | | | |
| chr2 | *SPOPL* | 3 | 84 | 10 | SPOPL | speckle type BTB/POZ protein | Other | other |
| chr1 | *RER1* | 4 | 214 | 10 | RER1 | retention in endoplasmic reticulum sorting receptor 1 | Cytoplasm | other |
| chr5 | *C5orf50* | 4 | 110 | 10 | SMIM23 | small integral membrane protein | Other | other |
| chr5 | *SKP1* | 3 | 98 | 10 | SKP1 | S-phase kinase associated | Nucleus | transcription regulator |
| chr2 | *R3HDM1* | 4 | 75 | 10 | R3HDM1 | R3H domain containing 1 | Other | other |
| chr14 | *PELI2* | 4 | 93 | 10 | PELI2 | pellino E3 ubiquitin protein ligase family member 2 | Cytoplasm | enzyme |
| chr8 | *GDAP1* | 4 | 96 | 10 | GDAP1 | ganglioside induced differentiation associated protein | Cytoplasm | other |
| chr3 | *SPINK8* | 5 | 91 | 10 | SPINK8 | serine peptidase inhibitor, Kazal type 8 (putative) | Other | other |
| chr10 | *LZTS2* | 5 | 281 | 10 | LZTS2 | leucine zipper tumor suppressor | Cytoplasm | other |
| chr7 | *C7orf72* | 5 | 121 | 10 | SPATA48 | spermatogenesis associated 48 | Other | other |
| chr17 | *TMEM235* | 5 | 180 | 10 | TMEM235 | transmembrane protein 235 | Other | other |
| chr16 | *RP11-150D5.2* | 4 | 91 | 10 | | | | |
| chr18 | *IL9RP4* | 4 | 279 | 10 | IL9RP4 | interleukin 9 receptor | Other | other |
| chr5 | *CDH6* | 5 | 267 | 10 | CDH6 | cadherin 6 | Plasma Membrane | other |
| chr15 | *RPL9P9* | 3 | 150 | 10 | | | | |
| chr16 | *UQCRC2* | 3 | 113 | 10 | UQCRC2 | ubiquinol-cytochrome c reductase core protein 2 | Cytoplasm | enzyme |
| chr15 | *CTD-2536I1.1* | 4 | 116 | 10 | | | | |
| chr8 | *RP11-369K17.1* | 4 | 93 | 10 | | | | |
| chr10 | *BBIP1* | 4 | 127 | 10 | BBIP1 | BBSome interacting protein 1 | Cytoplasm | other |
| chr19 | *ZNF724P* | 4 | 120 | 10 | ZNF724 | zinc finger protein 724 | Other | other |
| chr1 | *NBPF18P* | 5 | 152 | 10 | NBPF18P | NBPF member 18, pseudogene | Other | other |
| chr19 | *STK11* | 4 | 162 | 10 | STK11 | serine/threonine kinase 11 | Cytoplasm | kinase |

| | | | | | | | | | |
|---|---|---|---|---|---|---|---|---|---|
| chr14 | *PSMB5* | 4 | 114 | 10 | PSMB5 | proteasome subunit beta 5 | Cytoplasm | peptidase | bortezomib/dexamethasone/rituximab, bortezomib/thalidomide, ixazomib citrate, bortezomib/cladribine/rituximab, bortezomib/dexamethasone/pomalidomide, bortezomib/doxorubicin, bortezomib/dexamethasone/thalidomide, bortezomib/paclitaxel, bortezomib/fulvestrant, bortezomib/rituximab, bortezomib/sorafenib, carfilzomib, bortezomib/lenalidomide, carfilzomib/dexameth[illegible] |
| chr13 | *DGKH* | 4 | 83 | 10 | DGKH | diacylglycerol kinase eta | Cytoplasm | kinase | |
| chr9 | *C9orf91* | 4 | 122 | 10 | TMEM268 | transmembrane protein 268 | Cytoplasm | other | |
| chr6 | *BACH2* | 4 | 103 | 10 | BACH2 | BTB domain and CNC homolog 2 | Nucleus | transcription regulator | |
| chr8 | *SPATC1* | 5 | 156 | 10 | SPATC1 | spermatogenesis and centriole associated 1 | Cytoplasm | other | |
| chr7 | *ALKBH4* | 5 | 162 | 10 | ALKBH4 | alkB homolog 4, lysine | Cytoplasm | enzyme | |
| chr17 | *ICT1* | 5 | 115 | 10 | MRPL58 | mitochondrial ribosomal protein | Cytoplasm | enzyme | |
| chr8 | *CTD-2281E23.1* | 4 | 144 | 10 | | | | | |
| chr8 | *POMK* | 4 | 144 | 10 | POMK | protein O-mannose kinase | Cytoplasm | kinase | |
| chr7 | *RP11-212P7.2* | 5 | 135 | 10 | | | | | |
| chr2 | *DNAJC27-AS1* | 4 | 87 | 10 | DNAJC27-AS1 | DNAJC27 antisense RNA 1 | Other | other | |
| chr12 | *RP11-415I12.3* | 5 | 107 | 10 | | | | | |
| chr7 | *XRCC2* | 4 | 84 | 10 | XRCC2 | X-ray repair cross complementing | Nucleus | enzyme | |
| chr11 | *BATF2* | 4 | 195 | 10 | BATF2 | basic leucine zipper ATF-like transcription factor 2 | Other | other | |
| chr3 | *KIAA2018* | 4 | 121 | 10 | USF3 | upstream transcription factor family member 3 | Other | other | |
| chr3 | *PLCXD2* | 5 | 163 | 10 | PLCXD2 | phosphatidylinositol specific phospholipase C X domain containing 2 | Other | enzyme | |
| chr2 | *LINC00211* | 5 | 121 | 10 | LINC00211 | long intergenic non-protein coding RNA 211 | Other | other | |
| chr15 | *TBC1D21* | 4 | 124 | 10 | TBC1D21 | TBC1 domain family member 21 | Cytoplasm | other | |
| chr7 | *TSGA13* | 4 | 93 | 10 | TSGA13 | testis specific 13 | Other | other | |
| chr20 | *CD40* | 4 | 219 | 10 | CD40 | CD40 molecule | Plasma Membrane | transmembrane receptor | ABBV-927, APX005M, CP-870893, CFZ533, CDX-1140, SEA-CD40, FFP104, dacetuzumab, |
| chr9 | *CBWD3* | 5 | 123 | 10 | CBWD3/CBWD6 | COBW domain containing 3 | Other | other | |
| chr1 | *TTC24* | 4 | 255 | 10 | TTC24 | tetratricopeptide repeat domain | Other | other | |
| chr10 | *AL022344.4* | 5 | 144 | 10 | | | | | |
| chr1 | *RIT1* | 4 | 145 | 10 | RIT1 | Ras like without CAAX 1 | Plasma Membrane | enzyme | |
| chr19 | *ZNF383* | 4 | 107 | 10 | ZNF383 | zinc finger protein 383 | Nucleus | transcription regulator | |
| chr1 | *RP11-353N4.6* | 4 | 134 | 10 | | | | | |
| chr15 | *RP11-578F21.4* | 4 | 172 | 10 | | | | | |
| chr19 | *LINGO3* | 5 | 191 | 10 | LINGO3 | leucine rich repeat and Ig domain containing 3 | Other | other | |
| chrX | *WDR13* | 5 | 257 | 10 | WDR13 | WD repeat domain 13 | Nucleus | other | |
| chr18 | *RP11-267C16.1* | 3 | 89 | 9 | | | | | |
| chr12 | *LINC00937* | 4 | 112 | 9 | LINC00937 | long intergenic non-protein coding RNA 937 | Other | other | |
| chr1 | *SAMD11* | 5 | 245 | 9 | SAMD11 | sterile alpha motif domain containing 11 | Nucleus | other | |
| chr7 | *RP11-196D18.5* | 4 | 152 | 9 | | | | | |
| chr11 | *CTSC* | 4 | 115 | 9 | CTSC | cathepsin C | Cytoplasm | peptidase | |
| chr17 | *MNT* | 5 | 268 | 9 | MNT | MAX network transcriptional repressor | Nucleus | transcription regulator | |
| chr8 | *TTI2* | 5 | 109 | 9 | TTI2 | TELO2 interacting protein 2 | Cytoplasm | other | |
| chr19 | *ZNF569* | 5 | 124 | 9 | ZNF569 | zinc finger protein 569 | Nucleus | transcription regulator | |
| chr8 | *ADAM9* | 4 | 84 | 9 | ADAM9 | ADAM metallopeptidase domain | Plasma Membrane | peptidase | |
| chr17 | *CCDC137* | 4 | 181 | 9 | CCDC137 | coiled-coil domain containing 137 | Nucleus | other | |
| chr17 | *AURKB* | 4 | 192 | 9 | AURKB | aurora kinase B | Nucleus | kinase | chiauranib, ilorasertib, SNS 314, AT-9283, SKI-G-801, TTP607, CYC 116, tozasertib, AMG 900, danusertib, GSK1070916, |
| chr1 | *C1orf54* | 4 | 132 | 9 | C1orf54 | chromosome 1 open reading frame 54 | Other | other | |
| chr12 | *CDK2AP1* | 5 | 193 | 9 | CDK2AP1 | cyclin dependent kinase 2 associated protein 1 | Nucleus | other | |
| chr6 | *GTPBP2* | 4 | 165 | 9 | GTPBP2 | GTP binding protein 2 | Extracellular Space | enzyme | |

| | | | | | | | | | |
|---|---|---|---|---|---|---|---|---|---|
| chr8 | *SNTB1* | 3 | 96 | 9 | SNTB1 | syntrophin beta 1 | Plasma Membrane | other | |
| chr8 | *PDLIM2* | 5 | 293 | 9 | PDLIM2 | PDZ and LIM domain 2 | Cytoplasm | other | |
| chr15 | *ONECUT1* | 5 | 179 | 9 | ONECUT1 | one cut homeobox 1 | Nucleus | transcription regulator | |
| chr17 | *SPAG7* | 4 | 144 | 9 | SPAG7 | sperm associated antigen 7 | Nucleus | other | |
| chr1 | *MIB2* | 5 | 208 | 9 | MIB2 | mindbomb E3 ubiquitin protein ligase 2 | Nucleus | transcription regulator | |
| chr18 | *LIPG* | 4 | 125 | 9 | LIPG | lipase G, endothelial type | Extracellular Space | enzyme | |
| chr7 | *WBSCR27* | 5 | 227 | 9 | METTL27 | methyltransferase like 27 | Other | enzyme | |
| chr11 | *DNAJC24* | 4 | 94 | 9 | DNAJC24 | DnaJ heat shock protein family (Hsp40) member C24 | Other | other | |
| chrX | *TMEM187* | 4 | 116 | 9 | TMEM187 | transmembrane protein 187 | Cytoplasm | other | |
| chr13 | *PHF11* | 3 | 85 | 9 | PHF11 | PHD finger protein 11 | Nucleus | other | |
| chr15 | *GOLGA6B* | 4 | 247 | 9 | GA6A (includes ot | golgin A6 family member A | Other | other | |
| chr17 | *MIEF2* | 5 | 212 | 9 | MIEF2 | mitochondrial elongation factor 2 | Cytoplasm | other | |
| chr8 | *CCAR2* | 4 | 160 | 9 | CCAR2 | cell cycle and apoptosis regulator | Cytoplasm | peptidase | |
| chr20 | *BFSP1* | 4 | 128 | 9 | BFSP1 | beaded filament structural protein | Cytoplasm | enzyme | |
| chr14 | *MTHFD1* | 3 | 94 | 9 | MTHFD1 | methylenetetrahydrofolate dehydrogenase, cyclohydrolase and formyltetrahydrofolate | Cytoplasm | enzyme | |
| chr19 | *CYP2G2P* | 4 | 113 | 9 | CYP2G2P | cytochrome P450 family 2 subfamily G member 2, | Other | other | |
| chr1 | *FMOD* | 4 | 180 | 9 | FMOD | fibromodulin | Extracellular Space | other | |
| chrX | *NUP62CL* | 3 | 101 | 9 | NUP62CL | nucleoporin 62 C-terminal like | Other | other | |
| chr7 | *PON1* | 4 | 108 | 9 | PON1 | paraoxonase 1 | Extracellular Space | phosphatase | |
| chr14 | *ACIN1* | 3 | 113 | 9 | ACIN1 | apoptotic chromatin condensation inducer 1 | Nucleus | enzyme | |
| chr6 | *C6orf195* | 4 | 186 | 9 | LINC01600 | long intergenic non-protein coding RNA 1600 | Other | other | |
| chrX | *MSL3* | 4 | 124 | 9 | MSL3 | MSL complex subunit 3 | Nucleus | transcription regulator | |
| chr12 | *ING4* | 4 | 122 | 9 | ING4 | inhibitor of growth family member | Nucleus | transcription regulator | |
| chr3 | *POGLUT1* | 4 | 103 | 9 | POGLUT1 | protein O-glucosyltransferase 1 | Extracellular Space | enzyme | |
| chr19 | *TNPO2* | 4 | 190 | 9 | TNPO2 | transportin 2 | Nucleus | transporter | |
| chr9 | *ANAPC2* | 5 | 222 | 9 | ANAPC2 | anaphase promoting complex subunit 2 | Nucleus | other | |
| chr1 | *SH3GLB1* | 4 | 75 | 9 | SH3GLB1 | SH3 domain containing GRB2 like, endophilin B1 | Cytoplasm | enzyme | |
| chr4 | *EPHA5* | 4 | 108 | 9 | EPHA5 | EPH receptor A5 | Plasma Membrane | kinase | |
| chr4 | *RASGEF1B* | 4 | 98 | 9 | RASGEF1B | RasGEF domain family member | Other | other | |
| chr6 | *RNF217* | 3 | 81 | 9 | RNF217 | ring finger protein 217 | Cytoplasm | enzyme | |
| chr16 | *NPIPB15* | 4 | 106 | 9 | PB15 (includes oth | nuclear pore complex interacting protein family member B15 | Nucleus | other | |
| chr4 | *SLC30A9* | 4 | 102 | 9 | SLC30A9 | solute carrier family 30 member 9 | Nucleus | transporter | |
| chr8 | *MAFA* | 5 | 164 | 9 | MAFA | MAF bZIP transcription factor A | Nucleus | transcription regulator | |
| chr3 | *C3orf62* | 4 | 93 | 9 | C3orf62 | chromosome 3 open reading frame 62 | Other | other | |
| chr13 | *ATP4B* | 5 | 153 | 9 | ATP4B | ATPase H+/K+ transporting subunit beta | Plasma Membrane | transporter | ilaprazole, esomeprazole magnesium, aspirin/omeprazole, tenatoprazole, clarithromycin/omeprazole, lansoprazole, AR-H047108, diclofenac/omeprazole, dexlansoprazole, AGN 201904, magnesium hydroxide/omeprazole/sodium bicarbonate, omeprazole/sodium bicarbonate, pantoprazole, esomeprazole/napro |
| chr14 | *TXNDC16* | 4 | 103 | 9 | TXNDC16 | thioredoxin domain containing 16 | Extracellular Space | enzyme | |
| chr1 | *ELTD1* | 4 | 73 | 9 | ADGRL4 | adhesion G protein-coupled receptor L4 | Plasma Membrane | i-protein coupled recepto | |
| chr11 | *TRAF6* | 3 | 71 | 9 | TRAF6 | TNF receptor associated factor 6 | Cytoplasm | enzyme | |
| chr16 | *SNAI3-AS1* | 5 | 174 | 9 | SNAI3-AS1 | SNAI3 antisense RNA 1 | Other | other | |
| chr2 | *SUCLG1* | 4 | 118 | 9 | SUCLG1 | succinate-CoA ligase alpha | Cytoplasm | enzyme | |
| chr7 | *MIOS* | 4 | 96 | 9 | MIOS | meiosis regulator for oocyte development | Cytoplasm | other | |
| chr1 | *TTC22* | 5 | 169 | 9 | TTC22 | tetratricopeptide repeat domain | Other | other | |
| chr3 | *C3orf38* | 3 | 99 | 9 | C3orf38 | chromosome 3 open reading frame 38 | Nucleus | peptidase | |
| chr16 | *AC009120.6* | 4 | 157 | 9 | | | | | |
| chr2 | *KHK* | 4 | 157 | 9 | KHK | ketohexokinase | Cytoplasm | kinase | |
| chr14 | *SYNDIG1L* | 4 | 150 | 9 | SYNDIG1L | synapse differentiation inducing 1 | Cytoplasm | other | |
| chr17 | *AOC4P* | 5 | 166 | 9 | AOC4P | amine oxidase copper containing 4, pseudogene | Other | other | |
| chr19 | *ZNF418* | 4 | 133 | 9 | ZNF418 | zinc finger protein 418 | Nucleus | transcription regulator | |
| chr2 | *FEZ2* | 4 | 88 | 9 | FEZ2 | fasciculation and elongation protein zeta 2 | Cytoplasm | other | |
| chr3 | *AC018816.3* | 5 | 83 | 9 | | | | | |
| chr3 | *ECT2* | 4 | 84 | 9 | ECT2 | epithelial cell transforming 2 | Cytoplasm | other | |
| chr22 | *GRAP2* | 4 | 117 | 9 | GRAP2 | GRB2 related adaptor protein 2 | Cytoplasm | other | |
| chr21 | *SMIM11* | 4 | 129 | 9 | SMIM11A | small integral membrane protein | Plasma Membrane | other | |
| chr17 | *PSMB3* | 4 | 125 | 9 | PSMB3 | proteasome subunit beta 3 | Cytoplasm | peptidase | |
| chr6 | *AKAP7* | 3 | 81 | 9 | AKAP7 | A-kinase anchoring protein 7 | Plasma Membrane | other | |

| | | | | | | | | | |
|---|---|---|---|---|---|---|---|---|---|
| chrX | *ACRC* | 3 | 100 | 9 | GCNA | germ cell nuclear acidic | Nucleus | other | |
| chr15 | *RP11-1006G14.2* | 5 | 129 | 9 | | | | | |
| chr12 | *KRT85* | 5 | 349 | 9 | KRT85 | keratin 85 | Extracellular Space | other | |
| chr1 | *C1orf116* | 4 | 213 | 9 | C1orf116 | chromosome 1 open reading frame 116 | Cytoplasm | other | |
| chr19 | *SYNGR4* | 4 | 161 | 9 | SYNGR4 | synaptogyrin 4 | Plasma Membrane | other | |
| chr5 | *DDX4* | 4 | 117 | 9 | DDX4 | DEAD-box helicase 4 | Nucleus | enzyme | |
| chr12 | *RPL6* | 3 | 128 | 9 | RPL6 | ribosomal protein L6 | Nucleus | other | |
| chr8 | *KB-1615E4.2* | 4 | 96 | 9 | | | | | |
| chrX | *ATP6AP2* | 5 | 96 | 9 | ATP6AP2 | ATPase H+ transporting accessory protein 2 | Cytoplasm | transporter | |
| chr19 | *TMEM59L* | 5 | 220 | 9 | TMEM59L | transmembrane protein 59 like | Cytoplasm | other | |
| chr16 | *GLIS2* | 5 | 265 | 9 | GLIS2 | GLIS family zinc finger 2 | Nucleus | transcription regulator | |
| chr7 | *C7orf65* | 4 | 222 | 9 | C7orf65 | chromosome 7 open reading frame 65 | Other | other | |
| chr11 | *FOLH1B* | 4 | 129 | 9 | FOLH1B | folate hydrolase 1B | Cytoplasm | peptidase | |
| chr4 | *CORIN* | 4 | 80 | 9 | CORIN | corin, serine peptidase | Plasma Membrane | peptidase | |
| chr1 | *CD55* | 4 | 195 | 9 | CD55 | CD55 molecule (Cromer blood | Plasma Membrane | other | |
| chr19 | *CTD-2571L23.6* | 4 | 129 | 9 | | | | | |
| chr1 | *GPATCH3* | 4 | 126 | 9 | GPATCH3 | G-patch domain containing 3 | Other | other | |
| chr3 | *RP11-103G8.2* | 4 | 134 | 9 | | | | | |
| chr8 | *PURG* | 4 | 80 | 9 | PURG | purine rich element binding | Other | other | |
| chr7 | *RP11-328J2.1* | 4 | 83 | 9 | | | | | |
| chr3 | *TM4SF4* | 4 | 124 | 9 | TM4SF4 | transmembrane 4 L six family member 4 | Plasma Membrane | other | |
| chr2 | *WDR35* | 3 | 99 | 9 | WDR35 | WD repeat domain 35 | Cytoplasm | other | |
| chr20 | *ADRA1D* | 4 | 142 | 9 | ADRA1D | adrenoceptor alpha 1D | Plasma Membrane | G-protein coupled recepto | chlorpheniramine/methscopolamine/phenylephrine, apraclonidine, epinephrine/lidocaine, azatadine/pseudoephedrine, dobutamine, terazosine, quinidine, doxazosin, brexpiprazole, dihydroergotamine, guanethidine, tolazoline, brompheniramine/codeine/phenylpropanolamine, loratadine/pseudoephedrine, chlorpheniramine/ibuprofen/pseudoephedrine, brimonidine, dipivefrin, [illegible] |
| chr9 | *NUTM2F* | 4 | 154 | 9 | NUTM2F/NUTM2[illegible] | NUT family member 2F | Other | other | |
| chr22 | *CRYBB2P1* | 4 | 88 | 9 | CRYBB2P1 | crystallin beta B2 pseudogene 1 | Other | other | |
| chr10 | *GLUD1P2* | 5 | 213 | 9 | GLUD1P2 | glutamate dehydrogenase 1 pseudogene 2 | Other | other | |
| chr7 | *AC005281.1* | 4 | 120 | 9 | | | | | |
| chr19 | *ZNF44* | 4 | 114 | 9 | ZNF44 | zinc finger protein 44 | Nucleus | transcription regulator | |
| chr3 | *USP19* | 4 | 186 | 9 | USP19 | ubiquitin specific peptidase 19 | Cytoplasm | peptidase | |
| chr5 | *RNF145* | 4 | 174 | 9 | RNF145 | ring finger protein 145 | Other | other | |
| chr11 | *RP11-685M7.5* | 4 | 178 | 9 | | | | | |
| chrX | *CXorf22* | 4 | 99 | 9 | CFAP47 | cilia and flagella associated | Other | other | |
| chr14 | *RP11-33N16.3* | 4 | 91 | 9 | | | | | |
| chr19 | *CTC-360P9.1* | 4 | 80 | 9 | | | | | |
| chr19 | *SIGLEC6* | 4 | 133 | 9 | SIGLEC6 | sialic acid binding Ig like lectin 6 | Extracellular Space | other | |
| chr1 | *TROVE2* | 3 | 90 | 9 | RO60 | Ro60, Y RNA binding protein | Nucleus | other | |
| chr1 | *WASH7P* | 5 | 614 | 9 | WASH7P | WASP family homolog 7, pseudogene | Other | other | |
| chr2 | *OBSL1* | 4 | 196 | 9 | OBSL1 | obscurin like cytoskeletal adaptor | Cytoplasm | other | |
| chr4 | *ACSL1* | 3 | 95 | 9 | ACSL1 | acyl-CoA synthetase long chain family member 1 | Cytoplasm | enzyme | |
| chr12 | *ZNF740* | 4 | 244 | 9 | ZNF740 | zinc finger protein 740 | Other | other | |
| chr7 | *AC068610.5* | 4 | 97 | 9 | | | | | |
| chr17 | *SPATA20* | 4 | 278 | 9 | SPATA20 | spermatogenesis associated 20 | Other | other | |
| chr11 | *NAALADL1* | 4 | 187 | 9 | NAALADL1 | N-acetylated alpha-linked acidic dipeptidase like 1 | Plasma Membrane | peptidase | |
| chr15 | *STARD5* | 5 | 164 | 9 | STARD5 | StAR related lipid transfer domain containing 5 | Cytoplasm | transporter | |
| chr8 | *DDHD2* | 3 | 85 | 9 | DDHD2 | DDHD domain containing 2 | Cytoplasm | enzyme | |
| chr9 | *EPB41L4B* | 4 | 119 | 9 | EPB41L4B | erythrocyte membrane protein band 4.1 like 4B | Cytoplasm | transporter | |
| chr1 | *EIF3I* | 4 | 121 | 9 | EIF3I | eukaryotic translation initiation factor 3 subunit I | Cytoplasm | translation regulator | |
| chr3 | *CLDN16* | 3 | 72 | 9 | CLDN16 | claudin 16 | Plasma Membrane | transporter | |
| chr8 | *DEPTOR* | 4 | 79 | 9 | DEPTOR | DEP domain containing MTOR interacting protein | Cytoplasm | other | |
| chr2 | *CAPN14* | 4 | 106 | 9 | CAPN14 | calpain 14 | Other | other | |
| chrX | *RRAGB* | 4 | 122 | 9 | RRAGB | Ras related GTP binding B | Cytoplasm | enzyme | |
| chr2 | *CCDC150* | 4 | 108 | 9 | CCDC150 | coiled-coil domain containing 150 | Other | other | |

| | | | | | | | | | |
|---|---|---|---|---|---|---|---|---|---|
| chr2 | *KCNH7* | 3 | 97 | 9 | KCNH7 | potassium voltage-gated channel subfamily H member 7 | Plasma Membrane | ion channel | quinidine, procainamide, dextromethorphan/quinidine, dofetilide |
| chr15 | *RP11-403B2.7* | 4 | 124 | 9 | | | | | |
| chr1 | *RP11-533E19.2* | 3 | 103 | 9 | | | | | |
| chr15 | *CTD-2022H16.1* | 4 | 93 | 9 | | | | | |
| chr17 | *RP11-160E2.17* | 4 | 141 | 9 | | | | | |
| chr16 | *ZNF771* | 5 | 116 | 9 | ZNF771 | zinc finger protein 771 | Nucleus | transcription regulator | |
| chr2 | *STAT4* | 4 | 84 | 9 | STAT4 | signal transducer and activator of transcription 4 | Nucleus | transcription regulator | |
| chr7 | *CLEC2L* | 5 | 209 | 9 | CLEC2L | C-type lectin domain family 2 member L | Other | other | |
| chr2 | *FAHD2A* | 4 | 181 | 9 | FAHD2A | fumarylacetoacetate hydrolase domain containing 2A | Other | enzyme | |
| chr1 | *GBA* | 4 | 182 | 9 | GBA | glucosylceramidase beta | Cytoplasm | enzyme | |
| chr1 | *LGALS8* | 4 | 112 | 9 | LGALS8 | galectin 8 | Extracellular Space | other | |
| chr22 | *PES1* | 4 | 149 | 9 | PES1 | pescadillo ribosomal biogenesis factor 1 | Nucleus | other | |
| chr2 | *POTEI* | 4 | 142 | 9 | POTEI | POTE ankyrin domain family member I | Extracellular Space | other | |
| chr4 | *LRPAP1* | 5 | 228 | 9 | LRPAP1 | LDL receptor related protein associated protein 1 | Plasma Membrane | other | |
| chr2 | *MFSD9* | 4 | 176 | 9 | MFSD9 | major facilitator superfamily domain containing 9 | Other | transporter | |
| chrX | *CENPI* | 3 | 87 | 9 | CENPI | centromere protein I | Nucleus | other | |
| chr1 | *CHI3L2* | 5 | 131 | 9 | CHI3L2 | chitinase 3 like 2 | Extracellular Space | enzyme | |
| chr1 | *CNN3* | 5 | 115 | 9 | CNN3 | calponin 3 | Cytoplasm | other | |
| chr3 | *ARHGEF26-AS1* | 5 | 107 | 9 | ARHGEF26-AS1 | ARHGEF26 antisense RNA 1 | Other | other | |
| chr1 | *AKR7A2* | 4 | 189 | 9 | AKR7A2 | aldo-keto reductase family 7 member A2 | Cytoplasm | enzyme | |
| chr3 | *BAP1* | 4 | 224 | 9 | BAP1 | BRCA1 associated protein 1 | Nucleus | peptidase | |
| chr17 | *PIRT* | 4 | 106 | 9 | PIRT | phosphoinositide interacting regulator of transient receptor potential channels | Plasma Membrane | other | |
| chr1 | *PALMD* | 5 | 111 | 9 | PALMD | palmdelphin | Cytoplasm | other | |
| chr7 | *TRIM74* | 5 | 218 | 9 | TRIM73/TRIM74 | tripartite motif containing 74 | Cytoplasm | other | |
| chrX | *TEX28P1* | 4 | 136 | 9 | TEX28P1 | testis expressed 28 pseudogene | Other | other | |
| chr9 | *PSIP1* | 4 | 92 | 9 | PSIP1 | PC4 and SFRS1 interacting | Nucleus | transcription regulator | |
| chr14 | *RP11-433J8.2* | 4 | 102 | 9 | | | | | |
| chr19 | *ZNF473* | 4 | 118 | 9 | ZNF473 | zinc finger protein 473 | Nucleus | other | |
| chr20 | *SLC4A11* | 5 | 207 | 9 | SLC4A11 | solute carrier family 4 member 11 | Plasma Membrane | transporter | |
| chr11 | *RP11-430H10.2* | 4 | 182 | 9 | | | | | |
| chr2 | *AC010105.1* | 3 | 116 | 9 | | | | | |
| chr19 | *ZNF567* | 4 | 106 | 9 | ZNF567 | zinc finger protein 567 | Nucleus | transcription regulator | |
| chr22 | *ZDHHC8* | 5 | 237 | 9 | ZDHHC8 | zinc finger DHHC-type containing | Cytoplasm | enzyme | |
| chr14 | *SOCS4* | 4 | 108 | 9 | SOCS4 | suppressor of cytokine signaling | Cytoplasm | other | |
| chr7 | *ZNF107* | 4 | 100 | 9 | ZNF107 | zinc finger protein 107 | Nucleus | transcription regulator | |
| chr12 | *RP13-941N14.1* | 4 | 114 | 9 | | | | | |
| chr8 | *ZNF16* | 4 | 192 | 9 | ZNF16 | zinc finger protein 16 | Nucleus | transcription regulator | |
| chr6 | *KLC4* | 4 | 119 | 9 | KLC4 | kinesin light chain 4 | Other | other | |
| chr1 | *GFI1* | 4 | 165 | 9 | GFI1 | growth factor independent 1 transcriptional repressor | Nucleus | transcription regulator | |
| chr7 | *CASD1* | 3 | 123 | 9 | CASD1 | CAS1 domain containing 1 | Cytoplasm | enzyme | |
| chr14 | *RBM23* | 4 | 110 | 9 | RBM23 | RNA binding motif protein 23 | Nucleus | other | |
| chr19 | *LENG8* | 5 | 287 | 9 | LENG8 | leukocyte receptor cluster | Other | other | |
| chr11 | *TSPAN32* | 5 | 233 | 9 | TSPAN32 | tetraspanin 32 | Plasma Membrane | other | |
| chr6 | *MED20* | 4 | 117 | 9 | MED20 | mediator complex subunit 20 | Nucleus | transcription regulator | |
| chr20 | *RBM38* | 5 | 284 | 9 | RBM38 | RNA binding motif protein 38 | Nucleus | other | |
| chr19 | *CEACAM4* | 4 | 154 | 9 | CEACAM4 | CEA cell adhesion molecule 4 | Plasma Membrane | other | |
| chr19 | *POP4* | 4 | 189 | 9 | POP4 | POP4 homolog, ribonuclease P/MRP subunit | Nucleus | enzyme | |
| chr17 | *ANKRD40* | 4 | 126 | 9 | ANKRD40 | ankyrin repeat domain 40 | Other | other | |
| chr2 | *AC010145.4* | 4 | 121 | 9 | | | | | |
| chr10 | *PPIF* | 4 | 190 | 9 | PPIF | peptidylprolyl isomerase F | Cytoplasm | enzyme | |
| chr14 | *CTD-2277K2.1* | 5 | 107 | 9 | | | | | |
| chr3 | *RP11-190P13.2* | 3 | 134 | 9 | | | | | |
| chr8 | *WDYHV1* | 4 | 94 | 9 | WDYHV1 | WDYHV motif containing 1 | Cytoplasm | enzyme | |
| chr19 | *ZNF415* | 4 | 87 | 9 | ZNF415 | zinc finger protein 415 | Nucleus | transcription regulator | |
| chr17 | *CBX2* | 4 | 193 | 9 | CBX2 | chromobox 2 | Nucleus | transcription regulator | |
| chr3 | *RP11-550I24.2* | 4 | 127 | 9 | | | | | |
| chr20 | *CPNE1* | 4 | 167 | 9 | CPNE1 | copine 1 | Nucleus | transporter | |
| chr5 | *IL17B* | 4 | 134 | 9 | IL17B | interleukin 17B | Extracellular Space | cytokine | |
| chr15 | *NTRK3-AS1* | 5 | 134 | 9 | NTRK3-AS1 | NTRK3 antisense RNA 1 | Other | other | |
| chr6 | *SFT2D1* | 4 | 133 | 9 | SFT2D1 | SFT2 domain containing 1 | Other | other | |
| chr6 | *SRF* | 4 | 220 | 9 | SRF | serum response factor | Nucleus | transcription regulator | |
| chr16 | *ZNF500* | 4 | 184 | 9 | ZNF500 | zinc finger protein 500 | Nucleus | transcription regulator | |
| chr5 | *PCDH12* | 4 | 134 | 9 | PCDH12 | protocadherin 12 | Plasma Membrane | other | |
| chr10 | *GSTO2* | 4 | 132 | 9 | GSTO2 | glutathione S-transferase omega | Cytoplasm | enzyme | |
| chr12 | *RP11-625L16.1* | 3 | 91 | 9 | | | | | |
| chr8 | *RP11-37B2.1* | 4 | 108 | 9 | | | | | |
| chr11 | *ANKRD13D* | 5 | 219 | 9 | ANKRD13D | ankyrin repeat domain 13D | Plasma Membrane | other | |
| chr3 | *QARS* | 4 | 171 | 9 | | | | | |
| chr6 | *RP11-420G6.4* | 4 | 148 | 9 | | | | | |
| chr3 | *SERPINI2* | 3 | 84 | 9 | SERPINI2 | serpin family I member 2 | Extracellular Space | other | |
| chr16 | *FLYWCH2* | 5 | 220 | 9 | FLYWCH2 | FLYWCH family member 2 | Other | other | |
| chr9 | *SMC5-AS1* | 4 | 113 | 9 | SMC5-AS1 | SMC5 antisense RNA 1 (head to head) | Other | other | |
| chr16 | *INO80E* | 4 | 187 | 9 | INO80E | INO80 complex subunit E | Nucleus | other | |

| | | | | | | | | | |
|---|---|---|---|---|---|---|---|---|---|
| chr17 | *MYL4* | 4 | 95 | 9 | MYL4 | myosin light chain 4 | Cytoplasm | other | |
| chr2 | *IGFBP2* | 4 | 134 | 9 | IGFBP2 | insulin like growth factor binding protein 2 | Extracellular Space | other | |
| chr22 | *POM121L9P* | 4 | 201 | 9 | POM121L9P | POM121 transmembrane nucleoporin like 9, pseudogene | Other | other | |
| chr17 | *COIL* | 3 | 81 | 9 | COIL | coilin | Nucleus | enzyme | |
| chr16 | *CLUHP3* | 4 | 211 | 9 | CLUHP3 | clustered mitochondria homolog pseudogene 3 | Other | other | |
| chr22 | *SEPT3* | 4 | 98 | 9 | SEPTIN3 | septin 3 | Cytoplasm | enzyme | |
| chr16 | *NPIPA3* | 4 | 96 | 9 | PA1 (includes oth | nuclear pore complex interacting protein family member A3 | Nucleus | other | |
| chr1 | *RP11-269F19.2* | 4 | 144 | 9 | | | | | |
| chr12 | *FZD10-AS1* | 4 | 144 | 9 | FZD10-AS1 | FZD10 antisense divergent | Other | other | |
| chr6 | *RP3-429O6.1* | 4 | 127 | 9 | | | | | |
| chr17 | *TMC8* | 5 | 199 | 9 | TMC8 | transmembrane channel like 8 | Cytoplasm | other | |
| chr10 | *CUTC* | 3 | 111 | 9 | CUTC | cutC copper transporter | Cytoplasm | other | |
| chr2 | *AGFG1* | 3 | 81 | 9 | AGFG1 | ArfGAP with FG repeats 1 | Nucleus | other | |
| chr4 | *GPR125* | 3 | 79 | 9 | ADGRA3 | adhesion G protein-coupled receptor A3 | Plasma Membrane | i-protein coupled recepto | |
| chr6 | *RP11-392A23.4* | 4 | 110 | 9 | | | | | |
| chr16 | *CENPT* | 5 | 169 | 9 | CENPT | centromere protein T | Cytoplasm | transcription regulator | |
| chr3 | *KLHL24* | 4 | 103 | 9 | KLHL24 | kelch like family member 24 | Plasma Membrane | other | |
| chr14 | *NFATC4* | 4 | 156 | 9 | NFATC4 | nuclear factor of activated T cells | Nucleus | transcription regulator | |
| chr22 | *RP3-400N23.6* | 4 | 81 | 9 | | | | | |
| chr9 | *RP11-316P17.2* | 4 | 127 | 9 | | | | | |
| chr16 | *ESRP2* | 4 | 200 | 9 | ESRP2 | epithelial splicing regulatory | Nucleus | other | |
| chr21 | *LINC00159* | 4 | 102 | 9 | LINC00159 | long intergenic non-protein coding RNA 159 | Other | other | |
| chr9 | *CEL* | 4 | 222 | 9 | CEL | carboxyl ester lipase | Extracellular Space | enzyme | |
| chr22 | *DDX17* | 3 | 93 | 9 | DDX17 | DEAD-box helicase 17 | Nucleus | enzyme | |
| chr4 | *C4orf50* | 4 | 147 | 9 | C4orf50 | chromosome 4 open reading frame 50 | Other | other | |
| chr10 | *CUEDC2* | 4 | 156 | 9 | CUEDC2 | CUE domain containing 2 | Extracellular Space | other | |
| chr22 | *TSPO* | 4 | 230 | 9 | TSPO | translocator protein | Cytoplasm | transmembrane recepto | PK 11195, olanzapine, SSR180575, triazolam, flurazepam, flunitrazepam, flumazenil, chlordiazepoxide, midazolam, temazepam, alprazolam, estazolam, diazepinomicin, zolpidem, CGS-8216, fluoxetine/olanzapine, clorazepate, eszopiclone, clonazepam |
| chr12 | *DNAJC22* | 5 | 279 | 9 | DNAJC22 | DnaJ heat shock protein family (Hsp40) member C22 | Other | other | |
| chr7 | *RP11-44M6.1* | 4 | 111 | 9 | | | | | |
| chr5 | *CAMLG* | 4 | 112 | 9 | CAMLG | calcium modulating ligand | Cytoplasm | other | alemtuzumab/cyclosporin A, cyclosporin A, cyclosporine A/tacrolimus, cyclosporin A/methotrexate, cyclosporine A/sirolimus/tacrolimu |
| chr13 | *UBL3* | 4 | 85 | 9 | UBL3 | ubiquitin like 3 | Cytoplasm | other | |
| chr3 | *TXNRD3* | 4 | 105 | 9 | TXNRD3 | thioredoxin reductase 3 | Cytoplasm | enzyme | |
| chr11 | *ZBTB3* | 4 | 152 | 9 | ZBTB3 | zinc finger and BTB domain containing 3 | Nucleus | other | |
| chr19 | *XRCC1* | 4 | 165 | 9 | XRCC1 | X-ray repair cross complementing | Nucleus | other | |
| chr11 | *SLC29A2* | 4 | 367 | 9 | SLC29A2 | solute carrier family 29 member 2 | Plasma Membrane | transporter | |
| chr12 | *RP11-728G15.1* | 4 | 122 | 9 | | | | | |
| chr1 | *CSDE1* | 3 | 139 | 9 | CSDE1 | cold shock domain containing E1 | Cytoplasm | enzyme | |
| chr10 | *AGAP8* | 4 | 154 | 9 | AP5 (includes oth | ArfGAP with GTPase domain, ankyrin repeat and PH domain 5 | Other | other | |
| chr18 | *RP11-25L3.3* | 4 | 98 | 9 | | | | | |
| chr12 | *RP11-174G6.1* | 4 | 92 | 9 | | | | | |
| chr1 | *YIPF1* | 4 | 89 | 9 | YIPF1 | Yip1 domain family member 1 | Cytoplasm | other | |
| chr8 | *RP11-346L1.2* | 3 | 94 | 9 | | | | | |
| chrX | *OPN1LW* | 3 | 104 | 9 | OPN1LW | opsin 1, long wave sensitive | Plasma Membrane | i-protein coupled recepto | |
| chr7 | *BPGM* | 3 | 129 | 9 | BPGM | bisphosphoglycerate mutase | Extracellular Space | phosphatase | |
| chr12 | *ADIPOR2* | 3 | 72 | 9 | ADIPOR2 | adiponectin receptor 2 | Plasma Membrane | transmembrane recepto | |
| chr19 | *FAM71E1* | 4 | 159 | 9 | FAM71E1 | family with sequence similarity 71 member E1 | Other | other | |
| chr17 | *TCAM1P* | 4 | 165 | 9 | TCAM1P | testicular cell adhesion molecule 1, pseudogene | Other | other | |
| chr1 | *ERRFI1* | 4 | 148 | 9 | ERRFI1 | ERBB receptor feedback inhibitor | Cytoplasm | other | |
| chr12 | *RP11-25I15.2* | 5 | 138 | 9 | | | | | |
| chr1 | *DCDC2B* | 4 | 163 | 9 | DCDC2B | doublecortin domain containing | Other | other | |
| chr16 | *RP11-361L15.4* | 5 | 114 | 9 | | | | | |
| chr4 | *RP11-145G20.1* | 4 | 101 | 9 | | | | | |
| chr1 | *NES* | 4 | 231 | 9 | NES | nestin | Extracellular Space | other | |

| | | | | | | | | | |
|---|---|---|---|---|---|---|---|---|---|
| chr19 | *ZNF227* | 3 | 92 | 9 | ZNF227 | zinc finger protein 227 | Nucleus | transcription regulator | |
| chr16 | *FBXL16* | 5 | 290 | 9 | FBXL16 | F-box and leucine rich repeat protein 16 | Cytoplasm | other | |
| chr21 | *WRB* | 4 | 155 | 9 | GET1 | guided entry of tail-anchored proteins factor 1 | Extracellular Space | other | |
| chr17 | *ELAC2* | 5 | 226 | 9 | ELAC2 | elaC ribonuclease Z 2 | Nucleus | enzyme | |
| chr1 | *CPSF3L* | 5 | 245 | 9 | INTS11 | integrator complex subunit 11 | Nucleus | other | |
| chr9 | *RP11-399H11.2* | 4 | 223 | 9 | | | | | |
| chr16 | *ZNF267* | 4 | 106 | 9 | ZNF267 | zinc finger protein 267 | Nucleus | transcription regulator | |
| chr10 | *BNIP3* | 4 | 151 | 9 | BNIP3 | BCL2 interacting protein 3 | Cytoplasm | other | |
| chr6 | *SESN1* | 3 | 77 | 9 | SESN1 | sestrin 1 | Nucleus | other | |
| chr16 | *RP11-467I17.1* | 4 | 119 | 9 | | | | | |
| chr22 | *SF3A1* | 4 | 101 | 9 | SF3A1 | splicing factor 3a subunit 1 | Nucleus | other | |
| chr19 | *CTD-2369P2.5* | 4 | 99 | 9 | | | | | |
| chr2 | *KDM3A* | 4 | 77 | 9 | KDM3A | lysine demethylase 3A | Nucleus | transcription regulator | |
| chr17 | *NOL11* | 4 | 101 | 9 | NOL11 | nucleolar protein 11 | Nucleus | other | |
| chr22 | *AC011718.2* | 4 | 132 | 9 | | | | | |
| chr19 | *PRODH2* | 4 | 118 | 9 | PRODH2 | proline dehydrogenase 2 | Cytoplasm | enzyme | |
| chr4 | *CCDC158* | 4 | 89 | 9 | CCDC158 | coiled-coil domain containing 158 | Other | other | |
| chr5 | *MAN2A1* | 4 | 86 | 9 | MAN2A1 | mannosidase alpha class 2A member 1 | Cytoplasm | enzyme | |
| chr5 | *KIF3A* | 4 | 92 | 9 | KIF3A | kinesin family member 3A | Cytoplasm | enzyme | |
| chr6 | *SLC2A12* | 3 | 101 | 9 | SLC2A12 | solute carrier family 2 member 12 | Plasma Membrane | transporter | |
| chr19 | *PSG5* | 4 | 119 | 9 | PSG5 | pregnancy specific beta-1-glycoprotein 5 | Extracellular Space | other | |
| chr11 | *PEX16* | 5 | 305 | 9 | PEX16 | peroxisomal biogenesis factor 16 | Cytoplasm | other | |
| chr1 | *FBXO6* | 5 | 204 | 9 | FBXO6 | F-box protein 6 | Cytoplasm | other | |
| chr16 | *LA16c-306E5.3* | 5 | 137 | 9 | | | | | |
| chr19 | *TMEM145* | 4 | 143 | 9 | TMEM145 | transmembrane protein 145 | Other | other | |
| chrX | *RAP2C-AS1* | 3 | 90 | 9 | RAP2C-AS1 | RAP2C antisense RNA 1 | Other | other | |
| chr7 | *ZNF138* | 4 | 98 | 9 | ZNF138 | zinc finger protein 138 | Nucleus | transcription regulator | |
| chr15 | *GOLGA8J* | 3 | 134 | 9 | GA8K (includes ot | golgin A8 family member K | Other | other | |
| chr15 | *RP11-507B12.1* | 4 | 78 | 9 | | | | | |
| chr10 | *CCDC7* | 4 | 121 | 9 | CCDC7 | coiled-coil domain containing 7 | Other | other | |
| chr17 | *TOM1L1* | 3 | 87 | 9 | TOM1L1 | target of myb1 like 1 membrane trafficking protein | Cytoplasm | other | |
| chr6 | *PTCRA* | 4 | 176 | 9 | PTCRA | pre T cell antigen receptor alpha | Plasma Membrane | other | |
| chr7 | *CYP3A43* | 4 | 100 | 9 | CYP3A43 | cytochrome P450 family 3 subfamily A member 43 | Cytoplasm | enzyme | cobicistat, cobicistat/darunavir, cobicistat/elvitegravir /emtricitabine/tenofov ir disoproxil, atazanavir/cobicistat |
| chr20 | *NDUFAF5* | 4 | 83 | 9 | NDUFAF5 | NADH:ubiquinone oxidoreductase complex | Cytoplasm | other | |
| chr15 | *HDGFRP3* | 3 | 101 | 9 | | | | | |
| chr9 | *RP11-23J9.3* | 3 | 147 | 9 | | | | | |
| chr11 | *PIH1D2* | 3 | 72 | 9 | PIH1D2 | PIH1 domain containing 2 | Other | other | |
| chr5 | *RP11-114H21.2* | 4 | 89 | 9 | | | | | |
| chr20 | *NCOA5* | 4 | 152 | 9 | NCOA5 | nuclear receptor coactivator 5 | Nucleus | other | |
| chr9 | *RP11-383M4.6* | 3 | 123 | 9 | | | | | |
| chr19 | *CD70* | 4 | 146 | 9 | CD70 | CD70 molecule | Extracellular Space | cytokine | cusatuzumab, SGN-CD70A, vorsetuzumab |
| chr12 | *COPS7A* | 4 | 163 | 9 | COPS7A | COP9 signalosome subunit 7A | Cytoplasm | other | |
| chr11 | *API5* | 4 | 83 | 9 | API5 | apoptosis inhibitor 5 | Cytoplasm | other | |
| chr1 | *RP11-34P13.7* | 4 | 138 | 9 | | | | | |
| chr9 | *RP11-123K19.1* | 5 | 156 | 9 | | | | | |
| chr17 | *CEP95* | 4 | 83 | 9 | CEP95 | centrosomal protein 95 | Cytoplasm | other | |
| chr19 | *ZNF613* | 4 | 89 | 9 | ZNF613 | zinc finger protein 613 | Nucleus | transcription regulator | |
| chr22 | *SERHL* | 3 | 155 | 9 | SERHL | serine hydrolase like | Cytoplasm | other | |
| chr8 | *DGAT1* | 5 | 240 | 9 | DGAT1 | diacylglycerol O-acyltransferase 1 | Cytoplasm | enzyme | |
| chr4 | *TAPT1* | 3 | 85 | 9 | TAPT1 | transmembrane anterior posterior transformation 1 | Plasma Membrane | G-protein coupled recepto | |
| chr3 | *BCHE* | 4 | 76 | 9 | BCHE | butyrylcholinesterase | Plasma Membrane | enzyme | pyridostigmine, hexafluorenium, edrophonium, malathion, rivastigmine, pralidoxime, echothiophate, tacrine, dipivefrin, isoflurophate, |
| chr16 | *RP11-863P13.5* | 4 | 144 | 9 | | | | | |
| chr11 | *PACSIN3* | 5 | 494 | 9 | PACSIN3 | protein kinase C and casein kinase substrate in neurons 3 | Cytoplasm | other | |
| chrX | *RNF128* | 4 | 97 | 9 | RNF128 | ring finger protein 128 | Cytoplasm | enzyme | |
| chr21 | *CHODL-AS1* | 4 | 73 | 9 | CHODL-AS1 | CHODL antisense RNA 1 | Other | other | |
| chr13 | *VPS36* | 3 | 76 | 9 | VPS36 | vacuolar protein sorting 36 | Cytoplasm | other | |
| chr3 | *LINC00887* | 3 | 124 | 9 | LINC00887 | long intergenic non-protein coding RNA 887 | Other | other | |
| chr1 | *ERMAP* | 4 | 103 | 9 | ERMAP | erythroblast membrane associated protein (Scianna | Cytoplasm | other | |
| chr1 | *ETV3* | 4 | 169 | 9 | ETV3 | ETS variant transcription factor 3 | Nucleus | transcription regulator | |
| chr4 | *SLC9B1* | 4 | 97 | 9 | SLC9B1 | solute carrier family 9 member B1 | Plasma Membrane | transporter | |
| chr1 | *RP11-418J17.3* | 5 | 154 | 9 | | | | | |
| chr19 | *EBI3* | 4 | 143 | 9 | EBI3 | Epstein-Barr virus induced 3 | Extracellular Space | cytokine | |
| chr11 | *FOSL1* | 5 | 262 | 9 | FOSL1 | FOS like 1, AP-1 transcription factor subunit | Nucleus | transcription regulator | MORAb-202 |

| | | | | | | | | | |
|---|---|---|---|---|---|---|---|---|---|
| chr10 | *C10orf118* | 4 | 85 | 9 | CCDC186 | coiled-coil domain containing 186 | Other | other | |
| chr1 | *XKR8* | 4 | 197 | 9 | XKR8 | XK related 8 | Plasma Membrane | other | |
| chr5 | *HBEGF* | 4 | 118 | 9 | HBEGF | heparin binding EGF like growth factor | Extracellular Space | growth factor | |
| chr11 | *LMO2* | 5 | 135 | 9 | LMO2 | LIM domain only 2 | Nucleus | transcription regulator | |
| chr10 | *CTGLF8P* | 3 | 119 | 9 | | | | | |
| chr17 | *IFI35* | 4 | 152 | 9 | IFI35 | interferon induced protein 35 | Nucleus | other | |
| chr15 | *LINS* | 4 | 137 | 9 | LINS1 | lines homolog 1 | Other | other | |
| chr16 | *NPIPA2* | 4 | 113 | 9 | PA1 (includes oth | nuclear pore complex interacting protein family member A3 | Nucleus | other | |
| chr11 | *RP11-405K6.1* | 4 | 120 | 9 | | | | | |
| chr7 | *GARS* | 4 | 123 | 9 | GARS1 | glycyl-tRNA synthetase 1 | Cytoplasm | enzyme | |
| chr12 | *SMCO2* | 4 | 128 | 9 | SMCO2 | single-pass membrane protein with coiled-coil domains 2 | Other | other | |
| chr8 | *JRK* | 5 | 222 | 9 | JRK | Jrk helix-turn-helix protein | Nucleus | other | |
| chr19 | *PIN1* | 3 | 168 | 9 | PIN1 | peptidylprolyl cis/trans isomerase, NIMA-interacting 1 | Nucleus | enzyme | |
| chr22 | *CBX6* | 4 | 150 | 9 | CBX6 | chromobox 6 | Nucleus | other | |
| chr19 | *ZSCAN5B* | 4 | 109 | 9 | ZSCAN5B | zinc finger and SCAN domain containing 5B | Other | other | |
| chr1 | *ATXN7L2* | 4 | 289 | 9 | ATXN7L2 | ataxin 7 like 2 | Other | other | |
| chr11 | *ALG1L8P* | 4 | 187 | 9 | ALG1L8P | ALG1 like 8, pseudogene | Other | other | |
| chr21 | *GART* | 3 | 81 | 9 | GART | phosphoribosylglycinamide formyltransferase, phosphoribosylglycinamide synthetase, phosphoribosylaminoimidazole synthetase | Cytoplasm | enzyme | pemetrexed, bevacizumab/erlotinib/pemetrexed, bevacizumab/pemetrexed, pelitrexol, pembrolizumab/pemetrexed, gemcitabine/pemetrexed, bevacizumab/paclitaxel/pemetrexed, |
| chr9 | *TPRN* | 5 | 228 | 9 | TPRN | taperin | Extracellular Space | other | |
| chr1 | *GBP6* | 3 | 80 | 9 | GBP6 | guanylate binding protein family member 6 | Cytoplasm | enzyme | |
| chr14 | *GPR132* | 4 | 185 | 9 | GPR132 | G protein-coupled receptor 132 | Plasma Membrane | G-protein coupled recepto | |
| chr11 | *MTNR1B* | 4 | 163 | 9 | MTNR1B | melatonin receptor 1B | Plasma Membrane | G-protein coupled recepto | ramelteon, agomelatine, melatonin, amitriptyline/melatoni |
| chr2 | *AC079135.1* | 4 | 113 | 9 | | | | | |
| chr3 | *TFG* | 4 | 103 | 9 | TFG | trafficking from ER to golgi | Cytoplasm | other | |
| chr5 | *DNAJC21* | 4 | 252 | 9 | DNAJC21 | DnaJ heat shock protein family (Hsp40) member C21 | Other | other | |
| chr12 | *ZNF384* | 4 | 122 | 9 | ZNF384 | zinc finger protein 384 | Nucleus | transcription regulator | |
| chrX | *KIAA2022* | 4 | 85 | 9 | NEXMIF | neurite extension and migration | Nucleus | enzyme | |
| chr1 | *SELRC1* | 4 | 178 | 9 | COA7 | cytochrome c oxidase assembly factor 7 (putative) | Cytoplasm | other | |
| chr12 | *EIF2B1* | 4 | 94 | 9 | EIF2B1 | eukaryotic translation initiation factor 2B subunit alpha | Cytoplasm | translation regulator | |
| chr2 | *AC096669.2* | 5 | 88 | 9 | | | | | |
| chr16 | *OSGIN1* | 5 | 291 | 9 | OSGIN1 | oxidative stress induced growth inhibitor 1 | Other | growth factor | |
| chr2 | *AC008271.1* | 4 | 113 | 9 | | | | | |
| chr6 | *SAMD3* | 4 | 85 | 9 | SAMD3 | sterile alpha motif domain containing 3 | Other | other | |
| chr11 | *HEPACAM* | 4 | 126 | 9 | HEPACAM | hepatic and glial cell adhesion molecule | Plasma Membrane | other | |
| chr20 | *BPIFB6* | 5 | 187 | 9 | BPIFB6 | BPI fold containing family B member 6 | Extracellular Space | other | |
| chr3 | *RARRES1* | 4 | 148 | 9 | RARRES1 | retinoic acid receptor responder 1 | Plasma Membrane | other | |
| chr2 | *DGUOK* | 5 | 110 | 9 | DGUOK | deoxyguanosine kinase | Cytoplasm | kinase | |
| chr22 | *TXN2* | 4 | 119 | 9 | TXN2 | thioredoxin 2 | Cytoplasm | enzyme | |
| chr9 | *ZNF367* | 4 | 97 | 9 | ZNF367 | zinc finger protein 367 | Nucleus | transcription regulator | |
| chr11 | *CTD-2210P24.1* | 5 | 119 | 9 | | | | | |
| chr7 | *AC007128.1* | 4 | 99 | 9 | | | | | |
| chr19 | *CEA* | 3 | 104 | 9 | CEACAM5 | CEA cell adhesion molecule 5 | Plasma Membrane | other | labetuzumab govitecan, SAR408701, cergutuzumab |
| chr5 | *PRDM9* | 5 | 277 | 9 | PRDM9 | PR/SET domain 9 | Nucleus | enzyme | |
| chr12 | *RP11-753H16.3* | 4 | 144 | 9 | | | | | |
| chr2 | *FN1* | 3 | 81 | 9 | FN1 | fibronectin 1 | Extracellular Space | enzyme | L19-IL2 monoclonal antibody-cytokine fusion protein/L19TNFalpha, ocriplasmin, L19TNFalpha, L19-IL2 monoclonal antibody-cytokine |
| chr3 | *XIRP1* | 4 | 178 | 9 | XIRP1 | xin actin binding repeat | Plasma Membrane | other | |
| chr4 | *TMED11P* | 4 | 145 | 9 | TMED11P | transmembrane p24 trafficking protein 11, pseudogene | Other | other | |
| chr2 | *COPS7B* | 3 | 102 | 9 | COPS7B | COP9 signalosome subunit 7B | Cytoplasm | other | |
| chr22 | *SNAP29* | 4 | 97 | 9 | SNAP29 | synaptosome associated protein | Cytoplasm | other | |
| chr2 | *AC007365.1* | 4 | 160 | 9 | | | | | |
| chr11 | *POLR2G* | 4 | 142 | 9 | POLR2G | RNA polymerase II subunit G | Nucleus | enzyme | |
| chr16 | *SNX20* | 4 | 135 | 9 | SNX20 | sorting nexin 20 | Cytoplasm | other | |

| | | | | | | | | | |
|---|---|---|---|---|---|---|---|---|---|
| chr4 | *RP11-6C14.1* | 3 | 101 | 9 | | | | | |
| chr3 | *HLTF-AS1* | 4 | 168 | 9 | HLTF-AS1 | HLTF antisense RNA 1 | Other | other | |
| chr1 | *RP3-437I16.1* | 4 | 155 | 9 | | | | | |
| chr19 | *CEACAM20* | 3 | 93 | 9 | CEACAM20 | CEA cell adhesion molecule 20 | Plasma Membrane | other | |
| chr16 | *CORO7* | 4 | 156 | 9 | RO7/CORO7-PAI | coronin 7 | Cytoplasm | other | |
| chr3 | *COL6A4P2* | 4 | 141 | 9 | COL6A4P2 | collagen type VI alpha 4 pseudogene 2 | Other | other | |
| chr8 | *BAG4* | 3 | 81 | 9 | BAG4 | BCL2 associated athanogene 4 | Cytoplasm | other | |
| chr11 | *PAX6* | 4 | 202 | 9 | PAX6 | paired box 6 | Nucleus | transcription regulator | |
| chr2 | *EIF2AK3* | 4 | 81 | 9 | EIF2AK3 | eukaryotic translation initiation factor 2 alpha kinase 3 | Cytoplasm | kinase | |
| chr2 | *PLCD4* | 4 | 93 | 9 | PLCD4 | phospholipase C delta 4 | Cytoplasm | enzyme | |
| chr6 | *GCM1* | 4 | 92 | 9 | GCM1 | glial cells missing transcription | Nucleus | transcription regulator | |
| chr12 | *C12orf65* | 4 | 152 | 9 | C12orf65 | chromosome 12 open reading frame 65 | Other | other | |
| chr1 | *DUSP5P1* | 5 | 1965 | 9 | DUSP5P1 | dual specificity phosphatase 5 pseudogene 1 | Other | other | |
| chr7 | *AC003988.1* | 4 | 119 | 9 | | | | | |
| chr6 | *MRS2* | 4 | 79 | 9 | MRS2 | magnesium transporter MRS2 | Cytoplasm | transporter | |
| chr22 | *APOL4* | 4 | 113 | 9 | APOL4 | apolipoprotein L4 | Extracellular Space | other | |
| chr15 | *TUBGCP4* | 3 | 77 | 9 | TUBGCP4 | tubulin gamma complex associated protein 4 | Cytoplasm | other | |
| chr3 | *RP11-723O4.2* | 4 | 170 | 9 | | | | | |
| chr2 | *RP11-521D12.1* | 3 | 117 | 9 | | | | | |
| chr1 | *RP11-400N13.3* | 4 | 140 | 9 | | | | | |
| chr8 | *RP11-563N12.2* | 4 | 106 | 9 | | | | | |
| chr2 | *RP13-1039J1.4* | 4 | 253 | 9 | | | | | |
| chr9 | *RP11-424E7.3* | 4 | 330 | 9 | | | | | |
| chr12 | *DPY19L2* | 3 | 80 | 9 | DPY19L2 | dpy-19 like 2 | Nucleus | other | |
| chr16 | *ZNF213* | 5 | 218 | 9 | ZNF213 | zinc finger protein 213 | Nucleus | transcription regulator | |
| chr17 | *ABHD17AP6* | 4 | 78 | 9 | ABHD17AP6 | abhydrolase domain containing 17A pseudogene 6 | Other | other | |
| chr10 | *SLK* | 3 | 83 | 9 | SLK | STE20 like kinase | Nucleus | kinase | |
| chr10 | *ANKRD26* | 4 | 98 | 9 | ANKRD26 | ankyrin repeat domain 26 | Nucleus | transcription regulator | |
| chr2 | *AC013402.2* | 4 | 95 | 9 | | | | | |
| chr21 | *GRIK1* | 4 | 68 | 9 | GRIK1 | glutamate ionotropic receptor kainate type subunit 1 | Plasma Membrane | ion channel | topiramate, phentermine/topiram |
| chr18 | *FBXO15* | 4 | 101 | 9 | FBXO15 | F-box protein 15 | Cytoplasm | transcription regulator | |
| chr17 | *SLC16A3* | 5 | 119 | 9 | SLC16A3 | solute carrier family 16 member 3 | Plasma Membrane | transporter | |
| chr1 | *RP11-157N3.1* | 4 | 107 | 9 | | | | | |
| chr19 | *CTD-2621I17.6* | 4 | 143 | 9 | | | | | |
| chr19 | *SIGLEC8* | 4 | 128 | 9 | SIGLEC8 | sialic acid binding Ig like lectin 8 | Plasma Membrane | transmembrane receptoi | AK002 |
| chr6 | *ANKRD6* | 3 | 73 | 9 | ANKRD6 | ankyrin repeat domain 6 | Nucleus | transcription regulator | |
| chr2 | *MOB1A* | 4 | 108 | 9 | MOB1A | MOB kinase activator 1A | Plasma Membrane | other | |
| chr5 | *RP11-375B1.2* | 4 | 160 | 9 | | | | | |
| chr11 | *RP11-831H9.11* | 4 | 130 | 9 | | | | | |
| chr20 | *CCM2L* | 4 | 136 | 9 | CCM2L | CCM2 like scaffold protein | Other | other | |
| chr19 | *PRMT1* | 4 | 209 | 9 | PRMT1 | protein arginine | Nucleus | enzyme | GSK3368715 |
| chr22 | *SEC14L3* | 4 | 115 | 9 | SEC14L3 | SEC14 like lipid binding 3 | Extracellular Space | transporter | |
| chr16 | *ZNF778* | 4 | 152 | 9 | ZNF778 | zinc finger protein 778 | Nucleus | transcription regulator | |
| chr15 | *RP11-2I17.4* | 3 | 97 | 9 | | | | | |
| chr11 | *TRIM49C* | 3 | 146 | 9 | TRIM49/TRIM49C | tripartite motif containing 49 | Cytoplasm | other | |
| chr16 | *PRSS21* | 4 | 211 | 9 | PRSS21 | serine protease 21 | Cytoplasm | peptidase | |
| chr12 | *ASUN* | 3 | 126 | 9 | INTS13 | integrator complex subunit 13 | Cytoplasm | other | |
| chr19 | *UQCR11* | 4 | 167 | 9 | UQCR11 | ubiquinol-cytochrome c reductase, complex III subunit XI | Cytoplasm | enzyme | |
| chr11 | *AP003774.5* | 4 | 111 | 9 | | | | | |
| chr7 | *AC009403.2* | 4 | 105 | 9 | | | | | |
| chr16 | *AC138969.4* | 4 | 89 | 9 | | | | | |
| chr17 | *RP11-1055B8.3* | 4 | 135 | 9 | | | | | |
| chr5 | *SLC30A5* | 4 | 93 | 9 | SLC30A5 | solute carrier family 30 member 5 | Plasma Membrane | transporter | |
| chrX | *SCML1* | 4 | 123 | 9 | SCML1 | Scm polycomb group protein like | Nucleus | transcription regulator | |
| chrX | *NKAPP1* | 4 | 115 | 9 | NKAPP1 | NFKB activating protein pseudogene 1 | Other | other | |
| chr11 | *CCND1* | 5 | 230 | 9 | CCND1 | cyclin D1 | Nucleus | transcription regulator | arsenic trioxide/cytarabine/methotrexate, arsenic trioxide/idarubicin/tretinoin, arsenic trioxide/tretinoin, arsenic trioxide, arsenic trioxide/daunorubicin/tretinoin, arsenic trioxide/gemtuzumab |
| chr1 | *RP11-380J14.1* | 4 | 164 | 9 | | | | | |
| chr19 | *TPGS1* | 5 | 184 | 9 | TPGS1 | tubulin polyglutamylase complex subunit 1 | Cytoplasm | enzyme | |
| chr16 | *CTD-2009A10.1* | 4 | 114 | 9 | | | | | |
| chrX | *PIH1D3* | 3 | 70 | 9 | PIH1D3 | PIH1 domain containing 3 | Cytoplasm | other | |
| chr12 | *RP1-166H1.2* | 4 | 123 | 9 | | | | | |
| chr15 | *CSPG4P13* | 5 | 273 | 9 | CSPG4P13 | chondroitin sulfate proteoglycan 4 pseudogene 13 | Other | other | |
| chr17 | *GRB7* | 4 | 183 | 9 | GRB7 | growth factor receptor bound protein 7 | Plasma Membrane | other | |
| chr2 | *HADHA* | 3 | 84 | 9 | HADHA | hydroxyacyl-CoA dehydrogenase trifunctional multienzyme complex subunit alpha | Cytoplasm | enzyme | |
| chr1 | *RP11-63N8.3* | 4 | 199 | 9 | | | | | |

| | | | | | | | | | |
|---|---|---|---|---|---|---|---|---|---|
| chr3 | *COL6A4P1* | 5 | 139 | 9 | COL6A4P1 | collagen type VI alpha 4 pseudogene 1 | Other | other | |
| chr4 | *OTOP1* | 4 | 115 | 9 | OTOP1 | otopetrin 1 | Plasma Membrane | ion channel | |
| chr12 | *KRT77* | 4 | 236 | 9 | KRT77 | keratin 77 | Cytoplasm | other | |
| chr11 | *FERMT3* | 5 | 163 | 9 | FERMT3 | fermitin family member 3 | Cytoplasm | enzyme | |
| chrX | *XG* | 4 | 97 | 9 | XG | Xg glycoprotein (Xg blood group) | Plasma Membrane | other | |
| chr16 | *ERCC4* | 4 | 117 | 9 | ERCC4 | ERCC excision repair 4, endonuclease catalytic subunit | Nucleus | enzyme | |
| chr19 | *ZNF431* | 4 | 89 | 9 | ZNF431 | zinc finger protein 431 | Nucleus | transcription regulator | |
| chr11 | *RP11-64D24.2* | 4 | 97 | 9 | | | | | |
| chr10 | *ANKRD2* | 4 | 201 | 9 | ANKRD2 | ankyrin repeat domain 2 | Nucleus | transcription regulator | |
| chr14 | *VSX2* | 4 | 144 | 9 | VSX2 | visual system homeobox 2 | Nucleus | transcription regulator | |
| chr15 | *C15orf52* | 4 | 205 | 9 | CCDC9B | coiled-coil domain containing 9B | Other | other | |
| chr2 | *IL18R1* | 4 | 97 | 9 | IL18R1 | interleukin 18 receptor 1 | Plasma Membrane | transmembrane receptor | |
| chr8 | *DEFA1B* | 4 | 127 | 9 | FA1 (includes oth | defensin alpha 1 | Cytoplasm | other | |
| chr17 | *CSHL1* | 4 | 124 | 9 | CSHL1 | chorionic somatomammotropin hormone like 1 | Extracellular Space | growth factor | |
| chr12 | *TPH2* | 5 | 85 | 9 | TPH2 | tryptophan hydroxylase 2 | Plasma Membrane | enzyme | telotristat ethyl, telotristat |
| chrX | *MAGEC3* | 3 | 92 | 9 | MAGEC3 | MAGE family member C3 | Other | other | |
| chr1 | *ZFP69B* | 4 | 77 | 9 | ZFP69B | ZFP69 zinc finger protein B | Nucleus | transcription regulator | |
| chr17 | *KRT16P3* | 4 | 254 | 9 | KRT16P3 | keratin 16 pseudogene 3 | Other | other | |
| chr12 | *MRPS35* | 4 | 123 | 9 | MRPS35 | mitochondrial ribosomal protein | Cytoplasm | other | |
| chr8 | *AC138647.1* | 4 | 168 | 9 | | | | | |
| chr22 | *ZNF74* | 4 | 147 | 9 | ZNF74 | zinc finger protein 74 | Nucleus | transcription regulator | |
| chr11 | *SCN2B* | 5 | 164 | 9 | SCN2B | sodium voltage-gated channel beta subunit 2 | Plasma Membrane | ion channel | riluzole, zonisamide |
| chr11 | *AP000721.4* | 4 | 134 | 9 | | | | | |
| chr16 | *RP11-19N8.4* | 5 | 184 | 9 | | | | | |
| chr9 | *PSAT1* | 4 | 106 | 9 | PSAT1 | phosphoserine aminotransferase | Cytoplasm | enzyme | |
| chr20 | *SNAI1* | 4 | 236 | 9 | SNAI1 | snail family transcriptional | Nucleus | transcription regulator | |
| chr19 | *ASNA1* | 4 | 134 | 9 | GET3 | guided entry of tail-anchored proteins factor 3, ATPase | Nucleus | transporter | |
| chr11 | *FEZ1* | 4 | 101 | 9 | FEZ1 | fasciculation and elongation protein zeta 1 | Cytoplasm | other | |
| chr6 | *GINM1* | 3 | 68 | 9 | GINM1 | glycoprotein integral membrane 1 | Cytoplasm | other | |
| chr15 | *APH1B* | 4 | 79 | 9 | APH1B | aph-1 homolog B, gamma-secretase subunit | Cytoplasm | peptidase | |
| chr19 | *ODF3L2* | 4 | 226 | 9 | ODF3L2 | outer dense fiber of sperm tails 3 like 2 | Cytoplasm | other | |
| chr2 | *GULP1* | 4 | 107 | 9 | GULP1 | GULP PTB domain containing engulfment adaptor 1 | Cytoplasm | other | |
| chr22 | *PRAMENP* | 3 | 114 | 9 | PRAMENP | PRAME N-terminal like, | Other | other | |
| chr5 | *RP11-168A11.1* | 4 | 92 | 9 | | | | | |
| chr6 | *GPR110* | 4 | 91 | 9 | ADGRF1 | adhesion G protein-coupled receptor F1 | Plasma Membrane | -protein coupled recepto | |
| chr11 | *SLC22A12* | 4 | 202 | 9 | SLC22A12 | solute carrier family 22 member 12 | Plasma Membrane | transporter | verinurad, allopurinol/lesinurad, lesinurad, colchicine/probenecid, probenecid |
| chr10 | *MYOZ1* | 4 | 159 | 9 | MYOZ1 | myozenin 1 | Cytoplasm | other | |
| chr12 | *GLI1* | 4 | 115 | 9 | GLI1 | GLI family zinc finger 1 | Nucleus | transcription regulator | |
| chr19 | *ZNF554* | 4 | 99 | 9 | ZNF554 | zinc finger protein 554 | Nucleus | transcription regulator | |
| chr11 | *ALKBH8* | 4 | 91 | 9 | ALKBH8 | alkB homolog 8, tRNA methyltransferase | Cytoplasm | enzyme | |
| chr18 | *MBD2* | 3 | 74 | 9 | MBD2 | methyl-CpG binding domain | Nucleus | transcription regulator | |
| chr8 | *TONSL* | 5 | 193 | 9 | TONSL | tonsoku like, DNA repair protein | Cytoplasm | transcription regulator | |
| chr17 | *DHX33* | 4 | 126 | 9 | DHX33 | DEAH-box helicase 33 | Nucleus | enzyme | |
| chr10 | *ANKRD16* | 3 | 92 | 9 | ANKRD16 | ankyrin repeat domain 16 | Other | other | |
| chr16 | *FOPNL* | 4 | 143 | 9 | FOPNL | FGFR1OP N-terminal like | Cytoplasm | other | |
| chr1 | *GPR89B* | 3 | 104 | 9 | GPR89A/GPR89B | G protein-coupled receptor 89B | Cytoplasm | ion channel | |
| chr15 | *DNM1P51* | 4 | 122 | 9 | DNM1P51 | | Other | other | |
| chr1 | *TAL1* | 5 | 131 | 9 | TAL1 | TAL bHLH transcription factor 1, erythroid differentiation factor | Nucleus | transcription regulator | |
| chr15 | *GOLGA8H* | 4 | 172 | 9 | GA8K (includes ot | golgin A8 family member K | Other | other | |
| chr1 | *EMC1* | 4 | 101 | 9 | EMC1 | ER membrane protein complex subunit 1 | Plasma Membrane | other | |
| chr12 | *RASSF8-AS1* | 4 | 124 | 9 | RASSF8-AS1 | RASSF8 antisense RNA 1 | Other | other | |
| chrX | *FGF13* | 4 | 119 | 9 | FGF13 | fibroblast growth factor 13 | Extracellular Space | growth factor | |
| chr10 | *RP11-153K11.3* | 4 | 128 | 9 | | | | | |
| chr1 | *BATF3* | 5 | 160 | 9 | BATF3 | basic leucine zipper ATF-like transcription factor 3 | Nucleus | transcription regulator | |
| chr7 | *RP11-514P8.2* | 4 | 132 | 9 | | | | | |
| chr12 | *ITFG2* | 4 | 127 | 9 | ITFG2 | integrin alpha FG-GAP repeat containing 2 | Cytoplasm | other | |
| chr7 | *FSCN1* | 4 | 234 | 9 | FSCN1 | fascin actin-bundling protein 1 | Cytoplasm | other | |
| chr8 | *RP11-10A14.4* | 4 | 87 | 9 | | | | | |
| chr20 | *TNNC2* | 4 | 168 | 9 | TNNC2 | troponin C2, fast skeletal type | Cytoplasm | other | Ca2+ |
| chr8 | *GSDMD* | 5 | 243 | 9 | GSDMD | gasdermin D | Extracellular Space | other | |
| chr2 | *AC096649.3* | 3 | 84 | 9 | | | | | |
| chr9 | *PTRH1* | 5 | 172 | 9 | PTRH1 | peptidyl-tRNA hydrolase 1 | Cytoplasm | enzyme | |
| chr16 | *RP11-77K12.1* | 4 | 95 | 9 | | | | | |
| chr7 | *MDFIC* | 4 | 70 | 9 | MDFIC | MyoD family inhibitor domain containing | Nucleus | other | |
| chr8 | *ZFHX4-AS1* | 4 | 91 | 9 | ZFHX4-AS1 | ZFHX4 antisense RNA 1 | Other | other | |
| chr2 | *CENPA* | 4 | 91 | 9 | CENPA | centromere protein A | Nucleus | other | |
| chr6 | *RIOK1* | 4 | 105 | 9 | RIOK1 | RIO kinase 1 | Cytoplasm | kinase | |

| | | | | | | | | | |
|---|---|---|---|---|---|---|---|---|---|
| chr3 | *HTR3C* | 3 | 95 | 9 | HTR3C | 5-hydroxytryptamine receptor 3C | Plasma Membrane | ion channel | fenfluramine, dolasetron, ergotamine, palonosetron, alosetron, cisapride, granisetron, D-tubocurarine, |
| chr13 | *RB1* | 3 | 106 | 9 | RB1 | RB transcriptional corepressor 1 | Nucleus | transcription regulator | |
| chr14 | *RP11-517O13.3* | 3 | 80 | 9 | | | | | |
| chr2 | *AC018890.6* | 4 | 113 | 9 | | | | | |
| chr15 | *RP11-717I24.1* | 4 | 191 | 9 | | | | | |
| chr16 | *RP11-196G11.2* | 4 | 101 | 9 | | | | | |
| chr2 | *EFEMP1* | 4 | 109 | 9 | EFEMP1 | EGF containing fibulin extracellular matrix protein 1 | Extracellular Space | enzyme | |
| chr18 | *RP11-154H12.3* | 4 | 176 | 9 | | | | | |
| chr9 | *ZCCHC6* | 4 | 109 | 9 | TUT7 | terminal uridylyl transferase 7 | Cytoplasm | enzyme | |
| chr8 | *RP11-108E14.1* | 4 | 80 | 9 | | | | | |
| chr17 | *CNTROB* | 4 | 126 | 9 | CNTROB | centrobin, centriole duplication and spindle assembly protein | Cytoplasm | other | |
| chr4 | *CPE* | 3 | 69 | 9 | CPE | carboxypeptidase E | Cytoplasm | peptidase | |
| chr2 | *AOX1* | 5 | 92 | 9 | AOX1 | aldehyde oxidase 1 | Cytoplasm | enzyme | |
| chr9 | *GTF3C4* | 4 | 107 | 9 | GTF3C4 | general transcription factor IIIC subunit 4 | Nucleus | transcription regulator | |
| chr5 | *DHFR* | 3 | 85 | 9 | DHFR | dihydrofolate reductase | Nucleus | enzyme | iclaprim, pemetrexed, methotrexate/ofatumumab, cyclophosphamide/methotrexate/trastuzumab, cisplatin/doxorubicin/methotrexate, cytarabine/imatinib/methotrexate, methotrexate/secukinumab, infliximab/methotrexate, piritrexim, proguanil, folic acid, dasatinib/6-mercaptopurine/methotrexate, trimethoprim, cytarabine/methotrexate/mitoxantrone, bevacizumab/paclitaxel/pemetrexed, [illegible] |
| chr11 | *LAYN* | 4 | 100 | 9 | LAYN | layilin | Plasma Membrane | other | |
| chr16 | *ZNF689* | 5 | 126 | 9 | ZNF689 | zinc finger protein 689 | Nucleus | transcription regulator | |
| chr16 | *FHOD1* | 4 | 170 | 9 | FHOD1 | formin homology 2 domain containing 1 | Cytoplasm | other | |
| chr14 | *MAP3K9* | 3 | 119 | 9 | MAP3K9 | mitogen-activated protein kinase kinase kinase 9 | Cytoplasm | kinase | |
| chr6 | *SLC22A3* | 4 | 123 | 9 | SLC22A3 | solute carrier family 22 member 3 | Plasma Membrane | transporter | |
| chr7 | *AIMP2* | 4 | 141 | 9 | AIMP2 | aminoacyl tRNA synthetase complex interacting | Plasma Membrane | other | |
| chr3 | *TMPRSS7* | 4 | 120 | 9 | TMPRSS7 | transmembrane serine protease 7 | Plasma Membrane | peptidase | |
| chr5 | *CTB-131B5.5* | 4 | 110 | 9 | | | | | |
| chr17 | *FAM195B* | 4 | 215 | 9 | MCRIP1 | MAPK regulated corepressor interacting protein 1 | Nucleus | other | |
| chr9 | *RP11-276H19.2* | 4 | 99 | 9 | | | | | |
| chrX | *ATP11C* | 4 | 85 | 9 | ATP11C | ATPase phospholipid transporting 11C | Plasma Membrane | transporter | |
| chr11 | *RP11-624D11.2* | 4 | 126 | 9 | | | | | |
| chr16 | *SALL1* | 4 | 128 | 9 | SALL1 | spalt like transcription factor 1 | Nucleus | transcription regulator | |
| chr11 | *LINC00958* | 4 | 140 | 9 | LINC00958 | long intergenic non-protein coding RNA 958 | Other | other | |
| chr3 | *PEX5L* | 3 | 82 | 9 | PEX5L | peroxisomal biogenesis factor 5 | Cytoplasm | ion channel | |
| chr3 | *SELT* | 4 | 121 | 9 | | | | | |
| chr5 | *KCNN2* | 4 | 68 | 9 | KCNN2 | potassium calcium-activated channel subfamily N member 2 | Plasma Membrane | ion channel | |
| chr19 | *ZNF584* | 4 | 130 | 9 | ZNF584 | zinc finger protein 584 | Nucleus | transcription regulator | |
| chr22 | *AIFM3* | 4 | 164 | 9 | AIFM3 | apoptosis inducing factor mitochondria associated 3 | Cytoplasm | enzyme | |
| chr5 | *RP11-1280N14.3* | 4 | 83 | 9 | | | | | |
| chr12 | *RP13-735L24.1* | 3 | 107 | 9 | | | | | |
| chr14 | *EXOC3L4* | 4 | 198 | 9 | EXOC3L4 | exocyst complex component 3 | Other | other | |
| chr12 | *DCN* | 3 | 76 | 9 | DCN | decorin | Extracellular Space | other | |
| chr7 | *METTL2B* | 4 | 109 | 9 | METTL2B | methyltransferase like 2B | Other | enzyme | |
| chr2 | *C2orf62* | 4 | 108 | 9 | CATIP | ciliogenesis associated TTC17 interacting protein | Cytoplasm | other | |
| chr6 | *SLC22A1* | 3 | 144 | 9 | SLC22A1 | solute carrier family 22 member 1 | Plasma Membrane | transporter | |
| chr17 | *AC087294.2* | 4 | 164 | 9 | | | | | |
| chr15 | *ARRDC4* | 4 | 109 | 9 | ARRDC4 | arrestin domain containing 4 | Plasma Membrane | other | |
| chr11 | *SIPA1* | 5 | 312 | 9 | SIPA1 | signal-induced proliferation-associated 1 | Cytoplasm | other | |
| chr17 | *KRT222* | 3 | 73 | 9 | KRT222 | keratin 222 | Other | other | |
| chr10 | *LINC00959* | 4 | 91 | 9 | C10orf143 | chromosome 10 open reading frame 143 | Other | other | |

| | | | | | | | | | |
|---|---|---|---|---|---|---|---|---|---|
| chr1 | *HES2* | 4 | 221 | 9 | HES2 | hes family bHLH transcription | Nucleus | transcription regulator | |
| chr1 | *RP11-98G7.1* | 4 | 143 | 9 | | | | | |
| chr7 | *BCAP29* | 3 | 77 | 9 | BCAP29 | B cell receptor associated protein | Cytoplasm | transporter | |
| chr15 | *MTMR10* | 4 | 97 | 9 | MTMR10 | myotubularin related protein 10 | Cytoplasm | other | |
| chr17 | *HEATR6* | 4 | 83 | 9 | HEATR6 | HEAT repeat containing 6 | Other | other | |
| chr6 | *ZUFSP* | 3 | 113 | 9 | ZUP1 | zinc finger containing ubiquitin peptidase 1 | Other | other | |
| chr7 | *PSPHP1* | 4 | 132 | 9 | PSPHP1 | phosphoserine phosphatase pseudogene 1 | Other | other | |
| chrX | *GLOD5* | 4 | 95 | 9 | GLOD5 | glyoxalase domain containing 5 | Other | other | |
| chr12 | *FOXJ2* | 4 | 88 | 9 | FOXJ2 | forkhead box J2 | Nucleus | transcription regulator | |
| chr1 | *ZBTB48* | 4 | 227 | 9 | ZBTB48 | zinc finger and BTB domain containing 48 | Nucleus | transcription regulator | |
| chr4 | *TADA2B* | 4 | 171 | 9 | TADA2B | transcriptional adaptor 2B | Nucleus | transcription regulator | |
| chr1 | *RP11-439A17.4* | 3 | 159 | 9 | | | | | |
| chr13 | *TNFSF13B* | 3 | 108 | 9 | TNFSF13B | TNF superfamily member 13b | Extracellular Space | cytokine | tabalumab, |
| chr18 | *PQLC1* | 4 | 158 | 9 | SLC66A2 | solute carrier family 66 member 2 | Other | other | |
| chr3 | *RP11-93K22.6* | 5 | 148 | 9 | | | | | |
| chr8 | *ANXA13* | 3 | 75 | 9 | ANXA13 | annexin A13 | Plasma Membrane | other | |
| chr2 | *CAPN10* | 5 | 221 | 9 | CAPN10 | calpain 10 | Cytoplasm | peptidase | |
| chr10 | *ERLIN1* | 4 | 76 | 9 | ERLIN1 | ER lipid raft associated 1 | Plasma Membrane | other | |
| chr2 | *STAT1* | 4 | 73 | 9 | STAT1 | signal transducer and activator of transcription 1 | Nucleus | transcription regulator | |
| chr4 | *RP11-386B13.4* | 4 | 125 | 9 | | | | | |
| chr11 | *CTD-2140G10.2* | 4 | 110 | 9 | | | | | |
| chr2 | *TTC31* | 4 | 128 | 9 | TTC31 | tetratricopeptide repeat domain | Other | other | |
| chr11 | *HBG2* | 4 | 145 | 9 | HBG2 | hemoglobin subunit gamma 2 | Cytoplasm | other | |
| chr5 | *CTC-552D5.1* | 4 | 80 | 9 | | | | | |
| chr18 | *KCTD1* | 3 | 85 | 9 | KCTD1 | potassium channel tetramerization domain containing | Nucleus | transcription regulator | |
| chr17 | *HEXIM2* | 4 | 160 | 9 | HEXIM2 | HEXIM P-TEFb complex subunit | Nucleus | transcription regulator | |
| chr1 | *RP11-111I12.1* | 4 | 113 | 9 | | | | | |
| chr19 | *CEACAM3* | 4 | 119 | 9 | CEACAM3 | CEA cell adhesion molecule 3 | Other | other | |
| chr10 | *UBE2D1* | 5 | 120 | 9 | UBE2D1 | ubiquitin conjugating enzyme E2 | Cytoplasm | enzyme | |
| chr7 | *ZNF212* | 4 | 107 | 9 | ZNF212 | zinc finger protein 212 | Nucleus | transcription regulator | |
| chr17 | *UBE2Z* | 4 | 95 | 9 | UBE2Z | ubiquitin conjugating enzyme E2 | Nucleus | enzyme | |
| chr6 | *CLPSL1* | 4 | 125 | 9 | CLPSL1 | colipase like 1 | Other | other | |
| chr12 | *RP3-416H24.4* | 4 | 224 | 9 | | | | | |
| chr12 | *GIT2* | 4 | 111 | 9 | GIT2 | GIT ArfGAP 2 | Nucleus | other | |
| chr15 | *WDR72* | 3 | 101 | 9 | WDR72 | WD repeat domain 72 | Cytoplasm | other | |
| chr5 | *CTD-2215L10.1* | 4 | 232 | 9 | | | | | |
| chr6 | *RP11-428F8.2* | 3 | 69 | 9 | | | | | |
| chr1 | *SHC1* | 4 | 197 | 9 | SHC1 | SHC adaptor protein 1 | Cytoplasm | other | |
| chr1 | *MTMR9LP* | 4 | 121 | 9 | MTMR9LP | myotubularin related protein 9 like, pseudogene | Other | other | |
| chr16 | *CARHSP1* | 4 | 150 | 9 | CARHSP1 | calcium regulated heat stable protein 1 | Cytoplasm | transcription regulator | |
| chr18 | *RP11-683L23.1* | 4 | 148 | 9 | | | | | |
| chr12 | *HMGA2* | 4 | 89 | 9 | HMGA2 | high mobility group AT-hook 2 | Nucleus | enzyme | |
| chr1 | *S100A2* | 4 | 172 | 9 | S100A2 | S100 calcium binding protein A2 | Nucleus | other | |
| chr16 | *LINC00514* | 4 | 184 | 9 | LINC00514 | long intergenic non-protein coding RNA 514 | Other | other | |
| chr15 | *EMC4* | 3 | 100 | 9 | EMC4 | ER membrane protein complex subunit 4 | Cytoplasm | other | |
| chr17 | *AC126365.1* | 3 | 71 | 9 | | | | | |
| chr12 | *XPOT* | 4 | 99 | 9 | XPOT | exportin for tRNA | Nucleus | other | |
| chr2 | *AC114765.1* | 3 | 78 | 9 | | | | | |
| chr17 | *KAT2A* | 4 | 200 | 9 | KAT2A | lysine acetyltransferase 2A | Cytoplasm | enzyme | |
| chr7 | *AC007255.8* | 4 | 125 | 9 | | | | | |
| chr3 | *RP11-90K6.1* | 3 | 75 | 9 | | | | | |
| chr5 | *ERAP1* | 3 | 72 | 9 | ERAP1 | endoplasmic reticulum aminopeptidase 1 | Extracellular Space | peptidase | |
| chr5 | *ITK* | 4 | 96 | 9 | ITK | IL2 inducible T cell kinase | Cytoplasm | kinase | pazopanib, crizotinib/pazopanib, lapatinib/pazopanib |
| chr10 | *RP11-445P17.3* | 3 | 138 | 9 | | | | | |
| chr12 | *RP11-551L14.4* | 4 | 198 | 9 | | | | | |
| chr7 | *AC005682.6* | 4 | 131 | 9 | | | | | |
| chr13 | *TNFRSF19* | 3 | 69 | 9 | TNFRSF19 | TNF receptor superfamily | Plasma Membrane | transmembrane receptor | |
| chr4 | *RP11-148L24.1* | 4 | 112 | 9 | | | | | |
| chr12 | *HAL* | 4 | 79 | 9 | HAL | histidine ammonia-lyase | Cytoplasm | enzyme | |
| chr15 | *RP11-69H7.2* | 4 | 207 | 9 | | | | | |
| chr11 | *RP11-626H12.3* | 4 | 133 | 9 | | | | | |
| chr3 | *LSG1* | 4 | 84 | 9 | LSG1 | large 60S subunit nuclear export GTPase 1 | Cytoplasm | enzyme | |
| chr19 | *CTB-180A7.8* | 4 | 123 | 9 | | | | | |

| | | | | | | | | | |
|---|---|---|---|---|---|---|---|---|---|
| chr15 | *CHRM5* | 3 | 84 | 9 | CHRM5 | cholinergic receptor muscarinic 5 | Plasma Membrane | G-protein coupled recepto | dexamethasone/olanzapine, dicyclomine, cevimeline, mirabegron/solifenacin, olanzapine, aclidinium/formoterol, acetylcholine, atropine/pralidoxime, brompheniramine, umeclidinium, mepenzolic acid, propantheline, orphenadrine, aspirin/caffeine/orphenadrine, tolterodine, chlorpheniramine/methscopolamine/phenylephrine, trihexyphenidyl, oxybutynin, formoterol/glycopyrrolate, scopolamine, arecoline, [illegible] |
| chr3 | *TAGLN3* | 4 | 157 | 9 | TAGLN3 | transgelin 3 | Extracellular Space | other | |
| chr17 | *C17orf64* | 4 | 135 | 9 | C17orf64 | chromosome 17 open reading frame 64 | Other | other | |
| chr15 | *ULK3* | 4 | 171 | 9 | ULK3 | unc-51 like kinase 3 | Cytoplasm | kinase | |
| chr10 | *ZSWIM8* | 4 | 313 | 9 | ZSWIM8 | zinc finger SWIM-type containing | Extracellular Space | other | |
| chr7 | *STAG3L4* | 3 | 87 | 9 | STAG3L4 | stromal antigen 3-like 4 | Other | other | |
| chr14 | *BRMS1L* | 3 | 78 | 9 | BRMS1L | BRMS1 like transcriptional | Other | other | |
| chr3 | *CD86* | 4 | 115 | 9 | CD86 | CD86 molecule | Plasma Membrane | transmembrane recepto | belatacept, abatacept, |
| chr5 | *RP11-167P20.1* | 4 | 116 | 9 | | | | | |
| chr2 | *THNSL2* | 4 | 124 | 9 | THNSL2 | threonine synthase like 2 | Other | other | |
| chr16 | *SCNN1G* | 4 | 141 | 9 | SCNN1G | sodium channel epithelial 1 gamma subunit | Plasma Membrane | ion channel | hydrochlorothiazide/triamterene, amiloride, triamterene, amiloride/hydrochlorothiazide |
| chr5 | *CTB-49A3.2* | 4 | 85 | 9 | CTB-49A3.2 | | Other | other | |
| chr16 | *CRYM-AS1* | 4 | 119 | 9 | CRYM-AS1 | CRYM antisense RNA 1 | Other | other | |
| chr2 | *WDR43* | 3 | 69 | 9 | WDR43 | WD repeat domain 43 | Nucleus | transcription regulator | |
| chr15 | *RP11-1008C21.1* | 4 | 81 | 9 | | | | | |
| chr7 | *NUDT1* | 4 | 177 | 9 | NUDT1 | nudix hydrolase 1 | Extracellular Space | phosphatase | karonudib |
| chr1 | *BROX* | 3 | 97 | 9 | BROX | BRO1 domain and CAAX motif containing | Cytoplasm | other | |
| chr2 | *GPC1* | 5 | 242 | 9 | GPC1 | glypican 1 | Plasma Membrane | transmembrane recepto | |
| chr19 | *PALM3* | 4 | 151 | 9 | PALM3 | paralemmin 3 | Other | other | |
| chr3 | *PLSCR2* | 4 | 146 | 9 | PLSCR2 | phospholipid scramblase 2 | Other | other | |
| chr14 | *GTF2A1* | 3 | 87 | 9 | GTF2A1 | general transcription factor IIA subunit 1 | Cytoplasm | transcription regulator | |
| chr11 | *SYVN1* | 4 | 215 | 9 | SYVN1 | synoviolin 1 | Cytoplasm | transporter | |
| chr7 | *PMS2P4* | 4 | 97 | 9 | PMS2P4 | PMS1 homolog 2, mismatch repair system component | Other | other | |
| chr3 | *CX3CR1* | 4 | 116 | 9 | CX3CR1 | C-X3-C motif chemokine receptor | Plasma Membrane | G-protein coupled recepto | |
| chr15 | *SPATA31E2P* | 3 | 256 | 9 | SPATA31E2P | SPATA31 subfamily E member 2, pseudogene | Other | other | |
| chr3 | *TMEM39A* | 4 | 129 | 9 | TMEM39A | transmembrane protein 39A | Other | other | |
| chr2 | *SEPT2* | 3 | 73 | 9 | SEPTIN2 | septin 2 | Cytoplasm | enzyme | |
| chr9 | *INVS* | 3 | 89 | 9 | INVS | inversin | Nucleus | transcription regulator | |
| chr7 | *WASL* | 3 | 82 | 9 | WASL | WASP like actin nucleation promoting factor | Cytoplasm | other | |
| chr21 | *LINC00649* | 4 | 96 | 9 | LINC00649 | long intergenic non-protein coding RNA 649 | Other | other | |
| chr9 | *ZMYND19* | 4 | 345 | 9 | ZMYND19 | zinc finger MYND-type containing | Plasma Membrane | other | |
| chr12 | *VPS29* | 3 | 96 | 9 | VPS29 | VPS29 retromer complex | Cytoplasm | transporter | |
| chr22 | *MMP11* | 4 | 161 | 9 | MMP11 | matrix metallopeptidase 11 | Extracellular Space | peptidase | marimastat |
| chr19 | *MYPOP* | 4 | 97 | 9 | MYPOP | Myb related transcription factor, partner of profilin | Nucleus | transcription regulator | |
| chr8 | *RNF170* | 3 | 81 | 9 | RNF170 | ring finger protein 170 | Other | other | |
| chr1 | *ZSCAN20* | 4 | 139 | 9 | ZSCAN20 | zinc finger and SCAN domain containing 20 | Nucleus | transcription regulator | |
| chr1 | *FHL3* | 4 | 158 | 9 | FHL3 | four and a half LIM domains 3 | Plasma Membrane | other | |
| chr9 | *NAIF1* | 4 | 211 | 9 | NAIF1 | nuclear apoptosis inducing factor | Nucleus | other | |
| chr3 | *GYG1* | 5 | 98 | 9 | GYG1 | glycogenin 1 | Cytoplasm | enzyme | |
| chr21 | *U2AF1* | 3 | 77 | 9 | U2AF1/U2AF1L5 | U2 small nuclear RNA auxiliary factor 1 | Nucleus | other | |
| chr17 | *WDR81* | 4 | 132 | 9 | WDR81 | WD repeat domain 81 | Plasma Membrane | other | |
| chr12 | *TCP11L2* | 4 | 78 | 9 | TCP11L2 | t-complex 11 like 2 | Other | other | |
| chr11 | *SF1* | 4 | 161 | 9 | SF1 | splicing factor 1 | Nucleus | transcription regulator | |
| chr1 | *RP11-293P20.2* | 4 | 80 | 9 | | | | | |
| chr8 | *RP11-62H7.2* | 4 | 86 | 9 | | | | | |
| chr4 | *FBXW7* | 3 | 70 | 9 | FBXW7 | F-box and WD repeat domain containing 7 | Nucleus | enzyme | |
| chr1 | *ELK4* | 4 | 117 | 9 | ELK4 | ETS transcription factor ELK4 | Nucleus | transcription regulator | |
| chr11 | *TP53AIP1* | 4 | 231 | 9 | TP53AIP1 | tumor protein p53 regulated apoptosis inducing protein 1 | Cytoplasm | other | |
| chr2 | *NAGK* | 3 | 114 | 9 | NAGK | N-acetylglucosamine kinase | Cytoplasm | kinase | |

| | | | | | | | | | |
|---|---|---|---|---|---|---|---|---|---|
| chr6 | *C6orf52* | 4 | 83 | 9 | C6orf52 | chromosome 6 open reading frame 52 | Other | other | |
| chr1 | *UCHL5* | 3 | 103 | 9 | UCHL5 | ubiquitin C-terminal hydrolase L5 | Cytoplasm | peptidase | VLX1570 |
| chr7 | *CCDC136* | 4 | 122 | 9 | CCDC136 | coiled-coil domain containing 136 | Cytoplasm | other | |
| chr1 | *RP11-744H18.1* | 4 | 117 | 9 | | | | | |
| chr19 | *AC002511.1* | 4 | 94 | 9 | | | | | |
| chr15 | *RP11-720L8.1* | 3 | 106 | 9 | | | | | |
| chr1 | *OTUD3* | 4 | 97 | 9 | OTUD3 | OTU deubiquitinase 3 | Cytoplasm | peptidase | |
| chr2 | *KIAA1715* | 3 | 78 | 9 | LNPK | lunapark, ER junction formation | Cytoplasm | other | |
| chr6 | *TREML2* | 4 | 135 | 9 | TREML2 | triggering receptor expressed on myeloid cells like 2 | Plasma Membrane | other | |
| chr2 | *AC109826.1* | 4 | 131 | 9 | | | | | |
| chr14 | *KTN1-AS1* | 3 | 76 | 9 | KTN1-AS1 | KTN1 antisense RNA 1 | Other | other | |
| chr11 | *LDHAL6A* | 4 | 90 | 9 | LDHAL6A | lactate dehydrogenase A like 6A | Cytoplasm | enzyme | |
| chr13 | *MICU2* | 3 | 74 | 9 | MICU2 | mitochondrial calcium uptake 2 | Cytoplasm | other | |
| chr1 | *ZBTB41* | 4 | 83 | 9 | ZBTB41 | zinc finger and BTB domain containing 41 | Other | other | |
| chr11 | *PPP1CA* | 4 | 178 | 9 | PPP1CA | protein phosphatase 1 catalytic subunit alpha | Cytoplasm | phosphatase | |
| chr2 | *GCKR* | 3 | 83 | 9 | GCKR | glucokinase regulator | Nucleus | other | |
| chr15 | *PDCD7* | 4 | 70 | 9 | PDCD7 | programmed cell death 7 | Nucleus | other | |
| chr3 | *NCEH1* | 3 | 77 | 9 | NCEH1 | neutral cholesterol ester | Plasma Membrane | enzyme | |
| chr15 | *MAN2C1* | 4 | 172 | 9 | MAN2C1 | mannosidase alpha class 2C member 1 | Cytoplasm | enzyme | |
| chr10 | *ITGB1* | 4 | 110 | 9 | ITGB1 | integrin subunit beta 1 | Plasma Membrane | transmembrane receptor | |
| chr14 | *RP11-543C4.1* | 3 | 123 | 9 | | | | | |
| chr5 | *AC008697.1* | 4 | 97 | 9 | | | | | |
| chr16 | *CTD-2313J23.1* | 4 | 75 | 8 | | | | | |
| chr5 | *RP11-195E2.1* | 4 | 91 | 8 | | | | | |
| chr17 | *LINC00910* | 4 | 1431 | 8 | LINC00910 | long intergenic non-protein coding RNA 910 | Other | other | |
| chrX | *PNPLA4* | 3 | 76 | 8 | PNPLA4 | patatin like phospholipase domain containing 4 | Cytoplasm | enzyme | |
| chr7 | *NOBOX* | 4 | 146 | 8 | NOBOX | NOBOX oogenesis homeobox | Nucleus | transcription regulator | |
| chr17 | *AC004448.5* | 4 | 87 | 8 | | | | | |
| chr17 | *SGSH* | 4 | 200 | 8 | SGSH | N-sulfoglucosamine | Cytoplasm | enzyme | |
| chr16 | *RP11-876N24.2* | 4 | 115 | 8 | | | | | |
| chr8 | *EIF4EBP1* | 3 | 120 | 8 | EIF4EBP1 | eukaryotic translation initiation factor 4E binding protein 1 | Cytoplasm | translation regulator | |
| chr5 | *CTC-436P18.3* | 4 | 103 | 8 | | | | | |
| chr5 | *CTC-329D1.3* | 4 | 106 | 8 | | | | | |
| chr1 | *PDIK1L* | 4 | 99 | 8 | PDIK1L | PDLIM1 interacting kinase 1 like | Nucleus | kinase | |
| chr9 | *RP11-251M1.1* | 5 | 351 | 8 | | | | | |
| chr11 | *RP11-196E1.3* | 4 | 112 | 8 | | | | | |
| chr5 | *ARL10* | 4 | 110 | 8 | ARL10 | ADP ribosylation factor like GTPase 10 | Other | other | |
| chr11 | *EIF3M* | 4 | 126 | 8 | EIF3M | eukaryotic translation initiation factor 3 subunit M | Cytoplasm | other | |
| chr11 | *FAM86GP* | 4 | 370 | 8 | FAM86GP | family with sequence similarity 86, member A pseudogene | Other | other | |
| chr7 | *TBL2* | 4 | 174 | 8 | TBL2 | transducin beta like 2 | Plasma Membrane | other | |
| chr2 | *ACVR1* | 3 | 89 | 8 | ACVR1 | activin A receptor type 1 | Plasma Membrane | kinase | |
| chr1 | *SNRPE* | 4 | 128 | 8 | SNRPE | small nuclear ribonucleoprotein polypeptide E | Nucleus | other | |
| chr1 | *CAMK1G* | 4 | 150 | 8 | CAMK1G | calcium/calmodulin dependent protein kinase IG | Cytoplasm | kinase | |
| chr11 | *KCNJ1* | 4 | 165 | 8 | KCNJ1 | potassium inwardly rectifying channel subfamily J member 1 | Plasma Membrane | ion channel | glimepiride/pioglitazone, glimepiride/rosiglitazone, glimepiride, acetohexamide, |
| chr3 | *C3orf52* | 4 | 116 | 8 | C3orf52 | chromosome 3 open reading frame 52 | Other | other | |
| chr10 | *IL9RP2* | 4 | 155 | 8 | IL9RP2 | interleukin 9 receptor | Other | other | |
| chr14 | *TDP1* | 3 | 67 | 8 | TDP1 | tyrosyl-DNA phosphodiesterase 1 | Nucleus | enzyme | |
| chr20 | *R3HDML* | 4 | 158 | 8 | R3HDML | R3H domain containing like | Other | other | |
| chr3 | *TRIM42* | 4 | 166 | 8 | TRIM42 | tripartite motif containing 42 | Other | other | |
| chr16 | *RP11-719K4.3* | 4 | 133 | 8 | | | | | |
| chr1 | *CDCA8* | 4 | 110 | 8 | CDCA8 | cell division cycle associated 8 | Nucleus | other | |
| chr17 | *CACNG1* | 4 | 126 | 8 | CACNG1 | calcium voltage-gated channel auxiliary subunit gamma 1 | Plasma Membrane | ion channel | diltiazem/enalapril, lercanidipin, nitrendipine, |
| chr19 | *FAM90A28P* | 4 | 134 | 8 | FAM90A28P | family with sequence similarity 90 member A28, pseudogene | Other | other | |
| chr3 | *RP11-475N22.4* | 4 | 201 | 8 | | | | | |
| chr1 | *TRMT1L* | 4 | 87 | 8 | TRMT1L | tRNA methyltransferase 1 like | Other | enzyme | |
| chr7 | *BLACE* | 3 | 112 | 8 | BLACE | B cell acute lymphoblastic leukemia expressed | Other | other | |
| chr19 | *ANO8* | 4 | 184 | 8 | ANO8 | anoctamin 8 | Plasma Membrane | ion channel | |
| chr17 | *CTD-2047H16.4* | 4 | 106 | 8 | | | | | |
| chr16 | *NPIPA5* | 4 | 99 | 8 | PA1 (includes oth | nuclear pore complex interacting protein family member A3 | Nucleus | other | |
| chr1 | *SLC16A4* | 3 | 94 | 8 | SLC16A4 | solute carrier family 16 member 4 | Plasma Membrane | transporter | |
| chr1 | *CD160* | 4 | 131 | 8 | CD160 | CD160 molecule | Plasma Membrane | transmembrane receptor | |
| chr1 | *PLEKHO1* | 4 | 139 | 8 | PLEKHO1 | pleckstrin homology domain containing O1 | Plasma Membrane | other | |

| | | | | | | | | | |
|---|---|---|---|---|---|---|---|---|---|
| chr12 | *PRIM1* | 4 | 92 | 8 | PRIM1 | DNA primase subunit 1 | Nucleus | enzyme | alemtuzumab/cyclophosphamide/fludarabine phosphate, cytarabine/filgrastim/fludarabine phosphate/imatinib, fludarabine phosphate/rituximab, cytarabine/fludarabine phosphate/oxaliplatin/rituximab, dexamethasone/fludarabine phosphate/mitoxantrone/rituximab, cytarabine/fludarabine phosphate, cytarabine/filgrastim/fludarabine phosphate, cyclophosphamide/fludarabine [illegible] |
| chr11 | *RNH1* | 4 | 199 | 8 | RNH1 | ribonuclease/angiogenin inhibitor | Cytoplasm | other | |
| chr19 | *GIPR* | 4 | 150 | 8 | GIPR | gastric inhibitory polypeptide | Plasma Membrane | G-protein coupled recepto | |
| chr15 | *PDIA3* | 3 | 86 | 8 | PDIA3 | protein disulfide isomerase family A member 3 | Cytoplasm | peptidase | |
| chr8 | *TMEM66* | 3 | 111 | 8 | SARAF | store-operated calcium entry associated regulatory factor | Cytoplasm | other | |
| chr10 | *TYSND1* | 4 | 198 | 8 | TYSND1 | trypsin domain containing 1 | Cytoplasm | peptidase | |
| chr18 | *RP11-173L6.1* | 4 | 133 | 8 | | | | | |
| chr4 | *CTBP1* | 5 | 279 | 8 | CTBP1 | C-terminal binding protein 1 | Nucleus | enzyme | |
| chr5 | *RANBP3L* | 4 | 206 | 8 | RANBP3L | RAN binding protein 3 like | Other | other | |
| chr10 | *ENTPD1-AS1* | 4 | 87 | 8 | ENTPD1-AS1 | ENTPD1 antisense RNA 1 | Other | other | |
| chr3 | *BRK1* | 4 | 110 | 8 | BRK1 | BRICK1 subunit of SCAR/WAVE actin nucleating complex | Cytoplasm | other | |
| chr19 | *C19orf54* | 4 | 209 | 8 | C19orf54 | chromosome 19 open reading frame 54 | Other | other | |
| chr19 | *CTD-2525I3.6* | 3 | 85 | 8 | | | | | |
| chr2 | *NCL* | 4 | 162 | 8 | NCL | nucleolin | Nucleus | other | AGRO 100 |
| chr1 | *HDGF* | 4 | 207 | 8 | HDGF | heparin binding growth factor | Extracellular Space | growth factor | |
| chr16 | *RP11-1021N1.1* | 4 | 85 | 8 | | | | | |
| chr1 | *RP11-122M14.1* | 4 | 149 | 8 | | | | | |
| chr1 | *RP4-784A16.5* | 4 | 124 | 8 | | | | | |
| chr7 | *ATG9B* | 4 | 170 | 8 | ATG9B | autophagy related 9B | Cytoplasm | other | |
| chr19 | *ZNF575* | 4 | 122 | 8 | ZNF575 | zinc finger protein 575 | Nucleus | transcription regulator | |
| chr22 | *POM121L4P* | 3 | 155 | 8 | POM121L4P | POM121 transmembrane nucleoporin like 4, pseudogene | Other | other | |
| chr7 | *RABL5* | 4 | 128 | 8 | IFT22 | intraflagellar transport 22 | Extracellular Space | other | |
| chr11 | *LIN7C* | 3 | 159 | 8 | LIN7C | lin-7 homolog C, crumbs cell polarity complex component | Cytoplasm | other | |
| chr18 | *RP11-794M8.1* | 4 | 84 | 8 | | | | | |
| chr17 | *SARM1* | 4 | 152 | 8 | SARM1 | sterile alpha and TIR motif containing 1 | Plasma Membrane | transmembrane recepto | |
| chr5 | *RP11-779O18.3* | 4 | 318 | 8 | | | | | |
| chr11 | *RP11-262A12.1* | 4 | 71 | 8 | | | | | |
| chr2 | *PPIG* | 3 | 84 | 8 | PPIG | peptidylprolyl isomerase G | Nucleus | enzyme | |
| chr20 | *RP1-155G6.4* | 4 | 134 | 8 | | | | | |
| chr1 | *SEPN1* | 4 | 163 | 8 | SELENON | selenoprotein N | Cytoplasm | other | |
| chr5 | *RP11-357F12.1* | 4 | 176 | 8 | | | | | |
| chr10 | *RP11-324L3.1* | 4 | 110 | 8 | | | | | |
| chr1 | *DR1* | 4 | 91 | 8 | DR1 | down-regulator of transcription 1 | Nucleus | transcription regulator | |
| chr19 | *SLC1A5* | 3 | 120 | 8 | SLC1A5 | solute carrier family 1 member 5 | Plasma Membrane | transporter | |
| chr12 | *RP11-689B22.2* | 4 | 104 | 8 | | | | | |
| chr22 | *HMOX1* | 3 | 116 | 8 | HMOX1 | heme oxygenase 1 | Cytoplasm | enzyme | tin mesoporphyrin |
| chr17 | *CTD-2378E21.1* | 4 | 72 | 8 | | | | | |
| chr13 | *RP11-394A14.2* | 4 | 94 | 8 | | | | | |
| chr1 | *TADA1* | 3 | 68 | 8 | TADA1 | transcriptional adaptor 1 | Nucleus | transcription regulator | |
| chr16 | *RP11-434E6.2* | 4 | 71 | 8 | | | | | |
| chr1 | *RP11-496N12.6* | 4 | 277 | 8 | | | | | |
| chr14 | *NRL* | 3 | 94 | 8 | NRL | neural retina leucine zipper | Nucleus | transcription regulator | |
| chr7 | *AC004895.4* | 4 | 122 | 8 | | | | | |
| chr17 | *CPSF4L* | 4 | 144 | 8 | CPSF4L | cleavage and polyadenylation specific factor 4 like | Other | other | |
| chr2 | *ACSL3* | 4 | 68 | 8 | ACSL3 | acyl-CoA synthetase long chain family member 3 | Cytoplasm | enzyme | |
| chr17 | *RP11-357H14.17* | 4 | 92 | 8 | | | | | |
| chr16 | *KIAA0430* | 3 | 93 | 8 | MARF1 | meiosis regulator and mRNA stability factor 1 | Cytoplasm | other | |
| chr13 | *LINC00383* | 4 | 69 | 8 | LINC00383 | long intergenic non-protein coding RNA 383 | Other | other | |
| chr10 | *UCN3* | 3 | 86 | 8 | UCN3 | urocortin 3 | Extracellular Space | other | |
| chrX | *EGFL6* | 4 | 79 | 8 | EGFL6 | EGF like domain multiple 6 | Extracellular Space | other | |
| chr6 | *RP11-354I10.1* | 3 | 67 | 8 | | | | | |
| chr19 | *ISOC2* | 4 | 178 | 8 | ISOC2 | isochorismatase domain | Cytoplasm | enzyme | |
| chr19 | *PSG11* | 4 | 93 | 8 | PSG11 | pregnancy specific beta-1-glycoprotein 11 | Extracellular Space | other | |
| chr15 | *RP11-814P5.1* | 3 | 109 | 8 | | | | | |

| | | | | | | | | | |
|---|---|---|---|---|---|---|---|---|---|
| chr10 | *PLXDC2* | 3 | 88 | 8 | PLXDC2 | plexin domain containing 2 | Extracellular Space | other | |
| chr11 | *RP11-677M14.7* | 4 | 90 | 8 | | | | | |
| chr12 | *TMBIM4* | 4 | 108 | 8 | TMBIM4 | transmembrane BAX inhibitor motif containing 4 | Nucleus | other | |
| chr12 | *RAB21* | 3 | 87 | 8 | RAB21 | RAB21, member RAS oncogene family | Cytoplasm | enzyme | |
| chr3 | *AC136289.1* | 3 | 69 | 8 | | | | | |
| chr5 | *LINC00461* | 4 | 66 | 8 | LINC00461 | long intergenic non-protein coding RNA 461 | Other | other | |
| chr20 | *GID8* | 3 | 278 | 8 | GID8 | GID complex subunit 8 homolog | Nucleus | other | |
| chr2 | *AC009312.1* | 4 | 156 | 8 | | | | | |
| chr7 | *TSPAN12* | 4 | 93 | 8 | TSPAN12 | tetraspanin 12 | Plasma Membrane | other | |
| chr12 | *LTBR* | 4 | 149 | 8 | LTBR | lymphotoxin beta receptor | Plasma Membrane | transmembrane receptor | |
| chr12 | *RP11-150C16.1* | 3 | 93 | 8 | | | | | |
| chr17 | *DUSP3* | 4 | 152 | 8 | DUSP3 | dual specificity phosphatase 3 | Cytoplasm | phosphatase | |
| chr20 | *MATN4* | 4 | 185 | 8 | MATN4 | matrilin 4 | Extracellular Space | other | |
| chr1 | *C1orf87* | 3 | 69 | 8 | C1orf87 | chromosome 1 open reading frame 87 | Extracellular Space | other | |
| chr11 | *EIF3F* | 4 | 118 | 8 | EIF3F | eukaryotic translation initiation factor 3 subunit F | Cytoplasm | translation regulator | |
| chr6 | *ORC3* | 3 | 81 | 8 | ORC3 | origin recognition complex | Nucleus | other | |
| chr12 | *NUDT4* | 4 | 72 | 8 | NUDT4 | nudix hydrolase 4 | Cytoplasm | phosphatase | |
| chr1 | *RPF1* | 3 | 88 | 8 | RPF1 | ribosome production factor 1 | Nucleus | other | |
| chr6 | *NUS1* | 3 | 76 | 8 | NUS1 | NUS1 dehydrodolichyl diphosphate synthase subunit | Cytoplasm | enzyme | |
| chr7 | *GIMAP5* | 4 | 96 | 8 | GIMAP5 | GTPase, IMAP family member 5 | Cytoplasm | other | |
| chr19 | *C19orf45* | 4 | 145 | 8 | TEX45 | testis expressed 45 | Other | other | |
| chr4 | *DCLK2* | 4 | 96 | 8 | DCLK2 | doublecortin like kinase 2 | Cytoplasm | kinase | |
| chr8 | *FAM183CP* | 4 | 81 | 8 | LINC02209 | long intergenic non-protein coding RNA 2209 | Other | other | |
| chr19 | *FUZ* | 4 | 124 | 8 | FUZ | fuzzy planar cell polarity protein | Plasma Membrane | other | |
| chr3 | *QTRTD1* | 4 | 108 | 8 | QTRT2 | queuine tRNA-ribosyltransferase accessory subunit 2 | Cytoplasm | enzyme | |
| chr6 | *RP3-404K8.2* | 4 | 69 | 8 | | | | | |
| chr17 | *FAM57A* | 4 | 79 | 8 | TLCD3A | TLC domain containing 3A | Plasma Membrane | other | |
| chr10 | *DUSP13* | 4 | 249 | 8 | DUSP13 | dual specificity phosphatase 13 | Cytoplasm | phosphatase | |
| chr2 | *OSBPL6* | 3 | 62 | 8 | OSBPL6 | oxysterol binding protein like 6 | Cytoplasm | other | |
| chr3 | *MRPS25* | 4 | 118 | 8 | MRPS25 | mitochondrial ribosomal protein | Cytoplasm | other | |
| chr14 | *RP11-589M4.3* | 3 | 67 | 8 | | | | | |
| chr11 | *KLHL35* | 4 | 234 | 8 | KLHL35 | kelch like family member 35 | Other | other | |
| chr1 | *RP11-640M9.2* | 4 | 175 | 8 | | | | | |
| chr12 | *PTPRO* | 3 | 61 | 8 | PTPRO | protein tyrosine phosphatase receptor type O | Plasma Membrane | phosphatase | |
| chr7 | *WNT2* | 4 | 107 | 8 | WNT2 | Wnt family member 2 | Extracellular Space | cytokine | |
| chr19 | *PSG1* | 4 | 86 | 8 | PSG1 | pregnancy specific beta-1-glycoprotein 1 | Extracellular Space | other | |
| chr6 | *RP11-557H15.4* | 3 | 79 | 8 | | | | | |
| chr6 | *RP1-290I10.7* | 3 | 83 | 8 | | | | | |
| chr6 | *BAK1* | 3 | 159 | 8 | BAK1 | BCL2 antagonist/killer 1 | Cytoplasm | other | |
| chr4 | *LINC01088* | 3 | 94 | 8 | LINC01088 | long intergenic non-protein coding RNA 1088 | Other | other | |
| chrX | *RP11-348F1.2* | 4 | 97 | 8 | | | | | |
| chr12 | *SHMT2* | 4 | 164 | 8 | SHMT2 | serine hydroxymethyltransferase | Cytoplasm | enzyme | |
| chr2 | *ADAM23* | 3 | 71 | 8 | ADAM23 | ADAM metallopeptidase domain | Plasma Membrane | peptidase | |
| chr1 | *C1orf137* | 4 | 88 | 8 | C1orf137 | chromosome 1 open reading frame 137 (putative) | Other | other | |
| chr22 | *DESI1* | 3 | 82 | 8 | DESI1 | desumoylating isopeptidase 1 | Other | other | |
| chr3 | *DHX36* | 4 | 114 | 8 | DHX36 | DEAH-box helicase 36 | Cytoplasm | enzyme | |
| chr12 | *RP11-776A13.2* | 3 | 109 | 8 | | | | | |
| chr17 | *BZRAP1-AS1* | 4 | 158 | 8 | TSPOAP1-AS1 | TSPOAP1, SUPT4H1 and RNF43 antisense RNA 1 | Other | other | |
| chr8 | *ADAM2* | 3 | 80 | 8 | ADAM2 | ADAM metallopeptidase domain | Plasma Membrane | peptidase | |
| chr16 | *RP11-256I9.3* | 4 | 83 | 8 | | | | | |
| chr8 | *LYNX1* | 4 | 259 | 8 | LYNX1 | Ly6/neurotoxin 1 | Plasma Membrane | transporter | |
| chr13 | *EFNB2* | 4 | 99 | 8 | EFNB2 | ephrin B2 | Plasma Membrane | kinase | sEphB4-HSA |
| chr11 | *RAB39A* | 4 | 69 | 8 | RAB39A | RAB39A, member RAS oncogene family | Cytoplasm | enzyme | |
| chr10 | *RP11-275N1.1* | 3 | 72 | 8 | | | | | |
| chr16 | *PRSS27* | 4 | 224 | 8 | PRSS27 | serine protease 27 | Extracellular Space | peptidase | |
| chrX | *G6PD* | 4 | 129 | 8 | G6PD | glucose-6-phosphate | Cytoplasm | enzyme | |
| chr5 | *CTC-436P18.1* | 4 | 96 | 8 | | | | | |
| chr15 | *NARG2* | 3 | 84 | 8 | ICE2 | interactor of little elongation complex ELL subunit 2 | Nucleus | other | |
| chr19 | *KCNK6* | 4 | 117 | 8 | KCNK6 | potassium two pore domain channel subfamily K member 6 | Plasma Membrane | ion channel | |
| chr9 | *ERCC6L2* | 4 | 103 | 8 | ERCC6L2 | ERCC excision repair 6 like 2 | Other | enzyme | |
| chr22 | *AC002055.4* | 3 | 92 | 8 | | | | | |
| chr3 | *IMPG2* | 4 | 120 | 8 | IMPG2 | interphotoreceptor matrix proteoglycan 2 | Extracellular Space | other | |
| chrX | *RP5-964N17.1* | 4 | 75 | 8 | | | | | |
| chr16 | *TXNL4B* | 3 | 78 | 8 | TXNL4B | thioredoxin like 4B | Nucleus | enzyme | |
| chr20 | *MIR663A* | 4 | 141 | 8 | mir-663 | microRNA 663a | Cytoplasm | microRNA | |
| chr11 | *OTUB1* | 4 | 140 | 8 | OTUB1 | OTU deubiquitinase, ubiquitin aldehyde binding 1 | Cytoplasm | enzyme | |
| chr15 | *EIF3J* | 3 | 83 | 8 | EIF3J | eukaryotic translation initiation factor 3 subunit J | Cytoplasm | translation regulator | |

| | | | | | | | | | |
|---|---|---|---|---|---|---|---|---|---|
| chr2 | YWHAQ | 3 | 66 | 8 | YWHAQ | tyrosine 3-monooxygenase/tryptophan 5-monooxygenase activation protein theta | Cytoplasm | other | |
| chr3 | RP11-95M5.1 | 4 | 133 | 8 | | | | | |
| chr15 | AC103965.1 | 4 | 284 | 8 | | | | | |
| chr1 | SLC35D1 | 3 | 63 | 8 | SLC35D1 | solute carrier family 35 member | Cytoplasm | transporter | |
| chr8 | KCNB2 | 4 | 93 | 8 | KCNB2 | potassium voltage-gated channel subfamily B member 2 | Plasma Membrane | ion channel | dalfampridine |
| chr1 | RP11-114B7.6 | 3 | 123 | 8 | | | | | |
| chr10 | FAM21FP | 3 | 141 | 8 | FAM21FP | family with sequence similarity 21 member F, pseudogene | Other | other | |
| chrX | COL4A5 | 3 | 80 | 8 | COL4A5 | collagen type IV alpha 5 chain | Extracellular Space | other | collagenase clostridium |
| chr10 | RP11-399K21.11 | 5 | 145 | 8 | | | | | |
| chr18 | MPPE1 | 4 | 86 | 8 | MPPE1 | metallophosphoesterase 1 | Cytoplasm | enzyme | |
| chr1 | RP4-781K5.8 | 4 | 213 | 8 | | | | | |
| chr1 | TRIM17 | 4 | 235 | 8 | TRIM17 | tripartite motif containing 17 | Cytoplasm | enzyme | |
| chr11 | WEE1 | 4 | 108 | 8 | WEE1 | WEE1 G2 checkpoint kinase | Nucleus | kinase | adavosertib |
| chr1 | RP4-594I10.3 | 4 | 88 | 8 | | | | | |
| chr11 | LRP4-AS1 | 3 | 112 | 8 | LRP4-AS1 | LRP4 antisense RNA 1 | Other | other | |
| chr5 | SLC36A1 | 3 | 98 | 8 | SLC36A1 | solute carrier family 36 member 1 | Plasma Membrane | transporter | |
| chr11 | RP11-680F20.9 | 3 | 96 | 8 | | | | | |
| chr1 | CHTOP | 4 | 130 | 8 | CHTOP | chromatin target of PRMT1 | Nucleus | other | |
| chr16 | SLC6A10P | 4 | 212 | 8 | SLC6A10P | solute carrier family 6 member 10, pseudogene | Plasma Membrane | other | |
| chr16 | DPEP3 | 4 | 150 | 8 | DPEP3 | dipeptidase 3 | Other | peptidase | |
| chr8 | NRBP2 | 4 | 183 | 8 | NRBP2 | nuclear receptor binding protein 2 | Cytoplasm | kinase | |
| chr17 | NOTUM | 4 | 173 | 8 | NOTUM | notum, palmitoleoyl-protein carboxylesterase | Extracellular Space | enzyme | |
| chr12 | OR8S1 | 4 | 223 | 8 | OR8S1 | olfactory receptor family 8 subfamily S member 1 | Plasma Membrane | other | |
| chr15 | RP11-463I20.3 | 4 | 98 | 8 | | | | | |
| chr15 | GOLGA6D | 3 | 258 | 8 | GA6A (includes ot | golgin A6 family member A | Other | other | |
| chr8 | CTD-2292P10.4 | 4 | 110 | 8 | | | | | |
| chr2 | FAM132B | 4 | 197 | 8 | ERFE | erythroferrone | Extracellular Space | other | |
| chr10 | PNLIP | 3 | 64 | 8 | PNLIP | pancreatic lipase | Extracellular Space | enzyme | orlistat |
| chr7 | ANKRD7 | 3 | 93 | 8 | ANKRD7 | ankyrin repeat domain 7 | Nucleus | transcription regulator | |
| chr12 | SYT10 | 3 | 72 | 8 | SYT10 | synaptotagmin 10 | Cytoplasm | transporter | |
| chr21 | FAM3B | 3 | 125 | 8 | FAM3B | family with sequence similarity 3 member B | Extracellular Space | cytokine | |
| chr7 | AC004870.3 | 4 | 99 | 8 | | | | | |
| chr3 | DUSP7 | 4 | 190 | 8 | DUSP7 | dual specificity phosphatase 7 | Cytoplasm | phosphatase | |
| chr10 | ITIH2 | 3 | 103 | 8 | ITIH2 | inter-alpha-trypsin inhibitor heavy chain 2 | Extracellular Space | other | |
| chr1 | SYPL2 | 4 | 156 | 8 | SYPL2 | synaptophysin like 2 | Other | other | |
| chr2 | SULT1C4 | 4 | 116 | 8 | SULT1C4 | sulfotransferase family 1C | Cytoplasm | enzyme | |
| chr19 | DENND1C | 4 | 155 | 8 | DENND1C | DENN domain containing 1C | Cytoplasm | other | |
| chr11 | WTAPP1 | 4 | 97 | 8 | WTAPP1 | Wilms tumor 1 associated protein pseudogene 1 | Other | other | |
| chr11 | RP4-541C22.5 | 4 | 115 | 8 | | | | | |
| chr3 | PIGZ | 4 | 111 | 8 | PIGZ | phosphatidylinositol glycan anchor biosynthesis class Z | Cytoplasm | enzyme | |
| chr19 | AP2S1 | 4 | 151 | 8 | AP2S1 | adaptor related protein complex 2 subunit sigma 1 | Cytoplasm | transporter | |
| chr20 | ARFRP1 | 4 | 246 | 8 | ARFRP1 | ADP ribosylation factor related protein 1 | Cytoplasm | enzyme | |
| chr2 | STK39 | 4 | 66 | 8 | STK39 | serine/threonine kinase 39 | Nucleus | kinase | |
| chr3 | COPB2 | 3 | 83 | 8 | COPB2 | coatomer protein complex subunit beta 2 | Cytoplasm | transporter | |
| chr17 | KRT17P2 | 4 | 227 | 8 | KRT17P2 | keratin 17 pseudogene 2 | Other | other | |
| chr13 | LINC00348 | 4 | 74 | 8 | LINC00348 | long intergenic non-protein coding RNA 348 | Other | other | |
| chr1 | RP11-421L21.3 | 3 | 86 | 8 | | | | | |
| chr16 | ABCC6P1 | 4 | 88 | 8 | ABCC6P1 | ATP binding cassette subfamily C member 6 pseudogene 1 | Other | other | |
| chr1 | RP11-656D10.3 | 4 | 74 | 8 | | | | | |
| chr7 | ALG1L5P | 4 | 200 | 8 | ALG1L5P | ALG1 like 5, pseudogene | Other | other | |
| chr4 | HSD17B11 | 3 | 131 | 8 | HSD17B11 | hydroxysteroid 17-beta dehydrogenase 11 | Cytoplasm | enzyme | |
| chr3 | RP11-71E19.1 | 4 | 163 | 8 | | | | | |
| chr11 | NLRP14 | 4 | 138 | 8 | NLRP14 | NLR family pyrin domain containing 14 | Extracellular Space | other | |
| chr8 | FBXO32 | 4 | 103 | 8 | FBXO32 | F-box protein 32 | Cytoplasm | enzyme | |
| chr19 | ZNF432 | 4 | 77 | 8 | ZNF432 | zinc finger protein 432 | Nucleus | transcription regulator | |

| | | | | | | | | | |
|---|---|---|---|---|---|---|---|---|---|
| chr13 | *FLT3* | 4 | 86 | 8 | FLT3 | fms related tyrosine kinase 3 | Plasma Membrane | kinase | nintedanib, KW 2449, imatinib/sorafenib, IMC-EB10, gilteritinib, midostaurin, sorafenib, PIM/FLT3 kinase inhibitor SEL24, SB-1317, MRX-2843, sorafenib/sulindac/sunitinib, dovitinib, 4G8-SDIEM, imatinib/sunitinib, bevacizumab/sorafenib, dexamethasone/lenalidomide/sorafenib, sunitinib, 5-azacytidine/midostaurin, brigatinib, 5-azacytidine/sorafenib, tandutinib, XL999, quizartinib, ponatinib, [illegible] |
| chr16 | *CIAPIN1* | 4 | 85 | 8 | CIAPIN1 | cytokine induced apoptosis | Cytoplasm | other | |
| chr7 | *CROT* | 4 | 88 | 8 | CROT | carnitine O-octanoyltransferase | Cytoplasm | enzyme | |
| chr5 | *PANK3* | 3 | 105 | 8 | PANK3 | pantothenate kinase 3 | Cytoplasm | kinase | |
| chr11 | *RIC8A* | 3 | 166 | 8 | RIC8A | RIC8 guanine nucleotide exchange factor A | Cytoplasm | other | |
| chr19 | *SCAF1* | 4 | 175 | 8 | SCAF1 | SR-related CTD associated factor | Extracellular Space | other | |
| chr11 | *CPSF7* | 3 | 118 | 8 | CPSF7 | cleavage and polyadenylation specific factor 7 | Nucleus | other | |
| chr12 | *RP11-796E2.4* | 4 | 82 | 8 | | | | | |
| chr1 | *RP11-206L10.2* | 4 | 147 | 8 | | | | | |
| chr12 | *KRT3* | 4 | 281 | 8 | KRT3 | keratin 3 | Cytoplasm | other | |
| chr12 | *ACADS* | 4 | 137 | 8 | ACADS | acyl-CoA dehydrogenase short | Cytoplasm | enzyme | |
| chr5 | *POLR3G* | 3 | 66 | 8 | POLR3G | RNA polymerase III subunit G | Nucleus | enzyme | |
| chr11 | *ANKK1* | 4 | 141 | 8 | ANKK1 | ankyrin repeat and kinase domain containing 1 | Cytoplasm | kinase | |
| chr16 | *C16orf78* | 4 | 107 | 8 | C16orf78 | chromosome 16 open reading frame 78 | Nucleus | other | |
| chr19 | *PLEKHF1* | 4 | 138 | 8 | PLEKHF1 | pleckstrin homology and FYVE domain containing 1 | Cytoplasm | other | |
| chr10 | *PSTK* | 4 | 86 | 8 | PSTK | phosphoseryl-tRNA kinase | Cytoplasm | kinase | |
| chr8 | *ANKRD46* | 3 | 71 | 8 | ANKRD46 | ankyrin repeat domain 46 | Nucleus | transcription regulator | |
| chr15 | *RP11-566K19.6* | 4 | 112 | 8 | | | | | |
| chr2 | *AC018685.1* | 3 | 130 | 8 | | | | | |
| chr12 | *PPP1R1A* | 4 | 186 | 8 | PPP1R1A | protein phosphatase 1 regulatory inhibitor subunit 1A | Cytoplasm | phosphatase | |
| chr3 | *RP11-513G11.4* | 3 | 111 | 8 | | | | | |
| chr19 | *PAFAH1B3* | 4 | 161 | 8 | PAFAH1B3 | platelet activating factor acetylhydrolase 1b catalytic | Cytoplasm | enzyme | |
| chr20 | *RP4-550H1.4* | 4 | 109 | 8 | | | | | |
| chr6 | *SOBP* | 4 | 104 | 8 | SOBP | sine oculis binding protein | Nucleus | other | |
| chrX | *OPN1MW2* | 3 | 124 | 8 | 1MW (includes ot | opsin 1, medium wave sensitive | Plasma Membrane | -protein coupled recepto | |
| chr14 | *DDHD1* | 3 | 95 | 8 | DDHD1 | DDHD domain containing 1 | Extracellular Space | enzyme | |
| chr17 | *RP11-160E2.16* | 3 | 111 | 8 | | | | | |
| chr12 | *FAM109A* | 4 | 162 | 8 | PHETA1 | PH domain containing endocytic trafficking adaptor 1 | Cytoplasm | other | |
| chr1 | *MGST3* | 4 | 118 | 8 | MGST3 | microsomal glutathione S-transferase 3 | Cytoplasm | enzyme | |
| chr2 | *PRKAG3* | 3 | 125 | 8 | PRKAG3 | protein kinase AMP-activated non-catalytic subunit gamma 3 | Cytoplasm | other | |
| chr10 | *IDI2-AS1* | 3 | 77 | 8 | IDI2-AS1 | IDI2 antisense RNA 1 | Other | other | |
| chr7 | *ZC3HAV1L* | 4 | 130 | 8 | ZC3HAV1L | zinc finger CCCH-type containing, antiviral 1 like | Cytoplasm | other | |
| chr13 | *DHRS12* | 3 | 98 | 8 | DHRS12 | dehydrogenase/reductase 12 | Nucleus | other | |
| chr17 | *RP1-62O9.3* | 4 | 115 | 8 | | | | | |
| chr2 | *AC144449.1* | 3 | 69 | 8 | | | | | |
| chr1 | *PMVK* | 4 | 216 | 8 | PMVK | phosphomevalonate kinase | Cytoplasm | kinase | |
| chr17 | *KRT36* | 4 | 176 | 8 | KRT36 | keratin 36 | Cytoplasm | other | |
| chrX | *GAGE12I* | 4 | 344 | 8 | GE1 (includes oth | G antigen 1 | Other | other | |
| chr11 | *SMTNL1* | 4 | 157 | 8 | SMTNL1 | smoothelin like 1 | Cytoplasm | other | |
| chr1 | *RP11-48O20.4* | 4 | 148 | 8 | | | | | |
| chr1 | *PPP2R5A* | 4 | 98 | 8 | PPP2R5A | protein phosphatase 2 regulatory subunit B'alpha | Cytoplasm | phosphatase | |
| chr11 | *PDDC1* | 4 | 216 | 8 | GATD1 | glutamine amidotransferase like class 1 domain containing 1 | Cytoplasm | other | |
| chr11 | *RP11-712B9.2* | 4 | 226 | 8 | | | | | |
| chr17 | *PYCR1* | 4 | 246 | 8 | PYCR1 | pyrroline-5-carboxylate reductase | Cytoplasm | enzyme | |
| chr1 | *RP4-559A3.7* | 4 | 186 | 8 | | | | | |
| chr11 | *FOXR1* | 4 | 110 | 8 | FOXR1 | forkhead box R1 | Nucleus | transcription regulator | |
| chr10 | *CTA-109P11.1* | 3 | 91 | 8 | | | | | |
| chr21 | *SPATC1L* | 3 | 130 | 8 | SPATC1L | spermatogenesis and centriole associated 1 like | Other | other | |
| chr12 | *TARBP2* | 4 | 223 | 8 | TARBP2 | TARBP2 subunit of RISC loading complex | Nucleus | other | |
| chr7 | *NPTX2* | 4 | 190 | 8 | NPTX2 | neuronal pentraxin 2 | Extracellular Space | other | |
| chr2 | *TMEM178A* | 4 | 139 | 8 | TMEM178A | transmembrane protein 178A | Cytoplasm | other | |

| | | | | | | | | | |
|---|---|---|---|---|---|---|---|---|---|
| chr14 | *FRMD6* | 4 | 94 | 8 | FRMD6 | FERM domain containing 6 | Cytoplasm | other | |
| chr15 | *GOLGA8K* | 3 | 163 | 8 | GA8K (includes ot | golgin A8 family member K | Other | other | |
| chr4 | *DKK2* | 3 | 93 | 8 | DKK2 | dickkopf WNT signaling pathway inhibitor 2 | Extracellular Space | other | |
| chr3 | *SEC61A1* | 4 | 122 | 8 | SEC61A1 | SEC61 translocon alpha 1 | Cytoplasm | transporter | |
| chr5 | *RPL37* | 4 | 278 | 8 | RPL37 | ribosomal protein L37 | Cytoplasm | other | |
| chr9 | *RP11-204M4.2* | 3 | 105 | 8 | | | | | |
| chr16 | *CMTM3* | 4 | 207 | 8 | CMTM3 | CKLF like MARVEL transmembrane domain | Extracellular Space | cytokine | |
| chr12 | *GLS2* | 4 | 89 | 8 | GLS2 | glutaminase 2 | Cytoplasm | enzyme | |
| chr15 | *RP11-143J24.1* | 4 | 91 | 8 | | | | | |
| chr2 | *XIRP2* | 3 | 114 | 8 | XIRP2 | xin actin binding repeat | Other | other | |
| chr16 | *CTD-2288F12.1* | 4 | 94 | 8 | | | | | |
| chr2 | *AC016683.6* | 4 | 149 | 8 | | | | | |
| chr8 | *ASPH* | 4 | 72 | 8 | ASPH | aspartate beta-hydroxylase | Cytoplasm | enzyme | |
| chr3 | *RP11-538P18.2* | 3 | 76 | 8 | | | | | |
| chr16 | *RP11-958N24.1* | 4 | 207 | 8 | | | | | |
| chr20 | *FERMT1* | 3 | 82 | 8 | FERMT1 | fermitin family member 1 | Plasma Membrane | other | |
| chr4 | *RP11-217C7.1* | 3 | 85 | 8 | | | | | |
| chr1 | *PPIH* | 3 | 99 | 8 | PPIH | peptidylprolyl isomerase H | Nucleus | enzyme | |
| chr9 | *RP11-375O18.2* | 4 | 80 | 8 | | | | | |
| chr17 | *RPL21P122* | 3 | 77 | 8 | RPL21P122 | ribosomal protein L21 pseudogene 122 | Other | other | |
| chr2 | *PPP1CB* | 3 | 63 | 8 | PPP1CB | protein phosphatase 1 catalytic subunit beta | Cytoplasm | phosphatase | |
| chr22 | *HSCB* | 3 | 65 | 8 | HSCB | HscB mitochondrial iron-sulfur cluster cochaperone | Cytoplasm | other | |
| chr5 | *RP11-310P5.1* | 5 | 603 | 8 | | | | | |
| chr1 | *YBX1* | 3 | 95 | 8 | YBX1 | Y-box binding protein 1 | Nucleus | transcription regulator | |
| chr1 | *RP11-443B7.1* | 4 | 179 | 8 | | | | | |
| chr2 | *FAM228B* | 3 | 96 | 8 | FAM228B | family with sequence similarity 228 member B | Other | other | |
| chr8 | *MCMDC2* | 3 | 94 | 8 | MCMDC2 | minichromosome maintenance domain containing 2 | Other | other | |
| chr3 | *TRIM59* | 4 | 129 | 8 | TRIM59 | tripartite motif containing 59 | Cytoplasm | enzyme | |
| chr19 | *CTB-52I2.4* | 4 | 120 | 8 | | | | | |
| chr17 | *TMEM92* | 3 | 164 | 8 | TMEM92 | transmembrane protein 92 | Nucleus | other | |
| chr8 | *TEX15* | 4 | 68 | 8 | TEX15 | testis expressed 15, meiosis and synapsis associated | Extracellular Space | other | |
| chr9 | *RP11-406A20.4* | 4 | 87 | 8 | | | | | |
| chrX | *KLF8* | 3 | 121 | 8 | KLF8 | Kruppel like factor 8 | Nucleus | transcription regulator | |
| chr7 | *RP13-580B18.1* | 4 | 165 | 8 | | | | | |
| chr9 | *RP11-402N8.1* | 3 | 79 | 8 | | | | | |
| chr6 | *TMEM242* | 4 | 1243 | 8 | TMEM242 | transmembrane protein 242 | Other | other | |
| chr10 | *ZNF488* | 4 | 271 | 8 | ZNF488 | zinc finger protein 488 | Nucleus | transcription regulator | |
| chr19 | *SCN1B* | 4 | 157 | 8 | SCN1B | sodium voltage-gated channel beta subunit 1 | Plasma Membrane | ion channel | procaine, riluzole, articaine/epinephrine, prilocaine, lidocaine/prilocaine, penicillin G/procaine, articaine, epinephrine/lidocaine, lidocaine, fosphenytoin, zonisamide. |
| chr16 | *CCP110* | 4 | 65 | 8 | CCP110 | centriolar coiled-coil protein 110 | Cytoplasm | other | |
| chr19 | *FARSA* | 4 | 171 | 8 | FARSA | phenylalanyl-tRNA synthetase subunit alpha | Cytoplasm | enzyme | |
| chr11 | *RP11-428C19.4* | 4 | 123 | 8 | | | | | |
| chr1 | *C1orf158* | 4 | 104 | 8 | C1orf158 | chromosome 1 open reading frame 158 | Other | other | |
| chr1 | *ZNF436* | 4 | 108 | 8 | ZNF436 | zinc finger protein 436 | Nucleus | transcription regulator | |
| chr4 | *RP11-281P23.2* | 3 | 74 | 8 | | | | | |
| chr1 | *RP11-480I12.5* | 4 | 144 | 8 | | | | | |
| chr5 | *CTB-7E3.1* | 4 | 102 | 8 | | | | | |
| chr11 | *ZP1* | 3 | 216 | 8 | ZP1 | zona pellucida glycoprotein 1 | Extracellular Space | other | |
| chr6 | *RP1-137D17.1* | 3 | 125 | 8 | | | | | |
| chr19 | *HMHA1* | 4 | 195 | 8 | ARHGAP45 | Rho GTPase activating protein 45 | Cytoplasm | transporter | |
| chr13 | *KLF5* | 4 | 102 | 8 | KLF5 | Kruppel like factor 5 | Nucleus | transcription regulator | |
| chr10 | *PTER* | 3 | 69 | 8 | PTER | phosphotriesterase related | Cytoplasm | enzyme | |
| chr2 | *RND3* | 3 | 78 | 8 | RND3 | Rho family GTPase 3 | Cytoplasm | enzyme | |
| chr22 | *LINC00229* | 4 | 159 | 8 | LINC00229 | long intergenic non-protein coding RNA 229 | Other | other | |
| chr16 | *RP11-467J12.4* | 4 | 85 | 8 | | | | | |
| chr12 | *PUS7L* | 3 | 104 | 8 | PUS7L | pseudouridine synthase 7 like | Other | other | |
| chr16 | *CNEP1R1* | 3 | 79 | 8 | CNEP1R1 | CTD nuclear envelope phosphatase 1 regulatory subunit | Nucleus | other | |
| chr8 | *NSMAF* | 3 | 65 | 8 | NSMAF | neutral sphingomyelinase activation associated factor | Cytoplasm | other | |
| chr14 | *RP11-146E13.4* | 4 | 136 | 8 | | | | | |
| chr12 | *RP1-90J4.1* | 3 | 131 | 8 | | | | | |
| chr14 | *RP11-1070N10.5* | 3 | 79 | 8 | | | | | |
| chr4 | *GPM6A* | 4 | 80 | 8 | GPM6A | glycoprotein M6A | Plasma Membrane | ion channel | |
| chr19 | *CD177P1* | 4 | 155 | 8 | CD177P1 | CD177 molecule pseudogene 1 | Other | other | |
| chr19 | *RINL* | 4 | 144 | 8 | RINL | Ras and Rab interactor like | Other | other | |
| chr11 | *B3GAT3* | 4 | 191 | 8 | B3GAT3 | beta-1,3-glucuronyltransferase 3 | Cytoplasm | enzyme | |
| chr1 | *IFI16* | 3 | 69 | 8 | IFI16 | interferon gamma inducible | Nucleus | transcription regulator | |
| chr3 | *KCNMB3* | 3 | 74 | 8 | KCNMB3 | potassium calcium-activated channel subfamily M regulatory | Plasma Membrane | ion channel | tedisamil |

| | | | | | | | | | |
|---|---|---|---|---|---|---|---|---|---|
| chr20 | *TBC1D20* | 4 | 77 | 8 | TBC1D20 | TBC1 domain family member 20 | Nucleus | other | |
| chr8 | *RP11-1084E5.1* | 4 | 84 | 8 | | | | | |
| chr5 | *CTC-367J11.1* | 3 | 100 | 8 | | | | | |
| chr17 | *OTOP2* | 4 | 162 | 8 | OTOP2 | otopetrin 2 | Other | other | |
| chr1 | *PVRL4* | 4 | 235 | 8 | NECTIN4 | nectin cell adhesion molecule 4 | Plasma Membrane | other | anti-nectin-4 monoclonal antibody-drug conjugate AGS-22M6E, enfortumab vedotin |
| chr20 | *OSER1-AS1* | 5 | 128 | 8 | OSER1-DT | OSER1 divergent transcript | Other | other | |
| chr17 | *NOS2P3* | 4 | 175 | 8 | NOS2P3 | nitric oxide synthase 2 | Other | other | |
| chr5 | *RP11-241G9.3* | 4 | 90 | 8 | | | | | |
| chr7 | *TBRG4* | 3 | 166 | 8 | TBRG4 | transforming growth factor beta regulator 4 | Cytoplasm | other | |
| chr12 | *RP11-776A13.1* | 4 | 81 | 8 | | | | | |
| chr3 | *RP11-446H18.5* | 4 | 129 | 8 | | | | | |
| chr2 | *SGPP2* | 4 | 90 | 8 | SGPP2 | sphingosine-1-phosphate phosphatase 2 | Cytoplasm | phosphatase | |
| chr22 | *THOC5* | 3 | 89 | 8 | THOC5 | THO complex 5 | Cytoplasm | other | |
| chr19 | *WDR87* | 4 | 96 | 8 | WDR87 | WD repeat domain 87 | Other | other | |
| chr19 | *MAP1S* | 4 | 162 | 8 | MAP1S | microtubule associated protein | Cytoplasm | enzyme | |
| chr17 | *ATXN7L3* | 3 | 149 | 8 | ATXN7L3 | ataxin 7 like 3 | Nucleus | transcription regulator | |
| chr2 | *ANKRD20A8P* | 4 | 97 | 8 | ANKRD20A8P | ankyrin repeat domain 20 family member A8, pseudogene | Other | other | |
| chr5 | *MGAT1* | 4 | 199 | 8 | MGAT1 | mannosyl (alpha-1,3-)-glycoprotein beta-1,2-N-acetylglucosaminyltransferase | Cytoplasm | enzyme | |
| chr19 | *CTD-2162K18.5* | 4 | 66 | 8 | | | | | |
| chr17 | *ORMDL3* | 4 | 160 | 8 | ORMDL3 | ORMDL sphingolipid biosynthesis regulator 3 | Cytoplasm | other | |
| chr2 | *AC012494.1* | 4 | 80 | 8 | | | | | |
| chr16 | *Z97634.3* | 3 | 131 | 8 | | | | | |
| chr8 | *TMEM68* | 3 | 71 | 8 | TMEM68 | transmembrane protein 68 | Other | other | |
| chr8 | *CHRNB3* | 3 | 76 | 8 | CHRNB3 | cholinergic receptor nicotinic beta 3 subunit | Plasma Membrane | transmembrane receptor | succinylcholine, rapacuronium, doxacurium, vecuronium, pipecuronium, metocurine, mivacurium, D-tubocurarine, nicotine, arecoline, pancuronium, atracurium, isoflurane, cisatracurium, rocuronium, mecamylamine |
| chr12 | *CAPS2* | 3 | 68 | 8 | CAPS2 | calcyphosine 2 | Other | other | |
| chr1 | *RP11-442N24__B.* | 3 | 102 | 8 | | | | | |
| chr12 | *RP11-314D7.2* | 4 | 89 | 8 | | | | | |
| chr6 | *SIM1* | 3 | 121 | 8 | SIM1 | SIM bHLH transcription factor 1 | Nucleus | transcription regulator | |
| chr2 | *RP11-521D12.5* | 3 | 133 | 8 | | | | | |
| chr1 | *FAM72B* | 4 | 176 | 8 | FAM72B | family with sequence similarity 72 member B | Cytoplasm | other | |
| chr2 | *B3GNT2* | 4 | 80 | 8 | B3GNT2 | UDP-GlcNAc:betaGal beta-1,3-N-acetylglucosaminyltransferase 2 | Cytoplasm | enzyme | |
| chr7 | *AC007652.1* | 4 | 120 | 8 | | | | | |
| chr7 | *ZNF800* | 4 | 75 | 8 | ZNF800 | zinc finger protein 800 | Other | other | |
| chr8 | *SPAG11B* | 4 | 195 | 8 | SPAG11B | sperm associated antigen 11B | Extracellular Space | other | |
| chr16 | *ZNF646* | 4 | 176 | 8 | ZNF646 | zinc finger protein 646 | Nucleus | transcription regulator | |
| chr5 | *TRIM23* | 4 | 96 | 8 | TRIM23 | tripartite motif containing 23 | Nucleus | enzyme | |
| chr9 | *TTC16* | 4 | 232 | 8 | TTC16 | tetratricopeptide repeat domain | Other | other | |
| chr9 | *MAMDC4* | 5 | 220 | 8 | MAMDC4 | MAM domain containing 4 | Cytoplasm | other | |
| chr1 | *CYP4A11* | 4 | 113 | 8 | CYP4A11 | cytochrome P450 family 4 subfamily A member 11 | Cytoplasm | enzyme | |
| chr8 | *NPM2* | 4 | 155 | 8 | NPM2 | nucleophosmin/nucleoplasmin 2 | Nucleus | other | |
| chr11 | *RP11-613D13.5* | 4 | 72 | 8 | | | | | |
| chr13 | *LINC01044* | 3 | 153 | 8 | LINC01044 | long intergenic non-protein coding RNA 1044 | Other | other | |
| chr7 | *LRRC61* | 4 | 158 | 8 | LRRC61 | leucine rich repeat containing 61 | Other | other | |
| chr7 | *GIGYF1* | 4 | 242 | 8 | GIGYF1 | GRB10 interacting GYF protein 1 | Extracellular Space | other | |
| chr7 | *ASL* | 4 | 127 | 8 | ASL | argininosuccinate lyase | Cytoplasm | enzyme | |
| chr1 | *UTS2* | 4 | 95 | 8 | UTS2 | urotensin 2 | Extracellular Space | other | |
| chr12 | *LEPREL2* | 3 | 154 | 8 | P3H3 | prolyl 3-hydroxylase 3 | Nucleus | enzyme | |
| chr19 | *KLKP1* | 3 | 113 | 8 | KLKP1 | kallikrein pseudogene 1 | Other | other | |
| chr10 | *OBFC1* | 3 | 128 | 8 | STN1 | STN1 subunit of CST complex | Nucleus | other | |
| chr11 | *ANGPTL5* | 4 | 145 | 8 | ANGPTL5 | angiopoietin like 5 | Extracellular Space | other | |
| chr10 | *OPTN* | 3 | 79 | 8 | OPTN | optineurin | Cytoplasm | other | |
| chr6 | *PKIB* | 3 | 83 | 8 | PKIB | cAMP-dependent protein kinase inhibitor beta | Other | other | |
| chr2 | *BOLA3* | 3 | 133 | 8 | BOLA3 | bolA family member 3 | Cytoplasm | other | |
| chr10 | *PLAC9* | 4 | 137 | 8 | PLAC9 | placenta associated 9 | Other | other | |
| chr14 | *NEK9* | 3 | 72 | 8 | NEK9 | NIMA related kinase 9 | Nucleus | kinase | |
| chr9 | *RP11-15J10.8* | 4 | 147 | 8 | | | | | |
| chr12 | *PXN-AS1* | 4 | 121 | 8 | PXN-AS1 | PXN antisense RNA 1 | Other | other | |
| chr8 | *RP11-388G22.1* | 3 | 71 | 8 | | | | | |
| chr2 | *NMS* | 4 | 103 | 8 | NMS | neuromedin S | Extracellular Space | other | |
| chr17 | *TEX19* | 4 | 119 | 8 | TEX19 | testis expressed 19 | Nucleus | other | |

| | | | | | | | | | |
|---|---|---|---|---|---|---|---|---|---|
| chr16 | *RP11-17M15.2* | 4 | 126 | 8 | | | | | |
| chr3 | *PP2D1* | 4 | 102 | 8 | PP2D1 | protein phosphatase 2C like domain containing 1 | Other | phosphatase | |
| chr6 | *NRN1* | 4 | 129 | 8 | NRN1 | neuritin 1 | Cytoplasm | other | |
| chr21 | *LINC00313* | 4 | 180 | 8 | LINC00313 | long intergenic non-protein coding RNA 313 | Other | other | |
| chr4 | *TBC1D19* | 3 | 78 | 8 | TBC1D19 | TBC1 domain family member 19 | Other | other | |
| chr16 | *RP11-459F6.1* | 4 | 86 | 8 | | | | | |
| chr14 | *ABHD4* | 3 | 95 | 8 | ABHD4 | abhydrolase domain containing 4 | Cytoplasm | peptidase | |
| chr11 | *MS4A18* | 4 | 103 | 8 | MS4A18 | membrane spanning 4-domains | Other | other | |
| chr19 | *ZNF230* | 3 | 92 | 8 | ZNF230 | zinc finger protein 230 | Nucleus | transcription regulator | |
| chr2 | *SLC20A1* | 3 | 88 | 8 | SLC20A1 | solute carrier family 20 member 1 | Plasma Membrane | transporter | |
| chr2 | *NDUFS1* | 3 | 65 | 8 | NDUFS1 | NADH:ubiquinone oxidoreductase core subunit S1 | Cytoplasm | enzyme | |
| chr14 | *RP5-1021I20.1* | 3 | 77 | 8 | | | | | |
| chr11 | *ACY3* | 3 | 215 | 8 | ACY3 | aminoacylase 3 | Cytoplasm | enzyme | |
| chr7 | *ORC5* | 3 | 74 | 8 | ORC5 | origin recognition complex | Nucleus | other | |
| chr15 | *LOXL1-AS1* | 4 | 174 | 8 | LOXL1-AS1 | LOXL1 antisense RNA 1 | Other | other | |
| chr14 | *NOXRED1* | 3 | 61 | 8 | NOXRED1 | NADP dependent oxidoreductase domain containing 1 | Other | other | |
| chr3 | *HHLA2* | 4 | 100 | 8 | HHLA2 | HERV-H LTR-associating 2 | Other | other | |
| chr9 | *PTGES2* | 4 | 247 | 8 | PTGES2 | prostaglandin E synthase 2 | Cytoplasm | transcription regulator | |
| chr7 | *TMEM120A* | 4 | 334 | 8 | TMEM120A | transmembrane protein 120A | Other | other | |
| chr11 | *SSRP1* | 3 | 131 | 8 | SSRP1 | structure specific recognition | Nucleus | transcription regulator | |
| chr21 | *AF127577.8* | 3 | 106 | 8 | | | | | |
| chr3 | *ARPP21* | 4 | 73 | 8 | ARPP21 | cAMP regulated phosphoprotein | Cytoplasm | other | |
| chr7 | *AC073236.3* | 4 | 106 | 8 | | | | | |
| chr17 | *HLF* | 4 | 83 | 8 | HLF | HLF transcription factor, PAR bZIP family member | Nucleus | transcription regulator | |
| chr16 | *RP11-279O17.3* | 4 | 67 | 8 | | | | | |
| chr18 | *METTL4* | 3 | 95 | 8 | METTL4 | methyltransferase like 4 | Other | other | |
| chr17 | *CTD-2319I12.3* | 3 | 198 | 8 | | | | | |
| chr1 | *DPH5* | 4 | 95 | 8 | DPH5 | diphthamide biosynthesis 5 | Cytoplasm | enzyme | |
| chr5 | *THBS4* | 4 | 75 | 8 | THBS4 | thrombospondin 4 | Extracellular Space | other | |
| chr15 | *RP11-483E23.2* | 4 | 151 | 8 | | | | | |
| chr3 | *CTD-2021J15.1* | 3 | 65 | 8 | | | | | |
| chr2 | *PRKD3* | 3 | 63 | 8 | PRKD3 | protein kinase D3 | Nucleus | kinase | |
| chr22 | *APOL1* | 3 | 87 | 8 | APOL1 | apolipoprotein L1 | Extracellular Space | transporter | |
| chr22 | *LINC00898* | 3 | 126 | 8 | LINC00898 | long intergenic non-protein coding RNA 898 | Other | other | |
| chr18 | *KCNG2* | 4 | 183 | 8 | KCNG2 | potassium voltage-gated channel modifier subfamily G member 2 | Plasma Membrane | ion channel | amiodarone, nicorandil |
| chr11 | *TCIRG1* | 4 | 103 | 8 | TCIRG1 | T cell immune regulator 1, ATPase H+ transporting V0 | Plasma Membrane | enzyme | |
| chr16 | *RP11-58A18.1* | 4 | 100 | 8 | | | | | |
| chr11 | *PPP1R32* | 3 | 217 | 8 | PPP1R32 | protein phosphatase 1 regulatory subunit 32 | Other | other | |
| chr10 | *PROSER2* | 3 | 70 | 8 | PROSER2 | proline and serine rich 2 | Other | other | |
| chr11 | *COMMD9* | 4 | 157 | 8 | COMMD9 | COMM domain containing 9 | Cytoplasm | other | |
| chr20 | *SNPH* | 4 | 168 | 8 | SNPH | syntaphilin | Plasma Membrane | other | |
| chr5 | *RP11-232L2.2* | 4 | 96 | 8 | | | | | |
| chr6 | *CDKN1A* | 4 | 146 | 8 | CDKN1A | cyclin dependent kinase inhibitor | Nucleus | kinase | |
| chr10 | *NOLC1* | 4 | 110 | 8 | NOLC1 | nucleolar and coiled-body phosphoprotein 1 | Nucleus | transcription regulator | |
| chr16 | *RP4-536B24.4* | 3 | 158 | 8 | | | | | |
| chrX | *ARHGAP4* | 4 | 206 | 8 | ARHGAP4 | Rho GTPase activating protein 4 | Cytoplasm | other | |
| chrX | *GAGE2B* | 3 | 265 | 8 | GE2D (includes otl | G antigen 2D | Other | other | |
| chr20 | *RP4-610C12.4* | 4 | 431 | 8 | | | | | |
| chr1 | *ASCL5* | 4 | 171 | 8 | ASCL5 | achaete-scute family bHLH transcription factor 5 | Other | other | |
| chr5 | *RP11-373N22.3* | 4 | 69 | 8 | | | | | |
| chr10 | *FAM208B* | 3 | 80 | 8 | TASOR2 | transcription activation suppressor family member 2 | Nucleus | other | |
| chr1 | *SH2D2A* | 4 | 170 | 8 | SH2D2A | SH2 domain containing 2A | Cytoplasm | other | |
| chr17 | *RP11-353N14.2* | 3 | 168 | 8 | | | | | |
| chr15 | *LYSMD4* | 3 | 170 | 8 | LYSMD4 | LysM domain containing 4 | Other | other | |
| chr15 | *RP11-932O9.4* | 3 | 172 | 8 | | | | | |
| chr1 | *ETV3L* | 4 | 187 | 8 | ETV3L | ETS variant transcription factor 3 | Other | other | |
| chr13 | *LINC00402* | 3 | 89 | 8 | LINC00402 | long intergenic non-protein coding RNA 402 | Other | other | |
| chr17 | *CRYBA1* | 3 | 103 | 8 | CRYBA1 | crystallin beta A1 | Other | other | |
| chr1 | *RP11-306I1.2* | 4 | 123 | 8 | | | | | |
| chr1 | *ADIPOR1* | 4 | 96 | 8 | ADIPOR1 | adiponectin receptor 1 | Plasma Membrane | transmembrane receptoı | |
| chr5 | *MGAT4B* | 4 | 309 | 8 | MGAT4B | alpha-1,3-mannosyl-glycoprotein 4-beta-N-acetylglucosaminyltransferase B | Cytoplasm | enzyme | |
| chr5 | *FBXO4* | 3 | 188 | 8 | FBXO4 | F-box protein 4 | Nucleus | enzyme | |
| chr6 | *EPB41L2* | 3 | 67 | 8 | EPB41L2 | erythrocyte membrane protein band 4.1 like 2 | Plasma Membrane | other | |
| chr19 | *TNFSF14* | 4 | 155 | 8 | TNFSF14 | TNF superfamily member 14 | Extracellular Space | cytokine | |
| chr6 | *ZNF184* | 3 | 84 | 8 | ZNF184 | zinc finger protein 184 | Nucleus | transcription regulator | |
| chr17 | *RP11-1018N14.5* | 4 | 91 | 8 | | | | | |
| chr16 | *RP11-467M13.2* | 4 | 121 | 8 | | | | | |
| chr2 | *AC008268.2* | 4 | 221 | 8 | | | | | |
| chr2 | *AC009950.2* | 3 | 93 | 8 | | | | | |
| chr3 | *LSMEM2* | 4 | 182 | 8 | LSMEM2 | leucine rich single-pass membrane protein 2 | Other | other | |
| chr10 | *ARHGAP12* | 3 | 62 | 8 | ARHGAP12 | Rho GTPase activating protein 12 | Cytoplasm | other | |

| | | | | | | | | | |
|---|---|---|---|---|---|---|---|---|---|
| chr1 | POGK | 4 | 110 | 8 | POGK | pogo transposable element derived with KRAB domain | Nucleus | other | |
| chr6 | AKIRIN2 | 3 | 68 | 8 | AKIRIN2 | akirin 2 | Nucleus | other | |
| chr17 | LINC00674 | 3 | 84 | 8 | LINC00674 | long intergenic non-protein coding RNA 674 | Other | other | |
| chrX | HAUS7 | 4 | 190 | 8 | HAUS7 | HAUS augmin like complex | Cytoplasm | other | |
| chr7 | NACAD | 4 | 302 | 8 | NACAD | NAC alpha domain containing | Other | other | |
| chr7 | AHR | 3 | 88 | 8 | AHR | aryl hydrocarbon receptor | Nucleus | d-dependent nuclear rec | BAY2416964 |
| chr15 | KIF23 | 3 | 89 | 8 | KIF23 | kinesin family member 23 | Cytoplasm | other | |
| chr15 | LINC00593 | 3 | 142 | 8 | LINC00593 | long intergenic non-protein coding RNA 593 | Other | other | |
| chr3 | SEC22C | 3 | 73 | 8 | SEC22C | SEC22 homolog C, vesicle trafficking protein | Cytoplasm | other | |
| chr1 | RP11-191G24.1 | 4 | 72 | 8 | | | | | |
| chr14 | FAM177A1 | 3 | 66 | 8 | FAM177A1 | family with sequence similarity 177 member A1 | Other | other | |
| chr21 | USP16 | 3 | 68 | 8 | USP16 | ubiquitin specific peptidase 16 | Cytoplasm | peptidase | |
| chr1 | FAM72D | 3 | 202 | 8 | FAM72C/FAM72D | family with sequence similarity 72 member D | Nucleus | other | |
| chr19 | NCLN | 4 | 183 | 8 | NCLN | nicalin | Cytoplasm | peptidase | |
| chr7 | AC027269.2 | 4 | 91 | 8 | | | | | |
| chr17 | AC002539.1 | 3 | 83 | 8 | | | | | |
| chr19 | TMEM238 | 4 | 164 | 8 | TMEM238 | transmembrane protein 238 | Other | other | |
| chr10 | UCMA | 3 | 96 | 8 | UCMA | upper zone of growth plate and cartilage matrix associated | Extracellular Space | other | |
| chr8 | RP11-89M20.2 | 4 | 93 | 8 | | | | | |
| chr19 | ZNF845 | 4 | 101 | 8 | ZNF845 | zinc finger protein 845 | Other | other | |
| chr1 | TMEM183A | 4 | 104 | 8 | TMEM183A | transmembrane protein 183A | Other | other | |
| chr19 | STX10 | 4 | 120 | 8 | STX10 | syntaxin 10 | Cytoplasm | other | |
| chr6 | THBS2 | 4 | 135 | 8 | THBS2 | thrombospondin 2 | Extracellular Space | other | |
| chr17 | PTRH2 | 3 | 95 | 8 | PTRH2 | peptidyl-tRNA hydrolase 2 | Cytoplasm | enzyme | |
| chr16 | PALB2 | 3 | 89 | 8 | PALB2 | partner and localizer of BRCA2 | Nucleus | other | |
| chr8 | DUSP26 | 4 | 96 | 8 | DUSP26 | dual specificity phosphatase 26 | Cytoplasm | enzyme | |
| chr3 | RP11-767C1.1 | 4 | 144 | 8 | | | | | |
| chr19 | KXD1 | 3 | 160 | 8 | KXD1 | KxDL motif containing 1 | Cytoplasm | other | |
| chr20 | BPIFB9P | 3 | 161 | 8 | BPIFB9P | BPI fold containing family B member 9, pseudogene | Other | other | |
| chr1 | MEX3A | 4 | 177 | 8 | MEX3A | mex-3 RNA binding family | Cytoplasm | other | |
| chr6 | RP4-655C5.4 | 4 | 101 | 8 | | | | | |
| chr19 | SLC6A16 | 2 | 74 | 8 | SLC6A16 | solute carrier family 6 member 16 | Plasma Membrane | transporter | |
| chr17 | CD300C | 3 | 157 | 8 | CD300C | CD300c molecule | Plasma Membrane | transmembrane recepto | |
| chr22 | MED15P7 | 3 | 139 | 8 | MED15P7 | mediator complex subunit 15 pseudogene 7 | Other | other | |
| chr18 | MBD1 | 3 | 117 | 8 | MBD1 | methyl-CpG binding domain | Nucleus | transcription regulator | |
| chr15 | TM2D3 | 4 | 158 | 8 | TM2D3 | TM2 domain containing 3 | Other | other | |
| chr11 | SLC39A13 | 4 | 424 | 8 | SLC39A13 | solute carrier family 39 member | Cytoplasm | transporter | |
| chr8 | RP11-302J23.1 | 4 | 71 | 8 | | | | | |
| chr19 | CYP4F8 | 4 | 141 | 8 | CYP4F8 | cytochrome P450 family 4 subfamily F member 8 | Cytoplasm | enzyme | |
| chr1 | RP4-739H11.3 | 4 | 96 | 8 | | | | | |
| chr16 | RP11-65J21.4 | 4 | 75 | 8 | | | | | |
| chr9 | RP11-554F20.1 | 3 | 79 | 8 | | | | | |
| chr16 | MPHOSPH6 | 3 | 62 | 8 | MPHOSPH6 | M-phase phosphoprotein 6 | Nucleus | other | |
| chr16 | NPW | 4 | 171 | 8 | NPW | neuropeptide W | Extracellular Space | other | |
| chr2 | PELI1 | 3 | 67 | 8 | PELI1 | pellino E3 ubiquitin protein ligase | Cytoplasm | enzyme | |
| chr20 | JAG1 | 4 | 103 | 8 | JAG1 | jagged canonical Notch ligand 1 | Extracellular Space | growth factor | |
| chr16 | TAF1C | 4 | 212 | 8 | TAF1C | TATA-box binding protein associated factor, RNA | Nucleus | transcription regulator | |
| chr1 | GBAP1 | 5 | 328 | 8 | GBAP1 | glucosylceramidase beta pseudogene 1 | Other | other | |
| chr15 | THBS1 | 3 | 92 | 8 | THBS1 | thrombospondin 1 | Extracellular Space | other | |
| chr12 | KLRD1 | 3 | 68 | 8 | KLRD1 | killer cell lectin like receptor D1 | Plasma Membrane | transmembrane recepto | |
| chr10 | TRUB1 | 3 | 89 | 8 | TRUB1 | TruB pseudouridine synthase family member 1 | Other | enzyme | |
| chr12 | INHBC | 3 | 99 | 8 | INHBC | inhibin subunit beta C | Extracellular Space | growth factor | |
| chr3 | CCDC51 | 3 | 177 | 8 | CCDC51 | coiled-coil domain containing 51 | Cytoplasm | other | |
| chr11 | RP11-375D13.2 | 4 | 105 | 8 | | | | | |
| chr17 | RP11-599B13.7 | 3 | 98 | 8 | | | | | |
| chr3 | TMEM212 | 3 | 63 | 8 | TMEM212 | transmembrane protein 212 | Other | other | |
| chr2 | AGBL5 | 3 | 84 | 8 | AGBL5 | ATP/GTP binding protein like 5 | Cytoplasm | peptidase | |
| chr17 | MYOCD | 3 | 68 | 8 | MYOCD | myocardin | Nucleus | transcription regulator | |
| chr2 | KRCC1 | 3 | 63 | 8 | KRCC1 | lysine rich coiled-coil 1 | Extracellular Space | other | |
| chr8 | CSPP1 | 3 | 84 | 8 | CSPP1 | centrosome and spindle pole associated protein 1 | Cytoplasm | other | |
| chr22 | CES5AP1 | 4 | 155 | 8 | CES5AP1 | carboxylesterase 5A pseudogene | Other | other | |
| chr19 | ZNF880 | 4 | 88 | 8 | ZNF880 | zinc finger protein 880 | Other | other | |
| chr3 | EOGT | 3 | 75 | 8 | EOGT | EGF domain specific O-linked N-acetylglucosamine transferase | Extracellular Space | enzyme | |
| chr12 | RDH16 | 4 | 98 | 8 | RDH16 | retinol dehydrogenase 16 | Cytoplasm | enzyme | |
| chr9 | NSMF | 4 | 126 | 8 | NSMF | NMDA receptor synaptonuclear signaling and neuronal migration factor | Extracellular Space | other | |
| chr18 | RP11-403A21.1 | 3 | 78 | 8 | | | | | |
| chr12 | KRT6C | 4 | 217 | 8 | KRT6C | keratin 6C | Cytoplasm | other | |
| chr15 | STOML1 | 3 | 145 | 8 | STOML1 | stomatin like 1 | Other | other | |
| chr15 | GOLGA8UP | 3 | 130 | 8 | GOLGA8UP | golgin A8 family member U, pseudogene | Other | other | |
| chr11 | OSBP | 4 | 75 | 8 | OSBP | oxysterol binding protein | Cytoplasm | transporter | |
| chr1 | AL592494.5 | 3 | 124 | 8 | | | | | |

| | | | | | | | | | |
|---|---|---|---|---|---|---|---|---|---|
| chr7 | *IRF5* | 4 | 179 | 8 | IRF5 | interferon regulatory factor 5 | Nucleus | transcription regulator | |
| chr8 | *EPPK1* | 4 | 159 | 8 | EPPK1 | epiplakin 1 | Cytoplasm | other | |
| chr1 | *RP11-543D5.1* | 4 | 102 | 8 | | | | | |
| chr11 | *CCDC34* | 4 | 142 | 8 | CCDC34 | coiled-coil domain containing 34 | Other | other | |
| chr12 | *TSFM* | 3 | 92 | 8 | TSFM | Ts translation elongation factor, mitochondrial | Cytoplasm | translation regulator | |
| chr3 | *ACTBP13* | 3 | 158 | 8 | ACTBP13 | ACTB pseudogene 13 | Other | other | |
| chr19 | *LRRC25* | 4 | 130 | 8 | LRRC25 | leucine rich repeat containing 25 | Other | other | |
| chr17 | *C17orf97* | 4 | 104 | 8 | C17orf97 | chromosome 17 open reading frame 97 | Other | other | |
| chr14 | *TNFAIP2* | 4 | 201 | 8 | TNFAIP2 | TNF alpha induced protein 2 | Extracellular Space | other | |
| chr12 | *C12orf66* | 4 | 113 | 8 | C12orf66 | chromosome 12 open reading frame 66 | Plasma Membrane | other | |
| chr5 | *CKMT2-AS1* | 4 | 73 | 8 | CKMT2-AS1 | CKMT2 antisense RNA 1 | Other | other | |
| chr15 | *ODF3L1* | 3 | 196 | 8 | ODF3L1 | outer dense fiber of sperm tails 3 like 1 | Other | other | |
| chr19 | *AC018755.16* | 4 | 103 | 8 | | | | | |
| chr14 | *SMEK1* | 3 | 63 | 8 | PPP4R3A | protein phosphatase 4 regulatory subunit 3A | Plasma Membrane | other | |
| chr19 | *CTD-2224J9.8* | 3 | 109 | 8 | | | | | |
| chr7 | *RP5-965K10.3* | 3 | 115 | 8 | | | | | |
| chr7 | *ASB10* | 4 | 122 | 8 | ASB10 | ankyrin repeat and SOCS box containing 10 | Cytoplasm | other | |
| chr19 | *SIGLEC5* | 3 | 100 | 8 | SIGLEC5 | sialic acid binding Ig like lectin 5 | Plasma Membrane | other | |
| chr12 | *CKAP4* | 4 | 90 | 8 | CKAP4 | cytoskeleton associated protein 4 | Cytoplasm | other | |
| chr1 | *STMN1* | 3 | 114 | 8 | STMN1 | stathmin 1 | Cytoplasm | other | |
| chr3 | *NFKBIZ* | 3 | 83 | 8 | NFKBIZ | NFKB inhibitor zeta | Nucleus | transcription regulator | |
| chr6 | *ZNF451* | 3 | 115 | 8 | ZNF451 | zinc finger protein 451 | Nucleus | enzyme | |
| chrX | *CHIC1* | 3 | 80 | 8 | CHIC1 | cysteine rich hydrophobic domain | Plasma Membrane | other | |
| chr5 | *RP11-67M9.1* | 4 | 103 | 8 | | | | | |
| chr1 | *RP11-251P6.1* | 3 | 67 | 8 | | | | | |
| chr3 | *SEMA3B* | 4 | 172 | 8 | SEMA3B | semaphorin 3B | Extracellular Space | other | |
| chr15 | *GOLGA8N* | 3 | 121 | 8 | GA8K (includes ot | golgin A8 family member K | Other | other | |
| chr1 | *COA6* | 3 | 152 | 8 | COA6 | cytochrome c oxidase assembly factor 6 | Cytoplasm | other | |
| chr4 | *RP11-729M20.1* | 3 | 100 | 8 | | | | | |
| chr17 | *CTD-2006K23.1* | 3 | 134 | 8 | | | | | |
| chr13 | *TPTE2P2* | 3 | 99 | 8 | TPTE2P2 | transmembrane phosphoinositide 3-phosphatase and tensin homolog 2 pseudogene 2 | Other | other | |
| chr16 | *TP53TG3C* | 4 | 220 | 8 | TG3D (includes o | TP53 target 3D | Other | other | |
| chr8 | *AC145110.1* | 4 | 88 | 8 | | | | | |
| chr11 | *MTA2* | 4 | 108 | 8 | MTA2 | metastasis associated 1 family member 2 | Nucleus | transcription regulator | |
| chr19 | *PSMD8* | 4 | 108 | 8 | PSMD8 | proteasome 26S subunit, non-ATPase 8 | Cytoplasm | other | |
| chr11 | *AP000442.4* | 3 | 114 | 8 | | | | | |
| chr19 | *DHDH* | 3 | 133 | 8 | DHDH | dihydrodiol dehydrogenase | Other | enzyme | |
| chr15 | *FES* | 4 | 396 | 8 | FES | FES proto-oncogene, tyrosine | Cytoplasm | kinase | lorlatinib |
| chr1 | *AADACL3* | 4 | 79 | 8 | AADACL3 | arylacetamide deacetylase like 3 | Other | other | |
| chr15 | *CYP1A2* | 4 | 158 | 8 | CYP1A2 | cytochrome P450 family 1 subfamily A member 2 | Cytoplasm | enzyme | |
| chr11 | *MED19* | 3 | 110 | 8 | MED19 | mediator complex subunit 19 | Nucleus | other | |
| chr2 | *VSNL1* | 3 | 110 | 8 | VSNL1 | visinin like 1 | Cytoplasm | other | |
| chr19 | *LINC00905* | 4 | 124 | 8 | LINC00905 | long intergenic non-protein coding RNA 905 | Other | other | |
| chr9 | *IFT74* | 3 | 74 | 8 | IFT74 | intraflagellar transport 74 | Cytoplasm | other | |
| chr15 | *RP11-16E12.2* | 3 | 117 | 8 | | | | | |
| chr8 | *ASH2L* | 3 | 88 | 8 | ASH2L | ASH2 like, histone lysine methyltransferase complex | Nucleus | transcription regulator | |
| chr14 | *RP11-293M10.1* | 3 | 99 | 8 | | | | | |
| chr12 | *IRAK4* | 3 | 106 | 8 | IRAK4 | interleukin 1 receptor associated kinase 4 | Cytoplasm | kinase | PF-06650833, CA-4948 |
| chr1 | *MINOS1-NBL1* | 4 | 96 | 8 | COS10-NBL1/NB | NBL1, DAN family BMP | Nucleus | other | |
| chr15 | *PIF1* | 4 | 157 | 8 | PIF1 | PIF1 5'-to-3' DNA helicase | Nucleus | enzyme | |
| chr1 | *RAD54L* | 3 | 100 | 8 | RAD54L | RAD54 like | Nucleus | enzyme | |
| chr3 | *LINC00636* | 4 | 95 | 8 | LINC00636 | long intergenic non-protein coding RNA 636 | Other | other | |
| chr10 | *CHUK* | 3 | 82 | 8 | CHUK | component of inhibitor of nuclear factor kappa B kinase complex | Cytoplasm | kinase | bardoxolone methyl |
| chr1 | *MMACHC* | 3 | 101 | 8 | MMACHC | metabolism of cobalamin associated C | Cytoplasm | enzyme | |
| chr17 | *FAM64A* | 4 | 164 | 8 | PIMREG | PICALM interacting mitotic | Nucleus | other | |
| chr4 | *NDUFC1* | 3 | 57 | 8 | NDUFC1 | NADH:ubiquinone oxidoreductase subunit C1 | Cytoplasm | enzyme | |
| chr7 | *AC003991.3* | 4 | 89 | 8 | | | | | |
| chr7 | *C7orf63* | 3 | 84 | 8 | CFAP69 | cilia and flagella associated | Cytoplasm | other | |
| chr17 | *PSME3* | 3 | 90 | 8 | PSME3 | proteasome activator subunit 3 | Cytoplasm | peptidase | |
| chr20 | *FAM83D* | 4 | 121 | 8 | FAM83D | family with sequence similarity 83 member D | Cytoplasm | other | |
| chr3 | *ARGFX* | 4 | 85 | 8 | ARGFX | arginine-fifty homeobox | Other | other | |
| chr2 | *HOXD3* | 3 | 103 | 8 | HOXD3 | homeobox D3 | Nucleus | transcription regulator | |
| chr5 | *BTNL9* | 4 | 522 | 8 | BTNL9 | butyrophilin like 9 | Plasma Membrane | other | |
| chr6 | *HFE* | 3 | 95 | 8 | HFE | homeostatic iron regulator | Plasma Membrane | transmembrane receptor | |
| chr2 | *MTHFD2* | 4 | 83 | 8 | MTHFD2 | methylenetetrahydrofolate dehydrogenase (NADP+ dependent) 2, methenyltetrahydrofolate | Cytoplasm | enzyme | |
| chr5 | *RPL26L1* | 4 | 202 | 8 | RPL26L1 | ribosomal protein L26 like 1 | Cytoplasm | other | |

| | | | | | | | | | |
|---|---|---|---|---|---|---|---|---|---|
| chr17 | *MRPS23* | 3 | 101 | 8 | MRPS23 | mitochondrial ribosomal protein | Cytoplasm | other | |
| chr3 | *TTLL3* | 3 | 173 | 8 | TTLL3 | tubulin tyrosine ligase like 3 | Extracellular Space | enzyme | |
| chr1 | *C4BPA* | 4 | 103 | 8 | C4BPA | complement component 4 binding protein alpha | Extracellular Space | other | |
| chr4 | *RP11-659O3.1* | 3 | 89 | 8 | | | | | |
| chr5 | *SLC22A4* | 3 | 149 | 8 | SLC22A4 | solute carrier family 22 member 4 | Plasma Membrane | transporter | |
| chr17 | *FSCN2* | 4 | 263 | 8 | FSCN2 | fascin actin-bundling protein 2, | Cytoplasm | other | |
| chr15 | *RP13-996F3.5* | 3 | 139 | 8 | | | | | |
| chr14 | *XRCC3* | 4 | 203 | 8 | XRCC3 | X-ray repair cross complementing | Nucleus | enzyme | |
| chr6 | *DLK2* | 3 | 121 | 8 | DLK2 | delta like non-canonical Notch ligand 2 | Extracellular Space | other | |
| chr5 | *CTC-558O2.2* | 4 | 109 | 8 | | | | | |
| chr2 | *AC068057.2* | 3 | 147 | 8 | | | | | |
| chr15 | *RPUSD2* | 4 | 132 | 8 | RPUSD2 | RNA pseudouridine synthase domain containing 2 | Other | enzyme | |
| chr7 | *OR2A1-AS1* | 4 | 95 | 8 | OR2A1-AS1 | OR2A1 antisense RNA 1 | Other | other | |
| chr1 | *NTPCR* | 4 | 107 | 8 | NTPCR | nucleoside-triphosphatase, cancer-related | Cytoplasm | other | |
| chr3 | *RP11-644C3.1* | 3 | 78 | 8 | | | | | |
| chr3 | *CNBP* | 3 | 160 | 8 | CNBP | CCHC-type zinc finger nucleic acid binding protein | Nucleus | transcription regulator | |
| chr3 | *ROPN1* | 4 | 134 | 8 | ROPN1 | rhophilin associated tail protein 1 | Cytoplasm | other | |
| chrX | *TRPC5* | 3 | 71 | 8 | TRPC5 | transient receptor potential cation channel subfamily C member 5 | Plasma Membrane | ion channel | |
| chr19 | *CIRBP* | 4 | 225 | 8 | CIRBP | cold inducible RNA binding | Nucleus | translation regulator | |
| chr5 | *RP11-79P5.3* | 4 | 79 | 8 | | | | | |
| chr7 | *EPHB6* | 4 | 147 | 8 | EPHB6 | EPH receptor B6 | Plasma Membrane | kinase | |
| chr6 | *KLHL32* | 3 | 111 | 8 | KLHL32 | kelch like family member 32 | Other | other | |
| chr17 | *MFAP4* | 3 | 125 | 8 | MFAP4 | microfibril associated protein 4 | Extracellular Space | other | |
| chr1 | *SLC39A1* | 4 | 198 | 8 | SLC39A1 | solute carrier family 39 member 1 | Plasma Membrane | transporter | |
| chr2 | *AC104654.1* | 3 | 139 | 8 | | | | | |
| chr21 | *IFNAR1* | 3 | 72 | 8 | IFNAR1 | interferon alpha and beta receptor subunit 1 | Plasma Membrane | transmembrane receptor | interferon alfa-n1, pegintron/ribavirin, interferon alfacon-1/ribavirin, PEG-interferon alfa-2a/telaprevir, imatinib/peginterferon alfa-2a, IFNA2B/ribavirin, pentostatin/recombinant interferon, interferon alfacon-1, hydroxyurea/recombinant interferon, recombinant interferon, cytarabine/IFNA2B, PEG-interferon alfa-2a/ribavirin, interferon beta-1b, IFNA2, cladribine/recombinant interferon, [illegible] |
| chr7 | *RP5-1132H15.1* | 3 | 66 | 8 | | | | | |
| chr16 | *RP11-252A24.5* | 4 | 108 | 8 | | | | | |
| chr2 | *AC097724.3* | 3 | 73 | 8 | | | | | |
| chr5 | *SREK1* | 3 | 73 | 8 | SREK1 | splicing regulatory glutamic acid and lysine rich protein 1 | Nucleus | other | |
| chr4 | *TECRL* | 3 | 72 | 8 | TECRL | trans-2,3-enoyl-CoA reductase | Cytoplasm | other | |
| chr10 | *RP11-57H14.3* | 4 | 101 | 8 | | | | | |
| chr1 | *PEX10* | 4 | 256 | 8 | PEX10 | peroxisomal biogenesis factor 10 | Cytoplasm | other | |
| chr1 | *RUSC1* | 4 | 221 | 8 | RUSC1 | RUN and SH3 domain containing | Cytoplasm | other | |
| chr7 | *RP11-325F22.2* | 3 | 95 | 8 | | | | | |
| chr17 | *NAT9* | 3 | 140 | 8 | NAT9 | N-acetyltransferase 9 (putative) | Other | other | |
| chr3 | *OPA1* | 3 | 99 | 8 | OPA1 | OPA1 mitochondrial dynamin like GTPase | Cytoplasm | enzyme | |
| chr1 | *RAB4A* | 4 | 126 | 8 | RAB4A | RAB4A, member RAS oncogene family | Cytoplasm | enzyme | |
| chr2 | *PAX3* | 3 | 87 | 8 | PAX3 | paired box 3 | Nucleus | transcription regulator | |
| chr11 | *GYLTL1B* | 4 | 278 | 8 | LARGE2 | LARGE xylosyl- and glucuronyltransferase 2 | Cytoplasm | enzyme | |
| chr20 | *CSRP2BP* | 3 | 83 | 8 | KAT14 | lysine acetyltransferase 14 | Nucleus | enzyme | |
| chr1 | *FAM189B* | 4 | 211 | 8 | FAM189B | family with sequence similarity 189 member B | Other | other | |
| chr16 | *RP11-989E6.10* | 3 | 188 | 8 | | | | | |
| chr2 | *TMEM37* | 4 | 194 | 8 | TMEM37 | transmembrane protein 37 | Plasma Membrane | ion channel | Hu8F4 |
| chr7 | *ZNF783* | 4 | 180 | 8 | ZNF783 | zinc finger family member 783 | Other | other | |
| chrX | *CXorf38* | 4 | 76 | 8 | CXorf38 | chromosome X open reading frame 38 | Other | other | |
| chr1 | *DNALI1* | 3 | 152 | 8 | DNALI1 | dynein axonemal light intermediate chain 1 | Cytoplasm | other | |
| chr17 | *CCDC144NL* | 3 | 162 | 8 | CCDC144NL | coiled-coil domain containing 144 family, N-terminal like | Other | other | |
| chr12 | *AMDHD1* | 4 | 108 | 8 | AMDHD1 | amidohydrolase domain | Cytoplasm | enzyme | |
| chr12 | *RP13-653N12.1* | 4 | 81 | 8 | | | | | |
| chr8 | *RP11-706J10.1* | 3 | 134 | 8 | | | | | |
| chr7 | *RP11-458F8.1* | 3 | 82 | 8 | | | | | |

| | | | | | | | | | |
|---|---|---|---|---|---|---|---|---|---|
| chr19 | *PGLS* | 4 | 151 | 8 | PGLS | 6-phosphogluconolactonase | Cytoplasm | enzyme | |
| chr19 | *CTU1* | 4 | 112 | 8 | CTU1 | cytosolic thiouridylase subunit 1 | Cytoplasm | other | |
| chr10 | *TIAL1* | 2 | 67 | 8 | TIAL1 | TIA1 cytotoxic granule associated RNA binding protein like 1 | Nucleus | transcription regulator | |
| chr2 | *AMER3* | 3 | 144 | 8 | AMER3 | APC membrane recruitment | Other | other | |
| chr1 | *TUFT1* | 4 | 134 | 8 | TUFT1 | tuftelin 1 | Cytoplasm | other | |
| chr1 | *SCAMP3* | 4 | 237 | 8 | SCAMP3 | secretory carrier membrane | Cytoplasm | transporter | |
| chr19 | *RP11-678G14.2* | 4 | 93 | 8 | | | | | |
| chr14 | *WDHD1* | 3 | 67 | 8 | WDHD1 | WD repeat and HMG-box DNA binding protein 1 | Nucleus | transcription regulator | |
| chr6 | *RP11-716O23.2* | 4 | 83 | 8 | | | | | |
| chrX | *ZNF275* | 3 | 108 | 8 | ZNF275 | zinc finger protein 275 | Nucleus | transcription regulator | |
| chr4 | *SLAIN2* | 3 | 83 | 8 | SLAIN2 | SLAIN motif family member 2 | Cytoplasm | other | |
| chr5 | *FCHSD1* | 3 | 168 | 8 | FCHSD1 | FCH and double SH3 domains 1 | Other | other | |
| chr5 | *CTD-2118P12.1* | 3 | 204 | 8 | | | | | |
| chr21 | *AP001464.4* | 3 | 88 | 8 | | | | | |
| chr17 | *RP11-353N14.5* | 3 | 178 | 8 | | | | | |
| chr16 | *DYNC1LI2* | 4 | 87 | 8 | DYNC1LI2 | dynein cytoplasmic 1 light intermediate chain 2 | Cytoplasm | other | |
| chr11 | *ZW10* | 4 | 92 | 8 | ZW10 | zw10 kinetochore protein | Nucleus | other | |
| chr7 | *RP11-340I6.7* | 3 | 113 | 8 | | | | | |
| chr10 | *USMG5* | 3 | 85 | 8 | ATP5MD | ATP synthase membrane subunit DAPIT | Cytoplasm | other | |
| chr6 | *STMND1* | 4 | 115 | 8 | STMND1 | stathmin domain containing 1 | Other | other | |
| chr4 | *RP11-125O18.1* | 4 | 99 | 8 | | | | | |
| chr1 | *RP5-1066H13.4* | 4 | 74 | 8 | | | | | |
| chr12 | *RHOF* | 4 | 138 | 8 | RHOF | ras homolog family member F, filopodia associated | Cytoplasm | enzyme | |
| chr2 | *SEMA4C* | 4 | 258 | 8 | SEMA4C | semaphorin 4C | Plasma Membrane | other | |
| chrX | *CT45A6* | 3 | 126 | 8 | 45A1 (includes oth | cancer/testis antigen family 45 member A2 | Other | other | |
| chr5 | *RP11-661C8.2* | 4 | 489 | 8 | | | | | |
| chr16 | *ACSM5* | 3 | 76 | 8 | ACSM5 | acyl-CoA synthetase medium chain family member 5 | Other | enzyme | |
| chr10 | *RP11-298E9.6* | 3 | 91 | 8 | | | | | |
| chr21 | *AP000476.1* | 3 | 62 | 8 | | | | | |
| chr7 | *RBAK-RBAKDN* | 4 | 124 | 8 | RBAK-RBAKDN | RBAK-RBAKDN readthrough | Other | other | |
| chr5 | *AC138035.2* | 4 | 120 | 8 | | | | | |
| chr2 | *BOK-AS1* | 3 | 110 | 8 | BOK-AS1 | BOK antisense RNA 1 | Other | other | |
| chr7 | *ZYX* | 4 | 158 | 8 | ZYX | zyxin | Plasma Membrane | other | |
| chr19 | *LRFN3* | 4 | 92 | 8 | LRFN3 | leucine rich repeat and fibronectin type III domain containing 3 | Plasma Membrane | other | |
| chr11 | *RP11-672A2.1* | 4 | 267 | 8 | | | | | |
| chr22 | *SSTR3* | 3 | 153 | 8 | SSTR3 | somatostatin receptor 3 | Plasma Membrane | 3-protein coupled recepto | octreotide, everolimus/pasireotide, octreotide/prochlorperazine, haloperidol/octreotide |
| chr1 | *RP3-395M20.8* | 4 | 194 | 8 | | | | | |
| chr19 | *AC005614.5* | 3 | 61 | 8 | | | | | |
| chr1 | *NBPF14* | 4 | 296 | 8 | PF1 (includes othe | NBPF member 11 | Other | other | |
| chr17 | *RP1-56K13.5* | 4 | 78 | 8 | | | | | |
| chr12 | *ZBTB39* | 3 | 122 | 8 | ZBTB39 | zinc finger and BTB domain containing 39 | Nucleus | transcription regulator | |
| chr13 | *LINC00376* | 3 | 65 | 8 | LINC00376 | long intergenic non-protein coding RNA 376 | Other | other | |
| chr5 | *RP11-1379J22.5* | 4 | 154 | 8 | | | | | |
| chr19 | *ZNF439* | 4 | 88 | 8 | ZNF439 | zinc finger protein 439 | Nucleus | transcription regulator | |
| chr12 | *YARS2* | 3 | 105 | 8 | YARS2 | tyrosyl-tRNA synthetase 2 | Cytoplasm | enzyme | |
| chr12 | *C12orf50* | 3 | 75 | 8 | C12orf50 | chromosome 12 open reading frame 50 | Other | other | |
| chr2 | *AC012501.2* | 3 | 86 | 8 | | | | | |
| chr4 | *RP11-340B18.1* | 3 | 65 | 8 | | | | | |
| chr19 | *WASH5P* | 3 | 238 | 8 | WASH5P | WASP family homolog 5, pseudogene | Other | other | |
| chr2 | *SPATA3* | 4 | 104 | 8 | SPATA3 | spermatogenesis associated 3 | Other | other | |
| chr12 | *CCDC59* | 4 | 69 | 8 | CCDC59 | coiled-coil domain containing 59 | Nucleus | other | |
| chr9 | *RP11-216B9.6* | 3 | 118 | 8 | | | | | |
| chr8 | *RP11-258J10.1* | 4 | 78 | 8 | | | | | |
| chr3 | *ENPP7P2* | 3 | 100 | 8 | ENPP7P2 | ectonucleotide pyrophosphatase/phosphodiesterase 7 pseudogene 2 | Other | other | |
| chr15 | *NEIL1* | 4 | 206 | 8 | NEIL1 | nei like DNA glycosylase 1 | Nucleus | enzyme | |
| chr17 | *TBC1D3P2* | 3 | 275 | 8 | TBC1D3P2 | TBC1 domain family member 3 pseudogene 2 | Other | other | |
| chr20 | *ADAM33* | 4 | 179 | 8 | ADAM33 | ADAM metallopeptidase domain | Plasma Membrane | peptidase | |
| chr9 | *KIF27* | 4 | 68 | 8 | KIF27 | kinesin family member 27 | Cytoplasm | other | |
| chr11 | *FAM160A2* | 4 | 229 | 8 | FAM160A2 | family with sequence similarity 160 member A2 | Cytoplasm | other | |
| chr2 | *IL1RN* | 3 | 148 | 8 | IL1RN | interleukin 1 receptor antagonist | Extracellular Space | cytokine | |
| chr10 | *RP11-175I17.2* | 3 | 61 | 8 | | | | | |
| chr3 | *SLC22A13* | 3 | 111 | 8 | SLC22A13 | solute carrier family 22 member | Plasma Membrane | transporter | |
| chr8 | *RP11-962G15.1* | 4 | 66 | 8 | | | | | |
| chr8 | *RP11-468H14.2* | 4 | 67 | 8 | | | | | |
| chr3 | *PARP14* | 4 | 120 | 8 | PARP14 | poly(ADP-ribose) polymerase family member 14 | Cytoplasm | enzyme | |
| chr10 | *ENTPD1* | 4 | 66 | 8 | ENTPD1 | ectonucleoside triphosphate diphosphohydrolase 1 | Plasma Membrane | enzyme | TTX-030 |

| | | | | | | | | | |
|---|---|---|---|---|---|---|---|---|---|
| chr7 | *STRIP2* | 4 | 68 | 8 | STRIP2 | striatin interacting protein 2 | Cytoplasm | other | |
| chr17 | *RP11-1094H24.3* | 4 | 98 | 8 | | | | | |
| chr1 | *FDPS* | 4 | 135 | 8 | FDPS | farnesyl diphosphate synthase | Cytoplasm | enzyme | risedronic acid, zoledronic acid, minodronate, alendronate/cholecalciferol, pamidronic acid, alendronic acid, ibandronic acid |
| chr17 | *AC111186.1* | 3 | 147 | 8 | | | | | |
| chr12 | *KRT83* | 4 | 295 | 8 | KRT83 | keratin 83 | Cytoplasm | other | |
| chr19 | *KDELR1* | 3 | 118 | 8 | KDELR1 | KDEL endoplasmic reticulum protein retention receptor 1 | Cytoplasm | transporter | |
| chr5 | *PCYOX1L* | 3 | 86 | 8 | PCYOX1L | prenylcysteine oxidase 1 like | Other | other | |
| chr1 | *SASS6* | 3 | 85 | 8 | SASS6 | SAS-6 centriolar assembly | Cytoplasm | other | |
| chr17 | *KRT40* | 3 | 85 | 8 | KRT40 | keratin 40 | Cytoplasm | other | |
| chr4 | *RP11-565A3.2* | 3 | 77 | 8 | | | | | |
| chr2 | *MAP3K19* | 3 | 67 | 8 | MAP3K19 | mitogen-activated protein kinase kinase kinase 19 | Other | kinase | |
| chr10 | *MCM10* | 3 | 79 | 8 | MCM10 | minichromosome maintenance 10 replication initiation factor | Nucleus | other | |
| chr12 | *UNG* | 4 | 69 | 8 | UNG | uracil DNA glycosylase | Nucleus | enzyme | |
| chr5 | *AC010423.1* | 4 | 81 | 8 | | | | | |
| chr22 | *HORMAD2* | 3 | 57 | 8 | HORMAD2 | HORMA domain containing 2 | Nucleus | other | |
| chr14 | *RP11-1103G16.1* | 3 | 81 | 8 | | | | | |
| chr21 | *FTCD* | 3 | 154 | 8 | FTCD | formimidoyltransferase cyclodeaminase | Cytoplasm | enzyme | |
| chr3 | *IQCF1* | 4 | 97 | 8 | IQCF1 | IQ motif containing F1 | Cytoplasm | other | |
| chr2 | *AC079612.1* | 3 | 122 | 8 | | | | | |
| chr2 | *DUSP11* | 3 | 101 | 8 | DUSP11 | dual specificity phosphatase 11 | Nucleus | phosphatase | |
| chr19 | *ZNF534* | 4 | 84 | 8 | ZNF534 | zinc finger protein 534 | Other | other | |
| chr20 | *YWHAB* | 3 | 117 | 8 | YWHAB | tyrosine 3-monooxygenase/tryptophan 5-monooxygenase activation protein beta | Cytoplasm | other | |
| chr10 | *SFXN3* | 4 | 139 | 8 | SFXN3 | sideroflexin 3 | Cytoplasm | transporter | |
| chr2 | *EIF2AK2* | 3 | 68 | 8 | EIF2AK2 | eukaryotic translation initiation factor 2 alpha kinase 2 | Cytoplasm | kinase | mitapivat |
| chr9 | *ANKRD20A1* | 3 | 80 | 8 | D20A2 (includes c | ankyrin repeat domain 20 family member A2 | Plasma Membrane | other | |
| chr5 | *REEP2* | 3 | 126 | 8 | REEP2 | receptor accessory protein 2 | Plasma Membrane | other | |
| chr12 | *TMEM52B* | 3 | 71 | 8 | TMEM52B | transmembrane protein 52B | Cytoplasm | other | |
| chr10 | *EXOSC1* | 4 | 92 | 8 | EXOSC1 | exosome component 1 | Nucleus | enzyme | |
| chr11 | *AP000442.1* | 3 | 68 | 8 | | | | | |
| chr3 | *CAPN7* | 3 | 63 | 8 | CAPN7 | calpain 7 | Cytoplasm | peptidase | |
| chr3 | *GORASP1* | 3 | 173 | 8 | GORASP1 | golgi reassembly stacking protein | Cytoplasm | other | |
| chr20 | *WFDC8* | 3 | 91 | 8 | WFDC8 | WAP four-disulfide core domain 8 | Extracellular Space | other | |
| chr12 | *ATN1* | 3 | 157 | 8 | ATN1 | atrophin 1 | Nucleus | transcription regulator | |
| chr8 | *RP11-600K15.1* | 3 | 70 | 8 | | | | | |
| chr2 | *CCT7* | 4 | 125 | 8 | CCT7 | chaperonin containing TCP1 subunit 7 | Cytoplasm | other | |
| chr3 | *RP11-379B18.6* | 4 | 120 | 8 | | | | | |
| chr3 | *PARP3* | 3 | 160 | 8 | PARP3 | poly(ADP-ribose) polymerase family member 3 | Nucleus | enzyme | olaparib, rucaparib |
| chr12 | *ANAPC7* | 4 | 79 | 8 | ANAPC7 | anaphase promoting complex subunit 7 | Nucleus | other | |
| chr19 | *CYP4F24P* | 4 | 108 | 8 | CYP4F24P | cytochrome P450 family 4 subfamily F member 24, | Other | other | |
| chr3 | *WDR48* | 3 | 80 | 8 | WDR48 | WD repeat domain 48 | Cytoplasm | other | |
| chr19 | *AC005932.1* | 3 | 103 | 8 | | | | | |
| chr7 | *NAMPT* | 3 | 61 | 8 | NAMPT | nicotinamide phosphoribosyltransferase | Extracellular Space | cytokine | daporinad, KPT-9274 |
| chrX | *PIM2* | 3 | 128 | 8 | PIM2 | Pim-2 proto-oncogene, serine/threonine kinase | Nucleus | kinase | SGI 1776, PIM/FLT3 kinase inhibitor SEL24, LGH447, |
| chr12 | *SLCO1A2* | 3 | 93 | 8 | SLCO1A2 | solute carrier organic anion transporter family member 1A2 | Plasma Membrane | transporter | |
| chr17 | *RNFT1P3* | 3 | 63 | 8 | RNFT1P3 | ring finger protein, transmembrane 1 pseudogene 3 | Other | other | |
| chr15 | *CATSPER2P1* | 3 | 85 | 8 | CATSPER2P1 | cation channel sperm associated 2 pseudogene 1 | Other | other | |
| chr1 | *TSSK3* | 4 | 87 | 8 | TSSK3 | testis specific serine kinase 3 | Other | kinase | |
| chr9 | *UAP1L1* | 4 | 238 | 8 | UAP1L1 | UDP-N-acetylglucosamine pyrophosphorylase 1 like 1 | Other | other | |
| chr16 | *RP11-538I12.2* | 3 | 61 | 8 | | | | | |
| chr1 | *LINC00210* | 3 | 118 | 8 | LINC00210 | long intergenic non-protein coding RNA 210 | Other | other | |
| chr1 | *NUF2* | 3 | 84 | 8 | NUF2 | NUF2 component of NDC80 kinetochore complex | Nucleus | other | |
| chr1 | *C1orf172* | 4 | 118 | 8 | KDF1 | keratinocyte differentiation factor | Plasma Membrane | other | |
| chrX | *RP11-104D21.3* | 4 | 73 | 8 | | | | | |
| chr21 | *ZNF295-AS1* | 3 | 110 | 8 | ZNF295-AS1 | ZNF295 antisense RNA 1 | Other | other | |
| chr20 | *RP5-1164C1.2* | 3 | 91 | 8 | | | | | |
| chr1 | *UAP1* | 4 | 96 | 8 | UAP1 | UDP-N-acetylglucosamine pyrophosphorylase 1 | Nucleus | enzyme | |
| chr15 | *AP3B2* | 4 | 119 | 8 | AP3B2 | adaptor related protein complex 3 subunit beta 2 | Cytoplasm | other | |
| chr16 | *CYBA* | 4 | 150 | 8 | CYBA | cytochrome b-245 alpha chain | Cytoplasm | enzyme | |
| chr3 | *ZDHHC19* | 3 | 145 | 8 | ZDHHC19 | zinc finger DHHC-type containing | Cytoplasm | other | |

| | | | | | | | | |
|---|---|---|---|---|---|---|---|---|
| chr10 | ARMC3 | 3 | 69 | 8 | ARMC3 | armadillo repeat containing 3 | Cytoplasm | other |
| chr22 | IFT27 | 3 | 107 | 8 | IFT27 | intraflagellar transport 27 | Extracellular Space | enzyme |
| chr1 | S100A16 | 3 | 193 | 8 | S100A16 | S100 calcium binding protein A16 | Nucleus | other |
| chr2 | NCKAP1 | 3 | 72 | 8 | NCKAP1 | NCK associated protein 1 | Plasma Membrane | other |
| chr9 | FKTN | 3 | 98 | 7 | FKTN | fukutin | Extracellular Space | other |
| chr19 | ZNF525 | 4 | 73 | 7 | ZNF525 | zinc finger protein 525 | Nucleus | transcription regulator |
| chr17 | RAD51L3-RFFL | 3 | 105 | 7 | RAD51L3-RFFL | RAD51L3-RFFL readthrough | Other | other |
| chr11 | CCDC82 | 3 | 129 | 7 | CCDC82 | coiled-coil domain containing 82 | Other | other |
| chr1 | RP3-510D11.1 | 3 | 152 | 7 | | | | |
| chr7 | RP5-945F2.2 | 3 | 58 | 7 | | | | |
| chr4 | RP11-83C7.2 | 3 | 94 | 7 | | | | |
| chr1 | ZNF326 | 3 | 65 | 7 | ZNF326 | zinc finger protein 326 | Nucleus | transcription regulator |
| chr5 | RP11-497H16.2 | 2 | 167 | 7 | | | | |
| chr2 | UPP2 | 3 | 55 | 7 | UPP2 | uridine phosphorylase 2 | Cytoplasm | enzyme |
| chr17 | RP11-429O1.1 | 3 | 84 | 7 | | | | |
| chr12 | CD163L1 | 3 | 65 | 7 | CD163L1 | CD163 molecule like 1 | Plasma Membrane | transmembrane receptor |
| chr9 | NINJ1 | 3 | 149 | 7 | NINJ1 | ninjurin 1 | Plasma Membrane | other |
| chr1 | RP11-552D4.1 | 3 | 150 | 7 | | | | |
| chr16 | PRSS33 | 3 | 160 | 7 | PRSS33 | serine protease 33 | Extracellular Space | peptidase |
| chr12 | TROAP | 3 | 232 | 7 | TROAP | trophinin associated protein | Cytoplasm | peptidase |
| chr15 | RP11-467N20.5 | 3 | 201 | 7 | | | | |
| chr4 | RP11-290F5.2 | 3 | 85 | 7 | | | | |
| chr17 | KRT16 | 3 | 170 | 7 | KRT16 | keratin 16 | Cytoplasm | other |
| chrX | ABCD1 | 4 | 164 | 7 | ABCD1 | ATP binding cassette subfamily D member 1 | Plasma Membrane | transporter |
| chr1 | DMBX1 | 4 | 177 | 7 | DMBX1 | diencephalon/mesencephalon homeobox 1 | Nucleus | transcription regulator |
| chr14 | CCNK | 3 | 116 | 7 | CCNK | cyclin K | Nucleus | kinase |
| chr9 | MIR31HG | 3 | 79 | 7 | MIR31HG | MIR31 host gene | Other | other |
| chr1 | GS1-204I12.1 | 4 | 91 | 7 | | | | |
| chr11 | SLC36A4 | 3 | 136 | 7 | SLC36A4 | solute carrier family 36 member 4 | Plasma Membrane | transporter |
| chr20 | RP4-785G19.5 | 3 | 99 | 7 | | | | |
| chr20 | LINC00658 | 3 | 92 | 7 | LINC00658 | long intergenic non-protein coding RNA 658 | Other | other |
| chr5 | CSNK1G3 | 3 | 70 | 7 | CSNK1G3 | casein kinase 1 gamma 3 | Cytoplasm | kinase |
| chr10 | ST8SIA6 | 3 | 65 | 7 | ST8SIA6 | ST8 alpha-N-acetyl-neuraminide alpha-2,8-sialyltransferase 6 | Cytoplasm | enzyme |
| chr14 | LINC00871 | 3 | 68 | 7 | LINC00871 | long intergenic non-protein coding RNA 871 | Other | other |
| chrX | RS1 | 4 | 121 | 7 | RS1 | retinoschisin 1 | Extracellular Space | other |
| chr21 | SH3BGR | 3 | 89 | 7 | SH3BGR | SH3 domain binding glutamate rich protein | Cytoplasm | other |
| chr5 | RP11-32D16.1 | 4 | 95 | 7 | | | | |
| chr16 | RP11-178L8.4 | 4 | 200 | 7 | | | | |
| chr11 | RHOG | 4 | 279 | 7 | RHOG | ras homolog family member G | Cytoplasm | enzyme |
| chr11 | VWA5A | 3 | 79 | 7 | VWA5A | von Willebrand factor A domain containing 5A | Nucleus | other |
| chr19 | ZNF235 | 3 | 68 | 7 | ZNF235 | zinc finger protein 235 | Nucleus | transcription regulator |
| chr6 | CCDC162P | 3 | 80 | 7 | CCDC162P | coiled-coil domain containing 162, pseudogene | Other | other |
| chrX | GJB1 | 3 | 115 | 7 | GJB1 | gap junction protein beta 1 | Plasma Membrane | transporter |
| chr1 | RP11-396C23.2 | 3 | 139 | 7 | | | | |
| chr1 | FAM46B | 3 | 141 | 7 | TENT5B | terminal nucleotidyltransferase | Other | other |
| chr2 | MZT2B | 3 | 120 | 7 | MZT2B | mitotic spindle organizing protein | Cytoplasm | other |
| chr5 | MROH2B | 4 | 260 | 7 | MROH2B | maestro heat like repeat family member 2B | Cytoplasm | other |
| chr14 | MLH3 | 3 | 72 | 7 | MLH3 | mutL homolog 3 | Nucleus | other |
| chr19 | ZNF616 | 3 | 69 | 7 | ZNF616 | zinc finger protein 616 | Nucleus | transcription regulator |
| chr17 | AC015922.6 | 4 | 89 | 7 | | | | |
| chr8 | PROSC | 3 | 67 | 7 | PLPBP | pyridoxal phosphate binding | Cytoplasm | enzyme |
| chr7 | RP11-760D2.11 | 3 | 218 | 7 | | | | |
| chr2 | RQCD1 | 3 | 82 | 7 | CNOT9 | CCR4-NOT transcription complex subunit 9 | Cytoplasm | other |
| chr2 | WDR12 | 3 | 62 | 7 | WDR12 | WD repeat domain 12 | Cytoplasm | other |
| chr1 | NCDN | 4 | 205 | 7 | NCDN | neurochondrin | Cytoplasm | other |
| chr5 | MIR4454 | 4 | 159 | 7 | MIR4454 | microRNA 4454 | Cytoplasm | microRNA |
| chr19 | GTPBP3 | 4 | 158 | 7 | GTPBP3 | GTP binding protein 3, | Cytoplasm | enzyme |
| chr7 | FSCN3 | 4 | 98 | 7 | FSCN3 | fascin actin-bundling protein 3 | Cytoplasm | other |
| chr1 | RP1-65J11.1 | 3 | 106 | 7 | | | | |
| chr1 | MAGOH | 3 | 72 | 7 | MAGOH | mago homolog, exon junction complex subunit | Nucleus | other |
| chr8 | KIAA1875 | 4 | 145 | 7 | WDR97 | WD repeat domain 97 | Other | other |
| chr5 | SNX18 | 3 | 114 | 7 | SNX18 | sorting nexin 18 | Cytoplasm | transporter |
| chr19 | IGFL4 | 4 | 89 | 7 | IGFL4 | IGF like family member 4 | Extracellular Space | other |
| chr17 | SIRT7 | 4 | 148 | 7 | SIRT7 | sirtuin 7 | Nucleus | enzyme |
| chr11 | RP11-168K9.2 | 4 | 97 | 7 | | | | |
| chr14 | ASPG | 4 | 210 | 7 | ASPG | asparaginase | Cytoplasm | enzyme |
| chr3 | TRNT1 | 3 | 64 | 7 | TRNT1 | tRNA nucleotidyl transferase 1 | Cytoplasm | enzyme |
| chr15 | TICRR | 3 | 83 | 7 | TICRR | TOPBP1 interacting checkpoint and replication regulator | Nucleus | other |
| chr12 | RP11-313F23.4 | 4 | 120 | 7 | | | | |
| chr11 | GLYATL2 | 4 | 63 | 7 | GLYATL2 | glycine-N-acyltransferase like 2 | Cytoplasm | enzyme |
| chr19 | C19orf55 | 4 | 172 | 7 | PROSER3 | proline and serine rich 3 | Other | other |
| chr3 | SERP1 | 4 | 84 | 7 | SERP1 | stress associated endoplasmic reticulum protein 1 | Cytoplasm | other |
| chr1 | NDUFS2 | 4 | 141 | 7 | NDUFS2 | NADH:ubiquinone oxidoreductase core subunit S2 | Cytoplasm | enzyme |
| chr1 | EXO5 | 3 | 73 | 7 | EXO5 | exonuclease 5 | Nucleus | enzyme |
| chr17 | GRN | 4 | 156 | 7 | GRN | granulin precursor | Extracellular Space | growth factor |

| | | | | | | | | | |
|---|---|---|---|---|---|---|---|---|---|
| chr19 | *AC010642.1* | 3 | 107 | 7 | | | | | |
| chr3 | *DBR1* | 2 | 63 | 7 | DBR1 | debranching RNA lariats 1 | Nucleus | enzyme | |
| chr2 | *AC092168.3* | 4 | 145 | 7 | | | | | |
| chr11 | *RP11-159N11.3* | 3 | 125 | 7 | | | | | |
| chr3 | *LRRC58* | 3 | 120 | 7 | LRRC58 | leucine rich repeat containing 58 | Other | other | |
| chr3 | *SLC33A1* | 3 | 107 | 7 | SLC33A1 | solute carrier family 33 member 1 | Cytoplasm | transporter | |
| chr15 | *KIAA0101* | 3 | 99 | 7 | PCLAF | PCNA clamp associated factor | Nucleus | other | |
| chr17 | *TBC1D3P7* | 3 | 100 | 7 | TBC1D3P7 | | Other | other | |
| chr15 | *GDPGP1* | 4 | 168 | 7 | GDPGP1 | GDP-D-glucose phosphorylase 1 | Cytoplasm | enzyme | |
| chr12 | *KLRG1* | 3 | 66 | 7 | KLRG1 | killer cell lectin like receptor G1 | Plasma Membrane | other | |
| chr2 | *AC093901.1* | 4 | 193 | 7 | | | | | |
| chr5 | *GFRA3* | 3 | 78 | 7 | GFRA3 | GDNF family receptor alpha 3 | Plasma Membrane | transmembrane receptor | |
| chr17 | *CTB-22K21.2* | 3 | 146 | 7 | | | | | |
| chr7 | *ZFAND2A* | 4 | 187 | 7 | ZFAND2A | zinc finger AN1-type containing | Other | other | |
| chr19 | *BAX* | 3 | 111 | 7 | BAX | BCL2 associated X, apoptosis regulator | Cytoplasm | transporter | |
| chr14 | *RP11-7F17.7* | 3 | 100 | 7 | | | | | |
| chr1 | *METTL11B* | 3 | 95 | 7 | METTL11B | methyltransferase like 11B | Other | other | |
| chr9 | *XXyac-YM21GA2.* | 3 | 110 | 7 | | | | | |
| chr7 | *AC093627.12* | 4 | 270 | 7 | | | | | |
| chr22 | *SEC14L6* | 3 | 111 | 7 | SEC14L6 | SEC14 like lipid binding 6 | Other | other | |
| chr8 | *KCTD9* | 3 | 99 | 7 | KCTD9 | potassium channel tetramerization domain containing | Other | other | |
| chrX | *ATP7A* | 3 | 57 | 7 | ATP7A | ATPase copper transporting | Plasma Membrane | transporter | |
| chr7 | *MCM7* | 4 | 152 | 7 | MCM7 | minichromosome maintenance complex component 7 | Nucleus | enzyme | |
| chr5 | *RP11-404K5.2* | 3 | 376 | 7 | | | | | |
| chr4 | *EVC* | 3 | 69 | 7 | EVC | EvC ciliary complex subunit 1 | Cytoplasm | other | |
| chr5 | *RP11-1277A3.1* | 4 | 92 | 7 | | | | | |
| chr1 | *FCN3* | 3 | 130 | 7 | FCN3 | ficolin 3 | Extracellular Space | other | |
| chr1 | *RP5-865N13.2* | 3 | 119 | 7 | | | | | |
| chr14 | *KLC1* | 3 | 82 | 7 | KLC1 | kinesin light chain 1 | Cytoplasm | other | |
| chr3 | *PARP15* | 4 | 82 | 7 | PARP15 | poly(ADP-ribose) polymerase family member 15 | Other | other | |
| chr12 | *LAG3* | 4 | 156 | 7 | LAG3 | lymphocyte activating 3 | Plasma Membrane | transmembrane receptor | MK-4280, TSR-033, XmAb22841, LAG525, MGD013, GSK2831781, REGN3767, INCAGN02385, FS118, SYM022, BMS-986016 |
| chr17 | *HOXB7* | 4 | 161 | 7 | HOXB7 | homeobox B7 | Nucleus | transcription regulator | |
| chr16 | *ARMC5* | 4 | 144 | 7 | ARMC5 | armadillo repeat containing 5 | Cytoplasm | other | |
| chr17 | *MMD* | 4 | 99 | 7 | MMD | monocyte to macrophage differentiation associated | Plasma Membrane | kinase | |
| chr4 | *GALNT7* | 3 | 74 | 7 | GALNT7 | polypeptide N-acetylgalactosaminyltransferase | Cytoplasm | enzyme | |
| chr17 | *GEMIN4* | 3 | 190 | 7 | GEMIN4 | gem nuclear organelle associated protein 4 | Nucleus | other | |
| chr9 | *BARHL1* | 4 | 199 | 7 | BARHL1 | BarH like homeobox 1 | Nucleus | transcription regulator | |
| chr2 | *C2orf82* | 4 | 134 | 7 | SNORC | secondary ossification center associated regulator of chondrocyte maturation | Cytoplasm | other | |
| chr11 | *HINFP* | 3 | 143 | 7 | HINFP | histone H4 transcription factor | Nucleus | transcription regulator | |
| chr9 | *RP11-395P17.3* | 4 | 176 | 7 | | | | | |
| chr15 | *GOLGA8T* | 3 | 144 | 7 | GOLGA8T | golgin A8 family member T | Other | other | |
| chr8 | *DOK2* | 4 | 183 | 7 | DOK2 | docking protein 2 | Plasma Membrane | other | |
| chr16 | *PDZD9* | 4 | 62 | 7 | PDZD9 | PDZ domain containing 9 | Other | other | |
| chr7 | *AEBP1* | 4 | 163 | 7 | AEBP1 | AE binding protein 1 | Nucleus | peptidase | |
| chr3 | *CMTM6* | 3 | 68 | 7 | CMTM6 | CKLF like MARVEL transmembrane domain | Extracellular Space | cytokine | |
| chrX | *PRPS2* | 3 | 71 | 7 | PRPS2 | phosphoribosyl pyrophosphate synthetase 2 | Cytoplasm | kinase | |
| chrX | *GS1-594A7.3* | 3 | 76 | 7 | GS1-594A7.3 | uncharacterized LOC104798195 | Other | other | |
| chr17 | *GALK1* | 4 | 198 | 7 | GALK1 | galactokinase 1 | Cytoplasm | kinase | |
| chr2 | *AC007405.4* | 2 | 63 | 7 | | | | | |
| chr1 | *EDN2* | 3 | 193 | 7 | EDN2 | endothelin 2 | Extracellular Space | growth factor | |
| chr1 | *CRABP2* | 4 | 209 | 7 | CRABP2 | cellular retinoic acid binding | Cytoplasm | transporter | |
| chr7 | *CYP3A7* | 4 | 99 | 7 | CYP3A7 | cytochrome P450 family 3 subfamily A member 7 | Cytoplasm | enzyme | cobicistat, cobicistat/darunavir, cobicistat/elvitegravir/emtricitabine/tenofovir disoproxil, atazanavir/cobicistat |
| chr1 | *CENPF* | 4 | 127 | 7 | CENPF | centromere protein F | Nucleus | other | |
| chr17 | *DUS1L* | 4 | 153 | 7 | DUS1L | dihydrouridine synthase 1 like | Other | other | |
| chr19 | *RAD23A* | 3 | 173 | 7 | RAD23A | RAD23 homolog A, nucleotide excision repair protein | Nucleus | other | |
| chr20 | *CTSA* | 4 | 111 | 7 | CTSA | cathepsin A | Cytoplasm | peptidase | |
| chr5 | *UBLCP1* | 3 | 123 | 7 | UBLCP1 | ubiquitin like domain containing CTD phosphatase 1 | Nucleus | phosphatase | |
| chr2 | *SLC7A15P* | 3 | 101 | 7 | SLC7A15P | solute carrier family 7 member 15, pseudogene | Other | other | |
| chr9 | *GLT6D1* | 4 | 97 | 7 | GLT6D1 | glycosyltransferase 6 domain containing 1 | Other | other | |
| chr12 | *RP11-554D14.6* | 3 | 119 | 7 | | | | | |
| chr7 | *MOGAT3* | 4 | 123 | 7 | MOGAT3 | monoacylglycerol O- | Cytoplasm | enzyme | |

| | | | | | | | | | |
|---|---|---|---|---|---|---|---|---|---|
| chr1 | *ATP1B1* | 3 | 100 | 7 | ATP1B1 | ATPase Na+/K+ transporting subunit beta 1 | Plasma Membrane | transporter | |
| chr19 | *CTB-55O6.12* | 4 | 152 | 7 | | | | | |
| chr11 | *ACAD8* | 3 | 117 | 7 | ACAD8 | acyl-CoA dehydrogenase family member 8 | Cytoplasm | enzyme | |
| chr11 | *CRTAM* | 3 | 72 | 7 | CRTAM | cytotoxic and regulatory T cell molecule | Plasma Membrane | other | |
| chr7 | *AGR2* | 4 | 80 | 7 | AGR2 | anterior gradient 2, protein disulphide isomerase family | Extracellular Space | other | |
| chr3 | *KLHDC8B* | 4 | 200 | 7 | KLHDC8B | kelch domain containing 8B | Cytoplasm | other | |
| chr22 | *CECR3* | 3 | 97 | 7 | CECR3 | cat eye syndrome chromosome region, candidate 3 | Other | other | |
| chr12 | *KRT6B* | 4 | 200 | 7 | KRT6B | keratin 6B | Cytoplasm | other | |
| chr16 | *RP11-680G24.1* | 3 | 149 | 7 | | | | | |
| chr17 | *SLFN5* | 4 | 99 | 7 | SLFN5 | schlafen family member 5 | Nucleus | enzyme | |
| chr6 | *BCKDHB* | 3 | 63 | 7 | BCKDHB | branched chain keto acid dehydrogenase E1 subunit beta | Cytoplasm | enzyme | |
| chr15 | *LINC00929* | 3 | 86 | 7 | LINC00929 | long intergenic non-protein coding RNA 929 | Other | other | |
| chr17 | *PROCA1* | 3 | 151 | 7 | PROCA1 | protein interacting with cyclin A1 | Cytoplasm | other | |
| chr17 | *AC007952.4* | 3 | 114 | 7 | | | | | |
| chr1 | *AHCYL1* | 3 | 102 | 7 | AHCYL1 | adenosylhomocysteinase like 1 | Cytoplasm | enzyme | |
| chr11 | *LRRC56* | 4 | 172 | 7 | LRRC56 | leucine rich repeat containing 56 | Other | other | |
| chr13 | *SLITRK1* | 3 | 93 | 7 | SLITRK1 | SLIT and NTRK like family | Plasma Membrane | other | |
| chr11 | *ACP2* | 4 | 293 | 7 | ACP2 | acid phosphatase 2, lysosomal | Cytoplasm | phosphatase | |
| chr19 | *CD209* | 3 | 100 | 7 | CD209 | CD209 molecule | Plasma Membrane | other | |
| chr19 | *ZNF527* | 3 | 61 | 7 | ZNF527 | zinc finger protein 527 | Nucleus | other | |
| chr1 | *DIEXF* | 4 | 111 | 7 | UTP25 | UTP25 small subunit processor component | Nucleus | other | |
| chr2 | *COL5A2* | 3 | 59 | 7 | COL5A2 | collagen type V alpha 2 chain | Extracellular Space | other | collagenase clostridium |
| chr1 | *CCDC19* | 4 | 100 | 7 | CFAP45 | cilia and flagella associated | Nucleus | other | |
| chr9 | *RP11-203J24.9* | 4 | 113 | 7 | | | | | |
| chr19 | *ZNF585B* | 3 | 73 | 7 | ZNF585B | zinc finger protein 585B | Nucleus | other | |
| chr11 | *RP11-831A10.2* | 3 | 142 | 7 | | | | | |
| chr12 | *RP11-77I22.2* | 3 | 91 | 7 | | | | | |
| chr7 | *SLC13A1* | 4 | 64 | 7 | SLC13A1 | solute carrier family 13 member 1 | Plasma Membrane | transporter | |
| chr8 | *RP11-163N6.2* | 3 | 95 | 7 | | | | | |
| chr8 | *FAM87A* | 4 | 143 | 7 | FAM87A | family with sequence similarity 87 member A | Other | other | |
| chr9 | *ATP8B5P* | 4 | 74 | 7 | ATP8B5P | ATPase phospholipid transporting 8B5, pseudogene | Other | other | |
| chrX | *MIR1587* | 4 | 131 | 7 | MIR1587 | microRNA 1587 | Cytoplasm | microRNA | |
| chr18 | *RP11-128P17.3* | 3 | 70 | 7 | | | | | |
| chr11 | *MS4A4A* | 3 | 70 | 7 | MS4A4A | membrane spanning 4-domains | Cytoplasm | other | |
| chr5 | *C5orf47* | 3 | 169 | 7 | C5orf47 | chromosome 5 open reading frame 47 | Other | other | |
| chr17 | *LINC00482* | 4 | 134 | 7 | LINC00482 | long intergenic non-protein coding RNA 482 | Other | other | |
| chr5 | *RP11-332J15.1* | 4 | 227 | 7 | | | | | |
| chr16 | *RP11-343H19.1* | 4 | 80 | 7 | | | | | |
| chrX | *GAGE2C* | 3 | 188 | 7 | GE1 (includes oth | G antigen 1 | Other | other | |
| chr15 | *DNAJA4* | 3 | 155 | 7 | DNAJA4 | DnaJ heat shock protein family (Hsp40) member A4 | Nucleus | other | |
| chr17 | *RP11-387H17.4* | 4 | 82 | 7 | | | | | |
| chr6 | *SNRNP48* | 3 | 98 | 7 | SNRNP48 | small nuclear ribonucleoprotein U11/U12 subunit 48 | Nucleus | other | |
| chr3 | *LINC00501* | 2 | 64 | 7 | LINC00501 | long intergenic non-protein coding RNA 501 | Other | other | |
| chr2 | *GTF3C2* | 3 | 83 | 7 | GTF3C2 | general transcription factor IIIC subunit 2 | Nucleus | transcription regulator | |
| chr4 | *ADAM29* | 3 | 70 | 7 | ADAM29 | ADAM metallopeptidase domain | Plasma Membrane | peptidase | |
| chr19 | *MAP2K7* | 4 | 181 | 7 | MAP2K7 | mitogen-activated protein kinase kinase 7 | Cytoplasm | kinase | |
| chr5 | *GRM6* | 4 | 168 | 7 | GRM6 | glutamate metabotropic receptor | Plasma Membrane | G-protein coupled recepto | fasoracetam |
| chr19 | *U2AF2* | 4 | 158 | 7 | U2AF2 | U2 small nuclear RNA auxiliary factor 2 | Nucleus | other | |
| chr13 | *PDX1-AS1* | 3 | 87 | 7 | PLUT | PDX1 associated lncRNA, upregulator of transcription | Other | other | |
| chr15 | *PARP6* | 3 | 74 | 7 | PARP6 | poly(ADP-ribose) polymerase family member 6 | Other | other | |
| chr16 | *RP11-146F11.5* | 4 | 93 | 7 | | | | | |
| chr9 | *IDNK* | 3 | 75 | 7 | IDNK | IDNK gluconokinase | Cytoplasm | other | |
| chr9 | *RP11-197P3.4* | 3 | 76 | 7 | | | | | |
| chr3 | *RP11-71N10.1* | 3 | 78 | 7 | | | | | |
| chr3 | *UMPS* | 3 | 122 | 7 | UMPS | uridine monophosphate synthetase | Cytoplasm | enzyme | gimeracil/irinotecan/oxonic acid/tegafur, bevacizumab/gimeracil/irinotecan/oxonic acid/tegafur, gimeracil/oxonic acid/tegafur, pyrazofurin, cetuximab/gimeracil/irinotecan/oxonic acid/tegafur, oxonic acid |
| chr11 | *CD59* | 4 | 111 | 7 | CD59 | CD59 molecule (CD59 blood | Plasma Membrane | other | |
| chr1 | *RP4-781K5.4* | 4 | 186 | 7 | | | | | |

| | | | | | | | | | |
|---|---|---|---|---|---|---|---|---|---|
| chr10 | *AKR1E2* | 3 | 118 | 7 | AKR1E2 | aldo-keto reductase family 1 member E2 | Cytoplasm | enzyme | |
| chr12 | *NABP2* | 4 | 140 | 7 | NABP2 | nucleic acid binding protein 2 | Nucleus | other | |
| chr11 | *DUSP8* | 4 | 173 | 7 | DUSP8 | dual specificity phosphatase 8 | Nucleus | phosphatase | |
| chr1 | *ILF2* | 3 | 117 | 7 | ILF2 | interleukin enhancer binding | Nucleus | transcription regulator | |
| chr17 | *TAC4* | 3 | 100 | 7 | TAC4 | tachykinin precursor 4 | Extracellular Space | other | |
| chr12 | *KRT6A* | 3 | 201 | 7 | KRT6A | keratin 6A | Cytoplasm | other | |
| chr11 | *RPL27A* | 3 | 120 | 7 | RPL27A | ribosomal protein L27a | Cytoplasm | other | |
| chr9 | *RP11-542K23.7* | 4 | 104 | 7 | | | | | |
| chr9 | *LHX3* | 4 | 121 | 7 | LHX3 | LIM homeobox 3 | Nucleus | transcription regulator | |
| chr16 | *CASP16* | 4 | 176 | 7 | CASP16P | caspase 16, pseudogene | Other | other | |
| chr12 | *RP11-582E3.2* | 4 | 97 | 7 | | | | | |
| chr5 | *SNCB* | 4 | 166 | 7 | SNCB | synuclein beta | Cytoplasm | other | |
| chr15 | *DNM1P46* | 3 | 217 | 7 | DNM1P46 | dynamin 1 pseudogene 46 | Other | other | |
| chr4 | *KLHL2* | 3 | 58 | 7 | KLHL2 | kelch like family member 2 | Cytoplasm | other | |
| chr3 | *RPL39L* | 3 | 75 | 7 | RPL39L | ribosomal protein L39 like | Other | other | |
| chr9 | *CDK20* | 3 | 154 | 7 | CDK20 | cyclin dependent kinase 20 | Nucleus | kinase | |
| chr2 | *AC068490.2* | 3 | 60 | 7 | | | | | |
| chr1 | *ECM1* | 3 | 158 | 7 | ECM1 | extracellular matrix protein 1 | Extracellular Space | transporter | |
| chr17 | *AXIN2* | 4 | 122 | 7 | AXIN2 | axin 2 | Cytoplasm | other | |
| chr22 | *RNF185* | 3 | 65 | 7 | RNF185 | ring finger protein 185 | Cytoplasm | enzyme | |
| chr1 | *MYBPH* | 4 | 221 | 7 | MYBPH | myosin binding protein H | Cytoplasm | other | |
| chr6 | *HIST1H2AC* | 4 | 62 | 7 | H2AC6 | H2A clustered histone 6 | Nucleus | other | |
| chr19 | *LILRP2* | 3 | 151 | 7 | LILRP2 | leukocyte immunoglobulin-like receptor pseudogene 2 | Other | other | |
| chr12 | *PIANP* | 4 | 160 | 7 | PIANP | PILR alpha associated neural | Plasma Membrane | other | |
| chr17 | *CDK5RAP3* | 3 | 110 | 7 | CDK5RAP3 | CDK5 regulatory subunit associated protein 3 | Nucleus | other | |
| chr12 | *PCBP2* | 3 | 122 | 7 | PCBP2 | poly(rC) binding protein 2 | Nucleus | other | |
| chr1 | *LEMD1-AS1* | 4 | 96 | 7 | LEMD1-AS1 | LEMD1 antisense RNA 1 | Other | other | |
| chr7 | *VSTM2A* | 3 | 88 | 7 | VSTM2A | V-set and transmembrane domain containing 2A | Extracellular Space | other | |
| chr18 | *CXXC1* | 3 | 147 | 7 | CXXC1 | CXXC finger protein 1 | Nucleus | transcription regulator | |
| chr16 | *RP11-863P13.3* | 3 | 156 | 7 | | | | | |
| chr11 | *SAAL1* | 3 | 65 | 7 | SAAL1 | serum amyloid A like 1 | Other | other | |
| chr3 | *RP11-200A1.1* | 4 | 166 | 7 | | | | | |
| chr10 | *CUZD1* | 3 | 88 | 7 | CUZD1 | CUB and zona pellucida like domains 1 | Plasma Membrane | other | |
| chr16 | *MARVELD3* | 4 | 106 | 7 | MARVELD3 | MARVEL domain containing 3 | Plasma Membrane | other | |
| chr17 | *NEURL4* | 3 | 215 | 7 | NEURL4 | neuralized E3 ubiquitin protein ligase 4 | Other | other | |
| chr1 | *SH2D1B* | 4 | 105 | 7 | SH2D1B | SH2 domain containing 1B | Cytoplasm | other | |
| chr12 | *RP11-771K4.1* | 3 | 118 | 7 | | | | | |
| chr8 | *TGS1* | 2 | 53 | 7 | TGS1 | trimethylguanosine synthase 1 | Nucleus | enzyme | |
| chr2 | *EIF4E2* | 3 | 62 | 7 | EIF4E2 | eukaryotic translation initiation factor 4E family member 2 | Cytoplasm | translation regulator | |
| chr1 | *ACAP3* | 4 | 104 | 7 | ACAP3 | ArfGAP with coiled-coil, ankyrin repeat and PH domains 3 | Nucleus | transcription regulator | |
| chr10 | *SYNPO2L* | 3 | 197 | 7 | SYNPO2L | synaptopodin 2 like | Cytoplasm | other | |
| chr17 | *3C1D3P1-DHX40I* | 3 | 130 | 7 | 3C1D3P1-DHX40I | TBC1D3P1-DHX40P1 readthrough, transcribed | Other | other | |
| chrX | *TMSB15B* | 3 | 75 | 7 | TMSB15B | thymosin beta 15B | Other | other | |
| chr16 | *CTD-2135D7.3* | 2 | 57 | 7 | | | | | |
| chrX | *GAGE12F* | 3 | 186 | 7 | GE2D (includes otl | G antigen 2D | Other | other | |
| chrX | *BRWD3* | 3 | 63 | 7 | BRWD3 | bromodomain and WD repeat domain containing 3 | Other | other | |
| chr8 | *LY96* | 3 | 69 | 7 | LY96 | lymphocyte antigen 96 | Plasma Membrane | transmembrane recepto | |
| chr1 | *RP11-15I11.3* | 4 | 89 | 7 | | | | | |
| chr1 | *TPR* | 3 | 72 | 7 | TPR | translocated promoter region, nuclear basket protein | Nucleus | other | |
| chr5 | *GABRG2* | 3 | 64 | 7 | GABRG2 | gamma-aminobutyric acid type A receptor gamma2 subunit | Plasma Membrane | ion channel | [illegible] nitrazepam, chlordiazepoxide, sevoflurane, etomidate, alprazolam, pagoclone, zolpidem, phenobarbital, diazepam, isoflurane, halothane, pentobarbital, desflurane, adinazolam, SEP 174559, pregnenolone, zaleplon, tracazolate, enflurane, ezogabine, muscimol, flurazepam, flumazenil, metyrapone/oxazepam, isoniazid, [illegible] |
| chr10 | *LINC00202-1* | 3 | 120 | 7 | FAM238C | family with sequence similarity 238 member C | Other | other | |
| chr7 | *RP11-1217F2.15* | 4 | 101 | 7 | | | | | |
| chr11 | *LRRC32* | 4 | 239 | 7 | LRRC32 | leucine rich repeat containing 32 | Plasma Membrane | other | |

| | | | | | | | | | |
|---|---|---|---|---|---|---|---|---|---|
| chr5 | *CDKN2AIPNL* | 4 | 93 | 7 | CDKN2AIPNL | CDKN2A interacting protein N-terminal like | Other | other | |
| chr17 | *AC005863.1* | 3 | 84 | 7 | | | | | |
| chr1 | *C8A* | 4 | 86 | 7 | C8A | complement C8 alpha chain | Extracellular Space | other | |
| chr8 | *FAM83H-AS1* | 4 | 199 | 7 | FAM83H-AS1 | FAM83H antisense RNA 1 (head to head) | Other | other | |
| chr19 | *PSG9* | 4 | 99 | 7 | PSG9 | pregnancy specific beta-1-glycoprotein 9 | Extracellular Space | other | |
| chr19 | *CTC-360P9.5* | 3 | 66 | 7 | | | | | |
| chr17 | *MAFG-AS1* | 3 | 151 | 7 | MAFG-DT | MAFG divergent transcript | Other | other | |
| chr8 | *AGPAT6* | 3 | 79 | 7 | GPAT4 | glycerol-3-phosphate acyltransferase 4 | Cytoplasm | enzyme | |
| chr16 | *RP11-165E7.1* | 3 | 120 | 7 | | | | | |
| chr15 | *RP11-509A17.3* | 4 | 514 | 7 | | | | | |
| chr3 | *TMEM30C* | 3 | 68 | 7 | TMEM30CP | transmembrane protein 30C, pseudogene | Other | other | |
| chr5 | *LINC01021* | 3 | 202 | 7 | PURPL | p53 upregulated regulator of p53 levels | Other | other | |
| chrX | *SLC16A2* | 3 | 93 | 7 | SLC16A2 | solute carrier family 16 member 2 | Plasma Membrane | transporter | |
| chr3 | *NIT2* | 4 | 84 | 7 | NIT2 | nitrilase family member 2 | Cytoplasm | enzyme | |
| chr15 | *CTD-2116N17.1* | 3 | 82 | 7 | | | | | |
| chr19 | *POLRMT* | 4 | 114 | 7 | POLRMT | RNA polymerase mitochondrial | Cytoplasm | enzyme | |
| chr8 | *ZNF696* | 4 | 115 | 7 | ZNF696 | zinc finger protein 696 | Nucleus | transcription regulator | |
| chr11 | *NCR3LG1* | 4 | 112 | 7 | NCR3LG1 | natural killer cell cytotoxicity receptor 3 ligand 1 | Plasma Membrane | other | |
| chr17 | *SLFN12* | 3 | 88 | 7 | SLFN12 | schlafen family member 12 | Other | enzyme | |
| chr5 | *CTD-2091N23.1* | 2 | 62 | 7 | | | | | |
| chr1 | *DVL1* | 4 | 114 | 7 | DVL1 | dishevelled segment polarity | Cytoplasm | other | |
| chr5 | *FGF10* | 4 | 187 | 7 | FGF10 | fibroblast growth factor 10 | Extracellular Space | growth factor | |
| chr1 | *LDLRAD2* | 3 | 128 | 7 | LDLRAD2 | low density lipoprotein receptor class A domain containing 2 | Other | transporter | |
| chr2 | *AC109829.1* | 3 | 62 | 7 | | | | | |
| chr10 | *FXYD4* | 3 | 155 | 7 | FXYD4 | FXYD domain containing ion transport regulator 4 | Plasma Membrane | ion channel | |
| chr1 | *RP11-338C15.3* | 3 | 130 | 7 | | | | | |
| chr19 | *ZNF628* | 4 | 148 | 7 | ZNF628 | zinc finger protein 628 | Nucleus | transcription regulator | |
| chr9 | *SET* | 3 | 129 | 7 | SET | SET nuclear proto-oncogene | Nucleus | phosphatase | |
| chr2 | *TEX41* | 3 | 82 | 7 | TEX41 | testis expressed 41 | Other | other | |
| chr12 | *LINC00940* | 3 | 114 | 7 | LINC00940 | long intergenic non-protein coding RNA 940 | Other | other | |
| chr4 | *STIM2* | 3 | 72 | 7 | STIM2 | stromal interaction molecule 2 | Plasma Membrane | transporter | |
| chr4 | *EDNRA* | 3 | 60 | 7 | EDNRA | endothelin receptor type A | Plasma Membrane | transmembrane recepto | BSF 302146, sitaxsentan, ambrisentan/tadalafil, avosentan, bosentan, fandosentan, macitentan, SB 234551, zibotentan, clazosentan, atrasentan, YM598, |
| chr8 | *LINC00529* | 3 | 74 | 7 | LINC00529 | long intergenic non-protein coding RNA 529 | Other | other | |
| chr1 | *H3F3A* | 2 | 106 | 7 | H3-3A/H3-3B | H3.3 histone A | Nucleus | other | |
| chr5 | *TMEM173* | 3 | 111 | 7 | TMEM173 | transmembrane protein 173 | Cytoplasm | other | MIW815, MK-1454 |
| chr8 | *RP11-128L5.1* | 3 | 135 | 7 | | | | | |
| chr20 | *UCKL1* | 4 | 306 | 7 | UCKL1 | uridine-cytidine kinase 1 like 1 | Cytoplasm | kinase | |
| chr10 | *RP11-298H24.1* | 3 | 72 | 7 | | | | | |
| chr1 | *RP11-763B22.9* | 3 | 236 | 7 | | | | | |
| chr22 | *WI2-85898F10.1* | 4 | 178 | 7 | | | | | |
| chr21 | *NRIP1* | 3 | 60 | 7 | NRIP1 | nuclear receptor interacting | Nucleus | transcription regulator | |
| chr21 | *C21orf33* | 4 | 182 | 7 | GATD3A/GATD3E | glutamine amidotransferase like class 1 domain containing 3A | Cytoplasm | other | |
| chr11 | *CNGA4* | 3 | 189 | 7 | CNGA4 | cyclic nucleotide gated channel subunit alpha 4 | Plasma Membrane | ion channel | |
| chr9 | *ANP32B* | 3 | 65 | 7 | ANP32B | acidic nuclear phosphoprotein 32 family member B | Nucleus | other | |
| chr9 | *TOR1B* | 3 | 152 | 7 | TOR1B | torsin family 1 member B | Cytoplasm | enzyme | |
| chr17 | *TSR1* | 3 | 79 | 7 | TSR1 | TSR1 ribosome maturation factor | Nucleus | other | |
| chr12 | *NUP37* | 4 | 97 | 7 | NUP37 | nucleoporin 37 | Nucleus | transporter | |
| chr1 | *RP1-37C10.3* | 4 | 166 | 7 | | | | | |
| chrX | *ZMYM3* | 3 | 100 | 7 | ZMYM3 | zinc finger MYM-type containing | Nucleus | other | |
| chr11 | *RP11-168K9.1* | 3 | 62 | 7 | | | | | |
| chr19 | *SBSN* | 3 | 135 | 7 | SBSN | suprabasin | Cytoplasm | other | |
| chr9 | *EXOSC2* | 4 | 111 | 7 | EXOSC2 | exosome component 2 | Nucleus | enzyme | |
| chrX | *GAGE12E* | 3 | 175 | 7 | GE1 (includes oth | G antigen 1 | Other | other | |
| chr10 | *RP11-809M12.1* | 3 | 104 | 7 | | | | | |
| chr9 | *MAMDC2* | 4 | 74 | 7 | MAMDC2 | MAM domain containing 2 | Extracellular Space | other | |
| chr3 | *THOC7* | 3 | 65 | 7 | THOC7 | THO complex 7 | Nucleus | other | |
| chr3 | *C3orf27* | 3 | 179 | 7 | LINC01565 | long intergenic non-protein coding RNA 1565 | Other | other | |
| chr16 | *CTU2* | 4 | 142 | 7 | CTU2 | cytosolic thiouridylase subunit 2 | Cytoplasm | other | |
| chr5 | *PJA2* | 3 | 86 | 7 | PJA2 | praja ring finger ubiquitin ligase 2 | Cytoplasm | enzyme | |
| chr11 | *CYB561A3* | 3 | 134 | 7 | CYB561A3 | cytochrome b561 family member | Cytoplasm | other | |
| chr13 | *SMAD9* | 3 | 74 | 7 | SMAD9 | SMAD family member 9 | Nucleus | transcription regulator | |
| chr7 | *CAV1* | 3 | 147 | 7 | CAV1 | caveolin 1 | Plasma Membrane | transmembrane recepto | |
| chr19 | *CTB-147C22.9* | 4 | 73 | 7 | | | | | |
| chr2 | *AC007464.1* | 2 | 72 | 7 | | | | | |

| | | | | | | | | | |
|---|---|---|---|---|---|---|---|---|---|
| chr16 | *GDPD3* | 3 | 114 | 7 | GDPD3 | glycerophosphodiester phosphodiesterase domain containing 3 | Cytoplasm | enzyme | |
| chr3 | *RP11-167H9.5* | 4 | 102 | 7 | | | | | |
| chr17 | *TUBG2* | 3 | 143 | 7 | TUBG2 | tubulin gamma 2 | Cytoplasm | other | docetaxel/gemcitabine/vincristine, cyclophosphamide/epirubicin/5-fluorouracil/paclitaxel, bevacizumab/paclitaxel/topotecan, colchicine/probenecid, bevacizumab/docetaxel/prednisone/thalidomide/zoledronic acid, gemcitabine/paclitaxel, L-asparaginase/daunorubicin/dexamethasone/vincristine, prednisone/vincristine, docetaxel/5-fluorouracil, vinflunine, L-[illegible] |
| chr2 | *ANKRD30BL* | 3 | 146 | 7 | ANKRD30BL | ankyrin repeat domain 30B like | Other | other | |
| chr4 | *TMEM150C* | 3 | 53 | 7 | TMEM150C | transmembrane protein 150C | Plasma Membrane | ion channel | |
| chr8 | *RP11-252C15.1* | 3 | 93 | 7 | | | | | |
| chr12 | *ALKBH2* | 4 | 95 | 7 | ALKBH2 | alkB homolog 2, alpha-ketoglutarate dependent | Nucleus | enzyme | |
| chr1 | *GUK1* | 4 | 229 | 7 | GUK1 | guanylate kinase 1 | Cytoplasm | kinase | |
| chr3 | *LYZL4* | 3 | 99 | 7 | LYZL4 | lysozyme like 4 | Extracellular Space | enzyme | |
| chr19 | *NDUFB7* | 3 | 118 | 7 | NDUFB7 | NADH:ubiquinone oxidoreductase subunit B7 | Cytoplasm | enzyme | |
| chr1 | *HIST2H4B* | 3 | 134 | 7 | H4C15 | H4 clustered histone 15 | Nucleus | other | |
| chr17 | *RP11-385D13.1* | 3 | 99 | 7 | | | | | |
| chr18 | *FAM210A* | 3 | 82 | 7 | FAM210A | family with sequence similarity 210 member A | Cytoplasm | other | |
| chr11 | *RP11-646J21.2* | 3 | 144 | 7 | | | | | |
| chr11 | *RP11-21L23.4* | 4 | 169 | 7 | | | | | |
| chr1 | *PRRX1* | 4 | 77 | 7 | PRRX1 | paired related homeobox 1 | Nucleus | transcription regulator | |
| chr19 | *IZUMO2* | 4 | 93 | 7 | IZUMO2 | IZUMO family member 2 | Other | other | |
| chr15 | *UBL7-AS1* | 3 | 65 | 7 | UBL7-AS1 | UBL7 antisense RNA 1 (head to head) | Other | other | |
| chr8 | *RP11-13N12.2* | 3 | 115 | 7 | | | | | |
| chr11 | *CREBZF* | 3 | 172 | 7 | CREBZF | CREB/ATF bZIP transcription | Nucleus | transcription regulator | |
| chr17 | *CTD-2246P4.1* | 3 | 71 | 7 | | | | | |
| chr19 | *SIGLEC16* | 3 | 129 | 7 | SIGLEC16 | sialic acid binding Ig like lectin 16 (gene/pseudogene) | Plasma Membrane | other | |
| chr19 | *ZNF526* | 3 | 87 | 7 | ZNF526 | zinc finger protein 526 | Other | other | |
| chr19 | *ZNF175* | 3 | 67 | 7 | ZNF175 | zinc finger protein 175 | Nucleus | transcription regulator | |
| chr14 | *C14orf182* | 3 | 64 | 7 | LINC01588 | long intergenic non-protein coding RNA 1588 | Other | other | |
| chr19 | *CTD-2583A14.9* | 3 | 71 | 7 | | | | | |
| chrX | *GPC4* | 3 | 60 | 7 | GPC4 | glypican 4 | Plasma Membrane | transmembrane receptor | |
| chr11 | *RP11-98J9.1* | 4 | 75 | 7 | | | | | |
| chr14 | *ERO1L* | 3 | 61 | 7 | ERO1A | endoplasmic reticulum oxidoreductase 1 alpha | Cytoplasm | enzyme | |
| chrX | *RP4-562J12.2* | 3 | 65 | 7 | | | | | |
| chr6 | *ZNF292* | 3 | 75 | 7 | ZNF292 | zinc finger protein 292 | Nucleus | transcription regulator | |
| chr14 | *TMED8* | 2 | 61 | 7 | TMED8 | transmembrane p24 trafficking protein family member 8 | Other | other | |
| chrX | *SASH3* | 3 | 115 | 7 | SASH3 | SAM and SH3 domain containing | Cytoplasm | other | |
| chr16 | *CCL22* | 3 | 116 | 7 | CCL22 | C-C motif chemokine ligand 22 | Extracellular Space | cytokine | |
| chr13 | *MIPEP* | 3 | 95 | 7 | MIPEP | mitochondrial intermediate | Cytoplasm | peptidase | |
| chrX | *ARSE* | 3 | 67 | 7 | ARSL | arylsulfatase L | Cytoplasm | enzyme | |
| chr12 | *TMED2* | 3 | 75 | 7 | TMED2 | transmembrane p24 trafficking protein 2 | Cytoplasm | transporter | |
| chr14 | *RP3-414A15.10* | 3 | 63 | 7 | | | | | |
| chr9 | *TMEM8B* | 3 | 124 | 7 | TMEM8B | transmembrane protein 8B | Plasma Membrane | other | |
| chr6 | *TFAP2A* | 3 | 137 | 7 | TFAP2A | transcription factor AP-2 alpha | Nucleus | transcription regulator | |
| chr4 | *RAB28* | 3 | 81 | 7 | RAB28 | RAB28, member RAS oncogene family | Plasma Membrane | enzyme | |
| chr7 | *RNF216P1* | 3 | 87 | 7 | RNF216P1 | ring finger protein 216 | Other | other | |
| chr17 | *LINC00974* | 4 | 94 | 7 | LINC00974 | long intergenic non-protein coding RNA 974 | Other | other | |
| chr16 | *DCTN5* | 3 | 64 | 7 | DCTN5 | dynactin subunit 5 | Cytoplasm | other | |
| chr7 | *SGCE* | 3 | 113 | 7 | SGCE | sarcoglycan epsilon | Plasma Membrane | other | |
| chr4 | *KCTD8* | 3 | 98 | 7 | KCTD8 | potassium channel tetramerization domain containing | Other | other | |
| chr3 | *ANKUB1* | 3 | 92 | 7 | ANKUB1 | ankyrin repeat and ubiquitin domain containing 1 | Other | other | |
| chr13 | *N4BP2L2* | 3 | 68 | 7 | N4BP2L2 | NEDD4 binding protein 2 like 2 | Nucleus | transcription regulator | |
| chr19 | *UPK1A* | 3 | 133 | 7 | UPK1A | uroplakin 1A | Plasma Membrane | other | |
| chr17 | *PLEKHH3* | 4 | 250 | 7 | PLEKHH3 | pleckstrin homology, MyTH4 and FERM domain containing H3 | Extracellular Space | other | |

| | | | | | | | | | |
|---|---|---|---|---|---|---|---|---|---|
| chr11 | *RP11-347H15.4* | 3 | 90 | 7 | | | | | |
| chr17 | *AC005863.2* | 3 | 83 | 7 | | | | | |
| chr16 | *COX4I1* | 3 | 102 | 7 | COX4I1 | cytochrome c oxidase subunit 4I1 | Cytoplasm | enzyme | |
| chr5 | *SAP30L-AS1* | 3 | 85 | 7 | SAP30L-AS1 | SAP30L antisense RNA 1 (head to head) | Other | other | |
| chr5 | *ZNF879* | 4 | 91 | 7 | ZNF879 | zinc finger protein 879 | Other | other | |
| chr15 | *RP11-566K19.9* | 3 | 191 | 7 | | | | | |
| chr6 | *HACE1* | 3 | 86 | 7 | HACE1 | HECT domain and ankyrin repeat containing E3 ubiquitin protein ligase 1 | Cytoplasm | enzyme | |
| chr11 | *AP003419.11* | 3 | 114 | 7 | | | | | |
| chr15 | *GOLGA6L3* | 3 | 153 | 7 | GA6L9 (includes o | golgin A6 family like 9 | Other | other | |
| chr16 | *SULT1A2* | 3 | 118 | 7 | SULT1A2 | sulfotransferase family 1A | Cytoplasm | enzyme | |
| chr16 | *RP11-343C2.9* | 3 | 107 | 7 | | | | | |
| chr16 | *ZNF598* | 4 | 138 | 7 | ZNF598 | zinc finger protein 598 | Extracellular Space | enzyme | |
| chr22 | *MAPK8IP2* | 3 | 146 | 7 | MAPK8IP2 | mitogen-activated protein kinase 8 interacting protein 2 | Cytoplasm | other | |
| chr10 | *RP11-508N22.8* | 4 | 181 | 7 | | | | | |
| chr5 | *UTP15* | 2 | 59 | 7 | UTP15 | UTP15 small subunit processome component | Nucleus | other | |
| chr20 | *RP4-813D12.3* | 4 | 182 | 7 | | | | | |
| chr8 | *DPYS* | 3 | 77 | 7 | DPYS | dihydropyrimidinase | Cytoplasm | enzyme | |
| chr7 | *AC073071.1* | 3 | 70 | 7 | | | | | |
| chr12 | *GLIPR1L2* | 3 | 77 | 7 | GLIPR1L2 | GLIPR1 like 2 | Extracellular Space | other | |
| chr17 | *NXPH3* | 3 | 120 | 7 | NXPH3 | neurexophilin 3 | Extracellular Space | other | |
| chr3 | *TMEM43* | 3 | 110 | 7 | TMEM43 | transmembrane protein 43 | Nucleus | other | |
| chr2 | *GPR1* | 3 | 61 | 7 | GPR1 | G protein-coupled receptor 1 | Plasma Membrane | G-protein coupled recepto | |
| chr17 | *RP11-416I2.1* | 3 | 164 | 7 | | | | | |
| chrX | *FAM58A* | 3 | 126 | 7 | CCNQ | cyclin Q | Other | other | |
| chr10 | *RP11-179K3.2* | 3 | 111 | 7 | | | | | |
| chr11 | *TMEM151A* | 4 | 299 | 7 | TMEM151A | transmembrane protein 151A | Other | other | |
| chr1 | *FAIM3* | 4 | 166 | 7 | FCMR | Fc fragment of IgM receptor | Plasma Membrane | other | |
| chr7 | *ZNF853* | 4 | 201 | 7 | ZNF853 | zinc finger protein 853 | Other | other | |
| chr14 | *TEX22* | 3 | 112 | 7 | TEX22 | testis expressed 22 | Cytoplasm | other | |
| chr1 | *PRAMEF5* | 3 | 75 | 7 | MEF4 (includes ot | PRAME family member 4 | Other | other | |
| chr1 | *C1orf173* | 4 | 89 | 7 | ERICH3 | glutamate rich 3 | Other | other | |
| chr7 | *AC004014.3* | 3 | 93 | 7 | | | | | |
| chr1 | *THAP3* | 3 | 201 | 7 | THAP3 | THAP domain containing 3 | Nucleus | transcription regulator | |
| chr12 | *CAPRIN2* | 2 | 75 | 7 | CAPRIN2 | caprin family member 2 | Cytoplasm | other | |
| chr10 | *FAM25B* | 3 | 124 | 7 | FAM25BP | protein FAM25 | Other | other | |
| chr1 | *ATP13A2* | 3 | 138 | 7 | ATP13A2 | ATPase cation transporting 13A2 | Cytoplasm | transporter | |
| chr2 | *IL36B* | 4 | 139 | 7 | IL36B | interleukin 36 beta | Extracellular Space | cytokine | |
| chr4 | *HOPX* | 3 | 78 | 7 | HOPX | HOP homeobox | Nucleus | transcription regulator | |
| chr18 | *ZNF519* | 3 | 88 | 7 | ZNF519 | zinc finger protein 519 | Nucleus | transcription regulator | |
| chr18 | *PIK3C3* | 3 | 55 | 7 | PIK3C3 | phosphatidylinositol 3-kinase catalytic subunit type 3 | Cytoplasm | kinase | |
| chr3 | *COMMD2* | 3 | 111 | 7 | COMMD2 | COMM domain containing 2 | Other | other | |
| chr18 | *CDH20* | 3 | 77 | 7 | CDH20 | cadherin 20 | Plasma Membrane | other | |
| chr2 | *TMEM150A* | 3 | 151 | 7 | TMEM150A | transmembrane protein 150A | Plasma Membrane | other | |
| chr17 | *ENPP7* | 3 | 180 | 7 | ENPP7 | ectonucleotide pyrophosphatase/phosphodiesterase 7 | Plasma Membrane | enzyme | |
| chr8 | *SDC2* | 3 | 58 | 7 | SDC2 | syndecan 2 | Plasma Membrane | other | |
| chr9 | *TMEM261* | 3 | 74 | 7 | DMAC1 | distal membrane arm assembly complex 1 | Cytoplasm | other | |
| chr19 | *CTC-400I9.2* | 3 | 79 | 7 | | | | | |
| chr5 | *LSM11* | 3 | 81 | 7 | LSM11 | LSM11, U7 small nuclear RNA associated | Nucleus | other | |
| chr8 | *RP11-946L20.2* | 3 | 72 | 7 | | | | | |
| chr20 | *OGFR* | 4 | 242 | 7 | OGFR | opioid growth factor receptor | Plasma Membrane | other | enkephalin, |
| chr14 | *WDR89* | 3 | 85 | 7 | WDR89 | WD repeat domain 89 | Other | other | |
| chr1 | *CALR4P* | 3 | 66 | 7 | CALR4P | calreticulin 4, pseudogene | Other | other | |
| chr7 | *TNRC18P3* | 3 | 157 | 7 | TNRC18P3 | trinucleotide repeat containing 18 pseudogene 3 | Other | other | |
| chr1 | *C1QA* | 3 | 137 | 7 | C1QA | complement C1q A chain | Extracellular Space | other | |
| chr11 | *MRGPRX3* | 3 | 108 | 7 | MRGPRX3 | MAS related GPR family member | Plasma Membrane | G-protein coupled recepto | |
| chr21 | *AF064860.5* | 3 | 68 | 7 | | | | | |
| chr5 | *RP11-381K20.2* | 3 | 65 | 7 | | | | | |
| chrX | *LONRF3* | 3 | 63 | 7 | LONRF3 | LON peptidase N-terminal domain and ring finger 3 | Other | other | |
| chr9 | *EGFL7* | 4 | 199 | 7 | EGFL7 | EGF like domain multiple 7 | Extracellular Space | other | parsatuzumab |
| chr1 | *RP11-92G12.3* | 3 | 93 | 7 | | | | | |
| chr6 | *EPHA7* | 3 | 55 | 7 | EPHA7 | EPH receptor A7 | Plasma Membrane | kinase | |
| chr17 | *RP11-13K12.2* | 3 | 111 | 7 | | | | | |
| chr6 | *PAQR8* | 4 | 93 | 7 | PAQR8 | progestin and adipoQ receptor family member 8 | Plasma Membrane | other | |
| chr7 | *AC138783.11* | 3 | 100 | 7 | | | | | |
| chr22 | *RP5-1119A7.14* | 3 | 99 | 7 | | | | | |
| chr5 | *AC138035.1* | 3 | 185 | 7 | | | | | |
| chr3 | *PRSS50* | 3 | 150 | 7 | PRSS50 | serine protease 50 | Cytoplasm | peptidase | |
| chr15 | *HAUS2* | 3 | 65 | 7 | HAUS2 | HAUS augmin like complex | Cytoplasm | other | |
| chr12 | *STAC3* | 3 | 106 | 7 | STAC3 | SH3 and cysteine rich domain 3 | Extracellular Space | other | |
| chr1 | *HSD17B7* | 3 | 110 | 7 | HSD17B7 | hydroxysteroid 17-beta dehydrogenase 7 | Cytoplasm | enzyme | |
| chr1 | *CD2* | 3 | 103 | 7 | CD2 | CD2 molecule | Plasma Membrane | transmembrane receptor | alefacept, siplizumab |
| chr1 | *RP4-781K5.6* | 3 | 165 | 7 | | | | | |
| chr1 | *CDC42SE1* | 3 | 150 | 7 | CDC42SE1 | CDC42 small effector 1 | Plasma Membrane | other | |
| chr6 | *EEF1E1-BLOC1S* | 3 | 64 | 7 | EEF1E1-BLOC1S | EEF1E1-BLOC1S5 readthrough (NMD candidate) | Other | other | |

| | | | | | | | | | |
|---|---|---|---|---|---|---|---|---|---|
| chr2 | *AC013472.4* | 3 | 88 | 7 | | | | | |
| chr11 | *MEN1* | 3 | 174 | 7 | MEN1 | menin 1 | Nucleus | transcription regulator | SNDX-5613 |
| chr2 | *AC073479.1* | 2 | 68 | 7 | | | | | |
| chr6 | *PEX7* | 3 | 89 | 7 | PEX7 | peroxisomal biogenesis factor 7 | Cytoplasm | transporter | |
| chr6 | *MICAL1* | 3 | 110 | 7 | MICAL1 | microtubule associated monooxygenase, calponin and LIM domain containing 1 | Cytoplasm | enzyme | |
| chr2 | *KIDINS220* | 3 | 54 | 7 | KIDINS220 | kinase D interacting substrate | Nucleus | transcription regulator | |
| chr1 | *PINK1* | 3 | 134 | 7 | PINK1 | PTEN induced kinase 1 | Cytoplasm | kinase | |
| chr16 | *ZNF75A* | 3 | 64 | 7 | ZNF75A | zinc finger protein 75a | Nucleus | transcription regulator | |
| chr11 | *RP11-890B15.2* | 3 | 129 | 7 | | | | | |
| chr19 | *C19orf53* | 3 | 124 | 7 | C19orf53 | chromosome 19 open reading frame 53 | Nucleus | other | |
| chr5 | *ZNF354A* | 3 | 128 | 7 | ZNF354A | zinc finger protein 354A | Nucleus | transcription regulator | |
| chr2 | *AC104134.2* | 4 | 85 | 7 | | | | | |
| chr17 | *NT5C3B* | 3 | 113 | 7 | NT5C3B | 5'-nucleotidase, cytosolic IIIB | Cytoplasm | phosphatase | |
| chr16 | *NUDT7* | 3 | 64 | 7 | NUDT7 | nudix hydrolase 7 | Cytoplasm | enzyme | |
| chr16 | *FAM86A* | 4 | 280 | 7 | EEF2KMT | eukaryotic elongation factor 2 lysine methyltransferase | Cytoplasm | enzyme | |
| chr10 | *LYZL2* | 3 | 84 | 7 | LYZL2 | lysozyme like 2 | Other | other | |
| chr4 | *UGDH* | 3 | 60 | 7 | UGDH | UDP-glucose 6-dehydrogenase | Nucleus | enzyme | |
| chr15 | *LINC00277* | 3 | 104 | 7 | EWSAT1 | Ewing sarcoma associated transcript 1 | Other | other | |
| chr15 | *ZWILCH* | 3 | 72 | 7 | ZWILCH | zwilch kinetochore protein | Nucleus | other | |
| chr5 | *CLINT1* | 3 | 91 | 7 | CLINT1 | clathrin interactor 1 | Cytoplasm | other | |
| chr18 | *RP11-146N18.1* | 3 | 55 | 7 | | | | | |
| chr3 | *ZNF620* | 4 | 84 | 7 | ZNF620 | zinc finger protein 620 | Nucleus | transcription regulator | |
| chr8 | *RP11-38H17.1* | 4 | 61 | 7 | | | | | |
| chr15 | *AEN* | 3 | 206 | 7 | AEN | apoptosis enhancing nuclease | Nucleus | enzyme | |
| chr19 | *HCN2* | 4 | 184 | 7 | HCN2 | hyperpolarization activated cyclic nucleotide gated potassium and sodium channel 2 | Plasma Membrane | ion channel | |
| chr1 | *RFX5* | 3 | 135 | 7 | RFX5 | regulatory factor X5 | Nucleus | transcription regulator | |
| chr1 | *RSG1* | 3 | 157 | 7 | CPLANE2 | ciliogenesis and planar polarity effector 2 | Cytoplasm | other | |
| chr9 | *NPDC1* | 4 | 248 | 7 | NPDC1 | neural proliferation, differentiation and control 1 | Extracellular Space | other | |
| chrX | *RP4-769N13.6* | 3 | 89 | 7 | | | | | |
| chr18 | *NPIPB1P* | 3 | 74 | 7 | NPIPB1P | nuclear pore complex interacting protein family member B1, pseudogene | Other | other | |
| chr4 | *RP11-242J7.1* | 3 | 62 | 7 | | | | | |
| chr14 | *RABGGTA* | 3 | 112 | 7 | RABGGTA | Rab geranylgeranyltransferase subunit alpha | Extracellular Space | enzyme | |
| chr19 | *ZNF805* | 3 | 84 | 7 | ZNF805 | zinc finger protein 805 | Other | other | |
| chr15 | *OAZ2* | 3 | 83 | 7 | OAZ2 | ornithine decarboxylase antizyme | Cytoplasm | other | |
| chr8 | *FAM90A21P* | 3 | 227 | 7 | FAM90A21P | family with sequence similarity 90 member A21, pseudogene | Other | other | |
| chr2 | *CNOT11* | 3 | 94 | 7 | CNOT11 | CCR4-NOT transcription complex subunit 11 | Cytoplasm | other | |
| chr11 | *CCDC90B* | 2 | 81 | 7 | CCDC90B | coiled-coil domain containing 90B | Cytoplasm | other | |
| chr2 | *AC073254.1* | 3 | 75 | 7 | | | | | |
| chr11 | *RP11-159H22.2* | 3 | 146 | 7 | | | | | |
| chr22 | *RP1-85F18.5* | 3 | 56 | 7 | | | | | |
| chr13 | *TPP2* | 4 | 62 | 7 | TPP2 | tripeptidyl peptidase 2 | Cytoplasm | peptidase | |
| chr12 | *RP11-686F15.2* | 3 | 151 | 7 | | | | | |
| chr4 | *RP11-123O22.1* | 3 | 77 | 7 | | | | | |
| chr14 | *NOVA1-AS1* | 3 | 81 | 7 | LINC02588 | long intergenic non-protein coding RNA 2588 | Other | other | |
| chr19 | *CD33* | 4 | 73 | 7 | CD33 | CD33 molecule | Plasma Membrane | other | gemtuzumab ozogamicin, BI 836858, anti-CD33 monoclonal antibody, cytarabine/daunorubicin/etoposide/gemtuzumab ozogamicin, [225Ac]-FPI-1434, arsenic trioxide/gemtuzumab ozogamicin/tretinoin, cytarabine/gemtuzumab ozogamicin, GTB-3550, cytarabine/daunorubicin/gemtuzumab ozogamicin, actinium Ac 225 lintuzumab, vadastuximab talirine, cytarabine/daunorubicin/[illegible] |
| chr17 | *TBC1D27* | 3 | 135 | 7 | TBC1D27P | TBC1 domain family member 27, pseudogene | Other | other | |
| chr15 | *GREM1* | 3 | 72 | 7 | GREM1 | gremlin 1, DAN family BMP antagonist | Extracellular Space | other | |
| chr1 | *ARV1* | 3 | 83 | 7 | ARV1 | ARV1 homolog, fatty acid homeostasis modulator | Cytoplasm | transporter | |
| chr3 | *NICN1* | 3 | 112 | 7 | NICN1 | nicolin 1 | Nucleus | other | |
| chr15 | *FBXO22* | 4 | 90 | 7 | FBXO22 | F-box protein 22 | Cytoplasm | enzyme | |

| | | | | | | | | | |
|---|---|---|---|---|---|---|---|---|---|
| chr20 | *SNHG17* | 4 | 135 | 7 | SNHG17 | small nucleolar RNA host gene | Other | other | |
| chr12 | *AC156455.1* | 3 | 100 | 7 | | | | | |
| chr4 | *TACR3* | 3 | 82 | 7 | TACR3 | tachykinin receptor 3 | Plasma Membrane | G-protein coupled recepto | SSR-146977, |
| chrX | *ZNF75D* | 3 | 92 | 7 | ZNF75D | zinc finger protein 75D | Other | other | |
| chr15 | *FAM154B* | 3 | 78 | 7 | SAXO2 | stabilizer of axonemal | Nucleus | other | |
| chr3 | *ALG1L6P* | 3 | 138 | 7 | ALG1L6P | ALG1 like 6, pseudogene | Other | other | |
| chr16 | *RP11-626G11.4* | 3 | 68 | 7 | | | | | |
| chr18 | *SERPINB7* | 2 | 70 | 7 | SERPINB7 | serpin family B member 7 | Cytoplasm | other | |
| chr1 | *PPT1* | 3 | 69 | 7 | PPT1 | palmitoyl-protein thioesterase 1 | Cytoplasm | enzyme | |
| chr12 | *B4GALNT1* | 3 | 149 | 7 | B4GALNT1 | beta-1,4-N-acetyl-galactosaminyltransferase 1 | Cytoplasm | enzyme | |
| chr19 | *QPCTL* | 3 | 88 | 7 | QPCTL | glutaminyl-peptide cyclotransferase like | Cytoplasm | enzyme | |
| chr16 | *ERVK13-1* | 3 | 113 | 7 | ERVK13-1 | endogenous retrovirus group K13 member 1 | Other | other | |
| chr14 | *SRP54* | 3 | 64 | 7 | SRP54 | signal recognition particle 54 | Cytoplasm | other | |
| chr1 | *RP11-563D10.1* | 3 | 78 | 7 | | | | | |
| chr12 | *RP11-749H20.1* | 3 | 56 | 7 | | | | | |
| chr9 | *LPAR1* | 3 | 68 | 7 | LPAR1 | lysophosphatidic acid receptor 1 | Plasma Membrane | G-protein coupled recepto | |
| chr1 | *SCNM1* | 3 | 118 | 7 | SCNM1 | sodium channel modifier 1 | Nucleus | other | |
| chr15 | *SECISBP2L* | 3 | 58 | 7 | SECISBP2L | SECIS binding protein 2 like | Other | other | |
| chr19 | *RP11-255H23.4* | 3 | 55 | 7 | | | | | |
| chr5 | *CTC-236F12.4* | 4 | 71 | 7 | | | | | |
| chr19 | *AC007228.5* | 3 | 73 | 7 | | | | | |
| chr3 | *ALG1L2* | 3 | 139 | 7 | ALG1L2 | ALG1 chitobiosyldiphosphodolichol | Other | other | |
| chr5 | *CCNB1* | 3 | 74 | 7 | CCNB1 | cyclin B1 | Cytoplasm | kinase | |
| chr1 | *PLEKHN1* | 4 | 113 | 7 | PLEKHN1 | pleckstrin homology domain containing N1 | Cytoplasm | other | |
| chr1 | *LRRIQ3* | 3 | 67 | 7 | LRRIQ3 | leucine rich repeats and IQ motif containing 3 | Other | other | |
| chr13 | *PSPC1P1* | 3 | 74 | 7 | PSPC1P1 | paraspeckle component 1 pseudogene 1 | Other | other | |
| chr6 | *B3GAT2* | 3 | 83 | 7 | B3GAT2 | beta-1,3-glucuronyltransferase 2 | Cytoplasm | enzyme | |
| chr17 | *FTSJ3* | 3 | 91 | 7 | FTSJ3 | FtsJ RNA 2'-O-methyltransferase | Nucleus | enzyme | |
| chr7 | *TRPV5* | 3 | 97 | 7 | TRPV5 | transient receptor potential cation channel subfamily V member 5 | Plasma Membrane | ion channel | |
| chr3 | *NSUN3* | 4 | 93 | 7 | NSUN3 | NOP2/Sun RNA | Cytoplasm | enzyme | |
| chr9 | *FAM69B* | 4 | 186 | 7 | DIPK1B | divergent protein kinase domain | Other | other | |
| chr1 | *RP11-176D17.3* | 2 | 82 | 7 | | | | | |
| chr1 | *RP11-488L18.3* | 3 | 106 | 7 | | | | | |
| chr14 | *RP11-314P15.2* | 3 | 79 | 7 | | | | | |
| chr11 | *RP13-631K18.3* | 3 | 130 | 7 | | | | | |
| chr22 | *C1QTNF6* | 3 | 104 | 7 | C1QTNF6 | C1q and TNF related 6 | Extracellular Space | other | |
| chr21 | *MRAP* | 3 | 116 | 7 | MRAP | melanocortin 2 receptor accessory protein | Cytoplasm | other | |
| chr19 | *KLK2* | 4 | 139 | 7 | KLK2 | kallikrein related peptidase 2 | Extracellular Space | peptidase | |
| chr12 | *RP11-167N24.4* | 3 | 56 | 7 | | | | | |
| chr1 | *RP5-968D22.3* | 3 | 82 | 7 | | | | | |
| chr17 | *C17orf98* | 3 | 109 | 7 | C17orf98 | chromosome 17 open reading frame 98 | Other | other | |
| chr19 | *RAB3D* | 3 | 92 | 7 | RAB3D | RAB3D, member RAS oncogene family | Cytoplasm | enzyme | |
| chr9 | *RP11-399D6.2* | 3 | 53 | 7 | | | | | |
| chr9 | *GBA2* | 3 | 109 | 7 | GBA2 | glucosylceramidase beta 2 | Cytoplasm | enzyme | |
| chr7 | *FAM115C* | 3 | 99 | 7 | TCAF2 | TRPM8 channel associated factor | Plasma Membrane | other | |
| chr3 | *PLOD2* | 3 | 66 | 7 | PLOD2 | procollagen-lysine,2-oxoglutarate 5-dioxygenase 2 | Cytoplasm | enzyme | |
| chr6 | *ZNF391* | 3 | 121 | 7 | ZNF391 | zinc finger protein 391 | Other | other | |
| chr15 | *COMMD4* | 3 | 120 | 7 | COMMD4 | COMM domain containing 4 | Cytoplasm | other | |
| chr15 | *RP11-10O17.1* | 3 | 65 | 7 | | | | | |
| chr14 | *C14orf2* | 3 | 100 | 7 | ATP5MPL | ATP synthase membrane subunit 6.8PL | Cytoplasm | other | |
| chr17 | *RNMTL1* | 3 | 108 | 7 | MRM3 | mitochondrial rRNA methyltransferase 3 | Cytoplasm | enzyme | |
| chr19 | *SIRT6* | 3 | 122 | 7 | SIRT6 | sirtuin 6 | Nucleus | enzyme | |
| chr22 | *TBC1D10A* | 3 | 79 | 7 | TBC1D10A | TBC1 domain family member 10A | Plasma Membrane | other | |
| chr10 | *BMS1P7* | 3 | 128 | 7 | BMS1P7 | BMS1 pseudogene 7 | Other | other | |
| chr4 | *RP11-679C8.2* | 3 | 65 | 7 | | | | | |
| chr3 | *H1FX-AS1* | 3 | 185 | 7 | H1FX-AS1 | H1FX antisense RNA 1 | Other | other | |
| chr1 | *RTCA* | 4 | 80 | 7 | RTCA | RNA 3'-terminal phosphate | Nucleus | enzyme | |
| chr3 | *FBXO45* | 3 | 62 | 7 | FBXO45 | F-box protein 45 | Plasma Membrane | other | |
| chr19 | *KLK14* | 3 | 153 | 7 | KLK14 | kallikrein related peptidase 14 | Extracellular Space | peptidase | |
| chr7 | *AC002127.4* | 3 | 75 | 7 | | | | | |
| chr5 | *CTC-325J23.3* | 4 | 67 | 7 | | | | | |
| chr1 | *C1orf64* | 3 | 97 | 7 | SRARP | steroid receptor associated and regulated protein | Other | other | |
| chr11 | *CAT* | 4 | 134 | 7 | CAT | catalase | Cytoplasm | enzyme | fomepizole |
| chr22 | *SLC25A18* | 3 | 74 | 7 | SLC25A18 | solute carrier family 25 member | Cytoplasm | transporter | |
| chr4 | *PAPSS1* | 2 | 63 | 7 | PAPSS1 | 3'-phosphoadenosine 5'-phosphosulfate synthase 1 | Cytoplasm | enzyme | |
| chr11 | *BCO2* | 3 | 82 | 7 | BCO2 | beta-carotene oxygenase 2 | Cytoplasm | enzyme | |
| chr5 | *CTD-2023M8.1* | 3 | 198 | 7 | | | | | |
| chr16 | *DDX19B* | 3 | 102 | 7 | DDX19B | DEAD-box helicase 19B | Nucleus | enzyme | |
| chr19 | *EEF2* | 4 | 165 | 7 | EEF2 | eukaryotic translation elongation factor 2 | Cytoplasm | translation regulator | MDNA55 |
| chr1 | *RP11-6J21.2* | 3 | 100 | 7 | | | | | |
| chr10 | *TECTB* | 3 | 92 | 7 | TECTB | tectorin beta | Extracellular Space | other | |
| chr19 | *TEAD2* | 3 | 119 | 7 | TEAD2 | TEA domain transcription factor 2 | Nucleus | transcription regulator | |

| | | | | | | | | | |
|---|---|---|---|---|---|---|---|---|---|
| chr10 | *RP11-478K7.2* | 3 | 98 | 7 | | | | | |
| chr11 | *CABP4* | 4 | 198 | 7 | CABP4 | calcium binding protein 4 | Plasma Membrane | other | |
| chr19 | *ZNF257* | 4 | 84 | 7 | ZNF257 | zinc finger protein 257 | Nucleus | transcription regulator | |
| chr5 | *SLC45A2* | 3 | 306 | 7 | SLC45A2 | solute carrier family 45 member 2 | Plasma Membrane | transporter | |
| chr11 | *MSANTD4* | 3 | 97 | 7 | MSANTD4 | Myb/SANT DNA binding domain containing 4 with coiled-coils | Nucleus | other | |
| chr16 | *NPIPA8* | 3 | 95 | 7 | PA1 (includes oth | nuclear pore complex interacting protein family member A3 | Nucleus | other | |
| chrX | *MOSPD2* | 3 | 69 | 7 | MOSPD2 | motile sperm domain containing 2 | Plasma Membrane | other | |
| chr1 | *RP11-25K21.6* | 3 | 110 | 7 | | | | | |
| chr15 | *AC009878.2* | 3 | 66 | 7 | | | | | |
| chr19 | *TRIP10* | 3 | 170 | 7 | TRIP10 | thyroid hormone receptor interactor 10 | Cytoplasm | other | |
| chr2 | *DQX1* | 3 | 92 | 7 | DQX1 | DEAQ-box RNA dependent ATPase 1 | Other | enzyme | |
| chr1 | *EFNA1* | 3 | 235 | 7 | EFNA1 | ephrin A1 | Plasma Membrane | other | |
| chr17 | *RP11-640N20.6* | 3 | 106 | 7 | | | | | |
| chr12 | *GPR162* | 3 | 140 | 7 | GPR162 | G protein-coupled receptor 162 | Plasma Membrane | -protein coupled recepto | |
| chr9 | *CBWD6* | 3 | 81 | 7 | CBWD3/CBWD6 | COBW domain containing 3 | Other | other | |
| chr17 | *RP11-94L15.2* | 3 | 74 | 7 | | | | | |
| chr16 | *CES2* | 3 | 164 | 7 | CES2 | carboxylesterase 2 | Cytoplasm | enzyme | |
| chr8 | *POP1* | 3 | 65 | 7 | POP1 | POP1 homolog, ribonuclease P/MRP subunit | Nucleus | enzyme | |
| chr11 | *PANX3* | 3 | 100 | 7 | PANX3 | pannexin 3 | Plasma Membrane | transporter | |
| chr19 | *ZNF382* | 3 | 55 | 7 | ZNF382 | zinc finger protein 382 | Nucleus | transcription regulator | |
| chr19 | *AC007193.6* | 3 | 82 | 7 | | | | | |
| chr1 | *SCNN1D* | 4 | 149 | 7 | SCNN1D | sodium channel epithelial 1 delta subunit | Plasma Membrane | ion channel | hydrochlorothiazide/triamterene, amiloride, triamterene, amiloride/hydrochlorothiazide |
| chr2 | *SMYD5* | 4 | 110 | 7 | SMYD5 | SMYD family member 5 | Other | other | |
| chr11 | *SLC5A12* | 4 | 105 | 7 | SLC5A12 | solute carrier family 5 member 12 | Cytoplasm | transporter | |
| chr3 | *ETV5* | 3 | 60 | 7 | ETV5 | ETS variant transcription factor 5 | Nucleus | transcription regulator | |
| chr10 | *PITX3* | 3 | 77 | 7 | PITX3 | paired like homeodomain 3 | Nucleus | transcription regulator | |
| chr6 | *RP11-624M8.1* | 3 | 57 | 7 | | | | | |
| chr17 | *KRT43P* | 3 | 94 | 7 | KRT43P | keratin 43 pseudogene | Other | other | |
| chr1 | *RNF187* | 3 | 151 | 7 | RNF187 | ring finger protein 187 | Nucleus | enzyme | |
| chr11 | *BTG4* | 3 | 54 | 7 | BTG4 | BTG anti-proliferation factor 4 | Other | other | |
| chr5 | *SPATA9* | 3 | 55 | 7 | SPATA9 | spermatogenesis associated 9 | Other | other | |
| chr3 | *CLDN18* | 4 | 105 | 7 | CLDN18 | claudin 18 | Plasma Membrane | other | claudiximab |
| chr18 | *ATP5A1* | 3 | 67 | 7 | ATP5F1A | ATP synthase F1 subunit alpha | Cytoplasm | transporter | |
| chr13 | *LINC00355* | 2 | 84 | 7 | LINC00355 | long intergenic non-protein coding RNA 355 | Other | other | |
| chr16 | *PRSS41* | 3 | 113 | 7 | PRSS41 | serine protease 41 | Plasma Membrane | peptidase | |
| chr7 | *AC138783.12* | 3 | 66 | 7 | | | | | |
| chrX | *IDS* | 3 | 71 | 7 | IDS | iduronate 2-sulfatase | Cytoplasm | enzyme | |
| chr12 | *WNT10B* | 4 | 158 | 7 | WNT10B | Wnt family member 10B | Extracellular Space | other | |
| chr22 | *AP000345.1* | 3 | 69 | 7 | | | | | |
| chr4 | *TMEM175* | 4 | 154 | 7 | TMEM175 | transmembrane protein 175 | Cytoplasm | ion channel | |
| chr4 | *IL15* | 3 | 86 | 7 | IL15 | interleukin 15 | Extracellular Space | cytokine | ALT-803 |
| chr4 | *ZNF595* | 3 | 175 | 7 | ZNF595 | zinc finger protein 595 | Nucleus | transcription regulator | |
| chr18 | *RP11-567M16.1* | 4 | 165 | 7 | | | | | |
| chr7 | *DLD* | 2 | 63 | 7 | DLD | dihydrolipoamide dehydrogenase | Cytoplasm | enzyme | hexachlorophene |
| chr20 | *ZSWIM1* | 3 | 148 | 7 | ZSWIM1 | zinc finger SWIM-type containing | Nucleus | other | |
| chr10 | *ZRANB1* | 3 | 55 | 7 | ZRANB1 | zinc finger RANBP2-type containing 1 | Nucleus | peptidase | |
| chr9 | *LCN6* | 4 | 237 | 7 | LCN6 | lipocalin 6 | Extracellular Space | other | |
| chr13 | *RBM26-AS1* | 3 | 63 | 7 | RBM26-AS1 | RBM26 antisense RNA 1 | Other | other | |
| chr11 | *APOC3* | 3 | 119 | 7 | APOC3 | apolipoprotein C3 | Extracellular Space | transporter | |
| chr17 | *KRT14* | 3 | 143 | 7 | KRT14 | keratin 14 | Cytoplasm | other | |
| chr22 | *USP41* | 3 | 76 | 7 | USP41 | ubiquitin specific peptidase 41 | Other | other | |
| chr2 | *SESTD1* | 3 | 65 | 7 | SESTD1 | SEC14 and spectrin domain containing 1 | Extracellular Space | other | |
| chr20 | *RP5-1125A11.1* | 4 | 121 | 7 | | | | | |
| chr1 | *PLK3* | 3 | 123 | 7 | PLK3 | polo like kinase 3 | Nucleus | kinase | |
| chr1 | *C1QB* | 3 | 118 | 7 | C1QB | complement C1q B chain | Extracellular Space | other | |
| chr13 | *LINC00343* | 3 | 53 | 7 | LINC00343 | long intergenic non-protein coding RNA 343 | Other | other | |
| chr11 | *TMEM9B-AS1* | 3 | 95 | 7 | TMEM9B-AS1 | TMEM9B antisense RNA 1 | Other | other | |
| chr19 | *CTB-70G10.1* | 3 | 72 | 7 | | | | | |
| chr2 | *AC021188.4* | 4 | 147 | 7 | | | | | |
| chr19 | *MPV17L2* | 3 | 121 | 7 | MPV17L2 | MPV17 mitochondrial inner membrane protein like 2 | Cytoplasm | other | |
| chr5 | *PGGT1B* | 3 | 59 | 7 | PGGT1B | protein geranylgeranyltransferase type I subunit beta | Cytoplasm | enzyme | |
| chr22 | *KCTD17* | 3 | 151 | 7 | KCTD17 | potassium channel tetramerization domain containing | Cytoplasm | other | |
| chr14 | *SCFD1* | 3 | 127 | 7 | SCFD1 | sec1 family domain containing 1 | Cytoplasm | transporter | |
| chr12 | *KRT76* | 3 | 148 | 7 | KRT76 | keratin 76 | Cytoplasm | other | |
| chr11 | *MYRF* | 3 | 186 | 7 | MYRF | myelin regulatory factor | Nucleus | transcription regulator | |
| chr4 | *RP11-556I14.2* | 3 | 69 | 7 | | | | | |
| chr16 | *C16orf80* | 3 | 64 | 7 | CFAP20 | cilia and flagella associated | Cytoplasm | other | |
| chr1 | *SPTA1* | 3 | 63 | 7 | SPTA1 | spectrin alpha, erythrocytic 1 | Cytoplasm | other | |
| chr15 | *IQCH-AS1* | 3 | 68 | 7 | IQCH-AS1 | IQCH antisense RNA 1 | Other | other | |
| chr5 | *FBXO38* | 3 | 78 | 7 | FBXO38 | F-box protein 38 | Other | other | |
| chr13 | *COG3* | 2 | 60 | 7 | COG3 | component of oligomeric golgi complex 3 | Cytoplasm | other | |
| chr11 | *PTS* | 3 | 107 | 7 | PTS | 6-pyruvoyltetrahydropterin | Cytoplasm | enzyme | |

| | | | | | | | | | |
|---|---|---|---|---|---|---|---|---|---|
| chr10 | *WNT8B* | 3 | 64 | 7 | WNT8B | Wnt family member 8B | Extracellular Space | other | |
| chr21 | *PFKL* | 3 | 152 | 7 | PFKL | phosphofructokinase, liver type | Cytoplasm | kinase | |
| chr1 | *RP11-16L9.4* | 4 | 83 | 7 | | | | | |
| chr4 | *DDX60* | 3 | 67 | 7 | DDX60 | DExD/H-box helicase 60 | Cytoplasm | enzyme | |
| chr6 | *STX11* | 3 | 63 | 7 | STX11 | syntaxin 11 | Plasma Membrane | transporter | |
| chr19 | *USE1* | 3 | 152 | 7 | USE1 | unconventional SNARE in the ER | Cytoplasm | other | |
| chr13 | *ELF1* | 2 | 59 | 7 | ELF1 | E74 like ETS transcription factor | Nucleus | transcription regulator | |
| chr2 | *AC016700.3* | 3 | 77 | 7 | | | | | |
| chr5 | *OTP* | 3 | 137 | 7 | OTP | orthopedia homeobox | Nucleus | transcription regulator | |
| chrX | *FAM50A* | 4 | 141 | 7 | FAM50A | family with sequence similarity 50 member A | Nucleus | other | |
| chr3 | *ACAA1* | 3 | 141 | 7 | ACAA1 | acetyl-CoA acyltransferase 1 | Cytoplasm | enzyme | trimetazidine |
| chrX | *RP11-87M18.2* | 3 | 68 | 7 | | | | | |
| chr20 | *ADIG* | 3 | 162 | 7 | ADIG | adipogenin | Cytoplasm | other | |
| chr17 | *CCDC47* | 3 | 59 | 7 | CCDC47 | coiled-coil domain containing 47 | Extracellular Space | other | |
| chr1 | *ZC3H12A* | 3 | 121 | 7 | ZC3H12A | zinc finger CCCH-type containing 12A | Cytoplasm | enzyme | |
| chr7 | *SLC26A4* | 3 | 61 | 7 | SLC26A4 | solute carrier family 26 member 4 | Plasma Membrane | transporter | |
| chr16 | *CTD-2015G9.2* | 4 | 83 | 7 | | | | | |
| chr2 | *SNED1* | 3 | 155 | 7 | SNED1 | sushi, nidogen and EGF like domains 1 | Plasma Membrane | other | |
| chr1 | *TSPAN1* | 3 | 108 | 7 | TSPAN1 | tetraspanin 1 | Cytoplasm | other | |
| chrX | *RP5-1091N2.9* | 3 | 63 | 7 | | | | | |
| chr20 | *RP4-800J21.3* | 4 | 212 | 7 | | | | | |
| chr4 | *HERC5* | 3 | 59 | 7 | HERC5 | HECT and RLD domain containing E3 ubiquitin protein | Cytoplasm | enzyme | |
| chr16 | *RP11-297M9.1* | 3 | 66 | 7 | | | | | |
| chr11 | *SLC35C1* | 4 | 202 | 7 | SLC35C1 | solute carrier family 35 member | Cytoplasm | transporter | |
| chr19 | *TBXA2R* | 3 | 118 | 7 | TBXA2R | thromboxane A2 receptor | Plasma Membrane | G-protein coupled recepto | ifetroban, torsemide, ridogrel, S-18886 |
| chr3 | *CD200* | 3 | 115 | 7 | CD200 | CD200 molecule | Plasma Membrane | other | samalizumab |
| chr15 | *RP11-752G15.8* | 3 | 91 | 7 | | | | | |
| chr11 | *USP2* | 3 | 84 | 7 | USP2 | ubiquitin specific peptidase 2 | Cytoplasm | peptidase | |
| chr9 | *C9orf106* | 3 | 170 | 7 | C9orf106 | chromosome 9 open reading frame 106 (putative) | Other | other | |
| chr13 | *RP11-545P6.2* | 3 | 63 | 7 | | | | | |
| chr17 | *C1QL1* | 3 | 143 | 7 | C1QL1 | complement C1q like 1 | Extracellular Space | other | |
| chr2 | *DIS3L2P1* | 3 | 170 | 7 | DIS3L2P1 | DIS3 like 3'-5' exoribonuclease 2 pseudogene 1 | Other | other | |
| chr20 | *MMP9* | 3 | 148 | 7 | MMP9 | matrix metallopeptidase 9 | Extracellular Space | peptidase | glucosamine, prinomastat, GS-5745, rebimastat, marimastat, MMP2 |
| chrX | *AC152010.1* | 3 | 89 | 7 | | | | | |
| chr10 | *KAZALD1* | 3 | 181 | 7 | KAZALD1 | Kazal type serine peptidase inhibitor domain 1 | Extracellular Space | other | |
| chr11 | *RP11-448P19.1* | 3 | 54 | 7 | | | | | |
| chr3 | *FAM86JP* | 3 | 212 | 7 | FAM86JP | family with sequence similarity 86, member A pseudogene | Other | other | |
| chr1 | *CASQ1* | 3 | 131 | 7 | CASQ1 | calsequestrin 1 | Cytoplasm | other | |
| chr6 | *RP3-510O8.4* | 3 | 105 | 7 | | | | | |
| chr9 | *ERP44* | 3 | 92 | 7 | ERP44 | endoplasmic reticulum protein 44 | Cytoplasm | enzyme | |
| chr2 | *OTOS* | 3 | 140 | 7 | OTOS | otospiralin | Extracellular Space | other | |
| chr3 | *RP11-359D24.1* | 2 | 64 | 7 | | | | | |
| chr1 | *RP11-230B22.1* | 3 | 72 | 7 | | | | | |
| chr2 | *GAL3ST2* | 3 | 141 | 7 | GAL3ST2 | galactose-3-O-sulfotransferase 2 | Cytoplasm | enzyme | |
| chr11 | *WDR74* | 3 | 128 | 7 | WDR74 | WD repeat domain 74 | Nucleus | other | |
| chr11 | *IFITM10* | 3 | 117 | 7 | IFITM10 | interferon induced transmembrane protein 10 | Other | other | |
| chr16 | *PLK1* | 3 | 153 | 7 | PLK1 | polo like kinase 1 | Nucleus | kinase | lipid encapsulated anti-PLK1 siRNA TKM-080301, onvansertib, GSK461364, TAK-960, MK1496, |
| chr19 | *ANGPTL6* | 3 | 101 | 7 | ANGPTL6 | angiopoietin like 6 | Extracellular Space | other | |
| chr15 | *GOLGA6L2* | 3 | 148 | 7 | GOLGA6L2 | golgin A6 family like 2 | Extracellular Space | other | |
| chr16 | *RP11-403P17.5* | 3 | 91 | 7 | | | | | |
| chr17 | *ZPBP2* | 3 | 117 | 7 | ZPBP2 | zona pellucida binding protein 2 | Extracellular Space | other | |
| chr3 | *LNP1* | 3 | 96 | 7 | LNP1 | leukemia NUP98 fusion partner 1 | Other | other | |
| chr11 | *RP11-882G5.1* | 3 | 107 | 7 | | | | | |
| chr3 | *RP11-360P21.2* | 2 | 57 | 7 | | | | | |
| chr1 | *TAF1A* | 3 | 72 | 7 | TAF1A | TATA-box binding protein associated factor, RNA | Nucleus | transcription regulator | |
| chr15 | *ANP32AP1* | 3 | 70 | 7 | ANP32AP1 | acidic nuclear phosphoprotein 32 family member A pseudogene 1 | Other | other | |
| chr16 | *LPCAT2* | 2 | 50 | 7 | LPCAT2 | lysophosphatidylcholine acyltransferase 2 | Cytoplasm | enzyme | |
| chr6 | *TRAF3IP2-AS1* | 3 | 64 | 7 | TRAF3IP2-AS1 | TRAF3IP2 antisense RNA 1 | Other | other | |
| chr11 | *MSANTD2* | 2 | 57 | 7 | MSANTD2 | Myb/SANT DNA binding domain containing 2 | Other | other | |
| chr22 | *SNRPD3* | 3 | 71 | 7 | SNRPD3 | small nuclear ribonucleoprotein D3 polypeptide | Nucleus | other | |
| chr2 | *ANKAR* | 3 | 87 | 7 | ANKAR | ankyrin and armadillo repeat containing | Nucleus | transcription regulator | |
| chr16 | *NFATC2IP* | 3 | 71 | 7 | NFATC2IP | nuclear factor of activated T cells 2 interacting protein | Nucleus | other | |
| chr11 | *SF3B2* | 3 | 163 | 7 | SF3B2 | splicing factor 3b subunit 2 | Nucleus | other | |
| chr11 | *RP11-494M8.4* | 3 | 147 | 7 | | | | | |

| | | | | | | | | | |
|---|---|---|---|---|---|---|---|---|---|
| chr14 | *CTD-2555K7.2* | 3 | 55 | 7 | | | | | |
| chr18 | *GATA6* | 3 | 70 | 7 | GATA6 | GATA binding protein 6 | Nucleus | transcription regulator | |
| chr1 | *LBR* | 3 | 90 | 7 | LBR | lamin B receptor | Nucleus | enzyme | |
| chr19 | *CD79A* | 3 | 135 | 7 | CD79A | CD79a molecule | Plasma Membrane | transmembrane receptor | |
| chr12 | *C12orf73* | 3 | 89 | 7 | C12orf73 | chromosome 12 open reading frame 73 | Other | other | |
| chr8 | *NIPAL2* | 3 | 74 | 7 | NIPAL2 | NIPA like domain containing 2 | Other | other | |
| chr1 | *RP11-69E11.4* | 3 | 106 | 7 | | | | | |
| chr22 | *CECR5* | 3 | 83 | 7 | HDHD5 | haloacid dehalogenase like hydrolase domain containing 5 | Cytoplasm | other | |
| chr7 | *TMEM196* | 3 | 99 | 7 | TMEM196 | transmembrane protein 196 | Other | other | |
| chr22 | *PMM1* | 3 | 111 | 7 | PMM1 | phosphomannomutase 1 | Cytoplasm | enzyme | |
| chr12 | *DDX51* | 3 | 95 | 7 | DDX51 | DEAD-box helicase 51 | Other | enzyme | |
| chr22 | *MAFF* | 3 | 119 | 7 | MAFF | MAF bZIP transcription factor F | Nucleus | transcription regulator | |
| chr17 | *AC026954.6* | 3 | 141 | 7 | | | | | |
| chr1 | *CDCP2* | 3 | 150 | 7 | CDCP2 | CUB domain containing protein 2 | Other | transporter | |
| chr17 | *BCL6B* | 3 | 118 | 7 | BCL6B | BCL6B transcription repressor | Nucleus | transcription regulator | |
| chr1 | *EDEM3* | 3 | 65 | 7 | EDEM3 | ER degradation enhancing alpha-mannosidase like protein 3 | Cytoplasm | enzyme | |
| chr19 | *CCDC9* | 3 | 148 | 7 | CCDC9 | coiled-coil domain containing 9 | Extracellular Space | other | |
| chr10 | *ENO4* | 3 | 71 | 7 | ENO4 | enolase 4 | Extracellular Space | enzyme | |
| chr1 | *GDAP2* | 3 | 77 | 7 | GDAP2 | ganglioside induced differentiation associated protein | Cytoplasm | other | |
| chr20 | *C20orf85* | 3 | 162 | 7 | C20orf85 | chromosome 20 open reading frame 85 | Other | other | |
| chr10 | *AVPI1* | 3 | 110 | 7 | AVPI1 | arginine vasopressin induced 1 | Other | other | |
| chr3 | *U73166.2* | 3 | 110 | 7 | | | | | |
| chr2 | *AC010982.1* | 3 | 87 | 7 | | | | | |
| chrX | *RP11-265P11.2* | 4 | 72 | 7 | | | | | |
| chr16 | *FUS* | 3 | 80 | 7 | FUS | FUS RNA binding protein | Nucleus | transcription regulator | |
| chr3 | *HPS3* | 3 | 77 | 7 | HPS3 | HPS3 biogenesis of lysosomal organelles complex 2 subunit 1 | Cytoplasm | other | |
| chrX | *RP11-142G7.2* | 3 | 64 | 7 | | | | | |
| chr14 | *RP11-58E21.3* | 3 | 53 | 7 | | | | | |
| chr22 | *CHKB* | 3 | 86 | 7 | CHKB | choline kinase beta | Cytoplasm | kinase | |
| chr6 | *PREP* | 3 | 69 | 7 | PREP | prolyl endopeptidase | Cytoplasm | peptidase | Z 321 |
| chr17 | *RP11-642M2.1* | 3 | 74 | 7 | | | | | |
| chr12 | *PEX5* | 2 | 82 | 7 | PEX5 | peroxisomal biogenesis factor 5 | Cytoplasm | other | |
| chr12 | *RP11-153K16.2* | 3 | 95 | 7 | | | | | |
| chr22 | *LGALS1* | 3 | 105 | 7 | LGALS1 | galectin 1 | Extracellular Space | other | OTX008 |
| chr15 | *RPL29P30* | 3 | 90 | 7 | RPL29P30 | ribosomal protein L29 pseudogene 30 | Other | other | |
| chr7 | *RP11-166O4.1* | 3 | 105 | 7 | | | | | |
| chr1 | *CROCCP5* | 3 | 154 | 7 | CROCCP5 | CROCC pseudogene 5 | Other | other | |
| chr1 | *MPZ* | 3 | 119 | 7 | MPZ | myelin protein zero | Plasma Membrane | other | |
| chr8 | *TDH* | 3 | 73 | 7 | TDH | L-threonine dehydrogenase (pseudogene) | Cytoplasm | other | |
| chr10 | *MKI67* | 3 | 67 | 7 | MKI67 | marker of proliferation Ki-67 | Nucleus | other | |
| chr10 | *RP13-39P12.3* | 3 | 111 | 7 | | | | | |
| chr7 | *SPDYE7P* | 3 | 79 | 7 | SPDYE7P | speedy/RINGO cell cycle regulator family member E7, | Other | other | |
| chr1 | *HMGCS2* | 2 | 100 | 7 | HMGCS2 | 3-hydroxy-3-methylglutaryl-CoA synthase 2 | Cytoplasm | enzyme | |
| chr13 | *TRPC4* | 3 | 84 | 7 | TRPC4 | transient receptor potential cation channel subfamily C member 4 | Plasma Membrane | ion channel | |
| chrX | *CA5BP1* | 3 | 92 | 7 | CA5BP1 | carbonic anhydrase 5B pseudogene 1 | Other | other | |
| chr12 | *RP1-197B17.3* | 4 | 178 | 7 | | | | | |
| chr17 | *USH1G* | 3 | 159 | 7 | USH1G | USH1 protein network component sans | Cytoplasm | other | |
| chr3 | *HIGD1A* | 3 | 91 | 7 | HIGD1A | HIG1 hypoxia inducible domain family member 1A | Cytoplasm | other | |
| chr14 | *TC2N* | 3 | 53 | 7 | TC2N | tandem C2 domains, nuclear | Nucleus | transporter | |
| chr12 | *STYK1* | 3 | 62 | 7 | STYK1 | serine/threonine/tyrosine kinase 1 | Cytoplasm | kinase | |
| chr6 | *RP3-428L16.1* | 3 | 67 | 7 | | | | | |
| chr19 | *ZBTB32* | 3 | 126 | 7 | ZBTB32 | zinc finger and BTB domain containing 32 | Nucleus | transcription regulator | |
| chrX | *PASD1* | 3 | 61 | 7 | PASD1 | PAS domain containing repressor | Nucleus | other | |
| chrX | *GAGE10* | 3 | 111 | 7 | GAGE10 | G antigen 10 | Other | other | |
| chr3 | *UBTFL8* | 3 | 75 | 7 | UBTFL8 | UBTF like 8 (pseudogene) | Other | other | |
| chr17 | *RP5-1028K7.2* | 3 | 114 | 7 | | | | | |
| chr7 | *YKT6* | 3 | 134 | 7 | YKT6 | YKT6 v-SNARE homolog | Cytoplasm | enzyme | |
| chr15 | *AC135983.2* | 2 | 124 | 7 | | | | | |
| chr5 | *IK* | 3 | 58 | 7 | IK | IK cytokine | Extracellular Space | cytokine | |
| chrX | *MTM1* | 3 | 50 | 7 | MTM1 | myotubularin 1 | Cytoplasm | phosphatase | |
| chr5 | *F2RL1* | 3 | 101 | 7 | F2RL1 | F2R like trypsin receptor 1 | Plasma Membrane | G-protein coupled receptor | |
| chr3 | *GMNC* | 3 | 93 | 7 | GMNC | geminin coiled-coil domain | Nucleus | other | |
| chr10 | *DCLRE1A* | 3 | 67 | 7 | DCLRE1A | DNA cross-link repair 1A | Nucleus | enzyme | |
| chr7 | *AC096582.7* | 3 | 92 | 7 | | | | | |
| chr19 | *NFKBIB* | 3 | 141 | 7 | NFKBIB | NFKB inhibitor beta | Nucleus | transcription regulator | |
| chr2 | *SLC9B1P2* | 3 | 103 | 7 | SLC9B1P2 | solute carrier family 9 member B1 pseudogene 2 | Other | other | |
| chr14 | *FRMD6-AS2* | 2 | 48 | 7 | FRMD6-AS2 | FRMD6 antisense RNA 2 | Other | other | |
| chr1 | *RP11-27K13.3* | 3 | 126 | 7 | | | | | |
| chr6 | *UBE3D* | 3 | 69 | 7 | UBE3D | ubiquitin protein ligase E3D | Other | other | |
| chr2 | *PTPN18* | 3 | 108 | 7 | PTPN18 | protein tyrosine phosphatase non-receptor type 18 | Nucleus | phosphatase | |
| chr22 | *CDC42EP1* | 3 | 132 | 7 | CDC42EP1 | CDC42 effector protein 1 | Extracellular Space | other | |
| chr1 | *RP1-159A19.4* | 3 | 122 | 7 | | | | | |

| | | | | | | | | | |
|---|---|---|---|---|---|---|---|---|---|
| chr5 | *C5orf60* | 3 | 138 | 7 | C5orf60 | chromosome 5 open reading frame 60 | Other | other | |
| chr6 | *RP5-899B16.2* | 2 | 64 | 7 | | | | | |
| chr12 | *BCDIN3D-AS1* | 3 | 120 | 7 | BCDIN3D-AS1 | BCDIN3D antisense RNA 1 | Other | other | |
| chr17 | *CTD-2582D11.1* | 3 | 101 | 7 | | | | | |
| chr19 | *NDUFS7* | 3 | 151 | 7 | NDUFS7 | NADH:ubiquinone oxidoreductase core subunit S7 | Cytoplasm | enzyme | |
| chr17 | *AC003101.1* | 3 | 110 | 7 | | | | | |
| chr19 | *ZNF77* | 3 | 100 | 7 | ZNF77 | zinc finger protein 77 | Nucleus | transcription regulator | |
| chr15 | *RP11-97O12.2* | 3 | 109 | 7 | | | | | |
| chr11 | *TIRAP* | 3 | 69 | 7 | TIRAP | TIR domain containing adaptor protein | Cytoplasm | other | |
| chr14 | *SERPINA5* | 3 | 82 | 7 | SERPINA5 | serpin family A member 5 | Extracellular Space | other | |
| chr11 | *ANO9* | 4 | 150 | 7 | ANO9 | anoctamin 9 | Plasma Membrane | ion channel | |
| chr10 | *C10orf55* | 3 | 140 | 7 | C10orf55 | chromosome 10 open reading frame 55 | Other | other | |
| chr19 | *RAB3A* | 3 | 153 | 7 | RAB3A | RAB3A, member RAS oncogene family | Cytoplasm | enzyme | |
| chr1 | *RP11-190A12.7* | 4 | 118 | 7 | | | | | |
| chr1 | *FCRLB* | 4 | 168 | 7 | FCRLB | Fc receptor like B | Cytoplasm | other | |
| chr6 | *OPN5* | 3 | 60 | 7 | OPN5 | opsin 5 | Plasma Membrane | i-protein coupled recepto | |
| chr19 | *HAMP* | 3 | 103 | 7 | HAMP | hepcidin antimicrobial peptide | Extracellular Space | other | lexaptepid pegol |
| chr11 | *RP11-574M7.1* | 3 | 68 | 7 | | | | | |
| chr13 | *LINC00426* | 3 | 68 | 7 | LINC00426 | long intergenic non-protein coding RNA 426 | Other | other | |
| chr12 | *DTX3* | 3 | 122 | 7 | DTX3 | deltex E3 ubiquitin ligase 3 | Cytoplasm | other | |
| chr12 | *IGFBP6* | 3 | 160 | 7 | IGFBP6 | insulin like growth factor binding protein 6 | Extracellular Space | other | |
| chr8 | *MYC* | 3 | 139 | 7 | MYC | MYC proto-oncogene, bHLH transcription factor | Nucleus | transcription regulator | AVI-4126, MYC-targeting siRNA DCR-MYC |
| chr17 | *DGKE* | 3 | 90 | 7 | DGKE | diacylglycerol kinase epsilon | Cytoplasm | kinase | |
| chrX | *IKBKGP1* | 4 | 163 | 7 | IKBKGP1 | inhibitor of nuclear factor kappa B kinase subunit gamma | Other | other | |
| chr11 | *RP11-804A23.2* | 3 | 125 | 7 | | | | | |
| chr7 | *MEST* | 3 | 120 | 7 | MEST | mesoderm specific transcript | Cytoplasm | peptidase | |
| chr16 | *RP11-293N14.1* | 3 | 55 | 7 | | | | | |
| chr3 | *C3orf56* | 3 | 152 | 7 | C3orf56 | chromosome 3 open reading frame 56 | Other | other | |
| chrX | *RGN* | 3 | 56 | 7 | RGN | regucalcin | Nucleus | enzyme | |
| chr20 | *C20orf166-AS1* | 4 | 307 | 7 | MIR1-1HG-AS1 | MIR1-1HG antisense RNA 1 | Other | other | |
| chr1 | *PAQR7* | 3 | 95 | 7 | PAQR7 | progestin and adipoQ receptor family member 7 | Plasma Membrane | other | |
| chr14 | *RHOJ* | 2 | 60 | 7 | RHOJ | ras homolog family member J | Cytoplasm | enzyme | |
| chr4 | *RP11-326I19.3* | 3 | 85 | 7 | | | | | |
| chr21 | *RSPH1* | 3 | 88 | 7 | RSPH1 | radial spoke head component 1 | Nucleus | other | |
| chr17 | *AC025627.9* | 3 | 186 | 7 | | | | | |
| chr4 | *PPM1K* | 3 | 73 | 7 | PPM1K | protein phosphatase, Mg2+/Mn2+ dependent 1K | Cytoplasm | phosphatase | |
| chr8 | *FAM66A* | 3 | 114 | 7 | FAM66A | family with sequence similarity 66 member A | Other | other | |
| chr2 | *AC017104.6* | 3 | 125 | 7 | | | | | |
| chr17 | *BAIAP2-AS1* | 3 | 173 | 7 | BAIAP2-DT | BAIAP2 divergent transcript | Other | other | |
| chr1 | *CERS2* | 3 | 122 | 7 | CERS2 | ceramide synthase 2 | Nucleus | transcription regulator | |
| chr4 | *TTC29* | 3 | 75 | 7 | TTC29 | tetratricopeptide repeat domain | Other | other | |
| chr12 | *C12orf49* | 3 | 86 | 7 | C12orf49 | chromosome 12 open reading frame 49 | Other | other | |
| chr10 | *CYP26C1* | 3 | 88 | 7 | CYP26C1 | cytochrome P450 family 26 subfamily C member 1 | Cytoplasm | enzyme | |
| chr4 | *UCHL1-AS1* | 3 | 61 | 7 | UCHL1-AS1 | UCHL1 antisense RNA 1 (head to head) | Other | other | |
| chr11 | *SPDYC* | 3 | 137 | 7 | SPDYC | speedy/RINGO cell cycle regulator family member C | Nucleus | other | |
| chr19 | *TINCR* | 2 | 105 | 7 | TINCR | TINCR ubiquitin domain | Other | other | |
| chrX | *MAGEA3* | 3 | 99 | 7 | /IAGEA3/MAGEA( | MAGE family member A6 | Cytoplasm | other | |
| chr16 | *CMTM2* | 3 | 112 | 7 | CMTM2 | CKLF like MARVEL transmembrane domain | Extracellular Space | cytokine | |
| chr1 | *RP11-90C4.2* | 3 | 82 | 7 | | | | | |
| chr5 | *TCERG1* | 3 | 96 | 7 | TCERG1 | transcription elongation regulator | Nucleus | transcription regulator | |
| chr17 | *RP11-166P13.3* | 3 | 80 | 7 | | | | | |
| chr11 | *PSMD13* | 3 | 60 | 7 | PSMD13 | proteasome 26S subunit, non-ATPase 13 | Cytoplasm | peptidase | |
| chr1 | *TRAF3IP3* | 3 | 123 | 7 | TRAF3IP3 | TRAF3 interacting protein 3 | Other | other | |
| chr7 | *RP11-38M8.1* | 3 | 126 | 7 | | | | | |
| chr5 | *CTC-551A13.1* | 3 | 71 | 7 | | | | | |
| chr7 | *C7orf61* | 3 | 141 | 7 | C7orf61 | chromosome 7 open reading frame 61 | Nucleus | other | |
| chr16 | *CDIPT* | 3 | 153 | 7 | CDIPT | CDP-diacylglycerol--inositol 3-phosphatidyltransferase | Cytoplasm | enzyme | |
| chr10 | *RP11-168P8.5* | 3 | 115 | 7 | | | | | |
| chr8 | *RP11-231D20.2* | 2 | 55 | 7 | | | | | |
| chr1 | *PIAS3* | 3 | 148 | 7 | PIAS3 | protein inhibitor of activated | Nucleus | transcription regulator | |
| chr10 | *KLF6* | 2 | 62 | 7 | KLF6 | Kruppel like factor 6 | Nucleus | transcription regulator | |
| chr2 | *PDK1* | 3 | 65 | 7 | PDK1 | pyruvate dehydrogenase kinase 1 | Cytoplasm | kinase | dichloroacetic acid |
| chr3 | *HCLS1* | 4 | 84 | 7 | HCLS1 | hematopoietic cell-specific Lyn substrate 1 | Nucleus | other | |
| chr19 | *CTD-2586B10.1* | 3 | 97 | 7 | | | | | |
| chr9 | *C9orf16* | 3 | 192 | 7 | C9orf16 | chromosome 9 open reading frame 16 | Other | other | |

| | | | | | | | | | |
|---|---|---|---|---|---|---|---|---|---|
| chr12 | *RP11-493L12.5* | 3 | 86 | 7 | | | | | |
| chr15 | *PATL2* | 3 | 63 | 7 | PATL2 | PAT1 homolog 2 | Cytoplasm | translation regulator | |
| chrX | *MAGEA9B* | 3 | 107 | 7 | IAGEA9/MAGEA9 | MAGE family member A9 | Other | other | |
| chr7 | *IFRD1* | 3 | 92 | 7 | IFRD1 | interferon related developmental regulator 1 | Nucleus | other | |
| chr8 | *ESRP1* | 3 | 74 | 7 | ESRP1 | epithelial splicing regulatory | Nucleus | other | |
| chr16 | *RP11-616M22.10* | 3 | 130 | 7 | | | | | |
| chr5 | *CTC-205M6.5* | 3 | 163 | 7 | | | | | |
| chr19 | *AC004076.7* | 3 | 61 | 7 | | | | | |
| chrX | *SSX4B* | 3 | 89 | 7 | SSX4/SSX4B | SSX family member 4 | Nucleus | other | |
| chr11 | *MUC2* | 4 | 386 | 7 | MUC2 | mucin 2, oligomeric mucus/gel-forming | Extracellular Space | other | |
| chr7 | *UNC93B2* | 3 | 180 | 6 | UNC93B2 | unc-93 homolog B2, pseudogene | Other | other | |
| chr17 | *AC015849.2* | 3 | 86 | 6 | | | | | |
| chr7 | *RP11-220H4.2* | 3 | 177 | 6 | | | | | |
| chr1 | *RGS16* | 3 | 159 | 6 | RGS16 | regulator of G protein signaling | Cytoplasm | enzyme | AGS-16M18 |
| chr9 | *RP11-111F5.5* | 3 | 56 | 6 | | | | | |
| chrX | *ARHGAP36* | 3 | 110 | 6 | ARHGAP36 | Rho GTPase activating protein 36 | Cytoplasm | other | |
| chr7 | *PP13004* | 3 | 158 | 6 | | | | | |
| chr19 | *ZNF792* | 3 | 81 | 6 | ZNF792 | zinc finger protein 792 | Other | other | |
| chr10 | *PNLIPRP3* | 3 | 70 | 6 | PNLIPRP3 | pancreatic lipase related protein 3 | Other | enzyme | |
| chr1 | *ZNF687* | 3 | 128 | 6 | ZNF687 | zinc finger protein 687 | Nucleus | transcription regulator | |
| chr19 | *ZNF223* | 3 | 61 | 6 | ZNF223 | zinc finger protein 223 | Nucleus | other | |
| chr17 | *LRRC45* | 3 | 165 | 6 | LRRC45 | leucine rich repeat containing 45 | Cytoplasm | other | |
| chr13 | *LINC00448* | 2 | 61 | 6 | LINC00448 | long intergenic non-protein coding RNA 448 | Other | other | |
| chr1 | *PPIEL* | 3 | 85 | 6 | PPIEL | peptidylprolyl isomerase E like pseudogene | Other | other | |
| chr19 | *CYP2A13* | 3 | 143 | 6 | P2A6 (includes oth | cytochrome P450 family 2 subfamily A member 6 | Cytoplasm | enzyme | |
| chr22 | *BCRP3* | 3 | 100 | 6 | BCRP3 | BCR pseudogene 3 | Other | other | |
| chr8 | *IL7* | 3 | 85 | 6 | IL7 | interleukin 7 | Extracellular Space | cytokine | MK-1966 |
| chr18 | *RP11-411B10.5* | 3 | 145 | 6 | | | | | |
| chr13 | *DNAJC3-AS1* | 3 | 89 | 6 | DNAJC3-DT | DNAJC3 divergent transcript | Other | other | |
| chr11 | *DAK* | 3 | 181 | 6 | TKFC | triokinase and FMN cyclase | Cytoplasm | kinase | |
| chr4 | *MND1* | 3 | 51 | 6 | MND1 | meiotic nuclear divisions 1 | Nucleus | other | |
| chr9 | *LCN2* | 3 | 208 | 6 | LCN2 | lipocalin 2 | Extracellular Space | transporter | |
| chr7 | *WBSCR28* | 3 | 121 | 6 | TMEM270 | transmembrane protein 270 | Other | other | |
| chr7 | *AC093627.9* | 3 | 161 | 6 | | | | | |
| chr9 | *RP13-60M5.2* | 3 | 67 | 6 | | | | | |
| chr17 | *CD300LD* | 3 | 98 | 6 | CD300LD | CD300 molecule like family | Plasma Membrane | other | |
| chr6 | *RARS2* | 2 | 47 | 6 | RARS2 | arginyl-tRNA synthetase 2, mitochondrial | Cytoplasm | enzyme | |
| chr1 | *RP11-458D21.1* | 3 | 306 | 6 | | | | | |
| chr11 | *CYB5R2* | 3 | 183 | 6 | CYB5R2 | cytochrome b5 reductase 2 | Cytoplasm | enzyme | |
| chrX | *NKAP* | 2 | 56 | 6 | NKAP | NFKB activating protein | Nucleus | transcription regulator | |
| chr2 | *C2orf50* | 3 | 92 | 6 | C2orf50 | chromosome 2 open reading frame 50 | Other | other | |
| chr3 | *TUSC2* | 3 | 190 | 6 | TUSC2 | tumor suppressor 2, mitochondrial calcium regulator | Nucleus | other | |
| chr14 | *SLC10A1* | 2 | 52 | 6 | SLC10A1 | solute carrier family 10 member 1 | Plasma Membrane | transporter | |
| chrX | *PNCK* | 3 | 134 | 6 | PNCK | pregnancy up-regulated nonubiquitous CaM kinase | Other | kinase | |
| chrX | *LINC00632* | 3 | 73 | 6 | LINC00632 | long intergenic non-protein coding RNA 632 | Other | other | |
| chr10 | *LINC00263* | 3 | 97 | 6 | OLMALINC | oligodendrocyte maturation-associated long intergenic non-coding RNA | Other | other | |
| chr1 | *MUL1* | 2 | 90 | 6 | MUL1 | mitochondrial E3 ubiquitin protein ligase 1 | Cytoplasm | enzyme | |
| chr10 | *TM9SF3* | 3 | 68 | 6 | TM9SF3 | transmembrane 9 superfamily member 3 | Cytoplasm | transporter | |
| chr5 | *CTC-551A13.2* | 2 | 71 | 6 | | | | | |
| chr1 | *RP11-182I10.3* | 3 | 62 | 6 | | | | | |
| chr19 | *LILRB2* | 3 | 217 | 6 | LILRB2 | leukocyte immunoglobulin like receptor B2 | Plasma Membrane | transmembrane receptor | MK-4830 |
| chr7 | *AC004840.8* | 3 | 108 | 6 | | | | | |
| chr9 | *NMRK1* | 3 | 51 | 6 | NMRK1 | nicotinamide riboside kinase 1 | Cytoplasm | kinase | |
| chr1 | *RP11-540K16.1* | 3 | 104 | 6 | | | | | |
| chr20 | *RP5-1010E17.1* | 2 | 78 | 6 | | | | | |
| chr19 | *POLR2E* | 4 | 159 | 6 | POLR2E | RNA polymerase II subunit E | Nucleus | enzyme | |
| chr1 | *FUBP1* | 2 | 60 | 6 | FUBP1 | far upstream element binding protein 1 | Nucleus | transcription regulator | |
| chr1 | *GBP4* | 2 | 56 | 6 | GBP4 | guanylate binding protein 4 | Cytoplasm | enzyme | |
| chr19 | *ZNF574* | 3 | 144 | 6 | ZNF574 | zinc finger protein 574 | Other | other | |
| chrX | *FAM104B* | 2 | 74 | 6 | FAM104B | family with sequence similarity 104 member B | Other | other | |
| chrX | *MAGEB3* | 3 | 80 | 6 | MAGEB3 | MAGE family member B3 | Other | other | |
| chr2 | *SLC16A14* | 3 | 63 | 6 | SLC16A14 | solute carrier family 16 member | Other | other | |
| chr5 | *CTB-33O18.1* | 3 | 124 | 6 | | | | | |
| chr8 | *PPP1R3B* | 3 | 105 | 6 | PPP1R3B | protein phosphatase 1 regulatory subunit 3B | Cytoplasm | phosphatase | |
| chr10 | *HNRNPF* | 3 | 126 | 6 | HNRNPF | heterogeneous nuclear ribonucleoprotein F | Nucleus | other | |
| chr4 | *RP11-502M1.2* | 3 | 67 | 6 | | | | | |
| chr10 | *RP11-432J24.3* | 3 | 175 | 6 | | | | | |
| chr13 | *RP11-383H17.4* | 3 | 72 | 6 | | | | | |
| chr8 | *PNMA2* | 3 | 105 | 6 | PNMA2 | PNMA family member 2 | Nucleus | transporter | |

| | | | | | | | | | |
|---|---|---|---|---|---|---|---|---|---|
| chr19 | *RFXANK* | 3 | 93 | 6 | RFXANK | regulatory factor X associated ankyrin containing protein | Nucleus | transcription regulator | |
| chr2 | *AC073128.10* | 3 | 95 | 6 | | | | | |
| chr18 | *FECH* | 3 | 82 | 6 | FECH | ferrochelatase | Cytoplasm | enzyme | |
| chr1 | *PDZK1IP1* | 3 | 165 | 6 | PDZK1IP1 | PDZK1 interacting protein 1 | Extracellular Space | other | |
| chr3 | *ALG1L* | 3 | 186 | 6 | ALG1L | ALG1 chitobiosyldiphosphodolichol | Other | other | |
| chr2 | *ZNF2* | 3 | 87 | 6 | ZNF2 | zinc finger protein 2 | Other | other | |
| chr22 | *CRKL* | 3 | 68 | 6 | CRKL | CRK like proto-oncogene, adaptor protein | Cytoplasm | kinase | |
| chr12 | *SDR9C7* | 3 | 74 | 6 | SDR9C7 | short chain dehydrogenase/reductase family 9C member 7 | Other | enzyme | |
| chr7 | *C7orf41* | 3 | 96 | 6 | MTURN | maturin, neural progenitor differentiation regulator homolog | Other | other | |
| chr16 | *RP11-473M20.14* | 4 | 136 | 6 | | | | | |
| chr1 | *HPCA* | 3 | 119 | 6 | HPCA | hippocalcin | Cytoplasm | other | |
| chr1 | *LSM10* | 3 | 146 | 6 | LSM10 | LSM10, U7 small nuclear RNA associated | Nucleus | other | |
| chr6 | *TMEM14C* | 3 | 66 | 6 | TMEM14C | transmembrane protein 14C | Plasma Membrane | other | |
| chr3 | *RP11-246A10.1* | 3 | 76 | 6 | | | | | |
| chr1 | *TAS1R2* | 3 | 99 | 6 | TAS1R2 | taste 1 receptor member 2 | Plasma Membrane | G-protein coupled recepto | |
| chr19 | *ZNF610* | 3 | 62 | 6 | ZNF610 | zinc finger protein 610 | Nucleus | transcription regulator | |
| chr9 | *FAM163B* | 3 | 145 | 6 | FAM163B | family with sequence similarity 163 member B | Other | other | |
| chr3 | *CCR2* | 2 | 55 | 6 | CCR2 | C-C motif chemokine receptor 2 | Plasma Membrane | G-protein coupled recepto | propagermanium, BMS-813160, AZD2423, MLN1202, |
| chr3 | *KLHL40* | 3 | 108 | 6 | KLHL40 | kelch like family member 40 | Cytoplasm | other | |
| chr8 | *CTD-2547L16.1* | 3 | 68 | 6 | | | | | |
| chr5 | *RHOBTB3* | 2 | 50 | 6 | RHOBTB3 | Rho related BTB domain | Cytoplasm | enzyme | |
| chr13 | *MRPS31* | 2 | 62 | 6 | MRPS31 | mitochondrial ribosomal protein | Cytoplasm | other | |
| chr12 | *CSRP2* | 3 | 61 | 6 | CSRP2 | cysteine and glycine rich protein | Nucleus | other | |
| chr2 | *TPRKB* | 3 | 91 | 6 | TPRKB | TP53RK binding protein | Cytoplasm | other | |
| chr2 | *UBXN4* | 2 | 54 | 6 | UBXN4 | UBX domain protein 4 | Extracellular Space | other | |
| chr3 | *BSN-AS2* | 3 | 102 | 6 | BSN-DT | BSN divergent transcript | Other | other | |
| chr17 | *GP1BA* | 3 | 105 | 6 | GP1BA | glycoprotein Ib platelet subunit | Plasma Membrane | transmembrane recepto | |
| chr3 | *RP11-548O1.3* | 4 | 147 | 6 | | | | | |
| chr1 | *TMEM79* | 3 | 146 | 6 | TMEM79 | transmembrane protein 79 | Cytoplasm | other | |
| chr2 | *CHCHD5* | 3 | 142 | 6 | CHCHD5 | coiled-coil-helix-coiled-coil-helix domain containing 5 | Other | other | |
| chr17 | *CTD-2206N4.4* | 3 | 91 | 6 | | | | | |
| chr4 | *SRD5A3* | 2 | 57 | 6 | SRD5A3 | steroid 5 alpha-reductase 3 | Cytoplasm | enzyme | |
| chr9 | *SPATA31C1* | 3 | 156 | 6 | SPATA31C1 | SPATA31 subfamily C member 1 | Other | other | |
| chr12 | *RP11-162P23.2* | 3 | 85 | 6 | | | | | |
| chr1 | *IGSF8* | 3 | 213 | 6 | IGSF8 | immunoglobulin superfamily member 8 | Plasma Membrane | other | |
| chr1 | *RP11-66A2.2* | 3 | 72 | 6 | | | | | |
| chr6 | *CNPY3* | 3 | 114 | 6 | CNPY3 | canopy FGF signaling regulator 3 | Cytoplasm | other | |
| chr4 | *IGFBP7-AS1* | 3 | 71 | 6 | IGFBP7-AS1 | IGFBP7 antisense RNA 1 | Other | other | |
| chr11 | *UCP3* | 3 | 185 | 6 | UCP3 | uncoupling protein 3 | Cytoplasm | transporter | |
| chr19 | *TFPT* | 3 | 145 | 6 | TFPT | TCF3 fusion partner | Nucleus | other | |
| chr19 | *C19orf69* | 3 | 93 | 6 | ERICH4 | glutamate rich 4 | Other | other | |
| chr11 | *RAD9A* | 3 | 165 | 6 | RAD9A | RAD9 checkpoint clamp component A | Nucleus | enzyme | |
| chr19 | *LENG8-AS1* | 3 | 104 | 6 | LENG8-AS1 | | Other | other | |
| chr19 | *COPE* | 3 | 143 | 6 | COPE | coatomer protein complex subunit epsilon | Cytoplasm | transporter | |
| chr7 | *AC005022.1* | 3 | 100 | 6 | | | | | |
| chr6 | *CEP57L1* | 2 | 59 | 6 | CEP57L1 | centrosomal protein 57 like 1 | Other | other | |
| chr12 | *GPR19* | 3 | 64 | 6 | GPR19 | G protein-coupled receptor 19 | Plasma Membrane | G-protein coupled recepto | |
| chr1 | *RP11-174G17.2* | 3 | 66 | 6 | | | | | |
| chr20 | *EMILIN3* | 3 | 168 | 6 | EMILIN3 | elastin microfibril interfacer 3 | Extracellular Space | other | |
| chr19 | *KCNJ14* | 3 | 91 | 6 | KCNJ14 | potassium inwardly rectifying channel subfamily J member 14 | Plasma Membrane | ion channel | |
| chr16 | *EXOC3L1* | 3 | 165 | 6 | EXOC3L1 | exocyst complex component 3 | Cytoplasm | other | |
| chr17 | *SUMO2* | 3 | 77 | 6 | SUMO2 | small ubiquitin like modifier 2 | Nucleus | enzyme | |
| chr8 | *FAM90A8P* | 3 | 556 | 6 | | | | | |
| chr12 | *RP11-82C23.2* | 3 | 104 | 6 | | | | | |
| chr1 | *NBPF24* | 3 | 94 | 6 | PF1 (includes othe | NBPF member 11 | Other | other | |
| chr16 | *RP11-2C15.1* | 3 | 71 | 6 | | | | | |
| chr13 | *DLEU2* | 2 | 55 | 6 | DLEU2 | deleted in lymphocytic leukemia 2 | Other | other | |
| chr1 | *LPPR4* | 3 | 117 | 6 | | | | | |
| chr2 | *AC011286.1* | 2 | 69 | 6 | | | | | |
| chr17 | *METTL23* | 3 | 94 | 6 | METTL23 | methyltransferase like 23 | Cytoplasm | other | |
| chr11 | *LRTOMT* | 3 | 62 | 6 | LRTOMT | leucine rich transmembrane and O-methyltransferase domain | Plasma Membrane | enzyme | |
| chr15 | *TM6SF1* | 3 | 130 | 6 | TM6SF1 | transmembrane 6 superfamily member 1 | Plasma Membrane | other | |
| chr2 | *RMDN2* | 2 | 65 | 6 | RMDN2 | regulator of microtubule dynamics | Cytoplasm | other | |
| chr9 | *RP11-188C12.3* | 3 | 174 | 6 | | | | | |
| chr10 | *TIMM23* | 3 | 83 | 6 | TIMM23 | translocase of inner mitochondrial membrane 23 | Cytoplasm | transporter | |
| chr7 | *RP11-142A5.1* | 3 | 79 | 6 | | | | | |
| chr22 | *RPL3* | 3 | 85 | 6 | RPL3 | ribosomal protein L3 | Nucleus | other | omacetaxine mepesuccinate |
| chr17 | *SLFN13* | 3 | 146 | 6 | SLFN13 | schlafen family member 13 | Nucleus | enzyme | |
| chr22 | *AP000351.4* | 3 | 122 | 6 | | | | | |
| chr16 | *CTD-3203P2.2* | 3 | 58 | 6 | | | | | |

| | | | | | | | | | |
|---|---|---|---|---|---|---|---|---|---|
| chr3 | *RP11-543D10.2* | 3 | 118 | 6 | | | | | |
| chr20 | *NELFCD* | 3 | 154 | 6 | NELFCD | negative elongation factor complex member C/D | Nucleus | other | |
| chr4 | *LEF1* | 3 | 74 | 6 | LEF1 | lymphoid enhancer binding factor | Nucleus | transcription regulator | |
| chr1 | *KIAA0040* | 4 | 97 | 6 | KIAA0040 | KIAA0040 | Other | other | |
| chr2 | *WNT6* | 3 | 95 | 6 | WNT6 | Wnt family member 6 | Extracellular Space | other | |
| chr3 | *RRP9* | 3 | 122 | 6 | RRP9 | ribosomal RNA processing 9, U3 small nucleolar RNA binding | Nucleus | other | |
| chr9 | *GRHPR* | 3 | 101 | 6 | GRHPR | glyoxylate and hydroxypyruvate reductase | Cytoplasm | enzyme | |
| chr15 | *RP11-160C18.2* | 3 | 133 | 6 | | | | | |
| chr2 | *CCDC141* | 2 | 57 | 6 | CCDC141 | coiled-coil domain containing 141 | Cytoplasm | other | |
| chr1 | *SEC22B* | 3 | 231 | 6 | SEC22B | SEC22 homolog B, vesicle trafficking protein | Cytoplasm | other | |
| chr3 | *RP11-719N22.2* | 3 | 56 | 6 | | | | | |
| chr5 | *TMEM171* | 3 | 95 | 6 | TMEM171 | transmembrane protein 171 | Other | other | |
| chr13 | *LINC00382* | 3 | 89 | 6 | LINC00382 | long intergenic non-protein coding RNA 382 | Other | other | |
| chr17 | *RP5-1029F21.3* | 3 | 88 | 6 | | | | | |
| chr6 | *OSTCP1* | 3 | 71 | 6 | OSTCP1 | oligosaccharyltransferase complex subunit pseudogene 1 | Other | other | |
| chr12 | *RP11-428G5.5* | 3 | 68 | 6 | | | | | |
| chr16 | *GP2* | 3 | 68 | 6 | GP2 | glycoprotein 2 | Cytoplasm | other | |
| chr11 | *RP11-222N13.1* | 3 | 73 | 6 | | | | | |
| chr11 | *TRIM68* | 3 | 115 | 6 | TRIM68 | tripartite motif containing 68 | Cytoplasm | enzyme | |
| chr15 | *RP11-548M13.1* | 2 | 66 | 6 | | | | | |
| chr19 | *C19orf12* | 3 | 118 | 6 | C19orf12 | chromosome 19 open reading frame 12 | Cytoplasm | other | |
| chr12 | *GAS2L3* | 3 | 56 | 6 | GAS2L3 | growth arrest specific 2 like 3 | Other | other | |
| chr3 | *PPM1M* | 3 | 151 | 6 | PPM1M | protein phosphatase, Mg2+/Mn2+ dependent 1M | Nucleus | phosphatase | |
| chr2 | *FTCDNL1* | 2 | 52 | 6 | FTCDNL1 | formiminotransferase cyclodeaminase N-terminal like | Other | other | |
| chr7 | *TMEM243* | 3 | 85 | 6 | TMEM243 | transmembrane protein 243 | Other | other | |
| chr19 | *CTC-559E9.6* | 3 | 73 | 6 | | | | | |
| chr1 | *CYP4A22-AS1* | 3 | 74 | 6 | CYP4A22-AS1 | CYP4A22 antisense RNA 1 | Other | other | |
| chr3 | *RP11-432B6.3* | 3 | 92 | 6 | | | | | |
| chr2 | *AC016725.4* | 2 | 58 | 6 | | | | | |
| chr2 | *RP11-287D1.3* | 3 | 79 | 6 | | | | | |
| chr14 | *ARHGEF40* | 3 | 119 | 6 | ARHGEF40 | Rho guanine nucleotide exchange factor 40 | Other | other | |
| chr11 | *KDM4D* | 3 | 152 | 6 | KDM4D | lysine demethylase 4D | Nucleus | enzyme | |
| chr22 | *CRELD2* | 3 | 171 | 6 | CRELD2 | cysteine rich with EGF like | Other | other | |
| chr3 | *PLSCR1* | 3 | 104 | 6 | PLSCR1 | phospholipid scramblase 1 | Plasma Membrane | enzyme | |
| chr11 | *MS4A8* | 3 | 76 | 6 | MS4A8 | membrane spanning 4-domains | Plasma Membrane | other | |
| chr10 | *RP11-310E22.5* | 3 | 79 | 6 | | | | | |
| chr16 | *POLR3K* | 3 | 89 | 6 | POLR3K | RNA polymerase III subunit K | Nucleus | transcription regulator | |
| chr19 | *MEX3D* | 3 | 142 | 6 | MEX3D | mex-3 RNA binding family | Nucleus | other | |
| chr11 | *CTD-2655K5.1* | 3 | 107 | 6 | | | | | |
| chr1 | *OPN3* | 3 | 93 | 6 | OPN3 | opsin 3 | Plasma Membrane | G-protein coupled recepto | |
| chr22 | *ST13* | 2 | 48 | 6 | ST13 | ST13 Hsp70 interacting protein | Cytoplasm | other | |
| chr15 | *LMAN1L* | 3 | 123 | 6 | LMAN1L | lectin, mannose binding 1 like | Plasma Membrane | other | |
| chr1 | *RP11-242F24.1* | 3 | 117 | 6 | | | | | |
| chr16 | *COQ7* | 3 | 96 | 6 | COQ7 | coenzyme Q7, hydroxylase | Cytoplasm | enzyme | |
| chr6 | *RTN4IP1* | 3 | 59 | 6 | RTN4IP1 | reticulon 4 interacting protein 1 | Cytoplasm | enzyme | |
| chr5 | *RP11-332J15.3* | 3 | 350 | 6 | | | | | |
| chr3 | *ADPRH* | 3 | 91 | 6 | ADPRH | ADP-ribosylarginine hydrolase | Other | enzyme | |
| chr16 | *CORO1A* | 3 | 206 | 6 | CORO1A | coronin 1A | Cytoplasm | other | |
| chr6 | *LINC00240* | 3 | 63 | 6 | LINC00240 | long intergenic non-protein coding RNA 240 | Other | other | |
| chr1 | *PRDX1* | 3 | 87 | 6 | PRDX1 | peroxiredoxin 1 | Cytoplasm | enzyme | |
| chr19 | *ZNF121* | 3 | 71 | 6 | ZNF121 | zinc finger protein 121 | Nucleus | transcription regulator | |
| chrX | *CT45A4* | 3 | 108 | 6 | 45A1 (includes oth | cancer/testis antigen family 45 member A2 | Other | other | |
| chr11 | *CTSF* | 3 | 245 | 6 | CTSF | cathepsin F | Cytoplasm | peptidase | |
| chr15 | *RP11-138H8.3* | 2 | 58 | 6 | | | | | |
| chr4 | *SEC24D* | 2 | 57 | 6 | SEC24D | SEC24 homolog D, COPII coat complex component | Cytoplasm | transporter | |
| chr2 | *AC010880.1* | 2 | 62 | 6 | | | | | |
| chr11 | *NPAS4* | 3 | 241 | 6 | NPAS4 | neuronal PAS domain protein 4 | Nucleus | transcription regulator | |
| chr1 | *RP11-66D17.3* | 3 | 140 | 6 | | | | | |
| chr9 | *C9orf153* | 4 | 64 | 6 | C9orf153 | chromosome 9 open reading frame 153 | Other | other | |
| chr14 | *RP11-671J11.4* | 2 | 79 | 6 | | | | | |
| chr8 | *FER1L6* | 3 | 62 | 6 | FER1L6 | fer-1 like family member 6 | Cytoplasm | other | |
| chr20 | *LINC00261* | 3 | 97 | 6 | LINC00261 | long intergenic non-protein coding RNA 261 | Other | other | |
| chr17 | *AC022596.6* | 3 | 134 | 6 | | | | | |
| chr6 | *RHAG* | 3 | 70 | 6 | RHAG | Rh associated glycoprotein | Plasma Membrane | peptidase | |
| chr16 | *CTF2P* | 3 | 146 | 6 | CTF2P | cardiotrophin 2, pseudogene | Other | other | |
| chr8 | *RP11-10A14.5* | 3 | 57 | 6 | | | | | |
| chr2 | *DGUOK-AS1* | 3 | 89 | 6 | DGUOK-AS1 | DGUOK antisense RNA 1 | Other | other | |
| chr3 | *RP11-204C23.1* | 2 | 53 | 6 | | | | | |
| chr1 | *F3* | 3 | 69 | 6 | F3 | coagulation factor III, tissue factor | Plasma Membrane | transmembrane receptor | tisotumab vedotin, activated recombinant human |
| chr10 | *LIPK* | 3 | 59 | 6 | LIPK | lipase family member K | Extracellular Space | enzyme | |
| chr1 | *UTP11L* | 3 | 81 | 6 | UTP11 | UTP11 small subunit processome component | Nucleus | other | |

| | | | | | | | | | |
|---|---|---|---|---|---|---|---|---|---|
| chr16 | *SNN* | 3 | 181 | 6 | SNN | stannin | Plasma Membrane | other | |
| chr12 | *LRIG3* | 2 | 64 | 6 | LRIG3 | leucine rich repeats and immunoglobulin like domains 3 | Extracellular Space | other | |
| chr17 | *ZMYND15* | 3 | 152 | 6 | ZMYND15 | zinc finger MYND-type containing | Other | other | |
| chr10 | *GPAM* | 3 | 79 | 6 | GPAM | glycerol-3-phosphate acyltransferase, mitochondrial | Cytoplasm | enzyme | |
| chr14 | *FLVCR2* | 3 | 67 | 6 | FLVCR2 | feline leukemia virus subgroup C cellular receptor family member 2 | Plasma Membrane | transporter | |
| chr9 | *STOM* | 3 | 68 | 6 | STOM | stomatin | Plasma Membrane | other | |
| chr9 | *UBQLN1* | 3 | 67 | 6 | UBQLN1 | ubiquilin 1 | Cytoplasm | other | |
| chr16 | *WASH4P* | 2 | 165 | 6 | WASH4P | WASP family homolog 4, pseudogene | Other | other | |
| chr17 | *AC003665.1* | 3 | 70 | 6 | | | | | |
| chr17 | *RP11-316M20.1* | 3 | 103 | 6 | | | | | |
| chr22 | *AC008079.9* | 3 | 154 | 6 | | | | | |
| chr11 | *C11orf53* | 3 | 78 | 6 | C11orf53 | chromosome 11 open reading frame 53 | Other | other | |
| chr17 | *MIR22HG* | 3 | 192 | 6 | MIR22HG | MIR22 host gene | Other | other | |
| chr8 | *LAPTM4B* | 3 | 54 | 6 | LAPTM4B | lysosomal protein transmembrane 4 beta | Cytoplasm | other | |
| chr15 | *RP11-752G15.9* | 3 | 109 | 6 | | | | | |
| chr19 | *OSCAR* | 3 | 112 | 6 | OSCAR | osteoclast associated Ig-like | Plasma Membrane | other | |
| chr2 | *COBLL1* | 2 | 63 | 6 | COBLL1 | cordon-bleu WH2 repeat protein like 1 | Extracellular Space | other | |
| chr19 | *DOHH* | 3 | 149 | 6 | DOHH | deoxyhypusine hydroxylase | Cytoplasm | enzyme | |
| chr11 | *SBF2-AS1* | 3 | 107 | 6 | SBF2-AS1 | SBF2 antisense RNA 1 | Other | other | |
| chr1 | *C1orf204* | 3 | 97 | 6 | SNHG28 | small nucleolar RNA host gene | Other | other | |
| chr6 | *DOPEY1* | 2 | 47 | 6 | DOP1A | DOP1 leucine zipper like protein | Cytoplasm | other | |
| chr3 | *ACTR8* | 3 | 62 | 6 | ACTR8 | actin related protein 8 | Nucleus | other | |
| chr19 | *DAPK3* | 3 | 176 | 6 | DAPK3 | death associated protein kinase 3 | Cytoplasm | kinase | |
| chr14 | *RP11-1085N6.3* | 3 | 84 | 6 | | | | | |
| chr2 | *IL36RN* | 3 | 154 | 6 | IL36RN | interleukin 36 receptor antagonist | Extracellular Space | cytokine | |
| chr12 | *RP11-408B11.2* | 2 | 70 | 6 | | | | | |
| chr7 | *MNX1* | 3 | 114 | 6 | MNX1 | motor neuron and pancreas homeobox 1 | Nucleus | transcription regulator | |
| chr18 | *SLMO1* | 3 | 117 | 6 | PRELID3A | PRELI domain containing 3A | Cytoplasm | transporter | |
| chr16 | *GNPTG* | 3 | 162 | 6 | GNPTG | N-acetylglucosamine-1-phosphate transferase subunit | Cytoplasm | enzyme | |
| chr17 | *MYH13* | 3 | 65 | 6 | MYH13 | myosin heavy chain 13 | Cytoplasm | other | |
| chr17 | *AC087393.1* | 3 | 73 | 6 | | | | | |
| chr11 | *CD3E* | 3 | 83 | 6 | CD3E | CD3e molecule | Plasma Membrane | transmembrane receptor | muromonab-CD3, blinatumomab |
| chr8 | *RP11-594N15.2* | 3 | 68 | 6 | | | | | |
| chr12 | *KRT4* | 3 | 179 | 6 | KRT4 | keratin 4 | Cytoplasm | other | |
| chrX | *OCRL* | 2 | 50 | 6 | OCRL | OCRL inositol polyphosphate-5-phosphatase | Cytoplasm | phosphatase | |
| chr8 | *FAM90A17P* | 3 | 554 | 6 | | | | | |
| chr19 | *ANKLE1* | 3 | 122 | 6 | ANKLE1 | ankyrin repeat and LEM domain containing 1 | Cytoplasm | enzyme | |
| chr2 | *C2orf80* | 2 | 48 | 6 | C2orf80 | chromosome 2 open reading frame 80 | Other | other | |
| chr1 | *S100A13* | 3 | 132 | 6 | S100A13 | S100 calcium binding protein A13 | Cytoplasm | other | |
| chr3 | *PDE12* | 2 | 104 | 6 | PDE12 | phosphodiesterase 12 | Cytoplasm | enzyme | |
| chr2 | *NTSR2* | 3 | 96 | 6 | NTSR2 | neurotensin receptor 2 | Plasma Membrane | G-protein coupled receptor | levocabastine |
| chr19 | *FAM98C* | 3 | 110 | 6 | FAM98C | family with sequence similarity 98 member C | Other | other | |
| chr10 | *SLC39A12* | 3 | 62 | 6 | SLC39A12 | solute carrier family 39 member | Plasma Membrane | transporter | |
| chr4 | *OCIAD1* | 3 | 71 | 6 | OCIAD1 | OCIA domain containing 1 | Cytoplasm | other | |
| chr6 | *REPS1* | 2 | 46 | 6 | REPS1 | RALBP1 associated Eps domain containing 1 | Plasma Membrane | other | |
| chr4 | *SLBP* | 3 | 89 | 6 | SLBP | stem-loop binding protein | Nucleus | other | |
| chr22 | *TST* | 3 | 110 | 6 | TST | thiosulfate sulfurtransferase | Cytoplasm | enzyme | |
| chr1 | *RP11-451O13.1* | 3 | 113 | 6 | | | | | |
| chr18 | *RP11-739N10.1* | 3 | 69 | 6 | | | | | |
| chr11 | *TYR* | 3 | 115 | 6 | TYR | tyrosinase | Cytoplasm | enzyme | azelaic acid, hydroquinone |
| chr12 | *CREBL2* | 3 | 47 | 6 | CREBL2 | cAMP responsive element binding protein like 2 | Nucleus | transcription regulator | |
| chr12 | *AGAP2* | 3 | 128 | 6 | AGAP2 | ArfGAP with GTPase domain, ankyrin repeat and PH domain 2 | Nucleus | enzyme | |
| chr3 | *AMT* | 3 | 108 | 6 | AMT | aminomethyltransferase | Cytoplasm | enzyme | |
| chr9 | *LCN1* | 3 | 205 | 6 | LCN1 | lipocalin 1 | Extracellular Space | transporter | |
| chr2 | *TGOLN2* | 3 | 98 | 6 | TGOLN2 | trans-golgi network protein 2 | Cytoplasm | other | |
| chr6 | *IMPG1* | 2 | 46 | 6 | IMPG1 | interphotoreceptor matrix proteoglycan 1 | Extracellular Space | other | |
| chr3 | *DCBLD2* | 2 | 53 | 6 | DCBLD2 | discoidin, CUB and LCCL domain containing 2 | Plasma Membrane | other | |
| chr16 | *PRSS54* | 3 | 108 | 6 | PRSS54 | serine protease 54 | Cytoplasm | other | |
| chr2 | *AC009237.8* | 3 | 77 | 6 | | | | | |
| chr11 | *GAL3ST3* | 3 | 199 | 6 | GAL3ST3 | galactose-3-O-sulfotransferase 3 | Cytoplasm | enzyme | |
| chr1 | *NPHS2* | 3 | 91 | 6 | NPHS2 | NPHS2 stomatin family member, podocin | Plasma Membrane | other | |
| chr19 | *SYCE2* | 3 | 82 | 6 | SYCE2 | synaptonemal complex central element protein 2 | Nucleus | other | |
| chr13 | *COMMD6* | 3 | 102 | 6 | COMMD6 | COMM domain containing 6 | Other | other | |
| chr6 | *MEP1A* | 2 | 66 | 6 | MEP1A | meprin A subunit alpha | Plasma Membrane | peptidase | |
| chrX | *CLIC2* | 2 | 62 | 6 | CLIC2 | chloride intracellular channel 2 | Cytoplasm | enzyme | |
| chr17 | *CTD-2008P7.3* | 3 | 68 | 6 | | | | | |
| chr1 | *GSTM5* | 3 | 150 | 6 | GSTM5 | glutathione S-transferase mu 5 | Cytoplasm | enzyme | |

| | | | | | | | | | |
|---|---|---|---|---|---|---|---|---|---|
| chr2 | *ACVR1C* | 3 | 85 | 6 | ACVR1C | activin A receptor type 1C | Plasma Membrane | kinase | |
| chrX | *SATL1* | 3 | 65 | 6 | SATL1 | spermidine/spermine N1-acetyl transferase like 1 | Other | other | |
| chr10 | *TUBB8* | 3 | 133 | 6 | TUBB8 | tubulin beta 8 class VIII | Cytoplasm | other | |
| chr15 | *USP3* | 2 | 49 | 6 | USP3 | ubiquitin specific peptidase 3 | Cytoplasm | peptidase | |
| chr17 | *NKIRAS2* | 3 | 115 | 6 | NKIRAS2 | NFKB inhibitor interacting Ras | Cytoplasm | enzyme | |
| chr16 | *CBLN1* | 3 | 152 | 6 | CBLN1 | cerebellin 1 precursor | Cytoplasm | other | |
| chr1 | *RP11-12M5.4* | 3 | 84 | 6 | | | | | |
| chr19 | *FFAR2* | 3 | 101 | 6 | FFAR2 | free fatty acid receptor 2 | Plasma Membrane | G-protein coupled recepto | |
| chr2 | *GGCX* | 3 | 68 | 6 | GGCX | gamma-glutamyl carboxylase | Cytoplasm | enzyme | vitamin K3, |
| chr6 | *C6orf141* | 2 | 55 | 6 | C6orf141 | chromosome 6 open reading frame 141 | Other | other | |
| chr19 | *AC004076.9* | 3 | 69 | 6 | | | | | |
| chr3 | *RASSF1* | 3 | 116 | 6 | RASSF1 | Ras association domain family member 1 | Nucleus | other | |
| chr7 | *C7orf73* | 2 | 87 | 6 | STMP1 | short transmembrane mitochondrial protein 1 | Other | other | |
| chr7 | *AASS* | 2 | 82 | 6 | AASS | aminoadipate-semialdehyde synthase | Cytoplasm | enzyme | |
| chr19 | *ZNF865* | 3 | 106 | 6 | ZNF865 | zinc finger protein 865 | Other | other | |
| chr11 | *FOLR2* | 3 | 102 | 6 | FOLR2 | folate receptor beta | Plasma Membrane | transporter | |
| chr14 | *C14orf93* | 2 | 48 | 6 | C14orf93 | chromosome 14 open reading frame 93 | Other | other | |
| chr8 | *RP11-875O11.2* | 3 | 64 | 6 | | | | | |
| chr9 | *PTPDC1* | 3 | 79 | 6 | PTPDC1 | protein tyrosine phosphatase domain containing 1 | Extracellular Space | phosphatase | |
| chr1 | *GALE* | 3 | 180 | 6 | GALE | UDP-galactose-4-epimerase | Cytoplasm | enzyme | |
| chr8 | *ZNF705G* | 3 | 56 | 6 | ZNF705G | zinc finger protein 705G | Other | other | |
| chr6 | *WASF1* | 2 | 67 | 6 | WASF1 | WASP family member 1 | Nucleus | other | |
| chr7 | *RP4-604G5.1* | 3 | 102 | 6 | | | | | |
| chr19 | *FUT1* | 3 | 129 | 6 | FUT1 | fucosyltransferase 1 (H blood | Cytoplasm | enzyme | |
| chr7 | *EPO* | 3 | 110 | 6 | EPO | erythropoietin | Extracellular Space | cytokine | |
| chr9 | *FAM225A* | 2 | 95 | 6 | FAM225A | family with sequence similarity 225 member A | Other | other | |
| chr1 | *AMPD1* | 2 | 65 | 6 | AMPD1 | adenosine monophosphate deaminase 1 | Cytoplasm | enzyme | |
| chr11 | *RP11-804A23.1* | 3 | 98 | 6 | | | | | |
| chr11 | *CTD-2523D13.2* | 3 | 96 | 6 | | | | | |
| chr10 | *RP13-137A17.6* | 3 | 169 | 6 | | | | | |
| chr3 | *NGLY1* | 3 | 49 | 6 | NGLY1 | N-glycanase 1 | Cytoplasm | enzyme | |
| chr2 | *CNRIP1* | 3 | 55 | 6 | CNRIP1 | cannabinoid receptor interacting protein 1 | Extracellular Space | other | |
| chr7 | *AC095067.1* | 3 | 133 | 6 | | | | | |
| chr5 | *RP11-43F13.3* | 3 | 259 | 6 | | | | | |
| chr15 | *PEX11A* | 3 | 96 | 6 | PEX11A | peroxisomal biogenesis factor 11 alpha | Cytoplasm | other | |
| chr1 | *GJC2* | 3 | 180 | 6 | GJC2 | gap junction protein gamma 2 | Plasma Membrane | transporter | |
| chr10 | *RGR* | 3 | 100 | 6 | RGR | retinal G protein coupled receptor | Plasma Membrane | G-protein coupled recepto | |
| chr2 | *AC096554.1* | 2 | 107 | 6 | | | | | |
| chr17 | *AL353997.3* | 3 | 63 | 6 | | | | | |
| chr11 | *PHOX2A* | 3 | 169 | 6 | PHOX2A | paired like homeobox 2A | Nucleus | transcription regulator | |
| chr10 | *AKR1C1* | 2 | 67 | 6 | AKR1C1/AKR1C2 | aldo-keto reductase family 1 member C2 | Cytoplasm | enzyme | |
| chr19 | *CTB-180A7.3* | 3 | 81 | 6 | | | | | |
| chrX | *GAGE12D* | 2 | 91 | 6 | GE1 (includes oth | G antigen 1 | Other | other | |
| chr22 | *CBY1* | 3 | 51 | 6 | CBY1 | chibby family member 1, beta catenin antagonist | Cytoplasm | other | |
| chr12 | *GRASP* | 3 | 264 | 6 | GRASP | general receptor for phosphoinositides 1 associated scaffold protein | Plasma Membrane | other | |
| chr13 | *RNF17* | 2 | 58 | 6 | RNF17 | ring finger protein 17 | Cytoplasm | other | |
| chr14 | *OTX2-AS1* | 3 | 46 | 6 | OTX2-AS1 | OTX2 antisense RNA 1 (head to head) | Other | other | |
| chr1 | *PARS2* | 3 | 125 | 6 | PARS2 | prolyl-tRNA synthetase 2, mitochondrial | Cytoplasm | enzyme | |
| chr2 | *EMX1* | 3 | 133 | 6 | EMX1 | empty spiracles homeobox 1 | Nucleus | transcription regulator | |
| chrX | *RP1-30E17.2* | 3 | 67 | 6 | | | | | |
| chr17 | *RDM1* | 2 | 51 | 6 | RDM1 | RAD52 motif containing 1 | Nucleus | other | |
| chr7 | *SLC29A4P2* | 3 | 154 | 6 | C29A4P1/SLC29A | solute carrier family 29 member 4 pseudogene 1 | Other | other | |
| chr16 | *CDK10* | 3 | 132 | 6 | CDK10 | cyclin dependent kinase 10 | Nucleus | kinase | alvocidib |
| chr19 | *ZNF540* | 2 | 53 | 6 | ZNF540 | zinc finger protein 540 | Cytoplasm | translation regulator | |
| chr18 | *RIT2* | 2 | 46 | 6 | RIT2 | Ras like without CAAX 2 | Plasma Membrane | enzyme | |
| chrX | *RP11-706O15.3* | 2 | 53 | 6 | | | | | |
| chr7 | *NSUN5P1* | 3 | 128 | 6 | NSUN5P1 | NSUN5 pseudogene 1 | Other | other | |
| chr1 | *FBXO44* | 3 | 138 | 6 | FBXO44 | F-box protein 44 | Cytoplasm | enzyme | |
| chr10 | *FUT11* | 3 | 153 | 6 | FUT11 | fucosyltransferase 11 | Cytoplasm | enzyme | |
| chr16 | *RP11-305A4.3* | 3 | 110 | 6 | | | | | |
| chr10 | *C10orf53* | 3 | 90 | 6 | C10orf53 | chromosome 10 open reading frame 53 | Other | other | |
| chr20 | *FAM83C* | 3 | 169 | 6 | FAM83C | family with sequence similarity 83 member C | Other | other | |
| chr18 | *RIOK3* | 3 | 76 | 6 | RIOK3 | RIO kinase 3 | Cytoplasm | kinase | |
| chr12 | *C12orf80* | 3 | 210 | 6 | C12orf80 | chromosome 12 open reading frame 80 | Other | other | |
| chr11 | *BUD13* | 3 | 69 | 6 | BUD13 | BUD13 homolog | Nucleus | other | |
| chr17 | *MKS1* | 3 | 90 | 6 | MKS1 | MKS transition zone complex subunit 1 | Cytoplasm | other | |
| chr2 | *ZFAND2B* | 3 | 64 | 6 | ZFAND2B | zinc finger AN1-type containing | Cytoplasm | other | |

| | | | | | | | | | |
|---|---|---|---|---|---|---|---|---|---|
| chr8 | *TRIM55* | 3 | 62 | 6 | TRIM55 | tripartite motif containing 55 | Cytoplasm | other | |
| chr19 | *ZNF264* | 3 | 65 | 6 | ZNF264 | zinc finger protein 264 | Nucleus | transcription regulator | |
| chr7 | *BUD31* | 3 | 84 | 6 | BUD31 | BUD31 homolog | Nucleus | transcription regulator | |
| chr20 | *GFRA4* | 2 | 100 | 6 | GFRA4 | GDNF family receptor alpha 4 | Plasma Membrane | transmembrane recepto | |
| chr1 | *PROK1* | 3 | 106 | 6 | PROK1 | prokineticin 1 | Extracellular Space | growth factor | |
| chr19 | *ZNF528* | 3 | 59 | 6 | ZNF528 | zinc finger protein 528 | Other | other | |
| chr8 | *ZFPM2* | 2 | 49 | 6 | ZFPM2 | zinc finger protein, FOG family member 2 | Nucleus | transcription regulator | |
| chr15 | *RCCD1* | 3 | 171 | 6 | RCCD1 | RCC1 domain containing 1 | Other | other | |
| chr21 | *MX1* | 3 | 61 | 6 | MX1 | MX dynamin like GTPase 1 | Cytoplasm | enzyme | |
| chr2 | *OTX1* | 3 | 155 | 6 | OTX1 | orthodenticle homeobox 1 | Nucleus | transcription regulator | |
| chr19 | *CLEC4M* | 3 | 107 | 6 | CLEC4M | C-type lectin domain family 4 member M | Plasma Membrane | other | |
| chr1 | *UBE2U* | 3 | 50 | 6 | UBE2U | ubiquitin conjugating enzyme E2 | Other | enzyme | |
| chr16 | *MAP1LC3B* | 2 | 87 | 6 | MAP1LC3B | microtubule associated protein 1 light chain 3 beta | Cytoplasm | other | |
| chr19 | *C5AR2* | 3 | 76 | 6 | C5AR2 | complement component 5a receptor 2 | Plasma Membrane | -protein coupled recepto | |
| chr16 | *CCDC64B* | 3 | 177 | 6 | BICDL2 | BICD family like cargo adaptor 2 | Other | other | |
| chr2 | *RP11-762E8.1* | 3 | 72 | 6 | | | | | |
| chr16 | *NARFL* | 3 | 163 | 6 | CIAO3 | cytosolic iron-sulfur assembly component 3 | Cytoplasm | other | |
| chr8 | *THEM6* | 3 | 122 | 6 | THEM6 | thioesterase superfamily member | Other | other | |
| chr10 | *ATP5C1* | 2 | 73 | 6 | ATP5F1C | ATP synthase F1 subunit gamma | Cytoplasm | transporter | |
| chr18 | *TYMS* | 3 | 65 | 6 | TYMS | thymidylate synthetase | Nucleus | enzyme | pemetrexed, capecitabine/temozolomide, cyclophosphamide/epirubicin/5-fluorouracil/paclitaxel, BGC945, 5-fluorouracil/irinotecan/oxaliplatin, capecitabine/cetuximab/irinotecan, tegafur, capecitabine/irinotecan/oxaliplatin, trimethoprim, docetaxel/5-fluorouracil, cetuximab/gimeracil/irinotecan/oxonic acid/tegafur, floxuridine, gimeracil/irinotecan/oxonic acid/tegafur, [illegible] |
| chr4 | *TMEM192* | 2 | 57 | 6 | TMEM192 | transmembrane protein 192 | Cytoplasm | other | |
| chr10 | *DCLRE1C* | 3 | 51 | 6 | DCLRE1C | DNA cross-link repair 1C | Nucleus | enzyme | |
| chrX | *IDH3G* | 3 | 162 | 6 | IDH3G | isocitrate dehydrogenase (NAD(+)) 3 non-catalytic subunit | Cytoplasm | enzyme | |
| chr12 | *RP1-46F2.2* | 3 | 66 | 6 | | | | | |
| chr17 | *AC069363.1* | 3 | 74 | 6 | | | | | |
| chr7 | *AC005682.5* | 3 | 113 | 6 | | | | | |
| chr15 | *MTFMT* | 2 | 51 | 6 | MTFMT | mitochondrial methionyl-tRNA formyltransferase | Cytoplasm | enzyme | |
| chr15 | *RP11-152F13.3* | 3 | 80 | 6 | | | | | |
| chr12 | *RP11-818F20.4* | 3 | 60 | 6 | | | | | |
| chr19 | *FPR3* | 3 | 60 | 6 | FPR3 | formyl peptide receptor 3 | Plasma Membrane | other | |
| chr1 | *LINC00303* | 3 | 97 | 6 | LINC00303 | long intergenic non-protein coding RNA 303 | Other | other | |
| chr10 | *ZMYND11* | 2 | 54 | 6 | ZMYND11 | zinc finger MYND-type containing | Nucleus | transcription regulator | |
| chr17 | *FNDC8* | 3 | 86 | 6 | FNDC8 | fibronectin type III domain containing 8 | Nucleus | other | |
| chr3 | *NBPF21P* | 3 | 67 | 6 | NBPF21P | NBPF member 21, pseudogene | Other | other | |
| chr13 | *LRCH1* | 2 | 49 | 6 | LRCH1 | leucine rich repeats and calponin homology domain containing 1 | Cytoplasm | other | |
| chr11 | *FBXO3* | 3 | 71 | 6 | FBXO3 | F-box protein 3 | Cytoplasm | enzyme | |
| chr18 | *DOK6* | 3 | 58 | 6 | DOK6 | docking protein 6 | Cytoplasm | other | |
| chr19 | *PRPF31* | 3 | 147 | 6 | PRPF31 | pre-mRNA processing factor 31 | Nucleus | other | |
| chr2 | *MATN3* | 3 | 67 | 6 | MATN3 | matrilin 3 | Extracellular Space | other | |
| chr8 | *ATP6V1B2* | 3 | 72 | 6 | ATP6V1B2 | ATPase H+ transporting V1 subunit B2 | Cytoplasm | transporter | gallium nitrate |
| chr19 | *LYPD3* | 3 | 184 | 6 | LYPD3 | LY6/PLAUR domain containing 3 | Plasma Membrane | other | |
| chr12 | *RP11-1038A11.3* | 3 | 86 | 6 | | | | | |
| chr1 | *S100A9* | 2 | 161 | 6 | S100A9 | S100 calcium binding protein A9 | Cytoplasm | other | |
| chr5 | *CTD-3007L5.1* | 2 | 153 | 6 | | | | | |
| chr11 | *ROBO4* | 3 | 118 | 6 | ROBO4 | roundabout guidance receptor 4 | Plasma Membrane | other | |
| chr9 | *RP11-344B5.2* | 3 | 143 | 6 | | | | | |
| chr2 | *STK25* | 3 | 131 | 6 | STK25 | serine/threonine kinase 25 | Cytoplasm | kinase | |
| chr19 | *ZNF780B* | 2 | 70 | 6 | ZNF780B | zinc finger protein 780B | Other | other | |
| chr14 | *AMN* | 3 | 99 | 6 | AMN | amnion associated transmembrane protein | Plasma Membrane | other | |
| chr1 | *RP5-1070A16.1* | 2 | 59 | 6 | | | | | |
| chr1 | *METTL13* | 4 | 150 | 6 | EEF1AKNMT | eEF1A lysine and N-terminal methyltransferase | Other | enzyme | |
| chrX | *GS1-600G8.5* | 3 | 87 | 6 | | | | | |

| | | | | | | | | | |
|---|---|---|---|---|---|---|---|---|---|
| chr9 | C9orf24 | 3 | 92 | 6 | C9orf24 | chromosome 9 open reading frame 24 | Cytoplasm | other | |
| chr7 | LINC00996 | 3 | 95 | 6 | LINC00996 | long intergenic non-protein coding RNA 996 | Other | other | |
| chr6 | TUBB2B | 3 | 105 | 6 | TUBB2B | tubulin beta 2B class IIb | Cytoplasm | other | |
| chr19 | SIGLEC12 | 3 | 121 | 6 | SIGLEC12 | sialic acid binding Ig like lectin 12 (gene/pseudogene) | Other | other | |
| chr17 | POLDIP2 | 3 | 109 | 6 | POLDIP2 | DNA polymerase delta interacting protein 2 | Cytoplasm | other | |
| chr4 | SMIM20 | 2 | 60 | 6 | SMIM20 | small integral membrane protein | Cytoplasm | other | |
| chr19 | COX6B1 | 3 | 94 | 6 | COX6B1 | cytochrome c oxidase subunit | Cytoplasm | enzyme | |
| chr4 | RP11-586D19.1 | 3 | 109 | 6 | | | | | |
| chr17 | ZSWIM5P2 | 2 | 94 | 6 | ZSWIM5P2 | zinc finger SWIM-type containing 5 pseudogene 2 | Other | other | |
| chr17 | KRT31 | 3 | 93 | 6 | KRT31 | keratin 31 | Cytoplasm | other | |
| chr22 | C22orf23 | 3 | 76 | 6 | C22orf23 | chromosome 22 open reading frame 23 | Other | other | |
| chr10 | PNLIPRP2 | 3 | 99 | 6 | PNLIPRP2 | pancreatic lipase related protein 2 (gene/pseudogene) | Extracellular Space | enzyme | |
| chr17 | LINC00854 | 3 | 1102 | 6 | CCDC200 | coiled-coil domain containing 200 | Other | other | |
| chrX | GDI1 | 3 | 102 | 6 | GDI1 | GDP dissociation inhibitor 1 | Cytoplasm | other | |
| chr19 | CTD-2587H24.10 | 3 | 78 | 6 | | | | | |
| chr14 | BMP4 | 3 | 107 | 6 | BMP4 | bone morphogenetic protein 4 | Extracellular Space | growth factor | |
| chr9 | RP11-764K9.4 | 3 | 141 | 6 | | | | | |
| chr3 | SELK | 3 | 84 | 6 | | | | | |
| chr9 | ZNF462 | 3 | 79 | 6 | ZNF462 | zinc finger protein 462 | Nucleus | transcription regulator | |
| chr20 | PAK7 | 3 | 67 | 6 | PAK5 | p21 (RAC1) activated kinase 5 | Nucleus | kinase | |
| chr20 | CPXM1 | 3 | 131 | 6 | CPXM1 | carboxypeptidase X, M14 family member 1 | Extracellular Space | peptidase | |
| chr11 | RP11-35J10.5 | 3 | 122 | 6 | | | | | |
| chr1 | RP11-810H18.1 | 3 | 97 | 6 | | | | | |
| chr1 | MXRA8 | 3 | 88 | 6 | MXRA8 | matrix remodeling associated 8 | Cytoplasm | other | |
| chr6 | RP11-506B6.6 | 2 | 45 | 6 | | | | | |
| chr11 | RP11-467L20.10 | 3 | 169 | 6 | | | | | |
| chr11 | FAM86C1 | 3 | 82 | 6 | FAM86C1 | family with sequence similarity 86 member C1 | Other | other | |
| chr12 | KRT2 | 3 | 156 | 6 | KRT2 | keratin 2 | Cytoplasm | other | CIGB-300, |
| chr12 | CHPT1 | 3 | 62 | 6 | CHPT1 | choline phosphotransferase 1 | Cytoplasm | enzyme | |
| chr16 | RP11-719K4.6 | 3 | 108 | 6 | | | | | |
| chr5 | CTB-43E15.1 | 3 | 87 | 6 | | | | | |
| chr3 | ILDR1 | 3 | 116 | 6 | ILDR1 | immunoglobulin like domain containing receptor 1 | Plasma Membrane | transmembrane recepto | |
| chr8 | RP11-115J16.2 | 3 | 57 | 6 | | | | | |
| chrX | FAM122B | 2 | 56 | 6 | FAM122B | family with sequence similarity | Other | other | |
| chr6 | PPIL4 | 2 | 45 | 6 | PPIL4 | peptidylprolyl isomerase like 4 | Nucleus | enzyme | |
| chr17 | RP11-311F12.1 | 3 | 116 | 6 | | | | | |
| chr11 | CHRNA10 | 3 | 224 | 6 | CHRNA10 | cholinergic receptor nicotinic alpha 10 subunit | Plasma Membrane | transmembrane recepto | succinylcholine, rapacuronium, doxacurium, vecuronium, pipecuronium, metocurine, mivacurium, D-tubocurarine, nicotine, arecoline, pancuronium, atracurium, isoflurane, cisatracurium, rocuronium, mecamylamine, enflurane, ABT-089, methadone |
| chr9 | DEC1; | 3 | 71 | 6 | | | | | |
| chr19 | BEST2 | 3 | 142 | 6 | BEST2 | bestrophin 2 | Plasma Membrane | ion channel | |
| chr19 | AC007795.1 | 3 | 90 | 6 | | | | | |
| chr5 | ESM1 | 3 | 80 | 6 | ESM1 | endothelial cell specific molecule | Extracellular Space | growth factor | |
| chr17 | RP11-515E23.1 | 3 | 93 | 6 | | | | | |
| chr5 | IRX4 | 3 | 188 | 6 | IRX4 | iroquois homeobox 4 | Nucleus | transcription regulator | |
| chr19 | C19orf40 | 3 | 83 | 6 | FAAP24 | FA core complex associated protein 24 | Nucleus | other | |
| chr15 | RP13-996F3.4 | 2 | 116 | 6 | | | | | |
| chr10 | RP11-526P5.2 | 3 | 118 | 6 | | | | | |
| chr3 | KCNMB2 | 3 | 47 | 6 | KCNMB2 | potassium calcium-activated channel subfamily M regulatory | Plasma Membrane | ion channel | tedisamil |
| chr17 | AC022596.2 | 2 | 121 | 6 | | | | | |
| chrX | MAGEA9 | 3 | 101 | 6 | MAGEA9/MAGEA9 | MAGE family member A9 | Other | other | |
| chr11 | AP000439.5 | 3 | 80 | 6 | | | | | |
| chr8 | DCAF13 | 2 | 48 | 6 | DCAF13 | DDB1 and CUL4 associated factor 13 | Nucleus | other | |
| chr1 | ATPIF1 | 3 | 107 | 6 | ATP5IF1 | ATP synthase inhibitory factor subunit 1 | Cytoplasm | other | |
| chr4 | RP11-363G15.2 | 3 | 75 | 6 | | | | | |
| chr4 | RP11-722M1.1 | 2 | 52 | 6 | | | | | |
| chr11 | AP003062.1 | 2 | 114 | 6 | | | | | |
| chr9 | UCK1 | 3 | 203 | 6 | UCK1 | uridine-cytidine kinase 1 | Cytoplasm | kinase | |
| chr2 | AC016682.1 | 2 | 50 | 6 | | | | | |
| chr1 | VAMP3 | 3 | 76 | 6 | VAMP3 | vesicle associated membrane protein 3 | Plasma Membrane | other | |

| | | | | | | | | |
|---|---|---|---|---|---|---|---|---|
| chr9 | *FKBP15* | 2 | 53 | 6 | FKBP15 | FKBP prolyl isomerase 15 | Plasma Membrane | enzyme |
| chr4 | *ATP10D* | 2 | 48 | 6 | ATP10D | ATPase phospholipid transporting 10D (putative) | Cytoplasm | transporter |
| chr3 | *H1FOO* | 3 | 225 | 6 | H1-8 | H1.8 linker histone | Nucleus | other |
| chr3 | *ZDHHC23* | 3 | 111 | 6 | ZDHHC23 | zinc finger DHHC-type containing | Other | other |
| chr12 | *KRT73* | 3 | 204 | 6 | KRT73 | keratin 73 | Extracellular Space | other |
| chr1 | *SHISA4* | 3 | 159 | 6 | SHISA4 | shisa family member 4 | Other | other |
| chr16 | *CTD-2547E10.3* | 3 | 134 | 6 | | | | |
| chr14 | *KIF26A* | 3 | 136 | 6 | KIF26A | kinesin family member 26A | Cytoplasm | other |
| chr19 | *CNN2* | 3 | 165 | 6 | CNN2 | calponin 2 | Cytoplasm | other |
| chr7 | *NPSR1* | 3 | 73 | 6 | NPSR1 | neuropeptide S receptor 1 | Plasma Membrane | G-protein coupled recepto |
| chr3 | *OXNAD1* | 3 | 90 | 6 | OXNAD1 | oxidoreductase NAD binding domain containing 1 | Cytoplasm | transporter |
| chr1 | *LINC00339* | 3 | 74 | 6 | LINC00339 | long intergenic non-protein coding RNA 339 | Other | other |
| chr8 | *RAB11FIP1* | 3 | 70 | 6 | RAB11FIP1 | RAB11 family interacting protein | Cytoplasm | other |
| chrX | *PLXNA3* | 3 | 155 | 6 | PLXNA3 | plexin A3 | Plasma Membrane | transmembrane receptor |
| chr5 | *HMMR* | 3 | 69 | 6 | HMMR | hyaluronan mediated motility | Plasma Membrane | transmembrane receptor |
| chr10 | *RP11-462L8.1* | 3 | 64 | 6 | | | | |
| chr10 | *VENTX* | 3 | 86 | 6 | VENTX | VENT homeobox | Nucleus | transcription regulator |
| chr7 | *SBDSP1* | 2 | 58 | 6 | SBDSP1 | SBDS pseudogene 1 | Other | other |
| chr4 | *RP11-503N18.3* | 3 | 125 | 6 | | | | |
| chr1 | *RP11-488L18.8* | 2 | 83 | 6 | | | | |
| chr1 | *ATPAF1* | 3 | 58 | 6 | ATPAF1 | ATP synthase mitochondrial F1 complex assembly factor 1 | Cytoplasm | other |
| chr7 | *GIMAP8* | 3 | 78 | 6 | GIMAP8 | GTPase, IMAP family member 8 | Other | other |
| chr8 | *SCRT1* | 3 | 132 | 6 | SCRT1 | scratch family transcriptional repressor 1 | Nucleus | transcription regulator |
| chr20 | *WFDC5* | 3 | 158 | 6 | WFDC5 | WAP four-disulfide core domain 5 | Extracellular Space | other |
| chr11 | *RP11-531H8.2* | 3 | 67 | 6 | | | | |
| chr5 | *CTD-2203A3.1* | 3 | 55 | 6 | | | | |
| chr7 | *RP11-310H4.1* | 3 | 103 | 6 | | | | |
| chr19 | *SLC17A7* | 3 | 113 | 6 | SLC17A7 | solute carrier family 17 member 7 | Plasma Membrane | transporter |
| chr21 | *YBEY* | 3 | 99 | 6 | YBEY | ybeY metalloendoribonuclease | Cytoplasm | enzyme |
| chr17 | *TACO1* | 3 | 64 | 6 | TACO1 | translational activator of cytochrome c oxidase I | Cytoplasm | other |
| chr1 | *USP1* | 2 | 73 | 6 | USP1 | ubiquitin specific peptidase 1 | Cytoplasm | peptidase |
| chr19 | *FAM90A27P* | 3 | 110 | 6 | FAM90A27P | family with sequence similarity 90 member A27, pseudogene | Other | other |
| chr1 | *RP11-315I20.1* | 3 | 101 | 6 | | | | |
| chr5 | *GFM2* | 3 | 66 | 6 | GFM2 | G elongation factor mitochondrial | Cytoplasm | translation regulator |
| chr13 | *ZDHHC20* | 2 | 65 | 6 | ZDHHC20 | zinc finger DHHC-type containing | Plasma Membrane | enzyme |
| chr16 | *NDUFAB1* | 3 | 86 | 6 | NDUFAB1 | NADH:ubiquinone oxidoreductase subunit AB1 | Cytoplasm | enzyme |
| chr3 | *C3orf58* | 3 | 73 | 6 | DIPK2A | divergent protein kinase domain | Cytoplasm | other |
| chr3 | *CPOX* | 2 | 43 | 6 | CPOX | coproporphyrinogen oxidase | Cytoplasm | enzyme |
| chr1 | *RP4-781K5.7* | 3 | 165 | 6 | | | | |
| chr3 | *B3GALNT1* | 3 | 55 | 6 | B3GALNT1 | beta-1,3-N-acetylgalactosaminyltransferase 1 (globoside blood group) | Cytoplasm | enzyme |
| chr8 | *FAM83H* | 3 | 144 | 6 | FAM83H | family with sequence similarity 83 member H | Other | other |
| chr14 | *LRP10* | 3 | 122 | 6 | LRP10 | LDL receptor related protein 10 | Plasma Membrane | transmembrane receptor |
| chr1 | *AGT* | 3 | 167 | 6 | AGT | angiotensinogen | Extracellular Space | growth factor |
| chr16 | *CHTF18* | 3 | 100 | 6 | CHTF18 | chromosome transmission fidelity factor 18 | Nucleus | other |
| chr1 | *RP1-79C4.1* | 3 | 96 | 6 | | | | |
| chr3 | *RP11-379K17.4* | 2 | 88 | 6 | | | | |
| chr6 | *RAB23* | 3 | 64 | 6 | RAB23 | RAB23, member RAS oncogene family | Cytoplasm | enzyme |
| chr1 | *FCRL2* | 3 | 94 | 6 | FCRL2 | Fc receptor like 2 | Plasma Membrane | other |
| chr7 | *TRIM56* | 3 | 103 | 6 | TRIM56 | tripartite motif containing 56 | Cytoplasm | enzyme |
| chrX | *MPP1* | 2 | 48 | 6 | MPP1 | membrane palmitoylated protein | Plasma Membrane | kinase |
| chr22 | *DGCR6L* | 3 | 133 | 6 | DGCR6L | DiGeorge syndrome critical region gene 6 like | Nucleus | other |
| chr2 | *SERPINE2* | 3 | 70 | 6 | SERPINE2 | serpin family E member 2 | Extracellular Space | other |
| chr10 | *FAM25HP* | 3 | 145 | 6 | | | | |
| chr16 | *HS3ST6* | 3 | 119 | 6 | HS3ST6 | heparan sulfate-glucosamine 3-sulfotransferase 6 | Cytoplasm | enzyme |
| chr1 | *RP11-413P11.1* | 3 | 109 | 6 | | | | |
| chr1 | *GUCA2A* | 3 | 139 | 6 | GUCA2A | guanylate cyclase activator 2A | Extracellular Space | other |
| chr2 | *AC107070.1* | 2 | 62 | 6 | | | | |
| chr1 | *MRPL20* | 3 | 105 | 6 | MRPL20 | mitochondrial ribosomal protein | Cytoplasm | other |
| chr11 | *RP11-794P6.2* | 3 | 104 | 6 | | | | |
| chr19 | *ELOF1* | 3 | 85 | 6 | ELOF1 | elongation factor 1 homolog | Nucleus | other |
| chr1 | *LINGO4* | 3 | 153 | 6 | LINGO4 | leucine rich repeat and Ig domain containing 4 | Other | other |
| chr19 | *ZSCAN22* | 3 | 151 | 6 | ZSCAN22 | zinc finger and SCAN domain containing 22 | Nucleus | transcription regulator |
| chr1 | *SFPQ* | 3 | 92 | 6 | SFPQ | splicing factor proline and glutamine rich | Nucleus | other |
| chr16 | *ZNF668* | 3 | 70 | 6 | ZNF668 | zinc finger protein 668 | Cytoplasm | transcription regulator |
| chr14 | *OR4N2* | 2 | 56 | 6 | OR4N2 | olfactory receptor family 4 subfamily N member 2 | Plasma Membrane | G-protein coupled recepto |
| chr9 | *RP11-158D2.2* | 3 | 57 | 6 | | | | |
| chr5 | *KIAA1191* | 3 | 79 | 6 | KIAA1191 | KIAA1191 | Cytoplasm | other |
| chr10 | *RBP3* | 3 | 223 | 6 | RBP3 | retinol binding protein 3 | Extracellular Space | transporter |
| chr1 | *ELOVL1* | 3 | 119 | 6 | ELOVL1 | ELOVL fatty acid elongase 1 | Cytoplasm | enzyme |
| chr3 | *CDV3* | 3 | 71 | 6 | CDV3 | CDV3 homolog | Cytoplasm | other |

| | | | | | | | | | |
|---|---|---|---|---|---|---|---|---|---|
| chr10 | RP11-124O11.1 | 3 | 96 | 6 | | | | | |
| chr17 | KRT27 | 2 | 82 | 6 | KRT27 | keratin 27 | Cytoplasm | other | |
| chr1 | TLR12P | 2 | 95 | 6 | TLR12P | toll like receptor 12, pseudogene | Other | other | |
| chr17 | THA1P | 3 | 125 | 6 | THA1P | threonine aldolase 1, pseudogene | Other | other | |
| chr16 | RP11-118F19.1 | 3 | 85 | 6 | | | | | |
| chr16 | CTF1 | 3 | 81 | 6 | CTF1 | cardiotrophin 1 | Extracellular Space | cytokine | |
| chr7 | ING3 | 2 | 71 | 6 | ING3 | inhibitor of growth family member | Nucleus | other | |
| chrX | GRIA3 | 2 | 53 | 6 | GRIA3 | glutamate ionotropic receptor AMPA type subunit 3 | Plasma Membrane | ion channel | tezampanel, farampator, LY451395, |
| chr18 | CCDC102B | 3 | 65 | 6 | CCDC102B | coiled-coil domain containing | Extracellular Space | other | |
| chrX | MXRA5 | 2 | 60 | 6 | MXRA5 | matrix remodeling associated 5 | Extracellular Space | other | |
| chr21 | TFF2 | 3 | 96 | 6 | TFF2 | trefoil factor 2 | Extracellular Space | other | |
| chr2 | LYPD6B | 3 | 45 | 6 | LYPD6B | LY6/PLAUR domain containing | Other | other | |
| chr11 | RP5-1024C24.1 | 3 | 58 | 6 | | | | | |
| chr21 | TFF1 | 2 | 90 | 6 | TFF1 | trefoil factor 1 | Extracellular Space | other | |
| chrX | IRAK1 | 3 | 171 | 6 | IRAK1 | interleukin 1 receptor associated kinase 1 | Plasma Membrane | kinase | |
| chr9 | SOHLH1 | 3 | 206 | 6 | SOHLH1 | spermatogenesis and oogenesis specific basic helix-loop-helix 1 | Cytoplasm | transcription regulator | |
| chr2 | AC127383.1 | 2 | 79 | 6 | | | | | |
| chr15 | NF1P1 | 2 | 64 | 6 | NF1P1 | neurofibromin 1 pseudogene 1 | Other | other | |
| chr17 | AC004448.2 | 2 | 60 | 6 | | | | | |
| chr16 | UBFD1 | 3 | 66 | 6 | UBFD1 | ubiquitin family domain containing | Other | other | |
| chr2 | PRADC1 | 3 | 97 | 6 | PRADC1 | protease associated domain containing 1 | Extracellular Space | other | |
| chr16 | MMP25 | 3 | 129 | 6 | MMP25 | matrix metallopeptidase 25 | Plasma Membrane | peptidase | marimastat |
| chr16 | CES1P1 | 3 | 67 | 6 | CES1P1 | carboxylesterase 1 pseudogene 1 | Cytoplasm | other | |
| chr18 | SEH1L | 3 | 63 | 6 | SEH1L | SEH1 like nucleoporin | Cytoplasm | transporter | |
| chr16 | MSLN | 3 | 137 | 6 | MSLN | mesothelin | Extracellular Space | other | amatuximab, LMB-100, SS1(dsfv)PE38 |
| chr1 | AP006222.2 | 3 | 85 | 6 | | | | | |
| chr8 | PHF20L1 | 2 | 45 | 6 | PHF20L1 | PHD finger protein 20 like 1 | Other | other | |
| chr4 | ARAP2 | 2 | 59 | 6 | ARAP2 | ArfGAP with RhoGAP domain, ankyrin repeat and PH domain 2 | Cytoplasm | other | |
| chr3 | DPPA4 | 3 | 85 | 6 | DPPA4 | developmental pluripotency associated 4 | Nucleus | other | |
| chr15 | RP11-114H24.4 | 3 | 130 | 6 | | | | | |
| chr2 | AC108868.5 | 3 | 68 | 6 | | | | | |
| chr2 | CENPO | 2 | 59 | 6 | CENPO | centromere protein O | Nucleus | other | |
| chr10 | CALHM3 | 3 | 113 | 6 | CALHM3 | calcium homeostasis modulator 3 | Extracellular Space | other | |
| chr4 | ARL9 | 3 | 80 | 6 | ARL9 | ADP ribosylation factor like | Other | other | |
| chr5 | FAM114A2 | 3 | 50 | 6 | FAM114A2 | family with sequence similarity 114 member A2 | Other | other | |
| chr7 | MAFK | 3 | 173 | 6 | MAFK | MAF bZIP transcription factor K | Nucleus | transcription regulator | |
| chr11 | CDHR5 | 3 | 137 | 6 | CDHR5 | cadherin related family member 5 | Plasma Membrane | other | |
| chr6 | ZKSCAN3 | 3 | 74 | 6 | ZKSCAN3 | zinc finger with KRAB and SCAN domains 3 | Nucleus | transcription regulator | |
| chr9 | C9orf152 | 3 | 67 | 6 | C9orf152 | chromosome 9 open reading frame 152 | Other | other | |
| chr4 | FAM200B | 3 | 52 | 6 | FAM200B | family with sequence similarity 200 member B | Other | other | |
| chr16 | RP11-989E6.2 | 3 | 132 | 6 | | | | | |
| chr11 | PRPF19 | 3 | 119 | 6 | PRPF19 | pre-mRNA processing factor 19 | Nucleus | enzyme | |
| chr5 | CTD-2139B15.4 | 3 | 193 | 6 | | | | | |
| chrX | GEMIN8 | 3 | 85 | 6 | GEMIN8 | gem nuclear organelle associated protein 8 | Cytoplasm | other | |
| chr10 | AKR1C3 | 3 | 79 | 6 | AKR1C3 | aldo-keto reductase family 1 member C3 | Cytoplasm | enzyme | ASP9521 |
| chr20 | BPIFA3 | 3 | 92 | 6 | BPIFA3 | BPI fold containing family A member 3 | Other | other | |
| chr3 | ANAPC13 | 2 | 55 | 6 | ANAPC13 | anaphase promoting complex subunit 13 | Nucleus | other | |
| chr22 | NCF4 | 2 | 78 | 6 | NCF4 | neutrophil cytosolic factor 4 | Cytoplasm | enzyme | |
| chr10 | PRTFDC1 | 3 | 55 | 6 | PRTFDC1 | phosphoribosyl transferase domain containing 1 | Cytoplasm | enzyme | |
| chr16 | MSRB1 | 3 | 155 | 6 | MSRB1 | methionine sulfoxide reductase | Nucleus | enzyme | |
| chr20 | PCK1 | 3 | 172 | 6 | PCK1 | phosphoenolpyruvate carboxykinase 1 | Cytoplasm | kinase | |
| chrX | RP11-348F1.3 | 3 | 63 | 6 | | | | | |
| chr20 | WFDC9 | 3 | 65 | 6 | WFDC9 | WAP four-disulfide core domain 9 | Extracellular Space | other | |
| chr4 | IL21 | 2 | 44 | 6 | IL21 | interleukin 21 | Extracellular Space | cytokine | |
| chr1 | XXyac-YX155B6.[illegible] | 3 | 56 | 6 | | | | | |
| chr12 | LHX5 | 3 | 84 | 6 | LHX5 | LIM homeobox 5 | Nucleus | transcription regulator | |
| chr1 | RP4-788P17.1 | 3 | 46 | 6 | | | | | |
| chr20 | AURKA | 2 | 95 | 6 | AURKA | aurora kinase A | Nucleus | kinase | ilorasertib, MK 5108, SNS 314, AT-9283, alisertib, MLN8054, TTP607, CYC 116, TAS-119, tozasertib, AMG 900, danusertib |
| chr19 | ZNF99 | 3 | 61 | 6 | ZNF99 | zinc finger protein 99 | Other | other | |
| chr13 | LINC00284 | 3 | 64 | 6 | NRAD1 | non-coding RNA in the aldehyde dehydrogenase 1A pathway | Other | other | |
| chr22 | LGALS2 | 3 | 79 | 6 | LGALS2 | galectin 2 | Cytoplasm | other | |
| chr1 | TRNP1 | 3 | 86 | 6 | TRNP1 | TMF1 regulated nuclear protein 1 | Nucleus | other | |
| chr8 | KIFC2 | 3 | 142 | 6 | KIFC2 | kinesin family member C2 | Cytoplasm | other | |
| chr6 | SERPINB1 | 3 | 82 | 6 | SERPINB1 | serpin family B member 1 | Cytoplasm | other | |
| chr6 | WRNIP1 | 3 | 82 | 6 | WRNIP1 | WRN helicase interacting protein | Nucleus | enzyme | |

| | | | | | | | | | |
|---|---|---|---|---|---|---|---|---|---|
| chr1 | *ARTN* | 3 | 117 | 6 | ARTN | artemin | Extracellular Space | growth factor | |
| chr19 | *TMEM259* | 3 | 157 | 6 | TMEM259 | transmembrane protein 259 | Cytoplasm | other | |
| chr6 | *MRPS10* | 2 | 65 | 6 | MRPS10 | mitochondrial ribosomal protein | Cytoplasm | other | |
| chr1 | *C1orf174* | 3 | 146 | 6 | C1orf174 | chromosome 1 open reading frame 174 | Nucleus | other | |
| chr10 | *SFTPD* | 3 | 73 | 6 | SFTPD | surfactant protein D | Extracellular Space | other | |
| chr7 | *FAM115B* | 2 | 101 | 6 | TCAF1P1 | TRPM8 channel associated factor 1 pseudogene 1 | Other | other | |
| chr9 | *RP11-374M1.3* | 2 | 71 | 6 | | | | | |
| chr1 | *DENND2D* | 3 | 123 | 6 | DENND2D | DENN domain containing 2D | Cytoplasm | other | |
| chr22 | *AP000345.2* | 2 | 91 | 6 | | | | | |
| chr12 | *RP11-797M17.1* | 3 | 58 | 6 | | | | | |
| chr18 | *GALNT1* | 2 | 59 | 6 | GALNT1 | polypeptide N-acetylgalactosaminyltransferase | Cytoplasm | enzyme | |
| chr6 | *MMS22L* | 2 | 58 | 6 | MMS22L | MMS22 like, DNA repair protein | Nucleus | other | |
| chrX | *TRAPPC2* | 3 | 77 | 6 | TRAPPC2 | trafficking protein particle | Cytoplasm | other | |
| chr17 | *DVL2* | 3 | 165 | 6 | DVL2 | dishevelled segment polarity | Cytoplasm | other | |
| chr22 | *BPIFC* | 2 | 60 | 6 | BPIFC | BPI fold containing family C | Extracellular Space | transporter | |
| chr17 | *COPZ2* | 3 | 81 | 6 | COPZ2 | coatomer protein complex subunit zeta 2 | Cytoplasm | transporter | |
| chr8 | *CTD-2588J1.1* | 3 | 74 | 6 | | | | | |
| chr19 | *S1PR4* | 2 | 140 | 6 | S1PR4 | sphingosine-1-phosphate | Plasma Membrane | -protein coupled recepto | fingolimod phosphate |
| chr19 | *PTBP1* | 3 | 71 | 6 | PTBP1 | polypyrimidine tract binding | Nucleus | enzyme | |
| chr5 | *PFDN1* | 2 | 58 | 6 | PFDN1 | prefoldin subunit 1 | Cytoplasm | transcription regulator | |
| chr2 | *NDUFB3* | 3 | 56 | 6 | NDUFB3 | NADH:ubiquinone oxidoreductase subunit B3 | Cytoplasm | enzyme | |
| chr4 | *UGT8* | 2 | 63 | 6 | UGT8 | UDP glycosyltransferase 8 | Cytoplasm | enzyme | |
| chr9 | *DIRAS2* | 2 | 76 | 6 | DIRAS2 | DIRAS family GTPase 2 | Plasma Membrane | enzyme | |
| chr2 | *PSMD1* | 2 | 44 | 6 | PSMD1 | proteasome 26S subunit, non-ATPase 1 | Cytoplasm | other | bortezomib/cladribine/rituximab, bortezomib/dexamethasone/pomalidomide, bortezomib/doxorubicin, bortezomib/dexamethasone/thalidomide, bortezomib/paclitaxel, bortezomib/fulvestrant, bortezomib/rituximab, bortezomib/sorafenib, bortezomib/lenalidomide, bortezomib/dexamethasone/lenalidomide, bortezomib/dexamethasone/doxorubicin, bortezomib/predniso[illegible] |
| chr2 | *CNNM3* | 3 | 154 | 6 | CNNM3 | cyclin and CBS domain divalent metal cation transport mediator 3 | Other | other | |
| chr7 | *RP11-114L10.2* | 3 | 101 | 6 | | | | | |
| chr16 | *ZNF747* | 3 | 69 | 6 | ZNF747 | zinc finger protein 747 | Other | other | |
| chr2 | *GLS* | 2 | 51 | 6 | GLS | glutaminase | Cytoplasm | enzyme | CB-839 |
| chr14 | *TSHR* | 3 | 48 | 6 | TSHR | thyroid stimulating hormone | Plasma Membrane | -protein coupled recepto | ORG 41841 |
| chrX | *GAGE12G* | 3 | 82 | 6 | GE1 (includes oth | G antigen 1 | Other | other | |
| chr14 | *NID2* | 3 | 80 | 6 | NID2 | nidogen 2 | Extracellular Space | other | |
| chr1 | *GLTPD1* | 3 | 143 | 6 | CPTP | ceramide-1-phosphate transfer protein | Cytoplasm | transporter | |
| chr2 | *AC114783.1* | 3 | 123 | 6 | | | | | |
| chr18 | *KDSR* | 2 | 47 | 6 | KDSR | 3-ketodihydrosphingosine | Plasma Membrane | enzyme | |
| chr10 | *RP11-491H19.1* | 3 | 116 | 6 | | | | | |
| chr9 | *RABL6* | 3 | 244 | 6 | RABL6 | RAB, member RAS oncogene family like 6 | Cytoplasm | other | |
| chr1 | *RP11-84A19.2* | 3 | 81 | 6 | | | | | |
| chr11 | *HTATIP2* | 2 | 50 | 6 | HTATIP2 | HIV-1 Tat interactive protein 2 | Nucleus | transcription regulator | |
| chr22 | *CTA-992D9.7* | 2 | 47 | 6 | | | | | |
| chr20 | *RP11-112L6.2* | 3 | 118 | 6 | | | | | |
| chr4 | *PPARGC1A* | 2 | 46 | 6 | PPARGC1A | PPARG coactivator 1 alpha | Nucleus | transcription regulator | |
| chr19 | *ZNF66* | 3 | 81 | 6 | ZNF66 | zinc finger protein 66 | Other | other | |
| chr17 | *C17orf105* | 3 | 77 | 6 | CFAP97D1 | CFAP97 domain containing 1 | Other | other | |
| chr3 | *CAMP* | 3 | 87 | 6 | CAMP | cathelicidin antimicrobial peptide | Cytoplasm | other | |
| chr17 | *CXCL16* | 3 | 105 | 6 | CXCL16 | C-X-C motif chemokine ligand 16 | Extracellular Space | cytokine | |
| chr5 | *SPINK6* | 2 | 66 | 6 | SPINK6 | serine peptidase inhibitor, Kazal type 6 | Extracellular Space | other | |
| chr9 | *ELAVL2* | 2 | 45 | 6 | ELAVL2 | ELAV like RNA binding protein 2 | Cytoplasm | other | |
| chr17 | *ZACN* | 3 | 81 | 6 | ZACN | zinc activated ion channel | Plasma Membrane | ion channel | |
| chr22 | *TMEM211* | 2 | 67 | 6 | TMEM211 | transmembrane protein 211 | Other | other | |
| chr19 | *LILRB5* | 3 | 97 | 6 | LILRB5 | leukocyte immunoglobulin like receptor B5 | Plasma Membrane | transmembrane receptor | |
| chr7 | *RP11-153N17.1* | 2 | 55 | 6 | | | | | |
| chr9 | *PPP1R26* | 3 | 154 | 6 | PPP1R26 | protein phosphatase 1 regulatory subunit 26 | Nucleus | other | |
| chr7 | *RP5-884M6.1* | 3 | 53 | 6 | | | | | |
| chr8 | *RP11-378A12.1* | 3 | 53 | 6 | | | | | |
| chr6 | *TPD52L1* | 3 | 53 | 6 | TPD52L1 | TPD52 like 1 | Cytoplasm | other | |

| | | | | | | | | | |
|---|---|---|---|---|---|---|---|---|---|
| chr1 | *FCRL4* | 3 | 69 | 6 | FCRL4 | Fc receptor like 4 | Cytoplasm | other | |
| chr11 | *C1QTNF4* | 3 | 182 | 6 | C1QTNF4 | C1q and TNF related 4 | Extracellular Space | other | |
| chr5 | *PROP1* | 3 | 118 | 6 | PROP1 | PROP paired-like homeobox 1 | Nucleus | transcription regulator | |
| chr9 | *C9orf78* | 3 | 84 | 6 | C9orf78 | chromosome 9 open reading frame 78 | Other | other | |
| chr10 | *SEPHS1* | 3 | 76 | 6 | SEPHS1 | selenophosphate synthetase 1 | Other | enzyme | |
| chr9 | *RP11-391M20.1* | 3 | 285 | 6 | | | | | |
| chr16 | *RP11-883G14.1* | 3 | 53 | 6 | | | | | |
| chr11 | *CTD-2216M2.1* | 3 | 108 | 6 | | | | | |
| chr3 | *IL17RB* | 3 | 64 | 6 | IL17RB | interleukin 17 receptor B | Plasma Membrane | transmembrane recepto | |
| chr1 | *RP11-147C23.1* | 3 | 68 | 6 | | | | | |
| chr10 | *RP11-463P17.1* | 3 | 197 | 6 | | | | | |
| chr14 | *KCNK10* | 3 | 52 | 6 | KCNK10 | potassium two pore domain channel subfamily K member 10 | Plasma Membrane | ion channel | |
| chr1 | *SSX2IP* | 3 | 65 | 6 | SSX2IP | SSX family member 2 interacting protein | Plasma Membrane | other | |
| chr14 | *SAV1* | 2 | 42 | 6 | SAV1 | salvador family WW domain containing protein 1 | Cytoplasm | other | |
| chr1 | *RP11-10N16.3* | 3 | 68 | 6 | | | | | |
| chr2 | *KCNE4* | 3 | 58 | 6 | KCNE4 | potassium voltage-gated channel subfamily E regulatory subunit 4 | Plasma Membrane | ion channel | |
| chr7 | *ASNS* | 2 | 60 | 6 | ASNS | asparagine synthetase (glutamine-hydrolyzing) | Cytoplasm | enzyme | |
| chr5 | *RP11-308B16.2* | 2 | 102 | 6 | | | | | |
| chr16 | *RP11-429P3.3* | 3 | 50 | 6 | | | | | |
| chr12 | *A2M* | 2 | 47 | 6 | A2M | alpha-2-macroglobulin | Extracellular Space | transporter | |
| chr17 | *RP11-219A15.4* | 3 | 72 | 6 | | | | | |
| chr2 | *RPIA* | 2 | 47 | 6 | RPIA | ribose 5-phosphate isomerase A | Cytoplasm | enzyme | |
| chrX | *FHL1* | 2 | 68 | 6 | FHL1 | four and a half LIM domains 1 | Cytoplasm | other | |
| chr5 | *TRIM41* | 3 | 141 | 6 | TRIM41 | tripartite motif containing 41 | Cytoplasm | other | |
| chr1 | *DMAP1* | 3 | 103 | 6 | DMAP1 | DNA methyltransferase 1 associated protein 1 | Nucleus | transcription regulator | |
| chr15 | *RP11-346D14.1* | 2 | 61 | 6 | | | | | |
| chr15 | *RP11-154J22.1* | 3 | 50 | 6 | | | | | |
| chr15 | *ZFYVE19* | 2 | 115 | 6 | ZFYVE19 | zinc finger FYVE-type containing | Cytoplasm | other | |
| chr7 | *AC002383.2* | 3 | 77 | 6 | | | | | |
| chr17 | *NEUROD2* | 3 | 69 | 6 | NEUROD2 | neuronal differentiation 2 | Nucleus | transcription regulator | |
| chr1 | *ZRANB2-AS1* | 3 | 90 | 6 | ZRANB2-AS1 | ZRANB2 antisense RNA 1 | Other | other | |
| chr12 | *LEMD3* | 2 | 75 | 6 | LEMD3 | LEM domain containing 3 | Nucleus | other | |
| chr6 | *RPL7L1* | 3 | 67 | 6 | RPL7L1 | ribosomal protein L7 like 1 | Cytoplasm | transcription regulator | |
| chr16 | *SLC5A2* | 3 | 102 | 6 | SLC5A2 | solute carrier family 5 member 2 | Plasma Membrane | transporter | dapagliflozin, canagliflozin, ertugliflozin, ertugliflozin/sitagliptin, dapagliflozin/saxagliptin, SGLT2 inhibitor, empagliflozin, dapagliflozin/INS. |
| chr11 | *OR52A1* | 3 | 77 | 6 | OR52A1 | olfactory receptor family 52 subfamily A member 1 | Plasma Membrane | G-protein coupled recepto | |
| chr10 | *LRIT1* | 3 | 78 | 6 | LRIT1 | leucine rich repeat, Ig-like and transmembrane domains 1 | Cytoplasm | other | |
| chr2 | *AC092162.1* | 3 | 63 | 6 | | | | | |
| chr11 | *RP11-113D6.6* | 3 | 49 | 6 | | | | | |
| chr2 | *ECEL1* | 3 | 147 | 6 | ECEL1 | endothelin converting enzyme | Plasma Membrane | peptidase | |
| chr9 | *LCN15* | 3 | 213 | 6 | LCN15 | lipocalin 15 | Other | other | |
| chr20 | *DEFB123* | 3 | 109 | 6 | DEFB123 | defensin beta 123 | Extracellular Space | other | |
| chr2 | *DNAJC27* | 2 | 46 | 6 | DNAJC27 | DnaJ heat shock protein family (Hsp40) member C27 | Cytoplasm | enzyme | |
| chr7 | *AZGP1P1* | 3 | 116 | 6 | AZGP1P1 | AZGP1 pseudogene 1 | Other | other | |
| chr15 | *RP11-580I1.2* | 3 | 100 | 6 | | | | | |
| chr19 | *CTD-2081K17.2* | 3 | 54 | 6 | | | | | |
| chr3 | *WDR6* | 3 | 106 | 6 | WDR6 | WD repeat domain 6 | Cytoplasm | other | |
| chr22 | *CRYBB3* | 3 | 96 | 6 | CRYBB3 | crystallin beta B3 | Other | other | |
| chr13 | *LINC00598* | 3 | 61 | 6 | LINC00598 | long intergenic non-protein coding RNA 598 | Other | other | |
| chr4 | *HPSE* | 3 | 64 | 6 | HPSE | heparanase | Plasma Membrane | enzyme | 2-O,3-O-desulfated heparin, heparanase inhibitor PI-88, roneparstat, PG 545 |
| chr15 | *LYSMD2* | 3 | 46 | 6 | LYSMD2 | LysM domain containing 2 | Other | other | |
| chr9 | *MUSK* | 2 | 81 | 6 | MUSK | muscle associated receptor tyrosine kinase | Plasma Membrane | kinase | |
| chr19 | *BSPH1* | 2 | 45 | 6 | BSPH1 | binder of sperm protein homolog | Plasma Membrane | other | |
| chr10 | *FAM170B-AS1* | 3 | 129 | 6 | FAM170B-AS1 | FAM170B antisense RNA 1 | Other | other | |
| chr3 | *RP11-324C10.1* | 2 | 52 | 6 | | | | | |
| chr22 | *AP000525.9* | 3 | 47 | 6 | | | | | |
| chr11 | *MPZL3* | 2 | 67 | 6 | MPZL3 | myelin protein zero like 3 | Other | other | |
| chr19 | *EPHX3* | 3 | 97 | 6 | EPHX3 | epoxide hydrolase 3 | Other | enzyme | |
| chr17 | *RP11-700H6.2* | 3 | 77 | 6 | | | | | |
| chr13 | *LCP1* | 2 | 54 | 6 | LCP1 | lymphocyte cytosolic protein 1 | Cytoplasm | other | |
| chr12 | *RP11-644F5.16* | 3 | 122 | 6 | | | | | |
| chr4 | *MFSD8* | 2 | 57 | 6 | MFSD8 | major facilitator superfamily domain containing 8 | Nucleus | other | |
| chr19 | *MKNK2* | 3 | 162 | 6 | MKNK2 | MAPK interacting serine/threonine kinase 2 | Cytoplasm | kinase | ETC-1907206, tomivosertib |
| chr6 | *HIST1H2BN* | 2 | 85 | 6 | H2BC15 | H2B clustered histone 15 | Nucleus | other | |

| | | | | | | | | | |
|---|---|---|---|---|---|---|---|---|---|
| chrX | *CT45A3* | 2 | 113 | 6 | 45A1 (includes oth | cancer/testis antigen family 45 member A2 | Other | other | |
| chr11 | *TMEM9B* | 3 | 87 | 6 | TMEM9B | TMEM9 domain family member B | Other | other | |
| chr17 | *AC079325.5* | 3 | 128 | 6 | | | | | |
| chr5 | *PRRC1* | 3 | 73 | 6 | PRRC1 | proline rich coiled-coil 1 | Other | other | |
| chr12 | *RP11-654D12.2* | 2 | 58 | 6 | | | | | |
| chr12 | *RP13-507P19.1* | 3 | 109 | 6 | | | | | |
| chr1 | *RP5-1011O1.2* | 2 | 54 | 6 | | | | | |
| chr14 | *CTD-2200A16.1* | 2 | 71 | 6 | | | | | |
| chr4 | *AFAP1-AS1* | 2 | 98 | 6 | AFAP1-AS1 | AFAP1 antisense RNA 1 | Other | other | |
| chr17 | *SHC1P2* | 2 | 85 | 6 | SHC1P2 | SHC adaptor protein 1 | Other | other | |
| chr8 | *KB-173C10.2* | 2 | 58 | 6 | | | | | |
| chr6 | *RSPH4A* | 2 | 64 | 6 | RSPH4A | radial spoke head component 4A | Nucleus | other | |
| chr1 | *RP11-134P9.1* | 3 | 87 | 6 | | | | | |
| chr1 | *RP11-223J15.2* | 3 | 76 | 6 | | | | | |
| chr17 | *RP11-215E13.1* | 2 | 51 | 6 | | | | | |
| chrX | *MAGED2* | 2 | 127 | 6 | MAGED2 | MAGE family member D2 | Plasma Membrane | other | |
| chr12 | *RP11-547C5.2* | 2 | 58 | 6 | | | | | |
| chr20 | *RP4-614C15.2* | 3 | 146 | 6 | | | | | |
| chr2 | *CLIP4* | 3 | 45 | 6 | CLIP4 | CAP-Gly domain containing linker protein family member 4 | Other | other | |
| chr12 | *RP11-362K2.2* | 3 | 51 | 6 | | | | | |
| chr12 | *ARPC3* | 2 | 69 | 6 | ARPC3 | actin related protein 2/3 complex subunit 3 | Cytoplasm | other | |
| chr16 | *SRRM2* | 3 | 131 | 6 | SRRM2 | serine/arginine repetitive matrix 2 | Nucleus | other | |
| chr19 | *GALP* | 3 | 65 | 6 | GALP | galanin like peptide | Extracellular Space | other | |
| chr12 | *RP11-629N8.3* | 3 | 54 | 6 | | | | | |
| chr9 | *RP11-98L5.4* | 3 | 92 | 6 | | | | | |
| chr12 | *RP11-263K4.5* | 2 | 43 | 6 | | | | | |
| chr1 | *TAGLN2* | 3 | 169 | 6 | TAGLN2 | transgelin 2 | Cytoplasm | other | |
| chr14 | *CTD-2314B22.3* | 3 | 77 | 6 | | | | | |
| chr1 | *RP5-902P8.10* | 3 | 148 | 6 | | | | | |
| chr21 | *AF131217.1* | 3 | 43 | 6 | | | | | |
| chr5 | *AC008592.4* | 3 | 102 | 6 | | | | | |
| chr11 | *CNIH2* | 3 | 214 | 6 | CNIH2 | cornichon family AMPA receptor auxiliary protein 2 | Extracellular Space | other | |
| chr7 | *SLC35B4* | 2 | 63 | 6 | SLC35B4 | solute carrier family 35 member | Cytoplasm | transporter | |
| chr13 | *IL17D* | 3 | 103 | 6 | IL17D | interleukin 17D | Extracellular Space | cytokine | |
| chr7 | *TMEM176B* | 3 | 66 | 6 | TMEM176B | transmembrane protein 176B | Other | other | |
| chrX | *NXF2* | 2 | 77 | 6 | NXF2/NXF2B | nuclear RNA export factor 2 | Nucleus | transporter | |
| chr11 | *CTR9* | 3 | 106 | 6 | CTR9 | CTR9 homolog, Paf1/RNA polymerase II complex | Nucleus | other | |
| chr17 | *FAM171A2* | 3 | 131 | 6 | FAM171A2 | family with sequence similarity 171 member A2 | Other | other | |
| chr5 | *PTGER4* | 3 | 225 | 6 | PTGER4 | prostaglandin E receptor 4 | Plasma Membrane | G-protein coupled recepto | prostaglandin E2, grapiprant, BMS-986310, diclofenac/misoprostol, misoprostol, E7046, rivenprost, prostaglandin E1 |
| chr16 | *ZNF629* | 3 | 132 | 6 | ZNF629 | zinc finger protein 629 | Nucleus | transcription regulator | |
| chr11 | *RP11-680H20.2* | 3 | 483 | 6 | | | | | |
| chr19 | *GAMT* | 3 | 147 | 6 | GAMT | guanidinoacetate N-methyltransferase | Cytoplasm | enzyme | |
| chrX | *FAM155B* | 2 | 79 | 6 | FAM155B | family with sequence similarity 155 member B | Plasma Membrane | transmembrane receptor | |
| chr7 | *AC005029.1* | 2 | 64 | 6 | | | | | |
| chr11 | *DCUN1D5* | 3 | 54 | 6 | DCUN1D5 | defective in cullin neddylation 1 domain containing 5 | Other | other | |
| chr2 | *CYP27A1* | 3 | 81 | 6 | CYP27A1 | cytochrome P450 family 27 subfamily A member 1 | Cytoplasm | enzyme | |
| chr5 | *WNT8A* | 2 | 69 | 6 | WNT8A | Wnt family member 8A | Extracellular Space | other | |
| chr21 | *AP001626.1* | 2 | 61 | 6 | | | | | |
| chr17 | *GAS2L2* | 3 | 143 | 6 | GAS2L2 | growth arrest specific 2 like 2 | Cytoplasm | other | |
| chr19 | *CTC-326K19.6* | 3 | 79 | 6 | | | | | |
| chr15 | *PIGB* | 2 | 60 | 6 | PIGB | phosphatidylinositol glycan anchor biosynthesis class B | Cytoplasm | enzyme | |
| chr8 | *STMN4* | 3 | 108 | 6 | STMN4 | stathmin 4 | Cytoplasm | other | |
| chr17 | *HILS1* | 3 | 175 | 6 | H1-9 | H1.9 linker histone (pseudogene) | Nucleus | other | |
| chr16 | *RP11-481E4.2* | 3 | 51 | 6 | | | | | |
| chr4 | *RP11-422J15.1* | 3 | 59 | 6 | | | | | |
| chr5 | *CTB-113P19.1* | 3 | 62 | 6 | | | | | |
| chr20 | *PTK6* | 3 | 264 | 6 | PTK6 | protein tyrosine kinase 6 | Cytoplasm | kinase | everolimus/vandetanib, vandetanib |
| chr9 | *LRRC37A5P* | 3 | 45 | 6 | LRRC37A5P | leucine rich repeat containing 37 member A5, pseudogene | Other | other | |
| chr22 | *RP1-102D24.5* | 3 | 93 | 6 | | | | | |
| chr16 | *RP11-296I10.6* | 3 | 66 | 6 | | | | | |
| chr19 | *ALDH16A1* | 3 | 121 | 6 | ALDH16A1 | aldehyde dehydrogenase 16 family member A1 | Cytoplasm | enzyme | |
| chr12 | *RP11-981P6.1* | 3 | 64 | 6 | | | | | |
| chr19 | *SIGLEC20P* | 3 | 98 | 6 | SIGLEC20P | sialic acid binding Ig like lectin 20, pseudogene | Other | other | |
| chr2 | *CGREF1* | 2 | 76 | 6 | CGREF1 | cell growth regulator with EF-hand domain 1 | Extracellular Space | other | |
| chr6 | *RP1-152L7.1* | 3 | 60 | 6 | | | | | |
| chr18 | *HAUS1* | 2 | 49 | 6 | HAUS1 | HAUS augmin like complex | Nucleus | other | |

| | | | | | | | | | |
|---|---|---|---|---|---|---|---|---|---|
| chr20 | *FAM209B* | 3 | 128 | 6 | FAM209B | family with sequence similarity 209 member B | Nucleus | other | |
| chr19 | *PSG6* | 3 | 101 | 6 | PSG6 | pregnancy specific beta-1-glycoprotein 6 | Extracellular Space | other | |
| chr11 | *UNC93B5* | 3 | 136 | 6 | UNC93B5 | unc-93 homolog B5, pseudogene | Other | other | |
| chrX | *UPF3B* | 2 | 59 | 6 | UPF3B | UPF3B regulator of nonsense mediated mRNA decay | Nucleus | transporter | |
| chr18 | *C18orf42* | 3 | 126 | 6 | AKAIN1 | A-kinase anchor inhibitor 1 | Cytoplasm | other | |
| chr1 | *C1orf109* | 3 | 106 | 6 | C1orf109 | chromosome 1 open reading frame 109 | Nucleus | other | |
| chr6 | *CD83* | 2 | 79 | 6 | CD83 | CD83 molecule | Plasma Membrane | transmembrane receptor | |
| chr8 | *GPAA1* | 3 | 143 | 6 | GPAA1 | glycosylphosphatidylinositol anchor attachment 1 | Cytoplasm | enzyme | |
| chr1 | *MIR137HG* | 2 | 82 | 6 | MIR137HG | MIR137 host gene | Other | other | |
| chr20 | *RP4-764O22.1* | 2 | 61 | 6 | | | | | |
| chr6 | *PTP4A1* | 2 | 47 | 6 | PTP4A1 | protein tyrosine phosphatase 4A1 | Cytoplasm | phosphatase | |
| chr11 | *RP11-115E19.1* | 2 | 135 | 6 | | | | | |
| chr17 | *CTD-2319I12.4* | 3 | 100 | 6 | | | | | |
| chr20 | *STX16-NPEPL1* | 3 | 101 | 6 | STX16-NPEPL1 | STX16-NPEPL1 readthrough (NMD candidate) | Other | other | |
| chr8 | *LINC00208* | 3 | 112 | 6 | LINC00208 | long intergenic non-protein coding RNA 208 | Other | other | |
| chr10 | *TCTN3* | 3 | 65 | 6 | TCTN3 | tectonic family member 3 | Cytoplasm | other | |
| chr10 | *LINC00841* | 3 | 106 | 6 | LINC00841 | long intergenic non-protein coding RNA 841 | Other | other | |
| chrX | *NXF3* | 2 | 63 | 6 | NXF3 | nuclear RNA export factor 3 | Nucleus | transporter | |
| chr13 | *LINC00407* | 3 | 45 | 6 | LINC00407 | long intergenic non-protein coding RNA 407 | Other | other | |
| chr6 | *ENPP1* | 3 | 69 | 6 | ENPP1 | ectonucleotide pyrophosphatase/phosphodiesterase 1 | Plasma Membrane | enzyme | |
| chr15 | *RP11-2F9.4* | 3 | 256 | 6 | | | | | |
| chr12 | *KLHL42* | 3 | 102 | 6 | KLHL42 | kelch like family member 42 | Nucleus | other | |
| chr15 | *RP11-219B17.1* | 2 | 55 | 6 | | | | | |
| chr22 | *GSTT2* | 2 | 99 | 6 | GSTT2/GSTT2B | glutathione S-transferase theta 2 (gene/pseudogene) | Cytoplasm | enzyme | |
| chr19 | *ZNF266* | 2 | 68 | 6 | ZNF266 | zinc finger protein 266 | Nucleus | transcription regulator | |
| chr6 | *PHIP* | 2 | 62 | 6 | PHIP | pleckstrin homology domain interacting protein | Nucleus | other | |
| chr15 | *RP11-114H24.5* | 3 | 197 | 6 | | | | | |
| chr22 | *SUSD2* | 3 | 152 | 6 | SUSD2 | sushi domain containing 2 | Extracellular Space | other | |
| chr6 | *BTN2A3P* | 3 | 80 | 6 | BTN2A3P | butyrophilin subfamily 2 member A3, pseudogene | Other | other | |
| chr3 | *PHF7* | 3 | 78 | 6 | PHF7 | PHD finger protein 7 | Nucleus | other | |
| chr14 | *FAM161B* | 3 | 65 | 6 | FAM161B | FAM161 centrosomal protein B | Cytoplasm | other | |
| chr16 | *GDE1* | 3 | 46 | 6 | GDE1 | glycerophosphodiester phosphodiesterase 1 | Plasma Membrane | enzyme | |
| chr3 | *MST1* | 3 | 86 | 6 | MST1 | macrophage stimulating 1 | Extracellular Space | growth factor | |
| chr5 | *CTD-2046I8.1* | 3 | 261 | 6 | | | | | |
| chr12 | *XRCC6BP1* | 3 | 88 | 6 | ATP23 | ATP23 metallopeptidase and ATP synthase assembly factor | Plasma Membrane | kinase | |
| chr14 | *AE000661.37* | 3 | 52 | 6 | | | | | |
| chr3 | *PCNP* | 2 | 62 | 6 | PCNP | PEST proteolytic signal containing nuclear protein | Nucleus | other | |
| chrX | *ALAS2* | 3 | 80 | 6 | ALAS2 | 5'-aminolevulinate synthase 2 | Cytoplasm | enzyme | |
| chr19 | *APBA3* | 3 | 104 | 6 | APBA3 | amyloid beta precursor protein binding family A member 3 | Cytoplasm | transporter | |
| chr9 | *GRIN3A* | 2 | 65 | 6 | GRIN3A | glutamate ionotropic receptor NMDA type subunit 3A | Plasma Membrane | ion channel | besonprodil, delucemine, 1-aminocyclopropane-1-carboxylic acid, orphenadrine, dextromethorphan/promethazine, aspirin/caffeine/orphenadrine, phencyclidine, amitriptyline/ketamine, caffeine/ethanol, UK-240455, brompheniramine/dextromethorphan/pseudoephedrine, bicifadine, methadone, ethopropazine, procaine, nebostinel, amantadine, neramexane, felbamate, [illegible] |
| chr12 | *RP11-693J15.4* | 2 | 42 | 6 | | | | | |
| chrX | *SLC9A6* | 2 | 49 | 6 | SLC9A6 | solute carrier family 9 member A6 | Plasma Membrane | transporter | |
| chr10 | *C10orf67* | 2 | 73 | 6 | C10orf67 | chromosome 10 open reading frame 67 | Other | other | |
| chr6 | *RP11-145H9.3* | 2 | 104 | 6 | | | | | |
| chr17 | *TMEM98* | 2 | 78 | 6 | TMEM98 | transmembrane protein 98 | Cytoplasm | other | |
| chr6 | *PNLDC1* | 2 | 77 | 6 | PNLDC1 | PARN like, ribonuclease domain containing 1 | Cytoplasm | enzyme | |

| | | | | | | | | | |
|---|---|---|---|---|---|---|---|---|---|
| chr12 | *SOCS2-AS1* | 3 | 68 | 6 | SOCS2-AS1 | SOCS2 antisense RNA 1 | Other | other | |
| chr2 | *E2F6* | 2 | 44 | 6 | E2F6 | E2F transcription factor 6 | Nucleus | transcription regulator | |
| chr11 | *B4GALNT4* | 3 | 145 | 6 | B4GALNT4 | beta-1,4-N-acetyl-galactosaminyltransferase 4 | Extracellular Space | enzyme | |
| chr1 | *KIAA2013* | 3 | 67 | 6 | KIAA2013 | KIAA2013 | Other | other | |
| chr11 | *CHRM1* | 3 | 166 | 6 | CHRM1 | cholinergic receptor muscarinic 1 | Plasma Membrane | G-protein coupled recepto | dexamethasone/olanzapine, dicyclomine, cevimeline, mirabegron/solifenacin, olanzapine, aclidinium/formoterol, acetylcholine, atropine/pralidoxime, succinylcholine, brompheniramine, umeclidinium, mepenzolic acid, propantheline, orphenadrine, aspirin/caffeine/orphenadrine, tolterodine, chlorpheniramine/methscopolamine/phenylephrine, trihexyphenidyl, oxybutynin, formoterol/glycopyrrolate, scopolamine, [illegible] |
| chr14 | *SERPINA12* | 2 | 65 | 6 | SERPINA12 | serpin family A member 12 | Cytoplasm | other | |
| chr5 | *RP11-249M12.2* | 3 | 60 | 6 | | | | | |
| chr19 | *AC002511.3* | 3 | 80 | 6 | | | | | |
| chr20 | *FITM2* | 3 | 98 | 6 | FITM2 | fat storage inducing transmembrane protein 2 | Cytoplasm | other | |
| chr11 | *AP000439.2* | 3 | 137 | 6 | | | | | |
| chr9 | *CDKN2B-AS1* | 2 | 49 | 6 | CDKN2B-AS1 | CDKN2B antisense RNA 1 | Other | other | |
| chr10 | *FRMPD2P1* | 3 | 115 | 6 | FRMPD2B | FERM and PDZ domain containing 2B, pseudogene | Other | other | |
| chr12 | *RP11-443N24.2* | 2 | 62 | 6 | | | | | |
| chr1 | *RP1-69M21.2* | 3 | 86 | 6 | | | | | |
| chr6 | *MCHR2* | 2 | 62 | 6 | MCHR2 | melanin concentrating hormone receptor 2 | Plasma Membrane | G-protein coupled recepto | |
| chr16 | *BCKDK* | 3 | 119 | 6 | BCKDK | branched chain keto acid dehydrogenase kinase | Cytoplasm | kinase | |
| chr4 | *DCHS2* | 3 | 79 | 6 | DCHS2 | dachsous cadherin-related 2 | Extracellular Space | other | |
| chr12 | *KRT82* | 3 | 122 | 6 | KRT82 | keratin 82 | Cytoplasm | other | |
| chr2 | *APOB* | 2 | 41 | 6 | APOB | apolipoprotein B | Extracellular Space | transporter | mipomersen |
| chr10 | *RP11-107C16.2* | 3 | 111 | 6 | | | | | |
| chr19 | *GLTSCR2* | 3 | 95 | 6 | NOP53 | NOP53 ribosome biogenesis | Cytoplasm | other | |
| chr15 | *ELMO2P1* | 2 | 64 | 6 | ELMO2P1 | engulfment and cell motility 2 pseudogene 1 | Other | other | |
| chr21 | *DSCR9* | 3 | 106 | 6 | DSCR9 | Down syndrome critical region 9 | Other | other | |
| chr2 | *FAP* | 3 | 49 | 6 | FAP | fibroblast activation protein alpha | Cytoplasm | peptidase | RO6874813, RO6874281 |
| chr20 | *TSPY26P* | 3 | 153 | 6 | TSPY26P | testis specific protein Y-linked 26, pseudogene | Other | other | |
| chr7 | *ZNHIT1* | 3 | 117 | 6 | ZNHIT1 | zinc finger HIT-type containing 1 | Nucleus | other | |
| chr22 | *CABP7* | 3 | 100 | 6 | CABP7 | calcium binding protein 7 | Cytoplasm | other | |
| chr12 | *MKRN9P* | 2 | 68 | 6 | MKRN9P | makorin ring finger protein 9, pseudogene | Other | other | |
| chr9 | *C9orf114* | 3 | 105 | 6 | SPOUT1 | SPOUT domain containing methyltransferase 1 | Nucleus | other | |
| chr13 | *RP11-30L8.1* | 3 | 52 | 6 | | | | | |
| chr3 | *RP11-1020A11.2* | 3 | 80 | 6 | | | | | |
| chr19 | *ERVV-2* | 3 | 68 | 6 | ERVV-2 | endogenous retrovirus group V member 2, envelope | Other | other | |
| chr6 | *RP1-60O19.1* | 2 | 42 | 6 | | | | | |
| chr10 | *OR13A1* | 3 | 110 | 6 | OR13A1 | olfactory receptor family 13 subfamily A member 1 | Plasma Membrane | G-protein coupled recepto | |
| chr1 | *C1orf143* | 3 | 59 | 6 | C1orf143 | chromosome 1 open reading frame 143 (putative) | Other | other | |
| chr14 | *RP11-58E21.4* | 3 | 46 | 6 | | | | | |
| chr16 | *ROGDI* | 3 | 80 | 6 | ROGDI | rogdi atypical leucine zipper | Nucleus | other | |
| chrX | *EBP* | 3 | 105 | 6 | EBP | EBP cholestenol delta-isomerase | Cytoplasm | enzyme | SR 31747 |
| chr19 | *VSIG10L* | 3 | 117 | 6 | VSIG10L | V-set and immunoglobulin domain containing 10 like | Nucleus | other | |
| chr1 | *TBC1D3P6* | 3 | 100 | 6 | TBC1D3P6 | TBC1 domain family member 3 pseudogene 6 | Other | other | |
| chr16 | *TGFB1I1* | 3 | 125 | 6 | TGFB1I1 | transforming growth factor beta 1 induced transcript 1 | Nucleus | transcription regulator | |
| chr5 | *EFCAB9* | 3 | 82 | 6 | EFCAB9 | EF-hand calcium binding domain | Cytoplasm | other | |
| chr7 | *PAX4* | 2 | 82 | 6 | PAX4 | paired box 4 | Nucleus | transcription regulator | |
| chr1 | *KCNJ9* | 3 | 116 | 6 | KCNJ9 | potassium inwardly rectifying channel subfamily J member 9 | Plasma Membrane | ion channel | |
| chr15 | *UBE2Q2P1* | 3 | 59 | 6 | UBE2Q2P1 | ubiquitin conjugating enzyme E2 Q2 pseudogene 1 | Other | other | |
| chr9 | *CBWD5* | 2 | 65 | 6 | CBWD5 | COBW domain containing 5 | Other | other | |
| chr1 | *AMIGO1* | 3 | 124 | 6 | AMIGO1 | adhesion molecule with Ig like domain 1 | Plasma Membrane | other | |

| | | | | | | | | | |
|---|---|---|---|---|---|---|---|---|---|
| chr1 | *TARDBP* | 3 | 61 | 6 | TARDBP | TAR DNA binding protein | Nucleus | transcription regulator | |
| chr8 | *SLCO5A1* | 2 | 45 | 6 | SLCO5A1 | solute carrier organic anion transporter family member 5A1 | Plasma Membrane | transporter | |
| chr7 | *PIK3CG* | 3 | 58 | 6 | PIK3CG | phosphatidylinositol-4,5-bisphosphate 3-kinase catalytic subunit gamma | Cytoplasm | kinase | PX-866, SF 1126, pictilisib, CLR457, bimiralisib, buparlisib, PI3Kg inhibitor, dactolisib, tenalisib, pilaralisib |
| chr1 | *CREB3L4* | 2 | 100 | 6 | CREB3L4 | cAMP responsive element binding protein 3 like 4 | Nucleus | transcription regulator | |
| chr2 | *TMEM214* | 2 | 78 | 6 | TMEM214 | transmembrane protein 214 | Extracellular Space | other | |
| chr15 | *SNRPA1* | 3 | 115 | 6 | SNRPA1 | small nuclear ribonucleoprotein polypeptide A' | Nucleus | other | |
| chr16 | *CD2BP2* | 3 | 107 | 6 | CD2BP2 | CD2 cytoplasmic tail binding | Cytoplasm | other | |
| chr11 | *RBM4B* | 3 | 109 | 6 | RBM4B | RNA binding motif protein 4B | Nucleus | other | |
| chrX | *DRP2* | 3 | 75 | 6 | DRP2 | dystrophin related protein 2 | Plasma Membrane | other | |
| chr19 | *ZNF442* | 3 | 91 | 6 | ZNF442 | zinc finger protein 442 | Nucleus | transcription regulator | |
| chr2 | *CYP1B1-AS1* | 3 | 45 | 6 | CYP1B1-AS1 | CYP1B1 antisense RNA 1 | Other | other | |
| chr1 | *HTR6* | 3 | 86 | 6 | HTR6 | 5-hydroxytryptamine receptor 6 | Plasma Membrane | ;-protein coupled recepto | sertindole, iloperidone, WAY-181187, olanzapine, |
| chr5 | *CTD-2631K10.1* | 3 | 72 | 6 | | | | | |
| chr14 | *SLC8A3* | 3 | 50 | 6 | SLC8A3 | solute carrier family 8 member A3 | Plasma Membrane | transporter | |
| chr1 | *CCDC28B* | 3 | 128 | 6 | CCDC28B | coiled-coil domain containing 28B | Cytoplasm | other | |
| chr13 | *LINC00410* | 3 | 56 | 6 | LINC00410 | long intergenic non-protein coding RNA 410 | Other | other | |
| chr2 | *ABCD1P5* | 2 | 282 | 6 | ABCD1P5 | ATP binding cassette subfamily D member 1 pseudogene 5 | Other | other | |
| chr19 | *ELAVL3* | 3 | 102 | 6 | ELAVL3 | ELAV like RNA binding protein 3 | Nucleus | other | |
| chr10 | *C10orf2* | 3 | 93 | 6 | TWNK | twinkle mtDNA helicase | Cytoplasm | enzyme | |
| chr19 | *CTC-232P5.1* | 3 | 74 | 6 | | | | | |
| chr1 | *ADPRHL2* | 3 | 108 | 6 | ADPRHL2 | ADP-ribosylhydrolase like 2 | Cytoplasm | enzyme | |
| chr1 | *BTNL10* | 2 | 119 | 6 | BTNL10 | butyrophilin like 10 | Other | other | |
| chr19 | *DLL3* | 3 | 80 | 6 | DLL3 | delta like canonical Notch ligand | Extracellular Space | other | rovalpituzumab |
| chr22 | *RP5-983L19.2* | 2 | 90 | 6 | | | | | |
| chr10 | *RP11-464F9.21* | 3 | 118 | 6 | | | | | |
| chr19 | *UBE2S* | 3 | 111 | 6 | UBE2S | ubiquitin conjugating enzyme E2 | Nucleus | enzyme | |
| chr10 | *RP11-57H14.4* | 3 | 131 | 6 | | | | | |
| chr4 | *INTS12* | 3 | 73 | 6 | INTS12 | integrator complex subunit 12 | Nucleus | other | |
| chr19 | *ZNF85* | 3 | 60 | 6 | ZNF85 | zinc finger protein 85 | Nucleus | transcription regulator | |
| chr15 | *RP11-697E2.9* | 2 | 114 | 6 | | | | | |
| chr14 | *MED15P1* | 3 | 112 | 6 | MED15P1 | mediator complex subunit 15 pseudogene 1 | Other | other | |
| chr13 | *LINC01043* | 3 | 61 | 6 | LINC01043 | long intergenic non-protein coding RNA 1043 | Other | other | |
| chrX | *GAGE12J* | 2 | 135 | 6 | GE1 (includes oth | G antigen 1 | Other | other | |
| chr7 | *CHPF2* | 3 | 141 | 6 | CHPF2 | chondroitin polymerizing factor 2 | Cytoplasm | enzyme | |
| chr10 | *AL359195.1* | 3 | 70 | 6 | | | | | |
| chr22 | *NUP50* | 2 | 58 | 6 | NUP50 | nucleoporin 50 | Nucleus | transporter | |
| chr2 | *MAPRE3* | 2 | 77 | 5 | MAPRE3 | microtubule associated protein RP/EB family member 3 | Cytoplasm | enzyme | |
| chr16 | *HSF4* | 3 | 128 | 5 | HSF4 | heat shock transcription factor 4 | Nucleus | transcription regulator | |
| chr11 | *TBX10* | 3 | 88 | 5 | TBX10 | T-box transcription factor 10 | Nucleus | transcription regulator | |
| chr1 | *RP5-968D22.1* | 3 | 77 | 5 | | | | | |
| chr19 | *NCCRP1* | 3 | 98 | 5 | NCCRP1 | NCCRP1, F-box associated domain containing | Cytoplasm | other | |
| chr5 | *RP11-93O17.2* | 2 | 151 | 5 | | | | | |
| chr9 | *C9orf129* | 3 | 112 | 5 | C9orf129 | chromosome 9 open reading frame 129 | Other | other | |
| chr12 | *TMEM198B* | 3 | 157 | 5 | TMEM198B | transmembrane protein 198B (pseudogene) | Other | other | |
| chr5 | *CTB-11I22.1* | 3 | 71 | 5 | | | | | |
| chr17 | *GCGR* | 3 | 184 | 5 | GCGR | glucagon receptor | Plasma Membrane | ;-protein coupled recepto | MEDI0382, glucagon |
| chr19 | *ZNF324B* | 3 | 109 | 5 | ZNF324B | zinc finger protein 324B | Other | other | |
| chr17 | *RP11-13K12.5* | 2 | 92 | 5 | | | | | |
| chr3 | *RP11-372E1.4* | 2 | 73 | 5 | | | | | |
| chr22 | *RANBP1* | 2 | 109 | 5 | RANBP1 | RAN binding protein 1 | Nucleus | other | |
| chr19 | *CCDC106* | 3 | 148 | 5 | CCDC106 | coiled-coil domain containing 106 | Extracellular Space | other | |
| chr19 | *IRF3* | 3 | 114 | 5 | IRF3 | interferon regulatory factor 3 | Nucleus | transcription regulator | |
| chr1 | *GJA5* | 2 | 116 | 5 | GJA5 | gap junction protein alpha 5 | Plasma Membrane | transporter | |
| chr1 | *MED18* | 2 | 72 | 5 | MED18 | mediator complex subunit 18 | Nucleus | other | |
| chr9 | *ZBTB26* | 3 | 73 | 5 | ZBTB26 | zinc finger and BTB domain containing 26 | Nucleus | transcription regulator | |
| chr9 | *OBP2A* | 3 | 152 | 5 | OBP2A | odorant binding protein 2A | Extracellular Space | other | |
| chr19 | *RAVER1* | 3 | 87 | 5 | RAVER1 | ribonucleoprotein, PTB binding 1 | Nucleus | other | |
| chr1 | *DMRTB1* | 3 | 111 | 5 | DMRTB1 | DMRT like family B with proline rich C-terminal 1 | Nucleus | transcription regulator | |
| chr21 | *LINC00162* | 3 | 117 | 5 | PICSAR | P38 inhibited cutaneous squamous cell carcinoma | Other | other | |
| chr4 | *S100P* | 2 | 81 | 5 | S100P | S100 calcium binding protein P | Cytoplasm | other | |
| chr12 | *C1RL* | 2 | 72 | 5 | C1RL | complement C1r subcomponent | Extracellular Space | peptidase | |
| chr19 | *SF3A2* | 3 | 156 | 5 | SF3A2 | splicing factor 3a subunit 2 | Nucleus | other | |
| chr1 | *FCGR2A* | 3 | 110 | 5 | FCGR2A | Fc fragment of IgG receptor IIa | Plasma Membrane | transmembrane recepto | IgG |
| chr8 | *MMP16* | 3 | 67 | 5 | MMP16 | matrix metallopeptidase 16 | Extracellular Space | peptidase | marimastat |
| chr22 | *PPP1R26P2* | 2 | 137 | 5 | PPP1R26P2 | protein phosphatase 1 regulatory subunit 26 pseudogene 2 | Other | other | |
| chr3 | *FGD5P1* | 2 | 109 | 5 | FGD5P1 | FYVE, RhoGEF and PH domain containing 5 pseudogene 1 | Other | other | |

| | | | | | | | | | |
|---|---|---|---|---|---|---|---|---|---|
| chr6 | *IL20RA* | 2 | 55 | 5 | IL20RA | interleukin 20 receptor subunit | Plasma Membrane | transmembrane receptor | |
| chr17 | *CTD-2561B21.5* | 3 | 208 | 5 | | | | | |
| chr7 | *NSUN5* | 3 | 122 | 5 | NSUN5 | NOP2/Sun RNA | Nucleus | other | |
| chr17 | *C1QBP* | 3 | 90 | 5 | C1QBP | complement C1q binding protein | Cytoplasm | transcription regulator | |
| chr2 | *AC093642.5* | 2 | 51 | 5 | | | | | |
| chr8 | *VCPIP1* | 2 | 66 | 5 | VCPIP1 | valosin containing protein interacting protein 1 | Cytoplasm | peptidase | |
| chr11 | *MUC6* | 3 | 293 | 5 | MUC6 | mucin 6, oligomeric mucus/gel-forming | Extracellular Space | other | |
| chr2 | *TMEM18* | 2 | 143 | 5 | TMEM18 | transmembrane protein 18 | Nucleus | other | |
| chr11 | *OR51E2* | 2 | 65 | 5 | OR51E2 | olfactory receptor family 51 subfamily E member 2 | Plasma Membrane | G-protein coupled recepto | |
| chr2 | *KLF7* | 2 | 55 | 5 | KLF7 | Kruppel like factor 7 | Nucleus | transcription regulator | |
| chr1 | *DDOST* | 3 | 88 | 5 | DDOST | dolichyl-diphosphooligosaccharide--protein glycosyltransferase non- | Cytoplasm | enzyme | |
| chr5 | *SPATA24* | 3 | 76 | 5 | SPATA24 | spermatogenesis associated 24 | Nucleus | other | |
| chr1 | *RP4-663N10.2* | 3 | 84 | 5 | | | | | |
| chrX | *SLC6A8* | 3 | 136 | 5 | SLC6A8 | solute carrier family 6 member 8 | Cytoplasm | transporter | RGX-202-01 |
| chr15 | *PPIB* | 2 | 88 | 5 | PPIB | peptidylprolyl isomerase B | Cytoplasm | enzyme | |
| chrX | *FAM9A* | 2 | 105 | 5 | FAM9A | family with sequence similarity 9 member A | Nucleus | other | |
| chrX | *TCEAL3* | 2 | 73 | 5 | TCEAL3 | transcription elongation factor A | Other | other | |
| chr1 | *FAM19A3* | 3 | 157 | 5 | TAFA3 | TAFA chemokine like family member 3 | Extracellular Space | other | |
| chr12 | *PRKAB1* | 3 | 100 | 5 | PRKAB1 | protein kinase AMP-activated non-catalytic subunit beta 1 | Nucleus | kinase | |
| chr3 | *RP11-513G11.3* | 2 | 65 | 5 | | | | | |
| chr19 | *ZNF471* | 3 | 61 | 5 | ZNF471 | zinc finger protein 471 | Nucleus | transcription regulator | |
| chr3 | *EHHADH-AS1* | 3 | 62 | 5 | EHHADH-AS1 | EHHADH antisense RNA 1 | Other | other | |
| chr20 | *OGFR-AS1* | 3 | 186 | 5 | OGFR-AS1 | OGFR antisense RNA 1 | Other | other | |
| chr19 | *CACTIN* | 3 | 114 | 5 | CACTIN | cactin, spliceosome C complex subunit | Nucleus | other | |
| chr17 | *C17orf62* | 3 | 121 | 5 | CYBC1 | cytochrome b-245 chaperone 1 | Cytoplasm | other | |
| chr10 | *TMEM254* | 2 | 46 | 5 | TMEM254 | transmembrane protein 254 | Other | other | |
| chr4 | *CXCL13* | 3 | 47 | 5 | CXCL13 | C-X-C motif chemokine ligand 13 | Extracellular Space | cytokine | |
| chr19 | *GPR108* | 3 | 113 | 5 | GPR108 | G protein-coupled receptor 108 | Plasma Membrane | G-protein coupled recepto | |
| chr7 | *AC007566.10* | 2 | 66 | 5 | | | | | |
| chr3 | *RP11-475O23.3* | 3 | 83 | 5 | | | | | |
| chr3 | *ZMAT3* | 2 | 52 | 5 | ZMAT3 | zinc finger matrin-type 3 | Nucleus | other | |
| chr17 | *RP11-51L5.3* | 2 | 49 | 5 | | | | | |
| chr4 | *IGFBP7* | 2 | 54 | 5 | IGFBP7 | insulin like growth factor binding protein 7 | Extracellular Space | transporter | |
| chr7 | *UBE2D4* | 3 | 109 | 5 | UBE2D4 | ubiquitin conjugating enzyme E2 D4 (putative) | Other | enzyme | |
| chr20 | *C20orf96* | 3 | 92 | 5 | C20orf96 | chromosome 20 open reading frame 96 | Other | other | |
| chr17 | *KRT25* | 2 | 45 | 5 | KRT25 | keratin 25 | Cytoplasm | other | |
| chr11 | *FGF3* | 3 | 175 | 5 | FGF3 | fibroblast growth factor 3 | Extracellular Space | growth factor | |
| chr15 | *RAD51-AS1* | 3 | 53 | 5 | RAD51-AS1 | RAD51 antisense RNA 1 | Other | other | |
| chr19 | *EHD2* | 3 | 85 | 5 | EHD2 | EH domain containing 2 | Nucleus | other | |
| chr19 | *MRI1* | 3 | 107 | 5 | MRI1 | methylthioribose-1-phosphate isomerase 1 | Cytoplasm | translation regulator | |
| chr17 | *SCPEP1* | 2 | 52 | 5 | SCPEP1 | serine carboxypeptidase 1 | Cytoplasm | peptidase | |
| chr20 | *RP5-955M13.4* | 3 | 110 | 5 | | | | | |
| chr1 | *RP11-365O16.6* | 3 | 90 | 5 | | | | | |
| chr14 | *AKAP6* | 2 | 62 | 5 | AKAP6 | A-kinase anchoring protein 6 | Nucleus | other | |
| chr4 | *BOD1L1* | 2 | 51 | 5 | BOD1L1 | biorientation of chromosomes in cell division 1 like 1 | Nucleus | other | |
| chr9 | *TDRD7* | 2 | 40 | 5 | TDRD7 | tudor domain containing 7 | Cytoplasm | other | |
| chr6 | *DNPH1* | 3 | 139 | 5 | DNPH1 | 2'-deoxynucleoside 5'-phosphate N-hydrolase 1 | Nucleus | enzyme | |
| chr16 | *KIAA0895L* | 3 | 124 | 5 | KIAA0895L | KIAA0895 like | Other | other | |
| chr12 | *B3GNT4* | 3 | 73 | 5 | B3GNT4 | UDP-GlcNAc:betaGal beta-1,3-N-acetylglucosaminyltransferase 4 | Plasma Membrane | enzyme | |
| chr18 | *AQP4-AS1* | 2 | 43 | 5 | AQP4-AS1 | AQP4 antisense RNA 1 | Other | other | |
| chr12 | *RP11-25I15.3* | 3 | 104 | 5 | | | | | |
| chr1 | *HAX1* | 3 | 81 | 5 | HAX1 | HCLS1 associated protein X-1 | Cytoplasm | other | |
| chr10 | *RP11-432J24.5* | 3 | 154 | 5 | | | | | |
| chr1 | *RP4-683M8.2* | 3 | 80 | 5 | | | | | |
| chr10 | *GOLGA7B* | 3 | 88 | 5 | GOLGA7B | golgin A7 family member B | Other | other | |
| chr17 | *RSAD1* | 2 | 99 | 5 | RSAD1 | radical S-adenosyl methionine domain containing 1 | Cytoplasm | other | |
| chr20 | *REM1* | 3 | 117 | 5 | REM1 | RRAD and GEM like GTPase 1 | Other | enzyme | |
| chr2 | *TMEM177* | 3 | 131 | 5 | TMEM177 | transmembrane protein 177 | Cytoplasm | other | |
| chr4 | *C4orf47* | 2 | 49 | 5 | C4orf47 | chromosome 4 open reading frame 47 | Cytoplasm | other | |

| | | | | | | | | | |
|---|---|---|---|---|---|---|---|---|---|
| chrX | *AR* | 3 | 96 | 5 | AR | androgen receptor | Nucleus | d-dependent nuclear rec | galeterone, nandrolone phenpropionate, cyproterone acetate/ethinyl estradiol, testosterone propionate, testosterone cypionate, EPI-506, drospirenone, methyltestosterone, TRC253, AZD3514, bicalutamide/goserelin, apalutamide, fluoxymesterone/tamoxifen, ostarine, danazol, GSK2881078, testosterone enanthate, darolutamide, fluoxymesterone, [illegible] |
| chr6 | *FAXC* | 2 | 51 | 5 | FAXC | failed axon connections homolog, metaxin like GST domain | Other | other | |
| chr10 | *RP11-264E18.1* | 2 | 61 | 5 | | | | | |
| chrX | *DUSP9* | 3 | 166 | 5 | DUSP9 | dual specificity phosphatase 9 | Nucleus | phosphatase | |
| chr7 | *AC005077.5* | 2 | 140 | 5 | | | | | |
| chr17 | *KLHL11* | 3 | 70 | 5 | KLHL11 | kelch like family member 11 | Cytoplasm | other | |
| chr4 | *THEGL* | 2 | 57 | 5 | THEGL | theg spermatid protein like | Other | other | |
| chr12 | *ARL1* | 3 | 50 | 5 | ARL1 | ADP ribosylation factor like | Cytoplasm | enzyme | |
| chr12 | *HIGD1C* | 3 | 88 | 5 | HIGD1C | HIG1 hypoxia inducible domain family member 1C | Other | other | |
| chr7 | *INSIG1* | 2 | 65 | 5 | INSIG1 | insulin induced gene 1 | Cytoplasm | other | |
| chr1 | *PDC* | 2 | 76 | 5 | PDC | phosducin | Cytoplasm | other | |
| chr8 | *YWHAZ* | 2 | 63 | 5 | YWHAZ | tyrosine 3-monooxygenase/tryptophan 5-monooxygenase activation protein zeta | Cytoplasm | enzyme | |
| chr5 | *SPDL1* | 2 | 85 | 5 | SPDL1 | spindle apparatus coiled-coil | Nucleus | other | |
| chr2 | *GPBAR1* | 2 | 95 | 5 | GPBAR1 | G protein-coupled bile acid | Plasma Membrane | G-protein coupled recepto | |
| chrX | *MAP7D3* | 2 | 45 | 5 | MAP7D3 | MAP7 domain containing 3 | Cytoplasm | other | |
| chr16 | *TBL3* | 3 | 142 | 5 | TBL3 | transducin beta like 3 | Cytoplasm | peptidase | |
| chr12 | *C12orf44* | 3 | 229 | 5 | ATG101 | autophagy related 101 | Cytoplasm | other | |
| chr5 | *C5orf15* | 2 | 85 | 5 | C5orf15 | chromosome 5 open reading frame 15 | Other | other | |
| chr2 | *GGT8P* | 3 | 104 | 5 | GGT8P | gamma-glutamyltransferase 8 pseudogene | Other | other | |
| chr15 | *DNM1P32* | 2 | 96 | 5 | DNM1P32 | dynamin 1 pseudogene 32 | Other | other | |
| chr15 | *RP11-553E24.2* | 3 | 89 | 5 | | | | | |
| chr22 | *ABCD1P4* | 2 | 137 | 5 | ABCD1P4 | ATP binding cassette subfamily D member 1 pseudogene 4 | Other | other | |
| chr19 | *CTD-3105H18.16* | 2 | 59 | 5 | | | | | |
| chr2 | *HIBCH* | 2 | 61 | 5 | HIBCH | 3-hydroxyisobutyryl-CoA | Cytoplasm | enzyme | |
| chr10 | *INA* | 3 | 110 | 5 | INA | internexin neuronal intermediate filament protein alpha | Cytoplasm | other | |
| chr6 | *HDDC2* | 2 | 59 | 5 | HDDC2 | HD domain containing 2 | Cytoplasm | other | |
| chr5 | *NUDCD2* | 2 | 66 | 5 | NUDCD2 | NudC domain containing 2 | Cytoplasm | other | |
| chr3 | *SEC62* | 3 | 101 | 5 | SEC62 | SEC62 homolog, preprotein translocation factor | Cytoplasm | transporter | |
| chr8 | *RP11-661A12.12* | 2 | 104 | 5 | | | | | |
| chr19 | *CTD-3131K8.2* | 3 | 113 | 5 | | | | | |
| chr17 | *CSH1* | 3 | 95 | 5 | CSH1/CSH2 | chorionic somatomammotropin hormone 1 | Extracellular Space | other | |
| chr15 | *RP11-69G7.1* | 3 | 48 | 5 | | | | | |
| chr2 | *GPD2* | 2 | 40 | 5 | GPD2 | glycerol-3-phosphate dehydrogenase 2 | Cytoplasm | enzyme | |
| chr1 | *UBXN10* | 3 | 54 | 5 | UBXN10 | UBX domain protein 10 | Extracellular Space | other | |
| chr6 | *RP11-351J23.2* | 2 | 102 | 5 | | | | | |
| chr6 | *FAM46A* | 2 | 49 | 5 | TENT5A | terminal nucleotidyltransferase | Other | other | |
| chr2 | *RHBDD1* | 3 | 87 | 5 | RHBDD1 | rhomboid domain containing 1 | Extracellular Space | peptidase | |
| chrX | *GTPBP6* | 3 | 83 | 5 | GTPBP6 | GTP binding protein 6 (putative) | Other | other | |
| chr9 | *RP11-318K12.1* | 2 | 58 | 5 | | | | | |
| chr10 | *LINC00200* | 3 | 68 | 5 | LINC00200 | long intergenic non-protein coding RNA 200 | Other | other | |
| chr15 | *RP13-395E19.4* | 2 | 40 | 5 | | | | | |
| chr12 | *DEPDC4* | 2 | 41 | 5 | DEPDC4 | DEP domain containing 4 | Other | other | |
| chr15 | *RP11-403B2.6* | 3 | 111 | 5 | | | | | |
| chr1 | *PPOX* | 3 | 129 | 5 | PPOX | protoporphyrinogen oxidase | Cytoplasm | enzyme | |
| chr10 | *FAM204A* | 2 | 73 | 5 | FAM204A | family with sequence similarity 204 member A | Other | other | |
| chr13 | *SLITRK5* | 2 | 66 | 5 | SLITRK5 | SLIT and NTRK like family | Plasma Membrane | other | |
| chr1 | *RP4-798P15.3* | 3 | 74 | 5 | | | | | |
| chr2 | *AC098820.2* | 3 | 44 | 5 | | | | | |
| chr19 | *ZNF577* | 3 | 46 | 5 | ZNF577 | zinc finger protein 577 | Other | other | |
| chr7 | *COL1A2* | 3 | 62 | 5 | COL1A2 | collagen type I alpha 2 chain | Extracellular Space | other | collagenase clostridium |
| chr1 | *S100A14* | 2 | 130 | 5 | S100A14 | S100 calcium binding protein A14 | Cytoplasm | other | |

| | | | | | | | | | |
|---|---|---|---|---|---|---|---|---|---|
| chr1 | *RP11-545A16.3* | 3 | 162 | 5 | | | | | |
| chr11 | *MS4A12* | 3 | 61 | 5 | MS4A12 | membrane spanning 4-domains | Plasma Membrane | other | |
| chr15 | *RP11-32B5.7* | 3 | 109 | 5 | | | | | |
| chr3 | *RP11-368I23.2* | 3 | 86 | 5 | | | | | |
| chr2 | *TUBA4A* | 2 | 83 | 5 | TUBA4A | tubulin alpha 4a | Cytoplasm | other | docetaxel/gemcitabin e/vincristine, cyclophosphamide/e pirubicin/5-fluorouracil/paclitaxel, bevacizumab/paclitax el/topotecan, colchicine/probeneci d, bevacizumab/doceta xel/prednisone/thalid omide/zoledronic acid, gemcitabine/paclitax el, L-asparaginase/daunor ubicin/dexamethason e/vincristine, prednisone/vincristin e, docetaxel/5-fluorouracil, vinflunine, L-[illegible] |
| chr7 | *RP11-61L23.2* | 2 | 74 | 5 | | | | | |
| chr17 | *RP11-112H10.4* | 2 | 66 | 5 | | | | | |
| chrX | *ATP6AP1* | 2 | 97 | 5 | ATP6AP1 | ATPase H+ transporting accessory protein 1 | Cytoplasm | transporter | |
| chr1 | *RP11-46A10.2* | 2 | 119 | 5 | | | | | |
| chr2 | *AC093326.3* | 3 | 98 | 5 | | | | | |
| chr1 | *CELA2A* | 3 | 78 | 5 | CELA2A | chymotrypsin like elastase 2A | Extracellular Space | peptidase | |
| chr19 | *AES* | 3 | 96 | 5 | TLE5 | TLE family member 5, transcriptional modulator | Nucleus | transcription regulator | |
| chrX | *EIF1AX* | 2 | 44 | 5 | EIF1AX | eukaryotic translation initiation factor 1A X-linked | Cytoplasm | translation regulator | |
| chr17 | *AC004562.1* | 2 | 50 | 5 | | | | | |
| chr2 | *KLHL30* | 3 | 174 | 5 | KLHL30 | kelch like family member 30 | Other | other | |
| chr1 | *CCDC181* | 2 | 95 | 5 | CCDC181 | coiled-coil domain containing 181 | Nucleus | other | |
| chr3 | *TRMT10C* | 3 | 86 | 5 | TRMT10C | tRNA methyltransferase 10C, mitochondrial RNase P subunit | Cytoplasm | enzyme | |
| chr1 | *S100A4* | 3 | 105 | 5 | S100A4 | S100 calcium binding protein A4 | Cytoplasm | other | |
| chr19 | *SBK3* | 2 | 116 | 5 | SBK3 | SH3 domain binding kinase family member 3 | Other | other | |
| chr2 | *FAHD2CP* | 2 | 99 | 5 | FAHD2CP | fumarylacetoacetate hydrolase domain containing 2C, | Other | other | |
| chr5 | *RP11-2N5.1* | 2 | 44 | 5 | | | | | |
| chr9 | *RP11-473E2.2* | 3 | 170 | 5 | | | | | |
| chr7 | *POLR2J* | 3 | 172 | 5 | POLR2J | RNA polymerase II subunit J | Nucleus | enzyme | |
| chr11 | *BIRC3* | 3 | 164 | 5 | BIRC3 | baculoviral IAP repeat containing | Cytoplasm | enzyme | birinapant |
| chr3 | *LINC00880* | 3 | 74 | 5 | LINC00880 | long intergenic non-protein coding RNA 880 | Other | other | |
| chr1 | *C1orf51* | 2 | 101 | 5 | CIART | circadian associated repressor of transcription | Nucleus | other | |
| chr1 | *PAQR6* | 3 | 145 | 5 | PAQR6 | progestin and adipoQ receptor family member 6 | Plasma Membrane | other | |
| chrX | *SSX5* | 3 | 54 | 5 | SSX5 | SSX family member 5 | Other | other | |
| chr15 | *SPRED1* | 3 | 50 | 5 | SPRED1 | sprouty related EVH1 domain containing 1 | Plasma Membrane | other | |
| chr6 | *HCP5* | 2 | 48 | 5 | HCP5 | HLA complex P5 | Other | other | |
| chr17 | *RP11-849F2.5* | 2 | 62 | 5 | | | | | |
| chr11 | *CTD-2210P24.6* | 3 | 80 | 5 | | | | | |
| chrX | *GAB3* | 2 | 39 | 5 | GAB3 | GRB2 associated binding protein | Other | other | |
| chr2 | *AC023115.2* | 3 | 49 | 5 | | | | | |
| chr9 | *RP11-262H14.3* | 2 | 75 | 5 | | | | | |
| chr19 | *CTD-2207O23.3* | 3 | 70 | 5 | | | | | |
| chr1 | *S100A7* | 3 | 143 | 5 | S100A7 | S100 calcium binding protein A7 | Cytoplasm | other | |
| chr19 | *CTD-2325M2.1* | 3 | 53 | 5 | | | | | |
| chr19 | *PPP1R13L* | 3 | 106 | 5 | PPP1R13L | protein phosphatase 1 regulatory subunit 13 like | Nucleus | transcription regulator | |
| chr8 | *SHARPIN* | 3 | 117 | 5 | SHARPIN | SHANK associated RH domain interactor | Plasma Membrane | other | |
| chr13 | *LINC00565* | 3 | 72 | 5 | LINC00565 | long intergenic non-protein coding RNA 565 | Other | other | |
| chr15 | *POTEB2* | 3 | 117 | 5 | POTEB/POTEB2 | POTE ankyrin domain family member B2 | Other | other | |
| chr1 | *RP11-21J7.1* | 2 | 94 | 5 | | | | | |
| chr9 | *RP11-305L7.6* | 2 | 79 | 5 | | | | | |
| chr15 | *CHEK2P2* | 3 | 289 | 5 | CHEK2P2 | checkpoint kinase 2 pseudogene | Other | other | |
| chr1 | *MYOG* | 2 | 118 | 5 | MYOG | myogenin | Nucleus | transcription regulator | |
| chr2 | *SNRNP27* | 2 | 66 | 5 | SNRNP27 | small nuclear ribonucleoprotein U4/U6.U5 subunit 27 | Nucleus | other | |
| chr17 | *AOC2* | 3 | 108 | 5 | AOC2 | amine oxidase copper containing | Plasma Membrane | enzyme | |
| chr19 | *AC006262.6* | 2 | 70 | 5 | | | | | |

| | | | | | | | | | |
|---|---|---|---|---|---|---|---|---|---|
| chr8 | *GPIHBP1* | 3 | 157 | 5 | GPIHBP1 | glycosylphosphatidylinositol anchored high density lipoprotein binding protein 1 | Plasma Membrane | other | |
| chr10 | *SAR1A* | 3 | 106 | 5 | SAR1A | secretion associated Ras related GTPase 1A | Cytoplasm | enzyme | |
| chr8 | *LINC00964* | 3 | 40 | 5 | LINC00964 | long intergenic non-protein coding RNA 964 | Other | other | |
| chr3 | *RP11-483E7.1* | 2 | 64 | 5 | | | | | |
| chr17 | *H3F3B* | 2 | 59 | 5 | H3-3A/H3-3B | H3.3 histone A | Nucleus | other | |
| chr9 | *SURF2* | 3 | 107 | 5 | SURF2 | surfeit 2 | Nucleus | other | |
| chr5 | *RP11-1026M7.2* | 2 | 61 | 5 | | | | | |
| chr4 | *HELQ* | 2 | 44 | 5 | HELQ | helicase, POLQ like | Nucleus | enzyme | |
| chr11 | *NDUFS8* | 3 | 83 | 5 | NDUFS8 | NADH:ubiquinone oxidoreductase core subunit S8 | Cytoplasm | enzyme | |
| chr7 | *CTB-107G13.1* | 3 | 61 | 5 | | | | | |
| chr2 | *CIR1* | 2 | 43 | 5 | CIR1 | corepressor interacting with | Nucleus | transcription regulator | |
| chr3 | *MAP6D1* | 2 | 83 | 5 | MAP6D1 | MAP6 domain containing 1 | Cytoplasm | other | |
| chr4 | *C4orf51* | 2 | 45 | 5 | C4orf51 | chromosome 4 open reading frame 51 | Other | other | |
| chr7 | *STARD3NL* | 3 | 88 | 5 | STARD3NL | STARD3 N-terminal like | Cytoplasm | other | |
| chr7 | *AC006372.6* | 2 | 111 | 5 | | | | | |
| chr4 | *RP11-554D13.1* | 2 | 47 | 5 | | | | | |
| chr12 | *GPN3* | 3 | 56 | 5 | GPN3 | GPN-loop GTPase 3 | Cytoplasm | other | |
| chr1 | *RP3-395M20.7* | 3 | 190 | 5 | | | | | |
| chr11 | *AKIP1* | 2 | 98 | 5 | AKIP1 | A-kinase interacting protein 1 | Nucleus | other | |
| chr7 | *AZGP1* | 3 | 90 | 5 | AZGP1 | alpha-2-glycoprotein 1, zinc- | Extracellular Space | transporter | |
| chr19 | *NMRK2* | 3 | 98 | 5 | NMRK2 | nicotinamide riboside kinase 2 | Plasma Membrane | kinase | |
| chr9 | *RP11-381O7.3* | 3 | 87 | 5 | | | | | |
| chr7 | *AC099342.1* | 2 | 59 | 5 | | | | | |
| chr4 | *AASDH* | 2 | 66 | 5 | AASDH | aminoadipate-semialdehyde dehydrogenase | Other | enzyme | |
| chr1 | *CCDC23* | 3 | 52 | 5 | SVBP | small vasohibin binding protein | Other | other | |
| chr1 | *RP5-1166F10.1* | 2 | 69 | 5 | | | | | |
| chr9 | *ASB6* | 3 | 153 | 5 | ASB6 | ankyrin repeat and SOCS box containing 6 | Nucleus | transcription regulator | |
| chr11 | *RNF141* | 3 | 67 | 5 | RNF141 | ring finger protein 141 | Other | enzyme | |
| chr11 | *RP11-315O6.2* | 3 | 49 | 5 | | | | | |
| chr16 | *NPIPB9* | 2 | 73 | 5 | PB15 (includes oth | nuclear pore complex interacting protein family member B15 | Nucleus | other | |
| chr12 | *GDF11* | 3 | 166 | 5 | GDF11 | growth differentiation factor 11 | Extracellular Space | growth factor | |
| chr11 | *RP11-554A11.7* | 2 | 150 | 5 | | | | | |
| chr2 | *AC064853.3* | 3 | 45 | 5 | | | | | |
| chr19 | *HOMER3* | 3 | 79 | 5 | HOMER3 | homer scaffold protein 3 | Plasma Membrane | other | |
| chr4 | *C4orf36* | 2 | 49 | 5 | C4orf36 | chromosome 4 open reading frame 36 | Other | other | |
| chrX | *VBP1* | 2 | 72 | 5 | VBP1 | VHL binding protein 1 | Cytoplasm | other | |
| chr20 | *HAR1B* | 3 | 240 | 5 | HAR1B | highly accelerated region 1B | Other | other | |
| chr5 | *RP11-497H16.8* | 2 | 117 | 5 | | | | | |
| chr16 | *TBX6* | 3 | 86 | 5 | TBX6 | T-box transcription factor 6 | Nucleus | transcription regulator | |
| chr2 | *ZNF514* | 3 | 69 | 5 | ZNF514 | zinc finger protein 514 | Nucleus | transcription regulator | |
| chr13 | *RAP2A* | 2 | 84 | 5 | RAP2A | RAP2A, member of RAS oncogene family | Plasma Membrane | enzyme | |
| chr4 | *RP11-529E10.7* | 2 | 77 | 5 | | | | | |
| chr8 | *LETM2* | 2 | 64 | 5 | LETM2 | leucine zipper and EF-hand containing transmembrane | Other | other | |
| chr20 | *ZFAS1* | 3 | 62 | 5 | ZFAS1 | ZNFX1 antisense RNA 1 | Other | other | |
| chr8 | *RP11-162D9.3* | 2 | 54 | 5 | | | | | |
| chr17 | *RUNDC3A* | 3 | 95 | 5 | RUNDC3A | RUN domain containing 3A | Cytoplasm | other | |
| chr11 | *SPCS2* | 2 | 60 | 5 | SPCS2 | signal peptidase complex subunit | Cytoplasm | other | |
| chr17 | *CTC-296K1.4* | 2 | 71 | 5 | | | | | |
| chr11 | *MS4A1* | 3 | 60 | 5 | MS4A1 | membrane spanning 4-domains A1 | Plasma Membrane | other | bendamustine/ofatumumab, idelalisib/ofatumumab, IGM-2323, methotrexate/ofatumumab, bendamustine/obinutuzumab, cytarabine/fludarabine phosphate/oxaliplatin/rituximab, bendamustine/rituximab, GEN3013, chlorambucil/obinutuzumab, cyclophosphamide/dexamethasone/rituximab, anti-CD20 antibody, idelalisib/rituximab, chlorambucil/mitoxantrone/prednisone/ritu[illegible] |
| chr18 | *ONECUT2* | 2 | 78 | 5 | ONECUT2 | one cut homeobox 2 | Nucleus | transcription regulator | |
| chr19 | *DPRX* | 2 | 77 | 5 | DPRX | divergent-paired related | Other | other | |
| chr14 | *FLRT2* | 2 | 64 | 5 | FLRT2 | fibronectin leucine rich transmembrane protein 2 | Plasma Membrane | other | |
| chr19 | *CAPNS1* | 2 | 113 | 5 | CAPNS1 | calpain small subunit 1 | Cytoplasm | peptidase | |

| | | | | | | | | | |
|---|---|---|---|---|---|---|---|---|---|
| chr17 | *CBX4* | 3 | 123 | 5 | CBX4 | chromobox 4 | Nucleus | transcription regulator | |
| chr1 | *DDX20* | 2 | 72 | 5 | DDX20 | DEAD-box helicase 20 | Nucleus | transcription regulator | |
| chr8 | *RP11-363L24.3* | 2 | 41 | 5 | | | | | |
| chr1 | *PLA2G2A* | 3 | 67 | 5 | PLA2G2A | phospholipase A2 group IIA | Cytoplasm | enzyme | varespladib, indomethacin, varespladib methyl |
| chr9 | *AL360004.1* | 2 | 149 | 5 | | | | | |
| chr17 | *NGFR* | 3 | 61 | 5 | NGFR | nerve growth factor receptor | Plasma Membrane | transmembrane receptor | LM11A-31 |
| chr5 | *MCIDAS* | 3 | 65 | 5 | MCIDAS | multiciliate differentiation and DNA synthesis associated cell cycle protein | Nucleus | other | |
| chr6 | *RP11-753B14.1* | 3 | 58 | 5 | | | | | |
| chr13 | *IFT88* | 2 | 76 | 5 | IFT88 | intraflagellar transport 88 | Cytoplasm | other | |
| chr2 | *AC114788.2* | 2 | 79 | 5 | | | | | |
| chr1 | *RP5-1125N11.1* | 3 | 83 | 5 | | | | | |
| chr8 | *RPL23AP53* | 3 | 71 | 5 | RPL23AP53 | ribosomal protein L23a pseudogene 53 | Other | other | |
| chr7 | *RP11-196D18.1* | 3 | 80 | 5 | | | | | |
| chr7 | *EN2* | 3 | 166 | 5 | EN2 | engrailed homeobox 2 | Nucleus | transcription regulator | |
| chr1 | *PIGV* | 3 | 56 | 5 | PIGV | phosphatidylinositol glycan anchor biosynthesis class V | Cytoplasm | enzyme | |
| chr18 | *SERPINB11* | 3 | 39 | 5 | SERPINB11 | serpin family B member 11 (gene/pseudogene) | Cytoplasm | other | |
| chr6 | *BTN2A2* | 3 | 47 | 5 | BTN2A2 | butyrophilin subfamily 2 member | Cytoplasm | other | |
| chr1 | *RP5-1007M22.2* | 3 | 107 | 5 | | | | | |
| chr2 | *AC053503.11* | 3 | 101 | 5 | | | | | |
| chr17 | *CCL18* | 3 | 51 | 5 | CCL18 | C-C motif chemokine ligand 18 | Extracellular Space | cytokine | |
| chr3 | *RP11-174O3.4* | 2 | 90 | 5 | | | | | |
| chrX | *APEX2* | 3 | 111 | 5 | APEX2 | apurinic/apyrimidinic endodeoxyribonuclease 2 | Nucleus | enzyme | |
| chr11 | *RP11-286N22.10* | 3 | 131 | 5 | | | | | |
| chr16 | *RP11-505K9.3* | 2 | 125 | 5 | | | | | |
| chr22 | *SEPT5* | 3 | 161 | 5 | SEPTIN5 | septin 5 | Cytoplasm | enzyme | |
| chr1 | *WLS* | 2 | 45 | 5 | WLS | Wnt ligand secretion mediator | Cytoplasm | other | |
| chr3 | *SSR3* | 2 | 85 | 5 | SSR3 | signal sequence receptor subunit | Cytoplasm | other | |
| chr11 | *RP11-45A12.2* | 3 | 62 | 5 | | | | | |
| chr19 | *TBC1D17* | 3 | 112 | 5 | TBC1D17 | TBC1 domain family member 17 | Cytoplasm | other | |
| chr15 | *PHGR1* | 2 | 88 | 5 | PHGR1 | proline, histidine and glycine rich | Other | other | |
| chr6 | *TREM2* | 3 | 89 | 5 | TREM2 | triggering receptor expressed on myeloid cells 2 | Plasma Membrane | transmembrane receptor | anti-TREM2 antibody |
| chr1 | *RHOU* | 3 | 92 | 5 | RHOU | ras homolog family member U | Cytoplasm | enzyme | |
| chrX | *MAGIX* | 3 | 171 | 5 | MAGIX | MAGI family member, X-linked | Other | other | |
| chr3 | *MARK2P14* | 2 | 84 | 5 | MARK2P14 | microtubule affinity regulating kinase 2 pseudogene 14 | Other | other | |
| chr20 | *TUBB1* | 3 | 186 | 5 | TUBB1 | tubulin beta 1 class VI | Cytoplasm | other | paclitaxel/pertuzumab/trastuzumab, docetaxel/gemcitabine/vincristine, cyclophosphamide/epirubicin/5-fluorouracil/paclitaxel, bevacizumab/paclitaxel/topotecan, bortezomib/paclitaxel, colchicine/probenecid, bevacizumab/docetaxel/prednisone/thalidomide/zoledronic acid, gemcitabine/paclitaxel, simotaxel, L-asparaginase/daunorubicin/dexamethasone/vincristine, [illegible] |
| chr13 | *FREM2* | 2 | 63 | 5 | FREM2 | FRAS1 related extracellular | Extracellular Space | other | |
| chr1 | *NENF* | 3 | 74 | 5 | NENF | neudesin neurotrophic factor | Extracellular Space | growth factor | |
| chr5 | *RP11-184E9.1* | 3 | 113 | 5 | | | | | |
| chr11 | *LRRC55* | 2 | 98 | 5 | LRRC55 | leucine rich repeat containing 55 | Plasma Membrane | ion channel | |
| chr5 | *PTCD2* | 3 | 50 | 5 | PTCD2 | pentatricopeptide repeat domain | Cytoplasm | other | |
| chr2 | *AC105053.3* | 3 | 82 | 5 | | | | | |
| chr10 | *IKZF5* | 2 | 47 | 5 | IKZF5 | IKAROS family zinc finger 5 | Nucleus | transcription regulator | |
| chr1 | *ATP5F1* | 2 | 59 | 5 | ATP5PB | ATP synthase peripheral stalk-membrane subunit b | Cytoplasm | transporter | |
| chr11 | *IL10RA* | 2 | 71 | 5 | IL10RA | interleukin 10 receptor subunit | Plasma Membrane | transmembrane receptor | |
| chr17 | *TMEM97* | 2 | 57 | 5 | TMEM97 | transmembrane protein 97 | Extracellular Space | other | CT1812 |
| chr20 | *OCSTAMP* | 3 | 121 | 5 | OCSTAMP | osteoclast stimulatory transmembrane protein | Other | other | |
| chr17 | *MRM1* | 2 | 94 | 5 | MRM1 | mitochondrial rRNA methyltransferase 1 | Cytoplasm | enzyme | |
| chr8 | *FAM90A6P* | 2 | 117 | 5 | FAM90A6P | family with sequence similarity 90 member A6, pseudogene | Other | other | |
| chr17 | *YBX2* | 3 | 109 | 5 | YBX2 | Y-box binding protein 2 | Cytoplasm | translation regulator | |
| chr11 | *C11orf74* | 2 | 41 | 5 | C11orf74 | chromosome 11 open reading frame 74 | Extracellular Space | other | |
| chr10 | *ZNF503-AS2* | 3 | 118 | 5 | ZNF503-AS2 | ZNF503 antisense RNA 2 | Other | other | |

| | | | | | | | | | |
|---|---|---|---|---|---|---|---|---|---|
| chr19 | AC006277.2 | 2 | 58 | 5 | | | | | |
| chr20 | OSER1 | 2 | 53 | 5 | OSER1 | oxidative stress responsive serine rich 1 | Other | enzyme | |
| chr6 | KLHDC3 | 3 | 93 | 5 | KLHDC3 | kelch domain containing 3 | Cytoplasm | other | |
| chrX | NCBP2L | 2 | 49 | 5 | NCBP2L | nuclear cap binding protein subunit 2 like | Other | other | |
| chr17 | WFIKKN2 | 3 | 125 | 5 | WFIKKN2 | WAP, follistatin/kazal, immunoglobulin, kunitz and netrin domain containing 2 | Extracellular Space | other | |
| chr22 | SEC14L4 | 2 | 59 | 5 | SEC14L4 | SEC14 like lipid binding 4 | Other | transporter | |
| chr11 | CD81 | 3 | 140 | 5 | CD81 | CD81 molecule | Plasma Membrane | other | |
| chr16 | VPS9D1 | 3 | 82 | 5 | VPS9D1 | VPS9 domain containing 1 | Other | transporter | |
| chr16 | NME4 | 2 | 97 | 5 | NME4 | NME/NM23 nucleoside diphosphate kinase 4 | Cytoplasm | kinase | |
| chr12 | KRT5 | 3 | 187 | 5 | KRT5 | keratin 5 | Cytoplasm | other | |
| chr14 | DLK1 | 2 | 85 | 5 | DLK1 | delta like non-canonical Notch ligand 1 | Extracellular Space | other | |
| chr16 | ATXN2L | 3 | 62 | 5 | ATXN2L | ataxin 2 like | Nucleus | other | |
| chr7 | RP11-514P8.7 | 3 | 218 | 5 | | | | | |
| chr1 | DUSP12 | 3 | 68 | 5 | DUSP12 | dual specificity phosphatase 12 | Nucleus | phosphatase | |
| chr1 | CDKN2C | 2 | 45 | 5 | CDKN2C | cyclin dependent kinase inhibitor | Nucleus | transcription regulator | |
| chr11 | RP11-136I14.5 | 2 | 41 | 5 | | | | | |
| chr11 | RP11-264E20.1 | 3 | 107 | 5 | | | | | |
| chr10 | MSS51 | 2 | 88 | 5 | MSS51 | MSS51 mitochondrial translational activator | Other | enzyme | |
| chr9 | SURF6 | 3 | 92 | 5 | SURF6 | surfeit 6 | Nucleus | other | |
| chr4 | GPRIN3 | 3 | 65 | 5 | GPRIN3 | GPRIN family member 3 | Other | other | |
| chr12 | RP11-417L19.4 | 3 | 77 | 5 | | | | | |
| chr5 | CTD-2227I18.1 | 2 | 48 | 5 | | | | | |
| chr9 | RP11-326F20.5 | 3 | 87 | 5 | | | | | |
| chr1 | NEK2 | 2 | 58 | 5 | NEK2 | NIMA related kinase 2 | Cytoplasm | kinase | |
| chr13 | PCDH17 | 3 | 87 | 5 | PCDH17 | protocadherin 17 | Plasma Membrane | other | |
| chr1 | THEM5 | 2 | 113 | 5 | THEM5 | thioesterase superfamily member | Cytoplasm | enzyme | |
| chrX | FRMPD3 | 3 | 82 | 5 | FRMPD3 | FERM and PDZ domain | Cytoplasm | other | |
| chr6 | GPRC6A | 3 | 75 | 5 | GPRC6A | G protein-coupled receptor class C group 6 member A | Plasma Membrane | G-protein coupled recepto | |
| chr16 | RP11-473M20.5 | 2 | 75 | 5 | | | | | |
| chr16 | RP11-21B23.2 | 2 | 64 | 5 | | | | | |
| chr19 | SIGLEC9 | 2 | 82 | 5 | SIGLEC9 | sialic acid binding Ig like lectin 9 | Plasma Membrane | other | |
| chr14 | CTD-2509G16.5 | 2 | 61 | 5 | | | | | |
| chr22 | C22orf43 | 2 | 64 | 5 | DRICH1 | aspartate rich 1 | Other | other | |
| chrX | GUCY2F | 2 | 44 | 5 | GUCY2F | guanylate cyclase 2F, retinal | Plasma Membrane | kinase | |
| chr11 | CHORDC1 | 2 | 131 | 5 | CHORDC1 | cysteine and histidine rich domain containing 1 | Other | other | |
| chr19 | ZNF560 | 2 | 62 | 5 | ZNF560 | zinc finger protein 560 | Nucleus | transcription regulator | |
| chr21 | AP000472.2 | 2 | 63 | 5 | | | | | |
| chr11 | UCP2 | 3 | 98 | 5 | UCP2 | uncoupling protein 2 | Cytoplasm | transporter | |
| chr5 | NADK2 | 2 | 127 | 5 | NADK2 | NAD kinase 2, mitochondrial | Cytoplasm | kinase | |
| chr7 | AC073133.1 | 2 | 78 | 5 | | | | | |
| chr7 | SSMEM1 | 2 | 61 | 5 | SSMEM1 | serine rich single-pass membrane protein 1 | Other | other | |
| chr1 | PABPC4 | 2 | 76 | 5 | PABPC4 | poly(A) binding protein | Cytoplasm | translation regulator | |
| chrX | KRT8P14 | 2 | 90 | 5 | KRT8P14 | keratin 8 pseudogene 14 | Other | other | |
| chr3 | SHOX2 | 3 | 134 | 5 | SHOX2 | short stature homeobox 2 | Nucleus | transcription regulator | |
| chr6 | GNMT | 2 | 93 | 5 | GNMT | glycine N-methyltransferase | Cytoplasm | enzyme | |
| chr17 | RP11-47L3.1 | 2 | 84 | 5 | | | | | |
| chr19 | CTC-490G23.4 | 2 | 83 | 5 | | | | | |
| chr21 | AP000695.6 | 2 | 53 | 5 | | | | | |
| chr14 | SERPINA13P | 2 | 71 | 5 | SERPINA13P | serpin family A member 13, pseudogene | Extracellular Space | other | |
| chr2 | AC019330.1 | 2 | 54 | 5 | | | | | |
| chrX | TMEM27 | 3 | 64 | 5 | CLTRN | collectrin, amino acid transport regulator | Plasma Membrane | other | |
| chr20 | SRMP1 | 2 | 115 | 5 | SRMP1 | spermidine synthase pseudogene | Other | other | |
| chr10 | PIPSL | 2 | 64 | 5 | PIPSL | PIP5K1A and PSMD4 like (pseudogene) | Other | other | |
| chr1 | ACTRT2 | 3 | 100 | 5 | ACTRT2 | actin related protein T2 | Other | other | |
| chr17 | KRT35 | 3 | 82 | 5 | KRT35 | keratin 35 | Cytoplasm | other | |
| chr2 | MRPL35 | 2 | 61 | 5 | MRPL35 | mitochondrial ribosomal protein | Cytoplasm | other | |
| chr17 | KRT13 | 3 | 80 | 5 | KRT13 | keratin 13 | Cytoplasm | other | |
| chr22 | RRP7B | 3 | 141 | 5 | RRP7BP | ribosomal RNA processing 7 homolog B, pseudogene | Other | other | |
| chr10 | NFKB2 | 3 | 117 | 5 | NFKB2 | nuclear factor kappa B subunit 2 | Nucleus | transcription regulator | |
| chr11 | CARNS1 | 3 | 112 | 5 | CARNS1 | carnosine synthase 1 | Cytoplasm | enzyme | |
| chr18 | RP11-669I1.1 | 2 | 58 | 5 | | | | | |
| chr4 | CLOCK | 2 | 38 | 5 | CLOCK | clock circadian regulator | Nucleus | transcription regulator | |
| chr13 | GPR180 | 2 | 61 | 5 | GPR180 | G protein-coupled receptor 180 | Cytoplasm | other | |
| chr19 | CTD-2538C1.2 | 3 | 57 | 5 | | | | | |
| chr11 | RPS6KA4 | 3 | 136 | 5 | RPS6KA4 | ribosomal protein S6 kinase A4 | Cytoplasm | kinase | |
| chr2 | AC096574.5 | 2 | 55 | 5 | | | | | |
| chr1 | SFT2D2 | 3 | 77 | 5 | SFT2D2 | SFT2 domain containing 2 | Cytoplasm | other | |
| chr8 | LY6E | 3 | 80 | 5 | LY6E | lymphocyte antigen 6 family member E | Plasma Membrane | other | DLYE5953A |
| chr1 | AQP10 | 2 | 108 | 5 | AQP10 | aquaporin 10 | Plasma Membrane | transporter | |
| chr10 | SRGN | 2 | 93 | 5 | SRGN | serglycin | Cytoplasm | other | |
| chr22 | SYCE3 | 2 | 48 | 5 | SYCE3 | synaptonemal complex central element protein 3 | Nucleus | other | |
| chr11 | TMEM134 | 2 | 141 | 5 | TMEM134 | transmembrane protein 134 | Other | kinase | |

| | | | | | | | | | |
|---|---|---|---|---|---|---|---|---|---|
| chr9 | *ENTPD8* | 3 | 96 | 5 | ENTPD8 | ectonucleoside triphosphate diphosphohydrolase 8 | Plasma Membrane | enzyme | |
| chrX | *GYG2* | 2 | 60 | 5 | GYG2 | glycogenin 2 | Cytoplasm | enzyme | |
| chr1 | *PYGO2* | 3 | 107 | 5 | PYGO2 | pygopus family PHD finger 2 | Nucleus | other | |
| chr12 | *TAPBPL* | 2 | 84 | 5 | TAPBPL | TAP binding protein like | Cytoplasm | other | |
| chr11 | *C11orf57* | 3 | 66 | 5 | NKAPD1 | NKAP domain containing 1 | Other | other | |
| chr15 | *AP3S2* | 2 | 132 | 5 | AP3S2 | adaptor related protein complex 3 subunit sigma 2 | Cytoplasm | transporter | |
| chr4 | *NMU* | 2 | 58 | 5 | NMU | neuromedin U | Extracellular Space | other | |
| chr8 | *SNX16* | 2 | 38 | 5 | SNX16 | sorting nexin 16 | Cytoplasm | transporter | |
| chr19 | *IRGC* | 2 | 101 | 5 | IRGC | immunity related GTPase cinema | Other | other | |
| chr1 | *S100A12* | 2 | 73 | 5 | S100A12 | S100 calcium binding protein A12 | Cytoplasm | other | |
| chr7 | *FAM133B* | 2 | 96 | 5 | FAM133B | family with sequence similarity 133 member B | Other | other | |
| chr19 | *LSM7* | 3 | 128 | 5 | LSM7 | LSM7 homolog, U6 small nuclear RNA and mRNA degradation associated | Nucleus | other | |
| chr8 | *RAB2A* | 2 | 39 | 5 | RAB2A | RAB2A, member RAS oncogene family | Cytoplasm | enzyme | |
| chr21 | *AP000289.6* | 2 | 57 | 5 | | | | | |
| chr2 | *ITPRIPL1* | 2 | 126 | 5 | ITPRIPL1 | ITPRIP like 1 | Other | other | |
| chr6 | *RP11-154D6.1* | 2 | 62 | 5 | | | | | |
| chr11 | *TMEM216* | 3 | 115 | 5 | TMEM216 | transmembrane protein 216 | Extracellular Space | other | |
| chr7 | *AC004862.6* | 2 | 58 | 5 | | | | | |
| chr2 | *COL4A3* | 2 | 39 | 5 | COL4A3 | collagen type IV alpha 3 chain | Extracellular Space | other | collagenase clostridium |
| chr3 | *RP11-454C18.1* | 2 | 61 | 5 | | | | | |
| chr12 | *NDUFA4L2* | 3 | 126 | 5 | NDUFA4L2 | NDUFA4 mitochondrial complex associated like 2 | Other | enzyme | |
| chr2 | *AC074011.2* | 2 | 51 | 5 | | | | | |
| chr5 | *NPM1* | 2 | 63 | 5 | NPM1 | nucleophosmin 1 | Nucleus | transcription regulator | |
| chr22 | *GSTTP2* | 2 | 100 | 5 | GSTTP2 | glutathione S-transferase theta pseudogene 2 | Other | other | |
| chr9 | *ORM2* | 2 | 60 | 5 | ORM2 | orosomucoid 2 | Extracellular Space | other | |
| chr22 | *CYP2D8P1* | 3 | 113 | 5 | CYP2D8P | ccytochrome P450 family 2 subfamily D member 8, | Other | other | |
| chr1 | *CD164L2* | 2 | 92 | 5 | CD164L2 | CD164 molecule like 2 | Other | other | |
| chr12 | *C12orf10* | 3 | 103 | 5 | C12orf10 | chromosome 12 open reading frame 10 | Nucleus | other | |
| chr14 | *SSTR1* | 3 | 150 | 5 | SSTR1 | somatostatin receptor 1 | Plasma Membrane | G-protein coupled recepto | octreotide, everolimus/pasireotide, octreotide/prochlorperazine, haloperidol/octreotide |
| chr1 | *KCNJ10* | 3 | 96 | 5 | KCNJ10 | potassium inwardly rectifying channel subfamily J member 10 | Plasma Membrane | ion channel | |
| chr9 | *RP11-555F9.2* | 3 | 65 | 5 | | | | | |
| chr18 | *ME2* | 2 | 44 | 5 | ME2 | malic enzyme 2 | Cytoplasm | enzyme | |
| chr11 | *NDUFV1* | 3 | 115 | 5 | NDUFV1 | NADH:ubiquinone oxidoreductase core subunit V1 | Cytoplasm | enzyme | |
| chr17 | *ODF4* | 3 | 61 | 5 | ODF4 | outer dense fiber of sperm tails 4 | Cytoplasm | other | |
| chr15 | *SHF* | 2 | 83 | 5 | SHF | Src homology 2 domain | Other | other | |
| chr17 | *MLX* | 3 | 99 | 5 | MLX | MAX dimerization protein MLX | Nucleus | transcription regulator | |
| chr5 | *MXD3* | 3 | 108 | 5 | MXD3 | MAX dimerization protein 3 | Nucleus | transcription regulator | |
| chr16 | *RP11-189E14.5* | 3 | 70 | 5 | | | | | |
| chr18 | *LINC-ROR* | 2 | 46 | 5 | LINC-ROR | long intergenic non-protein coding RNA, regulator of reprogramming | Other | other | |
| chr11 | *RP11-56P9.10* | 3 | 125 | 5 | | | | | |
| chr2 | *EPHA4* | 3 | 38 | 5 | EPHA4 | EPH receptor A4 | Plasma Membrane | kinase | |
| chr19 | *SIGLEC7* | 2 | 104 | 5 | SIGLEC7 | sialic acid binding Ig like lectin 7 | Plasma Membrane | transmembrane recepto | |
| chr7 | *CLDN15* | 3 | 146 | 5 | CLDN15 | claudin 15 | Plasma Membrane | other | |
| chr9 | *ANKRD19P* | 2 | 44 | 5 | ANKRD19P | ankyrin repeat domain 19, pseudogene | Other | other | |
| chr10 | *PNLIPRP1* | 3 | 75 | 5 | PNLIPRP1 | pancreatic lipase related protein 1 | Extracellular Space | enzyme | |
| chr11 | *RP11-212D19.4* | 2 | 54 | 5 | | | | | |
| chr16 | *CTD-2542L18.1* | 3 | 106 | 5 | | | | | |
| chr7 | *AC007551.3* | 2 | 49 | 5 | | | | | |
| chr20 | *KCNS1* | 3 | 95 | 5 | KCNS1 | potassium voltage-gated channel modifier subfamily S member 1 | Plasma Membrane | ion channel | |
| chr2 | *CCNT2* | 2 | 41 | 5 | CCNT2 | cyclin T2 | Nucleus | transcription regulator | |
| chr2 | *AC007040.6* | 2 | 68 | 5 | | | | | |
| chr11 | *GAL* | 2 | 108 | 5 | GAL | galanin and GMAP prepropeptide | Extracellular Space | other | |
| chr7 | *GCC1* | 3 | 125 | 5 | GCC1 | GRIP and coiled-coil domain containing 1 | Cytoplasm | other | |
| chr8 | *RP11-77B22.2* | 2 | 38 | 5 | | | | | |
| chr20 | *RP11-290F20.2* | 2 | 144 | 5 | | | | | |
| chr2 | *AC011747.6* | 2 | 75 | 5 | | | | | |

| | | | | | | | | | |
|---|---|---|---|---|---|---|---|---|---|
| chr1 | *SERPINC1* | 2 | 59 | 5 | SERPINC1 | serpin family C member 1 | Extracellular Space | enzyme | dalteparin, glucuronyl glucosamine glycan sulfate, aspirin/enoxaparin, SR-123781A, heparin, fondaparinux/warfarin, fondaparinux, low molecular weight heparin, ardeparin, nadroparin, semuloparin, enoxaparin, |
| chr14 | *CTD-2561F5.1* | 2 | 58 | 5 | | | | | |
| chr5 | *HAND1* | 3 | 100 | 5 | HAND1 | heart and neural crest derivatives expressed 1 | Nucleus | transcription regulator | |
| chr17 | *TUBG1* | 2 | 102 | 5 | TUBG1 | tubulin gamma 1 | Cytoplasm | other | docetaxel/gemcitabine/vincristine, cyclophosphamide/epirubicin/5-fluorouracil/paclitaxel, bevacizumab/paclitaxel/topotecan, colchicine/probenecid, bevacizumab/docetaxel/prednisone/thalidomide/zoledronic acid, gemcitabine/paclitaxel, L-asparaginase/daunorubicin/dexamethasone/vincristine, prednisone/vincristine, docetaxel/5-fluorouracil, vinflunine, L-[illegible] |
| chr20 | *RP5-998H6.2* | 2 | 103 | 5 | | | | | |
| chr2 | *STON1-GTF2A1L* | 3 | 76 | 5 | STON1-GTF2A1L | STON1-GTF2A1L readthrough | Cytoplasm | other | |
| chr7 | *STEAP2* | 2 | 48 | 5 | STEAP2 | STEAP2 metalloreductase | Plasma Membrane | enzyme | |
| chr18 | *RP11-13N13.2* | 2 | 42 | 5 | | | | | |
| chr8 | *RP11-37N22.1* | 2 | 45 | 5 | | | | | |
| chr16 | *MLST8* | 2 | 85 | 5 | MLST8 | MTOR associated protein, LST8 homolog | Cytoplasm | other | |
| chr16 | *OGFOD1* | 2 | 47 | 5 | OGFOD1 | 2-oxoglutarate and iron dependent oxygenase domain | Cytoplasm | enzyme | |
| chr17 | *SGCA* | 3 | 130 | 5 | SGCA | sarcoglycan alpha | Plasma Membrane | other | |
| chr19 | *FFAR3* | 3 | 132 | 5 | FFAR3 | free fatty acid receptor 3 | Plasma Membrane | G-protein coupled receptor | |
| chr8 | *RP11-613H2.1* | 2 | 88 | 5 | | | | | |
| chr16 | *RP11-134D3.1* | 3 | 83 | 5 | | | | | |
| chr15 | *SLC51B* | 2 | 73 | 5 | SLC51B | solute carrier family 51 beta | Plasma Membrane | transporter | |
| chr9 | *EDF1* | 3 | 144 | 5 | EDF1 | endothelial differentiation related factor 1 | Nucleus | transcription regulator | |
| chr6 | *RMND1* | 2 | 40 | 5 | RMND1 | required for meiotic nuclear division 1 homolog | Cytoplasm | other | |
| chr5 | *DDX41* | 3 | 83 | 5 | DDX41 | DEAD-box helicase 41 | Nucleus | enzyme | |
| chr9 | *KLF9* | 2 | 59 | 5 | KLF9 | Kruppel like factor 9 | Nucleus | transcription regulator | |
| chr1 | *RP11-73M7.1* | 3 | 64 | 5 | | | | | |
| chr2 | *AC017074.2* | 2 | 61 | 5 | | | | | |
| chr16 | *NPIPB7* | 2 | 82 | 5 | NPIPB7 | nuclear pore complex interacting protein family member B7 | Nucleus | other | |
| chr6 | *RP11-13P5.1* | 2 | 46 | 5 | | | | | |
| chr11 | *FGF19* | 2 | 90 | 5 | FGF19 | fibroblast growth factor 19 | Extracellular Space | growth factor | |
| chr21 | *LINC00315* | 3 | 122 | 5 | LINC00315 | long intergenic non-protein coding RNA 315 | Other | other | |
| chrX | *SMARCA1* | 2 | 46 | 5 | SMARCA1 | SWI/SNF related, matrix associated, actin dependent regulator of chromatin, subfamily | Nucleus | transcription regulator | |
| chr9 | *INPP5E* | 3 | 153 | 5 | INPP5E | inositol polyphosphate-5-phosphatase E | Cytoplasm | phosphatase | |
| chr5 | *MIER3* | 2 | 71 | 5 | MIER3 | MIER family member 3 | Nucleus | transcription regulator | |
| chr16 | *UBALD1* | 3 | 111 | 5 | UBALD1 | UBA like domain containing 1 | Other | other | |
| chr17 | *RP11-260A9.4* | 2 | 84 | 5 | | | | | |
| chr16 | *SMIM22* | 2 | 68 | 5 | SMIM22 | small integral membrane protein | Other | other | |
| chr2 | *MYEOV2* | 2 | 84 | 5 | COPS9 | COP9 signalosome subunit 9 | Nucleus | other | |
| chr10 | *PLEKHS1* | 2 | 50 | 5 | PLEKHS1 | pleckstrin homology domain containing S1 | Other | other | |
| chr1 | *GPR89C* | 2 | 49 | 5 | GPR89A/GPR89B | G protein-coupled receptor 89B | Cytoplasm | ion channel | |
| chr19 | *MAMSTR* | 3 | 127 | 5 | MAMSTR | MEF2 activating motif and SAP domain containing transcriptional regulator | Nucleus | transcription regulator | |
| chr1 | *REG4* | 3 | 91 | 5 | REG4 | regenerating family member 4 | Extracellular Space | other | |
| chr20 | *SPTLC3* | 2 | 49 | 5 | SPTLC3 | serine palmitoyltransferase long chain base subunit 3 | Cytoplasm | enzyme | |
| chr14 | *TRIP11* | 2 | 63 | 5 | TRIP11 | thyroid hormone receptor interactor 11 | Cytoplasm | transcription regulator | |

| | | | | | | | | |
|---|---|---|---|---|---|---|---|---|
| chr22 | *CTA-217C2.1* | 2 | 46 | 5 | | | | |
| chr8 | *CTA-398F10.1* | 2 | 57 | 5 | | | | |
| chr15 | *PYGO1* | 3 | 51 | 5 | PYGO1 | pygopus family PHD finger 1 | Nucleus | other |
| chr2 | *CFLAR* | 2 | 76 | 5 | CFLAR | CASP8 and FADD like apoptosis regulator | Cytoplasm | other |
| chr15 | *GOLGA6L5* | 2 | 223 | 5 | GOLGA6L5P | golgin A6 family like 5, | Other | other |
| chr19 | *CTC-550B14.7* | 2 | 51 | 5 | | | | |
| chr11 | *KRTAP5-AS1* | 3 | 52 | 5 | KRTAP5-AS1 | KRTAP5-1/KRTAP5-2 antisense RNA 1 | Other | other |
| chr20 | *PROKR2* | 2 | 79 | 5 | PROKR2 | prokineticin receptor 2 | Plasma Membrane | G-protein coupled recepto |
| chr13 | *DIS3* | 3 | 45 | 5 | DIS3 | DIS3 homolog, exosome endoribonuclease and 3'-5' exoribonuclease | Nucleus | enzyme |
| chr14 | *POTEM* | 2 | 75 | 5 | TEH (includes oth | POTE ankyrin domain family member G | Other | other |
| chr16 | *KCTD13* | 2 | 94 | 5 | KCTD13 | potassium channel tetramerization domain containing | Nucleus | ion channel |
| chr19 | *CTD-2521M24.10* | 2 | 81 | 5 | | | | |
| chr7 | *GATAD1* | 2 | 68 | 5 | GATAD1 | GATA zinc finger domain | Nucleus | transcription regulator |
| chr2 | *EPT1* | 2 | 53 | 5 | SELENOI | selenoprotein I | Cytoplasm | enzyme |
| chr10 | *RP11-71J2.1* | 3 | 71 | 5 | | | | |
| chr17 | *IFT20* | 3 | 65 | 5 | IFT20 | intraflagellar transport 20 | Cytoplasm | other |
| chrX | *GLRA4* | 2 | 65 | 5 | GLRA4 | glycine receptor alpha 4 | Plasma Membrane | ion channel |
| chr10 | *RP11-472N13.3* | 2 | 55 | 5 | | | | |
| chr7 | *AC005077.7* | 2 | 71 | 5 | | | | |
| chr2 | *TUBA3E* | 2 | 77 | 5 | TUBA3E | tubulin alpha 3e | Cytoplasm | other |
| chr16 | *C16orf47* | 3 | 62 | 5 | ZFHX3 | zinc finger homeobox 3 | Nucleus | transcription regulator |
| chr6 | *LINC00518* | 2 | 41 | 5 | LINC00518 | long intergenic non-protein coding RNA 518 | Other | other |
| chr19 | *AC002398.5* | 3 | 132 | 5 | | | | |
| chr1 | *PHLDA3* | 3 | 138 | 5 | PHLDA3 | pleckstrin homology like domain family A member 3 | Plasma Membrane | other |
| chr19 | *MPND* | 2 | 100 | 5 | MPND | MPN domain containing | Other | other |
| chr19 | *CTD-2666L21.1* | 2 | 37 | 5 | | | | |
| chr15 | *ARPP19* | 2 | 71 | 5 | ARPP19 | cAMP regulated phosphoprotein | Cytoplasm | transporter |
| chr7 | *FTSJ2* | 3 | 94 | 5 | MRM2 | mitochondrial rRNA methyltransferase 2 | Nucleus | enzyme |
| chr13 | *PARP4P2* | 2 | 65 | 5 | PARP4P2 | poly(ADP-ribose) polymerase family member 4 pseudogene 2 | Other | other |
| chr7 | *INMT-FAM188B* | 3 | 65 | 5 | INMT-MINDY4 | INMT-MINDY4 readthrough (NMD candidate) | Other | other |
| chr19 | *KLK3* | 2 | 92 | 5 | KLK3 | kallikrein related peptidase 3 | Extracellular Space | peptidase |
| chr22 | *GCAT* | 2 | 82 | 5 | GCAT | glycine C-acetyltransferase | Cytoplasm | enzyme |
| chr19 | *PSG7* | 3 | 62 | 5 | PSG7 | pregnancy specific beta-1-glycoprotein 7 | Extracellular Space | other |
| chr7 | *AC007349.4* | 2 | 68 | 5 | | | | |
| chr22 | *POM121L10P* | 2 | 87 | 5 | POM121L10P | POM121 transmembrane nucleoporin like 10, pseudogene | Other | other |
| chr7 | *SMKR1* | 2 | 59 | 5 | SMKR1 | small lysine rich protein 1 | Other | other |
| chr1 | *RP11-465B22.5* | 3 | 81 | 5 | | | | |
| chr16 | *RP11-805I24.3* | 2 | 84 | 5 | | | | |
| chr2 | *AC007040.11* | 3 | 61 | 5 | | | | |
| chr9 | *RBM18* | 2 | 66 | 5 | RBM18 | RNA binding motif protein 18 | Other | other |
| chr2 | *C2orf72* | 3 | 47 | 5 | C2orf72 | chromosome 2 open reading frame 72 | Other | other |
| chr12 | *RP1-7G5.6* | 3 | 74 | 5 | | | | |
| chr5 | *CTD-2244C20.2* | 2 | 39 | 5 | | | | |
| chr3 | *NAA50* | 2 | 53 | 5 | NAA50 | N(alpha)-acetyltransferase 50, NatE catalytic subunit | Cytoplasm | enzyme |
| chr17 | *SP6* | 3 | 87 | 5 | SP6 | Sp6 transcription factor | Nucleus | transcription regulator |
| chr4 | *UGDH-AS1* | 2 | 46 | 5 | UGDH-AS1 | UGDH antisense RNA 1 | Other | other |
| chr1 | *VAMP4* | 3 | 87 | 5 | VAMP4 | vesicle associated membrane protein 4 | Cytoplasm | other |
| chr17 | *DLX3* | 3 | 110 | 5 | DLX3 | distal-less homeobox 3 | Nucleus | transcription regulator |
| chr2 | *ABCA12* | 2 | 37 | 5 | ABCA12 | ATP binding cassette subfamily A member 12 | Plasma Membrane | transporter |
| chr19 | *KLK4* | 2 | 95 | 5 | KLK4 | kallikrein related peptidase 4 | Extracellular Space | peptidase |
| chr7 | *SEPT7P3* | 2 | 71 | 5 | SEPTIN7P3 | septin 7 pseudogene 3 | Other | other |
| chr16 | *FAM195A* | 3 | 124 | 5 | MCRIP2 | MAPK regulated corepressor interacting protein 2 | Cytoplasm | other |
| chr19 | *HRC* | 2 | 95 | 5 | HRC | histidine rich calcium binding | Cytoplasm | other |
| chr12 | *SLCO1C1* | 2 | 47 | 5 | SLCO1C1 | solute carrier organic anion transporter family member 1C1 | Plasma Membrane | transporter |
| chr22 | *MIAT* | 2 | 66 | 5 | MIAT | myocardial infarction associated transcript | Other | other |
| chr12 | *RP5-1063M23.2* | 2 | 82 | 5 | | | | |
| chr10 | *ARMC4* | 3 | 44 | 5 | ARMC4 | armadillo repeat containing 4 | Extracellular Space | other |
| chrX | *CD99P1* | 2 | 44 | 5 | CD99P1 | CD99 molecule pseudogene 1 | Plasma Membrane | other |
| chr18 | *ADNP2* | 2 | 68 | 5 | ADNP2 | ADNP homeobox 2 | Other | other |
| chr17 | *WSB1* | 2 | 66 | 5 | WSB1 | WD repeat and SOCS box containing 1 | Cytoplasm | other |
| chr9 | *PES1P2* | 2 | 86 | 5 | PES1P2 | pescadillo ribosomal biogenesis factor 1 pseudogene 2 | Other | other |
| chr4 | *LCORL* | 2 | 57 | 5 | LCORL | ligand dependent nuclear receptor corepressor like | Nucleus | transcription regulator |
| chr19 | *AC084219.4* | 2 | 37 | 5 | | | | |
| chr1 | *PPCS* | 2 | 52 | 5 | PPCS | phosphopantothenoylcysteine synthetase | Cytoplasm | enzyme |
| chr7 | *RP11-611L7.1* | 2 | 93 | 5 | | | | |

| | | | | | | | | | |
|---|---|---|---|---|---|---|---|---|---|
| chrX | *TSPYL2* | 3 | 84 | 5 | TSPYL2 | TSPY like 2 | Nucleus | other | |
| chr2 | *AC140481.4* | 2 | 77 | 5 | | | | | |
| chr17 | *VEZF1* | 2 | 42 | 5 | VEZF1 | vascular endothelial zinc finger 1 | Nucleus | transcription regulator | |
| chr20 | *RP1-1J6.2* | 2 | 86 | 5 | | | | | |
| chr12 | *HOXC-AS5* | 2 | 147 | 5 | HOXC13-AS | HOXC13 antisense RNA | Other | other | |
| chr11 | *TMEM86A* | 2 | 101 | 5 | TMEM86A | transmembrane protein 86A | Other | other | |
| chr1 | *HENMT1* | 3 | 109 | 5 | HENMT1 | HEN methyltransferase 1 | Other | enzyme | |
| chr21 | *LINC00846* | 2 | 66 | 5 | | | | | |
| chr6 | *TREML4* | 2 | 55 | 5 | TREML4 | triggering receptor expressed on myeloid cells like 4 | Plasma Membrane | other | |
| chr15 | *AC011648.1* | 2 | 92 | 5 | | | | | |
| chr7 | *C7orf71* | 3 | 87 | 5 | LINC02860 | long intergenic non-protein coding RNA 2860 | Other | other | |
| chr16 | *RP11-252A24.7* | 2 | 70 | 5 | | | | | |
| chr20 | *FRG1B* | 2 | 313 | 5 | FRG1BP | FSHD region gene 1 family member B, pseudogene | Other | other | |
| chr4 | *HAND2-AS1* | 2 | 42 | 5 | HAND2-AS1 | HAND2 antisense RNA 1 | Other | other | |
| chr22 | *LINC00899* | 2 | 84 | 5 | LINC00899 | long intergenic non-protein coding RNA 899 | Other | other | |
| chr8 | *RP11-44N12.2* | 2 | 38 | 5 | | | | | |
| chr9 | *ISCA1* | 2 | 59 | 5 | ISCA1 | iron-sulfur cluster assembly 1 | Cytoplasm | other | |
| chr16 | *NOL3* | 3 | 133 | 5 | NOL3 | nucleolar protein 3 | Nucleus | other | |
| chr9 | *RP11-439K3.1* | 2 | 66 | 5 | | | | | |
| chr7 | *AC000370.2* | 2 | 51 | 5 | | | | | |
| chr8 | *RP13-467H17.1* | 3 | 113 | 5 | | | | | |
| chr22 | *AP000350.8* | 2 | 88 | 5 | | | | | |
| chr11 | *FOLR4* | 3 | 142 | 5 | IZUMO1R | IZUMO1 receptor, JUNO | Plasma Membrane | other | |
| chr7 | *RP5-905H7.9* | 2 | 51 | 5 | | | | | |
| chr4 | *PLK4* | 2 | 62 | 5 | PLK4 | polo like kinase 4 | Cytoplasm | kinase | CFI-400945 |
| chr3 | *RP11-77P16.4* | 2 | 70 | 5 | | | | | |
| chr19 | *AC005392.13* | 2 | 57 | 5 | | | | | |
| chr3 | *LINC00694* | 2 | 93 | 5 | C3orf86 | chromosome 3 open reading frame 86 | Other | other | |
| chr5 | *SPRY4* | 3 | 80 | 5 | SPRY4 | sprouty RTK signaling antagonist | Plasma Membrane | other | |
| chrX | *RP1-23K20.2* | 2 | 48 | 5 | | | | | |
| chr14 | *LRR1* | 3 | 47 | 5 | LRR1 | leucine rich repeat protein 1 | Cytoplasm | enzyme | |
| chr15 | *RP11-37C7.1* | 2 | 46 | 5 | | | | | |
| chr18 | *TPGS2* | 2 | 37 | 5 | TPGS2 | tubulin polyglutamylase complex subunit 2 | Nucleus | other | |
| chr3 | *CTD-2026G6.3* | 2 | 42 | 5 | | | | | |
| chr15 | *RP13-395E19.3* | 2 | 53 | 5 | | | | | |
| chr5 | *CTD-2004A9.1* | 3 | 183 | 5 | | | | | |
| chr17 | *KRT19* | 3 | 124 | 5 | KRT19 | keratin 19 | Cytoplasm | other | |
| chr2 | *AC010987.6* | 2 | 45 | 5 | | | | | |
| chr19 | *LYL1* | 3 | 75 | 5 | LYL1 | LYL1 basic helix-loop-helix family member | Nucleus | transcription regulator | |
| chr1 | *RP5-1126H10.2* | 2 | 103 | 5 | | | | | |
| chr1 | *RP11-277B15.2* | 2 | 47 | 5 | | | | | |
| chr17 | *RP5-1110E20.1* | 3 | 58 | 5 | | | | | |
| chr19 | *PLK5* | 3 | 77 | 5 | PLK5 | polo like kinase 5 | Cytoplasm | other | |
| chr8 | *ZBTB10* | 2 | 46 | 5 | ZBTB10 | zinc finger and BTB domain containing 10 | Nucleus | other | |
| chr17 | *ASB16-AS1* | 2 | 59 | 5 | ASB16-AS1 | ASB16 antisense RNA 1 | Other | other | |
| chrX | *RP4-646N3.1* | 2 | 40 | 5 | | | | | |
| chr19 | *CTD-3222D19.2* | 3 | 70 | 5 | | | | | |
| chr1 | *RP1-74M1.3* | 2 | 111 | 5 | | | | | |
| chr2 | *DNAJC10* | 2 | 38 | 5 | DNAJC10 | DnaJ heat shock protein family (Hsp40) member C10 | Cytoplasm | enzyme | |
| chr15 | *AC048382.4* | 2 | 63 | 5 | | | | | |
| chr16 | *ZKSCAN2* | 2 | 59 | 5 | ZKSCAN2 | zinc finger with KRAB and SCAN domains 2 | Nucleus | transcription regulator | |
| chr12 | *PGAM1P5* | 2 | 45 | 5 | PGAM1P5 | phosphoglycerate mutase 1 pseudogene 5 | Other | other | |
| chr10 | *AGAP4* | 2 | 84 | 5 | AP5 (includes oth | ArfGAP with GTPase domain, ankyrin repeat and PH domain 5 | Other | other | |
| chr1 | *GUCA2B* | 2 | 104 | 5 | GUCA2B | guanylate cyclase activator 2B | Extracellular Space | other | |
| chr15 | *RP11-653J6.1* | 2 | 47 | 5 | | | | | |
| chr5 | *RP11-506H20.1* | 3 | 61 | 5 | | | | | |
| chr17 | *TMUB2* | 2 | 92 | 5 | TMUB2 | transmembrane and ubiquitin like domain containing 2 | Other | other | |
| chr19 | *WDR18* | 3 | 72 | 5 | WDR18 | WD repeat domain 18 | Nucleus | other | |
| chr9 | *CELP* | 2 | 154 | 5 | CELP | carboxyl ester lipase pseudogene | Other | other | |
| chr22 | *TRABD* | 3 | 118 | 5 | TRABD | TraB domain containing | Extracellular Space | other | |
| chrX | *GS1-590J6.2* | 2 | 75 | 5 | | | | | |
| chr17 | *PTGES3L-AARSD* | 2 | 45 | 5 | PTGES3L-AARSD | PTGES3L-AARSD1 readthrough | Other | other | |
| chr22 | *BMS1P20* | 2 | 76 | 5 | BMS1P20 | BMS1 pseudogene 20 | Other | other | |
| chrX | *CT45A2* | 2 | 76 | 5 | 45A1 (includes oth | cancer/testis antigen family 45 member A2 | Other | other | |
| chr12 | *RP11-165P7.1* | 2 | 61 | 5 | | | | | |
| chr4 | *MGST2* | 2 | 39 | 5 | MGST2 | microsomal glutathione S-transferase 2 | Cytoplasm | enzyme | |
| chr4 | *LRAT* | 2 | 62 | 5 | LRAT | lecithin retinol acyltransferase | Cytoplasm | enzyme | |
| chr4 | *QDPR* | 2 | 52 | 5 | QDPR | quinoid dihydropteridine | Cytoplasm | enzyme | |
| chr19 | *CLPP* | 3 | 67 | 5 | CLPP | caseinolytic mitochondrial matrix peptidase proteolytic subunit | Cytoplasm | peptidase | |
| chr17 | *ENTHD2* | 3 | 123 | 5 | TEPSIN | TEPSIN adaptor related protein complex 4 accessory protein | Nucleus | other | |
| chr17 | *KRTAP3-3* | 1 | 41 | 5 | KRTAP3-3 | keratin associated protein 3-3 | Cytoplasm | other | |
| chr20 | *ISM1* | 2 | 40 | 5 | ISM1 | isthmin 1 | Other | other | |

| | | | | | | | | | |
|---|---|---|---|---|---|---|---|---|---|
| chr2 | *AC104088.1* | 2 | 104 | 5 | | | | | |
| chr11 | *CYP2R1* | 2 | 95 | 5 | CYP2R1 | cytochrome P450 family 2 subfamily R member 1 | Cytoplasm | enzyme | |
| chr11 | *PPFIA1* | 2 | 47 | 5 | PPFIA1 | PTPRF interacting protein alpha 1 | Plasma Membrane | phosphatase | |
| chr7 | *CCT6P1* | 2 | 49 | 5 | CCT6P1 | chaperonin containing TCP1 subunit 6 pseudogene 1 | Other | other | |
| chr1 | *TCHH* | 2 | 102 | 5 | TCHH | trichohyalin | Extracellular Space | other | |
| chr3 | *RP11-292E2.1* | 2 | 51 | 5 | | | | | |
| chr16 | *MT3* | 2 | 103 | 5 | MT3 | metallothionein 3 | Cytoplasm | other | |
| chr12 | *DYRK4* | 2 | 60 | 5 | DYRK4 | dual specificity tyrosine phosphorylation regulated kinase | Nucleus | kinase | |
| chr20 | *RP4-673D20.1* | 3 | 49 | 5 | | | | | |
| chr2 | *AC005042.5* | 2 | 46 | 5 | | | | | |
| chr9 | *TPM2* | 2 | 78 | 5 | TPM2 | tropomyosin 2 | Other | other | |
| chr14 | *DHRS1* | 2 | 88 | 5 | DHRS1 | dehydrogenase/reductase 1 | Cytoplasm | enzyme | |
| chr19 | *ZNF723* | 2 | 69 | 5 | ZNF723 | zinc finger protein 723 | Other | other | |
| chr1 | *GNAT2* | 2 | 63 | 5 | GNAT2 | G protein subunit alpha | Plasma Membrane | enzyme | |
| chr4 | *RP11-810P8.1* | 2 | 53 | 5 | | | | | |
| chr21 | *CLIC6* | 2 | 51 | 5 | CLIC6 | chloride intracellular channel 6 | Plasma Membrane | ion channel | |
| chr16 | *TRADD* | 3 | 156 | 5 | TRADD | TNFRSF1A associated via death domain | Cytoplasm | other | |
| chr8 | *FAM90A24P* | 2 | 459 | 5 | FAM90A24P | family with sequence similarity 90, member A24 | Other | other | |
| chr8 | *FAM90A13P* | 2 | 434 | 5 | 90A26 (includes ot | family with sequence similarity 90 member A26 | Other | other | |
| chr19 | *ZNF614* | 2 | 71 | 5 | ZNF614 | zinc finger protein 614 | Other | other | |
| chr22 | *CTA-929C8.8* | 2 | 44 | 5 | | | | | |
| chr16 | *RRN3P2* | 2 | 65 | 5 | RRN3P2 | RRN3 homolog, RNA polymerase I transcription factor pseudogene | Other | other | |
| chr1 | *DRAM2* | 2 | 56 | 5 | DRAM2 | DNA damage regulated autophagy modulator 2 | Cytoplasm | other | |
| chr2 | *RP11-363D14.1* | 2 | 127 | 5 | | | | | |
| chr1 | *FOXD2-AS1* | 2 | 75 | 5 | FOXD2-AS1 | FOXD2 adjacent opposite strand RNA 1 | Other | other | |
| chr5 | *CTC-255N20.1* | 2 | 68 | 5 | | | | | |
| chr10 | *PDCD4* | 2 | 48 | 5 | PDCD4 | programmed cell death 4 | Nucleus | other | |
| chr22 | *ARSA* | 3 | 117 | 5 | ARSA | arylsulfatase A | Cytoplasm | enzyme | |
| chr2 | *THUMPD2* | 2 | 70 | 5 | THUMPD2 | THUMP domain containing 2 | Other | enzyme | |
| chr20 | *RP11-103J8.1* | 2 | 40 | 5 | | | | | |
| chr6 | *RP1-214M20.2* | 3 | 53 | 5 | | | | | |
| chr2 | *MKI67IP* | 2 | 70 | 5 | NIFK | nucleolar protein interacting with the FHA domain of MKI67 | Nucleus | other | |
| chr17 | *CACNA1G-AS1* | 3 | 154 | 5 | CACNA1G-AS1 | CACNA1G antisense RNA 1 | Other | other | |
| chr11 | *PCF11* | 2 | 55 | 5 | PCF11 | PCF11 cleavage and polyadenylation factor subunit | Nucleus | other | |
| chr7 | *TOMM7* | 2 | 99 | 5 | TOMM7 | translocase of outer mitochondrial membrane 7 | Cytoplasm | transporter | |
| chr10 | *ZNF37A* | 2 | 57 | 5 | ZNF37A | zinc finger protein 37A | Nucleus | transcription regulator | |
| chr11 | *OR7E5P* | 2 | 59 | 5 | OR7E5P | olfactory receptor family 7 subfamily E member 5 | Other | other | |
| chr6 | *FILIP1* | 2 | 57 | 5 | FILIP1 | filamin A interacting protein 1 | Cytoplasm | other | |
| chr17 | *LINC00324* | 2 | 98 | 5 | LINC00324 | long intergenic non-protein coding RNA 324 | Other | other | |
| chr14 | *PLEK2* | 2 | 75 | 5 | PLEK2 | pleckstrin 2 | Plasma Membrane | other | |
| chr10 | *RPP30* | 2 | 48 | 5 | RPP30 | ribonuclease P/MRP subunit p30 | Nucleus | enzyme | |
| chr2 | *ANKRD36BP2* | 2 | 52 | 5 | ANKRD36BP2 | ankyrin repeat domain 36B pseudogene 2 | Other | other | |
| chr1 | *PFDN2* | 2 | 75 | 5 | PFDN2 | prefoldin subunit 2 | Cytoplasm | other | |
| chr6 | *RP1-167A14.2* | 2 | 41 | 5 | | | | | |
| chr15 | *RP11-467N20.6* | 2 | 63 | 5 | | | | | |
| chr10 | *SYT15* | 3 | 438 | 5 | SYT15 | synaptotagmin 15 | Cytoplasm | transporter | |
| chr1 | *AL450992.2* | 3 | 66 | 5 | | | | | |
| chr14 | *RP11-661D19.3* | 2 | 73 | 5 | | | | | |
| chr9 | *SOCS5P2* | 2 | 75 | 5 | SOCS5P2 | suppressor of cytokine signaling 5 pseudogene 2 | Other | other | |
| chr5 | *CTD-2010I22.2* | 3 | 147 | 5 | | | | | |
| chr3 | *SLITRK3* | 3 | 66 | 5 | SLITRK3 | SLIT and NTRK like family | Plasma Membrane | other | |
| chr8 | *ALG1L12P* | 2 | 92 | 5 | ALG1L12P | ALG1 like 12, pseudogene | Other | other | |
| chr9 | *RP11-195F19.9* | 2 | 51 | 5 | | | | | |
| chr3 | *LINC00635* | 2 | 52 | 5 | LINC00635 | long intergenic non-protein coding RNA 635 | Other | other | |
| chr1 | *RP5-1103B4.3* | 3 | 49 | 5 | | | | | |
| chr7 | *AC009262.2* | 2 | 54 | 5 | | | | | |
| chr11 | *NXPE1* | 3 | 61 | 5 | NXPE1 | neurexophilin and PC-esterase domain family member 1 | Other | other | |
| chr19 | *SIGLEC21P* | 2 | 90 | 5 | SIGLEC21P | sialic acid binding Ig like lectin 21, pseudogene | Other | other | |
| chr1 | *ZNF672* | 3 | 73 | 5 | ZNF672 | zinc finger protein 672 | Nucleus | transcription regulator | |
| chr2 | *AC067956.1* | 2 | 47 | 5 | | | | | |
| chr2 | *RP11-333I13.1* | 2 | 94 | 5 | | | | | |
| chr10 | *CYP17A1* | 2 | 58 | 5 | CYP17A1 | cytochrome P450 family 17 subfamily A member 1 | Cytoplasm | enzyme | abiraterone, abiraterone acetate, abiraterone/prednisolone, CFG-920, ketoconazole, orteronel, CYP17A1 inhibitor, seviteronel |
| chr17 | *HOXB13* | 3 | 106 | 5 | HOXB13 | homeobox B13 | Nucleus | transcription regulator | |
| chr8 | *PABPC1* | 2 | 41 | 5 | PABPC1 | poly(A) binding protein | Cytoplasm | translation regulator | |

| | | | | | | | | | |
|---|---|---|---|---|---|---|---|---|---|
| chr7 | *ARMC10* | 2 | 63 | 5 | ARMC10 | armadillo repeat containing 10 | Cytoplasm | other | |
| chr11 | *RP13-726E6.1* | 3 | 91 | 5 | | | | | |
| chr3 | *RP11-933H2.4* | 2 | 52 | 5 | | | | | |
| chr8 | *RP11-521M14.1* | 2 | 39 | 5 | | | | | |
| chr16 | *RPUSD1* | 3 | 89 | 5 | RPUSD1 | RNA pseudouridine synthase domain containing 1 | Other | enzyme | |
| chr14 | *DLST* | 2 | 42 | 5 | DLST | dihydrolipoamide S-succinyltransferase | Cytoplasm | enzyme | |
| chr13 | *GAS6-AS1* | 3 | 85 | 5 | GAS6-AS1 | GAS6 antisense RNA 1 | Other | other | |
| chr3 | *CSTA* | 2 | 80 | 5 | CSTA | cystatin A | Cytoplasm | other | |
| chr11 | *RP11-399J13.3* | 2 | 83 | 5 | | | | | |
| chr4 | *PI4K2B* | 2 | 37 | 5 | PI4K2B | phosphatidylinositol 4-kinase type 2 beta | Cytoplasm | kinase | |
| chr10 | *ZNF503-AS1* | 3 | 139 | 5 | ZNF503-AS1 | ZNF503 antisense RNA 1 | Other | other | |
| chr5 | *LINC01018* | 3 | 242 | 5 | LINC01018 | long intergenic non-protein coding RNA 1018 | Other | other | |
| chr2 | *OXER1* | 2 | 129 | 5 | OXER1 | oxoeicosanoid receptor 1 | Plasma Membrane | G-protein coupled recepto | |
| chr6 | *T* | 2 | 101 | 5 | TBXT | T-box transcription factor T | Nucleus | transcription regulator | |
| chr15 | *RP11-597K23.2* | 2 | 54 | 5 | | | | | |
| chr17 | *RP11-46I8.3* | 2 | 69 | 5 | | | | | |
| chr16 | *RP11-473M20.11* | 2 | 72 | 5 | | | | | |
| chr16 | *MPG* | 2 | 151 | 5 | MPG | N-methylpurine DNA glycosylase | Nucleus | enzyme | |
| chr5 | *GNB2L1* | 2 | 123 | 5 | RACK1 | receptor for activated C kinase 1 | Cytoplasm | enzyme | |
| chr13 | *N4BP2L1* | 2 | 40 | 5 | N4BP2L1 | NEDD4 binding protein 2 like 1 | Other | other | |
| chr17 | *SCRN2* | 2 | 128 | 5 | SCRN2 | secernin 2 | Extracellular Space | other | |
| chr11 | *AAMDC* | 2 | 75 | 5 | AAMDC | adipogenesis associated Mth938 domain containing | Cytoplasm | other | |
| chr19 | *AC005307.1* | 2 | 70 | 5 | | | | | |
| chr19 | *ZNF729* | 2 | 70 | 5 | ZNF729 | zinc finger protein 729 | Other | other | |
| chr2 | *IL1F10* | 2 | 107 | 5 | IL1F10 | interleukin 1 family member 10 | Extracellular Space | cytokine | |
| chr19 | *CABP5* | 2 | 45 | 5 | CABP5 | calcium binding protein 5 | Cytoplasm | other | |
| chr19 | *ZNF492* | 2 | 56 | 5 | ZNF492/ZNF98 | zinc finger protein 492 | Nucleus | transcription regulator | |
| chr4 | *PRIMPOL* | 2 | 43 | 5 | PRIMPOL | primase and DNA directed polymerase | Extracellular Space | enzyme | |
| chrX | *PHKA1* | 2 | 44 | 5 | PHKA1 | phosphorylase kinase regulatory subunit alpha 1 | Cytoplasm | kinase | |
| chr1 | *LYPLAL1* | 2 | 51 | 5 | LYPLAL1 | lysophospholipase like 1 | Cytoplasm | enzyme | |
| chr5 | *SKP2* | 2 | 120 | 5 | SKP2 | S-phase kinase associated | Nucleus | other | |
| chr1 | *PLA2G2E* | 2 | 104 | 5 | PLA2G2E | phospholipase A2 group IIE | Extracellular Space | enzyme | |
| chr17 | *CNTD1* | 2 | 52 | 5 | CNTD1 | cyclin N-terminal domain | Other | other | |
| chr1 | *DPH2* | 2 | 114 | 5 | DPH2 | diphthamide biosynthesis 2 | Cytoplasm | enzyme | |
| chr19 | *CTD-2525I3.2* | 2 | 52 | 5 | | | | | |
| chr10 | *LINC00858* | 2 | 64 | 5 | LINC00858 | long intergenic non-protein coding RNA 858 | Other | other | |
| chr5 | *NHP2* | 2 | 93 | 5 | NHP2 | NHP2 ribonucleoprotein | Nucleus | other | |
| chr1 | *MRPL24* | 2 | 101 | 5 | MRPL24 | mitochondrial ribosomal protein | Cytoplasm | other | |
| chr3 | *ISY1-RAB43* | 2 | 128 | 5 | ISY1-RAB43 | ISY1-RAB43 readthrough | Nucleus | other | |
| chr5 | *MRPS36* | 2 | 48 | 5 | MRPS36 | mitochondrial ribosomal protein | Cytoplasm | other | |
| chr5 | *KRT18P16* | 2 | 88 | 5 | KRT18P16 | keratin 18 pseudogene 16 | Other | other | |
| chrX | *PAGE1* | 2 | 69 | 5 | PAGE1 | PAGE family member 1 | Other | other | |
| chr14 | *TTC5* | 2 | 43 | 5 | TTC5 | tetratricopeptide repeat domain 5 | Nucleus | other | |
| chr1 | *RP11-54O7.3* | 3 | 168 | 5 | | | | | |
| chr11 | *CTD-2560E9.3* | 3 | 98 | 5 | | | | | |
| chr8 | *KB-1507C5.4* | 2 | 43 | 5 | | | | | |
| chr10 | *AL355490.1* | 2 | 51 | 5 | | | | | |
| chr9 | *RP11-384P7.6* | 2 | 49 | 5 | | | | | |
| chr14 | *TRMT61A* | 3 | 129 | 5 | TRMT61A | tRNA methyltransferase 61A | Nucleus | enzyme | |
| chr16 | *LA16c-321D4.2* | 2 | 112 | 5 | | | | | |
| chr15 | *RP11-20G13.1* | 2 | 219 | 5 | | | | | |
| chr17 | *AC145343.2* | 2 | 87 | 5 | | | | | |
| chr19 | *RETN* | 2 | 94 | 5 | RETN | resistin | Extracellular Space | other | |
| chr17 | *SLC35B1* | 2 | 44 | 5 | SLC35B1 | solute carrier family 35 member | Cytoplasm | transporter | |
| chr17 | *C17orf66* | 2 | 52 | 5 | HEATR9 | HEAT repeat containing 9 | Other | other | |
| chr18 | *RNF152* | 3 | 102 | 5 | RNF152 | ring finger protein 152 | Cytoplasm | enzyme | |
| chr3 | *AHSG* | 2 | 76 | 5 | AHSG | alpha 2-HS glycoprotein | Extracellular Space | other | |
| chr12 | *RFC5* | 2 | 75 | 5 | RFC5 | replication factor C subunit 5 | Nucleus | enzyme | |
| chr14 | *RP11-61O1.1* | 2 | 63 | 5 | | | | | |
| chr15 | *NF1P2* | 2 | 67 | 5 | NF1P2 | neurofibromin 1 pseudogene 2 | Other | other | |
| chr8 | *LRRC14* | 3 | 120 | 5 | LRRC14 | leucine rich repeat containing 14 | Cytoplasm | other | |
| chr19 | *CYP2T3P* | 3 | 134 | 5 | CYP2T3P | cytochrome P450 family 2 subfamily T member 3, | Other | other | |
| chr12 | *HSP90B1* | 2 | 44 | 5 | HSP90B1 | heat shock protein 90 beta family member 1 | Cytoplasm | other | cisplatin, retaspimycin, |
| chr11 | *RP11-655M14.13* | 3 | 63 | 5 | | | | | |
| chr2 | *CHRNG* | 2 | 74 | 5 | CHRNG | cholinergic receptor nicotinic gamma subunit | Plasma Membrane | transmembrane recepto | succinylcholine, rapacuronium, doxacurium, vecuronium, pipecuronium, metocurine, mivacurium, D-tubocurarine, nicotine, arecoline, pancuronium, atracurium, isoflurane, cisatracurium, rocuronium, mecamylamine |

| | | | | | | | | | |
|---|---|---|---|---|---|---|---|---|---|
| chr15 | *RP11-261B23.2* | 2 | 45 | 5 | | | | | |
| chr6 | *FAM184A* | 2 | 46 | 5 | FAM184A | family with sequence similarity 184 member A | Extracellular Space | other | |
| chr15 | *RP11-96O20.4* | 2 | 41 | 5 | | | | | |
| chr12 | *RP11-834C11.11* | 2 | 130 | 5 | | | | | |
| chr14 | *TM9SF1* | 2 | 59 | 5 | TM9SF1 | transmembrane 9 superfamily member 1 | Plasma Membrane | transporter | |
| chr5 | *HAPLN1* | 2 | 38 | 5 | HAPLN1 | hyaluronan and proteoglycan link protein 1 | Extracellular Space | other | |
| chr9 | *RP11-561O23.5* | 2 | 57 | 5 | | | | | |
| chr22 | *ADORA2A-AS1* | 2 | 61 | 5 | ADORA2A-AS1 | ADORA2A antisense RNA 1 | Other | other | |
| chr7 | *RP4-736H5.3* | 2 | 36 | 5 | | | | | |
| chr11 | *EIF1AD* | 2 | 98 | 5 | EIF1AD | eukaryotic translation initiation factor 1A domain containing | Nucleus | other | |
| chr10 | *FAM213A* | 2 | 68 | 5 | PRXL2A | peroxiredoxin like 2A | Extracellular Space | other | |
| chr1 | *ITLN2* | 2 | 99 | 5 | ITLN2 | intelectin 2 | Plasma Membrane | other | |
| chr19 | *IL11* | 2 | 73 | 5 | IL11 | interleukin 11 | Extracellular Space | cytokine | |
| chr11 | *PGA4* | 2 | 129 | 5 | GA5 (includes othe | pepsinogen A3 | Extracellular Space | peptidase | sucralfate |
| chr15 | *RP11-114H24.2* | 2 | 104 | 5 | | | | | |
| chr19 | *CTC-457E21.3* | 2 | 60 | 5 | | | | | |
| chr19 | *CTD-2616J11.14* | 2 | 68 | 5 | | | | | |
| chr3 | *RP11-48F14.1* | 2 | 40 | 5 | | | | | |
| chr16 | *RNPS1* | 2 | 63 | 5 | RNPS1 | RNA binding protein with serine rich domain 1 | Nucleus | other | |
| chr17 | *RP1-37N7.3* | 2 | 57 | 5 | | | | | |
| chr12 | *RP11-834C11.4* | 3 | 172 | 5 | | | | | |
| chr8 | *JPH1* | 2 | 36 | 5 | JPH1 | junctophilin 1 | Plasma Membrane | other | |
| chr5 | *AC005618.6* | 2 | 49 | 5 | | | | | |
| chr21 | *AP001471.1* | 2 | 93 | 5 | | | | | |
| chr19 | *TRIM28* | 2 | 145 | 5 | TRIM28 | tripartite motif containing 28 | Nucleus | transcription regulator | |
| chr8 | *ALG1L11P* | 2 | 149 | 5 | ALG1L11P | | Other | other | |
| chr11 | *YIF1A* | 2 | 180 | 5 | YIF1A | Yip1 interacting factor homolog A, membrane trafficking protein | Cytoplasm | other | |
| chr6 | *TFAP2D* | 2 | 61 | 5 | TFAP2D | transcription factor AP-2 delta | Nucleus | transcription regulator | |
| chr12 | *RP11-1028N23.3* | 2 | 52 | 5 | | | | | |
| chr8 | *RP11-1057B8.2* | 2 | 51 | 5 | | | | | |
| chr2 | *CLK1* | 2 | 44 | 5 | CLK1 | CDC like kinase 1 | Nucleus | kinase | |
| chr4 | *EMCN* | 2 | 43 | 5 | EMCN | endomucin | Extracellular Space | other | |
| chr16 | *CYLD* | 2 | 41 | 5 | CYLD | CYLD lysine 63 deubiquitinase | Nucleus | transcription regulator | |
| chr17 | *RP11-388C12.5* | 2 | 106 | 5 | | | | | |
| chr8 | *PDP1* | 2 | 42 | 5 | PDP1 | pyruvate dehyrogenase phosphatase catalytic subunit 1 | Cytoplasm | phosphatase | CPI-613 |
| chr13 | *RP11-189B4.6* | 2 | 53 | 5 | | | | | |
| chr17 | *MRPL10* | 2 | 64 | 5 | MRPL10 | mitochondrial ribosomal protein | Cytoplasm | other | |
| chr7 | *AC073333.8* | 2 | 54 | 5 | | | | | |
| chr5 | *DOK3* | 3 | 126 | 5 | DOK3 | docking protein 3 | Cytoplasm | other | |
| chr5 | *CTD-2533K21.4* | 3 | 94 | 5 | | | | | |
| chr17 | *RASL10B* | 2 | 92 | 5 | RASL10B | RAS like family 10 member B | Other | enzyme | |
| chrX | *BGN* | 3 | 135 | 5 | BGN | biglycan | Extracellular Space | other | |
| chr9 | *RP11-402G3.5* | 2 | 43 | 5 | | | | | |
| chr9 | *RP11-403N16.2* | 2 | 55 | 5 | | | | | |
| chr19 | *CTC-444N24.6* | 2 | 61 | 5 | | | | | |
| chr16 | *ABCC6P2* | 2 | 85 | 5 | ABCC6P2 | ATP binding cassette subfamily C member 6 pseudogene 2 | Other | other | |
| chr17 | *RP11-527L4.6* | 2 | 86 | 5 | | | | | |
| chr8 | *AF186192.5* | 2 | 89 | 5 | | | | | |
| chr1 | *AVPR1B* | 3 | 105 | 5 | AVPR1B | arginine vasopressin receptor 1B | Plasma Membrane | -protein coupled recepto | AVP, lypressin |
| chr1 | *RP11-286M16.1* | 3 | 75 | 5 | | | | | |
| chrX | *TIMM17B* | 2 | 66 | 5 | TIMM17B | translocase of inner mitochondrial membrane 17B | Cytoplasm | transporter | |
| chr17 | *VPS25* | 2 | 60 | 5 | VPS25 | vacuolar protein sorting 25 | Cytoplasm | other | |
| chr3 | *RPL32P3* | 2 | 101 | 5 | RPL32P3 | ribosomal protein L32 | Other | other | |
| chr2 | *MCFD2* | 2 | 90 | 5 | MCFD2 | multiple coagulation factor deficiency 2 | Cytoplasm | other | |
| chr7 | *ACHE* | 3 | 124 | 5 | ACHE | acetylcholinesterase (Cartwright blood group) | Plasma Membrane | enzyme | huperzine A, pyridostigmine, edrophonium, donepezil, rivastigmine, demecarium, tacrine, neostigmine, terestigmine, pralidoxime, donepezil/mefloquine, ZT-1, gallamine triethiodide, ambenonium, BW 284 C 51, physostigmine, 2,3 butanedione monoxime, donepezil/solifenacin, donepezil/memantine, galanthamine, atropine/neostigmine, echothiophate, caracemide, [illegible] |
| chr17 | *HEXIM1* | 2 | 84 | 5 | HEXIM1 | HEXIM P-TEFb complex subunit | Nucleus | transcription regulator | |

| | | | | | | | | | |
|---|---|---|---|---|---|---|---|---|---|
| chr17 | *PPP1R9B* | 2 | 109 | 5 | PPP1R9B | protein phosphatase 1 regulatory subunit 9B | Cytoplasm | enzyme | |
| chr17 | *STRA13* | 3 | 174 | 5 | | | | | |
| chr16 | *C16orf59* | 2 | 128 | 5 | TEDC2 | tubulin epsilon and delta complex | Other | other | |
| chr17 | *CD300LB* | 2 | 97 | 5 | CD300LB | CD300 molecule like family | Plasma Membrane | other | |
| chr20 | *UBE2C* | 3 | 116 | 5 | UBE2C | ubiquitin conjugating enzyme E2 | Cytoplasm | enzyme | |
| chr6 | *MRPS18A* | 2 | 101 | 5 | MRPS18A | mitochondrial ribosomal protein | Cytoplasm | other | |
| chrX | *RENBP* | 2 | 96 | 5 | RENBP | renin binding protein | Cytoplasm | enzyme | |
| chr14 | *PSMC1* | 2 | 35 | 5 | PSMC1 | proteasome 26S subunit, ATPase | Nucleus | peptidase | |
| chr10 | *RP11-452K12.4* | 2 | 56 | 5 | | | | | |
| chr15 | *C15orf57* | 2 | 65 | 5 | CCDC32 | coiled-coil domain containing 32 | Other | other | |
| chr1 | *RP11-40H20.4* | 2 | 52 | 5 | | | | | |
| chr17 | *RPL38* | 3 | 82 | 5 | RPL38 | ribosomal protein L38 | Cytoplasm | other | |
| chr2 | *RP11-60H5.1* | 2 | 75 | 5 | | | | | |
| chr5 | *AC008394.1* | 2 | 49 | 5 | | | | | |
| chr1 | *SIKE1* | 2 | 94 | 5 | SIKE1 | suppressor of IKBKE 1 | Cytoplasm | other | |
| chr12 | *ERGIC2* | 2 | 67 | 5 | ERGIC2 | ERGIC and golgi 2 | Cytoplasm | other | |
| chr5 | *CTC-210G5.1* | 2 | 53 | 5 | | | | | |
| chr8 | *COMMD5* | 2 | 96 | 5 | COMMD5 | COMM domain containing 5 | Nucleus | other | |
| chr6 | *MB21D1* | 2 | 38 | 5 | CGAS | cyclic GMP-AMP synthase | Cytoplasm | enzyme | |
| chr8 | *PPP1R42* | 2 | 45 | 5 | PPP1R42 | protein phosphatase 1 regulatory subunit 42 | Cytoplasm | other | |
| chr16 | *RP11-463O9.9* | 2 | 107 | 5 | | | | | |
| chr16 | *RP11-105C20.2* | 2 | 54 | 5 | | | | | |
| chr18 | *OACYLP* | 2 | 67 | 5 | OACYLP | O-acyltransferase like, | Other | other | |
| chr19 | *LGALS17A* | 2 | 66 | 5 | LGALS17A | galectin 14 pseudogene | Other | other | |
| chr5 | *SREK1IP1* | 2 | 38 | 5 | SREK1IP1 | SREK1 interacting protein 1 | Other | other | |
| chr11 | *AP000936.4* | 2 | 69 | 5 | | | | | |
| chr2 | *AC012363.13* | 2 | 147 | 5 | | | | | |
| chr6 | *UNC5CL* | 3 | 94 | 5 | UNC5CL | unc-5 family C-terminal like | Cytoplasm | peptidase | |
| chr11 | *IL18BP* | 2 | 84 | 5 | IL18BP | interleukin 18 binding protein | Extracellular Space | other | |
| chrX | *H2BFM* | 2 | 115 | 5 | H2BW2 | H2B.W histone 2 | Nucleus | other | |
| chr7 | *CICP14* | 2 | 72 | 5 | P14 (includes oth | capicua transcriptional repressor pseudogene 26 | Other | other | |
| chr7 | *AC096582.1* | 2 | 109 | 5 | | | | | |
| chr9 | *SLC2A6* | 3 | 201 | 5 | SLC2A6 | solute carrier family 2 member 6 | Plasma Membrane | transporter | |
| chr19 | *NDUFA3* | 2 | 97 | 5 | NDUFA3 | NADH:ubiquinone oxidoreductase subunit A3 | Cytoplasm | enzyme | |
| chr11 | *RP11-680F20.6* | 2 | 119 | 5 | | | | | |
| chr20 | *WFDC10B* | 2 | 79 | 5 | WFDC10B | WAP four-disulfide core domain | Extracellular Space | other | |
| chr17 | *GPS1* | 3 | 89 | 5 | GPS1 | G protein pathway suppressor 1 | Nucleus | other | |
| chrX | *RAB9A* | 2 | 43 | 5 | RAB9A | RAB9A, member RAS oncogene family | Cytoplasm | enzyme | |
| chr17 | *RP11-822E23.5* | 2 | 90 | 5 | | | | | |
| chr5 | *PLAC8L1* | 2 | 40 | 5 | PLAC8L1 | PLAC8 like 1 | Other | other | |
| chr1 | *GNG5* | 2 | 41 | 5 | GNG5 | G protein subunit gamma 5 | Plasma Membrane | other | |
| chr11 | *FIBP* | 3 | 164 | 5 | FIBP | FGF1 intracellular binding protein | Nucleus | other | |
| chr17 | *DHRS11* | 2 | 63 | 5 | DHRS11 | dehydrogenase/reductase 11 | Other | other | |
| chr19 | *FUT2* | 2 | 92 | 5 | FUT2 | fucosyltransferase 2 | Cytoplasm | enzyme | |
| chr18 | *RP11-397A16.1* | 2 | 48 | 5 | | | | | |
| chr1 | *RP1-13P20.6* | 2 | 60 | 5 | | | | | |
| chr17 | *CTD-3194G12.2* | 2 | 74 | 5 | | | | | |
| chr7 | *RP11-731K22.1* | 2 | 65 | 5 | | | | | |
| chr5 | *CTD-2297D10.2* | 3 | 175 | 5 | CTD-2297D10.2 | uncharacterized LOC101929176 | Other | other | |
| chr17 | *RNF222* | 2 | 57 | 5 | RNF222 | ring finger protein 222 | Other | other | |
| chrX | *BTK* | 2 | 41 | 5 | BTK | Bruton tyrosine kinase | Cytoplasm | kinase | ARQ 531, tirabrutinib, ACP-196/ACP-319, acalabrutinib, bendamustine/ibrutinib/rituximab, ibrutinib, SHR1459, ibrutinib/rituximab, vecabrutinib, BTK inhibitor, zanubrutinib, fenebrutinib, ibrutinib/obinutuzumab, M7583, ICP-022, BMS-986142, spebrutinib, TG- |
| chr11 | *RAB30* | 2 | 52 | 5 | RAB30 | RAB30, member RAS oncogene family | Cytoplasm | enzyme | |
| chr16 | *RP11-18F14.1* | 2 | 40 | 5 | | | | | |
| chr4 | *RP11-25H12.1* | 2 | 40 | 5 | | | | | |
| chr19 | *LPAR2* | 2 | 71 | 5 | LPAR2 | lysophosphatidic acid receptor 2 | Plasma Membrane | G-protein coupled recepto | |
| chr11 | *RP11-201M22.1* | 2 | 151 | 5 | | | | | |
| chr7 | *AP1S1* | 2 | 99 | 5 | AP1S1 | adaptor related protein complex 1 subunit sigma 1 | Cytoplasm | transporter | |

| | | | | | | | | | |
|---|---|---|---|---|---|---|---|---|---|
| chr4 | *PPAT* | 2 | 60 | 5 | PPAT | phosphoribosyl pyrophosphate amidotransferase | Cytoplasm | enzyme | L-asparaginase/cyclophosphamide/cytarabine/daunorubicin/6-mercaptopurine/prednisone/vincristine, 6-mercaptopurine/prednisone/thioguanine, cytarabine/daunorubicin/thioguanine, cyclophosphamide/cytarabine/6-mercaptopurine, cytarabine/thioguanine, 6-mercaptopurine, 6-mercaptopurine/vincristine, thioguanine, amidophosphoribosyltransferase inhibitor, hydroxyurea/6-mercaptopurine, 6-mercaptopurine/pred [illegible] |
| chr19 | *CYTH2* | 2 | 94 | 5 | CYTH2 | cytohesin 2 | Cytoplasm | other | |
| chr11 | *SCGB2A1* | 2 | 73 | 5 | SCGB2A1 | secretoglobin family 2A member | Extracellular Space | other | |
| chr19 | *MBD3L1* | 2 | 45 | 5 | MBD3L1 | methyl-CpG binding domain protein 3 like 1 | Nucleus | other | |
| chr22 | *RP4-539M6.14* | 2 | 59 | 5 | | | | | |
| chr4 | *AC141928.1* | 2 | 95 | 5 | | | | | |
| chr1 | *FAM72C* | 2 | 57 | 5 | FAM72C/FAM72D | family with sequence similarity 72 member D | Nucleus | other | |
| chr6 | *MKRNP2* | 2 | 75 | 5 | | | | | |
| chr7 | *MRPS17* | 2 | 55 | 5 | MRPS17 | mitochondrial ribosomal protein | Cytoplasm | other | |
| chr12 | *RP11-804F13.1* | 2 | 51 | 5 | | | | | |
| chr17 | *MAP3K14-AS1* | 2 | 127 | 5 | MAP3K14-AS1 | MAP3K14 antisense RNA 1 | Other | other | |
| chr3 | *LINC00606* | 2 | 117 | 5 | LINC00606 | long intergenic non-protein coding RNA 606 | Other | other | |
| chr16 | *HERPUD1* | 2 | 89 | 5 | HERPUD1 | homocysteine inducible ER protein with ubiquitin like domain | Cytoplasm | other | |
| chr11 | *RPUSD4* | 2 | 65 | 5 | RPUSD4 | RNA pseudouridine synthase D4 | Cytoplasm | other | |
| chr10 | *CALML3-AS1* | 2 | 68 | 5 | CALML3-AS1 | CALML3 antisense RNA 1 | Other | other | |
| chr11 | *CTD-2516F10.2* | 2 | 65 | 5 | | | | | |
| chr16 | *LCAT* | 2 | 79 | 5 | LCAT | lecithin-cholesterol | Extracellular Space | enzyme | |
| chr4 | *INTU* | 2 | 66 | 5 | INTU | inturned planar cell polarity | Cytoplasm | other | |
| chr3 | *RP11-460N16.1* | 2 | 60 | 5 | | | | | |
| chr2 | *PLGLB2* | 2 | 74 | 5 | PLGLB1/PLGLB2 | plasminogen like B2 | Extracellular Space | peptidase | |
| chr17 | *RP13-766D20.2* | 3 | 89 | 5 | | | | | |
| chr10 | *C10orf71-AS1* | 2 | 114 | 5 | C10orf71-AS1 | C10orf71 antisense RNA 1 | Other | other | |
| chr16 | *C1QTNF8* | 2 | 251 | 5 | C1QTNF8 | C1q and TNF related 8 | Extracellular Space | other | |
| chr11 | *KCTD21* | 3 | 101 | 5 | KCTD21 | potassium channel tetramerization domain containing | Other | other | |
| chr7 | *RP5-1159O4.1* | 2 | 100 | 5 | | | | | |
| chr6 | *SLC22A16* | 2 | 49 | 5 | SLC22A16 | solute carrier family 22 member | Plasma Membrane | transporter | |
| chr17 | *CD79B* | 2 | 117 | 5 | CD79B | CD79b molecule | Plasma Membrane | transmembrane receptor | polatuzumab vedotin, bendamustine/polatuzumab |
| chr9 | *C9orf62* | 2 | 132 | 5 | C9orf62 | chromosome 9 open reading frame 62 | Other | other | |
| chr18 | *TAF4B* | 2 | 47 | 5 | TAF4B | TATA-box binding protein associated factor 4b | Nucleus | transcription regulator | |
| chr12 | *RP11-503G7.1* | 3 | 125 | 5 | | | | | |
| chr10 | *NKX2-3* | 3 | 135 | 5 | NKX2-3 | NK2 homeobox 3 | Nucleus | transcription regulator | |
| chr14 | *NGDN* | 2 | 40 | 5 | NGDN | neuroguidin | Nucleus | other | |
| chr2 | *AL133249.1* | 2 | 51 | 5 | | | | | |
| chr4 | *CPEB2-AS1* | 2 | 35 | 5 | CPEB2-DT | CPEB2 divergent transcript | Other | other | |
| chr17 | *SNF8* | 2 | 78 | 5 | SNF8 | SNF8 subunit of ESCRT-II | Cytoplasm | enzyme | |
| chr14 | *RP11-964E11.2* | 2 | 93 | 5 | | | | | |
| chr12 | *ANKRD33* | 2 | 194 | 5 | ANKRD33 | ankyrin repeat domain 33 | Nucleus | transcription regulator | |
| chr12 | *RP11-1016B18.1* | 2 | 69 | 5 | | | | | |
| chr3 | *MTHFD2P1* | 2 | 63 | 5 | MTHFD2P1 | methylenetetrahydrofolate dehydrogenase (NADP+ dependent) 2, methenyltetrahydrofolate | Other | other | |
| chr19 | *GRWD1* | 2 | 91 | 5 | GRWD1 | glutamate rich WD repeat containing 1 | Nucleus | other | |
| chr9 | *DNAJB5* | 2 | 79 | 5 | DNAJB5 | DnaJ heat shock protein family (Hsp40) member B5 | Cytoplasm | other | |
| chr11 | *SCGB1D1* | 2 | 120 | 5 | SCGB1D1 | secretoglobin family 1D member | Extracellular Space | other | |
| chr17 | *HIGD1B* | 2 | 80 | 5 | HIGD1B | HIG1 hypoxia inducible domain family member 1B | Other | other | |
| chr16 | *RBBP6* | 2 | 37 | 5 | RBBP6 | RB binding protein 6, ubiquitin | Nucleus | enzyme | |
| chr11 | *STT3A-AS1* | 2 | 48 | 5 | STT3A-AS1 | STT3A antisense RNA 1 | Other | other | |
| chr17 | *AC003958.2* | 3 | 56 | 5 | | | | | |
| chr6 | *TREML5P* | 2 | 71 | 5 | TREML5P | triggering receptor expressed on myeloid cells like 5, pseudogene | Other | other | |
| chr3 | *RPSA* | 2 | 66 | 5 | RPSA | ribosomal protein SA | Cytoplasm | translation regulator | |
| chr14 | *RP11-384J4.2* | 2 | 40 | 5 | | | | | |
| chrX | *SSX2* | 2 | 72 | 5 | SSX2/SSX2B | SSX family member 2 | Nucleus | other | |
| chr12 | *RP11-968A15.8* | 2 | 94 | 5 | | | | | |

| | | | | | | | | | |
|---|---|---|---|---|---|---|---|---|---|
| chr17 | *MIF4GD* | 2 | 110 | 5 | MIF4GD | MIF4G domain containing | Other | other | |
| chr9 | *FAM205A* | 2 | 43 | 5 | FAM205A | family with sequence similarity 205 member A | Nucleus | other | |
| chr1 | *AP4B1-AS1* | 2 | 49 | 5 | AP4B1-AS1 | AP4B1 antisense RNA 1 | Other | other | |
| chr17 | *RAB34* | 2 | 102 | 5 | RAB34 | RAB34, member RAS oncogene family | Cytoplasm | enzyme | |
| chr12 | *RP11-598F7.6* | 2 | 80 | 5 | | | | | |
| chr16 | *RP11-96H17.1* | 2 | 73 | 5 | | | | | |
| chr7 | *RP4-630C24.3* | 2 | 43 | 5 | | | | | |
| chr1 | *KRT8P28* | 2 | 123 | 5 | KRT8P28 | keratin 8 pseudogene 28 | Other | other | |
| chr6 | *SCAF8* | 2 | 40 | 5 | SCAF8 | SR-related CTD associated factor | Nucleus | other | |
| chr12 | *PTMS* | 2 | 95 | 5 | PTMS | parathymosin | Nucleus | other | |
| chr9 | *SPATA31E1* | 2 | 73 | 5 | SPATA31E1 | SPATA31 subfamily E member 1 | Other | other | |
| chr19 | *DHPS* | 2 | 62 | 5 | DHPS | deoxyhypusine synthase | Cytoplasm | enzyme | dapsone |
| chr10 | *NMT2* | 2 | 61 | 5 | NMT2 | N-myristoyltransferase 2 | Cytoplasm | enzyme | |
| chr11 | *TMEM126A* | 2 | 102 | 5 | TMEM126A | transmembrane protein 126A | Cytoplasm | other | |
| chr19 | *C19orf25* | 3 | 117 | 5 | C19orf25 | chromosome 19 open reading frame 25 | Other | other | |
| chr9 | *SPATA31B1* | 2 | 78 | 5 | SPATA31B1P | SPATA31 subfamily B member 1, pseudogene | Other | other | |
| chr11 | *SAA3P* | 2 | 54 | 5 | SAA3P | serum amyloid A3, pseudogene | Other | other | |
| chr12 | *CCT2* | 2 | 39 | 5 | CCT2 | chaperonin containing TCP1 subunit 2 | Cytoplasm | kinase | |
| chrX | *PQBP1* | 2 | 53 | 5 | PQBP1 | polyglutamine binding protein 1 | Nucleus | transcription regulator | |
| chr19 | *CDC42EP5* | 3 | 92 | 5 | CDC42EP5 | CDC42 effector protein 5 | Cytoplasm | other | |
| chr17 | *MTND2P12* | 2 | 35 | 5 | MTND2P12 | MT-ND2 pseudogene 12 | Other | other | |
| chr7 | *CICP8* | 2 | 115 | 5 | CICP8 | capicua transcriptional repressor pseudogene 8 | Other | other | |
| chr17 | *MAPK7* | 2 | 105 | 5 | MAPK7 | mitogen-activated protein kinase | Cytoplasm | kinase | |
| chr19 | *CTC-518B2.10* | 2 | 43 | 5 | | | | | |
| chr8 | *FAM150A* | 2 | 41 | 5 | ALKAL1 | ALK and LTK ligand 1 | Other | other | |
| chr19 | *TBCB* | 3 | 70 | 5 | TBCB | tubulin folding cofactor B | Cytoplasm | other | |
| chr3 | *FXR1* | 2 | 34 | 5 | FXR1 | FMR1 autosomal homolog 1 | Cytoplasm | other | |
| chr22 | *SELM* | 2 | 38 | 5 | | | | | |
| chr20 | *SLC32A1* | 3 | 139 | 5 | SLC32A1 | solute carrier family 32 member 1 | Plasma Membrane | transporter | tiagabine |
| chr12 | *SLC38A4* | 3 | 128 | 5 | SLC38A4 | solute carrier family 38 member 4 | Plasma Membrane | transporter | |
| chr15 | *WHAMMP2* | 2 | 39 | 5 | WHAMMP2 | WAS protein homolog associated with actin, golgi membranes and microtubules pseudogene 2 | Other | other | |
| chrX | *RP11-571E6.1* | 2 | 61 | 5 | | | | | |
| chr18 | *CTC-782O7.1* | 2 | 34 | 5 | | | | | |
| chr10 | *RP11-57C13.4* | 2 | 50 | 5 | | | | | |
| chr10 | *HSD17B7P2* | 2 | 83 | 5 | HSD17B7P2 | hydroxysteroid 17-beta dehydrogenase 7 pseudogene 2 | Other | other | |
| chr7 | *RP11-460N20.4* | 2 | 55 | 5 | | | | | |
| chr6 | *CLPS* | 2 | 74 | 5 | CLPS | colipase | Extracellular Space | other | |
| chr8 | *FAM90A2P* | 2 | 129 | 5 | FAM90A2P | family with sequence similarity 90 member A2, pseudogene | Other | other | |
| chr5 | *RP11-321E2.2* | 2 | 1762 | 5 | | | | | |
| chr1 | *RP1-58B11.1* | 2 | 69 | 5 | | | | | |
| chrX | *RBBP7* | 2 | 48 | 5 | RBBP7 | RB binding protein 7, chromatin remodeling factor | Nucleus | transcription regulator | |
| chr1 | *RP11-5P18.10* | 2 | 197 | 5 | | | | | |
| chr2 | *MOGAT1* | 2 | 35 | 5 | MOGAT1 | monoacylglycerol O- | Cytoplasm | enzyme | |
| chr3 | *IL5RA* | 2 | 36 | 5 | IL5RA | interleukin 5 receptor subunit | Plasma Membrane | transmembrane recepto | MEDI-563 |
| chrX | *BMX* | 2 | 43 | 5 | BMX | BMX non-receptor tyrosine | Cytoplasm | kinase | |
| chr15 | *RP11-133K1.6* | 2 | 46 | 5 | | | | | |
| chr19 | *C19orf66* | 3 | 69 | 5 | SHFL | shiftless antiviral inhibitor of ribosomal frameshifting | Other | other | |
| chr20 | *RP4-694B14.5* | 2 | 34 | 5 | | | | | |
| chr19 | *CYP4F10P* | 2 | 52 | 5 | CYP4F10P | cytochrome P450 family 4 subfamily F member 10, | Other | other | |
| chr19 | *HIPK4* | 3 | 105 | 5 | HIPK4 | homeodomain interacting protein kinase 4 | Cytoplasm | kinase | |
| chrX | *RP11-66N11.7* | 2 | 79 | 5 | | | | | |
| chr18 | *POLI* | 2 | 63 | 5 | POLI | DNA polymerase iota | Nucleus | enzyme | |
| chr6 | *ULBP2* | 2 | 76 | 5 | ULBP2 | UL16 binding protein 2 | Plasma Membrane | transmembrane recepto | |
| chr19 | *ZNF784* | 2 | 80 | 5 | ZNF784 | zinc finger protein 784 | Other | other | |
| chr9 | *PTGDS* | 3 | 167 | 5 | PTGDS | prostaglandin D2 synthase | Cytoplasm | enzyme | |
| chr2 | *SLC5A7* | 2 | 56 | 5 | SLC5A7 | solute carrier family 5 member 7 | Plasma Membrane | transporter | |
| chr7 | *SPAM1* | 2 | 37 | 5 | SPAM1 | sperm adhesion molecule 1 | Plasma Membrane | enzyme | |
| chr1 | *RP11-181G12.4* | 3 | 102 | 5 | | | | | |
| chr12 | *LINC00485* | 3 | 64 | 5 | LINC00485 | long intergenic non-protein coding RNA 485 | Other | other | |
| chr21 | *CSTB* | 2 | 76 | 5 | CSTB | cystatin B | Cytoplasm | peptidase | |
| chr12 | *GALNT8* | 2 | 55 | 5 | GALNT8 | polypeptide N-acetylgalactosaminyltransferase | Cytoplasm | enzyme | |
| chr3 | *RP11-362A9.3* | 2 | 53 | 5 | | | | | |
| chr5 | *RP11-159F24.2* | 3 | 180 | 5 | | | | | |
| chr9 | *RP11-95K23.3* | 2 | 62 | 5 | | | | | |
| chr10 | *RP11-436D10.3* | 2 | 116 | 5 | | | | | |
| chr8 | *FAM85B* | 2 | 189 | 5 | FAM85B | | Other | other | |
| chr16 | *SSTR5-AS1* | 3 | 115 | 5 | SSTR5-AS1 | SSTR5 antisense RNA 1 | Other | other | |
| chr4 | *LINC00499* | 2 | 41 | 5 | LINC00499 | long intergenic non-protein coding RNA 499 | Other | other | |
| chr16 | *PABPN1L* | 2 | 121 | 5 | PABPN1L | PABPN1 like, cytoplasmic | Cytoplasm | other | |
| chr6 | *IL22RA2* | 2 | 35 | 5 | IL22RA2 | interleukin 22 receptor subunit alpha 2 | Plasma Membrane | transmembrane recepto | |

| | | | | | | | | | |
|---|---|---|---|---|---|---|---|---|---|
| chr2 | *ITGB1BP1* | 2 | 70 | 5 | ITGB1BP1 | integrin subunit beta 1 binding protein 1 | Plasma Membrane | other | |
| chr13 | *LINC00368* | 2 | 52 | 5 | LINC00368 | long intergenic non-protein coding RNA 368 | Other | other | |
| chr14 | *TBPL2* | 2 | 43 | 5 | TBPL2 | TATA-box binding protein like 2 | Nucleus | transcription regulator | |
| chr1 | *KRT8P21* | 2 | 97 | 5 | KRT8P21 | keratin 8 pseudogene 21 | Other | other | |
| chr1 | *RP11-365O16.3* | 2 | 74 | 5 | | | | | |
| chr19 | *C19orf48* | 3 | 57 | 5 | C19orf48 | chromosome 19 open reading frame 48 | Other | other | |
| chr8 | *FAM66D* | 3 | 71 | 5 | FAM66D | family with sequence similarity 66 member D | Other | other | |
| chr12 | *RP11-171L9.1* | 2 | 62 | 5 | | | | | |
| chr19 | *IZUMO1* | 2 | 71 | 5 | IZUMO1 | izumo sperm-egg fusion 1 | Cytoplasm | other | |
| chr12 | *LINC00944* | 2 | 56 | 5 | LINC00944 | long intergenic non-protein coding RNA 944 | Other | other | |
| chr19 | *LIM2* | 2 | 64 | 5 | LIM2 | lens intrinsic membrane protein 2 | Plasma Membrane | other | |
| chr17 | *AC104024.1* | 2 | 62 | 5 | | | | | |
| chr8 | *RP11-145O15.3* | 2 | 54 | 5 | | | | | |
| chr9 | *C9orf41* | 2 | 34 | 5 | CARNMT1 | carnosine N-methyltransferase 1 | Cytoplasm | enzyme | |
| chr9 | *TNFSF8* | 2 | 41 | 5 | TNFSF8 | TNF superfamily member 8 | Plasma Membrane | cytokine | |
| chr7 | *MEPCE* | 2 | 133 | 5 | MEPCE | methylphosphate capping | Other | enzyme | |
| chr8 | *MTMR9* | 2 | 38 | 5 | MTMR9 | myotubularin related protein 9 | Cytoplasm | phosphatase | |
| chr9 | *WDR38* | 2 | 150 | 5 | WDR38 | WD repeat domain 38 | Other | other | |
| chr12 | *TUBA1A* | 2 | 109 | 5 | TUBA1A | tubulin alpha 1a | Cytoplasm | other | docetaxel/gemcitabine/vincristine, cyclophosphamide/epirubicin/5-fluorouracil/paclitaxel, bevacizumab/paclitaxel/topotecan, colchicine/probenecid, bevacizumab/docetaxel/prednisone/thalidomide/zoledronic acid, gemcitabine/paclitaxel, L-asparaginase/daunorubicin/dexamethasone/vincristine, prednisone/vincristine, docetaxel/5-fluorouracil, vinflunine, [illegible] |
| chr17 | *KRT28* | 2 | 94 | 5 | KRT28 | keratin 28 | Cytoplasm | other | |
| chr12 | *RP11-338E21.1* | 2 | 67 | 5 | | | | | |
| chr7 | *CHCHD2* | 2 | 96 | 5 | CHCHD2 | coiled-coil-helix-coiled-coil-helix domain containing 2 | Cytoplasm | other | |
| chr2 | *MOGS* | 2 | 110 | 5 | MOGS | mannosyl-oligosaccharide glucosidase | Cytoplasm | enzyme | |
| chr12 | *NEDD1* | 2 | 34 | 5 | NEDD1 | NEDD1 gamma-tubulin ring complex targeting factor | Cytoplasm | other | |
| chrX | *MAGED4B* | 2 | 121 | 5 | IAGED4/MAGED4 | MAGE family member D4B | Other | other | |
| chr6 | *TREML3P* | 2 | 60 | 5 | TREML3P | triggering receptor expressed on myeloid cells like 3, pseudogene | Other | other | |
| chr17 | *LYZL6* | 2 | 51 | 5 | LYZL6 | lysozyme like 6 | Other | enzyme | |
| chr5 | *CTD-3080P12.3* | 3 | 342 | 5 | CTD-3080P12.3 | uncharacterized LOC101928857 | Other | other | |
| chr1 | *RP11-244H3.1* | 2 | 54 | 5 | | | | | |
| chr10 | *RP11-451M19.3* | 2 | 75 | 5 | | | | | |
| chr4 | *CFI* | 2 | 35 | 5 | CFI | complement factor I | Extracellular Space | peptidase | |
| chr11 | *RP11-687M24.4* | 2 | 55 | 5 | | | | | |
| chr11 | *C11orf86* | 2 | 151 | 5 | C11orf86 | chromosome 11 open reading frame 86 | Other | other | |
| chr12 | *RP11-1020M18.1[illegible]* | 2 | 137 | 5 | | | | | |
| chr16 | *RP11-347C12.8* | 2 | 112 | 5 | | | | | |
| chr8 | *CASC7* | 2 | 69 | 5 | AGO2 | argonaute RISC catalytic component 2 | Cytoplasm | translation regulator | |
| chr17 | *RP11-63A1.1* | 2 | 40 | 5 | | | | | |
| chr1 | *RRNAD1* | 2 | 130 | 5 | RRNAD1 | ribosomal RNA adenine dimethylase domain containing 1 | Other | other | |
| chr5 | *CTD-2029E14.1* | 2 | 170 | 5 | | | | | |
| chr17 | *RP11-6N17.4* | 2 | 56 | 5 | | | | | |
| chr1 | *RP11-122G18.5* | 2 | 44 | 5 | | | | | |
| chr15 | *RP11-89K11.1* | 2 | 90 | 5 | | | | | |
| chr17 | *RP11-400F19.8* | 2 | 92 | 5 | | | | | |
| chr5 | *GZMAP1* | 2 | 59 | 5 | GZMAP1 | granzyme A pseudogene 1 | Other | other | |
| chr19 | *FAM187B* | 2 | 97 | 5 | FAM187B | family with sequence similarity 187 member B | Other | other | |
| chr5 | *CCT5* | 3 | 140 | 5 | CCT5 | chaperonin containing TCP1 subunit 5 | Cytoplasm | other | |
| chr2 | *BOK* | 3 | 140 | 5 | BOK | BCL2 family apoptosis regulator | Cytoplasm | other | |
| chr19 | *SPRED3* | 2 | 69 | 5 | SPRED3 | sprouty related EVH1 domain containing 3 | Extracellular Space | cytokine | |
| chr1 | *THBS3* | 2 | 145 | 5 | THBS3 | thrombospondin 3 | Extracellular Space | other | |
| chr21 | *AIRE* | 3 | 108 | 5 | AIRE | autoimmune regulator | Nucleus | transcription regulator | |
| chr12 | *SPIC* | 2 | 49 | 5 | SPIC | Spi-C transcription factor | Nucleus | transcription regulator | |

| | | | | | | | | | |
|---|---|---|---|---|---|---|---|---|---|
| chr11 | *ART5* | 2 | 209 | 5 | ART5 | ADP-ribosyltransferase 5 | Extracellular Space | enzyme | |
| chr9 | *XXyac-YM21GA2.* | 2 | 62 | 5 | | | | | |
| chr14 | *RP11-597A11.1* | 2 | 41 | 5 | | | | | |
| chr15 | *MTHFS* | 2 | 42 | 5 | MTHFS | methenyltetrahydrofolate | Cytoplasm | enzyme | |
| chr15 | *DET1* | 2 | 58 | 5 | DET1 | DET1 partner of COP1 E3 ubiquitin ligase | Nucleus | other | |
| chr22 | *CHADL* | 2 | 105 | 4 | CHADL | chondroadherin like | Extracellular Space | other | |
| chr18 | *PSMG2* | 2 | 48 | 4 | PSMG2 | proteasome assembly chaperone | Nucleus | other | |
| chr7 | *FAM3C* | 2 | 41 | 4 | FAM3C | family with sequence similarity 3 member C | Extracellular Space | cytokine | |
| chr14 | *MIR381HG* | 2 | 60 | 4 | MIR381HG | MIR381 host gene | Other | other | |
| chr6 | *SLC35B2* | 2 | 89 | 4 | SLC35B2 | solute carrier family 35 member | Cytoplasm | transporter | |
| chr2 | *AC105342.1* | 3 | 100 | 4 | | | | | |
| chrX | *SYTL4* | 3 | 75 | 4 | SYTL4 | synaptotagmin like 4 | Cytoplasm | transporter | |
| chr4 | *RPL34-AS1* | 2 | 51 | 4 | RPL34-AS1 | RPL34 antisense RNA 1 (head to head) | Other | other | |
| chr22 | *KB-67B5.17* | 2 | 111 | 4 | | | | | |
| chr14 | *PSMB11* | 2 | 84 | 4 | PSMB11 | proteasome subunit beta 11 | Cytoplasm | peptidase | |
| chr19 | *DCAF15* | 3 | 131 | 4 | DCAF15 | DDB1 and CUL4 associated factor 15 | Other | other | |
| chr5 | *RP11-155L15.1* | 2 | 54 | 4 | | | | | |
| chr4 | *RP11-402J6.1* | 2 | 49 | 4 | | | | | |
| chr15 | *RP11-380D11.2* | 2 | 66 | 4 | | | | | |
| chr1 | *RP11-431K24.4* | 2 | 56 | 4 | | | | | |
| chr19 | *CTD-3018O17.3* | 2 | 52 | 4 | | | | | |
| chrX | *ZDHHC9* | 2 | 33 | 4 | ZDHHC9 | zinc finger DHHC-type containing | Cytoplasm | enzyme | |
| chr10 | *RP11-225H22.4* | 2 | 83 | 4 | | | | | |
| chr3 | *RP11-231E6.1* | 2 | 50 | 4 | | | | | |
| chr16 | *CTD-2014E2.5* | 2 | 52 | 4 | | | | | |
| chr17 | *NEK8* | 2 | 73 | 4 | NEK8 | NIMA related kinase 8 | Nucleus | kinase | |
| chr17 | *RNF112* | 2 | 108 | 4 | RNF112 | ring finger protein 112 | Nucleus | transcription regulator | |
| chr7 | *FAM131B* | 2 | 73 | 4 | FAM131B | family with sequence similarity 131 member B | Other | other | |
| chr3 | *NUP210P2* | 2 | 92 | 4 | NUP210P2 | nucleoporin 210 pseudogene 2 | Other | other | |
| chr5 | *RFESD* | 2 | 66 | 4 | RFESD | Rieske Fe-S domain containing | Other | other | |
| chr3 | *NDUFB4* | 2 | 78 | 4 | NDUFB4 | NADH:ubiquinone oxidoreductase subunit B4 | Cytoplasm | transporter | |
| chr2 | *RPL23AP7* | 2 | 53 | 4 | RPL23AP7 | ribosomal protein L23a pseudogene 7 | Other | other | |
| chr8 | *TCF24* | 2 | 77 | 4 | TCF24 | transcription factor 24 | Other | other | |
| chr15 | *RCN2* | 2 | 40 | 4 | RCN2 | reticulocalbin 2 | Cytoplasm | other | |
| chr18 | *SOCS6* | 2 | 51 | 4 | SOCS6 | suppressor of cytokine signaling | Cytoplasm | other | |
| chr11 | *AC068858.1* | 2 | 102 | 4 | | | | | |
| chr10 | *RP11-432J9.3* | 2 | 117 | 4 | | | | | |
| chr16 | *TPSP2* | 3 | 132 | 4 | TPSP2 | tryptase pseudogene 2 | Other | other | |
| chr17 | *INCA1* | 2 | 75 | 4 | INCA1 | inhibitor of CDK, cyclin A1 interacting protein 1 | Nucleus | other | |
| chr2 | *AC079779.4* | 2 | 73 | 4 | | | | | |
| chr17 | *ERAL1* | 2 | 49 | 4 | ERAL1 | Era like 12S mitochondrial rRNA chaperone 1 | Cytoplasm | other | |
| chr15 | *RP11-152F13.7* | 2 | 65 | 4 | | | | | |
| chr5 | *RP11-53O19.1* | 2 | 96 | 4 | | | | | |
| chr6 | *HMGA1* | 2 | 115 | 4 | HMGA1 | high mobility group AT-hook 1 | Nucleus | transcription regulator | |
| chr19 | *RP11-678G14.3* | 3 | 62 | 4 | | | | | |
| chr9 | *RP11-447M12.2* | 2 | 126 | 4 | | | | | |
| chr8 | *NKX2-6* | 3 | 113 | 4 | NKX2-6 | NK2 homeobox 6 | Nucleus | transcription regulator | |
| chr15 | *ALDH1A3* | 2 | 104 | 4 | ALDH1A3 | aldehyde dehydrogenase 1 family member A3 | Cytoplasm | enzyme | |
| chr6 | *OARD1* | 2 | 69 | 4 | OARD1 | O-acyl-ADP-ribose deacylase 1 | Nucleus | enzyme | |
| chr11 | *DEPDC7* | 2 | 50 | 4 | DEPDC7 | DEP domain containing 7 | Cytoplasm | other | |
| chr7 | *LRWD1* | 2 | 114 | 4 | LRWD1 | leucine rich repeats and WD repeat domain containing 1 | Nucleus | other | |
| chr1 | *RP11-86H7.7* | 2 | 101 | 4 | | | | | |
| chr20 | *NRSN2* | 2 | 87 | 4 | NRSN2 | neurensin 2 | Plasma Membrane | other | |
| chr19 | *AC011516.2* | 2 | 55 | 4 | | | | | |
| chr3 | *CD47* | 2 | 55 | 4 | CD47 | CD47 molecule | Plasma Membrane | transmembrane receptor | TJ011133, hu5F9-G4, TTI-622, AO-176, CC-90002, TTI-621, ALX148 |
| chr11 | *RP11-619A14.3* | 3 | 119 | 4 | | | | | |
| chr11 | *NXF1* | 2 | 65 | 4 | NXF1 | nuclear RNA export factor 1 | Nucleus | other | |
| chr1 | *CLDN19* | 2 | 124 | 4 | CLDN19 | claudin 19 | Plasma Membrane | other | |
| chr16 | *AC012322.1* | 2 | 38 | 4 | | | | | |
| chr19 | *ABHD17A* | 3 | 119 | 4 | ABHD17A | abhydrolase domain containing | Plasma Membrane | enzyme | |
| chr5 | *CTC-498J12.3* | 2 | 42 | 4 | | | | | |
| chr12 | *LINC00939* | 2 | 42 | 4 | LINC00939 | long intergenic non-protein coding RNA 939 | Other | other | |
| chr2 | *TNFAIP6* | 2 | 38 | 4 | TNFAIP6 | TNF alpha induced protein 6 | Extracellular Space | other | |
| chr16 | *RP11-598D12.3* | 2 | 94 | 4 | | | | | |
| chr7 | *AC002480.5* | 2 | 52 | 4 | | | | | |
| chr1 | *S100A7A* | 2 | 74 | 4 | S100A7A | S100 calcium binding protein A7A | Cytoplasm | other | |
| chr11 | *RP11-17A1.3* | 2 | 34 | 4 | | | | | |
| chr6 | *RP1-153P14.5* | 2 | 81 | 4 | | | | | |
| chr5 | *PART1* | 2 | 45 | 4 | PART1 | prostate androgen-regulated transcript 1 | Other | other | |
| chr19 | *CCNE1* | 2 | 47 | 4 | CCNE1 | cyclin E1 | Nucleus | transcription regulator | |
| chr1 | *C1orf115* | 3 | 68 | 4 | C1orf115 | chromosome 1 open reading frame 115 | Other | other | |
| chr5 | *CTD-2151A2.1* | 2 | 50 | 4 | | | | | |

| | | | | | | | | | |
|---|---|---|---|---|---|---|---|---|---|
| chr4 | *FAM198B* | 2 | 43 | 4 | GASK1B | golgi associated kinase 1B | Cytoplasm | other | |
| chr18 | *CABYR* | 2 | 49 | 4 | CABYR | calcium binding tyrosine phosphorylation regulated | Cytoplasm | other | |
| chr11 | *ZNF202* | 2 | 102 | 4 | ZNF202 | zinc finger protein 202 | Nucleus | transcription regulator | |
| chr7 | *CDK5* | 2 | 66 | 4 | CDK5 | cyclin dependent kinase 5 | Nucleus | kinase | CYC065, alvocidib, dinaciclib |
| chrX | *ARSF* | 2 | 34 | 4 | ARSF | arylsulfatase F | Extracellular Space | enzyme | |
| chr13 | *GTF3A* | 2 | 43 | 4 | GTF3A | general transcription factor IIIA | Nucleus | transcription regulator | |
| chr17 | *LRRC37A2* | 2 | 64 | 4 | C37A3 (includes o | leucine rich repeat containing 37 member A3 | Other | other | |
| chrX | *CFP* | 2 | 89 | 4 | CFP | complement factor properdin | Extracellular Space | other | |
| chrX | *SLC35A2* | 2 | 125 | 4 | SLC35A2 | solute carrier family 35 member | Cytoplasm | enzyme | |
| chr12 | *KITLG* | 2 | 43 | 4 | KITLG | KIT ligand | Extracellular Space | growth factor | |
| chr19 | *WDR83* | 2 | 61 | 4 | WDR83 | WD repeat domain 83 | Nucleus | other | |
| chr1 | *S100A10* | 2 | 76 | 4 | S100A10 | S100 calcium binding protein A10 | Cytoplasm | other | |
| chr2 | *AC016757.3* | 2 | 91 | 4 | | | | | |
| chr3 | *FANCD2OS* | 3 | 102 | 4 | FANCD2OS | FANCD2 opposite strand | Other | other | |
| chr10 | *RP11-351M16.3* | 2 | 33 | 4 | | | | | |
| chr11 | *MS4A5* | 2 | 73 | 4 | MS4A5 | membrane spanning 4-domains | Other | other | |
| chr6 | *ULBP1* | 2 | 67 | 4 | ULBP1 | UL16 binding protein 1 | Plasma Membrane | transmembrane receptor | |
| chr1 | *ZNF692* | 2 | 83 | 4 | ZNF692 | zinc finger protein 692 | Nucleus | transcription regulator | |
| chr9 | *RP11-83J21.3* | 2 | 134 | 4 | | | | | |
| chr7 | *GIMAP6* | 2 | 68 | 4 | GIMAP6 | GTPase, IMAP family member 6 | Cytoplasm | other | |
| chr14 | *MPP5* | 2 | 38 | 4 | MPP5 | membrane palmitoylated protein | Plasma Membrane | kinase | |
| chr1 | *HIST2H4A* | 2 | 58 | 4 | H4C14 | H4 clustered histone 14 | Nucleus | other | |
| chr1 | *MTFR1L* | 2 | 57 | 4 | MTFR1L | mitochondrial fission regulator 1 | Other | other | |
| chr3 | *GLYCTK-AS1* | 2 | 103 | 4 | GLYCTK-AS1 | GLYCTK antisense RNA 1 | Other | other | |
| chr12 | *HOXC-AS3* | 2 | 161 | 4 | HOXC-AS3 | HOXC cluster antisense RNA 3 | Other | other | |
| chr2 | *CD207* | 2 | 59 | 4 | CD207 | CD207 molecule | Plasma Membrane | other | |
| chr11 | *BRMS1* | 2 | 126 | 4 | BRMS1 | BRMS1 transcriptional repressor and anoikis regulator | Nucleus | transcription regulator | |
| chr16 | *MIR940* | 2 | 98 | 4 | mir-940 | microRNA 940 | Cytoplasm | microRNA | |
| chr1 | *RP11-398M15.1* | 2 | 37 | 4 | | | | | |
| chr17 | *KRT12* | 2 | 60 | 4 | KRT12 | keratin 12 | Cytoplasm | other | |
| chr17 | *LRRC59* | 2 | 60 | 4 | LRRC59 | leucine rich repeat containing 59 | Cytoplasm | other | |
| chr11 | *TMEM132A* | 3 | 174 | 4 | TMEM132A | transmembrane protein 132A | Cytoplasm | other | |
| chr1 | *PSRC1* | 2 | 154 | 4 | PSRC1 | proline and serine rich coiled-coil | Cytoplasm | other | |
| chr3 | *TNNC1* | 2 | 99 | 4 | TNNC1 | troponin C1, slow skeletal and cardiac type | Cytoplasm | other | levosimendan, Ca2+ |
| chr2 | *SPR* | 2 | 74 | 4 | SPR | sepiapterin reductase | Cytoplasm | enzyme | |
| chr4 | *EXOC1* | 2 | 49 | 4 | EXOC1 | exocyst complex component 1 | Cytoplasm | transporter | |
| chr10 | *ABCD1P2* | 2 | 115 | 4 | ABCD1P2 | ATP binding cassette subfamily D member 1 pseudogene 2 | Other | other | |
| chr19 | *AC024592.9* | 2 | 54 | 4 | | | | | |
| chr22 | *JOSD1* | 2 | 46 | 4 | JOSD1 | Josephin domain containing 1 | Cytoplasm | peptidase | |
| chr3 | *LMOD3* | 2 | 45 | 4 | LMOD3 | leiomodin 3 | Cytoplasm | other | |
| chr16 | *RP11-473M20.9* | 2 | 97 | 4 | | | | | |
| chr16 | *RNF151* | 2 | 111 | 4 | RNF151 | ring finger protein 151 | Nucleus | other | |
| chr17 | *CTB-131K11.1* | 2 | 63 | 4 | | | | | |
| chr17 | *UBALD2* | 2 | 70 | 4 | UBALD2 | UBA like domain containing 2 | Other | other | |
| chr5 | *GABRP* | 2 | 47 | 4 | GABRP | gamma-aminobutyric acid type A receptor pi subunit | Plasma Membrane | ion channel | nitrazepam, sevoflurane, etomidate, zolpidem, phenobarbital, diazepam, isoflurane, halothane, pentobarbital, desflurane, adinazolam, zaleplon, enflurane, flurazepam, metyrapone/oxazepam, isoniazid, midazolam, temazepam, diphenhydramine/midazolam, estazolam, felbamate, alphadolone, methoxyflurane, secobarbital, fluoxetine/olanzapine, eszopiclone, [illegible] |
| chr9 | *ACO1* | 2 | 70 | 4 | ACO1 | aconitase 1 | Cytoplasm | enzyme | |
| chr2 | *SLC19A3* | 2 | 39 | 4 | SLC19A3 | solute carrier family 19 member 3 | Plasma Membrane | transporter | |
| chr3 | *CD200R1* | 2 | 69 | 4 | CD200R1 | CD200 receptor 1 | Plasma Membrane | transmembrane receptor | |
| chr10 | *RP11-113E21.1* | 2 | 36 | 4 | | | | | |
| chr8 | *MAPK15* | 2 | 78 | 4 | MAPK15 | mitogen-activated protein kinase | Cytoplasm | kinase | |
| chr11 | *CTD-2531D15.4* | 2 | 126 | 4 | | | | | |
| chr17 | *CTD-2514K5.2* | 2 | 90 | 4 | | | | | |
| chr16 | *NUPR1* | 2 | 86 | 4 | NUPR1 | nuclear protein 1, transcriptional regulator | Nucleus | transcription regulator | |

| | | | | | | | | | |
|---|---|---|---|---|---|---|---|---|---|
| chr6 | *GABRR1* | 2 | 58 | 4 | GABRR1 | gamma-aminobutyric acid type A receptor rho1 subunit | Plasma Membrane | ion channel | triazolam, flurazepam, metyrapone/oxazepam, nitrazepam, diazepam, midazolam, temazepam, quazepam, diphenhydramine/mid |
| chr2 | *AC018804.6* | 2 | 48 | 4 | | | | | |
| chr8 | *NUDCD1* | 2 | 38 | 4 | NUDCD1 | NudC domain containing 1 | Nucleus | other | |
| chr15 | *ZSCAN29* | 2 | 48 | 4 | ZSCAN29 | zinc finger and SCAN domain containing 29 | Nucleus | transcription regulator | |
| chr9 | *VLDLR* | 2 | 54 | 4 | VLDLR | very low density lipoprotein | Plasma Membrane | transporter | |
| chr20 | *RP4-640H8.2* | 2 | 73 | 4 | | | | | |
| chr22 | *LA16c-83F12.6* | 2 | 89 | 4 | | | | | |
| chr9 | *PAEP* | 2 | 148 | 4 | PAEP | progestagen associated endometrial protein | Cytoplasm | other | |
| chr10 | *LINC00202-2* | 2 | 67 | 4 | FAM238B | family with sequence similarity 238 member B | Other | other | |
| chr20 | *LINC00489* | 2 | 67 | 4 | LINC00489 | long intergenic non-protein coding RNA 489 | Other | other | |
| chr12 | *C12orf45* | 2 | 84 | 4 | C12orf45 | chromosome 12 open reading frame 45 | Other | other | |
| chr9 | *C9orf116* | 2 | 116 | 4 | C9orf116 | chromosome 9 open reading frame 116 | Nucleus | other | |
| chr17 | *RP11-160O5.1* | 2 | 50 | 4 | | | | | |
| chrX | *TRMT2B* | 2 | 37 | 4 | TRMT2B | tRNA methyltransferase 2 | Cytoplasm | enzyme | |
| chr2 | *SCN3A* | 2 | 32 | 4 | SCN3A | sodium voltage-gated channel alpha subunit 3 | Plasma Membrane | ion channel | riluzole, zonisamide |
| chr3 | *MYLKP1* | 2 | 78 | 4 | MYLKP1 | myosin light chain kinase pseudogene 1 | Other | other | |
| chr11 | *RP11-452H21.1* | 2 | 75 | 4 | | | | | |
| chr19 | *AZU1* | 2 | 95 | 4 | AZU1 | azurocidin 1 | Cytoplasm | peptidase | |
| chr21 | *LINC00114* | 2 | 41 | 4 | LINC00114 | long intergenic non-protein coding RNA 114 | Other | other | |
| chrX | *SSR4* | 2 | 101 | 4 | SSR4 | signal sequence receptor subunit | Cytoplasm | other | |
| chr15 | *RP11-59H7.3* | 2 | 52 | 4 | | | | | |
| chr1 | *RP11-547D24.1* | 2 | 61 | 4 | | | | | |
| chr8 | *ZNF706* | 2 | 36 | 4 | ZNF706 | zinc finger protein 706 | Other | other | |
| chr6 | *TRDN* | 2 | 45 | 4 | TRDN | triadin | Cytoplasm | other | |
| chr1 | *SLC19A2* | 2 | 49 | 4 | SLC19A2 | solute carrier family 19 member 2 | Plasma Membrane | transporter | |
| chr19 | *ZNF846* | 2 | 72 | 4 | ZNF846 | zinc finger protein 846 | Other | other | |
| chr19 | *FXYD3* | 2 | 119 | 4 | FXYD3 | FXYD domain containing ion transport regulator 3 | Plasma Membrane | ion channel | |
| chr13 | *SMIM2-AS1* | 2 | 34 | 4 | SMIM2-AS1 | SMIM2 antisense RNA 1 | Other | other | |
| chr8 | *C8orf46* | 2 | 77 | 4 | VXN | vexin | Other | other | |
| chr2 | *AC104653.1* | 2 | 56 | 4 | | | | | |
| chr19 | *CLEC4GP1* | 2 | 63 | 4 | CLEC4GP1 | C-type lectin domain family 4 member G pseudogene 1 | Other | other | |
| chr1 | *SLAMF8* | 2 | 88 | 4 | SLAMF8 | SLAM family member 8 | Extracellular Space | other | |
| chr19 | *CTD-3220F14.1* | 2 | 230 | 4 | | | | | |
| chr14 | *RP11-305B6.3* | 2 | 34 | 4 | | | | | |
| chr7 | *GPR85* | 2 | 48 | 4 | GPR85 | G protein-coupled receptor 85 | Plasma Membrane | i-protein coupled recepto | |
| chr19 | *CTB-167G5.5* | 2 | 62 | 4 | | | | | |
| chr7 | *AC147651.3* | 2 | 135 | 4 | | | | | |
| chr8 | *REXO1L1* | 2 | 265 | 4 | REXO1L1P | REXO1 like 1, pseudogene | Other | other | |
| chr1 | *RP11-382D8.5* | 2 | 92 | 4 | | | | | |
| chr15 | *GCHFR* | 2 | 77 | 4 | GCHFR | GTP cyclohydrolase I feedback regulator | Cytoplasm | other | |
| chr5 | *RP11-260E18.1* | 2 | 114 | 4 | | | | | |
| chr1 | *TMEM69* | 2 | 55 | 4 | TMEM69 | transmembrane protein 69 | Other | other | |
| chr1 | *RP11-276H7.3* | 2 | 81 | 4 | | | | | |
| chr10 | *C10orf99* | 3 | 110 | 4 | C10orf99 | chromosome 10 open reading frame 99 | Extracellular Space | cytokine | |
| chr17 | *ABCA6* | 2 | 38 | 4 | ABCA6 | ATP binding cassette subfamily A member 6 | Plasma Membrane | transporter | |
| chr19 | *ATG4D* | 2 | 71 | 4 | ATG4D | autophagy related 4D cysteine peptidase | Cytoplasm | peptidase | |
| chr17 | *WDR45B* | 2 | 65 | 4 | WDR45B | WD repeat domain 45B | Cytoplasm | other | |
| chr16 | *RP11-347C12.3* | 2 | 125 | 4 | | | | | |
| chr18 | *TMX3* | 2 | 36 | 4 | TMX3 | thioredoxin related transmembrane protein 3 | Cytoplasm | enzyme | |
| chr10 | *RP11-120C12.3* | 2 | 60 | 4 | | | | | |
| chr15 | *AC007950.1* | 2 | 48 | 4 | | | | | |
| chr1 | *RP1-212P9.3* | 2 | 60 | 4 | | | | | |
| chr15 | *RP11-361M10.3* | 2 | 72 | 4 | | | | | |
| chr19 | *KLK1* | 2 | 75 | 4 | KLK1 | kallikrein 1 | Cytoplasm | peptidase | |
| chr15 | *DNM1P34* | 2 | 90 | 4 | DNM1P34 | dynamin 1 pseudogene 34 | Other | other | |
| chr1 | *ZNF669* | 2 | 49 | 4 | ZNF669 | zinc finger protein 669 | Other | other | |
| chr16 | *SDR42E1* | 2 | 74 | 4 | SDR42E1 | short chain dehydrogenase/reductase family 42E, member 1 | Other | enzyme | |
| chr6 | *RP1-140K8.5* | 2 | 79 | 4 | | | | | |
| chrX | *GS1-600G8.3* | 2 | 41 | 4 | GS1-600G8.3 | unknown transcript | Other | other | |
| chr2 | *RAPGEF4-AS1* | 2 | 63 | 4 | RAPGEF4-AS1 | RAPGEF4 antisense RNA 1 | Other | other | |
| chr10 | *PGAM1* | 2 | 50 | 4 | PGAM1 | phosphoglycerate mutase 1 | Cytoplasm | phosphatase | |
| chr11 | *MS4A3* | 2 | 53 | 4 | MS4A3 | membrane spanning 4-domains | Plasma Membrane | other | |

| | | | | | | | | | |
|---|---|---|---|---|---|---|---|---|---|
| chr3 | *UBA3* | 2 | 35 | 4 | UBA3 | ubiquitin like modifier activating enzyme 3 | Cytoplasm | enzyme | |
| chr1 | *BARHL2* | 2 | 112 | 4 | BARHL2 | BarH like homeobox 2 | Nucleus | transcription regulator | |
| chr2 | *GRB14* | 2 | 34 | 4 | GRB14 | growth factor receptor bound protein 14 | Plasma Membrane | other | |
| chr8 | *GEM* | 2 | 52 | 4 | GEM | GTP binding protein overexpressed in skeletal muscle | Plasma Membrane | enzyme | |
| chr2 | *AGXT* | 3 | 107 | 4 | AGXT | alanine--glyoxylate and serine--pyruvate aminotransferase | Cytoplasm | enzyme | |
| chr5 | *LINC00491* | 2 | 84 | 4 | LINC00491 | long intergenic non-protein coding RNA 491 | Other | other | |
| chr12 | *ERP29* | 2 | 52 | 4 | ERP29 | endoplasmic reticulum protein 29 | Cytoplasm | transporter | |
| chr19 | *SYNE4* | 2 | 87 | 4 | SYNE4 | spectrin repeat containing nuclear envelope family member 4 | Nucleus | other | |
| chr7 | *GPR37* | 2 | 67 | 4 | GPR37 | G protein-coupled receptor 37 | Plasma Membrane | G-protein coupled recepto | |
| chr5 | *PRR7* | 2 | 112 | 4 | PRR7 | proline rich 7, synaptic | Plasma Membrane | other | |
| chr13 | *PARP1P1* | 2 | 54 | 4 | PARP1P1 | poly(ADP-ribose) polymerase 1 pseudogene 1 | Other | other | |
| chr1 | *SH3BP5L* | 3 | 157 | 4 | SH3BP5L | SH3 binding domain protein 5 like | Other | other | |
| chr16 | *IL9RP3* | 2 | 127 | 4 | IL9RP3 | interleukin 9 receptor | Other | other | |
| chr10 | *ZDHHC16* | 2 | 89 | 4 | ZDHHC16 | zinc finger DHHC-type containing | Cytoplasm | enzyme | |
| chr8 | *KB-1732A1.1* | 2 | 78 | 4 | | | | | |
| chr2 | *BUB1* | 2 | 51 | 4 | BUB1 | BUB1 mitotic checkpoint serine/threonine kinase | Nucleus | kinase | |
| chr15 | *DUOXA1* | 2 | 78 | 4 | DUOXA1 | dual oxidase maturation factor 1 | Plasma Membrane | other | |
| chr14 | *PTPN21* | 2 | 53 | 4 | PTPN21 | protein tyrosine phosphatase non-receptor type 21 | Cytoplasm | phosphatase | |
| chr12 | *PRR4* | 2 | 49 | 4 | PRR4 | proline rich 4 | Extracellular Space | other | |
| chr11 | *RP4-594L9.2* | 2 | 72 | 4 | | | | | |
| chr11 | *MPEG1* | 2 | 67 | 4 | MPEG1 | macrophage expressed 1 | Cytoplasm | other | CFI-402257 |
| chr4 | *NSUN7* | 2 | 41 | 4 | NSUN7 | NOP2/Sun RNA methyltransferase family member | Other | other | |
| chr20 | *CYP24A1* | 2 | 73 | 4 | CYP24A1 | cytochrome P450 family 24 subfamily A member 1 | Cytoplasm | enzyme | |
| chr10 | *PTCHD3P1* | 2 | 41 | 4 | PTCHD3P1 | patched domain containing 3 pseudogene 1 | Other | other | |
| chr2 | *ABI2* | 2 | 33 | 4 | ABI2 | abl interactor 2 | Cytoplasm | other | |
| chr15 | *RP11-932O9.7* | 2 | 45 | 4 | | | | | |
| chr6 | *UQCC2* | 2 | 63 | 4 | UQCC2 | ubiquinol-cytochrome c reductase complex assembly factor 2 | Cytoplasm | other | |
| chr19 | *DMWD* | 2 | 58 | 4 | DMWD | DM1 locus, WD repeat containing | Nucleus | other | |
| chr22 | *GOLGA2P4* | 2 | 87 | 4 | GOLGA2P4 | GOLGA2 pseudogene 4 | Other | other | |
| chr1 | *FBXO2* | 2 | 122 | 4 | FBXO2 | F-box protein 2 | Cytoplasm | enzyme | |
| chr7 | *AKR1B10* | 2 | 50 | 4 | AKR1B10 | aldo-keto reductase family 1 member B10 | Cytoplasm | enzyme | |
| chr10 | *SFTPA1* | 2 | 68 | 4 | SFTPA1 | surfactant protein A1 | Extracellular Space | transporter | |
| chr4 | *RP11-506N2.1* | 2 | 34 | 4 | | | | | |
| chrX | *NXF2B* | 2 | 58 | 4 | NXF2/NXF2B | nuclear RNA export factor 2 | Nucleus | transporter | |
| chr6 | *RP11-359N11.2* | 2 | 65 | 4 | | | | | |
| chr11 | *AP006621.5* | 2 | 73 | 4 | | | | | |
| chr20 | *RP1-309K20.6* | 2 | 91 | 4 | | | | | |
| chr19 | *PNKP* | 2 | 60 | 4 | PNKP | polynucleotide kinase 3'- | Nucleus | kinase | |
| chr7 | *RP11-736E3.1* | 2 | 35 | 4 | | | | | |
| chr16 | *IL17C* | 2 | 119 | 4 | IL17C | interleukin 17C | Extracellular Space | cytokine | MOR106 |
| chr13 | *MTMR6* | 2 | 42 | 4 | MTMR6 | myotubularin related protein 6 | Cytoplasm | phosphatase | |
| chr16 | *FAHD1* | 2 | 55 | 4 | FAHD1 | fumarylacetoacetate hydrolase domain containing 1 | Cytoplasm | enzyme | |
| chr6 | *SRSF3* | 2 | 68 | 4 | SRSF3 | serine and arginine rich splicing factor 3 | Nucleus | other | |
| chr6 | *RP11-257K9.8* | 2 | 32 | 4 | | | | | |
| chr16 | *ZNF319* | 2 | 84 | 4 | ZNF319 | zinc finger protein 319 | Nucleus | transcription regulator | |
| chr10 | *RP11-144G6.4* | 2 | 61 | 4 | | | | | |
| chr17 | *RP11-1094M14.1* | 2 | 49 | 4 | | | | | |
| chr6 | *RP1-140C12.2* | 2 | 42 | 4 | | | | | |
| chr16 | *RP11-134D3.2* | 2 | 101 | 4 | | | | | |
| chr3 | *RP11-314M24.1* | 2 | 64 | 4 | | | | | |
| chr11 | *APOA5* | 2 | 112 | 4 | APOA5 | apolipoprotein A5 | Extracellular Space | transporter | |
| chr2 | *DARS* | 2 | 32 | 4 | DARS1 | aspartyl-tRNA synthetase 1 | Cytoplasm | enzyme | |
| chr22 | *POM121L7* | 2 | 79 | 4 | POM121L7P | POM121 transmembrane nucleoporin like 7 pseudogene | Other | other | |
| chr13 | *SETDB2* | 2 | 32 | 4 | SETDB2 | SET domain bifurcated histone lysine methyltransferase 2 | Nucleus | enzyme | |
| chr12 | *FGF6* | 2 | 41 | 4 | FGF6 | fibroblast growth factor 6 | Extracellular Space | growth factor | |
| chr7 | *LAMTOR4* | 2 | 113 | 4 | LAMTOR4 | late endosomal/lysosomal adaptor, MAPK and MTOR | Cytoplasm | other | |
| chr19 | *DACT3-AS1* | 2 | 36 | 4 | DACT3-AS1 | DACT3 antisense RNA 1 | Other | other | |
| chr15 | *RP11-16E12.1* | 2 | 58 | 4 | | | | | |
| chr1 | *NEXN-AS1* | 2 | 50 | 4 | NEXN-AS1 | NEXN antisense RNA 1 | Other | other | |
| chr2 | *POTEJ* | 2 | 48 | 4 | POTEJ | POTE ankyrin domain family member J | Extracellular Space | other | |
| chr1 | *IVL* | 2 | 93 | 4 | IVL | involucrin | Cytoplasm | other | |
| chr11 | *TRIM21* | 2 | 105 | 4 | TRIM21 | tripartite motif containing 21 | Nucleus | enzyme | |
| chr2 | *AC016722.4* | 2 | 67 | 4 | | | | | |
| chr5 | *CTD-2587M23.1* | 2 | 222 | 4 | | | | | |
| chr4 | *C4orf32* | 2 | 32 | 4 | FAM241A | family with sequence similarity 241 member A | Cytoplasm | other | |
| chr1 | *TUBB8P6* | 2 | 103 | 4 | TUBB8P6 | tubulin beta 8 class VIII pseudogene 6 | Other | other | |
| chr19 | *HCCAT3* | 2 | 51 | 4 | ZNF350-AS1 | ZNF350 antisense RNA 1 | Other | other | |

| | | | | | | | | | |
|---|---|---|---|---|---|---|---|---|---|
| chr6 | *RP3-448I9.1* | 2 | 53 | 4 | | | | | |
| chr2 | *INPP1* | 2 | 36 | 4 | INPP1 | inositol polyphosphate-1-phosphatase | Cytoplasm | phosphatase | |
| chr16 | *PRSS30P* | 2 | 111 | 4 | PRSS30P | serine protease 30, pseudogene | Other | other | |
| chr16 | *NTHL1* | 2 | 129 | 4 | NTHL1 | nth like DNA glycosylase 1 | Nucleus | enzyme | |
| chr3 | *RP11-875H7.1* | 2 | 44 | 4 | | | | | |
| chr8 | *CDH17* | 2 | 32 | 4 | CDH17 | cadherin 17 | Plasma Membrane | transporter | |
| chr21 | *LIPI* | 2 | 44 | 4 | LIPI | lipase I | Extracellular Space | enzyme | |
| chr13 | *TPTE2* | 2 | 52 | 4 | TPTE2 | transmembrane phosphoinositide 3-phosphatase and tensin | Cytoplasm | phosphatase | |
| chr2 | *CIAO1* | 2 | 115 | 4 | CIAO1 | cytosolic iron-sulfur assembly component 1 | Nucleus | transcription regulator | |
| chr4 | *USP53* | 2 | 70 | 4 | USP53 | ubiquitin specific peptidase 53 | Extracellular Space | enzyme | |
| chr7 | *RP5-1132H15.3* | 2 | 35 | 4 | | | | | |
| chr8 | *AC022182.1* | 2 | 68 | 4 | | | | | |
| chr1 | *SLC9C2* | 2 | 54 | 4 | SLC9C2 | solute carrier family 9 member C2 (putative) | Other | other | |
| chr3 | *ST3GAL6* | 2 | 32 | 4 | ST3GAL6 | ST3 beta-galactoside alpha-2,3-sialyltransferase 6 | Cytoplasm | enzyme | |
| chrX | *SSX2B* | 2 | 72 | 4 | SSX2/SSX2B | SSX family member 2 | Nucleus | other | |
| chr11 | *CFL1* | 2 | 71 | 4 | CFL1 | cofilin 1 | Nucleus | other | |
| chr1 | *AURKAIP1* | 2 | 78 | 4 | AURKAIP1 | aurora kinase A interacting | Nucleus | enzyme | |
| chr19 | *LENG1* | 2 | 103 | 4 | LENG1 | leukocyte receptor cluster | Other | other | |
| chr12 | *RP11-561P12.5* | 2 | 43 | 4 | | | | | |
| chr19 | *ZNF582* | 2 | 33 | 4 | ZNF582 | zinc finger protein 582 | Other | other | |
| chr6 | *RWDD1* | 2 | 38 | 4 | RWDD1 | RWD domain containing 1 | Cytoplasm | other | |
| chr1 | *C1orf210* | 2 | 99 | 4 | C1orf210 | chromosome 1 open reading frame 210 | Other | other | |
| chr15 | *CSNK1G1* | 2 | 49 | 4 | CSNK1G1 | casein kinase 1 gamma 1 | Cytoplasm | kinase | |
| chr6 | *ZKSCAN8* | 2 | 69 | 4 | ZKSCAN8 | zinc finger with KRAB and SCAN domains 8 | Nucleus | transcription regulator | |
| chr22 | *RP3-438O4.4* | 2 | 53 | 4 | | | | | |
| chr2 | *UGP2* | 2 | 64 | 4 | UGP2 | UDP-glucose pyrophosphorylase | Cytoplasm | enzyme | |
| chr14 | *CTD-2128A3.2* | 2 | 36 | 4 | | | | | |
| chr1 | *OLFML3* | 2 | 85 | 4 | OLFML3 | olfactomedin like 3 | Extracellular Space | other | |
| chr16 | *RP11-542M13.2* | 2 | 71 | 4 | | | | | |
| chr17 | *ASB16* | 2 | 72 | 4 | ASB16 | ankyrin repeat and SOCS box containing 16 | Cytoplasm | other | |
| chr11 | *ESRRA* | 2 | 80 | 4 | ESRRA | estrogen related receptor alpha | Nucleus | d-dependent nuclear rec | |
| chr4 | *KLKB1* | 2 | 54 | 4 | KLKB1 | kallikrein B1 | Extracellular Space | peptidase | ecallantide, aprotinin |
| chr11 | *MRGPRF* | 2 | 75 | 4 | MRGPRF | MAS related GPR family member | Plasma Membrane | G-protein coupled recepto | |
| chr17 | *SOST* | 2 | 65 | 4 | SOST | sclerostin | Extracellular Space | other | |
| chr3 | *KRT8P12* | 2 | 136 | 4 | KRT8P12 | keratin 8 pseudogene 12 | Other | other | |
| chr20 | *RP11-347D21.3* | 2 | 55 | 4 | | | | | |
| chr15 | *RP11-762H8.3* | 2 | 101 | 4 | | | | | |
| chr13 | *GGACT* | 2 | 138 | 4 | GGACT | gamma-glutamylamine cyclotransferase | Cytoplasm | enzyme | |
| chr8 | *RB1CC1* | 2 | 40 | 4 | RB1CC1 | RB1 inducible coiled-coil 1 | Nucleus | other | |
| chr6 | *IYD* | 2 | 48 | 4 | IYD | iodotyrosine deiodinase | Plasma Membrane | enzyme | |
| chr9 | *RP11-311H10.4* | 2 | 61 | 4 | | | | | |
| chrX | *ZDHHC15* | 2 | 36 | 4 | ZDHHC15 | zinc finger DHHC-type containing | Cytoplasm | enzyme | |
| chr5 | *RGMB* | 2 | 62 | 4 | RGMB | repulsive guidance molecule BMP co-receptor b | Plasma Membrane | other | |
| chr13 | *LINC00552* | 2 | 111 | 4 | LINC00552 | long intergenic non-protein coding RNA 552 | Other | other | |
| chrX | *RP11-716A19.3* | 2 | 67 | 4 | | | | | |
| chr16 | *PRR14* | 2 | 95 | 4 | PRR14 | proline rich 14 | Other | other | |
| chr3 | *TIMMDC1* | 2 | 78 | 4 | TIMMDC1 | translocase of inner mitochondrial membrane domain containing 1 | Cytoplasm | other | |
| chr3 | *RP11-702L6.1* | 2 | 46 | 4 | | | | | |
| chr7 | *AC093627.8* | 2 | 99 | 4 | | | | | |
| chr18 | *RP11-449D8.1* | 2 | 55 | 4 | | | | | |
| chr7 | *CYP3A4* | 2 | 49 | 4 | CYP3A4 | cytochrome P450 family 3 subfamily A member 4 | Cytoplasm | enzyme | stiripentol, cobicistat, cobicistat/darunavir, cobicistat/elvitegravir/emtricitabine/tenofovir disoproxil, ketoconazole, atazanavir/cobicistat |

| | | | | | | | | | |
|---|---|---|---|---|---|---|---|---|---|
| chr6 | *TUBB2A* | 2 | 111 | 4 | TUBB2A | tubulin beta 2A class IIa | Cytoplasm | other | docetaxel/gemcitabine/vincristine, batabulin, cyclophosphamide/epirubicin/5-fluorouracil/paclitaxel, bevacizumab/paclitaxel/topotecan, colchicine/probenecid, bevacizumab/docetaxel/prednisone/thalidomide/zoledronic acid, gemcitabine/paclitaxel, simotaxel, L-asparaginase/daunorubicin/dexamethasone/vincristine, prednisone/vincristine, docetaxel/5-fluorouracil, [illegible] |
| chrX | *PIGA* | 2 | 43 | 4 | PIGA | phosphatidylinositol glycan anchor biosynthesis class A | Cytoplasm | enzyme | |
| chr1 | *RP4-613A2.1* | 2 | 60 | 4 | | | | | |
| chr19 | *RP11-1137G4.4* | 2 | 66 | 4 | | | | | |
| chr10 | *RP11-262I2.2* | 2 | 62 | 4 | | | | | |
| chr4 | *AC092661.1* | 2 | 52 | 4 | | | | | |
| chr5 | *TMEM167A* | 2 | 53 | 4 | TMEM167A | transmembrane protein 167A | Other | other | |
| chr11 | *LRRN4CL* | 2 | 51 | 4 | LRRN4CL | LRRN4 C-terminal like | Other | other | |
| chr22 | *RRP7A* | 2 | 69 | 4 | RRP7A | ribosomal RNA processing 7 homolog A | Cytoplasm | other | |
| chr2 | *AC007364.1* | 2 | 53 | 4 | | | | | |
| chr14 | *CPNE6* | 2 | 97 | 4 | CPNE6 | copine 6 | Plasma Membrane | transporter | |
| chr14 | *RP11-472N19.3* | 1 | 38 | 4 | | | | | |
| chr4 | *AC195454.1* | 2 | 38 | 4 | | | | | |
| chr19 | *CTD-3162L10.1* | 2 | 712 | 4 | | | | | |
| chr17 | *AC108004.3* | 2 | 72 | 4 | | | | | |
| chr17 | *TIMM22* | 2 | 75 | 4 | TIMM22 | translocase of inner mitochondrial membrane 22 | Cytoplasm | transporter | |
| chr22 | *AP000350.7* | 2 | 53 | 4 | | | | | |
| chr11 | *RP11-697H9.3* | 2 | 49 | 4 | | | | | |
| chr10 | *TIMM23B* | 2 | 58 | 4 | TIMM23B | translocase of inner mitochondrial membrane 23 homolog B | Cytoplasm | other | |
| chr5 | *AC138517.4* | 2 | 41 | 4 | | | | | |
| chr8 | *RHPN1-AS1* | 2 | 66 | 4 | RHPN1-AS1 | RHPN1 antisense RNA 1 (head to head) | Other | other | |
| chr7 | *AC018464.3* | 2 | 33 | 4 | | | | | |
| chr9 | *RP11-791O21.5* | 2 | 73 | 4 | | | | | |
| chr3 | *SPTSSB* | 2 | 88 | 4 | SPTSSB | serine palmitoyltransferase small subunit B | Cytoplasm | enzyme | |
| chr11 | *TM7SF2* | 2 | 122 | 4 | TM7SF2 | transmembrane 7 superfamily member 2 | Cytoplasm | enzyme | |
| chr7 | *POLD2* | 2 | 71 | 4 | POLD2 | DNA polymerase delta 2, accessory subunit | Nucleus | enzyme | |
| chr1 | *GPR3* | 2 | 113 | 4 | GPR3 | G protein-coupled receptor 3 | Plasma Membrane | G-protein coupled receptor | |
| chr4 | *RP11-440L14.1* | 2 | 60 | 4 | | | | | |
| chr1 | *RP11-480I12.7* | 2 | 66 | 4 | | | | | |
| chr11 | *DBX1* | 2 | 89 | 4 | DBX1 | developing brain homeobox 1 | Nucleus | transcription regulator | |
| chr11 | *SLC37A4* | 2 | 87 | 4 | SLC37A4 | solute carrier family 37 member 4 | Cytoplasm | transporter | |
| chr16 | *CTD-2358C21.4* | 2 | 72 | 4 | | | | | |
| chr8 | *FAM90A25P* | 2 | 140 | 4 | FAM90A25P | family with sequence similarity 90 member A25, pseudogene | Other | other | |
| chr15 | *LPCAT4* | 2 | 62 | 4 | LPCAT4 | lysophosphatidylcholine acyltransferase 4 | Cytoplasm | enzyme | |
| chr8 | *CTD-3247F14.2* | 2 | 67 | 4 | | | | | |
| chr7 | *AP4M1* | 2 | 77 | 4 | AP4M1 | adaptor related protein complex 4 subunit mu 1 | Cytoplasm | other | |
| chr6 | *PAK1IP1* | 2 | 48 | 4 | PAK1IP1 | PAK1 interacting protein 1 | Nucleus | other | |
| chr19 | *PPP1R15A* | 2 | 80 | 4 | PPP1R15A | protein phosphatase 1 regulatory subunit 15A | Cytoplasm | other | |
| chr12 | *LRCOL1* | 2 | 95 | 4 | LRCOL1 | leucine rich colipase like 1 | Other | other | |
| chr8 | *RP11-587H10.2* | 2 | 33 | 4 | | | | | |
| chr2 | *C2orf40* | 2 | 58 | 4 | ECRG4 | ECRG4 augurin precursor | Extracellular Space | other | |
| chr9 | *DMRT2* | 2 | 102 | 4 | DMRT2 | doublesex and mab-3 related transcription factor 2 | Nucleus | transcription regulator | |
| chr5 | *RP11-798K23.3* | 2 | 139 | 4 | | | | | |
| chr8 | *AZIN1* | 2 | 39 | 4 | AZIN1 | antizyme inhibitor 1 | Cytoplasm | enzyme | |
| chr2 | *KCTD18* | 2 | 32 | 4 | KCTD18 | potassium channel tetramerization domain containing | Other | other | |
| chr1 | *GS1-279B7.1* | 2 | 49 | 4 | GS1-279B7.1 | microtubule associated protein 1 light chain 3 beta pseudogene | Other | other | |
| chr22 | *GSTT2B* | 2 | 79 | 4 | GSTT2/GSTT2B | glutathione S-transferase theta 2 (gene/pseudogene) | Cytoplasm | enzyme | |
| chr2 | *UBE2F-SCLY* | 2 | 55 | 4 | UBE2F-SCLY | UBE2F-SCLY readthrough (NMD candidate) | Other | other | |
| chr1 | *PRAMEF11* | 2 | 88 | 4 | PRAMEF11 | PRAME family member 11 | Extracellular Space | other | |

| | | | | | | | | | |
|---|---|---|---|---|---|---|---|---|---|
| chr1 | *TMEM240* | 2 | 77 | 4 | TMEM240 | transmembrane protein 240 | Plasma Membrane | other | |
| chr17 | *RP11-498C9.15* | 2 | 45 | 4 | | | | | |
| chr18 | *PSMA8* | 2 | 50 | 4 | PSMA8 | proteasome subunit alpha 8 | Nucleus | peptidase | |
| chr6 | *VGLL2* | 2 | 68 | 4 | VGLL2 | vestigial like family member 2 | Nucleus | transcription regulator | |
| chr17 | *TLCD2* | 2 | 116 | 4 | TLCD2 | TLC domain containing 2 | Plasma Membrane | other | |
| chr11 | *RP11-23F23.2* | 2 | 121 | 4 | | | | | |
| chr1 | *C1orf234* | 2 | 47 | 4 | TEX46 | testis expressed 46 | Other | other | |
| chr7 | *RP11-340I6.3* | 2 | 35 | 4 | | | | | |
| chr1 | *SLC50A1* | 2 | 86 | 4 | SLC50A1 | solute carrier family 50 member 1 | Plasma Membrane | transporter | |
| chr12 | *CLEC9A* | 2 | 42 | 4 | CLEC9A | C-type lectin domain containing | Plasma Membrane | other | |
| chr1 | *PRKAB2* | 2 | 52 | 4 | PRKAB2 | protein kinase AMP-activated non-catalytic subunit beta 2 | Cytoplasm | kinase | |
| chr11 | *SMPD1* | 2 | 144 | 4 | SMPD1 | sphingomyelin phosphodiesterase 1 | Cytoplasm | enzyme | acid sphingomyelinase |
| chr19 | *ZNF730* | 2 | 62 | 4 | ZNF730 | zinc finger protein 730 | Other | other | |
| chr4 | *PARM1* | 2 | 44 | 4 | PARM1 | prostate androgen-regulated mucin-like protein 1 | Extracellular Space | other | |
| chr1 | *UBE2Q1* | 2 | 123 | 4 | UBE2Q1 | ubiquitin conjugating enzyme E2 | Other | enzyme | |
| chr2 | *KANSL1L* | 2 | 33 | 4 | KANSL1L | KAT8 regulatory NSL complex subunit 1 like | Other | other | |
| chr7 | *PMPCB* | 2 | 32 | 4 | PMPCB | peptidase, mitochondrial processing beta subunit | Cytoplasm | peptidase | |
| chr19 | *LLNLR-246C6.1* | 2 | 58 | 4 | | | | | |
| chr6 | *C6orf118* | 2 | 67 | 4 | C6orf118 | chromosome 6 open reading frame 118 | Other | other | |
| chr20 | *SNX21* | 2 | 65 | 4 | SNX21 | sorting nexin family member 21 | Cytoplasm | transporter | |
| chr2 | *AC009502.4* | 2 | 46 | 4 | | | | | |
| chr12 | *FGF23* | 2 | 57 | 4 | FGF23 | fibroblast growth factor 23 | Extracellular Space | growth factor | burosumab |
| chr4 | *FBXL5* | 2 | 42 | 4 | FBXL5 | F-box and leucine rich repeat protein 5 | Cytoplasm | enzyme | |
| chr22 | *ADSL* | 2 | 62 | 4 | ADSL | adenylosuccinate lyase | Cytoplasm | enzyme | |
| chr19 | *ZNF69* | 2 | 55 | 4 | ZNF69 | zinc finger protein 69 | Nucleus | transcription regulator | |
| chr2 | *C1D* | 2 | 47 | 4 | C1D | C1D nuclear receptor | Nucleus | transcription regulator | |
| chr5 | *PITX1* | 2 | 71 | 4 | PITX1 | paired like homeodomain 1 | Nucleus | transcription regulator | |
| chr11 | *KRT18P59* | 2 | 39 | 4 | KRT18P59 | keratin 18 pseudogene 59 | Other | other | |
| chr5 | *SMAD5* | 2 | 42 | 4 | SMAD5 | SMAD family member 5 | Nucleus | transcription regulator | |
| chr13 | *LECT1* | 2 | 45 | 4 | CNMD | chondromodulin | Extracellular Space | other | |
| chr10 | *SUV39H2* | 2 | 33 | 4 | SUV39H2 | suppressor of variegation 3-9 homolog 2 | Nucleus | transcription regulator | |
| chr4 | *SNCA* | 2 | 32 | 4 | SNCA | synuclein alpha | Cytoplasm | enzyme | |
| chr7 | *OCM* | 2 | 65 | 4 | OCM | oncomodulin | Cytoplasm | other | |
| chr1 | *RP11-84O12.4* | 2 | 77 | 4 | | | | | |
| chr1 | *PRAMEF4* | 2 | 69 | 4 | MEF4 (includes ot | PRAME family member 4 | Other | other | |
| chr9 | *RP11-449H15.2* | 2 | 34 | 4 | | | | | |
| chr1 | *RP11-466F5.6* | 2 | 49 | 4 | | | | | |
| chr1 | *RP11-552D8.1* | 2 | 56 | 4 | | | | | |
| chr17 | *FOXJ1* | 2 | 74 | 4 | FOXJ1 | forkhead box J1 | Nucleus | transcription regulator | |
| chr9 | *LCN8* | 2 | 171 | 4 | LCN8 | lipocalin 8 | Extracellular Space | transporter | |
| chr3 | *DNAJB8-AS1* | 2 | 83 | 4 | DNAJB8-AS1 | DNAJB8 antisense RNA 1 | Other | other | |
| chr7 | *RP11-728K20.1* | 2 | 46 | 4 | | | | | |
| chr17 | *RP13-516M14.2* | 2 | 156 | 4 | | | | | |
| chr3 | *RP11-485G4.2* | 2 | 49 | 4 | | | | | |
| chr8 | *RP11-758M4.1* | 2 | 42 | 4 | | | | | |
| chr22 | *POM121L8P* | 2 | 130 | 4 | POM121L8P | POM121 transmembrane nucleoporin like 8, pseudogene | Other | other | |
| chr6 | *ABT1* | 2 | 80 | 4 | ABT1 | activator of basal transcription 1 | Nucleus | transcription regulator | |
| chr1 | *FAM72A* | 2 | 140 | 4 | FAM72A | family with sequence similarity 72 member A | Cytoplasm | other | |
| chr3 | *ZBTB11-AS1* | 2 | 76 | 4 | ZBTB11-AS1 | ZBTB11 antisense RNA 1 | Other | other | |
| chr11 | *AP000439.3* | 2 | 96 | 4 | | | | | |
| chrX | *SLC7A3* | 2 | 48 | 4 | SLC7A3 | solute carrier family 7 member 3 | Plasma Membrane | transporter | |
| chrX | *BHLHB9* | 2 | 49 | 4 | BHLHB9 | basic helix-loop-helix family member b9 | Cytoplasm | other | |
| chr17 | *AC079325.6* | 2 | 54 | 4 | | | | | |
| chr12 | *PRB1* | 2 | 42 | 4 | PRB1/PRB2 | proline rich protein BstNI subfamily 1 (gene/pseudogene) | Other | other | |
| chr7 | *ASB15* | 2 | 42 | 4 | ASB15 | ankyrin repeat and SOCS box containing 15 | Nucleus | transcription regulator | |
| chr9 | *RP11-323H21.3* | 2 | 75 | 4 | | | | | |
| chrX | *MUM1L1* | 2 | 33 | 4 | PWWP3B | PWWP domain containing 3B | Cytoplasm | other | |
| chr1 | *ARPC5* | 2 | 60 | 4 | ARPC5 | actin related protein 2/3 complex subunit 5 | Cytoplasm | other | |
| chr17 | *TMIGD1* | 2 | 40 | 4 | TMIGD1 | transmembrane and immunoglobulin domain | Other | other | |
| chr15 | *SPATA8-AS1* | 2 | 114 | 4 | SPATA8-AS1 | SPATA8 antisense RNA 1 (head to head) | Other | other | |
| chr11 | *CORO1B* | 2 | 77 | 4 | CORO1B | coronin 1B | Cytoplasm | other | |
| chr19 | *RPS5* | 2 | 102 | 4 | RPS5 | ribosomal protein S5 | Cytoplasm | other | |
| chr4 | *FHDC1* | 2 | 37 | 4 | FHDC1 | FH2 domain containing 1 | Cytoplasm | other | |
| chr19 | *CTBP2P7* | 2 | 71 | 4 | CTBP2P7 | C-terminal binding protein 2 pseudogene 7 | Other | other | |
| chrX | *MAGED4* | 2 | 74 | 4 | IAGED4/MAGED4 | MAGE family member D4B | Other | other | |
| chr15 | *DNM1P49* | 2 | 133 | 4 | DNM1P49 | dynamin 1 pseudogene 49 | Other | other | |
| chr10 | *RP11-508M1.7* | 2 | 137 | 4 | | | | | |
| chr5 | *RP11-826N14.2* | 2 | 80 | 4 | | | | | |
| chr2 | *NMUR1* | 2 | 106 | 4 | NMUR1 | neuromedin U receptor 1 | Plasma Membrane | i-protein coupled recepto | |
| chr1 | *RP4-798A10.2* | 2 | 53 | 4 | | | | | |
| chr3 | *GLT8D1* | 2 | 49 | 4 | GLT8D1 | glycosyltransferase 8 domain containing 1 | Cytoplasm | enzyme | |

| | | | | | | | | | |
|---|---|---|---|---|---|---|---|---|---|
| chr5 | *RP11-798K23.1* | 2 | 78 | 4 | | | | | |
| chr17 | *TMEM220* | 2 | 31 | 4 | TMEM220 | transmembrane protein 220 | Other | other | |
| chr22 | *RP3-434P1.6* | 2 | 75 | 4 | | | | | |
| chr4 | *LINC00290* | 2 | 36 | 4 | LINC00290 | long intergenic non-protein coding RNA 290 | Other | other | |
| chr19 | *AC025278.1* | 2 | 50 | 4 | | | | | |
| chr15 | *LINC00928* | 2 | 70 | 4 | LINC00928 | long intergenic non-protein coding RNA 928 | Other | other | |
| chr2 | *SULT1C2* | 2 | 57 | 4 | SULT1C2 | sulfotransferase family 1C | Cytoplasm | enzyme | |
| chr22 | *SH3BP1* | 2 | 71 | 4 | SH3BP1 | SH3 domain binding protein 1 | Plasma Membrane | other | |
| chr19 | *SCAMP4* | 2 | 81 | 4 | SCAMP4 | secretory carrier membrane | Cytoplasm | other | |
| chr11 | *RP11-626H12.2* | 2 | 80 | 4 | | | | | |
| chr10 | *XRCC6P1* | 2 | 41 | 4 | XRCC6P1 | X-ray repair cross complementing 6 pseudogene 1 | Other | other | |
| chr1 | *RP4-758J18.2* | 2 | 102 | 4 | | | | | |
| chr9 | *ZFAND5* | 1 | 31 | 4 | ZFAND5 | zinc finger AN1-type containing 5 | Nucleus | other | |
| chr4 | *NFKB1* | 2 | 30 | 4 | NFKB1 | nuclear factor kappa B subunit 1 | Nucleus | transcription regulator | triflusal, bortezomib/dexamethasone/thalidomide, dexamethasone/thalidomide, thalidomide, rituximab/thalidomide, prednisone/thalidomi |
| chr11 | *TBCEL* | 2 | 31 | 4 | TBCEL | tubulin folding cofactor E like | Other | other | |
| chr11 | *C11orf21* | 2 | 96 | 4 | C11orf21 | chromosome 11 open reading frame 21 | Cytoplasm | other | |
| chr3 | *ACVR2B-AS1* | 2 | 60 | 4 | ACVR2B-AS1 | ACVR2B antisense RNA 1 | Other | other | |
| chr19 | *ZNF45* | 2 | 51 | 4 | ZNF45 | zinc finger protein 45 | Nucleus | transcription regulator | |
| chr8 | *LINC00977* | 2 | 33 | 4 | LINC00977 | long intergenic non-protein coding RNA 977 | Other | other | |
| chr12 | *RERGL* | 2 | 42 | 4 | RERGL | RERG like | Other | other | |
| chr2 | *AC062020.1* | 1 | 43 | 4 | | | | | |
| chr6 | *ROS1* | 2 | 33 | 4 | ROS1 | ROS proto-oncogene 1, receptor tyrosine kinase | Plasma Membrane | kinase | ROS1 inhibitor, DS-6051b, crizotinib/vemurafenib, cabozantinib/nivolumab, TQ-B3139, alkotinib, brigatinib, cabozantinib, crizotinib/erlotinib, crizotinib/pazopanib, ceritinib, crizotinib/gefitinib, lorlatinib, cabozantinib/erlotinib |
| chr5 | *RP11-215P8.4* | 2 | 53 | 4 | | | | | |
| chr19 | *GPR42* | 2 | 103 | 4 | GPR42 | G protein-coupled receptor 42 (gene/pseudogene) | Plasma Membrane | ;-protein coupled recepto | |
| chr1 | *RP11-223A3.1* | 2 | 42 | 4 | | | | | |
| chr15 | *RP11-529J17.3* | 2 | 132 | 4 | | | | | |
| chr5 | *AQPEP* | 2 | 41 | 4 | | | | | |
| chr14 | *RP11-280K24.1* | 2 | 52 | 4 | | | | | |
| chr3 | *RP11-573D15.8* | 2 | 31 | 4 | | | | | |
| chr15 | *RP11-182J1.16* | 2 | 77 | 4 | | | | | |
| chr14 | *NOVA1* | 2 | 31 | 4 | NOVA1 | NOVA alternative splicing | Nucleus | other | |
| chr18 | *CDH19* | 2 | 34 | 4 | CDH19 | cadherin 19 | Plasma Membrane | other | |
| chr12 | *KRT75* | 2 | 111 | 4 | KRT75 | keratin 75 | Cytoplasm | other | |
| chr8 | *FAM90A5P* | 2 | 97 | 4 | 90A26 (includes o | family with sequence similarity 90 member A26 | Other | other | |
| chr11 | *CDC42EP2* | 2 | 172 | 4 | CDC42EP2 | CDC42 effector protein 2 | Plasma Membrane | other | |
| chr4 | *SLC4A4* | 2 | 47 | 4 | SLC4A4 | solute carrier family 4 member 4 | Plasma Membrane | transporter | |
| chr1 | *C1orf194* | 2 | 72 | 4 | C1orf194 | chromosome 1 open reading frame 194 | Cytoplasm | other | |
| chr7 | *DLX5* | 2 | 85 | 4 | DLX5 | distal-less homeobox 5 | Nucleus | transcription regulator | |
| chr12 | *NAB2* | 2 | 107 | 4 | NAB2 | NGFI-A binding protein 2 | Nucleus | transcription regulator | |
| chr14 | *VIPAS39* | 2 | 45 | 4 | VIPAS39 | VPS33B interacting protein, apical-basolateral polarity regulator, spe-39 homolog | Plasma Membrane | other | |
| chr6 | *RP3-380B8.4* | 2 | 66 | 4 | | | | | |
| chr21 | *AL157359.3* | 2 | 32 | 4 | | | | | |
| chr18 | *RP11-627G18.4* | 2 | 37 | 4 | | | | | |
| chr16 | *PKD1P5* | 2 | 76 | 4 | PKD1P5 | polycystin 1, transient receptor potential channel interacting pseudogene 5 | Other | other | |
| chr16 | *RP11-189E14.4* | 2 | 52 | 4 | | | | | |
| chr3 | *IFT57* | 2 | 45 | 4 | IFT57 | intraflagellar transport 57 | Cytoplasm | other | |
| chr7 | *ASB4* | 2 | 75 | 4 | ASB4 | ankyrin repeat and SOCS box containing 4 | Nucleus | transcription regulator | |
| chr3 | *DTX3L* | 2 | 61 | 4 | DTX3L | deltex E3 ubiquitin ligase 3L | Cytoplasm | enzyme | |
| chr1 | *RP11-439L8.3* | 2 | 80 | 4 | | | | | |
| chr6 | *L3MBTL3* | 2 | 33 | 4 | L3MBTL3 | L3MBTL histone methyl-lysine binding protein 3 | Nucleus | other | |
| chr17 | *EFNB3* | 2 | 99 | 4 | EFNB3 | ephrin B3 | Plasma Membrane | kinase | |
| chr8 | *RP11-653B10.1* | 2 | 48 | 4 | | | | | |
| chr1 | *RP1-65J11.5* | 2 | 67 | 4 | | | | | |
| chr21 | *LINC00319* | 2 | 117 | 4 | LINC00319 | long intergenic non-protein coding RNA 319 | Other | other | |

| | | | | | | | | |
|---|---|---|---|---|---|---|---|---|
| chr1 | *UBL4B* | 2 | 75 | 4 | UBL4B | ubiquitin like 4B | Other | other |
| chr11 | *GUCY2EP* | 2 | 136 | 4 | GUCY2EP | guanylate cyclase 2E, | Other | other |
| chr7 | *FAM200A* | 2 | 96 | 4 | FAM200A | family with sequence similarity 200 member A | Other | other |
| chr12 | *KRT18P20* | 2 | 50 | 4 | KRT18P20 | keratin 18 pseudogene 20 | Other | other |
| chr9 | *FAM27A* | 2 | 215 | 4 | FAM27C | family with sequence similarity 27 member C | Other | other |
| chr2 | *WDR54* | 2 | 89 | 4 | WDR54 | WD repeat domain 54 | Other | other |
| chr20 | *AP5S1* | 2 | 84 | 4 | AP5S1 | adaptor related protein complex 5 subunit sigma 1 | Cytoplasm | other |
| chr10 | *CUBNP2* | 2 | 65 | 4 | CUBNP2 | cubilin pseudogene 2 | Other | other |
| chrX | *KIAA1210* | 2 | 30 | 4 | KIAA1210 | KIAA1210 | Cytoplasm | other |
| chr16 | *SLC22A31* | 2 | 118 | 4 | SLC22A31 | solute carrier family 22 member | Other | other |
| chr20 | *TRIB3* | 2 | 71 | 4 | TRIB3 | tribbles pseudokinase 3 | Nucleus | kinase |
| chr9 | *RP11-145E5.5* | 2 | 89 | 4 | | | | |
| chr5 | *RP11-468D11.1* | 2 | 100 | 4 | | | | |
| chr2 | *GPR35* | 2 | 164 | 4 | GPR35 | G protein-coupled receptor 35 | Plasma Membrane | G-protein coupled recepto |
| chr20 | *WFDC11* | 2 | 70 | 4 | WFDC11 | WAP four-disulfide core domain | Extracellular Space | other |
| chr1 | *SLC30A2* | 2 | 94 | 4 | SLC30A2 | solute carrier family 30 member 2 | Plasma Membrane | transporter |
| chr7 | *CBX3* | 2 | 44 | 4 | CBX3 | chromobox 3 | Nucleus | transcription regulator |
| chr5 | *DHX29* | 2 | 45 | 4 | DHX29 | DExH-box helicase 29 | Cytoplasm | enzyme |
| chr9 | *RP11-244O19.1* | 2 | 68 | 4 | | | | |
| chr7 | *RP5-899E9.1* | 2 | 41 | 4 | | | | |
| chr3 | *KRT8P18* | 2 | 68 | 4 | KRT8P18 | keratin 8 pseudogene 18 | Other | other |
| chr17 | *ALOX12P1* | 2 | 60 | 4 | ALOX12P1 | arachidonate 12-lipoxygenase pseudogene 1 | Other | other |
| chr7 | *VPS37D* | 2 | 116 | 4 | VPS37D | VPS37D subunit of ESCRT-I | Cytoplasm | other |
| chr12 | *RP11-809C9.2* | 2 | 52 | 4 | | | | |
| chr19 | *C19orf67* | 2 | 59 | 4 | C19orf67 | chromosome 19 open reading frame 67 | Other | other |
| chr1 | *CHD1L* | 2 | 43 | 4 | CHD1L | chromodomain helicase DNA binding protein 1 like | Nucleus | enzyme |
| chr1 | *CYP2J2* | 2 | 58 | 4 | CYP2J2 | cytochrome P450 family 2 subfamily J member 2 | Cytoplasm | enzyme |
| chr19 | *SYT5* | 2 | 122 | 4 | SYT5 | synaptotagmin 5 | Cytoplasm | transporter |
| chr16 | *RNF166* | 2 | 109 | 4 | RNF166 | ring finger protein 166 | Other | other |
| chr2 | *CTDSP1* | 2 | 87 | 4 | CTDSP1 | CTD small phosphatase 1 | Nucleus | phosphatase |
| chr1 | *PEX11B* | 2 | 58 | 4 | PEX11B | peroxisomal biogenesis factor 11 beta | Cytoplasm | other |
| chr2 | *SULT1C2P1* | 2 | 66 | 4 | SULT1C2P1 | sulfotransferase family 1C member 2 pseudogene 1 | Other | other |
| chr1 | *C1orf85* | 2 | 96 | 4 | GLMP | glycosylated lysosomal membrane protein | Cytoplasm | transcription regulator |
| chr17 | *RP11-605F20.1* | 2 | 38 | 4 | | | | |
| chr15 | *DNM1P50* | 2 | 80 | 4 | DNM1P50 | dynamin 1 pseudogene 50 | Other | other |
| chr16 | *LINC00311* | 2 | 137 | 4 | LINC00311 | long intergenic non-protein coding RNA 311 | Other | other |
| chr2 | *MPHOSPH10* | 2 | 38 | 4 | MPHOSPH10 | M-phase phosphoprotein 10 | Nucleus | other |
| chr17 | *ZNF207* | 2 | 50 | 4 | ZNF207 | zinc finger protein 207 | Nucleus | transcription regulator |
| chr19 | *LINC00663* | 2 | 51 | 4 | LINC00663 | long intergenic non-protein coding RNA 663 | Other | other |
| chr14 | *RP11-566J3.2* | 2 | 99 | 4 | | | | |
| chr8 | *RP11-909N17.2* | 2 | 75 | 4 | | | | |
| chr12 | *RP11-551L14.7* | 2 | 128 | 4 | | | | |
| chr14 | *AL157871.2* | 2 | 50 | 4 | | | | |
| chr22 | *POM121L1P* | 2 | 86 | 4 | POM121L1P | POM121 transmembrane nucleoporin like 1, pseudogene | Other | other |
| chr11 | *RP11-867G2.2* | 2 | 127 | 4 | | | | |
| chr5 | *C5orf49* | 2 | 139 | 4 | C5orf49 | chromosome 5 open reading frame 49 | Other | other |
| chr8 | *RP11-1149O23.2* | 1 | 34 | 4 | | | | |
| chr10 | *CYP2C8* | 2 | 48 | 4 | CYP2C8 | cytochrome P450 family 2 subfamily C member 8 | Cytoplasm | enzyme |
| chr17 | *CDRT1* | 2 | 43 | 4 | CDRT1 | CMT1A duplicated region | Other | other |
| chr2 | *AC093388.3* | 2 | 40 | 4 | | | | |
| chr2 | *GTF3C3* | 2 | 44 | 4 | GTF3C3 | general transcription factor IIIC subunit 3 | Nucleus | transcription regulator |
| chr2 | *AC074091.13* | 2 | 47 | 4 | | | | |
| chr20 | *ANKRD60* | 2 | 82 | 4 | ANKRD60 | ankyrin repeat domain 60 | Other | other |
| chr1 | *NR1I3* | 2 | 62 | 4 | NR1I3 | nuclear receptor subfamily 1 group I member 3 | Nucleus | d-dependent nuclear rec |
| chr1 | *CCDC17* | 2 | 97 | 4 | CCDC17 | coiled-coil domain containing 17 | Other | other |
| chr19 | *CTD-2587H24.1* | 2 | 54 | 4 | | | | |
| chr12 | *SLC9A7P1* | 2 | 77 | 4 | SLC9A7P1 | solute carrier family 9 member 7 pseudogene 1 | Other | other |
| chr5 | *RP11-259O2.2* | 2 | 172 | 4 | | | | |
| chr5 | *CEP120* | 2 | 32 | 4 | CEP120 | centrosomal protein 120 | Cytoplasm | other |
| chr1 | *RP11-12M5.3* | 2 | 123 | 4 | | | | |
| chr7 | *TMED4* | 2 | 106 | 4 | TMED4 | transmembrane p24 trafficking protein 4 | Cytoplasm | transporter |
| chr1 | *U3* | 2 | 44 | 4 | | | | |
| chrX | *LRCH2* | 2 | 41 | 4 | LRCH2 | leucine rich repeats and calponin homology domain containing 2 | Other | other |
| chr4 | *RAB33B* | 2 | 31 | 4 | RAB33B | RAB33B, member RAS oncogene family | Cytoplasm | enzyme |
| chr12 | *ENO2* | 2 | 57 | 4 | ENO2 | enolase 2 | Cytoplasm | enzyme |
| chr2 | *RP11-734K21.2* | 2 | 55 | 4 | | | | |
| chr15 | *NPAP1* | 2 | 52 | 4 | NPAP1 | nuclear pore associated protein 1 | Cytoplasm | other |

| | | | | | | | | | |
|---|---|---|---|---|---|---|---|---|---|
| chr17 | *POLRMTP1* | 2 | 60 | 4 | POLRMTP1 | RNA polymerase mitochondrial pseudogene 1 | Other | other | |
| chr13 | *LINC00391* | 2 | 111 | 4 | LINC00391 | long intergenic non-protein coding RNA 391 | Other | other | |
| chr11 | *ZNF408* | 2 | 106 | 4 | ZNF408 | zinc finger protein 408 | Nucleus | transcription regulator | |
| chr10 | *CLRN3* | 2 | 69 | 4 | CLRN3 | clarin 3 | Cytoplasm | other | |
| chr17 | *SLC25A39* | 2 | 94 | 4 | SLC25A39 | solute carrier family 25 member | Cytoplasm | other | |
| chr17 | *SPDYE4* | 2 | 55 | 4 | SPDYE4 | speedy/RINGO cell cycle regulator family member E4 | Other | other | |
| chr14 | *MTA1* | 2 | 89 | 4 | MTA1 | metastasis associated 1 | Nucleus | transcription regulator | |
| chr4 | *TAPT1-AS1* | 2 | 30 | 4 | TAPT1-AS1 | TAPT1 antisense RNA 1 (head to head) | Other | other | |
| chr2 | *PREB* | 2 | 57 | 4 | PREB | prolactin regulatory element | Nucleus | transcription regulator | |
| chr6 | *RP11-274H24.1* | 2 | 41 | 4 | | | | | |
| chr16 | *ARHGDIG* | 2 | 75 | 4 | ARHGDIG | Rho GDP dissociation inhibitor gamma | Cytoplasm | other | |
| chr16 | *METRN* | 2 | 131 | 4 | METRN | meteorin, glial cell differentiation regulator | Extracellular Space | other | |
| chr21 | *ZBTB21* | 2 | 34 | 4 | ZBTB21 | zinc finger and BTB domain containing 21 | Nucleus | transcription regulator | |
| chr16 | *CLDN6* | 2 | 78 | 4 | CLDN6 | claudin 6 | Plasma Membrane | other | ASP1650 |
| chr5 | *CTD-2031P19.3* | 2 | 35 | 4 | | | | | |
| chr10 | *RP11-78A18.2* | 2 | 31 | 4 | | | | | |
| chr14 | *IGHG1* | 2 | 84 | 4 | IGHG1 | immunoglobulin heavy constant gamma 1 (G1m marker) | Extracellular Space | other | |
| chr20 | *LINC00851* | 2 | 51 | 4 | LINC00851 | long intergenic non-protein coding RNA 851 | Other | other | |
| chr8 | *FAM90A7P* | 1 | 102 | 4 | 90A26 (includes ot | family with sequence similarity 90 member A26 | Other | other | |
| chr5 | *ASS1P9* | 2 | 46 | 4 | ASS1P9 | argininosuccinate synthetase 1 pseudogene 9 | Other | other | |
| chr19 | *FUT6* | 2 | 59 | 4 | FUT6 | fucosyltransferase 6 | Cytoplasm | enzyme | |
| chr10 | *OAT* | 2 | 49 | 4 | OAT | ornithine aminotransferase | Cytoplasm | enzyme | |
| chr17 | *GGT6* | 2 | 124 | 4 | GGT6 | gamma-glutamyltransferase 6 | Plasma Membrane | peptidase | |
| chr3 | *AC090505.1* | 2 | 43 | 4 | | | | | |
| chr14 | *TTC9* | 2 | 41 | 4 | TTC9 | tetratricopeptide repeat domain 9 | Extracellular Space | other | |
| chr1 | *RP11-284F21.9* | 2 | 67 | 4 | | | | | |
| chr12 | *RP11-357K6.2* | 2 | 47 | 4 | | | | | |
| chr3 | *NPRL2* | 2 | 88 | 4 | NPRL2 | NPR2 like, GATOR1 complex | Cytoplasm | kinase | |
| chr17 | *CSH2* | 2 | 94 | 4 | CSH1/CSH2 | chorionic somatomammotropin hormone 1 | Extracellular Space | other | |
| chr2 | *AC016735.1* | 2 | 77 | 4 | | | | | |
| chr12 | *IGF1* | 2 | 39 | 4 | IGF1 | insulin like growth factor 1 | Extracellular Space | growth factor | BI 836845, MEDI- |
| chr22 | *PATZ1* | 2 | 46 | 4 | PATZ1 | POZ/BTB and AT hook containing zinc finger 1 | Nucleus | transcription regulator | |
| chr4 | *USP17L20* | 2 | 111 | 4 | 17L18 (includes ot | ubiquitin specific peptidase 17 like family member 5 | Nucleus | peptidase | |
| chr1 | *SERBP1* | 2 | 80 | 4 | SERBP1 | SERPINE1 mRNA binding protein | Cytoplasm | other | |
| chr4 | *RP11-701P16.4* | 2 | 46 | 4 | | | | | |
| chr22 | *AC008132.1* | 1 | 45 | 4 | | | | | |
| chr2 | *AC012507.3* | 2 | 55 | 4 | | | | | |
| chr12 | *UNC119B* | 2 | 42 | 4 | UNC119B | unc-119 lipid binding chaperone | Cytoplasm | other | |
| chr3 | *RPL24* | 2 | 53 | 4 | RPL24 | ribosomal protein L24 | Cytoplasm | other | |
| chr13 | *ESRRAP2* | 2 | 30 | 4 | | | | | |
| chr12 | *ZNF705A* | 2 | 44 | 4 | ZNF705A | zinc finger protein 705A | Nucleus | transcription regulator | |
| chr21 | *AP000475.2* | 2 | 34 | 4 | | | | | |
| chr15 | *PRKXP1* | 2 | 199 | 4 | PRKXP1 | PRKX pseudogene 1 | Other | other | |
| chr16 | *MT1H* | 2 | 98 | 4 | MT1H | metallothionein 1H | Other | other | |
| chr4 | *CLGN* | 2 | 53 | 4 | CLGN | calmegin | Cytoplasm | peptidase | |
| chr3 | *KRT18P17* | 2 | 81 | 4 | KRT18P17 | keratin 18 pseudogene 17 | Other | other | |
| chr3 | *CISH* | 2 | 77 | 4 | CISH | cytokine inducible SH2 containing protein | Cytoplasm | other | |
| chr15 | *CYP1A1* | 2 | 63 | 4 | CYP1A1 | cytochrome P450 family 1 subfamily A member 1 | Cytoplasm | enzyme | |
| chr9 | *TXNDC8* | 2 | 31 | 4 | TXNDC8 | thioredoxin domain containing 8 | Cytoplasm | transporter | |
| chr3 | *KRT8P25* | 2 | 74 | 4 | KRT8P25 | keratin 8 pseudogene 25 | Other | other | |
| chr15 | *POTEB* | 2 | 71 | 4 | POTEB/POTEB2 | POTE ankyrin domain family member B2 | Other | other | |
| chr14 | *STXBP6* | 2 | 29 | 4 | STXBP6 | syntaxin binding protein 6 | Cytoplasm | other | |
| chr17 | *RP11-141J13.3* | 2 | 49 | 4 | | | | | |
| chr5 | *CTD-2050E21.1* | 2 | 40 | 4 | | | | | |
| chr1 | *RP11-690C23.2* | 2 | 64 | 4 | | | | | |
| chr2 | *RNPEPL1* | 2 | 87 | 4 | RNPEPL1 | arginyl aminopeptidase like 1 | Other | peptidase | |
| chr15 | *EIF3J-AS1* | 2 | 44 | 4 | EIF3J-DT | EIF3J divergent transcript | Other | other | |
| chr9 | *RP11-87H9.2* | 2 | 227 | 4 | | | | | |
| chr1 | *RPL5* | 2 | 59 | 4 | RPL5 | ribosomal protein L5 | Cytoplasm | other | |
| chr4 | *RUFY3* | 2 | 34 | 4 | RUFY3 | RUN and FYVE domain | Plasma Membrane | other | |
| chr11 | *CTSW* | 2 | 99 | 4 | CTSW | cathepsin W | Cytoplasm | peptidase | |
| chr13 | *RP11-248G5.8* | 2 | 38 | 4 | | | | | |
| chr12 | *RP11-143E21.6* | 2 | 52 | 4 | | | | | |
| chr3 | *LSM3* | 2 | 50 | 4 | LSM3 | LSM3 homolog, U6 small nuclear RNA and mRNA degradation associated | Nucleus | other | |
| chr1 | *FCRL6* | 2 | 82 | 4 | FCRL6 | Fc receptor like 6 | Plasma Membrane | other | |
| chr8 | *RP11-150O12.5* | 2 | 49 | 4 | | | | | |
| chrX | *GATA1* | 2 | 89 | 4 | GATA1 | GATA binding protein 1 | Nucleus | transcription regulator | |
| chr9 | *POLR1E* | 2 | 38 | 4 | POLR1E | RNA polymerase I subunit E | Nucleus | enzyme | |
| chr2 | *AC092170.1* | 1 | 31 | 4 | | | | | |

| | | | | | | | | | |
|---|---|---|---|---|---|---|---|---|---|
| chr1 | *PTGS2* | 2 | 49 | 4 | PTGS2 | prostaglandin-endoperoxide synthase 2 | Cytoplasm | enzyme | aspirin/rivaroxaban/ticlopidine, phenacetin, balsalazide, sorafenib/sulindac/sunitinib, acetaminophen/hydrocodone, aspirin/dabigatran etexilate, acetaminophen/caffeine/dihydrocodeine, diclofenac/omeprazole, acetaminophen/butalbital/caffeine/codeine, diphenhydramine/ibuprofen, chlorpheniramine/ibuprofen/pseudoephedrine, aspirin/hydrocodone, [illegible] |
| chr2 | *ZEB2* | 2 | 31 | 4 | ZEB2 | zinc finger E-box binding | Nucleus | transcription regulator | |
| chr17 | *PNPO* | 2 | 63 | 4 | PNPO | pyridoxamine 5'-phosphate | Cytoplasm | enzyme | |
| chr7 | *TRPV6* | 2 | 71 | 4 | TRPV6 | transient receptor potential cation channel subfamily V member 6 | Plasma Membrane | ion channel | SOR-C13 |
| chr17 | *CTD-2561B21.10* | 2 | 92 | 4 | | | | | |
| chr1 | *HLX* | 2 | 123 | 4 | HLX | H2.0 like homeobox | Nucleus | transcription regulator | |
| chr6 | *CENPQ* | 2 | 67 | 4 | CENPQ | centromere protein Q | Nucleus | other | |
| chr15 | *RP11-247C2.1* | 2 | 101 | 4 | | | | | |
| chr1 | *HSP90B3P* | 2 | 43 | 4 | HSP90B3P | heat shock protein 90 beta family member 3, pseudogene | Other | other | |
| chr4 | *PGM2* | 2 | 29 | 4 | PGM2 | phosphoglucomutase 2 | Cytoplasm | enzyme | |
| chr2 | *RPL37A* | 2 | 30 | 4 | RPL37A | ribosomal protein L37a | Cytoplasm | other | |
| chr3 | *GTF2E1* | 2 | 49 | 4 | GTF2E1 | general transcription factor IIE subunit 1 | Nucleus | transcription regulator | |
| chr14 | *RP11-736N17.8* | 2 | 79 | 4 | | | | | |
| chr5 | *CTB-161M19.4* | 2 | 37 | 4 | | | | | |
| chr19 | *CTB-147C22.8* | 2 | 52 | 4 | | | | | |
| chr17 | *C17orf77* | 2 | 42 | 4 | C17orf77 | chromosome 17 open reading frame 77 | Other | other | |
| chrX | *PIN4* | 2 | 37 | 4 | PIN4 | peptidylprolyl cis/trans isomerase, NIMA-interacting 4 | Nucleus | enzyme | |
| chr19 | *CYP2A7P1* | 2 | 115 | 4 | CYP2A7P1 | cytochrome P450 family 2 subfamily A member 7 | Other | other | |
| chr16 | *TRAPPC2L* | 2 | 64 | 4 | TRAPPC2L | trafficking protein particle complex 2 like | Cytoplasm | other | |
| chr6 | *MYLIP* | 2 | 47 | 4 | MYLIP | myosin regulatory light chain interacting protein | Cytoplasm | enzyme | |
| chr19 | *SBK2* | 2 | 81 | 4 | SBK2 | SH3 domain binding kinase family member 2 | Other | other | |
| chr2 | *IL1A* | 2 | 45 | 4 | IL1A | interleukin 1 alpha | Extracellular Space | cytokine | MABp1 |
| chr12 | *RP11-253E3.3* | 2 | 52 | 4 | | | | | |
| chr10 | *C10orf126* | 2 | 32 | 4 | C10orf126 | chromosome 10 open reading frame 126 | Other | other | |
| chr3 | *GNAT1* | 2 | 109 | 4 | GNAT1 | G protein subunit alpha | Plasma Membrane | enzyme | |
| chr8 | *CNGB3* | 2 | 47 | 4 | CNGB3 | cyclic nucleotide gated channel subunit beta 3 | Plasma Membrane | ion channel | |
| chr7 | *TAF6* | 2 | 78 | 4 | TAF6 | TATA-box binding protein associated factor 6 | Nucleus | transcription regulator | |
| chr12 | *RHEBL1* | 2 | 107 | 4 | RHEBL1 | RHEB like 1 | Cytoplasm | enzyme | |
| chr3 | *RPL29* | 2 | 88 | 4 | RPL29 | ribosomal protein L29 | Cytoplasm | other | |
| chr11 | *TSPAN4* | 2 | 88 | 4 | TSPAN4 | tetraspanin 4 | Plasma Membrane | other | |
| chr14 | *PPP2R3C* | 2 | 39 | 4 | PPP2R3C | protein phosphatase 2 regulatory subunit B''gamma | Cytoplasm | other | |
| chr12 | *KRT1* | 2 | 83 | 4 | KRT1 | keratin 1 | Cytoplasm | other | |
| chr3 | *AC090945.1* | 2 | 65 | 4 | | | | | |
| chr14 | *RP11-488C13.4* | 2 | 73 | 4 | | | | | |
| chr11 | *RIN1* | 2 | 179 | 4 | RIN1 | Ras and Rab interactor 1 | Cytoplasm | other | |
| chr17 | *CTC-296K1.3* | 2 | 40 | 4 | | | | | |
| chr17 | *AC131056.5* | 2 | 70 | 4 | | | | | |
| chr17 | *ASGR1* | 2 | 59 | 4 | ASGR1 | asialoglycoprotein receptor 1 | Plasma Membrane | transmembrane receptor | |
| chr17 | *LRRC3C* | 2 | 105 | 4 | LRRC3C | leucine rich repeat containing 3C | Other | other | |
| chr19 | *CTD-3020H12.3* | 2 | 34 | 4 | | | | | |
| chr7 | *PSMC2* | 2 | 38 | 4 | PSMC2 | proteasome 26S subunit, ATPase | Nucleus | peptidase | |
| chr8 | *NUDT18* | 2 | 74 | 4 | NUDT18 | nudix hydrolase 18 | Cytoplasm | enzyme | |
| chr2 | *AC093609.1* | 2 | 91 | 4 | | | | | |
| chr17 | *FLJ45079* | 2 | 76 | 4 | | | | | |
| chr18 | *STARD6* | 2 | 29 | 4 | STARD6 | StAR related lipid transfer domain containing 6 | Other | transporter | |
| chr6 | *RP1-111C20.4* | 2 | 39 | 4 | | | | | |
| chr3 | *IGSF10* | 2 | 44 | 4 | IGSF10 | immunoglobulin superfamily member 10 | Extracellular Space | other | |
| chr7 | *GUSBP12* | 2 | 67 | 4 | GUSBP12 | GUSB pseudogene 12 | Other | other | |
| chr18 | *RP11-861E21.2* | 2 | 44 | 4 | | | | | |
| chr7 | *TMEM209* | 2 | 32 | 4 | TMEM209 | transmembrane protein 209 | Other | other | |
| chr16 | *BMS1P8* | 2 | 88 | 4 | BMS1P8 | BMS1 pseudogene 8 | Other | other | |

| | | | | | | | | | |
|---|---|---|---|---|---|---|---|---|---|
| chr19 | *RP11-15A1.2* | 2 | 33 | 4 | | | | | |
| chr12 | *AQP5* | 2 | 140 | 4 | AQP5 | aquaporin 5 | Plasma Membrane | transporter | |
| chr2 | *AC092687.3* | 2 | 37 | 4 | | | | | |
| chr4 | *UFSP2* | 2 | 34 | 4 | UFSP2 | UFM1 specific peptidase 2 | Other | enzyme | |
| chr6 | *TFAP2B* | 2 | 53 | 4 | TFAP2B | transcription factor AP-2 beta | Nucleus | transcription regulator | |
| chr14 | *IGHGP* | 2 | 102 | 4 | IGHGP | immunoglobulin heavy constant gamma P (non-functional) | Other | other | |
| chr6 | *LINC00336* | 2 | 105 | 4 | LINC00336 | long intergenic non-protein coding RNA 336 | Other | other | |
| chr9 | *BAG1* | 2 | 52 | 4 | BAG1 | BCL2 associated athanogene 1 | Cytoplasm | other | |
| chr1 | *S100A5* | 2 | 98 | 4 | S100A5 | S100 calcium binding protein A5 | Nucleus | other | |
| chr5 | *CTB-79E8.2* | 2 | 181 | 4 | | | | | |
| chr20 | *SRMS* | 2 | 120 | 4 | SRMS | src-related kinase lacking C-terminal regulatory tyrosine and N-terminal myristylation sites | Cytoplasm | kinase | cetuximab/vemurafenib, irinotecan/panitumumab/vemurafenib, cobimetinib/vemurafenib, trametinib/vemurafenib, dabrafenib/trametinib/vemurafenib, vemurafenib, crizotinib/vemurafenib, ipilimumab/vemurafe |
| chr6 | *GSTA4* | 2 | 33 | 4 | GSTA4 | glutathione S-transferase alpha 4 | Cytoplasm | enzyme | |
| chr16 | *RP11-7O14.1* | 2 | 48 | 4 | | | | | |
| chr17 | *SLC2A4* | 2 | 57 | 4 | SLC2A4 | solute carrier family 2 member 4 | Plasma Membrane | transporter | |
| chr2 | *GLB1L* | 2 | 54 | 4 | GLB1L | galactosidase beta 1 like | Extracellular Space | other | |
| chr4 | *SMARCAD1* | 2 | 35 | 4 | SMARCAD1 | SWI/SNF-related, matrix-associated actin-dependent regulator of chromatin, subfamily a, containing DEAD/H box 1 | Nucleus | enzyme | |
| chr10 | *VAX1* | 2 | 49 | 4 | VAX1 | ventral anterior homeobox 1 | Nucleus | transcription regulator | |
| chr11 | *ASCL3* | 2 | 57 | 4 | ASCL3 | achaete-scute family bHLH transcription factor 3 | Nucleus | transcription regulator | |
| chr2 | *AC079779.6* | 2 | 34 | 4 | | | | | |
| chr19 | *SIGLEC10* | 2 | 40 | 4 | SIGLEC10 | sialic acid binding Ig like lectin 10 | Plasma Membrane | other | |
| chr9 | *RP11-536O18.2* | 2 | 43 | 4 | | | | | |
| chr2 | *DTYMK* | 2 | 62 | 4 | DTYMK | deoxythymidylate kinase | Cytoplasm | kinase | |
| chr12 | *ACTR6* | 2 | 46 | 4 | ACTR6 | actin related protein 6 | Nucleus | transporter | |
| chr5 | *RP11-473L15.3* | 2 | 95 | 4 | | | | | |
| chr1 | *HTR1D* | 2 | 66 | 4 | HTR1D | 5-hydroxytryptamine receptor 1D | Plasma Membrane | ;-protein coupled recepto | ziprasidone, vortioxetine, ergotamine, eletriptan, frovatriptan, naratriptan, almotriptan, zolmitriptan, dihydroergotamine, fenfluramine, naproxen/sumatripta |
| chr6 | *AIM1* | 2 | 59 | 4 | CRYBG1 | crystallin beta-gamma domain containing 1 | Extracellular Space | other | |
| chr12 | *RP11-664H17.1* | 2 | 28 | 4 | | | | | |
| chr16 | *RP11-77H9.2* | 2 | 86 | 4 | | | | | |
| chr19 | *LILRP1* | 2 | 83 | 4 | LILRP1 | leukocyte immunoglobulin-like receptor pseudogene 1 | Other | other | |
| chr2 | *DLX1* | 2 | 89 | 4 | DLX1 | distal-less homeobox 1 | Nucleus | transcription regulator | |
| chr17 | *RPL12P38* | 2 | 38 | 4 | RPL12P38 | ribosomal protein L12 pseudogene 38 | Other | other | |
| chr1 | *TOE1* | 2 | 53 | 4 | TOE1 | target of EGR1, exonuclease | Nucleus | enzyme | |
| chr2 | *ITGA6* | 2 | 29 | 4 | ITGA6 | integrin subunit alpha 6 | Plasma Membrane | transmembrane receptor | |
| chr19 | *GSK3A* | 2 | 72 | 4 | GSK3A | glycogen synthase kinase 3 alpha | Nucleus | kinase | |
| chr1 | *NHLH1* | 2 | 102 | 4 | NHLH1 | nescient helix-loop-helix 1 | Nucleus | transcription regulator | |
| chr11 | *EFEMP2* | 2 | 125 | 4 | EFEMP2 | EGF containing fibulin extracellular matrix protein 2 | Extracellular Space | other | |
| chr1 | *KIAA1107* | 2 | 35 | 4 | KIAA1107 | KIAA1107 | Plasma Membrane | other | |
| chr1 | *PIGM* | 2 | 76 | 4 | PIGM | phosphatidylinositol glycan anchor biosynthesis class M | Cytoplasm | enzyme | |
| chr16 | *C16orf87* | 1 | 29 | 4 | C16orf87 | chromosome 16 open reading frame 87 | Other | other | |
| chr19 | *BST2* | 2 | 63 | 4 | BST2 | bone marrow stromal cell antigen | Plasma Membrane | other | |
| chr7 | *CICP22* | 2 | 116 | 4 | CICP22 | capicua transcriptional repressor pseudogene 22 | Other | other | |
| chr6 | *IBTK* | 2 | 49 | 4 | IBTK | inhibitor of Bruton tyrosine kinase | Cytoplasm | other | |
| chr22 | *SLC5A1* | 2 | 36 | 4 | SLC5A1 | solute carrier family 5 member 1 | Plasma Membrane | transporter | |
| chr6 | *RP1-90K10.4* | 2 | 48 | 4 | | | | | |
| chr7 | *KRT8P20* | 2 | 99 | 4 | KRT8P20 | keratin 8 pseudogene 20 | Other | other | |
| chr15 | *PSMA4* | 2 | 48 | 4 | PSMA4 | proteasome subunit alpha 4 | Cytoplasm | peptidase | |
| chr3 | *GATA2* | 2 | 141 | 4 | GATA2 | GATA binding protein 2 | Nucleus | transcription regulator | |
| chr16 | *DNAJA2* | 2 | 37 | 4 | DNAJA2 | DnaJ heat shock protein family (Hsp40) member A2 | Nucleus | enzyme | |
| chr13 | *PCDH20* | 2 | 47 | 4 | PCDH20 | protocadherin 20 | Other | other | |
| chr12 | *SLC5A8* | 2 | 47 | 4 | SLC5A8 | solute carrier family 5 member 8 | Plasma Membrane | transporter | |
| chr10 | *A1CF* | 2 | 54 | 4 | A1CF | APOBEC1 complementation | Nucleus | other | |

| | | | | | | | | | |
|---|---|---|---|---|---|---|---|---|---|
| chr11 | *HNRNPUL2-BSCL* | 2 | 42 | 4 | HNRNPUL2-BSCL | HNRNPUL2-BSCL2 readthrough (NMD candidate) | Other | other | |
| chr5 | *SRA1* | 2 | 75 | 4 | SRA1 | steroid receptor RNA activator 1 | Nucleus | transcription regulator | |
| chrX | *RP11-197B12.1* | 2 | 43 | 4 | | | | | |
| chr7 | *CASP2* | 2 | 41 | 4 | CASP2 | caspase 2 | Cytoplasm | peptidase | |
| chr14 | *CMTM5* | 2 | 65 | 4 | CMTM5 | CKLF like MARVEL transmembrane domain | Other | cytokine | |
| chr2 | *MTERFD2* | 2 | 49 | 4 | MTERF4 | mitochondrial transcription termination factor 4 | Cytoplasm | other | |
| chrX | *RP11-298A8.2* | 2 | 32 | 4 | | | | | |
| chr11 | *HYLS1* | 2 | 38 | 4 | HYLS1 | HYLS1 centriolar and ciliogenesis associated | Nucleus | other | |
| chr7 | *PAXIP1-AS2* | 2 | 47 | 4 | PAXIP1-AS2 | PAXIP1 antisense RNA 2 | Other | other | |
| chr22 | *MCHR1* | 2 | 72 | 4 | MCHR1 | melanin concentrating hormone receptor 1 | Plasma Membrane | -protein coupled recepto | SB 568849 |
| chr18 | *RP11-108P20.2* | 2 | 44 | 4 | | | | | |
| chrX | *RP11-524P6.1* | 2 | 31 | 4 | | | | | |
| chr10 | *RP11-135D11.2* | 2 | 126 | 4 | | | | | |
| chr16 | *RP11-170L3.7* | 2 | 50 | 4 | | | | | |
| chr1 | *RP11-307B6.3* | 2 | 75 | 4 | | | | | |
| chr1 | *RP11-12M5.1* | 2 | 54 | 4 | | | | | |
| chr12 | *KRT126P* | 2 | 99 | 4 | KRT126P | keratin 126 pseudogene | Other | other | |
| chr1 | *RP4-705F19.1* | 2 | 58 | 4 | | | | | |
| chr8 | *FAM90A15P* | 2 | 116 | 4 | 90A26 (includes ot | family with sequence similarity 90 member A26 | Other | other | |
| chr5 | *RP11-136K7.2* | 2 | 47 | 4 | | | | | |
| chr1 | *C1QC* | 2 | 89 | 4 | C1QC | complement C1q C chain | Extracellular Space | other | |
| chr16 | *FAM108A8P* | 2 | 105 | 4 | ABHD17AP7 | abhydrolase domain containing 17A pseudogene 7 | Other | other | |
| chr1 | *KLHDC7A* | 2 | 63 | 4 | KLHDC7A | kelch domain containing 7A | Other | other | |
| chr7 | *RP5-1185I7.1* | 2 | 60 | 4 | | | | | |
| chr6 | *SLC22A7* | 2 | 65 | 4 | SLC22A7 | solute carrier family 22 member 7 | Plasma Membrane | transporter | |
| chr1 | *CLCA4* | 2 | 32 | 4 | CLCA4 | chloride channel accessory 4 | Plasma Membrane | ion channel | |
| chr10 | *SEPT7P9* | 2 | 59 | 4 | SEPTIN7P9 | septin 7 pseudogene 9 | Other | other | |
| chr4 | *RP11-367J11.2* | 2 | 63 | 4 | | | | | |
| chr17 | *CTD-2319I12.5* | 2 | 115 | 4 | | | | | |
| chr16 | *AC009158.1* | 2 | 34 | 4 | | | | | |
| chr1 | *DNTTIP2* | 2 | 78 | 4 | DNTTIP2 | deoxynucleotidyltransferase terminal interacting protein 2 | Nucleus | other | |
| chr19 | *FKBP8* | 2 | 145 | 4 | FKBP8 | FKBP prolyl isomerase 8 | Cytoplasm | other | |
| chr19 | *CTD-3193O13.9* | 2 | 68 | 4 | | | | | |
| chr3 | *MFSD1* | 2 | 48 | 4 | MFSD1 | major facilitator superfamily domain containing 1 | Other | transporter | |
| chr11 | *HPX* | 2 | 82 | 4 | HPX | hemopexin | Extracellular Space | transporter | |
| chr11 | *RP11-317J19.1* | 2 | 76 | 4 | | | | | |
| chr22 | *EIF3D* | 2 | 56 | 4 | EIF3D | eukaryotic translation initiation factor 3 subunit D | Cytoplasm | other | |
| chr18 | *RP11-838N2.3* | 2 | 73 | 4 | | | | | |
| chr17 | *RP11-893F2.9* | 2 | 57 | 4 | | | | | |
| chr1 | *LINC00337* | 2 | 123 | 4 | LINC00337 | long intergenic non-protein coding RNA 337 | Other | other | |
| chr1 | *RNASEL* | 2 | 50 | 4 | RNASEL | ribonuclease L | Cytoplasm | enzyme | |
| chr19 | *AC092295.4* | 2 | 36 | 4 | | | | | |
| chr17 | *DERL2* | 2 | 34 | 4 | DERL2 | derlin 2 | Cytoplasm | other | |
| chr4 | *UBE2D3* | 2 | 51 | 4 | UBE2D3 | ubiquitin conjugating enzyme E2 | Cytoplasm | enzyme | |
| chr5 | *CD180* | 2 | 32 | 4 | CD180 | CD180 molecule | Plasma Membrane | other | |
| chr14 | *TRAPPC6B* | 2 | 41 | 4 | TRAPPC6B | trafficking protein particle | Cytoplasm | other | |
| chr10 | *RP11-342C24.8* | 2 | 83 | 4 | | | | | |
| chr18 | *RNMT* | 2 | 37 | 4 | RNMT | RNA guanine-7 | Nucleus | enzyme | |
| chr7 | *GNG11* | 1 | 36 | 4 | GNG11 | G protein subunit gamma 11 | Plasma Membrane | other | |
| chr17 | *LINC00672* | 2 | 44 | 4 | LINC00672 | long intergenic non-protein coding RNA 672 | Other | other | |
| chr9 | *LINC00094* | 2 | 94 | 4 | BRD3OS | BRD3 opposite strand | Other | other | |
| chrX | *VCX* | 1 | 92 | 4 | VCX/VCX3A | variable charge X-linked 3A | Nucleus | other | |
| chr19 | *CTC-471J1.8* | 2 | 29 | 4 | | | | | |
| chr1 | *RP5-968P14.2* | 2 | 50 | 4 | | | | | |
| chr7 | *RP11-73B2.4* | 2 | 40 | 4 | | | | | |
| chr1 | *RP11-54O7.16* | 2 | 137 | 4 | | | | | |
| chr2 | *IRS1* | 2 | 53 | 4 | IRS1 | insulin receptor substrate 1 | Cytoplasm | enzyme | |
| chr11 | *PELI3* | 2 | 97 | 4 | PELI3 | pellino E3 ubiquitin protein ligase family member 3 | Cytoplasm | enzyme | |
| chr3 | *MUC20* | 2 | 42 | 4 | MUC20 | mucin 20, cell surface associated | Plasma Membrane | other | |
| chr17 | *RP11-707O23.5* | 2 | 88 | 4 | | | | | |
| chr2 | *FAM228A* | 2 | 45 | 4 | FAM228A | family with sequence similarity 228 member A | Other | other | |
| chr11 | *RP11-770J1.4* | 2 | 55 | 4 | | | | | |
| chr17 | *RP11-353N14.3* | 2 | 86 | 4 | | | | | |
| chr1 | *RP11-465B22.3* | 2 | 111 | 4 | | | | | |
| chr11 | *ALKBH3* | 2 | 35 | 4 | ALKBH3 | alkB homolog 3, alpha-ketoglutaratedependent | Nucleus | enzyme | |
| chr17 | *ZNF232* | 2 | 45 | 4 | ZNF232 | zinc finger protein 232 | Nucleus | transcription regulator | |
| chr9 | *RP11-544A12.5* | 2 | 69 | 4 | | | | | |
| chr3 | *SALL4P5* | 2 | 89 | 4 | SALL4P5 | spalt like transcription factor 4 pseudogene 5 | Other | other | |
| chr14 | *EXD2* | 2 | 29 | 4 | EXD2 | exonuclease 3'-5' domain containing 2 | Cytoplasm | enzyme | |
| chr1 | *RP4-758J18.10* | 2 | 76 | 4 | | | | | |
| chr16 | *RP11-169E6.1* | 2 | 49 | 4 | | | | | |
| chr15 | *RP11-424I19.2* | 2 | 95 | 4 | | | | | |

| | | | | | | | | | |
|---|---|---|---|---|---|---|---|---|---|
| chr16 | *CHP2* | 2 | 59 | 4 | CHP2 | calcineurin like EF-hand protein 2 | Cytoplasm | other | |
| chr19 | *SYCN* | 2 | 53 | 4 | SYCN | syncollin | Extracellular Space | other | |
| chr3 | *HYAL3* | 2 | 109 | 4 | HYAL3 | hyaluronidase 3 | Cytoplasm | enzyme | diclofenac/hyaluronic acid, hyaluronic acid, hylan |
| chr19 | *MIR7-3HG* | 2 | 52 | 4 | MIR7-3HG | MIR7-3 host gene | Other | other | |
| chr3 | *TRAT1* | 2 | 35 | 4 | TRAT1 | T cell receptor associated transmembrane adaptor 1 | Plasma Membrane | kinase | |
| chr3 | *RP11-536I6.1* | 2 | 87 | 4 | | | | | |
| chr5 | *C5orf58* | 2 | 99 | 4 | C5orf58 | chromosome 5 open reading frame 58 | Other | other | |
| chr20 | *RP4-715N11.2* | 2 | 57 | 4 | | | | | |
| chr17 | *PHOSPHO1* | 2 | 97 | 4 | PHOSPHO1 | phosphoethanolamine/phosphocholine phosphatase 1 | Extracellular Space | enzyme | |
| chr2 | *C2orf54* | 2 | 79 | 4 | MAB21L4 | mab-21 like 4 | Other | other | |
| chr6 | *GLYATL3* | 2 | 51 | 4 | GLYATL3 | glycine-N-acyltransferase like 3 | Other | other | |
| chr12 | *CYP27B1* | 2 | 65 | 4 | CYP27B1 | cytochrome P450 family 27 subfamily B member 1 | Cytoplasm | enzyme | |
| chr16 | *SNRNP25* | 2 | 71 | 4 | SNRNP25 | small nuclear ribonucleoprotein U11/U12 subunit 25 | Nucleus | other | |
| chr1 | *RP1-212P9.2* | 2 | 48 | 4 | | | | | |
| chr15 | *CIB1* | 2 | 98 | 4 | CIB1 | calcium and integrin binding 1 | Nucleus | other | |
| chr6 | *TAF11* | 2 | 69 | 4 | TAF11 | TATA-box binding protein associated factor 11 | Nucleus | transcription regulator | |
| chr9 | *SNHG7* | 2 | 87 | 4 | SNHG7 | small nucleolar RNA host gene 7 | Other | other | |
| chr1 | *RP11-153F1.1* | 2 | 31 | 4 | | | | | |
| chr5 | *RP11-122C5.1* | 2 | 129 | 4 | | | | | |
| chr17 | *CBX8* | 2 | 87 | 4 | CBX8 | chromobox 8 | Nucleus | other | |
| chr11 | *CD3G* | 2 | 48 | 4 | CD3G | CD3g molecule | Plasma Membrane | transmembrane recepto | blinatumomab |
| chr8 | *FAM92A1* | 2 | 29 | 4 | FAM92A | family with sequence similarity 92 member A | Cytoplasm | other | |
| chr17 | *TCAP* | 2 | 118 | 4 | TCAP | titin-cap | Cytoplasm | other | |
| chr19 | *ZNF440* | 2 | 43 | 4 | ZNF440 | zinc finger protein 440 | Nucleus | transcription regulator | |
| chr2 | *CXCR2* | 2 | 42 | 4 | CXCR2 | C-X-C motif chemokine receptor 2 | Plasma Membrane | G-protein coupled recepto | SB-265610, SX-682, QBM076, navarixin, 2'-((4'-trifluoromethanesulfonyloxy)phenyl)-N-methanesulfonylpropionamide, repertaxin, AZD5069 |
| chr3 | *HSPD1P6* | 2 | 37 | 4 | HSPD1P6 | heat shock protein family D (Hsp60) member 1 pseudogene 6 | Other | other | |
| chr20 | *MRGBP* | 2 | 250 | 4 | MRGBP | MRG domain binding protein | Nucleus | other | |
| chr9 | *HEMGN* | 2 | 28 | 4 | HEMGN | hemogen | Nucleus | other | |
| chr4 | *ZEB2P1* | 2 | 31 | 4 | ZEB2P1 | zinc finger E-box binding homeobox 2 pseudogene 1 | Other | other | |
| chr3 | *ASB14* | 2 | 29 | 4 | ASB14 | ankyrin repeat and SOCS box containing 14 | Cytoplasm | other | |
| chr19 | *FAM32A* | 2 | 79 | 4 | FAM32A | family with sequence similarity 32 member A | Nucleus | other | |
| chr8 | *HTRA4* | 2 | 56 | 4 | HTRA4 | HtrA serine peptidase 4 | Other | peptidase | |
| chr5 | *AC112198.1* | 2 | 39 | 4 | | | | | |
| chr22 | *Z83851.1* | 2 | 55 | 4 | | | | | |
| chr17 | *LINC00483* | 2 | 57 | 4 | ANKRD40CL | ANKRD40 C-terminal like | Other | other | |
| chr22 | *DGCR6* | 2 | 90 | 4 | CR6/LOC102724 | DiGeorge syndrome critical region gene 6 | Nucleus | other | |
| chrX | *FAM47C* | 2 | 62 | 4 | FAM47C | family with sequence similarity 47 member C | Other | other | |
| chr3 | *CICP6* | 2 | 85 | 4 | CICP6 | capicua transcriptional repressor pseudogene 6 | Other | other | |
| chr16 | *TMEM186* | 2 | 51 | 4 | TMEM186 | transmembrane protein 186 | Cytoplasm | other | |
| chr16 | *ELMO3* | 2 | 64 | 4 | ELMO3 | engulfment and cell motility 3 | Cytoplasm | other | |
| chr10 | *BCCIP* | 2 | 75 | 4 | BCCIP | BRCA2 and CDKN1A interacting protein | Nucleus | other | |
| chr11 | *CST6* | 2 | 109 | 4 | CST6 | cystatin E/M | Extracellular Space | other | |
| chr18 | *RAB27B* | 2 | 49 | 4 | RAB27B | RAB27B, member RAS oncogene family | Cytoplasm | enzyme | |
| chr13 | *LINC00423* | 2 | 34 | 4 | LINC00423 | long intergenic non-protein coding RNA 423 | Other | other | |
| chr1 | *RP11-117D22.2* | 2 | 56 | 4 | | | | | |
| chr5 | *ECSCR* | 2 | 52 | 4 | ECSCR | endothelial cell surface expressed chemotaxis and | Other | other | |
| chr11 | *BSX* | 2 | 75 | 4 | BSX | brain specific homeobox | Cytoplasm | transcription regulator | |
| chr9 | *INIP* | 2 | 40 | 4 | INIP | INTS3 and NABP interacting | Nucleus | other | |
| chr2 | *AC097468.7* | 2 | 47 | 4 | | | | | |
| chr11 | *TRIM49* | 2 | 33 | 4 | TRIM49/TRIM49C | tripartite motif containing 49 | Cytoplasm | other | |
| chr6 | *CUL7* | 2 | 56 | 4 | CUL7 | cullin 7 | Cytoplasm | other | |
| chr5 | *RP11-826N14.1* | 2 | 74 | 4 | | | | | |
| chr3 | *RP11-3B7.1* | 2 | 50 | 4 | | | | | |
| chr12 | *SELPLG* | 2 | 74 | 4 | SELPLG | selectin P ligand | Plasma Membrane | other | |
| chr5 | *ENC1* | 2 | 48 | 4 | ENC1 | ectodermal-neural cortex 1 | Nucleus | peptidase | |
| chr8 | *CTD-2168K21.1* | 2 | 78 | 4 | | | | | |
| chr2 | *AC016735.3* | 2 | 77 | 4 | | | | | |
| chr17 | *RP11-387H17.6* | 2 | 51 | 4 | | | | | |
| chr4 | *NAAA* | 2 | 43 | 4 | NAAA | N-acylethanolamine acid amidase | Cytoplasm | enzyme | |
| chrX | *MAGEB2* | 2 | 62 | 4 | MAGEB2 | MAGE family member B2 | Cytoplasm | other | |
| chr11 | *TRPM5* | 2 | 112 | 4 | TRPM5 | transient receptor potential cation channel subfamily M member 5 | Plasma Membrane | ion channel | |

| | | | | | | | | |
|---|---|---|---|---|---|---|---|---|
| chrX | *PHKA2-AS1* | 2 | 52 | 4 | PHKA2-AS1 | PHKA2 antisense RNA 1 | Other | other |
| chr2 | *MPP4* | 2 | 28 | 4 | MPP4 | membrane palmitoylated protein | Cytoplasm | kinase |
| chr11 | *MPZL2* | 2 | 43 | 4 | MPZL2 | myelin protein zero like 2 | Plasma Membrane | other |
| chrX | *OTUD6A* | 2 | 50 | 4 | OTUD6A | OTU deubiquitinase 6A | Other | peptidase |
| chr9 | *NOXA1* | 2 | 91 | 4 | NOXA1 | NADPH oxidase activator 1 | Plasma Membrane | other |
| chr16 | *AC009093.1* | 2 | 62 | 4 | | | | |
| chr11 | *SLC17A6* | 2 | 91 | 4 | SLC17A6 | solute carrier family 17 member 6 | Plasma Membrane | transporter |
| chr5 | *CTD-2175A23.1* | 2 | 50 | 4 | | | | |
| chr19 | *STXBP2* | 2 | 68 | 4 | STXBP2 | syntaxin binding protein 2 | Plasma Membrane | transporter |
| chr17 | *AC006050.2* | 2 | 54 | 4 | | | | |
| chr15 | *RP11-697E2.7* | 2 | 64 | 4 | | | | |
| chrX | *ARHGEF6* | 2 | 32 | 4 | ARHGEF6 | Rac/Cdc42 guanine nucleotide exchange factor 6 | Cytoplasm | other |
| chr5 | *ZBED3* | 2 | 45 | 4 | ZBED3 | zinc finger BED-type containing 3 | Cytoplasm | transcription regulator |
| chr17 | *CTD-2319I12.2* | 2 | 42 | 4 | | | | |
| chr21 | *AP003900.6* | 2 | 110 | 4 | | | | |
| chr13 | *AL139319.1* | 1 | 38 | 4 | | | | |
| chr15 | *RP11-540O11.1* | 2 | 90 | 4 | | | | |
| chr17 | *SLC16A6P1* | 2 | 40 | 4 | SLC16A6P1 | SLC16A6 pseudogene 1 | Other | other |
| chr1 | *ISG20L2* | 2 | 65 | 4 | ISG20L2 | interferon stimulated exonuclease gene 20 like 2 | Nucleus | enzyme |
| chr1 | *RP1-10C16.1* | 2 | 39 | 4 | | | | |
| chr9 | *RP11-74C3.1* | 2 | 36 | 4 | | | | |
| chr1 | *HOOK1* | 2 | 56 | 4 | HOOK1 | hook microtubule tethering | Cytoplasm | other |
| chr11 | *AP001891.1* | 2 | 54 | 4 | | | | |
| chr16 | *DEXI* | 2 | 54 | 4 | DEXI | Dexi homolog | Other | other |
| chr11 | *RP11-727A23.5* | 2 | 54 | 4 | | | | |
| chr1 | *MRPS15* | 2 | 62 | 4 | MRPS15 | mitochondrial ribosomal protein | Cytoplasm | other |
| chr3 | *ZNF502* | 2 | 35 | 4 | ZNF502 | zinc finger protein 502 | Nucleus | transcription regulator |
| chr17 | *CTD-2561B21.11* | 2 | 56 | 4 | | | | |
| chr21 | *ANKRD30BP2* | 2 | 56 | 4 | ANKRD30BP2 | ankyrin repeat domain 30B pseudogene 2 | Other | other |
| chr9 | *SAPCD2* | 2 | 168 | 4 | SAPCD2 | suppressor APC domain | Nucleus | other |
| chr9 | *OR5C1* | 2 | 79 | 4 | OR5C1 | olfactory receptor family 5 subfamily C member 1 | Plasma Membrane | G-protein coupled recepto |
| chr4 | *RP11-367J11.3* | 2 | 67 | 4 | | | | |
| chr5 | *CTC-467M3.1* | 2 | 33 | 4 | | | | |
| chr1 | *RP4-714D9.5* | 2 | 36 | 4 | | | | |
| chr20 | *SOCS2P1* | 2 | 54 | 4 | SOCS2P1 | suppressor of cytokine signaling 2 pseudogene 1 | Other | other |
| chr5 | *PCDHA1* | 2 | 57 | 4 | PCDHA1 | protocadherin alpha 1 | Plasma Membrane | other |
| chr4 | *HPGDS* | 2 | 51 | 4 | HPGDS | hematopoietic prostaglandin D synthase | Cytoplasm | enzyme |
| chr17 | *GPR142* | 2 | 79 | 4 | GPR142 | G protein-coupled receptor 142 | Plasma Membrane | G-protein coupled recepto |
| chr12 | *C1QL4* | 2 | 95 | 4 | C1QL4 | complement C1q like 4 | Extracellular Space | other |
| chr10 | *RP11-564C4.6* | 2 | 100 | 4 | | | | |
| chr17 | *RP11-19P22.7* | 2 | 87 | 4 | | | | |
| chr16 | *RP11-483P21.2* | 2 | 51 | 4 | | | | |
| chr9 | *ANXA2P2* | 2 | 47 | 4 | ANXA2P2 | annexin A2 pseudogene 2 | Cytoplasm | other |
| chr13 | *SLC25A15* | 2 | 38 | 4 | SLC25A15 | solute carrier family 25 member | Cytoplasm | transporter |
| chr7 | *RP4-813F11.4* | 1 | 28 | 4 | | | | |
| chr9 | *ASS1P12* | 1 | 36 | 4 | ASS1P12 | argininosuccinate synthetase 1 pseudogene 12 | Other | other |
| chr17 | *RP11-1148O4.2* | 2 | 41 | 4 | | | | |
| chr1 | *FAM177B* | 1 | 42 | 4 | FAM177B | family with sequence similarity 177 member B | Other | other |
| chr4 | *RP11-496D24.2* | 2 | 28 | 4 | | | | |
| chr3 | *ARL6* | 2 | 60 | 4 | ARL6 | ADP ribosylation factor like | Cytoplasm | transporter |
| chr19 | *ZNF30* | 2 | 42 | 4 | ZNF30 | zinc finger protein 30 | Nucleus | transcription regulator |
| chr7 | *AC093627.10* | 2 | 108 | 4 | | | | |
| chr1 | *RP11-522D2.1* | 2 | 42 | 4 | | | | |
| chr7 | *CYCS* | 2 | 51 | 4 | CYCS | cytochrome c, somatic | Cytoplasm | transporter |
| chr12 | *MFSD5* | 2 | 103 | 4 | MFSD5 | major facilitator superfamily domain containing 5 | Other | transporter |
| chr7 | *SLC29A4P1* | 2 | 66 | 4 | C29A4P1/SLC29A | solute carrier family 29 member 4 pseudogene 1 | Other | other |
| chr10 | *LINC00707* | 2 | 47 | 4 | LINC00707 | long intergenic non-protein coding RNA 707 | Other | other |
| chr1 | *CHIA* | 2 | 59 | 4 | CHIA | chitinase acidic | Extracellular Space | enzyme |
| chr5 | *CTC-498M16.4* | 1 | 32 | 4 | | | | |
| chr15 | *RP11-198M11.2* | 2 | 29 | 4 | | | | |
| chr20 | *WI2-87327B8.1* | 2 | 127 | 4 | | | | |
| chr21 | *ERVH48-1* | 2 | 43 | 4 | ERVH48-1 | endogenous retrovirus group 48 member 1 | Extracellular Space | other |
| chr21 | *AP001255.2* | 1 | 27 | 4 | | | | |
| chr18 | *RP11-64C12.6* | 2 | 58 | 4 | | | | |
| chr7 | *RP11-212P7.3* | 1 | 38 | 4 | | | | |
| chr6 | *RP3-399L15.3* | 2 | 30 | 4 | | | | |
| chr2 | *FAM136A* | 2 | 45 | 4 | FAM136A | family with sequence similarity 136 member A | Cytoplasm | other |
| chr19 | *CHMP2A* | 2 | 55 | 4 | CHMP2A | charged multivesicular body protein 2A | Cytoplasm | other |
| chr15 | *SCAND2P* | 2 | 55 | 4 | SCAND2P | SCAN domain containing 2 pseudogene | Nucleus | other |
| chr12 | *OR7E47P* | 2 | 70 | 4 | OR7E47P | olfactory receptor family 7 subfamily E member 47 | Other | other |
| chr20 | *ZNF334* | 2 | 45 | 4 | ZNF334 | zinc finger protein 334 | Nucleus | transcription regulator |
| chr17 | *RP11-459C13.1* | 2 | 94 | 4 | | | | |

| | | | | | | | | | |
|---|---|---|---|---|---|---|---|---|---|
| chr17 | *KRT20* | 2 | 59 | 4 | KRT20 | keratin 20 | Cytoplasm | other | rituximab/temozolomide, alemtuzumab/rituximab, chlorambucil/rituximab, rituximab/thiotepa, brentuximab vedotin/rituximab, cyclophosphamide/pentostatin/rituximab, rituximab/thalidomide, paclitaxel/rituximab, lenalidomide/rituximab, rituximab, bortezomib/rituximab, bendamustine/ibrutinib/rituximab, methotrexate/rituximab/temozolomide, bendamustine/rituximab, bendamustine/idelalisib/rituximab, [illegible] |
| chr1 | *LRRC37A12P* | 2 | 80 | 4 | LRRC37A12P | leucine rich repeat containing 37 member A12, pseudogene | Other | other | |
| chr16 | *CTRB1* | 2 | 98 | 4 | CTRB1 | chymotrypsinogen B1 | Extracellular Space | peptidase | |
| chr2 | *AC098617.1* | 2 | 60 | 4 | | | | | |
| chr17 | *SYNGR2* | 2 | 60 | 4 | SYNGR2 | synaptogyrin 2 | Cytoplasm | other | |
| chr12 | *RP11-923I11.5* | 2 | 75 | 4 | | | | | |
| chr7 | *AC073150.6* | 2 | 82 | 4 | | | | | |
| chr1 | *RP11-431K24.2* | 2 | 44 | 4 | | | | | |
| chr14 | *RP11-986E7.7* | 2 | 47 | 4 | | | | | |
| chr7 | *AC091729.9* | 2 | 65 | 4 | | | | | |
| chr1 | *TNFRSF14* | 2 | 112 | 4 | TNFRSF14 | TNF receptor superfamily | Plasma Membrane | transmembrane receptor | |
| chr17 | *RP11-434D2.11* | 2 | 51 | 4 | | | | | |
| chr21 | *RWDD2B* | 2 | 55 | 4 | RWDD2B | RWD domain containing 2B | Other | other | |
| chr15 | *ADAL* | 2 | 27 | 4 | ADAL | adenosine deaminase like | Cytoplasm | enzyme | |
| chr2 | *ORC2* | 2 | 27 | 4 | ORC2 | origin recognition complex | Nucleus | other | |
| chr9 | *KIAA1161* | 2 | 54 | 4 | MYORG | myogenesis regulating glycosidase (putative) | Nucleus | other | |
| chr19 | *KCNA7* | 2 | 65 | 4 | KCNA7 | potassium voltage-gated channel subfamily A member 7 | Plasma Membrane | ion channel | dalfampridine, amiodarone, |
| chr17 | *SLC25A11* | 2 | 66 | 4 | SLC25A11 | solute carrier family 25 member | Cytoplasm | transporter | |
| chr6 | *HLA-DRB6* | 2 | 58 | 4 | HLA-DRB6 | major histocompatibility complex, class II, DR beta 6 (pseudogene) | Other | other | |
| chr4 | *RP11-571L19.7* | 2 | 27 | 4 | | | | | |
| chr2 | *CEBPZ* | 2 | 32 | 4 | CEBPZ | CCAAT enhancer binding protein zeta | Nucleus | transcription regulator | |
| chr3 | *ALDOAP1* | 2 | 59 | 4 | ALDOAP1 | ALDOA pseudogene 1 | Other | other | |
| chr6 | *MPC1* | 2 | 27 | 4 | MPC1 | mitochondrial pyruvate carrier 1 | Cytoplasm | other | |
| chr7 | *MPLKIP* | 2 | 58 | 4 | MPLKIP | M-phase specific PLK1 interacting protein | Nucleus | other | |
| chr1 | *RP1-43O17.2* | 1 | 47 | 4 | | | | | |
| chr4 | *RP11-103J17.2* | 2 | 45 | 4 | | | | | |
| chr17 | *GNGT2* | 2 | 53 | 4 | GNGT2 | G protein subunit gamma transducin 2 | Plasma Membrane | enzyme | |
| chr1 | *RP5-908D6.1* | 2 | 35 | 4 | | | | | |
| chr2 | *RP11-355B11.2* | 2 | 38 | 4 | | | | | |
| chrX | *WASF4P* | 2 | 43 | 4 | WASF4P | WASP family member 4, | Other | other | |
| chr22 | *AP000350.10* | 2 | 51 | 4 | | | | | |
| chr12 | *C12orf71* | 2 | 46 | 4 | C12orf71 | chromosome 12 open reading frame 71 | Other | other | |
| chr8 | *SLC26A7* | 2 | 32 | 4 | SLC26A7 | solute carrier family 26 member 7 | Plasma Membrane | transporter | |
| chrX | *CROCCP1* | 2 | 69 | 4 | CROCCP1 | CROCC pseudogene 1 | Other | other | |
| chr14 | *CTD-3035D6.2* | 2 | 38 | 4 | | | | | |
| chr5 | *AC136604.1* | 2 | 81 | 4 | | | | | |
| chr2 | *LINC00954* | 2 | 59 | 4 | LINC00954 | long intergenic non-protein coding RNA 954 | Other | other | |
| chr6 | *KHDC1* | 2 | 56 | 4 | KHDC1 | KH domain containing 1 | Other | other | |
| chr8 | *REXO1L10P* | 2 | 278 | 4 | REXO1L10P | REXO1 like 10, pseudogene | Other | other | |
| chr5 | *FAM53C* | 2 | 75 | 4 | FAM53C | family with sequence similarity 53 member C | Other | other | |
| chr19 | *ZNF225* | 2 | 35 | 4 | ZNF225 | zinc finger protein 225 | Nucleus | transcription regulator | |
| chr3 | *PDCD10* | 2 | 67 | 4 | PDCD10 | programmed cell death 10 | Cytoplasm | other | |
| chr6 | *TRIM38* | 2 | 38 | 4 | TRIM38 | tripartite motif containing 38 | Cytoplasm | other | |
| chr2 | *AC007970.1* | 2 | 38 | 4 | | | | | |
| chr5 | *CTC-529L17.2* | 1 | 32 | 4 | | | | | |
| chr17 | *RP11-927P21.6* | 2 | 59 | 4 | | | | | |
| chr9 | *XXyac-YM21GA2.* | 2 | 116 | 4 | | | | | |
| chr19 | *CRX* | 2 | 44 | 4 | CRX | cone-rod homeobox | Nucleus | transcription regulator | |
| chr18 | *ROCK1P1* | 2 | 5426 | 4 | ROCK1P1 | Rho associated coiled-coil containing protein kinase 1 | Other | other | |
| chrX | *SMPX* | 2 | 30 | 4 | SMPX | small muscle protein X-linked | Cytoplasm | other | |
| chr14 | *RP11-1152H15.1* | 2 | 35 | 4 | | | | | |
| chr11 | *RP11-800A3.3* | 2 | 77 | 4 | | | | | |
| chr13 | *PRR20A* | 2 | 251 | 4 | R20A (includes oth | proline rich 20A | Other | other | |
| chr8 | *RNF19A* | 2 | 30 | 4 | RNF19A | ring finger protein 19A, RBR E3 ubiquitin protein ligase | Nucleus | enzyme | |

| | | | | | | | | | |
|---|---|---|---|---|---|---|---|---|---|
| chr4 | *GLRB* | 2 | 33 | 4 | GLRB | glycine receptor beta | Plasma Membrane | ion channel | |
| chr19 | *ZNF773* | 2 | 29 | 4 | ZNF773 | zinc finger protein 773 | Other | other | |
| chr20 | *WFDC12* | 2 | 61 | 4 | WFDC12 | WAP four-disulfide core domain | Extracellular Space | other | |
| chr8 | *LRP12* | 2 | 32 | 4 | LRP12 | LDL receptor related protein 12 | Plasma Membrane | transmembrane receptor | |
| chr6 | *RP11-346D19.1* | 2 | 52 | 4 | | | | | |
| chr17 | *CISD3* | 2 | 90 | 4 | CISD3 | CDGSH iron sulfur domain 3 | Cytoplasm | other | |
| chr16 | *MT4* | 2 | 54 | 4 | MT4 | metallothionein 4 | Other | other | |
| chr11 | *FZD4* | 2 | 65 | 4 | FZD4 | frizzled class receptor 4 | Plasma Membrane | G-protein coupled recepto | |
| chr12 | *RP11-956E11.1* | 2 | 29 | 4 | | | | | |
| chr7 | *AGR3* | 2 | 43 | 4 | AGR3 | anterior gradient 3, protein disulphide isomerase family | Extracellular Space | other | |
| chr5 | *IL4* | 2 | 45 | 4 | IL4 | interleukin 4 | Extracellular Space | cytokine | |
| chr1 | *MAB21L3* | 2 | 55 | 4 | MAB21L3 | mab-21 like 3 | Other | other | |
| chr5 | *RP11-175K6.1* | 2 | 123 | 4 | | | | | |
| chr6 | *RP11-240M16.1* | 2 | 36 | 4 | | | | | |
| chr13 | *METTL21EP* | 2 | 31 | 4 | METTL21EP | methyltransferase like 21E, pseudogene | Other | other | |
| chr7 | *HAUS6P1* | 1 | 43 | 4 | HAUS6P1 | HAUS augmin like complex subunit 6 pseudogene 1 | Other | other | |
| chr21 | *SIK1* | 2 | 109 | 4 | SIK1/SIK1B | salt inducible kinase 1 | Nucleus | kinase | dabrafenib/ipilimumab/trametinib, dabrafenib, dabrafenib/panitumumab/trametinib, dabrafenib/trametinib, dabrafenib/trametinib/vemurafenib, cetuximab/dabrafenib/trametinib, |
| chr1 | *TRIM45* | 2 | 81 | 4 | TRIM45 | tripartite motif containing 45 | Cytoplasm | other | |
| chr6 | *ZNF204P* | 2 | 58 | 4 | ZNF204P | zinc finger protein 204, | Other | other | |
| chrX | *SEPHS1P4* | 1 | 28 | 4 | SEPHS1P4 | selenophosphate synthetase 1 pseudogene 4 | Other | other | |
| chr2 | *AC007381.3* | 2 | 55 | 4 | | | | | |
| chr18 | *RP11-64C12.3* | 2 | 56 | 4 | | | | | |
| chr10 | *RP11-464F9.20* | 2 | 90 | 4 | | | | | |
| chr16 | *RP11-143N13.1* | 2 | 37 | 4 | | | | | |
| chr2 | *AC104135.3* | 2 | 47 | 4 | | | | | |
| chr2 | *ADI1* | 2 | 45 | 4 | ADI1 | acireductone dioxygenase 1 | Nucleus | enzyme | |
| chr14 | *CTD-2142D14.1* | 2 | 46 | 4 | | | | | |
| chrX | *NONO* | 2 | 48 | 4 | NONO | non-POU domain containing octamer binding | Nucleus | transcription regulator | |
| chr19 | *C19orf24* | 2 | 126 | 4 | FAM174C | family with sequence similarity 174 member C | Other | other | |
| chr5 | *HARS2* | 2 | 35 | 4 | HARS2 | histidyl-tRNA synthetase 2, mitochondrial | Cytoplasm | enzyme | |
| chr19 | *OR1M1* | 2 | 56 | 4 | OR1M1 | olfactory receptor family 1 subfamily M member 1 | Plasma Membrane | G-protein coupled recepto | |
| chr14 | *SYT16* | 2 | 39 | 4 | SYT16 | synaptotagmin 16 | Other | transporter | |
| chr19 | *OCEL1* | 2 | 54 | 4 | OCEL1 | occludin/ELL domain containing 1 | Other | other | |
| chr1 | *RP11-328D5.1* | 2 | 54 | 4 | | | | | |
| chr1 | *TUBB8P10* | 2 | 90 | 4 | TUBB8P10 | tubulin beta 8 class VIII pseudogene 10 | Other | other | |
| chr12 | *C12orf23* | 2 | 57 | 4 | TMEM263 | transmembrane protein 263 | Other | other | |
| chr4 | *ANXA3* | 2 | 27 | 4 | ANXA3 | annexin A3 | Cytoplasm | enzyme | |
| chr3 | *DHFRL1* | 1 | 38 | 4 | DHFR2 | dihydrofolate reductase 2 | Cytoplasm | enzyme | |
| chr1 | *CICP5* | 2 | 105 | 4 | CICP5 | capicua transcriptional repressor pseudogene 5 | Other | other | |
| chr14 | *KLHL28* | 2 | 31 | 4 | KLHL28 | kelch like family member 28 | Other | other | |
| chr3 | *VN2R1P* | 2 | 59 | 4 | VN2R1P | vomeronasal 2 receptor 1 pseudogene | Other | other | |
| chr2 | *AHCYP3* | 2 | 61 | 4 | AHCYP3 | adenosylhomocysteinase pseudogene 3 | Other | other | |
| chr8 | *AC145123.2* | 2 | 59 | 4 | | | | | |
| chr20 | *NPEPL1* | 2 | 153 | 4 | NPEPL1 | aminopeptidase like 1 | Nucleus | peptidase | |
| chr15 | *CKMT1B* | 2 | 39 | 4 | CKMT1A/CKMT1E | creatine kinase, mitochondrial 1B | Cytoplasm | kinase | |
| chr1 | *C1orf52* | 2 | 42 | 4 | C1orf52 | chromosome 1 open reading frame 52 | Nucleus | other | |
| chr19 | *ISYNA1* | 2 | 120 | 4 | ISYNA1 | inositol-3-phosphate synthase 1 | Cytoplasm | enzyme | |
| chr9 | *RP11-536O18.1* | 2 | 46 | 4 | | | | | |
| chr5 | *MED10* | 2 | 148 | 4 | MED10 | mediator complex subunit 10 | Nucleus | other | |
| chr10 | *C10orf85* | 2 | 48 | 4 | LINC01561 | long intergenic non-protein coding RNA 1561 | Other | other | |
| chr7 | *HOTAIRM1* | 2 | 120 | 4 | HOTAIRM1 | HOXA transcript antisense RNA, myeloid-specific 1 | Other | other | |

| | | | | | | | | | |
|---|---|---|---|---|---|---|---|---|---|
| chr16 | *CD19* | 2 | 74 | 4 | CD19 | CD19 molecule | Plasma Membrane | transmembrane receptor | HD37-dgRTA, DI-B4, blinatumomab/dasatinib, SAR3419, tafasitamab, MDX-1342 antibody, MEDI-551, SGN-CD19A, AMG 562, blinatumomab/imatinib, blinatumomab, XmAb5871, combotox, DT2219ARL, anti-CD19/CD3 tetravalent antibody AFM11, RO7227166 |
| chr19 | *ZNF667-AS1* | 2 | 32 | 4 | ZNF667-AS1 | ZNF667 antisense RNA 1 (head to head) | Other | other | |
| chr11 | *KLC2* | 2 | 114 | 4 | KLC2 | kinesin light chain 2 | Cytoplasm | other | |
| chr1 | *PRAMEF12* | 2 | 72 | 4 | PRAMEF12 | PRAME family member 12 | Other | other | |
| chr18 | *RP11-888D10.3* | 2 | 42 | 4 | | | | | |
| chr11 | *CTBP2P6* | 2 | 60 | 4 | CTBP2P6 | C-terminal binding protein 2 pseudogene 6 | Other | other | |
| chr15 | *RP11-60L3.1* | 2 | 66 | 4 | | | | | |
| chr5 | *CTD-2296D1.5* | 2 | 83 | 4 | | | | | |
| chr1 | *PDZK1P1* | 2 | 44 | 4 | PDZK1P1 | PDZ domain containing 1 pseudogene 1 | Other | other | |
| chr17 | *RP11-599B13.6* | 2 | 67 | 4 | | | | | |
| chr8 | *AC104417.1* | 1 | 47 | 4 | | | | | |
| chr14 | *FAM71D* | 2 | 27 | 4 | FAM71D | family with sequence similarity 71 member D | Other | other | |
| chr2 | *AC013402.5* | 2 | 76 | 4 | | | | | |
| chr1 | *RP5-858B6.3* | 2 | 62 | 4 | | | | | |
| chr16 | *RP11-616M22.7* | 2 | 53 | 4 | | | | | |
| chr19 | *CTD-2525I3.3* | 2 | 47 | 4 | | | | | |
| chr20 | *SNHG11* | 2 | 72 | 4 | SNHG11 | small nucleolar RNA host gene | Other | other | |
| chr19 | *KLK6* | 2 | 45 | 4 | KLK6 | kallikrein related peptidase 6 | Extracellular Space | peptidase | |
| chr10 | *RP11-442O18.1* | 2 | 46 | 4 | | | | | |
| chr17 | *RPL23* | 2 | 46 | 4 | RPL23 | ribosomal protein L23 | Cytoplasm | other | |
| chr11 | *RP11-665E10.5* | 1 | 48 | 4 | | | | | |
| chr16 | *RP11-488I20.3* | 2 | 28 | 4 | | | | | |
| chr11 | *LRRC10B* | 2 | 87 | 4 | LRRC10B | leucine rich repeat containing | Other | other | |
| chr1 | *KPRP* | 2 | 88 | 4 | KPRP | keratinocyte proline rich protein | Cytoplasm | other | |
| chr22 | *NEFH* | 2 | 56 | 4 | NEFH | neurofilament heavy | Cytoplasm | other | |
| chr17 | *BTBD17* | 2 | 93 | 4 | BTBD17 | BTB domain containing 17 | Plasma Membrane | other | |
| chr9 | *RP11-426A6.5* | 2 | 131 | 4 | | | | | |
| chr11 | *PRG3* | 2 | 71 | 4 | PRG3 | proteoglycan 3, pro eosinophil major basic protein 2 | Extracellular Space | other | |
| chr16 | *RP11-680G10.1* | 2 | 90 | 4 | | | | | |
| chr7 | *RP9* | 2 | 46 | 4 | RP9 | RP9 pre-mRNA splicing factor | Nucleus | other | |
| chr14 | *NDRG2* | 2 | 62 | 4 | NDRG2 | NDRG family member 2 | Cytoplasm | other | |
| chr17 | *COPRS* | 2 | 38 | 4 | COPRS | coordinator of PRMT5 and differentiation stimulator | Nucleus | other | |
| chr4 | *SHISA3* | 2 | 48 | 4 | SHISA3 | shisa family member 3 | Other | other | |
| chr5 | *PCDHB11* | 2 | 36 | 4 | PCDHB11 | protocadherin beta 11 | Plasma Membrane | other | |
| chr2 | *AC007251.2* | 2 | 67 | 4 | | | | | |
| chr19 | *ZSCAN5C* | 2 | 42 | 4 | ZSCAN5C | zinc finger and SCAN domain containing 5C | Other | other | |
| chr1 | *TRMT13* | 1 | 34 | 4 | TRMT13 | tRNA methyltransferase 13 | Other | other | |
| chr16 | *THOC6* | 2 | 107 | 4 | THOC6 | THO complex 6 | Nucleus | other | |
| chr2 | *ACTR1B* | 2 | 88 | 4 | ACTR1B | actin related protein 1B | Cytoplasm | other | |
| chrX | *GS1-256O22.5* | 2 | 29 | 4 | | | | | |
| chr12 | *GS1-410F4.4* | 2 | 28 | 4 | | | | | |
| chr10 | *EIF5AL1* | 2 | 62 | 4 | EIF5AL1 | eukaryotic translation initiation factor 5A like 1 | Other | other | |
| chr17 | *AC005722.4* | 2 | 61 | 4 | | | | | |
| chr12 | *IL23A* | 2 | 67 | 4 | IL23A | interleukin 23 subunit alpha | Extracellular Space | cytokine | LY3074828, tildrakizumab, guselkumab |
| chr7 | *ARPC1B* | 2 | 109 | 4 | ARPC1B | actin related protein 2/3 complex subunit 1B | Cytoplasm | other | |
| chr1 | *RP11-439E19.9* | 2 | 45 | 4 | | | | | |
| chr1 | *RP11-261C10.2* | 2 | 77 | 4 | | | | | |
| chr6 | *RP11-570K4.1* | 2 | 49 | 4 | | | | | |
| chr11 | *OVOL1* | 2 | 105 | 4 | OVOL1 | ovo like transcriptional repressor | Nucleus | transcription regulator | |
| chr19 | *ICAM5* | 2 | 107 | 4 | ICAM5 | intercellular adhesion molecule 5 | Plasma Membrane | other | |

| | | | | | | | | | |
|---|---|---|---|---|---|---|---|---|---|
| chr2 | *PDCD1* | 2 | 82 | 4 | PDCD1 | programmed cell death 1 | Plasma Membrane | phosphatase | SCT-I10A, PF-06801591, GLS-010, RO7121661, MGD013, BI 754091, genolimzumab, XmAb20717, spartalizumab, MGD019, MEDI0680, camrelizumab, MEDI5752, ipilimumab/nivolumab, AK104, JNJ-63723283, pidilizumab, pembrolizumab/pemetrexed, pembrolizumab, anti PD-1 antibody, CS1003, HLX10, AGEN2034, BAT1306, dostarlimab, AMP-224, nivolumab, [illegible] |
| chr12 | *KRT127P* | 2 | 81 | 4 | KRT127P | keratin 127 pseudogene | Other | other | |
| chr5 | *FOXI1* | 2 | 64 | 4 | FOXI1 | forkhead box I1 | Nucleus | transcription regulator | |
| chr1 | *RP11-439L8.4* | 2 | 43 | 4 | | | | | |
| chr17 | *PSMB6* | 2 | 48 | 4 | PSMB6 | proteasome subunit beta 6 | Nucleus | peptidase | |
| chr1 | *BEST4* | 2 | 92 | 4 | BEST4 | bestrophin 4 | Plasma Membrane | other | |
| chr12 | *RP11-167N24.6* | 2 | 38 | 4 | | | | | |
| chr2 | *FRZB* | 2 | 52 | 4 | FRZB | frizzled related protein | Extracellular Space | other | |
| chr1 | *FMO2* | 2 | 38 | 4 | FMO2 | flavin containing dimethylaniline monoxygenase 2 | Cytoplasm | enzyme | |
| chr15 | *GABRA5* | 2 | 61 | 4 | GABRA5 | gamma-aminobutyric acid type A receptor alpha5 subunit | Plasma Membrane | ion channel | flunitrazepam, sevoflurane, alprazolam, butobarbital, atropine/hyoscyamine/phenobarbital/scopolamine, phenobarbital, diazepam, barbital, acetaminophen/butalbital/caffeine/codeine, hyoscyamine/phenobarbital, desflurane, adinazolam, SEP 174559, zaleplon, mephobarbital, ezogabine, flurazepam, butabarbital, midazolam, temazepam, diphenhydramine/mid[illegible] |
| chr19 | *CTC-454I21.3* | 2 | 31 | 4 | | | | | |
| chr7 | *AC024028.1* | 2 | 60 | 4 | | | | | |
| chr6 | *[illegible]Xyac-YX65C7_A.* | 2 | 48 | 4 | | | | | |
| chr4 | *CTD-2012I17.1* | 2 | 46 | 4 | | | | | |
| chr5 | *ST13P12* | 2 | 28 | 4 | ST13P12 | ST13, Hsp70 interacting protein pseudogene 12 | Other | other | |
| chr19 | *KLF1* | 2 | 61 | 4 | KLF1 | Kruppel like factor 1 | Nucleus | transcription regulator | |
| chr1 | *RP11-284F21.8* | 2 | 103 | 4 | | | | | |
| chr17 | *SLC16A13* | 2 | 72 | 4 | SLC16A13 | solute carrier family 16 member | Cytoplasm | other | |
| chr10 | *BMS1P1* | 1 | 26 | 4 | BMS1P1 | BMS1 pseudogene 1 | Other | other | |
| chr1 | *MAP1LC3C* | 2 | 62 | 4 | MAP1LC3C | microtubule associated protein 1 light chain 3 gamma | Cytoplasm | other | |
| chr17 | *RP11-1079K10.3* | 2 | 35 | 4 | | | | | |
| chr10 | *RP11-297A16.2* | 2 | 53 | 3 | | | | | |
| chr18 | *CXADRP3* | 2 | 39 | 3 | CXADRP3 | CXADR pseudogene 3 | Other | other | |
| chr13 | *TRIM13* | 2 | 37 | 3 | TRIM13 | tripartite motif containing 13 | Cytoplasm | enzyme | |
| chrX | *RP1-71L16.9* | 2 | 40 | 3 | | | | | |
| chr1 | *TMEM54* | 2 | 59 | 3 | TMEM54 | transmembrane protein 54 | Other | other | |
| chr15 | *RP11-665A22.1* | 2 | 43 | 3 | | | | | |
| chr4 | *LGI2* | 2 | 46 | 3 | LGI2 | leucine rich repeat LGI family member 2 | Extracellular Space | other | |
| chr7 | *AC226118.1* | 2 | 80 | 3 | | | | | |
| chr15 | *RP13-395E19.2* | 2 | 74 | 3 | | | | | |
| chr11 | *RP5-1027O15.1* | 2 | 33 | 3 | | | | | |
| chr4 | *AC226119.5* | 2 | 43 | 3 | | | | | |
| chr6 | *RP1-8B1.4* | 2 | 41 | 3 | | | | | |
| chr15 | *CTD-2147F2.1* | 2 | 41 | 3 | | | | | |
| chr14 | *RP11-813I20.2* | 2 | 30 | 3 | | | | | |
| chr11 | *TRAPPC4* | 2 | 41 | 3 | TRAPPC4 | trafficking protein particle | Cytoplasm | other | |
| chr1 | *RABIF* | 2 | 56 | 3 | RABIF | RAB interacting factor | Other | transporter | |
| chr16 | *MT1JP* | 2 | 100 | 3 | MT1JP | metallothionein 1J, pseudogene | Other | other | |
| chr17 | *GH2* | 2 | 75 | 3 | GH2 | growth hormone 2 | Extracellular Space | other | |

| | | | | | | | | |
|---|---|---|---|---|---|---|---|---|
| chr11 | *KAT5* | 2 | 91 | 3 | KAT5 | lysine acetyltransferase 5 | Nucleus | transcription regulator |
| chr9 | *ZBTB5* | 2 | 46 | 3 | ZBTB5 | zinc finger and BTB domain containing 5 | Nucleus | transcription regulator |
| chrX | *RP11-1M18.1* | 2 | 32 | 3 | | | | |
| chr19 | *GIPC3* | 2 | 73 | 3 | GIPC3 | GIPC PDZ domain containing family member 3 | Other | other |
| chr8 | *RP11-369E15.3* | 2 | 48 | 3 | | | | |
| chr19 | *ZNF480* | 2 | 36 | 3 | ZNF480 | zinc finger protein 480 | Nucleus | transcription regulator |
| chr16 | *RP11-1212A22.1* | 2 | 60 | 3 | | | | |
| chr20 | *BIRC7* | 2 | 166 | 3 | BIRC7 | baculoviral IAP repeat containing | Cytoplasm | enzyme |
| chr10 | *NPAP1P3* | 2 | 53 | 3 | NPAP1P3 | nuclear pore associated protein 1 pseudogene 3 | Other | other |
| chr19 | *FAM71E2* | 2 | 68 | 3 | FAM71E2 | family with sequence similarity 71 member E2 | Other | other |
| chr9 | *LINC00475* | 2 | 54 | 3 | LINC00475 | long intergenic non-protein coding RNA 475 | Other | other |
| chr8 | *KB-1562D12.1* | 2 | 32 | 3 | | | | |
| chr19 | *CTC-523E23.11* | 2 | 58 | 3 | | | | |
| chr16 | *LINC00304* | 2 | 81 | 3 | LINC00304 | long intergenic non-protein coding RNA 304 | Other | other |
| chr7 | *CICP24* | 2 | 55 | 3 | CICP24 | | Other | other |
| chr20 | *ADRM1* | 2 | 167 | 3 | ADRM1 | adhesion regulating molecule 1 | Plasma Membrane | other |
| chrX | *GAGE12H* | 2 | 64 | 3 | GE1 (includes oth | G antigen 1 | Other | other |
| chr11 | *AP000662.4* | 2 | 89 | 3 | | | | |
| chr1 | *MNDA* | 2 | 40 | 3 | MNDA | myeloid cell nuclear differentiation antigen | Nucleus | other |
| chr11 | *RP11-707G14.7* | 2 | 53 | 3 | | | | |
| chr6 | *FBXL4* | 2 | 42 | 3 | FBXL4 | F-box and leucine rich repeat protein 4 | Nucleus | other |
| chr11 | *RP11-736I10.2* | 1 | 36 | 3 | | | | |
| chr1 | *RP11-690C23.4* | 2 | 69 | 3 | | | | |
| chr12 | *TSPAN31* | 2 | 62 | 3 | TSPAN31 | tetraspanin 31 | Plasma Membrane | other |
| chr11 | *C11orf58* | 2 | 33 | 3 | C11orf58 | chromosome 11 open reading frame 58 | Other | other |
| chr8 | *RP11-775B15.2* | 2 | 41 | 3 | | | | |
| chr14 | *RP1-240K6.3* | 2 | 29 | 3 | | | | |
| chr7 | *COPS6* | 2 | 100 | 3 | COPS6 | COP9 signalosome subunit 6 | Nucleus | other |
| chr19 | *CTD-2588C8.8* | 2 | 54 | 3 | | | | |
| chr9 | *AL359091.2* | 2 | 64 | 3 | | | | |
| chr19 | *ZNF681* | 2 | 54 | 3 | ZNF681 | zinc finger protein 681 | Other | other |
| chr3 | *ACTRT3* | 2 | 78 | 3 | ACTRT3 | actin related protein T3 | Nucleus | other |
| chr4 | *USP17L5* | 2 | 66 | 3 | 17L18 (includes ot | ubiquitin specific peptidase 17 like family member 5 | Nucleus | peptidase |
| chr3 | *THRAP3P1* | 2 | 39 | 3 | THRAP3P1 | thyroid hormone receptor associated protein 3 pseudogene | Other | other |
| chr8 | *FAM90A14P* | 2 | 194 | 3 | | | | |
| chr17 | *COASY* | 2 | 76 | 3 | COASY | Coenzyme A synthase | Cytoplasm | kinase |
| chr11 | *RP11-554A11.6* | 2 | 86 | 3 | | | | |
| chr19 | *ZNF441* | 2 | 35 | 3 | ZNF441 | zinc finger protein 441 | Nucleus | transcription regulator |
| chr17 | *KRT223P* | 2 | 41 | 3 | KRT223P | keratin 223 pseudogene | Other | other |
| chr19 | *CTD-2554C21.2* | 2 | 33 | 3 | | | | |
| chr11 | *ANO1-AS2* | 2 | 73 | 3 | LINC02584 | long intergenic non-protein coding RNA 2584 | Other | other |
| chr2 | *SP110* | 1 | 27 | 3 | SP110 | SP110 nuclear body protein | Nucleus | transcription regulator |
| chr11 | *RP11-800A3.4* | 2 | 83 | 3 | | | | |
| chr18 | *RP11-243E13.2* | 1 | 31 | 3 | | | | |
| chr15 | *RP11-526I2.5* | 1 | 47 | 3 | | | | |
| chrX | *MID2* | 2 | 35 | 3 | MID2 | midline 2 | Cytoplasm | other |
| chr2 | *AC114814.4* | 2 | 38 | 3 | | | | |
| chr6 | *LMBRD1* | 2 | 36 | 3 | LMBRD1 | LMBR1 domain containing 1 | Cytoplasm | enzyme |
| chr2 | *IGFBP5* | 2 | 53 | 3 | IGFBP5 | insulin like growth factor binding protein 5 | Extracellular Space | other |
| chr1 | *SLC25A34* | 2 | 74 | 3 | SLC25A34 | solute carrier family 25 member | Cytoplasm | other |
| chr3 | *UPK1B* | 2 | 41 | 3 | UPK1B | uroplakin 1B | Plasma Membrane | other |
| chr5 | *RP11-114H7.2* | 2 | 70 | 3 | | | | |
| chr9 | *RP11-509J21.3* | 2 | 29 | 3 | | | | |
| chr22 | *CCDC117* | 2 | 40 | 3 | CCDC117 | coiled-coil domain containing 117 | Other | other |
| chr6 | *RP4-529N6.2* | 2 | 57 | 3 | | | | |
| chr19 | *AC016626.2* | 2 | 35 | 3 | | | | |
| chr2 | *AC011995.1* | 2 | 53 | 3 | | | | |
| chr17 | *TMEM106A* | 2 | 63 | 3 | TMEM106A | transmembrane protein 106A | Cytoplasm | other |
| chr1 | *C1orf110* | 2 | 76 | 3 | CCDC190 | coiled-coil domain containing 190 | Other | other |
| chr15 | *RP11-64K12.10* | 2 | 39 | 3 | | | | |
| chr2 | *TM4SF20* | 1 | 26 | 3 | TM4SF20 | transmembrane 4 L six family member 20 | Cytoplasm | other |
| chr15 | *C15orf48* | 2 | 45 | 3 | C15orf48 | chromosome 15 open reading frame 48 | Nucleus | other |
| chr9 | *ZNF883* | 1 | 32 | 3 | ZNF883 | zinc finger protein 883 | Other | other |
| chr19 | *ABHD8* | 2 | 74 | 3 | ABHD8 | abhydrolase domain containing 8 | Cytoplasm | enzyme |
| chr8 | *CTD-3064M3.1* | 2 | 70 | 3 | | | | |
| chr20 | *FAM217B* | 2 | 95 | 3 | FAM217B | family with sequence similarity 217 member B | Other | other |
| chr5 | *ABLIM3* | 2 | 39 | 3 | ABLIM3 | actin binding LIM protein family member 3 | Cytoplasm | other |
| chr11 | *OR10G7* | 2 | 39 | 3 | OR10G7 | olfactory receptor family 10 subfamily G member 7 | Plasma Membrane | i-protein coupled recepto |
| chr1 | *RP5-1061H20.5* | 2 | 63 | 3 | | | | |
| chr2 | *RP11-711M9.1* | 2 | 63 | 3 | | | | |
| chr7 | *RP11-340I6.8* | 2 | 29 | 3 | | | | |

| | | | | | | | | | |
|---|---|---|---|---|---|---|---|---|---|
| chr1 | *CFHR3* | 2 | 61 | 3 | CFHR3 | complement factor H related 3 | Extracellular Space | other | |
| chr20 | *POLR3F* | 2 | 29 | 3 | POLR3F | RNA polymerase III subunit F | Nucleus | enzyme | |
| chr12 | *RP11-955H22.1* | 2 | 42 | 3 | | | | | |
| chr1 | *RP11-214L19.1* | 2 | 72 | 3 | | | | | |
| chr5 | *KRT8P31* | 2 | 73 | 3 | KRT8P31 | keratin 8 pseudogene 31 | Other | other | |
| chr17 | *CTD-2535P7.1* | 2 | 32 | 3 | | | | | |
| chr9 | *RP11-363E7.3* | 2 | 70 | 3 | | | | | |
| chr10 | *RP11-20J15.3* | 2 | 74 | 3 | | | | | |
| chr19 | *CLEC4G* | 2 | 53 | 3 | CLEC4G | C-type lectin domain family 4 member G | Plasma Membrane | other | |
| chr15 | *RP11-8P11.3* | 2 | 50 | 3 | | | | | |
| chr1 | *RP5-902P8.12* | 2 | 67 | 3 | | | | | |
| chr11 | *RPS13* | 2 | 43 | 3 | RPS13 | ribosomal protein S13 | Cytoplasm | other | |
| chr5 | *CTD-2134P3.1* | 2 | 146 | 3 | | | | | |
| chr1 | *CDC20* | 2 | 53 | 3 | CDC20 | cell division cycle 20 | Nucleus | other | |
| chr10 | *RP11-773D16.1* | 2 | 31 | 3 | | | | | |
| chr12 | *LRRC10* | 2 | 59 | 3 | LRRC10 | leucine rich repeat containing 10 | Nucleus | other | |
| chr1 | *TUBB8P9* | 2 | 137 | 3 | TUBB8P9 | tubulin beta 8 class VIII pseudogene 9 | Other | other | |
| chr2 | *FLJ14816* | 2 | 80 | 3 | | | | | |
| chr6 | *HIST1H2BJ* | 2 | 38 | 3 | H2BC11 | H2B clustered histone 11 | Nucleus | other | |
| chr14 | *CCDC175* | 2 | 46 | 3 | CCDC175 | coiled-coil domain containing 175 | Other | other | |
| chr17 | *G6PC3* | 2 | 64 | 3 | G6PC3 | glucose-6-phosphatase catalytic subunit 3 | Cytoplasm | phosphatase | |
| chr12 | *MMP19* | 2 | 106 | 3 | MMP19 | matrix metallopeptidase 19 | Extracellular Space | peptidase | marimastat |
| chr6 | *NOX3* | 2 | 38 | 3 | NOX3 | NADPH oxidase 3 | Cytoplasm | enzyme | |
| chr5 | *TIGD6* | 1 | 28 | 3 | TIGD6 | tigger transposable element | Nucleus | other | |
| chr5 | *COX7C* | 2 | 52 | 3 | COX7C | cytochrome c oxidase subunit 7C | Cytoplasm | enzyme | |
| chr11 | *FAM111B* | 2 | 52 | 3 | FAM111B | family with sequence similarity 111 member B | Other | other | |
| chr19 | *DACT3* | 2 | 58 | 3 | DACT3 | dishevelled binding antagonist of beta catenin 3 | Cytoplasm | other | |
| chrX | *MCTS1* | 2 | 40 | 3 | MCTS1 | MCTS1 re-initiation and release | Cytoplasm | other | |
| chr22 | *PHF5A* | 2 | 33 | 3 | PHF5A | PHD finger protein 5A | Nucleus | transcription regulator | |
| chr4 | *DUX4L7* | 2 | 375 | 3 | X4L1 (includes oth | double homeobox 4 like 4 (pseudogene) | Other | transcription regulator | |
| chr4 | *RP3-368B9.2* | 2 | 107 | 3 | | | | | |
| chr22 | *LINC00634* | 2 | 58 | 3 | LINC00634 | long intergenic non-protein coding RNA 634 | Other | other | |
| chr1 | *CADM3* | 2 | 88 | 3 | CADM3 | cell adhesion molecule 3 | Plasma Membrane | other | |
| chr15 | *RP11-752G15.3* | 2 | 61 | 3 | | | | | |
| chr19 | *LILRA4* | 2 | 66 | 3 | LILRA4 | leukocyte immunoglobulin like receptor A4 | Plasma Membrane | other | |
| chr4 | *PACRGL* | 2 | 40 | 3 | PACRGL | parkin coregulated like | Other | other | |
| chr19 | *C19orf59* | 2 | 52 | 3 | MCEMP1 | mast cell expressed membrane protein 1 | Cytoplasm | other | |
| chr9 | *RP11-15J10.1* | 1 | 41 | 3 | | | | | |
| chr11 | *RCOR2* | 2 | 75 | 3 | RCOR2 | REST corepressor 2 | Nucleus | transcription regulator | |
| chr21 | *C21orf91* | 1 | 30 | 3 | C21orf91 | chromosome 21 open reading frame 91 | Other | other | |
| chr2 | *AC092570.2* | 2 | 74 | 3 | | | | | |
| chrX | *ARMCX6* | 2 | 36 | 3 | ARMCX6 | armadillo repeat containing X- | Other | other | |
| chr2 | *SCRN3* | 1 | 36 | 3 | SCRN3 | secernin 3 | Other | other | |
| chrX | *SSX7* | 2 | 51 | 3 | SSX7 | SSX family member 7 | Other | other | |
| chr2 | *AC113607.2* | 1 | 41 | 3 | | | | | |
| chr17 | *AMZ2P1* | 2 | 35 | 3 | AMZ2P1 | archaelysin family metallopeptidase 2 pseudogene 1 | Other | other | |
| chr20 | *SDCBP2-AS1* | 2 | 33 | 3 | SDCBP2-AS1 | SDCBP2 antisense RNA 1 | Other | other | |
| chr10 | *SPAG6* | 1 | 25 | 3 | SPAG6 | sperm associated antigen 6 | Cytoplasm | other | |
| chr1 | *RP11-275O4.3* | 2 | 63 | 3 | | | | | |
| chr19 | *ZNF763* | 2 | 50 | 3 | ZNF763 | zinc finger protein 763 | Cytoplasm | transcription regulator | |
| chr9 | *RP11-611D20.2* | 2 | 162 | 3 | | | | | |
| chr2 | *PTPRN* | 2 | 68 | 3 | PTPRN | protein tyrosine phosphatase receptor type N | Plasma Membrane | phosphatase | |
| chr17 | *SLC35G3* | 2 | 81 | 3 | SLC35G3 | solute carrier family 35 member | Other | enzyme | |
| chr7 | *GS1-124K5.3* | 2 | 49 | 3 | | | | | |
| chr8 | *ZNF395* | 2 | 53 | 3 | ZNF395 | zinc finger protein 395 | Cytoplasm | other | |
| chr3 | *GP9* | 2 | 77 | 3 | GP9 | glycoprotein IX platelet | Plasma Membrane | other | |
| chr7 | *DUS4L* | 1 | 28 | 3 | DUS4L | dihydrouridine synthase 4 like | Other | other | |
| chr19 | *UBXN6* | 2 | 46 | 3 | UBXN6 | UBX domain protein 6 | Cytoplasm | other | |
| chr6 | *FRS3* | 2 | 56 | 3 | FRS3 | fibroblast growth factor receptor substrate 3 | Plasma Membrane | other | |
| chr10 | *LINC00863* | 2 | 39 | 3 | LINC00863 | long intergenic non-protein coding RNA 863 | Other | other | |
| chr6 | *TNFAIP3* | 2 | 61 | 3 | TNFAIP3 | TNF alpha induced protein 3 | Nucleus | enzyme | |
| chr7 | *HTR5A* | 2 | 44 | 3 | HTR5A | 5-hydroxytryptamine receptor 5A | Plasma Membrane | i-protein coupled recepto | asenapine |
| chr15 | *RP11-26F2.1* | 1 | 37 | 3 | | | | | |
| chr5 | *CTD-2194D22.1* | 2 | 229 | 3 | | | | | |
| chr1 | *IFI44L* | 2 | 27 | 3 | IFI44L | interferon induced protein 44 like | Nucleus | other | |
| chr15 | *RP11-761I4.3* | 2 | 86 | 3 | | | | | |
| chr16 | *TCEB2* | 2 | 58 | 3 | ELOB | elongin B | Nucleus | transcription regulator | |
| chr2 | *LYPD6* | 2 | 25 | 3 | LYPD6 | LY6/PLAUR domain containing 6 | Extracellular Space | other | |
| chr17 | *CDRT15* | 1 | 55 | 3 | CDRT15 | CMT1A duplicated region transcript 15 | Other | other | |
| chr19 | *CTB-147N14.4* | 1 | 40 | 3 | | | | | |
| chr18 | *RP11-49I11.1* | 2 | 49 | 3 | | | | | |
| chr3 | *RP11-438J1.1* | 2 | 40 | 3 | | | | | |
| chr20 | *BAK1P1* | 2 | 72 | 3 | BAK1P1 | BCL2 antagonist/killer 1 pseudogene 1 | Other | other | |

| | | | | | | | | |
|---|---|---|---|---|---|---|---|---|
| chr8 | *RP11-1049H7.2* | 2 | 69 | 3 | | | | |
| chr3 | *RP11-71E19.2* | 2 | 80 | 3 | | | | |
| chr12 | *PZP* | 1 | 32 | 3 | PZP | PZP alpha-2-macroglobulin like | Extracellular Space | other |
| chr17 | *PHF23* | 2 | 63 | 3 | PHF23 | PHD finger protein 23 | Nucleus | other |
| chr6 | *SLC35B3* | 2 | 52 | 3 | SLC35B3 | solute carrier family 35 member | Cytoplasm | transporter |
| chr8 | *SLC35G5* | 2 | 60 | 3 | SLC35G5 | solute carrier family 35 member | Other | other |
| chr19 | *hsa-mir-125a* | 2 | 69 | 3 | | | | |
| chr20 | *VSX1* | 2 | 41 | 3 | VSX1 | visual system homeobox 1 | Nucleus | transcription regulator |
| chrX | *RP6-29D12.2* | 1 | 66 | 3 | | | | |
| chr2 | *AC096669.3* | 2 | 65 | 3 | | | | |
| chrX | *SPANXB1* | 2 | 103 | 3 | NXB1 (includes ot | SPANX family member C | Nucleus | other |
| chrX | *TAF7L* | 2 | 54 | 3 | TAF7L | TATA-box binding protein associated factor 7 like | Nucleus | transcription regulator |
| chr3 | *ZNF619* | 2 | 50 | 3 | ZNF619 | zinc finger protein 619 | Other | other |
| chr7 | *RP4-669B10.3* | 2 | 28 | 3 | | | | |
| chr11 | *RP5-1173A5.1* | 1 | 62 | 3 | | | | |
| chr9 | *DDX11L5* | 1 | 87 | 3 | DDX11L5 | DEAD/H-box helicase 11 like 5 | Other | other |
| chr14 | *RP11-219E7.4* | 2 | 36 | 3 | | | | |
| chr15 | *HERC2P7* | 2 | 86 | 3 | HERC2P7 | hect domain and RLD 2 pseudogene 7 | Other | other |
| chr15 | *HIGD2B* | 2 | 43 | 3 | HIGD2B | HIG1 hypoxia inducible domain family member 2B | Other | other |
| chr4 | *GRXCR1* | 2 | 27 | 3 | GRXCR1 | glutaredoxin and cysteine rich domain containing 1 | Extracellular Space | other |
| chr7 | *NAT16* | 2 | 144 | 3 | NAT16 | N-acetyltransferase 16 (putative) | Other | other |
| chr2 | *AC020601.1* | 2 | 31 | 3 | | | | |
| chr19 | *KLK11* | 2 | 74 | 3 | KLK11 | kallikrein related peptidase 11 | Extracellular Space | peptidase |
| chr7 | *CRYGN* | 2 | 82 | 3 | CRYGN | crystallin gamma N | Other | other |
| chr2 | *INHBB* | 2 | 89 | 3 | INHBB | inhibin subunit beta B | Extracellular Space | growth factor |
| chr12 | *RP5-1096D14.3* | 2 | 36 | 3 | | | | |
| chr15 | *RP11-483E23.3* | 1 | 44 | 3 | | | | |
| chrX | *MORC4* | 2 | 33 | 3 | MORC4 | MORC family CW-type zinc finger | Nucleus | other |
| chr14 | *RP11-857B24.1* | 1 | 28 | 3 | | | | |
| chrX | *ARX* | 2 | 69 | 3 | ARX | aristaless related homeobox | Nucleus | transcription regulator |
| chr20 | *KRT18P4* | 2 | 93 | 3 | KRT18P4 | keratin 18 pseudogene 4 | Other | other |
| chr9 | *RP11-555H7.2* | 2 | 75 | 3 | | | | |
| chr14 | *KLHL33* | 2 | 55 | 3 | KLHL33 | kelch like family member 33 | Other | other |
| chr11 | *CTD-2007L18.5* | 2 | 48 | 3 | | | | |
| chr17 | *RP11-1055B8.4* | 2 | 67 | 3 | | | | |
| chr4 | *NPFFR2* | 2 | 44 | 3 | NPFFR2 | neuropeptide FF receptor 2 | Plasma Membrane | G-protein coupled recepto |
| chr15 | *ULK4P2* | 2 | 40 | 3 | ULK4P2 | ULK4 pseudogene 2 | Other | other |
| chr19 | *PPM1N* | 2 | 77 | 3 | PPM1N | protein phosphatase, Mg2+/Mn2+ dependent 1N (putative) | Extracellular Space | other |
| chr1 | *RP11-542C10.1* | 1 | 32 | 3 | | | | |
| chr11 | *RP11-707M1.7* | 2 | 119 | 3 | | | | |
| chr22 | *C22orf24* | 2 | 38 | 3 | C22orf24 | chromosome 22 open reading frame 24 | Other | other |
| chr15 | *RP11-543G18.1* | 2 | 41 | 3 | | | | |
| chr2 | *AC140481.8* | 1 | 35 | 3 | | | | |
| chr12 | *METTL7A* | 2 | 70 | 3 | METTL7A | methyltransferase like 7A | Cytoplasm | other |
| chr13 | *LINC01054* | 2 | 47 | 3 | LINC01054 | long intergenic non-protein coding RNA 1054 | Other | other |
| chr2 | *AC005037.1* | 1 | 44 | 3 | | | | |
| chr5 | *SCGB3A2* | 2 | 51 | 3 | SCGB3A2 | secretoglobin family 3A member | Extracellular Space | other |
| chrX | *RP11-431J24.2* | 2 | 38 | 3 | | | | |
| chr11 | *FOXRED1* | 2 | 49 | 3 | FOXRED1 | FAD dependent oxidoreductase domain containing 1 | Cytoplasm | other |
| chr11 | *MIR210HG* | 2 | 77 | 3 | MIR210HG | MIR210 host gene | Other | other |
| chr4 | *MRFAP1* | 2 | 49 | 3 | MRFAP1 | Morf4 family associated protein 1 | Nucleus | other |
| chr15 | *LINC00933* | 2 | 52 | 3 | LINC00933 | long intergenic non-protein coding RNA 933 | Other | other |
| chr17 | *ACADVL* | 2 | 82 | 3 | ACADVL | acyl-CoA dehydrogenase very long chain | Cytoplasm | enzyme |
| chr17 | *NAGS* | 2 | 60 | 3 | NAGS | N-acetylglutamate synthase | Cytoplasm | enzyme |
| chr15 | *RP11-126F18.2* | 1 | 47 | 3 | | | | |
| chr17 | *PPP1R27* | 2 | 67 | 3 | PPP1R27 | protein phosphatase 1 regulatory subunit 27 | Nucleus | transcription regulator |
| chr13 | *PRR20C* | 2 | 240 | 3 | R20A (includes oth | proline rich 20A | Other | other |
| chr15 | *RHOV* | 2 | 76 | 3 | RHOV | ras homolog family member V | Plasma Membrane | enzyme |
| chr6 | *KCNK16* | 2 | 80 | 3 | KCNK16 | potassium two pore domain channel subfamily K member 16 | Plasma Membrane | ion channel |
| chr1 | *TOMM40L* | 2 | 74 | 3 | TOMM40L | translocase of outer mitochondrial membrane 40 like | Cytoplasm | other |
| chr19 | *ZNF649* | 2 | 50 | 3 | ZNF649 | zinc finger protein 649 | Nucleus | transcription regulator |
| chr17 | *RP11-527L4.5* | 2 | 52 | 3 | | | | |
| chrX | *XAGE1A* | 2 | 64 | 3 | XAGE1A/XAGE1E | X antigen family member 1B | Nucleus | other |
| chr7 | *IGFBP3* | 2 | 65 | 3 | IGFBP3 | insulin like growth factor binding protein 3 | Extracellular Space | other |
| chr1 | *RP11-147G16.1* | 2 | 45 | 3 | | | | |
| chr17 | *ARL16* | 2 | 58 | 3 | ARL16 | ADP ribosylation factor like GTPase 16 | Cytoplasm | other |
| chr2 | *DDX11L2* | 2 | 85 | 3 | DDX11L2 | DEAD/H-box helicase 11 like 2 | Other | other |
| chr22 | *RIMBP3* | 1 | 68 | 3 | IBP3 (includes oth | RIMS binding protein 3 | Nucleus | other |
| chr3 | *CTD-2013N24.2* | 1 | 25 | 3 | | | | |
| chr1 | *RP11-480I12.4* | 1 | 46 | 3 | | | | |
| chr17 | *RP11-219G17.6* | 1 | 54 | 3 | | | | |
| chr5 | *CTC-448D22.1* | 2 | 29 | 3 | | | | |
| chr10 | *FAM25D* | 2 | 66 | 3 | FAM25E | family with sequence similarity 25 member E | Other | other |

| | | | | | | | | | |
|---|---|---|---|---|---|---|---|---|---|
| chr8 | *TSTA3* | 2 | 155 | 3 | TSTA3 | tissue specific transplantation antigen P35B | Plasma Membrane | enzyme | |
| chr19 | *ZNF562* | 2 | 37 | 3 | ZNF562 | zinc finger protein 562 | Nucleus | transcription regulator | |
| chr9 | *IL6RP1* | 2 | 45 | 3 | IL6RP1 | interleukin 6 receptor | Other | other | |
| chr19 | *MBD3L2* | 2 | 93 | 3 | D3L2 (includes oth | methyl-CpG binding domain protein 3 like 2 | Other | other | |
| chr12 | *RP11-70F11.11* | 2 | 94 | 3 | | | | | |
| chr2 | *LINC00607* | 2 | 34 | 3 | LINC00607 | long intergenic non-protein coding RNA 607 | Other | other | |
| chr1 | *CPT2* | 2 | 53 | 3 | CPT2 | carnitine palmitoyltransferase 2 | Cytoplasm | enzyme | perhexiline |
| chr16 | *TSR3* | 2 | 72 | 3 | TSR3 | TSR3 ribosome maturation factor | Cytoplasm | enzyme | |
| chr6 | *FAM162B* | 1 | 25 | 3 | FAM162B | family with sequence similarity 162 member B | Other | other | |
| chr16 | *TPPP3* | 2 | 101 | 3 | TPPP3 | tubulin polymerization promoting protein family member 3 | Cytoplasm | other | |
| chr1 | *IL24* | 2 | 72 | 3 | IL24 | interleukin 24 | Extracellular Space | cytokine | |
| chr10 | *RP11-245J24.1* | 2 | 39 | 3 | | | | | |
| chr1 | *MROH7-TTC4* | 2 | 52 | 3 | MROH7-TTC4 | MROH7-TTC4 readthrough (NMD candidate) | Other | other | |
| chr2 | *AC093590.1* | 2 | 46 | 3 | | | | | |
| chr17 | *ZSWIM5P1* | 1 | 54 | 3 | ZSWIM5P1 | zinc finger SWIM-type containing 5 pseudogene 1 | Other | other | |
| chr1 | *NPPB* | 2 | 56 | 3 | NPPB | natriuretic peptide B | Extracellular Space | other | |
| chr5 | *CTD-2314G24.2* | 2 | 49 | 3 | | | | | |
| chr6 | *RP11-528A10.1* | 2 | 55 | 3 | | | | | |
| chr1 | *IFI6* | 2 | 61 | 3 | IFI6 | interferon alpha inducible protein | Cytoplasm | other | |
| chr6 | *CRIP3* | 2 | 44 | 3 | CRIP3 | cysteine rich protein 3 | Cytoplasm | other | |
| chr7 | *GS1-259H13.10* | 2 | 91 | 3 | | | | | |
| chr2 | *AC013727.1* | 2 | 42 | 3 | | | | | |
| chr19 | *AC004156.3* | 2 | 79 | 3 | | | | | |
| chr1 | *RP11-261C10.7* | 2 | 66 | 3 | | | | | |
| chr1 | *RP11-354K1.2* | 2 | 54 | 3 | | | | | |
| chr9 | *NDUFB6* | 1 | 39 | 3 | NDUFB6 | NADH:ubiquinone oxidoreductase subunit B6 | Cytoplasm | enzyme | |
| chr14 | *RP11-404P21.8* | 2 | 49 | 3 | | | | | |
| chr20 | *GPCPD1* | 2 | 29 | 3 | GPCPD1 | glycerophosphocholine phosphodiesterase 1 | Cytoplasm | enzyme | |
| chr3 | *IMPDH2* | 2 | 59 | 3 | IMPDH2 | inosine monophosphate dehydrogenase 2 | Cytoplasm | enzyme | mycophenolate mofetil, VX-944, pegintron/ribavirin, mycophenolate mofetil/prednisone, methylprednisolone/ mycophenolate mofetil, interferon alfacon-1/ribavirin, 6-mercaptopurine/pred nisone/thioguanine, cytarabine/daunorubi cin/thioguanine, cytarabine/thioguanin e, IFNA2B/ribavirin, ribavirin, mycophenolic acid |
| chr1 | *MAEL* | 2 | 30 | 3 | MAEL | maelstrom spermatogenic transposon silencer | Nucleus | transcription regulator | |
| chr1 | *S1PR1* | 1 | 61 | 3 | S1PR1 | sphingosine-1-phosphate | Plasma Membrane | G-protein coupled recepto | fingolimod phosphate |
| chr11 | *TMEM258* | 2 | 67 | 3 | TMEM258 | transmembrane protein 258 | Cytoplasm | other | |
| chrX | *SUV39H1* | 2 | 85 | 3 | SUV39H1 | suppressor of variegation 3-9 homolog 1 | Nucleus | enzyme | |
| chr11 | *RP11-358N4.6* | 2 | 50 | 3 | | | | | |
| chr2 | *IL37* | 2 | 71 | 3 | IL37 | interleukin 37 | Extracellular Space | cytokine | |
| chr5 | *RXFP3* | 2 | 195 | 3 | RXFP3 | relaxin family peptide receptor 3 | Plasma Membrane | G-protein coupled recepto | |
| chr15 | *AC123768.4* | 2 | 28 | 3 | | | | | |
| chr12 | *RP1-228P16.3* | 2 | 170 | 3 | | | | | |
| chr10 | *HPS6* | 2 | 65 | 3 | HPS6 | HPS6 biogenesis of lysosomal organelles complex 2 subunit 3 | Cytoplasm | other | |
| chr3 | *MYD88* | 2 | 60 | 3 | MYD88 | MYD88 innate immune signal transduction adaptor | Plasma Membrane | other | IMO-8400 |
| chr11 | *TIMM10B* | 2 | 49 | 3 | TIMM10B | translocase of inner mitochondrial membrane 10B | Cytoplasm | transporter | |
| chr14 | *RP6-114E22.1* | 2 | 38 | 3 | | | | | |
| chrX | *CXorf31* | 2 | 33 | 3 | LINC01545 | long intergenic non-protein coding RNA 1545 | Other | other | |
| chr7 | *AC008265.2* | 1 | 71 | 3 | | | | | |
| chrX | *XRCC6P5* | 2 | 55 | 3 | XRCC6P5 | X-ray repair cross complementing 6 pseudogene 5 | Other | other | |
| chr12 | *RP11-641A6.8* | 2 | 59 | 3 | | | | | |
| chrX | *MED12* | 2 | 46 | 3 | MED12 | mediator complex subunit 12 | Nucleus | transcription regulator | |
| chr12 | *FAM222A-AS1* | 2 | 64 | 3 | FAM222A-AS1 | FAM222A antisense RNA 1 | Other | other | |
| chr12 | *NEUROD4* | 2 | 58 | 3 | NEUROD4 | neuronal differentiation 4 | Nucleus | transcription regulator | |
| chr12 | *OVCH1-AS1* | 2 | 58 | 3 | OVCH1-AS1 | OVCH1 antisense RNA 1 | Other | other | |
| chr19 | *AC068499.10* | 2 | 49 | 3 | | | | | |
| chr5 | *AC011431.1* | 2 | 53 | 3 | | | | | |
| chr16 | *MLYCD* | 2 | 54 | 3 | MLYCD | malonyl-CoA decarboxylase | Cytoplasm | enzyme | |

| | | | | | | | | | |
|---|---|---|---|---|---|---|---|---|---|
| chr20 | *HRH3* | 2 | 114 | 3 | HRH3 | histamine receptor H3 | Plasma Membrane | G-protein coupled recepto | buclizine, pitolisant, betahistine, triprolidine, AZD5213, ABT-288, tesmilifene, |
| chr5 | *FAM159B* | 1 | 38 | 3 | SHISAL2B | shisa like 2B | Other | other | |
| chr1 | *LAMTOR2* | 2 | 66 | 3 | LAMTOR2 | late endosomal/lysosomal adaptor, MAPK and MTOR | Cytoplasm | other | |
| chr9 | *NCBP1* | 1 | 25 | 3 | NCBP1 | nuclear cap binding protein | Nucleus | other | |
| chr17 | *RP11-242D8.1* | 1 | 41 | 3 | | | | | |
| chr10 | *FAS* | 1 | 27 | 3 | FAS | Fas cell surface death receptor | Plasma Membrane | transmembrane recepto | APO010 |
| chr12 | *AC079630.4* | 1 | 35 | 3 | | | | | |
| chr3 | *RPL39P5* | 2 | 74 | 3 | RPL39P5 | ribosomal protein L39 | Other | other | |
| chr19 | *ASPDH* | 2 | 62 | 3 | ASPDH | aspartate dehydrogenase domain containing | Other | other | |
| chr19 | *KHSRP* | 2 | 72 | 3 | KHSRP | KH-type splicing regulatory | Nucleus | enzyme | |
| chr19 | *IFNL1* | 2 | 41 | 3 | IFNL1 | interferon lambda 1 | Extracellular Space | cytokine | |
| chr11 | *RP11-21L23.2* | 2 | 56 | 3 | | | | | |
| chr17 | *RP11-64J19.1* | 1 | 49 | 3 | | | | | |
| chr6 | *ZSCAN31* | 2 | 49 | 3 | ZSCAN31 | zinc finger and SCAN domain containing 31 | Nucleus | transcription regulator | |
| chr20 | *LINC00652* | 1 | 24 | 3 | LINC00652 | long intergenic non-protein coding RNA 652 | Other | other | |
| chr14 | *RP11-96D24.1* | 1 | 24 | 3 | | | | | |
| chr7 | *AC006372.1* | 2 | 50 | 3 | | | | | |
| chr15 | *LEO1* | 1 | 30 | 3 | LEO1 | LEO1 homolog, Paf1/RNA polymerase II complex | Nucleus | other | |
| chr22 | *CYP2D7P1* | 2 | 74 | 3 | CYP2D7 | cytochrome P450 family 2 subfamily D member 7 | Other | other | |
| chr19 | *CALR* | 2 | 49 | 3 | CALR | calreticulin | Cytoplasm | transcription regulator | |
| chr3 | *TIPARP* | 2 | 43 | 3 | TIPARP | TCDD inducible poly(ADP-ribose) polymerase | Nucleus | enzyme | RBN-2397 |
| chr10 | *ZNF239* | 2 | 31 | 3 | ZNF239 | zinc finger protein 239 | Nucleus | transcription regulator | |
| chrX | *VCX2* | 1 | 42 | 3 | CX2 (includes othe | variable charge Y-linked | Other | other | |
| chr4 | *RP11-404I7.2* | 1 | 37 | 3 | | | | | |
| chr10 | *GDF2* | 2 | 155 | 3 | GDF2 | growth differentiation factor 2 | Extracellular Space | growth factor | |
| chr11 | *CASP5* | 2 | 53 | 3 | CASP5 | caspase 5 | Cytoplasm | peptidase | |
| chr9 | *RP11-262H14.1* | 2 | 499 | 3 | | | | | |
| chr1 | *PPM1J* | 2 | 85 | 3 | PPM1J | protein phosphatase, Mg2+/Mn2+ dependent 1J | Other | phosphatase | |
| chr19 | *ETV2* | 2 | 78 | 3 | ETV2 | ETS variant transcription factor 2 | Nucleus | transcription regulator | |
| chr1 | *RPL11* | 1 | 51 | 3 | RPL11 | ribosomal protein L11 | Cytoplasm | other | |
| chr5 | *REEP5* | 2 | 31 | 3 | REEP5 | receptor accessory protein 5 | Extracellular Space | transporter | |
| chr8 | *RP11-664D7.4* | 1 | 24 | 3 | | | | | |
| chr2 | *C2orf81* | 2 | 54 | 3 | C2orf81 | chromosome 2 open reading frame 81 | Other | other | |
| chr22 | *IGLL5* | 2 | 53 | 3 | IGLL1/IGLL5 | immunoglobulin lambda like polypeptide 1 | Plasma Membrane | other | |
| chr3 | *RP11-439C8.1* | 1 | 53 | 3 | | | | | |
| chr17 | *EVPLL* | 2 | 99 | 3 | EVPLL | envoplakin like | Other | other | |
| chr16 | *NRN1L* | 2 | 50 | 3 | NRN1L | neuritin 1 like | Nucleus | other | |
| chr13 | *ALOX5AP* | 2 | 29 | 3 | ALOX5AP | arachidonate 5-lipoxygenase activating protein | Plasma Membrane | other | |
| chr3 | *CCDC71* | 2 | 73 | 3 | CCDC71 | coiled-coil domain containing 71 | Nucleus | other | |
| chr9 | *L09NC01-251B2.* | 2 | 70 | 3 | | | | | |
| chr1 | *RP5-1065J22.8* | 1 | 44 | 3 | | | | | |
| chr1 | *RP11-268J15.5* | 2 | 55 | 3 | | | | | |
| chr19 | *TSPAN16* | 2 | 32 | 3 | TSPAN16 | tetraspanin 16 | Other | other | |
| chr1 | *FAM58BP* | 2 | 77 | 3 | CCNQP1 | CCNQ pseudogene 1 | Other | other | |
| chr19 | *AC018755.1* | 2 | 42 | 3 | | | | | |
| chr7 | *NT5C3A* | 1 | 39 | 3 | NT5C3A | 5'-nucleotidase, cytosolic IIIA | Cytoplasm | phosphatase | |
| chr15 | *RP11-16B9.1* | 2 | 41 | 3 | | | | | |
| chr8 | *DEFA1* | 1 | 40 | 3 | FA1 (includes oth | defensin alpha 1 | Cytoplasm | other | |
| chr17 | *KRT33B* | 2 | 83 | 3 | KRT33B | keratin 33B | Extracellular Space | other | |
| chr17 | *KRTAP16-1* | 1 | 55 | 3 | KRTAP16-1 | keratin associated protein 16-1 | Cytoplasm | other | |
| chr12 | *RP1-228P16.4* | 2 | 65 | 3 | | | | | |
| chr1 | *C1orf216* | 2 | 59 | 3 | C1orf216 | chromosome 1 open reading frame 216 | Other | other | |
| chr17 | *RP11-893F2.5* | 2 | 73 | 3 | | | | | |
| chr6 | *GMNN* | 1 | 29 | 3 | GMNN | geminin DNA replication inhibitor | Nucleus | transcription regulator | |
| chrX | *SSX3* | 2 | 37 | 3 | SSX3 | SSX family member 3 | Other | other | |
| chr18 | *IMPACT* | 1 | 26 | 3 | IMPACT | impact RWD domain protein | Cytoplasm | other | |
| chr9 | *NR4A3* | 2 | 40 | 3 | NR4A3 | nuclear receptor subfamily 4 group A member 3 | Nucleus | d-dependent nuclear rec | |
| chr18 | *AKR1B1P6* | 1 | 49 | 3 | AKR1B1P6 | aldo-keto reductase family 1 member B1 pseudogene 6 | Other | other | |
| chr16 | *PRSS22* | 2 | 102 | 3 | PRSS22 | serine protease 22 | Extracellular Space | peptidase | |
| chr19 | *C19orf82* | 1 | 25 | 3 | ZNF561-AS1 | ZNF561 antisense RNA 1 (head to head) | Other | other | |
| chr15 | *RP11-4G2.1* | 2 | 58 | 3 | | | | | |
| chr15 | *AC105339.1* | 2 | 71 | 3 | | | | | |
| chr11 | *RP11-347H15.1* | 2 | 49 | 3 | | | | | |
| chr10 | *C10orf82* | 2 | 41 | 3 | C10orf82 | chromosome 10 open reading frame 82 | Other | other | |
| chr12 | *RP11-754N21.1* | 1 | 24 | 3 | | | | | |
| chr9 | *NDUFA5P3* | 1 | 28 | 3 | NDUFA5P3 | NADH:ubiquinone oxidoreductase subunit A5 | Other | other | |
| chr10 | *CLUHP5* | 2 | 148 | 3 | CLUHP5 | clustered mitochondria homolog pseudogene 5 | Other | other | |

| | | | | | | | | | |
|---|---|---|---|---|---|---|---|---|---|
| chr19 | CTD-2616J11.4 | 2 | 44 | 3 | | | | | |
| chr15 | CHAC1 | 2 | 68 | 3 | CHAC1 | ChaC glutathione specific gamma-glutamylcyclotransferase | Cytoplasm | enzyme | |
| chr8 | CYP7A1 | 1 | 24 | 3 | CYP7A1 | cytochrome P450 family 7 subfamily A member 1 | Cytoplasm | enzyme | |
| chr9 | RP11-229P13.22 | 2 | 69 | 3 | | | | | |
| chr5 | GUSBP8 | 1 | 44 | 3 | GUSBP8 | GUSB pseudogene 8 | Other | other | |
| chr16 | RP11-142G1.1 | 2 | 47 | 3 | | | | | |
| chr7 | RP5-912I13.1 | 2 | 61 | 3 | | | | | |
| chr14 | LINC00609 | 1 | 25 | 3 | LINC00609 | long intergenic non-protein coding RNA 609 | Other | other | |
| chr16 | RP11-715J22.2 | 2 | 99 | 3 | | | | | |
| chr8 | MRPL13 | 1 | 25 | 3 | MRPL13 | mitochondrial ribosomal protein | Cytoplasm | other | |
| chr19 | KLK10 | 2 | 47 | 3 | KLK10 | kallikrein related peptidase 10 | Extracellular Space | peptidase | |
| chr15 | RP11-475A13.1 | 1 | 24 | 3 | | | | | |
| chr9 | WDR31 | 2 | 47 | 3 | WDR31 | WD repeat domain 31 | Other | peptidase | |
| chr15 | DNM1P30 | 2 | 47 | 3 | DNM1P30 | dynamin 1 pseudogene 30 | Other | other | |
| chr1 | ODF2L | 2 | 58 | 3 | ODF2L | outer dense fiber of sperm tails 2 | Cytoplasm | other | |
| chr2 | ATG9A | 2 | 58 | 3 | ATG9A | autophagy related 9A | Cytoplasm | other | |
| chr8 | KIAA0196 | 2 | 27 | 3 | WASHC5 | WASH complex subunit 5 | Cytoplasm | other | |
| chrX | RP5-1139I1.2 | 1 | 25 | 3 | | | | | |
| chr1 | LMO4 | 2 | 42 | 3 | LMO4 | LIM domain only 4 | Nucleus | transcription regulator | |
| chr12 | RP11-272B17.1 | 2 | 45 | 3 | | | | | |
| chr7 | AC009945.4 | 1 | 70 | 3 | | | | | |
| chr1 | RP11-168O16.1 | 2 | 61 | 3 | | | | | |
| chr16 | HSD11B2 | 2 | 99 | 3 | HSD11B2 | hydroxysteroid 11-beta dehydrogenase 2 | Cytoplasm | enzyme | |
| chr1 | ISG15 | 2 | 106 | 3 | ISG15 | ISG15 ubiquitin like modifier | Extracellular Space | other | |
| chr1 | RAB7L1 | 2 | 59 | 3 | RAB29 | RAB29, member RAS oncogene family | Plasma Membrane | enzyme | |
| chr12 | MGST1 | 1 | 32 | 3 | MGST1 | microsomal glutathione S-transferase 1 | Cytoplasm | enzyme | |
| chr3 | EAF1-AS1 | 2 | 48 | 3 | EAF1-AS1 | EAF1 antisense RNA 1 | Other | other | |
| chr17 | RP11-666A8.8 | 2 | 44 | 3 | | | | | |
| chr11 | B3GNT6 | 2 | 73 | 3 | B3GNT6 | UDP-GlcNAc:betaGal beta-1,3-N-acetylglucosaminyltransferase 6 | Cytoplasm | enzyme | |
| chr19 | MIDN | 2 | 64 | 3 | MIDN | midnolin | Nucleus | other | |
| chr9 | ZNF510 | 2 | 37 | 3 | ZNF510 | zinc finger protein 510 | Nucleus | transcription regulator | |
| chr16 | ABCD1P3 | 2 | 156 | 3 | ABCD1P3 | ATP binding cassette subfamily D member 1 pseudogene 3 | Other | other | |
| chr1 | MUC1 | 2 | 137 | 3 | MUC1 | mucin 1, cell surface associated | Plasma Membrane | other | anti-TA-MUC1 antibody, GO-203-2C, pankomab, yttrium y 90 clivatuzumab tetraxetan, HuHMFG1, anti- |
| chr6 | EDN1 | 2 | 42 | 3 | EDN1 | endothelin 1 | Extracellular Space | cytokine | |
| chr11 | MRGPRX2 | 2 | 59 | 3 | MRGPRX2 | MAS related GPR family member | Plasma Membrane | i-protein coupled recepto | |
| chr2 | AC113618.1 | 1 | 38 | 3 | | | | | |
| chr4 | NUDT6 | 1 | 24 | 3 | NUDT6 | nudix hydrolase 6 | Extracellular Space | enzyme | |
| chr6 | RP3-468K18.5 | 1 | 30 | 3 | | | | | |
| chr17 | RP11-51L5.4 | 2 | 46 | 3 | | | | | |
| chr1 | CCDC163P | 2 | 47 | 3 | CCDC163 | coiled-coil domain containing 163 | Other | other | |
| chrX | CXorf49B | 2 | 59 | 3 | CXorf49/CXorf49E | chromosome X open reading frame 49 | Other | other | |
| chr17 | RP11-1109M24.1 | 2 | 1152 | 3 | | | | | |
| chr11 | AP002954.6 | 2 | 85 | 3 | | | | | |
| chr12 | SLCO1B1 | 2 | 43 | 3 | SLCO1B1 | solute carrier organic anion transporter family member 1B1 | Plasma Membrane | transporter | |
| chr2 | RP11-416L21.2 | 1 | 39 | 3 | | | | | |
| chr1 | CLCA1 | 2 | 34 | 3 | CLCA1 | chloride channel accessory 1 | Plasma Membrane | ion channel | crofelemer |
| chr19 | FOXA3 | 2 | 65 | 3 | FOXA3 | forkhead box A3 | Nucleus | transcription regulator | |
| chr4 | USP17L18 | 2 | 154 | 3 | 17L18 (includes ot | ubiquitin specific peptidase 17 like family member 5 | Nucleus | peptidase | |
| chr2 | AC019100.3 | 2 | 49 | 3 | | | | | |
| chr9 | GALT | 2 | 53 | 3 | GALT | galactose-1-phosphate uridylyltransferase | Cytoplasm | enzyme | |
| chr19 | KRT18P40 | 2 | 36 | 3 | KRT18P40 | keratin 18 pseudogene 40 | Other | other | |
| chr1 | CHIAP3 | 2 | 38 | 3 | CHIAP3 | chitinase, acidic pseudogene 3 | Other | other | |
| chr1 | RP11-38J22.3 | 2 | 56 | 3 | | | | | |
| chr9 | ZNF483 | 2 | 56 | 3 | ZNF483 | zinc finger protein 483 | Nucleus | transcription regulator | |
| chr7 | AC011290.4 | 2 | 32 | 3 | | | | | |
| chr11 | SCGB1D4 | 2 | 57 | 3 | SCGB1D4 | secretoglobin family 1D member | Extracellular Space | other | |
| chr1 | PKMP1 | 2 | 68 | 3 | PKMP1 | pyruvate kinase M1/2 | Other | other | |
| chr17 | CTB-175E5.7 | 2 | 43 | 3 | | | | | |
| chr18 | RP11-636O21.1 | 2 | 58 | 3 | | | | | |
| chr3 | JAGN1 | 2 | 53 | 3 | JAGN1 | jagunal homolog 1 | Cytoplasm | other | |
| chr3 | PCBP4 | 2 | 98 | 3 | PCBP4 | poly(rC) binding protein 4 | Nucleus | other | |
| chr20 | DZANK1 | 2 | 29 | 3 | DZANK1 | double zinc ribbon and ankyrin repeat domains 1 | Other | other | |
| chr7 | TRGV9 | 2 | 52 | 3 | TRGV9 | T cell receptor gamma variable 9 | Other | other | |
| chr17 | FAM27L | 2 | 59 | 3 | FAM27E5 | family with sequence similarity E5 | Other | other | |
| chr16 | FOXL1 | 2 | 55 | 3 | FOXL1 | forkhead box L1 | Nucleus | transcription regulator | |
| chr7 | RP11-458K10.1 | 2 | 32 | 3 | | | | | |
| chr1 | IPO9-AS1 | 1 | 56 | 3 | IPO9-AS1 | IPO9 antisense RNA 1 | Other | other | |
| chr14 | RP11-114N19.3 | 2 | 37 | 3 | | | | | |
| chr6 | RP1-90L14.1 | 1 | 24 | 3 | | | | | |
| chrX | KRT8P8 | 2 | 59 | 3 | KRT8P8 | keratin 8 pseudogene 8 | Other | other | |

| | | | | | | | | |
|---|---|---|---|---|---|---|---|---|
| chr22 | *PIK3IP1* | 2 | 37 | 3 | PIK3IP1 | phosphoinositide-3-kinase interacting protein 1 | Cytoplasm | other |
| chr11 | *VSIG2* | 2 | 48 | 3 | VSIG2 | V-set and immunoglobulin domain containing 2 | Plasma Membrane | other |
| chr19 | *SHD* | 2 | 47 | 3 | SHD | Src homology 2 domain containing transforming protein D | Cytoplasm | other |
| chr7 | *CLDN12* | 1 | 37 | 3 | CLDN12 | claudin 12 | Plasma Membrane | other |
| chr17 | *RP11-799D4.4* | 2 | 65 | 3 | | | | |
| chr12 | *EMP1* | 1 | 33 | 3 | EMP1 | epithelial membrane protein 1 | Plasma Membrane | other |
| chr1 | *RP5-855F14.1* | 2 | 91 | 3 | | | | |
| chr2 | *AC092687.4* | 1 | 36 | 3 | | | | |
| chr1 | *CD101* | 1 | 33 | 3 | CD101 | CD101 molecule | Plasma Membrane | other |
| chr6 | *C6orf1* | 2 | 68 | 3 | SMIM29 | small integral membrane protein | Other | other |
| chr6 | *PRSS16* | 2 | 43 | 3 | PRSS16 | serine protease 16 | Extracellular Space | peptidase |
| chr4 | *KIAA1430* | 2 | 27 | 3 | CFAP97 | cilia and flagella associated | Other | other |
| chr10 | *LINC00702* | 1 | 24 | 3 | LINC00702 | long intergenic non-protein coding RNA 702 | Other | other |
| chr3 | *GFM1* | 2 | 62 | 3 | GFM1 | G elongation factor mitochondrial | Cytoplasm | translation regulator |
| chr15 | *RP11-182J1.13* | 1 | 75 | 3 | | | | |
| chr13 | *CALM2P4* | 1 | 36 | 3 | CALM2P4 | calmodulin 2 pseudogene 4 | Other | other |
| chr18 | *RP11-268I9.3* | 2 | 38 | 3 | | | | |
| chr17 | *RP11-304F15.7* | 1 | 36 | 3 | | | | |
| chr16 | *RP11-44F14.1* | 2 | 50 | 3 | | | | |
| chr1 | *CD1A* | 1 | 45 | 3 | CD1A | CD1a molecule | Plasma Membrane | other |
| chr14 | *RP11-363J20.2* | 2 | 35 | 3 | | | | |
| chr1 | *TPRG1L* | 2 | 86 | 3 | TPRG1L | tumor protein p63 regulated 1 like | Cytoplasm | other |
| chr17 | *PCYT2* | 2 | 62 | 3 | PCYT2 | phosphate cytidylyltransferase 2, ethanolamine | Cytoplasm | enzyme |
| chr12 | *RP11-351O2.2* | 1 | 36 | 3 | | | | |
| chr1 | *RP11-206L10.10* | 2 | 31 | 3 | | | | |
| chr12 | *CTD-2021H9.3* | 2 | 52 | 3 | | | | |
| chr19 | *ZNF615* | 2 | 45 | 3 | ZNF615 | zinc finger protein 615 | Nucleus | transcription regulator |
| chrX | *GAGE1* | 2 | 111 | 3 | GE1 (includes oth | G antigen 1 | Other | other |
| chr5 | *RP11-455B3.1* | 2 | 34 | 3 | | | | |
| chr2 | *AC073257.1* | 2 | 52 | 3 | | | | |
| chr20 | *LINC00687* | 2 | 39 | 3 | LINC00687 | long intergenic non-protein coding RNA 687 | Other | other |
| chr1 | *RP11-527D7.1* | 2 | 53 | 3 | | | | |
| chr5 | *RP11-107N7.1* | 1 | 26 | 3 | | | | |
| chr19 | *AC104534.3* | 2 | 44 | 3 | | | | |
| chr11 | *RCE1* | 2 | 103 | 3 | RCE1 | Ras converting CAAX endopeptidase 1 | Cytoplasm | peptidase |
| chr3 | *CAV3* | 2 | 67 | 3 | CAV3 | caveolin 3 | Plasma Membrane | enzyme |
| chr13 | *RFC3* | 2 | 27 | 3 | RFC3 | replication factor C subunit 3 | Nucleus | enzyme |
| chr6 | *ZSCAN23* | 2 | 45 | 3 | ZSCAN23 | zinc finger and SCAN domain containing 23 | Cytoplasm | transcription regulator |
| chr17 | *SNORD3D* | 1 | 65 | 3 | SNORD3D | small nucleolar RNA, C/D box 3D | Other | other |
| chr12 | *PHC1P1* | 1 | 88 | 3 | PHC1P1 | polyhomeotic homolog 1 pseudogene 1 | Other | other |
| chr12 | *RP11-70F11.7* | 2 | 82 | 3 | | | | |
| chr16 | *RP11-266L9.5* | 2 | 43 | 3 | | | | |
| chr2 | *AC093162.5* | 1 | 52 | 3 | | | | |
| chrX | *TMEM185AP1* | 2 | 55 | 3 | TMEM185AP1 | transmembrane protein 185A pseudogene 1 | Other | other |
| chr7 | *SHH* | 2 | 58 | 3 | SHH | sonic hedgehog signaling | Extracellular Space | peptidase |
| chr19 | *CTC-360G5.1* | 2 | 66 | 3 | | | | |
| chr8 | *REXO1L9P* | 2 | 172 | 3 | REXO1L9P | REXO1 like 9, pseudogene | Other | other |
| chr11 | *RP11-867G23.13* | 2 | 92 | 3 | | | | |
| chr12 | *RP11-443N24.4* | 2 | 111 | 3 | | | | |
| chr11 | *KDELC2* | 2 | 40 | 3 | POGLUT3 | protein O-glucosyltransferase 3 | Other | other |
| chr7 | *CICP11* | 2 | 83 | 3 | CICP11 | capicua transcriptional repressor pseudogene 11 | Other | other |
| chr16 | *PKD1P1* | 2 | 96 | 3 | PKD1P1 | polycystin 1, transient receptor potential channel interacting pseudogene 1 | Other | other |
| chr5 | *ZMAT2* | 1 | 31 | 3 | ZMAT2 | zinc finger matrin-type 2 | Nucleus | other |
| chr10 | *PRF1* | 2 | 138 | 3 | PRF1 | perforin 1 | Cytoplasm | transporter |
| chr1 | *SNAPIN* | 2 | 54 | 3 | SNAPIN | SNAP associated protein | Plasma Membrane | other |
| chr3 | *RBMS3-AS3* | 2 | 31 | 3 | RBMS3-AS3 | RBMS3 antisense RNA 3 | Other | other |
| chr2 | *LINC00570* | 1 | 34 | 3 | LINC00570 | long intergenic non-protein coding RNA 570 | Other | other |
| chr7 | *AC005532.5* | 2 | 33 | 3 | | | | |
| chr4 | *NAA11* | 2 | 40 | 3 | NAA11 | N(alpha)-acetyltransferase 11, NatA catalytic subunit | Cytoplasm | enzyme |
| chr12 | *DYRK2* | 2 | 48 | 3 | DYRK2 | dual specificity tyrosine phosphorylation regulated kinase | Cytoplasm | kinase |
| chr19 | *AC092566.1* | 1 | 57 | 3 | | | | |
| chr19 | *PLIN5* | 2 | 60 | 3 | PLIN5 | perilipin 5 | Plasma Membrane | other |
| chr8 | *MROH6* | 2 | 73 | 3 | MROH6 | maestro heat like repeat family member 6 | Other | other |
| chr2 | *SNX17* | 2 | 47 | 3 | SNX17 | sorting nexin 17 | Cytoplasm | transporter |
| chr12 | *SPSB2* | 2 | 40 | 3 | SPSB2 | splA/ryanodine receptor domain and SOCS box containing 2 | Cytoplasm | other |
| chr11 | *ROM1* | 2 | 104 | 3 | ROM1 | retinal outer segment membrane protein 1 | Plasma Membrane | other |
| chr6 | *RP1-28O17.1* | 2 | 33 | 3 | | | | |
| chr12 | *RP11-268P4.4* | 1 | 34 | 3 | | | | |
| chr20 | *RP4-614C15.3* | 2 | 71 | 3 | | | | |

| | | | | | | | | | |
|---|---|---|---|---|---|---|---|---|---|
| chr2 | *CRIPT* | 1 | 41 | 3 | CRIPT | CXXC repeat containing interactor of PDZ3 domain | Cytoplasm | other | |
| chr16 | *RHOT2* | 2 | 74 | 3 | RHOT2 | ras homolog family member T2 | Cytoplasm | enzyme | |
| chr1 | *RN7SKP91* | 1 | 46 | 3 | RN7SKP91 | RN7SK pseudogene 91 | Other | other | |
| chr9 | *RP11-143M1.2* | 2 | 32 | 3 | | | | | |
| chr4 | *RP11-333E13.4* | 2 | 46 | 3 | | | | | |
| chr16 | *SYNGR3* | 2 | 106 | 3 | SYNGR3 | synaptogyrin 3 | Plasma Membrane | other | |
| chr12 | *FAM86FP* | 2 | 88 | 3 | FAM86FP | family with sequence similarity 86, member A pseudogene | Other | other | |
| chr1 | *PRPF38B* | 2 | 48 | 3 | PRPF38B | pre-mRNA processing factor 38B | Other | other | |
| chr10 | *CALHM2* | 2 | 92 | 3 | CALHM2 | calcium homeostasis modulator family member 2 | Other | other | |
| chr11 | *TRIM49D1* | 2 | 47 | 3 | RIM49D1/TRIM49I | tripartite motif containing 49D1 | Other | other | |
| chr17 | *PSMC3IP* | 2 | 65 | 3 | PSMC3IP | PSMC3 interacting protein | Nucleus | transcription regulator | |
| chr19 | *ZIM3* | 2 | 37 | 3 | ZIM3 | zinc finger imprinted 3 | Nucleus | transcription regulator | |
| chr20 | *RP11-199O14.1* | 2 | 31 | 3 | | | | | |
| chr6 | *RP11-686D16.1* | 2 | 27 | 3 | | | | | |
| chr5 | *CTC-338M12.9* | 2 | 75 | 3 | | | | | |
| chr12 | *RP5-1057I20.2* | 2 | 96 | 3 | | | | | |
| chr7 | *WI2-2373I1.2* | 2 | 80 | 3 | WI2-2373I1.2 | forkhead box L1-like | Other | other | |
| chr9 | *ARHGEF39* | 2 | 47 | 3 | ARHGEF39 | Rho guanine nucleotide exchange factor 39 | Plasma Membrane | other | |
| chr8 | *CYC1* | 2 | 68 | 3 | CYC1 | cytochrome c1 | Cytoplasm | enzyme | |
| chr7 | *RP11-325F22.5* | 1 | 25 | 3 | | | | | |
| chr7 | *HOTTIP* | 2 | 66 | 3 | HOTTIP | HOXA distal transcript antisense RNA | Other | other | |
| chr7 | *HSPB1* | 2 | 118 | 3 | HSPB1 | heat shock protein family B (small) member 1 | Cytoplasm | other | |
| chr6 | *RP3-420J14.1* | 1 | 28 | 3 | | | | | |
| chr16 | *PTX4* | 2 | 53 | 3 | PTX4 | pentraxin 4 | Other | other | |
| chr16 | *VKORC1* | 2 | 57 | 3 | VKORC1 | vitamin K epoxide reductase complex subunit 1 | Cytoplasm | enzyme | enoxaparin/warfarin, warfarin, acenocoumarol, vitamin K3, phenprocoumon, dicumarol, |
| chr20 | *RP5-1031J8.1* | 2 | 43 | 3 | | | | | |
| chr1 | *AL591704.9* | 2 | 62 | 3 | | | | | |
| chr11 | *APLNR* | 2 | 76 | 3 | APLNR | apelin receptor | Plasma Membrane | 3-protein coupled recepto | |
| chr19 | *VN1R2* | 1 | 26 | 3 | VN1R2 | vomeronasal 1 receptor 2 | Plasma Membrane | 3-protein coupled recepto | |
| chr9 | *RP11-29B9.1* | 1 | 28 | 3 | | | | | |
| chrX | *GAGE2E* | 2 | 161 | 3 | GE1 (includes oth | G antigen 1 | Other | other | |
| chr1 | *C1orf35* | 2 | 75 | 3 | C1orf35 | chromosome 1 open reading frame 35 | Other | other | |
| chr16 | *RP11-104C4.2* | 1 | 43 | 3 | | | | | |
| chr13 | *SPRY2* | 2 | 70 | 3 | SPRY2 | sprouty RTK signaling antagonist | Plasma Membrane | other | |
| chr3 | *RP11-699L21.1* | 1 | 30 | 3 | | | | | |
| chr15 | *GJD2* | 2 | 83 | 3 | GJD2 | gap junction protein delta 2 | Plasma Membrane | other | |
| chr7 | *AC010971.1* | 2 | 41 | 3 | | | | | |
| chr19 | *CTD-2545M3.8* | 1 | 48 | 3 | | | | | |
| chr6 | *HECA* | 1 | 24 | 3 | HECA | hdc homolog, cell cycle regulator | Nucleus | other | |
| chr3 | *TRH* | 2 | 74 | 3 | TRH | thyrotropin releasing hormone | Extracellular Space | other | |
| chr7 | *AC003092.1* | 2 | 39 | 3 | | | | | |
| chr9 | *LCN12* | 2 | 91 | 3 | LCN12 | lipocalin 12 | Extracellular Space | transporter | |
| chr10 | *ZNF485* | 2 | 46 | 3 | ZNF485 | zinc finger protein 485 | Nucleus | transcription regulator | |
| chr19 | *SNAPC2* | 2 | 66 | 3 | SNAPC2 | small nuclear RNA activating complex polypeptide 2 | Nucleus | transcription regulator | |
| chr3 | *ROPN1B* | 1 | 52 | 3 | ROPN1B | rhophilin associated tail protein | Cytoplasm | other | |
| chr19 | *CTD-2329C7.2* | 1 | 51 | 3 | | | | | |
| chr18 | *SALL3* | 2 | 57 | 3 | SALL3 | spalt like transcription factor 3 | Nucleus | transcription regulator | |
| chr19 | *CTD-2231E14.5* | 1 | 41 | 3 | | | | | |
| chr11 | *KRT8P7* | 2 | 81 | 3 | KRT8P7 | keratin 8 pseudogene 7 | Other | other | |
| chr17 | *CTD-2510F5.6* | 1 | 30 | 3 | | | | | |
| chr19 | *SPIB* | 2 | 43 | 3 | SPIB | Spi-B transcription factor | Nucleus | transcription regulator | |
| chr11 | *ARFIP2* | 2 | 97 | 3 | ARFIP2 | ADP ribosylation factor interacting protein 2 | Cytoplasm | other | |
| chr17 | *RP11-94C24.6* | 2 | 49 | 3 | | | | | |
| chr5 | *RP11-215P8.3* | 1 | 41 | 3 | | | | | |
| chr6 | *PDE7B* | 2 | 36 | 3 | PDE7B | phosphodiesterase 7B | Cytoplasm | enzyme | pentoxifylline, tolbutamide, anagrelide, theophylline, milrinone, |
| chr7 | *SNHG15* | 2 | 87 | 3 | SNHG15 | small nucleolar RNA host gene | Other | other | |
| chr2 | *C2orf27B* | 1 | 43 | 3 | C2orf27A/C2orf27I | chromosome 2 open reading frame 27B | Other | other | |
| chr1 | *DDX11L1* | 2 | 209 | 3 | DDX11L1/DDX11L | DEAD/H-box helicase 11 like 1 | Other | other | |
| chr20 | *RP5-850E9.3* | 2 | 34 | 3 | | | | | |
| chr17 | *TMEM99* | 2 | 42 | 3 | TMEM99 | transmembrane protein 99 | Other | other | |
| chr1 | *IRF6* | 2 | 44 | 3 | IRF6 | interferon regulatory factor 6 | Nucleus | transcription regulator | |
| chr18 | *ZBTB14* | 2 | 115 | 3 | ZBTB14 | zinc finger and BTB domain containing 14 | Nucleus | transcription regulator | |
| chr16 | *ORC6* | 2 | 44 | 3 | ORC6 | origin recognition complex | Nucleus | other | |
| chr9 | *ANKRD20A2* | 1 | 29 | 3 | D20A2 (includes c | ankyrin repeat domain 20 family member A2 | Plasma Membrane | other | |
| chr2 | *TMEM17* | 1 | 25 | 3 | TMEM17 | transmembrane protein 17 | Extracellular Space | other | |
| chr1 | *RP11-348H3.2* | 1 | 59 | 3 | | | | | |
| chr3 | *SLC35A5* | 2 | 47 | 3 | SLC35A5 | solute carrier family 35 member | Other | transporter | |
| chr12 | *CDCA3* | 2 | 34 | 3 | CDCA3 | cell division cycle associated 3 | Cytoplasm | other | |

| | | | | | | | | | |
|---|---|---|---|---|---|---|---|---|---|
| chr11 | *RP11-1036E20.9* | 2 | 36 | 3 | | | | | |
| chr2 | *AC115115.4* | 1 | 86 | 3 | | | | | |
| chr4 | *RP11-1406H17.1* | 1 | 30 | 3 | | | | | |
| chr4 | *UGT2B29P* | 1 | 29 | 3 | UGT2B29P | UDP glucuronosyltransferase family 2 member B29, | Other | other | |
| chr2 | *IDH1* | 1 | 23 | 3 | IDH1 | isocitrate dehydrogenase (NADP(+)) 1 | Cytoplasm | enzyme | olutasidenib, IDH305, ivosidenib, DS-1001, BAY1436032, IDH1 |
| chr18 | *RP11-128P17.2* | 1 | 33 | 3 | | | | | |
| chr1 | *RP11-467I20.3* | 1 | 42 | 3 | | | | | |
| chr4 | *UNC93B4* | 2 | 55 | 3 | UNC93B4 | unc-93 homolog B4, pseudogene | Other | other | |
| chr16 | *RP11-146F11.3* | 1 | 52 | 3 | | | | | |
| chr16 | *RP11-189E14.3* | 2 | 45 | 3 | | | | | |
| chr19 | *AC005794.1* | 1 | 38 | 3 | | | | | |
| chr2 | *FIGLA* | 2 | 37 | 3 | FIGLA | folliculogenesis specific bHLH transcription factor | Nucleus | transcription regulator | |
| chr3 | *RP11-803B1.2* | 2 | 30 | 3 | | | | | |
| chr16 | *RP11-417N10.3* | 1 | 33 | 3 | | | | | |
| chr7 | *LSMEM1* | 1 | 31 | 3 | LSMEM1 | leucine rich single-pass membrane protein 1 | Cytoplasm | other | |
| chr21 | *AP001631.10* | 2 | 73 | 3 | | | | | |
| chr2 | *CYTIP* | 1 | 25 | 3 | CYTIP | cytohesin 1 interacting protein | Cytoplasm | other | |
| chr2 | *ATF2* | 1 | 23 | 3 | ATF2 | activating transcription factor 2 | Nucleus | transcription regulator | |
| chr9 | *NPR2* | 2 | 35 | 3 | NPR2 | natriuretic peptide receptor 2 | Plasma Membrane | G-protein coupled recepto | nesiritide |
| chr22 | *LINC00528* | 2 | 40 | 3 | LINC00528 | long intergenic non-protein coding RNA 528 | Other | other | |
| chr10 | *KRT19P4* | 1 | 63 | 3 | KRT19P4 | keratin 19 pseudogene 4 | Other | other | |
| chr17 | *TMEM95* | 2 | 67 | 3 | TMEM95 | transmembrane protein 95 | Other | other | |
| chr8 | *ZNF705B* | 2 | 142 | 3 | ZNF705B/ZNF705I | zinc finger protein 705B | Other | other | |
| chr20 | *SNX5* | 1 | 24 | 3 | SNX5 | sorting nexin 5 | Cytoplasm | transporter | |
| chr12 | *TUBB8P5* | 2 | 97 | 3 | TUBB8P5 | tubulin beta 8 class VIII pseudogene 5 | Other | other | |
| chr5 | *RP11-432M8.1* | 1 | 478 | 3 | | | | | |
| chr17 | *RNASEH1P2* | 1 | 45 | 3 | RNASEH1P2 | ribonuclease H1 pseudogene 2 | Other | other | |
| chr10 | *PABPC1P12* | 2 | 61 | 3 | PABPC1P12 | poly(A) binding protein cytoplasmic 1 pseudogene 12 | Other | other | |
| chr2 | *RP11-440D17.1* | 2 | 74 | 3 | | | | | |
| chrX | *CXorf49* | 2 | 65 | 3 | CXorf49/CXorf49E | chromosome X open reading frame 49 | Other | other | |
| chr7 | *ASS1P11* | 2 | 62 | 3 | ASS1P11 | argininosuccinate synthetase 1 pseudogene 11 | Other | other | |
| chr6 | *RP1-40E16.2* | 2 | 49 | 3 | | | | | |
| chr17 | *COTL1P2* | 1 | 55 | 3 | COTL1P2 | COTL1 pseudogene 2 | Other | other | |
| chr16 | *RP11-586K12.11* | 1 | 34 | 3 | | | | | |
| chrX | *RP11-432N13.4* | 1 | 26 | 3 | | | | | |
| chr18 | *SERPINB5* | 2 | 36 | 3 | SERPINB5 | serpin family B member 5 | Extracellular Space | other | |
| chr11 | *AP000593.6* | 1 | 52 | 3 | | | | | |
| chr19 | *COX6B2* | 2 | 62 | 3 | COX6B2 | cytochrome c oxidase subunit | Cytoplasm | enzyme | |
| chr17 | *SNHG16* | 2 | 46 | 3 | SNHG16 | small nucleolar RNA host gene | Other | other | |
| chr7 | *GNB2* | 2 | 69 | 3 | GNB2 | G protein subunit beta 2 | Plasma Membrane | enzyme | |
| chr5 | *MARK2P5* | 2 | 168 | 3 | MARK2P5 | microtubule affinity regulating kinase 2 pseudogene 5 | Other | other | |
| chr21 | *AP001046.5* | 2 | 76 | 3 | | | | | |
| chr10 | *PCBD1* | 2 | 62 | 3 | PCBD1 | pterin-4 alpha-carbinolamine dehydratase 1 | Nucleus | transcription regulator | |
| chr2 | *AC007249.3* | 1 | 43 | 3 | | | | | |
| chr7 | *MRPS33* | 2 | 58 | 3 | MRPS33 | mitochondrial ribosomal protein | Cytoplasm | other | |
| chr20 | *SLMO2* | 2 | 53 | 3 | PRELID3B | PRELI domain containing 3B | Cytoplasm | other | |
| chr11 | *RP11-159D8.1* | 1 | 24 | 3 | | | | | |
| chr11 | *RP11-735A19.2* | 1 | 91 | 3 | | | | | |
| chr18 | *RP11-677O4.4* | 1 | 42 | 3 | | | | | |
| chr6 | *RP3-336K20__B.2* | 1 | 43 | 3 | | | | | |
| chr9 | *VN2R4P* | 2 | 35 | 3 | VN2R3P | vomeronasal 2 receptor 3, pseudogene | Other | other | |
| chr11 | *SRSF8* | 2 | 49 | 3 | SRSF8 | serine and arginine rich splicing factor 8 | Nucleus | other | |
| chr18 | *RP11-108P20.4* | 2 | 35 | 3 | | | | | |
| chr17 | *RP1-178F10.3* | 1 | 58 | 3 | | | | | |
| chr17 | *HES7* | 2 | 97 | 3 | HES7 | hes family bHLH transcription | Nucleus | transcription regulator | |
| chr10 | *SLC9A3P4* | 1 | 71 | 3 | SLC9A3P1 | solute carrier family 9 member 3 pseudogene 1 | Other | other | |
| chr17 | *USP32P3* | 2 | 36 | 3 | USP32P3 | ubiquitin specific peptidase 32 pseudogene 3 | Other | other | |
| chr10 | *RP11-184A2.3* | 2 | 29 | 3 | | | | | |
| chr5 | *MIR146A* | 2 | 41 | 3 | mir-146 | microRNA 146a | Cytoplasm | microRNA | |
| chr15 | *RP11-19J5.2* | 2 | 58 | 3 | | | | | |
| chr3 | *DUSP5P2* | 2 | 48 | 3 | DUSP5P2 | dual specificity phosphatase 5 pseudogene 2 | Other | other | |
| chr17 | *AC104024.2* | 1 | 51 | 3 | | | | | |
| chr3 | *POPDC2* | 2 | 46 | 3 | POPDC2 | popeye domain containing 2 | Plasma Membrane | other | |
| chr17 | *MIR2117* | 2 | 39 | 3 | MIR2117 | microRNA 2117 | Cytoplasm | microRNA | |
| chr17 | *C17orf76-AS1* | 2 | 54 | 3 | SNHG29 | small nucleolar RNA host gene | Other | other | |
| chr10 | *RP11-144G16.1* | 2 | 59 | 3 | | | | | |
| chr17 | *CTD-2349P21.1* | 1 | 35 | 3 | | | | | |
| chr1 | *GPX7* | 2 | 58 | 3 | GPX7 | glutathione peroxidase 7 | Cytoplasm | enzyme | |
| chr6 | *TREML1* | 2 | 51 | 3 | TREML1 | triggering receptor expressed on myeloid cells like 1 | Plasma Membrane | other | |
| chr14 | *RP11-930O11.1* | 1 | 28 | 3 | | | | | |

| | | | | | | | | | |
|---|---|---|---|---|---|---|---|---|---|
| chr19 | *OR10B1P* | 1 | 50 | 3 | OR10B1P | olfactory receptor family 10 subfamily B member 1 | Other | other | |
| chr1 | *OR2T33* | 1 | 59 | 3 | OR2T33 | olfactory receptor family 2 subfamily T member 33 | Plasma Membrane | other | |
| chr17 | *AC003958.6* | 1 | 54 | 3 | | | | | |
| chr20 | *NSFL1C* | 1 | 32 | 3 | NSFL1C | NSFL1 cofactor | Cytoplasm | other | |
| chr12 | *RP11-588G21.2* | 2 | 31 | 3 | | | | | |
| chr5 | *CTNNAP1* | 1 | 44 | 3 | CTNNA1P1 | catenin alpha 1 pseudogene 1 | Other | other | |
| chr4 | *LINC00989* | 1 | 27 | 3 | LINC00989 | long intergenic non-protein coding RNA 989 | Other | other | |
| chr10 | *AL133481.1* | 1 | 64 | 3 | | | | | |
| chr11 | *AP000479.1* | 2 | 40 | 3 | | | | | |
| chr8 | *VPS28* | 2 | 72 | 3 | VPS28 | VPS28 subunit of ESCRT-I | Cytoplasm | transporter | |
| chr6 | *YAP1P1* | 2 | 56 | 3 | YAP1P1 | Yes associated protein 1 pseudogene 1 | Other | other | |
| chr14 | *ACOT6* | 1 | 30 | 3 | ACOT6 | acyl-CoA thioesterase 6 | Cytoplasm | enzyme | |
| chr10 | *C10orf35* | 1 | 76 | 3 | FAM241B | family with sequence similarity 241 member B | Other | other | |
| chr3 | *VENTXP4* | 1 | 42 | 3 | VENTXP4 | VENT homeobox pseudogene 4 | Other | other | |
| chr21 | *AP001053.11* | 2 | 77 | 3 | | | | | |
| chr1 | *FMO11P* | 2 | 43 | 3 | FMO11P | flavin containing dimethylaniline monoxygenase 11, pseudogene | Other | other | |
| chr16 | *NTN3* | 2 | 69 | 3 | NTN3 | netrin 3 | Extracellular Space | other | |
| chr21 | *AP000230.1* | 1 | 22 | 3 | | | | | |
| chr12 | *RP4-816N1.6* | 1 | 31 | 3 | | | | | |
| chr10 | *MBL3P* | 2 | 39 | 3 | MBL3P | mannose-binding lectin family member 3, pseudogene | Other | other | |
| chr6 | *METTL24* | 2 | 53 | 3 | METTL24 | methyltransferase like 24 | Other | other | |
| chr12 | *RP11-486A14.1* | 1 | 36 | 3 | | | | | |
| chr13 | *LINC01070* | 1 | 49 | 3 | LINC01070 | long intergenic non-protein coding RNA 1070 | Other | other | |
| chr16 | *RP11-297D21.4* | 2 | 40 | 3 | | | | | |
| chr10 | *GLUD1P6* | 2 | 50 | 3 | GLUD1P6 | glutamate dehydrogenase 1 pseudogene 6 | Other | other | |
| chr9 | *SURF1* | 2 | 85 | 3 | SURF1 | SURF1 cytochrome c oxidase assembly factor | Cytoplasm | enzyme | |
| chr19 | *ZNF507* | 2 | 32 | 3 | ZNF507 | zinc finger protein 507 | Nucleus | transcription regulator | |
| chr1 | *RP11-344P13.4* | 1 | 43 | 3 | | | | | |
| chr17 | *SLFN11* | 2 | 65 | 3 | SLFN11 | schlafen family member 11 | Nucleus | enzyme | |
| chr15 | *RP11-114H24.3* | 2 | 47 | 3 | | | | | |
| chr6 | *FHL5* | 1 | 22 | 3 | FHL5 | four and a half LIM domains 5 | Nucleus | transcription regulator | |
| chr8 | *RAD54B* | 1 | 22 | 3 | RAD54B | RAD54 homolog B | Nucleus | enzyme | |
| chr8 | *RP11-383H13.1* | 2 | 29 | 3 | | | | | |
| chr1 | *NRAS* | 2 | 43 | 3 | NRAS | NRAS proto-oncogene, GTPase | Plasma Membrane | enzyme | |
| chr17 | *FAM101B* | 2 | 41 | 3 | RFLNB | refilin B | Other | other | |
| chr7 | *AC098592.8* | 1 | 44 | 3 | | | | | |
| chr10 | *RN7SL686P* | 1 | 31 | 3 | | | | | |
| chr17 | *AC087491.2* | 2 | 52 | 3 | | | | | |
| chr6 | *RP3-359N14.2* | 1 | 22 | 3 | | | | | |
| chr2 | *DRD5P1* | 1 | 184 | 3 | DRD5P1 | dopamine receptor D5 | Other | other | |
| chr19 | *PTOV1-AS1* | 1 | 36 | 3 | PTOV1-AS1 | PTOV1 antisense RNA 1 | Other | other | |
| chr1 | *C1orf170* | 2 | 66 | 3 | PERM1 | PPARGC1 and ESRR induced regulator, muscle 1 | Other | other | |
| chr20 | *PTMAP6* | 2 | 55 | 3 | PTMAP6 | prothymosin alpha pseudogene 6 | Other | other | |
| chr9 | *RP11-263F14.3* | 1 | 74 | 3 | | | | | |
| chr8 | *ZNF252P* | 1 | 23 | 3 | ZNF252P | zinc finger protein 252, | Other | other | |
| chr14 | *AL589743.1* | 2 | 41 | 3 | | | | | |
| chr9 | *TOR2A* | 2 | 141 | 3 | TOR2A | torsin family 2 member A | Extracellular Space | other | |
| chr13 | *PRR20E* | 2 | 144 | 3 | R20A (includes oth | proline rich 20A | Other | other | |
| chr19 | *DMPK* | 2 | 73 | 3 | DMPK | DM1 protein kinase | Cytoplasm | kinase | |
| chr19 | *TUBB4A* | 2 | 64 | 3 | TUBB4A | tubulin beta 4A class IVa | Cytoplasm | other | [illegible]e/vincristine, cyclophosphamide/epirubicin/5-fluorouracil/paclitaxel, bevacizumab/paclitaxel/topotecan, colchicine/probenecid, bevacizumab/docetaxel/prednisone/thalidomide/zoledronic acid, gemcitabine/paclitaxel, simotaxel, L-asparaginase/daunorubicin/dexamethasone/vincristine, prednisone/vincristine, docetaxel/5-fluorouracil, vinflunine, L-[illegible] |
| chr7 | *AC005027.3* | 2 | 41 | 3 | | | | | |
| chr10 | *FAM21D* | 1 | 34 | 3 | | | | | |
| chr6 | *LTV1* | 2 | 29 | 3 | LTV1 | LTV1 ribosome biogenesis factor | Other | other | |
| chrX | *BMP15* | 1 | 33 | 3 | BMP15 | bone morphogenetic protein 15 | Extracellular Space | growth factor | |
| chr1 | *RP4-706L14.2* | 1 | 53 | 3 | | | | | |

| | | | | | | | | | |
|---|---|---|---|---|---|---|---|---|---|
| chr19 | *SULT2A1* | 1 | 25 | 3 | SULT2A1 | sulfotransferase family 2A | Cytoplasm | enzyme | |
| chr20 | *MC3R* | 1 | 86 | 3 | MC3R | melanocortin 3 receptor | Plasma Membrane | G-protein coupled recepto | bremelanotide |
| chrX | *RP11-344N17.11* | 2 | 32 | 3 | | | | | |
| chr3 | *RP11-803B1.8* | 2 | 33 | 3 | | | | | |
| chrX | *USP26* | 1 | 22 | 3 | USP26 | ubiquitin specific peptidase 26 | Cytoplasm | peptidase | |
| chrX | *RP11-1114A5.4* | 1 | 22 | 3 | | | | | |
| chr1 | *CALML6* | 2 | 100 | 3 | CALML6 | calmodulin like 6 | Cytoplasm | other | |
| chr4 | *RP11-18H21.2* | 1 | 26 | 3 | | | | | |
| chr6 | *HIST1H2BI* | 1 | 42 | 3 | H2BC10 | H2B clustered histone 10 | Nucleus | other | |
| chr5 | *MSNP1* | 2 | 127 | 3 | MSNP1 | moesin pseudogene 1 | Other | other | |
| chr11 | *CRYAB* | 2 | 39 | 3 | CRYAB | crystallin alpha B | Nucleus | other | |
| chr11 | *ESAM* | 2 | 42 | 3 | ESAM | endothelial cell adhesion | Plasma Membrane | other | |
| chr5 | *RP11-461O14.3* | 1 | 57 | 3 | | | | | |
| chr12 | *KRT128P* | 2 | 77 | 3 | KRT128P | | Other | other | |
| chr2 | *AC068057.1* | 1 | 41 | 3 | | | | | |
| chr7 | *RP11-310H4.3* | 1 | 61 | 3 | | | | | |
| chr7 | *PRKAG2-AS1* | 1 | 55 | 3 | PRKAG2-AS1 | PRKAG2 antisense RNA 1 | Other | other | |
| chr18 | *HDHD2* | 1 | 30 | 3 | HDHD2 | haloacid dehalogenase like hydrolase domain containing 2 | Cytoplasm | other | |
| chr7 | *SLC25A5P3* | 1 | 55 | 3 | SLC25A5P3 | solute carrier family 25 member 5 pseudogene 3 | Other | other | |
| chr21 | *AP000688.8* | 1 | 25 | 3 | | | | | |
| chr22 | *MPST* | 2 | 46 | 3 | MPST | mercaptopyruvate | Cytoplasm | enzyme | |
| chr2 | *AC016738.4* | 2 | 70 | 3 | | | | | |
| chr2 | *RP11-734K21.5* | 1 | 37 | 3 | | | | | |
| chr3 | *AC106706.1* | 1 | 31 | 3 | | | | | |
| chr10 | *FRAT1* | 2 | 67 | 3 | FRAT1 | FRAT regulator of WNT signaling pathway 1 | Cytoplasm | other | |
| chr11 | *RP11-945A11.1* | 2 | 49 | 3 | | | | | |
| chr4 | *RP11-703G6.1* | 1 | 22 | 3 | | | | | |
| chr16 | *MOCS1P1* | 2 | 100 | 3 | MOCS1P1 | molybdenum cofactor synthesis 1 pseudogene 1 | Other | other | |
| chr20 | *RP11-112L6.3* | 2 | 50 | 3 | | | | | |
| chr7 | *MARK2P13* | 1 | 58 | 3 | MARK2P13 | microtubule affinity regulating kinase 2 pseudogene 13 | Other | other | |
| chr12 | *CPSF6* | 2 | 40 | 3 | CPSF6 | cleavage and polyadenylation specific factor 6 | Nucleus | other | |
| chr9 | *RP11-430K21.2* | 1 | 26 | 3 | | | | | |
| chr17 | *AC008088.4* | 1 | 27 | 3 | | | | | |
| chr6 | *RP3-441A12.1* | 1 | 27 | 3 | | | | | |
| chr9 | *FRRS1L* | 1 | 22 | 3 | FRRS1L | ferric chelate reductase 1 like | Cytoplasm | transporter | |
| chr15 | *RP11-23P13.6* | 2 | 70 | 3 | | | | | |
| chr11 | *C11orf71* | 1 | 30 | 3 | C11orf71 | chromosome 11 open reading frame 71 | Other | other | |
| chr9 | *IGKV1OR-3* | 1 | 57 | 3 | IGKV1OR-3 | immunoglobulin kappa variable 1/OR-3 (pseudogene) | Other | other | |
| chr1 | *RP5-1086K13.1* | 2 | 63 | 3 | | | | | |
| chr1 | *OR11L1* | 1 | 37 | 3 | OR11L1 | olfactory receptor family 11 subfamily L member 1 | Plasma Membrane | G-protein coupled recepto | |
| chr1 | *RP11-297H3.3* | 2 | 37 | 3 | | | | | |
| chr9 | *C9orf147* | 1 | 22 | 3 | C9orf147 | chromosome 9 open reading frame 147 | Other | other | |
| chr20 | *RP5-1069C8.2* | 1 | 22 | 3 | | | | | |
| chr16 | *RP11-419L9.1* | 1 | 51 | 3 | | | | | |
| chr10 | *RP11-38L15.3* | 1 | 138 | 3 | | | | | |
| chrX | *VCX3A* | 2 | 81 | 3 | VCX/VCX3A | variable charge X-linked 3A | Nucleus | other | |
| chr16 | *CTD-3032H12.2* | 2 | 55 | 3 | | | | | |
| chr21 | *KCNE2* | 1 | 22 | 3 | KCNE2 | potassium voltage-gated channel subfamily E regulatory subunit 2 | Plasma Membrane | ion channel | amiodarone, nicorandil |
| chr17 | *OR3A3* | 1 | 41 | 3 | OR3A3 | olfactory receptor family 3 subfamily A member 3 | Plasma Membrane | G-protein coupled recepto | |
| chr7 | *TAC1* | 2 | 118 | 3 | TAC1 | tachykinin precursor 1 | Extracellular Space | other | rolapitant |
| chr19 | *ZNF543* | 2 | 39 | 3 | ZNF543 | zinc finger protein 543 | Other | other | |
| chr1 | *RP11-541F9.1* | 1 | 27 | 3 | | | | | |
| chr19 | *CTD-3214H19.12* | 1 | 36 | 3 | | | | | |
| chr2 | *AC018865.8* | 1 | 35 | 3 | | | | | |
| chr12 | *RP11-273B20.3* | 1 | 22 | 3 | | | | | |
| chr18 | *RP11-739L10.1* | 1 | 22 | 3 | | | | | |
| chr5 | *LYSMD3* | 1 | 29 | 3 | LYSMD3 | LysM domain containing 3 | Other | other | |
| chr11 | *KBTBD3* | 1 | 35 | 3 | KBTBD3 | kelch repeat and BTB domain containing 3 | Extracellular Space | other | |
| chr11 | *RP11-108O10.2* | 1 | 24 | 3 | | | | | |
| chr12 | *DCP1B* | 1 | 36 | 3 | DCP1B | decapping mRNA 1B | Cytoplasm | other | |
| chr5 | *AC004237.1* | 1 | 25 | 3 | | | | | |
| chr8 | *RP11-589N15.1* | 1 | 39 | 3 | | | | | |
| chr1 | *SERTAD4* | 1 | 42 | 3 | SERTAD4 | SERTA domain containing 4 | Other | other | |
| chr20 | *EIF6* | 2 | 57 | 3 | EIF6 | eukaryotic translation initiation | Cytoplasm | translation regulator | |
| chr16 | *ENKD1* | 2 | 66 | 3 | ENKD1 | enkurin domain containing 1 | Cytoplasm | other | |
| chr12 | *RP11-490G8.1* | 1 | 40 | 3 | | | | | |
| chr7 | *FLJ20306* | 1 | 61 | 3 | | | | | |
| chr2 | *MED15P4* | 2 | 83 | 3 | MED15P4 | mediator complex subunit 15 pseudogene 4 | Other | other | |
| chr14 | *RP11-600F24.1* | 1 | 28 | 3 | | | | | |
| chr2 | *AC139712.2* | 2 | 71 | 3 | | | | | |
| chr12 | *RP11-407A16.7* | 1 | 27 | 3 | | | | | |
| chr11 | *C11orf88* | 1 | 22 | 3 | C11orf88 | chromosome 11 open reading frame 88 | Other | other | |
| chr5 | *AC008937.3* | 1 | 26 | 3 | | | | | |
| chr17 | *AC013248.2* | 1 | 22 | 3 | | | | | |

| | | | | | | | | | |
|---|---|---|---|---|---|---|---|---|---|
| chr1 | *RP11-435B5.6* | 1 | 38 | 3 | | | | | |
| chr5 | *CTD-2366F13.1* | 1 | 38 | 3 | | | | | |
| chr1 | *GJA8* | 2 | 78 | 3 | GJA8 | gap junction protein alpha 8 | Plasma Membrane | transporter | |
| chr19 | *AC092299.6* | 1 | 41 | 3 | | | | | |
| chr1 | *RP11-536C5.2* | 2 | 94 | 3 | | | | | |
| chrX | *SPIN3* | 2 | 61 | 3 | SPIN3 | spindlin family member 3 | Other | other | |
| chrX | *MAGEB1* | 2 | 51 | 3 | MAGEB1 | MAGE family member B1 | Nucleus | other | |
| chr6 | *HIST1H2BC* | 1 | 23 | 3 | H2BC4 | H2B clustered histone 4 | Nucleus | other | |
| chr1 | *RP13-221M14.6* | 1 | 27 | 3 | | | | | |
| chr4 | *MFAP3L* | 2 | 33 | 3 | MFAP3L | microfibril associated protein 3 | Nucleus | other | |
| chr14 | *ZBTB42* | 2 | 68 | 3 | ZBTB42 | zinc finger and BTB domain containing 42 | Cytoplasm | transcription regulator | |
| chr5 | *RP11-395C3.1* | 1 | 33 | 3 | | | | | |
| chr4 | *RP11-153M7.3* | 1 | 36 | 3 | | | | | |
| chr18 | *RP11-53I6.2* | 1 | 26 | 3 | | | | | |
| chr7 | *LINC01007* | 1 | 53 | 3 | LINC01007 | long intergenic non-protein coding RNA 1007 | Other | other | |
| chr19 | *RPL18A* | 2 | 66 | 3 | RPL18A | ribosomal protein L18a | Cytoplasm | other | |
| chr8 | *AF121898.3* | 1 | 29 | 3 | | | | | |
| chr4 | *RP11-367N14.3* | 1 | 22 | 3 | | | | | |
| chr9 | *OBP2B* | 2 | 81 | 3 | OBP2B | odorant binding protein 2B | Extracellular Space | transporter | |
| chr14 | *ATP5S* | 1 | 23 | 3 | DMAC2L | distal membrane arm assembly complex 2 like | Cytoplasm | transporter | |
| chr8 | *PYCRL* | 2 | 71 | 3 | PYCR3 | pyrroline-5-carboxylate reductase | Cytoplasm | enzyme | |
| chr15 | *DNM1P31* | 1 | 54 | 3 | DNM1P31 | dynamin 1 pseudogene 31 | Other | other | |
| chrX | *RP6-186E3.1* | 1 | 53 | 3 | | | | | |
| chr5 | *RNA5SP190* | 1 | 33 | 3 | RNA5SP190 | RNA, 5S ribosomal pseudogene | Other | other | |
| chr2 | *ABHD1* | 1 | 33 | 3 | ABHD1 | abhydrolase domain containing 1 | Other | other | |
| chr19 | *DBP* | 2 | 46 | 3 | DBP | D-box binding PAR bZIP transcription factor | Nucleus | transcription regulator | |
| chr16 | *Z84721.4* | 2 | 50 | 3 | | | | | |
| chr16 | *LRRC29* | 2 | 105 | 3 | LRRC29 | leucine rich repeat containing 29 | Other | enzyme | |
| chr7 | *RCC2P3* | 2 | 67 | 3 | RCC2P3 | regulator of chromosome condensation 2 pseudogene 3 | Other | other | |
| chr1 | *RP11-412H9.1* | 1 | 41 | 3 | | | | | |
| chr10 | *RP13-39P12.2* | 1 | 54 | 3 | | | | | |
| chr2 | *AC134915.1* | 1 | 36 | 3 | | | | | |
| chr11 | *RP11-810P12.1* | 1 | 64 | 3 | | | | | |
| chr4 | *HS3ST1* | 2 | 40 | 3 | HS3ST1 | heparan sulfate-glucosamine 3-sulfotransferase 1 | Cytoplasm | enzyme | |
| chr17 | *RP11-388C12.1* | 2 | 82 | 3 | | | | | |
| chr1 | *RP5-857K21.3* | 2 | 89 | 3 | | | | | |
| chr7 | *AC102953.4* | 2 | 122 | 3 | | | | | |
| chr11 | *AP005135.2* | 2 | 54 | 3 | | | | | |
| chr1 | *LCE3E* | 2 | 57 | 3 | LCE3E | late cornified envelope 3E | Other | other | |
| chr6 | *FGFR3P1* | 1 | 42 | 3 | FGFR3P1 | fibroblast growth factor receptor 3 pseudogene 1 | Other | other | |
| chr14 | *LINC00596* | 1 | 36 | 3 | LINC00596 | long intergenic non-protein coding RNA 596 | Other | other | |
| chr4 | *RP11-323J4.1* | 1 | 24 | 3 | | | | | |
| chr8 | *CA8* | 1 | 21 | 3 | CA8 | carbonic anhydrase 8 | Cytoplasm | enzyme | |
| chr2 | *AC009474.1* | 2 | 41 | 3 | | | | | |
| chr11 | *RP11-787P24.1* | 2 | 84 | 3 | | | | | |
| chr4 | *SLC25A4* | 1 | 29 | 3 | SLC25A4 | solute carrier family 25 member 4 | Cytoplasm | transporter | clodronic acid |
| chr22 | *HIRA* | 1 | 23 | 3 | HIRA | histone cell cycle regulator | Nucleus | transcription regulator | |
| chr12 | *RP11-429A20.4* | 1 | 27 | 3 | | | | | |
| chr11 | *COLCA1* | 2 | 43 | 3 | COLCA1 | colorectal cancer associated 1 | Other | other | |
| chr1 | *HSPB7* | 2 | 108 | 3 | HSPB7 | heat shock protein family B (small) member 7 | Cytoplasm | other | |
| chr5 | *RP11-321E2.9* | 1 | 848 | 3 | | | | | |
| chr2 | *INO80B* | 1 | 46 | 3 | INO80B | INO80 complex subunit B | Nucleus | other | |
| chr2 | *AC079790.2* | 1 | 23 | 3 | | | | | |
| chr16 | *TPSG1* | 2 | 70 | 3 | TPSG1 | tryptase gamma 1 | Extracellular Space | peptidase | |
| chr4 | *IDUA* | 2 | 46 | 3 | IDUA | alpha-L-iduronidase | Cytoplasm | enzyme | |
| chr14 | *ACOT2* | 2 | 59 | 3 | ACOT2 | acyl-CoA thioesterase 2 | Cytoplasm | enzyme | |
| chr17 | *C1QTNF1-AS1* | 1 | 40 | 3 | C1QTNF1-AS1 | C1QTNF1 antisense RNA 1 | Other | other | |
| chr6 | *HIST1H4G* | 1 | 39 | 3 | H4C7 | H4 clustered histone 7 | Nucleus | other | |
| chr2 | *CD28* | 1 | 27 | 3 | CD28 | CD28 molecule | Plasma Membrane | transmembrane recepto | lulizumab pegol, REGN5678 |
| chr22 | *THAP7* | 2 | 62 | 3 | THAP7 | THAP domain containing 7 | Nucleus | transcription regulator | |
| chr19 | *VN2R11P* | 1 | 37 | 3 | | | | | |
| chr2 | *ITGA4* | 1 | 21 | 3 | ITGA4 | integrin subunit alpha 4 | Plasma Membrane | transmembrane recepto | natalizumab, vedolizumab |
| chr4 | *RP11-789C2.1* | 1 | 22 | 3 | | | | | |
| chr1 | *EXTL2* | 1 | 36 | 3 | EXTL2 | exostosin like glycosyltransferase | Cytoplasm | enzyme | |
| chr14 | *LINC00341* | 1 | 33 | 3 | SYNE3 | spectrin repeat containing nuclear envelope family member 3 | Nucleus | other | |
| chrX | *CXorf65* | 2 | 27 | 3 | CXorf65 | chromosome X open reading frame 65 | Other | other | |
| chr14 | *RP11-1070N10.6* | 1 | 30 | 3 | | | | | |
| chr11 | *AP000640.2* | 1 | 27 | 3 | | | | | |
| chr6 | *LINC00266-3* | 1 | 26 | 3 | LINC00266-3 | long intergenic non-protein coding RNA 266-3 | Other | other | |
| chrX | *GPR112* | 1 | 21 | 3 | ADGRG4 | adhesion G protein-coupled receptor G4 | Plasma Membrane | -protein coupled recept | |
| chr17 | *RP11-46I8.1* | 1 | 32 | 3 | | | | | |
| chr1 | *RP11-245P10.6* | 1 | 34 | 3 | | | | | |
| chr21 | *LINC00112* | 1 | 36 | 3 | LINC00112 | long intergenic non-protein coding RNA 112 | Other | other | |

| | | | | | | | | | |
|---|---|---|---|---|---|---|---|---|---|
| chr4 | *PRKG2* | 1 | 27 | 3 | PRKG2 | protein kinase cGMP-dependent | Cytoplasm | kinase | |
| chr4 | *ENOPH1* | 1 | 28 | 3 | ENOPH1 | enolase-phosphatase 1 | Cytoplasm | enzyme | |
| chr5 | *CTD-2193G5.1* | 1 | 23 | 3 | | | | | |
| chr3 | *IQCF2* | 1 | 46 | 3 | IQCF2 | IQ motif containing F2 | Other | other | |
| chr13 | *DDX39AP1* | 1 | 26 | 3 | DDX39AP1 | DEAD-box helicase 39A pseudogene 1 | Other | other | |
| chr3 | *PRADC1P1* | 1 | 50 | 3 | PRADC1P1 | protease associated domain containing 1 pseudogene 1 | Other | other | |
| chr8 | *DEFT1P* | 1 | 34 | 3 | DEFT1P | defensin theta 1, pseudogene | Other | other | |
| chr7 | *AC004941.3* | 1 | 42 | 3 | | | | | |
| chr9 | *RP11-272P10.3* | 1 | 39 | 3 | | | | | |
| chr7 | *ZNF467* | 2 | 84 | 3 | ZNF467 | zinc finger protein 467 | Nucleus | transcription regulator | |
| chr3 | *RP11-15N24.4* | 1 | 32 | 3 | | | | | |
| chr3 | *RP11-280H21.1* | 1 | 30 | 3 | | | | | |
| chr7 | *PCOLCE* | 2 | 64 | 3 | PCOLCE | procollagen C-endopeptidase enhancer | Extracellular Space | other | |
| chr7 | *AC016831.6* | 1 | 50 | 3 | | | | | |
| chr10 | *RP11-106M7.1* | 1 | 29 | 3 | | | | | |
| chr2 | *UBTFL5* | 2 | 34 | 3 | TFL3 (includes oth | UBTF like 3 (pseudogene) | Other | other | |
| chr7 | *ASIC3* | 2 | 53 | 3 | ASIC3 | acid sensing ion channel subunit 3 | Plasma Membrane | ion channel | amiloride, amiloride/hydrochloro thiazide, agmatine |
| chr19 | *YIPF2* | 2 | 62 | 3 | YIPF2 | Yip1 domain family member 2 | Cytoplasm | other | |
| chr19 | *CTD-2616J11.16* | 2 | 41 | 3 | | | | | |
| chr3 | *CCRL2* | 1 | 32 | 3 | CCRL2 | C-C motif chemokine receptor | Plasma Membrane | -protein coupled recepto | |
| chr20 | *AL117380.1* | 1 | 75 | 3 | | | | | |
| chr2 | *AC114755.3* | 1 | 45 | 3 | | | | | |
| chr6 | *HIST1H1T* | 1 | 28 | 3 | H1-6 | H1.6 linker histone, cluster | Nucleus | other | |
| chr5 | *C5orf51* | 2 | 157 | 3 | C5orf51 | chromosome 5 open reading frame 51 | Other | other | |
| chr9 | *C9orf141* | 2 | 136 | 3 | LINC02692 | long intergenic non-protein coding RNA 2692 | Other | other | |
| chr9 | *SLC34A3* | 2 | 95 | 3 | SLC34A3 | solute carrier family 34 member 3 | Plasma Membrane | transporter | |
| chr8 | *RP11-17A4.4* | 1 | 28 | 3 | | | | | |
| chr6 | *SYNCRIP* | 1 | 28 | 3 | SYNCRIP | synaptotagmin binding cytoplasmic RNA interacting | Nucleus | other | supinoxin |
| chr2 | *AC108868.1* | 2 | 73 | 3 | | | | | |
| chr1 | *XX-FW84067D5.2* | 1 | 31 | 3 | | | | | |
| chr9 | *AL390778.1* | 2 | 98 | 3 | | | | | |
| chrX | *RP11-272G22.1* | 1 | 62 | 3 | | | | | |
| chr14 | *FBXO33* | 1 | 24 | 3 | FBXO33 | F-box protein 33 | Other | other | |
| chr17 | *RP11-663N22.1* | 1 | 42 | 3 | | | | | |
| chr1 | *RP11-345P4.9* | 2 | 65 | 3 | | | | | |
| chr11 | *CTD-2234N14.1* | 2 | 30 | 3 | | | | | |
| chr14 | *PROX2* | 1 | 38 | 3 | PROX2 | prospero homeobox 2 | Other | other | |
| chr9 | *OR13J1* | 1 | 37 | 3 | OR13J1 | olfactory receptor family 13 subfamily J member 1 | Plasma Membrane | -protein coupled recepto | |
| chr8 | *RN7SL178P* | 1 | 35 | 3 | RN7SL178P | | Other | other | |
| chr8 | *RP11-521M14.2* | 1 | 28 | 3 | | | | | |
| chr13 | *LINC01047* | 1 | 25 | 3 | LINC01047 | long intergenic non-protein coding RNA 1047 | Other | other | |
| chr17 | *VN1R71P* | 1 | 27 | 3 | VN1R71P | vomeronasal 1 receptor 71 pseudogene | Other | other | |
| chr17 | *RP13-890H12.2* | 1 | 68 | 3 | | | | | |
| chr1 | *HMGB1P48* | 1 | 31 | 3 | HMGB1P48 | high mobility group box 1 pseudogene 48 | Other | other | |
| chr19 | *ZNF414* | 2 | 67 | 3 | ZNF414 | zinc finger protein 414 | Other | other | |
| chr19 | *AC002128.4* | 1 | 48 | 3 | | | | | |
| chr21 | *PAXBP1* | 1 | 21 | 3 | PAXBP1 | PAX3 and PAX7 binding protein 1 | Nucleus | other | |
| chr2 | *NMI* | 1 | 25 | 3 | NMI | N-myc and STAT interactor | Cytoplasm | transcription regulator | |
| chr2 | *LINC01087* | 1 | 38 | 3 | LINC01087 | long intergenic non-protein coding RNA 1087 | Other | other | |
| chr12 | *METTL7B* | 1 | 79 | 3 | METTL7B | methyltransferase like 7B | Other | enzyme | |
| chr2 | *AC114776.3* | 1 | 25 | 3 | | | | | |
| chr11 | *RP11-428C19.5* | 2 | 27 | 3 | | | | | |
| chr17 | *MAFG* | 2 | 102 | 3 | MAFG | MAF bZIP transcription factor G | Nucleus | transcription regulator | |
| chr9 | *HSPA5* | 2 | 86 | 3 | HSPA5 | heat shock protein family A (Hsp70) member 5 | Cytoplasm | enzyme | |
| chr8 | *SDCBP* | 1 | 21 | 3 | SDCBP | syndecan binding protein | Plasma Membrane | enzyme | |
| chr15 | *RP11-253M7.4* | 1 | 31 | 3 | | | | | |
| chr20 | *DKKL1P1* | 1 | 71 | 3 | DKKL1P1 | DKKL1 pseudogene 1 | Other | other | |
| chr19 | *AC010642.2* | 1 | 55 | 3 | | | | | |
| chr12 | *RP11-662I13.2* | 1 | 31 | 3 | | | | | |
| chr5 | *RP11-524L6.2* | 1 | 49 | 3 | | | | | |
| chr2 | *B3GNT7* | 2 | 75 | 3 | B3GNT7 | UDP-GlcNAc:betaGal beta-1,3-N-acetylglucosaminyltransferase 7 | Cytoplasm | enzyme | |
| chr14 | *RP11-255G12.3* | 1 | 26 | 3 | | | | | |
| chr9 | *AL445665.1* | 1 | 79 | 3 | | | | | |
| chr19 | *CTC-523E23.5* | 2 | 64 | 3 | | | | | |
| chr15 | *RP11-463I20.2* | 1 | 27 | 3 | | | | | |
| chr19 | *CTD-2619J13.19* | 2 | 61 | 3 | | | | | |
| chr1 | *RP5-827C21.2* | 1 | 50 | 3 | | | | | |
| chr1 | *RP11-136B18.2* | 1 | 30 | 3 | | | | | |
| chr1 | *RP11-242O24.3* | 1 | 42 | 3 | | | | | |
| chr9 | *RP11-569G13.2* | 1 | 35 | 3 | | | | | |
| chrX | *TEX13B* | 1 | 34 | 3 | TEX13B | testis expressed 13B | Other | other | |
| chr19 | *MBD3L3* | 1 | 45 | 3 | D3L2 (includes oth | methyl-CpG binding domain protein 3 like 2 | Other | other | |
| chr15 | *DUOXA2* | 2 | 56 | 3 | DUOXA2 | dual oxidase maturation factor 2 | Cytoplasm | other | |

| | | | | | | | | | |
|---|---|---|---|---|---|---|---|---|---|
| chr22 | *RP11-12M9.4* | 1 | 44 | 3 | | | | | |
| chr20 | *CSTL1* | 1 | 31 | 3 | CSTL1 | cystatin like 1 | Extracellular Space | other | |
| chr18 | *RP11-27G24.1* | 1 | 22 | 3 | | | | | |
| chr18 | *RP11-794M8.2* | 1 | 35 | 3 | | | | | |
| chr1 | *DRD5P2* | 2 | 255 | 3 | DRD5P2 | dopamine receptor D5 | Other | other | |
| chr1 | *RP4-781L3.1* | 1 | 42 | 3 | | | | | |
| chr18 | *CCDC11* | 1 | 25 | 3 | CFAP53 | cilia and flagella associated | Other | other | |
| chr4 | *RP11-501E14.1* | 1 | 37 | 3 | | | | | |
| chr1 | *FAM167B* | 1 | 55 | 3 | FAM167B | family with sequence similarity 167 member B | Other | other | |
| chr3 | *FECHP1* | 1 | 25 | 3 | FECHP1 | ferrochelatase pseudogene 1 | Other | other | |
| chr12 | *OR10P1* | 1 | 74 | 3 | OR10P1 | olfactory receptor family 10 subfamily P member 1 | Plasma Membrane | G-protein coupled recepto | |
| chr8 | *MIR596* | 1 | 166 | 3 | mir-596 | microRNA 596 | Cytoplasm | microRNA | |
| chr19 | *CTB-167G5.6* | 1 | 49 | 3 | | | | | |
| chr12 | *HOXC13* | 1 | 91 | 3 | HOXC13 | homeobox C13 | Nucleus | transcription regulator | |
| chr19 | *CTC-412M14.6* | 1 | 30 | 3 | | | | | |
| chr19 | *PCSK4* | 2 | 44 | 3 | PCSK4 | proprotein convertase subtilisin/kexin type 4 | Extracellular Space | peptidase | |
| chr1 | *GBP1* | 1 | 21 | 3 | GBP1 | guanylate binding protein 1 | Cytoplasm | enzyme | |
| chr9 | *BMS1P10* | 2 | 98 | 3 | BMS1P10 | BMS1 pseudogene 10 | Other | other | |
| chr19 | *AC008992.1* | 1 | 21 | 3 | | | | | |
| chr9 | *TNFSF15* | 1 | 21 | 3 | TNFSF15 | TNF superfamily member 15 | Extracellular Space | cytokine | |
| chr19 | *RP11-618P17.4* | 2 | 35 | 3 | | | | | |
| chr1 | *USF1* | 2 | 57 | 3 | USF1 | upstream transcription factor 1 | Nucleus | transcription regulator | |
| chr11 | *SAC3D1* | 2 | 50 | 3 | SAC3D1 | SAC3 domain containing 1 | Cytoplasm | other | |
| chr20 | *PPDPF* | 2 | 92 | 3 | PPDPF | pancreatic progenitor cell differentiation and proliferation | Other | other | |
| chr14 | *RP11-44N21.1* | 2 | 73 | 3 | | | | | |
| chr15 | *RP11-483E23.4* | 2 | 119 | 3 | | | | | |
| chr22 | *RP1-293L6.1* | 1 | 35 | 3 | | | | | |
| chr9 | *PSMD5* | 1 | 36 | 3 | PSMD5 | proteasome 26S subunit, non-ATPase 5 | Other | other | |
| chr10 | *RP11-131N11.4* | 1 | 28 | 3 | | | | | |
| chrX | *RP11-535K18.2* | 1 | 25 | 3 | | | | | |
| chr2 | *AC072062.1* | 1 | 21 | 3 | | | | | |
| chr6 | *RP11-203H2.1* | 1 | 25 | 3 | | | | | |
| chr4 | *RP11-366M4.8* | 1 | 27 | 3 | | | | | |
| chr14 | *PSME1* | 1 | 34 | 3 | PSME1 | proteasome activator subunit 1 | Cytoplasm | other | |
| chr1 | *RN7SL480P* | 1 | 40 | 3 | RN7SL480P | RNA, 7SL, cytoplasmic 480, pseudogene | Other | other | |
| chr12 | *HNF1A-AS1* | 1 | 39 | 3 | HNF1A-AS1 | HNF1A antisense RNA 1 | Other | other | |
| chr19 | *ZNF428* | 1 | 38 | 3 | ZNF428 | zinc finger protein 428 | Other | other | |
| chr6 | *HIST1H2BL* | 1 | 42 | 3 | H2BC13 | H2B clustered histone 13 | Nucleus | other | |
| chr1 | *RP11-538D16.2* | 1 | 38 | 3 | | | | | |
| chr5 | *RP11-772E11.1* | 1 | 37 | 3 | | | | | |
| chr7 | *TP53TG1* | 2 | 70 | 3 | TP53TG1 | TP53 target 1 | Other | other | |
| chr4 | *FRG1* | 2 | 53 | 3 | FRG1 | FSHD region gene 1 | Nucleus | transporter | |
| chr17 | *SPHK1* | 2 | 64 | 3 | SPHK1 | sphingosine kinase 1 | Cytoplasm | kinase | |
| chr19 | *CTC-457E21.4* | 1 | 41 | 3 | | | | | |
| chr11 | *MRGPRG-AS1* | 2 | 125 | 3 | MRGPRG-AS1 | MRGPRG antisense RNA 1 | Other | other | |
| chr1 | *ZNF691* | 2 | 81 | 3 | ZNF691 | zinc finger protein 691 | Nucleus | transcription regulator | |
| chr17 | *KRTAP4-1* | 1 | 50 | 3 | KRTAP4-1 | keratin associated protein 4-1 | Cytoplasm | other | |
| chr19 | *IZUMO4* | 2 | 68 | 3 | IZUMO4 | IZUMO family member 4 | Nucleus | other | |
| chr9 | *RP11-296L22.7* | 1 | 36 | 3 | | | | | |
| chr3 | *MYNN* | 2 | 116 | 3 | MYNN | myoneurin | Nucleus | transcription regulator | |
| chr3 | *PRKAR2A-AS1* | 1 | 43 | 3 | PRKAR2A-AS1 | PRKAR2A antisense RNA 1 | Other | other | |
| chr18 | *LMAN1* | 1 | 21 | 3 | LMAN1 | lectin, mannose binding 1 | Cytoplasm | other | |
| chr22 | *RASL10A* | 1 | 42 | 3 | RASL10A | RAS like family 10 member A | Plasma Membrane | enzyme | |
| chr12 | *RP11-713N11.5* | 1 | 26 | 3 | | | | | |
| chr11 | *RP11-111M22.3* | 1 | 38 | 3 | | | | | |
| chr4 | *USP17L27* | 1 | 49 | 3 | 17L18 (includes ot | ubiquitin specific peptidase 17 like family member 5 | Nucleus | peptidase | |
| chr17 | *RP11-173M1.8* | 1 | 27 | 3 | | | | | |
| chr11 | *PTGDR2* | 2 | 56 | 3 | PTGDR2 | prostaglandin D2 receptor 2 | Plasma Membrane | G-protein coupled recepto | fevipiprant |
| chr9 | *RP11-237M21.1* | 1 | 25 | 3 | | | | | |
| chr17 | *KRTAP9-12P* | 1 | 45 | 3 | KRTAP9-12P | keratin associated protein 9-12, pseudogene | Other | other | |
| chr15 | *RP11-622C24.1* | 1 | 52 | 3 | | | | | |
| chr12 | *C12orf74* | 1 | 35 | 3 | C12orf74 | chromosome 12 open reading frame 74 | Other | other | |
| chr15 | *RP11-632K20.6* | 1 | 41 | 3 | | | | | |
| chr13 | *MRPS31P2* | 2 | 39 | 3 | MRPS31P2 | mitochondrial ribosomal protein S31 pseudogene 2 | Other | other | |
| chr7 | *RP11-758P17.3* | 1 | 62 | 3 | | | | | |
| chr5 | *CTD-2008N3.1* | 2 | 87 | 3 | | | | | |
| chr22 | *OSM* | 2 | 77 | 3 | OSM | oncostatin M | Extracellular Space | cytokine | |
| chr16 | *COQ9* | 1 | 36 | 3 | COQ9 | coenzyme Q9 | Cytoplasm | other | |
| chr1 | *RP11-567C20.3* | 1 | 31 | 3 | | | | | |
| chr7 | *WDR86-AS1* | 1 | 52 | 3 | WDR86-AS1 | WDR86 antisense RNA 1 | Other | other | |
| chr20 | *CST9* | 1 | 27 | 3 | CST9 | cystatin 9 | Extracellular Space | other | |
| chr12 | *RP11-474D1.4* | 1 | 43 | 3 | | | | | |
| chr15 | *NMB* | 2 | 64 | 3 | NMB | neuromedin B | Extracellular Space | other | |
| chr8 | *GOT1L1* | 1 | 26 | 3 | GOT1L1 | glutamic-oxaloacetic transaminase 1 like 1 | Other | enzyme | |
| chr18 | *RP11-64C12.9* | 1 | 35 | 3 | | | | | |
| chr12 | *FICD* | 1 | 60 | 3 | FICD | FIC domain containing | Nucleus | enzyme | |
| chr1 | *RP11-9L18.2* | 1 | 44 | 3 | | | | | |

| | | | | | | | | | |
|---|---|---|---|---|---|---|---|---|---|
| chr3 | *UBE2E2-AS1* | 2 | 57 | 3 | UBE2E2-AS1 | UBE2E2 antisense RNA 1 (head to head) | Other | other | |
| chr7 | *AC013436.6* | 1 | 52 | 3 | | | | | |
| chr1 | *CTSK* | 1 | 35 | 3 | CTSK | cathepsin K | Cytoplasm | peptidase | cathepsin K inhibitor, odanacatib, balicatib |
| chr3 | *RP11-723O4.9* | 1 | 44 | 3 | | | | | |
| chr11 | *SSU72P6* | 1 | 123 | 3 | SSU72P6 | SSU72 pseudogene 6 | Other | other | |
| chr21 | *C21orf91-OT1* | 1 | 36 | 3 | C21orf91-OT1 | C21orf91 overlapping transcript 1 | Other | other | |
| chr1 | *RAB13* | 1 | 47 | 3 | RAB13 | RAB13, member RAS oncogene family | Plasma Membrane | enzyme | |
| chr9 | *C9orf169* | 2 | 153 | 3 | CYSRT1 | cysteine rich tail 1 | Cytoplasm | other | |
| chr20 | *PDRG1* | 1 | 48 | 3 | PDRG1 | p53 and DNA damage regulated | Cytoplasm | other | |
| chr15 | *RP11-20G13.2* | 1 | 41 | 3 | | | | | |
| chr5 | *EEF1GP2* | 1 | 37 | 3 | EEF1GP2 | eukaryotic translation elongation factor 1 gamma pseudogene 2 | Other | other | |
| chr14 | *RN7SL356P* | 1 | 41 | 3 | RN7SL356P | | Other | other | |
| chr6 | *UBE2J1* | 1 | 20 | 3 | UBE2J1 | ubiquitin conjugating enzyme E2 | Cytoplasm | enzyme | |
| chr2 | *AC018731.3* | 1 | 20 | 3 | | | | | |
| chrY | *AJ271736.10* | 1 | 38 | 3 | | | | | |
| chr16 | *ZP2* | 1 | 25 | 3 | ZP2 | zona pellucida glycoprotein 2 | Extracellular Space | other | |
| chr15 | *BMS1P16* | 2 | 70 | 3 | BMS1P16 | BMS1 pseudogene 16 | Other | other | |
| chr8 | *RP11-697M17.2* | 1 | 23 | 3 | | | | | |
| chr1 | *RP11-168O16.2* | 1 | 64 | 3 | | | | | |
| chr1 | *GBP3* | 1 | 24 | 3 | GBP3 | guanylate binding protein 3 | Cytoplasm | enzyme | |
| chrX | *RP13-13A3.1* | 1 | 49 | 3 | | | | | |
| chr16 | *RP11-19N8.7* | 1 | 29 | 3 | | | | | |
| chr22 | *NDUFA6* | 1 | 50 | 3 | NDUFA6 | NADH:ubiquinone oxidoreductase subunit A6 | Cytoplasm | enzyme | |
| chr5 | *PCDHB1* | 1 | 46 | 3 | PCDHB1 | protocadherin beta 1 | Plasma Membrane | other | |
| chr4 | *CLUHP4* | 1 | 78 | 3 | CLUHP4 | clustered mitochondria homolog pseudogene 4 | Other | other | |
| chr3 | *RP11-58B17.2* | 1 | 53 | 3 | | | | | |
| chr10 | *GSTO1* | 2 | 42 | 3 | GSTO1 | glutathione S-transferase omega | Cytoplasm | enzyme | |
| chr3 | *KRT18P43* | 1 | 67 | 3 | KRT18P43 | keratin 18 pseudogene 43 | Other | other | |
| chr1 | *RP11-183G22.1* | 1 | 45 | 3 | | | | | |
| chr9 | *KCNV2* | 2 | 58 | 3 | KCNV2 | potassium voltage-gated channel modifier subfamily V member 2 | Plasma Membrane | ion channel | |
| chr17 | *AC005822.1* | 2 | 54 | 3 | | | | | |
| chr16 | *PPP1R1AP2* | 1 | 29 | 3 | PPP1R1AP2 | protein phosphatase 1 regulatory inhibitor subunit 1A pseudogene | Other | other | |
| chr9 | *CNN2P2* | 1 | 49 | 3 | CNN2P2 | calponin 2 pseudogene 2 | Other | other | |
| chr5 | *RP11-232L2.1* | 1 | 35 | 3 | | | | | |
| chr1 | *FAM110D* | 1 | 47 | 3 | FAM110D | family with sequence similarity 110 member D | Other | other | |
| chr19 | *CLEC11A* | 2 | 50 | 3 | CLEC11A | C-type lectin domain containing | Extracellular Space | growth factor | |
| chrX | *FAM47B* | 1 | 54 | 3 | FAM47B | family with sequence similarity 47 member B | Other | other | |
| chr1 | *ERVMER61-1* | 1 | 31 | 3 | | | | | |
| chr10 | *RP11-129J12.2* | 2 | 91 | 3 | | | | | |
| chrX | *PAK3* | 2 | 49 | 3 | PAK3 | p21 (RAC1) activated kinase 3 | Cytoplasm | kinase | |
| chr17 | *RP11-147L13.7* | 1 | 36 | 3 | | | | | |
| chr9 | *ZDHHC12* | 2 | 137 | 3 | ZDHHC12 | zinc finger DHHC-type containing | Other | enzyme | |
| chr17 | *RP11-690G19.3* | 1 | 28 | 3 | | | | | |
| chr3 | *RP11-408H1.3* | 1 | 20 | 3 | | | | | |
| chr19 | *CEACAMP3* | 1 | 35 | 3 | CEACAMP3 | CEA cell adhesion molecule pseudogene 3 | Other | other | |
| chr1 | *TEKT2* | 1 | 60 | 3 | TEKT2 | tektin 2 | Cytoplasm | other | |
| chr1 | *RP5-1057J7.1* | 1 | 49 | 3 | | | | | |
| chr3 | *SMIM4* | 2 | 60 | 3 | SMIM4 | small integral membrane protein | Cytoplasm | other | |
| chr12 | *METTL7AP1* | 1 | 72 | 3 | METTL7AP1 | methyltransferase like 7A pseudogene 1 | Other | other | |
| chr3 | *CRBN* | 1 | 20 | 3 | CRBN | cereblon | Cytoplasm | enzyme | dexamethasone/lenalidomide, iberdomide, bortezomib/thalidomide, lenalidomide, bendamustine/lenalidomide, CC-92480, CC-122, bortezomib/dexamethasone/pomalidomide, bortezomib/dexamethasone/thalidomide, thalidomide, rituximab/thalidomide, cereblon-binding agent, lenalidomide/rituximab, cetuximab/lenalidomide, erythropoietin/lenalidomide, bortezomib/lenalidom[illegible] |
| chr1 | *PRG4* | 1 | 49 | 3 | PRG4 | proteoglycan 4 | Extracellular Space | other | |
| chr3 | *NRADDP* | 1 | 49 | 3 | NRADDP | neurotrophin receptor associated death domain, pseudogene | Other | other | |

| | | | | | | | | | |
|---|---|---|---|---|---|---|---|---|---|
| chr5 | *TMED9* | 1 | 43 | 3 | TMED9 | transmembrane p24 trafficking protein 9 | Cytoplasm | transporter | |
| chr10 | *FAM35EP* | 1 | 24 | 3 | SHLD2P1 | shieldin complex subunit 2 pseudogene 1 | Other | other | |
| chr11 | *SLC25A22* | 2 | 81 | 3 | SLC25A22 | solute carrier family 25 member | Cytoplasm | transporter | |
| chr18 | *CPLX4* | 1 | 21 | 3 | CPLX4 | complexin 4 | Plasma Membrane | other | |
| chr10 | *WAC-AS1* | 1 | 25 | 3 | WAC-AS1 | WAC antisense RNA 1 (head to head) | Other | other | |
| chr21 | *AL035610.1* | 1 | 22 | 3 | | | | | |
| chr12 | *CLEC4D* | 1 | 39 | 3 | CLEC4D | C-type lectin domain family 4 member D | Plasma Membrane | other | |
| chr6 | *HIST1H2APS3* | 1 | 46 | 3 | H2AC9P | H2A clustered histone 9, pseudogene | Other | other | |
| chr11 | *KCNK7* | 2 | 111 | 3 | KCNK7 | potassium two pore domain channel subfamily K member 7 | Plasma Membrane | ion channel | |
| chr1 | *SCARNA18* | 1 | 39 | 3 | SCARNA18 | small Cajal body-specific RNA 18 | Other | other | |
| chr18 | *RP11-672L10.3* | 1 | 26 | 3 | | | | | |
| chr10 | *TLX1* | 2 | 95 | 3 | TLX1 | T cell leukemia homeobox 1 | Nucleus | transcription regulator | |
| chr12 | *RP11-297L6.1* | 1 | 39 | 3 | | | | | |
| chr9 | *C9orf163* | 1 | 79 | 3 | C9orf163 | chromosome 9 open reading frame 163 | Other | other | |
| chr9 | *QRFP* | 2 | 121 | 3 | QRFP | pyroglutamylated RFamide | Extracellular Space | other | |
| chr9 | *CA9* | 1 | 37 | 3 | CA9 | carbonic anhydrase 9 | Nucleus | enzyme | benzthiazide, I 131 chimeric G250, methazolamide, hydrochlorothiazide, lutetium Lu-177 girentuximab, DTP348, sulfacetamide, SLC-0111, topiramate, girentuximab, chlorothiazide, iodine I-124 girentuximab, chlorthalidone, acetazolamide, Y 90 chimeric G250 |
| chr7 | *C7orf49* | 1 | 64 | 3 | CYREN | cell cycle regulator of NHEJ | Other | other | |
| chr21 | *AF064858.11* | 1 | 34 | 3 | | | | | |
| chr9 | *C9orf69* | 2 | 138 | 3 | TMEM250 | transmembrane protein 250 | Nucleus | other | |
| chr22 | *PITPNB* | 1 | 20 | 3 | PITPNB | phosphatidylinositol transfer protein beta | Cytoplasm | transporter | |
| chr14 | *RP11-526N18.1* | 1 | 20 | 3 | | | | | |
| chr3 | *RP11-44H4.1* | 1 | 33 | 3 | | | | | |
| chr9 | *RP11-187C18.4* | 1 | 38 | 3 | | | | | |
| chr7 | *TRBV6-8* | 1 | 40 | 3 | TRBV6-8 | T cell receptor beta variable 6-8 | Other | other | |
| chr11 | *GSTP1* | 1 | 54 | 3 | GSTP1 | glutathione S-transferase pi 1 | Cytoplasm | enzyme | |
| chr2 | *CNN2P8* | 1 | 66 | 3 | CNN2P8 | calponin 2 pseudogene 8 | Other | other | |
| chr1 | *RP5-891H21.1* | 1 | 40 | 3 | | | | | |
| chr19 | *RPS18P13* | 1 | 50 | 3 | RPS18P13 | ribosomal protein S18 pseudogene 13 | Other | other | |
| chr2 | *GALNT5* | 1 | 22 | 3 | GALNT5 | polypeptide N-acetylgalactosaminyltransferase | Cytoplasm | enzyme | |
| chr10 | *RP11-295P9.3* | 1 | 20 | 3 | | | | | |
| chr1 | *RP11-439E19.7* | 1 | 36 | 3 | | | | | |
| chr7 | *MIR182* | 1 | 58 | 3 | mir-182 | microRNA 182 | Cytoplasm | microRNA | |
| chr1 | *RP4-782G3.1* | 1 | 25 | 3 | | | | | |
| chrX | *RP11-308D16.4* | 1 | 24 | 3 | | | | | |
| chr5 | *CH17-12M21.1* | 1 | 43 | 3 | | | | | |
| chr17 | *KPNA2* | 1 | 32 | 3 | KPNA2 | karyopherin subunit alpha 2 | Nucleus | transporter | |
| chr7 | *ATP6V0E2-AS1* | 1 | 49 | 3 | ATP6V0E2-AS1 | ATP6V0E2 antisense RNA 1 | Other | other | |
| chr5 | *RP11-436H11.1* | 1 | 34 | 3 | | | | | |
| chr1 | *RP11-319C21.1* | 1 | 33 | 3 | | | | | |
| chr15 | *RP11-621H8.2* | 1 | 49 | 3 | | | | | |
| chrX | *SRPX2* | 1 | 23 | 3 | SRPX2 | sushi repeat containing protein X-linked 2 | Cytoplasm | other | |
| chr2 | *PEBP1P2* | 1 | 45 | 3 | PEBP1P2 | phosphatidylethanolamine binding protein 1 pseudogene 2 | Other | other | |
| chr6 | *ENPP5* | 1 | 22 | 3 | ENPP5 | ectonucleotide pyrophosphatase/phosphodiesterase family member 5 | Extracellular Space | enzyme | |
| chr16 | *RP11-462G12.1* | 1 | 45 | 3 | | | | | |
| chrX | *GPRASP2* | 1 | 27 | 3 | X5-GPRASP2/GP | G protein-coupled receptor associated sorting protein 2 | Plasma Membrane | other | |
| chr7 | *RP11-796I2.1* | 1 | 29 | 3 | | | | | |
| chr1 | *RP4-591L5.2* | 1 | 45 | 3 | | | | | |
| chr19 | *TMED1* | 1 | 46 | 3 | TMED1 | transmembrane p24 trafficking protein 1 | Extracellular Space | transporter | |
| chr21 | *ABCC13* | 1 | 20 | 3 | ABCC13 | ATP binding cassette subfamily C member 13 (pseudogene) | Other | other | |
| chr18 | *RP11-672L10.4* | 1 | 29 | 3 | | | | | |
| chr1 | *PRPF38A* | 1 | 31 | 3 | PRPF38A | pre-mRNA processing factor 38A | Nucleus | other | |
| chr17 | *C17orf80* | 1 | 29 | 3 | C17orf80 | chromosome 17 open reading frame 80 | Cytoplasm | other | |
| chr2 | *AC104695.4* | 1 | 39 | 3 | | | | | |
| chr10 | *RPL13AP5* | 1 | 42 | 3 | RPL13AP5 | ribosomal protein L13a pseudogene 5 | Other | other | |
| chr12 | *RP11-603J24.4* | 1 | 55 | 3 | | | | | |

| | | | | | | | | | |
|---|---|---|---|---|---|---|---|---|---|
| chr4 | *LINC00575* | 1 | 29 | 3 | LINC00575 | long intergenic non-protein coding RNA 575 | Other | other | |
| chr12 | *RP11-467L13.5* | 1 | 40 | 3 | | | | | |
| chr11 | *RP11-672A2.5* | 1 | 72 | 3 | | | | | |
| chr19 | *ACPT* | 2 | 52 | 3 | ACP4 | acid phosphatase 4 | Plasma Membrane | phosphatase | |
| chr2 | *CRYGB* | 1 | 31 | 3 | CRYGB | crystallin gamma B | Nucleus | other | |
| chr17 | *ARHGDIA* | 2 | 104 | 3 | ARHGDIA | Rho GDP dissociation inhibitor | Cytoplasm | other | |
| chr11 | *AP000974.1* | 2 | 90 | 3 | | | | | |
| chr17 | *RP11-147L13.2* | 1 | 43 | 3 | | | | | |
| chr3 | *ALDH1L1-AS2* | 2 | 54 | 3 | ALDH1L1-AS2 | ALDH1L1 antisense RNA 2 | Other | other | |
| chr5 | *CTB-47B8.1* | 1 | 43 | 3 | | | | | |
| chr16 | *RP11-243A14.3* | 1 | 22 | 3 | | | | | |
| chr9 | *RP11-432J22.2* | 2 | 61 | 3 | | | | | |
| chrX | *TLR8-AS1* | 1 | 37 | 3 | TLR8-AS1 | TLR8 antisense RNA 1 | Other | other | |
| chr12 | *RP11-722P11.4* | 2 | 88 | 3 | | | | | |
| chr16 | *OR2C1* | 1 | 45 | 3 | OR2C1 | olfactory receptor family 2 subfamily C member 1 | Plasma Membrane | G-protein coupled recepto | |
| chr22 | *AC000067.1* | 1 | 35 | 3 | | | | | |
| chr9 | *C9orf170* | 2 | 63 | 3 | C9orf170 | chromosome 9 open reading frame 170 | Other | other | |
| chr11 | *AP003774.1* | 1 | 28 | 3 | | | | | |
| chr3 | *MRPL3* | 1 | 22 | 3 | MRPL3 | mitochondrial ribosomal protein | Cytoplasm | other | |
| chr10 | *ACTBP14* | 1 | 99 | 3 | ACTBP14 | ACTB pseudogene 14 | Other | other | |
| chr7 | *MUC3A* | 2 | 646 | 3 | MUC3A | mucin 3A, cell surface associated | Other | other | |
| chr7 | *FASTK* | 2 | 100 | 3 | FASTK | Fas activated serine/threonine | Cytoplasm | kinase | |
| chr2 | *MTX2* | 1 | 25 | 3 | MTX2 | metaxin 2 | Cytoplasm | transporter | |
| chr4 | *CNOT6L* | 1 | 28 | 3 | CNOT6L | CCR4-NOT transcription complex subunit 6 like | Cytoplasm | enzyme | |
| chr5 | *RP11-321E2.10* | 1 | 139 | 3 | | | | | |
| chr6 | *TDRD6* | 1 | 20 | 3 | TDRD6 | tudor domain containing 6 | Cytoplasm | other | |
| chr17 | *RAC3* | 2 | 67 | 3 | RAC3 | Rac family small GTPase 3 | Cytoplasm | enzyme | |
| chr5 | *RP11-497H16.7* | 1 | 26 | 3 | | | | | |
| chr15 | *RP11-317N6.1* | 1 | 25 | 3 | | | | | |
| chr14 | *SUPT16H* | 1 | 21 | 3 | SUPT16H | SPT16 homolog, facilitates chromatin remodeling subunit | Nucleus | transcription regulator | |
| chr7 | *RARRES2* | 2 | 47 | 3 | RARRES2 | retinoic acid receptor responder 2 | Plasma Membrane | transmembrane recepto | |
| chr1 | *GBP1P1* | 2 | 53 | 3 | GBP1P1 | guanylate binding protein 1 pseudogene 1 | Other | other | |
| chr14 | *RP11-218E20.2* | 1 | 23 | 3 | | | | | |
| chr3 | *CCR4* | 1 | 23 | 3 | CCR4 | C-C motif chemokine receptor 4 | Plasma Membrane | G-protein coupled recepto | FLX475, mogamulizumab |
| chr12 | *RPL22P19* | 1 | 37 | 3 | RPL22P19 | ribosomal protein L22 pseudogene 19 | Other | other | |
| chr16 | *RP11-327F22.2* | 1 | 28 | 3 | | | | | |
| chr19 | *ADAMTSL5* | 2 | 68 | 3 | ADAMTSL5 | ADAMTS like 5 | Extracellular Space | other | |
| chr15 | *AC112693.2* | 1 | 57 | 3 | | | | | |
| chr3 | *RP11-305F5.2* | 1 | 54 | 3 | | | | | |
| chr9 | *RP11-336N8.1* | 1 | 48 | 3 | | | | | |
| chr3 | *ATP13A5* | 1 | 22 | 3 | ATP13A5 | ATPase 13A5 | Plasma Membrane | transporter | |
| chr18 | *RP11-1030E3.1* | 1 | 23 | 3 | | | | | |
| chr1 | *RP5-893G23.1* | 1 | 44 | 3 | | | | | |
| chr9 | *DPP7* | 2 | 98 | 3 | DPP7 | dipeptidyl peptidase 7 | Cytoplasm | peptidase | |
| chr19 | *CNTD2* | 2 | 90 | 3 | CNTD2 | cyclin N-terminal domain | Other | other | |
| chr8 | *RP11-380I10.3* | 1 | 36 | 3 | | | | | |
| chr16 | *RP11-989E6.3* | 1 | 53 | 3 | | | | | |
| chr6 | *TXLNB* | 1 | 25 | 3 | TXLNB | taxilin beta | Cytoplasm | other | |
| chr1 | *OR2T12* | 1 | 38 | 3 | OR2T12 | olfactory receptor family 2 subfamily T member 12 | Plasma Membrane | other | |
| chr19 | *OR1I1* | 1 | 41 | 3 | OR1I1 | olfactory receptor family 1 subfamily I member 1 | Plasma Membrane | G-protein coupled recepto | |
| chr17 | *ZNHIT3* | 1 | 42 | 3 | ZNHIT3 | zinc finger HIT-type containing 3 | Nucleus | transcription regulator | |
| chr4 | *TMEM156* | 1 | 23 | 3 | TMEM156 | transmembrane protein 156 | Other | other | |
| chr4 | *AC007106.1* | 1 | 25 | 3 | | | | | |
| chr5 | *RP11-267A15.3* | 1 | 37 | 3 | | | | | |
| chr5 | *RP11-321E2.7* | 1 | 952 | 3 | | | | | |
| chr11 | *TRIM49B* | 1 | 44 | 3 | TRIM49B | tripartite motif containing 49B | Other | other | |
| chr1 | *RP4-798A10.4* | 2 | 88 | 3 | | | | | |
| chr2 | *NRBP1* | 1 | 25 | 3 | NRBP1 | nuclear receptor binding protein 1 | Nucleus | kinase | |
| chr1 | *LCE3C* | 1 | 41 | 3 | LCE3C | late cornified envelope 3C | Other | other | |
| chr1 | *WI2-925H4.1* | 1 | 43 | 3 | | | | | |
| chr11 | *RP11-81M19.3* | 2 | 160 | 3 | | | | | |
| chr1 | *U8* | 1 | 59 | 3 | | | | | |
| chr2 | *TRIM43* | 1 | 37 | 3 | TRIM43/TRIM43B | tripartite motif containing 43 | Other | other | |
| chr17 | *KRTAP9-8* | 1 | 52 | 3 | KRTAP9-8 | keratin associated protein 9-8 | Cytoplasm | other | |
| chr4 | *RP11-103J17.1* | 1 | 19 | 3 | | | | | |
| chr1 | *RP11-495P10.9* | 1 | 58 | 3 | | | | | |
| chr1 | *AC099791.1* | 1 | 38 | 3 | | | | | |
| chr15 | *RP11-261B23.1* | 1 | 61 | 3 | | | | | |
| chr11 | *PDZD3* | 1 | 55 | 3 | PDZD3 | PDZ domain containing 3 | Plasma Membrane | transporter | |
| chr5 | *RP11-661C8.3* | 2 | 130 | 3 | | | | | |
| chr8 | *RP11-267L5.1* | 1 | 40 | 3 | | | | | |
| chr16 | *HBZ* | 1 | 54 | 3 | HBZ | hemoglobin subunit zeta | Cytoplasm | transporter | |
| chr2 | *AC018804.7* | 1 | 57 | 3 | | | | | |
| chr11 | *RP11-787P24.4* | 1 | 21 | 3 | | | | | |
| chr12 | *DHH* | 2 | 105 | 3 | DHH | desert hedgehog signaling | Extracellular Space | peptidase | |
| chr3 | *RP4-613B23.1* | 1 | 45 | 3 | | | | | |
| chr1 | *RP4-803A2.1* | 1 | 49 | 3 | | | | | |
| chr2 | *MYCN* | 1 | 28 | 3 | MYCN | MYCN proto-oncogene, bHLH transcription factor | Nucleus | transcription regulator | |

| | | | | | | | | | |
|---|---|---|---|---|---|---|---|---|---|
| chr5 | *CTD-2199O4.1* | 1 | 85 | 3 | | | | | |
| chr19 | *SIGLEC18P* | 1 | 44 | 3 | SIGLEC18P | sialic acid binding Ig like lectin 18, pseudogene | Other | other | |
| chr3 | *KRT18P35* | 1 | 40 | 3 | KRT18P35 | keratin 18 pseudogene 35 | Other | other | |
| chrX | *FANCB* | 1 | 23 | 3 | FANCB | FA complementation group B | Nucleus | other | |
| chr1 | *TXNIP* | 1 | 47 | 3 | TXNIP | thioredoxin interacting protein | Cytoplasm | other | |
| chr5 | *STARD4* | 1 | 23 | 3 | STARD4 | StAR related lipid transfer domain containing 4 | Cytoplasm | transporter | |
| chr19 | *FBXL12* | 1 | 31 | 3 | FBXL12 | F-box and leucine rich repeat protein 12 | Cytoplasm | other | |
| chr10 | *RP11-336A10.4* | 1 | 36 | 3 | | | | | |
| chr16 | *LA16c-380H5.5* | 1 | 44 | 3 | | | | | |
| chr7 | *SEPT7P5* | 1 | 35 | 3 | SEPTIN7P5 | septin 7 pseudogene 5 | Other | other | |
| chr7 | *FZD9* | 1 | 56 | 3 | FZD9 | frizzled class receptor 9 | Plasma Membrane | -protein coupled recepto | |
| chrX | *SUPT20HL1* | 1 | 50 | 3 | SUPT20HL1 | SUPT20H like 1 | Other | other | |
| chr11 | *RP11-60C6.3* | 1 | 83 | 3 | | | | | |
| chrX | *RP11-552J9.11* | 1 | 37 | 3 | | | | | |
| chr2 | *CTD-2026C7.1* | 1 | 31 | 3 | | | | | |
| chr10 | *RP11-363N22.2* | 1 | 26 | 3 | | | | | |
| chr7 | *PNPT1P2* | 1 | 25 | 3 | PNPT1P2 | polyribonucleotide nucleotidyltransferase 1 pseudogene 2 | Other | other | |
| chr5 | *CTC-512J14.5* | 1 | 40 | 3 | | | | | |
| chr21 | *AP001065.15* | 1 | 37 | 3 | | | | | |
| chr9 | *TEX10* | 1 | 24 | 3 | TEX10 | testis expressed 10 | Nucleus | other | |
| chr19 | *ZNF426* | 1 | 26 | 3 | ZNF426 | zinc finger protein 426 | Nucleus | transcription regulator | |
| chr2 | *AC007163.6* | 1 | 20 | 3 | | | | | |
| chr16 | *RP11-77H9.8* | 1 | 42 | 3 | | | | | |
| chr10 | *GHITM* | 1 | 20 | 3 | GHITM | growth hormone inducible transmembrane protein | Cytoplasm | other | |
| chr15 | *CTD-2651B20.2* | 1 | 35 | 3 | | | | | |
| chr18 | *RP11-1151B14.2* | 1 | 32 | 3 | | | | | |
| chr17 | *RILP* | 1 | 80 | 3 | RILP | Rab interacting lysosomal protein | Cytoplasm | other | |
| chr1 | *CD1E* | 1 | 35 | 3 | CD1E | CD1e molecule | Cytoplasm | other | |
| chr16 | *RP11-80F22.7* | 1 | 39 | 3 | | | | | |
| chr19 | *IGFL1P2* | 1 | 36 | 3 | IGFL1P2 | IGF like family member 1 pseudogene 2 | Other | other | |
| chr5 | *AC005355.2* | 1 | 53 | 3 | | | | | |
| chr17 | *RP11-1096G20.5* | 1 | 35 | 3 | | | | | |
| chr4 | *RP11-18H21.3* | 1 | 26 | 3 | | | | | |
| chr5 | *SLC36A3* | 1 | 21 | 3 | SLC36A3 | solute carrier family 36 member 3 | Other | transporter | |
| chr13 | *PRR20B* | 2 | 255 | 3 | R20A (includes oth | proline rich 20A | Other | other | |
| chr20 | *TRERNA1* | 1 | 63 | 3 | TRERNA1 | translation regulatory long non-coding RNA 1 | Other | other | |
| chr1 | *RP11-404O13.1* | 1 | 33 | 3 | | | | | |
| chr12 | *RP11-662I13.1* | 1 | 40 | 3 | | | | | |
| chr13 | *LINC00396* | 1 | 40 | 3 | LINC00396 | long intergenic non-protein coding RNA 396 | Other | other | |
| chr1 | *RN7SL649P* | 1 | 41 | 3 | RN7SL649P | RNA, 7SL, cytoplasmic 649, pseudogene | Other | other | |
| chr2 | *AC013733.3* | 1 | 28 | 3 | | | | | |
| chr9 | *RP11-57C19.2* | 1 | 56 | 3 | | | | | |
| chr5 | *CTC-232H10.1* | 1 | 111 | 3 | | | | | |
| chr10 | *AHCYP1* | 1 | 71 | 3 | | | | | |
| chr16 | *AKTIP* | 2 | 52 | 3 | AKTIP | AKT interacting protein | Cytoplasm | other | |
| chr2 | *AC009965.2* | 1 | 31 | 3 | | | | | |
| chr2 | *AC012451.1* | 1 | 36 | 3 | | | | | |
| chr17 | *RP5-890E16.2* | 1 | 53 | 3 | | | | | |
| chr19 | *AC016586.1* | 2 | 46 | 3 | | | | | |
| chr22 | *CECR7* | 1 | 28 | 3 | CECR7 | cat eye syndrome chromosome region, candidate 7 | Other | other | |
| chr17 | *RPS20P35* | 1 | 45 | 3 | RPS20P35 | ribosomal protein S20 pseudogene 35 | Other | other | |
| chr20 | *RN7SL31P* | 1 | 36 | 3 | | | | | |
| chr10 | *TFAM* | 1 | 39 | 3 | TFAM | transcription factor A, | Cytoplasm | transcription regulator | |
| chr6 | *RP1-200K18.1* | 1 | 21 | 3 | | | | | |
| chr16 | *RP11-46D6.4* | 1 | 44 | 3 | | | | | |
| chr1 | *CFL1P6* | 1 | 32 | 3 | CFL1P6 | cofilin 1 pseudogene 6 | Other | other | |
| chr4 | *RP11-20I20.2* | 1 | 25 | 3 | | | | | |
| chr12 | *LYZ* | 1 | 22 | 3 | LYZ | lysozyme | Extracellular Space | enzyme | |
| chr9 | *ALDH1B1* | 1 | 37 | 3 | ALDH1B1 | aldehyde dehydrogenase 1 family member B1 | Cytoplasm | enzyme | |
| chr16 | *RNU6-416P* | 1 | 34 | 3 | RNU6-416P | RNA, U6 small nuclear 416, pseudogene | Other | other | |
| chr19 | *TMEM221* | 1 | 40 | 3 | TMEM221 | transmembrane protein 221 | Other | other | |
| chr7 | *GUSBP10* | 1 | 45 | 3 | GUSBP10 | GUSB pseudogene 10 | Other | other | |
| chr7 | *AC091705.1* | 1 | 36 | 3 | | | | | |
| chr16 | *MT1E* | 1 | 43 | 3 | MT1E | metallothionein 1E | Other | other | |
| chr4 | *RP11-1281K21.2* | 1 | 96 | 3 | | | | | |
| chr19 | *AC005336.5* | 1 | 55 | 3 | | | | | |
| chr6 | *CNR1* | 1 | 32 | 3 | CNR1 | cannabinoid receptor 1 | Plasma Membrane | -protein coupled recepto | delta-8-tetrahydrocannabinol, BAY 38-7271, SLV 319, delta-9-tetrahydrocannabinol, rimonabant, trans-(±)-nabilone |
| chr11 | *RP11-379P15.1* | 1 | 34 | 3 | | | | | |
| chr19 | *AC005789.11* | 1 | 46 | 3 | | | | | |

| | | | | | | | | |
|---|---|---|---|---|---|---|---|---|
| chr3 | *TATDN2* | 1 | 42 | 3 | TATDN2 | TatD DNase domain containing 2 | Nucleus | other |
| chr15 | *CTD-3094K11.1* | 1 | 104 | 3 | | | | |
| chr8 | *GS1-24F4.3* | 1 | 41 | 3 | | | | |
| chr7 | *RP11-395B7.7* | 1 | 39 | 3 | | | | |
| chr11 | *OR10S1* | 1 | 45 | 3 | OR10S1 | olfactory receptor family 10 subfamily S member 1 | Plasma Membrane | G-protein coupled recepto |
| chr17 | *SLFN14* | 1 | 25 | 3 | SLFN14 | schlafen family member 14 | Nucleus | enzyme |
| chr5 | *RP11-138J23.1* | 1 | 44 | 3 | | | | |
| chr1 | *RP11-335E6.2* | 1 | 23 | 3 | | | | |
| chr11 | *AP000593.5* | 1 | 76 | 3 | | | | |
| chr12 | *LINC00941* | 1 | 48 | 3 | LINC00941 | long intergenic non-protein coding RNA 941 | Other | other |
| chr11 | *PNPLA2* | 1 | 47 | 3 | PNPLA2 | patatin like phospholipase domain containing 2 | Cytoplasm | enzyme |
| chr2 | *AC098824.6* | 1 | 65 | 3 | | | | |
| chr16 | *FAM96B* | 1 | 44 | 3 | CIAO2B | cytosolic iron-sulfur assembly component 2B | Cytoplasm | other |
| chr14 | *TMEM260* | 1 | 20 | 3 | TMEM260 | transmembrane protein 260 | Other | other |
| chr6 | *RN7SL403P* | 1 | 34 | 3 | RN7SL403P | | Other | other |
| chr1 | *TSEN15* | 2 | 67 | 3 | TSEN15 | tRNA splicing endonuclease subunit 15 | Nucleus | enzyme |
| chr17 | *CSF3* | 1 | 55 | 3 | CSF3 | colony stimulating factor 3 | Extracellular Space | cytokine |
| chr11 | *RP13-631K18.5* | 1 | 50 | 3 | | | | |
| chr12 | *RP11-644F5.12* | 1 | 67 | 3 | | | | |
| chr5 | *RP11-43F13.4* | 2 | 140 | 3 | | | | |
| chr18 | *RP11-905K4.1* | 1 | 19 | 3 | | | | |
| chr3 | *UBQLN4P1* | 1 | 76 | 3 | UBQLN4P1 | ubiquilin 4 pseudogene 1 | Other | other |
| chr3 | *RP11-211G3.3* | 1 | 23 | 3 | | | | |
| chr1 | *RP11-61J19.3* | 1 | 37 | 3 | | | | |
| chr7 | *RN7SL76P* | 1 | 36 | 3 | RN7SL76P | RNA, 7SL, cytoplasmic 76, pseudogene | Other | other |
| chr2 | *AC017104.1* | 1 | 26 | 3 | | | | |
| chr11 | *RP11-702F3.2* | 1 | 69 | 3 | | | | |
| chr19 | *CTD-3105H18.18* | 1 | 26 | 3 | | | | |
| chr18 | *RPIAP1* | 1 | 26 | 3 | RPIAP1 | ribose 5-phosphate isomerase A pseudogene 1 | Other | other |
| chr2 | *AC007041.2* | 1 | 30 | 3 | | | | |
| chrX | *AFF2* | 1 | 19 | 3 | AFF2 | AF4/FMR2 family member 2 | Nucleus | other |
| chr6 | *QKI* | 1 | 19 | 3 | QKI | QKI, KH domain containing RNA binding | Nucleus | other |
| chr3 | *LINC00488* | 1 | 29 | 3 | LINC00488 | long intergenic non-protein coding RNA 488 | Other | other |
| chr16 | *RP11-159J3.1* | 1 | 43 | 3 | | | | |
| chr22 | *ZDHHC8P1* | 2 | 65 | 3 | ZDHHC8P1 | zinc finger DHHC-type containing 8 pseudogene 1 | Other | other |
| chr11 | *CCDC89* | 1 | 91 | 3 | CCDC89 | coiled-coil domain containing 89 | Nucleus | other |
| chr3 | *RP11-3K16.1* | 1 | 57 | 3 | | | | |
| chr1 | *RP11-126K1.9* | 1 | 47 | 3 | | | | |
| chr18 | *GALR1* | 1 | 36 | 3 | GALR1 | galanin receptor 1 | Plasma Membrane | G-protein coupled recepto |
| chr12 | *SNORA40* | 1 | 28 | 3 | SNORA40 | small nucleolar RNA, H/ACA box | Other | other |
| chr15 | *RP11-253M7.1* | 1 | 32 | 3 | | | | |
| chr14 | *IGHV3-33* | 1 | 28 | 3 | IGHV3-33 | immunoglobulin heavy variable 3- | Extracellular Space | other |
| chrX | *RP11-169L17.5* | 1 | 55 | 3 | | | | |
| chr1 | *UFC1* | 1 | 51 | 3 | UFC1 | ubiquitin-fold modifier conjugating enzyme 1 | Cytoplasm | enzyme |
| chr19 | *OR7G1* | 1 | 24 | 3 | OR7G1 | olfactory receptor family 7 subfamily G member 1 | Plasma Membrane | G-protein coupled recepto |
| chr1 | *HNRNPFP1* | 1 | 49 | 3 | HNRNPFP1 | heterogeneous nuclear ribonucleoprotein F pseudogene | Other | other |
| chr1 | *APH1A* | 1 | 52 | 3 | APH1A | aph-1 homolog A, gamma-secretase subunit | Cytoplasm | peptidase |
| chr20 | *RP4-705O1.1* | 1 | 34 | 3 | | | | |
| chr1 | *RP1-224A6.3* | 1 | 37 | 3 | | | | |
| chr19 | *AC004257.1* | 1 | 40 | 3 | | | | |
| chr8 | *EBAG9* | 1 | 19 | 3 | EBAG9 | estrogen receptor binding site associated antigen 9 | Cytoplasm | other |
| chr1 | *C1orf145* | 2 | 74 | 3 | OBSCN-AS1 | OBSCN antisense RNA 1 | Other | other |
| chr15 | *RP11-632K20.8* | 1 | 35 | 3 | | | | |
| chr7 | *AC005326.2* | 1 | 40 | 3 | | | | |
| chr9 | *TMEM215* | 1 | 32 | 3 | TMEM215 | transmembrane protein 215 | Other | other |
| chr10 | *RP11-563J2.2* | 1 | 22 | 3 | | | | |
| chr1 | *OR2B11* | 1 | 56 | 3 | OR2B11 | olfactory receptor family 2 subfamily B member 11 | Plasma Membrane | G-protein coupled recepto |
| chr7 | *PRSS3P3* | 1 | 26 | 3 | PRSS3P3 | PRSS3 pseudogene 3 | Other | other |
| chr19 | *SAMD1* | 1 | 52 | 3 | SAMD1 | sterile alpha motif domain containing 1 | Other | other |
| chr13 | *TEX26* | 1 | 35 | 3 | TEX26 | testis expressed 26 | Other | other |
| chr7 | *TMEM168* | 2 | 54 | 3 | TMEM168 | transmembrane protein 168 | Cytoplasm | other |
| chr7 | *WBP1LP1* | 1 | 38 | 3 | WBP1LP1 | | Other | other |
| chrX | *PDHA1* | 1 | 29 | 3 | PDHA1 | pyruvate dehydrogenase E1 alpha 1 subunit | Cytoplasm | enzyme |
| chr11 | *OR52W1* | 1 | 72 | 3 | OR52W1 | olfactory receptor family 52 subfamily W member 1 | Plasma Membrane | G-protein coupled recepto |
| chr14 | *RP11-1127D7.1* | 1 | 26 | 3 | | | | |
| chr2 | *NAB1* | 1 | 25 | 3 | NAB1 | NGFI-A binding protein 1 | Nucleus | transcription regulator |
| chr1 | *RP11-16L9.1* | 1 | 51 | 3 | | | | |
| chr16 | *TUBB8P7* | 1 | 85 | 3 | TUBB8P7 | tubulin beta 8 class VIII pseudogene 7 | Other | other |

| | | | | | | | | | |
|---|---|---|---|---|---|---|---|---|---|
| chr7 | *SLC25A5P5* | 1 | 43 | 3 | SLC25A5P5 | solute carrier family 25 member 5 pseudogene 5 | Other | other | |
| chr1 | *GEMIN8P4* | 1 | 53 | 3 | GEMIN8P4 | gem nuclear organelle associated protein 8 pseudogene 4 | Other | other | |
| chr2 | *AC068134.5* | 1 | 46 | 3 | | | | | |
| chr16 | *MT1M* | 1 | 46 | 3 | MT1M | metallothionein 1M | Other | other | |
| chr9 | *FAM205B* | 1 | 42 | 3 | FAM205BP | transmembrane protein C9orf144B pseudogene | Other | other | |
| chr12 | *CLIC1P1* | 1 | 27 | 3 | CLIC1P1 | chloride intracellular channel 1 pseudogene 1 | Other | other | |
| chr5 | *NSA2* | 1 | 27 | 2 | NSA2 | NSA2 ribosome biogenesis factor | Nucleus | other | |
| chr3 | *RP11-221E20.5* | 1 | 59 | 2 | | | | | |
| chr2 | *RP11-761B3.1* | 1 | 32 | 2 | | | | | |
| chr16 | *MIR3670-2* | 1 | 52 | 2 | mir-3670 | microRNA 3670-1 | Other | microRNA | |
| chr15 | *RP11-1360M22.2* | 1 | 103 | 2 | | | | | |
| chr20 | *RP5-968J1.1* | 1 | 30 | 2 | | | | | |
| chr19 | *ZNF582-AS1* | 1 | 47 | 2 | ZNF582-AS1 | ZNF582 antisense RNA 1 (head to head) | Other | other | |
| chrX | *RP11-552J9.15* | 1 | 38 | 2 | | | | | |
| chr12 | *RP11-1060J15.4* | 1 | 32 | 2 | | | | | |
| chr19 | *ZNF221* | 1 | 33 | 2 | ZNF221 | zinc finger protein 221 | Nucleus | transcription regulator | |
| chr3 | *RP11-259K5.1* | 1 | 46 | 2 | | | | | |
| chr5 | *CTC-498J12.1* | 1 | 29 | 2 | | | | | |
| chr5 | *AC106786.1* | 1 | 40 | 2 | | | | | |
| chr7 | *SBDS* | 1 | 21 | 2 | SBDS | SBDS ribosome maturation factor | Nucleus | other | |
| chr2 | *TFPI* | 1 | 34 | 2 | TFPI | tissue factor pathway inhibitor | Extracellular Space | other | dalteparin, BAY1093884 |
| chr1 | *RP11-46F15.2* | 1 | 46 | 2 | | | | | |
| chr11 | *RP11-1167A19.6* | 1 | 79 | 2 | | | | | |
| chr19 | *CTB-175P5.4* | 1 | 35 | 2 | | | | | |
| chr19 | *CDKN2D* | 1 | 38 | 2 | CDKN2D | cyclin dependent kinase inhibitor | Cytoplasm | transcription regulator | |
| chr10 | *LINC00710* | 1 | 23 | 2 | LINC00710 | long intergenic non-protein coding RNA 710 | Other | other | |
| chr12 | *RP11-81H14.4* | 1 | 42 | 2 | | | | | |
| chr7 | *TRBC2* | 1 | 69 | 2 | TRBC2 | T cell receptor beta constant 2 | Other | other | |
| chr6 | *GPR115* | 1 | 25 | 2 | ADGRF4 | adhesion G protein-coupled receptor F4 | Plasma Membrane | i-protein coupled recepto | |
| chr8 | *FAM90A11P* | 1 | 91 | 2 | FAM90A11P | family with sequence similarity 90, member A11 | Other | other | |
| chr9 | *RP11-399F4.2* | 1 | 97 | 2 | | | | | |
| chr1 | *CD52* | 1 | 44 | 2 | CD52 | CD52 molecule | Plasma Membrane | other | alemtuzumab/rituximab, alemtuzumab/cyclophosphamide/fludarabine phosphate/rituximab, alemtuzumab, alemtuzumab/fludarabine phosphate, alemtuzumab/bendamustine, alemtuzumab/cyclosporin A, alemtuzumab/cyclophosphamide/fludarabine phosphate, alemtuzumab/cyclophosphamide/fludarabine phosphate/mitoxantr |
| chr16 | *RP11-514D23.1* | 1 | 36 | 2 | | | | | |
| chr12 | *EEF1A1P33* | 1 | 28 | 2 | EEF1A1P33 | eukaryotic translation elongation factor 1 alpha 1 pseudogene 33 | Other | other | |
| chr20 | *RP4-753D10.3* | 1 | 31 | 2 | | | | | |
| chr16 | *snoU109* | 1 | 29 | 2 | | | | | |
| chr16 | *RP11-17M15.1* | 1 | 30 | 2 | | | | | |
| chr3 | *RP11-222K16.2* | 1 | 42 | 2 | | | | | |
| chr8 | *RP11-520P18.5* | 1 | 69 | 2 | | | | | |
| chr12 | *ALX1* | 1 | 38 | 2 | ALX1 | ALX homeobox 1 | Nucleus | transcription regulator | |
| chr15 | *CTD-3076O17.1* | 1 | 71 | 2 | | | | | |
| chr16 | *LA16c-325D7.1* | 1 | 58 | 2 | | | | | |
| chr10 | *SMNDC1* | 1 | 65 | 2 | SMNDC1 | survival motor neuron domain containing 1 | Nucleus | other | |
| chrX | *GAGE13* | 1 | 49 | 2 | iE2D (includes oth | G antigen 2D | Other | other | |
| chrX | *RP11-469E19.1* | 1 | 51 | 2 | | | | | |
| chr1 | *MRPL55* | 2 | 115 | 2 | MRPL55 | mitochondrial ribosomal protein | Cytoplasm | other | |
| chr16 | *HIRIP3* | 1 | 55 | 2 | HIRIP3 | HIRA interacting protein 3 | Nucleus | other | |

| | | | | | | | | | |
|---|---|---|---|---|---|---|---|---|---|
| chr4 | *GABRG1* | 1 | 21 | 2 | GABRG1 | gamma-aminobutyric acid type A receptor gamma1 subunit | Plasma Membrane | ion channel | triazolam, nitrazepam, chlordiazepoxide, sevoflurane, etomidate, pagoclone, zolpidem, phenobarbital, diazepam, isoflurane, halothane, pentobarbital, desflurane, adinazolam, SEP 174559, pregnenolone, zaleplon, tracazolate, enflurane, ezogabine, muscimol, flurazepam, metyrapone/oxazepam, isoniazid, midazolam, temazepam, [illegible] |
| chr10 | *RP11-442O18.2* | 1 | 35 | 2 | | | | | |
| chr5 | *AC008914.1* | 1 | 34 | 2 | | | | | |
| chr1 | *RP5-845O24.3* | 1 | 29 | 2 | | | | | |
| chr17 | *RP11-434D2.8* | 1 | 33 | 2 | | | | | |
| chrX | *SSXP3* | 1 | 26 | 2 | SSXP3 | SSX family pseudogene 3 | Other | other | |
| chrX | *NDP* | 1 | 36 | 2 | NDP | norrin cystine knot growth factor | Extracellular Space | growth factor | |
| chr19 | *HNRNPMP2* | 1 | 32 | 2 | HNRNPMP2 | heterogeneous nuclear ribonucleoprotein M pseudogene | Other | other | |
| chr1 | *PHBP3* | 1 | 53 | 2 | PHBP3 | prohibitin pseudogene 3 | Other | other | |
| chr1 | *RP11-117D22.1* | 1 | 45 | 2 | | | | | |
| chr1 | *SPCS2P4* | 1 | 25 | 2 | SPCS2P4 | signal peptidase complex subunit 2 pseudogene 4 | Other | other | |
| chr10 | *IDI2* | 1 | 28 | 2 | IDI2 | isopentenyl-diphosphate delta isomerase 2 | Cytoplasm | enzyme | |
| chr19 | *ARHGEF18* | 1 | 56 | 2 | ARHGEF18 | Rho/Rac guanine nucleotide exchange factor 18 | Cytoplasm | other | |
| chr10 | *GLUD1P8* | 1 | 65 | 2 | GLUD1P6 | glutamate dehydrogenase 1 pseudogene 6 | Other | other | |
| chr5 | *RNF14* | 1 | 31 | 2 | RNF14 | ring finger protein 14 | Cytoplasm | transcription regulator | |
| chr10 | *RP11-50E11.3* | 2 | 63 | 2 | | | | | |
| chr1 | *RP3-340N1.2* | 1 | 43 | 2 | | | | | |
| chr20 | *AL158091.1* | 1 | 88 | 2 | | | | | |
| chr11 | *RP11-313I2.7* | 1 | 58 | 2 | | | | | |
| chr15 | *DDX11L9* | 1 | 152 | 2 | DDX11L1/DDX11L | DEAD/H-box helicase 11 like 1 | Other | other | |
| chr7 | *CCT6A* | 1 | 34 | 2 | CCT6A | chaperonin containing TCP1 subunit 6A | Cytoplasm | other | |
| chr22 | *CTA-941F9.9* | 1 | 24 | 2 | | | | | |
| chrX | *CNGA2* | 1 | 24 | 2 | CNGA2 | cyclic nucleotide gated channel subunit alpha 2 | Plasma Membrane | ion channel | |
| chr16 | *RP11-510M2.4* | 1 | 43 | 2 | | | | | |
| chr12 | *RP11-497G19.3* | 1 | 33 | 2 | | | | | |
| chr16 | *RP11-67A1.2* | 1 | 42 | 2 | | | | | |
| chr11 | *RP11-1081L13.4* | 1 | 42 | 2 | | | | | |
| chr1 | *CICP26* | 1 | 75 | 2 | P14 (includes oth | capicua transcriptional repressor pseudogene 26 | Other | other | |
| chr14 | *RP11-111A21.1* | 1 | 26 | 2 | | | | | |
| chr15 | *MIR4508* | 1 | 31 | 2 | MIR4508 | microRNA 4508 | Cytoplasm | microRNA | |
| chr14 | *RP11-903H12.3* | 1 | 18 | 2 | | | | | |
| chr15 | *RP11-244F12.1* | 1 | 33 | 2 | | | | | |
| chr16 | *URAHP* | 1 | 52 | 2 | URAHP | urate (hydroxyiso-) hydrolase, pseudogene | Other | other | |
| chr11 | *KRTAP5-14P* | 1 | 48 | 2 | KRTAP5-14P | keratin associated protein 5-14, pseudogene | Other | other | |
| chr13 | *BIVM-ERCC5* | 1 | 23 | 2 | BIVM-ERCC5 | BIVM-ERCC5 readthrough | Other | other | |
| chr15 | *RP11-160C18.4* | 1 | 65 | 2 | | | | | |
| chr19 | *CTB-60E11.11* | 1 | 36 | 2 | | | | | |
| chr4 | *PPEF2* | 1 | 21 | 2 | PPEF2 | protein phosphatase with EF-hand domain 2 | Cytoplasm | phosphatase | |
| chr13 | *LINC00332* | 1 | 29 | 2 | LINC00332 | long intergenic non-protein coding RNA 332 | Other | other | |
| chr11 | *MMP12* | 1 | 26 | 2 | MMP12 | matrix metallopeptidase 12 | Extracellular Space | peptidase | marimastat |
| chr16 | *RP11-586K12.4* | 1 | 27 | 2 | | | | | |
| chr18 | *MYL12B* | 1 | 47 | 2 | MYL12B | myosin light chain 12B | Cytoplasm | other | |
| chrX | *UXT* | 1 | 28 | 2 | UXT | ubiquitously expressed prefoldin like chaperone | Cytoplasm | transcription regulator | |
| chr7 | *OR2A5* | 1 | 36 | 2 | OR2A5 | olfactory receptor family 2 subfamily A member 5 | Plasma Membrane | -protein coupled recept | |
| chr1 | *RP1-125I3.2* | 1 | 37 | 2 | | | | | |
| chr15 | *GRAMD4P5* | 1 | 55 | 2 | GRAMD4P5 | GRAM domain containing 4 pseudogene 5 | Other | other | |
| chr22 | *IGLC2* | 1 | 45 | 2 | IGLC2 | immunoglobulin lambda constant | Extracellular Space | other | |
| chr2 | *AC104395.1* | 1 | 36 | 2 | | | | | |
| chr10 | *FRG2B* | 1 | 50 | 2 | FRG2/FRG2B | FSHD region gene 2 | Other | other | |
| chr17 | *C17orf59* | 2 | 56 | 2 | BORCS6 | BLOC-1 related complex subunit | Cytoplasm | other | |
| chr1 | *RP11-344H11.4* | 1 | 32 | 2 | | | | | |

| | | | | | | | | | |
|---|---|---|---|---|---|---|---|---|---|
| chr20 | *RP13-379L11.1* | 1 | 47 | 2 | | | | | |
| chr1 | *HMGN1P5* | 1 | 38 | 2 | HMGN1P5 | high mobility group nucleosome binding domain 1 pseudogene 5 | Other | other | |
| chr7 | *RP11-775D22.2* | 1 | 49 | 2 | | | | | |
| chr21 | *MYL6P1* | 1 | 32 | 2 | MYL6P1 | myosin light chain 6 pseudogene | Other | other | |
| chr1 | *RN7SL729P* | 1 | 40 | 2 | | | | | |
| chr6 | *PLA2G7* | 1 | 25 | 2 | PLA2G7 | phospholipase A2 group VII | Extracellular Space | enzyme | darapladib |
| chr20 | *HAUS6P2* | 1 | 36 | 2 | HAUS6P2 | HAUS augmin like complex subunit 6 pseudogene 2 | Other | other | |
| chr2 | *CH17-132F21.1* | 1 | 110 | 2 | | | | | |
| chr5 | *RN7SL103P* | 1 | 27 | 2 | | | | | |
| chr4 | *ZSWIM5P3* | 1 | 26 | 2 | ZSWIM5P3 | zinc finger SWIM-type containing 5 pseudogene 3 | Other | other | |
| chr6 | *HIST1H4A* | 1 | 36 | 2 | H4C1 | H4 clustered histone 1 | Nucleus | other | |
| chr19 | *CTC-512J12.4* | 1 | 50 | 2 | | | | | |
| chr1 | *RP4-758J18.13* | 1 | 45 | 2 | | | | | |
| chr1 | *CFL1P2* | 1 | 34 | 2 | CFL1P2 | cofilin 1 pseudogene 2 | Other | other | |
| chr1 | *VHLL* | 1 | 49 | 2 | VHLL | VHL like | Other | other | |
| chr18 | *RP11-154H12.2* | 2 | 74 | 2 | | | | | |
| chr2 | *AC018892.9* | 1 | 69 | 2 | | | | | |
| chr10 | *RP11-523O18.5* | 1 | 57 | 2 | | | | | |
| chr12 | *RPL13AP20* | 1 | 51 | 2 | RPL13AP20 | ribosomal protein L13a pseudogene 20 | Other | other | |
| chr15 | *KRT8P23* | 1 | 73 | 2 | KRT8P23 | keratin 8 pseudogene 23 | Other | other | |
| chr2 | *AC016912.3* | 1 | 60 | 2 | | | | | |
| chr4 | *ZNF141* | 1 | 22 | 2 | ZNF141 | zinc finger protein 141 | Nucleus | transcription regulator | |
| chr1 | *AL590483.1* | 1 | 42 | 2 | | | | | |
| chr11 | *RP11-266A24.1* | 1 | 30 | 2 | | | | | |
| chr7 | *RP11-368M16.7* | 1 | 108 | 2 | | | | | |
| chr1 | *ANKRD65* | 1 | 46 | 2 | ANKRD65 | ankyrin repeat domain 65 | Other | other | |
| chrX | *PAGE2* | 1 | 53 | 2 | PAGE2/PAGE2B | PAGE family member 2 | Other | other | |
| chr1 | *BLZF1* | 1 | 38 | 2 | BLZF1 | basic leucine zipper nuclear | Cytoplasm | transcription regulator | |
| chr7 | *RP11-377H23.1* | 1 | 56 | 2 | | | | | |
| chr3 | *TADA3* | 1 | 48 | 2 | TADA3 | transcriptional adaptor 3 | Nucleus | transcription regulator | |
| chr11 | *RP11-713P14.1* | 1 | 36 | 2 | | | | | |
| chr19 | *MIR519A2* | 1 | 31 | 2 | mir-515 | microRNA 520c | Cytoplasm | microRNA | |
| chr7 | *FABP5P3* | 1 | 40 | 2 | FABP5P3 | fatty acid binding protein 5 pseudogene 3 | Other | other | |
| chr2 | *NPPC* | 1 | 36 | 2 | NPPC | natriuretic peptide C | Extracellular Space | other | |
| chr17 | *AC099850.1* | 1 | 25 | 2 | | | | | |
| chr10 | *RP11-150D20.5* | 1 | 77 | 2 | | | | | |
| chrX | *ZC3H12B* | 1 | 31 | 2 | ZC3H12B | zinc finger CCCH-type containing 12B | Other | other | |
| chr2 | *PIKFYVE* | 1 | 22 | 2 | PIKFYVE | phosphoinositide kinase, FYVE-type zinc finger containing | Cytoplasm | kinase | apilimod |
| chr1 | *RP11-389O22.1* | 1 | 36 | 2 | | | | | |
| chr6 | *AL590867.1* | 1 | 25 | 2 | | | | | |
| chrX | *HEPH* | 1 | 30 | 2 | HEPH | hephaestin | Plasma Membrane | transporter | |
| chr17 | *CDRT15P2* | 1 | 49 | 2 | CDRT15P2 | CMT1A duplicated region transcript 15 pseudogene 2 | Other | other | |
| chr14 | *RP11-16B13.1* | 1 | 18 | 2 | | | | | |
| chr17 | *RP11-640N20.9* | 1 | 36 | 2 | | | | | |
| chr16 | *RRAD* | 1 | 56 | 2 | RRAD | RRAD, Ras related glycolysis inhibitor and calcium channel regulator | Cytoplasm | enzyme | |
| chr20 | *WFDC2* | 1 | 45 | 2 | WFDC2 | WAP four-disulfide core domain 2 | Extracellular Space | other | |
| chr4 | *RP11-149A7.2* | 1 | 31 | 2 | | | | | |
| chr13 | *ARL11* | 1 | 35 | 2 | ARL11 | ADP ribosylation factor like GTPase 11 | Other | other | |
| chr7 | *RN7SL478P* | 1 | 50 | 2 | RN7SL478P | RNA, 7SL, cytoplasmic 478, pseudogene | Other | other | |
| chr22 | *RP3-508I15.9* | 1 | 27 | 2 | | | | | |
| chr19 | *CTC-523E23.8* | 1 | 46 | 2 | | | | | |
| chr5 | *GPX3* | 1 | 29 | 2 | GPX3 | glutathione peroxidase 3 | Extracellular Space | enzyme | |
| chr16 | *IL21R-AS1* | 1 | 35 | 2 | IL21R-AS1 | IL21R antisense RNA 1 | Other | other | |
| chr17 | *RP11-214C8.2* | 1 | 39 | 2 | | | | | |
| chr1 | *RP11-180D21.3* | 1 | 34 | 2 | | | | | |
| chr19 | *USP29* | 2 | 70 | 2 | USP29 | ubiquitin specific peptidase 29 | Other | other | |
| chr19 | *AC005777.3* | 1 | 44 | 2 | | | | | |
| chr4 | *PKD2* | 1 | 20 | 2 | PKD2 | polycystin 2, transient receptor potential cation channel | Plasma Membrane | ion channel | |
| chr1 | *MIR4417* | 1 | 38 | 2 | | | | | |
| chrX | *SPANXB2* | 1 | 81 | 2 | NXB1 (includes ot | SPANX family member C | Nucleus | other | |
| chr16 | *RP11-542M13.3* | 1 | 32 | 2 | | | | | |
| chr7 | *RP11-190G13.4* | 1 | 29 | 2 | | | | | |
| chr3 | *GS1-388B5.8* | 1 | 50 | 2 | | | | | |
| chr1 | *RP11-40H20.5* | 1 | 49 | 2 | | | | | |
| chr20 | *SYS1* | 1 | 43 | 2 | SYS1 | SYS1 golgi trafficking protein | Cytoplasm | other | |
| chr11 | *GPHA2* | 1 | 54 | 2 | GPHA2 | glycoprotein hormone subunit | Extracellular Space | other | |
| chr8 | *MYBL1* | 1 | 19 | 2 | MYBL1 | MYB proto-oncogene like 1 | Nucleus | transcription regulator | |
| chr20 | *C20orf195* | 2 | 152 | 2 | FNDC11 | fibronectin type III domain containing 11 | Cytoplasm | other | |
| chr11 | *RP11-864N7.2* | 1 | 53 | 2 | | | | | |
| chr6 | *RP3-483K16.4* | 1 | 26 | 2 | | | | | |
| chr5 | *OR2Y1* | 1 | 41 | 2 | OR2Y1 | olfactory receptor family 2 subfamily Y member 1 | Plasma Membrane | 3-protein coupled recepto | |
| chr1 | *IL10* | 1 | 33 | 2 | IL10 | interleukin 10 | Extracellular Space | cytokine | |
| chr12 | *RP11-946L16.1* | 1 | 34 | 2 | | | | | |
| chr6 | *RP11-55K22.5* | 1 | 36 | 2 | | | | | |

| | | | | | | | | | |
|---|---|---|---|---|---|---|---|---|---|
| chr6 | HIST1H1D | 1 | 41 | 2 | H1-3 | H1.3 linker histone, cluster | Nucleus | other | |
| chr12 | RP4-761J14.8 | 1 | 28 | 2 | | | | | |
| chr2 | ALPP | 1 | 40 | 2 | ALPP | alkaline phosphatase, placental | Plasma Membrane | phosphatase | zinc sulfate |
| chr9 | RP11-145E17.2 | 1 | 75 | 2 | | | | | |
| chr12 | CTD-2140B24.6 | 1 | 29 | 2 | | | | | |
| chr16 | RP11-510M2.8 | 1 | 42 | 2 | | | | | |
| chr3 | RP4-555D20.2 | 1 | 35 | 2 | | | | | |
| chr5 | CTC-487M23.8 | 1 | 30 | 2 | | | | | |
| chr1 | RN7SL131P | 1 | 33 | 2 | RN7SL131P | RNA, 7SL, cytoplasmic 131, pseudogene | Other | other | |
| chr16 | MT1L | 1 | 33 | 2 | MT1L | metallothionein 1L, pseudogene | Cytoplasm | other | |
| chr1 | FAM138A | 1 | 61 | 2 | FAM138A | family with sequence similarity 138 member A | Other | other | |
| chr11 | RP11-304C12.3 | 1 | 55 | 2 | | | | | |
| chr17 | PDLIM1P3 | 1 | 36 | 2 | PDLIM1P3 | PDZ and LIM domain 1 pseudogene 3 | Other | other | |
| chr16 | RP11-46C24.7 | 1 | 33 | 2 | | | | | |
| chr20 | TGIF2-C20orf24 | 1 | 56 | 2 | TGIF2-RAB5IF | TGIF2-RAB5IF readthrough | Other | other | |
| chr6 | XX-C2158C12.2 | 1 | 51 | 2 | | | | | |
| chr9 | RP11-404E6.1 | 1 | 48 | 2 | | | | | |
| chr17 | FTLP12 | 1 | 41 | 2 | FTLP12 | ferritin light chain pseudogene 12 | Other | other | |
| chr17 | MFSD6L | 1 | 49 | 2 | MFSD6L | major facilitator superfamily domain containing 6 like | Other | other | |
| chrX | HDX | 1 | 21 | 2 | HDX | highly divergent homeobox | Other | other | |
| chr12 | NANOG | 1 | 22 | 2 | NANOG | Nanog homeobox | Nucleus | transcription regulator | |
| chrX | SMEK3P | 1 | 34 | 2 | PPP4R3C | protein phosphatase 4 regulatory subunit 3C | Other | other | |
| chr7 | RP11-89N17.1 | 1 | 55 | 2 | | | | | |
| chr4 | GUSBP5 | 1 | 33 | 2 | GUSBP5 | GUSB pseudogene 5 | Other | other | |
| chr5 | ATG12 | 1 | 28 | 2 | ATG12 | autophagy related 12 | Cytoplasm | other | |
| chr17 | FTLP13 | 1 | 37 | 2 | FTLP13 | ferritin light chain pseudogene 13 | Other | other | |
| chr11 | SLC25A45 | 1 | 86 | 2 | SLC25A45 | solute carrier family 25 member | Cytoplasm | transporter | |
| chr17 | RP11-849F2.8 | 1 | 42 | 2 | | | | | |
| chr4 | FAM13A-AS1 | 1 | 18 | 2 | FAM13A-AS1 | FAM13A antisense RNA 1 | Other | other | |
| chr6 | RP11-184C23.4 | 1 | 47 | 2 | | | | | |
| chr8 | RP11-398H6.1 | 1 | 40 | 2 | | | | | |
| chr4 | RP11-696N14.3 | 1 | 24 | 2 | | | | | |
| chr2 | FTOP1 | 1 | 40 | 2 | FTOP1 | FTO pseudogene 1 | Other | other | |
| chr7 | FBXO24 | 1 | 61 | 2 | FBXO24 | F-box protein 24 | Other | enzyme | |
| chr8 | RP11-359E19.2 | 1 | 27 | 2 | | | | | |
| chr17 | RNA5SP448 | 1 | 34 | 2 | RNA5SP448 | RNA, 5S ribosomal pseudogene | Other | other | |
| chr17 | AC091199.1 | 1 | 26 | 2 | | | | | |
| chr1 | SDCCAG3P2 | 1 | 57 | 2 | ENTR1P2 | ENTR1 pseudogene 2 | Other | other | |
| chr6 | POM121L14P | 1 | 48 | 2 | POM121L14P | POM121 transmembrane nucleoporin like 14, pseudogene | Other | other | |
| chrX | BX842568.2 | 1 | 35 | 2 | | | | | |
| chr11 | GIF | 1 | 20 | 2 | CBLIF | cobalamin binding intrinsic factor | Extracellular Space | transporter | |
| chr9 | PHBP7 | 1 | 53 | 2 | PHBP7 | prohibitin pseudogene 7 | Other | other | |
| chr11 | BANF1 | 1 | 72 | 2 | BANF1 | barrier to autointegration factor 1 | Nucleus | other | |
| chr8 | TRIB1 | 1 | 32 | 2 | TRIB1 | tribbles pseudokinase 1 | Cytoplasm | kinase | |
| chr1 | U1 | 1 | 182 | 2 | | | | | |
| chr4 | SCARB2 | 1 | 19 | 2 | SCARB2 | scavenger receptor class B member 2 | Plasma Membrane | transmembrane receptor | |
| chr13 | LINC00390 | 1 | 22 | 2 | LINC00390 | | Other | other | |
| chr3 | RP11-202A13.1 | 1 | 65 | 2 | | | | | |
| chr16 | FBXL8 | 1 | 63 | 2 | FBXL8 | F-box and leucine rich repeat protein 8 | Cytoplasm | other | |
| chr17 | RP11-1113L8.6 | 1 | 25 | 2 | | | | | |
| chr17 | RP11-358B23.1 | 1 | 48 | 2 | | | | | |
| chr15 | RP11-315D16.4 | 1 | 43 | 2 | | | | | |
| chr10 | AL603965.1 | 1 | 89 | 2 | | | | | |
| chr4 | RP11-539L10.3 | 1 | 35 | 2 | | | | | |
| chr19 | CTB-60B18.6 | 1 | 33 | 2 | | | | | |
| chr17 | RNF167 | 1 | 46 | 2 | RNF167 | ring finger protein 167 | Cytoplasm | enzyme | |
| chr6 | TDP2 | 1 | 45 | 2 | TDP2 | tyrosyl-DNA phosphodiesterase 2 | Cytoplasm | transcription regulator | |
| chr15 | RP11-138H8.7 | 1 | 26 | 2 | | | | | |
| chr4 | TNIP3 | 1 | 19 | 2 | TNIP3 | TNFAIP3 interacting protein 3 | Cytoplasm | other | |
| chr3 | MBD4 | 1 | 35 | 2 | MBD4 | methyl-CpG binding domain 4, DNA glycosylase | Nucleus | enzyme | |
| chr8 | AP006547.3 | 1 | 63 | 2 | | | | | |
| chr4 | TRIML1 | 1 | 23 | 2 | TRIML1 | tripartite motif family like 1 | Other | other | |
| chr17 | RP11-557B23.3 | 1 | 41 | 2 | | | | | |
| chr5 | RP11-281O15.5 | 1 | 53 | 2 | | | | | |
| chrX | ATG4A | 1 | 19 | 2 | ATG4A | autophagy related 4A cysteine peptidase | Cytoplasm | peptidase | |
| chr17 | KRTAP2-5P | 1 | 42 | 2 | KRTAP2-5P | keratin associated protein 2-5, pseudogene | Other | other | |
| chr13 | RP11-199F6.4 | 1 | 45 | 2 | | | | | |
| chr1 | GAS5 | 1 | 42 | 2 | GAS5 | growth arrest specific 5 | Other | other | |
| chr1 | FLVCR1-AS1 | 1 | 35 | 2 | FLVCR1-DT | FLVCR1 divergent transcript | Other | other | |
| chr1 | RP11-380B22.1 | 1 | 36 | 2 | | | | | |
| chr1 | RP11-543B16.2 | 1 | 58 | 2 | | | | | |
| chr1 | CNN2P10 | 1 | 43 | 2 | CNN2P10 | calponin 2 pseudogene 10 | Other | other | |
| chr5 | CTB-27N1.1 | 1 | 53 | 2 | | | | | |
| chr7 | AC073094.4 | 2 | 112 | 2 | | | | | |
| chr7 | CTAGE15 | 1 | 56 | 2 | GE9 (includes oth | CTAGE family member 9 | Other | other | |
| chr4 | RP11-73G16.2 | 1 | 25 | 2 | | | | | |
| chr5 | RP11-432M8.5 | 1 | 1657 | 2 | | | | | |
| chr9 | RP11-307I14.4 | 1 | 33 | 2 | | | | | |

| | | | | | | | | |
|---|---|---|---|---|---|---|---|---|
| chr2 | *ST6GALNAC2P1* | 1 | 43 | 2 | ST6GALNAC2P1 | ST6 N-acetylgalactosaminide alpha-2,6-sialyltransferase 2 | Other | other |
| chr11 | *OR10Q2P* | 1 | 40 | 2 | OR10Q2P | olfactory receptor family 10 subfamily Q member 2 | Other | other |
| chr5 | *CCNG1* | 1 | 31 | 2 | CCNG1 | cyclin G1 | Nucleus | other |
| chr5 | *CTD-2201E18.5* | 1 | 91 | 2 | | | | |
| chr9 | *AK1* | 1 | 61 | 2 | AK1 | adenylate kinase 1 | Cytoplasm | kinase |
| chr6 | *ACTG1P9* | 1 | 43 | 2 | ACTG1P9 | actin gamma 1 pseudogene 9 | Other | other |
| chr20 | *RPS2P1* | 1 | 78 | 2 | RPS2P1 | ribosomal protein S2 pseudogene | Other | other |
| chr10 | *BLOC1S2* | 2 | 44 | 2 | BLOC1S2 | biogenesis of lysosomal organelles complex 1 subunit 2 | Cytoplasm | other |
| chr8 | *ADAM3A* | 1 | 31 | 2 | ADAM3A | ADAM metallopeptidase domain 3A (pseudogene) | Other | other |
| chr17 | *RP11-51F16.8* | 1 | 29 | 2 | | | | |
| chr5 | *GS1-39E22.1* | 1 | 31 | 2 | | | | |
| chr6 | *PM20D2* | 1 | 22 | 2 | PM20D2 | peptidase M20 domain containing | Nucleus | peptidase |
| chr7 | *TPM3P4* | 1 | 42 | 2 | TPM3P4 | tropomyosin 3 pseudogene 4 | Other | other |
| chr1 | *RP4-635E18.6* | 1 | 38 | 2 | | | | |
| chr22 | *IGLV3-9* | 1 | 24 | 2 | IGLV3-9 | immunoglobulin lambda variable 3-9 (gene/pseudogene) | Other | other |
| chr1 | *RP11-443B7.3* | 1 | 43 | 2 | | | | |
| chr12 | *DBX2* | 1 | 31 | 2 | DBX2 | developing brain homeobox 2 | Nucleus | transcription regulator |
| chr2 | *AC012360.1* | 1 | 40 | 2 | | | | |
| chr9 | *RP11-276H19.4* | 1 | 51 | 2 | | | | |
| chr11 | *OR56A3* | 1 | 40 | 2 | OR56A3 | olfactory receptor family 56 subfamily A member 3 | Plasma Membrane | G-protein coupled recepto |
| chr16 | *CTD-2144E22.10* | 1 | 28 | 2 | | | | |
| chr10 | *COMTD1* | 1 | 72 | 2 | COMTD1 | catechol-O-methyltransferase domain containing 1 | Plasma Membrane | enzyme |
| chr4 | *SOWAHB* | 1 | 38 | 2 | SOWAHB | sosondowah ankyrin repeat domain family member B | Nucleus | transcription regulator |
| chr16 | *RPL34P29* | 1 | 40 | 2 | RPL34P29 | ribosomal protein L34 pseudogene 29 | Other | other |
| chr12 | *LINC00943* | 1 | 28 | 2 | LINC00943 | long intergenic non-protein coding RNA 943 | Other | other |
| chr15 | *GABPB1-AS1* | 1 | 24 | 2 | GABPB1-AS1 | GABPB1 antisense RNA 1 | Other | other |
| chr20 | *LINC00657* | 1 | 35 | 2 | NORAD | non-coding RNA activated by DNA damage | Other | other |
| chr9 | *RPS10P3* | 1 | 32 | 2 | RPS10P3 | ribosomal protein S10 | Other | other |
| chr20 | *RP4-568F9.6* | 1 | 28 | 2 | | | | |
| chr19 | *LINC00904* | 1 | 23 | 2 | LINC00904 | long intergenic non-protein coding RNA 904 | Other | other |
| chr6 | *RPL35P2* | 1 | 41 | 2 | RPL35P2 | ribosomal protein L35 | Other | other |
| chr5 | *RP11-54F2.1* | 1 | 130 | 2 | | | | |
| chr8 | *RP11-593P24.4* | 1 | 40 | 2 | | | | |
| chr12 | *C12orf57* | 1 | 33 | 2 | C12orf57 | chromosome 12 open reading frame 57 | Cytoplasm | other |
| chr5 | *CTC-575D19.1* | 1 | 43 | 2 | | | | |
| chr2 | *AC009505.4* | 1 | 38 | 2 | | | | |
| chr1 | *RP11-422P22.1* | 1 | 28 | 2 | | | | |
| chrX | *GAPDHP65* | 1 | 42 | 2 | GAPDHP65 | glyceraldehyde 3 phosphate dehydrogenase pseudogene 65 | Other | other |
| chrX | *XAGE3* | 1 | 65 | 2 | XAGE3 | X antigen family member 3 | Other | other |
| chr10 | *RP11-343J3.2* | 1 | 75 | 2 | | | | |
| chrX | *TBX22* | 1 | 28 | 2 | TBX22 | T-box transcription factor 22 | Nucleus | transcription regulator |
| chr20 | *LZTS3* | 1 | 39 | 2 | LZTS3 | leucine zipper tumor suppressor family member 3 | Cytoplasm | other |
| chr5 | *ZFP62* | 1 | 30 | 2 | ZFP62 | ZFP62 zinc finger protein | Nucleus | transcription regulator |
| chr16 | *TMEM208* | 1 | 45 | 2 | TMEM208 | transmembrane protein 208 | Other | other |
| chr1 | *RP11-421E17.4* | 1 | 60 | 2 | | | | |
| chr15 | *RP11-64K10.1* | 1 | 34 | 2 | | | | |
| chr4 | *RP11-141E13.1* | 1 | 29 | 2 | | | | |
| chr15 | *AHCYP7* | 1 | 41 | 2 | AHCYP7 | adenosylhomocysteinase pseudogene 7 | Other | other |
| chr14 | *IGHV3-30* | 1 | 30 | 2 | IGHV3-30 | immunoglobulin heavy variable 3- | Extracellular Space | other |
| chr8 | *RP11-274M4.1* | 1 | 45 | 2 | | | | |
| chr19 | *C19orf35* | 1 | 38 | 2 | PEAK3 | PEAK family member 3 | Other | other |
| chr15 | *LACTB* | 1 | 37 | 2 | LACTB | lactamase beta | Cytoplasm | peptidase |
| chr22 | *IGLV9-49* | 1 | 32 | 2 | IGLV9-49 | immunoglobulin lambda variable | Other | other |
| chr2 | *AC022210.2* | 1 | 54 | 2 | | | | |
| chr15 | *RP11-38G5.3* | 1 | 47 | 2 | | | | |
| chr20 | *HDHD1P3* | 1 | 56 | 2 | PUDPP3 | pseudouridine 5'-phosphatase pseudogene 3 | Other | other |
| chr15 | *AC090519.2* | 1 | 30 | 2 | | | | |
| chr14 | *GOLGA5* | 1 | 17 | 2 | GOLGA5 | golgin A5 | Cytoplasm | kinase |
| chr1 | *RP11-122G18.7* | 1 | 55 | 2 | | | | |
| chr6 | *ZKSCAN4* | 1 | 26 | 2 | ZKSCAN4 | zinc finger with KRAB and SCAN domains 4 | Nucleus | transcription regulator |
| chr19 | *CTB-129P6.11* | 1 | 29 | 2 | | | | |
| chr8 | *KB-1460A1.1* | 1 | 27 | 2 | | | | |
| chr5 | *THG1L* | 1 | 53 | 2 | THG1L | tRNA-histidine guanylyltransferase 1 like | Cytoplasm | enzyme |
| chr3 | *GS1-388B5.1* | 1 | 72 | 2 | | | | |
| chr9 | *RP11-349E4.1* | 1 | 37 | 2 | | | | |
| chr2 | *RP11-254F7.2* | 1 | 33 | 2 | | | | |
| chr7 | *CICP17* | 1 | 69 | 2 | CICP17 | capicua transcriptional repressor pseudogene 17 | Other | other |
| chr7 | *DMTF1* | 1 | 34 | 2 | DMTF1 | cyclin D binding myb like transcription factor 1 | Nucleus | transcription regulator |

| | | | | | | | | | |
|---|---|---|---|---|---|---|---|---|---|
| chr1 | *AL021920.2* | 1 | 98 | 2 | | | | | |
| chr9 | *RP11-109M17.1* | 1 | 43 | 2 | | | | | |
| chr11 | *CTD-2562J17.2* | 1 | 44 | 2 | | | | | |
| chr11 | *RP11-540A21.2* | 1 | 57 | 2 | | | | | |
| chr7 | *PHKG1P4* | 1 | 61 | 2 | PHKG1P4 | phosphorylase kinase catalytic subunit gamma 1 pseudogene 4 | Other | other | |
| chr16 | *RP11-144N1.4* | 1 | 34 | 2 | | | | | |
| chr2 | *FOXI3* | 1 | 35 | 2 | FOXI3 | forkhead box I3 | Nucleus | transcription regulator | |
| chr6 | *C6orf62* | 1 | 22 | 2 | C6orf62 | chromosome 6 open reading frame 62 | Other | other | |
| chr7 | *AC006026.10* | 1 | 66 | 2 | | | | | |
| chr9 | *TSTD2* | 1 | 25 | 2 | TSTD2 | thiosulfate sulfurtransferase like domain containing 2 | Other | other | |
| chr2 | *IGKV1-39* | 1 | 25 | 2 | IGKV1-39 | immunoglobulin kappa variable 1-39 (gene/pseudogene) | Extracellular Space | other | |
| chr1 | *RP4-791M13.3* | 1 | 78 | 2 | | | | | |
| chr17 | *AC015815.5* | 1 | 35 | 2 | | | | | |
| chr20 | *RN7SL511P* | 1 | 48 | 2 | | | | | |
| chr20 | *RN7SL603P* | 1 | 53 | 2 | | | | | |
| chr12 | *NME2P1* | 1 | 43 | 2 | NME2P1 | NME/NM23 nucleoside diphosphate kinase 2 | Cytoplasm | other | |
| chr16 | *TPSAB1* | 1 | 77 | 2 | TPSAB1/TPSB2 | tryptase alpha/beta 1 | Extracellular Space | peptidase | |
| chr9 | *RP11-205K6.2* | 1 | 59 | 2 | | | | | |
| chr7 | *DGAT2L7P* | 1 | 48 | 2 | DGAT2L7P | diacylglycerol O-acyltransferase 2 like 7, pseudogene | Cytoplasm | other | |
| chr13 | *LIG4* | 1 | 37 | 2 | LIG4 | DNA ligase 4 | Nucleus | enzyme | |
| chr1 | *ZBTB18* | 1 | 51 | 2 | ZBTB18 | zinc finger and BTB domain containing 18 | Nucleus | transcription regulator | |
| chr2 | *CYBRD1* | 1 | 19 | 2 | CYBRD1 | cytochrome b reductase 1 | Cytoplasm | enzyme | |
| chr5 | *BTF3* | 1 | 19 | 2 | BTF3 | basic transcription factor 3 | Nucleus | transcription regulator | |
| chr22 | *DDTL* | 1 | 33 | 2 | DDTL | D-dopachrome tautomerase like | Cytoplasm | other | |
| chr1 | *SPRR1A* | 1 | 41 | 2 | SPRR1A | small proline rich protein 1A | Plasma Membrane | other | |
| chrX | *GAPDHP1* | 1 | 40 | 2 | GAPDHP1 | glyceraldehyde-3-phosphate dehydrogenase pseudogene 1 | Other | other | |
| chr5 | *RP11-432M8.4* | 1 | 1117 | 2 | | | | | |
| chr1 | *RN7SKP276* | 1 | 51 | 2 | RN7SKP276 | RN7SK pseudogene 276 | Other | other | |
| chr7 | *CTB-113D17.1* | 1 | 34 | 2 | | | | | |
| chr2 | *AQP12A* | 1 | 60 | 2 | AQP12A/AQP12B | aquaporin 12A | Cytoplasm | transporter | |
| chr2 | *AC010745.1* | 1 | 24 | 2 | | | | | |
| chr17 | *NOS2P2* | 1 | 56 | 2 | NOS2P2 | nitric oxide synthase 2 | Other | other | |
| chr12 | *RP11-155I9.1* | 1 | 49 | 2 | | | | | |
| chr21 | *LRRC3-AS1* | 1 | 31 | 2 | LRRC3-DT | LRRC3 divergent transcript | Other | other | |
| chr13 | *METTL21C* | 1 | 35 | 2 | METTL21C | methyltransferase like 21C | Nucleus | enzyme | |
| chr1 | *RP11-403I13.7* | 1 | 124 | 2 | | | | | |
| chr17 | *MRPS7* | 1 | 48 | 2 | MRPS7 | mitochondrial ribosomal protein | Cytoplasm | other | |
| chr11 | *CHRM4* | 1 | 79 | 2 | CHRM4 | cholinergic receptor muscarinic 4 | Plasma Membrane | G-protein coupled recepto | dexamethasone/olanzapine, dicyclomine, cevimeline, mirabegron/solifenacin, olanzapine, aclidinium/formoterol, acetylcholine, atropine/pralidoxime, brompheniramine, umeclidinium, mepenzolic acid, propantheline, orphenadrine, aspirin/caffeine/orphenadrine, tolterodine, chlorpheniramine/methscopolamine/phenylephrine, trihexyphenidyl, oxybutynin, formoterol/glycopyrrolate, scopolamine, arecoline, [illegible] |
| chr9 | *RP11-549A6.1* | 1 | 33 | 2 | | | | | |
| chr9 | *RP11-229P13.25* | 1 | 76 | 2 | | | | | |
| chrX | *HMGA1P1* | 1 | 43 | 2 | HMGA1P1 | high mobility group AT-hook 1 pseudogene 1 | Other | other | |
| chr13 | *BORA* | 1 | 17 | 2 | BORA | BORA aurora kinase A activator | Cytoplasm | other | |
| chr7 | *URGCP-MRPS24* | 1 | 52 | 2 | URGCP-MRPS24 | URGCP-MRPS24 readthrough | Other | other | |
| chr9 | *RP11-7G23.4* | 1 | 45 | 2 | | | | | |
| chr7 | *CCNJP1* | 1 | 45 | 2 | CCNJP1 | cyclin J pseudogene 1 | Other | other | |
| chr3 | *NDUFAF3* | 1 | 57 | 2 | NDUFAF3 | NADH:ubiquinone oxidoreductase complex | Nucleus | other | |
| chr3 | *ENPP7P3* | 1 | 52 | 2 | ENPP7P3 | | Other | other | |
| chr5 | *C5orf54* | 1 | 48 | 2 | ZBED8 | zinc finger BED-type containing 8 | Nucleus | other | |
| chr1 | *RP1-92O14.3* | 1 | 33 | 2 | | | | | |
| chrX | *RP11-169L17.3* | 1 | 27 | 2 | | | | | |
| chr7 | *AC102953.6* | 1 | 63 | 2 | | | | | |
| chr19 | *RLN3* | 1 | 54 | 2 | RLN3 | relaxin 3 | Extracellular Space | other | |
| chr5 | *RP11-417J1.1* | 1 | 46 | 2 | | | | | |
| chr16 | *APRT* | 1 | 62 | 2 | APRT | adenine | Cytoplasm | enzyme | |
| chr22 | *RP5-1177I5.3* | 1 | 32 | 2 | | | | | |
| chr1 | *RN7SL668P* | 1 | 45 | 2 | RN7SL668P | | Other | other | |

| | | | | | | | | | |
|---|---|---|---|---|---|---|---|---|---|
| chr9 | *LCN1P2* | 1 | 61 | 2 | LCN1P2 | lipocalin 1 pseudogene 2 | Other | other | |
| chr10 | *FRA10AC1* | 1 | 19 | 2 | FRA10AC1 | FRA10A associated CGG repeat | Nucleus | other | |
| chrX | *AMMECR1* | 1 | 28 | 2 | AMMECR1 | AMMECR nuclear protein 1 | Nucleus | other | |
| chr6 | *RP5-894D12.5* | 1 | 33 | 2 | | | | | |
| chr5 | *RP11-432M8.2* | 1 | 1780 | 2 | | | | | |
| chr4 | *CCKAR* | 1 | 24 | 2 | CCKAR | cholecystokinin A receptor | Plasma Membrane | G-protein coupled recepto | T-0632, dexloxiglumide, |
| chr9 | *KRT18P36* | 1 | 29 | 2 | KRT18P36 | keratin 18 pseudogene 36 | Other | other | |
| chr16 | *RP11-812E19.3* | 1 | 42 | 2 | | | | | |
| chr19 | *PSG4* | 1 | 45 | 2 | PSG4 | pregnancy specific beta-1-glycoprotein 4 | Extracellular Space | other | |
| chr1 | *FAM87B* | 1 | 36 | 2 | FAM87B | family with sequence similarity 87 member B | Other | other | |
| chr12 | *DDIT3* | 1 | 28 | 2 | DDIT3 | DNA damage inducible transcript | Nucleus | transcription regulator | |
| chr4 | *RP11-381N20.2* | 1 | 25 | 2 | | | | | |
| chr11 | *RP11-258O13.1* | 1 | 36 | 2 | | | | | |
| chr19 | *F2RL3* | 1 | 53 | 2 | F2RL3 | F2R like thrombin or trypsin receptor 3 | Plasma Membrane | G-protein coupled recepto | |
| chr15 | *CTD-2184D3.6* | 1 | 38 | 2 | | | | | |
| chr13 | *ARGLU1* | 1 | 21 | 2 | ARGLU1 | arginine and glutamate rich 1 | Other | other | |
| chr7 | *RP11-328P23.2* | 1 | 32 | 2 | | | | | |
| chr12 | *RP11-64B16.3* | 1 | 35 | 2 | | | | | |
| chr4 | *SLC25A31* | 1 | 45 | 2 | SLC25A31 | solute carrier family 25 member | Cytoplasm | transporter | |
| chr16 | *RP11-517A5.7* | 1 | 49 | 2 | | | | | |
| chr9 | *CYP4F59P* | 1 | 74 | 2 | | | | | |
| chr17 | *RP11-218M11.1* | 1 | 47 | 2 | | | | | |
| chr19 | *CTC-490E21.13* | 1 | 36 | 2 | | | | | |
| chr17 | *RP11-209M4.1* | 1 | 33 | 2 | | | | | |
| chr7 | *RP11-132A1.4* | 1 | 33 | 2 | | | | | |
| chr19 | *AC002116.7* | 1 | 38 | 2 | | | | | |
| chr11 | *RP11-89C3.4* | 1 | 25 | 2 | | | | | |
| chr1 | *RP11-90C4.1* | 1 | 25 | 2 | | | | | |
| chr11 | *TMEM136* | 1 | 28 | 2 | TLCD5 | TLC domain containing 5 | Other | other | |
| chr12 | *ZNF641* | 1 | 57 | 2 | ZNF641 | zinc finger protein 641 | Other | other | |
| chr11 | *CTD-3074O7.5* | 1 | 91 | 2 | | | | | |
| chr9 | *RP11-65J3.15* | 1 | 53 | 2 | | | | | |
| chr20 | *RP1-300I2.2* | 1 | 43 | 2 | | | | | |
| chr19 | *CEBPA-AS1* | 1 | 51 | 2 | CEBPA-DT | CEBPA divergent transcript | Other | other | |
| chr14 | *RP11-218E20.3* | 1 | 18 | 2 | | | | | |
| chr9 | *NPAP1P6* | 1 | 47 | 2 | NPAP1P6 | nuclear pore associated protein 1 pseudogene 6 | Other | other | |
| chr10 | *FUOM* | 1 | 56 | 2 | FUOM | fucose mutarotase | Cytoplasm | enzyme | |
| chr14 | *SIX1* | 1 | 33 | 2 | SIX1 | SIX homeobox 1 | Nucleus | transcription regulator | |
| chr5 | *CTD-2012J19.3* | 1 | 71 | 2 | | | | | |
| chr18 | *RP11-527H14.2* | 1 | 27 | 2 | | | | | |
| chr1 | *CD1B* | 1 | 34 | 2 | CD1B | CD1b molecule | Plasma Membrane | other | |
| chr16 | *RP11-626G11.5* | 1 | 28 | 2 | | | | | |
| chr8 | *REEP4* | 1 | 54 | 2 | REEP4 | receptor accessory protein 4 | Cytoplasm | other | |
| chr5 | *CTB-181F24.1* | 1 | 28 | 2 | | | | | |
| chr1 | *RP4-590F24.1* | 1 | 36 | 2 | | | | | |
| chr8 | *RECQL4* | 1 | 69 | 2 | RECQL4 | RecQ like helicase 4 | Nucleus | enzyme | |
| chr6 | *HIST1H2BE* | 1 | 39 | 2 | H2BC6 | H2B clustered histone 6 | Nucleus | other | |
| chr17 | *HOXB1* | 1 | 43 | 2 | HOXB1 | homeobox B1 | Nucleus | transcription regulator | |
| chr13 | *LRRC63* | 1 | 23 | 2 | LRRC63 | leucine rich repeat containing 63 | Other | transcription regulator | |
| chr6 | *DEFB112* | 1 | 35 | 2 | DEFB112 | defensin beta 112 | Extracellular Space | other | |
| chr11 | *OR52K3P* | 1 | 52 | 2 | OR52K3P | olfactory receptor family 52 subfamily K member 3 | Other | other | |
| chr8 | *FAM86B3P* | 1 | 91 | 2 | FAM86B3P | family with sequence similarity 86, member A pseudogene | Other | other | |
| chr19 | *AC010504.2* | 1 | 35 | 2 | | | | | |
| chr7 | *AC005102.1* | 1 | 59 | 2 | | | | | |
| chr7 | *NHP2P2* | 1 | 42 | 2 | NHP2P2 | NHP2 ribonucleoprotein pseudogene 2 | Other | other | |
| chr10 | *GLUD1P3* | 1 | 87 | 2 | GLUD1P3 | glutamate dehydrogenase 1 pseudogene 3 | Other | other | |
| chr7 | *C7orf34* | 1 | 33 | 2 | LLCFC1 | LLLL and CFNLAS motif | Other | other | |
| chr8 | *RP11-354A14.1* | 1 | 19 | 2 | | | | | |
| chr1 | *BECN1P1* | 1 | 74 | 2 | BECN2 | beclin 2 | Other | other | |
| chr5 | *CTD-2287O16.3* | 1 | 21 | 2 | | | | | |
| chr3 | *RP11-91K8.4* | 1 | 53 | 2 | | | | | |
| chr1 | *RP5-1056L3.1* | 1 | 23 | 2 | | | | | |
| chr11 | *RP11-169D4.1* | 1 | 53 | 2 | | | | | |
| chr2 | *AC008074.4* | 1 | 39 | 2 | | | | | |
| chr1 | *RP11-389O22.5* | 1 | 59 | 2 | | | | | |
| chr2 | *AL133245.2* | 1 | 25 | 2 | | | | | |
| chrX | *RP6-105D16.1* | 1 | 25 | 2 | | | | | |
| chr21 | *IGSF5* | 1 | 30 | 2 | IGSF5 | immunoglobulin superfamily member 5 | Plasma Membrane | other | |
| chr11 | *RP11-554A11.4* | 1 | 53 | 2 | | | | | |
| chr6 | *U91328.21* | 1 | 38 | 2 | | | | | |
| chr13 | *CCDC122* | 1 | 18 | 2 | CCDC122 | coiled-coil domain containing 122 | Other | other | |
| chr6 | *RP1-60O19.2* | 1 | 24 | 2 | | | | | |
| chr17 | *EIF4A1P9* | 1 | 32 | 2 | EIF4A1P9 | eukaryotic translation initiation factor 4A1 pseudogene 9 | Other | other | |
| chr22 | *RFPL2* | 1 | 23 | 2 | RFPL2 | ret finger protein like 2 | Other | other | |
| chr2 | *AC007248.6* | 1 | 67 | 2 | | | | | |
| chr1 | *CYB5R1* | 1 | 66 | 2 | CYB5R1 | cytochrome b5 reductase 1 | Cytoplasm | enzyme | |
| chr17 | *RND2* | 1 | 45 | 2 | RND2 | Rho family GTPase 2 | Cytoplasm | enzyme | |
| chr22 | *XXbac-B444P24.8* | 1 | 41 | 2 | | | | | |

| | | | | | | | | | |
|---|---|---|---|---|---|---|---|---|---|
| chr5 | *RAB5CP2* | 1 | 33 | 2 | RAB5CP2 | RAB5C, member RAS oncogene family pseudogene 2 | Other | other | |
| chr12 | *MARCH9;* | 1 | 49 | 2 | | | | | |
| chr2 | *AC022311.1* | 1 | 32 | 2 | | | | | |
| chr13 | *UBAC2-AS1* | 1 | 39 | 2 | UBAC2-AS1 | UBAC2 antisense RNA 1 | Other | other | |
| chr11 | *OR51A9P* | 1 | 62 | 2 | OR51A9P | olfactory receptor family 51 subfamily A member 9 | Other | other | |
| chr11 | *RP11-324J3.1* | 1 | 48 | 2 | | | | | |
| chr9 | *RP11-202G18.1* | 1 | 25 | 2 | | | | | |
| chr1 | *RP4-811H24.9* | 1 | 43 | 2 | | | | | |
| chr6 | *POPDC3* | 1 | 19 | 2 | POPDC3 | popeye domain containing 3 | Other | other | |
| chr16 | *MIR3670-1* | 1 | 55 | 2 | mir-3670 | microRNA 3670-1 | Other | microRNA | |
| chr6 | *TMEM244* | 1 | 24 | 2 | TMEM244 | transmembrane protein 244 | Other | other | |
| chr11 | *RP11-583F24.3* | 1 | 40 | 2 | | | | | |
| chr12 | *CLEC2D* | 1 | 28 | 2 | CLEC2D | C-type lectin domain family 2 member D | Plasma Membrane | transmembrane receptor | |
| chr20 | *RP5-1103G7.10* | 1 | 28 | 2 | | | | | |
| chrX | *P2RY4* | 1 | 35 | 2 | P2RY4 | pyrimidinergic receptor P2Y4 | Plasma Membrane | G-protein coupled recepto | |
| chr16 | *hsa-mir-3670-2* | 1 | 51 | 2 | | | | | |
| chr20 | *RP1-300I2.3* | 1 | 41 | 2 | | | | | |
| chr16 | *MT1DP* | 1 | 32 | 2 | MT1DP | metallothionein 1D, pseudogene | Other | other | |
| chr22 | *XXbac-B135H6.15* | 1 | 32 | 2 | | | | | |
| chr17 | *CTD-2116F7.1* | 1 | 45 | 2 | | | | | |
| chr4 | *DGKQ* | 1 | 40 | 2 | DGKQ | diacylglycerol kinase theta | Cytoplasm | kinase | |
| chr11 | *NLRP6* | 1 | 41 | 2 | NLRP6 | NLR family pyrin domain | Plasma Membrane | G-protein coupled recepto | |
| chr7 | *PEG10* | 1 | 64 | 2 | PEG10 | paternally expressed 10 | Nucleus | other | |
| chr1 | *KRT8P45* | 1 | 66 | 2 | KRT8P45 | keratin 8 pseudogene 45 | Other | other | |
| chr2 | *AC008278.3* | 1 | 27 | 2 | | | | | |
| chr19 | *MIR24-2* | 1 | 48 | 2 | mir-24 | microRNA 24-1 | Cytoplasm | microRNA | |
| chr1 | *RP11-547D24.3* | 1 | 67 | 2 | | | | | |
| chr1 | *RP11-396C23.3* | 1 | 64 | 2 | | | | | |
| chr5 | *RP11-1334A24.6* | 1 | 49 | 2 | | | | | |
| chr17 | *RP11-6N17.2* | 1 | 37 | 2 | | | | | |
| chr1 | *RP1-118J21.25* | 1 | 41 | 2 | | | | | |
| chr1 | *SMPDL3B* | 1 | 39 | 2 | SMPDL3B | sphingomyelin phosphodiesterase acid like 3B | Extracellular Space | enzyme | |
| chr17 | *RP11-666A8.11* | 1 | 27 | 2 | | | | | |
| chr19 | *CTD-2542C24.3* | 1 | 30 | 2 | | | | | |
| chr8 | *KB-1410C5.3* | 1 | 29 | 2 | | | | | |
| chr11 | *TPP1* | 1 | 65 | 2 | TPP1 | tripeptidyl peptidase 1 | Cytoplasm | peptidase | |
| chr12 | *RP11-486O12.2* | 1 | 40 | 2 | | | | | |
| chr1 | *RP4-665J23.4* | 1 | 31 | 2 | | | | | |
| chr4 | *RP11-302F12.3* | 1 | 18 | 2 | | | | | |
| chr9 | *CDK2AP2P1* | 1 | 179 | 2 | CDK2AP2P1 | cyclin dependent kinase 2 associated protein 2 pseudogene | Other | other | |
| chr10 | *RP11-564C4.4* | 1 | 66 | 2 | | | | | |
| chr2 | *AC005034.2* | 1 | 50 | 2 | | | | | |
| chr9 | *ALOX15P2* | 1 | 33 | 2 | ALOX15P2 | arachidonate 15-lipoxygenase pseudogene 2 | Other | other | |
| chr14 | *SLC22A17* | 1 | 41 | 2 | SLC22A17 | solute carrier family 22 member | Plasma Membrane | transporter | |
| chr7 | *OR2A14* | 1 | 31 | 2 | OR2A14 | olfactory receptor family 2 subfamily A member 14 | Plasma Membrane | G-protein coupled recepto | |
| chr3 | *RP11-605F14.2* | 1 | 63 | 2 | | | | | |
| chr8 | *GS1-24F4.1* | 1 | 21 | 2 | | | | | |
| chr22 | *TPTEP1* | 1 | 30 | 2 | TPTEP1 | TPTE pseudogene 1 | Other | other | |
| chr3 | *RP11-221E20.4* | 1 | 63 | 2 | | | | | |
| chr12 | *LDHAL6CP* | 1 | 42 | 2 | LDHAL6CP | lactate dehydrogenase A like 6C, pseudogene | Other | other | |
| chr16 | *RP11-354I13.1* | 1 | 61 | 2 | | | | | |
| chr20 | *SPEF1* | 1 | 38 | 2 | SPEF1 | sperm flagellar 1 | Cytoplasm | other | |
| chr1 | *RP5-1185H19.2* | 1 | 41 | 2 | | | | | |
| chr5 | *P4HA2-AS1* | 1 | 38 | 2 | P4HA2-AS1 | P4HA2 antisense RNA 1 | Other | other | |
| chr13 | *FBXL3* | 1 | 41 | 2 | FBXL3 | F-box and leucine rich repeat protein 3 | Nucleus | enzyme | |
| chr15 | *RP11-182J1.11* | 1 | 114 | 2 | | | | | |
| chr6 | *MAN1A1* | 1 | 28 | 2 | MAN1A1 | mannosidase alpha class 1A member 1 | Cytoplasm | enzyme | |
| chr17 | *TUFMP1* | 1 | 36 | 2 | TUFMP1 | Tu translation elongation factor, mitochondrial pseudogene 1 | Other | other | |
| chr12 | *RN7SL769P* | 1 | 34 | 2 | RN7SL769P | | Other | other | |
| chr9 | *C9orf142* | 1 | 79 | 2 | PAXX | PAXX non-homologous end joining factor | Cytoplasm | other | |
| chr2 | *AC073321.3* | 1 | 26 | 2 | | | | | |
| chrX | *TMEM185A* | 1 | 35 | 2 | TMEM185A | transmembrane protein 185A | Nucleus | other | |
| chr1 | *RP3-395M20.9* | 1 | 49 | 2 | | | | | |
| chr2 | *CXCR1* | 1 | 27 | 2 | CXCR1 | C-X-C motif chemokine receptor 1 | Plasma Membrane | G-protein coupled recepto | SB-265610, SX-682, 2'-((4'-trifluoromethanesulfonyloxy)phenyl)-N-methanesulfonylpropionamide, repertaxin |
| chr2 | *AC018696.2* | 1 | 103 | 2 | | | | | |
| chr11 | *POLD4* | 1 | 38 | 2 | POLD4 | DNA polymerase delta 4, accessory subunit | Nucleus | enzyme | |
| chr12 | *RN7SL380P* | 1 | 35 | 2 | RN7SL380P | | Other | other | |
| chr6 | *RP11-138M12.1* | 1 | 23 | 2 | | | | | |
| chr3 | *RP11-88I21.1* | 1 | 61 | 2 | | | | | |
| chrX | *MAGEB16* | 1 | 35 | 2 | MAGEB16 | MAGE family member B16 | Other | other | |
| chr1 | *RP11-533E19.5* | 1 | 37 | 2 | | | | | |

| | | | | | | | | | |
|---|---|---|---|---|---|---|---|---|---|
| chr17 | *LHX1* | 1 | 60 | 2 | LHX1 | LIM homeobox 1 | Nucleus | transcription regulator | |
| chr5 | *RP11-159F24.5* | 1 | 71 | 2 | | | | | |
| chr10 | *RP11-96C23.12* | 1 | 36 | 2 | | | | | |
| chr11 | *OR56A7P* | 1 | 46 | 2 | OR56A7P | olfactory receptor family 56 subfamily A member 7 | Other | other | |
| chr22 | *PANX2* | 1 | 42 | 2 | PANX2 | pannexin 2 | Plasma Membrane | transporter | |
| chr6 | *ULBP3* | 1 | 31 | 2 | ULBP3 | UL16 binding protein 3 | Plasma Membrane | transmembrane receptor | |
| chrX | *ZNF711* | 1 | 40 | 2 | ZNF711 | zinc finger protein 711 | Nucleus | transcription regulator | |
| chr6 | *PRDM1* | 1 | 39 | 2 | PRDM1 | PR/SET domain 1 | Nucleus | transcription regulator | |
| chr2 | *AC016700.5* | 1 | 43 | 2 | | | | | |
| chr14 | *AL117190.3* | 1 | 26 | 2 | | | | | |
| chr5 | *RP11-432M8.16* | 1 | 89 | 2 | | | | | |
| chr4 | *THAP6* | 1 | 22 | 2 | THAP6 | THAP domain containing 6 | Other | other | |
| chr2 | *AC007679.1* | 1 | 34 | 2 | | | | | |
| chr9 | *RP11-104G3.2* | 1 | 67 | 2 | | | | | |
| chr12 | *RNA5SP378* | 1 | 52 | 2 | RNA5SP378 | RNA, 5S ribosomal pseudogene | Other | other | |
| chr1 | *SLC25A38P1* | 1 | 49 | 2 | SLC25A38P1 | solute carrier family 25 member 38 pseudogene 1 | Other | other | |
| chr16 | *COG8* | 1 | 52 | 2 | COG8 | component of oligomeric golgi complex 8 | Cytoplasm | transporter | |
| chr11 | *RP11-358N4.3* | 1 | 44 | 2 | | | | | |
| chr1 | *RP5-934G17.6* | 1 | 61 | 2 | | | | | |
| chr9 | *RP11-344B5.3* | 1 | 47 | 2 | | | | | |
| chr19 | *FAM138F* | 1 | 73 | 2 | FAM138F | family with sequence similarity 138 member F | Other | other | |
| chr7 | *RP11-286H14.4* | 1 | 60 | 2 | | | | | |
| chr11 | *CTD-2530H12.4* | 1 | 60 | 2 | | | | | |
| chr9 | *SNX18P4* | 1 | 50 | 2 | SNX18P4 | sorting nexin 18 pseudogene 4 | Other | other | |
| chr22 | *APOL6* | 1 | 24 | 2 | APOL6 | apolipoprotein L6 | Extracellular Space | transporter | |
| chr5 | *RBMX2P5* | 1 | 33 | 2 | RBMX2P5 | RBMX2 pseudogene 5 | Other | other | |
| chr18 | *CTD-2008L17.2* | 1 | 24 | 2 | | | | | |
| chr12 | *RP11-38F22.1* | 1 | 24 | 2 | | | | | |
| chr16 | *RP11-331F4.4* | 1 | 56 | 2 | | | | | |
| chr10 | *RP11-561G1.3* | 1 | 34 | 2 | | | | | |
| chr4 | *CEP135* | 1 | 22 | 2 | CEP135 | centrosomal protein 135 | Cytoplasm | other | |
| chr10 | *RP11-479A21.1* | 1 | 64 | 2 | | | | | |
| chr11 | *C11orf42* | 1 | 69 | 2 | C11orf42 | chromosome 11 open reading frame 42 | Other | other | |
| chr18 | *RFWD2P1* | 1 | 55 | 2 | COP1P1 | COP1 pseudogene 1 | Other | other | |
| chr17 | *RP11-1407O15.3* | 1 | 115 | 2 | | | | | |
| chr7 | *RPS2P32* | 1 | 72 | 2 | RPS2P32 | ribosomal protein S2 pseudogene | Other | other | |
| chr3 | *XCR1* | 1 | 47 | 2 | XCR1 | X-C motif chemokine receptor 1 | Plasma Membrane | G-protein coupled recepto | |
| chr20 | *RP4-738P15.1* | 1 | 52 | 2 | | | | | |
| chr6 | *HIST1H2BK* | 1 | 41 | 2 | H2BC12 | H2B clustered histone 12 | Nucleus | other | |
| chr19 | *CTD-3137H5.1* | 1 | 73 | 2 | | | | | |
| chr11 | *RPS26P43* | 1 | 38 | 2 | RPS26P43 | ribosomal protein S26 pseudogene 43 | Other | other | |
| chr7 | *RP4-607J2.1* | 1 | 56 | 2 | | | | | |
| chr8 | *RP11-360L9.4* | 1 | 22 | 2 | | | | | |
| chr19 | *ZFP36* | 1 | 46 | 2 | ZFP36 | ZFP36 ring finger protein | Nucleus | transcription regulator | |
| chr7 | *FZD1* | 1 | 42 | 2 | FZD1 | frizzled class receptor 1 | Plasma Membrane | G-protein coupled recepto | vantictumab |
| chr11 | *OR4C4P* | 1 | 66 | 2 | OR4C4P | olfactory receptor family 4 subfamily C member 4 | Other | other | |
| chr11 | *CLCF1* | 1 | 47 | 2 | CLCF1 | cardiotrophin like cytokine factor | Extracellular Space | cytokine | |
| chr5 | *RP11-215G15.5* | 1 | 129 | 2 | | | | | |
| chr3 | *GPR79* | 1 | 56 | 2 | GPR79 | G protein-coupled receptor 79, pseudogene | Plasma Membrane | other | |
| chr6 | *HIST1H2BPS1* | 1 | 26 | 2 | H2BC2P | H2B clustered histone 2, pseudogene | Other | other | |
| chr10 | *RP1-251M9.2* | 1 | 19 | 2 | | | | | |
| chr8 | *NKX3-1* | 1 | 28 | 2 | NKX3-1 | NK3 homeobox 1 | Nucleus | transcription regulator | |
| chr7 | *CCT8L1P* | 1 | 51 | 2 | CCT8L1P | chaperonin containing TCP1 subunit 8 like 1, pseudogene | Other | other | |
| chr1 | *TMEM167B* | 1 | 42 | 2 | TMEM167B | transmembrane protein 167B | Other | other | |
| chr5 | *RP11-35O7.1* | 1 | 89 | 2 | | | | | |
| chrX | *NXF5* | 1 | 34 | 2 | NXF5 | nuclear RNA export factor 5 | Cytoplasm | other | |
| chr16 | *LDHD* | 1 | 40 | 2 | LDHD | lactate dehydrogenase D | Cytoplasm | enzyme | |
| chr2 | *AC013717.3* | 1 | 37 | 2 | | | | | |
| chr17 | *CTB-162E12.1* | 1 | 29 | 2 | | | | | |
| chr17 | *RP11-434D2.9* | 1 | 59 | 2 | | | | | |
| chr1 | *RP11-195M16.3* | 1 | 53 | 2 | | | | | |
| chrX | *KLHL4* | 1 | 25 | 2 | KLHL4 | kelch like family member 4 | Cytoplasm | other | |
| chr20 | *RP5-905G11.3* | 1 | 25 | 2 | | | | | |
| chr15 | *CD24P2* | 1 | 32 | 2 | CD24P2 | CD24 molecule pseudogene 2 | Plasma Membrane | other | |
| chr3 | *RHBDF1P1* | 1 | 63 | 2 | RHBDF1P1 | RHBDF1 pseudogene 1 | Other | other | |
| chr7 | *RP11-274B21.2* | 1 | 42 | 2 | | | | | |
| chr13 | *ERCC5* | 1 | 23 | 2 | ERCC5 | ERCC excision repair 5, endonuclease | Nucleus | enzyme | |
| chr7 | *AC069285.1* | 1 | 71 | 2 | | | | | |
| chr6 | *RP11-503C24.4* | 1 | 29 | 2 | | | | | |
| chr1 | *RP4-597N16.1* | 1 | 40 | 2 | | | | | |
| chr18 | *RP11-529J17.1* | 1 | 24 | 2 | | | | | |
| chr11 | *AP000997.2* | 1 | 35 | 2 | | | | | |
| chr12 | *AC140061.10* | 1 | 51 | 2 | | | | | |
| chr16 | *CTRB2* | 1 | 60 | 2 | CTRB2 | chymotrypsinogen B2 | Cytoplasm | peptidase | |
| chr18 | *CTD-2354A18.1* | 1 | 18 | 2 | | | | | |
| chr6 | *XXbac-BPG34I8.3* | 1 | 29 | 2 | | | | | |
| chr20 | *BPIFA1* | 1 | 44 | 2 | BPIFA1 | BPI fold containing family A member 1 | Extracellular Space | other | |

| | | | | | | | | |
|---|---|---|---|---|---|---|---|---|
| chr7 | *GS1-259H13.7* | 1 | 65 | 2 | | | | |
| chr5 | *KIF20A* | 1 | 36 | 2 | KIF20A | kinesin family member 20A | Cytoplasm | other |
| chr10 | *RP11-31L23.3* | 1 | 56 | 2 | | | | |
| chr1 | *ATP6V0B* | 1 | 61 | 2 | ATP6V0B | ATPase H+ transporting V0 | Cytoplasm | transporter |
| chr8 | *THAP1* | 1 | 29 | 2 | THAP1 | THAP domain containing 1 | Nucleus | transcription regulator |
| chr19 | *AC008753.3* | 1 | 37 | 2 | | | | |
| chr11 | *SCGB2A2* | 1 | 36 | 2 | SCGB2A2 | secretoglobin family 2A member | Extracellular Space | other |
| chr20 | *RPS2P7* | 1 | 67 | 2 | RPS2P7 | ribosomal protein S2 pseudogene | Other | other |
| chr17 | *RP11-498C9.17* | 1 | 58 | 2 | | | | |
| chr9 | *ZNF782* | 1 | 30 | 2 | ZNF782 | zinc finger protein 782 | Cytoplasm | transcription regulator |
| chr9 | *FAM27C* | 1 | 125 | 2 | FAM27C | family with sequence similarity 27 member C | Other | other |
| chrX | *RPS2P55* | 1 | 49 | 2 | RPS2P55 | ribosomal protein S2 pseudogene | Other | other |
| chr11 | *CIR1P3* | 1 | 29 | 2 | CIR1P3 | corepressor interacting with RBPJ, 1 pseudogene 3 | Other | other |
| chr19 | *TTC9B* | 1 | 50 | 2 | TTC9B | tetratricopeptide repeat domain | Other | other |
| chrX | *RP3-526F5.2* | 1 | 25 | 2 | | | | |
| chr12 | *DDX11L8* | 1 | 195 | 2 | DDX11L8 | DEAD/H-box helicase 11 like 8 | Other | other |
| chrX | *GAGE2D* | 1 | 125 | 2 | GE2D (includes ot | G antigen 2D | Other | other |
| chr10 | *PRKCQ-AS1* | 1 | 21 | 2 | PRKCQ-AS1 | PRKCQ antisense RNA 1 | Other | other |
| chr15 | *AC026150.8* | 1 | 26 | 2 | | | | |
| chr19 | *ZNF728* | 1 | 47 | 2 | ZNF728 | zinc finger protein 728 | Other | other |
| chr19 | *CTC-559E9.9* | 1 | 57 | 2 | | | | |
| chr2 | *ATP6V1B1* | 1 | 47 | 2 | ATP6V1B1 | ATPase H+ transporting V1 subunit B1 | Cytoplasm | transporter |
| chr18 | *RP11-720L2.4* | 1 | 33 | 2 | | | | |
| chr1 | *CAPN9* | 1 | 41 | 2 | CAPN9 | calpain 9 | Cytoplasm | peptidase |
| chr10 | *RP11-541N10.3* | 1 | 32 | 2 | | | | |
| chr16 | *RP11-558A11.1* | 1 | 36 | 2 | | | | |
| chr1 | *LCE5A* | 1 | 55 | 2 | LCE5A | late cornified envelope 5A | Other | other |
| chr5 | *CTB-14A14.2* | 1 | 63 | 2 | | | | |
| chr5 | *RP11-321E2.8* | 1 | 1185 | 2 | | | | |
| chr6 | *PRDM13* | 1 | 34 | 2 | PRDM13 | PR/SET domain 13 | Nucleus | enzyme |
| chr22 | *RP1-90L6.3* | 1 | 27 | 2 | | | | |
| chr17 | *RP5-1067M6.3* | 1 | 37 | 2 | | | | |
| chrX | *MYCLP2* | 1 | 54 | 2 | MYCLP2 | MYCL pseudogene 2 | Other | other |
| chr19 | *AC010649.1* | 1 | 19 | 2 | | | | |
| chr8 | *RP11-350F16.1* | 1 | 28 | 2 | | | | |
| chr11 | *CCDC87* | 1 | 73 | 2 | CCDC87 | coiled-coil domain containing 87 | Extracellular Space | other |
| chr16 | *RP11-46C24.3* | 1 | 57 | 2 | | | | |
| chr1 | *MIR34A* | 1 | 43 | 2 | mir-34 | microRNA 34a | Cytoplasm | microRNA |
| chr2 | *C2orf27A* | 1 | 38 | 2 | C2orf27A/C2orf27 | chromosome 2 open reading frame 27B | Other | other |
| chr12 | *PEBP1* | 1 | 22 | 2 | PEBP1 | phosphatidylethanolamine binding protein 1 | Cytoplasm | other |
| chr2 | *DLX2* | 1 | 41 | 2 | DLX2 | distal-less homeobox 2 | Nucleus | transcription regulator |
| chr16 | *C16orf82* | 1 | 35 | 2 | C16orf82 | chromosome 16 open reading frame 82 | Other | other |
| chr9 | *RP11-473E2.3* | 1 | 63 | 2 | | | | |
| chr11 | *RP11-163O19.8* | 1 | 44 | 2 | | | | |
| chr9 | *HMGA1P4* | 1 | 39 | 2 | HMGA1P4 | high mobility group AT-hook 1 pseudogene 4 | Other | other |
| chrX | *USP51* | 1 | 58 | 2 | USP51 | ubiquitin specific peptidase 51 | Other | peptidase |
| chr17 | *AC011193.1* | 1 | 26 | 2 | | | | |
| chr4 | *RP11-446J8.1* | 1 | 21 | 2 | | | | |
| chr1 | *MTMR11* | 1 | 40 | 2 | MTMR11 | myotubularin related protein 11 | Cytoplasm | other |
| chr11 | *GPR137* | 1 | 41 | 2 | GPR137 | G protein-coupled receptor 137 | Other | other |
| chr9 | *RP11-160N1.8* | 1 | 29 | 2 | | | | |
| chr5 | *CTD-2201E18.4* | 1 | 88 | 2 | | | | |
| chrX | *FTHL17* | 1 | 50 | 2 | FTHL17 | ferritin heavy chain like 17 | Other | other |
| chr6 | *HIST1H3J* | 1 | 43 | 2 | H3C12 | H3 clustered histone 12 | Nucleus | other |
| chr6 | *RAET1L* | 1 | 35 | 2 | RAET1L | retinoic acid early transcript 1L | Plasma Membrane | other |
| chr1 | *PIGPP2* | 1 | 32 | 2 | PIGPP2 | phosphatidylinositol glycan anchor biosynthesis class P | Other | other |
| chr7 | *SLC25A1P3* | 1 | 46 | 2 | SLC25A1P3 | solute carrier family 25 member 1 pseudogene 3 | Other | other |
| chr19 | *DUS3L* | 1 | 47 | 2 | DUS3L | dihydrouridine synthase 3 like | Other | other |
| chr10 | *RP11-20J15.2* | 1 | 68 | 2 | | | | |
| chr5 | *CTD-2313D3.1* | 1 | 98 | 2 | | | | |
| chr4 | *RP11-617D20.1* | 1 | 25 | 2 | | | | |
| chr1 | *PHBP11* | 1 | 53 | 2 | PHBP11 | prohibitin pseudogene 11 | Other | other |
| chrX | *XIST* | 1 | 24 | 2 | XIST | X inactive specific transcript | Nucleus | other |
| chr6 | *GSTA5* | 1 | 35 | 2 | GSTA5 | glutathione S-transferase alpha 5 | Cytoplasm | enzyme |
| chr17 | *KRTAP1-5* | 1 | 51 | 2 | KRTAP1-5 | keratin associated protein 1-5 | Cytoplasm | other |
| chr12 | *AL021546.6* | 1 | 22 | 2 | | | | |
| chr6 | *DPPA5* | 1 | 43 | 2 | DPPA5 | developmental pluripotency associated 5 | Cytoplasm | other |

| | | | | | | | | | |
|---|---|---|---|---|---|---|---|---|---|
| chr4 | *CHRNA9* | 1 | 28 | 2 | CHRNA9 | cholinergic receptor nicotinic alpha 9 subunit | Plasma Membrane | transmembrane receptor | succinylcholine, rapacuronium, doxacurium, vecuronium, pipecuronium, metocurine, mivacurium, D-tubocurarine, nicotine, arecoline, pancuronium, atracurium, isoflurane, cisatracurium, rocuronium, amobarbital, mecamylamine |
| chr1 | *EIF1AXP1* | 1 | 60 | 2 | EIF1AXP1 | EIF1AX pseudogene 1 | Other | other | |
| chr9 | *YBX1P10* | 1 | 24 | 2 | YBX1P10 | Y-box binding protein 1 pseudogene 10 | Other | other | |
| chr7 | *MOSPD3* | 1 | 40 | 2 | MOSPD3 | motile sperm domain containing 3 | Other | other | |
| chr11 | *C11orf83* | 1 | 45 | 2 | UQCC3 | ubiquinol-cytochrome c reductase complex assembly factor 3 | Extracellular Space | other | |
| chrX | *AC151961.1* | 1 | 42 | 2 | | | | | |
| chr2 | *DUSP2* | 1 | 67 | 2 | DUSP2 | dual specificity phosphatase 2 | Nucleus | phosphatase | |
| chr6 | *FAM136BP* | 1 | 37 | 2 | FAM136BP | family with sequence similarity 136 member B, pseudogene | Other | other | |
| chr11 | *RP11-56P9.5* | 1 | 77 | 2 | | | | | |
| chr10 | *RP11-481A12.2* | 1 | 60 | 2 | | | | | |
| chr15 | *AC026150.6* | 1 | 63 | 2 | | | | | |
| chr1 | *S100A7L2* | 1 | 47 | 2 | S100A7L2 | S100 calcium binding protein A7 like 2 | Extracellular Space | other | |
| chr22 | *CTA-342B11.2* | 1 | 41 | 2 | | | | | |
| chr11 | *AP5B1* | 1 | 70 | 2 | AP5B1 | adaptor related protein complex 5 subunit beta 1 | Cytoplasm | other | |
| chr17 | *RP13-20L14.1* | 1 | 60 | 2 | | | | | |
| chr12 | *RP1-102E24.6* | 1 | 39 | 2 | | | | | |
| chr21 | *AP001626.2* | 1 | 37 | 2 | | | | | |
| chr4 | *USP17L28* | 1 | 48 | 2 | 17L18 (includes ot | ubiquitin specific peptidase 17 like family member 5 | Nucleus | peptidase | |
| chr11 | *RP11-529A4.7* | 1 | 74 | 2 | | | | | |
| chr7 | *POP7* | 1 | 44 | 2 | POP7 | POP7 homolog, ribonuclease P/MRP subunit | Nucleus | enzyme | |
| chr3 | *RTP3* | 1 | 39 | 2 | RTP3 | receptor transporter protein 3 | Cytoplasm | other | |
| chr2 | *RP11-734K21.4* | 1 | 37 | 2 | | | | | |
| chr20 | *LAMP5* | 1 | 33 | 2 | LAMP5 | lysosomal associated membrane protein family member 5 | Cytoplasm | other | |
| chr15 | *CTD-2651B20.1* | 1 | 40 | 2 | | | | | |
| chr6 | *PLG* | 1 | 22 | 2 | PLG | plasminogen | Extracellular Space | peptidase | reteplase, PLAT, 6-aminocaproic acid, aprotinin, tenecteplase, |
| chr11 | *RP11-140L24.3* | 1 | 31 | 2 | | | | | |
| chr17 | *RP13-516M14.4* | 1 | 41 | 2 | | | | | |
| chr16 | *C16orf86* | 1 | 57 | 2 | C16orf86 | chromosome 16 open reading frame 86 | Other | other | |
| chr1 | *AL031602.1* | 1 | 34 | 2 | | | | | |
| chr19 | *ZNF324* | 1 | 50 | 2 | ZNF324 | zinc finger protein 324 | Nucleus | transcription regulator | |
| chr2 | *PLGLB1* | 1 | 21 | 2 | PLGLB1/PLGLB2 | plasminogen like B2 | Extracellular Space | peptidase | |
| chr7 | *EIF4HP1* | 1 | 59 | 2 | EIF4HP1 | eukaryotic translation initiation factor 4H pseudogene 1 | Other | other | |
| chr5 | *RP11-451H23.2* | 1 | 55 | 2 | | | | | |
| chr15 | *RP11-529J17.5* | 1 | 48 | 2 | | | | | |
| chr14 | *AL133168.3* | 1 | 23 | 2 | | | | | |
| chr8 | *TUBBP1* | 1 | 48 | 2 | TUBBP1 | tubulin beta pseudogene 1 | Other | other | |
| chr11 | *RP11-672A2.4* | 1 | 68 | 2 | | | | | |
| chr1 | *RP4-591L5.1* | 1 | 35 | 2 | | | | | |
| chr17 | *ALYREF* | 1 | 57 | 2 | ALYREF | Aly/REF export factor | Nucleus | transcription regulator | |
| chr5 | *RP11-432M8.8* | 1 | 77 | 2 | | | | | |
| chr1 | *RP4-561L24.3* | 1 | 63 | 2 | | | | | |
| chr2 | *AC010095.7* | 1 | 38 | 2 | | | | | |
| chr3 | *ARIH2OS* | 1 | 39 | 2 | ARIH2OS | ariadne RBR E3 ubiquitin protein ligase 2 opposite strand | Other | other | |
| chr15 | *KBTBD13* | 1 | 65 | 2 | KBTBD13 | kelch repeat and BTB domain containing 13 | Cytoplasm | other | |
| chr1 | *DEDD* | 1 | 37 | 2 | DEDD | death effector domain containing | Nucleus | other | |
| chr1 | *XX-FW84067D5.1* | 1 | 26 | 2 | | | | | |
| chr11 | *SRPR* | 1 | 34 | 2 | SRPRA | SRP receptor subunit alpha | Cytoplasm | other | |
| chr5 | *RP11-432M8.9* | 1 | 105 | 2 | | | | | |
| chr1 | *FAM43B* | 1 | 40 | 2 | FAM43B | family with sequence similarity 43 member B | Other | other | |
| chr7 | *HOXA11* | 1 | 58 | 2 | HOXA11 | homeobox A11 | Nucleus | transcription regulator | |
| chr4 | *SRP72* | 1 | 24 | 2 | SRP72 | signal recognition particle 72 | Nucleus | kinase | |
| chr17 | *ARL4D* | 1 | 47 | 2 | ARL4D | ADP ribosylation factor like GTPase 4D | Nucleus | enzyme | |
| chr18 | *RP11-861E21.1* | 1 | 37 | 2 | | | | | |
| chrX | *HSD17B10* | 1 | 51 | 2 | HSD17B10 | hydroxysteroid 17-beta dehydrogenase 10 | Cytoplasm | enzyme | |
| chr4 | *USP17L30* | 1 | 49 | 2 | 17L18 (includes ot | ubiquitin specific peptidase 17 like family member 5 | Nucleus | peptidase | |

| | | | | | | | | | |
|---|---|---|---|---|---|---|---|---|---|
| chr2 | *RP11-223I10.1* | 1 | 37 | 2 | | | | | |
| chr8 | *RP11-1C8.4* | 1 | 39 | 2 | | | | | |
| chr6 | *HIST1H3F* | 1 | 36 | 2 | H3C7 | H3 clustered histone 7 | Nucleus | other | |
| chr15 | *LINC00926* | 1 | 24 | 2 | LINC00926 | long intergenic non-protein coding RNA 926 | Other | other | |
| chrX | *GPR50-AS1* | 1 | 37 | 2 | GPR50-AS1 | GPR50 antisense RNA 1 | Other | other | |
| chr7 | *RP11-760D2.7* | 1 | 20 | 2 | | | | | |
| chr5 | *CTD-2139B15.1* | 1 | 115 | 2 | | | | | |
| chr5 | *RP11-321E2.6* | 1 | 99 | 2 | | | | | |
| chr10 | *TMEM14D* | 1 | 44 | 2 | TMEM14DP | transmembrane protein 14D, pseudogene | Other | other | |
| chr11 | *SSU72P4* | 1 | 129 | 2 | SSU72P4 | SSU72 pseudogene 4 | Other | other | |
| chr16 | *RP11-293B20.3* | 1 | 33 | 2 | | | | | |
| chr1 | *RP11-426L16.9* | 1 | 53 | 2 | | | | | |
| chr18 | *RP11-705O1.8* | 1 | 28 | 2 | | | | | |
| chr8 | *RP11-546K22.1* | 1 | 21 | 2 | | | | | |
| chr13 | *KDELC1* | 1 | 52 | 2 | POGLUT2 | protein O-glucosyltransferase 2 | Cytoplasm | enzyme | |
| chr11 | *RP11-390K5.1* | 1 | 53 | 2 | | | | | |
| chr6 | *RSPO3* | 1 | 34 | 2 | RSPO3 | R-spondin 3 | Extracellular Space | kinase | OMP-131R10 |
| chr1 | *RP11-504P24.8* | 1 | 74 | 2 | | | | | |
| chr19 | *CTD-3148I10.9* | 1 | 37 | 2 | | | | | |
| chr3 | *HYAL2* | 1 | 50 | 2 | HYAL2 | hyaluronidase 2 | Cytoplasm | enzyme | diclofenac/hyaluronic acid, hyaluronic acid, hylan |
| chr5 | *SLC36A2* | 1 | 22 | 2 | SLC36A2 | solute carrier family 36 member 2 | Plasma Membrane | transporter | |
| chr15 | *IGHV4OR15-8* | 1 | 86 | 2 | IGHV4OR15-8 | immunoglobulin heavy variable 4/OR15-8 (non-functional) | Other | other | |
| chr1 | *RP11-389O22.4* | 1 | 32 | 2 | | | | | |
| chr17 | *SCDP1* | 1 | 44 | 2 | SCDP1 | stearoyl-CoA desaturase pseudogene 1 | Other | other | |
| chr9 | *TLK1P1* | 1 | 24 | 2 | TLK1P1 | tousled like kinase 1 pseudogene | Other | other | |
| chr3 | *RP11-129B22.1* | 1 | 24 | 2 | | | | | |
| chr17 | *KRTAP9-2* | 1 | 52 | 2 | KRTAP9-2 | keratin associated protein 9-2 | Cytoplasm | other | |
| chr11 | *MRPL16* | 1 | 45 | 2 | MRPL16 | mitochondrial ribosomal protein | Cytoplasm | other | |
| chr16 | *AC137934.1* | 1 | 44 | 2 | | | | | |
| chr20 | *RP5-908M14.5* | 1 | 90 | 2 | | | | | |
| chr17 | *LIMD2* | 1 | 51 | 2 | LIMD2 | LIM domain containing 2 | Other | other | |
| chr11 | *SSSCA1* | 1 | 78 | 2 | ZNRD2 | zinc ribbon domain containing 2 | Other | other | |
| chr8 | *LYPD2* | 1 | 65 | 2 | LYPD2 | LY6/PLAUR domain containing 2 | Cytoplasm | other | |
| chr6 | *ANKRD66* | 1 | 35 | 2 | ANKRD66 | ankyrin repeat domain 66 | Other | other | |
| chr19 | *PRTN3* | 1 | 37 | 2 | PRTN3 | proteinase 3 | Extracellular Space | peptidase | |
| chr14 | *C14orf64* | 1 | 30 | 2 | LINC01550 | long intergenic non-protein coding RNA 1550 | Other | other | |
| chr6 | *RP11-732M18.3* | 1 | 42 | 2 | | | | | |
| chr10 | *RP11-332O19.5* | 1 | 27 | 2 | | | | | |
| chr17 | *UBBP4* | 1 | 39 | 2 | UBBP4 | ubiquitin B pseudogene 4 | Other | other | |
| chr3 | *RP11-67L2.2* | 1 | 37 | 2 | | | | | |
| chr14 | *DDX24* | 1 | 19 | 2 | DDX24 | DEAD-box helicase 24 | Nucleus | enzyme | |
| chr1 | *IFI44* | 1 | 23 | 2 | IFI44 | interferon induced protein 44 | Cytoplasm | other | |
| chr11 | *RP11-211G23.1* | 1 | 57 | 2 | | | | | |
| chr5 | *CTB-35F21.2* | 1 | 44 | 2 | | | | | |
| chrX | *XAGE1B* | 1 | 52 | 2 | XAGE1A/XAGE1E | X antigen family member 1B | Nucleus | other | |
| chr10 | *DCLRE1CP1* | 1 | 26 | 2 | DCLRE1CP1 | DNA cross-link repair 1C pseudogene 1 | Other | other | |
| chr12 | *RP11-627K11.1* | 1 | 59 | 2 | | | | | |
| chr7 | *GS1-124K5.9* | 1 | 54 | 2 | | | | | |
| chr14 | *RN7SL586P* | 1 | 22 | 2 | RN7SL586P | RNA, 7SL, cytoplasmic 586, pseudogene | Other | other | |
| chrX | *RP11-31H15.2* | 1 | 52 | 2 | | | | | |
| chr3 | *RP11-190C22.1* | 1 | 42 | 2 | | | | | |
| chr2 | *AC111200.1* | 1 | 47 | 2 | | | | | |
| chr19 | *GGN* | 1 | 49 | 2 | GGN | gametogenetin | Nucleus | other | |
| chr7 | *RP11-760D2.5* | 1 | 59 | 2 | | | | | |
| chr8 | *RP11-758M4.4* | 1 | 43 | 2 | | | | | |
| chr1 | *DPM3* | 1 | 64 | 2 | DPM3 | dolichyl-phosphate mannosyltransferase subunit 3, regulatory | Cytoplasm | enzyme | |
| chr19 | *OR10H5* | 1 | 53 | 2 | OR10H5 | olfactory receptor family 10 subfamily H member 5 | Plasma Membrane | -protein coupled recepto | |
| chr6 | *HIST1H2AA* | 1 | 28 | 2 | H2AC1 | H2A clustered histone 1 | Nucleus | other | |
| chr1 | *SMIM1* | 1 | 56 | 2 | SMIM1 | small integral membrane protein 1 (Vel blood group) | Cytoplasm | other | |
| chr5 | *CCNO* | 1 | 46 | 2 | CCNO | cyclin O | Nucleus | other | |
| chr19 | *ZNF563* | 1 | 36 | 2 | ZNF563 | zinc finger protein 563 | Nucleus | transcription regulator | |
| chr6 | *RP5-874C20.3* | 1 | 30 | 2 | | | | | |
| chrX | *SPANXD* | 1 | 38 | 2 | NXB1 (includes ot | SPANX family member C | Nucleus | other | |
| chr12 | *CERS5* | 1 | 43 | 2 | CERS5 | ceramide synthase 5 | Cytoplasm | transcription regulator | |
| chr17 | *RP11-259G18.2* | 1 | 69 | 2 | | | | | |
| chr7 | *TRGV4* | 1 | 43 | 2 | TRGV4 | T cell receptor gamma variable 4 | Other | other | |
| chr1 | *RP5-886K2.1* | 1 | 32 | 2 | | | | | |
| chrX | *ERAS* | 1 | 60 | 2 | ERAS | ES cell expressed Ras | Plasma Membrane | enzyme | |
| chr17 | *PSMD7P1* | 1 | 28 | 2 | PSMD7P1 | proteasome 26S subunit, non-ATPase 7 pseudogene 1 | Other | other | |
| chr12 | *RP11-497G19.2* | 1 | 22 | 2 | | | | | |
| chrX | *RP13-487C10.1* | 1 | 62 | 2 | | | | | |
| chr11 | *OR10Q1* | 1 | 55 | 2 | OR10Q1 | olfactory receptor family 10 subfamily Q member 1 | Plasma Membrane | -protein coupled recepto | |

| | | | | | | | | | |
|---|---|---|---|---|---|---|---|---|---|
| chr4 | *KDR* | 1 | 29 | 2 | KDR | kinase insert domain receptor | Plasma Membrane | kinase | CHIR-265, midostaurin, sorafenib/sulindac/sunitinib, apatinib, brivanib, imatinib/sunitinib, bevacizumab/sorafenib, tanibirumab, axitinib, motesanib, 5-azacytidine/sorafenib, tesevatinib, pazopanib, XL999, JI 101, PF 00337210, famitinib, SAR103168, telatinib, brivanib alaninate, henatinib, VEGFR2 inhibitor, pegdinetanib, rebastinib, orantinib, paclitaxel/pembrolizumab/ramucirumab, sulfatinib, vatalanib |
| chr8 | *TMEM249* | 1 | 58 | 2 | TMEM249 | transmembrane protein 249 | Other | other | |
| chr10 | *RP11-272J7.6* | 1 | 64 | 2 | | | | | |
| chr10 | *RP11-733D4.1* | 1 | 50 | 2 | | | | | |
| chr21 | *CRYAA* | 1 | 48 | 2 | CRYAA/CRYAA2 | crystallin alpha A | Cytoplasm | other | |
| chr11 | *CD151* | 1 | 44 | 2 | CD151 | CD151 molecule (Raph blood | Plasma Membrane | other | |
| chr8 | *CTD-2342N23.3* | 1 | 48 | 2 | | | | | |
| chr7 | *PQLC1P1* | 1 | 33 | 2 | PQLC1P1 | | Other | other | |
| chr20 | *RP13-379O24.2* | 1 | 104 | 2 | | | | | |
| chr11 | *RNASEH2C* | 1 | 80 | 2 | RNASEH2C | ribonuclease H2 subunit C | Other | other | |
| chr14 | *NPC2* | 1 | 23 | 2 | NPC2 | NPC intracellular cholesterol transporter 2 | Extracellular Space | transporter | |
| chr19 | *DIRAS1* | 1 | 36 | 2 | DIRAS1 | DIRAS family GTPase 1 | Plasma Membrane | enzyme | |
| chr14 | *DIO3* | 1 | 43 | 2 | DIO3 | iodothyronine deiodinase 3 | Plasma Membrane | enzyme | |
| chr12 | *RP11-495K9.9* | 1 | 66 | 2 | | | | | |
| chr5 | *YBX1P5* | 1 | 61 | 2 | YBX1P5 | Y-box binding protein 1 pseudogene 5 | Other | other | |
| chr16 | *HCFC1R1* | 1 | 60 | 2 | HCFC1R1 | host cell factor C1 regulator 1 | Nucleus | other | |
| chr5 | *FST* | 1 | 40 | 2 | FST | follistatin | Extracellular Space | other | |
| chr17 | *RP11-209D14.2* | 1 | 44 | 2 | | | | | |
| chr17 | *TMEM107* | 1 | 59 | 2 | TMEM107 | transmembrane protein 107 | Extracellular Space | other | |
| chr20 | *FOXS1* | 1 | 72 | 2 | FOXS1 | forkhead box S1 | Nucleus | transcription regulator | |
| chr5 | *RP11-310P5.2* | 1 | 137 | 2 | | | | | |
| chr8 | *EXOSC4* | 1 | 62 | 2 | EXOSC4 | exosome component 4 | Nucleus | enzyme | |
| chr17 | *RP1-56K13.6* | 1 | 24 | 2 | | | | | |
| chr5 | *RP11-289K10.1* | 1 | 33 | 2 | | | | | |
| chr19 | *RAB4B-EGLN2* | 1 | 41 | 2 | RAB4B-EGLN2 | RAB4B-EGLN2 readthrough (NMD candidate) | Other | other | |
| chr12 | *POP5* | 1 | 52 | 2 | POP5 | POP5 homolog, ribonuclease P/MRP subunit | Nucleus | enzyme | |
| chr16 | *RP11-388M20.1* | 1 | 32 | 2 | | | | | |
| chr9 | *CDK9* | 1 | 83 | 2 | CDK9 | cyclin dependent kinase 9 | Nucleus | kinase | alvocidib, AZD4573, riviciclib, roniciclib, dinaciclib, BAY1251152, AZD5438, CYC065, CDK9 inhibitor, SB-1317, BMS-387032, roscovitine |
| chr16 | *LA16c-380A1.1* | 1 | 59 | 2 | | | | | |
| chr19 | *HSPB6* | 1 | 52 | 2 | HSPB6 | heat shock protein family B (small) member 6 | Cytoplasm | other | |
| chr10 | *EMX2* | 1 | 40 | 2 | EMX2 | empty spiracles homeobox 2 | Nucleus | transcription regulator | |
| chr11 | *C11orf40* | 1 | 29 | 2 | C11orf40 | chromosome 11 open reading frame 40 (putative) | Other | other | |
| chr11 | *RP11-335F8.2* | 1 | 29 | 2 | | | | | |
| chr14 | *IGHG3* | 1 | 43 | 2 | IGHG3 | immunoglobulin heavy constant gamma 3 (G3m marker) | Extracellular Space | other | |
| chr8 | *RPL8* | 1 | 60 | 2 | RPL8 | ribosomal protein L8 | Cytoplasm | other | |
| chr1 | *RP11-474O21.5* | 1 | 32 | 2 | | | | | |
| chr9 | *RP11-216L13.17* | 1 | 57 | 2 | | | | | |
| chr11 | *RP11-215H18.5* | 1 | 44 | 2 | | | | | |
| chr1 | *PIFO* | 1 | 60 | 2 | PIFO | primary cilia formation | Cytoplasm | other | |
| chr1 | *PFN1P3* | 1 | 72 | 2 | PFN1P3 | profilin 1 pseudogene 3 | Other | other | |
| chr1 | *ZC3H11B* | 1 | 38 | 2 | ZC3H11B | zinc finger CCCH-type containing 11B | Other | other | |
| chr4 | *RP11-692E14.1* | 1 | 29 | 2 | | | | | |
| chr20 | *RPL7P3* | 1 | 80 | 2 | RPL7P3 | ribosomal protein L7 pseudogene | Other | other | |
| chr16 | *HSD3B7* | 1 | 49 | 2 | HSD3B7 | hydroxy-delta-5-steroid dehydrogenase, 3 beta- and steroid delta-isomerase 7 | Cytoplasm | enzyme | |
| chr19 | *JOSD2* | 1 | 46 | 2 | JOSD2 | Josephin domain containing 2 | Cytoplasm | enzyme | |
| chr16 | *RP11-2C24.4* | 1 | 46 | 2 | | | | | |
| chr3 | *VENTXP7* | 1 | 52 | 2 | VENTXP7 | VENT homeobox pseudogene 7 | Other | other | |
| chr13 | *CHAMP1* | 1 | 31 | 2 | CHAMP1 | chromosome alignment maintaining phosphoprotein 1 | Nucleus | other | |

| | | | | | | | | | |
|---|---|---|---|---|---|---|---|---|---|
| chr19 | *GPR32* | 1 | 44 | 2 | GPR32 | G protein-coupled receptor 32 | Plasma Membrane | G-protein coupled recepto | |
| chrX | *FTH1P18* | 1 | 42 | 2 | FTHL18 | ferritin heavy chain like 18 | Other | other | |
| chr9 | *NPAP1P4* | 1 | 36 | 2 | NPAP1P4 | nuclear pore associated protein 1 pseudogene 4 | Other | other | |
| chr19 | *APC2* | 1 | 57 | 2 | APC2 | APC regulator of WNT signaling pathway 2 | Cytoplasm | enzyme | |
| chr19 | *REEP6* | 1 | 48 | 2 | REEP6 | receptor accessory protein 6 | Plasma Membrane | other | |
| chr5 | *MZB1* | 1 | 58 | 2 | MZB1 | marginal zone B and B1 cell specific protein | Extracellular Space | other | |
| chr16 | *RP11-95H3.1* | 1 | 18 | 2 | | | | | |
| chr1 | *RP5-1061H20.4* | 1 | 30 | 2 | | | | | |
| chr5 | *SLC25A2* | 1 | 48 | 2 | SLC25A2 | solute carrier family 25 member 2 | Cytoplasm | transporter | |
| chr15 | *TVP23BP1* | 1 | 38 | 2 | TVP23BP1 | TVP23B pseudogene 1 | Other | other | |
| chr3 | *CNN2P6* | 1 | 53 | 2 | CNN2P6 | calponin 2 pseudogene 6 | Other | other | |
| chr2 | *C2orf83* | 1 | 33 | 2 | C2orf83 | chromosome 2 open reading frame 83 | Other | transporter | |
| chr6 | *PRELID1P1* | 1 | 30 | 2 | PRELID1P1 | PRELI domain containing 1 pseudogene 1 | Other | other | |
| chr10 | *EIF5AP4* | 1 | 24 | 2 | EIF5AP4 | eukaryotic translation initiation factor 5A pseudogene 4 | Other | other | |
| chr6 | *RP4-668J24.2* | 1 | 48 | 2 | | | | | |
| chr2 | *AC012065.7* | 1 | 22 | 2 | | | | | |
| chr1 | *GLUL* | 1 | 52 | 2 | GLUL | glutamate-ammonia ligase | Cytoplasm | enzyme | |
| chr1 | *PFN1P1* | 1 | 60 | 2 | PFN1P1 | profilin 1 pseudogene 1 | Other | other | |
| chr6 | *PTCHD3P3* | 1 | 45 | 2 | PTCHD3P3 | patched domain containing 3 pseudogene 3 | Other | other | |
| chr14 | *DACT1* | 1 | 37 | 2 | DACT1 | dishevelled binding antagonist of beta catenin 1 | Cytoplasm | other | |
| chr2 | *HTRA2* | 1 | 45 | 2 | HTRA2 | HtrA serine peptidase 2 | Cytoplasm | peptidase | |
| chr20 | *PSMA7* | 1 | 66 | 2 | PSMA7 | proteasome subunit alpha 7 | Cytoplasm | peptidase | |
| chr9 | *CLIC3* | 1 | 74 | 2 | CLIC3 | chloride intracellular channel 3 | Nucleus | ion channel | |
| chr5 | *LRRC14B* | 1 | 164 | 2 | LRRC14B | leucine rich repeat containing | Other | other | |
| chr16 | *RP11-467J12.1* | 1 | 41 | 2 | | | | | |
| chr1 | *LINC00982* | 1 | 53 | 2 | PRDM16-DT | PRDM16 divergent transcript | Other | other | |
| chr6 | *HIST1H1A* | 1 | 40 | 2 | H1-1 | H1.1 linker histone, cluster | Nucleus | other | |
| chr2 | *AC097532.2* | 1 | 59 | 2 | | | | | |
| chrX | *PAGE2B* | 1 | 53 | 2 | PAGE2/PAGE2B | PAGE family member 2 | Other | other | |
| chr1 | *SPRR4* | 1 | 30 | 2 | SPRR4 | small proline rich protein 4 | Cytoplasm | other | |
| chr1 | *CHCHD2P6* | 1 | 52 | 2 | CHCHD2P6 | coiled-coil-helix-coiled-coil-helix domain containing 2 pseudogene | Other | other | |
| chr5 | *KRT18P56* | 1 | 109 | 2 | KRT18P56 | keratin 18 pseudogene 56 | Other | other | |
| chr10 | *RP11-40F6.1* | 1 | 38 | 2 | | | | | |
| chr11 | *RP11-730K11.1* | 1 | 47 | 2 | | | | | |
| chr3 | *LINC00888* | 1 | 30 | 2 | LINC00888 | long intergenic non-protein coding RNA 888 | Other | other | |
| chr7 | *AC006026.12* | 1 | 41 | 2 | | | | | |
| chr11 | *KCNJ11* | 1 | 63 | 2 | KCNJ11 | potassium inwardly rectifying channel subfamily J member 11 | Plasma Membrane | ion channel | nateglinide, chlorpropamide, acetohexamide, thiamylal, tolazamide, repaglinide, glyburide, glipizide, glimepiride/pioglitazone, minoxidil, phentolamine, glimepiride/rosiglitaz |
| chr15 | *RP11-352D13.6* | 1 | 47 | 2 | | | | | |
| chr5 | *ADAMTS19-AS1* | 1 | 51 | 2 | ADAMTS19-AS1 | ADAMTS19 antisense RNA 1 | Other | other | |
| chr12 | *RP11-386G11.5* | 1 | 60 | 2 | | | | | |
| chr14 | *DHRS4L2* | 1 | 40 | 2 | DHRS4L2 | dehydrogenase/reductase 4 like 2 | Other | other | |
| chr1 | *RP11-740P5.2* | 1 | 39 | 2 | | | | | |
| chr19 | *RP11-157B13.3* | 1 | 30 | 2 | | | | | |
| chrX | *PRRG3* | 1 | 30 | 2 | PRRG3 | proline rich and Gla domain 3 | Plasma Membrane | other | |
| chr19 | *HMGN1P32* | 1 | 43 | 2 | HMGN1P32 | high mobility group nucleosome binding domain 1 pseudogene 32 | Other | other | |
| chr15 | *SPATA8* | 1 | 55 | 2 | SPATA8 | spermatogenesis associated 8 | Other | other | |
| chr16 | *MAF* | 1 | 40 | 2 | MAF | MAF bZIP transcription factor | Nucleus | transcription regulator | |
| chr8 | *RP11-661A12.14* | 1 | 59 | 2 | | | | | |
| chr3 | *RP11-278L15.3* | 1 | 51 | 2 | | | | | |
| chr5 | *CTC-573N18.1* | 1 | 53 | 2 | | | | | |
| chr20 | *RP3-461P17.9* | 1 | 57 | 2 | | | | | |
| chr3 | *RP11-413E6.7* | 1 | 5555 | 2 | | | | | |
| chrX | *LINC00086* | 1 | 35 | 2 | SMIM10L2A | small integral membrane protein 10 like 2A | Other | other | |
| chr17 | *KRTAP4-12* | 1 | 56 | 2 | KRTAP4-12 | keratin associated protein 4-12 | Other | other | |
| chr19 | *OR10H2* | 1 | 49 | 2 | OR10H2 | olfactory receptor family 10 subfamily H member 2 | Plasma Membrane | other | |
| chr19 | *hsa-mir-1199* | 1 | 44 | 2 | | | | | |
| chr2 | *MIR1285-2* | 1 | 33 | 2 | mir-1285 | microRNA 1285-2 | Cytoplasm | microRNA | |
| chr22 | *SLC25A1* | 1 | 35 | 2 | SLC25A1 | solute carrier family 25 member 1 | Other | transporter | |
| chr2 | *TMEM198* | 1 | 32 | 2 | TMEM198 | transmembrane protein 198 | Other | other | |
| chr5 | *CTC-534B23.1* | 1 | 44 | 2 | | | | | |
| chr11 | *AP003774.4* | 1 | 34 | 2 | | | | | |
| chr10 | *RP11-336A10.7* | 1 | 36 | 2 | | | | | |
| chr17 | *RP11-74E22.6* | 1 | 47 | 2 | | | | | |
| chr2 | *IHH* | 1 | 39 | 2 | IHH | Indian hedgehog signaling | Extracellular Space | enzyme | |
| chr15 | *KRT8P24* | 1 | 53 | 2 | KRT8P24 | keratin 8 pseudogene 24 | Other | other | |

| | | | | | | | | | |
|---|---|---|---|---|---|---|---|---|---|
| chr19 | *GTF2F1* | 1 | 39 | 2 | GTF2F1 | general transcription factor IIF subunit 1 | Nucleus | transcription regulator | |
| chr2 | *EMILIN1* | 1 | 40 | 2 | EMILIN1 | elastin microfibril interfacer 1 | Extracellular Space | other | |
| chr7 | *TMEM60* | 1 | 53 | 2 | TMEM60 | transmembrane protein 60 | Other | other | |
| chr3 | *HTR3D* | 1 | 34 | 2 | HTR3D | 5-hydroxytryptamine receptor 3D | Plasma Membrane | ion channel | fenfluramine, dolasetron, ergotamine, palonosetron, alosetron, cisapride, granisetron, D-tubocurarine, |
| chr19 | *OVOL3* | 1 | 44 | 2 | OVOL3 | ovo like zinc finger 3 | Other | other | |
| chr16 | *RP11-452L6.5* | 1 | 32 | 2 | | | | | |
| chr5 | *RP11-129I19.2* | 1 | 78 | 2 | | | | | |
| chr15 | *MIR1268A* | 1 | 79 | 2 | mir-1268 | microRNA 1268a | Cytoplasm | microRNA | |
| chr7 | *ELK1P1* | 1 | 68 | 2 | ELK1P1 | ELK1 pseudogene 1 | Other | other | |
| chr19 | *OR1AB1P* | 1 | 33 | 2 | OR1AB1P | olfactory receptor family 1 subfamily AB member 1 | Other | other | |
| chr1 | *RP11-385M4.2* | 1 | 63 | 2 | | | | | |
| chr16 | *TPSB2* | 1 | 58 | 2 | TPSAB1/TPSB2 | tryptase alpha/beta 1 | Extracellular Space | peptidase | |
| chr13 | *LINC00381* | 1 | 39 | 2 | LINC00381 | long intergenic non-protein coding RNA 381 | Other | other | |
| chr4 | *USP17L29* | 1 | 51 | 2 | 17L18 (includes ot | ubiquitin specific peptidase 17 like family member 5 | Nucleus | peptidase | |
| chr9 | *RNF224* | 1 | 78 | 2 | RNF224 | ring finger protein 224 | Other | other | |
| chr1 | *TMEM52* | 1 | 58 | 2 | TMEM52 | transmembrane protein 52 | Other | other | |
| chr2 | *SGOL2* | 1 | 38 | 2 | SGO2 | shugoshin 2 | Nucleus | other | |
| chr1 | *USP21* | 1 | 57 | 2 | USP21 | ubiquitin specific peptidase 21 | Cytoplasm | peptidase | |
| chr14 | *FOXA1* | 1 | 55 | 2 | FOXA1 | forkhead box A1 | Nucleus | transcription regulator | |
| chr10 | *RP11-179B15.6* | 1 | 68 | 2 | | | | | |
| chr3 | *RP11-501O2.3* | 1 | 25 | 2 | | | | | |
| chr4 | *HTT-AS* | 1 | 29 | 2 | HTT-AS | HTT antisense RNA | Other | other | |
| chr13 | *ZIC2* | 1 | 57 | 2 | ZIC2 | Zic family member 2 | Nucleus | transcription regulator | |
| chr7 | *KIAA0087* | 1 | 27 | 2 | KIAA0087 | KIAA0087 lncRNA | Other | other | |
| chr17 | *TSEN15P1* | 1 | 46 | 2 | TSEN15P1 | tRNA splicing endonuclease subunit 15 pseudogene 1 | Other | other | |
| chr7 | *RP4-701O16.5* | 1 | 34 | 2 | | | | | |
| chr7 | *RP11-90P13.1* | 1 | 65 | 2 | | | | | |
| chrX | *PAGE3* | 1 | 46 | 2 | PAGE3 | PAGE family member 3 | Other | other | |
| chr1 | *RXFP4* | 1 | 72 | 2 | RXFP4 | relaxin family peptide/INSL5 receptor 4 | Plasma Membrane | -protein coupled recepto | |
| chr12 | *RPS27P25* | 1 | 45 | 2 | RPS27P25 | ribosomal protein S27 pseudogene 25 | Other | other | |
| chr7 | *AC019117.1* | 1 | 43 | 2 | | | | | |
| chr8 | *TIGD5* | 1 | 65 | 2 | TIGD5 | tigger transposable element | Other | other | |
| chr14 | *LINC00221* | 1 | 59 | 2 | LINC00221 | long intergenic non-protein coding RNA 221 | Other | other | |
| chr19 | *CLDND2* | 1 | 49 | 2 | CLDND2 | claudin domain containing 2 | Other | other | |
| chr17 | *KRTAP4-3* | 1 | 57 | 2 | KRTAP4-3 | keratin associated protein 4-3 | Cytoplasm | other | |
| chr19 | *CTB-25J19.1* | 1 | 71 | 2 | | | | | |
| chr4 | *MFSD7* | 1 | 38 | 2 | SLC49A3 | solute carrier family 49 member 3 | Other | other | |
| chr17 | *KRTAP4-7* | 1 | 58 | 2 | KRTAP4-7 | keratin associated protein 4-7 | Other | other | |
| chr12 | *H3F3C* | 1 | 46 | 2 | H3-5 | H3.5 histone | Other | other | |
| chr1 | *MATN1-AS1* | 1 | 37 | 2 | MATN1-AS1 | MATN1 antisense RNA 1 | Other | other | |
| chr15 | *ABCB10P1* | 1 | 127 | 2 | ABCB10P1 | ABCB10 pseudogene 1 | Other | other | |
| chr2 | *FTH1P6* | 1 | 59 | 2 | FTH1P6 | | Other | other | |
| chr5 | *SNHG18* | 1 | 67 | 2 | SNHG18 | small nucleolar RNA host gene | Other | other | |
| chr16 | *NUDT16L1* | 1 | 54 | 2 | NUDT16L1 | nudix hydrolase 16 like 1 | Cytoplasm | other | |
| chr7 | *HNRNPA2B1* | 1 | 62 | 2 | HNRNPA2B1 | heterogeneous nuclear ribonucleoprotein A2/B1 | Nucleus | other | |
| chr9 | *LCN9* | 1 | 64 | 2 | LCN9 | lipocalin 9 | Extracellular Space | transporter | |
| chr4 | *RP11-1398P2.1* | 1 | 48 | 2 | | | | | |
| chr7 | *AC004854.4* | 1 | 71 | 2 | | | | | |
| chr10 | *FGF8* | 1 | 45 | 2 | FGF8 | fibroblast growth factor 8 | Extracellular Space | growth factor | |
| chr13 | *RNF6* | 1 | 23 | 2 | RNF6 | ring finger protein 6 | Nucleus | transcription regulator | |
| chr7 | *RP11-548K12.5* | 1 | 52 | 2 | | | | | |
| chr14 | *RDH11* | 1 | 27 | 2 | RDH11 | retinol dehydrogenase 11 | Cytoplasm | enzyme | |
| chr12 | *BCDIN3D* | 1 | 62 | 2 | BCDIN3D | BCDIN3 domain containing RNA methyltransferase | Cytoplasm | enzyme | |
| chr19 | *ZNF181* | 1 | 45 | 2 | ZNF181 | zinc finger protein 181 | Other | other | |
| chr1 | *BTG2* | 1 | 53 | 2 | BTG2 | BTG anti-proliferation factor 2 | Nucleus | transcription regulator | |
| chr22 | *TRMT2A* | 1 | 41 | 2 | TRMT2A | tRNA methyltransferase 2 | Other | kinase | |
| chr18 | *GRP* | 1 | 33 | 2 | GRP | gastrin releasing peptide | Extracellular Space | growth factor | |
| chr3 | *EOMES* | 1 | 47 | 2 | EOMES | eomesodermin | Nucleus | transcription regulator | |
| chr9 | *CDK2AP2P3* | 1 | 66 | 2 | CDK2AP2P3 | cyclin dependent kinase 2 associated protein 2 pseudogene | Other | other | |
| chr16 | *LA16c-444G7.2* | 1 | 48 | 2 | | | | | |
| chrX | *RP11-12D5.5* | 1 | 37 | 2 | | | | | |
| chr11 | *RP13-46H24.1* | 1 | 41 | 2 | | | | | |
| chr14 | *CKB* | 1 | 44 | 2 | CKB | creatine kinase B | Cytoplasm | kinase | |
| chr16 | *RP11-463O9.5* | 1 | 44 | 2 | | | | | |
| chrX | *HCCS* | 1 | 46 | 2 | HCCS | holocytochrome c synthase | Cytoplasm | enzyme | |
| chr19 | *C19orf43* | 1 | 41 | 2 | TRIR | telomerase RNA component interacting RNase | Nucleus | enzyme | |
| chr1 | *HPDL* | 1 | 50 | 2 | HPDL | 4-hydroxyphenylpyruvate dioxygenase like | Other | other | |
| chr1 | *HIST3H2BB* | 1 | 64 | 2 | H2BU1 | H2B.U histone 1 | Nucleus | other | |
| chr5 | *CTD-2013M15.1* | 1 | 129 | 2 | | | | | |
| chr1 | *LCE1B* | 1 | 42 | 2 | LCE1B | late cornified envelope 1B | Other | other | |

| | | | | | | | | | |
|---|---|---|---|---|---|---|---|---|---|
| chr2 | *IAH1* | 1 | 16 | 2 | IAH1 | isoamyl acetate hydrolyzing esterase 1 (putative) | Cytoplasm | other | |
| chrX | *FTH1P14* | 1 | 44 | 2 | FTH1P14 | ferritin heavy chain 1 pseudogene | Other | other | |
| chr9 | *ARID3C* | 1 | 31 | 2 | ARID3C | AT-rich interaction domain 3C | Nucleus | transcription regulator | |
| chr1 | *MANEAL* | 1 | 39 | 2 | MANEAL | mannosidase endo-alpha like | Other | other | |
| chr5 | *CTD-2233C11.3* | 1 | 107 | 2 | | | | | |
| chrX | *U82695.5* | 1 | 42 | 2 | | | | | |
| chr17 | *RP11-1109M24.1* | 1 | 39 | 2 | | | | | |
| chr17 | *PNMT* | 1 | 62 | 2 | PNMT | phenylethanolamine N-methyltransferase | Cytoplasm | enzyme | |
| chr5 | *EGR1* | 1 | 53 | 2 | EGR1 | early growth response 1 | Nucleus | transcription regulator | |
| chr17 | *AC144838.2* | 1 | 194 | 2 | | | | | |
| chr11 | *AP006285.7* | 1 | 40 | 2 | | | | | |
| chr1 | *RP11-7G12.2* | 1 | 47 | 2 | | | | | |
| chr9 | *RPL35* | 1 | 75 | 2 | RPL35 | ribosomal protein L35 | Cytoplasm | other | |
| chr5 | *CTD-2083E4.6* | 1 | 88 | 2 | | | | | |
| chr9 | *TMEM38B* | 1 | 39 | 2 | TMEM38B | transmembrane protein 38B | Nucleus | ion channel | |
| chr7 | *VGF* | 1 | 49 | 2 | VGF | VGF nerve growth factor | Extracellular Space | growth factor | |
| chr16 | *BOLA2B* | 1 | 60 | 2 | BOLA2/BOLA2B | bolA family member 2 | Cytoplasm | enzyme | |
| chr17 | *MIR3615* | 1 | 44 | 2 | mir-3615 | microRNA 3615 | Cytoplasm | microRNA | |
| chr6 | *TSPYL4* | 1 | 50 | 2 | TSPYL4 | TSPY like 4 | Nucleus | other | |
| chr12 | *RP11-123M21.1* | 1 | 41 | 2 | | | | | |
| chr1 | *ADAMTSL4* | 1 | 72 | 2 | ADAMTSL4 | ADAMTS like 4 | Extracellular Space | other | |
| chr2 | *HOXD-AS2* | 1 | 27 | 2 | HOXD-AS2 | HOXD cluster antisense RNA 2 | Other | other | |
| chr7 | *KMT2E-AS1* | 1 | 40 | 2 | KMT2E-AS1 | KMT2E antisense RNA 1 | Other | other | |
| chr15 | *NDN* | 1 | 53 | 2 | NDN | necdin, MAGE family member | Nucleus | transcription regulator | |
| chr12 | *P2RX2* | 1 | 52 | 2 | P2RX2 | purinergic receptor P2X 2 | Plasma Membrane | ion channel | |
| chr4 | *GUCY1A3* | 1 | 35 | 2 | GUCY1A1 | guanylate cyclase 1 soluble subunit alpha 1 | Cytoplasm | enzyme | nitroprusside, nitroglycerin, isosorbide dinitrate, isosorbide-5- |
| chrX | *LINC00629* | 1 | 42 | 2 | LINC00629 | long intergenic non-protein coding RNA 629 | Other | other | |
| chr14 | *KLHDC2* | 1 | 42 | 2 | KLHDC2 | kelch domain containing 2 | Cytoplasm | other | |
| chr17 | *KRTAP2-3* | 1 | 57 | 2 | RTAP2-3/KRTAP2 | keratin associated protein 2-3 | Cytoplasm | other | |
| chr10 | *AL031601.1* | 1 | 91 | 2 | | | | | |
| chr2 | *TMEM185B* | 1 | 74 | 2 | TMEM185B | transmembrane protein 185B | Other | other | |
| chrX | *SUPT20HL2* | 1 | 48 | 2 | SUPT20HL2 | SUPT20H like 2 | Extracellular Space | other | |
| chr6 | *LRRC73* | 1 | 39 | 2 | LRRC73 | leucine rich repeat containing 73 | Other | other | |
| chr8 | *FAM90A9P* | 1 | 113 | 2 | | | | | |
| chr10 | *PRLHR* | 1 | 55 | 2 | PRLHR | prolactin releasing hormone | Plasma Membrane | G-protein coupled recepto | |
| chr15 | *SPATA41* | 1 | 69 | 2 | SPATA41 | spermatogenesis associated 41 | Other | other | |
| chr22 | *GAS2L1* | 1 | 39 | 2 | GAS2L1 | growth arrest specific 2 like 1 | Cytoplasm | other | |
| chr11 | *AP000797.4* | 1 | 21 | 2 | | | | | |
| chr17 | *KCNAB3* | 1 | 49 | 2 | KCNAB3 | potassium voltage-gated channel subfamily A regulatory beta | Plasma Membrane | ion channel | |
| chr3 | *RBM15B* | 1 | 57 | 2 | RBM15B | RNA binding motif protein 15B | Nucleus | other | |
| chr9 | *RP11-350E12.5* | 1 | 50 | 2 | | | | | |
| chr20 | *RP11-112L6.4* | 1 | 56 | 2 | | | | | |
| chr18 | *CDH7* | 1 | 37 | 2 | CDH7 | cadherin 7 | Plasma Membrane | other | |
| chr12 | *RP11-357K6.1* | 1 | 37 | 2 | | | | | |
| chr4 | *SGMS2* | 1 | 32 | 2 | SGMS2 | sphingomyelin synthase 2 | Plasma Membrane | enzyme | |
| chr1 | *RP11-385M4.3* | 1 | 54 | 2 | | | | | |
| chr10 | *PTCHD3* | 1 | 32 | 2 | PTCHD3 | patched domain containing 3 | Cytoplasm | other | |
| chr5 | *RP11-432M8.10* | 1 | 128 | 2 | | | | | |
| chr19 | *CTD-3148I10.1* | 1 | 29 | 2 | | | | | |
| chr11 | *KRT8P26* | 1 | 92 | 2 | KRT8P26 | keratin 8 pseudogene 26 | Other | other | |
| chr11 | *CTD-2514A21.3* | 1 | 60 | 2 | | | | | |
| chr22 | *RP4-539M6.18* | 1 | 31 | 2 | | | | | |
| chr22 | *SELO* | 1 | 35 | 2 | | | | | |
| chr22 | *SHISA8* | 1 | 35 | 2 | SHISA8 | shisa family member 8 | Other | other | |
| chr12 | *COQ10A* | 1 | 50 | 2 | COQ10A | coenzyme Q10A | Cytoplasm | other | |
| chr6 | *SAYSD1* | 1 | 21 | 2 | SAYSD1 | SAYSVFN motif domain | Other | other | |
| chr17 | *DND1P1* | 1 | 88 | 2 | DND1P1 | DND microRNA-mediated repression inhibitor 1 pseudogene | Other | other | |
| chr9 | *DKFZP434A062* | 1 | 84 | 2 | DKFZP434A062 | uncharacterized LOC26102 | Other | other | |
| chr9 | *AL590708.2* | 1 | 73 | 2 | | | | | |
| chr13 | *LINC01053* | 1 | 20 | 2 | LINC01053 | long intergenic non-protein coding RNA 1053 | Other | other | |
| chr20 | *RP11-410N8.1* | 1 | 62 | 2 | | | | | |
| chr11 | *PHLDA2* | 1 | 100 | 2 | PHLDA2 | pleckstrin homology like domain family A member 2 | Cytoplasm | other | |
| chr17 | *KRTAP17-1* | 1 | 42 | 2 | KRTAP17-1 | keratin associated protein 17-1 | Cytoplasm | other | |
| chr11 | *RASSF7* | 1 | 38 | 2 | RASSF7 | Ras association domain family member 7 | Cytoplasm | other | |
| chr6 | *NR2E1* | 1 | 29 | 2 | NR2E1 | nuclear receptor subfamily 2 group E member 1 | Nucleus | d-dependent nuclear rec | |
| chr14 | *ZC2HC1C* | 1 | 34 | 2 | ZC2HC1C | zinc finger C2HC-type containing | Other | other | |
| chr1 | *RP11-316M1.3* | 1 | 54 | 2 | | | | | |
| chrX | *VENTXP1* | 1 | 37 | 2 | VENTXP1 | VENT homeobox pseudogene 1 | Other | other | |
| chr15 | *RP11-2F9.3* | 1 | 61 | 2 | | | | | |
| chr20 | *LINC00659* | 1 | 91 | 2 | LINC00659 | long intergenic non-protein coding RNA 659 | Other | other | |
| chr14 | *EIF5* | 1 | 38 | 2 | EIF5 | eukaryotic translation initiation | Cytoplasm | translation regulator | |
| chr14 | *RP11-536C10.4* | 1 | 37 | 2 | | | | | |
| chr17 | *DDX5* | 1 | 43 | 2 | DDX5 | DEAD-box helicase 5 | Nucleus | enzyme | |
| chr15 | *NR2F2* | 1 | 85 | 2 | NR2F2 | nuclear receptor subfamily 2 group F member 2 | Nucleus | d-dependent nuclear rec | |
| chr7 | *WNT16* | 1 | 43 | 2 | WNT16 | Wnt family member 16 | Extracellular Space | other | |

| | | | | | | | | | |
|---|---|---|---|---|---|---|---|---|---|
| chr11 | *RP11-676F20.2* | 1 | 79 | 2 | | | | | |
| chr13 | *PPP1R2P10* | 1 | 45 | 2 | PPP1R2P10 | protein phosphatase 1 regulatory inhibitor subunit 2 pseudogene 10 | Other | other | |
| chr16 | *PDIA2* | 1 | 53 | 2 | PDIA2 | protein disulfide isomerase family A member 2 | Cytoplasm | enzyme | |
| chrX | *GS1-421I3.2* | 1 | 32 | 2 | | | | | |
| chr16 | *YBX3P1* | 1 | 40 | 2 | YBX3P1 | Y-box binding protein 3 pseudogene 1 | Other | other | |
| chr9 | *RPL7A* | 1 | 70 | 2 | RPL7A | ribosomal protein L7a | Cytoplasm | other | |
| chr7 | *NAA38* | 1 | 39 | 2 | NAA38 | N(alpha)-acetyltransferase 38, NatC auxiliary subunit | Nucleus | other | |
| chr13 | *PRR20D* | 1 | 118 | 2 | R20A (includes oth | proline rich 20A | Other | other | |
| chr7 | *RP4-647J21.1* | 1 | 47 | 2 | | | | | |
| chr20 | *LINC00028* | 1 | 67 | 2 | LINC00028 | long intergenic non-protein coding RNA 28 | Other | other | |
| chr4 | *POU4F2* | 1 | 43 | 2 | POU4F2 | POU class 4 homeobox 2 | Nucleus | transcription regulator | |
| chr17 | *CHRNE* | 1 | 46 | 2 | CHRNE | cholinergic receptor nicotinic epsilon subunit | Plasma Membrane | transmembrane recepto | succinylcholine, rapacuronium, doxacurium, vecuronium, pipecuronium, metocurine, mivacurium, D-tubocurarine, nicotine, arecoline, pancuronium, atracurium, isoflurane, cisatracurium, rocuronium, amobarbital, mecamylamine |
| chr11 | *DRAP1* | 1 | 72 | 2 | DRAP1 | DR1 associated protein 1 | Nucleus | transcription regulator | |
| chr18 | *RP11-243E13.1* | 1 | 41 | 2 | | | | | |
| chr11 | *CEND1* | 1 | 39 | 2 | CEND1 | cell cycle exit and neuronal differentiation 1 | Other | other | |
| chr1 | *RP11-68I18.10* | 1 | 56 | 2 | | | | | |
| chr11 | *MRGPRX1* | 1 | 54 | 2 | MRGPRX1 | MAS related GPR family member | Plasma Membrane | G-protein coupled recepto | |
| chr19 | *B3GNT8* | 1 | 47 | 2 | B3GNT8 | UDP-GlcNAc:betaGal beta-1,3-N-acetylglucosaminyltransferase 8 | Cytoplasm | enzyme | |
| chr9 | *CTSL* | 1 | 45 | 2 | CTSL | cathepsin L | Cytoplasm | peptidase | cathepsin L inhibitor |
| chr1 | *RP11-438F14.3* | 1 | 39 | 2 | | | | | |
| chr1 | *C1orf65* | 1 | 70 | 2 | CCDC185 | coiled-coil domain containing 185 | Other | other | |
| chr17 | *RN7SL605P* | 1 | 33 | 2 | RN7SL605P | | Other | other | |
| chr13 | *CCR12P* | 1 | 63 | 2 | CCR12P | C-C motif chemokine receptor 12, pseudogene | Other | other | |
| chr8 | *RP11-481A20.11* | 1 | 47 | 2 | | | | | |
| chr11 | *SSSCA1-AS1* | 1 | 87 | 2 | SSSCA1-AS1 | SSSCA1 antisense RNA 1 (head to head) | Other | other | |
| chr21 | *AP001057.1* | 1 | 41 | 2 | | | | | |
| chr5 | *TSLP* | 1 | 42 | 2 | TSLP | thymic stromal lymphopoietin | Extracellular Space | cytokine | tezepelumab |
| chr10 | *LYZL1* | 1 | 29 | 2 | LYZL1 | lysozyme like 1 | Other | enzyme | |
| chr1 | *RP11-206L10.9* | 1 | 27 | 2 | | | | | |
| chr3 | *ZNF501* | 1 | 45 | 2 | ZNF501 | zinc finger protein 501 | Nucleus | transcription regulator | |
| chr2 | *DDX18* | 1 | 50 | 2 | DDX18 | DEAD-box helicase 18 | Nucleus | enzyme | |
| chr1 | *RP11-438H8.8* | 1 | 42 | 2 | | | | | |
| chr1 | *B3GALT6* | 1 | 59 | 2 | B3GALT6 | beta-1,3-galactosyltransferase 6 | Cytoplasm | enzyme | |
| chr19 | *CTD-2368P22.1* | 1 | 39 | 2 | | | | | |
| chr5 | *CTD-2244C20.1* | 1 | 29 | 2 | | | | | |
| chr13 | *LINC00567* | 1 | 60 | 2 | LINC00567 | long intergenic non-protein coding RNA 567 | Other | other | |
| chr8 | *SLC52A2* | 1 | 62 | 2 | SLC52A2 | solute carrier family 52 member 2 | Plasma Membrane | transporter | 4-hydroxybutanoic |
| chr12 | *GAPDH* | 1 | 52 | 2 | GAPDH | glyceraldehyde-3-phosphate dehydrogenase | Cytoplasm | enzyme | |
| chr1 | *HIST2H3A* | 1 | 71 | 2 | H3C15 | H3 clustered histone 15 | Nucleus | other | |
| chr19 | *AC010641.1* | 1 | 31 | 2 | | | | | |
| chr9 | *CDK2AP2P2* | 1 | 129 | 2 | CDK2AP2P2 | | Other | other | |
| chr7 | *ZDHHC4* | 1 | 55 | 2 | ZDHHC4 | zinc finger DHHC-type containing | Cytoplasm | other | |
| chr19 | *CTC-457E21.6* | 1 | 27 | 2 | | | | | |
| chr9 | *TMEM203* | 1 | 84 | 2 | TMEM203 | transmembrane protein 203 | Cytoplasm | other | |
| chr5 | *RPS23* | 1 | 39 | 2 | RPS23 | ribosomal protein S23 | Cytoplasm | translation regulator | |
| chr12 | *TBX5-AS1* | 1 | 45 | 2 | TBX5-AS1 | TBX5 antisense RNA 1 | Other | other | |
| chr4 | *FGFR3* | 1 | 41 | 2 | FGFR3 | fibroblast growth factor receptor 3 | Plasma Membrane | kinase | ASP5878, lapatinib/pazopanib, PRN1371, lenvatinib, 3D185, erdafitinib, debio 1347, lenvatinib/pembrolizumab, pazopanib, crizotinib/pazopanib, docetaxel/nintedanib, masitinib, everolimus/lenvatinib, SAR103168, LY3076226, pemigatinib, nintedanib, B-701 |
| chr20 | *ID1* | 1 | 107 | 2 | ID1 | inhibitor of DNA binding 1, HLH protein | Nucleus | transcription regulator | |

| | | | | | | | | | |
|---|---|---|---|---|---|---|---|---|---|
| chr1 | *RP4-597A16.3* | 1 | 36 | 2 | | | | | |
| chr10 | *RP11-432J24.2* | 1 | 47 | 2 | | | | | |
| chr11 | *CDK2AP2* | 1 | 48 | 2 | CDK2AP2 | cyclin dependent kinase 2 associated protein 2 | Nucleus | other | |
| chr11 | *AC109309.4* | 1 | 92 | 2 | | | | | |
| chr10 | *RP11-432J24.6* | 1 | 76 | 2 | | | | | |
| chr8 | *SOX7* | 1 | 47 | 2 | SOX7 | SRY-box transcription factor 7 | Nucleus | transcription regulator | |
| chr1 | *TMEM200B* | 1 | 47 | 2 | TMEM200B | transmembrane protein 200B | Other | other | |
| chr11 | *RP11-713P17.5* | 1 | 39 | 2 | | | | | |
| chr1 | *LINC00083* | 1 | 65 | 2 | CLEC20A | C-type lectin domain containing | Other | other | |
| chr11 | *HRAS* | 1 | 41 | 2 | HRAS | HRas proto-oncogene, GTPase | Plasma Membrane | enzyme | ISIS 2503 |
| chr19 | *AURKC* | 1 | 43 | 2 | AURKC | aurora kinase C | Nucleus | kinase | AMG 900, danusertib, GSK1070916, ilorasertib, SKI-G- |
| chr16 | *RP11-23E10.2* | 1 | 108 | 2 | | | | | |
| chr8 | *REXO1L3P* | 1 | 185 | 2 | REXO1L3P | REXO1 like 3, pseudogene | Other | other | |
| chr3 | *RP11-16N11.2* | 1 | 61 | 2 | | | | | |
| chr5 | *MRPS30* | 1 | 134 | 2 | MRPS30 | mitochondrial ribosomal protein | Cytoplasm | enzyme | |
| chr11 | *TBRG1* | 1 | 41 | 2 | TBRG1 | transforming growth factor beta regulator 1 | Nucleus | other | |
| chr17 | *RPL13P12* | 1 | 51 | 2 | RPL13P12 | ribosomal protein L13 pseudogene 12 | Other | other | |
| chr6 | *ZSCAN12P1* | 1 | 37 | 2 | ZSCAN12P1 | zinc finger and SCAN domain containing 12 pseudogene 1 | Other | other | |
| chr4 | *NAT8L* | 1 | 40 | 2 | NAT8L | N-acetyltransferase 8 like | Cytoplasm | enzyme | |
| chr15 | *RP11-100M12.3* | 1 | 45 | 2 | | | | | |
| chr14 | *CRIP2* | 1 | 43 | 2 | CRIP2 | cysteine rich protein 2 | Nucleus | other | |
| chr5 | *MTHFD2P6* | 1 | 73 | 2 | MTHFD2P6 | methylenetetrahydrofolate dehydrogenase (NADP+ dependent) 2, methenyltetrahydrofolate | Other | other | |
| chr1 | *TMEM125* | 1 | 56 | 2 | TMEM125 | transmembrane protein 125 | Other | other | |
| chr17 | *RAMP2* | 1 | 51 | 2 | RAMP2 | receptor activity modifying protein | Plasma Membrane | other | pramlintide |
| chr3 | *RP11-281P11.1* | 1 | 48 | 2 | | | | | |
| chr1 | *RP11-334L9.1* | 1 | 45 | 2 | | | | | |
| chr16 | *PARD6A* | 1 | 60 | 2 | PARD6A | par-6 family cell polarity regulator alpha | Plasma Membrane | other | |
| chr3 | *RP11-2A4.3* | 1 | 49 | 2 | | | | | |
| chr9 | *RP11-229P13.19* | 1 | 81 | 2 | | | | | |
| chr7 | *AC093627.11* | 1 | 73 | 2 | | | | | |
| chr8 | *REXO1L2P* | 1 | 216 | 2 | REXO1L2P | REXO1 like 2, pseudogene | Other | other | |
| chrX | *SPIN2B* | 1 | 47 | 2 | SPIN2A/SPIN2B | spindlin family member 2A | Other | other | |
| chr17 | *NEK4P2* | 1 | 61 | 2 | NEK4P2 | NIMA-related kinase 4 | Other | other | |
| chr8 | *EGR3* | 1 | 55 | 2 | EGR3 | early growth response 3 | Nucleus | transcription regulator | |
| chr6 | *RP3-431A14.4* | 1 | 64 | 2 | | | | | |

| Supplementary Table 6. Prior gene targets for cancer treatment identified in the literature and the genes damaged by cisplatin mapped in the present work. | | | | |
|---|---|---|---|---|
| Remarks:<br>Gene name in green represents this gene being the target of approved anticancer drugs in clinic or candidates in preclinical development;<br>Gene name in orange represents this gene being a poteintal target without drugs in clinical development but with evidence that support target tractability ;<br>Gene name in blank represents this gene being a poteintial target with no support or lack of information that could inform tractability. | | | | |
| Chromosome | Gene | Average number of identified peaks (n=3) | Average number of tags (n=3) | Average enrichment fold (n=3) |
| chr17 | *RPTOR* | 147 | 6474 | 351 |
| chr17 | *SMG6* | 73 | 2206 | 179 |
| chr3 | *STAG1* | 68 | 1921 | 173 |
| chr17 | *TBCD* | 75 | 3672 | 163 |
| chr15 | *IGF1R* | 68 | 3199 | 158 |
| chr17 | *JUP* | 55 | 2174 | 128 |
| chr6 | *FARS2* | 52 | 1445 | 125 |
| chr7 | *EGFR* | 52 | 2523 | 122 |
| chr8 | *PTK2* | 50 | 1337 | 122 |
| chr1 | *CAPZB* | 50 | 1742 | 117 |
| chr3 | *MRPS22* | 48 | 1947 | 111 |
| chr12 | *EP400* | 45 | 1893 | 107 |
| chr16 | *CREBBP* | 42 | 1326 | 106 |
| chr3 | *PIK3CB* | 39 | 1292 | 101 |
| chr7 | *TRRAP* | 41 | 1979 | 94 |
| chr17 | *KANSL1* | 34 | 977 | 94 |
| chr11 | *NUP98* | 37 | 1721 | 93 |
| chr3 | *ITGB5* | 38 | 1552 | 87 |
| chr3 | *WWTR1* | 35 | 1164 | 86 |
| chr17 | *VMP1* | 31 | 855 | 84 |
| chr13 | *ARHGEF7* | 33 | 1278 | 82 |
| chr22 | *MYH9* | 33 | 1390 | 82 |
| chr12 | *SFSWAP* | 30 | 1258 | 77 |
| chr19 | *DNM2* | 29 | 1155 | 77 |
| chr7 | *OGDH* | 34 | 1330 | 77 |
| chr17 | *CDK12* | 31 | 841 | 76 |
| chr4 | *RNF4* | 26 | 830 | 72 |
| chr1 | *MTOR* | 28 | 747 | 66 |
| chr16 | *GTF3C1* | 28 | 966 | 65 |
| chr10 | *BMPR1A* | 26 | 625 | 64 |
| chr17 | *METTL16* | 25 | 693 | 63 |
| chr11 | *RAB6A* | 25 | 796 | 62 |
| chr15 | *SIN3A* | 24 | 722 | 61 |
| chr10 | *FGFR2* | 26 | 769 | 61 |
| chr12 | *SBNO1* | 24 | 665 | 59 |
| chr16 | *PDPK1* | 26 | 1308 | 58 |
| chr12 | *ARID2* | 23 | 782 | 58 |
| chr17 | *GRB2* | 22 | 674 | 57 |
| chr9 | *TRAF2* | 24 | 1570 | 56 |
| chr15 | *INO80* | 22 | 570 | 56 |
| chr3 | *SCAP* | 23 | 718 | 55 |
| chr6 | *LYRM4* | 25 | 702 | 55 |
| chr17 | *ACLY* | 23 | 747 | 54 |
| chr11 | *FLI1* | 25 | 908 | 54 |
| chr13 | *TFDP1* | 25 | 1334 | 54 |
| chr17 | *STAT3* | 22 | 580 | 53 |
| chr3 | *RHOA* | 22 | 681 | 53 |
| chr1 | *LRP8* | 22 | 909 | 52 |
| chr5 | *ATP6V0E1* | 22 | 1183 | 52 |
| chr10 | *RRP12* | 22 | 935 | 52 |
| chr12 | *KNTC1* | 20 | 517 | 52 |
| chr3 | *FANCD2* | 20 | 512 | 52 |
| chr11 | *YAP1* | 20 | 992 | 51 |
| chr5 | *FBXW11* | 19 | 611 | 51 |
| chr11 | *HSD17B12* | 21 | 545 | 51 |
| chr19 | *DNMT1* | 19 | 681 | 50 |
| chr20 | *MYBL2* | 22 | 920 | 49 |

| | | | | |
|---|---|---|---|---|
| chr1 | *ZBTB8OS* | 21 | 605 | 49 |
| chr3 | *RPN1* | 20 | 756 | 48 |
| chr11 | *IPO7* | 19 | 740 | 48 |
| chr16 | *CDH1* | 21 | 550 | 45 |
| chr10 | *PDCD11* | 19 | 537 | 45 |
| chr1 | *DNAJC11* | 19 | 778 | 45 |
| chr1 | *INTS3* | 18 | 779 | 45 |
| chr17 | *SUZ12* | 18 | 472 | 44 |
| chr19 | *SAE1* | 16 | 476 | 43 |
| chr3 | *XRN1* | 17 | 524 | 43 |
| chr1 | *NUDC* | 18 | 567 | 43 |
| chr16 | *MBTPS1* | 18 | 827 | 43 |
| chr15 | *TRPM7* | 15 | 357 | 43 |
| chr1 | *UBR4* | 17 | 496 | 42 |
| chr19 | *SUGP1* | 20 | 746 | 42 |
| chr1 | *HDAC1* | 18 | 500 | 42 |
| chr16 | *AARS* | 17 | 599 | 42 |
| chr18 | *TGIF1* | 17 | 801 | 42 |
| chrX | *HUWE1* | 16 | 586 | 41 |
| chr16 | *NUP93* | 18 | 423 | 40 |
| chr3 | *ATR* | 15 | 474 | 40 |
| chr17 | *TRIM37* | 15 | 385 | 40 |
| chr20 | *TPX2* | 17 | 721 | 40 |
| chr7 | *ACTB* | 17 | 1068 | 40 |
| chr16 | *USP7* | 17 | 507 | 40 |
| chr7 | *CDK6* | 17 | 539 | 39 |
| chr12 | *PTPN11* | 17 | 386 | 38 |
| chr12 | *PPP1R12A* | 15 | 336 | 38 |
| chr2 | *PNPT1* | 15 | 507 | 37 |
| chr16 | *NDE1* | 17 | 461 | 37 |
| chr6 | *NUP153* | 14 | 366 | 37 |
| chr15 | *ARIH1* | 14 | 348 | 37 |
| chr1 | *MFN2* | 15 | 568 | 37 |
| chr22 | *SMARCB1* | 13 | 488 | 36 |
| chr5 | *TBCA* | 15 | 408 | 36 |
| chr1 | *MDM4* | 16 | 510 | 36 |
| chr22 | *GNB1L* | 15 | 657 | 36 |
| chr17 | *MED24* | 15 | 600 | 34 |
| chr15 | *CASC5* | 13 | 377 | 34 |
| chr3 | *COPG1* | 15 | 788 | 33 |
| chr16 | *SLC7A5* | 15 | 748 | 33 |
| chr2 | *XPO1* | 13 | 378 | 33 |
| chrX | *STAG2* | 12 | 316 | 33 |
| chr12 | *KMT2D* | 17 | 1141 | 33 |
| chr12 | *NCAPD2* | 12 | 415 | 33 |
| chr17 | *NUP85* | 13 | 442 | 32 |
| chr13 | *SLC7A1* | 11 | 410 | 32 |
| chr22 | *UBE2L3* | 13 | 427 | 32 |
| chr8 | *WRN* | 11 | 292 | 32 |
| chr16 | *POLR3E* | 14 | 568 | 31 |
| chr1 | *CMPK1* | 13 | 345 | 31 |
| chr6 | *PSMG4* | 13 | 473 | 31 |
| chr17 | *RAD51C* | 12 | 336 | 31 |
| chrX | *SMC1A* | 14 | 419 | 31 |
| chr19 | *SH3GL1* | 14 | 623 | 31 |
| chr18 | *ZNF407* | 11 | 273 | 30 |
| chr1 | *TRNAU1AP* | 13 | 354 | 30 |
| chr22 | *ACO2* | 12 | 417 | 30 |
| chr17 | *ERBB2* | 12 | 535 | 30 |
| chr4 | *WDR1* | 11 | 479 | 30 |
| chr8 | *FGFR1* | 11 | 406 | 30 |
| chr4 | *PDS5A* | 10 | 242 | 30 |
| chr11 | *POU2AF1* | 13 | 393 | 29 |
| chr1 | *NDC1* | 11 | 297 | 29 |
| chr19 | *LIG1* | 11 | 389 | 29 |
| chr18 | *BCL2* | 10 | 295 | 29 |
| chr2 | *RANBP2* | 11 | 381 | 29 |

| | | | | |
|---|---|---|---|---|
| chr20 | *CHMP4B* | 13 | 486 | 28 |
| chr3 | *PAK2* | 9 | 255 | 28 |
| chr7 | *RAC1* | 12 | 400 | 28 |
| chr3 | *GMPS* | 11 | 276 | 28 |
| chr11 | *DDX6* | 11 | 291 | 28 |
| chr8 | *EXT1* | 11 | 236 | 27 |
| chr9 | *HAUS6* | 12 | 307 | 27 |
| chr20 | *DDX27* | 11 | 507 | 27 |
| chr6 | *ESR1* | 11 | 273 | 27 |
| chr3 | *MBNL1* | 11 | 293 | 27 |
| chr16 | *CYB5B* | 9 | 299 | 27 |
| chr1 | *SLC2A1* | 12 | 446 | 27 |
| chr17 | *PFAS* | 11 | 516 | 26 |
| chr1 | *ADAR* | 11 | 420 | 26 |
| chr12 | *VEZT* | 11 | 278 | 26 |
| chr2 | *KANSL3* | 12 | 444 | 26 |
| chr18 | *ZNF236* | 9 | 301 | 26 |
| chr1 | *SMG5* | 11 | 483 | 25 |
| chr3 | *SLC4A7* | 10 | 285 | 25 |
| chr9 | *GTF3C5* | 11 | 609 | 25 |
| chr17 | *PPM1D* | 10 | 250 | 25 |
| chr10 | *ACTR1A* | 10 | 391 | 25 |
| chr17 | *NUP88* | 9 | 247 | 24 |
| chr9 | *SMC5* | 10 | 224 | 24 |
| chr1 | *SLC16A1* | 10 | 306 | 24 |
| chr9 | *SMARCA2* | 10 | 270 | 24 |
| chr17 | *ARMC7* | 10 | 429 | 24 |
| chr2 | *GEMIN6* | 10 | 316 | 24 |
| chr7 | *RFC2* | 10 | 334 | 23 |
| chr1 | *CELSR2* | 12 | 650 | 23 |
| chr6 | *SPDEF* | 10 | 504 | 23 |
| chr1 | *URB2* | 10 | 382 | 23 |
| chrX | *TAF1* | 8 | 217 | 23 |
| chr15 | *PKM* | 10 | 305 | 23 |
| chr19 | *RPL28* | 10 | 376 | 23 |
| chr6 | *E2F3* | 9 | 230 | 23 |
| chr1 | *KIF14* | 9 | 232 | 23 |
| chr21 | *UBE2G2* | 7 | 229 | 23 |
| chr19 | *ILF3* | 9 | 358 | 23 |
| chr15 | *FANCI* | 10 | 298 | 23 |
| chr2 | *TAF1B* | 8 | 244 | 22 |
| chr18 | *SMAD7* | 8 | 343 | 22 |
| chr17 | *UTP6* | 9 | 210 | 22 |
| chr2 | *ACTR3* | 8 | 273 | 22 |
| chr15 | *TUBGCP5* | 10 | 281 | 22 |
| chr3 | *PDCD6IP* | 8 | 208 | 22 |
| chr11 | *CDCA5* | 9 | 352 | 22 |
| chr10 | *SHOC2* | 8 | 251 | 22 |
| chr17 | *CLTC* | 8 | 240 | 22 |
| chr11 | *STX5* | 10 | 261 | 22 |
| chr7 | *RHEB* | 8 | 261 | 21 |
| chr15 | *SNAP23* | 8 | 228 | 21 |
| chr16 | *IRF8* | 9 | 351 | 21 |
| chr18 | *CEP192* | 8 | 228 | 21 |
| chr3 | *TAMM41* | 8 | 220 | 21 |
| chr7 | *MET* | 10 | 232 | 21 |
| chr1 | *PTPN7* | 9 | 512 | 21 |
| chr17 | *CPD* | 8 | 223 | 20 |
| chr5 | *APC* | 7 | 183 | 20 |
| chr20 | *RBM39* | 7 | 313 | 20 |
| chr20 | *BCL2L1* | 9 | 500 | 20 |
| chr3 | *PTPN23* | 9 | 272 | 19 |
| chr7 | *RBM28* | 8 | 211 | 19 |
| chr1 | *RBBP4* | 7 | 226 | 19 |
| chr8 | *CHMP7* | 8 | 320 | 19 |
| chr2 | *RAB10* | 7 | 173 | 19 |
| chr16 | *ALDOA* | 8 | 341 | 19 |

| | | | | |
|---|---|---|---|---|
| chr5 | *VDAC1* | 8 | 271 | 19 |
| chrX | *DDX3X* | 8 | 239 | 18 |
| chr1 | *UBE2J2* | 9 | 379 | 18 |
| chr8 | *GRHL2* | 7 | 191 | 18 |
| chr17 | *MYBBP1A* | 9 | 308 | 18 |
| chr1 | *CTPS1* | 9 | 258 | 18 |
| chr1 | *UBIAD1* | 9 | 246 | 18 |
| chr22 | *TUBGCP6* | 8 | 430 | 18 |
| chr3 | *ARF4* | 7 | 223 | 18 |
| chr1 | *SDHB* | 8 | 219 | 18 |
| chr19 | *CNOT3* | 8 | 414 | 18 |
| chr15 | *IDH3A* | 9 | 257 | 18 |
| chr16 | *DHX38* | 8 | 317 | 18 |
| chr1 | *PGD* | 8 | 211 | 18 |
| chr15 | *SNUPN* | 8 | 332 | 18 |
| chr10 | *MMS19* | 8 | 215 | 18 |
| chr3 | *UQCRC1* | 8 | 337 | 17 |
| chr9 | *FPGS* | 8 | 466 | 17 |
| chr16 | *ALG1* | 7 | 342 | 17 |
| chr8 | *UTP23* | 8 | 193 | 17 |
| chr20 | *CDS2* | 6 | 169 | 17 |
| chr11 | *SCYL1* | 8 | 501 | 17 |
| chr2 | *SLC39A10* | 6 | 161 | 17 |
| chr12 | *ERBB3* | 8 | 254 | 17 |
| chr1 | *RBBP5* | 7 | 193 | 17 |
| chr1 | *PPP1R8* | 7 | 201 | 17 |
| chr2 | *CUL3* | 5 | 134 | 17 |
| chr1 | *TIMM17A* | 6 | 267 | 16 |
| chr12 | *UBE2N* | 6 | 158 | 16 |
| chr5 | *IPO11* | 7 | 143 | 16 |
| chr17 | *SREBF1* | 7 | 464 | 16 |
| chr20 | *CSNK2A1* | 5 | 135 | 16 |
| chr7 | *RINT1* | 7 | 198 | 16 |
| chr1 | *PRCC* | 7 | 277 | 16 |
| chr19 | *ERCC2* | 7 | 275 | 16 |
| chr15 | *IREB2* | 7 | 247 | 16 |
| chr15 | *BUB1B* | 5 | 173 | 16 |
| chr12 | *MVK* | 6 | 358 | 15 |
| chr18 | *RBBP8* | 5 | 126 | 15 |
| chr14 | *TMED10* | 5 | 157 | 15 |
| chr8 | *PPP2R2A* | 6 | 170 | 15 |
| chr18 | *RTTN* | 5 | 168 | 15 |
| chr1 | *STXBP3* | 7 | 155 | 15 |
| chr13 | *IRS2* | 7 | 174 | 14 |
| chr17 | *CHMP6* | 7 | 348 | 14 |
| chr2 | *CAD* | 6 | 245 | 14 |
| chr1 | *ARF1* | 6 | 320 | 14 |
| chr22 | *USP18* | 7 | 160 | 14 |
| chr10 | *CEP55* | 6 | 158 | 14 |
| chr12 | *KRAS* | 5 | 172 | 14 |
| chr16 | *NOB1* | 6 | 216 | 14 |
| chr6 | *IRF4* | 6 | 179 | 13 |
| chr12 | *METAP2* | 5 | 156 | 13 |
| chr2 | *STAMBP* | 5 | 138 | 13 |
| chr1 | *IPO13* | 6 | 257 | 13 |
| chr20 | *RAE1* | 6 | 329 | 13 |
| chr19 | *CDC37* | 6 | 222 | 13 |
| chr17 | *PELP1* | 6 | 197 | 13 |
| chr3 | *TFRC* | 4 | 124 | 13 |
| chr7 | *SSBP1* | 5 | 160 | 13 |
| chr1 | *CDC42* | 6 | 131 | 13 |
| chr15 | *EMC7* | 4 | 150 | 13 |
| chr17 | *HGS* | 7 | 365 | 13 |
| chr9 | *NELFB* | 7 | 305 | 12 |
| chr15 | *OIP5* | 5 | 160 | 12 |
| chrX | *LAS1L* | 5 | 217 | 12 |
| chr1 | *TTF2* | 5 | 149 | 12 |

| | | | | |
|---|---|---|---|---|
| chr4 | *TACC3* | 5 | 252 | 12 |
| chr20 | *E2F1* | 6 | 341 | 12 |
| chr17 | *UTP18* | 4 | 129 | 12 |
| chr3 | *CCDC174* | 5 | 155 | 12 |
| chr19 | *PPP2R1A* | 6 | 144 | 12 |
| chr6 | *LEMD2* | 5 | 235 | 12 |
| chr17 | *FDXR* | 6 | 260 | 12 |
| chr16 | *MVD* | 6 | 271 | 12 |
| chr6 | *SOD2* | 4 | 128 | 12 |
| chr2 | *ITGAV* | 5 | 93 | 12 |
| chr19 | *FZR1* | 6 | 277 | 12 |
| chr20 | *CDC25B* | 4 | 213 | 12 |
| chr12 | *CCND2* | 4 | 133 | 12 |
| chr1 | *ENO1* | 5 | 172 | 12 |
| chr9 | *COQ4* | 6 | 240 | 11 |
| chr10 | *LDB1* | 6 | 226 | 11 |
| chr18 | *WDR7* | 4 | 116 | 11 |
| chr11 | *KIF18A* | 4 | 127 | 11 |
| chr1 | *SPRTN* | 4 | 168 | 11 |
| chr1 | *UBE2T* | 5 | 186 | 11 |
| chr5 | *HNRNPH1* | 5 | 220 | 11 |
| chr19 | *LONP1* | 6 | 321 | 11 |
| chr8 | *CPSF1* | 5 | 188 | 11 |
| chr19 | *HAUS5* | 5 | 228 | 11 |
| chr1 | *MAD2L2* | 5 | 186 | 11 |
| chr2 | *HJURP* | 4 | 153 | 11 |
| chr2 | *NFE2L2* | 4 | 87 | 11 |
| chr16 | *STX4* | 5 | 205 | 11 |
| chr12 | *USP5* | 4 | 181 | 10 |
| chr1 | *TAF13* | 5 | 148 | 10 |
| chr9 | *MED22* | 5 | 318 | 10 |
| chr10 | *NHLRC2* | 4 | 116 | 10 |
| chr2 | *SF3B1* | 4 | 97 | 10 |
| chr15 | *FURIN* | 6 | 459 | 10 |
| chr8 | *EFR3A* | 4 | 103 | 10 |
| chr16 | *DDX19A* | 5 | 166 | 10 |
| chr5 | *CSNK1A1* | 4 | 105 | 10 |
| chr19 | *DDX39A* | 5 | 210 | 10 |
| chr3 | *CTNNB1* | 4 | 91 | 10 |
| chr15 | *WDR73* | 5 | 151 | 10 |
| chr3 | *VHL* | 5 | 135 | 10 |
| chr5 | *DCP2* | 4 | 117 | 10 |
| chr1 | *RNPC3* | 4 | 117 | 10 |
| chr1 | *GGPS1* | 4 | 163 | 10 |
| chr1 | *ASPM* | 4 | 142 | 10 |
| chr12 | *ATP5B* | 4 | 147 | 10 |
| chr16 | *SLC7A6OS* | 4 | 139 | 10 |
| chr5 | *SKP1* | 3 | 98 | 10 |
| chr16 | *UQCRC2* | 3 | 113 | 10 |
| chr14 | *MTHFD1* | 3 | 94 | 9 |
| chr14 | *ACIN1* | 3 | 113 | 9 |
| chr7 | *MIOS* | 4 | 96 | 9 |
| chr17 | *ELAC2* | 5 | 226 | 9 |
| chr11 | *FOSL1* | 5 | 262 | 9 |
| chr11 | *CCND1* | 5 | 230 | 9 |
| chr19 | *ASNA1* | 4 | 134 | 9 |
| chr2 | *CENPA* | 4 | 91 | 9 |
| chr1 | *SHC1* | 4 | 197 | 9 |
| chr14 | *GTF2A1* | 3 | 87 | 9 |
| chr11 | *SYVN1* | 4 | 215 | 9 |
| chr12 | *PRIM1* | 4 | 92 | 8 |
| chr2 | *NCL* | 4 | 162 | 8 |
| chr2 | *ACSL3* | 4 | 68 | 8 |
| chr3 | *DHX36* | 4 | 114 | 8 |
| chr15 | *EIF3J* | 3 | 83 | 8 |
| chr20 | *ARFRP1* | 4 | 246 | 8 |
| chr10 | *PSTK* | 4 | 86 | 8 |

| | | | | |
|---|---|---|---|---|
| chr3 | *SEC61A1* | 4 | 122 | 8 |
| chr20 | *FERMT1* | 3 | 82 | 8 |
| chr22 | *HSCB* | 3 | 65 | 8 |
| chr13 | *KLF5* | 4 | 102 | 8 |
| chr16 | *TAF1C* | 4 | 212 | 8 |
| chr14 | *WDHD1* | 3 | 67 | 8 |
| chr3 | *WDR48* | 3 | 80 | 8 |
| chr7 | *NAMPT* | 3 | 61 | 8 |
| chr1 | *UAP1* | 4 | 96 | 8 |
| chr2 | *NCKAP1* | 3 | 72 | 8 |
| chr1 | *MAGOH* | 3 | 72 | 7 |
| chr1 | *NDUFS2* | 4 | 141 | 7 |
| chr17 | *GEMIN4* | 3 | 190 | 7 |
| chr1 | *AHCYL1* | 3 | 102 | 7 |
| chr2 | *GTF3C2* | 3 | 83 | 7 |
| chr3 | *UMPS* | 3 | 122 | 7 |
| chr1 | *TPR* | 3 | 72 | 7 |
| chr19 | *POLRMT* | 4 | 114 | 7 |
| chr17 | *TSR1* | 3 | 79 | 7 |
| chr9 | *EXOSC2* | 4 | 111 | 7 |
| chr3 | *THOC7* | 3 | 65 | 7 |
| chr1 | *GUK1* | 4 | 229 | 7 |
| chr19 | *NDUFB7* | 3 | 118 | 7 |
| chr11 | *MEN1* | 3 | 174 | 7 |
| chr9 | *LPAR1* | 3 | 68 | 7 |
| chr8 | *POP1* | 3 | 65 | 7 |
| chr18 | *ATP5A1* | 3 | 67 | 7 |
| chr7 | *DLD* | 2 | 63 | 7 |
| chr5 | *PGGT1B* | 3 | 59 | 7 |
| chrX | *FAM50A* | 4 | 141 | 7 |
| chr11 | *WDR74* | 3 | 128 | 7 |
| chr1 | *TAF1A* | 3 | 72 | 7 |
| chr5 | *IK* | 3 | 58 | 7 |
| chr8 | *MYC* | 3 | 139 | 7 |
| chr16 | *CDIPT* | 3 | 153 | 7 |
| chr1 | *FUBP1* | 2 | 60 | 6 |
| chr18 | *FECH* | 3 | 82 | 6 |
| chr22 | *CRKL* | 3 | 68 | 6 |
| chr17 | *SUMO2* | 3 | 77 | 6 |
| chr10 | *TIMM23* | 3 | 83 | 6 |
| chr20 | *NELFCD* | 3 | 154 | 6 |
| chr6 | *RTN4IP1* | 3 | 59 | 6 |
| chr1 | *PARS2* | 3 | 125 | 6 |
| chr10 | *ATP5C1* | 2 | 73 | 6 |
| chr18 | *TYMS* | 3 | 65 | 6 |
| chr19 | *COX6B1* | 3 | 94 | 6 |
| chr6 | *PPIL4* | 2 | 45 | 6 |
| chr18 | *SEH1L* | 3 | 63 | 6 |
| chrX | *GEMIN8* | 3 | 85 | 6 |
| chr19 | *PTBP1* | 3 | 71 | 6 |
| chr18 | *KDSR* | 2 | 47 | 6 |
| chr2 | *RPIA* | 2 | 47 | 6 |
| chr1 | *DMAP1* | 3 | 103 | 6 |
| chr11 | *CTR9* | 3 | 106 | 6 |
| chr19 | *UBE2S* | 3 | 111 | 6 |
| chr22 | *NUP50* | 2 | 58 | 6 |
| chr1 | *DDOST* | 3 | 88 | 5 |
| chr10 | *SAR1A* | 3 | 106 | 5 |
| chrX | *VBP1* | 2 | 72 | 5 |
| chr1 | *ATP5F1* | 2 | 59 | 5 |
| chr5 | *NPM1* | 2 | 63 | 5 |
| chr9 | *EDF1* | 3 | 144 | 5 |
| chr5 | *DDX41* | 3 | 83 | 5 |
| chr2 | *CFLAR* | 2 | 76 | 5 |
| chr4 | *PLK4* | 2 | 62 | 5 |
| chr14 | *LRR1* | 3 | 47 | 5 |
| chr19 | *WDR18* | 3 | 72 | 5 |

| | | | | |
|---|---|---|---|---|
| chr11 | *PCF11* | 2 | 55 | 5 |
| chr10 | *RPP30* | 2 | 48 | 5 |
| chr8 | *PABPC1* | 2 | 41 | 5 |
| chr5 | *SKP2* | 2 | 120 | 5 |
| chr5 | *NHP2* | 2 | 93 | 5 |
| chr12 | *HSP90B1* | 2 | 44 | 5 |
| chr11 | *EIF1AD* | 2 | 98 | 5 |
| chr20 | *UBE2C* | 3 | 116 | 5 |
| chr17 | *GPS1* | 3 | 89 | 5 |
| chr4 | *PPAT* | 2 | 60 | 5 |
| chr17 | *SNF8* | 2 | 78 | 5 |
| chr19 | *GRWD1* | 2 | 91 | 5 |
| chr19 | *DHPS* | 2 | 62 | 5 |
| chr19 | *TBCB* | 3 | 70 | 5 |
| chr3 | *NDUFB4* | 2 | 78 | 4 |
| chr19 | *CCNE1* | 2 | 47 | 4 |
| chr13 | *GTF3A* | 2 | 43 | 4 |
| chr10 | *PGAM1* | 2 | 50 | 4 |
| chr2 | *BUB1* | 2 | 51 | 4 |
| chr2 | *CIAO1* | 2 | 115 | 4 |
| chr11 | *CFL1* | 2 | 71 | 4 |
| chr19 | *LENG1* | 2 | 103 | 4 |
| chr6 | *PAK1IP1* | 2 | 48 | 4 |
| chr7 | *PMPCB* | 2 | 32 | 4 |
| chr22 | *ADSL* | 2 | 62 | 4 |
| chr2 | *ZEB2* | 2 | 31 | 4 |
| chr7 | *TAF6* | 2 | 78 | 4 |
| chr4 | *UBE2D3* | 2 | 51 | 4 |
| chr18 | *RNMT* | 2 | 37 | 4 |
| chr16 | *SNRNP25* | 2 | 71 | 4 |
| chr2 | *CEBPZ* | 2 | 32 | 4 |
| chrX | *NONO* | 2 | 48 | 4 |
| chr16 | *TCEB2* | 2 | 58 | 3 |
| chrX | *MED12* | 2 | 46 | 3 |
| chr3 | *TIPARP* | 2 | 43 | 3 |
| chr6 | *GMNN* | 1 | 29 | 3 |
| chr19 | *SNAPC2* | 2 | 66 | 3 |
| chr1 | *NRAS* | 2 | 43 | 3 |
| chr6 | *SYNCRIP* | 1 | 28 | 3 |
| chr2 | *MTX2* | 1 | 25 | 3 |
| chr2 | *NRBP1* | 1 | 25 | 3 |
| chr2 | *MYCN* | 1 | 28 | 3 |
| chr16 | *FAM96B* | 1 | 44 | 3 |
| chr7 | *SBDS* | 1 | 21 | 2 |
| chr5 | *THG1L* | 1 | 53 | 2 |
| chr13 | *ARGLU1* | 1 | 21 | 2 |
| chr16 | *COG8* | 1 | 52 | 2 |
| chr1 | *ATP6V0B* | 1 | 61 | 2 |
| chr16 | *MAF* | 1 | 40 | 2 |
| chr19 | *GTF2F1* | 1 | 39 | 2 |
| chr14 | *FOXA1* | 1 | 55 | 2 |
| chr17 | *DDX5* | 1 | 43 | 2 |
| chr20 | *ID1* | 1 | 107 | 2 |

| Supplementary Table 7. Kinase related genes damaged by cisplatin. | | | | | | | | |
|---|---|---|---|---|---|---|---|---|
| Chromosome | Gene | Average number of identified peaks (n=2) | Average number of reads (n=2) | Average enrichment fold (n=2) | Symbol in IPA | Entrez Gene Name | Location | Type(s) |
| chr6 | *RPS6KA2* | 122 | 5085 | 303 | RPS6KA2 | ribosomal protein S6 kinase A2 | Nucleus | protein kinase |
| chr11 | *DLG2* | 122 | 4844 | 282 | DLG2 | discs large MAGUK scaffold protein 2 | Plasma Membrane | kinase |
| chr7 | *PRKAG2* | 106 | 4667 | 259 | PRKAG2 | protein kinase AMP-activated non-catalytic subunit gamma 2 | Cytoplasm | protein kinase |
| chr3 | *EPHB1* | 109 | 3202 | 227 | EPHB1 | EPH receptor B1 | Plasma Membrane | protein kinase |
| chr5 | *TRIO* | 100 | 8450 | 223 | TRIO | trio Rho guanine nucleotide exchange factor | Cytoplasm | kinase |
| chr10 | *ADK* | 89 | 3118 | 220 | ADK | adenosine kinase | Nucleus | kinase |
| chr10 | *GRK5* | 89 | 3590 | 212 | GRK5 | G protein-coupled receptor kinase 5 | Plasma Membrane | protein kinase |
| chr2 | *PRKCE* | 99 | 3717 | 211 | PRKCE | protein kinase C epsilon | Cytoplasm | protein kinase |
| chr12 | *KSR2* | 92 | 2192 | 201 | KSR2 | kinase suppressor of ras 2 | Cytoplasm | protein kinase |
| chr3 | *KALRN* | 87 | 2677 | 189 | KALRN | kalirin RhoGEF kinase | Cytoplasm | protein kinase |
| chr1 | *EPHB2* | 79 | 3304 | 189 | EPHB2 | EPH receptor B2 | Plasma Membrane | protein kinase |
| chr7 | *MAGI2* | 79 | 2097 | 181 | MAGI2 | membrane associated guanylate kinase, WW and PDZ domain containing 2 | Plasma Membrane | kinase |
| chr17 | *PRKCA* | 78 | 3258 | 180 | PRKCA | protein kinase C alpha | Cytoplasm | protein kinase |
| chr7 | *PRKAR1B* | 77 | 4290 | 166 | PRKAR1B | protein kinase cAMP-dependent type I regulatory subunit beta | Cytoplasm | protein kinase |
| chr19 | *INSR* | 68 | 1990 | 163 | INSR | insulin receptor | Plasma Membrane | protein kinase |
| chr5 | *STK10* | 63 | 3064 | 153 | STK10 | serine/threonine kinase 10 | Cytoplasm | protein kinase |
| chr10 | *PRKG1* | 67 | 2338 | 152 | PRKG1 | protein kinase cGMP-dependent 1 | Cytoplasm | protein kinase |
| chr9 | *ABL1* | 57 | 2203 | 140 | ABL1 | ABL proto-oncogene 1, non-receptor tyrosine kinase | Nucleus | protein kinase |
| chr14 | *ITPK1* | 53 | 2299 | 135 | ITPK1 | inositol-tetrakisphosphate 1-kinase | Cytoplasm | kinase |
| chr16 | *PRKCB* | 61 | 1316 | 133 | PRKCB | protein kinase C beta | Cytoplasm | protein kinase |
| chrX | *CASK* | 50 | 1280 | 131 | CASK | calcium/calmodulin dependent serine protein kinase | Plasma Membrane | protein kinase |
| chr10 | *CAMK1D* | 49 | 1492 | 131 | CAMK1D | calcium/calmodulin dependent protein kinase ID | Cytoplasm | protein kinase |
| chr15 | *NTRK3* | 60 | 1913 | 126 | NTRK3 | neurotrophic receptor tyrosine kinase 3 | Plasma Membrane | protein kinase |
| chr3 | *MYLK* | 56 | 2341 | 123 | MYLK | myosin light chain kinase | Cytoplasm | protein kinase |
| chr7 | *EGFR* | 52 | 2523 | 122 | EGFR | epidermal growth factor receptor | Plasma Membrane | protein kinase |
| chr8 | *PTK2* | 50 | 1337 | 122 | PTK2 | protein tyrosine kinase 2 | Cytoplasm | protein kinase |
| chr17 | *KSR1* | 52 | 1961 | 121 | KSR1 | kinase suppressor of ras 1 | Cytoplasm | protein kinase |
| chr1 | *PRKCZ* | 56 | 2601 | 116 | PRKCZ | protein kinase C zeta | Cytoplasm | protein kinase |
| chr10 | *STK32C* | 53 | 2394 | 113 | STK32C | serine/threonine kinase 32C | Other | protein kinase |
| chr3 | *MAGI1* | 50 | 1085 | 113 | MAGI1 | membrane associated guanylate kinase, WW and PDZ domain containing 1 | Plasma Membrane | kinase |
| chr17 | *TAOK1* | 43 | 1095 | 113 | TAOK1 | TAO kinase 1 | Cytoplasm | protein kinase |
| chr7 | *CAMK2B* | 50 | 2965 | 113 | CAMK2B | calcium/calmodulin dependent protein kinase II beta | Cytoplasm | protein kinase |
| chr3 | *ULK4* | 47 | 1206 | 112 | ULK4 | unc-51 like kinase 4 | Other | protein kinase |
| chr7 | *SRPK2* | 45 | 1156 | 111 | SRPK2 | SRSF protein kinase 2 | Nucleus | protein kinase |
| chr1 | *MAST2* | 43 | 1202 | 109 | MAST2 | microtubule associated serine/threonine kinase 2 | Cytoplasm | protein kinase |
| chr3 | *EPHA6* | 46 | 1172 | 108 | EPHA6 | EPH receptor A6 | Plasma Membrane | protein kinase |
| chr9 | *AK8* | 47 | 2156 | 108 | AK8 | adenylate kinase 8 | Cytoplasm | kinase |
| chr9 | *DAPK1* | 46 | 1488 | 106 | DAPK1 | death associated protein kinase 1 | Cytoplasm | protein kinase |
| chr7 | *HIPK2* | 47 | 1916 | 106 | HIPK2 | homeodomain interacting protein kinase 2 | Nucleus | protein kinase |
| chr6 | *CCND3* | 41 | 1390 | 104 | CCND3 | cyclin D3 | Nucleus | kinase |
| chr7 | *CARD11* | 46 | 2242 | 103 | CARD11 | caspase recruitment domain family member 11 | Cytoplasm | kinase |
| chr9 | *ROR2* | 36 | 1313 | 103 | ROR2 | receptor tyrosine kinase like orphan receptor 2 | Plasma Membrane | protein kinase |
| chr7 | *CDK14* | 43 | 1338 | 102 | CDK14 | cyclin dependent kinase 14 | Nucleus | protein kinase |
| chr13 | *STK24* | 41 | 1902 | 101 | STK24 | serine/threonine kinase 24 | Cytoplasm | protein kinase |
| chr3 | *PIK3CB* | 39 | 1292 | 101 | PIK3CB | phosphatidylinositol-4,5-bisphosphate 3-kinase catalytic subunit | Cytoplasm | kinase |
| chr10 | *HK1* | 44 | 2237 | 98 | HK1 | hexokinase 1 | Cytoplasm | kinase |
| chr1 | *OBSCN* | 48 | 2810 | 98 | OBSCN | obscurin, cytoskeletal calmodulin and titin-interacting RhoGEF | Cytoplasm | protein kinase |
| chr11 | *SIK3* | 37 | 927 | 96 | SIK3 | SIK family kinase 3 | Cytoplasm | protein kinase |
| chr12 | *CIT* | 40 | 1039 | 94 | CIT | citron rho-interacting serine/threonine kinase | Cytoplasm | protein kinase |
| chr5 | *MAST4* | 42 | 913 | 94 | MAST4 | microtubule associated serine/threonine kinase family member | Other | protein kinase |
| chr22 | *BCR* | 33 | 1400 | 89 | BCR | BCR activator of RhoGEF and GTPase | Cytoplasm | protein kinase |
| chr15 | *DAPK2* | 39 | 1088 | 89 | DAPK2 | death associated protein kinase 2 | Cytoplasm | protein kinase |
| chr7 | *DGKB* | 36 | 995 | 88 | DGKB | diacylglycerol kinase beta | Cytoplasm | kinase |
| chr1 | *CDC42BPA* | 34 | 973 | 88 | CDC42BPA | CDC42 binding protein kinase alpha | Cytoplasm | protein kinase |
| chr17 | *SKAP1* | 35 | 843 | 87 | SKAP1 | src kinase associated phosphoprotein 1 | Cytoplasm | kinase |
| chr2 | *ALK* | 34 | 925 | 87 | ALK | ALK receptor tyrosine kinase | Plasma Membrane | protein kinase |
| chr3 | *PRKAR2A* | 34 | 912 | 87 | PRKAR2A | protein kinase cAMP-dependent type II regulatory subunit alpha | Cytoplasm | protein kinase |
| chr14 | *CDC42BPB* | 32 | 1229 | 86 | CDC42BPB | CDC42 binding protein kinase beta | Cytoplasm | protein kinase |
| chr17 | *BAIAP2* | 42 | 2153 | 85 | BAIAP2 | BAR/IMD domain containing adaptor protein 2 | Plasma Membrane | kinase |
| chr9 | *NEK6* | 38 | 2338 | 85 | NEK6 | NIMA related kinase 6 | Nucleus | protein kinase |
| chr17 | *TLK2* | 32 | 873 | 82 | TLK2 | tousled like kinase 2 | Cytoplasm | protein kinase |
| chr15 | *LRRK1* | 34 | 1949 | 81 | LRRK1 | leucine rich repeat kinase 1 | Cytoplasm | protein kinase |
| chr15 | *MAP2K5* | 34 | 855 | 80 | MAP2K5 | mitogen-activated protein kinase kinase 5 | Cytoplasm | protein kinase |
| chr2 | *NCK2* | 34 | 1197 | 79 | NCK2 | NCK adaptor protein 2 | Cytoplasm | protein kinase |
| chr3 | *RYK* | 34 | 1230 | 79 | RYK | receptor like tyrosine kinase | Plasma Membrane | protein kinase |
| chr1 | *TESK2* | 33 | 794 | 79 | TESK2 | testis associated actin remodelling kinase 2 | Nucleus | protein kinase |
| chr17 | *TEX14* | 34 | 851 | 79 | TEX14 | testis expressed 14, intercellular bridge forming factor | Plasma Membrane | protein kinase |
| chr7 | *HUS1* | 33 | 1292 | 78 | HUS1 | HUS1 checkpoint clamp component | Nucleus | kinase |
| chr12 | *TAOK3* | 27 | 760 | 77 | TAOK3 | TAO kinase 3 | Cytoplasm | protein kinase |
| chr3 | *IRAK2* | 32 | 1235 | 77 | IRAK2 | interleukin 1 receptor associated kinase 2 | Plasma Membrane | protein kinase |
| chr6 | *PTK7* | 32 | 1157 | 77 | PTK7 | protein tyrosine kinase 7 (inactive) | Plasma Membrane | protein kinase |
| chr17 | *CDK12* | 31 | 841 | 76 | CDK12 | cyclin dependent kinase 12 | Nucleus | protein kinase |
| chr19 | *BRD4* | 32 | 1190 | 75 | BRD4 | bromodomain containing 4 | Nucleus | protein kinase |
| chr1 | *TGFBR3* | 29 | 765 | 75 | TGFBR3 | transforming growth factor beta receptor 3 | Plasma Membrane | protein kinase |
| chrX | *WNK3* | 30 | 862 | 75 | WNK3 | WNK lysine deficient protein kinase 3 | Plasma Membrane | protein kinase |
| chr7 | *DGKI* | 30 | 916 | 74 | DGKI | diacylglycerol kinase iota | Cytoplasm | kinase |
| chr8 | *PTK2B* | 32 | 1120 | 72 | PTK2B | protein tyrosine kinase 2 beta | Cytoplasm | protein kinase |
| chr3 | *GSK3B* | 27 | 878 | 72 | GSK3B | glycogen synthase kinase 3 beta | Nucleus | protein kinase |
| chrX | *CDKL5* | 28 | 634 | 72 | CDKL5 | cyclin dependent kinase like 5 | Nucleus | protein kinase |
| chr6 | *PACSIN1* | 30 | 1158 | 71 | PACSIN1 | protein kinase C and casein kinase substrate in neurons 1 | Cytoplasm | protein kinase |
| chr9 | *WNK2* | 27 | 1257 | 69 | WNK2 | WNK lysine deficient protein kinase 2 | Cytoplasm | protein kinase |
| chr15 | *PEAK1* | 27 | 776 | 69 | PEAK1 | pseudopodium enriched atypical kinase 1 | Plasma Membrane | protein kinase |
| chr1 | *PIK3R3* | 27 | 749 | 67 | PIK3R3 | phosphoinositide-3-kinase regulatory subunit 3 | Cytoplasm | kinase |
| chr22 | *PI4KA* | 25 | 986 | 67 | PI4KA | phosphatidylinositol 4-kinase alpha | Cytoplasm | kinase |

| | | | | | | | | |
|---|---|---|---|---|---|---|---|---|
| chr1 | *ITPKB* | 29 | 1419 | 67 | ITPKB | inositol-trisphosphate 3-kinase B | Cytoplasm | kinase |
| chr12 | *CAMKK2* | 28 | 990 | 66 | CAMKK2 | calcium/calmodulin dependent protein kinase kinase 2 | Cytoplasm | protein kinase |
| chr2 | *MAP4K3* | 24 | 795 | 66 | MAP4K3 | mitogen-activated protein kinase kinase kinase kinase 3 | Other | protein kinase |
| chr1 | *MTOR* | 28 | 747 | 66 | MTOR | mechanistic target of rapamycin kinase | Nucleus | protein kinase |
| chr7 | *BRAF* | 24 | 627 | 65 | BRAF | B-Raf proto-oncogene, serine/threonine kinase | Cytoplasm | protein kinase |
| chr10 | *BMPR1A* | 26 | 625 | 64 | BMPR1A | bone morphogenetic protein receptor type 1A | Plasma Membrane | protein kinase |
| chr2 | *AAK1* | 26 | 663 | 63 | AAK1 | AP2 associated kinase 1 | Cytoplasm | protein kinase |
| chr11 | *PIK3C2A* | 25 | 613 | 63 | PIK3C2A | phosphatidylinositol-4-phosphate 3-kinase catalytic subunit type 2 alpha | Cytoplasm | kinase |
| chr2 | *MAP4K4* | 26 | 756 | 63 | MAP4K4 | mitogen-activated protein kinase kinase kinase kinase 4 | Cytoplasm | protein kinase |
| chr15 | *TTBK2* | 23 | 626 | 62 | TTBK2 | tau tubulin kinase 2 | Cytoplasm | protein kinase |
| chr7 | *LMTK2* | 26 | 1170 | 62 | LMTK2 | lemur tyrosine kinase 2 | Cytoplasm | protein kinase |
| chr1 | *ADCK3* | 29 | 1392 | 62 | COQ8A | coenzyme Q8A | Cytoplasm | protein kinase |
| chr7 | *TPK1* | 23 | 578 | 61 | TPK1 | thiamin pyrophosphokinase 1 | Cytoplasm | kinase |
| chr10 | *FGFR2* | 26 | 769 | 61 | FGFR2 | fibroblast growth factor receptor 2 | Plasma Membrane | protein kinase |
| chr1 | *MAGI3* | 24 | 989 | 61 | MAGI3 | membrane associated guanylate kinase, WW and PDZ domain containing 3 | Cytoplasm | kinase |
| chr7 | *CDK13* | 23 | 694 | 59 | CDK13 | cyclin dependent kinase 13 | Nucleus | protein kinase |
| chr4 | *STK32B* | 22 | 647 | 58 | STK32B | serine/threonine kinase 32B | Other | protein kinase |
| chr16 | *PDPK1* | 26 | 1308 | 58 | PDPK1 | 3-phosphoinositide dependent protein kinase 1 | Cytoplasm | protein kinase |
| chr1 | *PIK3CD* | 25 | 898 | 57 | PIK3CD | phosphatidylinositol-4,5-bisphosphate 3-kinase catalytic subunit | Cytoplasm | kinase |
| chr1 | *AKT3* | 22 | 654 | 57 | AKT3 | AKT serine/threonine kinase 3 | Cytoplasm | protein kinase |
| chr19 | *PAK4* | 24 | 1200 | 57 | PAK4 | p21 (RAC1) activated kinase 4 | Cytoplasm | protein kinase |
| chr1 | *RPS6KA1* | 23 | 977 | 56 | RPS6KA1 | ribosomal protein S6 kinase A1 | Cytoplasm | protein kinase |
| chr1 | *RPS6KC1* | 23 | 693 | 56 | RPS6KC1 | ribosomal protein S6 kinase C1 | Cytoplasm | protein kinase |
| chr7 | *PODXL* | 24 | 894 | 56 | PODXL | podocalyxin like | Plasma Membrane | kinase |
| chr2 | *MERTK* | 24 | 801 | 56 | MERTK | MER proto-oncogene, tyrosine kinase | Plasma Membrane | protein kinase |
| chr1 | *ABL2* | 24 | 756 | 55 | ABL2 | ABL proto-oncogene 2, non-receptor tyrosine kinase | Cytoplasm | protein kinase |
| chr20 | *SRC* | 26 | 1370 | 55 | SRC | SRC proto-oncogene, non-receptor tyrosine kinase | Cytoplasm | protein kinase |
| chr7 | *MMD2* | 25 | 937 | 54 | MMD2 | monocyte to macrophage differentiation associated 2 | Other | kinase |
| chr21 | *HUNK* | 22 | 613 | 54 | HUNK | hormonally up-regulated Neu-associated kinase | Cytoplasm | protein kinase |
| chr8 | *NDRG1* | 23 | 893 | 54 | NDRG1 | N-myc downstream regulated 1 | Nucleus | kinase |
| chr19 | *AKT2* | 22 | 1054 | 53 | AKT2 | AKT serine/threonine kinase 2 | Cytoplasm | protein kinase |
| chr3 | *VPRBP* | 23 | 588 | 53 | DCAF1 | DDB1 and CUL4 associated factor 1 | Nucleus | protein kinase |
| chr16 | *EEF2K* | 23 | 740 | 53 | EEF2K | eukaryotic elongation factor 2 kinase | Cytoplasm | protein kinase |
| chr3 | *DGKG* | 20 | 541 | 53 | DGKG | diacylglycerol kinase gamma | Cytoplasm | kinase |
| chr1 | *PIK3C2B* | 24 | 1093 | 53 | PIK3C2B | phosphatidylinositol-4-phosphate 3-kinase catalytic subunit type | Cytoplasm | kinase |
| chr9 | *SHC3* | 23 | 613 | 52 | SHC3 | SHC adaptor protein 3 | Cytoplasm | kinase |
| chr11 | *CHKA* | 22 | 659 | 52 | CHKA | choline kinase alpha | Cytoplasm | kinase |
| chr8 | *STK3* | 19 | 547 | 52 | STK3 | serine/threonine kinase 3 | Cytoplasm | protein kinase |
| chr10 | *CAMK2G* | 23 | 1455 | 52 | CAMK2G | calcium/calmodulin dependent protein kinase II gamma | Cytoplasm | protein kinase |
| chr5 | *FER* | 20 | 462 | 52 | FER | FER tyrosine kinase | Cytoplasm | protein kinase |
| chr15 | *MAP2K1* | 22 | 542 | 52 | MAP2K1 | mitogen-activated protein kinase kinase 1 | Cytoplasm | protein kinase |
| chr17 | *MINK1* | 22 | 865 | 51 | MINK1 | misshapen like kinase 1 | Cytoplasm | protein kinase |
| chr10 | *RET* | 23 | 1227 | 50 | RET | ret proto-oncogene | Plasma Membrane | protein kinase |
| chr11 | *DGKZ* | 24 | 1453 | 50 | DGKZ | diacylglycerol kinase zeta | Cytoplasm | kinase |
| chr17 | *SHPK* | 20 | 830 | 50 | SHPK | sedoheptulokinase | Cytoplasm | kinase |
| chr11 | *HIPK3* | 17 | 603 | 50 | HIPK3 | homeodomain interacting protein kinase 3 | Nucleus | protein kinase |
| chr17 | *PIK3R5* | 22 | 696 | 49 | PIK3R5 | phosphoinositide-3-kinase regulatory subunit 5 | Cytoplasm | kinase |
| chr1 | *DSTYK* | 21 | 636 | 49 | DSTYK | dual serine/threonine and tyrosine protein kinase | Cytoplasm | protein kinase |
| chr16 | *PHKB* | 20 | 527 | 49 | PHKB | phosphorylase kinase regulatory subunit beta | Cytoplasm | protein kinase |
| chr11 | *PAK1* | 18 | 565 | 49 | PAK1 | p21 (RAC1) activated kinase 1 | Cytoplasm | protein kinase |
| chr1 | *AK5* | 22 | 441 | 48 | AK5 | adenylate kinase 5 | Cytoplasm | kinase |
| chr19 | *MAST3* | 21 | 830 | 48 | MAST3 | microtubule associated serine/threonine kinase 3 | Other | protein kinase |
| chr19 | *PKN1* | 22 | 845 | 48 | PKN1 | protein kinase N1 | Cytoplasm | protein kinase |
| chr9 | *PIP5K1B* | 21 | 438 | 48 | PIP5K1B | phosphatidylinositol-4-phosphate 5-kinase type 1 beta | Cytoplasm | kinase |
| chr3 | *PRKCI* | 21 | 851 | 47 | PRKCI | protein kinase C iota | Cytoplasm | protein kinase |
| chr16 | *SMG1* | 19 | 515 | 47 | SMG1 | SMG1 nonsense mediated mRNA decay associated PI3K related kinase | Cytoplasm | protein kinase |
| chr10 | *PFKFB3* | 18 | 733 | 47 | PFKFB3 | 6-phosphofructo-2-kinase/fructose-2,6-biphosphatase 3 | Cytoplasm | kinase |
| chr17 | *ERN1* | 19 | 685 | 47 | ERN1 | endoplasmic reticulum to nucleus signaling 1 | Cytoplasm | protein kinase |
| chrX | *PHKA2* | 19 | 677 | 47 | PHKA2 | phosphorylase kinase regulatory subunit alpha 2 | Cytoplasm | protein kinase |
| chr20 | *HCK* | 20 | 735 | 47 | HCK | HCK proto-oncogene, Src family tyrosine kinase | Cytoplasm | protein kinase |
| chr7 | *LIMK1* | 21 | 835 | 46 | LIMK1 | LIM domain kinase 1 | Cytoplasm | protein kinase |
| chr17 | *MAP2K6* | 17 | 439 | 45 | MAP2K6 | mitogen-activated protein kinase kinase 6 | Cytoplasm | protein kinase |
| chr20 | *STK4* | 15 | 521 | 45 | STK4 | serine/threonine kinase 4 | Cytoplasm | protein kinase |
| chr3 | *RAF1* | 19 | 554 | 45 | RAF1 | Raf-1 proto-oncogene, serine/threonine kinase | Cytoplasm | protein kinase |
| chr1 | *ROR1* | 21 | 427 | 45 | ROR1 | receptor tyrosine kinase like orphan receptor 1 | Plasma Membrane | protein kinase |
| chr3 | *NEK11* | 19 | 536 | 44 | NEK11 | NIMA related kinase 11 | Nucleus | protein kinase |
| chr1 | *JAK1* | 17 | 542 | 44 | JAK1 | Janus kinase 1 | Cytoplasm | protein kinase |
| chr17 | *ULK2* | 18 | 467 | 44 | ULK2 | unc-51 like autophagy activating kinase 2 | Cytoplasm | protein kinase |
| chr11 | *SIK2* | 17 | 416 | 44 | SIK2 | salt inducible kinase 2 | Cytoplasm | protein kinase |
| chr4 | *GRK4* | 16 | 556 | 44 | GRK4 | G protein-coupled receptor kinase 4 | Plasma Membrane | protein kinase |
| chr3 | *PFKFB4* | 18 | 750 | 44 | PFKFB4 | 6-phosphofructo-2-kinase/fructose-2,6-biphosphatase 4 | Cytoplasm | kinase |
| chr16 | *MYLK3* | 20 | 700 | 44 | MYLK3 | myosin light chain kinase 3 | Cytoplasm | protein kinase |
| chr9 | *TJP2* | 18 | 455 | 43 | TJP2 | tight junction protein 2 | Plasma Membrane | kinase |
| chr3 | *IP6K2* | 18 | 554 | 43 | IP6K2 | inositol hexakisphosphate kinase 2 | Cytoplasm | kinase |
| chr3 | *PRKCD* | 18 | 857 | 43 | PRKCD | protein kinase C delta | Cytoplasm | protein kinase |
| chr5 | *CAMK2A* | 18 | 659 | 43 | CAMK2A | calcium/calmodulin dependent protein kinase II alpha | Cytoplasm | protein kinase |
| chr19 | *MARK4* | 17 | 647 | 43 | MARK4 | microtubule affinity regulating kinase 4 | Cytoplasm | protein kinase |
| chr15 | *TRPM7* | 15 | 357 | 43 | TRPM7 | transient receptor potential cation channel subfamily M member | Plasma Membrane | protein kinase |
| chr5 | *MAPK9* | 17 | 785 | 43 | MAPK9 | mitogen-activated protein kinase 9 | Cytoplasm | protein kinase |
| chr17 | *PIK3R6* | 18 | 627 | 43 | PIK3R6 | phosphoinositide-3-kinase regulatory subunit 6 | Cytoplasm | kinase |
| chr2 | *BMPR2* | 14 | 373 | 42 | BMPR2 | bone morphogenetic protein receptor type 2 | Plasma Membrane | protein kinase |
| chr2 | *MAP3K2* | 17 | 571 | 42 | MAP3K2 | mitogen-activated protein kinase kinase kinase 2 | Cytoplasm | protein kinase |
| chr6 | *TTBK1* | 19 | 808 | 42 | TTBK1 | tau tubulin kinase 1 | Other | protein kinase |
| chr1 | *EPHA8* | 20 | 965 | 42 | EPHA8 | EPH receptor A8 | Plasma Membrane | protein kinase |
| chr3 | *XYLB* | 19 | 540 | 42 | XYLB | xylulokinase | Cytoplasm | kinase |
| chr14 | *MARK3* | 14 | 372 | 42 | MARK3 | microtubule affinity regulating kinase 3 | Cytoplasm | protein kinase |
| chr2 | *DGKD* | 16 | 768 | 42 | DGKD | diacylglycerol kinase delta | Cytoplasm | kinase |
| chr3 | *IP6K1* | 17 | 508 | 42 | IP6K1 | inositol hexakisphosphate kinase 1 | Cytoplasm | kinase |
| chr6 | *MYLK4* | 17 | 433 | 41 | MYLK4 | myosin light chain kinase family member 4 | Cytoplasm | protein kinase |
| chr8 | *PRKDC* | 13 | 414 | 41 | PRKDC | protein kinase, DNA-activated, catalytic subunit | Nucleus | protein kinase |

| | | | | | | | | |
|---|---|---|---|---|---|---|---|---|
| chr5 | *PDGFRB* | 18 | 768 | 41 | PDGFRB | platelet derived growth factor receptor beta | Plasma Membrane | protein kinase |
| chr17 | *MAP3K14* | 17 | 838 | 41 | MAP3K14 | mitogen-activated protein kinase kinase kinase 14 | Cytoplasm | protein kinase |
| chr3 | *PXK* | 18 | 408 | 41 | PXK | PX domain containing serine/threonine kinase like | Cytoplasm | protein kinase |
| chr3 | *ATR* | 15 | 474 | 40 | ATR | ATR serine/threonine kinase | Nucleus | protein kinase |
| chr9 | *NTRK2* | 19 | 392 | 40 | NTRK2 | neurotrophic receptor tyrosine kinase 2 | Plasma Membrane | protein kinase |
| chr7 | *GCK* | 19 | 819 | 40 | GCK | glucokinase | Cytoplasm | kinase |
| chr3 | *NEK10* | 16 | 439 | 40 | NEK10 | NIMA related kinase 10 | Other | protein kinase |
| chr3 | *NEK4* | 15 | 452 | 39 | NEK4 | NIMA related kinase 4 | Nucleus | protein kinase |
| chr14 | *PRKCH* | 15 | 383 | 39 | PRKCH | protein kinase C eta | Cytoplasm | protein kinase |
| chr7 | *CDK6* | 17 | 539 | 39 | CDK6 | cyclin dependent kinase 6 | Nucleus | protein kinase |
| chr12 | *CDK17* | 15 | 361 | 39 | CDK17 | cyclin dependent kinase 17 | Cytoplasm | protein kinase |
| chr22 | *MAPK1* | 13 | 390 | 39 | MAPK1 | mitogen-activated protein kinase 1 | Cytoplasm | protein kinase |
| chr19 | *VRK3* | 14 | 547 | 39 | VRK3 | VRK serine/threonine kinase 3 | Nucleus | protein kinase |
| chr19 | *AXL* | 17 | 609 | 38 | AXL | AXL receptor tyrosine kinase | Plasma Membrane | protein kinase |
| chr1 | *PIP5K1A* | 16 | 586 | 38 | PIP5K1A | phosphatidylinositol-4-phosphate 5-kinase type 1 alpha | Cytoplasm | kinase |
| chr17 | *MAP3K3* | 16 | 495 | 38 | MAP3K3 | mitogen-activated protein kinase kinase kinase 3 | Cytoplasm | protein kinase |
| chr15 | *MAPK6* | 14 | 404 | 38 | MAPK6 | mitogen-activated protein kinase 6 | Cytoplasm | protein kinase |
| chr11 | *STK33* | 17 | 478 | 38 | STK33 | serine/threonine kinase 33 | Cytoplasm | protein kinase |
| chr15 | *EIF2AK4* | 14 | 372 | 38 | EIF2AK4 | eukaryotic translation initiation factor 2 alpha kinase 4 | Cytoplasm | kinase |
| chr3 | *EPHA3* | 15 | 421 | 38 | EPHA3 | EPH receptor A3 | Plasma Membrane | protein kinase |
| chr1 | *BRDT* | 14 | 370 | 38 | BRDT | bromodomain testis associated | Nucleus | kinase |
| chr15 | *ALPK3* | 19 | 703 | 38 | ALPK3 | alpha kinase 3 | Nucleus | protein kinase |
| chr1 | *EPHA2* | 17 | 831 | 38 | EPHA2 | EPH receptor A2 | Plasma Membrane | protein kinase |
| chr19 | *BRSK1* | 16 | 728 | 38 | BRSK1 | BR serine/threonine kinase 1 | Cytoplasm | protein kinase |
| chr22 | *CERK* | 16 | 663 | 38 | CERK | ceramide kinase | Plasma Membrane | kinase |
| chr8 | *BLK* | 16 | 574 | 38 | BLK | BLK proto-oncogene, Src family tyrosine kinase | Cytoplasm | protein kinase |
| chr3 | *NCK1* | 15 | 472 | 38 | NCK1 | NCK adaptor protein 1 | Cytoplasm | protein kinase |
| chr1 | *DDR2* | 17 | 454 | 37 | DDR2 | discoidin domain receptor tyrosine kinase 2 | Plasma Membrane | protein kinase |
| chr17 | *AATK* | 18 | 650 | 36 | AATK | apoptosis associated tyrosine kinase | Cytoplasm | protein kinase |
| chr19 | *PIP5K1C* | 18 | 794 | 36 | PIP5K1C | phosphatidylinositol-4-phosphate 5-kinase type 1 gamma | Plasma Membrane | kinase |
| chr11 | *MARK2* | 15 | 450 | 36 | MARK2 | microtubule affinity regulating kinase 2 | Cytoplasm | protein kinase |
| chr17 | *CAMKK1* | 16 | 767 | 35 | CAMKK1 | calcium/calmodulin dependent protein kinase kinase 1 | Cytoplasm | protein kinase |
| chr5 | *CSF1R* | 14 | 519 | 35 | CSF1R | colony stimulating factor 1 receptor | Plasma Membrane | protein kinase |
| chr17 | *NLK* | 13 | 348 | 35 | NLK | nemo like kinase | Nucleus | protein kinase |
| chr1 | *NTRK1* | 16 | 831 | 35 | NTRK1 | neurotrophic receptor tyrosine kinase 1 | Plasma Membrane | protein kinase |
| chr17 | *RPS6KB1* | 15 | 383 | 35 | RPS6KB1 | ribosomal protein S6 kinase B1 | Cytoplasm | protein kinase |
| chr5 | *EFNA5* | 15 | 339 | 35 | EFNA5 | ephrin A5 | Plasma Membrane | protein kinase |
| chr1 | *STK40* | 15 | 697 | 35 | STK40 | serine/threonine kinase 40 | Cytoplasm | protein kinase |
| chr14 | *MOK* | 13 | 466 | 34 | MOK | MOK protein kinase | Plasma Membrane | protein kinase |
| chr22 | *CSNK1E* | 14 | 531 | 34 | CSNK1E | casein kinase 1 epsilon | Cytoplasm | protein kinase |
| chr18 | *YES1* | 13 | 296 | 34 | YES1 | YES proto-oncogene 1, Src family tyrosine kinase | Cytoplasm | protein kinase |
| chr1 | *NOL9* | 15 | 597 | 34 | NOL9 | nucleolar protein 9 | Nucleus | kinase |
| chr15 | *GALK2* | 12 | 326 | 34 | GALK2 | galactokinase 2 | Cytoplasm | kinase |
| chr3 | *GK5* | 13 | 404 | 34 | GK5 | glycerol kinase 5 | Other | kinase |
| chr2 | *PASK* | 14 | 589 | 34 | PASK | PAS domain containing serine/threonine kinase | Cytoplasm | protein kinase |
| chr10 | *PI4K2A* | 13 | 362 | 33 | PI4K2A | phosphatidylinositol 4-kinase type 2 alpha | Cytoplasm | kinase |
| chr9 | *MELK* | 12 | 292 | 33 | MELK | maternal embryonic leucine zipper kinase | Cytoplasm | protein kinase |
| chr10 | *PFKP* | 13 | 526 | 33 | PFKP | phosphofructokinase, platelet | Cytoplasm | kinase |
| chr4 | *GAK* | 16 | 650 | 33 | GAK | cyclin G associated kinase | Nucleus | protein kinase |
| chr1 | *MAPKAPK2* | 13 | 695 | 33 | MAPKAPK2 | MAPK activated protein kinase 2 | Nucleus | protein kinase |
| chr16 | *PSKH1* | 14 | 515 | 32 | PSKH1 | protein serine kinase H1 | Nucleus | protein kinase |
| chrX | *PDK3* | 12 | 288 | 32 | PDK3 | pyruvate dehydrogenase kinase 3 | Cytoplasm | protein kinase |
| chr1 | *LCK* | 14 | 365 | 31 | LCK | LCK proto-oncogene, Src family tyrosine kinase | Cytoplasm | protein kinase |
| chr6 | *STK38* | 12 | 332 | 31 | STK38 | serine/threonine kinase 38 | Nucleus | protein kinase |
| chr6 | *CNKSR3* | 12 | 338 | 31 | CNKSR3 | CNKSR family member 3 | Plasma Membrane | kinase |
| chr1 | *CMPK1* | 13 | 345 | 31 | CMPK1 | cytidine/uridine monophosphate kinase 1 | Nucleus | kinase |
| chr12 | *MAPKAPK5* | 14 | 341 | 31 | MAPKAPK5 | MAPK activated protein kinase 5 | Cytoplasm | protein kinase |
| chr7 | *STK31* | 13 | 455 | 31 | STK31 | serine/threonine kinase 31 | Cytoplasm | protein kinase |
| chr3 | *TNIK* | 11 | 273 | 31 | TNIK | TRAF2 and NCK interacting kinase | Plasma Membrane | protein kinase |
| chr1 | *PI4KB* | 13 | 539 | 31 | PI4KB | phosphatidylinositol 4-kinase beta | Cytoplasm | kinase |
| chr2 | *HK2* | 14 | 423 | 31 | HK2 | hexokinase 2 | Cytoplasm | kinase |
| chr14 | *ADCK1* | 12 | 375 | 31 | ADCK1 | aarF domain containing kinase 1 | Cytoplasm | protein kinase |
| chr1 | *CDK18* | 14 | 956 | 30 | CDK18 | cyclin dependent kinase 18 | Cytoplasm | protein kinase |
| chr17 | *ERBB2* | 12 | 535 | 30 | ERBB2 | erb-b2 receptor tyrosine kinase 2 | Plasma Membrane | protein kinase |
| chr8 | *FGFR1* | 11 | 406 | 30 | FGFR1 | fibroblast growth factor receptor 1 | Plasma Membrane | protein kinase |
| chr1 | *NME7* | 13 | 439 | 29 | NME7 | NME/NM23 family member 7 | Cytoplasm | kinase |
| chr9 | *C9orf96* | 13 | 703 | 29 | STKLD1 | serine/threonine kinase like domain containing 1 | Other | protein kinase |
| chr12 | *STK38L* | 11 | 340 | 29 | STK38L | serine/threonine kinase 38 like | Cytoplasm | protein kinase |
| chr1 | *AK4* | 12 | 286 | 29 | AK4 | adenylate kinase 4 | Cytoplasm | kinase |
| chr16 | *SBK1* | 13 | 558 | 29 | SBK1 | SH3 domain binding kinase 1 | Other | protein kinase |
| chr3 | *MAP3K13* | 11 | 241 | 29 | MAP3K13 | mitogen-activated protein kinase kinase kinase 13 | Cytoplasm | protein kinase |
| chr6 | *IP6K3* | 12 | 561 | 29 | IP6K3 | inositol hexakisphosphate kinase 3 | Cytoplasm | kinase |
| chr6 | *RIPK1* | 12 | 320 | 29 | RIPK1 | receptor interacting serine/threonine kinase 1 | Plasma Membrane | protein kinase |
| chr18 | *MAPK4* | 12 | 297 | 29 | MAPK4 | mitogen-activated protein kinase 4 | Cytoplasm | protein kinase |
| chr6 | *SGK1* | 10 | 265 | 28 | SGK1 | serum/glucocorticoid regulated kinase 1 | Cytoplasm | protein kinase |
| chr9 | *TRPM6* | 12 | 309 | 28 | TRPM6 | transient receptor potential cation channel subfamily M member | Plasma Membrane | protein kinase |
| chr1 | *IKBKE* | 11 | 700 | 28 | IKBKE | inhibitor of nuclear factor kappa B kinase subunit epsilon | Cytoplasm | kinase |
| chr3 | *PAK2* | 9 | 255 | 28 | PAK2 | p21 (RAC1) activated kinase 2 | Cytoplasm | protein kinase |
| chr17 | *MPP2* | 12 | 549 | 28 | MPP2 | membrane palmitoylated protein 2 | Plasma Membrane | kinase |
| chr2 | *SPEG* | 13 | 563 | 28 | SPEG | striated muscle enriched protein kinase | Nucleus | protein kinase |
| chr17 | *DLG4* | 12 | 565 | 28 | DLG4 | discs large MAGUK scaffold protein 4 | Plasma Membrane | kinase |
| chr17 | *MAP2K3* | 13 | 1112 | 28 | MAP2K3 | mitogen-activated protein kinase kinase 3 | Cytoplasm | protein kinase |
| chr13 | *LATS2* | 10 | 307 | 28 | LATS2 | large tumor suppressor kinase 2 | Nucleus | protein kinase |
| chr17 | *PIP4K2B* | 12 | 377 | 28 | PIP4K2B | phosphatidylinositol-5-phosphate 4-kinase type 2 beta | Cytoplasm | kinase |
| chr21 | *PDXK* | 13 | 550 | 27 | PDXK | pyridoxal kinase | Cytoplasm | kinase |
| chr19 | *PRKACA* | 11 | 460 | 27 | PRKACA | protein kinase cAMP-activated catalytic subunit alpha | Cytoplasm | protein kinase |
| chr17 | *MPP3* | 11 | 433 | 27 | MPP3 | membrane palmitoylated protein 3 | Plasma Membrane | kinase |
| chr19 | *MAP2K2* | 12 | 479 | 27 | MAP2K2 | mitogen-activated protein kinase kinase 2 | Cytoplasm | protein kinase |
| chr19 | *JAK3* | 12 | 498 | 27 | JAK3 | Janus kinase 3 | Cytoplasm | protein kinase |
| chr6 | *SRPK1* | 10 | 276 | 27 | SRPK1 | SRSF protein kinase 1 | Nucleus | protein kinase |
| chr7 | *EIF2AK1* | 12 | 405 | 26 | EIF2AK1 | eukaryotic translation initiation factor 2 alpha kinase 1 | Cytoplasm | kinase |
| chr3 | *MAPKAPK3* | 12 | 425 | 26 | MAPKAPK3 | MAPK activated protein kinase 3 | Nucleus | protein kinase |
| chr5 | *ROPN1L* | 12 | 1199 | 26 | ROPN1L | rhophilin associated tail protein 1 like | Cytoplasm | kinase |

| | | | | | | | | |
|---|---|---|---|---|---|---|---|---|
| chr8 | *PDGFRL* | 11 | 331 | 26 | PDGFRL | platelet derived growth factor receptor like | Plasma Membrane | protein kinase |
| chr6 | *MAPK14* | 11 | 266 | 26 | MAPK14 | mitogen-activated protein kinase 14 | Cytoplasm | protein kinase |
| chr5 | *COL4A3BP* | 10 | 235 | 26 | CERT1 | ceramide transporter 1 | Cytoplasm | protein kinase |
| chr1 | *DYRK3* | 12 | 350 | 26 | DYRK3 | dual specificity tyrosine phosphorylation regulated kinase 3 | Nucleus | protein kinase |
| chr12 | *ACVR1B* | 10 | 537 | 26 | ACVR1B | activin A receptor type 1B | Plasma Membrane | protein kinase |
| chr4 | *NEK1* | 9 | 269 | 25 | NEK1 | NIMA related kinase 1 | Nucleus | protein kinase |
| chr15 | *TYRO3* | 10 | 368 | 25 | TYRO3 | TYRO3 protein tyrosine kinase | Plasma Membrane | protein kinase |
| chr5 | *GRK6* | 11 | 423 | 25 | GRK6 | G protein-coupled receptor kinase 6 | Plasma Membrane | protein kinase |
| chr19 | *MAP4K1* | 10 | 380 | 25 | MAP4K1 | mitogen-activated protein kinase kinase kinase kinase 1 | Cytoplasm | protein kinase |
| chr15 | *CLK3* | 10 | 506 | 25 | CLK3 | CDC like kinase 3 | Nucleus | protein kinase |
| chr10 | *IPMK* | 11 | 415 | 25 | IPMK | inositol polyphosphate multikinase | Nucleus | kinase |
| chr11 | *ADRBK1* | 11 | 524 | 24 | GRK2 | G protein-coupled receptor kinase 2 | Cytoplasm | protein kinase |
| chr8 | *IKBKB* | 9 | 370 | 24 | IKBKB | inhibitor of nuclear factor kappa B kinase subunit beta | Cytoplasm | kinase |
| chr4 | *BMPR1B* | 9 | 215 | 24 | BMPR1B | bone morphogenetic protein receptor type 1B | Plasma Membrane | protein kinase |
| chr1 | *NEK7* | 11 | 360 | 24 | NEK7 | NIMA related kinase 7 | Nucleus | protein kinase |
| chr6 | *MAP3K5* | 8 | 181 | 24 | MAP3K5 | mitogen-activated protein kinase kinase kinase 5 | Cytoplasm | protein kinase |
| chr2 | *ERBB4* | 9 | 242 | 24 | ERBB4 | erb-b2 receptor tyrosine kinase 4 | Plasma Membrane | protein kinase |
| chr20 | *RASSF2* | 9 | 294 | 24 | RASSF2 | Ras association domain family member 2 | Nucleus | kinase |
| chr10 | *MAPK8* | 9 | 374 | 24 | MAPK8 | mitogen-activated protein kinase 8 | Cytoplasm | protein kinase |
| chr16 | *FUK* | 11 | 415 | 24 | FCSK | fucose kinase | Cytoplasm | kinase |
| chr2 | *CDKL4* | 9 | 295 | 23 | CDKL4 | cyclin dependent kinase like 4 | Nucleus | protein kinase |
| chr3 | *DLG1* | 7 | 188 | 23 | DLG1 | discs large MAGUK scaffold protein 1 | Plasma Membrane | kinase |
| chr1 | *MARK1* | 9 | 291 | 23 | MARK1 | microtubule affinity regulating kinase 1 | Cytoplasm | protein kinase |
| chr19 | *PRKCG* | 11 | 399 | 23 | PRKCG | protein kinase C gamma | Cytoplasm | protein kinase |
| chr15 | *PKM* | 10 | 305 | 23 | PKM | pyruvate kinase M1/2 | Cytoplasm | kinase |
| chr1 | *TIE1* | 9 | 415 | 23 | TIE1 | tyrosine kinase with immunoglobulin like and EGF like domains | Plasma Membrane | protein kinase |
| chr7 | *MPP6* | 9 | 271 | 23 | MPP6 | membrane palmitoylated protein 6 | Plasma Membrane | kinase |
| chr2 | *ZAP70* | 11 | 613 | 23 | ZAP70 | zeta chain of T cell receptor associated protein kinase 70 | Plasma Membrane | protein kinase |
| chr1 | *PKN2* | 10 | 204 | 23 | PKN2 | protein kinase N2 | Cytoplasm | protein kinase |
| chr19 | *MAP3K10* | 11 | 359 | 23 | MAP3K10 | mitogen-activated protein kinase kinase kinase 10 | Cytoplasm | protein kinase |
| chr19 | *ITPKC* | 10 | 356 | 22 | ITPKC | inositol-trisphosphate 3-kinase C | Cytoplasm | kinase |
| chr15 | *CIB2* | 10 | 467 | 22 | CIB2 | calcium and integrin binding family member 2 | Cytoplasm | kinase |
| chr10 | *HKDC1* | 11 | 589 | 22 | HKDC1 | hexokinase domain containing 1 | Cytoplasm | kinase |
| chr17 | *MAP2K4* | 8 | 206 | 22 | MAP2K4 | mitogen-activated protein kinase kinase 4 | Cytoplasm | protein kinase |
| chr3 | *TNK2* | 9 | 474 | 22 | TNK2 | tyrosine kinase non receptor 2 | Cytoplasm | protein kinase |
| chr15 | *CSK* | 9 | 350 | 22 | CSK | C-terminal Src kinase | Cytoplasm | protein kinase |
| chr11 | *MAP4K2* | 9 | 417 | 22 | MAP4K2 | mitogen-activated protein kinase kinase kinase kinase 2 | Cytoplasm | protein kinase |
| chr3 | *ACVR2B* | 10 | 301 | 21 | ACVR2B | activin A receptor type 2B | Plasma Membrane | protein kinase |
| chr14 | *RPS6KA5* | 8 | 172 | 21 | RPS6KA5 | ribosomal protein S6 kinase A5 | Nucleus | protein kinase |
| chr19 | *PIK3R2* | 10 | 437 | 21 | PIK3R2 | phosphoinositide-3-kinase regulatory subunit 2 | Cytoplasm | kinase |
| chr2 | *TLK1* | 7 | 189 | 21 | TLK1 | tousled like kinase 1 | Nucleus | protein kinase |
| chr5 | *CDK7* | 9 | 216 | 21 | CDK7 | cyclin dependent kinase 7 | Nucleus | protein kinase |
| chr12 | *NUAK1* | 10 | 285 | 21 | NUAK1 | NUAK family kinase 1 | Nucleus | protein kinase |
| chr12 | *ULK1* | 11 | 382 | 21 | ULK1 | unc-51 like autophagy activating kinase 1 | Cytoplasm | protein kinase |
| chr18 | *ALPK2* | 7 | 203 | 21 | ALPK2 | alpha kinase 2 | Nucleus | protein kinase |
| chr7 | *MET* | 10 | 232 | 21 | MET | MET proto-oncogene, receptor tyrosine kinase | Plasma Membrane | protein kinase |
| chr9 | *SYK* | 7 | 252 | 21 | SYK | spleen associated tyrosine kinase | Cytoplasm | protein kinase |
| chr4 | *MAPK10* | 8 | 209 | 21 | MAPK10 | mitogen-activated protein kinase 10 | Cytoplasm | protein kinase |
| chr6 | *CDK19* | 7 | 187 | 21 | CDK19 | cyclin dependent kinase 19 | Nucleus | protein kinase |
| chr16 | *TAOK2* | 9 | 372 | 20 | TAOK2 | TAO kinase 2 | Cytoplasm | protein kinase |
| chr3 | *TGFBR2* | 9 | 241 | 20 | TGFBR2 | transforming growth factor beta receptor 2 | Plasma Membrane | protein kinase |
| chr18 | *ROCK1* | 7 | 170 | 20 | ROCK1 | Rho associated coiled-coil containing protein kinase 1 | Cytoplasm | protein kinase |
| chr11 | *CDC42BPG* | 10 | 525 | 20 | CDC42BPG | CDC42 binding protein kinase gamma | Cytoplasm | protein kinase |
| chr1 | *NADK* | 9 | 467 | 20 | NADK | NAD kinase | Cytoplasm | kinase |
| chr16 | *TK2* | 8 | 313 | 20 | TK2 | thymidine kinase 2 | Cytoplasm | kinase |
| chr6 | *FYN* | 8 | 183 | 20 | FYN | FYN proto-oncogene, Src family tyrosine kinase | Plasma Membrane | protein kinase |
| chr1 | *FGR* | 8 | 353 | 20 | FGR | FGR proto-oncogene, Src family tyrosine kinase | Nucleus | protein kinase |
| chr1 | *AK2* | 9 | 207 | 19 | AK2 | adenylate kinase 2 | Cytoplasm | kinase |
| chr5 | *STK32A* | 8 | 160 | 19 | STK32A | serine/threonine kinase 32A | Other | protein kinase |
| chr13 | *GRK1* | 8 | 560 | 19 | GRK1 | G protein-coupled receptor kinase 1 | Plasma Membrane | protein kinase |
| chr9 | *BRD3* | 10 | 477 | 19 | BRD3 | bromodomain containing 3 | Nucleus | protein kinase |
| chr17 | *FN3KRP* | 8 | 301 | 19 | FN3KRP | fructosamine 3 kinase related protein | Cytoplasm | kinase |
| chrX | *GK* | 7 | 173 | 19 | GK | glycerol kinase | Cytoplasm | kinase |
| chr11 | *GDPD4* | 8 | 250 | 19 | GDPD4 | glycerophosphodiester phosphodiesterase domain containing 4 | Other | kinase |
| chr14 | *CDKL1* | 7 | 195 | 19 | CDKL1 | cyclin dependent kinase like 1 | Nucleus | protein kinase |
| chr9 | *FXN* | 8 | 230 | 19 | FXN | frataxin | Cytoplasm | kinase |
| chr5 | *HK3* | 9 | 392 | 19 | HK3 | hexokinase 3 | Cytoplasm | kinase |
| chr8 | *ADCK5* | 8 | 518 | 19 | ADCK5 | aarF domain containing kinase 5 | Cytoplasm | protein kinase |
| chr10 | *MYO3A* | 7 | 169 | 18 | MYO3A | myosin IIIA | Cytoplasm | protein kinase |
| chr19 | *ADCK4* | 8 | 379 | 18 | COQ8B | coenzyme Q8B | Cytoplasm | protein kinase |
| chr3 | *MST1R* | 7 | 306 | 18 | MST1R | macrophage stimulating 1 receptor | Plasma Membrane | protein kinase |
| chr14 | *AK7* | 7 | 157 | 18 | AK7 | adenylate kinase 7 | Cytoplasm | kinase |
| chr22 | *ADRBK2* | 6 | 158 | 18 | GRK3 | G protein-coupled receptor kinase 3 | Cytoplasm | protein kinase |
| chr20 | *MYLK2* | 8 | 638 | 18 | MYLK2 | myosin light chain kinase 2 | Cytoplasm | protein kinase |
| chrX | *PRKX* | 7 | 175 | 18 | PRKX | protein kinase X-linked | Cytoplasm | protein kinase |
| chr7 | *EPHB4* | 8 | 378 | 18 | EPHB4 | EPH receptor B4 | Plasma Membrane | protein kinase |
| chr1 | *CDK11B* | 8 | 451 | 18 | CDK11B | cyclin dependent kinase 11B | Nucleus | protein kinase |
| chr1 | *MKNK1* | 8 | 206 | 18 | MKNK1 | MAPK interacting serine/threonine kinase 1 | Cytoplasm | protein kinase |
| chr12 | *LRRK2* | 8 | 208 | 18 | LRRK2 | leucine rich repeat kinase 2 | Cytoplasm | protein kinase |
| chr19 | *CSNK1G2* | 10 | 538 | 18 | CSNK1G2 | casein kinase 1 gamma 2 | Cytoplasm | protein kinase |
| chr11 | *BRSK2* | 8 | 514 | 18 | BRSK2 | BR serine/threonine kinase 2 | Cytoplasm | protein kinase |
| chr19 | *PRKD2* | 8 | 275 | 18 | PRKD2 | protein kinase D2 | Cytoplasm | protein kinase |
| chr17 | *GUCY2D* | 8 | 306 | 18 | GUCY2D | guanylate cyclase 2D, retinal | Plasma Membrane | kinase |
| chr19 | *LMTK3* | 8 | 265 | 18 | LMTK3 | lemur tyrosine kinase 3 | Other | protein kinase |
| chr5 | *PRKAA1* | 7 | 461 | 18 | PRKAA1 | protein kinase AMP-activated catalytic subunit alpha 1 | Cytoplasm | protein kinase |
| chr9 | *JAK2* | 7 | 197 | 17 | JAK2 | Janus kinase 2 | Cytoplasm | protein kinase |
| chr7 | *AGK* | 7 | 156 | 17 | AGK | acylglycerol kinase | Cytoplasm | kinase |
| chr13 | *FLT1* | 7 | 186 | 17 | FLT1 | fms related tyrosine kinase 1 | Plasma Membrane | protein kinase |
| chr2 | *ROCK2* | 6 | 138 | 17 | ROCK2 | Rho associated coiled-coil containing protein kinase 2 | Cytoplasm | protein kinase |
| chr5 | *CAMK4* | 8 | 165 | 17 | CAMK4 | calcium/calmodulin dependent protein kinase IV | Nucleus | protein kinase |
| chr19 | *TYK2* | 7 | 282 | 17 | TYK2 | tyrosine kinase 2 | Plasma Membrane | protein kinase |
| chrX | *RPS6KA3* | 7 | 150 | 17 | RPS6KA3 | ribosomal protein S6 kinase A3 | Cytoplasm | protein kinase |
| chr11 | *SCYL1* | 8 | 501 | 17 | SCYL1 | SCY1 like pseudokinase 1 | Cytoplasm | kinase |

| | | | | | | | | |
|---|---|---|---|---|---|---|---|---|
| chr17 | *WNK4* | 8 | 290 | 17 | WNK4 | WNK lysine deficient protein kinase 4 | Plasma Membrane | protein kinase |
| chr22 | *LIMK2* | 6 | 200 | 17 | LIMK2 | LIM domain kinase 2 | Cytoplasm | protein kinase |
| chr12 | *ERBB3* | 8 | 254 | 17 | ERBB3 | erb-b2 receptor tyrosine kinase 3 | Plasma Membrane | protein kinase |
| chr11 | *ATM* | 7 | 176 | 17 | ATM | ATM serine/threonine kinase | Nucleus | protein kinase |
| chr12 | *ACVRL1* | 8 | 622 | 17 | ACVRL1 | activin A receptor like type 1 | Plasma Membrane | protein kinase |
| chr9 | *PKN3* | 8 | 488 | 17 | PKN3 | protein kinase N3 | Nucleus | protein kinase |
| chr7 | *ADCK2* | 8 | 310 | 16 | ADCK2 | aarF domain containing kinase 2 | Cytoplasm | protein kinase |
| chr3 | *SNRK* | 7 | 176 | 16 | SNRK | SNF related kinase | Cytoplasm | protein kinase |
| chr12 | *PFKM* | 8 | 263 | 16 | PFKM | phosphofructokinase, muscle | Cytoplasm | kinase |
| chr1 | *ETNK2* | 7 | 302 | 16 | ETNK2 | ethanolamine kinase 2 | Cytoplasm | kinase |
| chr12 | *MAP3K12* | 7 | 352 | 16 | MAP3K12 | mitogen-activated protein kinase kinase kinase 12 | Cytoplasm | protein kinase |
| chr20 | *CSNK2A1* | 5 | 135 | 16 | CSNK2A1 | casein kinase 2 alpha 1 | Nucleus | protein kinase |
| chr12 | *TBK1* | 6 | 168 | 16 | TBK1 | TANK binding kinase 1 | Cytoplasm | protein kinase |
| chr1 | *EXOSC10* | 7 | 216 | 16 | EXOSC10 | exosome component 10 | Nucleus | kinase |
| chr20 | *SGK2* | 8 | 325 | 16 | SGK2 | serum/glucocorticoid regulated kinase 2 | Cytoplasm | protein kinase |
| chr15 | *BUB1B* | 5 | 173 | 16 | BUB1B | BUB1 mitotic checkpoint serine/threonine kinase B | Nucleus | protein kinase |
| chr16 | *CSNK2A2* | 7 | 151 | 16 | CSNK2A2 | casein kinase 2 alpha 2 | Cytoplasm | protein kinase |
| chr1 | *MAP3K6* | 7 | 412 | 16 | MAP3K6 | mitogen-activated protein kinase kinase kinase 6 | Other | protein kinase |
| chr17 | *CSNK1D* | 7 | 287 | 16 | CSNK1D | casein kinase 1 delta | Cytoplasm | protein kinase |
| chr12 | *MVK* | 6 | 358 | 15 | MVK | mevalonate kinase | Cytoplasm | kinase |
| chr3 | *OXSR1* | 7 | 206 | 15 | OXSR1 | oxidative stress responsive kinase 1 | Nucleus | protein kinase |
| chr10 | *PIP4K2A* | 6 | 176 | 15 | PIP4K2A | phosphatidylinositol-5-phosphate 4-kinase type 2 alpha | Cytoplasm | kinase |
| chr1 | *EFNA3* | 7 | 466 | 15 | EFNA3 | ephrin A3 | Plasma Membrane | protein kinase |
| chr16 | *MLKL* | 7 | 229 | 15 | MLKL | mixed lineage kinase domain like pseudokinase | Cytoplasm | protein kinase |
| chr10 | *PRKCQ* | 7 | 165 | 15 | PRKCQ | protein kinase C theta | Cytoplasm | protein kinase |
| chr2 | *NRP2* | 6 | 188 | 15 | NRP2 | neuropilin 2 | Plasma Membrane | kinase |
| chr3 | *GRK7* | 8 | 267 | 15 | GRK7 | G protein-coupled receptor kinase 7 | Cytoplasm | protein kinase |
| chr19 | *CKM* | 7 | 248 | 15 | CKM | creatine kinase, M-type | Cytoplasm | kinase |
| chr17 | *FN3K* | 7 | 265 | 15 | FN3K | fructosamine 3 kinase | Cytoplasm | kinase |
| chr2 | *CRIM1* | 6 | 147 | 15 | CRIM1 | cysteine rich transmembrane BMP regulator 1 | Extracellular Space | kinase |
| chr20 | *STK35* | 6 | 186 | 15 | STK35 | serine/threonine kinase 35 | Cytoplasm | protein kinase |
| chr15 | *HYKK* | 7 | 209 | 15 | HYKK | hydroxylysine kinase | Cytoplasm | kinase |
| chr6 | *AK9* | 6 | 167 | 14 | AK9 | adenylate kinase 9 | Nucleus | kinase |
| chr1 | *PFKFB2* | 7 | 268 | 14 | PFKFB2 | 6-phosphofructo-2-kinase/fructose-2,6-biphosphatase 2 | Cytoplasm | kinase |
| chr5 | *PPIP5K2* | 5 | 155 | 14 | PPIP5K2 | diphosphoinositol pentakisphosphate kinase 2 | Cytoplasm | kinase |
| chr13 | *DCLK1* | 7 | 169 | 14 | DCLK1 | doublecortin like kinase 1 | Plasma Membrane | protein kinase |
| chr7 | *PHKG1* | 7 | 332 | 14 | PHKG1 | phosphorylase kinase catalytic subunit gamma 1 | Cytoplasm | protein kinase |
| chr11 | *MAP3K11* | 7 | 533 | 14 | MAP3K11 | mitogen-activated protein kinase kinase kinase 11 | Cytoplasm | protein kinase |
| chr14 | *PRKD1* | 5 | 138 | 14 | PRKD1 | protein kinase D1 | Cytoplasm | protein kinase |
| chr19 | *TSKS* | 5 | 226 | 14 | TSKS | testis specific serine kinase substrate | Cytoplasm | kinase |
| chr6 | *ICK* | 6 | 136 | 14 | ICK | intestinal cell kinase | Cytoplasm | protein kinase |
| chr12 | *WNK1* | 5 | 117 | 14 | WNK1 | WNK lysine deficient protein kinase 1 | Cytoplasm | protein kinase |
| chr1 | *PANK4* | 8 | 373 | 14 | PANK4 | pantothenate kinase 4 | Cytoplasm | kinase |
| chr16 | *ERN2* | 6 | 176 | 14 | ERN2 | endoplasmic reticulum to nucleus signaling 2 | Cytoplasm | protein kinase |
| chr4 | *CDKL2* | 5 | 125 | 14 | CDKL2 | cyclin dependent kinase like 2 | Nucleus | protein kinase |
| chr9 | *PIP5KL1* | 6 | 437 | 13 | PIP5KL1 | phosphatidylinositol-4-phosphate 5-kinase like 1 | Plasma Membrane | kinase |
| chr2 | *VRK2* | 6 | 164 | 13 | VRK2 | VRK serine/threonine kinase 2 | Nucleus | protein kinase |
| chrX | *PFKFB1* | 6 | 255 | 13 | PFKFB1 | 6-phosphofructo-2-kinase/fructose-2,6-biphosphatase 1 | Cytoplasm | kinase |
| chr1 | *PKLR* | 6 | 372 | 13 | PKLR | pyruvate kinase L/R | Cytoplasm | kinase |
| chr2 | *MYO3B* | 6 | 116 | 13 | MYO3B | myosin IIIB | Plasma Membrane | protein kinase |
| chr5 | *MAP3K1* | 6 | 116 | 13 | MAP3K1 | mitogen-activated protein kinase kinase kinase 1 | Cytoplasm | protein kinase |
| chr1 | *NUAK2* | 6 | 308 | 13 | NUAK2 | NUAK family kinase 2 | Other | protein kinase |
| chr6 | *MAPK13* | 6 | 202 | 13 | MAPK13 | mitogen-activated protein kinase 13 | Cytoplasm | protein kinase |
| chr4 | *TBCK* | 5 | 172 | 13 | TBCK | TBC1 domain containing kinase | Cytoplasm | protein kinase |
| chr12 | *HSPB8* | 6 | 145 | 13 | HSPB8 | heat shock protein family B (small) member 8 | Cytoplasm | kinase |
| chrX | *IKBKG* | 6 | 252 | 13 | IKBKG | inhibitor of nuclear factor kappa B kinase regulatory subunit | Nucleus | kinase |
| chr1 | *CLK2* | 6 | 270 | 13 | CLK2 | CDC like kinase 2 | Nucleus | protein kinase |
| chr3 | *PIK3R4* | 5 | 174 | 13 | PIK3R4 | phosphoinositide-3-kinase regulatory subunit 4 | Cytoplasm | kinase |
| chr12 | *ETNK1* | 5 | 118 | 13 | ETNK1 | ethanolamine kinase 1 | Cytoplasm | kinase |
| chr1 | *NUCKS1* | 5 | 195 | 12 | NUCKS1 | nuclear casein kinase and cyclin dependent kinase substrate 1 | Nucleus | protein kinase |
| chr1 | *PRKAA2* | 5 | 112 | 12 | PRKAA2 | protein kinase AMP-activated catalytic subunit alpha 2 | Cytoplasm | protein kinase |
| chr10 | *PANK1* | 5 | 135 | 12 | PANK1 | pantothenate kinase 1 | Cytoplasm | kinase |
| chr17 | *PDK2* | 5 | 216 | 12 | PDK2 | pyruvate dehydrogenase kinase 2 | Cytoplasm | protein kinase |
| chr19 | *DYRK1B* | 6 | 231 | 12 | DYRK1B | dual specificity tyrosine phosphorylation regulated kinase 1B | Nucleus | protein kinase |
| chrX | *CDK16* | 5 | 232 | 12 | CDK16 | cyclin dependent kinase 16 | Cytoplasm | protein kinase |
| chr7 | *EPHA1* | 5 | 238 | 12 | EPHA1 | EPH receptor A1 | Plasma Membrane | protein kinase |
| chr9 | *TEK* | 4 | 97 | 12 | TEK | TEK receptor tyrosine kinase | Plasma Membrane | protein kinase |
| chr17 | *TNK1* | 6 | 197 | 12 | TNK1 | tyrosine kinase non receptor 1 | Cytoplasm | protein kinase |
| chr5 | *FGFR4* | 6 | 296 | 12 | FGFR4 | fibroblast growth factor receptor 4 | Plasma Membrane | protein kinase |
| chr3 | *DCLK3* | 5 | 158 | 12 | DCLK3 | doublecortin like kinase 3 | Cytoplasm | protein kinase |
| chr13 | *NEK5* | 5 | 108 | 12 | NEK5 | NIMA related kinase 5 | Other | protein kinase |
| chr1 | *UHMK1* | 4 | 136 | 12 | UHMK1 | U2AF homology motif kinase 1 | Nucleus | protein kinase |
| chr16 | *MAPK3* | 5 | 220 | 12 | MAPK3 | mitogen-activated protein kinase 3 | Cytoplasm | protein kinase |
| chr21 | *RIPK4* | 5 | 246 | 12 | RIPK4 | receptor interacting serine/threonine kinase 4 | Nucleus | protein kinase |
| chr7 | *PRKAR2B* | 5 | 106 | 12 | PRKAR2B | protein kinase cAMP-dependent type II regulatory subunit beta | Cytoplasm | protein kinase |
| chr12 | *IRAK3* | 4 | 116 | 12 | IRAK3 | interleukin 1 receptor associated kinase 3 | Cytoplasm | protein kinase |
| chr13 | *CAB39L* | 5 | 111 | 12 | CAB39L | calcium binding protein 39 like | Cytoplasm | kinase |
| chr2 | *PKDCC* | 5 | 264 | 12 | PKDCC | protein kinase domain containing, cytoplasmic | Cytoplasm | protein kinase |
| chr11 | *CHEK1* | 5 | 131 | 12 | CHEK1 | checkpoint kinase 1 | Nucleus | protein kinase |
| chrX | *DGKK* | 6 | 107 | 12 | DGKK | diacylglycerol kinase kappa | Cytoplasm | kinase |
| chr12 | *DGKA* | 6 | 351 | 12 | DGKA | diacylglycerol kinase alpha | Cytoplasm | kinase |
| chr19 | *MATK* | 6 | 216 | 12 | MATK | megakaryocyte-associated tyrosine kinase | Cytoplasm | protein kinase |
| chr19 | *FZR1* | 6 | 277 | 12 | FZR1 | fizzy and cell division cycle 20 related 1 | Nucleus | kinase |
| chr10 | *ALDH18A1* | 5 | 105 | 12 | ALDH18A1 | aldehyde dehydrogenase 18 family member A1 | Cytoplasm | kinase |
| chr1 | *SCYL3* | 4 | 166 | 12 | SCYL3 | SCY1 like pseudokinase 3 | Cytoplasm | kinase |
| chr6 | *MAP3K4* | 4 | 105 | 12 | MAP3K4 | mitogen-activated protein kinase kinase kinase 4 | Cytoplasm | protein kinase |
| chr1 | *PRKACB* | 5 | 142 | 12 | PRKACB | protein kinase cAMP-activated catalytic subunit beta | Cytoplasm | protein kinase |
| chr6 | *FRK* | 5 | 149 | 11 | FRK | fyn related Src family tyrosine kinase | Nucleus | protein kinase |
| chr20 | *PANK2* | 5 | 120 | 11 | PANK2 | pantothenate kinase 2 | Cytoplasm | kinase |
| chr1 | *HIPK1* | 6 | 165 | 11 | HIPK1 | homeodomain interacting protein kinase 1 | Nucleus | protein kinase |
| chrX | *PGK1* | 4 | 83 | 11 | PGK1 | phosphoglycerate kinase 1 | Cytoplasm | kinase |
| chr5 | *NME5* | 4 | 112 | 11 | NME5 | NME/NM23 family member 5 | Other | kinase |
| chr15 | *ADPGK* | 5 | 86 | 11 | ADPGK | ADP dependent glucokinase | Plasma Membrane | kinase |

| | | | | | | | | |
|---|---|---|---|---|---|---|---|---|
| chr1 | *UCK2* | 6 | 147 | 11 | UCK2 | uridine-cytidine kinase 2 | Cytoplasm | kinase |
| chr13 | *CDADC1* | 5 | 135 | 11 | CDADC1 | cytidine and dCMP deaminase domain containing 1 | Other | kinase |
| chrX | *DLG3* | 4 | 123 | 11 | DLG3 | discs large MAGUK scaffold protein 3 | Plasma Membrane | kinase |
| chr4 | *CAMK2D* | 4 | 92 | 11 | CAMK2D | calcium/calmodulin dependent protein kinase II delta | Cytoplasm | protein kinase |
| chr11 | *CLP1* | 4 | 164 | 11 | CLP1 | cleavage and polyadenylation factor I subunit 1 | Nucleus | kinase |
| chr22 | *CHEK2* | 4 | 100 | 11 | CHEK2 | checkpoint kinase 2 | Nucleus | protein kinase |
| chr14 | *RPS6KL1* | 5 | 212 | 11 | RPS6KL1 | ribosomal protein S6 kinase like 1 | Other | protein kinase |
| chr12 | *PIK3C2G* | 4 | 104 | 11 | PIK3C2G | phosphatidylinositol-4-phosphate 3-kinase catalytic subunit type 2 gamma | Cytoplasm | kinase |
| chr3 | *CAMKV* | 5 | 210 | 11 | CAMKV | CaM kinase like vesicle associated | Plasma Membrane | kinase |
| chr4 | *TEC* | 4 | 97 | 11 | TEC | tec protein tyrosine kinase | Cytoplasm | protein kinase |
| chr6 | *PRPF4B* | 4 | 92 | 10 | PRPF4B | pre-mRNA processing factor 4B | Nucleus | protein kinase |
| chr15 | *LTK* | 5 | 208 | 10 | LTK | leukocyte receptor tyrosine kinase | Plasma Membrane | protein kinase |
| chr2 | *STK36* | 4 | 111 | 10 | STK36 | serine/threonine kinase 36 | Cytoplasm | protein kinase |
| chr12 | *AMHR2* | 5 | 256 | 10 | AMHR2 | anti-Mullerian hormone receptor type 2 | Plasma Membrane | kinase |
| chr7 | *DBF4* | 3 | 89 | 10 | DBF4 | DBF4 zinc finger | Nucleus | kinase |
| chr17 | *TK1* | 5 | 179 | 10 | TK1 | thymidine kinase 1 | Cytoplasm | kinase |
| chr5 | *CSNK1A1* | 4 | 105 | 10 | CSNK1A1 | casein kinase 1 alpha 1 | Cytoplasm | protein kinase |
| chr8 | *PBK* | 3 | 125 | 10 | PBK | PDZ binding kinase | Cytoplasm | protein kinase |
| chr5 | *PIK3R1* | 5 | 109 | 10 | PIK3R1 | phosphoinositide-3-kinase regulatory subunit 1 | Cytoplasm | kinase |
| chr4 | *ALPK1* | 4 | 98 | 10 | ALPK1 | alpha kinase 1 | Cytoplasm | protein kinase |
| chr8 | *C8orf44-SGK3* | 4 | 95 | 10 | 8orf44-SGK3/SGK | serum/glucocorticoid regulated kinase family member 3 | Cytoplasm | protein kinase |
| chr3 | *ATRIP* | 5 | 112 | 10 | ATRIP | ATR interacting protein | Nucleus | kinase |
| chr5 | *CLK4* | 4 | 172 | 10 | CLK4 | CDC like kinase 4 | Nucleus | protein kinase |
| chr7 | *STK17A* | 4 | 112 | 10 | STK17A | serine/threonine kinase 17a | Nucleus | protein kinase |
| chr16 | *PKMYT1* | 5 | 257 | 10 | PKMYT1 | protein kinase, membrane associated tyrosine/threonine 1 | Cytoplasm | protein kinase |
| chr19 | *STK11* | 4 | 162 | 10 | STK11 | serine/threonine kinase 11 | Cytoplasm | protein kinase |
| chr13 | *DGKH* | 4 | 83 | 10 | DGKH | diacylglycerol kinase eta | Cytoplasm | kinase |
| chr8 | *POMK* | 4 | 144 | 10 | POMK | protein O-mannose kinase | Cytoplasm | protein kinase |
| chr17 | *AURKB* | 4 | 192 | 9 | AURKB | aurora kinase B | Nucleus | protein kinase |
| chr4 | *EPHA5* | 4 | 108 | 9 | EPHA5 | EPH receptor A5 | Plasma Membrane | protein kinase |
| chr2 | *KHK* | 4 | 157 | 9 | KHK | ketohexokinase | Cytoplasm | kinase |
| chr2 | *DGUOK* | 5 | 110 | 9 | DGUOK | deoxyguanosine kinase | Cytoplasm | kinase |
| chr2 | *EIF2AK3* | 4 | 81 | 9 | EIF2AK3 | eukaryotic translation initiation factor 2 alpha kinase 3 | Cytoplasm | kinase |
| chr10 | *SLK* | 3 | 83 | 9 | SLK | STE20 like kinase | Nucleus | protein kinase |
| chr6 | *RIOK1* | 4 | 105 | 9 | RIOK1 | RIO kinase 1 | Cytoplasm | protein kinase |
| chr14 | *MAP3K9* | 3 | 119 | 9 | MAP3K9 | mitogen-activated protein kinase kinase kinase 9 | Cytoplasm | protein kinase |
| chr5 | *ITK* | 4 | 96 | 9 | ITK | IL2 inducible T cell kinase | Cytoplasm | protein kinase |
| chr15 | *ULK3* | 4 | 171 | 9 | ULK3 | unc-51 like kinase 3 | Cytoplasm | protein kinase |
| chr2 | *NAGK* | 3 | 114 | 9 | NAGK | N-acetylglucosamine kinase | Cytoplasm | kinase |
| chr1 | *PDIK1L* | 4 | 99 | 8 | PDIK1L | PDLIM1 interacting kinase 1 like | Nucleus | protein kinase |
| chr2 | *ACVR1* | 3 | 89 | 8 | ACVR1 | activin A receptor type 1 | Plasma Membrane | protein kinase |
| chr1 | *CAMK1G* | 4 | 150 | 8 | CAMK1G | calcium/calmodulin dependent protein kinase IG | Cytoplasm | protein kinase |
| chr4 | *DCLK2* | 4 | 96 | 8 | DCLK2 | doublecortin like kinase 2 | Cytoplasm | protein kinase |
| chr13 | *EFNB2* | 4 | 99 | 8 | EFNB2 | ephrin B2 | Plasma Membrane | protein kinase |
| chr11 | *WEE1* | 4 | 108 | 8 | WEE1 | WEE1 G2 checkpoint kinase | Nucleus | protein kinase |
| chr8 | *NRBP2* | 4 | 183 | 8 | NRBP2 | nuclear receptor binding protein 2 | Cytoplasm | kinase |
| chr2 | *STK39* | 4 | 66 | 8 | STK39 | serine/threonine kinase 39 | Nucleus | protein kinase |
| chr13 | *FLT3* | 4 | 86 | 8 | FLT3 | fms related tyrosine kinase 3 | Plasma Membrane | protein kinase |
| chr5 | *PANK3* | 3 | 105 | 8 | PANK3 | pantothenate kinase 3 | Cytoplasm | kinase |
| chr11 | *ANKK1* | 4 | 141 | 8 | ANKK1 | ankyrin repeat and kinase domain containing 1 | Cytoplasm | protein kinase |
| chr10 | *PSTK* | 4 | 86 | 8 | PSTK | phosphoseryl-tRNA kinase | Cytoplasm | kinase |
| chr1 | *PMVK* | 4 | 216 | 8 | PMVK | phosphomevalonate kinase | Cytoplasm | kinase |
| chr14 | *NEK9* | 3 | 72 | 8 | NEK9 | NIMA related kinase 9 | Nucleus | protein kinase |
| chr2 | *PRKD3* | 3 | 63 | 8 | PRKD3 | protein kinase D3 | Nucleus | protein kinase |
| chr6 | *CDKN1A* | 4 | 146 | 8 | CDKN1A | cyclin dependent kinase inhibitor 1A | Nucleus | kinase |
| chr15 | *FES* | 4 | 396 | 8 | FES | FES proto-oncogene, tyrosine kinase | Cytoplasm | protein kinase |
| chr12 | *IRAK4* | 3 | 106 | 8 | IRAK4 | interleukin 1 receptor associated kinase 4 | Cytoplasm | protein kinase |
| chr10 | *CHUK* | 3 | 82 | 8 | CHUK | component of inhibitor of nuclear factor kappa B kinase complex | Cytoplasm | protein kinase |
| chr7 | *EPHB6* | 4 | 147 | 8 | EPHB6 | EPH receptor B6 | Plasma Membrane | protein kinase |
| chr2 | *MAP3K19* | 3 | 67 | 8 | MAP3K19 | mitogen-activated protein kinase kinase kinase 19 | Other | protein kinase |
| chr2 | *EIF2AK2* | 3 | 68 | 8 | EIF2AK2 | eukaryotic translation initiation factor 2 alpha kinase 2 | Cytoplasm | kinase |
| chrX | *PIM2* | 3 | 128 | 8 | PIM2 | Pim-2 proto-oncogene, serine/threonine kinase | Nucleus | protein kinase |
| chr1 | *TSSK3* | 4 | 87 | 8 | TSSK3 | testis specific serine kinase 3 | Other | protein kinase |
| chr14 | *CCNK* | 3 | 116 | 7 | CCNK | cyclin K | Nucleus | protein kinase |
| chr5 | *CSNK1G3* | 3 | 70 | 7 | CSNK1G3 | casein kinase 1 gamma 3 | Cytoplasm | protein kinase |
| chr17 | *MMD* | 4 | 99 | 7 | MMD | monocyte to macrophage differentiation associated | Plasma Membrane | kinase |
| chrX | *PRPS2* | 3 | 71 | 7 | PRPS2 | phosphoribosyl pyrophosphate synthetase 2 | Cytoplasm | kinase |
| chr17 | *GALK1* | 4 | 198 | 7 | GALK1 | galactokinase 1 | Cytoplasm | kinase |
| chr19 | *MAP2K7* | 4 | 181 | 7 | MAP2K7 | mitogen-activated protein kinase kinase 7 | Cytoplasm | protein kinase |
| chr9 | *CDK20* | 3 | 154 | 7 | CDK20 | cyclin dependent kinase 20 | Nucleus | protein kinase |
| chr20 | *UCKL1* | 4 | 306 | 7 | UCKL1 | uridine-cytidine kinase 1 like 1 | Cytoplasm | kinase |
| chr1 | *GUK1* | 4 | 229 | 7 | GUK1 | guanylate kinase 1 | Cytoplasm | kinase |
| chr18 | *PIK3C3* | 3 | 55 | 7 | PIK3C3 | phosphatidylinositol 3-kinase catalytic subunit type 3 | Cytoplasm | kinase |
| chr6 | *EPHA7* | 3 | 55 | 7 | EPHA7 | EPH receptor A7 | Plasma Membrane | protein kinase |
| chr1 | *PINK1* | 3 | 134 | 7 | PINK1 | PTEN induced kinase 1 | Cytoplasm | protein kinase |
| chr5 | *CCNB1* | 3 | 74 | 7 | CCNB1 | cyclin B1 | Cytoplasm | kinase |
| chr1 | *PLK3* | 3 | 123 | 7 | PLK3 | polo like kinase 3 | Nucleus | protein kinase |
| chr21 | *PFKL* | 3 | 152 | 7 | PFKL | phosphofructokinase, liver type | Cytoplasm | kinase |
| chr16 | *PLK1* | 3 | 153 | 7 | PLK1 | polo like kinase 1 | Nucleus | protein kinase |
| chr22 | *CHKB* | 3 | 86 | 7 | CHKB | choline kinase beta | Cytoplasm | kinase |
| chr12 | *STYK1* | 3 | 62 | 7 | STYK1 | serine/threonine/tyrosine kinase 1 | Cytoplasm | protein kinase |
| chr17 | *DGKE* | 3 | 90 | 7 | DGKE | diacylglycerol kinase epsilon | Cytoplasm | kinase |
| chr2 | *PDK1* | 3 | 65 | 7 | PDK1 | pyruvate dehydrogenase kinase 1 | Cytoplasm | protein kinase |
| chr11 | *DAK* | 3 | 181 | 6 | TKFC | triokinase and FMN cyclase | Cytoplasm | kinase |
| chrX | *PNCK* | 3 | 134 | 6 | PNCK | pregnancy up-regulated nonubiquitous CaM kinase | Other | protein kinase |
| chr9 | *NMRK1* | 3 | 51 | 6 | NMRK1 | nicotinamide riboside kinase 1 | Cytoplasm | kinase |
| chr22 | *CRKL* | 3 | 68 | 6 | CRKL | CRK like proto-oncogene, adaptor protein | Cytoplasm | protein kinase |
| chr19 | *DAPK3* | 3 | 176 | 6 | DAPK3 | death associated protein kinase 3 | Cytoplasm | protein kinase |
| chr2 | *ACVR1C* | 3 | 85 | 6 | ACVR1C | activin A receptor type 1C | Plasma Membrane | protein kinase |
| chr16 | *CDK10* | 3 | 132 | 6 | CDK10 | cyclin dependent kinase 10 | Nucleus | protein kinase |
| chr18 | *RIOK3* | 3 | 76 | 6 | RIOK3 | RIO kinase 3 | Cytoplasm | protein kinase |
| chr2 | *STK25* | 3 | 131 | 6 | STK25 | serine/threonine kinase 25 | Cytoplasm | protein kinase |

| | | | | | | | | |
|---|---|---|---|---|---|---|---|---|
| chr20 | *PAK7* | 3 | 67 | 6 | PAK5 | p21 (RAC1) activated kinase 5 | Nucleus | protein kinase |
| chr9 | *UCK1* | 3 | 203 | 6 | UCK1 | uridine-cytidine kinase 1 | Cytoplasm | kinase |
| chrX | *MPP1* | 2 | 48 | 6 | MPP1 | membrane palmitoylated protein 1 | Plasma Membrane | kinase |
| chrX | *IRAK1* | 3 | 171 | 6 | IRAK1 | interleukin 1 receptor associated kinase 1 | Plasma Membrane | protein kinase |
| chr20 | *PCK1* | 3 | 172 | 6 | PCK1 | phosphoenolpyruvate carboxykinase 1 | Cytoplasm | kinase |
| chr20 | *AURKA* | 2 | 95 | 6 | AURKA | aurora kinase A | Nucleus | protein kinase |
| chr9 | *MUSK* | 2 | 81 | 6 | MUSK | muscle associated receptor tyrosine kinase | Plasma Membrane | protein kinase |
| chr19 | *MKNK2* | 3 | 162 | 6 | MKNK2 | MAPK interacting serine/threonine kinase 2 | Cytoplasm | protein kinase |
| chr20 | *PTK6* | 3 | 264 | 6 | PTK6 | protein tyrosine kinase 6 | Cytoplasm | protein kinase |
| chr12 | *XRCC6BP1* | 3 | 88 | 6 | ATP23 | ATP23 metallopeptidase and ATP synthase assembly factor | Plasma Membrane | protein kinase |
| chr16 | *BCKDK* | 3 | 119 | 6 | BCKDK | branched chain keto acid dehydrogenase kinase | Cytoplasm | protein kinase |
| chr7 | *PIK3CG* | 3 | 58 | 6 | PIK3CG | phosphatidylinositol-4,5-bisphosphate 3-kinase catalytic subunit gamma | Cytoplasm | kinase |
| chr12 | *PRKAB1* | 3 | 100 | 5 | PRKAB1 | protein kinase AMP-activated non-catalytic subunit beta 1 | Nucleus | protein kinase |
| chr19 | *NMRK2* | 3 | 98 | 5 | NMRK2 | nicotinamide riboside kinase 2 | Plasma Membrane | kinase |
| chr16 | *NME4* | 2 | 97 | 5 | NME4 | NME/NM23 nucleoside diphosphate kinase 4 | Cytoplasm | kinase |
| chr1 | *NEK2* | 2 | 58 | 5 | NEK2 | NIMA related kinase 2 | Cytoplasm | protein kinase |
| chrX | *GUCY2F* | 2 | 44 | 5 | GUCY2F | guanylate cyclase 2F, retinal | Plasma Membrane | kinase |
| chr5 | *NADK2* | 2 | 127 | 5 | NADK2 | NAD kinase 2, mitochondrial | Cytoplasm | kinase |
| chr11 | *RPS6KA4* | 3 | 136 | 5 | RPS6KA4 | ribosomal protein S6 kinase A4 | Cytoplasm | protein kinase |
| chr11 | *TMEM134* | 2 | 141 | 5 | TMEM134 | transmembrane protein 134 | Other | kinase |
| chr2 | *EPHA4* | 3 | 38 | 5 | EPHA4 | EPH receptor A4 | Plasma Membrane | protein kinase |
| chr4 | *PLK4* | 2 | 62 | 5 | PLK4 | polo like kinase 4 | Cytoplasm | protein kinase |
| chr12 | *DYRK4* | 2 | 60 | 5 | DYRK4 | dual specificity tyrosine phosphorylation regulated kinase 4 | Nucleus | protein kinase |
| chr4 | *PI4K2B* | 2 | 37 | 5 | PI4K2B | phosphatidylinositol 4-kinase type 2 beta | Cytoplasm | kinase |
| chrX | *PHKA1* | 2 | 44 | 5 | PHKA1 | phosphorylase kinase regulatory subunit alpha 1 | Cytoplasm | protein kinase |
| chr2 | *CLK1* | 2 | 44 | 5 | CLK1 | CDC like kinase 1 | Nucleus | protein kinase |
| chrX | *BTK* | 2 | 41 | 5 | BTK | Bruton tyrosine kinase | Cytoplasm | protein kinase |
| chr12 | *CCT2* | 2 | 39 | 5 | CCT2 | chaperonin containing TCP1 subunit 2 | Cytoplasm | kinase |
| chr17 | *MAPK7* | 2 | 105 | 5 | MAPK7 | mitogen-activated protein kinase 7 | Cytoplasm | protein kinase |
| chrX | *BMX* | 2 | 43 | 5 | BMX | BMX non-receptor tyrosine kinase | Cytoplasm | protein kinase |
| chr19 | *HIPK4* | 3 | 105 | 5 | HIPK4 | homeodomain interacting protein kinase 4 | Cytoplasm | protein kinase |
| chr17 | *NEK8* | 2 | 73 | 4 | NEK8 | NIMA related kinase 8 | Nucleus | protein kinase |
| chr7 | *CDK5* | 2 | 66 | 4 | CDK5 | cyclin dependent kinase 5 | Nucleus | protein kinase |
| chr14 | *MPP5* | 2 | 38 | 4 | MPP5 | membrane palmitoylated protein 5 | Plasma Membrane | kinase |
| chr8 | *MAPK15* | 2 | 78 | 4 | MAPK15 | mitogen-activated protein kinase 15 | Cytoplasm | protein kinase |
| chr2 | *BUB1* | 2 | 51 | 4 | BUB1 | BUB1 mitotic checkpoint serine/threonine kinase | Nucleus | protein kinase |
| chr19 | *PNKP* | 2 | 60 | 4 | PNKP | polynucleotide kinase 3'-phosphatase | Nucleus | kinase |
| chr15 | *CSNK1G1* | 2 | 49 | 4 | CSNK1G1 | casein kinase 1 gamma 1 | Cytoplasm | protein kinase |
| chr1 | *PRKAB2* | 2 | 52 | 4 | PRKAB2 | protein kinase AMP-activated non-catalytic subunit beta 2 | Cytoplasm | protein kinase |
| chr6 | *ROS1* | 2 | 33 | 4 | ROS1 | ROS proto-oncogene 1, receptor tyrosine kinase | Plasma Membrane | protein kinase |
| chr17 | *EFNB3* | 2 | 99 | 4 | EFNB3 | ephrin B3 | Plasma Membrane | protein kinase |
| chr20 | *TRIB3* | 2 | 71 | 4 | TRIB3 | tribbles pseudokinase 3 | Nucleus | kinase |
| chr3 | *NPRL2* | 2 | 88 | 4 | NPRL2 | NPR2 like, GATOR1 complex subunit | Cytoplasm | kinase |
| chr20 | *SRMS* | 2 | 120 | 4 | SRMS | src-related kinase lacking C-terminal regulatory tyrosine and N-terminal myristylation sites | Cytoplasm | protein kinase |
| chr2 | *DTYMK* | 2 | 62 | 4 | DTYMK | deoxythymidylate kinase | Cytoplasm | kinase |
| chr19 | *GSK3A* | 2 | 72 | 4 | GSK3A | glycogen synthase kinase 3 alpha | Nucleus | protein kinase |
| chr3 | *TRAT1* | 2 | 35 | 4 | TRAT1 | T cell receptor associated transmembrane adaptor 1 | Plasma Membrane | kinase |
| chr2 | *MPP4* | 2 | 28 | 4 | MPP4 | membrane palmitoylated protein 4 | Cytoplasm | kinase |
| chr21 | *SIK1* | 2 | 109 | 4 | SIK1/SIK1B | salt inducible kinase 1 | Nucleus | protein kinase |
| chr15 | *CKMT1B* | 2 | 39 | 4 | CKMT1A/CKMT1E | creatine kinase, mitochondrial 1B | Cytoplasm | kinase |
| chr17 | *COASY* | 2 | 76 | 3 | COASY | Coenzyme A synthase | Cytoplasm | protein kinase |
| chr12 | *DYRK2* | 2 | 48 | 3 | DYRK2 | dual specificity tyrosine phosphorylation regulated kinase 2 | Cytoplasm | protein kinase |
| chr19 | *DMPK* | 2 | 73 | 3 | DMPK | DM1 protein kinase | Cytoplasm | protein kinase |
| chr4 | *PRKG2* | 1 | 27 | 3 | PRKG2 | protein kinase cGMP-dependent 2 | Cytoplasm | protein kinase |
| chr17 | *SPHK1* | 2 | 64 | 3 | SPHK1 | sphingosine kinase 1 | Cytoplasm | kinase |
| chrX | *PAK3* | 2 | 49 | 3 | PAK3 | p21 (RAC1) activated kinase 3 | Cytoplasm | protein kinase |
| chr7 | *FASTK* | 2 | 100 | 3 | FASTK | Fas activated serine/threonine kinase | Cytoplasm | protein kinase |
| chr2 | *NRBP1* | 1 | 25 | 3 | NRBP1 | nuclear receptor binding protein 1 | Nucleus | kinase |
| chr2 | *PIKFYVE* | 1 | 22 | 2 | PIKFYVE | phosphoinositide kinase, FYVE-type zinc finger containing | Cytoplasm | kinase |
| chr8 | *TRIB1* | 1 | 32 | 2 | TRIB1 | tribbles pseudokinase 1 | Cytoplasm | kinase |
| chr9 | *AK1* | 1 | 61 | 2 | AK1 | adenylate kinase 1 | Cytoplasm | kinase |
| chr14 | *GOLGA5* | 1 | 17 | 2 | GOLGA5 | golgin A5 | Cytoplasm | kinase |
| chr4 | *DGKQ* | 1 | 40 | 2 | DGKQ | diacylglycerol kinase theta | Cytoplasm | kinase |
| chr4 | *SRP72* | 1 | 24 | 2 | SRP72 | signal recognition particle 72 | Nucleus | kinase |
| chr6 | *RSPO3* | 1 | 34 | 2 | RSPO3 | R-spondin 3 | Extracellular Space | kinase |
| chr4 | *KDR* | 1 | 29 | 2 | KDR | kinase insert domain receptor | Plasma Membrane | protein kinase |
| chr9 | *CDK9* | 1 | 83 | 2 | CDK9 | cyclin dependent kinase 9 | Nucleus | protein kinase |
| chr22 | *TRMT2A* | 1 | 41 | 2 | TRMT2A | tRNA methyltransferase 2 homolog A | Other | kinase |
| chr14 | *CKB* | 1 | 44 | 2 | CKB | creatine kinase B | Cytoplasm | kinase |
| chr4 | *FGFR3* | 1 | 41 | 2 | FGFR3 | fibroblast growth factor receptor 3 | Plasma Membrane | protein kinase |
| chr19 | *AURKC* | 1 | 43 | 2 | AURKC | aurora kinase C | Nucleus | protein kinase |

Supplementary Table 8. Core signaling pathways with which cisplatin damaged genes are associated.

| Ingenuity Canonical Pathways | -log(p-value) | Ratio | z-score | Cisplatin damaged genes involved in |
|---|---|---|---|---|
| Sperm Motility | 2.20E+01 | 6.85E-01 | -8.185 | *AATK,ABHD3,ABL1,ABL2,ADCY10,ALK,ATP1A4,AXL,BLK,CACNA1G,CACNA1H,CAMK4,CATSPER1,CLK2,CLK3,CNGA3,CNGB1,CSF1R,CSF2RA,CSK,DDR2,DSTYK,DYRK3,EGFR,EPHA1,EPHA2,EPHA3,EPHA6,EPHA8,EPHB1,EPHB2,EPHB4,ERBB2,ERBB3,ERBB4,FER,FGFR1,FGFR2,FGFR4,FGR,FLT1,FLT4,FYN,GNAS,GUCY1A2,HCK,IGF1R,INSR,ITPR1,ITPR2,ITPR3,JAK1,JAK2,JAK3,LCK,LMTK2,LMTK3,MAP2K1,MAP2K3,MAP2K4,MAP2K5,MAP2K6,MAP3K11,MERTK,MET,MRAS,MST1R,NPR1,NTRK1,NTRK2,NTRK3,PDE1A,PDE1B,PDE1C,PDE2A,PDE4A,PDE4B,PDE4C,PDE4D,PDGFRB,PEAK1,PLA2G10,PLA2G12B,PLA2G2C,PLA2G2F,PLA2G4A,PLA2G4C,PLA2G4D,PLA2G4E,PLA2G4F,PLA2G5,PLA2G6,PLA2R1,PLAAT3,PLAAT4,PLAAT5,PLB1,PLCB1,PLCB2,PLCB3,PLCD3,PLCE1,PLCG1,PLCG2,PLCH1,PLCH2,PLCL1,PLCL2,PNPLA3,PNPLA8,PRKACA,PRKAG2,PRKAR1B,PRKAR2A,PRKCA,PRKCB,PRKCD,PRKCE,PRKCG,PRKCH,PRKCI,PRKCQ,PRKCZ,PRKD1,PRKG1,PTK2,PTK2B,PTK7,RET,ROR1,ROR2,RYK,SLC12A2,SLC9A5,SRC,SYK,TEK,TIE1,TNK1,TNK2,TYK2,TYRO3,YES1,ZAN,ZAP70,ZP3* |
| Synaptic Long Term Depression | 1.82E+01 | 6.78E-01 | -9.054 | *ABHD3,CACNA1A,CACNA1B,CACNA1C,CACNA1D,CACNA1E,CACNA1F,CACNA1G,CACNA1H,CACNA1I,CACNA1S,CACNA2D1,CACNA2D2,CACNA2D3,CACNA2D4,CACNB1,CACNB2,CACNB4,CACNG2,CACNG3,CACNG4,CACNG5,CACNG6,CACNG7,CACNG8,CRHR1,GNA11,GNA12,GNA14,GNAI1,GNAI2,GNAI3,GNAL,GNAO1,GNAQ,GNAS,GRIA1,GRIA2,GRIA4,GRID1,GRID2,GRM1,GRM3,GRM4,GRM5,GRM7,GRM8,GUCY1A2,GUCY2D,IGF1R,ITPR1,ITPR2,ITPR3,KRAS,MAP2K1,MAP2K2,MAPK1,MRAS,NOS1,NOS2,NOS3,NPR1,NPR3,PLA2G10,PLA2G12B,PLA2G2C,PLA2G2F,PLA2G4A,PLA2G4C,PLA2G4D,PLA2G4E,PLA2G4F,PLA2G5,PLA2G6,PLA2R1,PLAAT3,PLAAT4,PLAAT5,PLB1,PLCB1,PLCB2,PLCB3,PLCD3,PLCE1,PLCG1,PLCG2,PLCH1,PLCH2,PLCL1,PLCL2,PNPLA3,PNPLA8,PPM1L,PPP2CB,PPP2R1B,PPP2R2A,PPP2R2B,PPP2R2C,PPP2R3A,PPP2R5B,PPP2R5C,PPP2R5D,PPP2R5E,PRKCA,PRKCB,PRKCD,PRKCE,PRKCG,PRKCH,PRKCI,PRKCQ,PRKCZ,PRKD1,PRKG1,PTPA,RAF1,RALA,RALB,RAP1A,RAP1B,RRAS2,RYR1,RYR2,RYR3* |
| CREB Signaling in Neurons | 1.75E+01 | 6.54E-01 | -9.592 | *ADCY1,ADCY10,ADCY2,ADCY3,ADCY5,ADCY6,ADCY7,ADCY8,ADCY9,AKT2,AKT3,CACNA1A,CACNA1B,CACNA1C,CACNA1D,CACNA1E,CACNA1F,CACNA1G,CACNA1H,CACNA1I,CACNA1S,CACNA2D1,CACNA2D2,CACNA2D3,CACNA2D4,CACNB1,CACNB2,CACNB4,CACNG2,CACNG3,CACNG4,CACNG5,CACNG6,CACNG7,CACNG8,CAMK2A,CAMK2B,CAMK2G,CAMK4,CREB3L3,CREB5,CREBBP,EP300,GNA11,GNA12,GNA14,GNAI1,GNAI2,GNAI3,GNAL,GNAO1,GNAQ,GNAS,GNB1,GNB1L,GNB5,GNG12,GNG2,GNG4,GNG7,GRB2,GRIA1,GRIA2,GRIA4,GRID1,GRID2,GRIK3,GRIK4,GRIK5,GRIN1,GRIN2A,GRIN2B,GRIN2C,GRIN2D,GRM1,GRM3,GRM4,GRM5,GRM7,GRM8,GTF2B,ITPR1,ITPR2,ITPR3,KRAS,MAP2K1,MAP2K2,MAPK1,MRAS,PIK3C2A,PIK3C2B,PIK3CB,PIK3CD,PIK3R2,PIK3R3,PIK3R4,PIK3R5,PIK3R6,PLCB1,PLCB2,PLCB3,PLCD3,PLCE1,PLCG1,PLCG2,PLCH1,PLCH2,PLCL1,PLCL2,POLR2A,POLR2F,POLR2J2/POLR2J3,PRKACA,PRKAG2,PRKAR1B,PRKAR2A,PRKCA,PRKCB,PRKCD,PRKCE,PRKCG,PRKCH,PRKCI,PRKCQ,PRKCZ,PRKD1,RAF1,RALA,RALB,RAP1A,RAP1B,RPS6KA1,RRAS2,SOS1* |
| Role of NFAT in Cardiac Hypertrophy | 1.75E+01 | 6.49E-01 | -9.928 | *ADCY1,ADCY10,ADCY2,ADCY3,ADCY5,ADCY6,ADCY7,ADCY8,ADCY9,AKT2,AKT3,CABIN1,CACNA1A,CACNA1B,CACNA1C,CACNA1D,CACNA1E,CACNA1F,CACNA1G,CACNA1H,CACNA1I,CACNA1S,CACNA2D1,CACNA2D2,CACNA2D3,CACNA2D4,CACNB1,CACNB2,CACNB4,CACNG2,CACNG3,CACNG4,CACNG5,CACNG6,CACNG7,CACNG8,CAMK1D,CAMK2A,CAMK2B,CAMK2G,CAMK4,CHP1,EP300,GATA4,GNAI1,GNAI2,GNAI3,GNAQ,GNAS,GNB1,GNB1L,GNB5,GNG12,GNG2,GNG4,GNG7,GRB2,GSK3B,HDAC1,HDAC11,HDAC4,HDAC5,HDAC6,HDAC7,HDAC9,IGF1R,IL6ST,ITPR1,ITPR2,ITPR3,KRAS,MAP2K1,MAP2K2,MAP2K3,MAP2K4,MAP2K6,MAP3K1,MAPK1,MAPK10,MAPK13,MAPK14,MAPK8,MAPK9,MEF2A,MEF2D,MRAS,PIK3C2A,PIK3C2B,PIK3CB,PIK3CD,PIK3R2,PIK3R3,PIK3R4,PIK3R5,PIK3R6,PLCB1,PLCB2,PLCB3,PLCD3,PLCE1,PLCG1,PLCG2,PLCH1,PLCH2,PLCL1,PLCL2,PPP3CB,PPP3CC,PRKACA,PRKAG2,PRKAR1B,PRKAR2A,PRKCA,PRKCB,PRKCD,PRKCE,PRKCG,PRKCH,PRKCI,PRKCQ,PRKCZ,PRKD1,RAF1,RALA,RALB,RAP1A,RAP1B,RCAN2,RCAN3,RRAS2,SLC8A1,SLC8A2,SOS1,SRC,TGFB1,* |
| Opioid Signaling Pathway | 1.73E+01 | 6.27E-01 | -7.731 | *ADCY1,ADCY10,ADCY2,ADCY3,ADCY5,ADCY6,ADCY7,ADCY8,ADCY9,AKT2,AKT3,AP1B1,AP2A1,AP2A2,AP2B1,ARRB1,BLK,BRAF,CACNA1A,CACNA1B,CACNA1C,CACNA1D,CACNA1E,CACNA1F,CACNA1G,CACNA1H,CACNA1I,CACNA1S,CACNA2D1,CACNA2D2,CACNA2D3,CACNA2D4,CACNB1,CACNB2,CACNB4,CACNG2,CACNG3,CACNG4,CACNG5,CACNG6,CACNG7,CACNG8,CAMK1D,CAMK2A,CAMK2B,CAMK2G,CAMK4,CDC42,CLTA,CLTB,CLTC,CLTCL1,CREB3L3,CREB5,CREBBP,EP300,FGR,FYN,GNAI1,GNAI2,GNAI3,GNAL,GNAO1,GNAS,GNB1,GNG2,GNG7,GRIN1,GRIN2A,GRIN2B,GRIN2C,GRIN2D,GRK1,GRK2,GRK3,GRK4,GRK5,GRK6,GRK7,GSK3B,HCK,ITPR1,ITPR2,ITPR3,KCNJ5,KCNJ6,KRAS,LCK,MAP2K1,MAP2K2,MAP2K3,MAP2K4,MAP2K5,MAP2K6,MAPK1,MAPK4,MAPK6,MRAS,NOS1,NOS3,OPRD1,OPRL1,PDE1A,PDE1B,PDE1C,PLCB1,PLD2,PPP3CB,PPP3CC,PRKACA,PRKAG2,PRKAR1B,PRKAR2A,PRKCA,PRKCB,PRKCD,PRKCE,PRKCG,PRKCH,PRKCI,PRKCQ,PRKCZ,PRKD1,RAC1,RAC2,RAF1,RALA,RALB,RAP1A,RAP1B,RGS10,RGS12,RGS14,RGS20,RGS3,RGS5,RGS6,RGS7,RGS8,RGS9,RPS6KA1,RPS6KA2,RPS6KA3,RPS6KA5,RPS6KB1,RRAS2,RYR1,RYR2,RYR3,SLC12A5,SOS1,SRC,YES1* |

| | | | | |
|---|---|---|---|---|
| Axonal Guidance Signaling | 1.72E+01 | 5.49E-01 | NaN | ABL1,ABLIM1,ABLIM2,ACE,ACTR2,ACTR3,ADAM10,ADAM11,ADAM12,ADAM15,ADAM19,ADAM22,ADAM32,ADAMTS10,ADAMTS12,ADAMTS13,ADAMTS14,ADAMTS15,ADAMTS16,ADAMTS17,ADAMTS18,ADAMTS2,ADAMTS20,ADAMTS6,ADAMTS7,ADAMTS8,ADAMTS9,AFG3L2,AKT2,AKT3,ARHGEF11,ARHGEF12,ARHGEF15,ARHGEF7,ARPC1A,ARPC5L,BAIAP2,BCAR1,BMP1,BMP5,BMP6,BMP7,BMP8A,BMP8B,CDC42,CHP1,CRK,CXCL12,DCC,DOCK1,DPYSL2,DPYSL5,EFNA3,EFNA5,EPHA1,EPHA10,EPHA2,EPHA3,EPHA6,EPHA8,EPHB1,EPHB2,EPHB4,ERBB2,FARP2,FYN,FZD3,GLI2,GLI3,GLIS1,GNA11,GNA12,GNA14,GNAI1,GNAI2,GNAI3,GNAL,GNAO1,GNAQ,GNAS,GNB1,GNB1L,GNB5,GNG12,GNG2,GNG4,GNG7,GRB2,GSK3B,HERC2,ITGA2,ITGA3,KALRN,KCNJ12,KIF7,KRAS,L1CAM,LIMK1,LIMK2,LINGO1,LNPEP,LRRC4C,MAG,MAP2K1,MAP2K2,MAPK1,MET,MKNK1,MME,MMP14,MMP15,MMP17,MMP2,MMP21,MMP26,MMP28,MRAS,MYL10,MYL3,MYSM1,NCK1,NCK2,NFAT5,NFATC1,NFATC2,NFATC3,NGEF,NRP1,NRP2,NTF3,NTN1,NTN4,NTNG1,NTNG2,NTRK1,NTRK2,NTRK3,PAK1,PAK2,PAK4,PAPPA,PAPPA2,PDGFA,PDGFB,PDGFD,PFN2,PIK3C2A,PIK3C2B,PIK3CB,PIK3CD,PIK3R2,PIK3R3,PIK3R4,PIK3R5,PIK3R6,PLCB1,PLCB2,PLCB3,PLCD3,PLCE1,PLCG1,PLCG2,PLCH1,PLCH2,PLCL1,PLCL2,PLXNA1,PLXNA2,PLXNB1,PLXNB2,PLXNC1,PLXND1,PPP3CB,PPP3CC,PRKACA,PRKAG2,PRKAR1B,PRKAR2A,PRKCA,PRKCB,PRKCD,PRKCE,PRKCG,PRKCH,PRKCI,PRKCQ,PRKCZ,PRKD1,PTCH1,PTCH2,PTK2,PTPN11,PXN,RAC1,RAC2,RAF1,RALA,RALB,RAP1A,RAP1B,RASA1,RASSF5,RGS3,RHOA,RHOD,ROBO1,ROBO2,ROCK1,ROCK2,RRAS2,RTN4,RTN4R,SEMA3A,SEMA3C,SEMA3D,SEMA3E,SEMA3F,SEMA4A,SEMA4B,SEMA4D,SEMA4F,SEMA5A,SEMA6B,SEMA6C,SEMA6D,SEMA7A,SHANK2,SLIT2,SLIT3,SMO,SOS1,SRGAP1,SRGAP2,SRGAP3,SUFU,TUBA1C,TUBB3,UNC5A,UNC5B,UNC5C,UNC5D,VEGFA,VEGFD,WAS,WNT10A,WNT11,WNT2B,WNT3,WNT3A,WNT4,WNT5B,WNT7A,WNT7B,WNT9A,WNT9B,ZN |
| GNRH Signaling | 1.57E+01 | 6.65E-01 | -9.381 | ADCY1,ADCY10,ADCY2,ADCY3,ADCY5,ADCY6,ADCY7,ADCY8,ADCY9,CACNA1A,CACNA1B,CACNA1C,CACNA1D,CACNA1E,CACNA1F,CACNA1G,CACNA1H,CACNA1I,CACNA1S,CACNA2D1,CACNA2D2,CACNA2D3,CACNA2D4,CACNB1,CACNB2,CACNB4,CACNG2,CACNG3,CACNG4,CACNG5,CACNG6,CACNG7,CACNG8,CAMK2A,CAMK2B,CAMK2G,CAMK4,CDC42,CREB3L3,CREB5,CREBBP,EGFR,GNA11,GNA14,GNAI1,GNAI2,GNAI3,GNAQ,GNAS,GNB1,GNG2,GNG7,GRB2,ITPR1,ITPR2,ITPR3,KRAS,MAP2K1,MAP2K2,MAP2K3,MAP2K4,MAP2K6,MAP3K1,MAP3K10,MAP3K11,MAP3K12,MAP3K13,MAP3K14,MAP3K15,MAP3K2,MAP3K3,MAP3K5,MAP3K6,MAPK1,MAPK10,MAPK13,MAPK14,MAPK8,MAPK9,MMP2,MRAS,PAK1,PAK2,PAK4,PLCB1,PLCB2,PLCB3,PRKACA,PRKAG2,PRKAR1B,PRKAR2A,PRKCA,PRKCB,PRKCD,PRKCE,PRKCG,PRKCH,PRKCI,PRKCQ,PRKCZ,PRKD1,PTK2,PTK2B,PXN,RAC1,RAF1,RALA,RALB,RAP1A,RAP1B,RRAS2,SOS1,SRC |
| Synaptogenesis Signaling Pathway | 1.55E+01 | 5.83E-01 | -12.006 | ACTR2,ACTR3,ADCY1,ADCY10,ADCY2,ADCY3,ADCY5,ADCY6,ADCY7,ADCY8,ADCY9,AKT2,AKT3,AP1B1,AP2A1,AP2A2,AP2B1,ARHGEF7,ARPC1A,ARPC5L,BRAF,CACNA1B,CACNA2D1,CACNB1,CACNB4,CADM1,CAMK2A,CAMK2B,CAMK2G,CAMK4,CDC42,CDH1,CDH10,CDH11,CDH12,CDH13,CDH15,CDH16,CDH18,CDH22,CDH23,CDH26,CDH3,CDH4,CDH5,CDH8,CDH9,CHN1,CLASP2,CNTNAP2,CPLX1,CPLX2,CREB3L3,CREB5,CREBBP,CRK,DAB1,DLG4,DNAJC5,EFNA3,EFNA5,EPHA1,EPHA10,EPHA2,EPHA3,EPHA6,EPHA8,EPHB1,EPHB2,EPHB4,FARP1,FGR,FYN,GOSR1,GRB2,GRIA1,GRIA2,GRIA4,GRIN1,GRIN2A,GRIN2B,GRIN2C,GRIN2D,GRM1,GRM3,GRM4,GRM5,GRM7,GRM8,GSK3B,HCK,ITPR1,ITSN2,KALRN,KRAS,LCK,LIMK1,LRP1,LRP8,MAP1B,MAPK1,MAPK14,MAPT,MRAS,MTOR,NAP1L4,NECTIN1,NECTIN3,NLGN1,NLGN4X,NRXN1,NRXN2,NRXN3,NSF,NTRK2,PAFAH1B1,PAK1,PIK3C2A,PIK3C2B,PIK3CB,PIK3CD,PIK3R2,PIK3R3,PIK3R4,PIK3R5,PIK3R6,PLCG1,PLCG2,PRKACA,PRKAG2,PRKAR1B,PRKAR2A,PRKCD,PRKCE,RAB5B,RAC1,RAF1,RALA,RALB,RAP1A,RAP1B,RAPGEF1,RASGRF1,RELN,RHOA,RPS6KB1,RRAS2,SGTA,SHC2,SHC3,SHE,SOS1,SRC,STX1A,STX1B,STXBP1,STXBP3,STXBP4,STXBP5,SYN1,SYN2,SYN3,SYT1,SYT11,SYT12,SYT13,SYT14,SYT17,SYT2,SYT3,SYT6,SYT7,SYT9,TIAM1,TLN1,UNC13A,VTI1A,WAS,YES1 |
| Molecular Mechanisms of Cancer | 1.50E+01 | 5.55E-01 | NaN | ABL1,ADCY1,ADCY10,ADCY2,ADCY3,ADCY5,ADCY6,ADCY7,ADCY8,ADCY9,AKT2,AKT3,APAF1,APC,ARHGEF1,ARHGEF10,ARHGEF11,ARHGEF12,ARHGEF15,ARHGEF16,ARHGEF17,ARHGEF19,ARHGEF2,ARHGEF3,ARHGEF4,ARHGEF7,ARHGEF9,ATM,ATR,AXIN1,BCL2,BCL2L1,BCL2L11,BID,BIRC2,BMP1,BMP5,BMP6,BMP7,BMP8A,BMP8B,BMPR1A,BMPR1B,BMPR2,BRAF,BRCA1,CAMK2A,CAMK2B,CAMK2G,CASP8,CASP9,CBL,CCND3,CDC25A,CDC25C,CDC42,CDH1,CDK11B,CDK12,CDK13,CDK14,CDK16,CDK17,CDK18,CDK19,CDK6,CDK7,CREBBP,CRK,CTNNA1,CTNNA2,E2F1,E2F2,E2F3,E2F4,EP300,FANCD2,FNBP1,FOXO1,FYN,FZD3,GAB2,GNA11,GNA12,GNA14,GNAI1,GNAI2,GNAI3,GNAL,GNAO1,GNAQ,GNAS,GRB2,GSK3B,HHAT,HIPK2,ITGA2,ITGA3,JAK1,JAK2,JAK3,KRAS,LRP1,LRP5,LRP6,MAP2K1,MAP2K2,MAP2K3,MAP2K4,MAP2K6,MAP3K5,MAPK1,MAPK10,MAPK13,MAPK14,MAPK8,MAPK9,MDM2,MRAS,NAIP,NF1,NFKBID,NLK,NOTCH1,PAK1,PAK2,PAK4,PIK3C2A,PIK3C2B,PIK3CB,PIK3CD,PIK3R2,PIK3R3,PIK3R4,PIK3R5,PIK3R6,PLCB1,PLCB2,PLCB3,PRKACA,PRKAG2,PRKAR1B,PRKAR2A,PRKCA,PRKCB,PRKCD,PRKCE,PRKCG,PRKCH,PRKCI,PRKCQ,PRKCZ,PRKD1,PRKDC,PSEN1,PSEN2,PTCH1,PTCH2,PTK2,PTPN11,RAC1,RAC2,RAF1,RALA,RALB,RALBP1,RALGDS,RAP1A,RAP1B,RAPGEF1,RAPGEF3,RASA1,RASGRF1,RASGRF2,RBL1,RBPJ,RHOA,RHOBTB1,RHOD,RHOQ,RHOT1,RRAS2,SIN3A,SMAD3,SMAD6,SMAD7,SMO,SOS1,SRC,SUFU,TAB1,TAB2,TCF3,TCF4,TFDP1,TGFB1,TGFB2,TGFBR2,TP53,TYK2,WNT10A,WNT11,WNT2B,WNT3,WNT3A,WNT4,WNT5B,WNT7A,WNT7B,WNT9A,WNT9B,XIAP,ZBTB17 |
| Endocannabinoid Neuronal Synapse Pathway | 1.46E+01 | 7.01E-01 | -6.254 | ABHD6,ADCY1,ADCY10,ADCY2,ADCY3,ADCY5,ADCY6,ADCY7,ADCY8,ADCY9,CACNA1A,CACNA1B,CACNA1C,CACNA1D,CACNA1E,CACNA1F,CACNA1G,CACNA1H,CACNA1I,CACNA1S,CACNA2D1,CACNA2D2,CACNA2D3,CACNA2D4,CACNB1,CACNB2,CACNB4,CACNG2,CACNG3,CACNG4,CACNG5,CACNG6,CACNG7,CACNG8,CHP1,CNR2,DAGLA,FAAH,GNA11,GNA14,GNAI1,GNAI2,GNAI3,GNAL,GNAO1,GNAQ,GNB1,GNG2,GNG7,GRIA1,GRIA2,GRIA4,GRIN1,GRIN2A,GRIN2B,GRIN2C,GRIN2D,GRM1,GRM5,ITPR1,KCNJ5,KCNJ6,MAPK1,MAPK10,MAPK13,MAPK14,MAPK4,MAPK6,MAPK8,MAPK9,MGLL,PLCB1,PLCB2,PLCB3,PLCD3,PLCE1,PLCG1,PLCG2,PLCH1,PLCH2,PLCL1,PLCL2,PPP3CB,PPP3CC,PRKACA,PRKAG2,PRKAR1B,PRKAR2A,RIMS1 |
| GPCR-Mediated Nutrient Sensing in Enteroendocrine Cells | 1.36E+01 | 7.12E-01 | -8.213 | ADCY1,ADCY10,ADCY2,ADCY3,ADCY5,ADCY6,ADCY7,ADCY8,ADCY9,CACNA1A,CACNA1B,CACNA1C,CACNA1D,CACNA1E,CACNA1F,CACNA1G,CACNA1H,CACNA1I,CACNA1S,CACNA2D1,CACNA2D2,CACNA2D3,CACNA2D4,CACNB1,CACNB2,CACNB4,CACNG2,CACNG3,CACNG4,CACNG5,CACNG6,CACNG7,CACNG8,CASR,FFAR4,GIP,GNA11,GNA14,GNAI1,GNAI2,GNAI3,GNAQ,GNAS,GNG12,GNG2,GNG4,GNG7,ITPR1,ITPR2,ITPR3,LPAR5,PLCB1,PLCB2,PLCB3,PLCD3,PLCE1,PLCG1,PLCG2,PLCH1,PLCH2,PLCL1,PLCL2,PRKACA,PRKAG2,PRKAR1B,PRKAR2A,PRKCA,PRKCB,PRKCD,PRKCE,PRKCG,PRKCH,PRKCI,PRKCQ,PRKCZ,PRKD1,PYY,RAPGEF4,TAS1R1 |

| | | | | |
|---|---|---|---|---|
| Protein Kinase A Signaling | 1.36E+01 | 5.44E-01 | -2.729 | ADCY1,ADCY10,ADCY2,ADCY3,ADCY5,ADCY6,ADCY7,ADCY8,ADCY9,ADD1,ADD2,ADD3,AKAP1,AKAP10,AKAP12,AKAP13,AKAP8,AKAP9,ANAPC1,ANAPC5,ATF1,BRAF,CAMK2A,CAMK2B,CAMK2G,CAMK4,CDC14B,CDC16,CDC23,CDC25A,CDC25C,CDC27,CHP1,CNGA3,CNGB1,CREB3L3,CREB5,CREBBP,DCC,DUSP10,DUSP15,DUSP16,DUSP22,DUSP4,ENPP6,EPM2A,EYA2,EYA3,FLNA,FLNB,FLNC,GDPD1,GLI3,GNAI1,GNAI2,GNAI3,GNAQ,GNAS,GNB1,GNB1L,GNB5,GNG12,GNG2,GNG4,GNG7,GRK1,GRK7,GSK3B,GYS1,HHAT,HNF1A,ITPR1,ITPR2,ITPR3,MAP2K1,MAP2K2,MAP3K1,MAPK1,MPPED2,MTMR3,MYH10,MYL10,MYL3,MYLK,MYLK2,MYLK3,NFAT5,NFATC1,NFATC2,NFATC3,NFKBID,NOS3,NTN1,PDE10A,PDE11A,PDE1A,PDE1B,PDE1C,PDE2A,PDE3A,PDE3B,PDE4A,PDE4B,PDE4C,PDE4D,PDE6A,PDE6B,PDE6C,PDE6G,PDE8A,PDE8B,PDE9A,PHKA2,PHKB,PHKG1,PLCB1,PLCB2,PLCB3,PLCD3,PLCE1,PLCG1,PLCG2,PLCH1,PLCH2,PLCL1,PLCL2,PPP1R12A,PPP1R14C,PPP1R14D,PPP1R3A,PPP1R7,PPP3CB,PPP3CC,PRKACA,PRKAG2,PRKAR1B,PRKAR2A,PRKCA,PRKCB,PRKCD,PRKCE,PRKCG,PRKCH,PRKCI,PRKCQ,PRKCZ,PRKD1,PTCH1,PTCH2,PTEN,PTK2,PTK2B,PTPN1,PTPN11,PTPN12,PTPN14,PTPN2,PTPN22,PTPN23,PTPN3,PTPN4,PTPN5,PTPN7,PTPN9,PTPRA,PTPRB,PTPRC,PTPRD,PTPRE,PTPRF,PTPRG,PTPRH,PTPRJ,PTPRK,PTPRM,PTPRQ,PTPRR,PTPRS,PTPRT,PTPRU,PTPRZ1,PXN,PYGB,PYGM,RAF1,RAP1A,RAP1B,RHOA,ROCK1,ROCK2,RYR1,RYR2,RYR3,SIRPA,SMAD3,SMO,TCF3,TCF4,TCF7,TCF7L1,TCF7L2,TGFB1,TGFB2,TGFBR2,TIMM50,TNNI1,UBASH3B,YWHAE,YWHAG |
| Netrin Signaling | 1.26E+01 | 8.00E-01 | -6.934 | ABLIM1,ABLIM2,CACNA1A,CACNA1B,CACNA1C,CACNA1D,CACNA1E,CACNA1F,CACNA1G,CACNA1H,CACNA1I,CACNA1S,CACNA2D1,CACNA2D2,CACNA2D3,CACNA2D4,CACNB1,CACNB2,CACNB4,CACNG2,CACNG3,CACNG4,CACNG5,CACNG6,CACNG7,CACNG8,CHP1,DCC,ENAH,NCK1,NCK2,NFAT5,NFATC1,NFATC2,NFATC3,NTN1,PPP3CB,PPP3CC,PRKACA,PRKAG2,PRKAR1B,PRKAR2A,PRKG1,RAC1,RAC2,RYR1,RYR2,RYR3,UNC5A,UNC5B,UNC5C,UNC5D |
| GP6 Signaling Pathway | 1.14E+01 | 6.72E-01 | -8.379 | ADAM10,AKT2,AKT3,CAMK4,CERT1,COL11A1,COL13A1,COL15A1,COL16A1,COL17A1,COL18A1,COL1A1,COL20A1,COL21A1,COL22A1,COL23A1,COL24A1,COL25A1,COL26A1,COL27A1,COL28A1,COL2A1,COL4A1,COL4A2,COL4A4,COL4A6,COL5A1,COL5A3,COL6A3,COL6A5,COL6A6,COL7A1,COL8A1,COL8A2,COL9A2,COL9A3,FYN,GSK3B,ITGA2B,ITGB3,ITPR1,LAMA1,LAMA2,LAMA3,LAMA5,LAMB1,LAMB2,LAMB3,LAMC1,LAMC2,LAMC3,LCP2,PDPK1,PIK3C2A,PIK3C2B,PIK3CB,PIK3CD,PIK3R2,PIK3R3,PIK3R4,PIK3R5,PIK3R6,PLCG2,PRKCA,PRKCB,PRKCD,PRKCE,PRKCG,PRKCH,PRKCI,PRKCQ,PRKCZ,PRKD1,PTK2,RAC1,RAP1B,SYK,TLN1 |
| Reelin Signaling in Neurons | 1.14E+01 | 6.56E-01 | -8.835 | ACTR2,ACTR3,AKT2,AKT3,APBB1,APP,ARHGEF1,ARHGEF10,ARHGEF11,ARHGEF12,ARHGEF15,ARHGEF16,ARHGEF2,ARHGEF3,ARHGEF4,ARHGEF9,ARPC1A,ARPC5L,BLK,CAMK2A,CAMK2B,CAMK2G,CDC42,CNR2,CRK,DAB1,FGR,FYN,GRIN1,GRIN2A,GRIN2B,GRIN2C,GRIN2D,GSK3B,HCK,ITGA2,ITGA3,ITGAL,ITGB2,ITGB3,LCK,LIMK1,LRP8,MAP1B,MAP2K1,MAP2K2,MAP2K3,MAP2K4,MAP2K5,MAP2K6,MAP3K10,MAP3K11,MAP4K1,MAPK1,MAPK10,MAPK8,MAPK8IP1,MAPK8IP3,MAPK9,MAPT,MTOR,NDEL1,NECTIN3,PAFAH1B1,PAFAH1B2,PDK2,PDK3,PIK3C2A,PIK3C2B,PIK3CB,PIK3CD,PIK3R2,PIK3R3,PIK3R4,PIK3R5,PIK3R6,RAC1,RAP1A,RAP1B,RAPGEF1,RELN,RHOA,SRC,YES1 |
| G Beta Gamma Signaling | 1.08E+01 | 6.56E-01 | -8.605 | ADCY1,ADCY2,AKT2,AKT3,CACNA1A,CACNA1B,CACNA1C,CACNA1D,CACNA1E,CACNA1F,CACNA1G,CACNA1H,CACNA1I,CACNA1S,CACNA2D1,CACNA2D2,CACNA2D3,CACNA2D4,CACNB1,CACNB2,CACNB4,CACNG2,CACNG3,CACNG4,CACNG5,CACNG6,CACNG7,CACNG8,CDC42,DNM2,EGFR,GNA11,GNA12,GNA14,GNAI1,GNAI2,GNAI3,GNAL,GNAO1,GNAQ,GNAS,GNB1,GNB1L,GNB5,GNG12,GNG2,GNG4,GNG7,GRB2,KCNJ5,KCNJ6,KRAS,MAPK1,MRAS,PAK1,PDPK1,PLCG1,PLCG2,PRKACA,PRKAG2,PRKAR1B,PRKAR2A,PRKCA,PRKCB,PRKCD,PRKCE,PRKCG,PRKCH,PRKCI,PRKCQ,PRKCZ,PRKD1,RAF1,RALA,RALB,RAP1A,RAP1B,RRAS2,SOS1,SRC |
| Corticotropin Releasing Hormone Signaling | 1.06E+01 | 6.31E-01 | -3.491 | ADCY1,ADCY10,ADCY2,ADCY3,ADCY5,ADCY6,ADCY7,ADCY8,ADCY9,BRAF,CACNA1A,CACNA1B,CACNA1C,CACNA1D,CACNA1E,CACNA1F,CACNA1G,CACNA1H,CACNA1I,CACNA1S,CACNA2D1,CACNA2D2,CACNA2D3,CACNA2D4,CACNB1,CACNB2,CACNB4,CACNG2,CACNG3,CACNG4,CACNG5,CACNG6,CACNG7,CACNG8,CAMK4,CREB3L3,CREB5,CREBBP,CRHR1,CRHR2,GLI2,GLI3,GNAI1,GNAI2,GNAI3,GNAO1,GNAQ,GNAS,GUCY1A2,GUCY2D,ITPR1,ITPR2,ITPR3,MAP2K1,MAP2K2,MAPK1,MAPK13,MAPK14,MEF2A,MEF2D,NOS1,NOS2,NOS3,NPR1,NPR3,NR4A1,PLCG1,PLCG2,PRKACA,PRKAG2,PRKAR1B,PRKAR2A,PRKCA,PRKCB,PRKCD,PRKCE,PRKCG,PRKCH,PRKCI,PRKCQ,PRKCZ,PRKD1,PTCH1,PTCH2,RAF1,RAP1A,RAP1B,SMO,VEGFA |
| Calcium Signaling | 1.05E+01 | 5.86E-01 | -8.51 | ATP2A1,ATP2A2,ATP2A3,ATP2B1,ATP2B2,ATP2B3,ATP2B4,ATP2C1,CABIN1,CACNA1A,CACNA1B,CACNA1C,CACNA1D,CACNA1E,CACNA1F,CACNA1G,CACNA1H,CACNA1I,CACNA1S,CACNA2D1,CACNA2D2,CACNA2D3,CACNA2D4,CACNB1,CACNB2,CACNB4,CACNG2,CACNG3,CACNG4,CACNG5,CACNG6,CACNG7,CACNG8,CAMK1D,CAMK2A,CAMK2B,CAMK2G,CAMK4,CAMKK1,CAMKK2,CASQ2,CHP1,CHRFAM7A,CHRNA2,CHRNA4,CHRNA5,CHRNA7,CHRNB1,CHRNB2,CHRNB4,CREB3L3,CREB5,CREBBP,EP300,GRIA1,GRIA2,GRIA4,GRIN1,GRIN2A,GRIN2B,GRIN2C,GRIN2D,HDAC1,HDAC11,HDAC4,HDAC5,HDAC6,HDAC7,HDAC9,ITPR1,ITPR2,ITPR3,LETM1,MAPK1,MCU,MEF2A,MEF2D,MICU1,MYH10,MYH11,MYH14,MYH3,MYH6,MYH7,MYH7B,MYH9,MYL3,NFAT5,NFATC1,NFATC2,NFATC3,PPP3CB,PPP3CC,PRKACA,PRKAG2,PRKAR1B,PRKAR2A,RAP1A,RAP1B,RCAN2,RCAN3,RYR1,RYR2,RYR3,SLC8A1,SLC8A2,SLC8B1,TNNI1,TNNT1,TNNT2,TPM3,TPM4,TRPC1,TRPC6,TRPC7,TRPM8 |
| Paxillin Signaling | 1.03E+01 | 6.70E-01 | -6.804 | ACTB,ACTG2,ACTN1,ACTN2,ACTN4,ARF1,ARHGEF7,BCAR1,CDC42,CRK,CSK,DOCK1,GRB2,ITGA10,ITGA11,ITGA2,ITGA2B,ITGA3,ITGA7,ITGA9,ITGAD,ITGAE,ITGAL,ITGAM,ITGAX,ITGB2,ITGB3,ITGB4,ITGB5,ITGB6,ITGB7,ITGB8,KRAS,MAP2K4,MAPK1,MAPK10,MAPK13,MAPK14,MAPK8,MAPK9,MRAS,NCK1,NCK2,PAK1,PAK2,PAK4,PARVA,PIK3C2A,PIK3C2B,PIK3CB,PIK3CD,PIK3R2,PIK3R3,PIK3R4,PIK3R5,PIK3R6,PTK2,PTK2B,PTPN12,PXN,RAC1,RALA,RALB,RAP1A,RAP1B,RRAS2,SOS1,SRC,TLN1,TLN2,VCL |

| | | | | |
|---|---|---|---|---|
| Endothelin-1 Signaling | 1.01E+01 | 5.90E-01 | -8.1 | ABHD3,ADCY1,ADCY10,ADCY2,ADCY3,ADCY5,ADCY6,ADCY7,ADCY8,ADCY9,BRAF,CASP8,CASP9,ECE1,GNA11,GNA12,GNA14,GNAI1,GNAI2,GNAI3,GNAL,GNAO1,GNAQ,GNAS,GPLD1,GRB2,GUCY1A2,GUCY2D,ITPR1,ITPR2,ITPR3,KRAS,MAPK1,MAPK10,MAPK13,MAPK14,MAPK4,MAPK6,MAPK8,MAPK9,MRAS,NOS1,NOS2,NOS3,PIK3C2A,PIK3C2B,PIK3CB,PIK3CD,PIK3R2,PIK3R3,PIK3R4,PIK3R5,PIK3R6,PLA2G10,PLA2G12B,PLA2G2C,PLA2G2F,PLA2G4A,PLA2G4C,PLA2G4D,PLA2G4E,PLA2G4F,PLA2G5,PLA2G6,PLA2R1,PLAAT3,PLAAT4,PLAAT5,PLB1,PLCB1,PLCB2,PLCB3,PLCD3,PLCE1,PLCG1,PLCG2,PLCH1,PLCH2,PLCL1,PLCL2,PLD1,PLD2,PLD3,PLD5,PNPLA3,PNPLA8,PRKCA,PRKCB,PRKCD,PRKCE,PRKCG,PRKCH,PRKCI,PRKCQ,PRKCZ,PRKD1,PTGS1,RAF1,RALA,RALB,RAP1A,RAP1B,RRAS2,SHC2,SHC3,SHE,SOS1,SRC |
| Dopamine-DARPP32 Feedback in cAMP Signaling | 9.81E+00 | 6.04E-01 | -7.419 | ADCY1,ADCY10,ADCY2,ADCY3,ADCY5,ADCY6,ADCY7,ADCY8,ADCY9,ATF1,ATP2A1,ATP2A2,ATP2A3,CACNA1A,CACNA1C,CACNA1D,CACNA1E,CACNA1S,CAMK4,CAMKK1,CAMKK2,CHP1,CREB3L3,CREB5,CREBBP,CSNK1D,CSNK1E,CSNK1G2,DRD2,DRD3,GNAI1,GNAI2,GNAI3,GNAQ,GNAS,GRIN1,GRIN2A,GRIN2B,GRIN2C,GRIN2D,GUCY1A2,ITPR1,ITPR2,ITPR3,KCNJ12,KCNJ15,KCNJ4,KCNJ5,KCNJ6,NOS1,PAWR,PLCB1,PLCB2,PLCB3,PLCD3,PLCE1,PLCG1,PLCG2,PLCH1,PLCH2,PLCL1,PLCL2,PPM1L,PPP1R12A,PPP1R14C,PPP1R14D,PPP1R3A,PPP1R7,PPP2CB,PPP2R1B,PPP2R2A,PPP2R2B,PPP2R2C,PPP2R3A,PPP2R5B,PPP2R5C,PPP2R5D,PPP2R5E,PPP3CB,PPP3CC,PRKACA,PRKAG2,PRKAR1B,PRKAR2A,PRKCA,PRKCB,PRKCD,PRKCE,PRKCG,PRKCH,PRKCI,PRKCQ,PRKCZ,PRKD1,PRKG1,PTPA |
| IL-15 Production | 9.77E+00 | 6.44E-01 | -8.718 | AATK,ABL1,ABL2,ALK,AXL,BLK,CLK2,CLK3,CSF1R,CSF2RA,CSK,DDR2,DSTYK,DYRK3,EGFR,EPHA1,EPHA2,EPHA3,EPHA6,EPHA8,EPHB1,EPHB2,EPHB4,ERBB2,ERBB3,ERBB4,FER,FGFR1,FGFR2,FGFR4,FGR,FLT1,FLT4,FYN,HCK,IGF1R,INSR,JAK1,JAK2,JAK3,LCK,LMTK2,LMTK3,MAP2K1,MAP2K3,MAP2K4,MAP2K5,MAP2K6,MAP3K11,MERTK,MET,MST1R,NTRK1,NTRK2,NTRK3,PDGFRB,PEAK1,PRKCI,PRKCZ,PTK2,PTK2B,PTK7,RET,ROR1,ROR2,RYK,SRC,SYK,TEK,TIE1,TNK1,TNK2,TYK2,TYRO3,YES1,ZAP70 |
| Thrombin Signaling | 9.56E+00 | 5.71E-01 | -7.839 | ADCY1,ADCY10,ADCY2,ADCY3,ADCY5,ADCY6,ADCY7,ADCY8,ADCY9,AKT2,AKT3,ARHGEF1,ARHGEF10,ARHGEF11,ARHGEF12,ARHGEF15,ARHGEF16,ARHGEF2,ARHGEF3,ARHGEF4,ARHGEF9,CAMK1D,CAMK2A,CAMK2B,CAMK2G,CAMK4,CDC42,EGFR,F2,FNBP1,GATA4,GNA11,GNA12,GNA14,GNAI1,GNAI2,GNAI3,GNAL,GNAO1,GNAQ,GNAS,GNB1,GNB1L,GNB5,GNG12,GNG2,GNG4,GNG7,GRB2,IKBKB,ITPR1,ITPR2,ITPR3,KRAS,MAP2K1,MAP2K2,MAPK1,MAPK13,MAPK14,MPRIP,MRAS,MYL10,MYL3,MYLK,PDPK1,PIK3C2A,PIK3C2B,PIK3CB,PIK3CD,PIK3R2,PIK3R3,PIK3R4,PIK3R5,PIK3R6,PLCB1,PLCB2,PLCB3,PLCD3,PLCE1,PLCG1,PLCG2,PLCH1,PLCH2,PLCL1,PLCL2,PPP1R12A,PPP1R12B,PRKCA,PRKCB,PRKCD,PRKCE,PRKCG,PRKCH,PRKCI,PRKCQ,PRKCZ,PRKD1,PTK2,RAC1,RAC2,RAF1,RALA,RALB,RAP1A,RAP1B,RHOA,RHOBTB1,RHOD,RHOQ,RHOT1,ROCK1,ROCK2,RPS6KB1,RRAS2,SOS1,SRC |
| Fcγ Receptor-mediated Phagocytosis in Macrophages and Monocytes | 9.56E+00 | 6.77E-01 | -7.366 | ACTB,ACTG2,ACTR2,ACTR3,AKT2,AKT3,ARPC1A,ARPC5L,CBL,CDC42,CRK,DGKB,DOCK1,EZR,FGR,FYB1,FYN,GAB2,GPLD1,HCK,INPP5D,LCP2,MAPK1,MYO5A,NCK1,NCK2,PAK1,PIK3R2,PIK3R3,PIP5K1A,PLA2G6,PLCG1,PLD1,PLD2,PLD3,PLD5,PRKCA,PRKCB,PRKCD,PRKCE,PRKCG,PRKCH,PRKCI,PRKCQ,PRKCZ,PRKD1,PTEN,PTK2B,PXN,RAB11A,RAB11B,RAC1,RAC2,RPS6KB1,SRC,SYK,TLN1,TLN2,VAV1,VAV2,VAV3,WAS,YES1 |
| CCR3 Signaling in Eosinophils | 9.26E+00 | 6.33E-01 | -7.28 | CAMK4,CCL24,CCL26,CCR3,GNAI1,GNAI2,GNAI3,GNAS,GNB1,GNB1L,GNB5,GNG12,GNG2,GNG4,GNG7,ITPR1,ITPR2,ITPR3,KRAS,LIMK1,LIMK2,MAP2K1,MAP2K2,MAPK1,MAPK13,MAPK14,MPRIP,MRAS,MYLK,PAK1,PAK2,PAK4,PIK3C2A,PIK3C2B,PIK3CB,PIK3CD,PIK3R2,PIK3R3,PIK3R4,PIK3R5,PIK3R6,PLA2G10,PLA2G12B,PLA2G2F,PLA2G4A,PLA2G4C,PLA2G4D,PLA2G4E,PLA2G4F,PLA2G5,PLA2G6,PLCB1,PLCB2,PLCB3,PPP1R12A,PPP1R12B,PRKCA,PRKCB,PRKCD,PRKCE,PRKCG,PRKCH,PRKCI,PRKCQ,PRKCZ,PRKD1,RAC1,RAF1,RALA,RALB,RAP1A,RAP1B,RHOA,ROCK1,ROCK2,RRAS2 |
| Neuropathic Pain Signaling In Dorsal Horn Neurons | 9.24E+00 | 6.60E-01 | -8.124 | CAMK1D,CAMK2A,CAMK2B,CAMK2G,CAMK4,GRIA1,GRIA2,GRIA4,GRIN1,GRIN2A,GRIN2B,GRIN2C,GRIN2D,GRM1,GRM3,GRM4,GRM5,GRM7,GRM8,ITPR1,ITPR2,ITPR3,KCNH2,KCNN1,KCNN3,KCNN4,KCNQ2,KCNQ3,MAPK1,NTRK2,PIK3C2A,PIK3C2B,PIK3CB,PIK3CD,PIK3R2,PIK3R3,PIK3R4,PIK3R5,PIK3R6,PLCB1,PLCB2,PLCB3,PLCD3,PLCE1,PLCG1,PLCG2,PLCH1,PLCH2,PLCL1,PLCL2,PRKACA,PRKAG2,PRKAR1B,PRKAR2A,PRKCA,PRKCB,PRKCD,PRKCE,PRKCG,PRKCH,PRKCI,PRKCQ,PRKCZ,PRKD1,SRC,TACR1 |
| Integrin Signaling | 9.21E+00 | 5.66E-01 | -9.238 | ABL1,ACTB,ACTG2,ACTN1,ACTN2,ACTN4,ACTR2,ACTR3,AKT2,AKT3,ARF1,ARF4,ARHGAP26,ARHGEF7,ARPC1A,ARPC5L,ASAP1,BCAR1,BCAR3,BRAF,CAPN1,CAPN11,CAPN2,CAPN5,CAPN8,CDC42,CRK,CTTN,DOCK1,FNBP1,FYN,GRB2,GSK3B,GSN,ITGA10,ITGA11,ITGA2,ITGA2B,ITGA3,ITGA7,ITGA9,ITGAD,ITGAE,ITGAL,ITGAM,ITGAX,ITGB2,ITGB3,ITGB4,ITGB5,ITGB6,ITGB7,ITGB8,KRAS,LIMS1,MAP2K1,MAP2K2,MAP2K4,MAP3K11,MAPK1,MAPK8,MPRIP,MRAS,MYLK,MYLK2,MYLK3,NCK1,NCK2,NEDD9,PAK1,PAK2,PAK4,PARVA,PARVB,PDGFB,PFN2,PIK3C2A,PIK3C2B,PIK3CB,PIK3CD,PIK3R2,PIK3R3,PIK3R4,PIK3R5,PIK3R6,PLCG1,PLCG2,PPP1R12A,PPP1R12B,PTEN,PTK2,PXN,RAC1,RAC2,RAF1,RALA,RALB,RAP1A,RAP1B,RAPGEF1,RHOA,RHOBTB1,RHOD,RHOQ,RHOT1,ROCK1,RRAS2,SOS1,SRC,TLN1,TLN2,TNK2,TSPAN3,TSPAN7,VCL,WAS |
| Signaling by Rho Family GTPases | 9.10E+00 | 5.48E-01 | -9.725 | ACTB,ACTG2,ACTR2,ACTR3,ARHGEF1,ARHGEF10,ARHGEF11,ARHGEF12,ARHGEF15,ARHGEF16,ARHGEF17,ARHGEF19,ARHGEF2,ARHGEF3,ARHGEF4,ARHGEF7,ARHGEF9,ARPC1A,ARPC5L,BAIAP2,CDC42,CDC42EP4,CDH1,CDH10,CDH11,CDH12,CDH13,CDH15,CDH16,CDH18,CDH22,CDH23,CDH26,CDH3,CDH4,CDH5,CDH8,CDH9,CIT,CLIP1,CYFIP1,DIAPH3,EZR,FNBP1,GFAP,GNA11,GNA12,GNA14,GNAI1,GNAI2,GNAI3,GNAL,GNAO1,GNAQ,GNAS,GNB1,GNB1L,GNB5,GNG12,GNG2,GNG4,GNG7,IQGAP1,ITGA2,ITGA3,LIMK1,LIMK2,MAP2K1,MAP2K2,MAP2K4,MAP3K10,MAP3K11,MAP3K12,MAPK1,MAPK10,MAPK8,MAPK9,MRAS,MSN,MYL10,MYL3,MYLK,NEDD4,NOX4,PAK1,PAK2,PAK4,PARD3,PI4KA,PIK3C2A,PIK3C2B,PIK3CB,PIK3CD,PIK3R2,PIK3R3,PIK3R4,PIK3R5,PIK3R6,PIP4K2B,PIP5K1A,PIP5K1B,PIP5K1C,PIP5KL1,PKN1,PLD1,PPP1R12A,PPP1R12B,PPP1R12C,PRKCI,PRKCZ,PTK2,PTK2B,RAC1,RAC2,RAF1,RDX,RHOA,RHOBTB1,RHOD,RHOQ,RHOT1,ROCK1,ROCK2,SEPTIN10,SEPTIN14,SEPTIN6,SEPTIN7,SEPTIN8,SEPTIN9,SLC9A1,WAS,WASF3 |

| | | | | |
|---|---|---|---|---|
| ErbB Signaling | 8.99E+00 | 6.67E-01 | -7.298 | *CDC42,EGFR,ERBB2,ERBB3,ERBB4,FOXO1,GRB2,GSK3B,KRAS,MAP2K1,MAP2K2,MAP2K3,MAP2K4,MAP2K6,MAPK1,MAPK10,MAPK13,MAPK14,MAPK8,MAPK9,MRAS,MTOR,NCK1,NCK2,NRG1,NRG2,NRG3,NRG4,PAK1,PAK2,PAK4,PDPK1,PIK3C2A,PIK3C2B,PIK3CB,PIK3CD,PIK3R2,PIK3R3,PIK3R4,PIK3R5,PIK3R6,PLCG1,PLCG2,PRKCA,PRKCB,PRKCD,PRKCE,PRKCG,PRKCH,PRKCI,PRKCQ,PRKCZ,PRKD1,RAF1,RALA,RALB,RAP1A,RAP1B,RPS6KB1,RRAS2,SOS1,TGFA* |
| nNOS Signaling in Skeletal Muscle Cells | 8.83E+00 | 8.25E-01 | NaN | *CACNA1A,CACNA1B,CACNA1C,CACNA1D,CACNA1E,CACNA1F,CACNA1G,CACNA1H,CACNA1I,CACNA1S,CACNA2D1,CACNA2D2,CACNA2D3,CACNA2D4,CACNB1,CACNB2,CACNB4,CACNG2,CACNG3,CACNG4,CACNG5,CACNG6,CACNG7,CACNG8,CAMK4,DAG1,DMD,NOS1,RYR1,RYR2,RYR3,SNTA1,SNTB2* |
| Cellular Effects of Sildenafil (Viagra) | 8.81E+00 | 6.19E-01 | NaN | *ACTB,ACTG2,ADCY1,ADCY10,ADCY2,ADCY3,ADCY5,ADCY6,ADCY7,ADCY8,ADCY9,CACNA1A,CACNA1C,CACNA1D,CACNA1E,CACNA1S,CACNG2,CACNG3,CACNG4,CACNG5,CACNG6,CACNG7,CACNG8,CAMK4,GNAS,GUCY1A2,GUCY2D,ITPR1,ITPR2,ITPR3,KCNH2,KCNN1,KCNN3,KCNN4,KCNQ2,KCNQ3,MPRIP,MYH10,MYH11,MYH14,MYH3,MYH6,MYH7,MYH7B,MYH9,MYL10,MYL3,MYLK,NOS3,PDE1A,PDE1B,PDE1C,PDE2A,PDE3A,PDE3B,PDE4A,PDE4B,PDE4C,PDE4D,PLCB1,PLCB2,PLCB3,PLCD3,PLCE1,PLCG1,PLCG2,PLCH1,PLCH2,PLCL1,PLCL2,PPP1R12A,PPP1R12B,PRKACA,PRKAG2,PRKAR1B,PRKAR2A,PRKG1,SLC4A10* |
| α-Adrenergic Signaling | 8.71E+00 | 6.60E-01 | -6.325 | *ADCY1,ADCY10,ADCY2,ADCY3,ADCY5,ADCY6,ADCY7,ADCY8,ADCY9,ADRA1A,CAMK4,EPM2A,GNAI1,GNAI2,GNAI3,GNAQ,GNAS,GNB1,GNB1L,GNB5,GNG12,GNG2,GNG4,GNG7,GYS1,ITPR1,ITPR2,ITPR3,KRAS,MAP2K1,MAP2K2,MAPK1,MRAS,PHKA2,PHKB,PHKG1,PLCG1,PLCG2,PRKACA,PRKAG2,PRKAR1B,PRKAR2A,PRKCA,PRKCB,PRKCD,PRKCE,PRKCG,PRKCH,PRKCI,PRKCQ,PRKCZ,PRKD1,PYGB,PYGM,RAF1,RALA,RALB,RAP1A,RAP1B,RRAS2,SLC8A1,SLC8A2* |
| Actin Cytoskeleton Signaling | 8.60E+00 | 5.54E-01 | -8.178 | *ACTB,ACTG2,ACTN1,ACTN2,ACTN4,ACTR2,ACTR3,APC,ARHGAP24,ARHGAP35,ARHGEF1,ARHGEF12,ARHGEF4,ARHGEF7,ARPC1A,ARPC5L,BAIAP2,BCAR1,CDC42,CRK,CSK,CYFIP1,CYFIP2,DIAPH1,DIAPH2,DIAPH3,DOCK1,EZR,F2,FGD1,FGD3,FGF1,FGF12,FGF14,FGF18,FLNA,GNA12,GNG12,GRB2,GSN,IQGAP1,IQGAP2,IQGAP3,ITGA2,ITGA3,KRAS,LBP,LIMK1,LIMK2,MAP2K1,MAP2K2,MAPK1,MPRIP,MRAS,MSN,MYH10,MYH11,MYH14,MYH3,MYH6,MYH7,MYH7B,MYH9,MYL10,MYL3,MYLK,MYLK2,MYLK3,PAK1,PAK2,PAK4,PDGFA,PDGFB,PDGFD,PFN2,PIK3C2A,PIK3C2B,PIK3CB,PIK3CD,PIK3R2,PIK3R3,PIK3R4,PIK3R5,PIK3R6,PIP5K1A,PIP5K1B,PIP5K1C,PPP1R12A,PPP1R12B,PTK2,PXN,RAC1,RAC2,RAF1,RALA,RALB,RAP1A,RAP1B,RDX,RHOA,ROCK1,ROCK2,RRAS2,SLC9A1,SOS1,SSH1,SSH2,TIAM1,TIAM2,TLN1,TLN2,TRIO,VAV1,VAV2,VAV3,VCL,WAS,WASF2* |
| AMPK Signaling | 8.60E+00 | 5.54E-01 | -3.989 | *ACACA,ACACB,ACTB,ACTL6B,ADRA1A,ADRA1B,AK2,AK4,AK5,AK7,AK8,AK9,AKT2,AKT3,ARID1A,ARID2,CAB39,CAMKK2,CFTR,CHRM3,CHRNA2,CHRNA4,CHRNA5,CHRNA7,CHRNB1,CHRNB2,CHRNB4,CKM,CPT1A,CREB3L3,CREB5,CREBBP,EEF2K,ELAVL1,EP300,EPM2A,FOXO1,FOXO3,FOXO6,GNAS,GNB1L,GYS1,HLTF,HNF4A,INSR,IRS2,KAT2B,MAP2K3,MAPK1,MAPK13,MAPK14,MRAS,MTOR,NOS3,ORAI1,ORAI2,PBRM1,PFKFB1,PFKFB2,PFKFB3,PFKFB4,PFKM,PFKP,PHLPP1,PHLPP2,PIK3C2A,PIK3C2B,PIK3CB,PIK3CD,PIK3R2,PIK3R3,PIK3R4,PIK3R5,PIK3R6,PPM1B,PPM1D,PPM1E,PPM1F,PPM1H,PPM1L,PPP2CB,PPP2R1B,PPP2R2A,PPP2R2B,PPP2R2C,PPP2R3A,PPP2R5B,PPP2R5C,PPP2R5D,PPP2R5E,PRKAA1,PRKAA2,PRKACA,PRKAG2,PRKAR1B,PRKAR2A,PTPA,RAB11A,RAB1A,RAB22A,RAB27A,RAB6A,RAB7A,RPS6KB1,RPTOR,SIRT1,SLC2A1,SMARCA2,SMARCA4,SMARCB1,SMARCC1,SMARCD1,SMARCD3,STIM1,TBC1D1,TSC1,TSC2,ULK1* |
| Type II Diabetes Mellitus Signaling | 8.49E+00 | 6.00E-01 | -5.308 | *ABCC8,ACSBG1,ACSL4,ACSL5,ACSL6,AKT2,AKT3,CACNA1A,CACNA1B,CACNA1C,CACNA1D,CACNA1E,CACNA1F,CACNA1G,CACNA1H,CACNA1I,CACNA1S,CACNA2D1,CACNA2D2,CACNA2D3,CACNA2D4,CACNB1,CACNB2,CACNB4,CACNG2,CACNG3,CACNG4,CACNG5,CACNG6,CACNG7,CACNG8,CD36,GCK,IKBKB,IKBKE,IKBKG,INSR,IRS2,MAP2K4,MAP3K1,MAP3K14,MAP3K5,MAPK1,MAPK10,MAPK8,MAPK9,MTOR,NFKBID,PDPK1,PIK3C2A,PIK3C2B,PIK3CB,PIK3CD,PIK3R2,PIK3R3,PIK3R4,PIK3R5,PIK3R6,PKLR,PKM,PPARG,PRKAA1,PRKAA2,PRKAG2,PRKCA,PRKCB,PRKCD,PRKCE,PRKCG,PRKCH,PRKCI,PRKCQ,PRKCZ,PRKD1,SLC27A1,SLC27A4,SLC27A5,SLC27A6,SMPD3,SMPD4,SOCS5,SOCS7,TNFRSF1B,TRAF2* |
| Cardiac Hypertrophy Signaling (Enhanced) | 8.48E+00 | 4.85E-01 | -13.908 | *ACE,ACVR1B,ACVR2B,ADCY1,ADCY10,ADCY2,ADCY3,ADCY5,ADCY6,ADCY7,ADCY8,ADCY9,ADRA1A,ADRA1B,AKAP13,AKT2,AKT3,ATF6,ATP2A1,ATP2A2,ATP2A3,BMPR2,CACNA1A,CACNA1C,CACNA1D,CACNA1E,CACNA1S,CAMK2A,CAMK2B,CAMK2G,CAMK4,CHP1,CSF2RB,DIAPH1,DIAPH2,DIAPH3,DLG1,EDA,EIF2B3,EIF2B5,ENPP6,EP300,FGF1,FGF12,FGF14,FGF18,FGFR1,FGFR2,FGFR4,FZD3,GATA4,GDPD1,GHR,GNA11,GNA12,GNA14,GNAI1,GNAI2,GNAI3,GNAQ,GNAS,GNB1,GNG2,GNG7,GSK3B,HDAC1,HDAC11,HDAC4,HDAC5,HDAC6,HDAC7,HDAC9,IFNLR1,IGF1R,IKBKB,IKBKE,IKBKG,IL12RB1,IL12RB2,IL15RA,IL17RA,IL17RC,IL17RD,IL1R1,IL21R,IL22RA1,IL27RA,IL2RA,IL2RB,IL31RA,IL3RA,IL4R,IL6R,IL6ST,INPP5F,ITGA2,ITGA3,ITPR1,ITPR2,ITPR3,JAK2,KRAS,MAP2K1,MAP2K2,MAP2K3,MAP2K4,MAP2K5,MAP2K6,MAP3K1,MAP3K10,MAP3K11,MAP3K12,MAP3K13,MAP3K14,MAP3K15,MAP3K2,MAP3K3,MAP3K5,MAP3K6,MAPK1,MAPK10,MAPK13,MAPK14,MAPK8,MAPK9,MAPKAPK2,MAPKAPK3,MEF2A,MEF2D,MKNK1,MPPED2,MRAS,MTOR,NFAT5,NFATC1,NFATC2,NFATC3,NPR1,PDE10A,PDE11A,PDE1A,PDE1B,PDE1C,PDE2A,PDE3A,PDE3B,PDE4A,PDE4B,PDE4C,PDE4D,PDE6A,PDE6B,PDE6C,PDE6G,PDE8A,PDE8B,PDE9A,PIK3C2A,PIK3C2B,PIK3CB,PIK3CD,PIK3R2,PIK3R3,PIK3R4,PIK3R5,PIK3R6,PKN1,PLCB1,PLCB2,PLCB3,PLCD3,PLCE1,PLCG1,PLCG2,PLCH1,PLCH2,PLCL1,PLCL2,PPP3CB,PPP3CC,PRKACA,PRKAG2,PRKAR1B,PRKAR2A,PRKCA,PRKCB,PRKCD,PRKCE,PRKCG,PRKCH,PRKCI,PRKCQ,PRKCZ,PRKD1,PRKG1,PTEN,PTK2,RAF1,RALA,RALB,RAP1A,RAP1B,RHOA,ROCK1,ROCK2,RPS6KA5,RPS6KB1,RRAS2,RYR1,RYR2,RYR3,SMO,STAT3,TG,TGFB1,TGFB2,TGFBR2,TGFBR3,TNFRSF1B,TSC2,WNT10A,WNT11,WNT2B,WNT3,WNT3A,WNT4,WNT5B,WNT7A,WNT7B,WNT9A,WNT9B* |
| CXCR4 Signaling | 8.38E+00 | 5.79E-01 | -6.932 | *ADCY1,ADCY10,ADCY2,ADCY3,ADCY5,ADCY6,ADCY7,ADCY8,ADCY9,AKT2,AKT3,ARHGEF11,BCAR1,CD4,CDC42,CRK,CXCL12,DOCK1,ELMO1,ELMO2,FNBP1,GNA11,GNA12,GNA14,GNAI1,GNAI2,GNAI3,GNAL,GNAO1,GNAQ,GNAS,GNB1,GNB1L,GNB5,GNG12,GNG2,GNG4,GNG7,ITPR1,ITPR2,ITPR3,KRAS,MAP2K1,MAP2K2,MAP2K4,MAPK1,MAPK10,MAPK8,MAPK9,MRAS,MYL10,MYL3,PAK1,PAK2,PAK4,PIK3C2A,PIK3C2B,PIK3CB,PIK3CD,PIK3R2,PIK3R3,PIK3R4,PIK3R5,PIK3R6,PLCB1,PLCB2,PLCB3,PRKCA,PRKCB,PRKCD,PRKCE,PRKCG,PRKCH,PRKCI,PRKCQ,PRKCZ,PRKD1,PTK2,PXN,RAC1,RAC2,RAF1,RALA,RALB,RAP1A,RAP1B,RHOA,RHOBTB1,RHOD,RHOQ,RHOT1,ROCK1,ROCK2,RRAS2,SRC* |

| | | | | |
|---|---|---|---|---|
| Synaptic Long Term Potentiation | 8.34E+00 | 6.11E-01 | -7.506 | ADCY1,ADCY8,CACNA1C,CAMK2A,CAMK2B,CAMK2G,CAMK4,CHP1,CREB3L3,CREB5,CREBBP,EP300,GNA11,GNA14,GNAQ,GRIA1,GRIA2,GRIA4,GRIN1,GRIN2A,GRIN2B,GRIN2C,GRIN2D,GRM1,GRM3,GRM4,GRM5,GRM7,GRM8,ITPR1,ITPR2,ITPR3,KRAS,MAP2K1,MAP2K2,MAPK1,MRAS,PLCB1,PLCB2,PLCB3,PLCD3,PLCE1,PLCG1,PLCG2,PLCH1,PLCH2,PLCL1,PLCL2,PPP1R12A,PPP1R14C,PPP1R14D,PPP1R3A,PPP1R7,PPP3CB,PPP3CC,PRKACA,PRKAG2,PRKAR1B,PRKAR2A,PRKCA,PRKCB,PRKCD,PRKCE,PRKCG,PRKCH,PRKCI,PRKCQ,PRKCZ,PRKD1,RAF1,RALA,RALB,RAP1A,RAP1B,RAPGEF3,R |
| HGF Signaling | 8.22E+00 | 6.27E-01 | -8 | AKT2,AKT3,CDC42,DOCK1,ELF2,ELF4,ELF5,ELK3,ETS1,GRB2,HGF,ITGA2,ITGA3,KRAS,MAP2K1,MAP2K2,MAP2K4,MAP3K1,MAP3K10,MAP3K11,MAP3K12,MAP3K13,MAP3K14,MAP3K15,MAP3K2,MAP3K3,MAP3K5,MAP3K6,MAPK1,MAPK10,MAPK8,MAPK9,MET,MRAS,PAK1,PIK3C2A,PIK3C2B,PIK3CB,PIK3CD,PIK3R2,PIK3R3,PIK3R4,PIK3R5,PIK3R6,PLCG1,PLCG2,PRKCA,PRKCB,PRKCD,PRKCE,PRKCG,PRKCH,PRKCI,PRKCQ,PRKCZ,PRKD1,PTK2,PTPN11,PXN,RAC1,RAF1,RALA,RALB,RAP1A,RAP1B,RAP |
| Estrogen Receptor Signaling | 8.12E+00 | 5.11E-01 | -10.574 | ADCY1,ADCY10,ADCY2,ADCY3,ADCY5,ADCY6,ADCY7,ADCY8,ADCY9,AKT2,AKT3,BCL2,CACNA1A,CACNA1C,CACNA1D,CACNA1E,CACNA1S,CARM1,CREB3L3,CREB5,CREBBP,CTBP2,DLG4,EGFR,EIF2B3,EIF2B5,EP300,ESR1,ESR2,FOXO1,FOXO3,FOXO6,GNA11,GNA12,GNA14,GNAI1,GNAI2,GNAI3,GNAL,GNAO1,GNAQ,GNAS,GNB1,GNG2,GNG7,GRB2,GSK3B,HSP90AA1,IGF1R,IGF2R,JAK1,JAK2,JAK3,KRAS,LEPR,LIMK1,LIMK2,MAP2K1,MAP2K2,MAPK1,MCU,MED1,MED12L,MED13,MED13L,MED14,MED15,MED16,MED17,MED24,MED27,MMP14,MMP15,MMP17,MMP2,MMP21,MMP26,MMP28,MPRIP,MRAS,MTOR,MYL10,MYL3,NCOA1,NCOA3,NCOR1,NCOR2,NOS3,NOTCH1,NR3C1,NRF1,PAK1,PELP1,PGR,PIK3C2A,PIK3C2B,PIK3CB,PIK3CD,PIK3R2,PIK3R3,PIK3R4,PIK3R5,PIK3R6,PLCB1,PLCB2,PLCB3,PLCD3,PLCE1,PLCG1,PLCG2,PLCH1,PLCH2,PLCL1,PLCL2,PPP1R12A,PPP1R12B,PRKAA1,PRKAA2,PRKACA,PRKAG2,PRKAR1B,PRKAR2A,PRKCA,PRKCB,PRKCD,PRKCE,PRKCG,PRKCH,PRKCI,PRKCQ,PRKCZ,PRKD1,PRKDC,PTEN,RAF1,RALA,RALB,RAP1A,RAP1B,RBFOX2,RHOA,ROCK1,ROCK2,RPS6KB1,RRAS2,RUNX2,SDHC,SDHD,SETD7,SHC2,SHC3,SHF,SOS1,SP1,SRC,TBL1XR1,THRAP3,TP53,T |
| Endocannabinoid Developing Neuron Pathway | 8.01E+00 | 6.17E-01 | -5.253 | ADCY1,ADCY10,ADCY2,ADCY3,ADCY5,ADCY6,ADCY7,ADCY8,ADCY9,AKT2,AKT3,BRAF,BRCA1,CNR2,CREB3L3,CREB5,CREBBP,GNAI1,GNAI2,GNAI3,GNAL,GNAO1,GNB1,GNB1L,GNG2,GNG7,GSK3B,KRAS,MAP2K1,MAP2K2,MAP2K3,MAP2K4,MAP2K5,MAP2K6,MAPK1,MAPK10,MAPK13,MAPK14,MAPK4,MAPK6,MAPK8,MAPK9,MGLL,MRAS,MTOR,PIK3C2A,PIK3C2B,PIK3CB,PIK3CD,PIK3R2,PIK3R3,PIK3R4,PIK3R5,PIK3R6,PRKACA,PRKAG2,PRKAR1B,PRKAR2A,RAC1,RAC2,RAF1,RALA,RALB,RAP1A,RAP1B,RAP1GAP,RHOA,RPTOR,RRAS2,SRC,STAT3 |
| FcγRIIB Signaling in B Lymphocytes | 7.95E+00 | 6.80E-01 | -4.811 | BLNK,CACNA1A,CACNA1B,CACNA1C,CACNA1D,CACNA1E,CACNA1F,CACNA1G,CACNA1H,CACNA1I,CACNA1S,CACNA2D1,CACNA2D2,CACNA2D3,CACNA2D4,CACNB1,CACNB2,CACNB4,CACNG2,CACNG3,CACNG4,CACNG5,CACNG6,CACNG7,CACNG8,GRB2,INPP5D,KRAS,MAP2K4,MAPK10,MAPK8,MAPK9,MRAS,PDPK1,PIK3C2A,PIK3C2B,PIK3CB,PIK3CD,PIK3R2,PIK3R3,PIK3R4,PIK3R5,PIK3R6,PLCG2,RALA,RALB,RAP1A,RAP1B,RRAS2,SOS1,SYK |
| PTEN Signaling | 7.89E+00 | 6.03E-01 | 6.203 | AKT2,AKT3,BCAR1,BCL2,BCL2L1,BCL2L11,BMPR1A,BMPR1B,BMPR2,CASP9,CBL,CDC42,CNKSR3,CSNK2A1,CSNK2A2,EGFR,FGFR1,FGFR2,FGFR4,FLT1,FLT4,FOXO1,FOXO3,FOXO6,GHR,GRB2,GSK3B,IGF1R,IGF2R,IKBKB,IKBKE,IKBKG,INPP5B,INPP5D,INPP5F,INPP5K,INPPL1,INSR,ITGA2,ITGA3,KRAS,MAGI1,MAGI2,MAGI3,MAP2K1,MAP2K2,MAPK1,MAST2,MRAS,NTRK1,NTRK2,NTRK3,PDGFRB,PDPK1,PIK3CB,PIK3CD,PIK3R2,PIK3R3,PIK3R5,PRKCZ,PTEN,PTK2,RAC1,RAC2,RAF1,RALA,RALB,RAP1A,RAP1B,RPS6KB1,RRAS2,SOS1,SYNJ2,TGFBR2,TGFBR3,TNFRSF11A |
| Ephrin Receptor Signaling | 7.82E+00 | 5.62E-01 | -8.605 | ABL1,ACTR2,ACTR3,ADAM10,AKT2,AKT3,ARHGEF15,ARPC1A,ARPC5L,AXIN1,BCAR1,CDC42,CREB3L3,CREB5,CREBBP,CRK,CXCL12,EFNA3,EFNA5,EPHA1,EPHA10,EPHA2,EPHA3,EPHA6,EPHA8,EPHB1,EPHB2,EPHB4,FGF1,FYN,GNA11,GNA12,GNA14,GNAI1,GNAI2,GNAI3,GNAL,GNAO1,GNAQ,GNAS,GNB1,GNB1L,GNB5,GNG12,GNG2,GNG4,GNG7,GRB2,GRIN1,GRIN2A,GRIN2B,GRIN2C,GRIN2D,ITGA2,ITGA3,JAK2,KALRN,KRAS,LIMK1,LIMK2,MAP2K1,MAP2K2,MAP3K14,MAP4K4,MAPK1,MRAS,NCK1,NCK2,NGEF,PAK1,PAK2,PAK4,PDGFA,PDGFB,PDGFD,PTK2,PTPN11,PXN,RAC1,RAC2,RAF1,RALA,RALB,RAP1A,RAP1B,RAPGEF1,RASA1,RGS3,RHOA,ROCK1,ROCK2,RRAS2,SH2D3C,SORBS1,SOS1,SRC |
| Androgen Signaling | 7.77E+00 | 5.93E-01 | -6.581 | CACNA1A,CACNA1B,CACNA1C,CACNA1D,CACNA1E,CACNA1F,CACNA1G,CACNA1H,CACNA1I,CACNA1S,CACNA2D1,CACNA2D2,CACNA2D3,CACNA2D4,CACNB1,CACNB2,CACNB4,CACNG2,CACNG3,CACNG4,CACNG5,CACNG6,CACNG7,CACNG8,CAMK4,CDK7,CREBBP,EP300,ERCC2,ERCC3,GNA11,GNA12,GNA14,GNAI1,GNAI2,GNAI3,GNAL,GNAO1,GNAQ,GNAS,GNB1,GNB1L,GNB5,GNG12,GNG2,GNG4,GNG7,GTF2B,GTF2E2,GTF2H1,GTF2H3,HSP90AA1,HSPA4,KAT2B,KAT7,MAPK1,MNAT1,MRAS,NCOA1,NCOA4,POLR2A,POLR2F,POLR2J2/POLR2J3,PRKACA,PRKAG2,PRKAR1B,PRKAR2A,PRKCA,PRKCB,PRKCD,PRKCE,PRKCG,PRKCH,PRKCI,PRKCQ,PRKCZ,PRKD1,SHBG,SMAD3,SRC |
| Apelin Endothelial Signaling Pathway | 7.77E+00 | 6.14E-01 | -5.737 | ADCY1,ADCY10,ADCY2,ADCY3,ADCY5,ADCY6,ADCY7,ADCY8,ADCY9,AKT2,AKT3,ARNT,CAMK4,GNA11,GNAI1,GNAI2,GNAI3,GNAL,GNAO1,GNAS,HDAC4,HDAC5,KRAS,MAP2K1,MAP2K2,MAP2K4,MAPK1,MAPK10,MAPK8,MAPK9,MEF2A,MEF2D,MRAS,MTOR,NOS3,PIK3C2A,PIK3C2B,PIK3CB,PIK3CD,PIK3R2,PIK3R3,PIK3R4,PIK3R5,PIK3R6,PLCB1,PLCB2,PLCB3,PRKAA1,PRKAA2,PRKAG2,PRKCA,PRKCB,PRKCD,PRKCE,PRKCG,PRKCH,PRKCI,PRKCQ,PRKCZ,PRKD1,RAF1,RALA,RALB,RAP1A,RAP1B,RPS |
| RAR Activation | 7.69E+00 | 5.52E-01 | NaN | ACTB,ACTL6B,ADCY1,ADCY10,ADCY2,ADCY3,ADCY5,ADCY6,ADCY7,ADCY8,ADCY9,AKR1B1,AKT2,AKT3,ALDH1A1,ALDH1A2,ARID1A,ARID2,CARM1,CDK7,CREBBP,CSF2RB,CSK,CSNK2A1,CSNK2A2,DHRS3,DHRS7C,EP300,ERCC2,ERCC3,GTF2H1,GTF2H3,HLTF,IL3RA,JAK2,KAT2B,MAP2K1,MAP2K4,MAP3K1,MAP3K5,MAPK1,MAPK10,MAPK13,MAPK14,MAPK8,MAPK9,MAPKAPK2,MED1,MNAT1,NCOA1,NCOR1,NCOR2,NSD1,PARP1,PBRM1,PDPK1,PIK3CB,PIK3CD,PIK3R2,PIK3R3,PML,PRKACA,PRKAG2,PRKAR1B,PRKAR2A,PRKCA,PRKCB,PRKCD,PRKCE,PRKCG,PRKCH,PRKCI,PRKCQ,PRKCZ,PRKD1,PTEN,RAC1,RARA,RARB,RARG,RBP7,RDH13,REL,RELB,RXRA,RXRG,SMAD3,SMAD6,SMAD7,SMARCA2,SMARCA4,SMARCB1,SMARCC1,SMARCD1,SMARCD3,SORBS3,SRC,STAT5A,STAT5B,TAF4,TGFB |
| Cardiac Hypertrophy Signaling | 7.64E+00 | 5.32E-01 | -9.501 | ADCY1,ADCY10,ADCY2,ADCY3,ADCY5,ADCY6,ADCY7,ADCY8,ADCY9,ADRA1A,ADRA1B,CACNA1A,CACNA1C,CACNA1D,CACNA1E,CACNA1S,CAMK4,CDC42,CHP1,CREBBP,EIF2B3,EIF2B5,EP300,FNBP1,GATA4,GNA11,GNA12,GNA14,GNAI1,GNAI2,GNAI3,GNAL,GNAO1,GNAQ,GNAS,GNB1,GNB1L,GNB5,GNG12,GNG2,GNG4,GNG7,GRB2,GSK3B,IGF1R,IL6R,KRAS,MAP2K1,MAP2K2,MAP2K3,MAP2K4,MAP2K6,MAP3K1,MAP3K10,MAP3K11,MAP3K12,MAP3K13,MAP3K14,MAP3K15,MAP3K2,MAP3K3,MAP3K5,MAP3K6,MAPK1,MAPK10,MAPK13,MAPK14,MAPK8,MAPK9,MAPKAPK2,MAPKAPK3,MEF2A,MEF2D,MRAS,MTOR,MYL10,MYL3,PIK3C2A,PIK3C2B,PIK3CB,PIK3CD,PIK3R2,PIK3R3,PIK3R4,PIK3R5,PIK3R6,PLCB1,PLCB2,PLCB3,PLCD3,PLCE1,PLCG1,PLCG2,PLCH1,PLCH2,PLCL1,PLCL2,PPP3CB,PPP3CC,PRKACA,PRKAG2,PRKAR1B,PRKAR2A,RAC1,RAC2,RAF1,RALA,RALB,RAP1A,RAP1B,RHOA,RHOBTB1,RHOD,RHOQ,RHOT1,ROCK1,ROCK2,RPS6KA1,RPS6KB1,RRAS2,SOS1,TAB1,TGFB1,TGFB2,TGFB |

| | | | | |
|---|---|---|---|---|
| GABA Receptor Signaling | 7.64E+00 | 6.38E-01 | NaN | ABAT,ADCY1,ADCY10,ADCY2,ADCY3,ADCY5,ADCY6,ADCY7,ADCY8,ADCY9,AP1B1,AP2A1,AP2A2,AP2B1,CACNA1A,CACNA1B,CACNA1C,CACNA1D,CACNA1E,CACNA1F,CACNA1G,CACNA1H,CACNA1I,CACNA1S,CACNA2D1,CACNA2D2,CACNA2D3,CACNA2D4,CACNB1,CACNB2,CACNB4,CACNG2,CACNG3,CACNG4,CACNG5,CACNG6,CACNG7,CACNG8,DNM1,GABBR2,GABRA3,GABRA6,GABRB2,GABRB3,GABRG3,GABRR2,GNAS,GPHN,KCNH2,KCNN1,KCNN3,KCNN4,KCNQ2,KCNQ3,MRAS,NSF,SLC6A |
| CCR5 Signaling in Macrophages | 7.63E+00 | 6.41E-01 | -5.099 | CACNA1A,CACNA1B,CACNA1C,CACNA1D,CACNA1E,CACNA1F,CACNA1G,CACNA1H,CACNA1I,CACNA1S,CACNA2D1,CACNA2D2,CACNA2D3,CACNA2D4,CACNB1,CACNB2,CACNB4,CACNG2,CACNG3,CACNG4,CACNG5,CACNG6,CACNG7,CACNG8,CAMK4,CD247,CD4,GNAI1,GNAI2,GNAI3,GNAS,GNB1,GNB1L,GNB5,GNG12,GNG2,GNG4,GNG7,MAP2K4,MAPK1,MAPK10,MAPK13,MAPK14,MAPK8,MAPK9,MRAS,PLCG1,PLCG2,PRKCA,PRKCB,PRKCD,PRKCE,PRKCG,PRKCH,PRKCI,PRKCQ,PRKCZ,PRKD1, |
| HER-2 Signaling in Breast Cancer | 7.63E+00 | 6.55E-01 | NaN | AKT2,AKT3,CASP9,CDC42,CDK6,EGFR,ERBB2,ERBB3,FOXO1,GRB2,GSK3B,ITGB2,ITGB3,ITGB4,ITGB5,ITGB6,ITGB7,ITGB8,KRAS,MAP3K5,MDM2,MMP2,MRAS,NRG1,PARD3,PARD6B,PIK3C2A,PIK3C2B,PIK3CB,PIK3CD,PIK3R2,PIK3R3,PIK3R4,PIK3R5,PIK3R6,PLCG1,PRKCA,PRKCB,PRKCD,PRKCE,PRKCG,PRKCH,PRKCI,PRKCQ,PRKCZ,PRKD1,RALA,RALB,RAP1A,RAP1B,RRAS2,SOS1,TP53,TSC1,TSC2 |
| RhoGDI Signaling | 7.61E+00 | 5.59E-01 | 7.826 | ACTB,ACTG2,ACTR2,ACTR3,ARHGAP1,ARHGAP35,ARHGAP6,ARHGEF1,ARHGEF10,ARHGEF11,ARHGEF12,ARHGEF15,ARHGEF16,ARHGEF17,ARHGEF19,ARHGEF2,ARHGEF3,ARHGEF4,ARHGEF7,ARHGEF9,ARPC1A,ARPC5L,CD44,CDC42,CDH1,CDH10,CDH11,CDH12,CDH13,CDH15,CDH16,CDH18,CDH22,CDH23,CDH26,CDH3,CDH4,CDH5,CDH8,CDH9,CREBBP,DGKZ,DLC1,EP300,ESR1,ESR2,EZR,FNBP1,GNA11,GNA12,GNA14,GNAI1,GNAI2,GNAI3,GNAL,GNAO1,GNAQ,GNAS,GNB1,GNB1L,GNB5,GNG12,GNG2,GNG4,GNG7,GRIP1,ITGA2,ITGA3,LIMK1,LIMK2,MRAS,MSN,MYL10,MYL3,PAK1,PAK2,PAK4,PI4KA,PIP4K2B,PIP5K1A,PIP5K1B,PIP5K1C,PIP5KL1,PPP1R12A,PPP1R12B,PPP1R12C,PRKCA,RAC1,RAC2,RDX,RHOA,RHOBTB1,RHOD,RHOQ,RHOT1,ROCK1,ROCK2,SRC,WASF2 |
| Renin-Angiotensin Signaling | 7.56E+00 | 6.07E-01 | -7.81 | ACE,ADCY1,ADCY10,ADCY2,ADCY3,ADCY5,ADCY6,ADCY7,ADCY8,ADCY9,GNAQ,GRB2,ITPR1,ITPR2,ITPR3,JAK2,KRAS,MAP2K1,MAP2K2,MAP2K4,MAP3K1,MAPK1,MAPK10,MAPK13,MAPK14,MAPK8,MAPK9,MRAS,PAK1,PAK2,PAK4,PIK3C2A,PIK3C2B,PIK3CB,PIK3CD,PIK3R2,PIK3R3,PIK3R4,PIK3R5,PIK3R6,PLCG1,PLCG2,PRKACA,PRKAG2,PRKAR1B,PRKAR2A,PRKCA,PRKCB,PRKCD,PRKCE,PRKCG,PRKCH,PRKCI,PRKCQ,PRKCZ,PRKD1,PTK2,PTK2B,RAC1,RAF1,RALA,RALB,RAP1A,RAP1B,REN,RRAS2 |
| p70S6K Signaling | 7.47E+00 | 5.94E-01 | -5.735 | AKT2,AKT3,BCAP31,EEF2K,EGFR,F2,GNAI1,GNAI2,GNAI3,GNAQ,GRB2,IL4R,JAK1,KRAS,MAP2K1,MAP2K2,MAPK1,MAPT,MRAS,MTOR,PDPK1,PIK3C2A,PIK3C2B,PIK3CB,PIK3CD,PIK3R2,PIK3R3,PIK3R4,PIK3R5,PIK3R6,PLCB1,PLCB2,PLCB3,PLCD3,PLCE1,PLCG1,PLCG2,PLCH1,PLCH2,PLCL1,PLCL2,PLD1,PPM1L,PPP2CB,PPP2R1B,PPP2R2A,PPP2R2B,PPP2R2C,PPP2R3A,PPP2R5B,PPP2R5C,PPP2R5D,PPP2R5E,PRKCA,PRKCB,PRKCD,PRKCE,PRKCG,PRKCH,PRKCI,PRKCQ,PRKCZ,PRKD1,PTPA,RAF1,RALA,RALB,RAP1A,RAP1B,RPS6KB1,RRAS2,SOS1,SRC,SYK,YWHAE,YWHAG |
| Phospholipase C Signaling | 7.38E+00 | 5.24E-01 | -9.164 | ADCY1,ADCY10,ADCY2,ADCY3,ADCY5,ADCY6,ADCY7,ADCY8,ADCY9,AHNAK,ARHGEF1,ARHGEF10,ARHGEF11,ARHGEF12,ARHGEF15,ARHGEF16,ARHGEF17,ARHGEF19,ARHGEF2,ARHGEF3,ARHGEF4,ARHGEF7,ARHGEF9,BLNK,CAMK4,CD247,CDC42,CHP1,CREB3L3,CREB5,CREBBP,EP300,FNBP1,FYN,GNAQ,GNAS,GNB1,GNB1L,GNB5,GNG12,GNG2,GNG4,GNG7,GPLD1,GRB2,HDAC1,HDAC11,HDAC4,HDAC5,HDAC6,HDAC7,HDAC9,ITGA2,ITGA3,ITPR1,ITPR2,ITPR3,KRAS,LCK,LCP2,MAP2K1,MAP2K2,MAPK1,MEF2A,MEF2D,MPRIP,MRAS,MYL10,MYL3,NFAT5,NFATC1,NFATC2,NFATC3,PLA2G10,PLA2G12B,PLA2G2F,PLA2G4A,PLA2G4C,PLA2G4D,PLA2G4E,PLA2G4F,PLA2G5,PLA2G6,PLCB1,PLCB2,PLCB3,PLCD3,PLCE1,PLCG1,PLCG2,PLD1,PLD2,PLD3,PLD5,PPP1R12A,PPP1R12B,PPP3CB,PPP3CC,PRKCA,PRKCB,PRKCD,PRKCE,PRKCG,PRKCH,PRKCI,PRKCQ,PRKCZ,PRKD1,RAC1,RAC2,RAF1,RALA,RALB,RALGDS,RAP1A,RAP1B,RAPGEF3,RHOA,RHOBTB1,RHOD,RHOQ,RHOT1,RPS6KA3,RRA |
| PKCθ Signaling in T Lymphocytes | 7.37E+00 | 5.73E-01 | -7.81 | CACNA1A,CACNA1B,CACNA1C,CACNA1D,CACNA1E,CACNA1F,CACNA1G,CACNA1H,CACNA1I,CACNA1S,CACNA2D1,CACNA2D2,CACNA2D3,CACNA2D4,CACNB1,CACNB2,CACNB4,CACNG2,CACNG3,CACNG4,CACNG5,CACNG6,CACNG7,CACNG8,CAMK2A,CAMK2B,CAMK2G,CARD11,CD247,CD4,CHP1,FYN,GRB2,IKBKB,IKBKE,IKBKG,KRAS,LCK,LCP2,MAP2K4,MAP3K1,MAP3K10,MAP3K11,MAP3K12,MAP3K13,MAP3K14,MAP3K15,MAP3K2,MAP3K3,MAP3K5,MAP3K6,MAPK1,MAPK8,MRAS,NFAT5,NFATC1,NFATC2,NFATC3,NFKBID,PIK3C2A,PIK3C2B,PIK3CB,PIK3CD,PIK3R2,PIK3R3,PIK3R4,PIK3R5,PIK3R6,PLCG1,PLCG2,POU2F1,PPP3CB,PPP3CC,PRKCQ,RAC1,RAC2,RALA,RALB,RAP1A,RAP1B,RRAS2, |
| Gαq Signaling | 7.22E+00 | 5.69E-01 | -8.03 | ADRA1A,ADRA1B,AKT2,AKT3,CALCR,CAMK4,CDC42,CHP1,CHRM3,CSK,EPM2A,FNBP1,GNA11,GNA14,GNAQ,GNAS,GNB1,GNB1L,GNB5,GNG12,GNG2,GNG4,GNG7,GPLD1,GRK2,GRM1,GRM5,GSK3B,GYS1,HRH1,HTR2C,IKBKB,IKBKE,IKBKG,ITPR1,ITPR2,ITPR3,MAP2K1,MAP2K2,MAPK1,MRAS,NFATC1,NFATC2,NFATC3,NFKBID,PIK3C2A,PIK3C2B,PIK3CB,PIK3CD,PIK3R2,PIK3R3,PIK3R4,PIK3R5,PIK3R6,PLCB1,PLCB2,PLCB3,PLCG1,PLCG2,PLD1,PLD2,PLD3,PLD5,PPP3CB,PPP3CC,PRKCA,PRKCB,PRKCD,PRKCE,PRKCG,PRKCH,PRKCI,PRKCQ,PRKCZ,PRKD1,PTK2B,RAC1,RAC2,RAF1,RGS7,RHOA,RH |
| Cardiac β-adrenergic Signaling | 7.21E+00 | 5.79E-01 | -3.457 | ADCY1,ADCY10,ADCY2,ADCY3,ADCY5,ADCY6,ADCY7,ADCY8,ADCY9,AKAP1,AKAP10,AKAP12,AKAP13,AKAP8,AKAP9,ATP2A1,ATP2A2,ATP2A3,CACNA1A,CACNA1C,CACNA1D,CACNA1E,CACNA1S,ENPP6,GDPD1,GNAS,GNB1,GNB1L,GNB5,GNG12,GNG2,GNG4,GNG7,GRK2,GRK3,MPPED2,MRAS,PDE10A,PDE11A,PDE1A,PDE1B,PDE1C,PDE2A,PDE3A,PDE3B,PDE4A,PDE4B,PDE4C,PDE4D,PDE6A,PDE6B,PDE6C,PDE6G,PDE8A,PDE8B,PDE9A,PKIG,PPM1L,PPP1R12A,PPP1R14C,PPP1R14D,PPP1R3A,PPP1R7,PPP2CB,PPP2R1B,PPP2R2A,PPP2R2B,PPP2R2C,PPP2R3A,PPP2R5B,PPP2R5C,PPP2R5D,PPP2R5E,PRKACA,PRKAG2,PRKAR1B,PRKAR2A,PTPA,RYR2,SLC8A1,SLC8A2 |
| Apelin Cardiomyocyte Signaling Pathway | 7.16E+00 | 6.22E-01 | -7.353 | AKT2,AKT3,ARNT,ATP2A1,ATP2A2,ATP2A3,GNA11,GNAI1,GNAI2,GNAI3,ITPR1,MAPK1,MAPK10,MAPK13,MAPK14,MAPK4,MAPK6,MAPK8,MAPK9,MYL10,MYL3,MYLK,NOS3,PIK3C2A,PIK3C2B,PIK3CB,PIK3CD,PIK3R2,PIK3R3,PIK3R4,PIK3R5,PIK3R6,PLCB1,PLCB2,PLCB3,PLCD3,PLCE1,PLCG1,PLCG2,PLCH1,PLCH2,PLCL1,PLCL2,PRKCA,PRKCB,PRKCD,PRKCE,PRKCG,PRKCH,PRKCI,PRKCQ,PRKCZ,PRKD1,SLC8A1,SLC8A2,SLC9A1,SLC9A2,SLC9A3,SLC9A5,SLC9A8,TGFB1 |
| B Cell Receptor Signaling | 7.12E+00 | 5.50E-01 | -7.49 | ABL1,AKT2,AKT3,BCL2L1,BLNK,CAMK2A,CAMK2B,CAMK2G,CAMK4,CARD10,CD22,CDC42,CREB3L3,CREB5,CREBBP,CSK,EBF1,ETS1,FOXO1,GAB2,GRB2,GSK3B,IKBKB,IKBKE,IKBKG,INPP5B,INPP5D,INPP5F,INPP5K,INPPL1,KRAS,MAP2K1,MAP2K2,MAP2K3,MAP2K4,MAP2K6,MAP3K1,MAP3K10,MAP3K11,MAP3K12,MAP3K13,MAP3K14,MAP3K15,MAP3K2,MAP3K3,MAP3K5,MAP3K6,MAPK1,MAPK13,MAPK14,MAPK8,MAPK9,MRAS,MTOR,NFAT5,NFATC1,NFATC2,NFATC3,NFKBID,PAX5,PDPK1,PIK3AP1,PIK3C2A,PIK3C2B,PIK3CB,PIK3CD,PIK3R2,PIK3R3,PIK3R4,PIK3R5,PIK3R6,PLCG2,POU2F2,PPP3CB,PPP3CC,PRKCB,PRKCQ,PTEN,PTK2,PTK2B,PTPN11,PTPRC,RAC1,RAC2,RAF1,RALA,RALB,RAP1A,RAP1B,RASSF5,RPS6KB1,RRAS2,SOS1,SYK,SYNJ2,TCF3,VAV1,VAV2,VAV3 |

| | | | | |
|---|---|---|---|---|
| Fc Epsilon RI Signaling | 7.12E+00 | 5.98E-01 | -6.932 | AKT2,AKT3,FYN,GRB2,INPP5B,INPP5D,INPP5F,INPP5K,INPPL1,KRAS,LCP2,MAP2K1,MAP2K2,MAP2K3,MAP2K4,MAP2K6,MAPK1,MAPK10,MAPK13,MAPK14,MAPK8,MAPK9,MRAS,PDPK1,PIK3C2A,PIK3C2B,PIK3CB,PIK3CD,PIK3R2,PIK3R3,PIK3R4,PIK3R5,PIK3R6,PLA2G10,PLA2G12B,PLA2G2F,PLA2G4A,PLA2G4C,PLA2G4D,PLA2G4E,PLA2G4F,PLA2G5,PLA2G6,PLCG1,PLCG2,PRKCA,PRKCB,PRKCD,PRKCE,PRKCG,PRKCH,PRKCI,PRKCQ,PRKCZ,PRKD1,PTPN11,RAC1,RAC2,RAF1,RALA,RALB,RAP1A,RAP1B,RRAS2,SOS1,SYK,SYNJ2,VAV1,VAV2,VAV3 |
| Macropinocytosis Signaling | 7.09E+00 | 6.58E-01 | -6.083 | ACTN4,ANKFY1,CDC42,CSF1,CSF1R,HGF,ITGB2,ITGB3,ITGB4,ITGB5,ITGB6,ITGB7,ITGB8,KRAS,MET,MRAS,PAK1,PDGFA,PDGFB,PDGFD,PIK3C2A,PIK3C2B,PIK3CB,PIK3CD,PIK3R2,PIK3R3,PIK3R4,PIK3R5,PIK3R6,PLCG1,PLCG2,PRKCA,PRKCB,PRKCD,PRKCE,PRKCG,PRKCH,PRKCI,PRKCQ,PRKCZ,PRKD1,RAC1,RALA,RALB,RAP1A,RAP1B,RHOA,RRAS2,SRC,USP6NL |
| PPARα/RXRα Activation | 7.09E+00 | 5.51E-01 | 1.709 | ABCA1,ACOX1,ACVR1B,ACVR2B,ADCY1,ADCY10,ADCY2,ADCY3,ADCY5,ADCY6,ADCY7,ADCY8,ADCY9,AP2A2,BMPR2,CD36,CHD5,CKAP5,CREBBP,CYP2C19,EP300,GHR,GK,GNA11,GNA14,GNAQ,GNAS,GOT2,GPD1,GRB2,HELZ2,HSP90AA1,IKBKB,IKBKE,IKBKG,IL1R1,IL1RAP,IL1RAPL1,IL1RAPL2,INSR,ITGB5,JAK2,KRAS,LPL,MAP2K1,MAP2K2,MAP2K3,MAP2K4,MAP2K6,MAP3K14,MAP4K4,MAPK1,MAPK14,MAPK8,MED1,MED24,MRAS,NCOA3,NCOA6,NCOR1,NCOR2,NFKBID,NR2C2,PLCB1,PLCB2,PLCB3,PLCD3,PLCE1,PLCG1,PLCG2,PLCH1,PLCH2,PLCL1,PLCL2,PPARA,PRKAA1,PRKAA2,PRKACA,PRKAG2,PRKAR1B,PRKAR2A,PRKCA,PRKCB,RAF1,RALA,RALB,RAP1A,RAP1B,RRAS2,RXRA,SLC27A1,SMAD3,SOS1,STAT5B,TGFB1,TGFB2,TGFBR2,TGFBR3 |
| NGF Signaling | 7.07E+00 | 6.02E-01 | -8.124 | AKT2,AKT3,CDC42,CREB3L3,CREB5,CREBBP,CRK,GRB2,IKBKB,IKBKE,IKBKG,KRAS,MAP2K1,MAP2K2,MAP2K4,MAP3K1,MAP3K10,MAP3K11,MAP3K12,MAP3K13,MAP3K14,MAP3K15,MAP3K2,MAP3K3,MAP3K5,MAP3K6,MAPK1,MAPK10,MAPK8,MAPK9,MRAS,NTRK1,PDPK1,PIK3C2A,PIK3C2B,PIK3CB,PIK3CD,PIK3R2,PIK3R3,PIK3R4,PIK3R5,PIK3R6,PLCG1,PLCG2,PRKCD,PRKCZ,PTPN11,RAC1,RAF1,RALA,RALB,RAP1A,RAP1B,RHOA,ROCK1,ROCK2,RPS6KA1,RPS6KA2,RPS6KA3,RPS6KA5,RPS6KB1,RPS6KC1,RRAS2,SMPD3,SMPD4,SOS1,TP53,TRIO |
| P2Y Purigenic Receptor Signaling Pathway | 7.02E+00 | 5.87E-01 | -7.519 | ADCY1,ADCY10,ADCY2,ADCY3,ADCY5,ADCY6,ADCY7,ADCY8,ADCY9,AKT2,AKT3,CREB3L3,CREB5,CREBBP,GNAI1,GNAI2,GNAI3,GNAQ,GNB1,GNB1L,GNB5,GNG12,GNG2,GNG4,GNG7,ITGA2B,ITGB3,KRAS,MAP2K1,MAP2K2,MAPK1,MRAS,P2RY2,P2RY6,PIK3C2A,PIK3C2B,PIK3CB,PIK3CD,PIK3R2,PIK3R3,PIK3R4,PIK3R5,PIK3R6,PLCB1,PLCB2,PLCB3,PLCD3,PLCE1,PLCG1,PLCG2,PLCH1,PLCH2,PLCL1,PLCL2,PRKACA,PRKAG2,PRKAR1B,PRKAR2A,PRKCA,PRKCB,PRKCD,PRKCE,PRKCG,PRKCH,PRKCI,PRKCQ,PRKCZ,PRKD1,RAF1,RALA,RALB,RAP1A,RAP1B,RRAS2 |
| Virus Entry via Endocytic Pathways | 7.01E+00 | 6.07E-01 | NaN | ABL1,ACTB,ACTG2,AP1B1,AP1G1,AP1M1,AP2A1,AP2A2,AP2B1,AP3B1,AP3M1,CDC42,CLTA,CLTB,CLTC,CLTCL1,CXADR,DNM1,DNM2,FLNA,FLNB,FLNC,FYN,ITGA2,ITGA3,ITGAL,ITGB2,ITGB3,ITGB4,ITGB5,ITGB6,ITGB7,ITGB8,KRAS,MRAS,PIK3C2A,PIK3C2B,PIK3CB,PIK3CD,PIK3R2,PIK3R3,PIK3R4,PIK3R5,PIK3R6,PLCG1,PLCG2,PRKCA,PRKCB,PRKCD,PRKCE,PRKCG,PRKCH,PRKCI,PRKCQ,PRKCZ,PRKD1,RAC1,RAC2,RALA,RALB,RAP1A,RAP1B,RRAS2,SRC,TFRC |
| IL-3 Signaling | 6.82E+00 | 6.46E-01 | -6.581 | AKT2,AKT3,CHP1,CSF2RB,FOXO1,GAB2,GRB2,IL3RA,INPP5D,JAK1,JAK2,KRAS,MAP2K1,MAP2K2,MAPK1,MRAS,PAK1,PIK3C2A,PIK3C2B,PIK3CB,PIK3CD,PIK3R2,PIK3R3,PIK3R4,PIK3R5,PIK3R6,PPP3CB,PPP3CC,PRKCA,PRKCB,PRKCD,PRKCE,PRKCG,PRKCH,PRKCI,PRKCQ,PRKCZ,PRKD1,RAC1,RAF1,RALA,RALB,RAP1A,RAP1B,RAPGEF1,RRAS2,SOS1,STAT3,STAT5A,STAT5B,STAT6 |
| Senescence Pathway | 6.71E+00 | 5.07E-01 | -7.544 | ACVR1B,ACVR2B,AKT2,AKT3,ANAPC1,ANAPC5,ASXL1,ASXL2,ATF3,ATM,ATR,BMPR2,BRAF,CACNA1A,CACNA1C,CACNA1D,CACNA1E,CACNA1S,CAMK4,CAPN1,CAPN11,CAPN2,CAPN5,CAPN8,CBX7,CCNB3,CDC16,CDC23,CDC25A,CDC25C,CDC27,CDK6,CHP1,CREBBP,DHCR24,DLAT,E2F1,E2F2,E2F3,E2F4,ELF2,ELF4,ELF5,ELK3,EP300,EP400,ETS1,EZH2,FOXO3,GATA4,HIPK2,IKBKB,IKBKE,IKBKG,ITPR2,ITPR3,ITSN2,KAT2B,KRAS,MAP2K1,MAP2K2,MAP2K3,MAP2K4,MAP2K5,MAP2K6,MAPK1,MAPK14,MAPK4,MAPK6,MAPKAPK2,MAPKAPK3,MAPKAPK5,MCU,MDM2,MRAS,MRE11,MTOR,NF1,NFAT5,NFATC1,NFATC2,NFATC3,PARP1,PCGF2,PCGF6,PDHX,PDK2,PDK3,PHF19,PIK3C2A,PIK3C2B,PIK3CB,PIK3CD,PIK3R2,PIK3R3,PIK3R4,PIK3R5,PIK3R6,PML,PPM1L,PPP2CB,PPP2R1B,PPP2R2A,PPP2R2B,PPP2R2C,PPP2R3A,PPP2R5B,PPP2R5C,PPP2R5D,PPP2R5E,PPP3CB,PPP3CC,PRKCZ,PTEN,PTPA,RAD50,RAF1,RALA,RALB,RAP1A,RAP1B,RASSF5,RBL1,RNF2,RPS6KA5,RRAS2,SERPINE1,SIRT1,SMAD3,SMAD6,SMAD7,SQSTM1,TBK1,TGFB1,TGFB2,TGFBR2,TGFBR3,TP53 |
| Factors Promoting Cardiogenesis in Vertebrates | 6.70E+00 | 5.66E-01 | -8.213 | ACVR1B,ACVR2B,APC,AXIN1,BMP1,BMP5,BMP6,BMP7,BMP8A,BMP8B,BMPR1A,BMPR1B,BMPR2,CAMK2A,CAMK2B,CAMK2G,CREB3L3,CREB5,CREBBP,FZD3,GATA4,GSK3B,HNF1A,LRP1,LRP5,LRP6,MAP2K4,MAPK10,MAPK14,MAPK8,MAPK9,MYH6,MYH7,NOX4,PLCB1,PLCB2,PLCB3,PLCD3,PLCE1,PLCG1,PLCG2,PLCH1,PLCH2,PLCL1,PLCL2,PRKCA,PRKCB,PRKCD,PRKCE,PRKCG,PRKCH,PRKCI,PRKCQ,PRKCZ,PRKD1,RHOA,ROCK1,ROCK2,SCN5A,SMO,TBX5,TCF3,TCF4,TCF7,TCF7L1,TCF7L2,TDGF1,TGFB1,TGFB2,TGFBR2,TGFBR3,WNT10A,WNT11,WNT2B,WNT3,WNT3A,WNT4,WNT5B,WNT7A,WNT |
| ATM Signaling | 6.67E+00 | 6.15E-01 | -0.283 | ABL1,ATF1,ATM,ATR,BID,BLM,BRAT1,BRCA1,CBX1,CBX5,CCNB3,CDC25A,CDC25C,CREB3L3,CREB5,CREBBP,FANCD2,HERC2,HP1BP3,MAP2K4,MAPK10,MAPK13,MAPK14,MAPK8,MAPK9,MDM2,MDM4,MRE11,PPM1D,PPM1L,PPP2CB,PPP2R1B,PPP2R2A,PPP2R2B,PPP2R2C,PPP2R3A,PPP2R5B,PPP2R5C,PPP2R5D,PPP2R5E,PTPA,RAD17,RAD50,RAD51,RBBP8,RNF168,RNF8,SMC1A,SMC1B,SMC3,TLK1,TLK2,TOPBP1,TP53,TP53BP1,TP73,TRRAP,USP7,ZNF420 |
| Superpathway of Inositol Phosphate Compounds | 6.66E+00 | 5.36E-01 | -10.1 | ACP6,ALPL,CDC25A,CDC25C,CILP,DOT1L,DUSP10,DUSP14,DUSP15,DUSP16,DUSP27,EPHX2,EYA4,HACD2,INPP5A,INPP5B,INPP5D,INPP5F,INPP5K,INPPL1,IP6K1,IP6K2,IP6K3,IPMK,ITPK1,ITPKB,ITPKC,MTMR4,MTMR7,PALD1,PAWR,PI4K2A,PI4KA,PI4KB,PIK3C2A,PIK3C2B,PIK3CB,PIK3CD,PIK3R2,PIK3R3,PIK3R4,PIK3R5,PIK3R6,PIP4K2A,PIP4K2B,PIP5K1A,PIP5K1B,PIP5K1C,PIP5KL1,PLCB1,PLCB2,PLCB3,PLCD3,PLCE1,PLCG1,PLCG2,PLCH1,PLPP7,PPFIA2,PPFIA3,PPFIA4,PPFIBP2,PPIP5K1,PPIP5K2,PPM1H,PPP1R12A,PPP1R12C,PPP1R13B,PPP1R16B,PPP1R1C,PPP1R7,PPP1R8,PPP2R3A,PPP2R5B,PPP2R5D,PPP2R5E,PPP4C,PPP5C,PPTC7,PRUNE1,PTEN,PTPA,PTPN1,PTPN11,PTPN12,PTPN2,PTPN20,PTPN22,PTPN23,PTPN7,PTPRC,PTPRF,PTPRH,PTPRJ,PTPRM,PXYLP1,RASA1,SEC16A,SIRPA,S |
| ERK/MAPK Signaling | 6.66E+00 | 5.36E-01 | -6.2 | ATF1,BCAR1,BRAF,CREB3L3,CREB5,CREBBP,CRK,DOCK1,DUSP4,ELF2,ELF4,ELF5,ELK3,ESR1,ETS1,FYN,GRB2,ITGA2,ITGA3,KRAS,KSR1,MAP2K1,MAP2K2,MAPK1,MAPKAPK5,MKNK1,MRAS,NFATC1,PAK1,PAK2,PAK4,PIK3C2A,PIK3C2B,PIK3CB,PIK3CD,PIK3R2,PIK3R3,PIK3R4,PIK3R5,PIK3R6,PLA2G10,PLA2G12B,PLA2G2F,PLA2G4A,PLA2G4C,PLA2G4D,PLA2G4E,PLA2G4F,PLA2G5,PLA2G6,PLCG1,PLCG2,PPARG,PPM1L,PPP1R12A,PPP1R14C,PPP1R14D,PPP1R3A,PPP1R7,PPP2CB,PPP2R1B,PPP2R2A,PPP2R2B,PPP2R2C,PPP2R3A,PPP2R5B,PPP2R5C,PPP2R5D,PPP2R5E,PRKACA,PRKAG2,PRKAR1B,PRKAR2A,PRKCA,PRKCB,PRKCD,PRKCE,PRKCG,PRKCI,PTK2,PTK2B,PTPA,PXN,RAC1,RAC2,RAF1,RALA,RALB,RAP1A,RAP1B,RAPGEF1,RAPGEF3,RAPGEF4,RPS6KA1,RPS6KA5,RRAS2,SOS1,SRC, |

| | | | | |
|---|---|---|---|---|
| FAK Signaling | 6.66E+00 | 6.17E-01 | NaN | ACTB,ACTG2,AKT2,AKT3,ARHGAP26,ARHGEF7,ASAP1,BCAR1,CAPN1,CAPN11,CAPN2,CAPN5,CAPN8,CRK,CSK,DOCK1,EGFR,FYN,GRB2,ITGA2,ITGA3,KRAS,MAP2K1,MAP2K2,MAPK1,MRAS,PAK1,PAK2,PAK4,PDPK1,PIK3C2A,PIK3C2B,PIK3CB,PIK3CD,PIK3R2,PIK3R3,PIK3R4,PIK3R5,PIK3R6,PLCG1,PLCG2,PTEN,PTK2,PXN,RAC1,RAF1,RALA,RALB,RAP1A,RAP1B,RRAS2,SOS1,SRC,TLN1,TLN2,TNS1,V |
| RANK Signaling in Osteoclasts | 6.60E+00 | 6.28E-01 | -7.211 | AKT2,AKT3,BIRC2,CAMK4,CBL,CHP1,GSN,IKBKB,IKBKE,IKBKG,MAP2K1,MAP2K2,MAP2K4,MAP2K6,MAP3K1,MAP3K10,MAP3K11,MAP3K12,MAP3K13,MAP3K14,MAP3K15,MAP3K2,MAP3K3,MAP3K5,MAP3K6,MAPK1,MAPK10,MAPK13,MAPK14,MAPK8,MAPK9,MITF,NFATC1,NFATC2,NFKBID,PIK3C2A,PIK3C2B,PIK3CB,PIK3CD,PIK3R2,PIK3R3,PIK3R4,PIK3R5,PIK3R6,PPP3CB,PPP3CC,PTK2B,RAF1,SRC,TAB2,TNFRSF11A,TRAF2,TRAF5,XIAP |
| VEGF Family Ligand-Receptor Interactions | 6.59E+00 | 6.31E-01 | -7.141 | AKT2,AKT3,FLT1,FLT4,GRB2,KRAS,MAP2K1,MAP2K2,MAPK1,MRAS,NOS3,NRP1,NRP2,PIK3C2A,PIK3C2B,PIK3CB,PIK3CD,PIK3R2,PIK3R3,PIK3R4,PIK3R5,PIK3R6,PLA2G10,PLA2G12B,PLA2G2F,PLA2G4A,PLA2G4C,PLA2G4D,PLA2G4E,PLA2G4F,PLA2G5,PLA2G6,PLCG1,PLCG2,PRKCA,PRKCB,PRKCD,PRKCE,PRKCG,PRKCH,PRKCI,PRKCQ,PRKCZ,PRKD1,RAF1,RALA,RALB,RAP1A,RAP1B,RRAS2,SOS1, |
| Melatonin Signaling | 6.55E+00 | 6.57E-01 | -5.729 | BRAF,CAMK2A,CAMK2B,CAMK2G,CAMK4,GNAI1,GNAI2,GNAI3,GNAO1,GNAQ,MAP2K1,MAP2K2,MAP2K3,MAP2K4,MAP2K5,MAP2K6,MAPK1,PLCB1,PLCB2,PLCB3,PLCD3,PLCE1,PLCG1,PLCG2,PLCH1,PLCH2,PLCL1,PLCL2,PRKACA,PRKAG2,PRKAR1B,PRKAR2A,PRKCA,PRKCB,PRKCD,PRKCE,PRKCG,PRKCH,PRKCI,PRKCQ,PRKCZ,PRKD1,RAF1,RORA,RORB,RORC |
| PI3K/AKT Signaling | 6.47E+00 | 5.43E-01 | -1.588 | AKT2,AKT3,BCL2,BCL2L1,CDC37,CSF2RB,EPM2A,FOXO1,FOXO3,GAB2,GDF15,GHR,GRB2,GSK3B,GYS1,HSP90AA1,IFNLR1,IKBKB,IKBKE,IKBKG,IL12RB1,IL12RB2,IL15RA,IL17RA,IL17RC,IL17RD,IL1R1,IL21R,IL22RA1,IL27RA,IL2RA,IL2RB,IL31RA,IL3RA,IL4R,IL6R,IL6ST,INPP5B,INPP5D,INPP5F,INPP5K,INPPL1,ITGA2,ITGA3,JAK1,JAK2,JAK3,KRAS,LIMS1,MAP2K1,MAP2K2,MAP3K5,MAPK1,MAPK8IP1,MDM2,MRAS,MTOR,NFKBID,NOS3,PDPK1,PIK3CB,PIK3CD,PIK3R2,PIK3R3,PPM1L,PPP2CB,PPP2R1B,PPP2R2A,PPP2R2B,PPP2R2C,PPP2R3A,PPP2R5B,PPP2R5C,PPP2R5D,PPP2R5E,PTEN,PTPA,RAF1,RALA,RALB,RAP1A,RAP1B,RHEB,RPS6KB1,RRAS2,SOS1,SYNJ2,THEM4,TP53,TSC1,TSC2,TYK2,YWHAE, |
| Insulin Receptor Signaling | 6.36E+00 | 5.66E-01 | -4.417 | ACLY,AKT2,AKT3,ASIC1,ASIC2,CBL,CRK,EIF2B3,EIF2B5,EPM2A,FOXO1,FOXO3,FYN,GRB10,GRB2,GSK3B,GYS1,INPP5B,INPP5D,INPP5F,INPP5K,INPPL1,INSR,IRS2,JAK1,JAK2,KRAS,MAP2K1,MAP2K2,MAPK1,MAPK8,MRAS,MTOR,NCK1,PDE3B,PDPK1,PIK3C2A,PIK3C2B,PIK3CB,PIK3CD,PIK3R2,PIK3R3,PIK3R4,PIK3R5,PIK3R6,PPP1R12A,PPP1R14C,PPP1R14D,PPP1R3A,PPP1R7,PRKACA,PRKAG2,PRKAR1B,PRKAR2A,PRKCI,PRKCZ,PTEN,PTPN1,PTPN11,PTPRF,RAF1,RALA,RALB,RAP1A,RAP1B,RAPGEF1,RHOQ,RPS6KB1,RPTOR,RRAS2,SCNN1B,SGK1,SOS1,STXBP4,SYNJ2,TSC1,TSC2 |
| Telomerase Signaling | 6.30E+00 | 5.96E-01 | -3.286 | ABL1,AKT2,AKT3,E2F1,EGFR,ELF2,ELF4,ELF5,ELK3,ETS1,GRB2,HDAC1,HDAC11,HDAC4,HDAC5,HDAC6,HDAC7,HDAC9,HSP90AA1,IL2RA,IL2RB,KRAS,MAP2K1,MAP2K2,MAPK1,MRAS,PDPK1,PIK3C2A,PIK3C2B,PIK3CB,PIK3CD,PIK3R2,PIK3R3,PIK3R4,PIK3R5,PIK3R6,PPM1L,PPP2CB,PPP2R1B,PPP2R2A,PPP2R2B,PPP2R2C,PPP2R3A,PPP2R5B,PPP2R5C,PPP2R5D,PPP2R5E,PTGES3,PTPA,RAF1,RALA,RALB,RAP1A,RAP1B,RRAS2,SOS1,SP1,TEP1,TERF2,TERF2IP,TERT,TP53 |
| Thrombopoietin Signaling | 6.30E+00 | 6.67E-01 | -6.481 | GAB2,GRB2,IRS2,JAK2,KRAS,MAP2K1,MAP2K2,MAPK1,MPL,MRAS,PIK3C2A,PIK3C2B,PIK3CB,PIK3CD,PIK3R2,PIK3R3,PIK3R4,PIK3R5,PIK3R6,PLCG1,PLCG2,PRKCA,PRKCB,PRKCD,PRKCE,PRKCG,PRKCH,PRKCI,PRKCQ,PRKCZ,PRKD1,PTPN11,RAF1,RALA,RALB,RAP1A,RAP1B,RRAS2,SOS1,STAT3,ST |
| Non-Small Cell Lung Cancer Signaling | 6.29E+00 | 6.44E-01 | -4.644 | ABL1,AKT2,AKT3,CASP9,CDK6,E2F1,EGFR,ERBB2,FHIT,FOXO3,GRB2,ITPR1,ITPR2,ITPR3,KRAS,MAP2K1,MAP2K2,MAPK1,MRAS,PDPK1,PIK3C2A,PIK3C2B,PIK3CB,PIK3CD,PIK3R2,PIK3R3,PIK3R4,PIK3R5,PIK3R6,PLCG1,PRKCA,RAF1,RALA,RALB,RAP1A,RAP1B,RARB,RASSF5,RRAS2,RXRA,RXRG,SIN3A,SOS1,STK4,TFDP1,TGFA,TP53 |
| Nitric Oxide Signaling in the Cardiovascular System | 6.19E+00 | 6.06E-01 | -6.337 | AKT2,AKT3,ATP2A1,ATP2A2,ATP2A3,CACNA1A,CACNA1C,CACNA1D,CACNA1E,CACNA1S,CAMK4,FLT1,FLT4,GUCY1A2,GUCY2D,HSP90AA1,ITPR1,ITPR2,ITPR3,MAP2K1,MAP2K2,MAPK1,NOS3,PDE1A,PDE1B,PDE1C,PDE2A,PDE3B,PIK3C2A,PIK3C2B,PIK3CB,PIK3CD,PIK3R2,PIK3R3,PIK3R4,PIK3R5,PIK3R6,PRKAA1,PRKACA,PRKAG2,PRKAR1B,PRKAR2A,PRKCA,PRKCB,PRKCD,PRKCE,PRKCG,PRKCH,PRKCI,PRKCQ,PRKCZ,PRKD1,PRKG1,RYR2,SLC7A1,VEGFA,VEGFD |
| STAT3 Pathway | 6.16E+00 | 5.63E-01 | -7.181 | BCL2,BMP6,BMPR1A,BMPR1B,BMPR2,CDC25A,CSF2RB,EGFR,FGFR1,FGFR2,FGFR4,FLT1,FLT4,GHR,HGF,IFNLR1,IGF1R,IGF2R,IL12RB1,IL12RB2,IL15RA,IL17RA,IL17RC,IL17RD,IL1R1,IL21R,IL22RA1,IL27RA,IL2RA,IL2RB,IL31RA,IL3RA,IL4R,IL6R,IL6ST,INSR,JAK2,KRAS,MAP2K1,MAP2K2,MAP2K4,MAP3K10,MAP3K11,MAP3K12,MAPK1,MAPK10,MAPK13,MAPK14,MAPK8,MAPK9,MRAS,NTRK1,NTRK2,NTRK3,PDGFB,PDGFRB,PTPN2,RAC1,RAF1,RALA,RALB,RAP1A,RAP1B,RRAS2,SOCS5,SOCS7,SRC,STAT3,TGFA,TGFB1,TGFB2,TGFBR2,TGFBR3,TNFRSF11A,TYK2,VEGFA |
| ErbB4 Signaling | 6.01E+00 | 6.52E-01 | -6.325 | ERBB4,GRB2,KRAS,MAP2K1,MAP2K2,MAPK1,MRAS,NRG1,NRG2,NRG3,NRG4,PDPK1,PIK3C2A,PIK3C2B,PIK3CB,PIK3CD,PIK3R2,PIK3R3,PIK3R4,PIK3R5,PIK3R6,PLCG1,PLCG2,PRKCA,PRKCB,PRKCD,PRKCE,PRKCG,PRKCH,PRKCI,PRKCQ,PRKCZ,PRKD1,PSEN1,PSEN2,RAF1,RALA,RALB,RAP1A,RAP1 |
| Colorectal Cancer Metastasis Signaling | 5.91E+00 | 5.04E-01 | -8.178 | ADCY1,ADCY10,ADCY2,ADCY3,ADCY5,ADCY6,ADCY7,ADCY8,ADCY9,AKT2,AKT3,APC,APPL1,ARRB1,AXIN1,BCL2L1,BRAF,CASP9,CDC42,CDH1,DCC,E2F4,EGFR,FNBP1,FZD3,GNAS,GNB1,GNB1L,GNB5,GNG12,GNG2,GNG4,GNG7,GRB2,GRK2,GRK3,GSK3B,HNF1A,IL6R,IL6ST,JAK1,JAK2,JAK3,KRAS,LRP1,LRP5,LRP6,MAP2K1,MAP2K2,MAP2K4,MAPK1,MAPK10,MAPK8,MAPK9,MLH1,MMP14,MMP15,MMP17,MMP2,MMP21,MMP26,MMP28,MRAS,MSH2,MSH3,MSH6,NOS2,PIK3C2A,PIK3C2B,PIK3CB,PIK3CD,PIK3R2,PIK3R3,PIK3R4,PIK3R5,PIK3R6,PRKACA,PRKAG2,PRKAR1B,PRKAR2A,PTGER3,RAC1,RAC2,RALA,RALB,RALGDS,RAP1A,RAP1B,RHOA,RHOBTB1,RHOD,RHOQ,RHOT1,RRAS2,SIAH1,SMAD3,SMO,SOS1,SRC,STAT3,TCF3,TCF4,TCF7,TCF7L1,TCF7L2,TGFB1,TGFB2,TGFBR2,TLR5,TP53,TYK2,VEGFA,VEGFD,WNT10A,WNT11,WNT2B,WNT3,WNT3A,WNT4,WNT5B,WNT7A,WNT7B,WNT9A,WNT9 |
| Epithelial Adherens Junction Signaling | 5.88E+00 | 5.47E-01 | NaN | ACTB,ACTG2,ACTN1,ACTN2,ACTN4,ACTR2,ACTR3,ACVR1B,ACVR2B,AKT2,AKT3,APC,ARPC1A,ARPC5L,BAIAP2,BMPR2,CDC42,CDH1,CLIP1,CRK,CTNNA1,CTNNA2,EGFR,EPN1,EPN2,FARP2,FER,FGF1,FGFR1,HGF,HNF1A,IQGAP1,JUP,KEAP1,KRAS,MAGI1,MAGI2,MET,MRAS,MYH10,MYH11,MYH14,MYH3,MYH6,MYH7,MYH7B,MYH9,MYL3,MYO7A,NECTIN1,NECTIN2,NECTIN3,NOTCH1,NOTCH2,NOTCH3,PARD3,PTEN,PTPRM,RAC1,RALA,RALB,RAP1A,RAP1B,RAPGEF1,RHOA,RRAS2,SORBS1,SRC,TCF3,TCF4,TCF7,TCF7L1,TCF7L2,TGFB2,TGFBR2,TGFBR3,TUBA1C,TUBB3,VAV2,VCL,WAS,YES1 |

| | | | | |
|---|---|---|---|---|
| G-Protein Coupled Receptor Signaling | 5.86E+00 | 4.96E-01 | NaN | *ADCY1,ADCY10,ADCY2,ADCY3,ADCY5,ADCY6,ADCY7,ADCY8,ADCY9,ADORA1,ADORA2B,ADORA3,ADRA1A,ADRA1B,AKT2,AKT3,BRAF,CALCR,CAMK2A,CAMK2B,CAMK2G,CAMK4,CHRM3,CNR2,CREB3L3,CREB5,CREBBP,CRHR1,DRD2,DRD3,DUSP4,ENPP6,FYN,GABBR2,GDPD1,GLP1R,GNA11,GNA14,GNAI1,GNAI2,GNAI3,GNAL,GNAO1,GNAQ,GNAS,GRB2,GRK2,GRK4,GRM1,GRM3,GRM4,GRM5,GRM7,GRM8,HRH1,HRH2,HTR2C,HTR4,HTR7,IKBKB,IKBKE,IKBKG,KRAS,MAP2K1,MAP2K2,MAPK1,MPPED2,MRAS,NFKBID,NPR3,OPRD1,OPRL1,PDE10A,PDE11A,PDE1A,PDE1B,PDE1C,PDE2A,PDE3A,PDE3B,PDE4A,PDE4B,PDE4C,PDE4D,PDE6A,PDE6B,PDE6C,PDE6G,PDE8A,PDE8B,PDE9A,PDPK1,PIK3C2A,PIK3C2B,PIK3CB,PIK3CD,PIK3R2,PIK3R3,PIK3R4,PIK3R5,PIK3R6,PLCB1,PLCB2,PLCB3,PRKACA,PRKAG2,PRKAR1B,PRKAR2A,PRKCA,PRKCB,PRKCE,PRKCG,PTGER3,PTH1R,PTK2B,RAF1,RALA,RALB,RAP1A,RAP1B,RAP1GAP,RAPGEF3,RAPGEF4,RASA1,RGS10,RGS12,RGS14,RGS7,RPS6KA1,RRA* |
| White Adipose Tissue Browning Pathway | 5.85E+00 | 5.62E-01 | -8.189 | *ADCY1,ADCY10,ADCY2,ADCY3,ADCY5,ADCY6,ADCY7,ADCY8,ADCY9,BMP7,CACNA1A,CACNA1B,CACNA1C,CACNA1D,CACNA1E,CACNA1F,CACNA1G,CACNA1H,CACNA1I,CACNA1S,CACNA2D1,CACNA2D2,CACNA2D3,CACNA2D4,CACNB1,CACNB2,CACNB4,CACNG2,CACNG3,CACNG4,CACNG5,CACNG6,CACNG7,CACNG8,CAMKK2,CREB3L3,CREB5,CREBBP,CTBP2,FGFR1,FGFR2,FGFR4,GNAS,GUCY1A2,HIF1AN,LDHA,LDHC,MAPK1,MAPK13,MAPK14,NPR1,NRF1,PPARA,PPARG,PRDM16,PRKAA1,PRKAA2,PRKACA,PRKAG2,PRKAR1B,PRKAR2A,PRKG1,RARA,RARB,RARG,RXRA,RXRG,SIRT1,SLC16* |
| Ephrin A Signaling | 5.85E+00 | 7.02E-01 | NaN | *ADAM10,BCAR1,CDC42,EFNA3,EFNA5,EPHA1,EPHA10,EPHA2,EPHA3,EPHA6,EPHA8,FYN,LIMK1,NGEF,PAK1,PIK3C2A,PIK3C2B,PIK3CB,PIK3CD,PIK3R2,PIK3R3,PIK3R4,PIK3R5,PIK3R6,PTK2,PTPN11,RAC1,RHOA,ROCK1,ROCK2,VAV1,VAV2,VAV3* |
| Leptin Signaling in Obesity | 5.78E+00 | 6.30E-01 | -4.264 | *ADCY1,ADCY10,ADCY2,ADCY3,ADCY5,ADCY6,ADCY7,ADCY8,ADCY9,AKT2,AKT3,FOXO1,GRB2,JAK2,LEPR,MAP2K1,MAP2K2,MAPK1,PDE3A,PDE3B,PIK3C2A,PIK3C2B,PIK3CB,PIK3CD,PIK3R2,PIK3R3,PIK3R4,PIK3R5,PIK3R6,PLCB1,PLCB2,PLCB3,PLCD3,PLCE1,PLCG1,PLCG2,PLCH1,PLCH2,PLCL1,PLCL2,PRKACA,PRKAG2,PRKAR1B,PRKAR2A,PTPN11,STAT3* |
| Glioblastoma Multiforme Signaling | 5.76E+00 | 5.37E-01 | -6.952 | *AKT2,AKT3,APC,AXIN1,CDC42,CDK6,E2F1,E2F2,E2F3,E2F4,EGFR,FNBP1,FOXO1,FZD3,GRB2,GSK3B,IGF1R,ITPR1,ITPR2,ITPR3,KRAS,MAP2K1,MAP2K2,MAPK1,MDM2,MRAS,MTOR,NF1,NF2,PDGFA,PDGFB,PDGFD,PDGFRB,PIK3C2A,PIK3C2B,PIK3CB,PIK3CD,PIK3R2,PIK3R3,PIK3R4,PIK3R5,PIK3R6,PLCB1,PLCB2,PLCB3,PLCD3,PLCE1,PLCG1,PLCG2,PLCH1,PLCH2,PLCL1,PLCL2,PRKCD,PTEN,RAC1,RAC2,RAF1,RALA,RALB,RAP1A,RAP1B,RHOA,RHOBTB1,RHOD,RHOQ,RHOT1,RPS6KB1,RRAS2,SMO,SOS1,SRC,TCF3,TP53,TSC1,TSC2,WNT10A,WNT11,WNT2B,WNT3,WNT3A,WNT4,WNT5B,WNT7A,* |
| Glioma Signaling | 5.74E+00 | 5.78E-01 | -6.861 | *ABL1,AKT2,AKT3,CAMK1D,CAMK2A,CAMK2B,CAMK2G,CAMK4,CDK6,E2F1,E2F2,E2F3,E2F4,EGFR,GRB2,IDH2,IGF1R,IGF2R,KRAS,MAP2K1,MAP2K2,MAPK1,MDM2,MRAS,MTOR,PDGFA,PDGFB,PDGFD,PDGFRB,PIK3C2A,PIK3C2B,PIK3CB,PIK3CD,PIK3R2,PIK3R3,PIK3R4,PIK3R5,PIK3R6,PLCG1,PLCG2,PRKCA,PRKCB,PRKCD,PRKCE,PRKCG,PRKCH,PRKCI,PRKCQ,PRKCZ,PRKD1,PTEN,RAF1,RALA,RALB,RAP1A,RAP1B,RBL1,RRAS2,SIN3A,SOS1,TFDP1,TGFA,TP53* |
| Endocannabinoid Cancer Inhibition Pathway | 5.73E+00 | 5.49E-01 | 2.065 | *ADCY1,ADCY10,ADCY2,ADCY3,ADCY5,ADCY6,ADCY7,ADCY8,ADCY9,AKT2,AKT3,ATF3,CAMKK2,CASP8,CASP9,CCND3,CDH1,CNR2,CREB3L3,CREB5,CREBBP,GNAI1,GNAI2,GNAI3,GNAL,GNAO1,GNAQ,GNB1L,GPR55,GSK3B,HNF1A,MAP2K1,MAP2K2,MAP2K3,MAP2K4,MAP2K5,MAP2K6,MAPK1,MAPK14,MMP2,MTOR,NOS1,NOS2,NOS3,PIK3C2A,PIK3C2B,PIK3CB,PIK3CD,PIK3R2,PIK3R3,PIK3R4,PIK3R5,PIK3R6,PRKAA1,PRKAA2,PRKACA,PRKAG2,PRKAR1B,PRKAR2A,PTK2,RAF1,RHOA,ROCK1,ROCK2,RPTOR,SMPD3,SMPD4,SPTLC1,SPTLC2,SRC,TCF3,TCF4,TCF7,TCF7L1,TCF7L2,TWIST2,VEGFA,VE* |
| CDK5 Signaling | 5.71E+00 | 5.79E-01 | -4.296 | *ABL1,ADCY1,ADCY10,ADCY2,ADCY3,ADCY5,ADCY6,ADCY7,ADCY8,ADCY9,CABLES1,CACNA1A,CAPN1,GNAL,GNAS,ITGA2,ITGA3,KRAS,LAMA1,LAMA5,LAMB1,LAMC1,MAP2K1,MAP2K2,MAPK1,MAPK10,MAPK13,MAPK14,MAPK4,MAPK6,MAPK8,MAPK9,MAPT,MRAS,NTRK2,PPM1L,PPP1R12A,PPP1R14C,PPP1R14D,PPP1R3A,PPP1R7,PPP2CB,PPP2R1B,PPP2R2A,PPP2R2B,PPP2R2C,PPP2R3A,PPP2R5B,PPP2R5C,PPP2R5D,PPP2R5E,PRKACA,PRKAG2,PRKAR1B,PRKAR2A,PTPA,RAF1,RALA,RALB,RAP1* |
| Adrenomedullin signaling pathway | 5.70E+00 | 5.21E-01 | -9.037 | *ADCY1,ADCY10,ADCY2,ADCY3,ADCY5,ADCY6,ADCY7,ADCY8,ADCY9,AKT2,AKT3,ARNT,BCL2,BRAF,C3,CAMK4,CFH,CSK,GNA11,GNA14,GNAQ,GNAS,GRB2,GSK3B,GUCY1A2,GUCY2D,ITPR1,ITPR2,ITPR3,KCNH2,KCNN1,KCNN3,KCNN4,KCNQ2,KCNQ3,KRAS,MAP2K1,MAP2K2,MAP2K3,MAP2K4,MAP2K5,MAP2K6,MAPK1,MAPK10,MAPK13,MAPK14,MAPK4,MAPK6,MAPK8,MAPK9,MMP2,MRAS,MYLK,MYLK2,MYLK3,NOS3,NPR1,NPR3,PIK3C2A,PIK3C2B,PIK3CB,PIK3CD,PIK3R2,PIK3R3,PIK3R4,PIK3R5,PIK3R6,PLCB1,PLCB2,PLCB3,PLCD3,PLCE1,PLCG1,PLCG2,PLCH1,PLCH2,PLCL1,PLCL2,PPARG,PRKACA,PRKAG2,PRKAR1B,PRKAR2A,PRKG1,PTK2,PTK2B,RAF1,RALA,RALB,RAMP3,RAP1A,RAP1B,RRAS2,* |
| Germ Cell-Sertoli Cell Junction Signaling | 5.51E+00 | 5.30E-01 | NaN | *ACTB,ACTG2,ACTN1,ACTN2,ACTN4,AXIN1,BCAR1,CDC42,CDH1,CTNNA1,CTNNA2,EPN1,EPN2,FER,FNBP1,GSN,IQGAP1,ITGA2,ITGA3,JUP,KEAP1,KRAS,LAMC3,LIMK1,LIMK2,MAP2K1,MAP2K2,MAP2K3,MAP2K4,MAP2K6,MAP3K1,MAP3K10,MAP3K11,MAP3K12,MAP3K13,MAP3K14,MAP3K15,MAP3K2,MAP3K3,MAP3K5,MAP3K6,MAPK1,MAPK10,MAPK14,MAPK8,MAPK9,MRAS,MYO7A,NECTIN2,NECTIN3,PAK1,PAK2,PAK4,PDPK1,PIK3C2A,PIK3C2B,PIK3CB,PIK3CD,PIK3R2,PIK3R3,PIK3R4,PIK3R5,PIK3R6,PLS1,PTK2,PXN,RAB8B,RAC1,RAC2,RALA,RALB,RAP1A,RAP1B,RHOA,RHOBTB1,RHOD,RHOQ,RHOT1,RRAS2,SORBS1,SRC,TGFB1,TGFB2,TGFBR2,TJP1,TUBA1C,TUBB3,VCL* |
| Tec Kinase Signaling | 5.39E+00 | 5.31E-01 | -8.062 | *ACTB,ACTG2,BLK,CDC42,FGR,FNBP1,FYN,GNA11,GNA12,GNA14,GNAI1,GNAI2,GNAI3,GNAL,GNAO1,GNAQ,GNAS,GNB1,GNB1L,GNB5,GNG12,GNG2,GNG4,GNG7,GTF2I,HCK,ITGA2,ITGA3,JAK1,JAK2,JAK3,LCK,MAP2K4,MAPK10,MAPK8,MAPK9,MRAS,PAK1,PAK2,PAK4,PIK3C2A,PIK3C2B,PIK3CB,PIK3CD,PIK3R2,PIK3R3,PIK3R4,PIK3R5,PIK3R6,PLCG1,PLCG2,PRKCA,PRKCB,PRKCD,PRKCE,PRKCG,PRKCH,PRKCI,PRKCQ,PRKCZ,PRKD1,PTK2,PTK2B,RAC1,RAC2,RHOA,RHOBTB1,RHOD,RHOQ,RHOT1,SRC,STAT2,STAT3,STAT5A,STAT5B,STAT6,TNFRSF10A,TNFRSF10B,TNFRSF21,TYK2,VAV1,VAV2,VA* |
| Rac Signaling | 5.38E+00 | 5.68E-01 | -7.681 | *ACTR2,ACTR3,ANK1,ARPC1A,ARPC5L,BAIAP2,CD44,CDC42,CYFIP1,CYFIP2,IQGAP1,IQGAP2,IQGAP3,ITGA2,ITGA3,KRAS,LIMK1,LIMK2,MAP2K1,MAP2K2,MAP2K4,MAP3K1,MAP3K11,MAPK1,MAPK8,MCF2L,MRAS,NOX4,PAK1,PAK2,PAK4,PARD3,PI4KA,PIK3C2A,PIK3C2B,PIK3CB,PIK3CD,PIK3R2,PIK3R3,PIK3R4,PIK3R5,PIK3R6,PIP4K2B,PIP5K1A,PIP5K1B,PIP5K1C,PIP5KL1,PLD1,PRKCI,PRKCZ,PTK2,PTK2B,RAC1,RAF1,RALA,RALB,RAP1A,RAP1B,RHOA,RPS6KB1,RRAS2,SH3RF1,TIAM1* |
| Prolactin Signaling | 5.37E+00 | 6.05E-01 | -6.193 | *CREBBP,EP300,FYN,GRB2,JAK2,KRAS,MAP2K1,MAP2K2,MAPK1,MRAS,NR3C1,PDPK1,PIK3C2A,PIK3C2B,PIK3CB,PIK3CD,PIK3R2,PIK3R3,PIK3R4,PIK3R5,PIK3R6,PLCG1,PLCG2,PRKCA,PRKCB,PRKCD,PRKCE,PRKCG,PRKCH,PRKCI,PRKCQ,PRKCZ,PRKD1,PRLR,PTPN11,RAF1,RALA,RALB,RAP1A,RAP1B,RRAS2,SOCS5,SOCS7,SOS1,SP1,STAT3,STAT5A,STAT5B,TCF7* |

| | | | | |
|---|---|---|---|---|
| Sertoli Cell-Sertoli Cell Junction Signaling | 5.33E+00 | 5.19E-01 | NaN | ACTB,ACTG2,ACTN1,ACTN2,ACTN4,ADCY10,AKT2,AKT3,AXIN1,BCAR1,CDC42,CDH1,CGN,CLDN10,CLDN11,CLDN14,CLDN4,CTNNA1,CTNNA2,DLG1,EPB41,EPN1,EPN2,F11R,GSK3B,GUCY1A2,ITGA2,ITGA3,JAM3,JUP,KEAP1,KRAS,MAGI2,MAP2K1,MAP2K2,MAP2K3,MAP2K4,MAP3K1,MAP3K10,MAP3K11,MAP3K12,MAP3K13,MAP3K14,MAP3K15,MAP3K2,MAP3K3,MAP3K5,MAP3K6,MAPK1,MAPK10,MAPK13,MAPK14,MAPK8,MAPK9,MPP6,MRAS,MYO7A,NECTIN1,NECTIN2,NECTIN3,NOS1,NOS2,NOS3,PLS1,PRKACA,PRKAG2,PRKAR1B,PRKAR2A,PRKG1,PTEN,RAB8B,RAC1,RAF1,RALA,RALB,RAP1A,RAP1B,RRAS2,SORBS1,SPTAN1,SPTB,SPTBN1,SPTBN2,SRC,SYMPK,TGFBR3,TJAP1,TJP1,TJP2,TJP3,T |
| Relaxin Signaling | 5.31E+00 | 5.37E-01 | -6.332 | ADCY1,ADCY10,ADCY2,ADCY3,ADCY5,ADCY6,ADCY7,ADCY8,ADCY9,AKT2,AKT3,BRAF,ENPP6,GDPD1,GNA11,GNA12,GNA14,GNAI1,GNAI2,GNAI3,GNAL,GNAO1,GNAQ,GNAS,GNB1,GNB1L,GNB5,GNG12,GNG2,GNG4,GNG7,GUCY1A2,GUCY2D,MAP2K1,MAPK1,MPPED2,MRAS,NFKBID,NOS2,NOS3,NPR1,NPR3,PDE10A,PDE11A,PDE1A,PDE1B,PDE1C,PDE2A,PDE3A,PDE3B,PDE4A,PDE4B,PDE4C,PDE4D,PDE6A,PDE6B,PDE6C,PDE6G,PDE8A,PDE8B,PDE9A,PIK3C2A,PIK3C2B,PIK3CB,PIK3CD,PIK3R2,PIK3R3,PIK3R4,PIK3R5,PIK3R6,PRKACA,PRKAG2,PRKAR1B,PRKAR2A,PRKCZ,RAP1A,RAP1B,RXFP1,V |
| Leukocyte Extravasation Signaling | 5.28E+00 | 5.13E-01 | -7.862 | ABL1,ACTB,ACTG2,ACTN1,ACTN2,ACTN4,ARHGAP1,ARHGAP35,ARHGAP6,BCAR1,CD44,CD99,CDC42,CDH5,CLDN10,CLDN11,CLDN14,CLDN4,CRK,CTNNA1,CTNNA2,CTNNA3,CTTN,CXCL12,DLC1,EDIL3,EZR,F11R,FER,GNAI1,GNAI2,GNAI3,ITGA2,ITGA3,ITGAL,ITGAM,ITGB2,ITGB3,JAM3,MAP2K2,MAP2K4,MAPK1,MAPK10,MAPK13,MAPK14,MAPK8,MAPK9,MMP14,MMP15,MMP17,MMP2,MMP21,MMP26,MMP28,MSN,PIK3C2A,PIK3C2B,PIK3CB,PIK3CD,PIK3R2,PIK3R3,PIK3R4,PIK3R5,PIK3R6,PLCG1,PLCG2,PRKCA,PRKCB,PRKCD,PRKCE,PRKCG,PRKCH,PRKCI,PRKCQ,PRKCZ,PRKD1,PTK2,PTK2B,PTPN11,PXN,RAC1,RAC2,RAP1A,RAP1B,RAP1GAP,RAPGEF3,RAPGEF4,RASSF5,RDX,RHOA,ROCK1,ROC |
| Role of Tissue Factor in Cancer | 5.27E+00 | 5.60E-01 | NaN | AKT2,AKT3,ARRB1,BCL2L1,BLK,CDC42,CSF1,EGFR,F10,F2,FGR,FYN,GNA11,GNA12,GNA14,GNAQ,HCK,ITGA3,ITGB3,ITGB5,JAK2,KRAS,LCK,LIMK1,LIMK2,MAPK1,MAPK13,MAPK14,MRAS,MTOR,P4HB,PAK1,PDIA6,PIK3C2A,PIK3C2B,PIK3CB,PIK3CD,PIK3R2,PIK3R3,PIK3R4,PIK3R5,PIK3R6,PLAUR,PLCB1,PRKCA,PTEN,PTK2B,PTPN11,RAC1,RALA,RALB,RAP1A,RAP1B,RPS6KA1,RPS6KA2,RPS6KA3,RPS6KA5,RPS6KB1,RRAS2,SRC,STAT5A,STAT5B,TP53,VEGFA,YES1 |
| PI3K Signaling in B Lymphocytes | 5.23E+00 | 5.45E-01 | -7.584 | ABL1,AKT2,AKT3,ATF1,ATF3,ATF6,BLK,BLNK,C3,CAMK2A,CAMK2B,CAMK2G,CAMK4,CARD10,CBL,CHP1,FOXO3,FYN,IKBKB,IKBKE,IKBKG,IL4R,INPP5D,IRS2,ITPR1,ITPR2,ITPR3,KRAS,MAP2K1,MAP2K2,MAPK1,MRAS,NFAT5,NFATC1,NFATC2,NFATC3,NFKBID,PDPK1,PIK3AP1,PIK3CB,PIK3CD,PIK3R2,PLCB1,PLCB2,PLCB3,PLCD3,PLCE1,PLCG1,PLCG2,PLCH1,PLCH2,PLCL1,PLCL2,PLEKHA2,PLEKHA4,PPP3CB,PPP3CC,PRKCB,PRKCI,PRKCZ,PTEN,PTPRC,RAC1,RAF1,RALA,RALB,RAP1A,RAP1B,RRAS2, |
| UVC-Induced MAPK Signaling | 5.21E+00 | 6.67E-01 | -5.745 | ATR,BRAF,EGFR,KRAS,MAP2K1,MAP2K2,MAP2K4,MAPK1,MAPK10,MAPK13,MAPK14,MAPK8,MAPK9,MRAS,PRKCA,PRKCB,PRKCD,PRKCE,PRKCG,PRKCH,PRKCI,PRKCQ,PRKCZ,PRKD1,RAF1,RALA,RALB,RAP1A,RAP1B,RRAS2,SMPD3,SMPD4,SRC,TP53 |
| PDGF Signaling | 5.21E+00 | 5.93E-01 | -6.861 | ABL1,ABL2,CRK,CSNK2A1,CSNK2A2,GRB2,INPP5B,INPP5D,INPP5F,INPP5K,INPPL1,JAK1,JAK2,JAK3,KRAS,MAP2K1,MAP2K2,MAP2K4,MAP3K1,MAPK1,MAPK8,MRAS,PDGFA,PDGFB,PDGFD,PDGFRB,PIK3C2A,PIK3C2B,PIK3CB,PIK3CD,PIK3R2,PIK3R3,PIK3R4,PIK3R5,PIK3R6,PLCG1,PLCG2,PRKCA,PRKCB,RAF1,RALA,RALB,RAP1A,RAP1B,RASA1,RRAS2,SOS1,SRC,STAT3,SYNJ2,TYK2 |
| LPS-stimulated MAPK Signaling | 5.16E+00 | 5.98E-01 | -6.856 | ATF1,CDC42,IKBKB,IKBKE,IKBKG,KRAS,LBP,MAP2K1,MAP2K2,MAP2K3,MAP2K4,MAP2K6,MAP3K14,MAP3K5,MAPK1,MAPK10,MAPK13,MAPK14,MAPK8,MAPK9,MRAS,NFKBID,PAK1,PIK3C2A,PIK3C2B,PIK3CB,PIK3CD,PIK3R2,PIK3R3,PIK3R4,PIK3R5,PIK3R6,PRKCA,PRKCB,PRKCD,PRKCE,PRKCG,PRKCH,PRKCI,PRKCQ,PRKCZ,PRKD1,RAC1,RAF1,RALA,RALB,RAP1A,RAP1B,RRAS2 |
| UVA-Induced MAPK Signaling | 5.16E+00 | 5.77E-01 | -5.916 | ATM,BCL2L1,CASP9,EGFR,KRAS,MAP2K4,MAPK1,MAPK10,MAPK13,MAPK14,MAPK8,MAPK9,MRAS,MTOR,PARP1,PARP10,PARP12,PARP16,PARP8,PIK3C2A,PIK3C2B,PIK3CB,PIK3CD,PIK3R2,PIK3R3,PIK3R4,PIK3R5,PIK3R6,PLCB1,PLCB2,PLCB3,PLCD3,PLCE1,PLCG1,PLCG2,PLCH1,PLCH2,PLCL1,PLCL2,PRKCA,RALA,RALB,RAP1A,RAP1B,RPS6KA1,RPS6KA2,RPS6KA3,RPS6KA5,RPS6KB1,RPS6KC1,RRAS2,SMPD3,SMPD4,TNKS,TNKS2,TP53 |
| VEGF Signaling | 5.16E+00 | 5.77E-01 | -5.774 | ACTB,ACTG2,ACTN1,ACTN2,ACTN4,AKT2,AKT3,ARNT,BCL2,BCL2L1,EIF2B3,EIF2B5,EIF2S3,ELAVL1,FLT1,FLT4,FOXO1,FOXO3,GRB2,KRAS,MAP2K1,MAP2K2,MAPK1,MRAS,NOS3,PIK3C2A,PIK3C2B,PIK3CB,PIK3CD,PIK3R2,PIK3R3,PIK3R4,PIK3R5,PIK3R6,PLCG1,PLCG2,PRKCA,PRKCB,PTK2,PTK2B,PTPN11,PXN,RAF1,RALA,RALB,RAP1A,RAP1B,ROCK1,ROCK2,RRAS2,SOS1,SRC,VCL,VEGFA,VEGFD,Y |
| Agrin Interactions at Neuromuscular Junction | 5.09E+00 | 6.05E-01 | -6.325 | ACTB,ACTG2,AGRN,ARHGEF7,CDC42,CTTN,DAG1,EGFR,ERBB2,ERBB3,ERBB4,GABPB1,ITGA2,ITGA3,ITGAL,ITGB2,ITGB3,KRAS,LAMA2,LAMB1,LAMC1,MAP2K4,MAPK1,MAPK10,MAPK8,MAPK9,MRAS,NRG1,NRG2,NRG3,NRG4,PAK1,PAK2,PAK4,PKLR,PTK2,PXN,RAC1,RAC2,RALA,RALB,RAP1A,RAP1B, |
| fMLP Signaling in Neutrophils | 5.07E+00 | 5.57E-01 | -7.488 | ACTR2,ACTR3,ARPC1A,ARPC5L,CAMK4,CDC42,CHP1,GNAI1,GNAI2,GNAI3,GNAS,GNB1,GNB1L,GNB5,GNG12,GNG2,GNG4,GNG7,ITPR1,ITPR2,ITPR3,KRAS,MAP2K1,MAP2K2,MAPK1,MRAS,NFAT5,NFATC1,NFATC2,NFATC3,NFKBID,NOX4,PIK3C2A,PIK3C2B,PIK3CB,PIK3CD,PIK3R2,PIK3R3,PIK3R4,PIK3R5,PIK3R6,PLCB1,PLCB2,PLCB3,PPP3CB,PPP3CC,PRKCA,PRKCB,PRKCD,PRKCE,PRKCG,PRKCH,PRKCI,PRKCQ,PRKCZ,PRKD1,RAC1,RAF1,RALA,RALB,RAP1A,RAP1B,RRAS2,WAS |
| Gustation Pathway | 5.06E+00 | 5.30E-01 | NaN | ADCY1,ADCY10,ADCY2,ADCY3,ADCY5,ADCY6,ADCY7,ADCY8,ADCY9,ASIC1,ASIC2,CACNA1A,CACNA1B,CACNA1C,CACNA1D,CACNA1E,CACNA1F,CACNA1G,CACNA1H,CACNA1I,CACNA1S,CACNA2D1,CACNA2D2,CACNA2D3,CACNA2D4,CACNB1,CACNB2,CACNB4,CACNG2,CACNG3,CACNG4,CACNG5,CACNG6,CACNG7,CACNG8,ENPP6,GDPD1,GNB1,GNG2,GNG7,ITPR1,ITPR2,ITPR3,MPPED2,P2RX1,P2RX3,P2RX4,P2RX7,P2RY2,P2RY6,P2RY8,PANX1,PDE10A,PDE11A,PDE1A,PDE1B,PDE1C,PDE2A,PDE3A,PDE3B,PDE4A,PDE4B,PDE4C,PDE4D,PDE6A,PDE6B,PDE6C,PDE6G,PDE8A,PDE8B,PDE9A,PKD1L3,PKD2L1,PLCB2,PRKACA,PRKAG2,PRKAR1B,PRKAR2A,SCNN1B,TAS1R1 |
| CNTF Signaling | 4.94E+00 | 6.43E-01 | -6 | CNTFR,GRB2,IL6ST,JAK1,JAK2,KRAS,LIFR,MAP2K1,MAP2K2,MAPK1,MRAS,MTOR,PIK3C2A,PIK3C2B,PIK3CB,PIK3CD,PIK3R2,PIK3R3,PIK3R4,PIK3R5,PIK3R6,PTPN11,RAF1,RALA,RALB,RAP1A,RAP1B,RPS6KA1,RPS6KA2,RPS6KA3,RPS6KA5,RPS6KB1,RRAS2,SOS1,STAT3,TYK2 |
| Neuregulin Signaling | 4.91E+00 | 5.74E-01 | -6.429 | AKT2,AKT3,CRK,DLG4,EGFR,ERBB2,ERBB3,ERBB4,ERBIN,GRB2,HSP90AA1,ITGA2,ITGA3,KRAS,MAP2K1,MAP2K2,MAPK1,MRAS,MTOR,NRG1,NRG2,NRG3,NRG4,PDPK1,PIK3R2,PIK3R3,PLCG1,PLCG2,PRKCA,PRKCB,PRKCD,PRKCE,PRKCG,PRKCH,PRKCI,PRKCQ,PRKCZ,PRKD1,PSEN1,PTEN,PTPN11,RAF1,RALA,RALB,RAP1A,RAP1B,RNF41,RPS6KB1,RRAS2,SOS1,SRC,STAT5A,STAT5B,TGFA |
| Melanocyte Development and Pigmentation Signaling | 4.91E+00 | 5.74E-01 | -7.28 | ADCY1,ADCY10,ADCY2,ADCY3,ADCY5,ADCY6,ADCY7,ADCY8,ADCY9,BCL2,CREB3L3,CREB5,CREBBP,CRK,EP300,GNAS,GRB2,KRAS,MAP2K1,MAP2K2,MAPK1,MITF,MRAS,PIK3C2A,PIK3C2B,PIK3CB,PIK3CD,PIK3R2,PIK3R3,PIK3R4,PIK3R5,PIK3R6,PLCG1,PLCG2,PRKACA,PRKAG2,PRKAR1B,PRKAR2A,PTPN11,RAF1,RALA,RALB,RAP1A,RAP1B,RPS6KA1,RPS6KA2,RPS6KA3,RPS6KA5,RPS6KB1,RPS6KC1,RRAS2,SH2B2,SOS1,SRC |

| | | | | |
|---|---|---|---|---|
| Clathrin-mediated Endocytosis Signaling | 4.83E+00 | 5.05E-01 | NaN | AAK1,ACTB,ACTG2,ACTR2,ACTR3,AMPH,AP1B1,AP1G1,AP1M1,AP2A1,AP2A2,AP2B1,AP3B1,AP3M1,ARPC1A,ARPC5L,ARRB1,CBL,CD2AP,CDC42,CHP1,CLTA,CLTB,CLTC,CLTCL1,CLU,CSNK2A1,CSNK2A2,CTTN,DAB2,DNM1,DNM1L,DNM2,DNM3,EPHB2,EPN1,EPS15,F2,FGF1,FGF12,FGF14,FGF18,GAK,GRB2,HGS,HIP1,HIP1R,ITGB2,ITGB3,ITGB4,ITGB5,ITGB6,ITGB7,ITGB8,LDLR,LDLRAP1,LPA,MDM2,MET,MYO1E,MYO6,NUMB,PCYOX1,PDGFA,PDGFB,PDGFD,PICALM,PIK3C2A,PIK3C2B,PIK3CB,PIK3CD,PIK3R2,PIK3R3,PIK3R4,PIK3R5,PIK3R6,PIP5K1C,PPP3CB,PPP3CC,RAB11A,RAB11B,RAB5B,RAB7A,RAC1,SH3BP4,SH3GL1,SH3GL2,SH3GL3,SH3GLB2,SH3KBP1,SNX9,SRC,TFRC,TSG101,USP9X,VEGF |
| 14-3-3-mediated Signaling | 4.74E+00 | 5.40E-01 | -6.379 | AKT2,AKT3,CBL,EDC3,FOXO1,GFAP,GRB2,GSK3B,KRAS,MAP2K1,MAP2K2,MAP2K4,MAP3K5,MAPK1,MAPK10,MAPK8,MAPK9,MAPT,MRAS,PDCD6IP,PIK3C2A,PIK3C2B,PIK3CB,PIK3CD,PIK3R2,PIK3R3,PIK3R4,PIK3R5,PIK3R6,PLCB1,PLCB2,PLCB3,PLCD3,PLCE1,PLCG1,PLCG2,PLCH1,PLCH2,PLCL1,PLCL2,PRKCA,PRKCB,PRKCD,PRKCE,PRKCG,PRKCH,PRKCI,PRKCQ,PRKCZ,PRKD1,RAF1,RALA,RALB,RAP1A,RAP1B,RPS6KA1,RRAS2,SRC,SRPK2,TP73,TRAF2,TSC1,TSC2,TUBA1C,TUBB3,YAP1,YWHAE, |
| Xenobiotic Metabolism Signaling | 4.72E+00 | 4.80E-01 | NaN | ABCB1,ABCC2,ABCC3,AHRR,ALDH1A1,ALDH1A2,ALDH1L1,ALDH1L2,ALDH2,ALDH3A1,ALDH3A2,ALDH3B1,ALDH7A1,ARNT,CAMK1D,CAMK2A,CAMK2B,CAMK2G,CAMK4,CES1,CES3,CES4A,CES5A,CHST1,CHST10,CHST11,CHST12,CHST15,CHST3,CHST7,CREBBP,CUL3,CYP2C19,DNAJC7,EP300,GCLC,GRIP1,HDAC4,HDAC5,HS2ST1,HS3ST2,HS3ST3A1,HS3ST3B1,HS3ST4,HS6ST1,HS6ST2,HS6ST3,HSP90AA1,IL4I1,KEAP1,KRAS,MAOA,MAOB,MAP2K1,MAP2K2,MAP2K3,MAP2K4,MAP2K5,MAP2K6,MAP3K1,MAP3K10,MAP3K11,MAP3K12,MAP3K13,MAP3K14,MAP3K15,MAP3K2,MAP3K3,MAP3K5,MAP3K6,MAPK1,MAPK13,MAPK14,MAPK8,MAPK9,MED1,MGMT,MRAS,NCOA1,NCOR2,NDST1,NOS2,NQO1,NQO2,NR1I2,PIK3C2A,PIK3C2B,PIK3CB,PIK3CD,PIK3R2,PIK3R3,PIK3R4,PIK3R5,PIK3R6,PPM1L,PPP2CB,PPP2R1B,PPP2R2A,PPP2R2B,PPP2R2C,PPP2R3A,PPP2R5B,PPP2R5C,PPP2R5D,PPP2R5E,PRKCA,PRKCB,PRKCD,PRKCE,PRKCG,PRKCH,PRKCI,PRKCQ,PRKCZ,PRKD1,PTGES3,PTPA,RAF1,RALA,RALB,RAP1A,RAP1B,RRAS2,RXRA,SMOX,SULT1A1,SULT2B1,SULT4A1,UST |
| Phospholipases | 4.70E+00 | 6.27E-01 | -5.916 | ABHD3,GPLD1,LIPC,PLA1A,PLA2G10,PLA2G12B,PLA2G2C,PLA2G2F,PLA2G4A,PLA2G4C,PLA2G4D,PLA2G4E,PLA2G4F,PLA2G5,PLA2G6,PLA2R1,PLAAT3,PLAAT4,PLAAT5,PLB1,PLCB1,PLCB2,PLCB3,PLCD3,PLCE1,PLCG1,PLCG2,PLCH1,PLCH2,PLCL1,PLCL2,PLD1,PLD2,PLD3,PLD5,PNPLA3,PNPLA8 |
| Antioxidant Action of Vitamin C | 4.69E+00 | 5.58E-01 | 7 | ABHD3,CSF2RA,CSF2RB,GPLD1,IKBKB,IKBKE,IKBKG,JAK2,MAP2K4,MAPK1,MAPK10,MAPK13,MAPK14,MAPK8,MAPK9,NFKBID,NXN,PLA2G10,PLA2G12B,PLA2G2C,PLA2G2F,PLA2G4A,PLA2G4C,PLA2G4D,PLA2G4E,PLA2G4F,PLA2G5,PLA2G6,PLA2R1,PLAAT3,PLAAT4,PLAAT5,PLB1,PLCB1,PLCB2,PLCB3,PLCD3,PLCE1,PLCG1,PLCG2,PLCH1,PLCH2,PLCL1,PLCL2,PLD1,PLD2,PLD3,PLD5,PNPLA3,PNPLA8,SLC23A2,SLC2A1,SLC2A14,SLC2A5,STAT5A,STAT5B,TXNRD1,TXNRD2 |
| EGF Signaling | 4.68E+00 | 6.36E-01 | -5.578 | AKT2,AKT3,CSNK2A1,CSNK2A2,EGFR,GRB2,ITPR1,ITPR2,ITPR3,JAK1,MAP2K1,MAP2K4,MAP3K1,MAPK1,MAPK13,MAPK14,MAPK8,MTOR,PIK3C2A,PIK3C2B,PIK3CB,PIK3CD,PIK3R2,PIK3R3,PIK3R4,PIK3R5,PIK3R6,PLCG1,PRKCA,RAF1,RASA1,RPS6KB1,SOS1,SRC,STAT3 |
| Wnt/β-catenin Signaling | 4.67E+00 | 5.12E-01 | -0.585 | ACVR1B,ACVR2B,AKT2,AKT3,APC,APPL1,APPL2,AXIN1,BCL9,BMPR2,BTRC,CD44,CDH1,CDH12,CDH3,CDH5,CREBBP,CSNK1D,CSNK1E,CSNK1G2,CSNK2A1,CSNK2A2,DKK3,EP300,FZD3,GNAO1,GNAQ,GSK3B,HDAC1,HNF1A,KREMEN1,LRP1,LRP5,LRP6,MAP4K1,MARK2,MDM2,NLK,NR5A2,PPARD,PPM1L,PPP2CB,PPP2R1B,PPP2R2A,PPP2R2B,PPP2R2C,PPP2R3A,PPP2R5B,PPP2R5C,PPP2R5D,PPP2R5E,PTPA,RARA,RARB,RARG,RUVBL2,SFRP1,SMO,SOX13,SOX5,SOX6,SRC,TAB1,TCF3,TCF4,TCF7,TCF7L1,TCF7L2,TGFB1,TGFB2,TGFBR2,TGFBR3,TLE1,TLE3,TLE4,TP53,WNT10A,WNT11,WNT2B,WNT3,WNT3A,WNT4,WNT5B,WNT7A,WNT7B,WNT9A,WNT9B |
| IL-1 Signaling | 4.66E+00 | 5.71E-01 | -4.811 | ADCY1,ADCY10,ADCY2,ADCY3,ADCY5,ADCY6,ADCY7,ADCY8,ADCY9,GNA11,GNA12,GNA14,GNAI1,GNAI2,GNAI3,GNAL,GNAO1,GNAQ,GNAS,GNB1,GNB1L,GNB5,GNG12,GNG2,GNG4,GNG7,IKBKB,IKBKE,IKBKG,IL1R1,IL1RAP,IRAK2,MAP2K3,MAP2K4,MAP2K6,MAP3K1,MAP3K14,MAPK1,MAPK10,MAPK13,MAPK14,MAPK8,MAPK9,MRAS,NFKBID,PRKACA,PRKAG2,PRKAR1B,PRKAR2A,TAB1,TAB2,TOLLI |
| Chronic Myeloid Leukemia Signaling | 4.66E+00 | 5.59E-01 | NaN | ABL1,AKT2,AKT3,BCL2L1,CDK6,CRK,CTBP2,E2F1,E2F2,E2F3,E2F4,GAB2,GRB2,HDAC1,HDAC11,HDAC4,HDAC5,HDAC6,HDAC7,HDAC9,IKBKB,IKBKE,IKBKG,KRAS,MAP2K1,MAP2K2,MAPK1,MDM2,MECOM,MRAS,PIK3C2A,PIK3C2B,PIK3CB,PIK3CD,PIK3R2,PIK3R3,PIK3R4,PIK3R5,PIK3R6,PTPN11,RAF1,RALA,RALB,RAP1A,RAP1B,RBL1,RRAS2,SIN3A,SMAD3,SOS1,STAT5A,STAT5B,TFDP1,TGFB1,TGFB2 |
| IL-8 Signaling | 4.63E+00 | 5.00E-01 | -9.487 | AKT2,AKT3,BCL2,BCL2L1,BRAF,CCND3,CDC42,CDH1,EGFR,FLT1,FLT4,FNBP1,GNA12,GNAI1,GNAI2,GNAI3,GNAS,GNB1,GNB1L,GNB5,GNG12,GNG2,GNG4,GNG7,GPLD1,IKBKB,IKBKE,IKBKG,IQGAP1,IRAK2,ITGAM,ITGAX,ITGB2,ITGB3,ITGB5,KRAS,LASP1,LIMK1,LIMK2,MAP2K1,MAP2K2,MAP2K4,MAP4K4,MAPK1,MAPK10,MAPK8,MAPK9,MMP2,MRAS,MTOR,NOX4,PAK2,PIK3C2A,PIK3C2B,PIK3CB,PIK3CD,PIK3R2,PIK3R3,PIK3R4,PIK3R5,PIK3R6,PLCB2,PLD1,PLD2,PLD3,PLD5,PRKCA,PRKCB,PRKCD,PRKCE,PRKCG,PRKCH,PRKCI,PRKCQ,PRKCZ,PRKD1,PTK2,PTK2B,RAC1,RAC2,RAF1,RALA,RALB,RAP1A,RAP1B,RHOA,RHOBTB1,RHOD,RHOQ,RHOT1,ROCK1,ROCK2,RPS6KB1,RRAS2,SRC,TEK,VEGFA, |
| Hepatic Fibrosis Signaling Pathway | 4.57E+00 | 4.60E-01 | -10.261 | ACVR1B,ACVR2B,AKT2,AKT3,AXIN1,BCL2,BMPR2,BRAF,BRD4,CACNA1A,CACNA1C,CACNA1D,CACNA1E,CACNA1S,CAMK4,CCDC88A,CDC42,COL18A1,COL1A1,COL2A1,COL5A3,CREB3L3,CREB5,CREBBP,EZH2,FGFR1,FLT1,FLT4,FNBP1,FOXO1,FZD3,GLI2,GLI3,GLIS1,GNAI1,GNAI2,GNAI3,GSK3B,HNF1A,IKBKB,IKBKE,IKBKG,IL17RA,IL1R1,IL1RAP,IL1RAPL1,IL1RAPL2,IRAK2,IRS2,ITGA2,ITGA3,ITGB3,JAK1,JAK2,KRAS,LEPR,LRP5,LRP6,MAP2K1,MAP2K2,MAP2K3,MAP2K4,MAP2K5,MAP2K6,MAPK1,MAPK10,MAPK13,MAPK14,MAPK8,MAPK9,MRAS,MTOR,MYL10,MYL3,MYLK,MYLK2,MYLK3,NFKBID,NOX4,PDGFA,PDGFB,PDGFD,PDGFRB,PIK3C2A,PIK3C2B,PIK3CB,PIK3CD,PIK3R2,PIK3R3,PIK3R4,PIK3R5,PIK3R6,PLCG1,PLCG2,PPARG,PRKACA,PRKAG2,PRKAR1B,PRKAR2A,PRKCA,PRKCB,PRKCD,PRKCE,PRKCG,PRKCH,PRKCI,PRKCQ,PRKCZ,PRKD1,PTCH1,PTCH2,PTEN,PTK2,RAC1,RAC2,RAF1,RALA,RALB,RAP1A,RAP1B,RHOA,RHOBTB1,RHOD,RHOQ,RHOT1,RIPK1,ROCK1,ROCK2,RPS6KB1,RRAS2,SDHA,SDHB,SERPINE1,SMAD3,SMAD7,SMO,SOS1,SP1,STAT3,TCF3,TCF4,TCF7,TCF7L1,TCF7L2,TFRC,TGFB1,TGFB2,TGFBR2,TGFBR3,TNFRSF1B,TRPM7,VEGFA,VEGFD,WNT10A,WNT11,WNT2B, |
| Gα12/13 Signaling | 4.48E+00 | 5.31E-01 | -4.123 | AKT2,AKT3,ARHGEF1,CDC42,CDH1,CDH10,CDH11,CDH12,CDH13,CDH15,CDH16,CDH18,CDH22,CDH23,CDH26,CDH3,CDH4,CDH5,CDH8,CDH9,F2,GNA12,IKBKB,IKBKE,IKBKG,KRAS,LPAR3,LPAR5,MAP2K1,MAP2K2,MAP2K4,MAP3K1,MAP3K5,MAPK1,MAPK10,MAPK8,MAPK9,MEF2A,MEF2D,MRAS,MYL10,MYL3,NFKBID,PIK3C2A,PIK3C2B,PIK3CB,PIK3CD,PIK3R2,PIK3R3,PIK3R4,PIK3R5,PIK3R6,PTK2,PTK2B,PXN,RAF1,RALA,RALB,RAP1A,RAP1B,RASA1,RHOA,ROCK1,ROCK2,RRAS2,SRC,VAV1,VAV2,VAV |
| Glutamate Receptor Signaling | 4.43E+00 | 6.25E-01 | -4 | CAMK4,DLG4,GNB1,GNG2,GNG7,GRIA1,GRIA2,GRIA4,GRID1,GRID2,GRIK3,GRIK4,GRIK5,GRIN1,GRIN2A,GRIN2B,GRIN2C,GRIN2D,GRIP1,GRM1,GRM3,GRM4,GRM5,GRM7,GRM8,HOMER1,HOMER2,SLC17A8,SLC1A1,SLC1A2,SLC1A3,SLC1A4,SLC1A6,SLC1A7,SLC38A1 |
| Acute Myeloid Leukemia Signaling | 4.42E+00 | 5.68E-01 | -5.126 | AKT2,AKT3,BRAF,CSF1R,CSF2RA,CSF2RB,CSF3R,GRB2,HNF1A,IDH2,JUP,KRAS,MAP2K1,MAP2K2,MAP2K3,MAP2K4,MAP2K5,MAP2K6,MAPK1,MRAS,MTOR,PIK3C2A,PIK3C2B,PIK3CB,PIK3CD,PIK3R2,PIK3R3,PIK3R4,PIK3R5,PIK3R6,PML,RAF1,RALA,RALB,RAP1A,RAP1B,RARA,RPS6KB1,RRAS2,RUNX1,SOS1,SPI1,STAT3,STAT5A,STAT5B,TCF3,TCF4,TCF7,TCF7L1,TCF7L2 |

| | | | | |
|---|---|---|---|---|
| UVB-Induced MAPK Signaling | 4.41E+00 | 6.35E-01 | -5.745 | EGFR,MAP2K1,MAP2K4,MAPK1,MAPK10,MAPK13,MAPK14,MAPK8,MAPK9,MTOR,PIK3C2A,PIK3C2B,PIK3CB,PIK3CD,PIK3R2,PIK3R3,PIK3R4,PIK3R5,PIK3R6,PRKCA,PRKCB,PRKCD,PRKCE,PRKCG,PRKCH,PRKCI,PRKCQ,PRKCZ,PRKD1,RPS6KA3,RPS6KA5,RPS6KB1,TP53 |
| Pancreatic Adenocarcinoma Signaling | 4.40E+00 | 5.46E-01 | -5.898 | ABL1,AKT2,AKT3,BCL2,BCL2L1,CASP9,CDC42,E2F1,E2F2,E2F3,E2F4,EGFR,ERBB2,GPLD1,GRB2,JAK1,JAK2,JAK3,KRAS,MAP2K1,MAP2K2,MAP2K4,MAPK1,MAPK10,MAPK8,MAPK9,MDM2,NOTCH1,PIK3C2A,PIK3C2B,PIK3CB,PIK3CD,PIK3R2,PIK3R3,PIK3R4,PIK3R5,PIK3R6,PLD1,PLD2,PLD3,PLD5,RAC1,RAD51,RAF1,RALA,RALBP1,RALGDS,SIN3A,SMAD3,STAT3,TFDP1,TGFA,TGFB1,TGFB2,TGFBR2,TP |
| nNOS Signaling in Neurons | 4.39E+00 | 6.52E-01 | -3.464 | CAMK2A,CAMK4,CAPN1,CAPN11,CAPN2,CAPN5,CAPN8,CHP1,DLG2,DLG4,GRIN1,GRIN2A,GRIN2B,GRIN2C,GRIN2D,NOS1,NOS1AP,PFKM,PPP3CB,PPP3CC,PRKCA,PRKCB,PRKCD,PRKCE,PRKCG,PRKCH,PRKCI,PRKCQ,PRKCZ,PRKD1 |
| Ovarian Cancer Signaling | 4.33E+00 | 5.22E-01 | -5.916 | ABL1,AKT2,AKT3,APC,ARRB1,AXIN1,BCL2,BRAF,BRCA1,CD44,E2F1,EGFR,FZD3,GSK3B,HNF1A,KRAS,MAP2K1,MAP2K2,MAPK1,MLH1,MMP2,MRAS,MSH2,MSH6,MTOR,PIK3C2A,PIK3C2B,PIK3CB,PIK3CD,PIK3R2,PIK3R3,PIK3R4,PIK3R5,PIK3R6,PMS2,PRKACA,PRKAG2,PRKAR1B,PRKAR2A,PTEN,PTGS1,RAD51,RAF1,RALA,RALB,RAP1A,RAP1B,RPS6KB1,RRAS2,SIN3A,SMO,SRC,TCF3,TCF4,TCF7,TCF7L1,TCF7L2,TFDP1,TP53,VEGFA,VEGFD,WNT10A,WNT11,WNT2B,WNT3,WNT3A,WNT4,WNT5B,WNT7A |
| IL-2 Signaling | 4.24E+00 | 6.07E-01 | -5.831 | AKT2,AKT3,CSNK2A1,CSNK2A2,GRB2,IL2RA,IL2RB,JAK1,JAK3,KRAS,LCK,MAP2K1,MAP2K2,MAPK1,MAPK8,MRAS,PIK3C2A,PIK3C2B,PIK3CB,PIK3CD,PIK3R2,PIK3R3,PIK3R4,PIK3R5,PIK3R6,PTK2B,PTPN11,RAF1,RALA,RALB,RAP1A,RAP1B,RRAS2,SOS1,STAT5A,STAT5B,SYK |
| Erythropoietin Signaling | 4.23E+00 | 5.79E-01 | NaN | AKT2,AKT3,CBL,GRB2,JAK2,KRAS,MAP2K1,MAP2K2,MAPK1,MRAS,NFKBID,PDPK1,PIK3C2A,PIK3C2B,PIK3CB,PIK3CD,PIK3R2,PIK3R3,PIK3R4,PIK3R5,PIK3R6,PLCG1,PLCG2,PRKCA,PRKCB,PRKCD,PRKCE,PRKCG,PRKCH,PRKCI,PRKCQ,PRKCZ,PRKD1,RAF1,RALA,RALB,RAP1A,RAP1B,RPS6KB1,RRAS2,SOS1,SRC,STAT5A,STAT5B |
| 3-phosphoinositide Biosynthesis | 4.18E+00 | 5.06E-01 | -8.944 | ACP6,ALPL,CDC25A,CDC25C,CILP,DOT1L,DUSP10,DUSP14,DUSP15,DUSP16,DUSP27,EPHX2,EYA4,HACD2,INPP5B,INPP5F,ITPK1,MTMR4,MTMR7,PALD1,PAWR,PI4K2A,PI4KA,PI4KB,PIK3C2A,PIK3C2B,PIK3CB,PIK3CD,PIK3R2,PIK3R3,PIK3R4,PIK3R5,PIK3R6,PIP4K2B,PIP5K1A,PIP5K1B,PIP5K1C,PIP5KL1,PLPP7,PPFIA2,PPFIA3,PPFIA4,PPFIBP2,PPIP5K1,PPM1H,PPP1R12A,PPP1R12C,PPP1R13B,PPP1R16B,PPP1R1C,PPP1R7,PPP1R8,PPP2R3A,PPP2R5B,PPP2R5D,PPP2R5E,PPP4C,PPP5C,PPTC7,PRUNE1,PTEN,PTPA,PTPN1,PTPN11,PTPN12,PTPN2,PTPN20,PTPN22,PTPN23,PTPN7,PTPRC,PTPRF,PTPRH,PTPRJ,PTPRM,PXYLP1,RASA1,SIRPA,STYXL1,TNS3,TPTE |
| TGF-β Signaling | 4.18E+00 | 5.52E-01 | -4.919 | ACVR1B,ACVR2B,BCL2,BMP7,BMPR1A,BMPR1B,BMPR2,CDC42,CREBBP,EP300,GRB2,HDAC1,HNF4A,KRAS,MAP2K1,MAP2K2,MAP2K3,MAP2K4,MAP2K6,MAP4K1,MAPK1,MAPK13,MAPK14,MAPK8,MAPK9,MRAS,PIAS4,PMEPA1,RAF1,RALA,RALB,RAP1A,RAP1B,RNF111,RRAS2,RUNX2,RUNX3,SERPINE1,SKI,SMAD3,SMAD6,SMAD7,SMURF1,SMURF2,SOS1,TAB1,TGFB1,TGFB2,TGFBR2,TGIF1,VDR,ZFY |
| Ephrin B Signaling | 4.17E+00 | 5.83E-01 | -5.488 | AXIN1,CBL,CDC42,CXCL12,EPHB1,EPHB2,EPHB4,GNA11,GNA12,GNA14,GNAI1,GNAI2,GNAI3,GNAL,GNAO1,GNAQ,GNAS,GNB1,GNB1L,GNB5,GNG12,GNG2,GNG4,GNG7,ITSN2,KALRN,LIMK1,MAPK1,MRAS,NCK2,PAK1,PTK2,PXN,RAC1,RAC2,RGS3,RHOA,ROCK1,ROCK2,VAV1,VAV2,VAV3 |
| Huntington's Disease Signaling | 4.15E+00 | 4.79E-01 | -4.431 | AKT2,AKT3,AP2A2,APAF1,BCL2L1,CACNA1B,CAPN1,CAPN11,CAPN2,CAPN5,CAPN8,CASP8,CASP9,CLTA,CLTB,CLTC,CLTCL1,CPLX2,CREB3L3,CREB5,CREBBP,DCTN1,DLG4,DNAJC5,DNM1,DNM1L,DNM2,DNM3,EGFR,EP300,GNA11,GNA14,GNAQ,GNB1,GNB1L,GNB5,GNG12,GNG2,GNG4,GNG7,GOSR1,GRB2,GRIN2B,GRM1,GRM5,HDAC1,HDAC11,HDAC4,HDAC5,HDAC6,HDAC7,HDAC9,HIP1,HSPA4,HTT,IGF1R,ITPR1,MAP2K4,MAP3K10,MAPK1,MAPK8,MAPK9,MTOR,NCOR1,NCOR2,NSF,NTRK1,PACSIN1,PDPK1,PIK3C2A,PIK3C2B,PIK3CB,PIK3CD,PIK3R2,PIK3R3,PIK3R4,PIK3R5,PIK3R6,PLCB1,PLCB2,PLCB3,POLR2A,POLR2F,POLR2J2/POLR2J3,PRKCA,PRKCB,PRKCD,PRKCE,PRKCG,PRKCH,PRKCI,PRKCQ,PRKCZ,PRKD1,PSME4,RASA1,RCOR1,RCOR3,RPH3A,SDHA,SDHB,SGK1,SH3GL3,SIN3A,SO |
| RhoA Signaling | 4.15E+00 | 5.29E-01 | -4.99 | ABL2,ACTB,ACTG2,ACTR2,ACTR3,ANLN,ARHGAP1,ARHGAP35,ARHGAP6,ARHGEF1,ARHGEF11,ARHGEF12,ARPC1A,ARPC5L,BAIAP2,CDC42EP4,CIT,DLC1,EPHA1,EZR,GNA12,IGF1R,LIMK1,LIMK2,LPAR3,LPAR5,MPRIP,MSN,MYL10,MYL3,MYLK,MYLK2,MYLK3,NEDD4,NGEF,NRP2,PFN2,PI4KA,PIP4K2B,PIP5K1A,PIP5K1B,PIP5K1C,PIP5KL1,PKN1,PLD1,PLEKHG5,PLXNA1,PPP1R12A,PPP1R12B,PTK2,PTK2B,RAPGEF2,RDX,RHOA,RHPN2,ROCK1,ROCK2,SEMA3F,SEPTIN10,SEPTIN14,SEPTIN6,SEPTIN7,S |
| FLT3 Signaling in Hematopoietic Progenitor Cells | 4.07E+00 | 5.70E-01 | -6.41 | AKT2,AKT3,CBL,CREB3L3,CREB5,CREBBP,GAB2,GRB2,INPP5D,KRAS,MAP2K1,MAP2K2,MAPK1,MAPK13,MAPK14,MRAS,MTOR,PDPK1,PIK3C2A,PIK3C2B,PIK3CB,PIK3CD,PIK3R2,PIK3R3,PIK3R4,PIK3R5,PIK3R6,PTPN11,RAF1,RALA,RALB,RAP1A,RAP1B,RPS6KA1,RPS6KA2,RPS6KA3,RPS6KA5,RPS6KB1,RRAS2,SOS1,STAT2,STAT3,STAT5A,STAT5B,STAT6 |
| Gap Junction Signaling | 4.07E+00 | 4.90E-01 | NaN | ACTB,ACTG2,ADCY1,ADCY10,ADCY2,ADCY3,ADCY5,ADCY6,ADCY7,ADCY8,ADCY9,AKT2,AKT3,CSNK1D,CSNK1E,CSNK1G2,DBN1,DRD2,EGFR,GJC1,GNAI1,GNAI2,GNAI3,GNAQ,GNAS,GRB2,GRIA1,GRIA2,GRIA4,GRIK3,GUCY1A2,GUCY2D,HTR2C,ITPR1,ITPR2,ITPR3,KRAS,MAP2K1,MAP2K2,MAP2K5,MAP3K2,MAPK1,MRAS,NPR1,NPR3,PIK3C2A,PIK3C2B,PIK3CB,PIK3CD,PIK3R2,PIK3R3,PIK3R4,PIK3R5,PIK3R6,PLCB1,PLCB2,PLCB3,PLCD3,PLCE1,PLCG1,PLCG2,PLCH1,PLCH2,PLCL1,PLCL2,PPP3CB,PPP3CC,PRKACA,PRKAG2,PRKAR1B,PRKAR2A,PRKCA,PRKCB,PRKCD,PRKCE,PRKCG,PRKCH,PRKCI,PRKCQ,PRKCZ,PRKD1,PRKG1,RAF1,RALA,RALB,RAP1A,RAP1B,RRAS2,SOS1,SP1,SRC,TJP1,TJP2,T |
| ErbB2-ErbB3 Signaling | 4.05E+00 | 5.94E-01 | -5.191 | ERBB2,ERBB3,FOXO1,GRB2,GSK3B,JAK3,KRAS,MAP2K1,MAP2K2,MAPK1,MRAS,NRG1,NRG2,NRG3,NRG4,PDPK1,PIK3C2A,PIK3C2B,PIK3CB,PIK3CD,PIK3R2,PIK3R3,PIK3R4,PIK3R5,PIK3R6,PTEN,RAF1,RALA,RALB,RAP1A,RAP1B,RRAS2,SOS1,SP1,STAT3,STAT5A,STAT5B,TYK2 |
| eNOS Signaling | 4.02E+00 | 5.07E-01 | -7.5 | ADCY1,ADCY10,ADCY2,ADCY3,ADCY5,ADCY6,ADCY7,ADCY8,ADCY9,AKT2,AKT3,AQP1,AQP7,AQP8,CAMK4,CASP8,CASP9,CHRM3,CHRNA2,CHRNA4,CHRNA5,CHRNA7,CHRNB1,CHRNB2,CHRNB4,CNGA3,CNGB1,DNM2,ESR1,ESR2,FLT1,FLT4,GNAQ,GNAS,GUCY1A2,HSP90AA1,HSPA4,ITPR1,ITPR2,ITPR3,LPAR3,LPAR5,MIP,NOS3,NOSIP,PDPK1,PIK3C2A,PIK3C2B,PIK3CB,PIK3CD,PIK3R2,PIK3R3,PIK3R4,PIK3R5,PIK3R6,PLCG1,PLCG2,PRKAA1,PRKAA2,PRKACA,PRKAG2,PRKAR1B,PRKAR2A,PRKCA,PRKCB,PRKCD,PRKCE,PRKCG,PRKCH,PRKCI,PRKCQ,PRKCZ,PRKD1,PRKG1,SLC7A1,VEGFA,VEGFD |
| Endometrial Cancer Signaling | 4.00E+00 | 6.00E-01 | -3.772 | AKT2,AKT3,AXIN1,CASP9,CDH1,CTNNA1,CTNNA2,ERBB2,FOXO3,GRB2,GSK3B,KRAS,MAP2K1,MAP2K2,MAPK1,MLH1,MRAS,PDPK1,PIK3C2A,PIK3C2B,PIK3CB,PIK3CD,PIK3R2,PIK3R3,PIK3R4,PIK3R5,PIK3R6,PTEN,RAF1,RALA,RALB,RAP1A,RAP1B,RRAS2,SOS1,TP53 |
| Caveolar-mediated Endocytosis Signaling | 3.97E+00 | 5.75E-01 | NaN | ABL1,ACTB,ACTG2,ARCN1,CAVIN1,COPA,COPB1,COPG1,COPG2,DNM2,DYRK3,EGFR,FLNA,FLNB,FLNC,FLOT2,FYN,INSR,ITGA10,ITGA11,ITGA2,ITGA2B,ITGA3,ITGA7,ITGA9,ITGAD,ITGAE,ITGAL,ITGAM,ITGAX,ITGB2,ITGB3,ITGB4,ITGB5,ITGB6,ITGB7,ITGB8,MAP3K2,PRKCA,PTPN1,RAB5B,SRC |
| HIPPO signaling | 3.97E+00 | 5.60E-01 | 2.556 | BTRC,CD44,CRB1,CSNK1D,CSNK1E,CUL1,DLG1,DLG2,DLG4,DLG5,ITCH,LATS2,LLGL1,NF2,PARD3,PATJ,PPM1L,PPP1R12A,PPP1R14C,PPP1R14D,PPP1R3A,PPP1R7,PPP2CB,PPP2R1B,PPP2R2A,PPP2R2B,PPP2R2C,PPP2R3A,PPP2R5B,PPP2R5C,PPP2R5D,PPP2R5E,PTPA,SCRIB,SMAD3,STK3,STK4,TEAD1,TEAD3,TEAD4,TJP2,TP53BP2,WWC1,WWTR1,YAP1,YWHAE,YWHAG |

| | | | | |
|---|---|---|---|---|
| NF-κB Activation by Viruses | 3.93E+00 | 5.61E-01 | -6.782 | AKT2,AKT3,CD4,IKBKB,IKBKE,IKBKG,ITGA2,ITGA3,ITGAL,ITGB2,ITGB3,ITGB5,KRAS,LCK,MAP3K1,MAP3K14,MAPK1,MRAS,NFKBID,PIK3C2A,PIK3C2B,PIK3CB,PIK3CD,PIK3R2,PIK3R3,PIK3R4,PIK3R5,PIK3R6,PRKCA,PRKCB,PRKCD,PRKCE,PRKCG,PRKCH,PRKCI,PRKCQ,PRKCZ,PRKD1,RAF1,RALA,RALB,RAP1A,RAP1B,RIPK1,RRAS2,TRAF2 |
| PEDF Signaling | 3.93E+00 | 5.61E-01 | -4.919 | AKT2,AKT3,ARHGAP22,BCL2,BCL2L1,CASP8,DOCK3,HNF1A,HNF1B,IKBKB,IKBKE,IKBKG,KRAS,MAPK1,MAPK13,MAPK14,MRAS,NFKBID,PIK3C2A,PIK3C2B,PIK3CB,PIK3CD,PIK3R2,PIK3R3,PIK3R4,PIK3R5,PIK3R6,PPARG,RAC1,RAF1,RALA,RALB,RAP1A,RAP1B,RHOA,ROCK1,ROCK2,RRAS2,SERPINF1,TCF12,TCF4,TCF7,TCF7L1,TCF7L2,TP53,WASF2 |
| PFKFB4 Signaling Pathway | 3.87E+00 | 6.30E-01 | -4.158 | CREB3L3,CREB5,CREBBP,FBP1,GCK,GNAS,GPI,HK1,HK2,HK3,MAP2K1,MAP2K2,MAP2K3,MAP2K4,MAP2K5,MAP2K6,MAPK1,NCOA3,PFKFB4,PFKM,PRKACA,PRKAG2,PRKAR1B,PRKAR2A,RPS6KA5,TGFB1,TGFB2,TKT,TP53 |
| Renal Cell Carcinoma Signaling | 3.86E+00 | 5.64E-01 | -5.916 | AKT2,AKT3,ARNT,CDC42,CREBBP,CRK,EGLN3,EP300,ETS1,GRB2,HGF,KRAS,MAP2K1,MAP2K2,MAPK1,MET,MRAS,PAK1,PAK2,PAK4,PDGFB,PIK3C2A,PIK3C2B,PIK3CB,PIK3CD,PIK3R2,PIK3R3,PIK3R4,PIK3R5,PIK3R6,PTPN11,RAC1,RAF1,RALA,RALB,RAP1A,RAP1B,RAPGEF1,RRAS2,SLC2A1,SOS1,TG |
| Aldosterone Signaling in Epithelial Cells | 3.78E+00 | 5.00E-01 | -7.147 | AHCY,ASIC1,ASIC2,DNAJB12,DNAJB6,DNAJC1,DNAJC11,DNAJC12,DNAJC13,DNAJC16,DNAJC17,DNAJC18,DNAJC3,DNAJC5,DNAJC6,DNAJC7,DNAJC8,HSP90AA1,HSPA12A,HSPA12B,HSPA4,HSPB8,ITPR1,ITPR2,ITPR3,KRAS,MAP2K1,MAP2K2,MAPK1,NEDD4,NR3C2,PDPK1,PI4KA,PIK3C2A,PIK3C2B,PIK3CB,PIK3CD,PIK3R2,PIK3R3,PIK3R4,PIK3R5,PIK3R6,PIP4K2B,PIP5K1A,PIP5K1B,PIP5K1C,PIP5KL1,PLCB1,PLCB2,PLCB3,PLCD3,PLCE1,PLCG1,PLCG2,PLCH1,PLCH2,PLCL1,PLCL2,PRKCA,PRKCB,PRKCD,PRKCE,PRKCG,PRKCH,PRKCI,PRKCQ,PRKCZ,PRKD1,RAF1,SACS,SCNN1B,SGK1,SLC12A2,SL |
| ILK Signaling | 3.76E+00 | 4.86E-01 | -4.838 | ACTB,ACTG2,ACTN1,ACTN2,ACTN4,AKT2,AKT3,CDC42,CDH1,CREB3L3,CREB5,CREBBP,DOCK1,DSP,FBLIM1,FLNA,FLNB,FLNC,FNBP1,GSK3B,IRS2,ITGB2,ITGB3,ITGB4,ITGB5,ITGB6,ITGB7,ITGB8,LIMS1,LIMS2,MAP2K4,MAP2K6,MAPK1,MAPK10,MAPK8,MAPK9,MTOR,MYH10,MYH11,MYH14,MYH3,MYH6,MYH7,MYH7B,MYH9,MYL3,NACA,NCK2,NOS2,PARVA,PARVB,PDPK1,PIK3C2A,PIK3C2B,PIK3CB,PIK3CD,PIK3R2,PIK3R3,PIK3R4,PIK3R5,PIK3R6,PPM1L,PPP1R12A,PPP2CB,PPP2R1B,PPP2R2A,PPP2R2B,PPP2R2C,PPP2R3A,PPP2R5B,PPP2R5C,PPP2R5D,PPP2R5E,PTEN,PTK2,PTPA,PXN,RAC1,RAC2,RHOA,RHOBTB1,RHOD,RHOQ,RHOT1,RICTOR,RPS6KA5,RSU1,VCL,VEGFA,VEGFD |
| SAPK/JNK Signaling | 3.75E+00 | 5.35E-01 | -6.102 | CDC42,CRK,DUSP10,DUSP4,GNA12,GNB1,GNG2,GNG7,GRB2,KRAS,LCK,MAP2K4,MAP3K1,MAP3K10,MAP3K11,MAP3K12,MAP3K13,MAP3K2,MAP3K3,MAP3K5,MAP4K1,MAP4K2,MAP4K3,MAP4K4,MAPK10,MAPK8,MAPK8IP1,MAPK8IP3,MAPK9,MINK1,MRAS,NFATC1,NFATC3,PIK3C2A,PIK3C2B,PIK3CB,PIK3CD,PIK3R2,PIK3R3,PIK3R4,PIK3R5,PIK3R6,RAC1,RAC2,RALA,RALB,RAP1A,RAP1B,RIPK1,RRAS |
| GM-CSF Signaling | 3.73E+00 | 5.71E-01 | -5.754 | AKT2,AKT3,BCL2L1,CAMK2A,CAMK2B,CAMK2G,CSF2RA,CSF2RB,ETS1,GRB2,HCK,JAK2,KRAS,MAP2K1,MAP2K2,MAPK1,MRAS,PIK3C2A,PIK3C2B,PIK3CB,PIK3CD,PIK3R2,PIK3R3,PIK3R4,PIK3R5,PIK3R6,PPP3CB,PPP3CC,PRKCB,PTPN11,RAF1,RALA,RALB,RAP1A,RAP1B,RRAS2,RUNX1,SOS1,STAT3,S |
| Chemokine Signaling | 3.66E+00 | 5.58E-01 | -5.642 | CAMK1D,CAMK2A,CAMK2B,CAMK2G,CAMK4,CCL24,CCR3,CXCL12,GNAI1,GNAI2,GNAI3,GNAQ,KRAS,LIMK1,LIMK2,MAP2K1,MAP2K2,MAPK1,MAPK13,MAPK14,MAPK8,MPRIP,MRAS,PLCB1,PLCB2,PLCB3,PLCG1,PLCG2,PPP1R12A,PPP1R12B,PRKCA,PRKCB,PTK2,PTK2B,RAF1,RALA,RALB,RAP1A,RAP1B,RHOA,ROCK2,RRAS2,SRC |
| T Cell Receptor Signaling | 3.65E+00 | 5.29E-01 | NaN | CAMK4,CARD11,CBL,CD247,CD4,CD8B,CSK,FYN,GRB2,IKBKB,IKBKE,IKBKG,KRAS,LCK,LCP2,MAP2K1,MAP2K2,MAP2K4,MAP3K1,MAPK1,MAPK8,MRAS,NFAT5,NFATC1,NFATC2,NFATC3,PIK3C2A,PIK3C2B,PIK3CB,PIK3CD,PIK3R2,PIK3R3,PIK3R4,PIK3R5,PIK3R6,PLCG1,PPP3CB,PPP3CC,PRKCQ,PTPN7,PTPRC,PTPRH,RAC1,RAF1,RALA,RALB,RAP1A,RAP1B,RASA1,RRAS2,SOS1,VAV1,VAV2,VAV3,ZAP7 |
| Hereditary Breast Cancer Signaling | 3.57E+00 | 5.04E-01 | NaN | ACTB,ACTL6B,AKT2,AKT3,ARID1A,ARID2,ATM,ATR,BLM,BRCA1,CDC25C,CDK6,CREBBP,DDB2,E2F1,EP300,FAAP100,FANCA,FANCC,FANCD2,FANCL,HDAC1,HDAC11,HDAC4,HDAC5,HDAC6,HDAC7,HDAC9,HLTF,KRAS,MLH1,MRAS,MRE11,MSH2,MSH6,PBRM1,PIK3C2A,PIK3C2B,PIK3CB,PIK3CD,PIK3R2,PIK3R3,PIK3R4,PIK3R5,PIK3R6,PMS2,POLR2A,POLR2F,POLR2J2/POLR2J3,PTEN,RAD50,RAD51,RALA,RALB,RAP1A,RAP1B,RFC1,RFC2,RPA1,RRAS2,SLC19A1,SMARCA2,SMARCA4,SMARCB1,SMARCC1,SMARCD1,SMARCD3,TP53,XPC |
| HOTAIR Regulatory Pathway | 3.56E+00 | 4.94E-01 | -6.496 | AEBP2,AGO1,AGO2,AGO3,AGO4,AKT2,AKT3,ATG3,ATG7,ATXN1,BRD4,CD44,CDH1,COL1A1,CREBBP,DNMT3B,ELAVL1,EP300,ERBB2,ESR1,ESR2,EZH2,HDAC1,HNF1A,HSF1,JARID2,KMT2A,KMT2C,MDM2,MET,MMP14,MMP15,MMP17,MMP2,MMP21,MMP26,MMP28,MRTFA,MTF2,PIK3C2A,PIK3C2B,PIK3CB,PIK3CD,PIK3R2,PIK3R3,PIK3R4,PIK3R5,PIK3R6,PTEN,RBBP4,RCOR1,ROCK1,ROCK2,SETDB1,SNUPN,STAT3,STK38,SUZ12,TCF3,TCF4,TCF7,TCF7L1,TCF7L2,TGFB1,TWIST2,WNT10A,WNT11,WNT2B,WNT3,WNT3A,WNT4,WNT5B,WNT7A,WNT7B,WNT9A,WNT9B,XIAP |
| Sphingosine-1-phosphate Signaling | 3.54E+00 | 5.18E-01 | -4.258 | ACER1,ACER2,ACER3,ADCY1,ADCY10,ADCY2,ADCY3,ADCY5,ADCY6,ADCY7,ADCY8,ADCY9,AKT2,AKT3,CASP8,CASP9,CDC42,FNBP1,GNA12,GNAI1,GNAI2,GNAI3,GNAQ,MAPK1,PDGFA,PDGFB,PDGFD,PDGFRB,PIK3C2A,PIK3C2B,PIK3CB,PIK3CD,PIK3R2,PIK3R3,PIK3R4,PIK3R5,PIK3R6,PLCB1,PLCB2,PLCB3,PLCD3,PLCE1,PLCG1,PLCG2,PLCH1,PLCH2,PLCL1,PLCL2,PTK2,PTK2B,RAC1,RAC2,RHOA,RHOBTB1,RHOD,RHOQ,RHOT1,SMPD3,SMPD4 |
| Role of Macrophages, Fibroblasts and Endothelial Cells in Rheumatoid Arthritis | 3.53E+00 | 4.53E-01 | NaN | AKT2,AKT3,APC,AXIN1,C5,C5AR1,CAMK2A,CAMK2B,CAMK2G,CAMK4,CHP1,CREB3L3,CREB5,CREBBP,CSF1,CXCL12,DKK3,FZD3,GNAO1,GNAQ,GSK3B,HNF1A,IKBKB,IKBKE,IKBKG,IL16,IL17RA,IL17RC,IL1R1,IL1RAP,IL1RAPL1,IL1RAPL2,IL32,IL6R,IL6ST,IRAK2,JAK2,KRAS,LRP1,LRP5,LRP6,MAP2K1,MAP2K2,MAP2K3,MAP2K4,MAP2K6,MAP3K14,MAPK1,MAPK14,MAPK9,MAPKAPK2,MRAS,NFAT5,NFATC1,NFATC2,NFATC3,NFKBID,NLK,NOS2,PDGFA,PDGFB,PDGFD,PIK3C2A,PIK3C2B,PIK3CB,PIK3CD,PIK3R2,PIK3R3,PIK3R4,PIK3R5,PIK3R6,PLCB1,PLCB2,PLCB3,PLCD3,PLCE1,PLCG1,PLCG2,PLCH1,PLCH2,PLCL1,PLCL2,PPP3CB,PPP3CC,PRKCA,PRKCB,PRKCD,PRKCE,PRKCG,PRKCH,PRKCI,PRKCQ,PRKCZ,PRKD1,RAC1,RAF1,RALA,RALB,RAP1A,RAP1B,RHOA,RIPK1,ROCK1,ROCK2,ROR2,RRAS2,RYK,SFRP1,SMO,SRC,STAT3,TCF3,TCF4,TCF7,TCF7L1,TCF7L2,TGFB1,TLR5,TNFRSF1B,TRAF1,TRAF2,TRAF3,TRAF5,VEGFA,VEGFD,WNT10A,WNT11,WNT2B,WNT3,WNT3A,WNT4,WNT5B,WNT7A,WNT7B |
| Role of BRCA1 in DNA Damage Response | 3.53E+00 | 5.50E-01 | -1.46 | ACTB,ACTL6B,ARID1A,ARID2,ATF1,ATM,ATR,BABAM2,BLM,BRCA1,BRIP1,E2F1,E2F2,E2F3,E2F4,FAAP100,FANCA,FANCC,FANCD2,FANCL,HLTF,MLH1,MRE11,MSH2,MSH6,PBRM1,POU2F1,RAD50,RAD51,RBBP8,RBL1,RFC1,RFC2,RPA1,SLC19A1,SMARCA2,SMARCA4,SMARCB1,SMARCC1,SMARCD1,SMARCD3,TOPBP1,TP53,UIMC1 |
| Production of Nitric Oxide and Reactive Oxygen Species in Macrophages | 3.52E+00 | 4.81E-01 | -5.522 | AKT2,AKT3,CDC42,CLU,CREBBP,FNBP1,IFNGR2,IKBKB,IKBKE,IKBKG,IRF8,JAK1,JAK2,JAK3,LPA,MAP2K1,MAP2K4,MAP3K1,MAP3K10,MAP3K11,MAP3K12,MAP3K13,MAP3K14,MAP3K15,MAP3K2,MAP3K3,MAP3K5,MAP3K6,MAPK1,MAPK10,MAPK13,MAPK14,MAPK8,MAPK9,NFKBID,NOS2,PCYOX1,PIK3C2A,PIK3C2B,PIK3CB,PIK3CD,PIK3R2,PIK3R3,PIK3R4,PIK3R5,PIK3R6,PLCG1,PLCG2,PPARA,PPM1L,PPP1R12A,PPP1R14C,PPP1R14D,PPP1R3A,PPP1R7,PPP2CB,PPP2R1B,PPP2R2A,PPP2R2B,PPP2R2C,PPP2R3A,PPP2R5B,PPP2R5C,PPP2R5D,PPP2R5E,PRKCA,PRKCB,PRKCD,PRKCE,PRKCG,PRKCH,PRKCI,PRKCQ,PRKCZ,PRKD1,PTPA,RAC1,RAC2,RAP1A,RAP1B,RHOA,RHOBTB1,RHOD,RHOQ,RHOT |

| | | | | |
|---|---|---|---|---|
| Amyotrophic Lateral Sclerosis Signaling | 3.49E+00 | 5.31E-01 | -4.621 | AKT3,APAF1,BCL2,BCL2L1,BID,BIRC2,CACNA1A,CACNA1C,CACNA1D,CACNA1E,CACNA1S,CAPN1,CAPN11,CAPN2,CAPN5,CAPN8,CASP9,CCS,GRIA1,GRIA2,GRIA4,GRID1,GRID2,GRIK3,GRIK4,GRIK5,GRIN1,GRIN2A,GRIN2B,GRIN2C,GRIN2D,HECW1,NAIP,NOS1,PAK1,PIK3C2A,PIK3C2B,PIK3CB,PIK3CD,PIK3R2,PIK3R3,PIK3R4,PIK3R5,PIK3R6,RAB5B,RAC1,SLC1A2,TP53,VEGFA,VEGFD,XIAP |
| PAK Signaling | 3.49E+00 | 5.31E-01 | -6.788 | ARHGAP10,ARHGEF7,CDC42,DSCAM,EPHA3,GRB2,ITGA2,ITGA3,KRAS,LIMK1,LIMK2,MAP2K1,MAP2K2,MAP2K4,MAPK1,MAPK10,MAPK8,MAPK9,MRAS,MYL10,MYL3,MYLK,NCK1,NCK2,PAK1,PAK2,PAK4,PDGFA,PDGFB,PDGFD,PDGFRB,PIK3C2A,PIK3C2B,PIK3CB,PIK3CD,PIK3R2,PIK3R3,PIK3R4,PIK3R5,PIK3R6,PTK2,PTK2B,PXN,RAC1,RAF1,RALA,RALB,RAP1A,RAP1B,RRAS2,SOS1 |
| Role of MAPK Signaling in the Pathogenesis of Influenza | 3.49E+00 | 5.51E-01 | NaN | ABHD3,AKT2,AKT3,BCL2,KRAS,MAP2K1,MAP2K2,MAP2K3,MAP2K4,MAP2K6,MAP3K5,MAPK1,MAPK10,MAPK13,MAPK14,MAPK8,MAPK9,MRAS,PLA2G10,PLA2G12B,PLA2G2C,PLA2G2F,PLA2G4A,PLA2G4C,PLA2G4D,PLA2G4E,PLA2G4F,PLA2G5,PLA2G6,PLA2R1,PLAAT3,PLAAT4,PLAAT5,PLB1,PNPLA3,PNPLA8,PRKCA,RAF1,RALA,RALB,RAP1A,RAP1B,RRAS2 |
| CD27 Signaling in Lymphocytes | 3.46E+00 | 5.96E-01 | -4.017 | APAF1,BCL2L1,BID,CASP8,CASP9,IKBKB,IKBKE,IKBKG,MAP2K1,MAP2K2,MAP2K3,MAP2K4,MAP2K5,MAP2K6,MAP3K1,MAP3K10,MAP3K11,MAP3K12,MAP3K13,MAP3K14,MAP3K15,MAP3K2,MAP3K3,MAP3K5,MAP3K6,MAPK10,MAPK8,MAPK9,NFKBID,TRAF2,TRAF5 |
| GDNF Family Ligand-Receptor Interactions | 3.45E+00 | 5.53E-01 | -6.083 | CDC42,DOK4,DOK5,FRS2,GFRA1,GFRA2,GRB2,IRS2,ITPR1,ITPR2,ITPR3,KRAS,MAP2K1,MAP2K2,MAP2K4,MAPK1,MAPK10,MAPK8,MAPK9,MRAS,NCK1,PIK3C2A,PIK3C2B,PIK3CB,PIK3CD,PIK3R2,PIK3R3,PIK3R4,PIK3R5,PIK3R6,PLCG1,PLCG2,RAC1,RAF1,RALA,RALB,RAP1A,RAP1B,RASA1,RET,RRAS2, |
| Neurotrophin/TRK Signaling | 3.45E+00 | 5.53E-01 | -6.083 | CDC42,CREB3L3,CREB5,CREBBP,FRS2,GRB2,KRAS,MAP2K1,MAP2K2,MAP2K3,MAP2K4,MAP2K5,MAP2K6,MAP3K5,MAPK1,MAPK8,MRAS,NTF3,NTRK1,NTRK2,NTRK3,PDPK1,PIK3C2A,PIK3C2B,PIK3CB,PIK3CD,PIK3R2,PIK3R3,PIK3R4,PIK3R5,PIK3R6,PLCG1,PTPN11,RAF1,RALA,RALB,RAP1A,RAP1B,RPS6KA1,RRAS2,SORCS1,SOS1 |
| Dopamine Receptor Signaling | 3.45E+00 | 5.53E-01 | -3.153 | ADCY1,ADCY10,ADCY2,ADCY3,ADCY5,ADCY6,ADCY7,ADCY8,ADCY9,COMT,DDC,DRD2,DRD3,IL4I1,MAOA,MAOB,NCS1,PPM1L,PPP1R12A,PPP1R14C,PPP1R14D,PPP1R3A,PPP1R7,PPP2CB,PPP2R1B,PPP2R2A,PPP2R2B,PPP2R2C,PPP2R3A,PPP2R5B,PPP2R5C,PPP2R5D,PPP2R5E,PRKACA,PRKAG2,PRKAR1B,PRKAR2A,PTPA,SLC18A1,SLC18A2,SLC6A3,SMOX |
| Role of NANOG in Mammalian Embryonic Stem Cell Pluripotency | 3.45E+00 | 5.13E-01 | -5.145 | AKT2,AKT3,APC,AXIN1,BMP1,BMP5,BMP6,BMP7,BMP8A,BMP8B,BMPR1A,BMPR1B,BMPR2,FZD3,GATA4,GRB2,GSK3B,IL6ST,JAK1,JAK2,JAK3,KRAS,LIFR,MAP2K1,MAP2K2,MAPK1,MRAS,PIK3C2A,PIK3C2B,PIK3CB,PIK3CD,PIK3R2,PIK3R3,PIK3R4,PIK3R5,PIK3R6,RAF1,RALA,RALB,RAP1A,RAP1B,RRAS2,SALL4,SMO,SOS1,STAT3,TCF7L1,TP53,TYK2,WNT10A,WNT11,WNT2B,WNT3,WNT3A,WNT4,WNT5B,WNT7A,WNT7B,WNT9A,WNT9B |
| Role of Osteoblasts, Osteoclasts and Chondrocytes in Rheumatoid Arthritis | 3.43E+00 | 4.70E-01 | NaN | AKT2,AKT3,ALPL,APC,AXIN1,BCL2,BIRC2,BMP1,BMP5,BMP6,BMP7,BMP8A,BMP8B,BMPR1A,BMPR2,CALCR,CAMK4,CASP9,CBL,CHP1,COL1A1,CSF1,CSF1R,DKK3,FOXO1,FZD3,GSK3B,GSN,HNF1A,IKBKB,IKBKE,IKBKG,IL1R1,IL1RAP,IL1RAPL1,IL1RAPL2,ITGA2,ITGA3,ITGB3,LRP1,LRP5,LRP6,MAP2K3,MAP2K4,MAP2K6,MAP3K14,MAP3K5,MAPK1,MAPK10,MAPK14,MAPK8,MAPK9,MMP14,NAIP,NFAT5,NFATC1,NFATC2,NFATC3,NFKBID,PIK3C2A,PIK3C2B,PIK3CB,PIK3CD,PIK3R2,PIK3R3,PIK3R4,PIK3R5,PIK3R6,PPP3CB,PPP3CC,PTK2B,RUNX2,SFRP1,SMAD6,SMO,SMURF1,SP7,SRC,TAB2,TCF3,TCF4,TCF7,TCF7L1,TCF7L2,TGFB1,TNFRSF11A,TNFRSF1B,TRAF2,TRAF5,WNT10A,WNT11,WNT2B,WNT3,WNT3A.WNT4.WNT5B.WNT7A.WNT7B.WNT9A.WNT9B.XIAP |
| Angiopoietin Signaling | 3.41E+00 | 5.54E-01 | -3.413 | AKT2,AKT3,ANGPT4,CASP9,CRK,FOXO1,GRB2,IKBKB,IKBKE,IKBKG,KRAS,MRAS,NCK1,NFKBID,NOS3,PAK1,PAK2,PAK4,PIK3C2A,PIK3C2B,PIK3CB,PIK3CD,PIK3R2,PIK3R3,PIK3R4,PIK3R5,PIK3R6,PTK2,PTPN11,RALA,RALB,RAP1A,RAP1B,RASA1,RRAS2,SOS1,STAT5A,STAT5B,TEK,TIE1,TNIP1 |
| IGF-1 Signaling | 3.34E+00 | 5.19E-01 | -5.774 | AKT2,AKT3,CASP9,CSNK2A1,CSNK2A2,FOXO1,FOXO3,GRB10,GRB2,IGF1R,IGFBP4,IRS2,JAK1,JAK2,KRAS,MAP2K1,MAP2K2,MAPK1,MAPK8,MRAS,NEDD4,PDPK1,PIK3C2A,PIK3C2B,PIK3CB,PIK3CD,PIK3R2,PIK3R3,PIK3R4,PIK3R5,PIK3R6,PRKACA,PRKAG2,PRKAR1B,PRKAR2A,PRKCI,PRKCZ,PTK2,PTPN11,PXN,RAF1,RALA,RALB,RAP1A,RAP1B,RASA1,RPS6KB1,RRAS2,SOCS5,SOCS7,SOS1,STAT3,YW |
| Semaphorin Signaling in Neurons | 3.32E+00 | 5.79E-01 | NaN | ARHGAP1,ARHGEF12,CDC42,CRMP1,DPYSL2,DPYSL3,DPYSL4,DPYSL5,FNBP1,FYN,LIMK1,LIMK2,MAPK1,MET,NRP1,PAK1,PAK2,PAK4,PLXNA1,PLXNB1,PTK2,RAC1,RAC2,RHOA,RHOBTB1,RHOD,RHOQ,RHOT1,ROCK1,ROCK2,SEMA3A,SEMA4D,SEMA7A |
| Role of CHK Proteins in Cell Cycle Checkpoint Control | 3.32E+00 | 5.79E-01 | -2.2 | ATM,ATR,BRCA1,CDC25A,CDC25C,CLSPN,E2F1,E2F2,E2F3,E2F4,HUS1,MRE11,PPM1L,PPP2CB,PPP2R1B,PPP2R2A,PPP2R2B,PPP2R2C,PPP2R3A,PPP2R5B,PPP2R5C,PPP2R5D,PPP2R5E,PTPA,RAD17,RAD50,RFC1,RFC2,RPA1,SLC19A1,TLK1,TLK2,TP53 |
| Thyroid Cancer Signaling | 3.29E+00 | 5.45E-01 | -5.246 | AKT2,AKT3,ALK,BRAF,CDH1,FOXO1,GSK3B,HNF1A,INSR,IRS2,KRAS,MAP2K1,MAP2K2,MAPK1,MRAS,MTOR,NTF3,NTRK1,NTRK2,NTRK3,PIK3C2A,PIK3C2B,PIK3CB,PIK3CD,PIK3R2,PIK3R3,PIK3R4,PIK3R5,PIK3R6,PTEN,RALA,RALB,RAP1A,RAP1B,RET,RRAS2,TCF3,TCF4,TCF7,TCF7L1,TCF7L2,TP53 |
| G Protein Signaling Mediated by Tubby | 3.28E+00 | 6.67E-01 | NaN | ABL1,GNAQ,GNAS,GNB1,GNB1L,GNB5,GNG12,GNG2,GNG4,GNG7,INSR,JAK2,LCK,MRAS,PLCB1,PLCB2,PLCB3,PLCG1,PLCG2,TUB |
| TR/RXR Activation | 3.25E+00 | 5.36E-01 | NaN | ACACA,AKT2,AKT3,ATP2A1,CAMK4,COL6A3,DIO1,EP300,F10,G6PC,LDLR,MDM2,MED1,MTOR,NCOA1,NCOA3,NCOA4,NCOA6,NCOR1,NCOR2,PDE3B,PFKP,PIK3C2A,PIK3C2B,PIK3CB,PIK3CD,PIK3R2,PIK3R3,PIK3R4,PIK3R5,PIK3R6,RAB3B,RCAN2,RXRA,RXRG,SCARB1,SLC2A1,SREBF1,SREBF2,STRBP,SYT12,SYT2,TBL1XR1,THRA,THRB |
| FGF Signaling | 3.25E+00 | 5.36E-01 | -6.708 | AKT2,AKT3,CREB3L3,CREB5,CREBBP,CRK,FGF1,FGF12,FGF14,FGF18,FGFR1,FGFR2,FGFR4,FRS2,GRB2,HGF,ITPR1,MAP2K1,MAP2K3,MAP2K6,MAP3K1,MAP3K5,MAPK1,MAPK13,MAPK14,MAPK8,MAPKAPK2,MET,PIK3C2A,PIK3C2B,PIK3CB,PIK3CD,PIK3R2,PIK3R3,PIK3R4,PIK3R5,PIK3R6,PLCG1,PRKCA,PTPN11,RAC1,RAF1,RPS6KA5,SOS1,STAT3 |
| IL-15 Signaling | 3.25E+00 | 5.47E-01 | -6.403 | AKT2,AKT3,BCL2L1,GNB1L,IL15RA,IL2RB,JAK1,JAK2,JAK3,KRAS,LCK,MAP2K1,MAP2K2,MAPK1,MAPK13,MAPK14,MRAS,MTOR,PIK3C2A,PIK3C2B,PIK3CB,PIK3CD,PIK3R2,PIK3R3,PIK3R4,PIK3R5,PIK3R6,RAF1,RALA,RALB,RAP1A,RAP1B,RPS6KB1,RPTOR,RRAS2,STAT3,STAT5A,STAT5B,STAT6,SYK,TYK |
| CD40 Signaling | 3.23E+00 | 5.62E-01 | -5.145 | ATF1,FCER2,IKBKB,IKBKE,IKBKG,JAK3,MAP2K1,MAP2K2,MAP2K3,MAP2K4,MAP2K5,MAP2K6,MAP3K14,MAPK1,MAPK10,MAPK13,MAPK14,MAPK8,MAPK9,MAPKAPK2,NFKBID,PIK3C2A,PIK3C2B,PIK3CB,PIK3CD,PIK3R2,PIK3R3,PIK3R4,PIK3R5,PIK3R6,PTGS1,STAT3,TRAF1,TRAF2,TRAF3,TRAF5 |
| Pyridoxal 5'-phosphate Salvage Pathway | 3.23E+00 | 5.62E-01 | -6 | AKT2,BRAF,CDK18,CDK6,CDK7,CSNK1D,DAPK1,FAM20B,G6PC,GRK4,GRK5,GRK6,LIMK1,LIMK2,MAP2K1,MAP2K2,MAP2K3,MAP2K4,MAP2K6,MAP3K6,MAPK1,MAPK6,MAPK8,MAPK9,PAK1,PAK2,PDXK,PKN1,PRKAA1,PRKAA2,PRKCD,PRKCE,PRKCH,PRKCQ,PRKX,SGK1 |
| p53 Signaling | 3.21E+00 | 5.20E-01 | -0.469 | ADGRB1,AKT2,AKT3,APAF1,ATM,ATR,BCL2,BCL2L1,BRCA1,COQ8A,CSNK1D,DRAM1,E2F1,EP300,GML,GSK3B,HDAC1,HDAC9,HIPK2,JMY,KAT2B,MAPK14,MAPK8,MDM2,MDM4,MED1,PIAS1,PIK3C2A,PIK3C2B,PIK3CB,PIK3CD,PIK3R2,PIK3R3,PIK3R4,PIK3R5,PIK3R6,PLAGL1,PML,PPP1R13B,PRKDC,PTEN,SIRT1,STAG1,TNFRSF10A,TNFRSF10B,TOPBP1,TP53,TP53BP2,TP73,TRIM29,WT1 |
| Calcium Transport I | 3.21E+00 | 9.00E-01 | -3 | ATP2A1,ATP2A2,ATP2A3,ATP2B1,ATP2B2,ATP2B3,ATP2B4,ATP2C1,ATP2C2 |

| | | | | |
|---|---|---|---|---|
| Xenobiotic Metabolism General Signaling Pathway | 3.19E+00 | 4.93E-01 | -7.632 | ABCA2,ABCA4,AKT2,AKT3,ARNT,CREBBP,CUL3,EP300,GCLC,KEAP1,KRAS,MAP2K1,MAP2K2,MAP2K3,MAP2K4,MAP2K5,MAP2K6,MAP3K1,MAP3K10,MAP3K11,MAP3K12,MAP3K13,MAP3K14,MAP3K15,MAP3K2,MAP3K3,MAP3K5,MAP3K6,MAPK1,MAPK13,MAPK14,MAPK8,MAPK9,MRAS,NQO1,NQO2,NR1H2,NR1H3,NR1H4,NR1I2,PIK3C2A,PIK3C2B,PIK3CB,PIK3CD,PIK3R2,PIK3R3,PIK3R4,PIK3R5,PIK3R6,PPARA,PRKCA,PRKCB,PRKCD,PRKCE,PRKCG,PRKCH,PRKCI,PRKCQ,PRKCZ,PRKD1,RAF1,RALA,RALB,RAP1A,RAP1B,RRAS2,RXRA,SLC9C1 |
| Regulation of the Epithelial-Mesenchymal Transition Pathway | 3.18E+00 | 4.71E-01 | NaN | AKT2,AKT3,APC,AXIN1,BCL9,BRAF,CDH1,CDH12,EGFR,ETS1,FGF1,FGF12,FGF14,FGF18,FGFR1,FGFR2,FGFR4,FRS2,FZD3,GRB2,GSK3B,HGF,HNF1A,JAK1,JAK2,JAK3,KRAS,MAML1,MAP2K1,MAP2K2,MAP2K3,MAP2K4,MAP2K5,MAP2K6,MAPK1,MET,MMP2,MRAS,NOTCH1,NOTCH2,NOTCH3,PARD6B,PDGFD,PDGFRB,PIK3C2A,PIK3C2B,PIK3CB,PIK3CD,PIK3R2,PIK3R3,PIK3R4,PIK3R5,PIK3R6,PSEN1,PSEN2,PTPN11,RAF1,RALA,RALB,RAP1A,RAP1B,RBPJ,RHOA,RRAS2,SMAD3,SMO,SMURF1,SOS1,STAT3,TCF3,TCF4,TCF7,TCF7L1,TCF7L2,TGFB1,TGFB2,TGFBR2,TWIST2,TYK2,WNT10A,WNT11,WNT2B,WNT3,WNT3A,WNT4,WNT5B,WNT7A,WNT7B,WNT9A,WNT9B |
| RAN Signaling | 3.15E+00 | 7.65E-01 | -3.606 | CSE1L,IPO5,KPNA1,KPNA3,KPNA4,KPNA6,KPNA7,KPNB1,RANBP2,RANGAP1,RCC1,TNPO1,XPO1 |
| Mechanisms of Viral Exit from Host Cells | 3.09E+00 | 6.10E-01 | NaN | ACTB,ACTG2,CHMP4B,CHMP6,LMNB1,LMNB2,NEDD4,PDCD6IP,PRKCA,PRKCB,PRKCD,PRKCE,PRKCG,PRKCH,PRKCI,PRKCQ,PRKCZ,PRKD1,SH3GL1,SH3GL2,SH3GL3,SH3GLB2,TSG101,VPS37C,XPO |
| D-myo-inositol-5-phosphate Metabolism | 3.08E+00 | 4.83E-01 | -8.485 | ACP6,ALPL,CDC25A,CDC25C,CILP,DOT1L,DUSP10,DUSP14,DUSP15,DUSP16,DUSP27,EPHX2,EYA4,HACD2,INPP5B,INPP5F,ITPK1,MTMR4,MTMR7,PALD1,PAWR,PIP4K2A,PLCB1,PLCB2,PLCB3,PLCD3,PLCE1,PLCG1,PLCG2,PLCH1,PLPP7,PPFIA2,PPFIA3,PPFIA4,PPFIBP2,PPIP5K1,PPM1H,PPP1R12A,PPP1R12C,PPP1R13B,PPP1R16B,PPP1R1C,PPP1R7,PPP1R8,PPP2R3A,PPP2R5B,PPP2R5D,PPP2R5E,PPP4C,PPP5C,PPTC7,PRUNE1,PTEN,PTPA,PTPN1,PTPN11,PTPN12,PTPN2,PTPN20,PTPN22,PTPN23,PTPN7,PTPRC,PTPRF,PTPRH,PTPRJ,PTPRM,PXYLP1,RASA1,SIRPA,STYXL1,TNS3,TPTE |
| CTLA4 Signaling in Cytotoxic T Lymphocytes | 3.01E+00 | 5.23E-01 | NaN | AKT2,AKT3,AP1B1,AP1G1,AP1M1,AP1S3,AP2A1,AP2A2,AP2B1,CD247,CD8B,CLTA,CLTB,CLTC,CLTCL1,FYN,GRB2,JAK2,LCK,LCP2,PIK3C2A,PIK3C2B,PIK3CB,PIK3CD,PIK3R2,PIK3R3,PIK3R4,PIK3R5,PIK3R6,PLCG1,PPM1L,PPP2CB,PPP2R1B,PPP2R2A,PPP2R2B,PPP2R2C,PPP2R3A,PPP2R5B,PPP2R5C,PPP2R5D,PPP2R5E,PTPA,PTPN11,PTPN22,SYK,ZAP70 |
| Role of JAK family kinases in IL-6-type Cytokine Signaling | 3.01E+00 | 6.80E-01 | NaN | IL6R,IL6ST,JAK1,JAK2,MAP2K4,MAPK1,MAPK10,MAPK13,MAPK14,MAPK8,MAPK9,OSMR,PTPN11,STAT3,STAT5A,STAT5B,TYK2 |
| D-myo-inositol (1,4,5)-Trisphosphate Biosynthesis | 3.01E+00 | 6.80E-01 | -4.123 | PI4K2A,PI4KA,PI4KB,PIP4K2A,PIP4K2B,PIP5K1A,PIP5K1B,PIP5K1C,PIP5KL1,PLCB1,PLCB2,PLCB3,PLCD3,PLCE1,PLCG1,PLCG2,PLCH1 |
| Melanoma Signaling | 3.00E+00 | 5.80E-01 | -4.707 | AKT2,AKT3,BRAF,CDH1,E2F1,KRAS,MAP2K1,MAP2K2,MAPK1,MDM2,MITF,MRAS,PIK3C2A,PIK3C2B,PIK3CB,PIK3CD,PIK3R2,PIK3R3,PIK3R4,PIK3R5,PIK3R6,PTEN,RAF1,RALA,RALB,RAP1A,RAP1B,RRA |
| Mouse Embryonic Stem Cell Pluripotency | 3.00E+00 | 5.10E-01 | -6.021 | AKT2,AKT3,APC,AXIN1,BMPR1A,BMPR2,CREBBP,FZD3,GRB2,GSK3B,HNF1A,IL6ST,JAK1,JAK2,JAK3,KRAS,LIFR,MAP2K1,MAP2K2,MAPK1,MAPK13,MAPK14,MRAS,PIK3C2A,PIK3C2B,PIK3CB,PIK3CD,PIK3R2,PIK3R3,PIK3R4,PIK3R5,PIK3R6,PTPN11,RAF1,RALA,RALB,RAP1A,RAP1B,RRAS2,SMO,SOS1,STAT3,TAB1,TCF3,TCF4,TCF7,TCF7L1,TCF7L2,TP53,TYK2,WNT3A,XIAP |
| 3-phosphoinositide Degradation | 2.94E+00 | 4.80E-01 | -8.426 | ACP6,ALPL,CDC25A,CDC25C,CILP,DOT1L,DUSP10,DUSP14,DUSP15,DUSP16,DUSP27,EPHX2,EYA4,HACD2,INPP4A,INPP4B,INPP5B,INPP5D,INPP5F,INPP5K,INPPL1,ITPK1,MTMR14,MTMR3,MTMR4,MTMR7,PALD1,PAWR,PLPP7,PPFIA2,PPFIA3,PPFIA4,PPFIBP2,PPIP5K1,PPM1H,PPP1R12A,PPP1R12C,PPP1R13B,PPP1R16B,PPP1R1C,PPP1R7,PPP1R8,PPP2R3A,PPP2R5B,PPP2R5D,PPP2R5E,PPP4C,PPP5C,PPTC7,PRUNE1,PTEN,PTPA,PTPN1,PTPN11,PTPN12,PTPN2,PTPN20,PTPN22,PTPN23,PTPN7,PTPRC,PTPRF,PTPRH,PTPRJ,PTPRM,PXYLP1,RASA1,SIRPA,STYXL1,SYNJ2,TNS3,TPTE |
| Apoptosis Signaling | 2.91E+00 | 5.10E-01 | 0.866 | APAF1,BCL2,BCL2L1,BCL2L11,BID,BIRC2,BIRC6,CAPN1,CAPN11,CAPN2,CAPN5,CAPN8,CASP8,CASP9,DFFA,DFFB,IKBKB,IKBKE,IKBKG,KRAS,LMNA,MAP2K1,MAP2K2,MAP2K4,MAP3K14,MAP3K5,MAP4K4,MAPK1,MAPK8,MRAS,NAIP,NFKBID,PARP1,PLCG1,PLCG2,PRKCA,PRKCE,PRKCQ,RAF1,RALA,RALB,RAP1A,RAP1B,ROCK1,RPS6KA1,RRAS2,SPTAN1,TNFRSF1B,TP53,XIAP |
| Cholecystokinin/Gastrin-mediated Signaling | 2.89E+00 | 4.96E-01 | -7.348 | BCAR1,CDC42,EGFR,FNBP1,GNA12,GNAQ,GRB2,ITPR1,ITPR2,ITPR3,KRAS,MAP2K1,MAP2K2,MAP2K3,MAP2K4,MAP2K5,MAP2K6,MAPK1,MAPK10,MAPK14,MAPK8,MAPK9,MEF2A,MEF2D,MRAS,PLCB1,PLCB2,PLCB3,PRKCA,PRKCB,PRKCD,PRKCE,PRKCG,PRKCH,PRKCI,PRKCQ,PRKCZ,PRKD1,PTK2,PTK2B,PXN,RAC1,RAC2,RAF1,RALA,RALB,RAP1A,RAP1B,RHOA,RHOBTB1,RHOD,RHOQ,RHOT1,ROCK1,ROCK2,RRAS2,SOS1,SRC |
| DNA Double-Strand Break Repair by Homologous | 2.89E+00 | 7.86E-01 | NaN | ABL1,ATM,ATRX,BRCA1,LIG1,MRE11,POLA1,RAD50,RAD51,RAD52,RPA1 |
| CD28 Signaling in T Helper Cells | 2.84E+00 | 4.96E-01 | -6.877 | ACTR2,ACTR3,AKT2,AKT3,ARPC1A,ARPC5L,CAMK4,CARD11,CD247,CD4,CDC42,CHP1,CSK,FYN,GRB2,IKBKB,IKBKE,IKBKG,ITPR1,ITPR2,ITPR3,LCK,LCP2,MAP2K1,MAP2K2,MAP2K4,MAP3K1,MAPK10,MAPK8,MAPK9,NFAT5,NFATC1,NFATC2,NFATC3,NFKBID,PAK1,PDPK1,PIK3C2A,PIK3C2B,PIK3CB,PIK3CD,PIK3R2,PIK3R3,PIK3R4,PIK3R5,PIK3R6,PLCG1,PPP3CB,PPP3CC,PRKCQ,PTPN11,PTPRC,RAC |
| JAK/Stat Signaling | 2.84E+00 | 5.25E-01 | -4.938 | AKT2,AKT3,BCL2L1,GNAQ,GRB2,JAK1,JAK2,JAK3,KRAS,MAP2K1,MAP2K2,MAPK1,MRAS,MTOR,PIAS1,PIAS4,PIK3C2A,PIK3C2B,PIK3CB,PIK3CD,PIK3R2,PIK3R3,PIK3R4,PIK3R5,PIK3R6,PTPN1,PTPN11,RAF1,RALA,RALB,RAP1A,RAP1B,RRAS2,SOCS5,SOCS7,SOS1,STAT2,STAT3,STAT5A,STAT5B,STAT6, |
| Small Cell Lung Cancer Signaling | 2.83E+00 | 5.35E-01 | -4.041 | ABL1,AKT2,AKT3,APAF1,BCL2,BCL2L1,BID,BIRC2,CASP9,CDK6,E2F1,FHIT,IKBKB,IKBKE,IKBKG,NFKBID,NOS1,PIK3C2A,PIK3C2B,PIK3CB,PIK3CD,PIK3R2,PIK3R3,PIK3R4,PIK3R5,PIK3R6,PTEN,PTK2,RARB,RXRA,RXRG,SIN3A,TFDP1,TP53,TRAF1,TRAF2,TRAF3,TRAF5 |
| Docosahexaenoic Acid (DHA) Signaling | 2.82E+00 | 6.05E-01 | NaN | AKT2,AKT3,ALOX15,APAF1,APP,BCL2,BCL2L1,BID,BIK,CASP9,FOXO1,GSK3B,PDPK1,PIK3C2A,PIK3C2B,PIK3CB,PIK3CD,PIK3R2,PIK3R3,PIK3R4,PIK3R5,PIK3R6,SERPINF1 |
| Human Embryonic Stem Cell Pluripotency | 2.81E+00 | 4.85E-01 | NaN | AKT2,AKT3,APC,AXIN1,BMP1,BMP5,BMP6,BMP7,BMP8A,BMP8B,BMPR1A,BMPR2,FGFR1,FGFR2,FGFR4,FOXO1,FZD3,GNAS,GSK3B,HNF1A,MRAS,NTF3,NTRK1,NTRK2,NTRK3,PDGFA,PDGFB,PDGFD,PDGFRB,PDPK1,PIK3C2A,PIK3C2B,PIK3CB,PIK3CD,PIK3R2,PIK3R3,PIK3R4,PIK3R5,PIK3R6,SALL4,SMAD3,SMAD6,SMAD7,SMO,TCF3,TCF4,TCF7,TCF7L1,TCF7L2,TDGF1,TGFB1,TGFB2,TGFBR2,WNT10A,WNT11,WNT2B,WNT3,WNT3A,WNT4,WNT5B,WNT7A,WNT7B,WNT9A,WNT9B |
| Calcium-induced T Lymphocyte Apoptosis | 2.79E+00 | 5.50E-01 | -5.112 | ATP2A1,ATP2A2,ATP2A3,CABIN1,CAMK4,CAPN2,CD247,CD4,CHP1,EP300,HDAC1,ITPR1,ITPR2,ITPR3,LCK,MEF2D,NFATC2,NR4A1,ORAI1,PLCG1,PPP3CB,PPP3CC,PRKCA,PRKCB,PRKCD,PRKCE,PRKCG,PRKCH,PRKCI,PRKCQ,PRKCZ,PRKD1,ZAP70 |
| Role of NFAT in Regulation of the Immune Response | 2.77E+00 | 4.66E-01 | -7.276 | AKT2,AKT3,BLNK,CABIN1,CAMK4,CD247,CD4,CHP1,CSNK1D,CSNK1E,CSNK1G2,FYN,GATA4,GNA11,GNA12,GNA14,GNAI1,GNAI2,GNAI3,GNAL,GNAO1,GNAQ,GNAS,GNB1,GNB1L,GNB5,GNG12,GNG2,GNG4,GNG7,GRB2,GSK3B,IKBKB,IKBKE,IKBKG,ITPR1,ITPR2,ITPR3,KRAS,LCK,LCP2,MAP2K1,MAP2K2,MAPK1,MEF2A,MEF2D,MRAS,NFAT5,NFATC1,NFATC2,NFATC3,NFKBID,ORAI1,PIK3C2A,PIK3C2B,PIK3CB,PIK3CD,PIK3R2,PIK3R3,PIK3R4,PIK3R5,PIK3R6,PLCB1,PLCB2,PLCB3,PLCG1,PLCG2,PPP3CB,PPP3CC,PRKCQ,RAF1,RALA,RALB,RAP1A,RAP1B,RCAN2,RCAN3,RRAS2,SOS1,SYK,XPO1,ZAP70 |

| | | | | |
|---|---|---|---|---|
| BMP signaling pathway | 2.75E+00 | 5.18E-01 | -5.466 | BMP1,BMP5,BMP6,BMP7,BMP8A,BMP8B,BMPR1A,BMPR1B,BMPR2,CAMK4,CREBBP,GRB2,KRAS,MAGED1,MAP2K1,MAP2K2,MAP2K4,MAPK1,MAPK10,MAPK13,MAPK14,MAPK8,MAPK9,MRAS,PRKACA,PRKAG2,PRKAR1B,PRKAR2A,RAF1,RALA,RALB,RAP1A,RAP1B,RRAS2,RUNX2,SMAD6,SMAD7,SMURF1,SOS1,SOSTDC1,TAB1,XIAP,ZNF423 |
| Trehalose Degradation II (Trehalase) | 2.69E+00 | 1.00E+00 | -2.236 | GCK,HK1,HK2,HK3,HKDC1,TREH |
| GPCR-Mediated Integration of Enteroendocrine Signaling Exemplified by an L Cell | 2.68E+00 | 5.28E-01 | -0.649 | ADCY1,ADCY10,ADCY2,ADCY3,ADCY5,ADCY6,ADCY7,ADCY8,ADCY9,GIP,GLP1R,GLP2R,GNA11,GNA14,GNAI1,GNAI2,GNAI3,GNAQ,GNAS,ITPR1,ITPR2,ITPR3,PLCB1,PLCB2,PLCB3,PLCD3,PLCE1,PLCG1,PLCG2,PLCH1,PLCH2,PLCL1,PLCL2,PRKACA,PRKAG2,PRKAR1B,PRKAR2A,PYY |
| D-myo-inositol (1,4,5,6)-Tetrakisphosphate Biosynthesis | 2.65E+00 | 4.78E-01 | -8 | ACP6,ALPL,CDC25A,CDC25C,CILP,DOT1L,DUSP10,DUSP14,DUSP15,DUSP16,DUSP27,EPHX2,EYA4,HACD2,INPP5B,INPP5F,IPMK,ITPK1,MTMR4,MTMR7,PALD1,PAWR,PLPP7,PPFIA2,PPFIA3,PPFIA4,PPFIBP2,PPIP5K1,PPM1H,PPP1R12A,PPP1R12C,PPP1R13B,PPP1R16B,PPP1R1C,PPP1R7,PPP1R8,PPP2R3A,PPP2R5B,PPP2R5D,PPP2R5E,PPP4C,PPP5C,PPTC7,PRUNE1,PTEN,PTPA,PTPN1,PTPN11,PTPN12,PTPN2,PTPN20,PTPN22,PTPN23,PTPN7,PTPRC,PTPRF,PTPRH,PTPRJ,PTPRM,PXYLP1,RASA1, |
| D-myo-inositol (3,4,5,6)-tetrakisphosphate Biosynthesis | 2.65E+00 | 4.78E-01 | -8 | ACP6,ALPL,CDC25A,CDC25C,CILP,DOT1L,DUSP10,DUSP14,DUSP15,DUSP16,DUSP27,EPHX2,EYA4,HACD2,INPP5B,INPP5F,IPMK,ITPK1,MTMR4,MTMR7,PALD1,PAWR,PLPP7,PPFIA2,PPFIA3,PPFIA4,PPFIBP2,PPIP5K1,PPM1H,PPP1R12A,PPP1R12C,PPP1R13B,PPP1R16B,PPP1R1C,PPP1R7,PPP1R8,PPP2R3A,PPP2R5B,PPP2R5D,PPP2R5E,PPP4C,PPP5C,PPTC7,PRUNE1,PTEN,PTPA,PTPN1,PTPN11,PTPN12,PTPN2,PTPN20,PTPN22,PTPN23,PTPN7,PTPRC,PTPRF,PTPRH,PTPRJ,PTPRM,PXYLP1,RASA1, |
| Basal Cell Carcinoma Signaling | 2.64E+00 | 5.29E-01 | -3.272 | APC,AXIN1,BMP1,BMP5,BMP6,BMP7,BMP8A,BMP8B,FZD3,GLI2,GLI3,GLIS1,GSK3B,HNF1A,KIF7,PTCH1,PTCH2,SMO,SUFU,TCF3,TCF4,TCF7,TCF7L1,TCF7L2,TP53,WNT10A,WNT11,WNT2B,WNT3,WNT3A,WNT4,WNT5B,WNT7A,WNT7B,WNT9A,WNT9B,ZNF875 |
| cAMP-mediated signaling | 2.62E+00 | 4.49E-01 | -7.155 | ADCY1,ADCY10,ADCY2,ADCY3,ADCY5,ADCY6,ADCY7,ADCY8,ADCY9,ADORA1,ADORA2B,ADORA3,AKAP1,AKAP10,AKAP12,AKAP13,AKAP8,AKAP9,BRAF,CAMK1D,CAMK2A,CAMK2B,CAMK2G,CAMK4,CHRM3,CNGA3,CNGB1,CNR2,CREB3L3,CREB5,CREBBP,CRHR1,DRD2,DRD3,DUSP4,ENPP6,GABBR2,GDPD1,GLP1R,GNAI1,GNAI2,GNAI3,GNAL,GNAO1,GNAS,GRK2,GRK4,GRM3,GRM4,GRM7,GRM8,HRH2,HTR4,HTR7,MAP2K1,MAP2K2,MAPK1,MPPED2,NPR3,OPRD1,OPRL1,PDE10A,PDE11A,PDE1A,PDE1B,PDE1C,PDE2A,PDE3A,PDE3B,PDE4A,PDE4B,PDE4C,PDE4D,PDE6A,PDE6B,PDE6C,PDE6G,PDE8A,PDE8B,PDE9A,PKIG,PPP3CB,PPP3CC,PRKACA,PRKAR1B,PRKAR2A,PTGER3,PTH1R,RAF1,RAP1A,RAP1GAP,RAPGEF3,RAPGEF4,RGS10,RGS12,RGS14,RGS7,RPS6KA1,SRC,STAT3,VIPR2 |
| IL-4 Signaling | 2.58E+00 | 5.12E-01 | NaN | AKT2,AKT3,FCER2,GRB2,IL4R,INPP5B,INPP5D,INPP5F,INPP5K,INPPL1,IRF4,JAK1,JAK2,JAK3,KRAS,MRAS,MTOR,NFAT5,NFATC1,NFATC2,NFATC3,NR3C1,NR3C2,PIK3C2A,PIK3C2B,PIK3CB,PIK3CD,PIK3R2,PIK3R3,PIK3R4,PIK3R5,PIK3R6,RALA,RALB,RAP1A,RAP1B,RPS6KB1,RRAS2,SOS1,STAT6,SYN |
| NF-κB Signaling | 2.57E+00 | 4.60E-01 | -8.333 | AKT2,AKT3,BMPR1A,BMPR1B,BMPR2,BRAF,BTRC,CARD10,CARD11,CASP8,CREBBP,CSNK2A1,CSNK2A2,EGFR,EP300,FGFR1,FGFR2,FGFR4,FLT1,FLT4,GHR,GSK3B,HDAC1,IGF1R,IGF2R,IKBKB,IKBKG,IL1R1,INSR,KRAS,LCK,MAP2K6,MAP3K1,MAP3K14,MAP3K3,MAP4K4,MAPK8,MRAS,NFKBID,NTRK1,NTRK2,NTRK3,PDGFRB,PIK3C2A,PIK3C2B,PIK3CB,PIK3CD,PIK3R2,PIK3R3,PIK3R4,PIK3R5,PIK3R6,PLCG2,PRKACA,PRKCB,PRKCQ,PRKCZ,RAF1,RALA,RALB,RAP1A,RAP1B,RELB,RIPK1,RRAS2,TAB1,TAB2,TAB3,TBK1,TGFA,TGFBR2,TGFBR3,TLR5,TNFRSF11A,TNFRSF1B,TNIP1,TRAF2,TRAF3,TRAF5 |
| Heparan Sulfate Biosynthesis | 2.56E+00 | 5.30E-01 | -5.916 | AARS1,B3GAT1,B4GALT7,CES3,CHST1,CHST10,CHST11,CHST12,CHST15,CHST3,CHST7,EXT1,EXT2,EXTL1,EXTL3,GLCE,HS2ST1,HS3ST2,HS3ST3A1,HS3ST3B1,HS3ST4,HS6ST1,HS6ST2,HS6ST3,NDST1,PLA1A,PNPLA7,PPME1,SULT1A1,SULT2B1,SULT4A1,TARS2,UST,XYLT1,XYLT2 |
| Natural Killer Cell Signaling | 2.55E+00 | 4.55E-01 | -7.826 | AKT2,AKT3,CD244,CD247,CDC42,COL18A1,COL1A1,COL2A1,COL5A3,FYN,GRB2,HSPA4,IL12RB1,IL12RB2,IL2RB,ITGAL,JAK2,JAK3,KRAS,LAIR1,LCK,LCP2,LILRB1,LIMK1,LIMK2,MAP2K1,MAP2K2,MAP2K3,MAP3K1,MAP3K10,MAP3K11,MAP3K12,MAP3K13,MAP3K14,MAP3K15,MAP3K2,MAP3K3,MAP3K5,MAP3K6,MAPK1,MAPK13,MAPK14,MRAS,MTOR,NCK1,NCR2,NECTIN2,NFAT5,NFATC1,NFATC2,NFATC3,PAK1,PAK2,PAK4,PIK3C2A,PIK3C2B,PIK3CB,PIK3CD,PIK3R2,PIK3R3,PIK3R4,PIK3R5,PIK3R6,PLCG1,PLCG2,PRKCQ,PTK2B,PTPN11,PXN,RAC1,RAC2,RAF1,RALA,RALB,RAP1A,RAP1B,RASSF5,ROCK1,RRAS2,SOS1,SYK,TYK2,VAV1,VAV2,VAV3,WAS,ZAP70 |
| Cyclins and Cell Cycle Regulation | 2.54E+00 | 5.12E-01 | 1.512 | ABL1,ATM,ATR,BTRC,CCNB3,CCND3,CDC25A,CDK6,CDK7,CUL1,E2F1,E2F2,E2F3,E2F4,GSK3B,HDAC1,HDAC11,HDAC4,HDAC5,HDAC6,HDAC7,HDAC9,MYT1,PPM1L,PPP2CB,PPP2R1B,PPP2R2A,PPP2R2B,PPP2R2C,PPP2R3A,PPP2R5B,PPP2R5C,PPP2R5D,PPP2R5E,PTPA,RAF1,SIN3A,TFDP1,TGFB1,T |
| Gαs Signaling | 2.53E+00 | 4.91E-01 | -5.898 | ADCY1,ADCY10,ADCY2,ADCY3,ADCY5,ADCY6,ADCY7,ADCY8,ADCY9,ADD1,ADD2,ADD3,ADORA2B,BRAF,CHRM3,CNGA3,CNGB1,CREB3L3,CREB5,CREBBP,CRHR1,GLP1R,GNAS,GNB1,GNB1L,GNB5,GNG12,GNG2,GNG4,GNG7,HCK,HRH2,HTR4,HTR7,MAP2K1,MAP2K2,MAPK1,MRAS,PRKACA,PRKAG2,PRKAR1B,PRKAR2A,PTH1R,RAP1A,RAPGEF2,RAPGEF3,RAPGEF4,RYR1,RYR2,RYR3,SRC,VIPR2 |
| Growth Hormone Signaling | 2.50E+00 | 5.21E-01 | -5.333 | GHR,IGF1R,JAK2,MAPK1,PDPK1,PIK3C2A,PIK3C2B,PIK3CB,PIK3CD,PIK3R2,PIK3R3,PIK3R4,PIK3R5,PIK3R6,PLCG1,PLCG2,PRKCA,PRKCB,PRKCD,PRKCE,PRKCG,PRKCH,PRKCI,PRKCQ,PRKCZ,PRKD1,RPS6KA1,RPS6KA2,RPS6KA3,RPS6KA5,RPS6KB1,RPS6KC1,SOCS5,SOCS7,STAT3,STAT5A,STAT5B |
| BEX2 Signaling Pathway | 2.49E+00 | 5.13E-01 | -0.48 | AKT2,AKT3,BCL2,ERBB2,GSK3B,HNF1A,ITGB6,MAP2,MAP2K4,MAPK10,MAPK8,MAPK9,MMP2,MTOR,PLXNC1,PPM1L,PPP2CB,PPP2R1B,PPP2R2A,PPP2R2B,PPP2R2C,PPP2R3A,PPP2R5B,PPP2R5C,PPP2R5D,PPP2R5E,PTEN,PTPA,SMARCB1,STAT3,SWAP70,TCF3,TCF4,TCF7,TCF7L1,TCF7L2,TSC1,TSC |
| mTOR Signaling | 2.47E+00 | 4.50E-01 | -4.331 | AKT2,AKT3,ATG13,CDC42,DGKZ,EIF3A,EIF3B,EIF3H,EIF3K,EIF3L,EIF4G3,FNBP1,GNB1L,GPLD1,INSR,KRAS,MAPK1,MAPKAP1,MRAS,MTOR,PDPK1,PIK3C2A,PIK3C2B,PIK3CB,PIK3CD,PIK3R2,PIK3R3,PIK3R4,PIK3R5,PIK3R6,PLD1,PLD2,PLD3,PLD5,PPM1L,PPP2CB,PPP2R1B,PPP2R2A,PPP2R2B,PPP2R2C,PPP2R3A,PPP2R5B,PPP2R5C,PPP2R5D,PPP2R5E,PRKAA1,PRKAA2,PRKAG2,PRKCA,PRKCB,PRKCD,PRKCE,PRKCG,PRKCH,PRKCI,PRKCQ,PRKCZ,PRKD1,PRR5,PRR5L,PTPA,RAC1,RAC2,RALA,RALB,RAP1A,RAP1B,RHEB,RHOA,RHOBTB1,RHOD,RHOQ,RHOT1,RICTOR,RPS19,RPS24,RPS3,RPS6KA1,RPS6KA2,RPS6KA3,RPS6KA5,RPS6KB1,RPS6KC1,RPS9,RPTOR,RRAS2,TSC1,TSC2,ULK1,VEGFA, |
| Lymphotoxin β Receptor Signaling | 2.47E+00 | 5.47E-01 | -4.796 | AKT2,AKT3,APAF1,BCL2L1,BIRC2,CASP9,CREBBP,EP300,IKBKB,IKBKE,IKBKG,MAP3K14,MAPK1,NFKBID,PDPK1,PIK3C2A,PIK3C2B,PIK3CB,PIK3CD,PIK3R2,PIK3R3,PIK3R4,PIK3R5,PIK3R6,RELB,TRAF1,TRAF2,TRAF3,TRAF5 |
| Phagosome Formation | 2.44E+00 | 4.82E-01 | NaN | C5AR1,CDC42,CR1,FCAMR,FCAR,FCER2,FNBP1,ITGA2,ITGA3,ITGAM,ITGAX,ITGB2,MARCO,MRC2,MSR1,PIK3C2A,PIK3C2B,PIK3CB,PIK3CD,PIK3R2,PIK3R3,PIK3R4,PIK3R5,PIK3R6,PLCB1,PLCB2,PLCB3,PLCD3,PLCE1,PLCG1,PLCG2,PLCH1,PLCH2,PLCL1,PLCL2,PRKCA,PRKCB,PRKCD,PRKCE,PRKCG,PRKCH,PRKCI,PRKCQ,PRKCZ,PRKD1,RAC1,RAC2,RHOA,RHOBTB1,RHOD,RHOQ,RHOT1,SCARA3,SY |
| Tryptophan Degradation X (Mammalian, via Tryptamine) | 2.44E+00 | 6.84E-01 | -3.606 | AKR1A1,ALDH1A1,ALDH1A2,ALDH2,ALDH3A1,ALDH3A2,ALDH3B1,ALDH7A1,DDC,IL4I1,MAOA,MAOB,SMOX |

| | | | | |
|---|---|---|---|---|
| Role of JAK1 and JAK3 in γc Cytokine Signaling | 2.42E+00 | 5.22E-01 | NaN | BLNK,GRB2,IL15RA,IL21R,IL2RA,IL2RB,IL4R,IRS2,JAK1,JAK2,JAK3,KRAS,MAPK1,MRAS,PIK3C2A,PIK3C2B,PIK3CB,PIK3CD,PIK3R2,PIK3R3,PIK3R4,PIK3R5,PIK3R6,PTK2B,PTPN11,RALA,RALB,RAP1A,RAP1B,RRAS2,STAT3,STAT5A,STAT5B,STAT6,SYK |
| Regulation of IL-2 Expression in Activated and Anergic T Lymphocytes | 2.42E+00 | 5.00E-01 | NaN | CAMK4,CARD11,CD247,CHP1,FYN,GRB2,IKBKB,IKBKE,IKBKG,KRAS,MAP2K1,MAP2K2,MAP2K4,MAP3K1,MAPK1,MAPK10,MAPK8,MAPK9,MRAS,NFAT5,NFATC1,NFATC2,NFATC3,NFKBID,PLCG1,PLCG2,PPP3CB,PPP3CC,RAC1,RAF1,RALA,RALB,RAP1A,RAP1B,RRAS2,SMAD3,SOS1,TGFB1,TGFB2,TGFBR2,VAV1,VAV2,VAV3,ZAP70 |
| Adipogenesis pathway | 2.34E+00 | 4.70E-01 | NaN | AGPAT2,ARNTL,ATG5,ATG7,BMP7,BMPR1A,BMPR2,CDK7,CTBP2,DGKD,EBF1,EGR2,ERCC2,ERCC3,EZH2,FGF1,FGFR1,FGFR2,FGFR4,FOXO1,FZD3,GTF2H1,GTF2H3,HDAC1,HDAC11,HDAC4,HDAC5,HDAC6,HDAC7,HDAC9,KAT2B,KAT6B,KAT7,KMT2B,LPIN1,LPL,MNAT1,NR1D2,PAXIP1,PER2,PPARG,PIP5K1,RBBP4,RPS6KA1,RPS6KB1,RPS6KC1,SAP130,SENP2,SETDB1,SIN3A,SIN3B,SIRT1,SIRT2,SMAD3,SMO,SREBF1,STAT5B,TBL1XR1,TCF7,TGFB1,TP53,ZNF423 |
| Sumoylation Pathway | 2.29E+00 | 4.85E-01 | 0.845 | CDC42,CDH1,CREBBP,CTBP2,DNMT3A,EP300,ETS1,FNBP1,HDAC1,ISG20,MAP2K4,MAP3K5,MAPK10,MAPK8,MAPK9,MDM2,NR3C1,PIAS1,PIAS4,PML,RAC1,RAC2,RANBP2,RANGAP1,RCC1,RCOR1,RFC1,RFC2,RHOA,RHOBTB1,RHOD,RHOQ,RHOT1,RNF4,RPA1,SAE1,SENP1,SENP2,SENP5,SENP7,SIRT1,SLC19A1,SP1,TDG,TP53,UBA2,UBE2I,XIAP,ZNF217 |
| Inhibition of Angiogenesis by TSP1 | 2.29E+00 | 5.94E-01 | -4 | AKT2,AKT3,CD36,FYN,GUCY1A2,HSPG2,MAP2K4,MAPK1,MAPK10,MAPK13,MAPK14,MAPK8,MAPK9,NOS3,SDC1,TGFB1,TGFBR2,TP53,VEGFA |
| Hepatic Fibrosis / Hepatic Stellate Cell Activation | 2.29E+00 | 4.51E-01 | NaN | BCL2,CCR7,CERT1,COL11A1,COL13A1,COL15A1,COL16A1,COL17A1,COL18A1,COL1A1,COL20A1,COL21A1,COL22A1,COL23A1,COL24A1,COL25A1,COL26A1,COL27A1,COL28A1,COL2A1,COL4A1,COL4A2,COL4A4,COL4A6,COL5A1,COL5A3,COL6A3,COL6A5,COL6A6,COL7A1,COL8A1,COL8A2,COL9A2,COL9A3,CSF1,ECE1,EGFR,FGF1,FGFR1,FGFR2,FLT1,FLT4,HGF,IFNGR2,IGF1R,IGFBP4,IL1R1,IL1RAP,IL1RAPL1,IL1RAPL2,IL4R,IL6R,LAMA1,LBP,LEPR,LHX2,MET,MMP2,MYH10,MYH11,MYH14,MYH3,MYH6,MYH7,MYH7B,MYH9,MYL3,PDGFA,PDGFB,PDGFD,PDGFRB,SERPINE1,SMAD3,SMAD7,TGFA,TGFB1,TGFB2,TGFBR2,TIMP2,TNFRSF1B,VEGFA,VEGFD |
| Chondroitin Sulfate Biosynthesis | 2.27E+00 | 5.38E-01 | -5.292 | B3GAT1,B4GALT7,CHST1,CHST10,CHST11,CHST12,CHST15,CHST3,CHST7,CHSY1,CHSY3,CSGALNACT1,CSGALNACT2,HS2ST1,HS3ST2,HS3ST3A1,HS3ST3B1,HS3ST4,HS6ST1,HS6ST2,HS6ST3,NDST1,SULT1A1,SULT2B1,SULT4A1,UST,XYLT1,XYLT2 |
| HIF1α Signaling | 2.26E+00 | 4.77E-01 | NaN | AKT2,AKT3,ARNT,CREBBP,EGLN1,EGLN3,EP300,HSP90AA1,KRAS,LDHA,LDHC,MAPK1,MAPK10,MAPK13,MAPK14,MAPK4,MAPK6,MAPK8,MAPK9,MDM2,MMP14,MMP15,MMP17,MMP2,MMP21,MMP26,MMP28,MRAS,NCOA1,NOS1,NOS2,NOS3,P4HTM,PIK3C2A,PIK3C2B,PIK3CB,PIK3CD,PIK3R2,PIK3R3,PIK3R4,PIK3R5,PIK3R6,RALA,RALB,RAP1A,RAP1B,RRAS2,SLC2A1,SLC2A14,SLC2A5,TP53,VEGFA,V |
| Heparan Sulfate Biosynthesis (Late Stages) | 2.25E+00 | 5.25E-01 | -5.568 | AARS1,CES3,CHST1,CHST10,CHST11,CHST12,CHST15,CHST3,CHST7,EXT1,EXT2,EXTL1,EXTL3,GLCE,HS2ST1,HS3ST2,HS3ST3A1,HS3ST3B1,HS3ST4,HS6ST1,HS6ST2,HS6ST3,NDST1,PLA1A,PNPLA7,PPME1,SULT1A1,SULT2B1,SULT4A1,TARS2,UST |
| NAD Biosynthesis from 2-amino-3-carboxymuconate | 2.24E+00 | 1.00E+00 | -2.236 | NADSYN1,NMNAT1,NMNAT2,NMNAT3,QPRT |
| Protein Citrullination | 2.24E+00 | 1.00E+00 | -2.236 | PADI1,PADI2,PADI3,PADI4,PADI6 |
| Amyloid Processing | 2.23E+00 | 5.40E-01 | -2.673 | AKT2,AKT3,APP,BACE1,BACE2,CAPN1,CAPN11,CAPN2,CAPN5,CAPN8,CSNK1D,CSNK1E,CSNK2A1,CSNK2A2,GSK3B,MAPK1,MAPK13,MAPK14,MAPT,MARK1,PRKACA,PRKAG2,PRKAR1B,PRKAR2A,PRKCE,PSEN1,PSEN2 |
| April Mediated Signaling | 2.20E+00 | 5.64E-01 | -4.264 | IKBKB,IKBKE,IKBKG,MAP2K4,MAP3K1,MAP3K14,MAPK1,MAPK10,MAPK13,MAPK14,MAPK8,MAPK9,NFAT5,NFATC1,NFATC2,NFATC3,NFKBID,TNFRSF13B,TRAF1,TRAF2,TRAF3,TRAF5 |
| IL-22 Signaling | 2.18E+00 | 6.25E-01 | -3.873 | AKT2,AKT3,IL22RA1,JAK1,MAP2K4,MAPK1,MAPK10,MAPK13,MAPK14,MAPK8,MAPK9,STAT3,STAT5A,STAT5B,TYK2 |
| Dermatan Sulfate Biosynthesis | 2.17E+00 | 5.27E-01 | -5.385 | B3GAT1,B4GALT7,CHST1,CHST10,CHST11,CHST12,CHST15,CHST3,CHST7,CHSY1,CHSY3,CSGALNACT1,CSGALNACT2,DSE,HS2ST1,HS3ST2,HS3ST3A1,HS3ST3B1,HS3ST4,HS6ST1,HS6ST2,HS6ST3,NDST1,SULT1A1,SULT2B1,SULT4A1,UST,XYLT1,XYLT2 |
| Ceramide Signaling | 2.15E+00 | 4.89E-01 | -2.402 | AKT2,AKT3,BCL2,CERK,CNKSR1,KRAS,KSR1,MAP2K1,MAP2K4,MAP3K1,MAPK8,MRAS,PIK3C2A,PIK3C2B,PIK3CB,PIK3CD,PIK3R2,PIK3R3,PIK3R4,PIK3R5,PIK3R6,PPM1L,PPP2CB,PPP2R1B,PPP2R2A,PPP2R2B,PPP2R2C,PPP2R3A,PPP2R5B,PPP2R5C,PPP2R5D,PPP2R5E,PRKCZ,PTPA,RAF1,RALA,RALB,RAP1A,RAP1B,RRAS2,SMPD3,SMPD4,TNFRSF1B |
| Granzyme B Signaling | 2.15E+00 | 6.88E-01 | -2.714 | APAF1,BID,CASP8,CASP9,DFFA,DFFB,LMNB1,LMNB2,NUMA1,PARP1,PRKDC |
| Mismatch Repair in Eukaryotes | 2.15E+00 | 6.88E-01 | NaN | EXO1,MLH1,MSH2,MSH3,MSH6,PMS2,POLD1,RFC1,RFC2,RPA1,SLC19A1 |
| 1D-myo-inositol Hexakisphosphate Biosynthesis | 2.14E+00 | 6.67E-01 | -3.464 | INPP5A,INPP5B,INPP5D,INPP5F,INPP5K,INPPL1,IPMK,ITPK1,ITPKB,ITPKC,SEC16A,SYNJ2 |
| D-myo-inositol (1,3,4)-trisphosphate Biosynthesis | 2.14E+00 | 6.67E-01 | -3.464 | INPP5A,INPP5B,INPP5D,INPP5F,INPP5K,INPPL1,IPMK,ITPKB,ITPKC,PTEN,SEC16A,SYNJ2 |
| Chondroitin Sulfate Biosynthesis (Late Stages) | 2.11E+00 | 5.45E-01 | -4.899 | CHST1,CHST10,CHST11,CHST12,CHST15,CHST3,CHST7,CHSY1,CHSY3,CSGALNACT1,CSGALNACT2,HS2ST1,HS3ST2,HS3ST3A1,HS3ST3B1,HS3ST4,HS6ST1,HS6ST2,HS6ST3,NDST1,SULT1A1,SULT2B1,SULT4A1,UST |
| Tight Junction Signaling | 2.10E+00 | 4.49E-01 | NaN | ACTB,ACTG2,AKT2,AKT3,ARHGEF2,CASK,CDC42,CGN,CLDN10,CLDN11,CLDN14,CLDN4,CNKSR3,CPSF3,CSTF3,CTNNA1,EPB41,F11R,GOSR1,HSF1,JAM3,LLGL1,MAGI2,MARK2,MPDZ,MYH10,MYH11,MYH14,MYH3,MYH6,MYH7,MYH7B,MYH9,MYL3,MYLK,NECTIN1,NECTIN2,NECTIN3,NSF,PATJ,PPM1L,PPP2CB,PPP2R1B,PPP2R2A,PPP2R2B,PPP2R2C,PPP2R3A,PPP2R5B,PPP2R5C,PPP2R5D,PPP2R5E,PRKACA,PRKAG2,PRKAR1B,PRKAR2A,PRKCI,PRKCZ,PTEN,PTPA,RAC1,RHOA,SAFB,SMURF1,SPTAN1,SYMPK,TGFB1,TGFB2,TGFBR2,TIAM1,TJP1,TJP2,TJP3,TNFRSF1B,VCL,VTI1A |
| IL-17A Signaling in Airway Cells | 2.07E+00 | 5.08E-01 | -4.596 | AKT2,AKT3,GSK3B,IKBKB,IKBKE,IKBKG,IL17RA,IL17RC,JAK1,JAK2,JAK3,MAP2K1,MAP2K2,MAP2K4,MAPK1,MAPK10,MAPK13,MAPK14,MAPK8,MAPK9,NFKBID,PIK3C2A,PIK3C2B,PIK3CB,PIK3CD,PIK3R2,PIK3R3,PIK3R4,PIK3R5,PIK3R6,PTEN,STAT3,TYK2 |
| Prostate Cancer Signaling | 2.04E+00 | 4.83E-01 | NaN | ABL1,AKT2,AKT3,BCL2,CASP9,CREB3L3,CREB5,CREBBP,E2F1,FOXO1,GRB2,GSK3B,HSP90AA1,KRAS,MAP2K1,MAP2K2,MAPK1,MDM2,MRAS,MTOR,NFKBID,PDPK1,PIK3C2A,PIK3C2B,PIK3CB,PIK3CD,PIK3R2,PIK3R3,PIK3R4,PIK3R5,PIK3R6,PTEN,RAF1,RALA,RALB,RAP1A,RAP1B,RRAS2,SIN3A,SOS1, |
| Systemic Lupus Erythematosus In T Cell Signaling Pathway | 2.02E+00 | 4.35E-01 | -4.796 | AKT2,AKT3,CAMK4,CASP8,CASP9,CBL,CD247,CD44,CDC42,CHP1,CREB3L3,CREB5,CREBBP,DNMT1,ESR1,EZR,FNBP1,FOXP3,GNA12,GNAI1,GNAI2,GNAI3,GNB1L,GRB2,IRF4,ITGAL,ITPR1,KRAS,LEPR,MAP2K1,MAP2K2,MAP2K3,MAP2K4,MAP2K5,MAP2K6,MAPK1,MRAS,MSN,MTOR,NFATC2,NOS1,NOS2,NOS3,ORAI1,PIK3C2A,PIK3C2B,PIK3CB,PIK3CD,PIK3R2,PIK3R3,PIK3R4,PIK3R5,PIK3R6,PLCG1,PPM1L,PPP2CB,PPP2R1B,PPP2R2A,PPP2R2B,PPP2R2C,PPP2R3A,PPP2R5B,PPP2R5C,PPP2R5D,PPP2R5E,PPP3CB,PPP3CC,PTK2,PTPA,RAC1,RAC2,RALA,RALB,RAP1A,RAP1B,RDX,RHOA,RHOBTB1,RHOD,RHOQ,RHOT1,ROCK1,ROCK2,RORC,RPS6KB1,RPTOR,RRAS2,SOS1,SP1,STAT3,STIM1,SYK,VA |

| | | | | |
|---|---|---|---|---|
| Inositol Pyrophosphates Biosynthesis | 2.00E+00 | 8.57E-01 | -2.449 | *IP6K1,IP6K2,IP6K3,IPMK,PPIP5K1,PPIP5K2* |
| Iron homeostasis signaling pathway | 2.00E+00 | 4.58E-01 | NaN | *ABCB10,ACO2,ARNT,ATP6AP1L,ATP6V0A1,ATP6V0A2,ATP6V0A4,ATP6V0D1,ATP6V0E1,ATP6V1A,ATP6V1C2,ATP6V1E1,ATP6V1E2,BMP1,BMP5,BMP6,BMP7,BMP8A,BMP8B,BMPR1A,BMPR1B,BMPR2,BTRC,CIAO2A,CP,CREB3L3,CUL1,EGFR,EPAS1,FLVCR1,FXN,GDF15,HMOX2,IL6R,IREB2,JAK1,JAK2,JAK3,LRP1,LYRM4,MAPK1,MMS19,NFS1,PDGFA,PDGFB,PDGFRB,SLC11A1,SLC11A2,SLC25A37,SLC39A14,SLC48A1,SMAD3,STAT3,STAT5A,STAT5B,STEAP3,TFR2,TFRC,TMPRSS6,TYK2* |
| IL-7 Signaling Pathway | 1.96E+00 | 4.93E-01 | -5.303 | *AKT2,AKT3,BCL2,CCND3,CDC25A,EBF1,FOXO1,FOXO3,FOXO6,FYN,GRB2,GSK3B,HGF,JAK1,JAK3,MAPK1,MAPK13,MAPK14,MET,NFATC1,PAX5,PDPK1,PIK3C2A,PIK3C2B,PIK3CB,PIK3CD,PIK3R2,PIK3R3,PIK3R4,PIK3R5,PIK3R6,PTK2,SLC2A1,SOS1,STAT5A,STAT5B* |
| Superpathway of D-myo-inositol (1,4,5)-trisphosphate | 1.93E+00 | 6.09E-01 | -3.742 | *BPNT1,IMPA2,INPP5A,INPP5B,INPP5D,INPP5F,INPP5K,INPPL1,IPMK,ITPKB,ITPKC,PTEN,SEC16A,SYNJ2* |
| Salvage Pathways of Pyrimidine Ribonucleotides | 1.92E+00 | 4.74E-01 | -6.708 | *AK4,AK5,AK7,AK8,AK9,AKT2,APOBEC3G,BRAF,CDA,CDK18,CDK6,CDK7,CMPK1,CSNK1D,DAPK1,FAM20B,G6PC,GRK4,GRK5,GRK6,LIMK1,LIMK2,MAP2K1,MAP2K2,MAP2K3,MAP2K4,MAP2K6,MAP3K6,MAPK1,MAPK6,MAPK8,MAPK9,NME7,PAK1,PAK2,PKN1,PRKAA1,PRKAA2,PRKCD,PRKCE,PRKCH,PRK* |
| Oncostatin M Signaling | 1.91E+00 | 5.35E-01 | -4.69 | *EPAS1,GRB2,IL6ST,JAK1,JAK2,JAK3,KRAS,MAP2K1,MAP2K2,MAPK1,MRAS,OSMR,RAF1,RALA,RALB,RAP1A,RAP1B,RRAS2,SOS1,STAT3,STAT5A,STAT5B,TYK2* |
| tRNA Splicing | 1.91E+00 | 5.35E-01 | -4.69 | *ENPP6,GDPD1,MPPED2,PDE10A,PDE11A,PDE1A,PDE1B,PDE1C,PDE2A,PDE3A,PDE3B,PDE4A,PDE4B,PDE4C,PDE4D,PDE6A,PDE6B,PDE6C,PDE6G,PDE8A,PDE8B,PDE9A,TSEN2* |
| Leucine Degradation I | 1.91E+00 | 7.78E-01 | -2.646 | *ACADM,BCAT1,HMGCL,HMGCLL1,IVD,MCCC1,MCCC2* |
| Gαi Signaling | 1.88E+00 | 4.56E-01 | -1.941 | *ADCY1,ADCY10,ADCY2,ADCY3,ADCY5,ADCY6,ADCY7,ADCY8,ADCY9,ADORA1,ADORA3,CNR2,DRD2,DRD3,GABBR2,GNAI1,GNAI2,GNAI3,GNAS,GNB1,GNB1L,GNB5,GNG12,GNG2,GNG4,GNG7,GRB2,GRM3,GRM4,GRM7,GRM8,KRAS,MAPK1,MRAS,NPR3,OPRD1,OPRL1,PRKACA,PRKAG2,PRKAR1B,PRKAR2A,PTGER3,RAF1,RALA,RALB,RALGDS,RAP1A,RAP1B,RAP1GAP,RGS10,RGS12,RGS14,RGS7,R* |
| IL-6 Signaling | 1.88E+00 | 4.56E-01 | -6.949 | *ABCB1,AKT2,AKT3,COL1A1,CSNK2A1,CSNK2A2,GRB2,IKBKB,IKBKE,IKBKG,IL1R1,IL1RAP,IL1RAPL1,IL1RAPL2,IL6R,IL6ST,JAK2,KRAS,LBP,MAP2K1,MAP2K2,MAP2K3,MAP2K4,MAP2K6,MAP3K14,MAP4K4,MAPK1,MAPK10,MAPK13,MAPK14,MAPK8,MAPK9,MAPKAPK2,MRAS,NFKBID,PIK3C2A,PIK3C2B,PIK3CB,PIK3CD,PIK3R2,PIK3R3,PIK3R4,PIK3R5,PIK3R6,PTPN11,RAF1,RALA,RALB,RAP1A,RAP1B,RRAS2,SOS1,STAT3,TAB1,TNFRSF1B,TRAF2,VEGFA* |
| B Cell Activating Factor Signaling | 1.87E+00 | 5.37E-01 | -4.583 | *IKBKB,IKBKE,IKBKG,MAP2K4,MAP3K1,MAP3K14,MAPK1,MAPK10,MAPK13,MAPK14,MAPK8,MAPK9,NFAT5,NFATC1,NFATC2,NFATC3,NFKBID,TNFRSF13B,TRAF1,TRAF2,TRAF3,TRAF5* |
| Role of JAK2 in Hormone-like Cytokine Signaling | 1.87E+00 | 5.62E-01 | NaN | *GHR,HLTF,IRS2,JAK1,JAK2,PRLR,PTPN1,PTPN11,SH2B1,SH2B2,SH2B3,SIRPA,SOCS5,SOCS7,STAT3,STAT5A,STAT5B,TYK2* |
| NAD biosynthesis II (from tryptophan) | 1.86E+00 | 7.27E-01 | -2.828 | *AFMID,HAAO,KMO,NADSYN1,NMNAT1,NMNAT2,NMNAT3,QPRT* |
| Putrescine Degradation III | 1.86E+00 | 6.47E-01 | -3.317 | *ALDH1A1,ALDH1A2,ALDH2,ALDH3A1,ALDH3A2,ALDH3B1,ALDH7A1,IL4I1,MAOA,MAOB,SMOX* |
| Oxidative Ethanol Degradation | 1.85E+00 | 6.67E-01 | -3.162 | *ACSS1,ACSS2,ACSS3,ALDH1A1,ALDH1A2,ALDH2,ALDH3A1,ALDH3A2,ALDH3B1,ALDH7A1* |
| iCOS-iCOSL Signaling in T Helper Cells | 1.81E+00 | 4.62E-01 | -5.466 | *AKT2,AKT3,CAMK2A,CAMK2B,CAMK2G,CAMK4,CD247,CD4,CHP1,CSK,GAB2,GRB2,IKBKB,IKBKE,IKBKG,IL2RA,IL2RB,INPP5D,ITPR1,ITPR2,ITPR3,LCK,LCP2,NFAT5,NFATC1,NFATC2,NFATC3,NFKBID,PDPK1,PIK3C2A,PIK3C2B,PIK3CB,PIK3CD,PIK3R2,PIK3R3,PIK3R4,PIK3R5,PIK3R6,PLCG1,PLEKHA2,PLEKHA4,PPP3CB,PPP3CC,PRKCQ,PTEN,PTPRC,RAC1,VAV1,ZAP70* |
| Melatonin Degradation II | 1.79E+00 | 1.00E+00 | -2 | *IL4I1,MAOA,MAOB,SMOX* |
| Cancer Drug Resistance By Drug Efflux | 1.77E+00 | 5.00E-01 | NaN | *ABCB1,ABCC1,ABCC10,ABCC2,ABCG2,AKT2,AKT3,BRAF,FOXO1,FOXO3,FOXO6,KRAS,MAP2K1,MAP2K2,MAPK1,MDM2,MRAS,PIK3CB,PIK3CD,PIK3R2,PIK3R3,PTEN,RAF1,RALA,RALB,RAP1A,RAP1B,RR* |
| Actin Nucleation by ARP-WASP Complex | 1.76E+00 | 4.86E-01 | -4.271 | *ACTR2,ACTR3,ARPC1A,ARPC5L,BAIAP2,CDC42,FNBP1,GNA12,GRB2,ITGA2,ITGA3,KRAS,MRAS,NCK1,NCK2,PPP1R12A,PPP1R12B,PPP1R12C,RAC1,RAC2,RALA,RALB,RAP1A,RAP1B,RHOA,RHOBTB1,RHOD,RHOQ,RHOT1,ROCK1,ROCK2,RRAS2,SOS1,WAS* |
| Wnt/Ca+ pathway | 1.70E+00 | 4.92E-01 | -5.477 | *AXIN1,CAMK2A,CREB3L3,CREB5,CREBBP,FZD3,GSK3B,NFAT5,NFATC1,NFATC2,NFATC3,PDE6A,PDE6B,PDE6C,PDE6G,PLCB1,PLCB2,PLCB3,PLCD3,PLCE1,PLCG1,PLCG2,PLCH1,PLCH2,PLCL1,PLCL2,PRKCA,ROR1,SMO,WNT5B* |
| Remodeling of Epithelial Adherens Junctions | 1.68E+00 | 4.85E-01 | -3.606 | *ACTB,ACTG2,ACTN1,ACTN2,ACTN4,ACTR2,ACTR3,APC,ARPC1A,ARPC5L,CDH1,CLIP1,CTNNA1,CTNNA2,DNM1,DNM1L,DNM2,DNM3,EXOC2,HGF,HGS,IQGAP1,MAPRE1,MAPRE2,MET,RAB5B,RAB7A,RALA,SRC,TUBA1C,TUBB3,VCL* |
| Xenobiotic Metabolism CAR Signaling Pathway | 1.65E+00 | 4.33E-01 | -4.185 | *ABCB1,ABCC1,ABCC2,ABCC3,ALDH1A1,ALDH1A2,ALDH1L1,ALDH1L2,ALDH2,ALDH3A1,ALDH3A2,ALDH3B1,ALDH7A1,CHST1,CHST10,CHST11,CHST12,CHST15,CHST3,CHST7,CREBBP,CYP2C19,DNAJC7,EGFR,EP300,GRIP1,HS2ST1,HS3ST2,HS3ST3A1,HS3ST3B1,HS3ST4,HS6ST1,HS6ST2,HS6ST3,HSP90AA1,MAP2K1,MAP2K2,MAP2K3,MAP2K4,MAP2K5,MAP2K6,MAPK1,MED1,NCOA1,NDST1,NOS2,PPM1L,PPP2CB,PPP2R1B,PPP2R2A,PPP2R2B,PPP2R2C,PPP2R3A,PPP2R5B,PPP2R5C,PPP2R5D,PPP2R5E,PRKCA,PRKCB,PRKCD,PRKCE,PRKCG,PRKCH,PRKCI,PRKCQ,PRKCZ,PRKD1,PTPA,RXRA,SR* |
| tRNA Charging | 1.63E+00 | 5.26E-01 | -4.472 | *AARS1,CARS1,CARS2,DARS2,EARS2,EPRS1,FARS2,HARS1,IARS1,IARS2,LARS1,LARS2,NARS2,RARS1,SARS1,TARS1,TARS2,TARS3,WARS2,YARS1* |
| Regulation of Cellular Mechanics by Calpain Protease | 1.63E+00 | 4.84E-01 | -4.025 | *ACTN1,ACTN2,ACTN4,CAPN1,CAPN11,CAPN2,CAPN5,CAPN8,CAST,CDK6,CNGA3,CNGB1,EGFR,EZR,GRB2,ITGA2,ITGA3,KRAS,MAPK1,MRAS,PTK2,PXN,RALA,RALB,RAP1A,RAP1B,RRAS2,SRC,TLN1,T* |
| Cdc42 Signaling | 1.57E+00 | 4.42E-01 | -5.689 | *ACTR2,ACTR3,APC,ARPC1A,ARPC5L,BAIAP2,CD247,CDC42,CDC42BPA,CLIP1,DIAPH1,DIAPH3,EXOC2,EXOC3,EXOC4,EXOC6,EXOC7,FGD1,FGD3,FNBP1L,GSK3B,IQGAP1,IQGAP2,IQGAP3,ITGA2,ITGA3,LIMK1,LIMK2,LLGL1,MAP2K4,MAP3K11,MAPK1,MAPK10,MAPK13,MAPK14,MAPK8,MAPK9,MPRIP,MYL10,MYL3,MYLK,PAK1,PAK2,PAK4,PARD3,PPP1R12A,PPP1R12B,PRKCI,PRKCZ,RAF1,RALA,RASA1,* |
| DNA Double-Strand Break Repair by Non-Homologous | 1.56E+00 | 6.43E-01 | NaN | *ATM,LIG3,MRE11,PARP1,PRKDC,RAD50,WRN,XRCC4,XRCC6* |
| Phenylalanine Degradation IV (Mammalian, via Side Chain) | 1.56E+00 | 6.43E-01 | -3 | *ACSF3,ALDH2,ALDH3A2,GOT2,HPD,IL4I1,MAOA,MAOB,SMOX* |
| TNFR1 Signaling | 1.55E+00 | 5.00E-01 | -3.674 | *APAF1,BID,BIRC2,CASP8,CASP9,CDC42,CRADD,IKBKB,IKBKE,IKBKG,MADD,MAP2K4,MAP3K1,MAP3K14,MAP4K2,MAPK8,NAIP,NFKBID,PAK1,PAK2,PAK4,RIPK1,TRAF2,XIAP* |
| Autophagy | 1.52E+00 | 4.91E-01 | NaN | *ATG10,ATG13,ATG16L1,ATG3,ATG4B,ATG4C,ATG5,ATG7,BCL2,CTSB,CTSH,CTSS,LAMP1,MAP1LC3A,MAP1LC3B2,MTOR,NBR1,PIK3R4,SQSTM1,ULK1,VPS11,VPS18,VPS33B,VPS39,VPS41,WDFY3* |
| Stearate Biosynthesis I (Animals) | 1.50E+00 | 5.00E-01 | -4.583 | *ACOT11,ACOT7,ACOT8,ACOT9,ACSBG1,ACSL4,ACSL5,ACSL6,DBT,DHCR24,ELOVL2,ELOVL6,GNPAT,HNF4A,MBOAT7,PORCN,SLC27A1,SLC27A4,SLC27A5,SLC27A6,TBXAS1,TECR,THEM4* |
| 4-1BB Signaling in T Lymphocytes | 1.50E+00 | 5.31E-01 | -4 | *IKBKB,IKBKE,IKBKG,MAP2K1,MAP2K2,MAP2K4,MAP3K14,MAP3K5,MAPK1,MAPK10,MAPK13,MAPK14,MAPK8,MAPK9,NFKBID,TRAF1,TRAF2* |

| | | | | |
|---|---|---|---|---|
| Xenobiotic Metabolism PXR Signaling Pathway | 1.50E+00 | 4.26E-01 | -4.734 | ABCB1,ABCC2,ABCC3,ALDH1A1,ALDH1A2,ALDH1L1,ALDH1L2,ALDH2,ALDH3A1,ALDH3A2,ALDH3B1,ALDH7A1,CAMK2A,CAMK2B,CAMK2G,CES1,CES3,CES4A,CES5A,CHST1,CHST10,CHST11,CHST12,CHST15,CHST3,CHST7,CREBBP,CYP2C19,DNAJC7,EP300,GRIP1,HS2ST1,HS3ST2,HS3ST3A1,HS3ST3B1,HS3ST4,HS6ST1,HS6ST2,HS6ST3,HSP90AA1,IL4I1,MAOA,MAOB,MED1,NCOA1,NCOR1,NCOR2,NDST1,NOS2,NR1I2,PPP1R12A,PPP1R14C,PPP1R14D,PPP1R3A,PPP1R7,PRKACA,PRKAG2,PRKAR1B,PRKAR2A,PRKCA,PRKCB,PRKCD,PRKCE,PRKCG,PRKCH,PRKCI,PRKCQ,PRKCZ,PRKD1,RXRA,SM |
| IL-17 Signaling | 1.50E+00 | 4.62E-01 | NaN | AKT2,AKT3,GSK3B,IL17RA,IL17RC,JAK1,JAK2,KRAS,MAP2K1,MAP2K2,MAP2K3,MAP2K4,MAP2K6,MAP3K14,MAPK1,MAPK10,MAPK13,MAPK14,MAPK8,MAPK9,MAPKAPK2,MRAS,NOS2,PIK3C2A,PIK3C2B,PIK3CB,PIK3CD,PIK3R2,PIK3R3,PIK3R4,PIK3R5,PIK3R6,RALA,RALB,RAP1A,RAP1B,RRAS2 |
| Triacylglycerol Biosynthesis | 1.49E+00 | 5.13E-01 | -4.359 | ABHD5,AGPAT2,AGPAT3,AGPAT4,AGPAT5,DBT,DGAT2,ELOVL2,ELOVL6,LCLAT1,LPIN1,LPIN2,LPIN3,MBOAT2,MBOAT7,PLPP1,PLPP3,PLPP4,PNPLA3,PORCN |
| Dopamine Degradation | 1.49E+00 | 5.65E-01 | -3.606 | ALDH1A1,ALDH1A2,ALDH2,ALDH3A1,ALDH3A2,ALDH3B1,ALDH7A1,COMT,IL4I1,MAOA,MAOB,SMOX,SULT1A1 |
| Estrogen-Dependent Breast Cancer Signaling | 1.46E+00 | 4.66E-01 | -5.385 | AKT2,AKT3,CREB3L3,CREB5,CREBBP,EGFR,ESR1,HSD17B12,HSD17B14,HSD17B3,HSD17B4,IGF1R,KRAS,MAPK1,MRAS,PIK3C2A,PIK3C2B,PIK3CB,PIK3CD,PIK3R2,PIK3R3,PIK3R4,PIK3R5,PIK3R6,RALA,RALB,RAP1A,RAP1B,RRAS2,SP1,SRC,STAT5A,STAT5B,TERT |
| T Cell Exhaustion Signaling Pathway | 1.46E+00 | 4.26E-01 | -2.889 | ACVR1B,ACVR2B,AKT2,AKT3,BMPR2,BTLA,FOXO1,FOXP1,HAVCR2,IL12RB1,IL12RB2,IL6R,IRF4,JAK1,JAK2,JAK3,KRAS,LGALS9,MAP2K4,MAPK1,MAPK10,MAPK8,MAPK9,MGAT5,MRAS,MTOR,NFAT5,NFATC1,NFATC2,NFATC3,PDCD1LG2,PIK3C2A,PIK3C2B,PIK3CB,PIK3CD,PIK3R2,PIK3R3,PIK3R4,PIK3R5,PIK3R6,PLCG1,PLCG2,PPM1L,PPP2CB,PPP2R1B,PPP2R2A,PPP2R2B,PPP2R2C,PPP2R3A,PPP2R5B,PPP2R5C,PPP2R5D,PPP2R5E,PRKCQ,PTPA,PTPN11,RALA,RALB,RAP1A,RAP1B,RRAS2,SMAD3,STAT2,STAT3,TBX21,TCF7,TGFB1,TGFBR2,TGFBR3,TYK2,VEGFA,ZAP70 |
| SPINK1 General Cancer Pathway | 1.44E+00 | 4.75E-01 | -5.385 | AKT2,AKT3,EGFR,IL6R,JAK1,JAK2,JAK3,KRAS,MAP2K1,MAP2K2,MAPK1,MRAS,PIK3C2A,PIK3C2B,PIK3CB,PIK3CD,PIK3R2,PIK3R3,PIK3R4,PIK3R5,PIK3R6,RAF1,RALA,RALB,RAP1A,RAP1B,RRAS2,STAT |
| Dermatan Sulfate Biosynthesis (Late Stages) | 1.41E+00 | 5.00E-01 | -4.583 | CHST1,CHST10,CHST11,CHST12,CHST15,CHST3,CHST7,DSE,HS2ST1,HS3ST2,HS3ST3A1,HS3ST3B1,HS3ST4,HS6ST1,HS6ST2,HS6ST3,NDST1,SULT1A1,SULT2B1,SULT4A1,UST |
| Serotonin Receptor Signaling | 1.41E+00 | 5.00E-01 | NaN | ADCY1,ADCY10,ADCY2,ADCY3,ADCY5,ADCY6,ADCY7,ADCY8,ADCY9,DDC,HTR2C,HTR3B,HTR4,HTR7,IL4I1,MAOA,MAOB,SLC18A1,SLC18A2,SLC6A4,SMOX |
| PPAR Signaling | 1.41E+00 | 4.46E-01 | 4.621 | CREBBP,EP300,GRB2,HSP90AA1,IKBKB,IKBKE,IKBKG,IL1R1,IL1RAP,IL1RAPL1,IL1RAPL2,INSR,KRAS,MAP2K1,MAP2K2,MAP3K14,MAP4K4,MAPK1,MED1,MRAS,NCOA1,NCOR1,NCOR2,NFKBID,NR1H3,PDGFA,PDGFB,PDGFD,PDGFRB,PPARA,PPARD,PPARG,RAF1,RALA,RALB,RAP1A,RAP1B,RRAS2,RXRA,SOS1,STAT5A,STAT5B,TAB1,TNFRSF1B,TRAF2 |
| NRF2-mediated Oxidative Stress Response | 1.37E+00 | 4.19E-01 | -6.272 | ABCC1,ABCC2,ABCC4,ACTB,ACTG2,AKR1A1,BACH1,CREBBP,CUL3,DNAJA3,DNAJB12,DNAJB6,DNAJC1,DNAJC11,DNAJC13,DNAJC16,DNAJC17,DNAJC18,DNAJC3,DNAJC5,DNAJC6,DNAJC7,DNAJC8,EP300,EPHX1,FKBP5,GCLC,GCLM,GSK3B,GSR,HACD3,HSPB8,KEAP1,KRAS,MAP2K1,MAP2K2,MAP2K3,MAP2K4,MAP2K5,MAP2K6,MAP3K1,MAP3K5,MAPK1,MAPK14,MAPK8,MAPK9,MRAS,NQO1,NQO2,PIK3C2A,PIK3C2B,PIK3CB,PIK3CD,PIK3R2,PIK3R3,PIK3R4,PIK3R5,PIK3R6,PRKCA,PRKCB,PRKCD,PRKCE,PRKCG,PRKCH,PRKCI,PRKCQ,PRKCZ,PRKD1,RAF1,RALA,RALB,RAP1A,RAP1B,RRAS2,SCARB1, |
| Antiproliferative Role of Somatostatin Receptor 2 | 1.37E+00 | 4.59E-01 | -5.099 | BRAF,GNB1,GNB1L,GNB5,GNG12,GNG2,GNG4,GNG7,GUCY1A2,GUCY2D,KRAS,MAP2K1,MAP2K2,MAPK1,MRAS,NOS1,NPR1,NPR3,PIK3C2A,PIK3C2B,PIK3CB,PIK3CD,PIK3R2,PIK3R3,PIK3R4,PIK3R5,PIK3R6,PTPN11,RALA,RALB,RAP1A,RAP1B,RRAS2,SRC |
| γ-linolenate Biosynthesis II (Animals) | 1.37E+00 | 5.88E-01 | -3.162 | ACSBG1,ACSL4,ACSL5,ACSL6,CYB5R3,FADS2,SLC27A1,SLC27A4,SLC27A5,SLC27A6 |
| D-myo-inositol (1,4,5)-trisphosphate Degradation | 1.37E+00 | 5.88E-01 | -3.162 | BPNT1,IMPA2,INPP5A,INPP5B,INPP5D,INPP5F,INPP5K,INPPL1,SEC16A,SYNJ2 |
| Circadian Rhythm Signaling | 1.36E+00 | 5.15E-01 | NaN | ADCYAP1R1,ARNTL,CREB3L3,CREB5,CREBBP,CRY1,CRY2,CSNK1D,CSNK1E,GRIN1,GRIN2A,GRIN2B,GRIN2C,GRIN2D,PER2,PER3,VIPR2 |
| Glutathione Biosynthesis | 1.35E+00 | 1.00E+00 | NaN | GCLC,GCLM,GSS |
| Biotin-carboxyl Carrier Protein Assembly | 1.35E+00 | 1.00E+00 | NaN | ACACA,ACACB,HLCS |
| ERK5 Signaling | 1.33E+00 | 4.58E-01 | -5.396 | CREB3L3,CREB5,CREBBP,EGFR,FOXO3,GNA12,GNAQ,IL6ST,KRAS,MAP2K5,MAP3K2,MAP3K3,MEF2A,MEF2D,MRAS,NTRK1,PTPN11,RALA,RALB,RAP1A,RAP1B,RPS6KA1,RPS6KA2,RPS6KA3,RPS6KA5,RPS6KB1,RPS6KC1,RRAS2,SGK1,SRC,WNK1,YWHAE,YWHAG |
| Glioma Invasiveness Signaling | 1.33E+00 | 4.58E-01 | -5.014 | CD44,CDC42,FNBP1,ITGB3,ITGB5,KRAS,MAPK1,MMP2,MRAS,PIK3C2A,PIK3C2B,PIK3CB,PIK3CD,PIK3R2,PIK3R3,PIK3R4,PIK3R5,PIK3R6,PLAUR,PTK2,RAC1,RAC2,RALA,RALB,RAP1A,RAP1B,RHOA,RHOBTB1,RHOD,RHOQ,RHOT1,RRAS2,TIMP2 |
| Hypoxia Signaling in the Cardiovascular System | 1.33E+00 | 4.58E-01 | -1.604 | ARNT,ATM,BIRC6,CREB3L3,CREB5,CREBBP,CSNK1D,EP300,HIF1AN,HSP90AA1,LDHA,MDM2,NFKBID,NOS3,NQO1,P4HB,PTEN,TP53,UBE2D2,UBE2E1,UBE2E2,UBE2G1,UBE2G2,UBE2H,UBE2I,UBE2J2,UBE2K,UBE2L3,UBE2N,UBE2O,UBE2Q2,UBE2R2,VEGFA |
| Role of IL-17A in Arthritis | 1.31E+00 | 4.73E-01 | NaN | IL17RA,IL17RC,MAP2K1,MAP2K2,MAP2K3,MAP2K4,MAP2K6,MAPK1,MAPK10,MAPK13,MAPK14,MAPK8,MAPK9,MAPKAPK2,NFKBID,NOS2,PIK3C2A,PIK3C2B,PIK3CB,PIK3CD,PIK3R2,PIK3R3,PIK3R4,PIK3R5,PIK3R6,RPS6KA1 |

| spplementary table 9. cisplatin damaged genes associated with sperm motility signaling pathway. | | | | | | | | | | | |
|---|---|---|---|---|---|---|---|---|---|---|---|
| **Symbol** | **Entrez Gene Name** | **Gene Symbol - human (HUGO / HGNC / Entrez Gene)** | **Expr Fold Change** | **Expected** | **Location** | **Type(s)** | **Biomarker Application(s)** | **Drug(s)** | **Entrez Gene ID for Human** | **Entrez Gene ID for Mouse** | **Entrez Gene ID for Rat** |
| *ATP1A4* | ATPase Na+K+ transporting subunit alpha 4 | ATP1A4 | -16.396 | | Plasma Membrane | transporter | | digoxin,ethacrynic acid,perphenazine | 480 | 27222 | 29132 |
| *SLC12A2* | solute carrier family 12 member 2 | SLC12A2 | -13.835 | | Plasma Membrane | transporter | | bumetanide,quinethazon | 6558 | 20496 | 83629 |
| *SLC9A5* | solute carrier family 9 member A5 | SLC9A5 | -23.144 | | Plasma Membrane | transporter | | | 6553 | 277973 | 192215 |
| *CSF2RA* | colony stimulating factor 2 receptor subunit alpha | CSF2RA | -34.989 | | Plasma Membrane | transmembrane receptor | | EPO/sargramostim,mavrilimumab,rituximab/sargramostim,sargramostim | 1438 | 12982 | 652957 |
| *FLT4* | fms related receptor tyrosine kinase 4 | FLT4 | -42.337 | | Plasma Membrane | transmembrane receptor | diagnosis,disease progression,efficacy,prognosis | 5-azacytidine/sorafenib,anlotinib,avelumab/axitinib,axitinib,axitinib/pembrolizumab,bevacizumab/sorafenib,BMS-690514,bortezomib/sorafenib,cabozantinib,cabozantinib/erlotinib,cabozantinib/nivolumab,CEP 7055,clofarabine/cytarabine/sorafenib,crizotinib/pazopanib,cyta[illegible] | 2324 | 14257 | 114110 |
| *IGF1R* | insulin like growth factor 1 receptor | IGF1R | -157.868 | | Plasma Membrane | transmembrane receptor | diagnosis,efficacy,prognosis,unspecified application | [111In]-FPI-1547,[225Ac]-FPI-1434,[I-124]-CPD-1028,AVE1642,BIIB022,BMS-754807,brigatinib,ceritinib,cixutumumab,dalotuzumab,ganitumab,IGF-1R inhibitor,IGF-methotrexate conjugate,IGF1,istiratumab,KW-2450,linsitinib,[illegible] | 3480 | 16001 | 25718 |
| *PLA2R1* | phospholipase A2 receptor 1 | PLA2R1 | -12.26 | Up | Plasma Membrane | transmembrane receptor | diagnosis | | 22925 | 18779 | 295631 |
| *ZAN* | zonadhesin (genepseudogene) | ZAN | -82.585 | | Plasma Membrane | other | | | 7455 | 22635 | 304379 |
| *ZP3* | zona pellucida glycoprotein 3 | ZP3 | -36.465 | Up | Extracellular Space | other | | | 7784 | 22788 | 114639 |
| *AATK* | apoptosis associated tyrosine kinase | AATK | -36.213 | | Cytoplasm | kinase | | | 9625 | 11302 | 690853 |
| *ABL1* | ABL proto-oncogene 1, non-receptor tyrosine kinase | ABL1 | -139.505 | | Nucleus | kinase | efficacy,response to therapy,unspecified application | bosutinib,bosutinib/hydroxyurea/ponatinib,bosutinib/imatinib,bosutinib/ponatinib,bosutinib/rituximab,clofarabine/cyclophosphamide/etoposide/imatinib,cyclophosphamide/daunorubicin/imatinib/prednisolone/vincristine,cyclophosphamide/daunorubicin/imatinib/prednisone/vincristine,cytarabin[illegible] | 25 | 11350 | 311860\|100909750 |
| *ABL2* | ABL proto-oncogene 2, non-receptor tyrosine kinase | ABL2 | -54.958 | | Cytoplasm | kinase | | dasatinib,nilotinib | 27 | 11352 | 304883 |
| *ALK* | ALK receptor tyrosine kinase | ALK | -87.244 | | Plasma Membrane | kinase | efficacy,prognosis,unspecified application | alectinib,ALK inhibitor,alkotinib,ASP3026,belizatinib,brigatinib,CEP-37440,ceritinib,conteltinib,crizotinib,crizotinib/erlotinib,crizotinib/gefitinib,crizotinib/gemcitabine,crizotinib/pazopanib,crizotinib/prednisolone,crizotinib/vemurafenib,ensartinib,entrectinib,gilteritinib,lorlatinib[illegible] | 238 | 11682 | 266802 |

| | | | | | | | | | | | |
|---|---|---|---|---|---|---|---|---|---|---|---|
| *AXL* | AXL receptor tyrosine kinase | AXL | -38.37 | | Plasma Membrane | kinase | | [illegible]ADCT-601,BA3011,bemcentinib,bosutinib,bosutinib/imatinib,bosutinib/ponatinib,bosutinib/rituximab,BPI-9016M,cabozantinib,cabozantinib/erlotinib,cabozantinib/nivolumab,DS-1205c,gilteritinib,humax-AXL-ADC,INCB081776,ningetinib,ONO-7475,RXDX-106,sitravati[illegible] | 558 | 26362 | 308444 |
| *BLK* | BLK proto-oncogene, Src family tyrosine kinase | BLK | -37.686 | | Cytoplasm | kinase | | afatinib/osimertinib,erlotinib/osimertinib,gefitinib/osimertinib,osi[illegible] | 640 | 12143 | 364403 |
| *CAMK4* | calciumcalmodulin dependent protein kinase IV | CAMK4 | -16.909 | Up | Nucleus | kinase | | | 814 | 12326 | 25050 |
| *CLK2* | CDC like kinase 2 | CLK2 | -12.662 | | Nucleus | kinase | | | 1196 | 12748 | 365842 |
| *CLK3* | CDC like kinase 3 | CLK3 | -24.902 | | Nucleus | kinase | | | 1198 | 102414 | 171305 |
| *CSF1R* | colony stimulating factor 1 receptor | CSF1R | -35.112 | | Plasma Membrane | kinase | prognosis | [illegible]3D185,AMG820,ARRY-382,blinatumomab/imatinib,BLZ-945,bosutinib,bosutinib/imatinib,bosutinib/ponatinib,bosutinib/rituximab,c-FMS inhibitor,crizotinib/pazopanib,dasatinib/nilotinib,DCC-3014,decitabine/imatinib,edicotinib,emactuzumab,erdafitinib,FPA008,imati[illegible] | 1436 | 12978 | 307403 |
| *CSK* | C-terminal Src kinase | CSK | -21.582 | | Cytoplasm | kinase | | bosutinib,bosutinib/imatinib,bosutinib/ponatinib,bosutinib/rituxi[illegible] | 1445 | 12988 | 315707 |
| *DDR2* | discoidin domain receptor tyrosine kinase 2 | DDR2 | -36.795 | | Plasma Membrane | kinase | | blinatumomab/imatinib,bosutinib/imatinib,dacomitinib,decitabine/imatinib,imatinib,imatinib/inotuzumab ozogamicin,imatinib/nilotinib,imatinib/sirolimus,imatinib/sorafenib,imatinib/sunitinib,regorafenib,sitravatin[illegible] | 4921 | 18214 | 685781 |
| *DSTYK* | dual serinethreonine and tyrosine protein kinase | DSTYK | -48.71 | | Cytoplasm | kinase | | | 25778 | 213452 | 304791 |
| *DYRK3* | dual specificity tyrosine phosphorylation regulated kinase 3 | DYRK3 | -25.833 | | Nucleus | kinase | | | 8444 | 226419 | 304775 |
| *EGFR* | epidermal growth factor receptor | EGFR | -122.314 | | Plasma Membrane | kinase | diagnosis,disease progression,efficacy,prognosis,response to therapy,safety,unspecified application | [illegible]osimertinib,18F-PEG6-IPQA,89Zr-DFO-nimotuzumab,89Zr-panitumumab,ABBV-221,ABBV-321,ABT-414,ABY-029,AC0010MA,AEE788,afatinib,afatinib/cetuximab,afatinib/osimertinib,afatinib/paclitaxel,AFM24,alflutinib,allitinib,AMG596,anastroz[illegible] | 1956 | 13649 | 24329 |
| *EPHA1* | EPH receptor A1 | EPHA1 | -12.107 | | Plasma Membrane | kinase | | | 2041 | 13835 | 312279 |

| | | | | | | | | | | | |
|---|---|---|---|---|---|---|---|---|---|---|---|
| *EPHA2* | EPH receptor A2 | EPHA2 | -37.832 | | Plasma Membrane | kinase | disease progression | blinatumomab/dasatinib,BT5528,cytarabine/dasatinib,cytarabine/dasatinib/dexamethasone/methotrexate,cytarabine/dasatinib/filgrastim/fludarabine phosphate,cytarabine/dasatinib/filgrastim/fludarabine phosphate/idarubicin,dasatinib,dasatinib/6-mercaptopurine/methotre[illegible] | 1969 | 13836 | 366492 |
| *EPHA3* | EPH receptor A3 | EPHA3 | -37.998 | | Plasma Membrane | kinase | | anti-EphA3 monoclonal antibody KB004 | 2042 | 13837 | 29210 |
| *EPHA6* | EPH receptor A6 | EPHA6 | -108.268 | | Plasma Membrane | kinase | | dacomitinib | 285220 | 13840 | 29202 |
| *EPHA8* | EPH receptor A8 | EPHA8 | -42.264 | | Plasma Membrane | kinase | | | 2046 | 13842 | 60589 |
| *EPHB1* | EPH receptor B1 | EPHB1 | -227.414 | | Plasma Membrane | kinase | | | 2047 | 270190 | 24338 |
| *EPHB2* | EPH receptor B2 | EPHB2 | -188.625 | | Plasma Membrane | kinase | unspecified application | | 2048 | 13844 | 313633 |
| *EPHB4* | EPH receptor B4 | EPHB4 | -18.155 | | Plasma Membrane | kinase | | JI 101,tesevatinib | 2050 | 13846 | 686310 |
| *ERBB2* | erb-b2 receptor tyrosine kinase 2 | ERBB2 | -30.05 | | Plasma Membrane | kinase | diagnosis,disease progression,efficacy,prognosis,safety,unspecified application | 68GaNOTA-anti-HER2 VHH1,89Zr-trastuzumab,99m-Tc-NM-02,99mTc-HPArk2,[68Ga]ABY-025,A166,ADCT-502,afatinib,afatinib/cetuximab,afatinib/osimertinib,afatinib/paclitaxel,allitinib,alpelisib/trastuzumab emtansine,anastrozole/lapatinib,anastrozole/lapatinib/trastuzu[illegible] | 2064 | 13866 | 24337 |
| *ERBB3* | erb-b2 receptor tyrosine kinase 3 | ERBB3 | -16.752 | | Plasma Membrane | kinase | efficacy,prognosis,safety | afatinib,afatinib/cetuximab,afatinib/osimertinib,afatinib/paclitaxel,BDTX-189,CDX-3379,elgemtumab,erlotinib/osimertinib,gefitinib/osimertinib,GSK2849330,istiratumab,ISU104,KTN3379,lumretuzumab,MCLA-128,MEHD7945A,MM-111,osimertinib,patritumab,REGN1400,sapitinib,s[illegible] | 2065 | 13867 | 29496 |
| *ERBB4* | erb-b2 receptor tyrosine kinase 4 | ERBB4 | -23.891 | | Plasma Membrane | kinase | efficacy,safety | afatinib,afatinib/cetuximab,afatinib/osimertinib,afatinib/paclitaxel,allitinib,BMS-599626,dacomitinib,erlotinib/osimertinib,gefitinib/osimertinib,JNJ-26483327,neratinib,neratinib/paclitaxel,osimertinib[illegible] | 2066 | 13869 | 59323 |
| *FER* | FER tyrosine kinase | FER | -51.814 | | Cytoplasm | kinase | | lorlatinib | 2241 | 14158 | 301737 |

| | | | | | | | | | | | |
|---|---|---|---|---|---|---|---|---|---|---|---|
| *FGFR1* | fibroblast growth factor receptor 1 | FGFR1 | -29.745 | | Plasma Membrane | kinase | diagnosis,prognosis,unspecified application | SD185,5-azacytidine/sorafenib,ASP5878,bortezomib/sorafenib,clofarabine/cytarabine/sorafenib,crizotinib/pazopanib,cytarabine/idarubicin/sorafenib,debio 1347,decitabine/sorafenib,derazantinib,dexamethasone/lenalidomide/sorafenib,docetaxel/nintedanib,erdafitinib,everolimus/lenva[illegible] | 2260 | 14182 | 79114 |
| *FGFR2* | fibroblast growth factor receptor 2 | FGFR2 | -60.714 | | Plasma Membrane | kinase | diagnosis,response to therapy,unspecified application | SD185,alofanib,aprutumab ixadotin,ASP5878,BAY1179470,bortezomib/dexamethasone/thalidomide,bortezomib/thalidomide,debio 1347,derazantinib,dexamethasone/thalidomide,docetaxel/nintedanib,erdafitinib,everolimus/lenvatinib,FPA144,lenvatinib,lenvatinib/pembroli[illegible] | 2263 | 14183 | 25022 |
| *FGFR4* | fibroblast growth factor receptor 4 | FGFR4 | -12.018 | | Plasma Membrane | kinase | unspecified application | ASP5878,BLU-554,erdafitinib,everolimus/lenvatinib,H3B-6527,INCB062079,lenvatinib,lenvatinib/pembrolizumab,LY2874455,PRN137[illegible] | 2264 | 14186 | 25114 |
| *FGR* | FGR proto-oncogene, Src family tyrosine kinase | FGR | -19.596 | | Nucleus | kinase | | cetuximab/irinotecan/vemurafenib,cetuximab/vemurafenib,cobimetinib/vemurafenib,crizotinib/vemurafenib,dabrafenib/trametinib/vemurafenib,ipilimumab/vemurafenib,irinotecan/panitumumab/vemurafenib,pembrolizumab/vemurafenib,rebastinib,trametinib/ve[illegible] | 2268 | 14191 | 79113 |
| *FLT1* | fms related receptor tyrosine kinase 1 | FLT1 | -17.099 | | Plasma Membrane | kinase | diagnosis,disease progression,efficacy,prognosis,response to therapy,safety | 5-azacytidine/sorafenib,avelumab/axitinib,axitinib,axitinib/pembrolizumab,bevacizumab/sorafenib,BMS-690514,bortezomib/sorafenib,cabozantinib,cabozantinib/erlotinib,cabozantinib/nivolumab,CEP 7055,clofarabine/cytarabine/sorafenib,crizotinib/pazopanib,cyta[illegible] | 2321 | 14254 | |
| *FYN* | FYN proto-oncogene, Src family tyrosine kinase | FYN | -19.656 | | Plasma Membrane | kinase | efficacy | blinatumomab/dasatinib,dasatinib,dasatinib/erlotinib,dasatinib/filgrastim,dasatinib/inotuzumab ozogamicin,dasatinib/nilotinib,JNJ-[illegible] | 2534 | 14360 | 25150 |

| | | | | | | | | | | | |
|---|---|---|---|---|---|---|---|---|---|---|---|
| *HCK* | HCK proto-oncogene, Src family tyrosine kinase | HCK | -46.631 | | Cytoplasm | kinase | | bosutinib,bosutinib/hydroxyurea/ponatinib,bosutinib/imatinib,bosutinib/ponatinib,bosutinib/rituximab,re | 3055 | 15162 | 25734 |
| *INSR* | insulin receptor | INSR | -162.937 | | Plasma Membrane | kinase | prognosis | alogliptin/INS,BMS-754807,canakinumab/INS,ceritinib,dapagliflozin/INS,dulaglutide/INS,INS,INS/pioglitazone,INS/protamine zinc insulin,INS/saxagliptin,INS/sitagliptin,INS/vildagliptin,insulin aspart,insulin aspart/insulin degludec,insulin aspart/insuli | 3643 | 16337 | 24954 |
| *JAK1* | Janus kinase 1 | JAK1 | -44.331 | | Cytoplasm | kinase | efficacy | AZD4205,baricitinib,baricitinib/methotrexate,brepocitinib,erlotinib/ruxolitinib,filgotinib,INCB052793,itacitinib,JAK1 inhibitor,jaktinib,methotrexate/tofacitinib,momelotinib,ruxolitinib,t | 3716 | 16451 | 84598 |
| *JAK2* | Janus kinase 2 | JAK2 | -17.307 | | Cytoplasm | kinase | efficacy,prognosis,unspecified application | AZD1480,baricitinib,baricitinib/methotrexate,BMS-911543,entrectinib,erlotinib/ruxolitinib,fedratinib,gandotinib,jak2 inhibitor,jaktinib,methotrexate/tofacitinib,momelotinib,NS-018,pacritinib,ruxolitinib,SB-1317,tofacitinib,upadacitinib,WP1066 | 3717 | 16452 | 24514 |
| *JAK3* | Janus kinase 3 | JAK3 | -26.841 | | Cytoplasm | kinase | efficacy | methotrexate/tofacitinib,PF-06651600,R- | 3718 | 16453 | 25326 |
| *LCK* | LCK proto-oncogene, Src family tyrosine kinase | LCK | -31.474 | | Cytoplasm | kinase | diagnosis | blinatumomab/dasatinib,crizotinib/pazopanib,dacomitinib,dasatinib,dasatinib/erlotinib,dasatinib/filgrastim,dasatinib/inotuzumab ozogamicin,dasatinib/nilotinib,docetaxel/nintedanib,JNJ-26483327,lapatinib/pazo | 3932 | 16818 | 313050 |
| *LMTK2* | lemur tyrosine kinase 2 | LMTK2 | -61.921 | | Cytoplasm | kinase | | | 22853 | 231876 | 304286 |
| *LMTK3* | lemur tyrosine kinase 3 | LMTK3 | -17.668 | | Other | kinase | | | 114783 | 381983 | 691056 |
| *MAP2K1* | mitogen-activated protein kinase kinase 1 | MAP2K1 | -51.67 | | Cytoplasm | kinase | | ARRY-424704,AS703988,binimetinib,binimetinib/cetuximab/encorafenib,binimetinib/encorafenib,binimetinib/encorafenib/panitumumab,cetuximab/dabrafenib/trametinib,cobimetinib,cobimetinib/vemurafenib,dabrafenib/ipilimumab/trametinib,dabrafenib/panitumumab/trametinib,dabrafenib/t | 5604 | 26395 | 170851 |
| *MAP2K3* | mitogen-activated protein kinase | MAP2K3 | -27.584 | | Cytoplasm | kinase | | | 5606 | 26397 | 303200 |

| | | | | | | | | | | | |
|---|---|---|---|---|---|---|---|---|---|---|---|
| *MAP2K4* | mitogen-activated protein kinase kinase 4 | MAP2K4 | -22.124 | | Cytoplasm | kinase | | binimetinib/cetuximab/encorafenib,binimetinib/encorafenib,binimetinib/encorafenib/panitumumab,cetuximab/encorafenib,enco | 6416 | 26398 | 287398 |
| *MAP2K5* | mitogen-activated protein kinase | MAP2K5 | -80.45 | | Cytoplasm | kinase | | | 5607 | 23938 | 29568 |
| *MAP2K6* | mitogen-activated protein kinase | MAP2K6 | -45.468 | | Cytoplasm | kinase | prognosis | | 5608 | 26399 | 114495 |
| *MAP3K11* | mitogen-activated protein kinase kinase kinase 11 | MAP3K11 | -14.294 | Up | Cytoplasm | kinase | | K-252 | 4296 | 26403 | 309168 |
| *MERTK* | MER proto-oncogene, tyrosine kinase | MERTK | -55.647 | | Plasma Membrane | kinase | | cabozantinib,cabozantinib/erlotinib,cabozantinib/nivolumab,INCB081776,MRX-2843,ningetinib,ONO-7475,RXDX-106,sitravati | 10461 | 17289 | 65037 |
| *MET* | MET proto-oncogene, receptor tyrosine kinase | MET | -20.804 | | Plasma Membrane | kinase | diagnosis,disease progression,efficacy,prognosis,response to therapy,unspecified application | ABN401,ABT-700,AL2846,altiratinib,AMG 208,AMG 337,amuvatinib,antibody-drug conjugate ABBV-399,ARGX-111,BMS 777607,bozitinib,BPI-9016M,c-MET inhibitor,cabozantinib,cabozantinib/erlotinib,cabozantinib/nivolumab,capmatinib,crizotinib | 4233 | 17295 | 24553 |
| *MST1R* | macrophage stimulating 1 receptor | MST1R | -18.294 | Up | Plasma Membrane | kinase | | crizotinib,crizotinib/erlotinib,crizotinib/gefitinib,crizotinib/pazopanib,crizotinib/vemurafenib,glesatinib, | 4486 | 19882 | 300999 |
| *NTRK1* | neurotrophic receptor tyrosine kinase 1 | NTRK1 | -34.9 | | Plasma Membrane | kinase | | belizatinib,DS-6051b,entrectinib,larotrectinib,lorlatinib,LOXO-195,milciclib,NOV1601,PLX7486,regorafenib,repotrectinib,TRK tyrosine kinase inhibitor VM | 4914 | 18211 | 59109 |
| *NTRK2* | neurotrophic receptor tyrosine kinase 2 | NTRK2 | -39.759 | | Plasma Membrane | kinase | diagnosis,disease progression,response to therapy | belizatinib,cabozantinib,cabozantinib/erlotinib,cabozantinib/nivolumab,DS-6051b,entrectinib,larotrectinib,lorlatinib,LOXO-195,NOV1601,ONO-7579,PLX7486,repotrecti | 4915 | 18212 | 25054 |
| *NTRK3* | neurotrophic receptor tyrosine kinase 3 | NTRK3 | -125.683 | | Plasma Membrane | kinase | | belizatinib,DS-6051b,entrectinib,larotrectinib,lorlatinib,LOXO-195,NOV1601,ONO-7579,PLX74 | 4916 | 18213 | 29613 |
| *PDGFRB* | platelet derived growth factor receptor beta | PDGFRB | -41.086 | | Plasma Membrane | kinase | prognosis,response to therapy,unspecified application | 5-azacytidine/midostaurin,5-azacytidine/sorafenib,axitinib,becaplermin,bevacizumab/sorafenib,blinatumomab/dasatinib,blinatumomab/imatinib,bortezomib/sorafenib,bosutinib/imatinib,clofarabine/cytarabine/sorafenib,crizotinib/pazopanib,cytarabine/idarubicin/sorafenib, | 5159 | 18596 | 24629 |
| *PEAK1* | pseudopodium enriched atypical | PEAK1 | -68.641 | | Plasma Membrane | kinase | | | 79834 | 244895 | 315686 |
| *PRKACA* | protein kinase cAMP-activated catalytic subunit alpha | PRKACA | -27.162 | Up | Cytoplasm | kinase | | | 5566 | 18747 | 25636 |

| | | | | | | | | | | | |
|---|---|---|---|---|---|---|---|---|---|---|---|
| *PRKAG2* | protein kinase AMP-activated non-catalytic subunit gamma 2 | PRKAG2 | -259.153 | Up | Cytoplasm | kinase | | | 51422 | 108099 | 373545 |
| *PRKAR1B* | protein kinase cAMP-dependent type I regulatory subunit beta | PRKAR1B | -165.776 | Up | Cytoplasm | kinase | | | 5575 | 19085 | 25521 |
| *PRKAR2A* | protein kinase cAMP-dependent type II regulatory subunit alpha | PRKAR2A | -86.523 | Up | Cytoplasm | kinase | | | 5576 | 19087 | 29699 |
| *PRKCA* | protein kinase C alpha | PRKCA | -179.666 | Up | Cytoplasm | kinase | | aprinocarsen,ingenol mebutate,L-threo- | 5578 | 18750 | 24680 |
| *PRKCB* | protein kinase C beta | PRKCB | -132.786 | Up | Cytoplasm | kinase | diagnosis,efficacy,unspecified application | enzastaurin,ingenol mebutate,MS-553,ruboxistaurin,sotrast | 5579 | 18751 | 25023 |
| *PRKCD* | protein kinase C delta | PRKCD | -43.302 | Up | Cytoplasm | kinase | | ingenol mebutate | 5580 | 18753 | 170538 |
| *PRKCE* | protein kinase C epsilon | PRKCE | -211.147 | Up | Cytoplasm | kinase | disease progression | ingenol mebutate,protein kinase C-epsilon activator | 5581 | 18754 | 29340 |
| *PRKCG* | protein kinase C gamma | PRKCG | -23.094 | Up | Cytoplasm | kinase | | ingenol mebutate | 5582 | 18752 | 24681 |
| *PRKCH* | protein kinase C eta | PRKCH | -39.168 | Up | Cytoplasm | kinase | diagnosis | ingenol mebutate | 5583 | 18755 | 81749 |
| *PRKCI* | protein kinase C iota | PRKCI | -47.31 | Up | Cytoplasm | kinase | unspecified application | ingenol mebutate | 5584 | 18759 | 84006 |
| *PRKCQ* | protein kinase C theta | PRKCQ | -15.21 | Up | Cytoplasm | kinase | diagnosis | ingenol mebutate,sotrastaurin | 5588 | 18761 | 85420 |
| *PRKCZ* | protein kinase C zeta | PRKCZ | -115.662 | Up | Cytoplasm | kinase | | ingenol mebutate,pkc-zeta | 5590 | 18762 | 25522 |
| *PRKD1* | protein kinase D1 | PRKD1 | -13.884 | Up | Cytoplasm | kinase | | | 5587 | 18760 | 85421 |
| *PRKG1* | protein kinase cGMP-dependent 1 | PRKG1 | -152.425 | Up | Cytoplasm | kinase | | | 5592 | 19091 | 54286 |
| *PTK2* | protein tyrosine kinase 2 | PTK2 | -121.776 | Up | Cytoplasm | kinase | diagnosis,disease progression,efficacy,prognosis | BI 853520,conteltinib,lorlatinib,repotrectin | 5747 | 14083 | 25614 |
| *PTK7* | protein tyrosine kinase 7 (inactive) | PTK7 | -76.712 | Up | Plasma Membrane | kinase | | cofetuzumab pelidotin | 5754 | 71461 | |
| *PTK2B* | protein tyrosine kinase 2 beta | PTK2B | -72.357 | Up | Cytoplasm | kinase | | conteltinib,lorlatinib,PF-562271 | 2185 | 19229 | 50646 |
| *RET* | ret proto-oncogene | RET | -50.391 | | Plasma Membrane | kinase | efficacy,response to therapy,unspecified application | 5-azacytidine/sorafenib,alectinib,amuvatinib,bevacizumab/sorafenib,blinatumomab/imatinib,bortezomib/sorafenib,BOS172738,bosutinib/hydroxyurea/ponatinib,bosutinib/imatinib,bosutinib/ponatinib,cabozantinib,cabozantinib/erlotinib,cabozantinib/nivolumab,CEP-32496,clofar[illegible] | 5979 | 19713 | 24716 |
| *ROR1* | receptor tyrosine kinase like orphan receptor 1 | ROR1 | -45.163 | | Plasma Membrane | kinase | | cirmtuzumab,VLS-101 | 4919 | 26563 | 362550 |
| *ROR2* | receptor tyrosine kinase like orphan receptor 2 | ROR2 | -102.598 | | Plasma Membrane | kinase | | conditionally active biologic anti-ROR2 antibody | 4920 | 26564 | 306782 |
| *RYK* | receptor like tyrosine kinase | RYK | -78.893 | | Plasma Membrane | kinase | | | 6259 | 20187 | 140585 |
| *SRC* | SRC proto-oncogene, non-receptor tyrosine kinase | SRC | -54.912 | | Cytoplasm | kinase | diagnosis,disease progression,efficacy,prognosis,response to therapy | AZD0424,AZM-475271,blinatumomab/dasatinib,bosutinib,bosutinib/hydroxyurea/ponatinib,bosutinib/imatinib,bosutinib/ponatinib,bosutinib/rituximab,dasatinib,dasatinib/erlotinib,dasatinib/filgrastim,dasatinib/inotuzumab ozogamicin,dasatinib/nilotinib,docetaxel/nintedanib,KX-[illegible] | 6714 | 20779 | 83805 |
| *SYK* | spleen associated tyrosine kinase | SYK | -20.783 | | Cytoplasm | kinase | efficacy,response to therapy | cerdulatinib,entospletinib,fostamatinib,GSK2646264,HMPL-523,lanraplenib,R-348,SKI-O-592,TAK- | 6850 | 20963 | 25155 |

| | | | | | | | | | | | |
|---|---|---|---|---|---|---|---|---|---|---|---|
| *TEK* | TEK receptor tyrosine kinase | TEK | -12.058 | | Plasma Membrane | kinase | efficacy | altiratinib,bosutinib/hydroxyurea/ponatinib,bosutinib/ponatinib,cabozantinib,cabozantinib/erlotinib,cabozantinib/nivolumab,CEP-11981,everolimus/vandetanib,glesatinib,pexmetinib,ponatinib,rebastinib,regorafenib,SAR103168,TIE2 inhibitor van[illegible] | 7010 | 21687 | 89804 |
| *TIE1* | tyrosine kinase with immunoglobulin like and EGF like domains 1 | TIE1 | -22.887 | | Plasma Membrane | kinase | | | 7075 | 21846 | 89806 |
| *TNK1* | tyrosine kinase non receptor 1 | TNK1 | -12.052 | | Cytoplasm | kinase | | | 8711 | 83813 | 303247 |
| *TNK2* | tyrosine kinase non receptor 2 | TNK2 | -21.61 | | Cytoplasm | kinase | | afatinib/osimertinib,cetuximab/irinotecan/vemurafenib,cetuximab/vemurafenib,cobimetinib/vemurafenib,crizotinib/vemurafenib,dabrafenib/trametinib/vemurafenib,entrectinib,erlotinib/osimertinib,gefitinib/osimertinib,ipilimumab/vemurafenib,irinotecan/panitumumab/vemurafenib,lorlatinib,osimerti[illegible] | 10188 | 51789 | 303882 |
| *TYK2* | tyrosine kinase 2 | TYK2 | -16.854 | | Plasma Membrane | kinase | | brepocitinib,ruxolitinib | 7297 | 54721 | 100361294 |
| *TYRO3* | TYRO3 protein tyrosine kinase | TYRO3 | -25.158 | | Plasma Membrane | kinase | | cabozantinib,cabozantinib/erlotinib,cabozantinib/nivolumab,RX | 7301 | 22174 | 25232 |
| *YES1* | YES proto-oncogene 1, Src family tyrosine kinase | YES1 | -34.09 | | Cytoplasm | kinase | | blinatumomab/dasatinib,cytarabine/dasatinib,cytarabine/dasatinib/dexamethasone/methotrexate,cytarabine/dasatinib/filgrastim/fludarabine phosphate,cytarabine/dasatinib/filgrastim/fludarabine phosphate/idarubicin,dasatinib,dasatinib/6-mercaptopurine/methotrexate,dasatini[illegible] | 7525 | 22612 | 24884 |
| *ZAP70* | zeta chain of T cell receptor associated protein kinase 70 | ZAP70 | -22.697 | | Plasma Membrane | kinase | prognosis,response to therapy | | 7535 | 22637 | 301348 |
| *CACNA1G* | calcium voltage-gated channel subunit alpha1 G | CACNA1G | -66.246 | | Plasma Membrane | ion channel | diagnosis | cinnarizine,ethosuximide,flunarizine,mibefradil,trimethadione,zonisamide | 8913 | 12291 | 29717 |
| *CACNA1H* | calcium voltage-gated channel subunit alpha1 H | CACNA1H | -37.527 | | Plasma Membrane | ion channel | | bepridil,cinnarizine,flunarizine,isradipine,mibefradil,zonisamide | 8912 | 58226 | 114862 |
| *CATSPER1* | cation channel sperm associated 1 | CATSPER1 | -12.958 | | Plasma Membrane | ion channel | | | 117144 | 225865 | 689349 |
| *CNGA3* | cyclic nucleotide gated channel subunit alpha 3 | CNGA3 | -31.984 | | Plasma Membrane | ion channel | unspecified application | | 1261 | 12790 | 85257 |
| *CNGB1* | cyclic nucleotide gated channel subunit beta 1 | CNGB1 | -77.73 | | Plasma Membrane | ion channel | | | 1258 | | 83686 |
| *ITPR1* | inositol 1,4,5-trisphosphate receptor type 1 | ITPR1 | -106.32 | Up | Cytoplasm | ion channel | | | 3708 | 16438 | 25262 |
| *ITPR2* | inositol 1,4,5-trisphosphate receptor type 2 | ITPR2 | -73.803 | Up | Cytoplasm | ion channel | | | 3709 | 16439 | 81678 |
| *ITPR3* | inositol 1,4,5-trisphosphate receptor type 3 | ITPR3 | -64.379 | Up | Cytoplasm | ion channel | | | 3710 | 16440 | 25679 |
| *ABHD3* | abhydrolase domain containing 3, phospholipase | ABHD3 | -23.165 | Up | Plasma Membrane | enzyme | | | 171586 | 106861 | 291793 |
| *ADCY10* | adenylate cyclase 10 | ADCY10 | -32.939 | Up | Cytoplasm | enzyme | | | 55811 | 271639 | 59320 |
| *GNAS* | GNAS complex locus | GNAS | -14.33 | Up | Plasma Membrane | enzyme | unspecified application | | 2778 | 14683 | 24896 |
| *GUCY1A2* | guanylate cyclase 1 soluble subunit alpha 2 | GUCY1A2 | -26.264 | Up | Cytoplasm | enzyme | | isosorbide dinitrate,isosorbide-5-mononitrate,nitroglycerin,nitroprusside | 2977 | 234889 | 66012 |
| *MRAS* | muscle RAS oncogene homolog | MRAS | -43.426 | Up | Plasma Membrane | enzyme | | | 22808 | 17532 | 25482 |
| *NPR1* | natriuretic peptide receptor 1 | NPR1 | -14.041 | Up | Plasma Membrane | enzyme | | hydralazine/isosorbide dinitrate,isosorbide dinitrate,nesiritide,nitroglycerin,nitroprusside | 4881 | 18160 | 24603 |
| *PDE1A* | phosphodiesterase 1A | PDE1A | -18.503 | Up | Cytoplasm | enzyme | | | 5136 | 18573 | 81529 |

| | | | | | | | | | | | |
|---|---|---|---|---|---|---|---|---|---|---|---|
| *PDE1B* | phosphodiesterase 1B | PDE1B | -13.937 | Up | Cytoplasm | enzyme | | | 5153 | 18574 | 29691 |
| *PDE1C* | phosphodiesterase 1C | PDE1C | -115.401 | Up | Cytoplasm | enzyme | | | 5137 | 18575 | 81742 |
| *PDE2A* | phosphodiesterase 2A | PDE2A | -82.23 | Up | Cytoplasm | enzyme | | | 5138 | 207728 | 81743 |
| *PDE4A* | phosphodiesterase 4A | PDE4A | -31.165 | Up | Cytoplasm | enzyme | | amrinone,anagrelide,apremilast,arofylline,aspirin/dipyridamole,aspirin/dipyridamole/telmisartan,caffeine,caffeine/ergotamine,cilomilast,dipyridamole,drotaverin,dyphylline,enprofylline,ibudilast,ketotifen,L 869298,L-826,141,milrinone,pentoxifylline,roflumi[illegible] | 5141 | 18577 | 25638 |
| *PDE4B* | phosphodiesterase 4B | PDE4B | -59.351 | Up | Cytoplasm | enzyme | | anagrelide,apremilast,arofylline,caffeine,caffeine/ergotamine,cilomilast,dipyridamole,dyphylline,enprofylline,ibudilast,ketotifen,L 869298,L-826,141,milrinone,nitroglycerin,papaverine,pentoxifylline,roflumilast,tetomilast,theophyllin[illegible] | 5142 | 18578 | 24626 |
| *PDE4C* | phosphodiesterase 4C | PDE4C | -31.789 | Up | Cytoplasm | enzyme | | anagrelide,arofylline,caffeine,caffeine/ergotamine,cilomilast,dipyridamole,dyphylline,ibudilast,ketotifen,L 869298,L-826,141,milrinone,nitroglycerin,pentoxifylline,roflumilast,tetomilast,theophylli[illegible] | 5143 | 110385 | 290646 |
| *PDE4D* | phosphodiesterase 4D | PDE4D | -177.988 | Up | Cytoplasm | enzyme | | anagrelide,apremilast,arofylline,caffeine,caffeine/ergotamine,cilomilast,dipyridamole,dyphylline,ketotifen,L 869298,L-826,141,milrinone,nitroglycerin,pentoxifylline,roflumilast,tetomilast,theophylli[illegible] | 5144 | | 24627 |
| *PLA2G5* | phospholipase A2 group V | PLA2G5 | -23.138 | Up | Extracellular Space | enzyme | | varespladib, varespladib methyl | 5322 | 18784 | 29354 |
| *PLA2G6* | phospholipase A2 group VI | PLA2G6 | -70.359 | Up | Cytoplasm | enzyme | | quinacrine | 8398 | 53357 | 360426 |
| *PLA2G10* | phospholipase A2 group X | PLA2G10 | -21.889 | Up | Cytoplasm | enzyme | | varespladib, varespladib methyl | 8399 | 26565 | 29359 |
| *PLA2G12B* | phospholipase A2 group XIIB | PLA2G12B | -15.469 | Up | Extracellular Space | enzyme | | | 84647 | 69836 | 367415 |
| *PLA2G2C* | phospholipase A2 group IIC | PLA2G2C | -15.056 | Up | Extracellular Space | enzyme | | | 391013 | 18781 | 29387 |
| *PLA2G2F* | phospholipase A2 group IIF | PLA2G2F | -13.394 | Up | Extracellular Space | enzyme | | | 64600 | 26971 | 690388 |
| *PLA2G4A* | phospholipase A2 group IVA | PLA2G4A | -20.5 | Up | Cytoplasm | enzyme | | cytosolic phospholipase A2 inhibitor,quinacrine | 5321 | 18783 | 24653 |
| *PLA2G4C* | phospholipase A2 group IVC | PLA2G4C | -20.111 | Up | Plasma Membrane | enzyme | | | 8605 | 232889 | |
| *PLA2G4D* | phospholipase A2 group IVD | PLA2G4D | -22.743 | Up | Cytoplasm | enzyme | | | 283748 | 78390 | 691905 |
| *PLA2G4E* | phospholipase A2 group IVE | PLA2G4E | -34.33 | Up | Cytoplasm | enzyme | | | 123745 | 329502 | 296091 |
| *PLA2G4F* | phospholipase A2 group IVF | PLA2G4F | -16.706 | Up | Cytoplasm | enzyme | | | 255189 | 271844 | 691907 |
| *PLAAT3* | phospholipase A and acyltransferase | PLA2G16 | -27.134 | Up | Nucleus | enzyme | | | 11145 | 225845 | 24913 |
| *PLAAT4* | phospholipase A and acyltransferase | RARRES3 | -12.735 | Up | Cytoplasm | enzyme | | | 5920 | | |
| *PLAAT5* | phospholipase A and acyltransferase | HRASLS5 | -14.643 | | Other | enzyme | | | 117245 | 66727 | 293711 |
| *PLB1* | phospholipase B1 | PLB1 | -58.33 | Up | Cytoplasm | enzyme | | | 151056 | 665270 | 192259 |
| *PLCB1* | phospholipase C beta 1 | PLCB1 | -22.627 | Up | Cytoplasm | enzyme | | | 23236 | 18795 | 24654 |
| *PLCB2* | phospholipase C beta 2 | PLCB2 | -25.707 | Up | Cytoplasm | enzyme | | | 5330 | 18796 | 85240 |
| *PLCB3* | phospholipase C beta 3 | PLCB3 | -13.203 | Up | Cytoplasm | enzyme | | | 5331 | 18797 | 29322 |
| *PLCD3* | phospholipase C delta 3 | PLCD3 | -20.815 | Up | Cytoplasm | enzyme | | | 113026 | 72469 | 287745 |
| *PLCE1* | phospholipase C epsilon 1 | PLCE1 | -70.317 | Up | Cytoplasm | enzyme | | | 51196 | 74055 | 114633 |
| *PLCG1* | phospholipase C gamma 1 | PLCG1 | -31.388 | Up | Cytoplasm | enzyme | | | 5335 | 18803 | 25738 |
| *PLCG2* | phospholipase C gamma 2 | PLCG2 | -122.094 | Up | Cytoplasm | enzyme | | | 5336 | 234779 | 29337 |
| *PLCH1* | phospholipase C eta 1 | PLCH1 | -65.089 | Up | Cytoplasm | enzyme | | | 23007 | 269437 | 310463 |
| *PLCH2* | phospholipase C eta 2 | PLCH2 | -51.407 | Up | Cytoplasm | enzyme | | | 9651 | 269615 | 313756 |
| *PLCL1* | phospholipase C like 1 (inactive) | PLCL1 | -17.168 | Up | Cytoplasm | enzyme | | quinacrine | 5334 | 227120 | 84587 |
| *PLCL2* | phospholipase C like 2 | PLCL2 | -54.778 | Up | Cytoplasm | enzyme | | | 23228 | 224860 | 301173 |
| *PNPLA3* | patatin like phospholipase domain containing 3 | PNPLA3 | -13.534 | Up | Cytoplasm | enzyme | | | 80339 | 116939 | 362972 |
| *PNPLA8* | patatin like phospholipase domain containing 8 | PNPLA8 | -16.863 | Up | Cytoplasm | enzyme | | | 50640 | 67452 | 314075 |

| Symbol | Entrez Gene Name | Gene Symbol - human (HUGO / HGNC / Entrez Gene) | Expr Fold Change | Expected | Location | Type(s) | Biomarker Application(s) | Drug(s) | Entrez Gene ID for Human | Entrez Gene ID for Mouse | Entrez Gene ID for Rat |
|---|---|---|---|---|---|---|---|---|---|---|---|
| | **Supplementary Table 10. Cisplatin damaged genes asscoiated with protein kinase A signaling pathway.** | | | | | | | | | | |
| *PTPRD* | protein tyrosine phosphatase receptor type D | PTPRD | -286.303 | Down | Plasma Membrane | phosphatase | | | 5789 | | 313278 |
| *PRKAG2* | protein kinase AMP-activated non-catalytic subunit gamma 2 | PRKAG2 | -259.153 | Up | Cytoplasm | kinase | | | 51422 | 108099 | 373545 |
| *PTPRT* | protein tyrosine phosphatase receptor type T | PTPRT | -246.937 | Down | Plasma Membrane | phosphatase | | | 11122 | 19281 | |
| *PRKCE* | protein kinase C epsilon | PRKCE | -211.147 | Up | Cytoplasm | kinase | disease progression | ingenol mebutate,protein kinase C-epsilon | 5581 | 18754 | 29340 |
| *PTPRM* | protein tyrosine phosphatase receptor type M | PTPRM | -191.957 | Down | Plasma Membrane | phosphatase | | | 5797 | 19274 | 29616 |
| *RYR2* | ryanodine receptor 2 | RYR2 | -190.216 | Up | Plasma Membrane | ion channel | | | 6262 | 20191 | 689560 |
| *PRKCA* | protein kinase C alpha | PRKCA | -179.666 | Up | Cytoplasm | kinase | | aprinocarsen,ingenol mebutate,L-threo-safingol | 5578 | 18750 | 24680 |
| *PDE4D* | phosphodiesterase 4D | PDE4D | -177.988 | Up | Cytoplasm | enzyme | | anagrelide,apremilast,arofylline,caffeine,caffeine/ergotamine,cilomilast,dipyridamole,dyphylline,ketotifen,L 869298,L-826,141,milrinone,nitroglycerin,pentoxifylline,roflumila[illegible] | 5144 | | 24627 |
| *PRKAR1B* | protein kinase cAMP-dependent type I regulatory subunit beta | PRKAR1B | -165.776 | Up | Cytoplasm | kinase | | | 5575 | 19085 | 25521 |
| *NTN1* | netrin 1 | NTN1 | -158.204 | Up | Extracellular Space | growth factor | | NP137 | 9423 | 18208 | 114523 |
| *EYA2* | EYA transcriptional coactivator and phosphatase 2 | EYA2 | -150.029 | Down | Nucleus | phosphatase | diagnosis | | 2139 | 14049 | 156826 |
| *ADCY5* | adenylate cyclase 5 | ADCY5 | -142.07 | Down | Plasma Membrane | enzyme | | | 111 | 224129 | 64532 |
| *RYR1* | ryanodine receptor 1 | RYR1 | -136.21 | Up | Cytoplasm | ion channel | | dantrolene | 6261 | 20190 | 114207 |
| *ADCY9* | adenylate cyclase 9 | ADCY9 | -134.385 | Up | Plasma Membrane | enzyme | | | 115 | 11515 | 302950 |
| *PRKCB* | protein kinase C beta | PRKCB | -132.786 | Up | Cytoplasm | kinase | diagnosis,efficacy,unspecified application | enzastaurin,ingenol mebutate,MS-553,ruboxistaurin,sotrastaurin | 5579 | 18751 | 25023 |
| *PTPRJ* | protein tyrosine phosphatase receptor type J | PTPRJ | -131.217 | Down | Plasma Membrane | phosphatase | | | 5795 | 19271 | 29645 |
| *NFATC2* | nuclear factor of activated T cells 2 | NFATC2 | -129.516 | Up | Nucleus | transcription regulator | | | 4773 | 18019 | 311658 |
| *TCF7L1* | transcription factor 7 like 1 | TCF7L1 | -127.988 | | Nucleus | transcription regulator | | | 83439 | 21415 | 312451 |
| *MYLK* | myosin light chain kinase | MYLK | -123.172 | | Cytoplasm | kinase | prognosis | K-252 | 4638 | 107589 | 288057 |
| *PLCG2* | phospholipase C gamma 2 | PLCG2 | -122.094 | Up | Cytoplasm | enzyme | | | 5336 | 234779 | 29337 |
| *PDE10A* | phosphodiesterase 10A | PDE10A | -121.859 | Up | Cytoplasm | enzyme | | aspirin/dipyridamole,aspirin/dipyridamole/telmisartan,dipyridamole.triflu[illegible] | 10846 | 23984 | 63885 |
| *PTK2* | protein tyrosine kinase 2 | PTK2 | -121.776 | Up | Cytoplasm | kinase | diagnosis,disease progression,efficacy,prognosis | BI 853520,conteltinib,lorlatinib,repotrectinib | 5747 | 14083 | 25614 |
| *AKAP13* | A-kinase anchoring protein 13 | AKAP13 | -120.012 | | Cytoplasm | other | | | 11214 | 75547 | 293024 |
| *PTPRF* | protein tyrosine phosphatase receptor type F | PTPRF | -117.676 | Down | Plasma Membrane | phosphatase | | | 5792 | 19268 | 360406 |
| *PRKCZ* | protein kinase C zeta | PRKCZ | -115.662 | Up | Cytoplasm | kinase | | ingenol mebutate,pkc-zeta inhibitor | 5590 | 18762 | 25522 |
| *PDE1C* | phosphodiesterase 1C | PDE1C | -115.401 | Up | Cytoplasm | enzyme | | | 5137 | 18575 | 81742 |
| *ADCY1* | adenylate cyclase 1 | ADCY1 | -115.053 | Down | Plasma Membrane | enzyme | | | 107 | 432530 | 305509 |
| *GNG7* | G protein subunit gamma 7 | GNG7 | -112.559 | Up | Plasma Membrane | enzyme | | | 2788 | 14708 | 58979 |
| *CAMK2B* | calciumcalmodulin dependent protein kinase II beta | CAMK2B | -112.552 | Up | Cytoplasm | kinase | | | 816 | 12323 | 24245 |
| *PTPRG* | protein tyrosine phosphatase receptor type G | PTPRG | -106.717 | Down | Plasma Membrane | phosphatase | safety | | 5793 | 19270 | 171357 |
| *ITPR1* | inositol 1,4,5-trisphosphate receptor type 1 | ITPR1 | -106.32 | Up | Cytoplasm | ion channel | | | 3708 | 16438 | 25262 |
| *CREBBP* | CREB binding protein | CREBBP | -106.296 | Up | Nucleus | transcription regulator | | CCS1477,PRI-724 | 1387 | 12914 | 54244 |
| *ADCY2* | adenylate cyclase 2 | ADCY2 | -104.281 | Up | Plasma Membrane | enzyme | | | 108 | 210044 | 81636 |
| *SMAD3* | SMAD family member 3 | SMAD3 | -103.02 | Up | Nucleus | transcription regulator | | | 4088 | 17127 | 25631 |
| *RYR3* | ryanodine receptor 3 | RYR3 | -100.923 | Up | Plasma Membrane | ion channel | unspecified application | | 6263 | 20192 | 170546 |

| | | | | | | | | | | | |
|---|---|---|---|---|---|---|---|---|---|---|---|
| *PTPRE* | protein tyrosine phosphatase receptor type E | PTPRE | -94.021 | Down | Plasma Membrane | phosphatase | | | 5791 | 19267 | 114767 |
| *CREB5* | cAMP responsive element binding protein 5 | CREB5 | -93.373 | Up | Nucleus | transcription regulator | | | 9586 | 231991 | 500131 |
| *FLNB* | filamin B | FLNB | -91.075 | | Cytoplasm | other | unspecified application | | 2317 | 286940 | 306204 |
| *PTPRS* | protein tyrosine phosphatase receptor type S | PTPRS | -87.297 | Down | Plasma Membrane | phosphatase | | | 5802 | 19280 | 25529 |
| *PRKAR2A* | protein kinase cAMP-dependent type II regulatory subunit alpha | PRKAR2A | -86.523 | Up | Cytoplasm | kinase | | | 5576 | 19087 | 29699 |
| *TCF7L2* | transcription factor 7 like 2 | TCF7L2 | -84.909 | | Nucleus | transcription regulator | efficacy | | 6934 | 21416 | 679869 |
| *PDE9A* | phosphodiesterase 9A | PDE9A | -82.384 | Up | Cytoplasm | enzyme | | | 5152 | 18585 | 191569 |
| *PDE2A* | phosphodiesterase 2A | PDE2A | -82.23 | Up | Cytoplasm | enzyme | | | 5138 | 207728 | 81743 |
| *GNG4* | G protein subunit gamma 4 | GNG4 | -80.319 | | Plasma Membrane | enzyme | | | 2786 | 14706 | 114118 |
| *PTPRU* | protein tyrosine phosphatase receptor type U | PTPRU | -79.681 | Down | Plasma Membrane | phosphatase | | | 10076 | 19273 | 116680 |
| *CNGB1* | cyclic nucleotide gated channel subunit beta 1 | CNGB1 | -77.73 | | Plasma Membrane | ion channel | | | 1258 | | 83686 |
| *PDE8B* | phosphodiesterase 8B | PDE8B | -77.337 | Up | Cytoplasm | enzyme | | ketotife | 8622 | 218461 | 309962 |
| *NFAT5* | nuclear factor of activated T cells 5 | NFAT5 | -76.395 | Up | Nucleus | transcription regulator | | | 10725 | 54446 | 307820 |
| *UBASH3B* | ubiquitin associated and SH3 domain containing B | UBASH3B | -75.672 | Down | Other | enzyme | | | 84959 | 72828 | 315579 |
| *ITPR2* | inositol 1,4,5-trisphosphate receptor type 2 | ITPR2 | -73.803 | Up | Cytoplasm | ion channel | | | 3709 | 16439 | 81678 |
| *PTK2B* | protein tyrosine kinase 2 beta | PTK2B | -72.357 | Up | Cytoplasm | kinase | | contelti nib,lorlatinib,PF-562271 | 2185 | 19229 | 50646 |
| *GSK3B* | glycogen synthase kinase 3 beta | GSK3B | -71.824 | Down | Nucleus | kinase | efficacy | enzastaurin,glycogen synthase kinase-3beta | 2932 | 56637 | 84027 |
| *PLCE1* | phospholipase C epsilon 1 | PLCE1 | -70.317 | Up | Cytoplasm | enzyme | | | 51196 | 74055 | 114633 |
| *ANAPC5* | anaphase promoting complex subunit 5 | ANAPC5 | -70.061 | Down | Nucleus | other | | | 51433 | 59008 | 288671 |
| *PTPN4* | protein tyrosine phosphatase non-receptor type 4 | PTPN4 | -69.135 | Down | Cytoplasm | phosphatase | | | 5775 | 19258 | 246116 |
| *GLI3* | GLI family zinc finger 3 | GLI3 | -68.255 | | Nucleus | transcription regulator | efficacy | | 2737 | 14634 | 140588 |
| *ADD2* | adducin 2 | ADD2 | -67.042 | Down | Cytoplasm | other | | | 119 | 11519 | 24171 |
| *PTPN14* | protein tyrosine phosphatase non-receptor type 14 | PTPN14 | -66.559 | Down | Cytoplasm | phosphatase | | | 5784 | 19250 | |
| *HHAT* | hedgehog acyltransferase | HHAT | -65.412 | Down | Cytoplasm | enzyme | | | 55733 | 226861 | 289344 |
| *PLCH1* | phospholipase C eta 1 | PLCH1 | -65.089 | Up | Cytoplasm | enzyme | | | 23007 | 269437 | 310463 |
| *BRAF* | B-Raf proto-oncogene, serinethreonine kinase | BRAF | -64.862 | Up | Cytoplasm | kinase | diagnosis,prognosis,response to therapy,unspecified application | 5-azacytidine/sorafenib,ARQ 736,belvarafenib,bevacizumab/sorafenib,BGB-3245,binimetinib/cetuximab/encorafenib,binimetinib/encorafenib,binimetinib/encorafenib/pa[illegible] | 673 | 109880 | 114486 |
| *ADD3* | adducin 3 | ADD3 | -64.623 | Down | Cytoplasm | other | | | 120 | 27360 | 25230 |
| *ITPR3* | inositol 1,4,5-trisphosphate receptor type 3 | ITPR3 | -64.379 | Up | Cytoplasm | ion channel | | | 3710 | 16440 | 25679 |
| *ANAPC1* | anaphase promoting complex subunit 1 | ANAPC1 | -63.491 | Down | Nucleus | other | | | 64682 | 17222 | 311412 |
| *AKAP9* | A-kinase anchoring protein 9 | AKAP9 | -62.142 | | Cytoplasm | other | | | 10142 | | |
| *GNB1* | G protein subunit beta 1 | GNB1 | -61.474 | Up | Plasma Membrane | enzyme | | | 2782 | 14688 | 24400 |
| *PDE4B* | phosphodiesterase 4B | PDE4B | -59.351 | Up | Cytoplasm | enzyme | | anagrelide,apremilast,arofylline,caffeine,caffeine/ergotamine,cilomilast,dipyridamole,dyphylline,enprofylline,ibudilast,ketotifen,L 869298,L-826,141,milrinone,nitroglycerin[illegible] | 5142 | 18578 | 24626 |
| *ADCY3* | adenylate cyclase 3 | ADCY3 | -59.244 | Up | Plasma Membrane | enzyme | | | 109 | 104111 | 64508 |
| *YWHAE* | tyrosine 3-monooxygenasetryptophan 5-monooxygenase activation protein epsilon | YWHAE | -58.942 | Up | Cytoplasm | other | diagnosis | | 7531 | 22627 | 29753 |
| *PPP3CC* | protein phosphatase 3 catalytic subunit gamma | PPP3CC | -58.648 | Up | Cytoplasm | phosphatase | | cyclosporin A,cyclosporin A/methotrexate,methylprednisolone/tacrolimus,pimecrolimus,prednisone/tacrolimus,tacrolimus[illegible] | 5533 | 19057 | 171378 |

| | | | | | | | | | | | |
|---|---|---|---|---|---|---|---|---|---|---|---|
| *PDE8A* | phosphodiesterase 8A | PDE8A | -57.523 | Up | Cytoplasm | enzyme | | anagrelide,dipyridamole,dyphylline,ketotifen,milrinone,nitroglycerin,pentoxifylline,theophylline,t | 5151 | 18584 | 308776 |
| *ADCY7* | adenylate cyclase 7 | ADCY7 | -56.373 | Up | Plasma Membrane | enzyme | | | 113 | 11513 | 84420 |
| *GNAQ* | G protein subunit alpha q | GNAQ | -55.251 | Up | Plasma Membrane | enzyme | | | 2776 | 14682 | 81666 |
| *NFATC3* | nuclear factor of activated T cells 3 | NFATC3 | -55.002 | Up | Nucleus | transcription regulator | | | 4775 | 18021 | 361400 |
| *PLCL2* | phospholipase C like 2 | PLCL2 | -54.778 | Up | Cytoplasm | enzyme | | | 23228 | 224860 | 301173 |
| *RHOA* | ras homolog family member A | RHOA | -52.591 | Up | Cytoplasm | enzyme | | | 387 | 11848 | 117273 |
| *CAMK2G* | calciumcalmodulin dependent protein kinase II gamma | CAMK2G | -52.234 | Up | Cytoplasm | kinase | | | 818 | 12325 | 171140 |
| *MAP2K1* | mitogen-activated protein kinase kinase 1 | MAP2K1 | -51.67 | Up | Cytoplasm | kinase | | ARRY-424704,AS703988,binimetinib,binimetinib/cetuximab/encorafenib,binimetinib/encorafenib,binimetinib/encorafenib/panitumumab,cetuximab/dabrafenib/trametinib,cobimeti[illegible] | 5604 | 26395 | 170851 |
| *PLCH2* | phospholipase C eta 2 | PLCH2 | -51.407 | Up | Cytoplasm | enzyme | | | 9651 | 269615 | 313756 |
| *PHKB* | phosphorylase kinase regulatory subunit beta | PHKB | -48.668 | Up | Cytoplasm | kinase | | | 5257 | 102093 | 361377 |
| *PTPRR* | protein tyrosine phosphatase receptor type R | PTPRR | -48.363 | Down | Plasma Membrane | phosphatase | | | 5801 | 19279 | 94202 |
| *TGFB2* | transforming growth factor beta 2 | TGFB2 | -47.63 | Up | Extracellular Space | growth factor | efficacy,unspecified application | dalantercept,fresolimumab,trabederse | 7042 | 21808 | 81809 |
| *PTPN1* | protein tyrosine phosphatase non-receptor type 1 | PTPN1 | -47.354 | Down | Cytoplasm | phosphatase | prognosis | protein tyrosine phosphatase 1B inhibitor,trodusq | 5770 | 19246 | 24697 |
| *PRKCI* | protein kinase C iota | PRKCI | -47.31 | Up | Cytoplasm | kinase | unspecified application | ingenol mebutate | 5584 | 18759 | 84006 |
| *PHKA2* | phosphorylase kinase regulatory subunit alpha 2 | PHKA2 | -46.759 | Up | Cytoplasm | kinase | | K-252 | 5256 | 110094 | 678739 |
| *EYA3* | EYA transcriptional coactivator and phosphatase 3 | EYA3 | -46.596 | Down | Nucleus | phosphatase | | | 2140 | 14050 | 313027 |
| *RAF1* | Raf-1 proto-oncogene, serinethreonine kinase | RAF1 | -45.385 | Up | Cytoplasm | kinase | efficacy | 5-azacytidine/sorafenib,belvarafenib,bevacizumab/sorafenib,binimetinib/cetuximab/encorafenib,binimetinib/encorafenib,binimetinib/encorafenib/panitumumab,bortezomib/sor[illegible] | 5894 | 110157 | 24703 |
| *MYH10* | myosin heavy chain 10 | MYH10 | -45.172 | Down | Cytoplasm | enzyme | unspecified application | | 4628 | 77579 | 79433 |
| *MYLK3* | myosin light chain kinase 3 | MYLK3 | -43.609 | | Cytoplasm | kinase | | | 91807 | 213435 | 291926 |
| *PRKCD* | protein kinase C delta | PRKCD | -43.302 | Up | Cytoplasm | kinase | | ingenol mebutate | 5580 | 18753 | 170538 |
| *RAP1A* | RAP1A, member of RAS oncogene family | RAP1A | -43.262 | Up | Cytoplasm | enzyme | | | 5906 | 109905 | 295347 |
| *CAMK2A* | calciumcalmodulin dependent protein kinase II alpha | CAMK2A | -43.185 | Up | Cytoplasm | kinase | | | 815 | 12322 | 25400 |
| *TCF7* | transcription factor 7 | TCF7 | -42.748 | | Nucleus | transcription regulator | | | 6932 | | |
| *ATF1* | activating transcription factor 1 | ATF1 | -41.718 | Up | Nucleus | transcription regulator | | | 466 | 11908\|100040260 | 315305 |
| *NFATC1* | nuclear factor of activated T cells 1 | NFATC1 | -41.099 | Up | Nucleus | transcription regulator | | | 4772 | 18018 | 100361818 |
| *PRKCH* | protein kinase C eta | PRKCH | -39.168 | Up | Cytoplasm | kinase | diagnosis | ingenol mebutate | 5583 | 18755 | 81749 |

| | | | | | | | | | | | |
|---|---|---|---|---|---|---|---|---|---|---|---|
| *MAPK1* | mitogen-activated protein kinase 1 | MAPK1 | -38.855 | Up | Cytoplasm | kinase | efficacy | ASN007,ASTX029,binimetinib,KO-947,LTT462,LY-3007113,LY3214996,MAP kinase1 inhibitor,pexmetinib,ulix | 5594 | 26413 | 116590 |
| *PTPN11* | protein tyrosine phosphatase non-receptor type 11 | PTPN11 | -38.48 | Down | Cytoplasm | phosphatase | | JAB-3068,RLY-1971,RMC-4630,TNO155 | 5781 | 19247 | 25622 |
| *PPP1R12A* | protein phosphatase 1 regulatory subunit 12A | PPP1R12A | -37.984 | Down | Cytoplasm | phosphatase | | | 4659 | 17931 | |
| *PXN* | paxillin | PXN | -37.245 | | Cytoplasm | other | efficacy,prognosis,unspecified application | | 5829 | 19303 | 360820 |

| Supplementary Table 10. Cisplatin damaged genes asscoiated with molecular mechanisms of cancer singaling pathway. | | | | | | | | | | | |
|---|---|---|---|---|---|---|---|---|---|---|---|
| Symbol | Entrez Gene Name | Gene Symbol - human (HUGO / HGNC / Entrez Gene) | Expr Fold Change | Expected | Location | Type(s) | Biomarker Application(s) | Drug(s) | Entrez Gene ID for Human | Entrez Gene ID for Mouse | Entrez Gene ID for Rat |
| PRKAG2 | protein kinase AMP-activated non-catalytic subunit gamma 2 | *PRKAG2* | -259.153 | | Cytoplasm | kinase | | | 51422 | 108099 | 373545 |
| PRKCE | protein kinase C epsilon | *PRKCE* | -211.147 | | Cytoplasm | kinase | disease progression | ingenol mebutate,protein kinase C-epsilon activator | 5581 | 18754 | 29340 |
| PRKCA | protein kinase C alpha | *PRKCA* | -179.666 | | Cytoplasm | kinase | | aprinocarsen,ingenol mebutate,L-threo-safingol | 5578 | 18750 | 24680 |
| PRKAR1B | protein kinase cAMP-dependent type I regulatory subunit beta | *PRKAR1B* | -165.776 | | Cytoplasm | kinase | | | 5575 | 19085 | 25521 |
| CTNNA2 | catenin alpha 2 | *CTNNA2* | -164.443 | | Plasma Membrane | other | | | 1496 | 12386 | 297357 |
| CTNNA1 | catenin alpha 1 | *CTNNA1* | -159.914 | | Plasma Membrane | other | diagnosis | | 1495 | 12385 | 307505 |
| ADCY5 | adenylate cyclase 5 | *ADCY5* | -142.07 | | Plasma Membrane | enzyme | | | 111 | 224129 | 64532 |
| ABL1 | ABL proto-oncogene 1, non-receptor tyrosine kinase | *ABL1* | -139.505 | | Nucleus | kinase | efficacy,response to therapy,unspecified application | bosutinib,bosutinib/hydroxyurea/ponatinib,bosutinib/imatinib,bosutinib/ponatinib,bosutinib/rituximab,clofarabine/cyclophosphamide/etoposide/imatinib,cyclophosphamide/daunorubicin/imatinib/prednisolone/vincristine,cyclophosphamide/daunorubicin/imatinib/prednisone/vincristine,cytarabine/etoposide/imatinib/mitoxantrone,cytarabine/filgrastim/fludarabine phosphate/imatinib,cytarabine/imatinib,dasatinib,dasatinib/nilotinib,dexamethasone/imatinib,dexamethasone/imatinib/vincristine,etoposide/imatinib/mitoxantrone,filgrastim/imatinib,HQP1351,hydrocortisone/imatinib,imatinib,imatinib/nilotinib,imatinib/nilotinib/pegintron,imatinib/prednisone,imatinib/thioguanine,KW 2449,L- | 25 | 11350 | 311860\|100909750 |
| LRP5 | LDL receptor related protein 5 | *LRP5* | -138.982 | | Plasma Membrane | transmembrane receptor | diagnosis | | 4041 | 16973 | 293649 |
| RAPGEF1 | Rap guanine nucleotide exchange factor 1 | *RAPGEF1* | -137.024 | | Cytoplasm | other | | | 2889 | 107746 | 63881 |
| ADCY9 | adenylate cyclase 9 | *ADCY9* | -134.385 | | Plasma Membrane | enzyme | | | 115 | 11515 | 302950 |
| PRKCB | protein kinase C beta | *PRKCB* | -132.786 | | Cytoplasm | kinase | diagnosis,efficacy,unspecified application | enzastaurin,ingenol mebutate,MS- | 5579 | 18751 | 25023 |
| FNBP1 | formin binding protein 1 | *FNBP1* | -123.266 | | Nucleus | enzyme | | | 23048 | 14269 | 192348 |
| PTK2 | protein tyrosine kinase 2 | *PTK2* | -121.776 | | Cytoplasm | kinase | diagnosis,disease progression,efficacy,prognosis | BI 853520,conteltinib,lorlatinib,repotrectinib | 5747 | 14083 | 25614 |
| ARHGEF3 | Rho guanine nucleotide exchange factor 3 | *ARHGEF3* | -121.042 | | Cytoplasm | other | | | 50650 | 71704 | 290541 |
| PRKCZ | protein kinase C zeta | *PRKCZ* | -115.662 | | Cytoplasm | kinase | | ingenol mebutate,pkc-zeta inhibitor | 5590 | 18762 | 25522 |
| ADCY1 | adenylate cyclase 1 | *ADCY1* | -115.053 | | Plasma Membrane | enzyme | | | 107 | 432530 | 305509 |
| CAMK2B | calciumcalmodulin dependent protein kinase II beta | *CAMK2B* | -112.552 | | Cytoplasm | kinase | | | 816 | 12323 | 24245 |
| CREBBP | CREB binding protein | *CREBBP* | -106.296 | | Nucleus | transcription regulator | | CCS1477,PRI-724 | 1387 | 12914 | 54244 |
| HIPK2 | homeodomain interacting protein kinase 2 | *HIPK2* | -105.809 | | Nucleus | kinase | | | 28996 | 15258 | 362342 |
| ADCY2 | adenylate cyclase 2 | *ADCY2* | -104.281 | | Plasma Membrane | enzyme | | | 108 | 210044 | 81636 |
| CCND3 | cyclin D3 | *CCND3* | -104.136 | | Nucleus | kinase | prognosis | | 896 | 12445 | 25193 |
| SMAD3 | SMAD family member 3 | *SMAD3* | -103.02 | | Nucleus | transcription regulator | | | 4088 | 17127 | 25631 |
| CDK14 | cyclin dependent kinase 14 | *CDK14* | -101.934 | | Nucleus | kinase | | | 5218 | 18647 | 362316 |
| PIK3CB | hosphatidylinositol-4,5-bisphosphate 3-kinase catalytic subunit be | *PIK3CB* | -100.595 | | Cytoplasm | kinase | | acalisib,AZD8186,BAY1082439,bimiralisib,buparlisib,CLR457,dactolisib,GSK2636771,KA2237,pictilisib,pilaralisib,PX-866,SAR260301,SF 1126 | 5291 | 74769 | 85243 |
| SUFU | SUFU negative regulator of hedgehog signaling | *SUFU* | -100.386 | | Nucleus | transcription regulator | | | 51684 | 24069 | 361769 |
| GNAO1 | G protein subunit alpha o1 | *GNAO1* | -98.356 | | Plasma Membrane | enzyme | | | 2775 | 14681 | 50664 |
| NF1 | neurofibromin 1 | *NF1* | -88.589 | | Cytoplasm | other | | | 4763 | 18015 | 24592 |
| ARHGEF4 | Rho guanine nucleotide exchange factor 4 | *ARHGEF4* | -88.334 | | Cytoplasm | other | | | 50649 | 226970 | 301334 |
| PRKAR2A | protein kinase cAMP-dependent type II regulatory subunit alpha | *PRKAR2A* | -86.523 | | Cytoplasm | kinase | | | 5576 | 19087 | 29699 |
| RASGRF1 | Ras protein specific guanine nucleotide releasing factor 1 | *RASGRF1* | -84.955 | | Cytoplasm | other | | | 5923 | 19417 | 192213 |
| ARHGEF7 | Rho guanine nucleotide exchange factor 7 | *ARHGEF7* | -82.434 | | Cytoplasm | other | | | 8874 | 54126 | 114559 |
| ARHGEF10 | Rho guanine nucleotide exchange factor 10 | *ARHGEF10* | -80.186 | | Cytoplasm | enzyme | | | 9639 | 234094 | 306618 |
| LRP1 | LDL receptor related protein 1 | *LRP1* | -79.42 | | Plasma Membrane | transmembrane receptor | | paclitaxel-angiopep-2 conjugate,serpin peptide 16 | 4035 | 16971 | 299858 |
| CDK12 | cyclin dependent kinase 12 | *CDK12* | -76.344 | | Nucleus | kinase | | | 51755 | 69131 | 192350 |
| SMAD6 | SMAD family member 6 | *SMAD6* | -73.945 | | Nucleus | transcription regulator | unspecified application | | 4091 | 17130 | 367100 |
| AXIN1 | axin 1 | *AXIN1* | -73.121 | | Cytoplasm | other | | | 8312 | 12005 | 79257 |
| GSK3B | glycogen synthase kinase 3 beta | *GSK3B* | -71.824 | | Nucleus | kinase | efficacy | enzastaurin,glycogen synthase kinase-3beta | 2932 | 56637 | 84027 |
| PIK3R3 | phosphoinositide-3-kinase regulatory subunit 3 | *PIK3R3* | -67.266 | | Cytoplasm | kinase | | | 8503 | 18710 | 60664 |
| GNAL | G protein subunit alpha L | *GNAL* | -65.921 | | Cytoplasm | enzyme | | | 2774 | 14680 | 24611 |
| HHAT | hedgehog acyltransferase | *HHAT* | -65.412 | | Cytoplasm | enzyme | | | 55733 | 226861 | 289344 |
| BRAF | B-Raf proto-oncogene, serinethreonine kinase | *BRAF* | -64.862 | | Cytoplasm | kinase | diagnosis,prognosis,response to therapy,unspecified application | 5-azacytidine/sorafenib,ARQ 736,belvarafenib,bevacizumab/sorafenib,BGB-3245,binimetinib/cetuximab/encorafenib,binimetinib/encorafenib,binimetinib/encorafenib/panitumumab,bortezomib/sorafenib,BRAF inhibitor,CEP-32496,cetuximab/dabrafenib/trametinib,cetuximab/encorafenib,cetuximab/irinotecan/vemurafenib,cetuximab/vemurafenib,clofarabine/cytarabine/sorafenib,cobimetinib/vemurafenib,crizotinib/vemurafenib,cytarabine/idarubicin/sorafenib,dabrafenib,dabrafenib/ipilimumab/trametinib,dabrafenib/panitumumab,dabrafenib/panitumumab/trametinib,dabrafenib/trametinib,dabrafenib/trametinib/vemurafenib,decitabine/sorafenib,dexamethasone/lenalidomide/sor | 673 | 109880 | 114486 |
| BMPR1A | bone morphogenetic protein receptor type 1A | *BMPR1A* | -64.295 | | Plasma Membrane | kinase | diagnosis,unspecified | | 657 | 12166 | 81507 |
| BMP7 | bone morphogenetic protein 7 | *BMP7* | -63.42 | | Extracellular Space | growth factor | diagnosis | | 655 | 12162 | 85272 |
| PIK3C2A | osphatidylinositol-4-phosphate 3-kinase catalytic subunit type 2 alp | *PIK3C2A* | -62.915 | | Cytoplasm | kinase | | | 5286 | 18704 | 361632 |
| SIN3A | SIN3 transcription regulator family member A | *SIN3A* | -61.177 | | Nucleus | transcription regulator | | | 25942 | 20466 | 363067 |
| ADCY3 | adenylate cyclase 3 | *ADCY3* | -59.244 | | Plasma Membrane | enzyme | | | 109 | 104111 | 64508 |
| CDK13 | cyclin dependent kinase 13 | *CDK13* | -58.918 | | Nucleus | kinase | | | 8621 | 69562 | 306998 |
| WNT7A | Wnt family member 7A | *WNT7A* | -58.166 | | Extracellular Space | cytokine | | | 7476 | 22421 | 114850 |
| BMP6 | bone morphogenetic protein 6 | *BMP6* | -58.075 | | Extracellular Space | growth factor | | | 654 | 12161 | 25644 |
| PIK3CD | hosphatidylinositol-4,5-bisphosphate 3-kinase catalytic subunit del | *PIK3CD* | -57.218 | | Cytoplasm | kinase | | acalisib,ACP-196/ACP-319,ACP-319,bendamustine/idelalisib/rituximab,bimiralisib,buparlisib,CLR457,copanlisib,dactolisib,dezapelisib,duvelisib,GS-9901,idelalisib,idelalisib/ofatumumab,idelalisib/rituximab,KA2237,nemiralisib,parsaclisib,pictilisib,pilaralisib,PWT143,PX-866,SF | 5293 | 18707 | 366508 |
| RBL1 | RB transcriptional corepressor like 1 | *RBL1* | -57.19 | | Nucleus | transcription regulator | | | 5933 | 19650 | 680111 |
| AKT3 | AKT serinethreonine kinase 3 | *AKT3* | -56.867 | | Cytoplasm | kinase | efficacy | enzastaurin,GSK690693,triciribine,triciribine phosphate | 10000 | 23797 | 29414 |
| PAK4 | p21 (RAC1) activated kinase 4 | *PAK4* | -56.64 | | Cytoplasm | kinase | | KPT-9274,PF-3758309 | 10298 | 70584 | 292756 |
| GRB2 | growth factor receptor bound protein 2 | *GRB2* | -56.618 | | Cytoplasm | other | | liposome-incorporated Grb2 antisense | 2885 | 14784 | 81504 |
| ADCY7 | adenylate cyclase 7 | *ADCY7* | -56.373 | | Plasma Membrane | enzyme | | | 113 | 11513 | 84420 |
| RASGRF2 | Ras protein specific guanine nucleotide releasing factor 2 | *RASGRF2* | -55.709 | | Cytoplasm | other | | | 5924 | 19418 | 114513 |
| WNT3 | Wnt family member 3 | *WNT3* | -55.699 | | Extracellular Space | other | | | 7473 | 22415 | 24882 |

| | | | | | | | | | | | |
|---|---|---|---|---|---|---|---|---|---|---|---|
| ARHGEF17 | Rho guanine nucleotide exchange factor 17 | *ARHGEF17* | -55.63 | | Cytoplasm | other | | | 9828 | 207212 | 308862 |
| GNAQ | G protein subunit alpha q | *GNAQ* | -55.251 | | Plasma Membrane | enzyme | | | 2776 | 14682 | 81666 |
| SRC | SRC proto-oncogene, non-receptor tyrosine kinase | *SRC* | -54.912 | | Cytoplasm | kinase | diagnosis,disease progression,efficacy,prognosis,response to therapy | AZD0424,AZM-475271,blinatumomab/dasatinib,bosutinib,bosutinib/hydroxyurea/ponatinib,bosutinib/imatinib,bosutinib/ponatinib,bosutinib/rituximab,dasatinib,dasatinib/erlotinib,dasatinib/filgrastim,dasatinib/inotuzumab ozogamicin,dasatinib/nilotinib,docetaxel/nintedanib,KX-01,nintedanib,pexmetinib,repo | 6714 | 20779 | 83805 |
| BRCA1 | BRCA1 DNA repair associated | *BRCA1* | -54.566 | | Nucleus | transcription regulator | diagnosis,efficacy,response to therapy | | 672 | 12189 | 497672 |
| TFDP1 | transcription factor Dp-1 | *TFDP1* | -53.504 | | Nucleus | transcription regulator | | | 7027 | 21781 | 361178 |
| AKT2 | AKT serinethreonine kinase 2 | *AKT2* | -53.338 | | Cytoplasm | kinase | | BAY1125976,enzastaurin,GSK690693,triciribine,triciribine phosphate | 208 | 11652 | 25233 |
| RALGDS | ral guanine nucleotide dissociation stimulator | *RALGDS* | -53.207 | | Cytoplasm | other | | | 5900 | 19730 | 29622 |
| RHOA | ras homolog family member A | *RHOA* | -52.591 | | Cytoplasm | enzyme | | | 387 | 11848 | 117273 |
| PIK3C2B | osphatidylinositol-4-phosphate 3-kinase catalytic subunit type 2 be | *PIK3C2B* | -52.575 | | Cytoplasm | kinase | | | 5287 | 240752 | 289021 |
| RHOT1 | ras homolog family member T1 | *RHOT1* | -52.347 | | Cytoplasm | enzyme | | | 55288 | 59040 | 303351 |
| CAMK2G | calciumcalmodulin dependent protein kinase II gamma | *CAMK2G* | -52.234 | | Cytoplasm | kinase | | | 818 | 12325 | 171140 |
| MAP2K1 | mitogen-activated protein kinase kinase 1 | *MAP2K1* | -51.67 | | Cytoplasm | kinase | | ARRY-424704,AS703988,binimetinib,binimetinib/cetuximab/encorafenib,binimetinib/encorafenib,binimetinib/encorafenib/panitumumab,cetuximab/dabrafenib/trametinib,cobimetinib,cobimetinib/vemurafenib,dabrafenib/ipilimumab/trametinib,dabrafenib/panitumumab/trametinib,dabrafenib/trametinib,dabrafenib/trametinib/vemurafenib,E 6201,FCN-159,MKK1 inhibitor,PD184352,pimasertib,refametinib,selumetinib,TAK 733,trametinib,trametinib/vem | 5604 | 26395 | 170851 |
| FANCD2 | FA complementation group D2 | *FANCD2* | -51.614 | | Nucleus | other | | | 2177 | 211651 | 312641 |
| PIK3R5 | phosphoinositide-3-kinase regulatory subunit 5 | *PIK3R5* | -49.401 | | Cytoplasm | kinase | | | 23533 | 320207 | 497931 |
| WNT9B | Wnt family member 9B | *WNT9B* | -49.098 | | Extracellular Space | other | | | 7484 | 22412 | 303586 |
| GNA12 | G protein subunit alpha 12 | *GNA12* | -48.833 | | Plasma Membrane | enzyme | response to therapy | | 2768 | 14673 | 81663 |
| PAK1 | p21 (RAC1) activated kinase 1 | *PAK1* | -48.568 | | Cytoplasm | kinase | | K-252 | 5058 | 18479 | 29431 |
| TGFB2 | transforming growth factor beta 2 | *TGFB2* | -47.63 | | Extracellular Space | growth factor | efficacy,unspecified application | dalantercept,fresolimumab,trabedersen | 7042 | 21808 | 81809 |
| PRKCI | protein kinase C iota | *PRKCI* | -47.31 | | Cytoplasm | kinase | unspecified application | ingenol mebutate | 5584 | 18759 | 84006 |
| GAB2 | GRB2 associated binding protein 2 | *GAB2* | -46.911 | | Cytoplasm | other | | | 9846 | 14389 | 84477 |
| CBL | Cbl proto-oncogene | *CBL* | -46.77 | | Nucleus | transcription regulator | | | 867 | 12402 | 500985 |
| BMP1 | bone morphogenetic protein 1 | *BMP1* | -46.219 | | Extracellular Space | peptidase | efficacy | | 649 | 12153 | 83470 |
| ARHGEF2 | RhoRac guanine nucleotide exchange factor 2 | *ARHGEF2* | -45.642 | | Cytoplasm | other | | | 9181 | 16800 | 310635 |
| ARHGEF11 | Rho guanine nucleotide exchange factor 11 | *ARHGEF11* | -45.49 | | Cytoplasm | other | | | 9826 | 213498 | 78966 |
| MAP2K6 | mitogen-activated protein kinase kinase 6 | *MAP2K6* | -45.468 | | Cytoplasm | kinase | prognosis | | 5608 | 26399 | 114495 |
| RAF1 | Raf-1 proto-oncogene, serinethreonine kinase | *RAF1* | -45.385 | | Cytoplasm | kinase | efficacy | 5-azacytidine/sorafenib,belvarafenib,bevacizumab/sorafenib,binimetinib/cetuximab/encorafenib,binimetinib/encorafenib,binimetinib/encorafenib/panitumumab,bortezomib/sorafenib,c-raf antisense oligonucleotide liposome,cetuximab/dabrafenib/trametinib,cetuximab/encorafenib,cetuximab/irinotecan/vemurafenib,cetuximab/vemurafenib,CHIR-265,clofarabine/cytarabine/sorafenib,cobimetinib/vemurafenib,crizotinib/vemurafenib,cytarabine/idarubicin/sorafenib,dabrafenib,dabrafenib/ipilimumab/trametinib,dabrafenib/panitumumab,dabrafenib/panitumumab/trametinib,dabrafenib/trametinib,dabrafenib/trametinib/vemurafenib,decitabine/sorafenib,de | 5894 | 110157 | 24703 |
| CDH1 | cadherin 1 | *CDH1* | -45.053 | | Plasma Membrane | other | diagnosis,disease progression,efficacy,prognosis | | 999 | 12550 | 83502 |
| CRK | CRK proto-oncogene, adaptor protein | *CRK* | -44.635 | | Cytoplasm | other | | | 1398 | 12928 | 54245 |
| JAK1 | Janus kinase 1 | *JAK1* | -44.331 | | Cytoplasm | kinase | efficacy | AZD4205,baricitinib,baricitinib/methotrexate,brepocitinib,erlotinib/ruxolitinib,filgotinib,INCB052793,itacitinib,JAK1 inhibitor,jaktinib,methotrexate/tofacitinib,momelotinib,ruxolitinib,tofacitinib,upadacitinib | 3716 | 16451 | 84598 |
| MRAS | muscle RAS oncogene homolog | *MRAS* | -43.426 | | Plasma Membrane | enzyme | | | 22808 | 17532 | 25482 |
| PRKCD | protein kinase C delta | *PRKCD* | -43.302 | | Cytoplasm | kinase | | ingenol mebutate | 5580 | 18753 | 170538 |
| RAP1A | RAP1A, member of RAS oncogene family | *RAP1A* | -43.262 | | Cytoplasm | enzyme | | | 5906 | 109905 | 295347 |
| CAMK2A | calciumcalmodulin dependent protein kinase II alpha | *CAMK2A* | -43.185 | | Cytoplasm | kinase | | | 815 | 12322 | 25400 |
| SOS1 | SOS RasRac guanine nucleotide exchange factor 1 | *SOS1* | -42.685 | | Cytoplasm | other | | BI 1701963 | 6654 | 20662 | 313845 |
| MAPK9 | mitogen-activated protein kinase 9 | *MAPK9* | -42.643 | | Cytoplasm | kinase | | binimetinib/cetuximab/encorafenib,binimetinib/encorafenib,binimetinib/encorafenib/panitumumab,cetuximab/encorafeni | 5601 | 26420 | 50658 |
| PIK3R6 | phosphoinositide-3-kinase regulatory subunit 6 | *PIK3R6* | -42.633 | | Cytoplasm | kinase | | | 146850 | 104709 | 497932 |
| BMPR2 | bone morphogenetic protein receptor type 2 | *BMPR2* | -42.479 | | Plasma Membrane | kinase | diagnosis,unspecified | | 659 | 12168 | 140590 |
| PRKDC | protein kinase, DNA-activated, catalytic subunit | *PRKDC* | -41.35 | | Nucleus | kinase | | CC- | 5591 | 19090 | 360748 |
| ATR | ATR serinethreonine kinase | *ATR* | -40.438 | | Nucleus | kinase | | BAY1895344,berzosertib,ceralasertib,VX-803 | 545 | 245000 | 685055 |
| WNT3A | Wnt family member 3A | *WNT3A* | -40.325 | | Extracellular Space | cytokine | | | 89780 | 22416 | 303181 |
| PRKCH | protein kinase C eta | *PRKCH* | -39.168 | | Cytoplasm | kinase | diagnosis | ingenol mebutate | 5583 | 18755 | 81749 |
| CDK6 | cyclin dependent kinase 6 | *CDK6* | -39.081 | | Nucleus | kinase | | abemaciclib,abemaciclib/anastrozole,abemaciclib/aromatase inhibitor,abemaciclib/aromatase inhibitor/GnRH analog,abemaciclib/exemestane,abemaciclib/fulvestrant,abemaciclib/letrozole,alvocidib,anastrozole/palbociclib,anastrozole/ribociclib,everolimus/ribociclib,exemestane/palbociclib,exemestane/ribociclib,FCN-437,FLX925,FN-1501,fulvestrant/palbociclib,fulvestrant/ribociclib,HS-10342,lerociclib,letrozole/palbociclib,letrozole/ribociclib,palb | 1021 | 12571 | 114483 |
| CDK17 | cyclin dependent kinase 17 | *CDK17* | -39.005 | | Cytoplasm | kinase | | | 5128 | 237459 | 314743 |
| MAPK1 | mitogen-activated protein kinase 1 | *MAPK1* | -38.855 | | Cytoplasm | kinase | efficacy | ASN007,ASTX029,binimetinib,KO-947,LTT462,LY-3007113,LY3214996,MAP kinase1 inhibitor,pexmetinib,ulixertinib | 5594 | 26413 | 116590 |
| PTPN11 | protein tyrosine phosphatase non-receptor type 11 | *PTPN11* | -38.48 | | Cytoplasm | phosphatase | | JAB-3068,RLY-1971,RMC-4630,TNO155 | 5781 | 19247 | 25622 |
| NOTCH1 | notch receptor 1 | *NOTCH1* | -36.184 | | Plasma Membrane | transcription regulator | diagnosis,efficacy | OMP-52M51 | 4851 | 18128 | 25496 |
| GNAI2 | G protein subunit alpha i2 | *GNAI2* | -35.922 | | Plasma Membrane | enzyme | | | 2771 | 14678 | 81664 |
| NLK | nemo like kinase | *NLK* | -34.98 | | Nucleus | kinase | | | 51701 | 18099 | 497961 |
| ADCY8 | adenylate cyclase 8 | *ADCY8* | -34.766 | | Plasma Membrane | enzyme | | | 114 | 11514 | 29241 |

| | | | | | | | | | | | |
|---|---|---|---|---|---|---|---|---|---|---|---|
| TGFB1 | transforming growth factor beta 1 | *TGFB1* | -34.598 | | Extracellular Space | growth factor | diagnosis,efficacy,prognosis,safety,unspecified application | AVID200,dalantercept,fresolimumab,LY3200882,MSB0011359C,NIS793,YL-13027 | 7040 | 21803 | 59086 |
| ZBTB17 | zinc finger and BTB domain containing 17 | *ZBTB17* | -34.013 | | Nucleus | transcription regulator | | | 7709 | 22642 | 313666 |
| WNT7B | Wnt family member 7B | *WNT7B* | -33.954 | | Extracellular Space | other | | | 7477 | 22422 | 315196 |
| GNA14 | G protein subunit alpha 14 | *GNA14* | -33.289 | | Plasma Membrane | enzyme | | | 9630 | 14675 | 309242 |
| ADCY10 | adenylate cyclase 10 | *ADCY10* | -32.939 | | Cytoplasm | enzyme | | | 55811 | 271639 | 59320 |
| RALB | RAS like proto-oncogene B | *RALB* | -32.499 | | Cytoplasm | enzyme | | | 5899 | 64143 | 116546 |
| CASP9 | caspase 9 | *CASP9* | -32.26 | | Cytoplasm | peptidase | diagnosis,efficacy | caspase-9 inhibitor | 842 | 12371 | 58918 |
| WNT5B | Wnt family member 5B | *WNT5B* | -31.642 | | Extracellular Space | other | | | 81029 | 22419 | 282582 |
| BID | BH3 interacting domain death agonist | *BID* | -31.47 | | Cytoplasm | other | | | 637 | 12122 | 64625 |
| TAB2 | TGF-beta activated kinase 1 (MAP3K7) binding protein 2 | *TAB2* | -31.433 | | Cytoplasm | other | | | 23118 | 68652 | 308267 |
| ARHGEF1 | Rho guanine nucleotide exchange factor 1 | *ARHGEF1* | -31.052 | | Cytoplasm | other | unspecified application | | 9138 | 16801 | 60323 |
| WNT2B | Wnt family member 2B | *WNT2B* | -30.65 | | Extracellular Space | other | | | 7482 | 22414 | 116466 |
| CDK18 | cyclin dependent kinase 18 | *CDK18* | -30.483 | | Cytoplasm | kinase | | | 5129 | 18557 | 289019 |
| E2F2 | E2F transcription factor 2 | *E2F2* | -30.249 | | Nucleus | transcription regulator | | | 1870 | 242705 | 684111 |
| ARHGEF12 | Rho guanine nucleotide exchange factor 12 | *ARHGEF12* | -30.167 | | Cytoplasm | other | | | 23365 | 69632 | 367072 |
| TP53 | tumor protein p53 | *TP53* | -29.92 | | Nucleus | transcription regulator | diagnosis,disease progression,efficacy,prognosis,response to therapy,unspecified application | ALT-801,APR-246,azurin 50-77,BI 907828,cenersen,CGM097,COTI-2,kevetrin | 7157 | 22059 | 24842 |
| EP300 | E1A binding protein p300 | *EP300* | -29.364 | | Nucleus | transcription regulator | | CCS1477 | 2033 | 328572 | 170915 |
| RAP1B | RAP1B, member of RAS oncogene family | *RAP1B* | -29.292 | | Cytoplasm | enzyme | unspecified application | | 5908 | 215449 | 171337 |
| BCL2 | BCL2 apoptosis regulator | *BCL2* | -28.943 | | Cytoplasm | transporter | diagnosis,efficacy,prognosis,unspecified application | (-)-gossypol,5-azacytidine/venetoclax,afatinib/paclitaxel,APG-1252,APG-2575,BCL-2 blocker,BCL201,bortezomib/paclitaxel,chelerythrine,decitabine/venetoclax,doxorubicin/lapatinib/paclitaxel/trastuzumab,doxorubicin/paclitaxel,epirubicin/paclitaxel,everolimus/paclitaxel,gemcitabine/paclitaxel,lapatinib/paclitaxel,levodopa/rasagiline,navitoclax,obatoclax,obinutuzumab/venetoclax,oblimersen,paclitaxel,paclitaxel/pembrolizumab/ramucirumab,paclitaxel/pertuzumab/trastuzumab,paclitaxel/ramucirumab,paclitaxel/rituximab,paclitaxel/topot[illegible] | 596 | 12043 | 24224 |
| PAK2 | p21 (RAC1) activated kinase 2 | *PAK2* | -28.075 | | Cytoplasm | kinase | | | 5062 | 224105 | 100910732 |
| ITGA3 | integrin subunit alpha 3 | *ITGA3* | -27.914 | | Plasma Membrane | other | prognosis | | 3675 | 16400 | 360606 |
| RAC1 | Rac family small GTPase 1 | *RAC1* | -27.807 | | Plasma Membrane | enzyme | | | 5879 | 19353 | 363875 |
| MAP2K3 | mitogen-activated protein kinase kinase 3 | *MAP2K3* | -27.584 | | Cytoplasm | kinase | | | 5606 | 26397 | 303200 |
| PRKACA | protein kinase cAMP-activated catalytic subunit alpha | *PRKACA* | -27.162 | | Cytoplasm | kinase | | | 5566 | 18747 | 25636 |
| MAP2K2 | mitogen-activated protein kinase kinase 2 | *MAP2K2* | -27.026 | | Cytoplasm | kinase | | AS703988,binimetinib,binimetinib/cetuximab/encorafenib,binimetinib/encorafenib,binimetinib/encorafenib/panitumumab,cetuximab/dabrafenib/trametinib,cobimetinib,cobimetinib/vemurafenib,dabrafenib/ipilimumab/trametinib,dabrafenib/panitumumab/trametinib,dabrafenib/trametinib,dabrafenib/trametinib/vemurafenib,FCN-159,PD184352,pimasertib,selumetinib,TAK 733,trametinib,trametinib/vem | 5605 | 26396 | 58960 |
| JAK3 | Janus kinase 3 | *JAK3* | -26.841 | | Cytoplasm | kinase | efficacy | methotrexate/tofacitinib,PF-06651600,R-348,tofacitinib | 3718 | 16453 | 25326 |
| RBPJ | combination signal binding protein for immunoglobulin kappa J reg | *RBPJ* | -26.402 | | Nucleus | transcription regulator | | | 3516 | 19664 | 679028 |
| RALA | RAS like proto-oncogene A | *RALA* | -26.023 | | Cytoplasm | enzyme | | | 5898 | 56044 | 81757 |
| MAPK14 | mitogen-activated protein kinase 14 | *MAPK14* | -25.994 | | Cytoplasm | kinase | efficacy | p38 MAP kinase inhibitor,ralimetinib,RO-3201195,talmapimod | 1432 | 26416 | 81649 |
| PLCB2 | phospholipase C beta 2 | *PLCB2* | -25.707 | | Cytoplasm | enzyme | | | 5330 | 18796 | 85240 |
| PTCH2 | patched 2 | *PTCH2* | -25.653 | | Plasma Membrane | transmembrane receptor | | | 8643 | 19207 | |
| WNT4 | Wnt family member 4 | *WNT4* | -25.57 | | Extracellular Space | cytokine | | | 54361 | 22417 | 84426 |
| PTCH1 | patched 1 | *PTCH1* | -25.368 | | Plasma Membrane | transmembrane receptor | efficacy | | 5727 | 19206 | 89830 |
| APAF1 | apoptotic peptidase activating factor 1 | *APAF1* | -24.701 | | Cytoplasm | other | diagnosis | | 317 | 11783 | 78963 |
| BMPR1B | bone morphogenetic protein receptor type 1B | *BMPR1B* | -24.068 | | Plasma Membrane | kinase | unspecified application | | 658 | 12167 | |
| MAP3K5 | mitogen-activated protein kinase kinase kinase 5 | *MAP3K5* | -23.947 | | Cytoplasm | kinase | | selonsertib | 4217 | 26408 | 365057 |
| MAPK8 | mitogen-activated protein kinase 8 | *MAPK8* | -23.872 | | Cytoplasm | kinase | | aplidine,binimetinib/cetuximab/encorafenib,binimetinib/encorafenib,binimetinib/encorafenib/panitumumab,cetuximab/encorafenib,encorafenib | 5599 | 26419 | 116554 |
| PRKCG | protein kinase C gamma | *PRKCG* | -23.094 | | Cytoplasm | kinase | | ingenol mebutate | 5582 | 18752 | 24681 |
| E2F3 | E2F transcription factor 3 | *E2F3* | -23.017 | | Nucleus | transcription regulator | unspecified application | | 1871 | 13557 | 291105 |
| NAIP | NLR family apoptosis inhibitory protein | *NAIP* | -22.723 | | Cytoplasm | other | | | 4671 | 17948 | |
| PLCB1 | phospholipase C beta 1 | *PLCB1* | -22.627 | | Cytoplasm | enzyme | | | 23236 | 18795 | 24654 |
| ADCY6 | adenylate cyclase 6 | *ADCY6* | -22.619 | | Plasma Membrane | enzyme | efficacy | | 112 | 11512 | 25289 |
| SMAD7 | SMAD family member 7 | *SMAD7* | -22.453 | | Nucleus | transcription regulator | | | 4092 | 17131 | 81516 |
| MAP2K4 | mitogen-activated protein kinase kinase 4 | *MAP2K4* | -22.124 | | Cytoplasm | kinase | | binimetinib/cetuximab/encorafenib,binimetinib/encorafenib,binimetinib/encorafenib/panitumumab,cetuximab/encorafeni | 6416 | 26398 | 287398 |
| RHOBTB1 | Rho related BTB domain containing 1 | *RHOBTB1* | -22.104 | | Other | enzyme | | | 9886 | 69288 | 309722 |
| BMP8A | bone morphogenetic protein 8a | *BMP8A* | -21.949 | | Extracellular Space | cytokine | | | 353500 | 12163 | 680931 |
| TAB1 | TGF-beta activated kinase 1 (MAP3K7) binding protein 1 | *TAB1* | -21.517 | | Cytoplasm | enzyme | | | 10454 | 66513 | 315139 |
| PIK3R2 | phosphoinositide-3-kinase regulatory subunit 2 | *PIK3R2* | -21.309 | | Cytoplasm | kinase | | | 5296 | 18709 | 29741 |
| PSEN2 | presenilin 2 | *PSEN2* | -21.273 | | Cytoplasm | peptidase | diagnosis | tarenflurbil | 5664 | 19165 | 81751 |
| GNAI1 | G protein subunit alpha i1 | *GNAI1* | -21.245 | | Plasma Membrane | enzyme | | | 2770 | 14677 | 25686 |
| CDK7 | cyclin dependent kinase 7 | *CDK7* | -21.208 | | Nucleus | kinase | | alvocidib,BMS-387032,CT7001,roscovitine,SB-1317,SY-1365 | 1022 | 12572 | 171150 |
| CDC25C | cell division cycle 25C | *CDC25C* | -21.206 | | Nucleus | phosphatase | | | 995 | 12532 | 307511 |
| TCF3 | transcription factor 3 | *TCF3* | -20.783 | | Nucleus | transcription regulator | unspecified application | | 6929 | 21423 | 171046 |
| MAPK10 | mitogen-activated protein kinase 10 | *MAPK10* | -20.704 | | Cytoplasm | kinase | | binimetinib/cetuximab/encorafenib,binimetinib/encorafenib,binimetinib/encorafenib/panitumumab,cetuximab/encorafeni | 5602 | 26414 | 25272 |
| RASA1 | RAS p21 protein activator 1 | *RASA1* | -20.703 | | Cytoplasm | transporter | | | 5921 | 218397 | 25676 |
| CDK19 | cyclin dependent kinase 19 | *CDK19* | -20.614 | | Nucleus | kinase | | alvocidib,BCD-115,SEL120 | 23097 | 78334 | 309804 |
| TGFBR2 | transforming growth factor beta receptor 2 | *TGFBR2* | -20.319 | | Plasma Membrane | kinase | | IMC-TR1 | 7048 | 21813 | 81810 |
| APC | APC regulator of WNT signaling pathway | *APC* | -19.766 | | Nucleus | enzyme | diagnosis,disease progression,efficacy,prognosis,s | | 324 | 11789 | 24205 |
| FYN | FYN proto-oncogene, Src family tyrosine kinase | *FYN* | -19.656 | | Plasma Membrane | kinase | efficacy | blinatumomab/dasatinib,dasatinib,dasatinib/erlotinib,dasatinib/filgrastim,dasatinib/inotuzumab ozogamicin,dasatinib/nilotinib, | 2534 | 14360 | 25150 |
| BCL2L1 | BCL2 like 1 | *BCL2L1* | -19.527 | | Cytoplasm | other | efficacy,prognosis | | 598 | 12048 | 24888 |
| GNA11 | G protein subunit alpha 11 | *GNA11* | -18.879 | | Plasma Membrane | enzyme | | | 2767 | 14672 | 81662 |
| CDK11B | cyclin dependent kinase 11B | *CDK11B* | -18.143 | | Nucleus | kinase | | | 984 | | |
| RHOD | ras homolog family member D | *RHOD* | -17.716 | | Cytoplasm | enzyme | | | 29984 | 11854 | 293660 |
| BMP8B | bone morphogenetic protein 8b | *BMP8B* | -17.679 | | Extracellular Space | growth factor | | | 656 | | |
| BCL2L11 | BCL2 like 11 | *BCL2L11* | -17.562 | | Cytoplasm | other | efficacy | | 10018 | 12125 | 64547 |
| PSEN1 | presenilin 1 | *PSEN1* | -17.415 | | Plasma Membrane | peptidase | diagnosis,unspecified | tarenflurbil | 5663 | 19164 | 29192 |
| JAK2 | Janus kinase 2 | *JAK2* | -17.307 | | Cytoplasm | kinase | efficacy,prognosis,unspecified application | AZD1480,baricitinib,baricitinib/methotrexate,BMS-911543,entrectinib,erlotinib/ruxolitinib,fedratinib,gandotinib,jak2 inhibitor,jaktinib,methotrexate/tofacitinib,momelotinib,NS-018,pacritinib,ruxolitinib,SB-1317,tofacitinib,upadacitinib, | 3717 | 16452 | 24514 |
| FOXO1 | forkhead box O1 | *FOXO1* | -17.132 | | Nucleus | transcription regulator | | | 2308 | 56458 | 84482 |
| TYK2 | tyrosine kinase 2 | *TYK2* | -16.854 | | Plasma Membrane | kinase | | brepocitinib,ruxolitinib | 7297 | 54721 | 100361294 |

| | | | | | | | | | | | |
|---|---|---|---|---|---|---|---|---|---|---|---|
| TCF4 | transcription factor 4 | *TCF4* | -16.752 | | Nucleus | transcription regulator | | | 6925 | 21413 | 84382 |
| ATM | ATM serinethreonine kinase | *ATM* | -16.635 | | Nucleus | kinase | diagnosis | AZD0156,AZD1390,M3541 | 472 | 11920 | 300711 |
| CDC25A | cell division cycle 25A | *CDC25A* | -16.149 | | Nucleus | phosphatase | | | 993 | 12530 | 171102 |
| GNAI3 | G protein subunit alpha i3 | *GNAI3* | -16.035 | | Cytoplasm | enzyme | | | 2773 | 14679 | 25643 |
| ARHGEF16 | Rho guanine nucleotide exchange factor 16 | *ARHGEF16* | -15.972 | | Cytoplasm | other | | | 27237 | 230972 | 687105 |
| RRAS2 | RAS related 2 | *RRAS2* | -15.85 | | Plasma Membrane | enzyme | | | 22800 | 66922 | 365355 |
| WNT9A | Wnt family member 9A | *WNT9A* | -15.787 | | Extracellular Space | other | | | 7483 | 216795 | 287357 |
| RALBP1 | ralA binding protein 1 | *RALBP1* | -15.615 | | Cytoplasm | enzyme | | | 10928 | 19765 | 84014 |
| ITGA2 | integrin subunit alpha 2 | *ITGA2* | -15.488 | | Plasma Membrane | transmembrane receptor | | | 3673 | 16398 | 170921 |
| XIAP | X-linked inhibitor of apoptosis | *XIAP* | -15.279 | | Cytoplasm | enzyme | diagnosis,efficacy | ASTX660,BI-891065,birinapant,GDC-0152,GDC-0917,idronoxil,XIAP antisense oligonucleotide AEG35156 | 331 | 11798 | 63879 |
| RAC2 | Rac family small GTPase 2 | *RAC2* | -15.21 | | Cytoplasm | enzyme | | | 5880 | 19354 | 366957 |
| PRKCQ | protein kinase C theta | *PRKCQ* | -15.21 | | Cytoplasm | kinase | diagnosis | ingenol mebutate,sotrastaurin | 5588 | 18761 | 85420 |
| FZD3 | frizzled class receptor 3 | *FZD3* | -15.056 | | Plasma Membrane | G-protein coupled receptor | | | 7976 | 14365 | 266715 |
| RHOQ | ras homolog family member Q | *RHOQ* | -14.89 | | Plasma Membrane | enzyme | | | 23433 | 104215 | 85428 |
| WNT11 | Wnt family member 11 | *WNT11* | -14.587 | | Extracellular Space | other | | | 7481 | 22411 | 140584 |
| MDM2 | MDM2 proto-oncogene | *MDM2* | -14.33 | | Nucleus | transcription regulator | disease progression,efficacy,unspecified application | ALRN-6924,AMG-232,APG-115,ASTX295,BI 907828,CGM097,HDM201,idasanutlin,KRT-232,MI-773,milademetan,MK-8242,RO-5045337,RO6839921,serdem | 4193 | 17246 | 314856 |
| GNAS | GNAS complex locus | *GNAS* | -14.33 | | Plasma Membrane | enzyme | unspecified application | | 2778 | 14683 | 24896 |
| BIRC2 | baculoviral IAP repeat containing 2 | *BIRC2* | -14.043 | | Cytoplasm | enzyme | | ASTX660,BI-891065,birinapant,GDC-0152,GDC-0917 | 329 | 11797 | 60371 |
| ARHGEF19 | Rho guanine nucleotide exchange factor 19 | *ARHGEF19* | -13.97 | | Cytoplasm | other | | | 128272 | 213649 | 362648 |
| PRKD1 | protein kinase D1 | *PRKD1* | -13.884 | | Cytoplasm | kinase | | | 5587 | 18760 | 85421 |
| NFKBID | NFKB inhibitor delta | *NFKBID* | -13.719 | | Nucleus | transcription regulator | | | 84807 | 243910 | 308496 |
| KRAS | KRAS proto-oncogene, GTPase | *KRAS* | -13.663 | | Cytoplasm | enzyme | diagnosis,efficacy,prognosis,response to therapy,unspecified | AZD4785,chelerythrine,JNJ-74699157,LY3499446,MRTX8 | 3845 | 16653 | 24525 |
| ARHGEF9 | Cdc42 guanine nucleotide exchange factor 9 | *ARHGEF9* | -13.632 | | Cytoplasm | other | | | 23229 | 236915 | 66013 |
| RAPGEF3 | Rap guanine nucleotide exchange factor 3 | *RAPGEF3* | -13.361 | | Nucleus | other | | | 10411 | 223864 | 59326 |
| BMP5 | bone morphogenetic protein 5 | *BMP5* | -13.257 | | Extracellular Space | growth factor | | | 653 | 12160 | 315824 |
| CASP8 | caspase 8 | *CASP8* | -13.228 | | Nucleus | peptidase | diagnosis,efficacy,prognosis | | 841 | 12370 | 64044 |
| LRP6 | LDL receptor related protein 6 | *LRP6* | -13.208 | | Plasma Membrane | transmembrane receptor | | | 4040 | 16974 | 312781 |
| PLCB3 | phospholipase C beta 3 | *PLCB3* | -13.203 | | Cytoplasm | enzyme | | | 5331 | 18797 | 29322 |
| MAPK13 | mitogen-activated protein kinase 13 | *MAPK13* | -12.894 | | Cytoplasm | kinase | | talmapimod | 5603 | 26415 | 29513 |
| CDC42 | cell division cycle 42 | *CDC42* | -12.835 | | Cytoplasm | enzyme | | | 998 | | |
| SMO | smoothened, frizzled class receptor | *SMO* | -12.812 | | Plasma Membrane | G-protein coupled receptor | response to therapy | BMS-833923,erismodegib,erlotinib/vismodegib,gemcitabine/vismodegib,glasdegib,NVP-LEQ-506,saridegib,TAK-441,taladegib,vismodegib,ZSP1602 | 6608 | 319757 | 25273 |
| E2F4 | E2F transcription factor 4 | *E2F4* | -12.662 | | Nucleus | transcription regulator | | | 1874 | 104394 | 100360427 |
| PIK3R4 | phosphoinositide-3-kinase regulatory subunit 4 | *PIK3R4* | -12.655 | | Cytoplasm | kinase | | | 30849 | 75669 | 363131 |
| CDK16 | cyclin dependent kinase 16 | *CDK16* | -12.328 | | Cytoplasm | kinase | | | 5127 | 18555 | 81741 |
| ARHGEF15 | Rho guanine nucleotide exchange factor 15 | *ARHGEF15* | -12.287 | | Cytoplasm | other | | | 22899 | 442801 | 287418 |
| WNT10A | Wnt family member 10A | *WNT10A* | -12.181 | | Extracellular Space | other | | | 80326 | 22409 | 316527 |
| E2F1 | E2F transcription factor 1 | *E2F1* | -12.134 | | Nucleus | transcription regulator | disease progression,prognosis | | 1869 | 13555 | 399489 |

| Supplementary Table 12. Diseases and functions with which cisplatin damaged genes are associated. | |
|---|---|
| **Category** | **p-value** |
| Cancer | 8.04E-313 |
| Organismal Injury and Abnormalities | 8.04E-313 |
| Endocrine System Disorders | 8.04E-313 |
| Gastrointestinal Disease | 8.04E-313 |
| Dermatological Diseases and Conditions | 2.32E-222 |
| Reproductive System Disease | 8.93E-162 |
| Hepatic System Disease | 1.86E-108 |
| Renal and Urological Disease | 2.03E-45 |
| Hematological Disease | 1.65E-41 |
| Respiratory Disease | 5.81E-36 |
| Immunological Disease | 7.49E-32 |
| Neurological Disease | 1.69E-29 |
| Cellular Assembly and Organization | 1.94E-29 |
| Cellular Function and Maintenance | 1.94E-29 |
| Hereditary Disorder | 3.05E-29 |
| Molecular Transport | 2.99E-22 |
| Developmental Disorder | 1.49E-19 |
| Cell Morphology | 2.53E-18 |
| Post-Translational Modification | 1.64E-16 |
| Gene Expression | 6.34E-16 |
| Cardiovascular Disease | 4.83E-15 |
| Tissue Development | 4.23E-13 |
| Connective Tissue Disorders | 4.46E-13 |
| Cell Signaling | 4.78E-13 |
| Protein Synthesis | 1.51E-12 |
| Psychological Disorders | 1.61E-12 |
| Skeletal and Muscular Disorders | 1.28E-11 |
| Infectious Diseases | 9.78E-11 |
| Cellular Movement | 5.07E-10 |
| Nervous System Development and Function | 5.07E-10 |
| Cell Death and Survival | 2.45E-09 |
| Cell-To-Cell Signaling and Interaction | 3.53E-09 |
| Cellular Development | 5.16E-09 |
| Cellular Growth and Proliferation | 5.16E-09 |
| Organismal Development | 5.16E-09 |
| Carbohydrate Metabolism | 2.08E-08 |
| Metabolic Disease | 2.11E-08 |
| Hair and Skin Development and Function | 5.25E-08 |
| Organismal Survival | 6.59E-08 |
| Inflammatory Disease | 6.83E-08 |
| Ophthalmic Disease | 8.66E-08 |
| Cell Cycle | 2.73E-07 |
| DNA Replication, Recombination, and Repair | 2.73E-07 |
| Lipid Metabolism | 3.95E-07 |
| Small Molecule Biochemistry | 3.95E-07 |

| Supplementary Table 13. Cancers and functions with which cisplatin damaged genes are associated. | | | | | |
|---|---|---|---|---|---|
| Diseases or Functions | p-value | Predicted Activation State | Activation z-score | # Molecules | *SPAG9* |
| Cell transformation | 9.46E-09 | Decreased | -3.565 | 82 | |
| Cancer | 8.85E-154 | Decreased | -3.149 | 6981 | YES |
| Neoplasia of cells | 5.43E-81 | Decreased | -2.475 | 3247 | YES |
| Cancer of cells | 2.27E-74 | Decreased | -2.102 | 2855 | YES |
| Hematopoietic neoplasm | 3.62E-41 | | -1.907 | 1820 | YES |
| Digestive organ tumor | 4.79E-206 | | -1.72 | 6357 | YES |
| Lymphohematopoietic neoplasia | 3.00E-41 | | -1.568 | 1827 | YES |
| Digestive system cancer | 1.14E-211 | | -1.517 | 6292 | YES |
| Carcinoma | 5.73E-218 | | -1.448 | 6901 | YES |
| Tumorigenesis of tissue | 2.89E-212 | | -1.401 | 6911 | YES |
| Gastrointestinal tract cancer | 2.40E-235 | | -1.332 | 5958 | YES |
| Solid tumor | 2.77E-148 | | -1.274 | 6984 | YES |
| Squamous-cell carcinoma | 5.90E-68 | | -1.172 | 2265 | |
| Acute leukemia | 4.46E-28 | | -1.154 | 954 | |
| Head and neck tumor | 0.00E+00 | | -1.109 | 6190 | YES |
| Extracranial solid tumor | 1.16E-155 | | -1.103 | 6968 | YES |
| Cancer of secretory structure | 5.40E-310 | | -1.091 | 6311 | YES |
| Epithelial neoplasm | 3.79E-217 | | -1.087 | 6905 | YES |
| Malignant solid tumor | 8.81E-158 | | -1.084 | 6976 | YES |
| Non-melanoma solid tumor | 2.10E-205 | | -1.077 | 6920 | YES |
| Gastrointestinal tumor | 1.70E-236 | | -1 | 5975 | YES |
| Extrapulmonary squamous cell carcinoma | 1.33E-72 | | -0.849 | 2149 | |
| Breast or colorectal cancer | 1.78E-111 | | -0.83 | 3945 | YES |
| Leukemia | 7.49E-32 | | -0.812 | 1208 | YES |
| Head and neck cancer | 0.00E+00 | | -0.762 | 6159 | YES |
| Head and neck carcinoma | 0.00E+00 | | -0.762 | 6149 | YES |
| Malignant neoplasm of large intestine | 3.00E-249 | | -0.747 | 5789 | YES |
| Intestinal cancer | 7.87E-249 | | -0.747 | 5790 | YES |
| Colorectal cancer | 7.96E-103 | | -0.747 | 3398 | YES |
| Nonhematologic malignant neoplasm | 3.99E-198 | | -0.736 | 6945 | YES |
| Non-hematological solid tumor | 3.79E-193 | | -0.713 | 6953 | YES |
| Breast or pancreatic cancer | 2.78E-122 | | -0.655 | 3241 | YES |
| Anogenital cancer | 5.80E-161 | | -0.594 | 4547 | YES |
| Mammary tumor | 5.05E-30 | | -0.58 | 1556 | YES |
| Abdominal cancer | 1.06E-205 | | -0.558 | 6569 | YES |
| Intrathoracic malignant tumor | 3.35E-22 | | -0.557 | 1493 | |
| Lung cancer | 3.67E-22 | | -0.557 | 1487 | |
| Respiratory system tumor | 4.84E-35 | | -0.466 | 1864 | |
| Liquid tumor | 2.37E-30 | | -0.454 | 1276 | YES |
| Breast cancer | 1.43E-34 | | -0.431 | 1532 | YES |
| Large intestine neoplasm | 2.24E-250 | | -0.394 | 5806 | YES |
| Intestinal tumor | 3.88E-250 | | -0.394 | 5808 | YES |
| Colorectal tumor | 2.63E-104 | | -0.394 | 3445 | YES |
| Breast or gynecological cancer | 4.02E-103 | | -0.311 | 3491 | YES |
| Breast or ovarian cancer | 1.77E-74 | | -0.311 | 2376 | YES |
| Hematologic cancer | 5.97E-41 | | -0.308 | 1796 | YES |
| Development of malignant tumor | 7.57E-194 | | -0.307 | 4858 | YES |
| Lymphoreticular neoplasm | 5.05E-37 | | -0.307 | 1554 | |
| Urogenital cancer | 5.12E-163 | | -0.222 | 4859 | YES |
| Malignant genitourinary solid tumor | 8.76E-163 | | -0.222 | 4858 | YES |
| Lung carcinoma | 1.01E-22 | | -0.2 | 1420 | |
| Central nervous system tumor | 1.69E-29 | | -0.185 | 904 | |
| Nervous system neoplasm | 1.95E-29 | | -0.185 | 966 | |
| Central nervous system solid tumor | 3.25E-29 | | -0.185 | 902 | |
| Genitourinary tumor | 6.46E-155 | | -0.172 | 4896 | YES |
| Lymphocytic leukemia | 1.05E-13 | | -0.152 | 496 | YES |
| Extraadrenal retroperitoneal tumor | 1.18E-92 | | -0.137 | 2303 | |
| Malignant neoplasm of retroperitoneum | 1.37E-86 | | -0.137 | 2144 | |
| Abdominal neoplasm | 1.13E-201 | | -0.135 | 6617 | YES |
| Frequency of tumor | 4.91E-194 | | -0.126 | 4896 | YES |
| Incidence of tumor | 1.67E-182 | | -0.126 | 4985 | YES |

| | | | | | |
|---|---|---|---|---|---|
| Thoracic neoplasm | 2.78E-23 | | -0.084 | 1520 | |
| Lung tumor | 2.87E-23 | | -0.084 | 1513 | |
| Myeloid or lymphoid neoplasm | 1.65E-41 | | -0.027 | 1808 | YES |
| Neoplasia of blood cells | 2.98E-41 | | -0.027 | 1806 | YES |
| Neuroepithelial tumor | 2.29E-26 | | 0.033 | 787 | |
| Development of neuroepithelial tumor | 3.18E-26 | | 0.033 | 786 | |
| Glioma | 3.77E-26 | | 0.033 | 782 | |
| Lymphohematopoietic cancer | 4.46E-41 | | 0.061 | 1804 | YES |
| Gastrointestinal carcinoma | 3.69E-288 | | 0.159 | 5876 | YES |
| Urinary tract cancer | 2.03E-45 | | 0.333 | 1478 | |
| Urinary tract tumor | 7.38E-43 | | 0.333 | 1496 | |
| Renal tumor | 4.00E-21 | | 0.333 | 826 | |
| Renal cancer | 1.11E-20 | | 0.333 | 809 | |
| Neoplasia of leukocytes | 6.43E-28 | | 0.342 | 1179 | YES |
| Lymphocytic cancer | 6.54E-28 | | 0.342 | 1175 | YES |
| Lymphocytic neoplasm | 8.61E-28 | | 0.342 | 1176 | YES |
| Hematologic cancer of cells | 4.33E-27 | | 0.342 | 1117 | YES |
| Melanoma | 7.90E-231 | | 0.386 | 4984 | YES |
| Lymphatic system tumor | 7.78E-28 | | 0.396 | 1198 | YES |
| Abdominal carcinoma | 1.15E-252 | | 0.434 | 6503 | YES |
| Development of carcinoma | 8.79E-195 | | 0.659 | 4829 | YES |
| Lymphoid cancer | 3.25E-28 | | 0.726 | 1186 | YES |
| Tumorigenesis of epithelial neoplasm | 8.81E-194 | | 0.968 | 4863 | YES |
| Pelvic tumor | 1.25E-167 | | 1 | 4461 | YES |
| Genital tumor | 1.28E-161 | | 1 | 4197 | YES |
| Tumorigenesis of lymphocytes | 1.06E-23 | | 1.067 | 1081 | YES |
| Lymphoma | 5.32E-19 | | 1.067 | 819 | |
| Nonpituitary endocrine tumor | 0.00E+00 | | | 6007 | YES |
| Endocrine cancer | 0.00E+00 | | | 5983 | YES |
| Endocrine carcinoma | 0.00E+00 | | | 5978 | YES |
| Thyroid carcinoma | 0.00E+00 | | | 5971 | YES |
| Thyroid gland tumor | 0.00E+00 | | | 5983 | YES |
| Thyroid cancer | 0.00E+00 | | | 5975 | YES |
| Neck neoplasm | 0.00E+00 | | | 6023 | YES |
| Endocrine gland tumor | 0.00E+00 | | | 6013 | YES |
| Large intestine carcinoma | 1.45E-307 | | | 5697 | YES |
| Intestinal carcinoma | 1.99E-307 | | | 5698 | YES |
| Gastrointestinal adenocarcinoma | 1.43E-302 | | | 5757 | YES |
| Abdominal adenocarcinoma | 2.33E-285 | | | 6285 | YES |
| Adenocarcinoma | 1.96E-269 | | | 6351 | YES |
| Skin cancer | 2.32E-222 | | | 4559 | YES |
| Skin tumor | 3.67E-222 | | | 4561 | YES |
| Malignant solid organ tumor | 3.70E-222 | | | 4612 | YES |
| Cutaneous melanoma | 1.29E-213 | | | 4364 | YES |
| Pelvic adenocarcinoma | 4.00E-190 | | | 3725 | YES |
| Genitourinary adenocarcinoma | 1.60E-182 | | | 4005 | YES |
| Genitourinary carcinoma | 1.06E-175 | | | 4392 | YES |
| Pelvic cancer | 2.42E-169 | | | 4415 | YES |
| Genital tract cancer | 8.93E-162 | | | 4145 | YES |
| Colorectal adenocarcinoma | 1.64E-154 | | | 3115 | YES |
| Development of digestive organ tumor | 4.28E-152 | | | 3464 | YES |
| Development of colorectal tumor | 1.81E-151 | | | 3172 | YES |
| Colorectal carcinoma | 2.18E-151 | | | 3171 | YES |
| Prostatic adenocarcinoma | 3.33E-151 | | | 2362 | YES |
| Prostatic carcinoma | 4.00E-150 | | | 2439 | YES |
| Colon adenocarcinoma | 4.43E-148 | | | 3001 | YES |
| Colon carcinoma | 2.37E-145 | | | 3028 | YES |
| Colon tumor | 5.34E-139 | | | 3149 | YES |
| Colon cancer | 1.66E-136 | | | 3120 | YES |
| Prostate cancer | 1.24E-135 | | | 2688 | YES |
| Malignant neoplasm of male genital organ | 2.06E-135 | | | 2712 | YES |
| Prostatic tumor | 1.58E-134 | | | 2690 | YES |
| Male genital neoplasm | 3.50E-134 | | | 2716 | YES |

| | | | | | |
|---|---|---|---|---|---|
| Hepatobiliary carcinoma | 4.16E-110 | | | 3048 | YES |
| Hepatobiliary system cancer | 1.86E-108 | | | 3094 | YES |
| Liver tumor | 6.80E-108 | | | 3351 | YES |
| Liver carcinoma | 6.09E-101 | | | 2872 | YES |
| Liver cancer | 2.82E-99 | | | 2920 | YES |
| Breast or ovarian carcinoma | 1.46E-88 | | | 2057 | YES |
| Non-colon gastrointestinal cancer | 1.20E-87 | | | 2248 | YES |
| Upper gastrointestinal carcinoma | 6.92E-87 | | | 1995 | YES |
| Upper gastrointestinal tract cancer | 4.63E-78 | | | 2050 | YES |
| Upper gastrointestinal tract tumor | 3.28E-77 | | | 2053 | YES |
| Gastroesophageal adenocarcinoma | 4.68E-53 | | | 1126 | YES |
| Gastro-esophageal carcinoma | 2.16E-49 | | | 1229 | YES |
| Gastric adenocarcinoma | 1.93E-42 | | | 918 | YES |
| Breast carcinoma | 3.79E-41 | | | 1189 | YES |
| Gastroesophageal cancer | 4.13E-41 | | | 1287 | YES |
| Gastric carcinoma | 6.34E-40 | | | 947 | YES |
| Gastric cancer | 6.44E-31 | | | 1004 | YES |
| Stomach tumor | 4.13E-30 | | | 1007 | YES |
| B-cell neoplasm | 6.25E-28 | | | 925 | YES |
| Mature B-cell neoplasm | 6.35E-25 | | | 857 | YES |
| Rectum tumor | 7.79E-21 | | | 404 | YES |
| Rectal adenocarcinoma | 1.38E-20 | | | 382 | YES |
| Distal colorectal carcinoma | 2.10E-20 | | | 384 | YES |
| Rectum cancer | 2.27E-20 | | | 399 | YES |
| Rectum carcinoma | 2.31E-20 | | | 383 | YES |
| Connective tissue tumor | 2.86E-20 | | | 811 | YES |
| Soft tissue neoplasm | 4.04E-20 | | | 686 | YES |
| Connective or soft tissue tumor | 5.90E-20 | | | 934 | YES |
| Mature lymphocytic neoplasm | 1.14E-19 | | | 981 | YES |
| Sarcoma | 5.91E-19 | | | 760 | YES |
| Malignant connective or soft tissue neoplasm | 3.08E-18 | | | 768 | YES |
| Malignant soft tissue neoplasm | 8.98E-18 | | | 654 | YES |
| Soft tissue sarcoma | 1.20E-17 | | | 653 | YES |
| Blue round small cell tumor | 5.90E-15 | | | 401 | YES |
| Small blue round cell sarcoma | 1.16E-11 | | | 260 | YES |
| Rhabdomyosarcoma | 1.28E-11 | | | 259 | YES |
| Myosarcoma | 7.87E-11 | | | 274 | YES |
| Chronic leukemia | 2.08E-10 | | | 357 | YES |
| Tumor in striated muscle | 2.24E-10 | | | 231 | YES |
| Rhabdomyosarcoma in striated muscle | 3.48E-10 | | | 229 | YES |
| Embryonal rhabdomyosarcoma | 1.00E-09 | | | 231 | YES |
| Embryonal rhabdomyosarcoma in striated | 2.31E-09 | | | 214 | YES |
| Muscle tumor | 6.78E-09 | | | 401 | YES |
| Chronic form of lymphocytic leukemia | 4.43E-07 | | | 283 | YES |
| Hairy cell leukemia | 4.86E-07 | | | 169 | YES |
| Large intestine adenocarcinoma | 1.25E-312 | | | 5675 | YES |
| Intestinal adenocarcinoma | 8.04E-313 | | | 5676 | YES |
| Development of adenocarcinoma | 1.02E-129 | | | 3208 | |
| Female genital tract adenocarcinoma | 1.11E-107 | | | 2439 | |
| Pancreatobiliary tumor | 8.07E-103 | | | 2090 | |
| Endometrioid carcinoma | 1.31E-102 | | | 2080 | |
| Development of genital tumor | 2.16E-94 | | | 2793 | |
| Tumorigenesis of reproductive tract | 3.03E-92 | | | 2879 | |
| Female genital tract cancer | 3.15E-92 | | | 2808 | |
| Female genital neoplasm | 4.85E-92 | | | 2884 | |
| Ductal carcinoma | 7.15E-92 | | | 1864 | |
| Pancreatic carcinoma | 6.03E-90 | | | 1577 | |
| Pancreatic ductal adenocarcinoma | 2.32E-87 | | | 1471 | |
| Pancreatic tumor | 7.67E-87 | | | 1809 | |
| Pancreatic adenocarcinoma | 1.80E-84 | | | 1492 | |
| Pancreatic cancer | 4.86E-81 | | | 1637 | |
| Endometrial carcinoma | 3.21E-77 | | | 1884 | |
| Uterine endometrioid carcinoma | 2.92E-76 | | | 1617 | |

| | | | | |
|---|---|---|---|---|
| Endometrioid endometrial adenocarcinoma | 2.24E-74 | | | 1587 |
| Endometrial adenocarcinoma | 2.19E-73 | | | 1655 |
| Ovarian carcinoma | 4.55E-70 | | | 1233 |
| Endometrium tumor | 2.50E-69 | | | 1972 |
| Endometrial cancer | 6.04E-69 | | | 1967 |
| Upper aerodigestive tract carcinoma | 1.30E-68 | | | 1564 |
| Head and neck squamous cell carcinoma | 4.15E-68 | | | 1753 |
| Ovarian adenocarcinoma | 5.61E-68 | | | 1173 |
| Malignant neoplasm of aerodigestive tract | 9.85E-67 | | | 1588 |
| Uterine carcinoma | 3.84E-65 | | | 1948 |
| Ovarian cancer | 2.49E-60 | | | 1316 |
| Gonadal tumor | 1.77E-59 | | | 1365 |
| Ovarian tumor | 8.95E-59 | | | 1327 |
| Uterine cancer | 2.74E-58 | | | 2060 |
| Uterine tumor | 1.40E-57 | | | 2139 |
| Upper aero-digestive squamous cell carcinoma | 6.43E-53 | | | 1234 |
| Mixed neoplasia | 2.34E-52 | | | 1077 |
| Cancer of head | 1.39E-46 | | | 969 |
| Ovarian endometrioid carcinoma | 1.52E-45 | | | 724 |
| Oral cavity carcinoma | 5.26E-45 | | | 861 |
| Oral squamous cell carcinoma | 4.75E-44 | | | 831 |
| Adenosquamous ovarian carcinoma | 5.53E-43 | | | 689 |
| Mixed ovarian carcinoma | 6.57E-43 | | | 705 |
| Adenosquamous carcinoma | 6.90E-43 | | | 708 |
| Oral cancer | 7.49E-43 | | | 869 |
| Oral tumor | 8.17E-43 | | | 871 |
| Mixed cell carcinoma | 1.29E-41 | | | 726 |
| Skin carcinoma | 3.85E-40 | | | 921 |
| Bladder carcinoma | 4.00E-39 | | | 870 |
| Bladder cancer | 2.63E-36 | | | 918 |
| Lung adenocarcinoma | 5.81E-36 | | | 1112 |
| Nonsquamous non-small cell lung carcinoma | 1.86E-35 | | | 1116 |
| Bladder tumor | 1.29E-32 | | | 922 |
| Serous neoplasm | 5.27E-30 | | | 677 |
| Serous adenocarcinoma | 1.15E-29 | | | 672 |
| Ovarian serous tumor | 2.93E-29 | | | 552 |
| Myeloid neoplasm | 8.45E-28 | | | 1006 |
| Myeloid leukemia | 1.03E-27 | | | 894 |
| Non-melanoma skin cancer | 2.11E-27 | | | 575 |
| Bone marrow neoplasm | 2.52E-27 | | | 997 |
| Skin squamous cell carcinoma | 3.86E-27 | | | 566 |
| Central nervous system cancer | 4.28E-27 | | | 790 |
| Acute myeloid leukemia | 4.63E-27 | | | 868 |
| Malignant myeloid neoplasm | 6.39E-27 | | | 969 |
| Non-small cell lung carcinoma | 8.86E-27 | | | 1315 |
| Bone marrow cancer | 1.78E-26 | | | 973 |
| Serous adenocarcinoma in ovary | 4.65E-26 | | | 438 |
| B-cell lymphoma | 1.13E-24 | | | 615 |
| Glioma cancer | 1.19E-24 | | | 684 |
| B cell cancer | 1.57E-24 | | | 798 |
| Brain tumor | 7.23E-24 | | | 609 |
| Intracranial neoplasm | 7.62E-24 | | | 611 |
| B-cell non-Hodgkin lymphoma | 7.71E-24 | | | 593 |
| Renal clear cell adenocarcinoma | 9.27E-24 | | | 609 |
| Clear-cell adenocarcinoma | 9.67E-24 | | | 636 |
| Biliary tract cancer | 1.17E-23 | | | 475 |
| Brain glioma | 2.50E-23 | | | 511 |
| Grade 3-4 glioma cancer | 3.36E-23 | | | 482 |
| Clear cell neoplasm | 4.25E-23 | | | 638 |
| Biliary tract carcinoma | 4.52E-23 | | | 467 |
| Adenocarcinoma in right upper lobe of lung | 5.70E-23 | | | 340 |
| Grade 3-4 glioma | 5.88E-23 | | | 485 |
| Mature B cell malignant tumor | 1.42E-22 | | | 673 |

| | | | | |
|---|---|---|---|---|
| Grade 3-4 tumor | 3.08E-22 | | | 513 |
| Astrocytoma | 1.87E-21 | | | 651 |
| Upper airway cancer | 4.24E-21 | | | 598 |
| High grade glioma | 5.32E-21 | | | 624 |
| High grade astrocytoma | 7.33E-21 | | | 623 |
| Diffuse lymphoma | 1.00E-20 | | | 488 |
| Bile duct cancer | 2.70E-20 | | | 436 |
| Kidney adenocarcinoma | 3.34E-20 | | | 733 |
| Bile duct carcinoma | 3.61E-20 | | | 433 |
| Renal cell carcinoma | 3.67E-20 | | | 732 |
| Kidney carcinoma | 2.40E-19 | | | 737 |
| Pancreatic carcinoid tumor | 2.75E-19 | | | 266 |
| Carcinoid tumor | 1.30E-18 | | | 282 |
| Laryngeal cancer | 2.22E-18 | | | 387 |
| Diffuse B-cell lymphoma | 2.53E-18 | | | 444 |
| Laryngeal squamous cell carcinoma | 5.71E-18 | | | 380 |
| Abdominal neuroendocrine tumor | 6.87E-18 | | | 322 |
| Grade 1-4 astrocytoma | 8.79E-18 | | | 431 |
| Cecum adenocarcinoma | 1.02E-17 | | | 371 |
| Pancreatic endocrine tumor | 1.10E-17 | | | 285 |
| Non-Hodgkin lymphoma | 1.72E-17 | | | 747 |
| Grade 4 high grade glioma | 2.26E-17 | | | 405 |
| Grade 4 malignant glioma | 2.26E-17 | | | 405 |
| Grade 4 astrocytoma | 2.26E-17 | | | 405 |
| Extrapancreatic neuroendocrine tumor | 2.44E-17 | | | 415 |
| Neuroendocrine gastroenteropancreatic tumor | 3.46E-17 | | | 287 |
| Esophageal carcinoma | 3.68E-17 | | | 408 |
| Esophageal cancer | 7.08E-17 | | | 430 |
| Esophageal adenocarcinoma | 1.75E-16 | | | 297 |
| Brain astrocytoma | 4.49E-16 | | | 408 |
| Low-grade lymphoma | 6.00E-16 | | | 384 |
| Diffuse small-cell lymphoma | 1.03E-15 | | | 373 |
| Basal cell carcinoma | 1.38E-14 | | | 414 |
| Neuroendocrine tumor | 2.92E-14 | | | 425 |
| Mature B-cell lymphoma | 5.74E-14 | | | 341 |
| Oligodendroglioma | 3.03E-13 | | | 140 |
| HER2 expressing carcinoma | 3.23E-13 | | | 139 |
| HER2 positive carcinoma | 3.43E-13 | | | 138 |
| Indolent non-Hodgkin lymphoma | 3.81E-13 | | | 291 |
| Small B-cell lymphocytic lymphoma | 3.99E-13 | | | 282 |
| HER2 expressing cancer | 4.58E-13 | | | 147 |
| HER2 expressing solid tumor | 4.58E-13 | | | 147 |
| Low-grade B-cell lymphoma | 6.09E-13 | | | 286 |
| HER2 expressing solid malignant tumor | 7.51E-13 | | | 146 |
| HER2 positive solid malignant tumor | 8.06E-13 | | | 145 |
| Embryonal tumor | 9.91E-13 | | | 412 |
| Indolent B-cell non-Hodgkin lymphoma | 1.36E-12 | | | 283 |
| Carcinosarcoma | 2.49E-12 | | | 386 |
| Soft tissue sarcoma of trunk | 8.15E-12 | | | 391 |
| Malignant neoplasm of connective or soft tissue of abdomen | 8.97E-12 | | | 401 |
| High grade lymphocytic cancer | 1.16E-11 | | | 243 |
| Sarcoma of the female genital tract | 1.19E-11 | | | 389 |
| Grade 3 malignant glioma | 1.40E-11 | | | 105 |
| Chronic lymphocytic leukemia / small lymphocytic lymphoma | 1.64E-11 | | | 381 |
| Uterine sarcoma | 1.68E-11 | | | 383 |
| Mucosal melanoma | 2.01E-11 | | | 96 |
| High-grade lymphoma | 2.12E-11 | | | 248 |
| Chronic lymphocytic leukemia or small lymphocytic lymphoma or prolymphocytic leukemia | 2.12E-11 | | | 387 |

| | | | | |
|---|---|---|---|---|
| Grade 3 glioma | 3.18E-11 | | | 108 |
| Brain oligodendroglioma | 3.53E-11 | | | 95 |
| Sun-shielded melanoma | 4.78E-11 | | | 100 |
| Grade 2-3 glioma | 5.57E-11 | | | 132 |
| Blastoma | 6.02E-11 | | | 287 |
| Malignant mixed Mullerian tumor | 6.78E-11 | | | 374 |
| Pelvic carcinosarcoma | 9.00E-11 | | | 373 |
| Aggressive mature B-cell lymphoma | 9.53E-11 | | | 229 |
| Aggressive non-Hodgkin lymphoma | 1.47E-10 | | | 230 |
| Uterine carcinosarcoma | 2.25E-10 | | | 367 |
| HER2-positive carcinoma of breast | 2.54E-10 | | | 117 |
| Malignant mixed Mullerian tumor in endometrium | 2.97E-10 | | | 353 |
| Glioblastoma | 3.12E-10 | | | 279 |
| Aggressive cancer | 3.85E-10 | | | 236 |
| Acute lymphoblastic leukemia | 3.88E-10 | | | 211 |
| Anaplastic glioma | 3.99E-10 | | | 286 |
| Endometrial mixed tumor | 4.66E-10 | | | 363 |
| HER2-positive breast cancer | 4.96E-10 | | | 124 |
| Non-M3 acute myeloid leukemia | 7.75E-10 | | | 93 |
| Endometrial carcinosarcoma | 8.80E-10 | | | 360 |
| Myeloproliferative neoplasm | 8.83E-10 | | | 178 |
| Transitional-cell carcinoma | 1.01E-09 | | | 212 |
| Urothelial transitional cell carcinoma | 1.05E-09 | | | 206 |
| Endometrial sarcoma | 1.16E-09 | | | 361 |
| Urothelial neoplasm | 1.41E-09 | | | 222 |
| Cardiovascular neoplasm | 2.09E-09 | | | 129 |
| Urothelial cancer | 2.17E-09 | | | 220 |
| Vascular tumor | 2.28E-09 | | | 128 |
| T-cell leukemia | 4.74E-09 | | | 184 |
| Urothelial bladder carcinoma | 6.16E-09 | | | 198 |
| Angioma | 9.50E-09 | | | 101 |
| Hemangioma | 1.10E-08 | | | 99 |
| Primitive neuroectodermal tumor | 1.46E-08 | | | 144 |
| Fibrous tissue tumor | 1.50E-08 | | | 89 |
| Benign Tumors | 1.68E-08 | | | 432 |
| Neuroectodermal tumor | 2.02E-08 | | | 147 |
| Development of cardiovascular neoplasm | 2.10E-08 | | | 64 |
| Capillary hemangioma | 2.29E-08 | | | 60 |
| Benign solid tumor | 2.34E-08 | | | 426 |
| Development of angioma | 3.74E-08 | | | 63 |
| Small intestine cancer | 4.14E-08 | | | 59 |
| Unresectable cutaneous melanoma | 4.37E-08 | | | 53 |
| Mesenchymal tumor | 4.54E-08 | | | 96 |
| Chronic myeloproliferative neoplasm | 4.81E-08 | | | 147 |
| Brain cancer | 8.39E-08 | | | 144 |
| B-lymphoblastic leukemia/lymphoma | 1.34E-07 | | | 116 |
| T-lymphoblastic leukemia/lymphoma | 1.80E-07 | | | 123 |
| Acinar lung adenocarcinoma | 2.47E-07 | | | 66 |
| Occult primary tumor | 2.48E-07 | | | 79 |
| Large-cell lymphoma | 2.53E-07 | | | 171 |
| Gastrointestinal stromal tumor | 2.79E-07 | | | 46 |
| Female genital tract serous carcinoma | 2.84E-07 | | | 253 |
| Acinar-cell carcinoma | 2.95E-07 | | | 82 |

| Supplementary Table 14. Toxicity or Functions with which cisplatin damaged genes are associated. | |
|---|---|
| **Category** | **p-value** |
| Liver Hyperplasia/Hyperproliferation | 6.80E-108 |
| Cardiac Arteriopathy | 4.83E-15 |
| Heart Failure | 6.59E-08 |
| Cardiac Arrythmia | 1.76E-07 |
| Nephrosis | 1.12E-06 |
| Renal Proliferation | 3.86E-06 |
| Hepatocellular carcinoma | 6.66E-06 |
| Tachycardia | 2.71E-05 |
| Cardiac Dysfunction | 4.76E-05 |
| Congenital Heart Anomaly | 2.40E-04 |
| Renal Necrosis/Cell Death | 2.55E-04 |
| Glomerular Injury | 5.23E-04 |
| Cardiac Dilation | 9.74E-04 |
| Cardiac Enlargement | 9.74E-04 |
| Cardiac Congestive Cardiac Failure | 1.73E-03 |
| Cardiac Infarction | 7.16E-03 |
| Liver Inflammation/Hepatitis | 1.38E-02 |
| Kidney Failure | 1.59E-02 |
| Liver Damage | 2.58E-02 |
| Renal Inflammation | 2.58E-02 |
| Renal Nephritis | 2.58E-02 |
| Renal Dysplasia | 4.30E-02 |
| Cardiac Stenosis | 4.51E-02 |
| Liver Failure | 4.51E-02 |

| Supplementary Table 15. Renal necrosis and renal proliferation with which cisplatin damaged genes are associated. | | | | | |
|---|---|---|---|---|---|
| Diseases or Functions | p-value | Predicted Activation State | Activation z-score | Molecules | # Molecules |
| Cell proliferation of kidney cell lines | 3.86E-06 | | -0.092 | *ABCC5,ABCG2,ADCY1,AFF1,ALOX15,ARNTL2,BAIAP2L1,BCL2,BICC1,CABLES1,CALCR,CDCP1,CHKA,CNKSR1,CSNK1E,CTCF,DAPK1,DDR2,DLC1,E2F1,EGFR,ERBB4,ESR1,FBLN2,FBN2,FGF1,FGFR2,FGFR4,FOXO1,GAK,HGF,HNF4A,HSP90AA1,HSPA4,IGF2BP3,IQGAP2,KMT2A,LARP1,MAPK14,MAPK9,MTOR,MYOC,NINL,NTN1,OSBP2,P2RX7,PEMT,PIK3CD,PKD1,PLAAT4,PLAUR,PLD1,PLD2,PMP22,POLG,PPP2R1B,PTCH1,PTGER3,PTGES,PTP4A3,RALA,RASSF5,RBM25,RCC1,RET,RHOA,RNF144B,RXRA,RYBP,SALL4,SCUBE3,SDHB,SLC22A6,SPDYA,SPOCK3,STAT3,STT3B,SYK,TEAD4,TERT,TFDP1,TGFB1,TSC1,TSC22D3,TTLL4,TXNRD1,UBE2I,UNC5A,UNC5B,UNC5C,VDAC1,VTCN1,YAP1,ZMIZ2,ZNF512B,ZSCAN18* | 96 |
| Cell death of kidney cell lines | 2.55E-04 | Decreased | -4.652 | *AAK1,AATF,ABCC1,ABCG2,ALDH3B1,AMOTL1,APP,ATF6,ATG5,ATG7,ATP1A4,ATP2B2,BCL2,BCL2L1,BCL2L11,BCL2L14,BID,BIK,BIRC2,BNIP3L,BRAF,BRCA1,CA4,CABLES1,CALCR,CARD14,CASP8,CASP9,CASR,CD44,CDC42,CDON,CHMP6,CIDEC,CNNM4,DCC,EMP2,EPHB2,ERN1,F2,FGF1,FGFR4,FOXO1,FYN,GAS7,GRB10,GRIN2B,GRK2,GRM1,GSK3B,GSN,HDAC6,HGF,HIPK2,HK1,HK2,HTT,IGF1R,IKBKG,IL17RD,IP6K2,ITCH,ITPR1,JAK2,KCNB1,KRAS,KYAT1,LDHA,MAP2K1,MAP2K4,MAP2K6,MAP3K1,MAP3K5,MAPK8,MAPT,MAVS,MEFV,MELK,MIB1,MLKL,MYOC,NDEL1,NF2,NLRP11,NMNAT3,NOS2,NTRK3,P2RX7,PAK2,PARK7,PAWR,PAX2,PDCD6IP,PDPK1,PIAS1,PKD1,PKN2,PLA2G4A,PLCB1,PMP22,PPARD,PPARG,PRKAA1,PRKCB,PRKDC,PSEN1,PTCH1,PTGIS,RAC1,RAD51,RASSF4,RASSF5,RHOA,RHOT1,RIPK1,RNF216,SDHC,SGPL1,SH3RF1,SIAH1,SIRT2,SIRT4,SLC47A1,SREBF2,STAT3,STK35,STK4,STRA6,TCF12,TCF3,TCF4,TERT,TFRC,TGFB1,TIAM1,TICAM1,TJP2,TNFRSF10A,TNFRSF10B,TNFRSF1B,TP53,TP53BP2,TP73,TRAF1,TRPM2,TRPM7,TRPS1,UMOD,UNC13B,UNC5B,VDAC1,VOPP1,XIAP,YAP1,ZBP1,ZNF512B* | 156 |
| Cell death of kidney cells | 4.10E-04 | Decreased | -4.552 | *AAK1,AATF,ABCC1,ABCG2,ALDH3B1,AMOTL1,APP,ATF6,ATG5,ATG7,ATP1A4,ATP2B2,BCL2,BCL2L1,BCL2L11,BCL2L14,BID,BIK,BIRC2,BNIP3L,BRAF,BRCA1,CA4,CABLES1,CALCR,CARD14,CASP8,CASP9,CASR,CD44,CDC42,CDON,CHMP6,CIDEC,CNNM4,DCC,EMP2,EPHB2,ERN1,F2,FGF1,FGFR4,FOXO1,FYN,GAS7,GRB10,GRIN2B,GRK2,GRM1,GSK3B,GSN,HDAC6,HGF,HIPK2,HK1,HK2,HTT,IGF1R,IKBKG,IL17RD,IP6K2,ITCH,ITPR1,JAK2,KCNB1,KRAS,KYAT1,LDHA,MAP2K1,MAP2K4,MAP2K6,MAP3K1,MAP3K5,MAPK8,MAPT,MAVS,MEFV,MELK,MIB1,MLKL,MYOC,NDEL1,NF2,NLRP11,NMNAT3,NOS2,NPHS1,NTRK3,P2RX7,PAK2,PARK7,PAWR,PAX2,PDCD6IP,PDPK1,PIAS1,PKD1,PKN2,PLA2G4A,PLCB1,PMP22,PPARD,PPARG,PRKAA1,PRKCB,PRKDC,PROC,PSEN1,PTCH1,PTGIS,RAC1,RAD51,RASSF4,RASSF5,RHOA,RHOT1,RIPK1,RNF216,SDHC,SGPL1,SH3RF1,SIAH1,SIRT2,SIRT4,SLC47A1,SREBF2,STAT3,STK35,STK4,STRA6,TCF12,TCF3,TCF4,TERT,TFRC,TGFB1,TIAM1,TICAM1,TJP2,TNFRSF10A,TNFRSF10B,TNFRSF1B,TP53,TP53BP2,TP73,TRAF1,TRPM2,TRPM7,TRPS1,UMOD,UNC13B,UNC5B,VDAC1,VOPP1,XIAP,YAP1,ZBP1,ZNF512B* | 158 |
| Apoptosis of kidney cell lines | 5.21E-03 | Decreased | -3.809 | *AATF,ABCC1,AMOTL1,APP,ATF6,ATP2B2,BCL2,BCL2L1,BCL2L11,BCL2L14,BID,BIK,BIRC2,BNIP3L,BRAF,BRCA1,CA4,CABLES1,CALCR,CARD14,CASP8,CASP9,CASR,CD44,CDON,CIDEC,DCC,F2,FGF1,FGFR4,FOXO1,GAS7,GRB10,GRIN2B,GRK2,GRM1,GSK3B,GSN,HGF,HIPK2,HTT,IL17RD,ITCH,ITPR1,JAK2,KRAS,MAP2K1,MAP2K4,MAP2K6,MAP3K1,MAP3K5,MAPK8,MAPT,MEFV,MELK,MYOC,NF2,NLRP11,NOS2,P2RX7,PAK2,PAWR,PAX2,PDPK1,PIAS1,PKD1,PKN2,PPARD,PRKAA1,PTCH1,PTGIS,RAC1,RASSF4,RASSF5,RHOT1,RIPK1,RNF216,SDHC,SGPL1,SH3RF1,SIAH1,SREBF2,STAT3,STK35,STK4,STRA6,TCF12,TCF3,TCF4,TERT,TFRC,TGFB1,TIAM1,TICAM1,TJP2,TNFRSF10A,TNFRSF10B,TP53,TP53BP2,TP73,TRAF1,TRPS1,UMOD,UNC13B,VDAC1,VOPP1,XIAP,YAP1,ZNF512B* | 109 |
| Apoptosis of podocytes | 4.51E-02 | | | *NPHS1,PROC,TGFB1* | 3 |

| Supplementary Table 16. Genes selected for Western Blot Assays. | | | | | | | | | | |
|---|---|---|---|---|---|---|---|---|---|---|
| Gene name | Chromosome | Average number of identified peaks (n=2) | Average number of tags (n=2) | Average enrichment fold (n=2) | Protein name | Type | Cell type for WB | Cisplatin dosage | Catalog number of antibody (Abcam) | Role in cell |
| EGFR | chr7 | 52 | 2523 | 122 | epidermal growth factor receptor | kinase | A549 | 50 µM | ab52894 | proliferation, expression in, growth, apoptosis, phosphorylation in, activation in, migration, survival, cell death, invasion |
| PRKCZ | chr1 | 56 | 2601 | 116 | protein kinase C zeta | kinase | A549 | 50 µM | ab59364 | apoptosis, proliferation, phosphorylation in, activation in, survival, growth, expression in, morphology, motility, formation |
| PDE4D | chr5 | 73 | 1774 | 178 | phosphodiesterase 4D | enzyme | A549 | 50 µM | ab171750 | expression in, apoptosis, proliferation, phosphorylation in, production in, response, response by, ubiquitination in, binding in, growth |
| PRKCI | chr3 | 21 | 851 | 47 | protein kinase C iota | kinase | A549 | 50 µM | ab5282 | expression in, growth, apoptosis, invasion, migration, production in, transformation, invasion by, replication in, activation in |
| PTK2 | chr8 | 50 | 1337 | 122 | protein tyrosine kinase 2 | kinase | A549 | 50 µM | ab131435 | migration, proliferation, apoptosis, expression in, activation in, phosphorylation in, adhesion, growth, cell spreading, invasion |
| PTK2B | chr8 | 32 | 1120 | 72 | protein tyrosine kinase 2 beta | kinase | A549 | 50 µM | ab226798 | migration, apoptosis, proliferation, adhesion, activation in, phosphorylation in, growth, response by, tyrosine phosphorylation in, cell spreading |
| SRC | chr20 | 26 | 1370 | 55 | SRC proto-oncogene, non-receptor tyrosine kinase | kinase | A549 | 50 µM | ab109381 | transformation, phosphorylation in, proliferation, apoptosis, tyrosine phosphorylation in, migration, activation in, expression in, growth, adhesion |
| PDE1C | chr7 | 52 | 1526 | 115 | phosphodiesterase 1C | enzyme | HepG2 | 50 µM | ab98194 | regulates proliferation and migration of vascular smooth muscle cells, and neointimal hyperplasia, plays a role in pathological vascular remodeling by regulating the stability of growth factor receptors, such as PDGF-receptor-beta |
| EPHB1 | chr3 | 109 | 3202 | 227 | EPH receptor B1 | kinase | HeLa | 25 µM | ab129103 | guidance, chemotaxis, adhesion, migration, abnormal morphology, pruning, attachment, refinement, formation, proliferation |
| EPHB2 | chr1 | 79 | 3304 | 189 | EPH receptor B2 | kinase | NCCIT | 12 µM | ab252935 | guidance, abnormal morphology, outgrowth, expression in, adhesion, tyrosine dephosphorylation in, phosphorylation in, tyrosine phosphorylation in, plasticity, interaction in |
| PLCG2 | chr16 | 52 | 1602 | 122 | phospholipase C gamma 2 | enzyme | NCCIT | 12 µM | ab109267 | differentiation, number, apoptosis, function, proliferation, maturation, development, activation, activation in, expression in |
| MERTK | chr2 | 24 | 801 | 56 | MER proto-oncogene, tyrosine kinase | kinase | NCCIT | 12 µM | ab52968 | phagocytosis by, proliferation, apoptosis, expression in, activation, degeneration, phagocytosis, abnormal morphology, phosphorylation in, number |
| PDE2A | chr11 | 37 | 2165 | 82 | phosphodiesterase 2A | enzyme | NCCIT | 12 µM | ab224616 | response by, response, cell death, organization, differentiation, angiogenesis by, activation in, production in, proliferation, migration |

| | | | | | | | | | | |
|---|---|---|---|---|---|---|---|---|---|---|
| PRKCA | chr17 | 78 | 3258 | 180 | protein kinase C alpha | kinase | NCCIT | 12 μM | ab32376 | phosphorylation in, apoptosis, expression in, proliferation, migration, activation in, growth, transcription in, binding in, adhesion |
| LMTK2 | chr7 | 26 | 1170 | 62 | lemur tyrosine kinase 2 | kinase | NCCIT | 12 μM | ab172622 | guidance |
| IGF1R | chr15 | 68 | 3199 | 158 | insulin like growth factor 1 receptor | transmembrane receptor | NCCIT | 12 μM | ab182408 | proliferation, apoptosis, expression in, growth, migration, phosphorylation in, activation in, transformation, survival, cell death |
| INSR | chr19 | 68 | 1990 | 163 | insulin receptor | kinase | NCCIT | 12 μM | ab137747 | proliferation, growth, differentiation, migration, apoptosis, mass, mitosis, mitogenesis, activation in, expression in |
| PRKG1 | chr10 | 67 | 2338 | 152 | protein kinase cGMP-dependent 1 | kinase | NCCIT | 12 μM | ab97339 | phosphorylation in, aggregation, migration, expression in, activation in, clearance, abnormal morphology, number, proliferation, apoptosis |
| SPAG9 | chr17 | 23 | 606 | 61 | sperm associated antigen 9 | other | NCCIT | 12 μM | ab12331 | differentiation, migration, localization, growth, activation in |
| AKT3 | chr1 | 22 | 654 | 57 | AKT serine/threonine kinase 3 | kinase | A549 | 50 μM | ab152157 | proliferation, growth, apoptosis, migration, expression in, necrosis, invasion, autophagy by, aneuploidy, activation in |
